# Protein coevolution and physicochemical adaptation in the APAF-1/apoptosome: structural and functional implications

**DOI:** 10.1101/2020.11.24.392506

**Authors:** José A Sánchez-Borbón, Steven E Massey, José D Hernández-Martich

## Abstract

The apoptosome is involved in the mitochondrial pathway of apoptosis, consisting of APAF-1, caspase 9 and cytochrome c, forming a heptamer that activates effector caspases, causing cell death. This protein complex has also been characterized in *Drosophila melanogaster* (DARK) and *Caenorhabditis elegans* (CED-4). Here we present an evolutionary guided *in silico* characterization of the APAF-1/apoptosome. The evolutionary history of the apoptosome was determined, taking the possible orthologs of the APAF-1 version, and executing a protein coevolution and a positive selection analysis, to make structural and functional inferences and identify residues under destabilizing changes, respectively. Results suggests that the APAF-1/apoptosome is not unique to vertebrates, but also some basal invertebrates could possess orthologous copies. New possible versions were also detected in other taxa. Not all insects and other arthropods have the DARK version, just as not all nematodes have the CED-4 version. In the APAF-1 version, amino acid clusters with coevolution signal located in the interior, gave more insights on new potential interactions, allowing us to infer a more detailed model that includes allosterism, of how cytochrome c associates with β propellers during APAF-1 activation, as well as interactions essential for nucleotide exchange, activation of CASP9, the molecular timer and other pathways in mitochondria to induce apoptosis. Residues on the surface under destabilizing changes have guided the protein complex in adaptations necessary for conformational changes, interactions and folding.

## Introduction

Under certain physiological and pathological conditions, multicellular organisms must eliminate unnecessary or undesirable cells through regulated mechanisms, to maintain homeostasis. Regulated cell death is the product of the activation of signal transduction, which can be pharmacologically or genetically modulated (Galluzi et al., 2018). Among its main mechanisms is apoptosis (Kerr et al., 1972). One of the main pathways of apoptosis is the intrinsic or mitochondrial, which can be initiated by perturbations of both the extracellular and intracellular environments (Galluzi et al. 2018). These perturbations cause the permeabilization of the outer membrane of the mitochondria due to the formation of BAX / BAK pores, releasing pro-apoptotic proteins such as cytochrome c (CYCS). Once released into the cytoplasm, CYCS associates with APAF-1, which associates with CASP9, forming a heptamer called apoptosome. Through the apoptosome, CASP9 activates CASP3, which in turn activates caspases 6 and 7, causing cell death (Elmore, 2007). The protein complex has also been characterized in the fruit fly *Drosophila melanogaster* and the nematode *Caenorhabditis* elegans, which have the apoptosomes made up of DARK and CED-4, respectively. Despite being homologous, they have differences in their structure and functional dynamics. The *Drosophila* apoptosome is an octamer of DARK (Yu et al., 2006). Despite having the regulatory region, CYCS is not necessary for the activation of the protein, and there is another, as yet unknown, ligand that associates with β propellers (Dorstyn et al., 2002). In *C. elegans* the protein complex is formed by an octamer of CED-4 (Qi et al., 2010). CYCS is also not involved in its activation, due to the absence of β propellers in it (Lettre and Hengartner, 2006).

One of the phenomena by which proteins evolve is coevolution. This process occurs when a substitution of an amino acid located in a specific site, generates a compensatory substitution of another amino acid located in another site (Altschuh et al., 1987; Altschuh et al., 1988; Taylor and Hatrick, 1994). These series of mutations can occur in speciation events. There are various evolutionary forces or constraints that influence how these compensatory changes occur. One of them is the structural restriction due to orthosteric interactions between residues (Göbel et al., 1994; Shindyalov et al., 1994). The other type of restriction is functional, in which the amino acids under coevolution may be in distant positions in the protein, this due to their binding to the same ligand or due to allosteric interactions (Süel et al., 2003; Anishchenko et al., 2017). The amino acids under coevolution can be located in the same protein or in different proteins:intraprotein and interprotein coevolution, respectively (Pazos et al., 1997).

Changes in amino acids are caused by mutations in their codons. In synonymous mutations, a nucleotide substitution generates a different codon, but it still codes for the same amino acid. In contrast, in the non-synonymous mutations, the new codon produced codes for an amino acid different from the original one (Pál et al., 2006). When non-synonymous changes produce a beneficial genetic variant, it quickly spreads through the population. This is known as positive selection (Pál et al., 2006). In some cases of positive selection, the substitutions can code for amino acids with physicochemical properties very different from the original, being “selected” by drastic evolutionary pressures, and producing radical changes in the structure and function of that protein region. All this contributes to the new protein being able to adapt physicochemically to new cellular microenvironments. McClellan et al. (2005) call this phenomenon “destabilizing positive selection”.

In this research we carried out an *in silico* characterization of the APAF-1/apoptosome, through evolutionary analysis of its sequences and three-dimensional structures. Focusing on the sequences homologous to humans, the evolutionary history of the apoptosome in Metazoa was determined taking as reference the protein sequences of the apoptosomes formed by APAF-1, DARK and CED-4, and identifying their putative orthologs in other taxa. From these sequences, the possible orthologues of the APAF-1/apoptosome were taken, identifying amino acids with coevolution signal and those under destabilizing positive selection. We use this information to make structural and functional inferences about the protein complex.

## Results

Homologous sequences similar to APAF-1/apoptosome in Metazoa were searched in GenBank (Supplementary file 1), detecting putative orthologs to human sequences. For this, similarity analyses were made through the Ortho-Profile method (Szklarczyk et al., 2012), in which the human sequences against invertebrates were subjected to a series of BLASTs and HMMs in three stages with increased sensitivity, sequence against sequence, profile against sequence and HMM against HMM. If in one of the phases, the sequences show reciprocity with e-values less than 0.01, they are considered orthologs. Orthology was also detected through reconciliation of phylogenies, also using as reference the DARK (*D. melanogaster*) and CED-4 (*C. elegans*) apoptosomes.

### Proteins orthologous to human sequences identified by similarity

Of the homologous proteins in invertebrates, only 20 passed the test (Supplementary file 2). In the first phase, six proteins were detected; three in the cnidarians *Acropora millepora*, *Hydra vulgaris* and *Orbicella faveolata*; the echinoderm *Strongylocentrotus purpuratus*; the hemichord *Saccoglossus kowalevskii* and the cephalochord *Branchiostoma floridae*. In the second phase, 14 proteins were identified; one in the poriferous *Amphimedon queenslandica* and the other 13 within Protostomia; three in the nematodes *Trichinella nativa*, *T. papuae* and *T. zimbabwensis*; 10 in arthropods distributed in two branchiopods, *Daphnia magna* and *D. pulex*; the arachnid *Parasteatoda tepidariorum*; the entognatha *Orchesella cincta* and six insects, *Amyelois transitella*, *Athalia rosae*, *Neodiprion lecontei*, *Ooceraea biroi*, *Trachymyrmex cornetzi* and *Trichogramma pretiosum*. In the third phase, for CASP9, in *Lingula anatina* and *S. purpuratus*, possible orthologs were detected, but they did not coincide with the homologous proteins.

### Proteins orthologous to human sequences identified by reconciliation of phylogenies

The APAF-1 phylogeny reconciled with its species tree shows 10 duplications. Close to *H. sapiens* there are four duplication events (Figure 1). One that gives rise to two speciation events; to entognatha and another clade that includes Porífera, Priapúlida, Xiphosura, Plathelminthes, Enoplea and Arachnida. Branchiopoda arises from another duplication. Upon reaching deuterostomes, another duplication occurs with two speciation events. One goes to Cnidaria, Echinodermata, Hemichordata and Cephalochordata and the other to Vertebrata. Taking into account that in the unreconciled phylogeny, these phyla are found in the same clade (Figure 1-figure supplement 1), we interpret that the sequences involved in this duplication event would be co-orthologous. In birds there is a duplication where Falconiformes, Passeriformes and Apodiformes diverge. In insects there are six duplication events (Figure 2). One of them gave rise to Isoptera and Hemiptera. Another duplication gives rise to the coleopteran families Cerambycidae, Nitidulidae and Tenebrionadae. Within coleoptera another duplication originates the family Silphidae. A duplication is in the node that originates Hymenoptera and Lepidoptera. Within Hymenoptera, in *Trachymirmex* there is a duplication in between *T. cornetzi* and *T. septentrionalis*. The clade of Hymenoptera and Lepidoptera also includes the beetle *Nicrophorus vespilloides* of the family Buprestidae. Diptera originates from a unique duplication event. In relation to Nematoda (Figure 3), Enoplea and Plathelminthes arise from the same duplication event. *C. elegans* originates from a unique duplication. The analysis of conserved domains in Pfam in the nematodes *Trichuris suis*, *T. nativa* and *T. papuae* and Plathelminthes weakly detected WD40 repeats that form the β propellers (Figure 3-figure supplement 1), so we decided to do more detailed analysis of conserved domains in the NCBI, confirming that unlike *C. elegans*, these species have the WD40 repeats (Figure 3-figure supplement 2).

**Figure 1.**
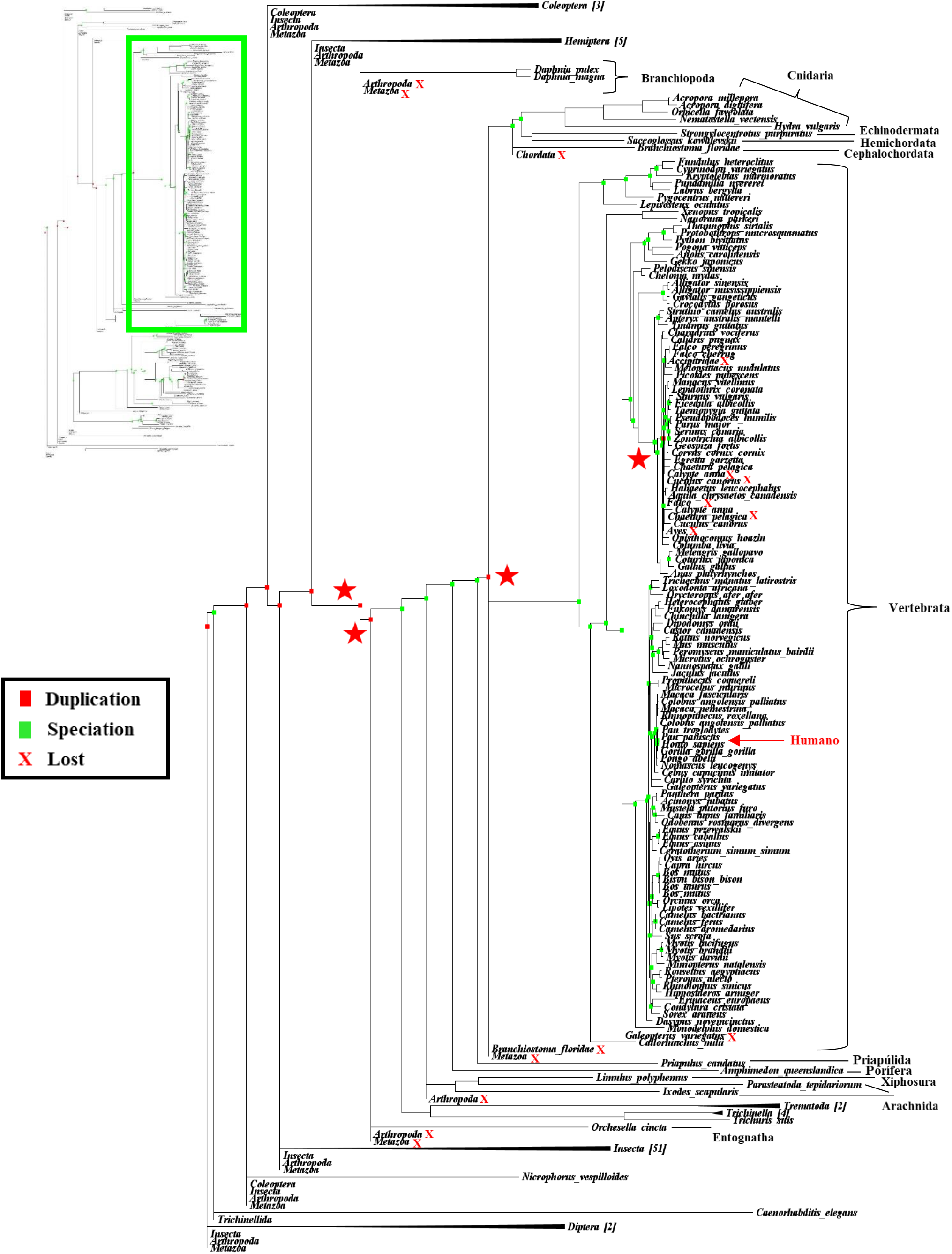
Emphasis on regions close to the human sequence of the APAF-1 phylogeny reconciled with its species tree. The stars indicate the four nodes that show duplication in the evolution of the protein during speciation events. The other nodes are collapsed. See also Figure 1-figure supplement 1.

**Figure 2.**
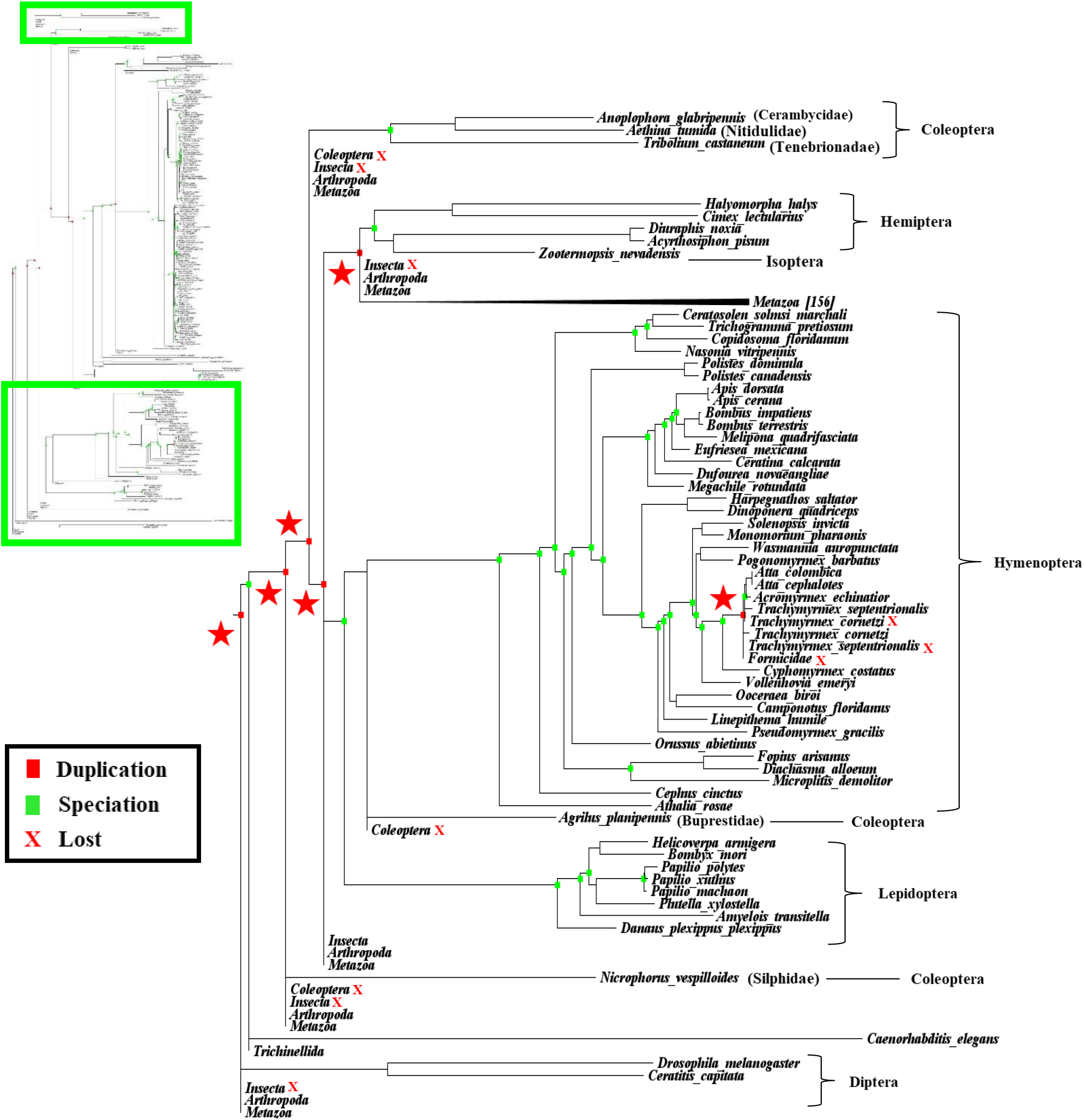
Emphasis on insects of the APAF-1 phylogeny reconciled with its species tree. The stars indicate the six nodes that show duplication in the evolution of the protein during speciation events in insects. The nodes of the other taxa are collapsed. See also Figure 2-figure supplement 1.

**Figure 3.**
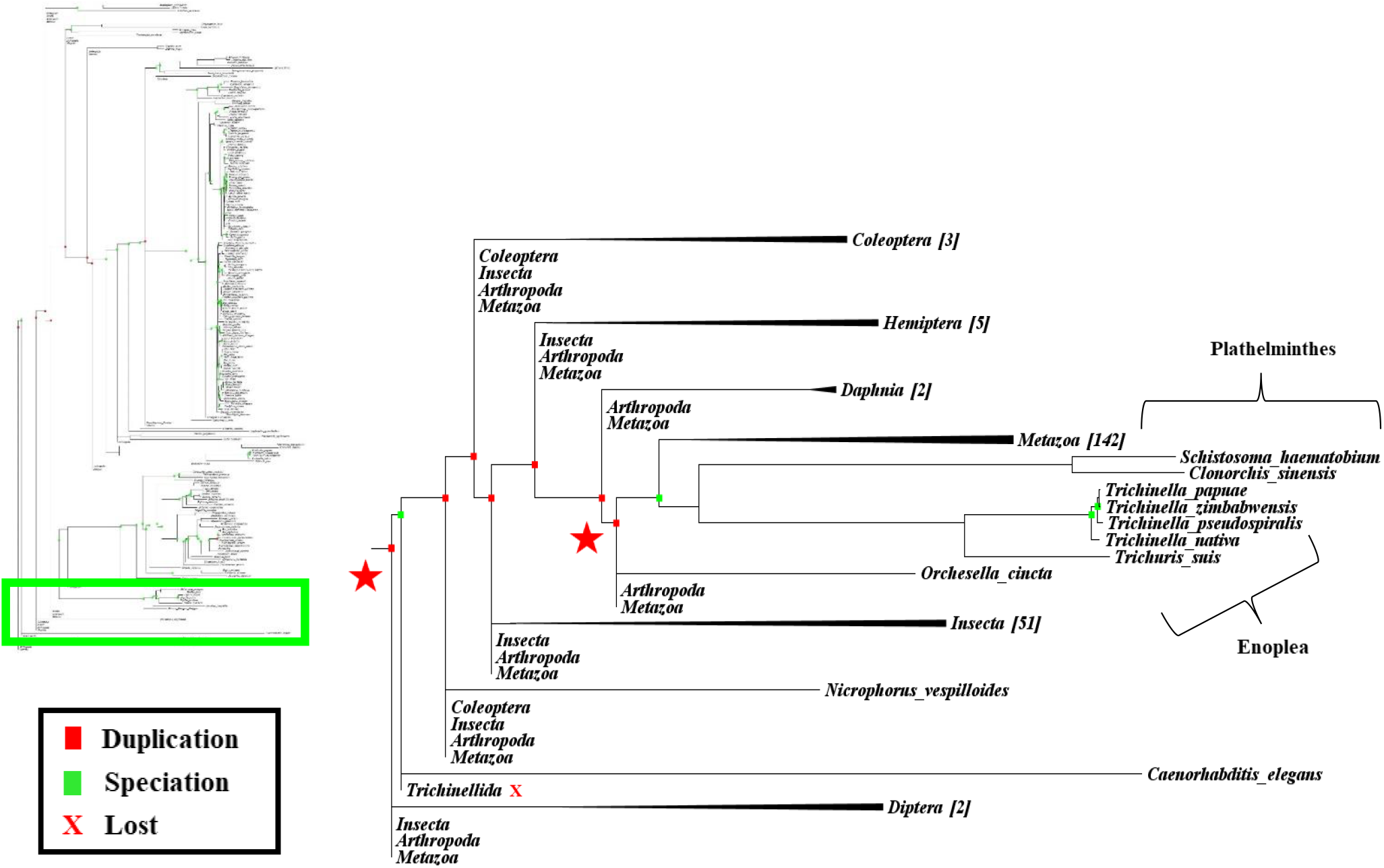
Emphasis on nematodes of the APAF-1 phylogeny reconciled with its species tree. The stars highlight the two nodes that show duplication in protein evolution during speciation events in Nematoda. The Enoplea clade also includes Plathelminthes. The nodes of the other taxa are collapsed. See also Figure 3-figures supplements 1 and 2.

The phylogeny of CASP9 reconciled with its species tree presents six duplications (Figure 4). *H. sapiens* is included in a duplication event at the node where deuterostomes originate with a speciation event encompassing echinoderms, *B. floridae*, and other vertebrates. In the latter there are three duplication events; one at the node where reptiles and birds emerge with two speciation events, one towards Squamata and the other towards Crocodylia, Testudines and Aves. A duplication occurs at the node where the Hominidae family originates with two speciation events, one going towards *Pongo abelii*, *H. sapiens*, *Pan Paniscus* and the other towards *P. troglodytes*. The other duplication event within vertebrates is at the node common to Eulipotyhla, separating *Sorex araneus* from the rest of the order. The clade of the deuterostomes also includes Porífera, the arachnid *P. tepidariorum* and the mollusk *Octopus bimaculoides*. *Branchiostoma belcheri* is in a unique speciation event. The *Aplysia calyfornica* mollusk and the lingulate *L. anatina* are in the same speciation event. At the node where Diptera arises there is a duplication with two branches of speciation. One goes to *D. melanogaster* and the other covers the other Diptera *D. kikkawai*, *D. takahashii* and *Bactrocera oleae*. Lepidoptera arises from a unique duplication event far removed from other insects. Plathelminthes are in a unique speciation event.

**Figure 4.**
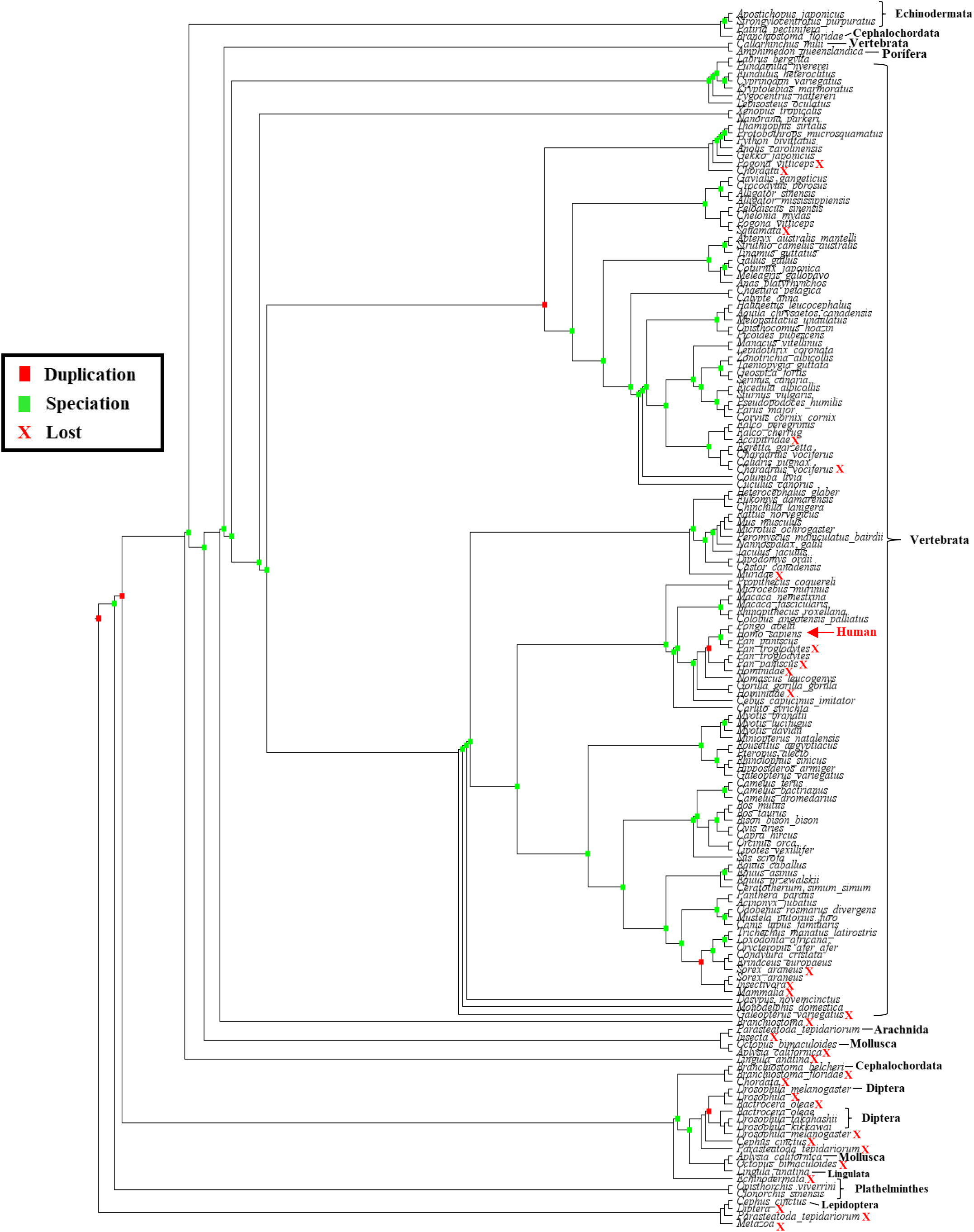
CASP9 phylogeny reconciled with its species tree. The six nodes that show duplication in the evolution of the protein during speciation events in invertebrates and vertebrates are shown. See also Figure 4-figure supplement 1.

The automatically generated species trees in the NCBI taxonomy database are polytomous. Another aspect to take into account is that these trees are cladograms, not phylogenies as such, so the branches do not have the real evolutionary distances. Taking the previous statements as caveats, they could have generated the following possible errors in the reconciliations; the duplication events in both proteins within vertebrates. As a support of this, APAF-1 has not been duplicated in vertebrates (Zmasek et al., 2007). The duplication event in APAF-1/DARK homologues found within Hymenoptera, and coleopterans outside their respective clade. The APAF-1 of Porifera and Priapulida should originate in the same duplication event as cnidarians, echinoderms, hemicordates, and cephalochords. CASP9 of *O. bimaculoides* and *P. tepidariorum* must be in a clade separate from that of vertebrates. If the phylogenies of species had been made by ourselves, the results would be more accurate. There are also the following possible errors that we believe are the result of false positives during the search for CASP9 homologs; the possible cause of *B. belcheri* being in a node other than *B. floridae* is that it is identified as caspase 7 in the database. The sequences of the Diptera, *D. kikkawai*, *D. takahashii* and *B. oleae* according to the analysis of conserved domains are not well characterized, so we suggest that in this case they may also be caspases other than CASP9/DRONC. This may be the reason why these sequences originate from a node other than *D. melanogaster*.

The results of the first phase of Ortho-Profile were very similar to the unreconciled phylogeny of APAF-1. In the second phase, only the APAF-1 of *A. queenslandica* matches the phylogeny (Supplementary file 2 and Figure 1-supplement figure 1). Considering the low precision of the second phase, we think that the third phase would have even less precision because it has more flexible restrictions, which suggests that they are mostly false positives.

From the putative orthologs detected, we selected the species that have the two proteins available, to apply protein coevolution and destabilizing positive selection analyses. In invertebrates these were the poriferous *A. queenslandica*, the echinoderm *S. purpuratus* and the cephalochord *B. floridae*, plus the 115 species of vertebrates including the reference species *H. sapiens*, making a total of 119 species. For each of them, the CYCS sequences were obtained to complete the proteins that make up the APAF-1/apoptosome to include them in the analyses (Supplementary file 1).

### Amino acids with coevolution signal in the APAF-1/apoptosome

The concatenated alignment with the proteins of the selected species is 5,426 positions long including gaps; together with its respective phylogeny were uploaded to the BIS2Analyzer server (http://www.lcqb.upmc.fr/BIS2Analyzer/) (Oteri et al., 2017), which detects amino acids with a coevolution signal. The program detected 23 amino acid clusters distributed in the different protein domains (Figure 5). Clusters 21, 22, and 23 have symmetric and environmental values of 0.55, 0.83, 0.55, 0.65, and 0.73, 0.5, respectively. Clusters 1 to 20 have values 1, 1, with a perfect coevolution signal (Supplementary files 3 and 4 and Figure 5-table supplement 1).

**Figure 5.**
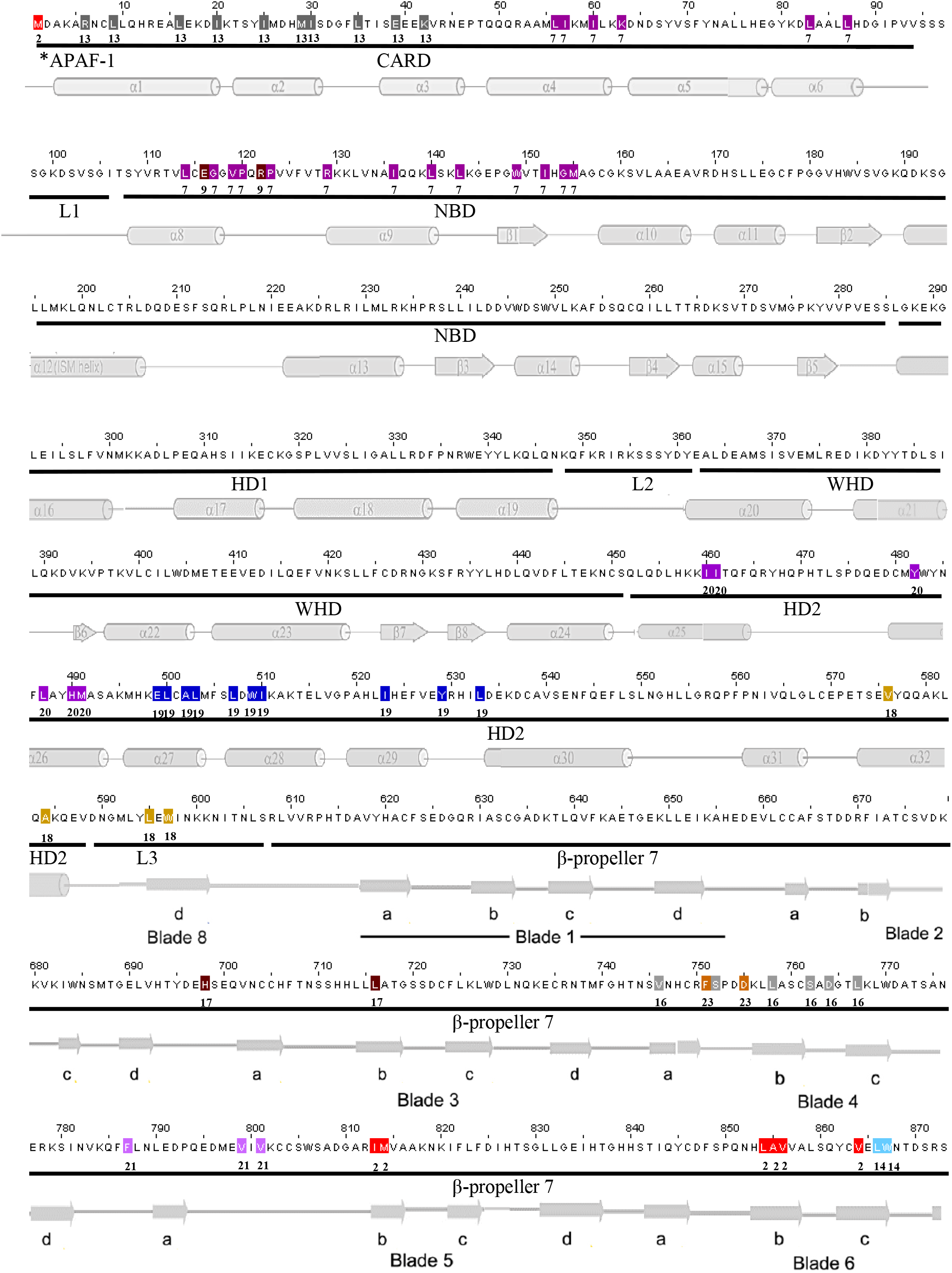
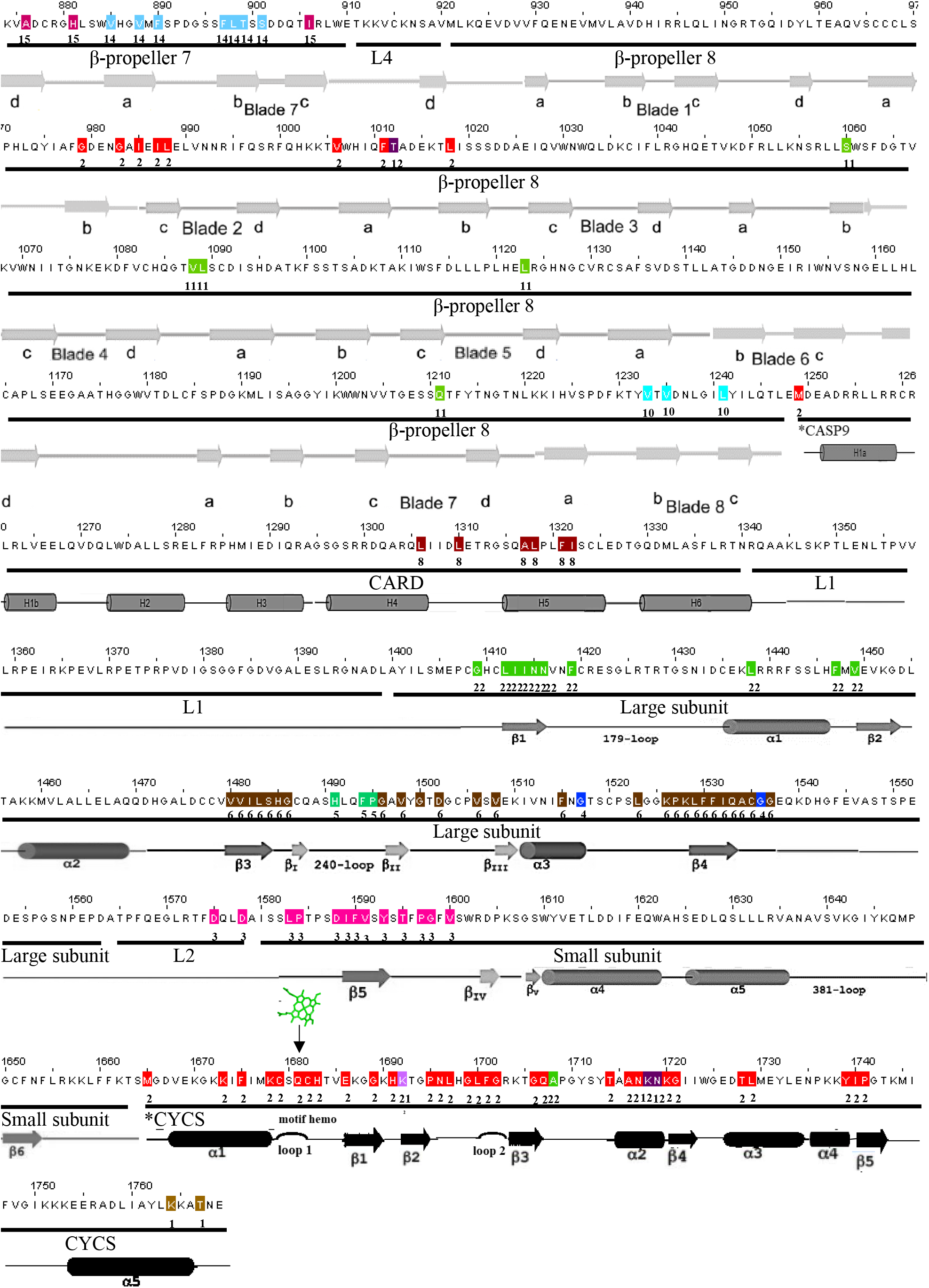
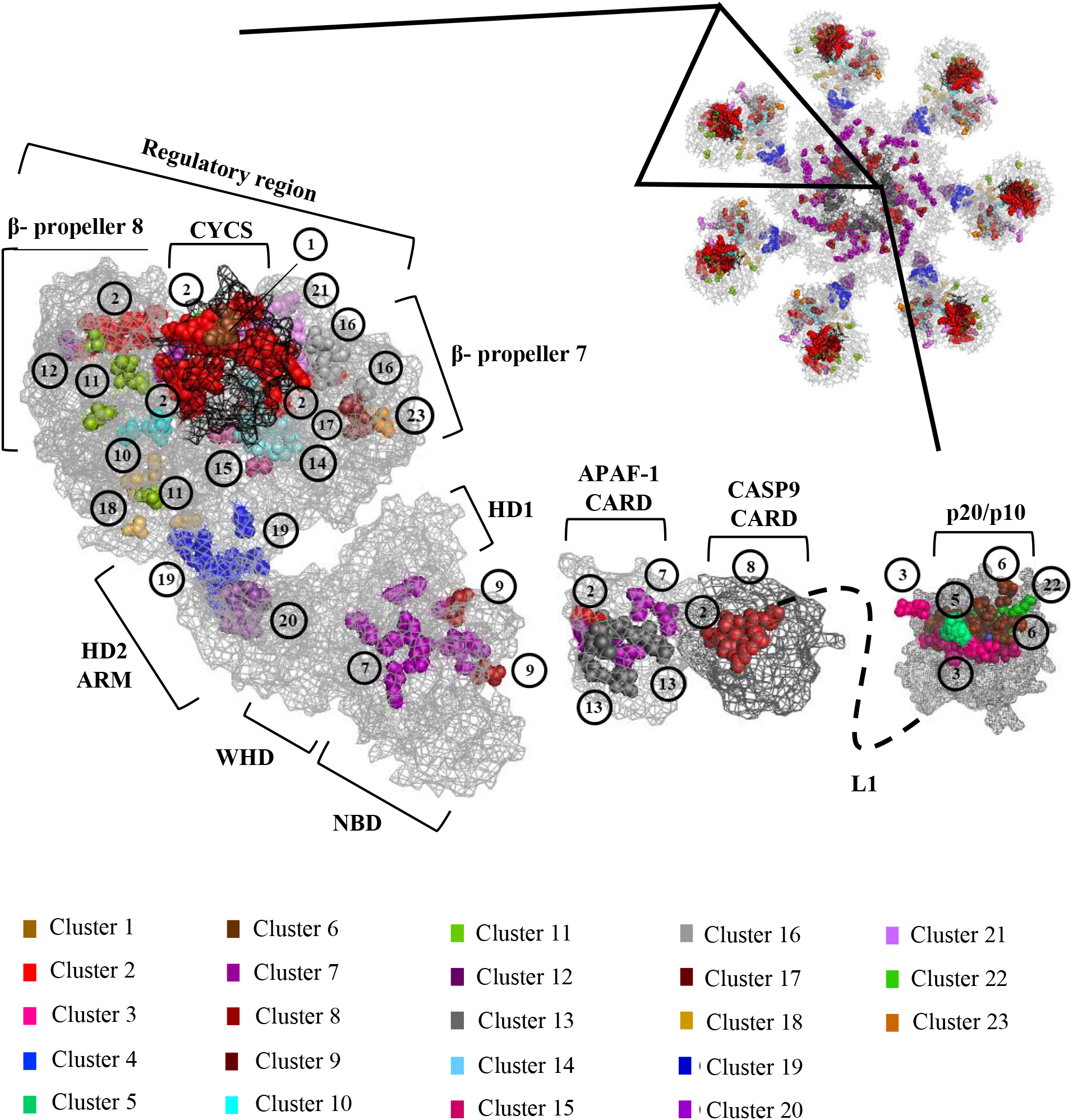
Clusters of amino acids with coevolution signal mapped in the concatenated human sequences and a single unit of the APAF-1/apoptosome. **A)** The protein sequences of *H. sapiens* are shown. The numbers above it indicate the amino acid positions within the sequences. The lines below delimit which protein and domain these residues belong to. Asterisks indicate where a protein begins. Secondary structures are shown below in their respective colors, gray for APAF-1, dark gray for CASP9, and black for CYCS. The heme group is shown as green sticks located in its motif in loop 1. The amino acids within each cluster were marked with a specific color. To facilitate the visualization of these groupings, the numbers of the clusters to which they belong were placed under their respective residues. For original positions of the CASP9 and CYCS residues see Figure 5-table supplement 1. **B)** A single unit is shown inside the PDB 5JUY heptameter in “mesh” format, which allows visualization of the residues mapped in the interior. The catalytic domain of CASP9 PDB 1JXQ was attached, representing L1 with intermittent lines. The amino acids within each cluster were marked with their respective colors. To facilitate the visualization of these groupings, the numbers of the clusters to which they belong were placed near their respective residues. See also Figure 5-figure supplement 1.

### Amino acids affected by destabilizing positive selection in the APAF-1/apoptosome

A concatenated alignment with the coding DNA of the components of the apoptosome in the selected species and their respective phylogeny was supplied to TreeSAAP (Woolley et al., 2003), to detect amino acids under positive selection. The program identified 3,929 codon substitutions during phylogeny. During these substitutions, taking as reference the 1,769 amino acids in the concatenated human sequences, 1,164 residues are under positive selection in all categories of magnitude (1-8), distributed in all protein domains. Of these, 996 residues are under destabilizing changes in the most radical categories 6, 7 and 8. In the three-dimensional structures, residues under coevolution are located inside the proteins, close to active sites and other regions involved in conformational changes, specifically in the regulatory region, the arm, the nucleotide exchange zone, the CARD domains of both proteins and the catalytic domain of CASP9, while those that are under destabilizing changes are found in the periphery (Figure 6). A few residues were detected in both analyzes; on the APAF-1, R/122, V/799, V/801, T/1012 and CYCS K/1692 and K/1718.

**Figure 6.**
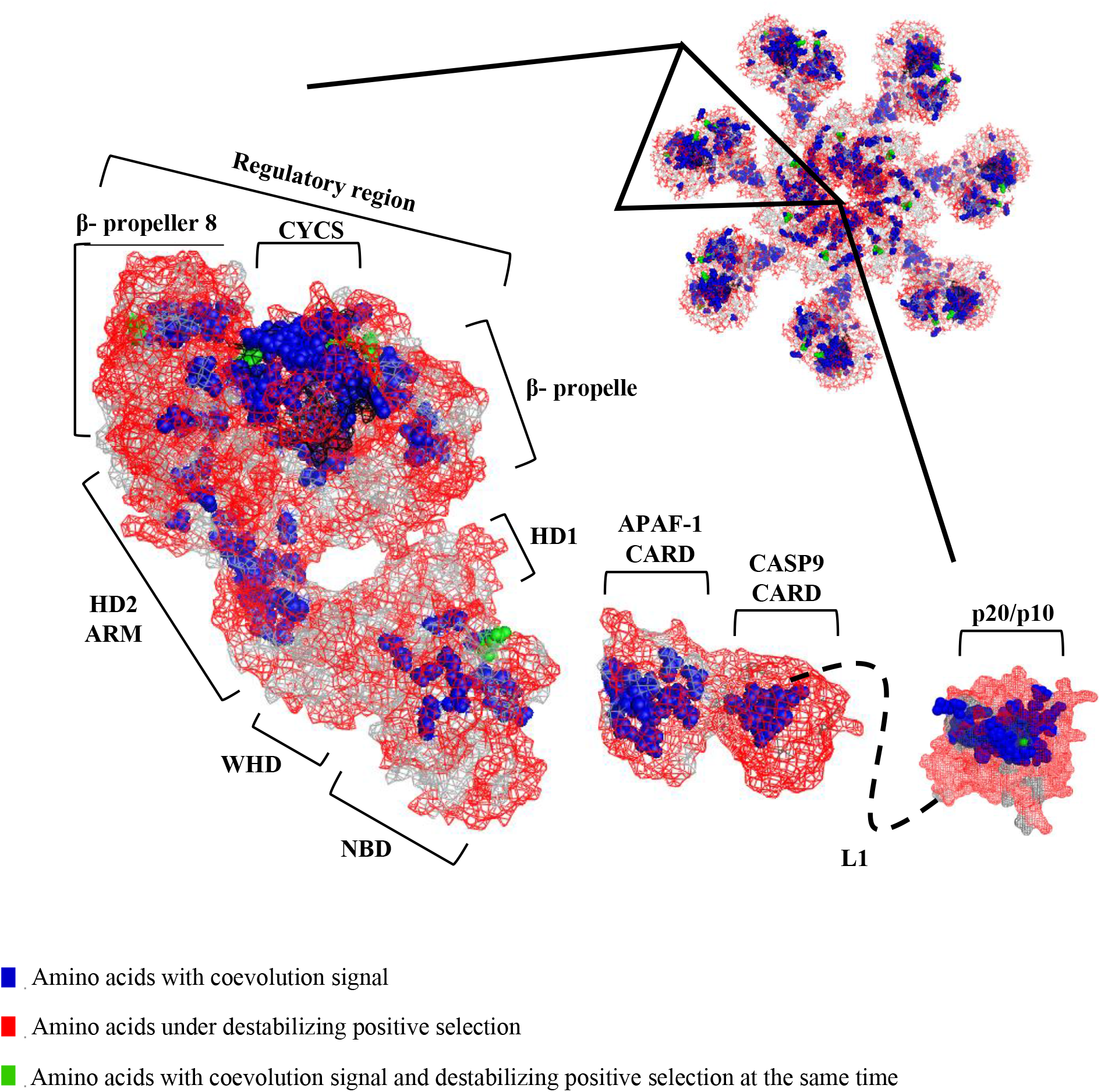
Amino acids under destabilizing positive selection mapped in a single unit of the APAF-1/apoptosome. A single unit is shown within the PDB 5JUY heptameter in “mesh” format, which allows visualization of residues in the interior. The catalytic domain of CASP9 PDB 1JXQ was attached, representing L1 with intermittent lines. The residues under destabilizing positive selection in categories 6, 7 and 8 are shown as red sticks, whose z scores are + 3.09 and −3.09. To facilitate the visualization of these amino acids, the 23 clusters under coevolution were colored blue. Green spheres indicate residues detected in both analysis. It can be seen that amino acids under coevolution are in the interior, close to active sites and other regions involved in conformational changes, while those that are under destabilizing positive selection are located in the periphery. See also Figure 6-figure supplement 1 and Supplementary file 5.

**Figure 6.**
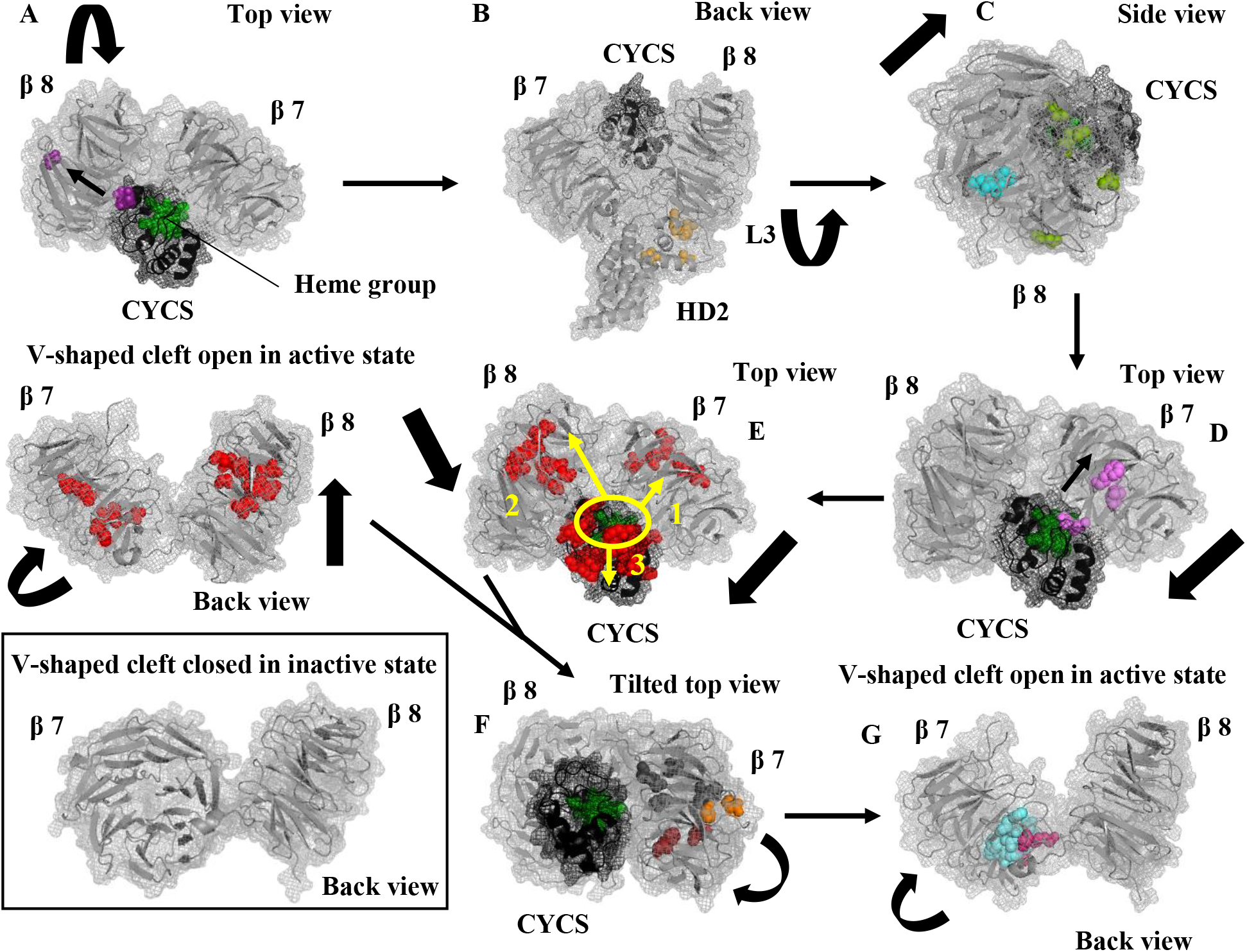
Tentative model based on clusters with coevolution signal, of sequence of events in regulatory region during APAF-1 activation. **A - G)** A hierarchical sequence of the order (arrows) is shown in how the clusters cause the conformational changes (thick arrows) during the activation of APAF-1. Clusters are shown separately or in groups with a common function for clarity. The heme group in CYCS appears as green spheres. **A)** (Cluster 12) The interaction between CYCS and the regulatory region of APAF-1 starts when CYCS residues K/54 and N/55 interact directly with D/1147 in β propeller 8; These through allosteric interactions influence T/1012 (indicated by the arrow between the residues), initiating the upward rotation of this protein domain. **B)** (Cluster 18) HD2 residues V/576, A/584 and L3 L/595, W/597 detach β propeller 8 from HD2 in the lower region. **C)** (Clusters 10 and 11) A ligand other than CYCS, by allosteric regulation interacts with β propeller 8 V/1233, V/1235 and L/1241. Independently, another ligand allosterically regulates S/1060, V/1088, L/1089, L/1123 and Q/1211, initiating the formation of the V shaped cleft at the base and finishing the conformational changes that rotate β propeller 8 upwards, allowing interaction with β propeller 7; **D)** (Cluster 21) which occurs when CYCS K/28 interacts with this domain, associating these allosterically with β propeller 7 F/787, V/799 and V/801 (indicated by the arrow between the residues), initiating the rotation and the turning of this protein domain and the formation of the V shaped cleft in the upper region. Then, **E)** (Cluster 2), more interactions are produced that cover the entire length of the contact zone, through a triple allosteric regulation with the participation of heme as a second ligand (circled and indicated by arrows), they influence CYCS K/9, K/14, E/22, G/25, H/27, P/31, N/32, L/33, G/35, L/36, F/37, G/38, G/42, T/50, A/52, N/53, T/64, L/65, Y/75 and I/76, β propeller 7 I/813, M/814, L/854, A/855, V/856 and V/864 and β propeller 8 G/979, G/983, I/985, I/987, L/988, V/1006, F/1011 and L/1018, causing conformational changes that assist L4 to continue the formation of the V shaped cleft, causing CYCS to start to fit into it. The CYCS was removed in order to clearly visualize the cleft. To compare the cleft with the inactive/closed state, the regulatory region is shown in its inactive PDB 3SFZ state. **F)** (Clusters 16, 17 and 23) Some ligand other than CYCS by allosteric interactions regulates β propeller 7 V/746, S/752, L/758, S/762, D/764, L/767, H/698, L/716, F/751 and D/755, which completes the wide turn in which β propeller 7 “hugs” the CYCS, staying this protein between both β propellers. **G)** (Clusters 14 and 15) Some ligand binds allosterically to β propeller 7 L/866, W/867, V/885, V/888, F/890, F/897, L/898, T/899, S/901, A/876, H/881 and I/906, causing conformational changes in the base of β propeller 7, finishing the formation of the V shaped cleft, making the CYCS fit completely in it. Some of the residues of these clusters break the salt bridge that binds NBD K/192 and β propeller 7 D/616, causing the alterations that release CARD for the activation of CASP9, finishing the conformational changes between CYCS and β propellers during APAF-1 activation. See also Video 1.

The destabilizing substitutions have significantly affected nine physicochemical properties in the following protein domains, taking into account regions under strong selection (protein regions with a large agglomeration of affected residues):helical contact area (*Ca*) (Richmond and Richards, 1978) (APAF-1 HD2, β propellers 7 and 8, CASP9 and CYCS), compressibility (*K0*) (Gromiha and Ponnuswamy, 1993) (APAF-1 β propellers 7 and 8, CASP9 and CYCS), hydrophobicity by thermodynamic transfer (*Ht*) (Zimmerman *et al*., 1968) (APAF-1 NOD, HD2 and β propeller 7, CASP9 p20/p10 and CYCS), chromatographic index (*RF*) (Zimmerman *et al*., 1968), which affects the majority of residues in the protein complex (Table 1) (APAF-1 CARD, NOD, HD2, β propellers 7 and 8, CASP9 and CYCS), refractive index (*μ*) (Prabhakaran and Ponnuswamy, 1979) (APAF-1 WHD, HD2, β propellers 7 and 8, CASP9 p20/p10 and CYCS), molecular weight (*Mw*) (Fasman, 1976) (APAF-1 HD2 and β propeller 7, CASP9 CARD), bulkiness (*Bl*) (Zimmerman *et al*., 1968) ( APAF-1 β propeller 8 and CASP9), partial specific volume (*V0*) (Gromiha and Ponnuswamy, 1993) (APAF-1 β propeller 7, CASP9 CARD) and molecular volume (*Mv*) (Grantam, 1974) (APAF-1 β propeller 8, CASP9 and CYCS) (See sliding windows in Figure 6-figure supplement 1). The same amino acid can be involved in several categories of magnitude, tendencies and physicochemical properties at the same time, because homologues to that residue can have substitutions of different natures at different nodes (Supplementary file 5).

**Table 1.**
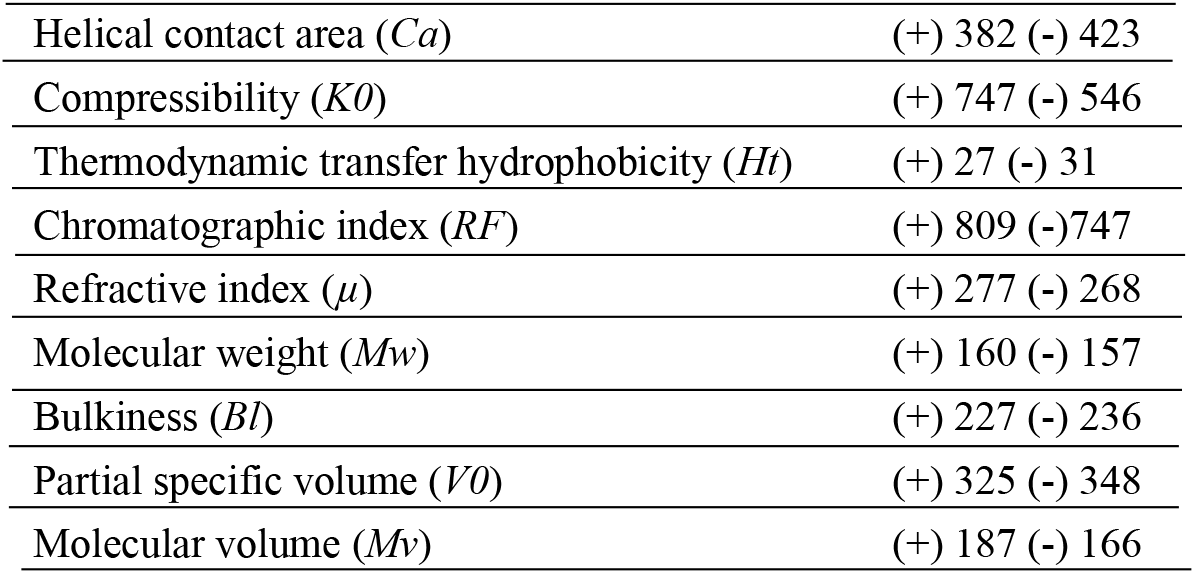
Number of destabilizing residues divided into positive and negative tendencies.

|  |  |
| --- | --- |
| Helical contact area (*Ca*) | (+) 382 (-) 423 |
| Compressibility (*K0*) | (+) 747 (-) 546 |
| Thermodynamic transfer hydrophobicity (*Ht*) | (+) 27 (-) 31 |
| Chromatographic index (*RF*) | (+) 809 (-) 747 |
| Refractive index ($\mu$) | (+) 277 (-) 268 |
| Molecular weight (*M<sub>w</sub>*) | (+) 160 (-) 157 |
| Bulkiness (*Bl*) | (+) 227 (-) 236 |
| Partial specific volume (*V0*) | (+) 325 (-) 348 |
| Molecular volume (*M<sub>v</sub>*) | (+) 187 (-) 166 |

## Discussion

### Evolutionary history of the apoptosome in Metazoa

**The APAF-1/apoptosome arises in Porifera and is present in other basal invertebrates**

Zmasek et al. (2007), proposed that the common ancestor of Cnidaria - Bilateria had a protein similar to human APAF-1. Zmasek and Godzik (2013), proposed that the CARD domain arose at the base of Metazoa, putting *A. queenslandica* as a model species. This is confirmed by our results having the APAF-1 sequence of this poriferous the CARD domain. Similarity and phylogenetic analyses suggest that not only the CARD domain as such, but also the APAF-1 version of vertebrates was already present in Porifera (Figure 1-figure supplement 1 and Supplementary file 2), so this phylum would be the common ancestor at the base of Metazoa, where the APAF-1/apoptosome arose. CASP9 from *A. queenslandica* is also included within vertebrates (Figure 4 and Figure 4-figure supplement 1). This codivergence in the evolutionary history of both proteins would also be evidence that the interaction and therefore the coadaptation between them already existed since the emergence of Metazoa and that as a result of this, they have coevolved until reaching the vertebrates. Similarity analyzes and unreconciled phylogenies of APAF-1 in the phyla Cnidaria, Echinodermata and the subphylum Cephalochordata (Figure 1-figure supplement 1 and Supplementary file 2), coincide with those of other investigations in which genes and proteins have been detected in these phyla similar to APAF-1 of vertebrates, specifically in the cnidarians *N. vectensis* (Zmasek et al. 2007) and *A. acropora* (Moya et al. 2016), in the echinoderm *S. purpuratus* (Robertson et al. 2006; Zmasek et al. 2007) and in the cephalochord *B. floridae* (Zmasek et al. 2007). In the phylogeny generated by Zmasek et al. (2007), some of these species have several copies of the gene, each having exclusive copies of its group and a copy similar to APAF-1 of vertebrates. In *S. purpuratus*, all copies are found in the clade of vertebrates. Based on this; as in the reconciled phylogeny these phyla are in a separate speciation event from that of vertebrates, we suggest that the sequences in these basal invertebrates would be co-orthologous to the vertebrate proteins; apparently in this scenario with all these genes duplicated at the base of Metazoa, the only gene similar to APAF-1 in turn also began to duplicate and in the course of evolution when the gene system was simplified, vertebrates inherited one of those copies.

### Several taxa have apoptosomes other than APAF-1, DARK, and CED-4

In the similarity analyses performed by Menze et al. (2010), the APAF-1 of the crustacean *D. pulex* was close to the human sequence, showing that not all species in the phylum Arthropoda must be similar to DARK of *D. melanogaster*. Our phylogenetic analyses agree with this result, in the sense that the proteins of the genus *Daphnia* are found in a node close to that of vertebrates, but not within the same clade. The origins of the genus *Daphnia* and the entognate *O. cincta* in unique duplication events confirms the existence of apoptosomes different from those formed by APAF-1, DARK and CED-4 in Arthropoda (Figure 1).

A caspase similar to human CASP9 was detected in the mollusc *M. provincialis* (Romero et al. 2011). Our results differ in this aspect, since molluscs have multiple origins distant from those of vertebrates (Figure 4), but we also did not rule out the possibility that in *M. provincialis* there is an apoptosome similar to that of vertebrates, which suggests that within Mollusca there are different versions of the protein complex that are common to Brachiopoda and Chelicerata, so the common ancestor of Lophotrochozoa and Ecdyzosoa probably had all these versions.

### Not all insects have the DARK/apoptosome

There is a paradigm that the version of the DARK/apoptosome in *Drosophila* is common to all insects (Lee y Fairlie, 2012). From our phylogenetic results, we speculate that in the course of the evolution of Insecta, APAF-1/DARK duplicated several times, with the different orders acquiring different versions of the protein (Figure 2), which challenges this paradigm. CASP9/DRONC has a similar divergence, where *D. melanogaster* originates from a single node and is distant from the hymenopter *C. cinctus* (Figure 4), suggesting that the different versions of APAF-1/DARK and CASP9/DRONC within Insecta, by interacting and forming different apoptosomes in the different orders, have coevolved in the course of their evolutionary histories.

### Not all nematodes have the CED-4/apoptosome

The version of the protein complex of *C. elegans* has been generalized to the other nematodes (Oberst *et al*., 2008). Our phylogenetic analyzes suggest that the version of APAF-1/CED-4 in nematodes of the genera *Trichinella* and *Trichuris* is different from that of *C. elegans* (Figure 3), which contradicts the mentioned paradigm. The only caspase present in *C. elegans* is CED-3, homologous to CASP3 (Yuan et al., 1993). As APAF-1/CED-4 from Plathelminthes are possible orthologous to *Trichinella* and *Trichuris*, the detection of CASP9-like proteins in Plathelminthes suggests the presence of this protein, as well as other initiator caspases in other nematodes. In the apoptosome characterized in *C. elegans*, CED-4 lacks the WD40 domains that form the β propellers at the carboxyl terminal (Qi et al., 2010). In the genera *Trichinella*, *Trichuris* and the Plathelminthes, the presence of the WD40 domains (Figure 3-figure supplement 2) suggests that, similar to APAF-1, the protein is regulated by CYCS or another ligand. Bender et al. (2012), demonstrated that in the Plathelminthe *Schmidtea mediterránea* apoptosis is regulated by MOMP, so it may be dependent on CYCS. They suggested that the mitochondrial pathway originated before vertebrates. They assumed that the mitochondrial pathway was lost in some lineages like Nematoda. In contrast to these authors, we suggest that the mitochondrial pathway as well as the WD40 repeats were not lost in most nematodes. Apparently, due to some evolutionary pressure, there was only loss and divergence in the genus *Caenorhabditis*.

Coevolution and destabilizing positive selection analyses were performed on the possible orthologs of APAF-1, CASP9 and CYCS in the species that have the three available proteins (Supplementary file 1), to predict possible aspects of the structure and functional dynamics of the protein complex.

### Structural and functional inferences in the APAF-1/apoptosome, guided by amino acid clusters with coevolution signal

**Regulatory region**

**Allosteric regulation in interaction CYCS and β propellers during APAF-1 activation**

The first step during the activation of APAF-1 is the association of CYCS with the regulatory region, to cause conformational changes in APAF-1 that release the CARD domain. The area of interaction between CYCS and β propeller 8 is greater than with β propeller 7, ~ 1,242 Å^2^ and ~ 712 Å^2^, respectively (Zhou et al., 2015), so CYCS will probably first contact APAF-1 through β propeller 8 (Cheng et al., 2016). One of the interactions identified by Shalaeva et al. (2015), is between CYCS K/54 and β propeller 8 D/1147. In our results, only K/54 was detected in cluster 12, together with CYCS N/55 and β propeller 8 T/1012, located on β-sheet “a” of blade 2 on the back face of β propeller 8 (Figures 5 – A-figure supplement 1 - E). K/54 and N/55 are associated in the CYCS in the area of contact with β propeller 8. After CYCS binding, β propeller 8 rotates slightly upward to approach β propeller 7 (Cheng et al., 2016). This suggests that the interaction of K/54 and N/55 with D/1147, through an allosteric interaction influence T/1012, which causes a conformational change that would initiate this rotation, specifically in the upper part of the protein domain, moving β-sheet “a” of blade 2 forward and up (Figure 6 - A). Therefore, cluster 12 residues would be involved in the initiation of CYCS interaction with APAF-1.

After CYCS binding during APAF-1 activation, α29-32 helices in HD2 rotate slightly upward along with β propeller 8 (Cheng et al., 2016). Residues V/576 and A/584 in cluster 18 are located in the α-helix 32 (Figure 5 – A-figure supplement 1 B and C), suggesting that these are part of the specific amino acids involved in conformational changes at the base of the protein domain during rotation (Figure 6 - B). Residues in HD2 L/593, L/595, W/597 and I/603 follow a hydrophobic corridor on the surface of β propeller 8 (Zhou et al., 2015). Two of these residues, L/595 and W/597 are located in cluster 18. These are in linker 3 (Figure 5 – A), which suggests that the linker is also involved in the rotation (Figure 6 - B).

The amino acids in clusters 10 and 11 are unique to β propeller 8 (Figure 5 – A-figure supplement 1 B and C). The distance between the residues in the structure suggests that they are involved in allosteric changes that cause alterations in β-sheet “d” of blade 4, β-sheets “a” and “d” of blade 5, β-sheet “c” of blade 7 and β-sheet “b”of blade 8, finalizing the conformational changes that rotate upwards to β propeller 8 (Figure 6 - C). Clusters 10 and 11 are independent of CYCS, suggesting that conformational changes in β propeller 8 are also caused by other ligands. The participation of residues V/1235 and L/1241 of cluster 10 in the rotation would initiate the formation of the V shaped cleft at the base.

The next step during APAF-1 activation is the contact of CYCS with β propeller 7. Cheng et al. (2016), identified CYCS K/28 as an important residue for the interaction with the regulatory region of APAF-1. Shalaeva et al. (2015), determined that this residue interacts with β propeller 8 E/925. In our results K/28 was located in cluster 21 along with other residues in β propeller 7 (Figure 5 – A-figure supplement 1 - E), which makes more sense since K/28 is on the surface of the CYCS in contact with this domain. This suggests that CYCS contacts β propeller 7 through these residues. After contact, β propeller 7 spins and rotates to “hug” CYCS in the V shaped cleft (Cheng et al., 2016), which suggests that K/28 through allosteric regulation interacts with F/787, V/799 and V/801, causing alterations in the β-sheet “a” of blade 4, which would initiate the rotation and spin of this protein domain and the formation in the upper region of the V shaped cleft in which the CYCS is coupled (Figure 6 - D).

Now the CYCS is in interaction with both β propellers. Other interactions in the contact area encompassing CYCS and β propellers include: CYCS M/13 and β propeller 7 E/700, CYCS Q/43 and β propeller 8 E/1045, CYCS G/57 and β propeller 8 T/1087, CYCS P/77 and β propeller 8 W/1179 (Zhou et al., 2015), CYCS K/56 and β propeller 8 E/1045 (Shalaeva et al., 2015). The residue CYCS K/56 was also identified by Cheng et al. (2016) as important in the interaction with the APAF-1 regulatory region. Our results reveal that the majority of CYCS residues involved in these interactions are connected through cluster 2 with more residues away from the contact area between both proteins, encompassing areas around the heme group in CYCS (Figure 5 – A-figure supplement 1 – G) and the central and posterior area of the β propellers (Figures 5 – A-figure supplement 1 - E). This suggests that the heme and/or residues within cluster 2 located in the contact zone, through a triple allosteric regulation, influence the other amino acids within the cluster located far from the contact zone, affecting three domains, causing conformational changes in β-sheet “b” of blade 5 and β-sheets “b” and “c” of blade 6 in β propeller 7, β-sheets “b” and “c” of blade 2 and β-sheets “a” and “b” of blade 3 in β propeller 8 and in the upper region of CYCS (Figure 6-E). In APAF-1 the residues in the β propellers far from the contact zone are located longitudinally around the V shaped cleft (Figure 6 - E), which suggests that the conformational changes in these amino acids continue with the formation of the cleft longitudinally, assisting linker 4, causing the CYCS to begin to fit into the cleft.

Clusters 14-17 and 23 are unique of β propeller 7 (Figure 5 – A-figure supplement 1 – B, D and E). Clusters 16, 17, and 23 would come into play first. During conformational changes in the regulatory region, β propeller 7 rotates ~ 60° towards β propeller 8 (Zhou et al., 2015), “hugging” CYCS making it fit into the V shaped cleft (Cheng et al., 2016), which suggests that this turn is caused by allosterism in amino acids in clusters 16, 17 and 23 causing alterations in β-sheet “d” of blade 2, β-sheet “b” of blade 3 and β-sheets “a", “b” and “c” of blade 4, also contributing to the formation of the V shaped cleft in the upper region (Figure 6 - F).

Clusters 14 and 15 would then come into play, located at the base of the posterior face just below the V shaped cleft (Figure 5 – A-figure supplement 1 – B, D and E). These would cause conformational changes in β-sheets “c” and “d” of blade 6 and β-sheets “a”, “b” and “I” of blade 7, assisting linkers 3 and 4, finishing the formation of the V shaped cleft at the base, making the CYCS fully fit into the cleft (Figure 6 - G). Clusters 14-17 and 23 are also independent of CYCS, so the existence of other ligands that contribute to conformational changes in β propeller 7 is possible. In inactive APAF-1, NBD K/192 interacts with β propeller 7 D/616 through a salt bridge, keeping CARD hidden (Reubold et al., 2011). During protein activation, when β propeller 7 spins, it disconnects this salt bridge, releasing CARD (Reubold et al., 2011). The residues in clusters 14 and 15 are close to K/192 and D/616, so they could contribute to breaking this salt bridge. See animated morph with sequence of events described so far in Video 1.

### Arm and platform

Cluster 19 is unique to the HD2 domain, encompassing residues in α27 – α29 helices (Figure 5 –A-figure supplement 1 - B). They are probably important to confer stability to the domain during conformational changes.

There are a series of interactions between domains that help maintain APAF-1 in its extended or active conformation, among these WHD-HD2 (Zhou et al., 2015). WHD associates with HD2 through Van der Waals interactions, where residues WHD I/388, V/393, V/399 and L/403 rest on the hydrophobic surface formed by HD2 H/457, I/460, H/490, Y/482, W/483, F/486 and Y/489 (Zhou et al., 2015). Cluster 20 has in common several of the residues in HD2 involved in these interactions, I/460 and I/461 in α25, Y/482 and H/490 in the α26. Other amino acids in the cluster are L/487 and M/491 in the α26 helix (Figure 5 – A-figure supplement 1 - B).

The α8 linker helix in NBD associates with a NBD of an adjacent APAF-1 during the oligomerization process (Yuan et al., 2010). In these interactions NBD Q/121 donates a hydrogen bridge to Q/257 of the adjacent NBD (Zhou et al., 2015). E/116 and R/122 (next to Q/121) located in the α8 helix of the NBD belong to cluster 9 (Figure 5). Hence we presumed that cluster 9 participates in the oligomerization process.

### Specific residues in NBD and APAF-1 CARD open hydrophobic pocket for exchange and stabilization of nucleotides

At the center of the interface in the interactions between WHD-NBD-HD1, there is an access channel or hydrophobic pocket that stores and keeps ADP stable in the inactive state of APAF-1 (Riedl et al., 2005; Cheng et al., 2016). This is blocked by CARD, but opens when this domain is removed, acquiring its extended conformation, allowing exchange for ATP (Riedl et al., 2005). Two of the amino acids that give structural support stabilizing ADP/ATP are P/123 and R/129 in NBD (Riedl et al., 2005; Cheng et al., 2016). These two residues belong to cluster 7 along with others located in α4, α6 helices at the carboxyl terminal of the CARD domain; in α8, α9 helices and in β1 sheet at the amino terminal of the NBD (Figure 5). Unlike P/123 and R/129, the other amino acids in the cluster probably participate in the conformational changes necessary for the ADP/ATP exchange. Within these changes, the cluster residues located at the carboxyl terminal of the CARD would be essential for blocking and opening the hydrophobic pocket where the nucleotide is found. (Figure 7 and Video 2).

**Figure 7.**
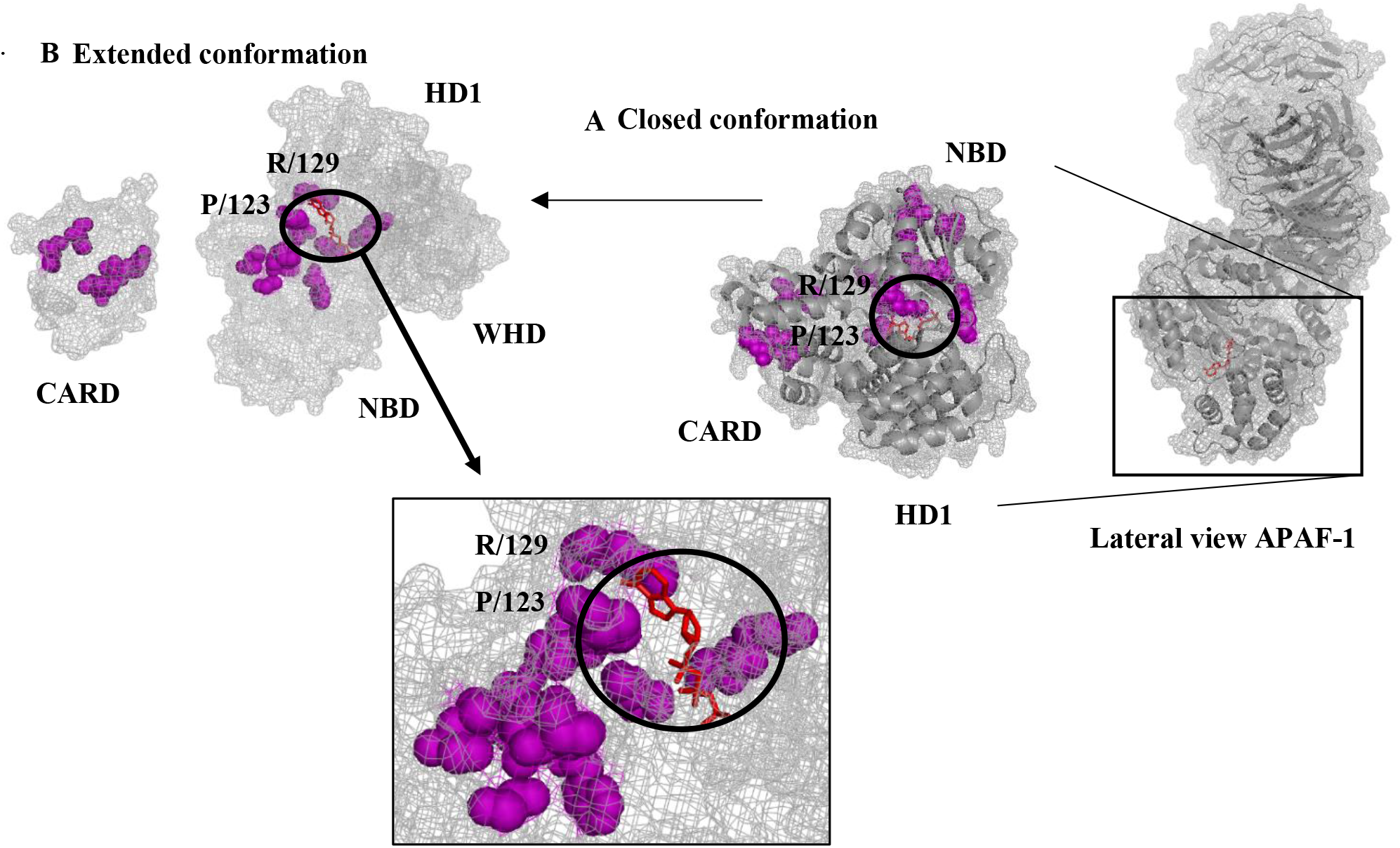
Role of cluster 7 in hydrophobic pocket aperture for nucleotide exchange. Side view of APAF-1 in inactive state PDB 3SFZ. **A)** Close-up of the closed hydrophobic pocket that stores ADP (red sticks circled) in structure PDB 1Z6T, showing the residues in cluster 7 located between CARD and NBD. **B)** The residues P/123 and R/129 remain below and above the adenine, giving it stability. The other residues of the cluster would participate in conformational changes that remove CARD, remaining in its extended conformation, opening the hydrophobic pocket (augmented region and indicated with an arrow) for nucleotide exchange. After the removal of the CARD, the open hydrophobic pocket can be seen, the nucleotide being most visible. Structure PDB 5JUY. See also Video 2.

### Disk

**Large subunit of CASP9 also contributes to homodimerization**

When catalytic domains homodimerize, they do it by their carboxyl terminals through motif 402 GCFNF 407 in the small subunit (Wu et al., 2016). Renatus et al., (2001) identified in the large subunit a loop of 7 residues inserted at position 240 between the beta 3 and 3A sheets that interact through the interface of the dimer, therefore both the small and large subunits participate in the homodimerization process. Our results corroborate this statement, since of these 7 residues, H/243, F/246 and P/247 that belong to cluster 5 (Figure 5 - A-figure supplement 1 - F), in the dimer structure they are in direct contact (Figure 8).

**Figure 8.**
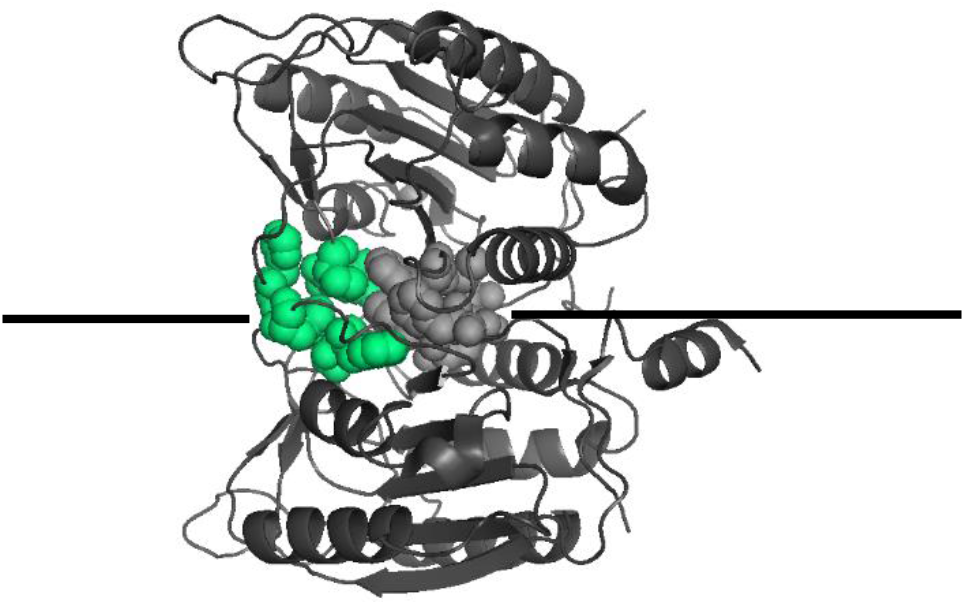
Residues of cluster 5 participate in CASP9 homodimerization. Catalytic domains p20/p10 are shown as dimer (delimited by horizontal black line) PDB 1JXQ. Amino acids of Cluster 5 were mapped as lime green spheres located in the large subunit. The residues as gray spheres belong to motif 402 GCFNF 407 (not detected in coevolution analysis) in the small subunit. In the dimer, all these residues are in direct contact, so the large subunit also contributes to homodimerization.

### Conglomerate residues throughout the catalytic domain would transmit the signal to the active site C/287 of CASP9 and would participate in inhibition mediated by Zinc

In the process of activation of CASP9, cleavages are caused in E/306, D/315 and D/330 that expose the active site C/287 (Bratton et al., 2001). In our results, residue C/287 located in β-sheet 4 belongs to cluster 6 together with another 25 amino acids in the large subunit, sheets β1, β2 of 240-loop, β3, β4 and helix α3 ( Figure 5-A). Renatus et al. (2001) propose that conformational changes during homodimerization transmit the activation signal to the residues surrounding the active site C/287. In the three-dimensional structure, cluster 6 residues are in direct contact throughout the large subunit (Figure 5-figure supplement 1 - F), and may be those that participate in the conformational changes that transmit this signal. The catalytic domain is also involved in the inhibition of CASP9 mediated by Zinc, having two binding regions to this inhibitor, one of them encompassing the active site C/287 and two residues that in the three-dimensional structure are around it, H/237 and C/239 (Huber and Hardy, 2012). Of these residues, apart from the active site, H/237 belongs to cluster 6 (Figure 5 - A-figure supplement 1 - F), so we can infer that the residues in cluster 6 are involved in both the activation and inhibition mediated by Zinc of CASP9.

### The molecular timer

Wu and Bratton (2017) state that as part of the molecular timer, the cleavage in D/315 during the activation of CASP9 destabilizes the dimers causing them to dissociate from the complex, losing catalytic activity. If the protein complex requires more functioning time, CASP3 avoids the destabilization of the dimers through a positive feedback, carrying out a cleavage at D/330, removing the linker from the intersubunit, which gives more stability to the dimers‥ In our results, D/330 belongs to cluster 3 along with other residues located in linker 2 and β-sheet 5 (Figure 5 – A-figure supplement 1 - F). Apparently they are responsible for giving stability to CASP9 after cleavage at D/330 by CASP3.

Another of the regulatory mechanisms in these processes to remove caspases from the apoptosome is through a caspase-mediated cleavage in APAF-1 that removes H1 from the CARD domain, turning it into an 84 or 130 kDa fragment, not being able to interact more with caspases (Lauber et al., 2001). Possibly the cleavage occurs in motif 16 LEKD 19 at the amino terminal of CARD, and other amino acids at positions 37-52 and 12-28 may also be involved in the generation of fragments p84 and p130, respectively (Lauber et al., 2001). Several residues inside this range fell within cluster 13: L/16 of motif 16 LEKD 19, along with other residues in α1, α2 and α3 helices (Figure 5), giving a more precise mapping of the specific residues that could be involved in the cleavage process. This suggests that residues in cluster 13 would have an inhibitory function, blocking the functioning of the apoptosome as part of the molecular timer.

### As an alternate mechanism, the catalytic domain of CASP9 could interact with CYCS in the regulatory region to expose active site C/287

Gasperin - Sánchez et al. (2020), made molecular dynamics simulations in order to describe conformational changes in pc9 during its activation, where in all these the active site C/287 was always hidden, so the authors propose a different model than the holoenzyme, in which the catalytic domains interact with other proteins of the apoptosome, causing the exposure of the active site, and then homodimerize. In this model, in one of the conformations, the catalytic domains p20/p10 through linker 1 remain in contact with the regulatory region. In our results, cluster 22 encompasses residues in β sheets 1 and 2 and helix α1 in the catalytic domain of CASP9 and CYCS A/44 (Figure 5 - A-figure supplement 1 – F and G), suggesting that this possible interaction between CASP9 and the regulatory region, specifically it would be between the catalytic domain of CASP9 and CYCS. In this hypothetical scenario, after pc9 anchors itself to the apoptosome in the disc, as an alternate mechanism, p20/p10 interacts through linker 1 with CYCS in the regulatory region through these residues, exposing the active site C/287, so that later the catalytic domains can homodimerize (Figure 9).

**Figure 9.**
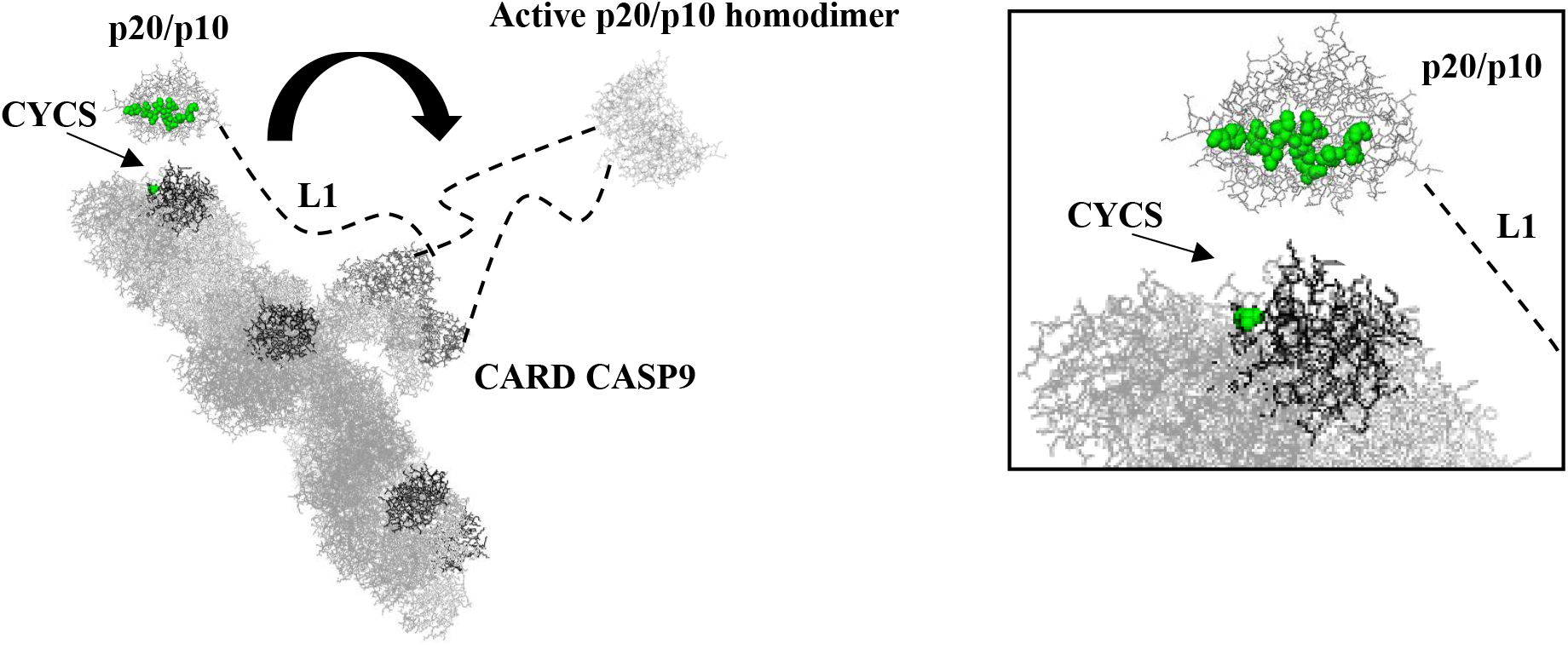
Exposure of the active site C/287 in pc9 caused by interaction with CYCS in regulatory region. Once pc9 is anchored to the disk of the apoptosome (PDB 5JUY), the p20/p10 domains (PDB 1JXQ) interact through L1 with CYCS in the regulatory region, to expose the active site C/287, and then homodimerize.

### Apoptosome-independent apoptotic functional inferences

**CASP9 catalytic domain could be myristoylated to interact with mitochondrial outer membrane**

CASP9 also interacts with the outer membrane of the mitochondria, producing cleavages in the anti-apoptotic BCL-2 proteins, amplifying the release of more pro-apoptotic proteins (Guerrero et al., 2012). Cluster 4 of CASP9 has two glycine residues, G/269 in the α3 helix and G/288 in the β4 sheet of the large subunit, located at each end of the catalytic domain (Figure 5 – A-figure supplement 1 - F). To complement the results of Guerrero et al., 2012, what we propose is that the interaction of CASP9 with the outer membrane of the mitochondrion is carried out through these two residues. In this hypothetical scenario, some caspase or other protease would produce a cleavage in CASP9, exposing the two glycine residues, which are myristoylated, then interacting with the membrane.

### pc9 could dimerize inside mitochondria

Inside the mitochondria, the amino terminal of pc9 has been reported to enter the matrix (Milisav and Šuput, 2007). Cluster 8 residues exclusive of pc9/CASP9 are hydrophobic, located within the protein domain (Figure 5-figure supplement 1 – B and E), so they may serve as an anchor point for pc9 to associate with the inner membrane of mitochondria. Once anchored, it is possible that it dimerizes with another anchored pc9 to become activated, with the possibility that it interacts with other substrates including CASP3.

We have as a caveat the following statements. Amino acids with a coevolution signal do not necessarily have to be involved in structural or functional interactions. Other restrictions by which they can be detected are: negative design, codon effects and errors in phylogenetic analyzes (Anishchenko et al., 2017). Another aspect to take into account of these detected residues is that the interaction may not be in that particular protein, but that they could reflect interactions in other members of the protein family. For example, amino acids that coevolve in distant positions in a protein, is because their homologous residues can be in direct contact in other proteins of the family (Anishchenko et al., 2017). Therefore, this restriction is due to amino acids having a common evolutionary history (Chakrabarti and Panchenko, 2009). Another reason why the residues may be distant is due to errors in the determined structure of the protein, which may suggest that these distant amino acids are actually in direct interaction (Anishchenko et al., 2017), which in our specific case could be reflected in some regions of the APAF-1/apoptosome that have been structurally characterized at high resolutions, not being of good quality. For these reasons, certain caution must be taken when taking as valid the structural and functional inferences of the protein complex guided by amino acids with coevolution signals, which are only models. Even so, these potential interactions detected will serve as a guide to be validated experimentally.

### Influence of destabilizing changes in structure and function of the APAF-1/apoptosome

In general, molecular adaptation does not work on the specific amino acids necessary for a catalytic reaction or conformational change. On the contrary, adaptation results from changes in those amino acids that surround active/functional residues to represent a slight change in the way in which these sites perform their function (Baer, 2007), so we presume that the substitutions in amino acids with coevolution signal are more conservative because they are in active sites and involved in conformational changes, so that the functionality of the protein is maintained. On the other hand, the substitutions in amino acids located in the periphery are destabilizing, since their function is not so essential, being more apt to undergo radical changes. In some cases, residues in the interior near active sites may require radical adaptations, these changes being guided in a subtle way by residues in the periphery.

Compressibility is characteristic of globular proteins (Gromiha and Ponnuswamy, 1993), which in turn is characteristic of enzymatic proteins with regulatory function. For this reason, it makes sense that the regions most affected by this property in the apoptosome are globular regions, encompassing in APAF-1 the β propellers that form the regulatory region, CYCS and CASP9 throughout their length, whose function is enzymatic. Compressibility is related to volume, partial specific volume, molecular volume, and molecular weight. These physicochemical properties influence the amino acid packing density of proteins (Karshikoff and Ladenstein, 1998 and Fischer et al., 2004).

The helical contact area and the hydrophobicity by thermodynamic transfer favor the sequestration of residues into the protein (Harris and Pettitte, 2016) and the folding of globular proteins (Dill, 1990). These have served to guide the folding, conglomeration and associations of the hydrophobic amino acids within the affected regions.

Proteins affected by refractive index and chromatographic index possess stability in external fields, which is required during conformational changes and interactions between enzymes (Pace et al., 1990; Vörös, 2004; Hong et al., 2012), such as processes of oligomerization (Folta-Stogniew, 2006). It is relevant to note that one of the regions most affected by these two properties was the platform of the protein complex, which is essential for the oligomerization process. Chromatographic index indicates the most affected residues (Table 1). This makes sense since the apoptosome is very dynamic, having many conformational changes and interactions. The stability given by these properties may be required in the regulatory region for association with CYCS and the conformational changes caused by this interaction during APAF-1 activation; In NOD, during nucleotide exchange and oligomerization during the heptamer formation; on HD2, during the spin to extend the CARD domain of APAF-1; in the CARDs domains of both proteins, during the CARD-CARD interaction for the recruitment of CASP9 in the disk; and in the catalytic domain of pc9 during the cleavage and activation processes of this protein.

Apparently, the globular nature and the dynamic complexity of the apoptosome were the evolutionary constraints on natural selection to favor these significant physicochemical properties that caused destabilizing changes in the protein complex. These drastic changes were essential to keep the apoptosome functioning throughout its evolution.

## Materials and methods

### Obtention of the genes, messenger RNAs and proteins of the human apoptosome

*H. sapiens* sequences were obtained from GenBank (Benson et al., 1993; Sayers et al., 2019). The gene codes are 317 for APAF-1 and 842 for CASP9. For each of these sequences in the section “gene table” there are several isoforms of transcripts. The canonical isoform and its translated protein were selected for each. Here the codes for the messenger RNAs and proteins, respectively, APAF-1: NM_181861.1 and NP_863651.1, CASP9: NM_001229.4 and NP_001220.2.

### Identification of proteins homologous to the human apoptosome in invertebrates

The sequences were obtained in two ways on 06/25/2017. First, a general search was made in GenBank using as reference the names of the human sequences, selecting the genes within invertebrates. To avoid the exclusion of sequences that could be selected, genes were searched using all synonymous names. The longest protein and its transcript were selected from each gene. Second, within the GenBank reference sequence database (RefSeq), a BLASTP search was run on human protein sequences against the more general invertebrate taxa: Porifera, Coelenterata, Protostomes, and Deuterostomes (excluding vertebrates). A single iteration per taxon was made, selecting the sequences whose scores were greater than 200. For each one its transcript was obtained. As a result, sequences of APAF-1 and CASP9 were obtained in 92 species of invertebrates (Supplementary file 1). Not all species have the two proteins in common. Of these, five sequences were not under the name CED-4 but APAF-1 in the nematodes *Trichinella nativa*, *T. papuae*, *T. pseudospiralis*, *T. zimbabwensis* and *Trichuris suis*, while 59 were not under the name DARK but APAF-1 in six orders of insects, Isoptera, Hemiptera, Coleoptera, Diptera, Lepidoptera and Hymenoptera.

### Identification by similarity of orthologs to the human apoptosome in Metazoa

The Ortho-Profile method (Szklarczyk et al., 2012) was applied to the species within invertebrates in which homologs were detected, designed to detect orthologous genes and proteins in highly divergent sequences. This consists of three phases with an increase in sensitivity in each of them. If the possible ortholog sought is not detected in the first phase, it goes to the second with more sensitivity, and if it is not detected in this, it goes to the third, which has even more sensitivity. In each of the phases, for the similarity to be significant, both sequences must present reciprocity and have an e-value of less than 0.01. In the first phase, a single iteration of reciprocal BLASTP to the human proteins against the selected species was executed, one by one, sequence against sequence, in the RefSeq database. In the second phase, a PSI-BLAST (Altschul et al., 1997) was also applied reciprocally, in the GenBank non-redundant reference database (RefSeq nr), against the remaining species, one by one, profile versus sequence with a profile inclusion threshold equal to 0.005. Up to three iterations were done to see if it detected significant similarities. In the third and final phase, the profile-based HMM was used. On the HMMER online server (http://www.ebi.ac.uk/Tools/hmmer/), the algorithm was run against the human sequences against the reference proteome database and against the remaining species one by one.

In the complete GenBank report for each human apoptosome gene, in the general information tab, the putative orthologous genes of 115 species of vertebrates were obtained through the annotation pipeline (Thibaud-Nissen et al., 2016). For each gene, the protein with the longest length was selected with its respective transcript (Supplementary file 1).

### Identification by phylogeny of orthologs to the human apoptosome in Metazoa

For the phylogenies, apart from the human sequences, the DARK and DRONC proteins from *D. melanogaster* and CED-4 from *C. elegans* were also included, to be used as a reference and to determine in relation to the apoptosomes formed by APAF-1, DARK and CED-4, where the others are located. In GenBank, for *D. melanogaster* and *C. elegans*, the canonical protein sequences, DARK: NP_725637.1, DRONC: NP_524017.1 and CED-4: NP_001021202.1, were selected. To obtain the architecture of the protein domains of the reference sequences and other invertebrates and vertebrates, analyzes of conserved domains were made in Pfam 32.0 (https://pfam.xfam.org/) (Sonnhammer et al., 1997; El-Gebali et al., 2019). Each protein family was aligned separately in MAFFT v7.380 (Katoh et al., 2002; Katoh and Standley, 2013), under the “–auto” mode, which selected the strategies “FFT-NS-i” and the method of iterative refinement, doing two iterations for the APAF-1 alignment, and “L-INS-i” and the iterative refinement method, doing an iteration with information of local pairs, for the CASP9 alignment. The produced alignments were inspected and edited in AliView 1.20 (Larsson, 2014), subsequently removing sequences with amino acids of unknown nature. To increase the quality of the alignments, the poorly aligned regions were removed using trimAL 1.2rev57 (Capella-Gutiérrez et al., 2009). Once edited, they were run separately in phylip format in ProtTest 3.4.2 (Abascal et al., 2005; Darriba et al., 2011), determining the following best models of amino acid substitutions during evolution, “JTT + G + F “, assigning to the gamma distribution the value of 1.639 for the APAF-1 and “JTT + I + G “, assigning to the gamma distribution and proportion of invariant sites the values of 1.186 and 0.028 for the CASP9. Based on these models, the phylogenetic reconstructions were made in phyML 3.1 (Guindon and Gascuel, 2003; Guindon et al., 2010), under the principle of maximum likelihood (Felsenstein, 1981), using the non-parametric method Shimodaira - Hasegawa (Shimodaira and Hasegawa, 1999), generating 3 phylogenetic trees corresponding to each protein family. These were rooted in Seaview 4 (Gouy et al., 2009) and visualized in FigTree 1.4.3 (Rambaut, 2017) (https://tree.bio.ed.ac.uk/software/figtree).

Species-level phylogenies were generated automatically in the NCBI taxonomy database (Federhen, 2012). A species tree was generated for each protein phylogeny. For the reconciliation of protein and species phylogenetic trees, Notung Version 2.9 (Chen et al., 2000) was used. All of our protein phylogenies are binary, but our species trees are polytomous, which is why they were reconciled in the program under the parsimony algorithm of gene duplication, transfer, loss, and incomplete lineage sorting (Stolzer et al., 2012). The reconciliations were visualized in Archeopteryx version 0.9921 beta (Zmasek, 2015).

### Obtention of the CYCS sequences

Once proteins orthologs to the human sequences were detected and selected, for each species with the two orthologs available, the CYCS sequences were obtained, completing the proteins that make up the apoptosome. Using as reference the protein sequence of the human CYCS (Protein ID: NP_061820.1), putative orthologous proteins were obtained in vertebrates through the annotation pipeline (Thibaud-Nissen et al., 2016). For invertebrates, a BLASTP search was applied to the human sequence against the selected species (Supplementary file 1). CYCS sequences were aligned in MAFFT v7.380 (Katoh et al., 2002; Katoh and Standley, 2013), under the “–auto” mode, which selected the strategy “L-INS-i” and the iterative refinement method, iterating with local pair alignment information.

### Protein coevolution analysis

The complete alignments of orthologous sequences (not trimmed with trimAL) were concatenated in Mesquite Version 3.31 (Maddison and Maddison, 2017) (http://www.mesquiteproject.org), again determining the best amino acid substitution model and a phylogeny, using ProtTest 3.4.2 (Abascal et al., 2005; Darriba et al., 2011) and phyML 3.1 (Guindon and Gascuel, 2003; Guindon et al., 2010), respectively. In Seaview 4 (Gouy et al., 2009) the phylogeny generated was rooted in the node that separates invertebrates from vertebrates.

For the coevolution analysis, the alignment and its respective phylogeny were supplied to the BIS2Analyzer server (http://www.lcqb.upmc.fr/BIS2Analyzer/) (Oteri et al., 2017). The API was calculated for our concatenated alignment in Geneious 11.1.3 (Kearse et al., 2012), obtaining a score of 67.3%, this having a moderate divergence. Therefore, once uploaded to the server, the option “Alphabet-Reduction” was selected, with a maximum dimension (D) of 2. The results were mapped into the concatenated linear sequences and three-dimensional structures available in the database of proteins (PDB) using Jalview 2.10.4 (Clamp et al., 2004; Waterhouse et al., 2009) and PyMOL (DeLano, 2014) (https://pymol.org/2/), respectively. Clusters were also mapped on morphs (transformation from inactive to active protein state) in UCSF Chimera 1.13.1 rc (Pettersen et al., 2004).

### Positive selection analysis

In GenBank, from the messenger RNA isoforms of the selected orthologs, only the coding regions were extracted, then aligned in TranslatorX Version 1.1 (Abascal et al., 2010), under the MAFFT algorithm, using as reference the same alignments of the proteins generated in the coevolution part. The alignments were concatenated in Mesquite Version 3.31 (Maddison and Maddison, 2017) (http://www.mesquiteproject.org), then executed in phylip format in jModelTest 2.1.10 (Posada, 2008; Darriba et al., 2012), determining the best model for nucleotide substitution during evolution to “TIM3 + G”, assigning the gamma distribution a value of 0.7160. Based on this model, the phylogenetic reconstruction was done in phyML 3.1 (Guindon and Gascuel, 2003; Guindon et al., 2010). In Seaview 4 (Gouy et al., 2010) the phylogeny generated was rooted in the node that separates invertebrates from vertebrates.

To detect positive selection, TreeSAAP Version 3.2 (Woolley et al., 2003) was used. The input file with the alignment of the coding DNA in nexus format and the phylogeny in newick format were supplied to the program. A first run was carried out under the predetermined parameters, taking into account the 31 physicochemical properties of the amino acids available in the program, identifying which are the most significant. Using these preliminary data as a reference, another run was made, this time taking into account only the significant physicochemical properties, giving the sliding windows an average length of 20. The definitive results, focusing on the residues under destabilizing positive selection (categories 6 - 8 and z scores: −3.09 and +3.09), were visualized in IMPACT_S (Maldonado et al., 2014), then mapping the residues in three-dimensional structures in PyMOL (DeLano, 2014) (https://pymol.org/2/).

## Supporting information

Supplementary file 1-Species selected for analysis

Supplementary file 2-Ortho-Profile results for APAF-1 and CASP9

Figure 1-figure supplement 1-Non reconciliated APAF-1 phylogeny showing proteins nearby and within the human clade

Figure 2-figure supplement 1-Non reconciliated APAF-1 phylogeny showing the relationship between insect sequences

Figure 3-figure supplement 1-Non reconciliated APAF-1 phylogeny showing the relationship between nematode sequences

Figure 3-figure supplement 2-Analysis of conserved domains obtained in the NCBI of homologues to APAF-1CED-4 in Nematoda and Plathelminthes

Figure 4-figure supplement 1-Non reconciliated CASP9 phylogeny

Suplementary file 3-Output file BIS2Analyzer with alignment and hits associated to dimensions and blocks

Supplementary file 4-Output file BIS2Analyzer in table form with hits associated to dimensions and blocks

Figure 5-table supplement 1-Clusters of amino acids with coevolution signal

Figure 5-figure supplement 1-Clusters of aminoacids with coevolution signal mapped in tridimensional structures

Figure 6-figure supplement 1-TreeSAAP sliding windows aminoacids under destabilizing positive selection

Supplementary file 5-TreeSAAP amino acids under non synonymous substitutions

Video 1-Model of APAF-1 activation based on clusters of amino acids with coevolution signal

Video 2-Role cluster 7 in NBD-CARD nucleotide exchange

## Acknowledgements

We thank Ruth Bastardo, Altagracia Espinosa, Santo Navarro, Candy Ramírez and América Sánchez of the Institute of Botanical and Zoological Research Prof. Rafael M. Moscoso (IIBZ) of the Autonomous University of Santo Domingo, for their guidance on how to organize voluminous data products of this research. José A. Sánchez - Borbón thanks Dr. Yamixa Delgado - Reyes from the San Juan Bautista School of Medicine, Caguas, Puerto Rico, for teaching the basics of the PyMOL program; Willy Maurer from the Specialized Institute for Higuer Education - Loyola, San Cristobal, Dominican Republic, and Fritz Pichardo - Marcano from the IIBZ, for recommending and providing technical support with the use of some of the programs to run the analyzes.

## Authors’ contributions

José A. Sánchez - Borbón, conceptualization, design of the methodology, execution of the analyzes, interpretation of the results, writing of the original draft; Steven E. Massey, conceptualization, methodology design, supervision, original draft writing; José D. Hernández - Martich, supervision, writing of the original draft.

