## Supplementary file 1-Species selected for analysis for "Protein coevolution and physicochemical adaptation in the APAF-1/apoptosome: structural and functional implications"

**Supplementary file 1. Species selected for analysis.** Species GenBank IDs of the genes, messenger RNAs, and proteins are shown, indicating whether they are putative orthologs to the human sequences and whether they qualified for coevolution and positive selection analyzes. The check indicates “yes” and X “no”.

| **#** | **Genus** | **Species** | **Class** | **Gen** | **Gen ID** | **ID transcript/mRNA variant** | **ID protein isoform** | **Putative ortholog** | **coevolution/positive** |
| --- | --- | --- | --- | --- | --- | --- | --- | --- | --- |
|  |  |  |  |  |  |  |  |  | **Selection analyzes** |
| **Species used as reference** | | |  |  |  |  |  |  |  |
| 1 | *Caenorhabditis* | *elegans* | Chromadorea | CED4 | [175643](https://www.ncbi.nlm.nih.gov/sites/entrez?db=gene&cmd=Retrieve&dopt=full_report&list_uids=175643) | [-](https://www.ncbi.nlm.nih.gov/nuccore/NM_001026031.2) | [NP 001021202.1](https://www.ncbi.nlm.nih.gov/nuccore/NP_001021202.1) | X | X |
| 2 | *Drosophila* | *melanogaster* | Insecta | DARK | [36914](https://www.ncbi.nlm.nih.gov/sites/entrez?db=gene&cmd=Retrieve&dopt=full_report&list_uids=36914) | [-](https://www.ncbi.nlm.nih.gov/nuccore/NM_166207.3) | [NP 725637.1](https://www.ncbi.nlm.nih.gov/nuccore/NP_725637.1) | X | X |
|  |  |  |  | DRONC | [39173](https://www.ncbi.nlm.nih.gov/sites/entrez?db=gene&cmd=Retrieve&dopt=full_report&list_uids=39173) | [-](https://www.ncbi.nlm.nih.gov/nuccore/NM_079293.4) | [NP 524017.1](https://www.ncbi.nlm.nih.gov/nuccore/NP_524017.1) | X | X |
|  |  |  |  | APAF-1 | [317](https://www.ncbi.nlm.nih.gov/sites/entrez?db=gene&cmd=Retrieve&dopt=full_report&list_uids=317) | [NM 181861.1](https://www.ncbi.nlm.nih.gov/nuccore/NM_181861.1) | [NP 863651.1](https://www.ncbi.nlm.nih.gov/protein/32483359) | ✓ | ✓ |
| 3 | *Homo* | *sapiens* | Mammalia | CASP9 | [842](https://www.ncbi.nlm.nih.gov/sites/entrez?db=gene&cmd=Retrieve&dopt=full_report&list_uids=842) | [NM 001229.4](https://www.ncbi.nlm.nih.gov/nuccore/NM_001229.4) | [NP 001220.2](https://www.ncbi.nlm.nih.gov/protein/14790124) | ✓ | ✓ |
|  |  |  |  | CYCS | [54205](https://www.ncbi.nlm.nih.gov/sites/entrez?db=gene&cmd=Retrieve&dopt=full_report&list_uids=54205) | [NM 018947.5](https://www.ncbi.nlm.nih.gov/nuccore/NM_018947.5) | [NP 061820.1](https://www.ncbi.nlm.nih.gov/protein/11128019) | ✓ | ✓ |
| **Porifera** | |  |  |  |  |  |  |  |  |
|  |  |  |  | APAF-1 | [105313814](https://www.ncbi.nlm.nih.gov/gene/105313814) | [XM 020000155.1](https://www.ncbi.nlm.nih.gov/nuccore/XM_020000155.1) | [XP 019855714.1](https://www.ncbi.nlm.nih.gov/protein/1133471758?report=genbank&log$=prottop&blast_rank=1&RID=P9MNVGTG014) | ✓ | ✓ |
| 4 | *Amphimedon* | *queenslandica* | Demospongiae | CASP9 | [109585977](https://www.ncbi.nlm.nih.gov/gene/109585977) | [XM 020002120.1](https://www.ncbi.nlm.nih.gov/nuccore/XM_020002120.1) | [XP 019857679.1](https://www.ncbi.nlm.nih.gov/sites/entrez?cmd=Search&db=protein&term=XP_019857679.1&dopt=GenBank) | ✓ | ✓ |
|  |  |  |  | CYCS | [100636040](https://www.ncbi.nlm.nih.gov/gene/100636040) | [XM 003388889.3](https://www.ncbi.nlm.nih.gov/nuccore/XM_003388889.3) | [XP 003388937.1](https://www.ncbi.nlm.nih.gov/sites/entrez?cmd=Search&db=protein&term=XP_003388937.1&dopt=GenBank) | ✓ | ✓ |
| **Cnidaria** | |  |  |  |  |  |  |  |  |
| 5 | *Acropora* | *digitifera* | Anthozoa | APAF-1 | [107354394](https://www.ncbi.nlm.nih.gov/gene/107354394) | - | [XP 015776356.1](https://www.ncbi.nlm.nih.gov/protein/1005479259?report=genbank&log$=prottop&blast_rank=7&RID=P9NE5SJ4015) | ✓ | X |
| 6 | *Acropora* | *millepora* | Anthozoa | APAF-1 | n/a | - | [AJG37574.1](https://www.ncbi.nlm.nih.gov/protein/752445179?report=genbank&log$=prottop&blast_rank=1&RID=P9NE5SJ4015) | ✓ | X |
| 7 | *Hydra* | *vulgaris* | Hydrozoa | APAF-1 | [100210403](https://www.ncbi.nlm.nih.gov/gene/100210403) | - | [XP 012566478.1](https://www.ncbi.nlm.nih.gov/protein/XP_012566478.1) | ✓ | X |
| 8 | *Nematostella* | *vectensis* | Anthozoa | APAF-1 | [5519825](https://www.ncbi.nlm.nih.gov/gene/5519825) | - | [XP 001639696.1](https://www.ncbi.nlm.nih.gov/protein/156402636?report=genbank&log$=prottop&blast_rank=2&RID=P9NE5SJ4015) | ✓ | X |
| 9 | *Orbicella* | *faveolata* | Anthozoa | APAF-1 | [110058482](https://www.ncbi.nlm.nih.gov/gene/110058482) | - | [XP 020620792.1](https://www.ncbi.nlm.nih.gov/protein/1176100192?report=genbank&log$=prottop&blast_rank=3&RID=P9NE5SJ4015) | ✓ | X |
| **Protostomia** | |  |  |  |  |  |  |  |  |
| 10 | *Acromyrmex* | *echinatior* | Insecta | APAF-1 | [105154033](https://www.ncbi.nlm.nih.gov/gene/105154033) | - | [XP 011067575.1](https://www.ncbi.nlm.nih.gov/protein/746872606?report=genbank&log$=prottop&blast_rank=61&RID=PCFXDSUZ015) | X | X |
| 11 | *Acyrthosiphon* | *pisum* | Insecta | APAF-1 | [100571592](https://www.ncbi.nlm.nih.gov/gene/100571592) | - | [XP 008186216.1](https://www.ncbi.nlm.nih.gov/protein/641673160?report=genbank&log$=prottop&blast_rank=1&RID=PCFXDSUZ015) | X | X |
| 12 | *Aethina* | *tumida* | Insecta | APAF-1 | [109599977](https://www.ncbi.nlm.nih.gov/gene/109599977) | - | [XP 019871594.1](https://www.ncbi.nlm.nih.gov/protein/1133431172?report=genbank&log$=prottop&blast_rank=78&RID=PCFXDSUZ015) | X | X |
| 13 | *Agrilus* | *planipennis* | Insecta | APAF-1 | [108738371](https://www.ncbi.nlm.nih.gov/gene/108738371) | - | [XP 018327269.1](https://www.ncbi.nlm.nih.gov/protein/1069806250?report=genbank&log$=prottop&blast_rank=3&RID=PCFXDSUZ015) | X | X |
| 14 | *Amyelois* | *transitella* | Insecta | APAF-1 | [106136964](https://www.ncbi.nlm.nih.gov/gene/106136964) | - | [XP 013193147.1](https://www.ncbi.nlm.nih.gov/protein/913298637?report=genbank&log$=prottop&blast_rank=42&RID=PCFXDSUZ015) | X | X |
| 15 | *Anoplophora* | *glabripennis* | Insecta | APAF-1 | [108905670](https://www.ncbi.nlm.nih.gov/gene/108905670) | - | [XP 018564158.1](https://www.ncbi.nlm.nih.gov/protein/1080045866?report=genbank&log$=prottop&blast_rank=16&RID=PCFXDSUZ015) | X | X |
| 16 | *Apis* | *cerana* | Insecta | APAF-1 | [107996897](https://www.ncbi.nlm.nih.gov/gene/107996897) | - | [XP 016910708.1](https://www.ncbi.nlm.nih.gov/protein/1035606348?report=genbank&log$=prottop&blast_rank=104&RID=PCFXDSUZ015) | X | X |
| 17 | *Apis* | *dorsata* | Insecta | APAF-1 | [102681002](https://www.ncbi.nlm.nih.gov/gene/102681002) | - | [XP 006622914.1](https://www.ncbi.nlm.nih.gov/protein/572314483?report=genbank&log$=prottop&blast_rank=105&RID=PCFXDSUZ015) | X | X |
| 18 | *Aplysia* | *californica* | Gastropoda | CASP9 | [101846323](https://www.ncbi.nlm.nih.gov/sites/entrez?db=gene&cmd=Retrieve&dopt=full_report&list_uids=101846323) | - | [XP 012945424.1](https://www.ncbi.nlm.nih.gov/nuccore/XP_012945424.1) | X | X |
| 19 | *Athalia* | *rosae* | Insecta | APAF-1 | [105691126](https://www.ncbi.nlm.nih.gov/gene/105691126) | - | [XP 012264834.1](https://www.ncbi.nlm.nih.gov/protein/817084371?report=genbank&log$=prottop&blast_rank=20&RID=PCFXDSUZ015) | X | X |
| 20 | *Atta* | *cephalotes* | Insecta | APAF-1 | [105617990](https://www.ncbi.nlm.nih.gov/gene/105617990) | - | [XP 012054917.1](https://www.ncbi.nlm.nih.gov/protein/801383505?report=genbank&log$=prottop&blast_rank=27&RID=PCFXDSUZ015) | X | X |
| 21 | *Atta* | *colombica* | Insecta | APAF-1 | [108686929](https://www.ncbi.nlm.nih.gov/gene/108686929) | - | [XP 018047941.1](https://www.ncbi.nlm.nih.gov/protein/1068399944?report=genbank&log$=prottop&blast_rank=26&RID=PCFXDSUZ015) | X | X |
| 22 | *Bactrocera* | *oleae* | Insecta | CASP9 | [106625175](https://www.ncbi.nlm.nih.gov/sites/entrez?db=gene&cmd=Retrieve&dopt=full_report&list_uids=106625175) | - | [XP 014100464.1](https://www.ncbi.nlm.nih.gov/protein/929380181) | X | X |
| 23 | *Bombus* | *terrestris* | Insecta | APAF-1 | [100651239](https://www.ncbi.nlm.nih.gov/gene/100651239) | - | [XP 012175672.1](https://www.ncbi.nlm.nih.gov/protein/808116450?report=genbank&log$=prottop&blast_rank=120&RID=PCFXDSUZ015) | X | X |
| 24 | *Bombus* | *impatiens* | Insecta | APAF-1 | [100748962](https://www.ncbi.nlm.nih.gov/gene/100748962) | - | [XP 012240134.1](https://www.ncbi.nlm.nih.gov/protein/815908967?report=genbank&log$=prottop&blast_rank=114&RID=PCFXDSUZ015) | X | X |
| 25 | *Bombyx* | *mori* | Insecta | APAF-1 | [100529085](https://www.ncbi.nlm.nih.gov/gene/100529085) | - | [XP 012553167.2](https://www.ncbi.nlm.nih.gov/protein/1200712491?report=genbank&log$=prottop&blast_rank=86&RID=PCFXDSUZ015) | X | X |
| 26 | *Camponotus* | *floridanus* | Insecta | APAF-1 | [105257852](https://www.ncbi.nlm.nih.gov/gene/105257852) | - | [XP 019885106.1](https://www.ncbi.nlm.nih.gov/protein/1134640328?report=genbank&log$=prottop&blast_rank=97&RID=PCFXDSUZ015) | X | X |
| 27 | *Cephus* | *cinctus* | Insecta | APAF-1 | [107274732](https://www.ncbi.nlm.nih.gov/gene/107274732) | - | [XP 015609643.1](https://www.ncbi.nlm.nih.gov/sites/entrez?cmd=Search&db=protein&term=XP_015609643.1&dopt=GenBank) | X | X |
|  |  |  |  | CASP9 | [107266595](https://www.ncbi.nlm.nih.gov/gene/107266595) | - | [XP 015592710.1](https://www.ncbi.nlm.nih.gov/protein/1000740228?report=genbank&log$=prottop&blast_rank=1&RID=PA6CYHSY015) | X | X |
| 28 | *Cerapachys* | *biroi* | Insecta | APAF-1 | [105282532](https://www.ncbi.nlm.nih.gov/sites/entrez?db=gene&cmd=Retrieve&dopt=full_report&list_uids=105282532) | - | [XP 011342858.1](https://www.ncbi.nlm.nih.gov/nuccore/XP_011342858.1) | X | X |
| 29 | *Ceratina* | *calcarata* | Insecta | APAF-1 | [108627465](https://www.ncbi.nlm.nih.gov/gene/108627465) | - | [XP 017884242.1](https://www.ncbi.nlm.nih.gov/protein/1061116035?report=genbank&log$=prottop&blast_rank=55&RID=PCFXDSUZ015) | X | X |
| 30 | *Ceratitis* | *capitata* | Insecta | APAF-1 | [101448318](https://www.ncbi.nlm.nih.gov/sites/entrez?db=gene&cmd=Retrieve&dopt=full_report&list_uids=101448318) | - | [XP 012161874.1](https://www.ncbi.nlm.nih.gov/nuccore/XP_012161874.1) | X | X |
| 31 | *Ceratosolen* | *solmsi marchali* | Insecta | APAF-1 | [105362014](https://www.ncbi.nlm.nih.gov/gene/105362014) | - | [XP 011497642.1](https://www.ncbi.nlm.nih.gov/protein/766930965?report=genbank&log$=prottop&blast_rank=18&RID=PCFXDSUZ015) | X | X |
| 32 | *Cimex* | *lectularius* | Insecta | APAF-1 | [106668095](https://www.ncbi.nlm.nih.gov/gene/106668095) | - | [XP 014252009.1](https://www.ncbi.nlm.nih.gov/protein/939264041?report=genbank&log$=prottop&blast_rank=4&RID=PCFXDSUZ015) | X | X |
| 33 | *Clonorchis* | *sinensis* | Trematoda | APAF-1 | n/a | - | [GAA49379.1](https://www.ncbi.nlm.nih.gov/sites/entrez?cmd=Search&db=protein&term=GAA49379.1&dopt=GenBank) | X | X |
|  |  |  |  | CASP9 | n/a | - | [GAA35878.1](https://www.ncbi.nlm.nih.gov/sites/entrez?cmd=Search&db=protein&term=GAA35878.1&dopt=GenBank) | X | X |
| 34 | *Copidosoma* | *floridanum* | Insecta | APAF-1 | [106635991](https://www.ncbi.nlm.nih.gov/gene/106635991) | - | [XP 014203691.1](https://www.ncbi.nlm.nih.gov/protein/936577791?report=genbank&log$=prottop&blast_rank=119&RID=PCFXDSUZ015) | X | X |
| 35 | *Cyphomyrmex* | *costatus* | Insecta | APAF-1 | [108779435](https://www.ncbi.nlm.nih.gov/gene/108779435) | - | [XP 018402370.1](https://www.ncbi.nlm.nih.gov/protein/1070609203?report=genbank&log$=prottop&blast_rank=35&RID=PCFXDSUZ015) | X | X |
| 36 | *Danaus* | *plexippus plexippus* | Insecta | APAF-1 | n/a | - | [OWR45425.1](https://www.ncbi.nlm.nih.gov/protein/1209691073?report=genbank&log$=prottop&blast_rank=10&RID=PCFXDSUZ015) | X | X |
| 37 | *Daphnia* | *magna* | Branchiopoda | APAF-1 | n/a | - | [KZS20735.1](https://www.ncbi.nlm.nih.gov/protein/1022777150?report=genbank&log$=prottop&blast_rank=4&RID=P9R4B8UX014) | X | X |
| 38 | *Daphnia* | *pulex* | Branchiopoda | APAF-1 | n/a | - | [EFX74325.1](https://www.ncbi.nlm.nih.gov/protein/321463308?report=genbank&log$=prottop&blast_rank=3&RID=P9R4B8UX014) | X | X |
| 39 | *Diachasma* | *alloeum* | Insecta | APAF-1 | [107041411](https://www.ncbi.nlm.nih.gov/gene/107041411) | - | [XP 015117459.1](https://www.ncbi.nlm.nih.gov/protein/970903243?report=genbank&log$=prottop&blast_rank=87&RID=PCFXDSUZ015) | X | X |
| 40 | *Dinoponera* | *quadriceps* | Insecta | APAF-1 | [106744477](https://www.ncbi.nlm.nih.gov/gene/106744477) | - | [XP 014474785.1](https://www.ncbi.nlm.nih.gov/protein/951549664?report=genbank&log$=prottop&blast_rank=70&RID=PCFXDSUZ015) | X | X |
| 41 | *Diuraphis* | *noxia* | Insecta | APAF-1 | [107164576](https://www.ncbi.nlm.nih.gov/gene/107164576) | - | [XP 015367927.1](https://www.ncbi.nlm.nih.gov/protein/985400504?report=genbank&log$=prottop&blast_rank=2&RID=PCFXDSUZ015) | X | X |
| 42 | *Drosophila* | *kikkawai* | Insecta | CASP9 | [108071389](https://www.ncbi.nlm.nih.gov/sites/entrez?db=gene&cmd=Retrieve&dopt=full_report&list_uids=108071389) | - | [XP 017017622.1](https://www.ncbi.nlm.nih.gov/protein/1036749293) | X | X |
| 43 | *Drosophila* | *takahashii* | Insecta | CASP9 | [108068849](https://www.ncbi.nlm.nih.gov/sites/entrez?db=gene&cmd=Retrieve&dopt=full_report&list_uids=108068849) | - | [XP 017014169.1](https://www.ncbi.nlm.nih.gov/protein/1037084886) | X | X |
| 44 | *Dufourea* | *novaeangliae* | Insecta | APAF-1 | [107185829](https://www.ncbi.nlm.nih.gov/gene/107185829) | - | [XP 015429082.1](https://www.ncbi.nlm.nih.gov/protein/987907140?report=genbank&log$=prottop&blast_rank=68&RID=PCFXDSUZ015) | X | X |
| 45 | *Eufriesea* | *mexicana* | Insecta | APAF-1 | [108550513](https://www.ncbi.nlm.nih.gov/gene/108550513) | - | [XP 017759741.1](https://www.ncbi.nlm.nih.gov/protein/1059228679?report=genbank&log$=prottop&blast_rank=66&RID=PCFXDSUZ015) | X | X |
| 46 | *Fopius* | *arisanus* | Insecta | APAF-1 | [105266039](https://www.ncbi.nlm.nih.gov/gene/105266039) | - | [XP 011302222.1](https://www.ncbi.nlm.nih.gov/protein/755952824?report=genbank&log$=prottop&blast_rank=36&RID=PCFXDSUZ015) | X | X |
| 47 | *Halyomorpha* | *halys* | Insecta | APAF-1 | [106684447](https://www.ncbi.nlm.nih.gov/gene/106684447) | - | [XP 014282024.1](https://www.ncbi.nlm.nih.gov/protein/939670511?report=genbank&log$=prottop&blast_rank=8&RID=PCFXDSUZ015) | X | X |
| 48 | *Harpegnathos* | *saltator* | Insecta | APAF-1 | [105183168](https://www.ncbi.nlm.nih.gov/gene/105183168) | - | [XP 011139410.1](https://www.ncbi.nlm.nih.gov/protein/749753994?report=genbank&log$=prottop&blast_rank=49&RID=PCFXDSUZ015) | X | X |
| 49 | *Helicoverpa* | *armigera* | Insecta | APAF-1 | [110370241](https://www.ncbi.nlm.nih.gov/gene/110370241) | - | [XP 021181657.1](https://www.ncbi.nlm.nih.gov/protein/1199374883?report=genbank&log$=prottop&blast_rank=9&RID=PCFXDSUZ015) | X | X |
| 50 | *Ixodes* | *scapularis* | Chelicerata | APAF-1 | [8027224](https://www.ncbi.nlm.nih.gov/gene/8027224) | - | [XP 002407629.1](https://www.ncbi.nlm.nih.gov/protein/241155731?report=genbank&log$=prottop&blast_rank=2&RID=P9R4B8UX014) | X | X |
| 51 | *Limulus* | *polyphemus* | Xiphosura | APAF-1 | [106465856](https://www.ncbi.nlm.nih.gov/gene/106465856) | - | [XP 013781556.1](https://www.ncbi.nlm.nih.gov/protein/926609100?report=genbank&log$=prottop&blast_rank=1&RID=P9R4B8UX014) | X | X |
| 52 | *Linepithema* | *humile* | Insecta | APAF-1 | [105672289](https://www.ncbi.nlm.nih.gov/gene/105672289) | - | [XP 012222555.1](https://www.ncbi.nlm.nih.gov/protein/815803703?report=genbank&log$=prottop&blast_rank=57&RID=PCFXDSUZ015) | X | X |
| 53 | *Lingula* | *anatina* | Lingulata | CASP9 | [106170245](https://www.ncbi.nlm.nih.gov/gene/106170245) | - | [XP 013405490.1](https://www.ncbi.nlm.nih.gov/protein/919044613?report=genbank&log$=prottop&blast_rank=2&RID=PAHHEE9G014) | X | X |
| 54 | *Megachile* | *rotundata* | Insecta | APAF-1 | [100880336](https://www.ncbi.nlm.nih.gov/gene/100880336) | - | [XP 012135177.1](https://www.ncbi.nlm.nih.gov/protein/805766121?report=genbank&log$=prottop&blast_rank=99&RID=PCFXDSUZ015) | X | X |
| 55 | *Melipona* | *quadrifasciata* | Insecta | APAF-1 | n/a | - | [KOX73754.1](https://www.ncbi.nlm.nih.gov/protein/925676551?report=genbank&log$=prottop&blast_rank=69&RID=PCFXDSUZ015) | X | X |
| 56 | *Microplitis* | *demolitor* | Insecta | APAF-1 | [106693462](https://www.ncbi.nlm.nih.gov/gene/106693462) | - | [XP 014296699.1](https://www.ncbi.nlm.nih.gov/protein/939653891?report=genbank&log$=prottop&blast_rank=94&RID=PCFXDSUZ015) | X | X |
| 57 | *Monomorium* | *pharaonis* | Insecta | APAF-1 | [105830371](https://www.ncbi.nlm.nih.gov/gene/105830371) | - | [XP 012525103.1](https://www.ncbi.nlm.nih.gov/protein/826420360?report=genbank&log$=prottop&blast_rank=23&RID=PCFXDSUZ015) | X | X |
| 58 | *Nasonia* | *vitripennis* | Insecta | APAF-1 | [100679184](https://www.ncbi.nlm.nih.gov/gene/100679184) | - | [XP 016843733.1](https://www.ncbi.nlm.nih.gov/protein/1032736606?report=genbank&log$=prottop&blast_rank=79&RID=PCFXDSUZ015) | X | X |
| 59 | *Neodiprion* | *lecontei* | Insecta | APAF-1 | [107218403](https://www.ncbi.nlm.nih.gov/gene/107218403) | - | [XP 015511747.1](https://www.ncbi.nlm.nih.gov/protein/998503297?report=genbank&log$=prottop&blast_rank=38&RID=PCFXDSUZ015) | X | X |
| 60 | *Nicrophorus* | *vespilloides* | Insecta | APAF-1 | [108557857](https://www.ncbi.nlm.nih.gov/gene/108557857) | - | [XP 017770032.1](https://www.ncbi.nlm.nih.gov/protein/1059390408?report=genbank&log$=prottop&blast_rank=64&RID=PCFXDSUZ015) | X | X |
| 61 | *Octopus* | *bimaculoides* | Cephalopoda | CASP9 | [106867725](https://www.ncbi.nlm.nih.gov/sites/entrez?db=gene&cmd=Retrieve&dopt=full_report&list_uids=106867725) | - | [XP 014768177.1](https://www.ncbi.nlm.nih.gov/protein/XP_014768177.1) | X | X |
| 62 | *Ooceraea* | *biroi* | Insecta | APAF-1 | [105282532](https://www.ncbi.nlm.nih.gov/gene/105282532) | - | [XP 011342858.1](https://www.ncbi.nlm.nih.gov/protein/759066792?report=genbank&log$=prottop&blast_rank=19&RID=PCFXDSUZ015) | X | X |
| 63 | *Opisthorchis* | *viverrini* | Trematoda | CASP9 | [20314953](https://www.ncbi.nlm.nih.gov/gene/20314953) | - | [XP 009162878.1](https://www.ncbi.nlm.nih.gov/sites/entrez?cmd=Search&db=protein&term=XP_009162878.1&dopt=GenBank) | X | X |
| 64 | *Orchesella* | *cincta* | Entognatha | APAF-1 | n/a | - | [ODN02648.1](https://www.ncbi.nlm.nih.gov/protein/1061485518?report=genbank&log$=prottop&blast_rank=1&RID=PABGVBER014) | X | X |
| 65 | *Orussus* | *abietinus* | Insecta | APAF-1 | [105698105](https://www.ncbi.nlm.nih.gov/gene/105698105) | - | [XP 012277485.1](https://www.ncbi.nlm.nih.gov/protein/817203331?report=genbank&log$=prottop&blast_rank=11&RID=PCFXDSUZ015) | X | X |
| 66 | *Papilio* | *machaon* | Insecta | APAF-1 | [106711546](https://www.ncbi.nlm.nih.gov/gene/106711546) | - | [XP 014359375.1](https://www.ncbi.nlm.nih.gov/protein/943952996?report=genbank&log$=prottop&blast_rank=15&RID=PCFXDSUZ015) | X | X |
| 67 | *Papilio* | *polytes* | Insecta | APAF-1 | [106103143](https://www.ncbi.nlm.nih.gov/gene/106103143) | - | [XP 013138265.1](https://www.ncbi.nlm.nih.gov/protein/909565856?report=genbank&log$=prottop&blast_rank=12&RID=PCFXDSUZ015) | X | X |
| 68 | *Papilio* | *xuthus* | Insecta | APAF-1 | n/a | - | [KPI94114.1](https://www.ncbi.nlm.nih.gov/protein/930651772?report=genbank&log$=prottop&blast_rank=24&RID=PCFXDSUZ015) | X | X |
| 69 | *Parasteatoda* | *tepidariorum* | Arachnida | APAF-1 | [107449760](https://www.ncbi.nlm.nih.gov/gene/107449760) | - | [XP 015920891.1](https://www.ncbi.nlm.nih.gov/protein/1009578872?report=genbank&log$=prottop&blast_rank=10&RID=P9R4B8UX014) | X | X |
|  |  |  |  | CASP9 | [107448186](https://www.ncbi.nlm.nih.gov/gene/107448186) | - | [XP 015918787.1](https://www.ncbi.nlm.nih.gov/sites/entrez?cmd=Search&db=protein&term=XP_015918787.1&dopt=GenBank) | X | X |
| 70 | *Plutella* | *xylostella* | Insecta | APAF-1 | [105383941](https://www.ncbi.nlm.nih.gov/gene/105383941) | - | [XP 011552382.1](https://www.ncbi.nlm.nih.gov/protein/768422996?report=genbank&log$=prottop&blast_rank=39&RID=PCFXDSUZ015) | X | X |
| 71 | *Pogonomyrmex* | *barbatus* | Insecta | APAF-1 | [105432976](https://www.ncbi.nlm.nih.gov/gene/105432976) | - | [XP 011646318.1](https://www.ncbi.nlm.nih.gov/protein/769868547?report=genbank&log$=prottop&blast_rank=41&RID=PCFXDSUZ015) | X | X |
| 72 | *Polistes* | *canadensis* | Insecta | APAF-1 | [106788106](https://www.ncbi.nlm.nih.gov/gene/106788106) | - | [XP 014606549.1](https://www.ncbi.nlm.nih.gov/protein/954565222?report=genbank&log$=prottop&blast_rank=33&RID=PCFXDSUZ015) | X | X |
| 73 | *Polistes* | *dominula* | Insecta | APAF-1 | [107073095](https://www.ncbi.nlm.nih.gov/gene/107073095) | - | [XP 015189029.1](https://www.ncbi.nlm.nih.gov/protein/972213100?report=genbank&log$=prottop&blast_rank=25&RID=PCFXDSUZ015) | X | X |
| 74 | *Priapulus* | *caudatus* | Priapulida | APAF-1 | [106811529](https://www.ncbi.nlm.nih.gov/gene/106811529) | - | [XP 014670675.1](https://www.ncbi.nlm.nih.gov/protein/957840721?report=genbank&log$=prottop&blast_rank=1&RID=PAB4B6XD015) | X | X |
| 75 | *Pseudomyrmex* | *gracilis* | Insecta | APAF-1 | [109852961](https://www.ncbi.nlm.nih.gov/gene/109852961) | - | [XP 020280208.1](https://www.ncbi.nlm.nih.gov/protein/1153716385?report=genbank&log$=prottop&blast_rank=46&RID=PCFXDSUZ015) | X | X |
| 76 | *Schistosoma* | *haematobium* | Trematoda | APAF-1 | [24595980](https://www.ncbi.nlm.nih.gov/sites/entrez?db=gene&cmd=Retrieve&dopt=full_report&list_uids=24595980) | - | [XP 012800052.1](https://www.ncbi.nlm.nih.gov/protein/844869532) | X | X |
| 77 | *Solenopsis* | *invicta* | Insecta | APAF-1 | [105193154](https://www.ncbi.nlm.nih.gov/gene/105193154) | - | [XP 011155807.1](https://www.ncbi.nlm.nih.gov/protein/751206563?report=genbank&log$=prottop&blast_rank=21&RID=PCFXDSUZ015) | X | X |
| 78 | *Trachymyrmex* | *cornetzi* |  | APAF-1 | [108765102](https://www.ncbi.nlm.nih.gov/gene/108765102) | - | [XP 018369153.1](https://www.ncbi.nlm.nih.gov/protein/1070198688?report=genbank&log$=prottop&blast_rank=47&RID=PCFXDSUZ015) | X | X |
| 79 | *Trachymyrmex* | *septentrionalis* | Insecta | APAF-1 | [108749296](https://www.ncbi.nlm.nih.gov/gene/108749296) | - | [XP 018343397.1](https://www.ncbi.nlm.nih.gov/protein/1070162844?report=genbank&log$=prottop&blast_rank=65&RID=PCFXDSUZ015) | X | X |
| 80 | *Trachymyrmex* | *zeteki* | Insecta | APAF-1 | [108730745](https://www.ncbi.nlm.nih.gov/gene/108730745) | - | [XP 018316145.1](https://www.ncbi.nlm.nih.gov/protein/1069719760?report=genbank&log$=prottop&blast_rank=121&RID=PCFXDSUZ015) | X | X |
| 81 | *Tribolium* | *castaneum* | Insecta | APAF-1 | [662893](https://www.ncbi.nlm.nih.gov/gene/662893) | - | [XP 008201360.1](https://www.ncbi.nlm.nih.gov/protein/642913035?report=genbank&log$=prottop&blast_rank=90&RID=PCFXDSUZ015) | X | X |
| 82 | *Trichinella* | *nativa* | Enoplea | APAF-1 | n/a | - | [OUC48424.1](https://www.ncbi.nlm.nih.gov/protein/1194519247?report=genbank&log$=prottop&blast_rank=28&RID=PCRTX35P014) | X | X |
| 83 | *Trichinella* | *papuae* | Enoplea | APAF-1 | n/a | - | [KRZ71961.1](https://www.ncbi.nlm.nih.gov/protein/954594743?report=genbank&log$=prottop&blast_rank=36&RID=PCRTX35P014) | X | X |
| 84 | *Trichinella* | *pseudospiralis* | Enoplea | APAF-1 | n/a | - | [KRX99873.1](https://www.ncbi.nlm.nih.gov/protein/954312189?report=genbank&log$=prottop&blast_rank=2&RID=PABWSVZN014) | X | X |
| 85 | *Trichinella* | *zimbabwensis* | Enoplea | APAF-1 | n/a | - | [KRZ19337.1](https://www.ncbi.nlm.nih.gov/sites/entrez?cmd=Search&db=protein&term=KRZ19337.1&dopt=GenBank) | X | X |
| 86 | *Trichogramma* | *pretiosum* | Insecta | APAF-1 | n/a | - | [XP 014234781.1](https://www.ncbi.nlm.nih.gov/protein/936713638?report=genbank&log$=prottop&blast_rank=48&RID=PCFXDSUZ015) | X | X |
| 87 | *Trichuris* | *suis* | Enoplea | APAF-1 | n/a | - | KFD49941.1 | X | X |
| 88 | *Vollenhovia* | *emeryi* | Insecta | APAF-1 | [105566676](https://www.ncbi.nlm.nih.gov/gene/105566676) | - | [XP 011876259.1](https://www.ncbi.nlm.nih.gov/protein/795075663?report=genbank&log$=prottop&blast_rank=29&RID=PCFXDSUZ015) | X | X |
| 89 | *Wasmannia* | *auropunctata* | Insecta | APAF-1 | [105451514](https://www.ncbi.nlm.nih.gov/gene/105451514) | - | [XP 011690307.1](https://www.ncbi.nlm.nih.gov/protein/780650725?report=genbank&log$=prottop&blast_rank=72&RID=PCFXDSUZ015) | X | X |
| 90 | *Zootermopsis* | *nevadensis* | Insecta | APAF-1 | n/a | - | [KDR23380.1](https://www.ncbi.nlm.nih.gov/protein/646722367?report=genbank&log$=prottop&blast_rank=7&RID=PCFXDSUZ015) | X | X |
| **Deuterostomia** | |  |  |  |  |  |  |  |  |
| **Hemichordata** | |  |  |  |  |  |  |  |  |
| 91 | *Saccoglossus* | *kowalevskii* | Enteropneusta | APAF-1 | 102804745 | XM 006818234.1 | XP 006818297.1 | ✓ | X |
| **Echinodermata** | |  |  | APAF-1 | 591503 | [XM 011684681.1](https://www.ncbi.nlm.nih.gov/nuccore/XM_011684681.1) | XP 011682983.1 | ✓ | ✓ |
| 92 | *Strongylocentrotus* | *purpuratus* | Echinoidea | CASP9 | 594735 | [XM 011662940.1](https://www.ncbi.nlm.nih.gov/nuccore/XM_011662940.1) | [XP 011661242.1](https://www.ncbi.nlm.nih.gov/protein/780004918?report=genbank&log$=prottop&blast_rank=2&RID=P9WVD56Y015) | ✓ | ✓ |
|  |  |  |  | CYCS | 575347 | [XM 775754.4](https://www.ncbi.nlm.nih.gov/nuccore/XM_775754.4) | [XP 780847.1](https://www.ncbi.nlm.nih.gov/protein/72007473?report=genbank&log$=prottop&blast_rank=1&RID=PD4VEWMN015) | ✓ | ✓ |
| 93 | *Patiria* | *pectinifera* | Asteroidea | CASP9 |  |  | [ACM46824.1](https://www.ncbi.nlm.nih.gov/protein/222145982?report=genbank&log$=prottop&blast_rank=1&RID=P9WVD56Y015) |  |  |
| 94 | *Apostichopus* | *japonicus* | Holothuroidea | CASP9 |  |  | [AOR82888.1](https://www.ncbi.nlm.nih.gov/protein/1070088506?report=genbank&log$=prottop&blast_rank=3&RID=P9WVD56Y015) |  |  |
| **Cephalochordata** | |  |  |  |  |  |  |  |  |
|  |  |  |  | APAF-1 | 7221615 | [XM 002598319.1](https://www.ncbi.nlm.nih.gov/nuccore/XM_002598319.1) | [XP 002598365.1](https://www.ncbi.nlm.nih.gov/protein/260806987?report=genbank&log$=prottop&blast_rank=1&RID=PD0A15RC014) | ✓ | ✓ |
| 95 | *Branchiostoma* | *floridae* | Leptocardii | CASP9 | 7225967 | [XM 002586499.1](https://www.ncbi.nlm.nih.gov/nuccore/XM_002586499.1) | [XP 002586545.1](https://www.ncbi.nlm.nih.gov/sites/entrez?cmd=Search&db=protein&term=XP_002586545.1&dopt=GenBank) | ✓ | ✓ |
|  |  |  |  | CYCS | 7206558 | [XM 002596877.1](https://www.ncbi.nlm.nih.gov/nuccore/XM_002596877.1) | [XP 002596923.1](https://www.ncbi.nlm.nih.gov/sites/entrez?cmd=Search&db=protein&term=XP_002596923.1&dopt=GenBank) | ✓ | ✓ |
| 96 | *Branchiostoma* | *belcheri* | Leptocardii | CASP9 | [109469526](https://www.ncbi.nlm.nih.gov/gene/109469526) | [XM 019768053.1](https://www.ncbi.nlm.nih.gov/nuccore/XM_019768053.1) | [XP 019623612.1](https://www.ncbi.nlm.nih.gov/sites/entrez?cmd=Search&db=protein&term=XP_019623612.1&dopt=GenBank) | ✓ | X |
| **Vertebrata** | |  |  |  |  |  |  |  |  |
|  |  |  |  | APAF-1 | [106973746](https://www.ncbi.nlm.nih.gov/sites/entrez?db=gene&cmd=Retrieve&dopt=full_report&list_uids=106973746) | [XM 015070845.1](https://www.ncbi.nlm.nih.gov/nuccore/961731800) | [XP 014926331.1](https://www.ncbi.nlm.nih.gov/protein/961731801) | ✓ | ✓ |
| 97 | *Acinonyx* | *jubatus* | Mammalia | CASP9 | [106975594](https://www.ncbi.nlm.nih.gov/sites/entrez?db=gene&cmd=Retrieve&dopt=full_report&list_uids=106975594) | [XM 015072927.1](https://www.ncbi.nlm.nih.gov/nuccore/XM_015072927.1) | [XP 014928413.1](https://www.ncbi.nlm.nih.gov/protein/961735855) | ✓ | ✓ |
|  |  |  |  | CYCS | [106982840](https://www.ncbi.nlm.nih.gov/sites/entrez?db=gene&cmd=Retrieve&dopt=full_report&list_uids=106982840) | [XM 015081013.1](https://www.ncbi.nlm.nih.gov/nuccore/XM_015081013.1) | [XP 014936499.1](https://www.ncbi.nlm.nih.gov/protein/961751748) | ✓ | ✓ |
|  |  |  |  | APAF-1 | [102560580](https://www.ncbi.nlm.nih.gov/sites/entrez?db=gene&cmd=Retrieve&dopt=full_report&list_uids=102560580) | [XM 019480848.1](https://www.ncbi.nlm.nih.gov/nuccore/XM_019480848.1) | [XP 019336393.1](https://www.ncbi.nlm.nih.gov/nuccore/XP_019336393.1) | ✓ | ✓ |
| 98 | *Alligator* | *mississippiensis* | Reptilia | CASP9 | [102573968](https://www.ncbi.nlm.nih.gov/sites/entrez?db=gene&cmd=Retrieve&dopt=full_report&list_uids=102573968) | [XM 019499976.1](https://www.ncbi.nlm.nih.gov/nuccore/XM_019499976.1) | [XP 019355521.1](https://www.ncbi.nlm.nih.gov/nuccore/XP_019355521.1) | ✓ | ✓ |
|  |  |  |  | CYCS | [102563299](https://www.ncbi.nlm.nih.gov/sites/entrez?db=gene&cmd=Retrieve&dopt=full_report&list_uids=102563299) | [XM 006275120.3](https://www.ncbi.nlm.nih.gov/nuccore/XM_006275120.3) | [XP 006275182.1](https://www.ncbi.nlm.nih.gov/protein/564236557) | ✓ | ✓ |
|  |  |  |  | APAF-1 | [102371819](https://www.ncbi.nlm.nih.gov/sites/entrez?db=gene&cmd=Retrieve&dopt=full_report&list_uids=102371819) | [XM 006023636.2](https://www.ncbi.nlm.nih.gov/nuccore/XM_006023636.2) | [XP 006023698.1](https://www.ncbi.nlm.nih.gov/nuccore/XP_006023698.1) | ✓ | ✓ |
| 99 | *Alligator* | *sinensis* | Reptilia | CASP9 | [102384999](https://www.ncbi.nlm.nih.gov/sites/entrez?db=gene&cmd=Retrieve&dopt=full_report&list_uids=102384999) | [XM 006037682.2](https://www.ncbi.nlm.nih.gov/nuccore/XM_006037682.2) | [XP 006037744.1](https://www.ncbi.nlm.nih.gov/protein/557329687) | ✓ | ✓ |
|  |  |  |  | CYCS | [102383053](https://www.ncbi.nlm.nih.gov/sites/entrez?db=gene&cmd=Retrieve&dopt=full_report&list_uids=102383053) | [XM 006027034.2](https://www.ncbi.nlm.nih.gov/nuccore/XM_006027034.2) | [XP 006027096.1](https://www.ncbi.nlm.nih.gov/nuccore/XP_006027096.1) | ✓ | ✓ |
|  |  |  |  | APAF-1 | [101799443](https://www.ncbi.nlm.nih.gov/sites/entrez?db=gene&cmd=Retrieve&dopt=full_report&list_uids=101799443) | [XM 005024526.3](https://www.ncbi.nlm.nih.gov/nuccore/XM_005024526.3) | [XP 005024583.1](https://www.ncbi.nlm.nih.gov/nuccore/XP_005024583.1) | ✓ | ✓ |
| 100 | *Anas* | *platyrhynchos* | Aves | CASP9 | [101795386](https://www.ncbi.nlm.nih.gov/sites/entrez?db=gene&cmd=Retrieve&dopt=full_report&list_uids=101795386) | [XM 021269206.1](https://www.ncbi.nlm.nih.gov/nuccore/XM_021269206.1) | [XP 021124881.1](https://www.ncbi.nlm.nih.gov/nuccore/XP_021124881.1) | ✓ | ✓ |
|  |  |  |  | CYCS | [101801398](https://www.ncbi.nlm.nih.gov/sites/entrez?db=gene&cmd=Retrieve&dopt=full_report&list_uids=101801398) | [XM 005024272.3](https://www.ncbi.nlm.nih.gov/nuccore/XM_005024272.3) | [XP 005024329.1](https://www.ncbi.nlm.nih.gov/protein/514765930) | ✓ | ✓ |
|  |  |  |  | APAF-1 | [100556318](https://www.ncbi.nlm.nih.gov/sites/entrez?db=gene&cmd=Retrieve&dopt=full_report&list_uids=100556318) | [XM 003221102.3](https://www.ncbi.nlm.nih.gov/nuccore/XM_003221102.3) | [XP 003221150.1](https://www.ncbi.nlm.nih.gov/protein/327272756) | ✓ | ✓ |
| 101 | *Anolis* | *carolinensis* | Reptilia | CASP9 | [100560103](https://www.ncbi.nlm.nih.gov/sites/entrez?db=gene&cmd=Retrieve&dopt=full_report&list_uids=100560103) | [XM 008125614.2](https://www.ncbi.nlm.nih.gov/nuccore/XM_008125614.2) | [XP 008123821.1](https://www.ncbi.nlm.nih.gov/protein/637381247) | ✓ | ✓ |
|  |  |  |  | CYCS | [100560655](https://www.ncbi.nlm.nih.gov/sites/entrez?db=gene&cmd=Retrieve&dopt=full_report&list_uids=100560655) | [XM 003222032.3](https://www.ncbi.nlm.nih.gov/nuccore/XM_003222032.3) | [XP 003222080.1](https://www.ncbi.nlm.nih.gov/protein/327274631) | ✓ | ✓ |
|  |  |  |  | APAF-1 | [106488625](https://www.ncbi.nlm.nih.gov/sites/entrez?db=gene&cmd=Retrieve&dopt=full_report&list_uids=106488625) | [XM 013947512.1](https://www.ncbi.nlm.nih.gov/nuccore/XM_013947512.1) | [XP 013802966.1](https://www.ncbi.nlm.nih.gov/protein/926508126) | ✓ | ✓ |
| 102 | *Apteryx* | *australis mantelli* | Aves | CASP9 | [106493414](https://www.ncbi.nlm.nih.gov/sites/entrez?db=gene&cmd=Retrieve&dopt=full_report&list_uids=106493414) | [XM 013953454.1](https://www.ncbi.nlm.nih.gov/nuccore/XM_013953454.1) | [XP 013808908.1](https://www.ncbi.nlm.nih.gov/protein/926519616) | ✓ | ✓ |
|  |  |  |  | CYCS | [106483447](https://www.ncbi.nlm.nih.gov/sites/entrez?db=gene&cmd=Retrieve&dopt=full_report&list_uids=106483447) | [XM 013941367.1](https://www.ncbi.nlm.nih.gov/nuccore/XM_013941367.1) | [XP 013796821.1](https://www.ncbi.nlm.nih.gov/nuccore/XP_013796821.1) | ✓ | ✓ |
|  |  |  |  | APAF-1 | [105400874](https://www.ncbi.nlm.nih.gov/sites/entrez?db=gene&cmd=Retrieve&dopt=full_report&list_uids=105400874) | [XM 011574798.1](https://www.ncbi.nlm.nih.gov/nuccore/XM_011574798.1) | [XP 011573100.1](https://www.ncbi.nlm.nih.gov/protein/768352359) | ✓ | ✓ |
| 103 | *Aquila* | *chrysaetos canadensis* | Aves | CASP9 | [105406545](https://www.ncbi.nlm.nih.gov/sites/entrez?db=gene&cmd=Retrieve&dopt=full_report&list_uids=105406545) | [XM 011585168.1](https://www.ncbi.nlm.nih.gov/nuccore/XM_011585168.1) | [XP 011583470.1](https://www.ncbi.nlm.nih.gov/protein/768372041) | ✓ | ✓ |
|  |  |  |  | CYCS | [105406742](https://www.ncbi.nlm.nih.gov/sites/entrez?db=gene&cmd=Retrieve&dopt=full_report&list_uids=105406742) | [XM 011585528.1](https://www.ncbi.nlm.nih.gov/nuccore/XM_011585528.1) | [XP 011583830.1](https://www.ncbi.nlm.nih.gov/protein/768372734) | ✓ | ✓ |
|  |  |  |  | APAF-1 | [104982240](https://www.ncbi.nlm.nih.gov/sites/entrez?db=gene&cmd=Retrieve&dopt=full_report&list_uids=104982240) | [XM 010831575.1](https://www.ncbi.nlm.nih.gov/nuccore/XM_010831575.1) | [XP 010829877.1](https://www.ncbi.nlm.nih.gov/protein/742235536) | ✓ | ✓ |
| 104 | *Bison* | *bison bison* | Mammalia | CASP9 | [104995965](https://www.ncbi.nlm.nih.gov/sites/entrez?db=gene&cmd=Retrieve&dopt=full_report&list_uids=104995965) | [XM 010850045.1](https://www.ncbi.nlm.nih.gov/nuccore/XM_010850045.1) | [XP 010848347.1](https://www.ncbi.nlm.nih.gov/protein/742164953) | ✓ | ✓ |
|  |  |  |  | CYCS | [104999778](https://www.ncbi.nlm.nih.gov/sites/entrez?db=gene&cmd=Retrieve&dopt=full_report&list_uids=104999778) | [XM 010855376.1](https://www.ncbi.nlm.nih.gov/nuccore/XM_010855376.1) | [XP 010853678.1](https://www.ncbi.nlm.nih.gov/protein/742184520) | ✓ | ✓ |
|  |  |  |  | APAF-1 | [102285660](https://www.ncbi.nlm.nih.gov/sites/entrez?db=gene&cmd=Retrieve&dopt=full_report&list_uids=102285660) | [XM 005905528.2](https://www.ncbi.nlm.nih.gov/nuccore/XM_005905528.2) | [XP 005905590.1](https://www.ncbi.nlm.nih.gov/protein/555987479) | ✓ | ✓ |
| 105 | *Bos* | *mutus* | Mammalia | CASP9 | [102268660](https://www.ncbi.nlm.nih.gov/sites/entrez?db=gene&cmd=Retrieve&dopt=full_report&list_uids=102268660) | [XM 005888921.1](https://www.ncbi.nlm.nih.gov/nuccore/XM_005888921.1) | [XP 005888983.1](https://www.ncbi.nlm.nih.gov/protein/555953621) | ✓ | ✓ |
|  |  |  |  | CYCS | [102277620](https://www.ncbi.nlm.nih.gov/sites/entrez?db=gene&cmd=Retrieve&dopt=full_report&list_uids=102277620) | [XM 005902881.2](https://www.ncbi.nlm.nih.gov/nuccore/XM_005902881.2) | [XP 005902943.1](https://www.ncbi.nlm.nih.gov/nuccore/XP_005902943.1) | ✓ | ✓ |
|  |  |  |  | APAF-1 | [537782](https://www.ncbi.nlm.nih.gov/sites/entrez?db=gene&cmd=Retrieve&dopt=full_report&list_uids=537782) | [NM 001191507.1](https://www.ncbi.nlm.nih.gov/nuccore/NM_001191507.1) | [NP 001178436.1](https://www.ncbi.nlm.nih.gov/nuccore/NP_001178436.1) | ✓ | ✓ |
| 106 | *Bos* | *taurus* | Mammalia | CASP9 | [100140945](https://www.ncbi.nlm.nih.gov/sites/entrez?db=gene&cmd=Retrieve&dopt=full_report&list_uids=100140945) | [NM 001205504.1](https://www.ncbi.nlm.nih.gov/nuccore/NM_001205504.1) | [NP 001192433.1](https://www.ncbi.nlm.nih.gov/protein/329664762) | ✓ | ✓ |
|  |  |  |  | CYCS | [510767](https://www.ncbi.nlm.nih.gov/sites/entrez?db=gene&cmd=Retrieve&dopt=full_report&list_uids=510767) | [NM 001046061.2](https://www.ncbi.nlm.nih.gov/nuccore/NM_001046061.2) | [NP 001039526.1](https://www.ncbi.nlm.nih.gov/nuccore/NP_001039526.1) | ✓ | ✓ |
|  |  |  |  | APAF-1 | [106901477](https://www.ncbi.nlm.nih.gov/sites/entrez?db=gene&cmd=Retrieve&dopt=full_report&list_uids=106901477) | [XM 014964268.1](https://www.ncbi.nlm.nih.gov/nuccore/XM_014964268.1) | [XP 014819754.1](https://www.ncbi.nlm.nih.gov/protein/961007451) | ✓ | ✓ |
| 107 | *Calidris* | *pugnax* | Aves | CASP9 | [106893821](https://www.ncbi.nlm.nih.gov/sites/entrez?db=gene&cmd=Retrieve&dopt=full_report&list_uids=106893821) | [XM 014952053.1](https://www.ncbi.nlm.nih.gov/nuccore/XM_014952053.1) | [XP 014807539.1](https://www.ncbi.nlm.nih.gov/protein/960984552) | ✓ | ✓ |
|  |  |  |  | CYCS | [106885583](https://www.ncbi.nlm.nih.gov/sites/entrez?db=gene&cmd=Retrieve&dopt=full_report&list_uids=106885583) | [XM 014937889.1](https://www.ncbi.nlm.nih.gov/nuccore/XM_014937889.1) | [XP 014793375.1](https://www.ncbi.nlm.nih.gov/protein/960957520) | ✓ | ✓ |
|  |  |  |  | APAF-1 | [103187971](https://www.ncbi.nlm.nih.gov/sites/entrez?db=gene&cmd=Retrieve&dopt=full_report&list_uids=103187971) | [XM 007907743.1](https://www.ncbi.nlm.nih.gov/nuccore/XM_007907743.1) | [XP 007905934.1](https://www.ncbi.nlm.nih.gov/nuccore/XP_007905934.1) | ✓ | ✓ |
| 108 | *Callorhinchus* | *milii* | Chondrichthyes | CASP9 | [103187647](https://www.ncbi.nlm.nih.gov/sites/entrez?db=gene&cmd=Retrieve&dopt=full_report&list_uids=103187647) | [XM 007907235.1](https://www.ncbi.nlm.nih.gov/nuccore/XM_007907235.1) | [XP 007905426.1](https://www.ncbi.nlm.nih.gov/protein/632977578) | ✓ | ✓ |
|  |  |  |  | CYCS | [103183182](https://www.ncbi.nlm.nih.gov/sites/entrez?db=gene&cmd=Retrieve&dopt=full_report&list_uids=103183182) | [XM 007900519.1](https://www.ncbi.nlm.nih.gov/nuccore/XM_007900519.1) | [XP 007898710.1](https://www.ncbi.nlm.nih.gov/protein/632965073) | ✓ | ✓ |
|  |  |  |  | APAF-1 | [103533299](https://www.ncbi.nlm.nih.gov/sites/entrez?db=gene&cmd=Retrieve&dopt=full_report&list_uids=103533299) | [XM 008499140.1](https://www.ncbi.nlm.nih.gov/nuccore/XM_008499140.1) | [XP 008497362.1](https://www.ncbi.nlm.nih.gov/protein/663278060) | ✓ | ✓ |
| 109 | *Calypte* | *anna* | Aves | CASP9 | [103526208](https://www.ncbi.nlm.nih.gov/sites/entrez?db=gene&cmd=Retrieve&dopt=full_report&list_uids=103526208) | [XM 008491238.1](https://www.ncbi.nlm.nih.gov/nuccore/XM_008491238.1) | [XP 008489460.1](https://www.ncbi.nlm.nih.gov/protein/663253026) | ✓ | ✓ |
|  |  |  |  | CYCS | [103528817](https://www.ncbi.nlm.nih.gov/sites/entrez?db=gene&cmd=Retrieve&dopt=full_report&list_uids=103528817) | [XM 008494215.1](https://www.ncbi.nlm.nih.gov/nuccore/XM_008494215.1) | [XP 008492437.1](https://www.ncbi.nlm.nih.gov/protein/663262273) | ✓ | ✓ |
|  |  |  |  | APAF-1 | [105075026](https://www.ncbi.nlm.nih.gov/sites/entrez?db=gene&cmd=Retrieve&dopt=full_report&list_uids=105075026) | [XM 010962769.1](https://www.ncbi.nlm.nih.gov/nuccore/XM_010962769.1) | [XP 010961071.1](https://www.ncbi.nlm.nih.gov/protein/743733685) | ✓ | ✓ |
| 110 | *Camelus* | *bactrianus* | Mammalia | CASP9 | [105070945](https://www.ncbi.nlm.nih.gov/sites/entrez?db=gene&cmd=Retrieve&dopt=full_report&list_uids=105070945) | [XM 010957501.1](https://www.ncbi.nlm.nih.gov/nuccore/XM_010957501.1) | [XP 010955803.1](https://www.ncbi.nlm.nih.gov/protein/743723764) | ✓ | ✓ |
|  |  |  |  | CYCS | [105079817](https://www.ncbi.nlm.nih.gov/sites/entrez?db=gene&cmd=Retrieve&dopt=full_report&list_uids=105079817) | [XM 010968634.1](https://www.ncbi.nlm.nih.gov/nuccore/XM_010968634.1) | [XP 010966936.1](https://www.ncbi.nlm.nih.gov/nuccore/XP_010966936.1) | ✓ | ✓ |
|  |  |  |  | APAF-1 | [102516966](https://www.ncbi.nlm.nih.gov/sites/entrez?db=gene&cmd=Retrieve&dopt=full_report&list_uids=102516966) | [XM 006186553.2](https://www.ncbi.nlm.nih.gov/nuccore/XM_006186553.2) | [XP 006186615.1](https://www.ncbi.nlm.nih.gov/protein/560920809) | ✓ | ✓ |
| 111 | *Camelus* | *ferus* | Mammalia | CASP9 | [102519816](https://www.ncbi.nlm.nih.gov/sites/entrez?db=gene&cmd=Retrieve&dopt=full_report&list_uids=102519816) | [XM 014562275.1](https://www.ncbi.nlm.nih.gov/nuccore/XM_014562275.1) | [XP 014417761.1](https://www.ncbi.nlm.nih.gov/protein/946661561) | ✓ | ✓ |
|  |  |  |  | CYCS | [102524306](https://www.ncbi.nlm.nih.gov/sites/entrez?db=gene&cmd=Retrieve&dopt=full_report&list_uids=102524306) | [XM 014551494.1](https://www.ncbi.nlm.nih.gov/nuccore/XM_014551494.1) | [XP 014406980.1](https://www.ncbi.nlm.nih.gov/nuccore/XP_014406980.1) | ✓ | ✓ |
|  |  |  |  | APAF-1 | [105097442](https://www.ncbi.nlm.nih.gov/sites/entrez?db=gene&cmd=Retrieve&dopt=full_report&list_uids=105097442) | [XM 010989989.1](https://www.ncbi.nlm.nih.gov/nuccore/XM_010989989.1) | [XP 010988291.1](https://www.ncbi.nlm.nih.gov/protein/744594278) | ✓ | ✓ |
| 112 | *Camelus* | *dromedarius* | Mammalia | CASP9 | [105107056](https://www.ncbi.nlm.nih.gov/sites/entrez?db=gene&cmd=Retrieve&dopt=full_report&list_uids=105107056) | [XM 011000878.1](https://www.ncbi.nlm.nih.gov/nuccore/XM_011000878.1) | [XP 010999180.1](https://www.ncbi.nlm.nih.gov/protein/744547014) | ✓ | ✓ |
|  |  |  |  | CYCS | [105095080](https://www.ncbi.nlm.nih.gov/sites/entrez?db=gene&cmd=Retrieve&dopt=full_report&list_uids=105095080) | [XM 010987199.1](https://www.ncbi.nlm.nih.gov/nuccore/XM_010987199.1) | [XP 010985501.1](https://www.ncbi.nlm.nih.gov/nuccore/XP_010985501.1) | ✓ | ✓ |
|  |  |  |  | APAF-1 | [482620](https://www.ncbi.nlm.nih.gov/sites/entrez?db=gene&cmd=Retrieve&dopt=full_report&list_uids=482620) | [XM 022403382.1](https://www.ncbi.nlm.nih.gov/nuccore/XM_022403382.1) | [XP 022259090.1](https://www.ncbi.nlm.nih.gov/nuccore/XP_022259090.1) | ✓ | ✓ |
| 113 | *Canis* | *lupus familiaris* | Mammalia | CASP9 | [487432](https://www.ncbi.nlm.nih.gov/sites/entrez?db=gene&cmd=Retrieve&dopt=full_report&list_uids=487432) | [NM 001031633.1](https://www.ncbi.nlm.nih.gov/nuccore/NM_001031633.1) | [NP 001026803.1](https://www.ncbi.nlm.nih.gov/nuccore/NP_001026803.1) | ✓ | ✓ |
|  |  |  |  | CYCS | [475258](https://www.ncbi.nlm.nih.gov/sites/entrez?db=gene&cmd=Retrieve&dopt=full_report&list_uids=475258) | [NM 001197045.1](https://www.ncbi.nlm.nih.gov/nuccore/NM_001197045.1) | [NP 001183974.1](https://www.ncbi.nlm.nih.gov/protein/308081835) | ✓ | ✓ |
|  |  |  |  | APAF-1 | [102188830](https://www.ncbi.nlm.nih.gov/sites/entrez?db=gene&cmd=Retrieve&dopt=full_report&list_uids=102188830) | [XM 013963999.2](https://www.ncbi.nlm.nih.gov/nuccore/XM_013963999.2) | [XP 013819453.1](https://www.ncbi.nlm.nih.gov/protein/926692117) | ✓ | ✓ |
| 114 | *Capra* | *hircus* | Mammalia | CASP9 | [102174681](https://www.ncbi.nlm.nih.gov/sites/entrez?db=gene&cmd=Retrieve&dopt=full_report&list_uids=102174681) | [XM 005690814.3](https://www.ncbi.nlm.nih.gov/nuccore/XM_005690814.3) | [XP 005690871.2](https://www.ncbi.nlm.nih.gov/protein/926711536) | ✓ | ✓ |
|  |  |  |  | CYCS | [102173997](https://www.ncbi.nlm.nih.gov/sites/entrez?db=gene&cmd=Retrieve&dopt=full_report&list_uids=102173997) | [XM 018047078.1](https://www.ncbi.nlm.nih.gov/nuccore/XM_018047078.1) | [XP 017902567.1](https://www.ncbi.nlm.nih.gov/protein/1062954568) | ✓ | ✓ |
|  |  |  |  | APAF-1 | [109690039](https://www.ncbi.nlm.nih.gov/sites/entrez?db=gene&cmd=Retrieve&dopt=full_report&list_uids=109690039) | [XM 020169152.1](https://www.ncbi.nlm.nih.gov/nuccore/XM_020169152.1) | [XP 020024741.1](https://www.ncbi.nlm.nih.gov/protein/XP_020024741.1) | ✓ | ✓ |
| 115 | *Castor* | *canadensis* | Mammalia | CASP9 | [109692548](https://www.ncbi.nlm.nih.gov/sites/entrez?db=gene&cmd=Retrieve&dopt=full_report&list_uids=109692548) | [XM 020173208.1](https://www.ncbi.nlm.nih.gov/nuccore/XM_020173208.1) | [XP 020028797.1](https://www.ncbi.nlm.nih.gov/protein/XP_020028797.1) | ✓ | ✓ |
|  |  |  |  | CYCS | [109697493](https://www.ncbi.nlm.nih.gov/sites/entrez?db=gene&cmd=Retrieve&dopt=full_report&list_uids=109697493) | [XM 020181133.1](https://www.ncbi.nlm.nih.gov/nuccore/XM_020181133.1) | [XP 020036722.1](https://www.ncbi.nlm.nih.gov/protein/1147375043) | ✓ | ✓ |
|  |  |  |  | APAF-1 | [108290860](https://www.ncbi.nlm.nih.gov/sites/entrez?db=gene&cmd=Retrieve&dopt=full_report&list_uids=108290860) | [XM 017510886.1](https://www.ncbi.nlm.nih.gov/nuccore/XM_017510886.1) | [XP 017366375.1](https://www.ncbi.nlm.nih.gov/protein/XP_017366375.1) | ✓ | ✓ |
| 116 | *Cebus* | *capucinus imitator* | Mammalia | CASP9 | [108286842](https://www.ncbi.nlm.nih.gov/sites/entrez?db=gene&cmd=Retrieve&dopt=full_report&list_uids=108286842) | [XM 017504959.1](https://www.ncbi.nlm.nih.gov/nuccore/XM_017504959.1) | [XP 017360448.1](https://www.ncbi.nlm.nih.gov/protein/XP_017360448.1) | ✓ | ✓ |
|  |  |  |  | CYCS | [108303234](https://www.ncbi.nlm.nih.gov/sites/entrez?db=gene&cmd=Retrieve&dopt=full_report&list_uids=108303234) | [XM 017527009.1](https://www.ncbi.nlm.nih.gov/nuccore/XM_017527009.1) | [XP 017382498.1](https://www.ncbi.nlm.nih.gov/nuccore/XP_017382498.1) | ✓ | ✓ |
|  |  |  |  | APAF-1 | [101394006](https://www.ncbi.nlm.nih.gov/sites/entrez?db=gene&cmd=Retrieve&dopt=full_report&list_uids=101394006) | [XM 014780154.1](https://www.ncbi.nlm.nih.gov/nuccore/XM_014780154.1) | [XP 014635640.1](https://www.ncbi.nlm.nih.gov/protein/XP_014635640.1) | ✓ | ✓ |
| 117 | *Ceratotherium* | *simum simum* | Mammalia | CASP9 | [101398346](https://www.ncbi.nlm.nih.gov/sites/entrez?db=gene&cmd=Retrieve&dopt=full_report&list_uids=101398346) | [XM 014783888.1](https://www.ncbi.nlm.nih.gov/nuccore/XM_014783888.1) | [XP 014639374.1](https://www.ncbi.nlm.nih.gov/protein/XP_014639374.1) | ✓ | ✓ |
|  |  |  |  | CYCS | [101396758](https://www.ncbi.nlm.nih.gov/sites/entrez?db=gene&cmd=Retrieve&dopt=full_report&list_uids=101396758) | [XM 004418907.2](https://www.ncbi.nlm.nih.gov/nuccore/XM_004418907.2) | [XP 004418964.1](https://www.ncbi.nlm.nih.gov/protein/478489168) | ✓ | ✓ |
|  |  |  |  | APAF-1 | [104397996](https://www.ncbi.nlm.nih.gov/sites/entrez?db=gene&cmd=Retrieve&dopt=full_report&list_uids=104397996) | [XM 010008038.1](https://www.ncbi.nlm.nih.gov/nuccore/XM_010008038.1) | [XP 010006340.1](https://www.ncbi.nlm.nih.gov/protein/701361304) | ✓ | ✓ |
| 118 | *Chaetura* | *pelagica* | Aves | CASP9 | [104395515](https://www.ncbi.nlm.nih.gov/sites/entrez?db=gene&cmd=Retrieve&dopt=full_report&list_uids=104395515) | [XM 010005223.1](https://www.ncbi.nlm.nih.gov/nuccore/XM_010005223.1) | [XP 010003525.1](https://www.ncbi.nlm.nih.gov/protein/701429727) | ✓ | ✓ |
|  |  |  |  | CYCS | [104397577](https://www.ncbi.nlm.nih.gov/sites/entrez?db=gene&cmd=Retrieve&dopt=full_report&list_uids=104397577) | [XM 010007571.1](https://www.ncbi.nlm.nih.gov/nuccore/XM_010007571.1) | [XP 010005873.1](https://www.ncbi.nlm.nih.gov/protein/701357993) | ✓ | ✓ |
|  |  |  |  | APAF-1 | [104284312](https://www.ncbi.nlm.nih.gov/sites/entrez?db=gene&cmd=Retrieve&dopt=full_report&list_uids=104284312) | [XM 009882637.1](https://www.ncbi.nlm.nih.gov/nuccore/XM_009882637.1) | [XP 009880939.1](https://www.ncbi.nlm.nih.gov/protein/699653811) | ✓ | ✓ |
| 119 | *Charadrius* | *vociferus* | Aves | CASP9 | [104290815](https://www.ncbi.nlm.nih.gov/sites/entrez?db=gene&cmd=Retrieve&dopt=full_report&list_uids=104290815) | [XM 009889981.1](https://www.ncbi.nlm.nih.gov/nuccore/XM_009889981.1) | [XP 009888283.1](https://www.ncbi.nlm.nih.gov/protein/699687576) | ✓ | ✓ |
|  |  |  |  | CYCS | [104281523](https://www.ncbi.nlm.nih.gov/sites/entrez?db=gene&cmd=Retrieve&dopt=full_report&list_uids=104281523) | [XM 009879534.1](https://www.ncbi.nlm.nih.gov/nuccore/XM_009879534.1) | [XP 009877836.1](https://www.ncbi.nlm.nih.gov/protein/699639288) | ✓ | ✓ |
|  |  |  |  | APAF-1 | [102940699](https://www.ncbi.nlm.nih.gov/sites/entrez?db=gene&cmd=Retrieve&dopt=full_report&list_uids=102940699) | [XM 007069209.1](https://www.ncbi.nlm.nih.gov/nuccore/XM_007069209.1) | [XP 007069271.1](https://www.ncbi.nlm.nih.gov/nuccore/XP_007069271.1) | ✓ | ✓ |
| 120 | *Chelonia* | *mydas* | Reptilia | CASP9 | [102939753](https://www.ncbi.nlm.nih.gov/sites/entrez?db=gene&cmd=Retrieve&dopt=full_report&list_uids=102939753) | [XM 007067738.1](https://www.ncbi.nlm.nih.gov/nuccore/XM_007067738.1) | [XP 007067800.1](https://www.ncbi.nlm.nih.gov/protein/591386760) | ✓ | ✓ |
|  |  |  |  | CYCS | 102937007 | [XM 007071560.1](https://www.ncbi.nlm.nih.gov/nuccore/XM_007071560.1) | [XP 007071622.1](https://www.ncbi.nlm.nih.gov/nuccore/XP_007071622.1) | ✓ | ✓ |
|  |  |  |  | APAF-1 | [102018463](https://www.ncbi.nlm.nih.gov/sites/entrez?db=gene&cmd=Retrieve&dopt=full_report&list_uids=102018463) | [XM 013518590.1](https://www.ncbi.nlm.nih.gov/nuccore/XM_013518590.1) | [XP 013374044.1](https://www.ncbi.nlm.nih.gov/protein/918578196) | ✓ | ✓ |
| 121 | *Chinchilla* | *lanigera* | Mammalia | CASP9 | [102012799](https://www.ncbi.nlm.nih.gov/sites/entrez?db=gene&cmd=Retrieve&dopt=full_report&list_uids=102012799) | [XM 013507423.1](https://www.ncbi.nlm.nih.gov/nuccore/XM_013507423.1) | [XP 013362877.1](https://www.ncbi.nlm.nih.gov/protein/918650228) | ✓ | ✓ |
|  |  |  |  | CYCS | [102008080](https://www.ncbi.nlm.nih.gov/sites/entrez?db=gene&cmd=Retrieve&dopt=full_report&list_uids=102008080) | [XM 005378936.2](https://www.ncbi.nlm.nih.gov/nuccore/XM_005378936.2) | [XP 005378993.1](https://www.ncbi.nlm.nih.gov/protein/533127666) | ✓ | ✓ |
|  |  |  |  | APAF-1 | [105506411](https://www.ncbi.nlm.nih.gov/sites/entrez?db=gene&cmd=Retrieve&dopt=full_report&list_uids=105506411) | [XM 011934545.1](https://www.ncbi.nlm.nih.gov/nuccore/XM_011934545.1) | [XP 011789935.1](https://www.ncbi.nlm.nih.gov/protein/795095073) | ✓ | ✓ |
| 122 | *Colobus* | *angolensis palliatus* | Mammalia | CASP9 | [105510236](https://www.ncbi.nlm.nih.gov/sites/entrez?db=gene&cmd=Retrieve&dopt=full_report&list_uids=105510236) | [XM 011939909.1](https://www.ncbi.nlm.nih.gov/nuccore/XM_011939909.1) | [XP 011795299.1](https://www.ncbi.nlm.nih.gov/protein/795154287) | ✓ | ✓ |
|  |  |  |  | CYCS | [105509351](https://www.ncbi.nlm.nih.gov/sites/entrez?db=gene&cmd=Retrieve&dopt=full_report&list_uids=105509351) | [XM 011938558.1](https://www.ncbi.nlm.nih.gov/nuccore/XM_011938558.1) | [XP 011793948.1](https://www.ncbi.nlm.nih.gov/protein/795147739) | ✓ | ✓ |
|  |  |  |  | APAF-1 | [102095228](https://www.ncbi.nlm.nih.gov/sites/entrez?db=gene&cmd=Retrieve&dopt=full_report&list_uids=102095228) | [XM 005511705.2](https://www.ncbi.nlm.nih.gov/nuccore/XM_005511705.2) | [XP 005511762.1](https://www.ncbi.nlm.nih.gov/nuccore/XP_005511762.1) | ✓ | ✓ |
| 123 | *Columba* | *livia* | Aves | CASP9 | [102093506](https://www.ncbi.nlm.nih.gov/sites/entrez?db=gene&cmd=Retrieve&dopt=full_report&list_uids=102093506) | [XM 021295213.1](https://www.ncbi.nlm.nih.gov/nuccore/XM_021295213.1) | [XP 021150888.1](https://www.ncbi.nlm.nih.gov/nuccore/XP_021150888.1) | ✓ | ✓ |
|  |  |  |  | CYCS | [102085712](https://www.ncbi.nlm.nih.gov/sites/entrez?db=gene&cmd=Retrieve&dopt=full_report&list_uids=102085712) | [XM 005513477.3](https://www.ncbi.nlm.nih.gov/nuccore/XM_005513477.3) | [XP 005513534.2](https://www.ncbi.nlm.nih.gov/nuccore/XP_005513534.2) | ✓ | ✓ |
|  |  |  |  | APAF-1 | [101630342](https://www.ncbi.nlm.nih.gov/sites/entrez?db=gene&cmd=Retrieve&dopt=full_report&list_uids=101630342) | [XM 004675779.2](https://www.ncbi.nlm.nih.gov/nuccore/XM_004675779.2) | [XP 004675836.1](https://www.ncbi.nlm.nih.gov/protein/507929177) | ✓ | ✓ |
| 124 | *Condylura* | *cristata* | Mammalia | CASP9 | [101634533](https://www.ncbi.nlm.nih.gov/sites/entrez?db=gene&cmd=Retrieve&dopt=full_report&list_uids=101634533) | [XM 004678524.2](https://www.ncbi.nlm.nih.gov/nuccore/XM_004678524.2) | [XP 004678581.1](https://www.ncbi.nlm.nih.gov/protein/507934698) | ✓ | ✓ |
|  |  |  |  | CYCS | [101625994](https://www.ncbi.nlm.nih.gov/sites/entrez?db=gene&cmd=Retrieve&dopt=full_report&list_uids=101625994) | [XM 004676786.2](https://www.ncbi.nlm.nih.gov/nuccore/XM_004676786.2) | [XP 004676843.1](https://www.ncbi.nlm.nih.gov/protein/507931200) | ✓ | ✓ |
|  |  |  |  | APAF-1 | [104696424](https://www.ncbi.nlm.nih.gov/sites/entrez?db=gene&cmd=Retrieve&dopt=full_report&list_uids=104696424) | [XM 010410780.3](https://www.ncbi.nlm.nih.gov/nuccore/XM_010410780.3) | [XP 010409082.2](https://www.ncbi.nlm.nih.gov/protein/XP_010409082.2) | ✓ | ✓ |
| 125 | *Corvus* | *cornix cornix* | Aves | CASP9 | [104697549](https://www.ncbi.nlm.nih.gov/sites/entrez?db=gene&cmd=Retrieve&dopt=full_report&list_uids=104697549) | [XM 010412475.3](https://www.ncbi.nlm.nih.gov/nuccore/XM_010412475.3) | [XP 010410777.3](https://www.ncbi.nlm.nih.gov/nuccore/XP_010410777.3) | ✓ | ✓ |
|  |  |  |  | CYCS | [104689098](https://www.ncbi.nlm.nih.gov/sites/entrez?db=gene&cmd=Retrieve&dopt=full_report&list_uids=104689098) | [XM 010399026.3](https://www.ncbi.nlm.nih.gov/nuccore/XM_010399026.3) | [XP 010397328.1](https://www.ncbi.nlm.nih.gov/protein/727015845) | ✓ | ✓ |
|  |  |  |  | APAF-1 | [107311930](https://www.ncbi.nlm.nih.gov/sites/entrez?db=gene&cmd=Retrieve&dopt=full_report&list_uids=107311930) | [XM 015858858.1](https://www.ncbi.nlm.nih.gov/nuccore/XM_015858858.1) | [XP 015714344.1](https://www.ncbi.nlm.nih.gov/protein/1003714339) | ✓ | ✓ |
| 126 | *Coturnix* | *japonica* | Aves | CASP9 | [107323418](https://www.ncbi.nlm.nih.gov/sites/entrez?db=gene&cmd=Retrieve&dopt=full_report&list_uids=107323418) | [XM 015882460.1](https://www.ncbi.nlm.nih.gov/nuccore/XM_015882460.1) | [XP 015737946.1](https://www.ncbi.nlm.nih.gov/protein/1003979844) | ✓ | ✓ |
|  |  |  |  | CYCS | [107309196](https://www.ncbi.nlm.nih.gov/sites/entrez?db=gene&cmd=Retrieve&dopt=full_report&list_uids=107309196) | [XM 015853794.1](https://www.ncbi.nlm.nih.gov/nuccore/XM_015853794.1) | [XP 015709280.1](https://www.ncbi.nlm.nih.gov/protein/1003750424) | ✓ | ✓ |
|  |  |  |  | APAF-1 | [109316780](https://www.ncbi.nlm.nih.gov/sites/entrez?db=gene&cmd=Retrieve&dopt=full_report&list_uids=109316780) | [XM 019545096.1](https://www.ncbi.nlm.nih.gov/nuccore/XM_019545096.1) | [XP 019400641.1](https://www.ncbi.nlm.nih.gov/nuccore/XP_019400641.1) | ✓ | ✓ |
| 127 | *Crocodylus* | *porosus* | Reptilia | CASP9 | [109314378](https://www.ncbi.nlm.nih.gov/sites/entrez?db=gene&cmd=Retrieve&dopt=full_report&list_uids=109314378) | [XM 019541524.1](https://www.ncbi.nlm.nih.gov/nuccore/XM_019541524.1) | [XP 019397069.1](https://www.ncbi.nlm.nih.gov/protein/XP_019397069.1) | ✓ | ✓ |
|  |  |  |  | CYCS | [109309929](https://www.ncbi.nlm.nih.gov/sites/entrez?db=gene&cmd=Retrieve&dopt=full_report&list_uids=109309929) | [XM 019534831.1](https://www.ncbi.nlm.nih.gov/nuccore/XM_019534831.1) | [XP 019390376.1](https://www.ncbi.nlm.nih.gov/protein/1121799880) | ✓ | ✓ |
|  |  |  |  | APAF-1 | [104055179](https://www.ncbi.nlm.nih.gov/sites/entrez?db=gene&cmd=Retrieve&dopt=full_report&list_uids=104055179) | [XM 009556277.1](https://www.ncbi.nlm.nih.gov/nuccore/XM_009556277.1) | [XP 009554572.1](https://www.ncbi.nlm.nih.gov/protein/696967979) | ✓ | ✓ |
| 128 | *Cuculus* | *canorus* | Aves | CASP9 | [104060614](https://www.ncbi.nlm.nih.gov/sites/entrez?db=gene&cmd=Retrieve&dopt=full_report&list_uids=104060614) | [XM 009562422.1](https://www.ncbi.nlm.nih.gov/nuccore/XM_009562422.1) | [XP 009560717.1](https://www.ncbi.nlm.nih.gov/protein/696985138) | ✓ | ✓ |
|  |  |  |  | CYCS | [104061004](https://www.ncbi.nlm.nih.gov/sites/entrez?db=gene&cmd=Retrieve&dopt=full_report&list_uids=104061004) | [XM 009562866.1](https://www.ncbi.nlm.nih.gov/nuccore/XM_009562866.1) | [XP 009561161.1](https://www.ncbi.nlm.nih.gov/protein/696986353) | ✓ | ✓ |
|  |  |  |  | APAF-1 | [107101831](https://www.ncbi.nlm.nih.gov/sites/entrez?db=gene&cmd=Retrieve&dopt=full_report&list_uids=107101831) | [XM 015400893.1](https://www.ncbi.nlm.nih.gov/nuccore/XM_015400893.1) | [XP 015256379.1](https://www.ncbi.nlm.nih.gov/protein/974114065) | ✓ | ✓ |
| 129 | *Cyprinodon* | *variegatus* | Actinopterygii | CASP9 | [107091512](https://www.ncbi.nlm.nih.gov/sites/entrez?db=gene&cmd=Retrieve&dopt=full_report&list_uids=107091512) | [XM 015385479.1](https://www.ncbi.nlm.nih.gov/nuccore/XM_015385479.1) | [XP 015240965.1](https://www.ncbi.nlm.nih.gov/nuccore/XP_015240965.1) | ✓ | ✓ |
|  |  |  |  | CYCS | 107084350 | [XM 015374151.1](https://www.ncbi.nlm.nih.gov/nuccore/XM_015374151.1) | [XP 015229637.1](https://www.ncbi.nlm.nih.gov/nuccore/XP_015229637.1) | ✓ | ✓ |
|  |  |  |  | APAF-1 | [101432735](https://www.ncbi.nlm.nih.gov/sites/entrez?db=gene&cmd=Retrieve&dopt=full_report&list_uids=101432735) | [XM 004464146.2](https://www.ncbi.nlm.nih.gov/nuccore/XM_004464146.2) | [XP 004464203.1](https://www.ncbi.nlm.nih.gov/protein/488545345) | ✓ | ✓ |
| 130 | *Dasypus* | *novemcinctus* | Mammalia | CASP9 | [101432519](https://www.ncbi.nlm.nih.gov/sites/entrez?db=gene&cmd=Retrieve&dopt=full_report&list_uids=101432519) | [XM 004472170.2](https://www.ncbi.nlm.nih.gov/nuccore/XM_004472170.2) | [XP 004472227.1](https://www.ncbi.nlm.nih.gov/protein/488565085) | ✓ | ✓ |
|  |  |  |  | CYCS | [101413106](https://www.ncbi.nlm.nih.gov/sites/entrez?db=gene&cmd=Retrieve&dopt=full_report&list_uids=101413106) | [XM 004447556.2](https://www.ncbi.nlm.nih.gov/nuccore/XM_004447556.2) | [XP 004447613.1](https://www.ncbi.nlm.nih.gov/protein/488511069) | ✓ | ✓ |
|  |  |  |  | APAF-1 | [105987659](https://www.ncbi.nlm.nih.gov/sites/entrez?db=gene&cmd=Retrieve&dopt=full_report&list_uids=105987659) | [XM 013019001.1](https://www.ncbi.nlm.nih.gov/nuccore/XM_013019001.1) | [XP 012874455.1](https://www.ncbi.nlm.nih.gov/protein/852728729) | ✓ | ✓ |
| 131 | *Dipodomys* | *ordii* | Mammalia | CASP9 | [105989961](https://www.ncbi.nlm.nih.gov/sites/entrez?db=gene&cmd=Retrieve&dopt=full_report&list_uids=105989961) | [XM 013022232.1](https://www.ncbi.nlm.nih.gov/nuccore/XM_013022232.1) | [XP 012877686.1](https://www.ncbi.nlm.nih.gov/protein/852766470) | ✓ | ✓ |
|  |  |  |  | CYCS | [105991774](https://www.ncbi.nlm.nih.gov/sites/entrez?db=gene&cmd=Retrieve&dopt=full_report&list_uids=105991774) | [XM 013024494.1](https://www.ncbi.nlm.nih.gov/nuccore/XM_013024494.1) | [XP 012879948.1](https://www.ncbi.nlm.nih.gov/protein/852773041) | ✓ | ✓ |
|  |  |  |  | APAF-1 | [104132316](https://www.ncbi.nlm.nih.gov/sites/entrez?db=gene&cmd=Retrieve&dopt=full_report&list_uids=104132316) | [XM 009645591.1](https://www.ncbi.nlm.nih.gov/nuccore/XM_009645591.1) | [XP 009643886.1](https://www.ncbi.nlm.nih.gov/protein/697847034) | ✓ | ✓ |
| 132 | *Egretta* | *garzetta* | Aves | CASP9 | [104133155](https://www.ncbi.nlm.nih.gov/sites/entrez?db=gene&cmd=Retrieve&dopt=full_report&list_uids=104133155) | [XM 009646525.1](https://www.ncbi.nlm.nih.gov/nuccore/XM_009646525.1) | [XP 009644820.1](https://www.ncbi.nlm.nih.gov/protein/697848893) | ✓ | ✓ |
|  |  |  |  | CYCS | [104126145](https://www.ncbi.nlm.nih.gov/sites/entrez?db=gene&cmd=Retrieve&dopt=full_report&list_uids=104126145) | [XM 009638661.1](https://www.ncbi.nlm.nih.gov/nuccore/XM_009638661.1) | [XP 009636956.1](https://www.ncbi.nlm.nih.gov/protein/697833192) | ✓ | ✓ |
|  |  |  |  | APAF-1 | [106827252](https://www.ncbi.nlm.nih.gov/sites/entrez?db=gene&cmd=Retrieve&dopt=full_report&list_uids=106827252) | [XM 014834996.1](https://www.ncbi.nlm.nih.gov/nuccore/XM_014834996.1) | [XP 014690482.1](https://www.ncbi.nlm.nih.gov/protein/958822192) | ✓ | ✓ |
| 133 | *Equus* | *asinus* | Mammalia | CASP9 | [106828005](https://www.ncbi.nlm.nih.gov/sites/entrez?db=gene&cmd=Retrieve&dopt=full_report&list_uids=106828005) | [XM 014836232.1](https://www.ncbi.nlm.nih.gov/nuccore/XM_014836232.1) | [XP 014691718.1](https://www.ncbi.nlm.nih.gov/protein/958679302) | ✓ | ✓ |
|  |  |  |  | CYCS | [106829744](https://www.ncbi.nlm.nih.gov/sites/entrez?db=gene&cmd=Retrieve&dopt=full_report&list_uids=106829744) | [XM 014839000.1](https://www.ncbi.nlm.nih.gov/nuccore/XM_014839000.1) | [XP 014694486.1](https://www.ncbi.nlm.nih.gov/protein/958689300) | ✓ | ✓ |
|  |  |  |  | APAF-1 | [100052552](https://www.ncbi.nlm.nih.gov/sites/entrez?db=gene&cmd=Retrieve&dopt=full_report&list_uids=100052552) | [XM 005606533.2](https://www.ncbi.nlm.nih.gov/nuccore/XM_005606533.2) | [XP 005606590.1](https://www.ncbi.nlm.nih.gov/protein/545206047) | ✓ | ✓ |
| 134 | *Equus* | *caballus* | Mammalia | CASP9 | [100054331](https://www.ncbi.nlm.nih.gov/sites/entrez?db=gene&cmd=Retrieve&dopt=full_report&list_uids=100054331) | [XM 005607504.2](https://www.ncbi.nlm.nih.gov/nuccore/XM_005607504.2) | [XP 005607561.1](https://www.ncbi.nlm.nih.gov/protein/545209028) | ✓ | ✓ |
|  |  |  |  | CYCS | [100053958](https://www.ncbi.nlm.nih.gov/sites/entrez?db=gene&cmd=Retrieve&dopt=full_report&list_uids=100053958) | [XM 005609174.2](https://www.ncbi.nlm.nih.gov/nuccore/XM_005609174.2) | [XP 005609231.1](https://www.ncbi.nlm.nih.gov/nuccore/XP_005609231.1) | ✓ | ✓ |
|  |  |  |  | APAF-1 | [103550811](https://www.ncbi.nlm.nih.gov/sites/entrez?db=gene&cmd=Retrieve&dopt=full_report&list_uids=103550811) | [XM 008519890.1](https://www.ncbi.nlm.nih.gov/nuccore/XM_008519890.1) | [XP 008518112.1](https://www.ncbi.nlm.nih.gov/protein/664722098) | ✓ | ✓ |
| 135 | *Equus* | *przewalskii* | Mammalia | CASP9 | [103565016](https://www.ncbi.nlm.nih.gov/sites/entrez?db=gene&cmd=Retrieve&dopt=full_report&list_uids=103565016) | [XM 008540964.1](https://www.ncbi.nlm.nih.gov/nuccore/XM_008540964.1) | [XP 008539186.1](https://www.ncbi.nlm.nih.gov/protein/664763068) | ✓ | ✓ |
|  |  |  |  | CYCS | [103561978](https://www.ncbi.nlm.nih.gov/sites/entrez?db=gene&cmd=Retrieve&dopt=full_report&list_uids=103561978) | [XM 008536872.1](https://www.ncbi.nlm.nih.gov/nuccore/XM_008536872.1) | [XP 008535094.1](https://www.ncbi.nlm.nih.gov/protein/664755230) | ✓ | ✓ |
|  |  |  |  | APAF-1 | [103112598](https://www.ncbi.nlm.nih.gov/sites/entrez?db=gene&cmd=Retrieve&dopt=full_report&list_uids=103112598) | [XM 016187661.1](https://www.ncbi.nlm.nih.gov/nuccore/XM_016187661.1) | [XP 016043147.1](https://www.ncbi.nlm.nih.gov/protein/1016640515) | ✓ | ✓ |
| 136 | *Erinaceus* | *europaeus* | Mammalia | CASP9 | [103114081](https://www.ncbi.nlm.nih.gov/sites/entrez?db=gene&cmd=Retrieve&dopt=full_report&list_uids=103114081) | [XM 007523705.2](https://www.ncbi.nlm.nih.gov/nuccore/XM_007523705.2) | [XP 007523767.1](https://www.ncbi.nlm.nih.gov/protein/617605447) | ✓ | ✓ |
|  |  |  |  | CYCS | [103118841](https://www.ncbi.nlm.nih.gov/sites/entrez?db=gene&cmd=Retrieve&dopt=full_report&list_uids=103118841) | [XM 007529055.2](https://www.ncbi.nlm.nih.gov/nuccore/XM_007529055.2) | [XP 007529117.1](https://www.ncbi.nlm.nih.gov/protein/617629004) | ✓ | ✓ |
|  |  |  |  | APAF-1 | [102046505](https://www.ncbi.nlm.nih.gov/sites/entrez?db=gene&cmd=Retrieve&dopt=full_report&list_uids=102046505) | [XM 005444044.2](https://www.ncbi.nlm.nih.gov/nuccore/XM_005444044.2) | [XP 005444101.1](https://www.ncbi.nlm.nih.gov/protein/541984764) | ✓ | ✓ |
| 137 | *Falco* | *cherrug* | Aves | CASP9 | [102054465](https://www.ncbi.nlm.nih.gov/sites/entrez?db=gene&cmd=Retrieve&dopt=full_report&list_uids=102054465) | [XM 014283929.1](https://www.ncbi.nlm.nih.gov/nuccore/XM_014283929.1) | [XP 014139404.1](https://www.ncbi.nlm.nih.gov/protein/929482200) | ✓ | ✓ |
|  |  |  |  | CYCS | [102046053](https://www.ncbi.nlm.nih.gov/sites/entrez?db=gene&cmd=Retrieve&dopt=full_report&list_uids=102046053) | [XM 014285497.1](https://www.ncbi.nlm.nih.gov/nuccore/XM_014285497.1) | [XP 014140972.1](https://www.ncbi.nlm.nih.gov/protein/929493962) | ✓ | ✓ |
|  |  |  |  | APAF-1 | [101911538](https://www.ncbi.nlm.nih.gov/sites/entrez?db=gene&cmd=Retrieve&dopt=full_report&list_uids=101911538) | [XM 005242672.2](https://www.ncbi.nlm.nih.gov/nuccore/XM_005242672.2) | [XP 005242729.1](https://www.ncbi.nlm.nih.gov/protein/529446043) | ✓ | ✓ |
| 138 | *Falco* | *peregrinus* | Aves | CASP9 | [101914844](https://www.ncbi.nlm.nih.gov/sites/entrez?db=gene&cmd=Retrieve&dopt=full_report&list_uids=101914844) | [XM 013299959.1](https://www.ncbi.nlm.nih.gov/nuccore/XM_013299959.1) | [XP 013155413.1](https://www.ncbi.nlm.nih.gov/protein/909782159) | ✓ | ✓ |
|  |  |  |  | CYCS | [101923643](https://www.ncbi.nlm.nih.gov/sites/entrez?db=gene&cmd=Retrieve&dopt=full_report&list_uids=101923643) | [XM 005237191.2](https://www.ncbi.nlm.nih.gov/nuccore/XM_005237191.2) | [XP 005237248.1](https://www.ncbi.nlm.nih.gov/protein/529434940) | ✓ | ✓ |
|  |  |  |  | APAF-1 | [101817511](https://www.ncbi.nlm.nih.gov/sites/entrez?db=gene&cmd=Retrieve&dopt=full_report&list_uids=101817511) | [XM 005039553.1](https://www.ncbi.nlm.nih.gov/nuccore/XM_005039553.1) | [XP 005039610.1](https://www.ncbi.nlm.nih.gov/protein/524983223) | ✓ | ✓ |
| 139 | *Ficedula* | *albicollis* | Aves | CASP9 | [101808292](https://www.ncbi.nlm.nih.gov/sites/entrez?db=gene&cmd=Retrieve&dopt=full_report&list_uids=101808292) | [XM 005057750.2](https://www.ncbi.nlm.nih.gov/nuccore/XM_005057750.2) | [XP 005057807.1](https://www.ncbi.nlm.nih.gov/protein/525020585) | ✓ | ✓ |
|  |  |  |  | CYCS | [101811954](https://www.ncbi.nlm.nih.gov/sites/entrez?db=gene&cmd=Retrieve&dopt=full_report&list_uids=101811954) | [XM 005040982.1](https://www.ncbi.nlm.nih.gov/nuccore/XM_005040982.1) | [XP 005041039.1](https://www.ncbi.nlm.nih.gov/protein/524986178) | ✓ | ✓ |
|  |  |  |  | APAF-1 | [104865574](https://www.ncbi.nlm.nih.gov/sites/entrez?db=gene&cmd=Retrieve&dopt=full_report&list_uids=104865574) | [XM 010629144.1](https://www.ncbi.nlm.nih.gov/nuccore/XM_010629144.1) | [XP 010627446.1](https://www.ncbi.nlm.nih.gov/protein/731226870) | ✓ | ✓ |
| 140 | *Fukomys* | *damarensis* | Mammalia | CASP9 | [104852834](https://www.ncbi.nlm.nih.gov/sites/entrez?db=gene&cmd=Retrieve&dopt=full_report&list_uids=104852834) | [XM 010610619.1](https://www.ncbi.nlm.nih.gov/nuccore/XM_010610619.1) | [XP 010608921.1](https://www.ncbi.nlm.nih.gov/protein/731280210) | ✓ | ✓ |
|  |  |  |  | CYCS | [104864952](https://www.ncbi.nlm.nih.gov/sites/entrez?db=gene&cmd=Retrieve&dopt=full_report&list_uids=104864952) | [XM 010628250.2](https://www.ncbi.nlm.nih.gov/nuccore/XM_010628250.2) | [XP 010626552.1](https://www.ncbi.nlm.nih.gov/nuccore/XP_010626552.1) | ✓ | ✓ |
|  |  |  |  | APAF-1 | [105931177](https://www.ncbi.nlm.nih.gov/sites/entrez?db=gene&cmd=Retrieve&dopt=full_report&list_uids=105931177) | [XM 012869749.2](https://www.ncbi.nlm.nih.gov/nuccore/XM_012869749.2) | [XP 012725203.1](https://www.ncbi.nlm.nih.gov/protein/831542690) | ✓ | ✓ |
| 141 | *Fundulus* | *heteroclitus* | Actinopterygii | CASP9 | [105929117](https://www.ncbi.nlm.nih.gov/sites/entrez?db=gene&cmd=Retrieve&dopt=full_report&list_uids=105929117) | [XM 012866755.2](https://www.ncbi.nlm.nih.gov/nuccore/XM_012866755.2) | [XP 012722209.1](https://www.ncbi.nlm.nih.gov/nuccore/XP_012722209.1) | ✓ | ✓ |
|  |  |  |  | CYCS | [105934284](https://www.ncbi.nlm.nih.gov/sites/entrez?db=gene&cmd=Retrieve&dopt=full_report&list_uids=105934284) | [XM 012874196.2](https://www.ncbi.nlm.nih.gov/nuccore/XM_012874196.2) | [XP 012729650.1](https://www.ncbi.nlm.nih.gov/protein/831555265) | ✓ | ✓ |
|  |  |  |  | APAF-1 | [103590490](https://www.ncbi.nlm.nih.gov/sites/entrez?db=gene&cmd=Retrieve&dopt=full_report&list_uids=103590490) | [XM 008572720.1](https://www.ncbi.nlm.nih.gov/nuccore/XM_008572720.1) | [XP 008570942.1](https://www.ncbi.nlm.nih.gov/protein/667271234) | ✓ | ✓ |
| 142 | *Galeopterus* | *variegatus* | Mammalia | CASP9 | [103590595](https://www.ncbi.nlm.nih.gov/sites/entrez?db=gene&cmd=Retrieve&dopt=full_report&list_uids=103590595) | [XM 008572853.1](https://www.ncbi.nlm.nih.gov/nuccore/XM_008572853.1) | [XP 008571075.1](https://www.ncbi.nlm.nih.gov/protein/667271649) | ✓ | ✓ |
|  |  |  |  | CYCS | [103610678](https://www.ncbi.nlm.nih.gov/sites/entrez?db=gene&cmd=Retrieve&dopt=full_report&list_uids=103610678) | [XM 008594791.1](https://www.ncbi.nlm.nih.gov/nuccore/XM_008594791.1) | [XP 008593013.1](https://www.ncbi.nlm.nih.gov/protein/667335809) | ✓ | ✓ |
|  |  |  |  | APAF-1 | [417926](https://www.ncbi.nlm.nih.gov/sites/entrez?db=gene&cmd=Retrieve&dopt=full_report&list_uids=417926) | [XM 416167.5](https://www.ncbi.nlm.nih.gov/nuccore/XM_416167.5) | [XP 416167.3](https://www.ncbi.nlm.nih.gov/protein/363727703) | ✓ | ✓ |
| 143 | *Gallus* | *gallus* | Aves | CASP9 | [426970](https://www.ncbi.nlm.nih.gov/sites/entrez?db=gene&cmd=Retrieve&dopt=full_report&list_uids=426970) | [XM 424580.5](https://www.ncbi.nlm.nih.gov/nuccore/XM_424580.5) | [XP 424580.5](https://www.ncbi.nlm.nih.gov/protein/971429488) | ✓ | ✓ |
|  |  |  |  | CYCS | [420624](https://www.ncbi.nlm.nih.gov/sites/entrez?db=gene&cmd=Retrieve&dopt=full_report&list_uids=420624) | [XM 015281452.1](https://www.ncbi.nlm.nih.gov/nuccore/XM_015281452.1) | [XP 015136938.1](https://www.ncbi.nlm.nih.gov/nuccore/XP_015136938.1) | ✓ | ✓ |
|  |  |  |  | APAF-1 | [109295891](https://www.ncbi.nlm.nih.gov/sites/entrez?db=gene&cmd=Retrieve&dopt=full_report&list_uids=109295891) | [XM 019515020.1](https://www.ncbi.nlm.nih.gov/nuccore/XM_019515020.1) | [XP 019370565.1](https://www.ncbi.nlm.nih.gov/nuccore/XP_019370565.1) | ✓ | ✓ |
| 144 | *Gavialis* | *gangeticus* | Reptilia | CASP9 | [109293786](https://www.ncbi.nlm.nih.gov/sites/entrez?db=gene&cmd=Retrieve&dopt=full_report&list_uids=109293786) | [XM 019511889.1](https://www.ncbi.nlm.nih.gov/nuccore/XM_019511889.1) | [XP 019367434.1](https://www.ncbi.nlm.nih.gov/protein/XP_019367434.1) | ✓ | ✓ |
|  |  |  |  | CYCS | [109291564](https://www.ncbi.nlm.nih.gov/sites/entrez?db=gene&cmd=Retrieve&dopt=full_report&list_uids=109291564) | [XM 019508604.1](https://www.ncbi.nlm.nih.gov/nuccore/XM_019508604.1) | [XP 019364149.1](https://www.ncbi.nlm.nih.gov/protein/1117287307) | ✓ | ✓ |
|  |  |  |  | APAF-1 | [107118861](https://www.ncbi.nlm.nih.gov/sites/entrez?db=gene&cmd=Retrieve&dopt=full_report&list_uids=107118861) | [XM 015421246.1](https://www.ncbi.nlm.nih.gov/nuccore/XM_015421246.1) | [XP 015276732.1](https://www.ncbi.nlm.nih.gov/protein/975089234) | ✓ | ✓ |
| 145 | *Gekko* | *japonicus* | Reptilia | CASP9 | [107124463](https://www.ncbi.nlm.nih.gov/sites/entrez?db=gene&cmd=Retrieve&dopt=full_report&list_uids=107124463) | [XM 015427936.1](https://www.ncbi.nlm.nih.gov/nuccore/XM_015427936.1) | [XP 015283422.1](https://www.ncbi.nlm.nih.gov/protein/975090435) | ✓ | ✓ |
|  |  |  |  | CYCS | [107126216](https://www.ncbi.nlm.nih.gov/sites/entrez?db=gene&cmd=Retrieve&dopt=full_report&list_uids=107126216) | [XM 015429744.1](https://www.ncbi.nlm.nih.gov/nuccore/XM_015429744.1) | [XP 015285230.1](https://www.ncbi.nlm.nih.gov/nuccore/XP_015285230.1) | ✓ | ✓ |
|  |  |  |  | APAF-1 | [102039129](https://www.ncbi.nlm.nih.gov/sites/entrez?db=gene&cmd=Retrieve&dopt=full_report&list_uids=102039129) | [XM 005421613.2](https://www.ncbi.nlm.nih.gov/nuccore/XM_005421613.2) | [XP 005421670.1](https://www.ncbi.nlm.nih.gov/protein/543262240) | ✓ | ✓ |
| 146 | *Geospiza* | *fortis* | Aves | CASP9 | [102039030](https://www.ncbi.nlm.nih.gov/sites/entrez?db=gene&cmd=Retrieve&dopt=full_report&list_uids=102039030) | [XM 014309984.1](https://www.ncbi.nlm.nih.gov/nuccore/XM_014309984.1) | [XP 014165459.1](https://www.ncbi.nlm.nih.gov/protein/930241878) | ✓ | ✓ |
|  |  |  |  | CYCS | [102033495](https://www.ncbi.nlm.nih.gov/sites/entrez?db=gene&cmd=Retrieve&dopt=full_report&list_uids=102033495) | [XM 005421748.1](https://www.ncbi.nlm.nih.gov/nuccore/XM_005421748.1) | [XP 005421805.1](https://www.ncbi.nlm.nih.gov/protein/543262656) | ✓ | ✓ |
|  |  |  |  | APAF-1 | [101135722](https://www.ncbi.nlm.nih.gov/sites/entrez?db=gene&cmd=Retrieve&dopt=full_report&list_uids=101135722) | [XM 019038859.1](https://www.ncbi.nlm.nih.gov/nuccore/XM_019038859.1) | [XP 018894404.1](https://www.ncbi.nlm.nih.gov/protein/1099122362) | ✓ | ✓ |
| 147 | *Gorilla* | *gorilla* | Mammalia | CASP9 | [101130414](https://www.ncbi.nlm.nih.gov/sites/entrez?db=gene&cmd=Retrieve&dopt=full_report&list_uids=101130414) | [XM 019011435.1](https://www.ncbi.nlm.nih.gov/nuccore/XM_019011435.1) | [XP 018866980.1](https://www.ncbi.nlm.nih.gov/protein/1099079084) | ✓ | ✓ |
|  |  |  |  | CYCS | [101135431](https://www.ncbi.nlm.nih.gov/sites/entrez?db=gene&cmd=Retrieve&dopt=full_report&list_uids=101135431) | [XM 019031021.1](https://www.ncbi.nlm.nih.gov/nuccore/XM_019031021.1) | [XP 018886566.1](https://www.ncbi.nlm.nih.gov/protein/1099290894) | ✓ | ✓ |
|  |  |  |  | APAF-1 | [104828305](https://www.ncbi.nlm.nih.gov/sites/entrez?db=gene&cmd=Retrieve&dopt=full_report&list_uids=104828305) | [XM 010561913.1](https://www.ncbi.nlm.nih.gov/nuccore/XM_010561913.1) | [XP 010560215.1](https://www.ncbi.nlm.nih.gov/nuccore/XP_010560215.1) | ✓ | ✓ |
| 148 | *Haliaeetus* | *leucocephalus* | Aves | CASP9 | [104837865](https://www.ncbi.nlm.nih.gov/sites/entrez?db=gene&cmd=Retrieve&dopt=full_report&list_uids=104837865) | [XM 010577285.1](https://www.ncbi.nlm.nih.gov/nuccore/XM_010577285.1) | [XP 010575587.1](https://www.ncbi.nlm.nih.gov/protein/729757482) | ✓ | ✓ |
|  |  |  |  | CYCS | [104838060](https://www.ncbi.nlm.nih.gov/sites/entrez?db=gene&cmd=Retrieve&dopt=full_report&list_uids=104838060) | [XM 010577589.1](https://www.ncbi.nlm.nih.gov/nuccore/XM_010577589.1) | [XP 010575891.1](https://www.ncbi.nlm.nih.gov/protein/729758069) | ✓ | ✓ |
|  |  |  |  | APAF-1 | [101706721](https://www.ncbi.nlm.nih.gov/sites/entrez?db=gene&cmd=Retrieve&dopt=full_report&list_uids=101706721) | [XM 021258426.1](https://www.ncbi.nlm.nih.gov/nuccore/XM_021258426.1) | [XP 021114085.1](https://www.ncbi.nlm.nih.gov/nuccore/XP_021114085.1) | ✓ | ✓ |
| 149 | *Heterocephalus* | *glaber* | Mammalia | CASP9 | [101702898](https://www.ncbi.nlm.nih.gov/sites/entrez?db=gene&cmd=Retrieve&dopt=full_report&list_uids=101702898) | [XM 021262839.1](https://www.ncbi.nlm.nih.gov/nuccore/XM_021262839.1) | [XP 021118498.1](https://www.ncbi.nlm.nih.gov/nuccore/XP_021118498.1) | ✓ | ✓ |
|  |  |  |  | CYCS | [101718508](https://www.ncbi.nlm.nih.gov/sites/entrez?db=gene&cmd=Retrieve&dopt=full_report&list_uids=101718508) | [XM 004839538.3](https://www.ncbi.nlm.nih.gov/nuccore/XM_004839538.3) | [XP 004839595.1](https://www.ncbi.nlm.nih.gov/nuccore/XP_004839595.1) | ✓ | ✓ |
|  |  |  |  | APAF-1 | [109382336](https://www.ncbi.nlm.nih.gov/sites/entrez?db=gene&cmd=Retrieve&dopt=full_report&list_uids=109382336) | [XM 019641738.1](https://www.ncbi.nlm.nih.gov/nuccore/XM_019641738.1) | [XP 019497283.1](https://www.ncbi.nlm.nih.gov/protein/XP_019497283.1) | ✓ | ✓ |
| 150 | *Hipposideros* | *armiger* | Mammalia | CASP9 | [109381302](https://www.ncbi.nlm.nih.gov/sites/entrez?db=gene&cmd=Retrieve&dopt=full_report&list_uids=109381302) | [XM 019639839.1](https://www.ncbi.nlm.nih.gov/nuccore/XM_019639839.1) | [XP 019495384.1](https://www.ncbi.nlm.nih.gov/nuccore/XP_019495384.1) | ✓ | ✓ |
|  |  |  |  | CYCS | [109381354](https://www.ncbi.nlm.nih.gov/sites/entrez?db=gene&cmd=Retrieve&dopt=full_report&list_uids=109381354) | [XM 019639932.1](https://www.ncbi.nlm.nih.gov/nuccore/XM_019639932.1) | [XP 019495477.1](https://www.ncbi.nlm.nih.gov/protein/1121776862) | ✓ | ✓ |
|  |  |  |  | APAF-1 | [101617070](https://www.ncbi.nlm.nih.gov/sites/entrez?db=gene&cmd=Retrieve&dopt=full_report&list_uids=101617070) | [XM 004650202.1](https://www.ncbi.nlm.nih.gov/nuccore/XM_004650202.1) | [XP 004650259.1](https://www.ncbi.nlm.nih.gov/protein/507532372) | ✓ | ✓ |
| 151 | *Jaculus* | *jaculus* | Mammalia | CASP9 | [101594658](https://www.ncbi.nlm.nih.gov/sites/entrez?db=gene&cmd=Retrieve&dopt=full_report&list_uids=101594658) | [XM 004657624.1](https://www.ncbi.nlm.nih.gov/nuccore/XM_004657624.1) | [XP 004657681.1](https://www.ncbi.nlm.nih.gov/protein/507547702) | ✓ | ✓ |
|  |  |  |  | CYCS | [101596283](https://www.ncbi.nlm.nih.gov/sites/entrez?db=gene&cmd=Retrieve&dopt=full_report&list_uids=101596283) | [XM 004652799.1](https://www.ncbi.nlm.nih.gov/nuccore/XM_004652799.1) | [XP 004652856.1](https://www.ncbi.nlm.nih.gov/protein/507537665) | ✓ | ✓ |
|  |  |  |  | APAF-1 | [108233243](https://www.ncbi.nlm.nih.gov/sites/entrez?db=gene&cmd=Retrieve&dopt=full_report&list_uids=108233243) | [XM 017411553.1](https://www.ncbi.nlm.nih.gov/nuccore/XM_017411553.1) | [XP 017267042.1](https://www.ncbi.nlm.nih.gov/protein/XP_017267042.1) | ✓ | ✓ |
| 152 | *Kryptolebias* | *marmoratus* | Actinopterygii | CASP9 | [108233046](https://www.ncbi.nlm.nih.gov/sites/entrez?db=gene&cmd=Retrieve&dopt=full_report&list_uids=108233046) | [XM 017411220.1](https://www.ncbi.nlm.nih.gov/nuccore/XM_017411220.1) | [XP 017266709.1](https://www.ncbi.nlm.nih.gov/protein/XP_017266709.1) | ✓ | ✓ |
|  |  |  |  | CYCS | [108238381](https://www.ncbi.nlm.nih.gov/sites/entrez?db=gene&cmd=Retrieve&dopt=full_report&list_uids=108238381) | [XM 017420405.1](https://www.ncbi.nlm.nih.gov/nuccore/XM_017420405.1) | [XP 017275894.1](https://www.ncbi.nlm.nih.gov/protein/1041062623) | ✓ | ✓ |
|  |  |  |  | APAF-1 | [109998782](https://www.ncbi.nlm.nih.gov/sites/entrez?db=gene&cmd=Retrieve&dopt=full_report&list_uids=109998782) | [XM 020653658.1](https://www.ncbi.nlm.nih.gov/nuccore/XM_020653658.1) | [XP 020509314.1](https://www.ncbi.nlm.nih.gov/nuccore/XP_020509314.1) | ✓ | ✓ |
| 153 | *Labrus* | *bergylta* | Actinopterygii | CASP9 | [109996636](https://www.ncbi.nlm.nih.gov/sites/entrez?db=gene&cmd=Retrieve&dopt=full_report&list_uids=109996636) | [XM 020650873.1](https://www.ncbi.nlm.nih.gov/nuccore/XM_020650873.1) | [XP 020506529.1](https://www.ncbi.nlm.nih.gov/nuccore/XP_020506529.1) | ✓ | ✓ |
|  |  |  |  | CYCS | [109992886](https://www.ncbi.nlm.nih.gov/sites/entrez?db=gene&cmd=Retrieve&dopt=full_report&list_uids=109992886) | [XM 020645659.1](https://www.ncbi.nlm.nih.gov/nuccore/XM_020645659.1) | [XP 020501315.1](https://www.ncbi.nlm.nih.gov/nuccore/XP_020501315.1) | ✓ | ✓ |
|  |  |  |  | APAF-1 | [108492239](https://www.ncbi.nlm.nih.gov/sites/entrez?db=gene&cmd=Retrieve&dopt=full_report&list_uids=108492239) | [XM 017804308.1](https://www.ncbi.nlm.nih.gov/nuccore/XM_017804308.1) | [XP 017659797.1](https://www.ncbi.nlm.nih.gov/nuccore/XP_017659797.1) | ✓ | ✓ |
| 154 | *Lepidothrix* | *coronata* | Aves | CASP9 | [108505693](https://www.ncbi.nlm.nih.gov/sites/entrez?db=gene&cmd=Retrieve&dopt=full_report&list_uids=108505693) | [XM 017831950.1](https://www.ncbi.nlm.nih.gov/nuccore/XM_017831950.1) | [XP 017687439.1](https://www.ncbi.nlm.nih.gov/protein/XP_017687439.1) | ✓ | ✓ |
|  |  |  |  | CYCS | [108501705](https://www.ncbi.nlm.nih.gov/sites/entrez?db=gene&cmd=Retrieve&dopt=full_report&list_uids=108501705) | [XM 017823870.1](https://www.ncbi.nlm.nih.gov/nuccore/XM_017823870.1) | [XP 017679359.1](https://www.ncbi.nlm.nih.gov/protein/1051192710) | ✓ | ✓ |
|  |  |  |  | APAF-1 | [102694419](https://www.ncbi.nlm.nih.gov/sites/entrez?db=gene&cmd=Retrieve&dopt=full_report&list_uids=102694419) | [XM 015351932.1](https://www.ncbi.nlm.nih.gov/nuccore/XM_015351932.1) | [XP 015207418.1](https://www.ncbi.nlm.nih.gov/nuccore/XP_015207418.1) | ✓ | ✓ |
| 155 | *Lepisosteus* | *oculatus* | Actinopterygii | CASP9 | [102691575](https://www.ncbi.nlm.nih.gov/sites/entrez?db=gene&cmd=Retrieve&dopt=full_report&list_uids=102691575) | [XM 006641900.2](https://www.ncbi.nlm.nih.gov/nuccore/XM_006641900.2) | [XP 006641963.1](https://www.ncbi.nlm.nih.gov/protein/573908489) | ✓ | ✓ |
|  |  |  |  | CYCS | [102685547](https://www.ncbi.nlm.nih.gov/sites/entrez?db=gene&cmd=Retrieve&dopt=full_report&list_uids=102685547) | [XM 006636487.2](https://www.ncbi.nlm.nih.gov/nuccore/XM_006636487.2) | [XP 006636550.1](https://www.ncbi.nlm.nih.gov/protein/573897629) | ✓ | ✓ |
|  |  |  |  | APAF-1 | [103069785](https://www.ncbi.nlm.nih.gov/sites/entrez?db=gene&cmd=Retrieve&dopt=full_report&list_uids=103069785) | [XM 007467163.1](https://www.ncbi.nlm.nih.gov/nuccore/XM_007467163.1) | [XP 007467225.1](https://www.ncbi.nlm.nih.gov/protein/602714014) | ✓ | ✓ |
| 156 | *Lipotes* | *vexillifer* | Mammalia | CASP9 | [103086745](https://www.ncbi.nlm.nih.gov/sites/entrez?db=gene&cmd=Retrieve&dopt=full_report&list_uids=103086745) | [XM 007459290.1](https://www.ncbi.nlm.nih.gov/nuccore/XM_007459290.1) | [XP 007459352.1](https://www.ncbi.nlm.nih.gov/nuccore/XP_007459352.1) | ✓ | ✓ |
|  |  |  |  | CYCS | [103075495](https://www.ncbi.nlm.nih.gov/sites/entrez?db=gene&cmd=Retrieve&dopt=full_report&list_uids=103075495) | [XM 007466066.1](https://www.ncbi.nlm.nih.gov/nuccore/XM_007466066.1) | [XP 007466128.1](https://www.ncbi.nlm.nih.gov/protein/602711786) | ✓ | ✓ |
|  |  |  |  | APAF-1 | [100665091](https://www.ncbi.nlm.nih.gov/sites/entrez?db=gene&cmd=Retrieve&dopt=full_report&list_uids=100665091) | [XM 003405278.2](https://www.ncbi.nlm.nih.gov/nuccore/XM_003405278.2) | [XP 003405326.1](https://www.ncbi.nlm.nih.gov/protein/344266516) | ✓ | ✓ |
| 157 | *Loxodonta* | *africana* | Mammalia | CASP9 | [100656597](https://www.ncbi.nlm.nih.gov/sites/entrez?db=gene&cmd=Retrieve&dopt=full_report&list_uids=100656597) | [XM 010593097.1](https://www.ncbi.nlm.nih.gov/nuccore/XM_010593097.1) | [XP 010591399.1](https://www.ncbi.nlm.nih.gov/protein/731491650) | ✓ | ✓ |
|  |  |  |  | CYCS | [100657132](https://www.ncbi.nlm.nih.gov/sites/entrez?db=gene&cmd=Retrieve&dopt=full_report&list_uids=100657132) | [XM 003407070.2](https://www.ncbi.nlm.nih.gov/nuccore/XM_003407070.2) | [XP 003407118.1](https://www.ncbi.nlm.nih.gov/protein/344270572) | ✓ | ✓ |
|  |  |  |  | APAF-1 | [102137473](https://www.ncbi.nlm.nih.gov/sites/entrez?db=gene&cmd=Retrieve&dopt=full_report&list_uids=102137473) | [XM 005571970.2](https://www.ncbi.nlm.nih.gov/nuccore/XM_005571970.2) | [XP 005572027.1](https://www.ncbi.nlm.nih.gov/protein/544472342) | ✓ | ✓ |
| 158 | *Macaca* | *fascicularis* | Mammalia | CASP9 | [102126830](https://www.ncbi.nlm.nih.gov/sites/entrez?db=gene&cmd=Retrieve&dopt=full_report&list_uids=102126830) | [XM 005544723.2](https://www.ncbi.nlm.nih.gov/nuccore/XM_005544723.2) | [XP 005544780.1](https://www.ncbi.nlm.nih.gov/protein/544408889) | ✓ | ✓ |
|  |  |  |  | CYCS | [102120205](https://www.ncbi.nlm.nih.gov/sites/entrez?db=gene&cmd=Retrieve&dopt=full_report&list_uids=102120205) | [XM 005549954.2](https://www.ncbi.nlm.nih.gov/nuccore/XM_005549954.2) | [XP 005550011.1](https://www.ncbi.nlm.nih.gov/protein/544420868) | ✓ | ✓ |
|  |  |  |  | APAF-1 | [105463369](https://www.ncbi.nlm.nih.gov/sites/entrez?db=gene&cmd=Retrieve&dopt=full_report&list_uids=105463369) | [XM 011710429.1](https://www.ncbi.nlm.nih.gov/nuccore/XM_011710429.1) | [XP 011708731.1](https://www.ncbi.nlm.nih.gov/protein/795492582) | ✓ | ✓ |
| 159 | *Macaca* | *nemestrina* | Mammalia | CASP9 | [105473212](https://www.ncbi.nlm.nih.gov/sites/entrez?db=gene&cmd=Retrieve&dopt=full_report&list_uids=105473212) | [XM 011726879.1](https://www.ncbi.nlm.nih.gov/nuccore/XM_011726879.1) | [XP 011725181.1](https://www.ncbi.nlm.nih.gov/protein/795591802) | ✓ | ✓ |
|  |  |  |  | CYCS | [105475711](https://www.ncbi.nlm.nih.gov/sites/entrez?db=gene&cmd=Retrieve&dopt=full_report&list_uids=105475711) | [XM 011731203.1](https://www.ncbi.nlm.nih.gov/nuccore/XM_011731203.1) | [XP 011729505.1](https://www.ncbi.nlm.nih.gov/nuccore/XP_011729505.1) | ✓ | ✓ |
|  |  |  |  | APAF-1 | [103754448](https://www.ncbi.nlm.nih.gov/sites/entrez?db=gene&cmd=Retrieve&dopt=full_report&list_uids=103754448) | [XM 018080695.1](https://www.ncbi.nlm.nih.gov/nuccore/XM_018080695.1) | [XP 017936184.1](https://www.ncbi.nlm.nih.gov/protein/1062849883) | ✓ | ✓ |
| 160 | *Manacus* | *vitellinus* | Aves | CASP9 | [103762142](https://www.ncbi.nlm.nih.gov/sites/entrez?db=gene&cmd=Retrieve&dopt=full_report&list_uids=103762142) | [XM 018079324.1](https://www.ncbi.nlm.nih.gov/nuccore/XM_018079324.1) | [XP 017934813.1](https://www.ncbi.nlm.nih.gov/protein/1062842843) | ✓ | ✓ |
|  |  |  |  | CYCS | [103754555](https://www.ncbi.nlm.nih.gov/sites/entrez?db=gene&cmd=Retrieve&dopt=full_report&list_uids=103754555) | [XM 018075633.1](https://www.ncbi.nlm.nih.gov/nuccore/XM_018075633.1) | [XP 017931122.1](https://www.ncbi.nlm.nih.gov/protein/1062823869) | ✓ | ✓ |
|  |  |  |  | APAF-1 | [100549497](https://www.ncbi.nlm.nih.gov/sites/entrez?db=gene&cmd=Retrieve&dopt=full_report&list_uids=100549497) | [XM 010709407.2](https://www.ncbi.nlm.nih.gov/nuccore/XM_010709407.2) | [XP 010707709.1](https://www.ncbi.nlm.nih.gov/nuccore/XP_010707709.1) | ✓ | ✓ |
| 161 | *Meleagris* | *gallopavo* | Aves | CASP9 | [100550410](https://www.ncbi.nlm.nih.gov/sites/entrez?db=gene&cmd=Retrieve&dopt=full_report&list_uids=100550410) | [XM 010723032.2](https://www.ncbi.nlm.nih.gov/nuccore/XM_010723032.2) | [XP 010721334.2](https://www.ncbi.nlm.nih.gov/protein/1121946123) | ✓ | ✓ |
|  |  |  |  | CYCS | [100549365](https://www.ncbi.nlm.nih.gov/sites/entrez?db=gene&cmd=Retrieve&dopt=full_report&list_uids=100549365) | [XM 010712788.1](https://www.ncbi.nlm.nih.gov/nuccore/XM_010712788.1) | [XP 010711090.1](https://www.ncbi.nlm.nih.gov/nuccore/XP_010711090.1) | ✓ | ✓ |
|  |  |  |  | APAF-1 | [101881512](https://www.ncbi.nlm.nih.gov/sites/entrez?db=gene&cmd=Retrieve&dopt=full_report&list_uids=101881512) | [XM 005143130.1](https://www.ncbi.nlm.nih.gov/nuccore/XM_005143130.1) | [XP 005143187.1](https://www.ncbi.nlm.nih.gov/protein/XP_005143187.1) | ✓ | ✓ |
| 162 | *Melopsittacus* | *undulatus* | Aves | CASP9 | [101880973](https://www.ncbi.nlm.nih.gov/sites/entrez?db=gene&cmd=Retrieve&dopt=full_report&list_uids=101880973) | [XM 005145478.1](https://www.ncbi.nlm.nih.gov/nuccore/XM_005145478.1) | [XP 005145535.1](https://www.ncbi.nlm.nih.gov/protein/XP_005145535.1) | ✓ | ✓ |
|  |  |  |  | CYCS | [101870620](https://www.ncbi.nlm.nih.gov/sites/entrez?db=gene&cmd=Retrieve&dopt=full_report&list_uids=101870620) | [XM 005152576.1](https://www.ncbi.nlm.nih.gov/nuccore/XM_005152576.1) | [XP 005152633.1](https://www.ncbi.nlm.nih.gov/protein/527269230) | ✓ | ✓ |
|  |  |  |  | APAF-1 | [105883940](https://www.ncbi.nlm.nih.gov/sites/entrez?db=gene&cmd=Retrieve&dopt=full_report&list_uids=105883940) | [XM 012787617.2](https://www.ncbi.nlm.nih.gov/nuccore/XM_012787617.2) | [XP 012643071.2](https://www.ncbi.nlm.nih.gov/protein/XP_012643071.2) | ✓ | ✓ |
| 163 | *Microcebus* | *murinus* | Mammalia | CASP9 | [105870442](https://www.ncbi.nlm.nih.gov/sites/entrez?db=gene&cmd=Retrieve&dopt=full_report&list_uids=105870442) | [XM 012763045.2](https://www.ncbi.nlm.nih.gov/nuccore/XM_012763045.2) | [XP 012618499.1](https://www.ncbi.nlm.nih.gov/protein/829758244) | ✓ | ✓ |
|  |  |  |  | CYCS | [105875906](https://www.ncbi.nlm.nih.gov/sites/entrez?db=gene&cmd=Retrieve&dopt=full_report&list_uids=105875906) | [XM 012773332.2](https://www.ncbi.nlm.nih.gov/nuccore/XM_012773332.2) | [XP 012628786.1](https://www.ncbi.nlm.nih.gov/protein/829809017) | ✓ | ✓ |
|  |  |  |  | APAF-1 | [102002666](https://www.ncbi.nlm.nih.gov/sites/entrez?db=gene&cmd=Retrieve&dopt=full_report&list_uids=102002666) | [XM 005358175.2](https://www.ncbi.nlm.nih.gov/nuccore/XM_005358175.2) | [XP 005358232.1](https://www.ncbi.nlm.nih.gov/protein/532030577) | ✓ | ✓ |
| 164 | *Microtus* | *ochrogaster* | Mammalia | CASP9 | [101999156](https://www.ncbi.nlm.nih.gov/sites/entrez?db=gene&cmd=Retrieve&dopt=full_report&list_uids=101999156) | [XM 013348328.1](https://www.ncbi.nlm.nih.gov/nuccore/XM_013348328.1) | [XP 013203782.1](https://www.ncbi.nlm.nih.gov/nuccore/XP_013203782.1) | ✓ | ✓ |
|  |  |  |  | CYCS | [101980101](https://www.ncbi.nlm.nih.gov/sites/entrez?db=gene&cmd=Retrieve&dopt=full_report&list_uids=101980101) | [XM 005366616.2](https://www.ncbi.nlm.nih.gov/nuccore/XM_005366616.2) | [XP 005366673.1](https://www.ncbi.nlm.nih.gov/protein/532047785) | ✓ | ✓ |
|  |  |  |  | APAF-1 | [107545129](https://www.ncbi.nlm.nih.gov/sites/entrez?db=gene&cmd=Retrieve&dopt=full_report&list_uids=107545129) | [XM 016223182.1](https://www.ncbi.nlm.nih.gov/nuccore/XM_016223182.1) | [XP 016078668.1](https://www.ncbi.nlm.nih.gov/nuccore/XP_016078668.1) | ✓ | ✓ |
| 165 | *Miniopterus* | *natalensis* | Mammalia | CASP9 | [107531542](https://www.ncbi.nlm.nih.gov/sites/entrez?db=gene&cmd=Retrieve&dopt=full_report&list_uids=107531542) | [XM 016205640.1](https://www.ncbi.nlm.nih.gov/nuccore/XM_016205640.1) | [XP 016061126.1](https://www.ncbi.nlm.nih.gov/protein/XP_016061126.1) | ✓ | ✓ |
|  |  |  |  | CYCS | [107540285](https://www.ncbi.nlm.nih.gov/sites/entrez?db=gene&cmd=Retrieve&dopt=full_report&list_uids=107540285) | [XM 016217000.1](https://www.ncbi.nlm.nih.gov/nuccore/XM_016217000.1) | [XP 016072486.1](https://www.ncbi.nlm.nih.gov/protein/1016701661) | ✓ | ✓ |
|  |  |  |  | APAF-1 | [100013051](https://www.ncbi.nlm.nih.gov/sites/entrez?db=gene&cmd=Retrieve&dopt=full_report&list_uids=100013051) | [XM 007503259.1](https://www.ncbi.nlm.nih.gov/nuccore/XM_007503259.1) | [XP 007503321.1](https://www.ncbi.nlm.nih.gov/nuccore/XP_007503321.1) | ✓ | ✓ |
| 166 | *Monodelphis* | *domestica* | Mammalia | CASP9 | [100027338](https://www.ncbi.nlm.nih.gov/sites/entrez?db=gene&cmd=Retrieve&dopt=full_report&list_uids=100027338) | [XM 001377622.3](https://www.ncbi.nlm.nih.gov/nuccore/XM_001377622.3) | [XP 001377659.3](https://www.ncbi.nlm.nih.gov/protein/612025239) | ✓ | ✓ |
|  |  |  |  | CYCS | [100010620](https://www.ncbi.nlm.nih.gov/sites/entrez?db=gene&cmd=Retrieve&dopt=full_report&list_uids=100010620) | [XM 007505338.1](https://www.ncbi.nlm.nih.gov/nuccore/XM_007505338.1) | [XP 007505400.1](https://www.ncbi.nlm.nih.gov/protein/612057009) | ✓ | ✓ |
|  |  |  |  | APAF-1 | [11783](https://www.ncbi.nlm.nih.gov/sites/entrez?db=gene&cmd=Retrieve&dopt=full_report&list_uids=11783) | [NM 001042558.1](https://www.ncbi.nlm.nih.gov/nuccore/NM_001042558.1) | [NP 001036023.1](https://www.ncbi.nlm.nih.gov/protein/110347465) | ✓ | ✓ |
| 167 | *Mus* | *musculus* | Mammalia | CASP9 | [12371](https://www.ncbi.nlm.nih.gov/sites/entrez?db=gene&cmd=Retrieve&dopt=full_report&list_uids=12371) | [NM 015733.5](https://www.ncbi.nlm.nih.gov/nuccore/NM_015733.5) | [NP 056548.2](https://www.ncbi.nlm.nih.gov/nuccore/NP_056548.2) | ✓ | ✓ |
|  |  |  |  | CYCS | [13063](https://www.ncbi.nlm.nih.gov/sites/entrez?db=gene&cmd=Retrieve&dopt=full_report&list_uids=13063) | [NM 007808.4](https://www.ncbi.nlm.nih.gov/nuccore/NM_007808.4) | [NP 031834.1](https://www.ncbi.nlm.nih.gov/protein/6681095) | ✓ | ✓ |
|  |  |  |  | APAF-1 | [101691091](https://www.ncbi.nlm.nih.gov/sites/entrez?db=gene&cmd=Retrieve&dopt=full_report&list_uids=101691091) | [XM 013051503.1](https://www.ncbi.nlm.nih.gov/nuccore/XM_013051503.1) | [XP 012906957.1](https://www.ncbi.nlm.nih.gov/nuccore/XP_012906957.1) | ✓ | ✓ |
| 168 | *Mustela* | *putorius furo* | Mammalia | CASP9 | [101681290](https://www.ncbi.nlm.nih.gov/sites/entrez?db=gene&cmd=Retrieve&dopt=full_report&list_uids=101681290) | [XM 004741419.2](https://www.ncbi.nlm.nih.gov/nuccore/XM_004741419.2) | [XP 004741476.1](https://www.ncbi.nlm.nih.gov/protein/511833670) | ✓ | ✓ |
|  |  |  |  | CYCS | [101690488](https://www.ncbi.nlm.nih.gov/sites/entrez?db=gene&cmd=Retrieve&dopt=full_report&list_uids=101690488) | [XM 004762520.2](https://www.ncbi.nlm.nih.gov/nuccore/XM_004762520.2) | [XP 004762577.1](https://www.ncbi.nlm.nih.gov/protein/511885529) | ✓ | ✓ |
|  |  |  |  | APAF-1 | [102242936](https://www.ncbi.nlm.nih.gov/sites/entrez?db=gene&cmd=Retrieve&dopt=full_report&list_uids=102242936) | [XM 005882353.2](https://www.ncbi.nlm.nih.gov/nuccore/XM_005882353.2) | [XP 005882415.1](https://www.ncbi.nlm.nih.gov/protein/554581863) | ✓ | ✓ |
| 169 | *Myotis* | *brandtii* | Mammalia | CASP9 | [102254877](https://www.ncbi.nlm.nih.gov/sites/entrez?db=gene&cmd=Retrieve&dopt=full_report&list_uids=102254877) | [XM 014533288.1](https://www.ncbi.nlm.nih.gov/nuccore/XM_014533288.1) | [XP 014388774.1](https://www.ncbi.nlm.nih.gov/protein/946809467) | ✓ | ✓ |
|  |  |  |  | CYCS | [102248212](https://www.ncbi.nlm.nih.gov/sites/entrez?db=gene&cmd=Retrieve&dopt=full_report&list_uids=102248212) | [XM 005862978.2](https://www.ncbi.nlm.nih.gov/nuccore/XM_005862978.2) | [XP 005863040.1](https://www.ncbi.nlm.nih.gov/nuccore/XP_005863040.1) | ✓ | ✓ |
|  |  |  |  | APAF-1 | [102773861](https://www.ncbi.nlm.nih.gov/sites/entrez?db=gene&cmd=Retrieve&dopt=full_report&list_uids=102773861) | [XM 006773338.2](https://www.ncbi.nlm.nih.gov/nuccore/XM_006773338.2) | [XP 006773401.1](https://www.ncbi.nlm.nih.gov/protein/584057759) | ✓ | ✓ |
| 170 | *Myotis* | *davidii* | Mammalia | CASP9 | [102759481](https://www.ncbi.nlm.nih.gov/sites/entrez?db=gene&cmd=Retrieve&dopt=full_report&list_uids=102759481) | [XM 006766783.2](https://www.ncbi.nlm.nih.gov/nuccore/XM_006766783.2) | [XP 006766846.1](https://www.ncbi.nlm.nih.gov/protein/584043975) | ✓ | ✓ |
|  |  |  |  | CYCS | [102752458](https://www.ncbi.nlm.nih.gov/sites/entrez?db=gene&cmd=Retrieve&dopt=full_report&list_uids=102752458) | [XM 006778997.2](https://www.ncbi.nlm.nih.gov/nuccore/XM_006778997.2) | [XP 006779060.1](https://www.ncbi.nlm.nih.gov/nuccore/XP_006779060.1) | ✓ | ✓ |
|  |  |  |  | APAF-1 | [102435183](https://www.ncbi.nlm.nih.gov/sites/entrez?db=gene&cmd=Retrieve&dopt=full_report&list_uids=102435183) | [XM 006084110.2](https://www.ncbi.nlm.nih.gov/nuccore/XM_006084110.2) | [XP 006084172.1](https://www.ncbi.nlm.nih.gov/protein/558104231) | ✓ | ✓ |
| 171 | *Myotis* | *lucifugus* | Mammalia | CASP9 | [102428235](https://www.ncbi.nlm.nih.gov/sites/entrez?db=gene&cmd=Retrieve&dopt=full_report&list_uids=102428235) | [XM 006104756.2](https://www.ncbi.nlm.nih.gov/nuccore/XM_006104756.2) | [XP 006104818.1](https://www.ncbi.nlm.nih.gov/protein/558197885) | ✓ | ✓ |
|  |  |  |  | CYCS | [102424392](https://www.ncbi.nlm.nih.gov/sites/entrez?db=gene&cmd=Retrieve&dopt=full_report&list_uids=102424392) | [XM 006088785.2](https://www.ncbi.nlm.nih.gov/nuccore/XM_006088785.2) | [XP 006088847.1](https://www.ncbi.nlm.nih.gov/nuccore/XP_006088847.1) | ✓ | ✓ |
|  |  |  |  | APAF-1 | [103733246](https://www.ncbi.nlm.nih.gov/sites/entrez?db=gene&cmd=Retrieve&dopt=full_report&list_uids=103733246) | [XM 008832051.2](https://www.ncbi.nlm.nih.gov/nuccore/XM_008832051.2) | [XP 008830273.1](https://www.ncbi.nlm.nih.gov/protein/674048303) | ✓ | ✓ |
| 172 | *Nannospalax* | *galili* | Mammalia | CASP9 | [103725336](https://www.ncbi.nlm.nih.gov/sites/entrez?db=gene&cmd=Retrieve&dopt=full_report&list_uids=103725336) | [XM 008822668.2](https://www.ncbi.nlm.nih.gov/nuccore/XM_008822668.2) | [XP 008820890.1](https://www.ncbi.nlm.nih.gov/protein/674095138) | ✓ | ✓ |
|  |  |  |  | CYCS | [103740800](https://www.ncbi.nlm.nih.gov/sites/entrez?db=gene&cmd=Retrieve&dopt=full_report&list_uids=103740800) | [XM 008841488.2](https://www.ncbi.nlm.nih.gov/nuccore/XM_008841488.2) | [XP 008839710.1](https://www.ncbi.nlm.nih.gov/protein/674065619) | ✓ | ✓ |
|  |  |  |  | APAF-1 | [108798110](https://www.ncbi.nlm.nih.gov/sites/entrez?db=gene&cmd=Retrieve&dopt=full_report&list_uids=108798110) | [XM 018569826.1](https://www.ncbi.nlm.nih.gov/nuccore/XM_018569826.1) | [XP 018425328.1](https://www.ncbi.nlm.nih.gov/protein/XP_018425328.1) | ✓ | ✓ |
| 173 | *Nanorana* | *parkeri* | Amphibia | CASP9 | [108797190](https://www.ncbi.nlm.nih.gov/sites/entrez?db=gene&cmd=Retrieve&dopt=full_report&list_uids=108797190) | [XM 018568765.1](https://www.ncbi.nlm.nih.gov/nuccore/XM_018568765.1) | [XP 018424267.1](https://www.ncbi.nlm.nih.gov/protein/XP_018424267.1) | ✓ | ✓ |
|  |  |  |  | CYCS | [108802029](https://www.ncbi.nlm.nih.gov/sites/entrez?db=gene&cmd=Retrieve&dopt=full_report&list_uids=108802029) | [XM 018574075.1](https://www.ncbi.nlm.nih.gov/nuccore/XM_018574075.1) | [XP 018429577.1](https://www.ncbi.nlm.nih.gov/nuccore/XP_018429577.1) | ✓ | ✓ |
|  |  |  |  | APAF-1 | [100590369](https://www.ncbi.nlm.nih.gov/sites/entrez?db=gene&cmd=Retrieve&dopt=full_report&list_uids=100590369) | [XM 003259706.2](https://www.ncbi.nlm.nih.gov/nuccore/XM_003259706.2) | [XP 003259754.1](https://www.ncbi.nlm.nih.gov/protein/332221211) | ✓ | ✓ |
| 174 | *Nomascus* | *leucogenys* | Mammalia | CASP9 | [100597173](https://www.ncbi.nlm.nih.gov/sites/entrez?db=gene&cmd=Retrieve&dopt=full_report&list_uids=100597173) | [XM 003279950.3](https://www.ncbi.nlm.nih.gov/nuccore/XM_003279950.3) | [XP 003279998.1](https://www.ncbi.nlm.nih.gov/protein/332261889) | ✓ | ✓ |
|  |  |  |  | CYCS | [100580069](https://www.ncbi.nlm.nih.gov/sites/entrez?db=gene&cmd=Retrieve&dopt=full_report&list_uids=100580069) | [XM 003270424.3](https://www.ncbi.nlm.nih.gov/nuccore/XM_003270424.3) | [XP 003270472.1](https://www.ncbi.nlm.nih.gov/protein/332242599) | ✓ | ✓ |
|  |  |  |  | APAF-1 | [101383104](https://www.ncbi.nlm.nih.gov/sites/entrez?db=gene&cmd=Retrieve&dopt=full_report&list_uids=101383104) | [XM 004403875.1](https://www.ncbi.nlm.nih.gov/nuccore/XM_004403875.1) | [XP 004403932.1](https://www.ncbi.nlm.nih.gov/protein/472368818) | ✓ | ✓ |
| 175 | *Odobenus* | *rosmarus divergens* | Mammalia | CASP9 | [101362812](https://www.ncbi.nlm.nih.gov/sites/entrez?db=gene&cmd=Retrieve&dopt=full_report&list_uids=101362812) | [XM 012561249.1](https://www.ncbi.nlm.nih.gov/nuccore/XM_012561249.1) | [XP 012416703.1](https://www.ncbi.nlm.nih.gov/protein/823400761) | ✓ | ✓ |
|  |  |  |  | CYCS | [101386944](https://www.ncbi.nlm.nih.gov/sites/entrez?db=gene&cmd=Retrieve&dopt=full_report&list_uids=101386944) | [XM 004397349.2](https://www.ncbi.nlm.nih.gov/nuccore/XM_004397349.2) | [XP 004397406.1](https://www.ncbi.nlm.nih.gov/protein/472355508) | ✓ | ✓ |
|  |  |  |  | APAF-1 | [104334484](https://www.ncbi.nlm.nih.gov/sites/entrez?db=gene&cmd=Retrieve&dopt=full_report&list_uids=104334484) | [XM 009940121.1](https://www.ncbi.nlm.nih.gov/nuccore/XM_009940121.1) | [XP 009938423.1](https://www.ncbi.nlm.nih.gov/protein/700395648) | ✓ | ✓ |
| 176 | *Opisthocomus* | *hoazin* | Aves | CASP9 | [104336695](https://www.ncbi.nlm.nih.gov/sites/entrez?db=gene&cmd=Retrieve&dopt=full_report&list_uids=104336695) | [XM 009942545.1](https://www.ncbi.nlm.nih.gov/nuccore/XM_009942545.1) | [XP 009940847.1](https://www.ncbi.nlm.nih.gov/protein/700400028) | ✓ | ✓ |
|  |  |  |  | CYCS | [104335603](https://www.ncbi.nlm.nih.gov/sites/entrez?db=gene&cmd=Retrieve&dopt=full_report&list_uids=104335603) | [XM 009941349.1](https://www.ncbi.nlm.nih.gov/nuccore/XM_009941349.1) | [XP 009939651.1](https://www.ncbi.nlm.nih.gov/nuccore/XP_009939651.1) | ✓ | ✓ |
|  |  |  |  | APAF-1 | [101280303](https://www.ncbi.nlm.nih.gov/sites/entrez?db=gene&cmd=Retrieve&dopt=full_report&list_uids=101280303) | [XM 004269502.2](https://www.ncbi.nlm.nih.gov/nuccore/XM_004269502.2) | [XP 004269550.1](https://www.ncbi.nlm.nih.gov/protein/466007654) | ✓ | ✓ |
| 177 | *Orcinus* | *orca* | Mammalia | CASP9 | [101271677](https://www.ncbi.nlm.nih.gov/sites/entrez?db=gene&cmd=Retrieve&dopt=full_report&list_uids=101271677) | [XM 004272425.2](https://www.ncbi.nlm.nih.gov/nuccore/XM_004272425.2) | [XP 004272473.1](https://www.ncbi.nlm.nih.gov/protein/466022285) | ✓ | ✓ |
|  |  |  |  | CYCS | [101283052](https://www.ncbi.nlm.nih.gov/sites/entrez?db=gene&cmd=Retrieve&dopt=full_report&list_uids=101283052) | [XM 004265648.2](https://www.ncbi.nlm.nih.gov/nuccore/XM_004265648.2) | [XP 004265696.1](https://www.ncbi.nlm.nih.gov/nuccore/XP_004265696.1) | ✓ | ✓ |
|  |  |  |  | APAF-1 | [103202343](https://www.ncbi.nlm.nih.gov/sites/entrez?db=gene&cmd=Retrieve&dopt=full_report&list_uids=103202343) | [XM 007947213.1](https://www.ncbi.nlm.nih.gov/nuccore/XM_007947213.1) | [XP 007945404.1](https://www.ncbi.nlm.nih.gov/protein/634865983) | ✓ | ✓ |
| 178 | *Orycteropus* | *afer afer* | Mammalia | CASP9 | [103205454](https://www.ncbi.nlm.nih.gov/sites/entrez?db=gene&cmd=Retrieve&dopt=full_report&list_uids=103205454) | [XM 007950760.1](https://www.ncbi.nlm.nih.gov/nuccore/XM_007950760.1) | [XP 007948951.1](https://www.ncbi.nlm.nih.gov/protein/634875904) | ✓ | ✓ |
|  |  |  |  | CYCS | [103201153](https://www.ncbi.nlm.nih.gov/sites/entrez?db=gene&cmd=Retrieve&dopt=full_report&list_uids=103201153) | [XM 007945786.1](https://www.ncbi.nlm.nih.gov/nuccore/XM_007945786.1) | [XP 007943977.1](https://www.ncbi.nlm.nih.gov/protein/634824746) | ✓ | ✓ |
|  |  |  |  | APAF-1 | [100135699](https://www.ncbi.nlm.nih.gov/sites/entrez?db=gene&cmd=Retrieve&dopt=full_report&list_uids=100135699) | [XM 012174846.2](https://www.ncbi.nlm.nih.gov/nuccore/XM_012174846.2) | [XP 012030236.2](https://www.ncbi.nlm.nih.gov/nuccore/XP_012030236.2) | ✓ | ✓ |
| 179 | *Ovis* | *aries* | Mammalia | CASP9 | [101110953](https://www.ncbi.nlm.nih.gov/sites/entrez?db=gene&cmd=Retrieve&dopt=full_report&list_uids=101110953) | [XM 015099300.1](https://www.ncbi.nlm.nih.gov/nuccore/XM_015099300.1) | [XP 014954786.1](https://www.ncbi.nlm.nih.gov/nuccore/XP_014954786.1) | ✓ | ✓ |
|  |  |  |  | CYCS | [106990092](https://www.ncbi.nlm.nih.gov/sites/entrez?db=gene&cmd=Retrieve&dopt=full_report&list_uids=106990092) | [XM 004007950.2](https://www.ncbi.nlm.nih.gov/nuccore/XM_004007950.2) | [XP 004007999.1](https://www.ncbi.nlm.nih.gov/protein/426227792) | ✓ | ✓ |
|  |  |  |  | APAF-1 | [452153](https://www.ncbi.nlm.nih.gov/sites/entrez?db=gene&cmd=Retrieve&dopt=full_report&list_uids=452153) | [XM 003313880.3](https://www.ncbi.nlm.nih.gov/nuccore/XM_003313880.3) | [XP 003313928.1](https://www.ncbi.nlm.nih.gov/protein/332840131) | ✓ | ✓ |
| 180 | *Pan* | *troglodytes* | Mammalia | CASP9 | [456457](https://www.ncbi.nlm.nih.gov/sites/entrez?db=gene&cmd=Retrieve&dopt=full_report&list_uids=456457) | [XM 009448820.2](https://www.ncbi.nlm.nih.gov/nuccore/XM_009448820.2) | [XP 009447095.1](https://www.ncbi.nlm.nih.gov/nuccore/XP_009447095.1) | ✓ | ✓ |
|  |  |  |  | CYCS | [744779](https://www.ncbi.nlm.nih.gov/sites/entrez?db=gene&cmd=Retrieve&dopt=full_report&list_uids=744779) | [NM 001071821.1](https://www.ncbi.nlm.nih.gov/nuccore/NM_001071821.1) | [NP 001065289.1](https://www.ncbi.nlm.nih.gov/protein/115392119) | ✓ | ✓ |
|  |  |  |  | APAF-1 | [100995524](https://www.ncbi.nlm.nih.gov/sites/entrez?db=gene&cmd=Retrieve&dopt=full_report&list_uids=100995524) | [XM 003832607.2](https://www.ncbi.nlm.nih.gov/nuccore/XM_003832607.2) | [XP 003832655.1](https://www.ncbi.nlm.nih.gov/protein/397525393) | ✓ | ✓ |
| 181 | *Pan* | *paniscus* | Mammalia | CASP9 | [100992567](https://www.ncbi.nlm.nih.gov/sites/entrez?db=gene&cmd=Retrieve&dopt=full_report&list_uids=100992567) | [XM 003806257.2](https://www.ncbi.nlm.nih.gov/nuccore/XM_003806257.2) | [XP 003806305.1](https://www.ncbi.nlm.nih.gov/protein/397469313) | ✓ | ✓ |
|  |  |  |  | CYCS | [100986481](https://www.ncbi.nlm.nih.gov/sites/entrez?db=gene&cmd=Retrieve&dopt=full_report&list_uids=100986481) | [XM 003807922.2](https://www.ncbi.nlm.nih.gov/nuccore/XM_003807922.2) | [XP 003807970.1](https://www.ncbi.nlm.nih.gov/protein/397472902) | ✓ | ✓ |
|  |  |  |  | APAF-1 | [109265383](https://www.ncbi.nlm.nih.gov/sites/entrez?db=gene&cmd=Retrieve&dopt=full_report&list_uids=109265383) | [XM 019446520.1](https://www.ncbi.nlm.nih.gov/nuccore/XM_019446520.1) | [XP 019302065.1](https://www.ncbi.nlm.nih.gov/protein/XP_019302065.1) | ✓ | ✓ |
| 182 | *Panthera* | *pardus* | Mammalia | CASP9 | [109274153](https://www.ncbi.nlm.nih.gov/sites/entrez?db=gene&cmd=Retrieve&dopt=full_report&list_uids=109274153) | [XM 019461464.1](https://www.ncbi.nlm.nih.gov/nuccore/XM_019461464.1) | [XP 019317009.1,](https://www.ncbi.nlm.nih.gov/nuccore/XP_019317009.1) | ✓ | ✓ |
|  |  |  |  | CYCS | [109263846](https://www.ncbi.nlm.nih.gov/sites/entrez?db=gene&cmd=Retrieve&dopt=full_report&list_uids=109263846) | [XM 019443636.1](https://www.ncbi.nlm.nih.gov/nuccore/XM_019443636.1) | [XP 019299181.1](https://www.ncbi.nlm.nih.gov/protein/1111106096) | ✓ | ✓ |
|  |  |  |  | APAF-1 | [107204470](https://www.ncbi.nlm.nih.gov/sites/entrez?db=gene&cmd=Retrieve&dopt=full_report&list_uids=107204470) | [XM 015627640.2](https://www.ncbi.nlm.nih.gov/nuccore/XM_015627640.2) | [XP 015483126.1](https://www.ncbi.nlm.nih.gov/nuccore/XP_015483126.1) | ✓ | ✓ |
| 183 | *Parus* | *major* | Aves | CASP9 | [107213687](https://www.ncbi.nlm.nih.gov/sites/entrez?db=gene&cmd=Retrieve&dopt=full_report&list_uids=107213687) | [XM 015648388.2](https://www.ncbi.nlm.nih.gov/nuccore/XM_015648388.2) | [XP 015503874.1](https://www.ncbi.nlm.nih.gov/protein/998731362) | ✓ | ✓ |
|  |  |  |  | CYCS | [107200910](https://www.ncbi.nlm.nih.gov/sites/entrez?db=gene&cmd=Retrieve&dopt=full_report&list_uids=107200910) | [XM 015620015.2](https://www.ncbi.nlm.nih.gov/nuccore/XM_015620015.2) | [XP 015475501.1](https://www.ncbi.nlm.nih.gov/protein/998675914) | ✓ | ✓ |
|  |  |  |  | APAF-1 | [102451576](https://www.ncbi.nlm.nih.gov/sites/entrez?db=gene&cmd=Retrieve&dopt=full_report&list_uids=102451576) | [XM 006118275.2](https://www.ncbi.nlm.nih.gov/nuccore/XM_006118275.2) | [XP 006118337.1](https://www.ncbi.nlm.nih.gov/protein/558140177) | ✓ | ✓ |
| 184 | *Pelodiscus* | *sinensis* | Reptilia | CASP9 | [102443524](https://www.ncbi.nlm.nih.gov/sites/entrez?db=gene&cmd=Retrieve&dopt=full_report&list_uids=102443524) | [XM 006132727.2](https://www.ncbi.nlm.nih.gov/nuccore/XM_006132727.2) | [XP 006132789.1](https://www.ncbi.nlm.nih.gov/protein/558206762) | ✓ | ✓ |
|  |  |  |  | CYCS | [102455941](https://www.ncbi.nlm.nih.gov/sites/entrez?db=gene&cmd=Retrieve&dopt=full_report&list_uids=102455941) | [XM 006138376.1](https://www.ncbi.nlm.nih.gov/nuccore/XM_006138376.1) | [XP 006138438.1](https://www.ncbi.nlm.nih.gov/protein/558229181) | ✓ | ✓ |
|  |  |  |  | APAF-1 | [102905376](https://www.ncbi.nlm.nih.gov/sites/entrez?db=gene&cmd=Retrieve&dopt=full_report&list_uids=102905376) | [XM 006979814.2](https://www.ncbi.nlm.nih.gov/nuccore/XM_006979814.2) | [XP 006979876.1](https://www.ncbi.nlm.nih.gov/nuccore/XP_006979876.1) | ✓ | ✓ |
| 185 | *Peromyscus* | *maniculatus bairdii* | Mammalia | CASP9 | [102906175](https://www.ncbi.nlm.nih.gov/sites/entrez?db=gene&cmd=Retrieve&dopt=full_report&list_uids=102906175) | [XM 015994341.1](https://www.ncbi.nlm.nih.gov/nuccore/XM_015994341.1) | [XP 015849827.1](https://www.ncbi.nlm.nih.gov/protein/1008747714) | ✓ | ✓ |
|  |  |  |  | CYCS | [102913936](https://www.ncbi.nlm.nih.gov/sites/entrez?db=gene&cmd=Retrieve&dopt=full_report&list_uids=102913936) | [XM 006982189.2](https://www.ncbi.nlm.nih.gov/nuccore/XM_006982189.2) | [XP 006982251.1](https://www.ncbi.nlm.nih.gov/nuccore/XP_006982251.1) | ✓ | ✓ |
|  |  |  |  | APAF-1 | [104308028](https://www.ncbi.nlm.nih.gov/sites/entrez?db=gene&cmd=Retrieve&dopt=full_report&list_uids=104308028) | XM 009909153.1 | XP 009907455.1 | ✓ | ✓ |
| 186 | *Picoides* | *pubescens* | Aves | CASP9 | [104305903](https://www.ncbi.nlm.nih.gov/sites/entrez?db=gene&cmd=Retrieve&dopt=full_report&list_uids=104305903) | [XM 009906822.1](https://www.ncbi.nlm.nih.gov/nuccore/XM_009906822.1) | [XP 009905124.1](https://www.ncbi.nlm.nih.gov/protein/699691418) | ✓ | ✓ |
|  |  |  |  | CYCS | [104307617](https://www.ncbi.nlm.nih.gov/sites/entrez?db=gene&cmd=Retrieve&dopt=full_report&list_uids=104307617) | [XM 009908700.1](https://www.ncbi.nlm.nih.gov/nuccore/XM_009908700.1) | [XP 009907002.1](https://www.ncbi.nlm.nih.gov/protein/699699906) | ✓ | ✓ |
|  |  |  |  | APAF-1 | [110084192](https://www.ncbi.nlm.nih.gov/sites/entrez?db=gene&cmd=Retrieve&dopt=full_report&list_uids=110084192) | [XM 020803318.1](https://www.ncbi.nlm.nih.gov/nuccore/XM_020803318.1) | [XP 020658977.1](https://www.ncbi.nlm.nih.gov/nuccore/XP_020658977.1) | ✓ | ✓ |
| 187 | *Pogona* | *vitticeps* | Reptilia | CASP9 | [110072690](https://www.ncbi.nlm.nih.gov/sites/entrez?db=gene&cmd=Retrieve&dopt=full_report&list_uids=110072690) | [XM 020781235.1](https://www.ncbi.nlm.nih.gov/nuccore/XM_020781235.1) | [XP 020636894.1](https://www.ncbi.nlm.nih.gov/nuccore/XP_020636894.1) | ✓ | ✓ |
|  |  |  |  | CYCS | [110072471](https://www.ncbi.nlm.nih.gov/sites/entrez?db=gene&cmd=Retrieve&dopt=full_report&list_uids=110072471) | [XM 020780885.1](https://www.ncbi.nlm.nih.gov/nuccore/XM_020780885.1) | [XP 020636544.1](https://www.ncbi.nlm.nih.gov/nuccore/XP_020636544.1) | ✓ | ✓ |
|  |  |  |  | APAF-1 | [100459074](https://www.ncbi.nlm.nih.gov/sites/entrez?db=gene&cmd=Retrieve&dopt=full_report&list_uids=100459074) | [XM 002823623.3](https://www.ncbi.nlm.nih.gov/nuccore/XM_002823623.3) | [XP 002823669.1](https://www.ncbi.nlm.nih.gov/protein/297692687) | ✓ | ✓ |
| 188 | *Pongo* | *abelii* | Mammalia | CASP9 | [100460459](https://www.ncbi.nlm.nih.gov/sites/entrez?db=gene&cmd=Retrieve&dopt=full_report&list_uids=100460459) | [XM 002811445.3](https://www.ncbi.nlm.nih.gov/nuccore/XM_002811445.3) | [XP 002811491.1](https://www.ncbi.nlm.nih.gov/protein/297666354) | ✓ | ✓ |
|  |  |  |  | CYCS | [100171479](https://www.ncbi.nlm.nih.gov/sites/entrez?db=gene&cmd=Retrieve&dopt=full_report&list_uids=100171479) | [NM 001131167.2](https://www.ncbi.nlm.nih.gov/nuccore/NM_001131167.2) | [NP 001124639.1](https://www.ncbi.nlm.nih.gov/protein/197099570) | ✓ | ✓ |
|  |  |  |  | APAF-1 | [105826075](https://www.ncbi.nlm.nih.gov/sites/entrez?db=gene&cmd=Retrieve&dopt=full_report&list_uids=105826075) | [XM 012663987.1](https://www.ncbi.nlm.nih.gov/nuccore/XM_012663987.1) | [XP 012519441.1](https://www.ncbi.nlm.nih.gov/protein/826349396) | ✓ | ✓ |
| 189 | *Propithecus* | *coquereli* | Mammalia | CASP9 | [105825008](https://www.ncbi.nlm.nih.gov/sites/entrez?db=gene&cmd=Retrieve&dopt=full_report&list_uids=105825008) | [XM 012662665.1](https://www.ncbi.nlm.nih.gov/nuccore/XM_012662665.1) | [XP 012518119.1](https://www.ncbi.nlm.nih.gov/protein/826345589) | ✓ | ✓ |
|  |  |  |  | CYCS | [105807375](https://www.ncbi.nlm.nih.gov/sites/entrez?db=gene&cmd=Retrieve&dopt=full_report&list_uids=105807375) | [XM 012640754.1](https://www.ncbi.nlm.nih.gov/nuccore/XM_012640754.1) | [XP 012496208.1](https://www.ncbi.nlm.nih.gov/protein/826283600) | ✓ | ✓ |
|  |  |  |  | APAF-1 | [107290550](https://www.ncbi.nlm.nih.gov/sites/entrez?db=gene&cmd=Retrieve&dopt=full_report&list_uids=107290550) | [XM 015819437.1](https://www.ncbi.nlm.nih.gov/nuccore/XM_015819437.1) | [XP 015674923.1](https://www.ncbi.nlm.nih.gov/protein/1002592278) | ✓ | ✓ |
| 190 | *Protobothrops* | *mucrosquamatus* | Reptilia | CASP9 | [107291376](https://www.ncbi.nlm.nih.gov/sites/entrez?db=gene&cmd=Retrieve&dopt=full_report&list_uids=107291376) | [XM 015820373.1](https://www.ncbi.nlm.nih.gov/nuccore/XM_015820373.1) | [XP 015675859.1](https://www.ncbi.nlm.nih.gov/protein/1002593976) | ✓ | ✓ |
|  |  |  |  | CYCS | [107282765](https://www.ncbi.nlm.nih.gov/sites/entrez?db=gene&cmd=Retrieve&dopt=full_report&list_uids=107282765) | [XM 015810583.1](https://www.ncbi.nlm.nih.gov/nuccore/XM_015810583.1) | [XP 015666069.1](https://www.ncbi.nlm.nih.gov/protein/1002574085) | ✓ | ✓ |
|  |  |  |  | APAF-1 | [102103178](https://www.ncbi.nlm.nih.gov/sites/entrez?db=gene&cmd=Retrieve&dopt=full_report&list_uids=102103178) | [XM 005519454.2](https://www.ncbi.nlm.nih.gov/nuccore/XM_005519454.2) | [XP 005519511.1](https://www.ncbi.nlm.nih.gov/protein/543348200) | ✓ | ✓ |
| 191 | *Pseudopodoces* | *humilis* | Aves | CASP9 | [102108950](https://www.ncbi.nlm.nih.gov/sites/entrez?db=gene&cmd=Retrieve&dopt=full_report&list_uids=102108950) | [XM 014257210.1](https://www.ncbi.nlm.nih.gov/nuccore/XM_014257210.1) | [XP 014112685.1](https://www.ncbi.nlm.nih.gov/nuccore/XP_014112685.1) | ✓ | ✓ |
|  |  |  |  | CYCS | [102113796](https://www.ncbi.nlm.nih.gov/sites/entrez?db=gene&cmd=Retrieve&dopt=full_report&list_uids=102113796) | [XM 014248787.1](https://www.ncbi.nlm.nih.gov/nuccore/XM_014248787.1) | [XP 014104262.1](https://www.ncbi.nlm.nih.gov/protein/929418400) | ✓ | ✓ |
|  |  |  |  | APAF-1 | [102897687](https://www.ncbi.nlm.nih.gov/sites/entrez?db=gene&cmd=Retrieve&dopt=full_report&list_uids=102897687) | [XM 006914569.2](https://www.ncbi.nlm.nih.gov/nuccore/XM_006914569.2) | [XP 006914631.2](https://www.ncbi.nlm.nih.gov/protein/989963459) | ✓ | ✓ |
| 192 | *Pteropus* | *alecto* | Mammalia | CASP9 | [102878565](https://www.ncbi.nlm.nih.gov/sites/entrez?db=gene&cmd=Retrieve&dopt=full_report&list_uids=102878565) | [XM 006914334.2](https://www.ncbi.nlm.nih.gov/nuccore/XM_006914334.2) | [XP 006914396.1](https://www.ncbi.nlm.nih.gov/protein/586562277) | ✓ | ✓ |
|  |  |  |  | CYCS | [102879159](https://www.ncbi.nlm.nih.gov/sites/entrez?db=gene&cmd=Retrieve&dopt=full_report&list_uids=102879159) | [XM 006912003.2](https://www.ncbi.nlm.nih.gov/nuccore/XM_006912003.2) | [XP 006912065.1](https://www.ncbi.nlm.nih.gov/nuccore/XP_006912065.1) | ✓ | ✓ |
|  |  |  |  | APAF-1 | [102199079](https://www.ncbi.nlm.nih.gov/sites/entrez?db=gene&cmd=Retrieve&dopt=full_report&list_uids=102199079) | [XM 005733502.1](https://www.ncbi.nlm.nih.gov/nuccore/XM_005733502.1) | [XP 005733559.1](https://www.ncbi.nlm.nih.gov/protein/548376665) | ✓ | ✓ |
| 193 | *Pundamilia* | *nyererei* | Actinopterygii | CASP9 | [102210622](https://www.ncbi.nlm.nih.gov/sites/entrez?db=gene&cmd=Retrieve&dopt=full_report&list_uids=102210622) | [XM 005730723.1](https://www.ncbi.nlm.nih.gov/nuccore/XM_005730723.1) | [XP 005730780.1](https://www.ncbi.nlm.nih.gov/nuccore/XP_005730780.1) | ✓ | ✓ |
|  |  |  |  | CYCS | [102195352](https://www.ncbi.nlm.nih.gov/sites/entrez?db=gene&cmd=Retrieve&dopt=full_report&list_uids=102195352) | [XM 005723515.2](https://www.ncbi.nlm.nih.gov/nuccore/XM_005723515.2) | [XP 005723572.1](https://www.ncbi.nlm.nih.gov/nuccore/XP_005723572.1) | ✓ | ✓ |
|  |  |  |  | APAF-1 | [108438498](https://www.ncbi.nlm.nih.gov/sites/entrez?db=gene&cmd=Retrieve&dopt=full_report&list_uids=108438498) | [XM 017716343.1](https://www.ncbi.nlm.nih.gov/nuccore/XM_017716343.1) | [XP 017571832.1](https://www.ncbi.nlm.nih.gov/nuccore/XP_017571832.1) | ✓ | ✓ |
| 194 | *Pygocentrus* | *nattereri* | Actinopterygii | CASP9 | [108425555](https://www.ncbi.nlm.nih.gov/sites/entrez?db=gene&cmd=Retrieve&dopt=full_report&list_uids=108425555) | [XM 017694276.1](https://www.ncbi.nlm.nih.gov/nuccore/XM_017694276.1) | [XP 017549765.1](https://www.ncbi.nlm.nih.gov/nuccore/XP_017549765.1) | ✓ | ✓ |
|  |  |  |  | CYCS | [108430847](https://www.ncbi.nlm.nih.gov/sites/entrez?db=gene&cmd=Retrieve&dopt=full_report&list_uids=108430847) | [XM 017703637.1](https://www.ncbi.nlm.nih.gov/nuccore/XM_017703637.1) | [XP 017559126.1](https://www.ncbi.nlm.nih.gov/protein/1049240461) | ✓ | ✓ |
|  |  |  |  | APAF-1 | [103057709](https://www.ncbi.nlm.nih.gov/sites/entrez?db=gene&cmd=Retrieve&dopt=full_report&list_uids=103057709) | [XM 007420190.2](https://www.ncbi.nlm.nih.gov/nuccore/XM_007420190.2) | [XP 007420252.1](https://www.ncbi.nlm.nih.gov/protein/602626596) | ✓ | ✓ |
| 195 | *Python* | *bivittatus* | Reptilia | CASP9 | [103063403](https://www.ncbi.nlm.nih.gov/sites/entrez?db=gene&cmd=Retrieve&dopt=full_report&list_uids=103063403) | [XM 015891586.1](https://www.ncbi.nlm.nih.gov/nuccore/XM_015891586.1) | [XP 015747072.1](https://www.ncbi.nlm.nih.gov/protein/1004664451) | ✓ | ✓ |
|  |  |  |  | CYCS | [103054542](https://www.ncbi.nlm.nih.gov/sites/entrez?db=gene&cmd=Retrieve&dopt=full_report&list_uids=103054542) | [XM 007420820.2](https://www.ncbi.nlm.nih.gov/nuccore/XM_007420820.2) | [XP 007420882.1](https://www.ncbi.nlm.nih.gov/protein/602627881) | ✓ | ✓ |
|  |  |  |  | APAF-1 | [78963](https://www.ncbi.nlm.nih.gov/sites/entrez?db=gene&cmd=Retrieve&dopt=full_report&list_uids=78963) | [NM 023979.1](https://www.ncbi.nlm.nih.gov/nuccore/NM_023979.1) | [NP 076469.1](https://www.ncbi.nlm.nih.gov/protein/13027436) | ✓ | ✓ |
| 196 | *Rattus* | *norvegicus* | Mammalia | CASP9 | [58918](https://www.ncbi.nlm.nih.gov/sites/entrez?db=gene&cmd=Retrieve&dopt=full_report&list_uids=58918) | [NM 031632.1](https://www.ncbi.nlm.nih.gov/nuccore/NM_031632.1) | [NP 113820.1](https://www.ncbi.nlm.nih.gov/protein/13928872) | ✓ | ✓ |
|  |  |  |  | CYCS | [25309](https://www.ncbi.nlm.nih.gov/sites/entrez?db=gene&cmd=Retrieve&dopt=full_report&list_uids=25309) | [NM 012839.2](https://www.ncbi.nlm.nih.gov/nuccore/NM_012839.2) | [NP 036971.1](https://www.ncbi.nlm.nih.gov/protein/6978725) | ✓ | ✓ |
|  |  |  |  | APAF-1 | [109447611](https://www.ncbi.nlm.nih.gov/sites/entrez?db=gene&cmd=Retrieve&dopt=full_report&list_uids=109447611) | [XM 019732133.1](https://www.ncbi.nlm.nih.gov/nuccore/XM_019732133.1) | [XP 019587692.1](https://www.ncbi.nlm.nih.gov/nuccore/XP_019587692.1) | ✓ | ✓ |
| 197 | *Rhinolophus* | *sinicus* | Mammalia | CASP9 | [109459000](https://www.ncbi.nlm.nih.gov/sites/entrez?db=gene&cmd=Retrieve&dopt=full_report&list_uids=109459000) | [XM 019753131.1](https://www.ncbi.nlm.nih.gov/nuccore/XM_019753131.1) | [XP 019608690.1](https://www.ncbi.nlm.nih.gov/nuccore/XP_019608690.1) | ✓ | ✓ |
|  |  |  |  | CYCS | [109444615](https://www.ncbi.nlm.nih.gov/sites/entrez?db=gene&cmd=Retrieve&dopt=full_report&list_uids=109444615) | [XM 019726126.1](https://www.ncbi.nlm.nih.gov/nuccore/XM_019726126.1) | [XP 019581685.1](https://www.ncbi.nlm.nih.gov/protein/1123935603) | ✓ | ✓ |
|  |  |  |  | APAF-1 | [104656342](https://www.ncbi.nlm.nih.gov/sites/entrez?db=gene&cmd=Retrieve&dopt=full_report&list_uids=104656342) | [XM 010355937.1](https://www.ncbi.nlm.nih.gov/nuccore/XM_010355937.1) | [XP 010354239.1](https://www.ncbi.nlm.nih.gov/protein/724960160) | ✓ | ✓ |
| 198 | *Rhinopithecus* | *roxellana* | Mammalia | CASP9 | [104677696](https://www.ncbi.nlm.nih.gov/sites/entrez?db=gene&cmd=Retrieve&dopt=full_report&list_uids=104677696) | [XM 010382817.1](https://www.ncbi.nlm.nih.gov/nuccore/XM_010382817.1) | [XP 010381119.1](https://www.ncbi.nlm.nih.gov/protein/724919485) | ✓ | ✓ |
|  |  |  |  | CYCS | [104672188](https://www.ncbi.nlm.nih.gov/sites/entrez?db=gene&cmd=Retrieve&dopt=full_report&list_uids=104672188) | [XM 010375926.1](https://www.ncbi.nlm.nih.gov/nuccore/XM_010375926.1) | [XP 010374228.1](https://www.ncbi.nlm.nih.gov/protein/724887305) | ✓ | ✓ |
|  |  |  |  | APAF-1 | [107517482](https://www.ncbi.nlm.nih.gov/sites/entrez?db=gene&cmd=Retrieve&dopt=full_report&list_uids=107517482) | [XM 016158232.1](https://www.ncbi.nlm.nih.gov/nuccore/XM_016158232.1) | [XP 016013718.1](https://www.ncbi.nlm.nih.gov/protein/XP_016013718.1) | ✓ | ✓ |
| 199 | *Rousettus* | *aegyptiacus* | Mammalia | CASP9 | [107501830](https://www.ncbi.nlm.nih.gov/sites/entrez?db=gene&cmd=Retrieve&dopt=full_report&list_uids=107501830) | [XM 016128385.1](https://www.ncbi.nlm.nih.gov/nuccore/XM_016128385.1) | [XP 015983871.1](https://www.ncbi.nlm.nih.gov/protein/XP_015983871.1) | ✓ | ✓ |
|  |  |  |  | CYCS | [107518128](https://www.ncbi.nlm.nih.gov/sites/entrez?db=gene&cmd=Retrieve&dopt=full_report&list_uids=107518128) | [XM 016159536.1](https://www.ncbi.nlm.nih.gov/nuccore/XM_016159536.1) | [XP 016015022.1](https://www.ncbi.nlm.nih.gov/protein/1012209953) | ✓ | ✓ |
|  |  |  |  | APAF-1 | [103813383](https://www.ncbi.nlm.nih.gov/sites/entrez?db=gene&cmd=Retrieve&dopt=full_report&list_uids=103813383) | [XM 009086639.2](https://www.ncbi.nlm.nih.gov/nuccore/XM_009086639.2) | [XP 009084887.1](https://www.ncbi.nlm.nih.gov/protein/683902598) | ✓ | ✓ |
| 200 | *Serinus* | *canaria* | Aves | CASP9 | [103824350](https://www.ncbi.nlm.nih.gov/sites/entrez?db=gene&cmd=Retrieve&dopt=full_report&list_uids=103824350) | [XM 018924683.1](https://www.ncbi.nlm.nih.gov/nuccore/XM_018924683.1) | [XP 018780228.1](https://www.ncbi.nlm.nih.gov/nuccore/XP_018780228.1) | ✓ | ✓ |
|  |  |  |  | CYCS | [103812888](https://www.ncbi.nlm.nih.gov/sites/entrez?db=gene&cmd=Retrieve&dopt=full_report&list_uids=103812888) | [XM 009086055.2](https://www.ncbi.nlm.nih.gov/nuccore/XM_009086055.2) | [XP 009084303.1](https://www.ncbi.nlm.nih.gov/protein/683900983) | ✓ | ✓ |
|  |  |  |  | APAF-1 | [101537432](https://www.ncbi.nlm.nih.gov/sites/entrez?db=gene&cmd=Retrieve&dopt=full_report&list_uids=101537432) | [XM 004602843.1](https://www.ncbi.nlm.nih.gov/nuccore/XM_004602843.1) | [XP 004602900.1](https://www.ncbi.nlm.nih.gov/protein/505777861) | ✓ | ✓ |
| 201 | *Sorex* | *araneus* | Mammalia | CASP9 | [101537803](https://www.ncbi.nlm.nih.gov/sites/entrez?db=gene&cmd=Retrieve&dopt=full_report&list_uids=101537803) | [XM 004603653.1](https://www.ncbi.nlm.nih.gov/nuccore/XM_004603653.1) | [XP 004603710.1](https://www.ncbi.nlm.nih.gov/protein/505781110) | ✓ | ✓ |
|  |  |  |  | CYCS | [101558408](https://www.ncbi.nlm.nih.gov/sites/entrez?db=gene&cmd=Retrieve&dopt=full_report&list_uids=101558408) | [XM 004604069.1](https://www.ncbi.nlm.nih.gov/nuccore/XM_004604069.1) | [XP 004604126.1](https://www.ncbi.nlm.nih.gov/nuccore/XP_004604126.1) | ✓ | ✓ |
|  |  |  |  | APAF-1 | [104143811](https://www.ncbi.nlm.nih.gov/sites/entrez?db=gene&cmd=Retrieve&dopt=full_report&list_uids=104143811) | [XM 009674519.1](https://www.ncbi.nlm.nih.gov/nuccore/XM_009674519.1) | [XP 009672814.1](https://www.ncbi.nlm.nih.gov/protein/697481956) | ✓ | ✓ |
| 202 | *Struthio* | *camelus australis* | Aves | CASP9 | [104153653](https://www.ncbi.nlm.nih.gov/sites/entrez?db=gene&cmd=Retrieve&dopt=full_report&list_uids=104153653) | [XM 009689617.1](https://www.ncbi.nlm.nih.gov/nuccore/XM_009689617.1) | [XP 009687912.1](https://www.ncbi.nlm.nih.gov/protein/697437121) | ✓ | ✓ |
|  |  |  |  | CYCS | [104145105](https://www.ncbi.nlm.nih.gov/sites/entrez?db=gene&cmd=Retrieve&dopt=full_report&list_uids=104145105) | [XM 009676445.1](https://www.ncbi.nlm.nih.gov/nuccore/XM_009676445.1) | [XP 009674740.1](https://www.ncbi.nlm.nih.gov/nuccore/XP_009674740.1) | ✓ | ✓ |
|  |  |  |  | APAF-1 | [106861312](https://www.ncbi.nlm.nih.gov/sites/entrez?db=gene&cmd=Retrieve&dopt=full_report&list_uids=106861312) | [XM 014889612.1](https://www.ncbi.nlm.nih.gov/nuccore/XM_014889612.1) | [XP 014745098.1](https://www.ncbi.nlm.nih.gov/protein/959040583) | ✓ | ✓ |
| 203 | *Sturnus* | *vulgaris* | Aves | CASP9 | [106856421](https://www.ncbi.nlm.nih.gov/sites/entrez?db=gene&cmd=Retrieve&dopt=full_report&list_uids=106856421) | [XM 014881727.1](https://www.ncbi.nlm.nih.gov/nuccore/XM_014881727.1) | [XP 014737213.1](https://www.ncbi.nlm.nih.gov/protein/959067856) | ✓ | ✓ |
|  |  |  |  | CYCS | [106864714](https://www.ncbi.nlm.nih.gov/sites/entrez?db=gene&cmd=Retrieve&dopt=full_report&list_uids=106864714) | [XM 014894321.1](https://www.ncbi.nlm.nih.gov/nuccore/XM_014894321.1) | [XP 014749807.1](https://www.ncbi.nlm.nih.gov/protein/959041540) | ✓ | ✓ |
|  |  |  |  | APAF-1 | [100513283](https://www.ncbi.nlm.nih.gov/sites/entrez?db=gene&cmd=Retrieve&dopt=full_report&list_uids=100513283) | [XM 003481742.4](https://www.ncbi.nlm.nih.gov/nuccore/XM_003481742.4) | [XP 003481790.2](https://www.ncbi.nlm.nih.gov/nuccore/XP_003481790.2) | ✓ | ✓ |
| 204 | *Sus* | *scrofa* | Mammalia | CASP9 | [100518913](https://www.ncbi.nlm.nih.gov/sites/entrez?db=gene&cmd=Retrieve&dopt=full_report&list_uids=100518913) | [XM 003127618.4](https://www.ncbi.nlm.nih.gov/nuccore/XM_003127618.4) | [XP 003127666.1](https://www.ncbi.nlm.nih.gov/nuccore/XP_003127666.1) | ✓ | ✓ |
|  |  |  |  | CYCS | [100170131](https://www.ncbi.nlm.nih.gov/sites/entrez?db=gene&cmd=Retrieve&dopt=full_report&list_uids=100170131) | [NM 001129970.1](https://www.ncbi.nlm.nih.gov/nuccore/NM_001129970.1) | [NP 001123442.1](https://www.ncbi.nlm.nih.gov/protein/194018698) | ✓ | ✓ |
|  |  |  |  | APAF-1 | [100222477](https://www.ncbi.nlm.nih.gov/sites/entrez?db=gene&cmd=Retrieve&dopt=full_report&list_uids=100222477) | [XM 002190092.2](https://www.ncbi.nlm.nih.gov/nuccore/XM_002190092.2) | [XP 002190128.1](https://www.ncbi.nlm.nih.gov/protein/224094450) | ✓ | ✓ |
| 205 | *Taeniopygia* | *guttata* | Aves | CASP9 | [100227981](https://www.ncbi.nlm.nih.gov/sites/entrez?db=gene&cmd=Retrieve&dopt=full_report&list_uids=100227981) | [XM 002190655.3](https://www.ncbi.nlm.nih.gov/nuccore/XM_002190655.3) | [XP 002190691.2](https://www.ncbi.nlm.nih.gov/protein/823475502) | ✓ | ✓ |
|  |  |  |  | CYCS | [100223555](https://www.ncbi.nlm.nih.gov/sites/entrez?db=gene&cmd=Retrieve&dopt=full_report&list_uids=100223555) | [NM 001143673.2](https://www.ncbi.nlm.nih.gov/nuccore/NM_001143673.2) | [NP 001137145.2](https://www.ncbi.nlm.nih.gov/protein/379642588) | ✓ | ✓ |
|  |  |  |  | APAF-1 | [103251017](https://www.ncbi.nlm.nih.gov/sites/entrez?db=gene&cmd=Retrieve&dopt=full_report&list_uids=103251017) | [XM 008049613.1](https://www.ncbi.nlm.nih.gov/nuccore/XM_008049613.1) | [XP 008047804.1](https://www.ncbi.nlm.nih.gov/protein/XP_008047804.1) | ✓ | ✓ |
| 206 | *Tarsius* | *syrichta* | Mammalia | CASP9 | [103260483](https://www.ncbi.nlm.nih.gov/sites/entrez?db=gene&cmd=Retrieve&dopt=full_report&list_uids=103260483) | [XM 021712373.1](https://www.ncbi.nlm.nih.gov/nuccore/XM_021712373.1) | [XP 021568048.1](https://www.ncbi.nlm.nih.gov/nuccore/XP_021568048.1) | ✓ | ✓ |
|  |  |  |  | CYCS | [103274909](https://www.ncbi.nlm.nih.gov/sites/entrez?db=gene&cmd=Retrieve&dopt=full_report&list_uids=103274909) | [XM 008072323.1](https://www.ncbi.nlm.nih.gov/nuccore/XM_008072323.1) | [XP 008070514.1](https://www.ncbi.nlm.nih.gov/protein/640826701) | ✓ | ✓ |
|  |  |  |  | APAF-1 | [106547522](https://www.ncbi.nlm.nih.gov/sites/entrez?db=gene&cmd=Retrieve&dopt=full_report&list_uids=106547522) | [XM 014064734.1](https://www.ncbi.nlm.nih.gov/nuccore/XM_014064734.1) | [XP 013920209.1](https://www.ncbi.nlm.nih.gov/protein/927163363) | ✓ | ✓ |
| 207 | *Thamnophis* | *sirtalis* | Reptilia | CASP9 | [106547434](https://www.ncbi.nlm.nih.gov/sites/entrez?db=gene&cmd=Retrieve&dopt=full_report&list_uids=106547434) | [XM 014064609.1](https://www.ncbi.nlm.nih.gov/nuccore/XM_014064609.1) | [XP 013920084.1](https://www.ncbi.nlm.nih.gov/protein/927162973) | ✓ | ✓ |
|  |  |  |  | CYCS | [106539539](https://www.ncbi.nlm.nih.gov/sites/entrez?db=gene&cmd=Retrieve&dopt=full_report&list_uids=106539539) | [XM 014054339.1](https://www.ncbi.nlm.nih.gov/nuccore/XM_014054339.1) | [XP 013909814.1](https://www.ncbi.nlm.nih.gov/nuccore/XP_013909814.1) | ✓ | ✓ |
|  |  |  |  | APAF-1 | [104573948](https://www.ncbi.nlm.nih.gov/sites/entrez?db=gene&cmd=Retrieve&dopt=full_report&list_uids=104573948) | [XM 010221185.1](https://www.ncbi.nlm.nih.gov/nuccore/XM_010221185.1) | [XP 010219487.1](https://www.ncbi.nlm.nih.gov/protein/719771850) | ✓ | ✓ |
| 208 | *Tinamus* | *guttatus* | Aves | CASP9 | [104580482](https://www.ncbi.nlm.nih.gov/sites/entrez?db=gene&cmd=Retrieve&dopt=full_report&list_uids=104580482) | [XM 010227991.1](https://www.ncbi.nlm.nih.gov/nuccore/XM_010227991.1) | [XP 010226293.1](https://www.ncbi.nlm.nih.gov/protein/719737471) | ✓ | ✓ |
|  |  |  |  | CYCS | [104573152](https://www.ncbi.nlm.nih.gov/sites/entrez?db=gene&cmd=Retrieve&dopt=full_report&list_uids=104573152) | [XM 010220333.1](https://www.ncbi.nlm.nih.gov/nuccore/XM_010220333.1) | [XP 010218635.1](https://www.ncbi.nlm.nih.gov/nuccore/XP_010218635.1) | ✓ | ✓ |
|  |  |  |  | APAF-1 | [101349393](https://www.ncbi.nlm.nih.gov/sites/entrez?db=gene&cmd=Retrieve&dopt=full_report&list_uids=101349393) | [XM 004381254.1](https://www.ncbi.nlm.nih.gov/nuccore/XM_004381254.1) | [XP 004381311.1](https://www.ncbi.nlm.nih.gov/protein/XP_004381311.1) | ✓ | ✓ |
| 209 | *Trichechus* | *manatus latirostris* | Mammalia | CASP9 | [101355103](https://www.ncbi.nlm.nih.gov/sites/entrez?db=gene&cmd=Retrieve&dopt=full_report&list_uids=101355103) | [XM 004377285.1](https://www.ncbi.nlm.nih.gov/nuccore/XM_004377285.1) | [XP 004377342.1](https://www.ncbi.nlm.nih.gov/protein/XP_004377342.1) | ✓ | ✓ |
|  |  |  |  | CYCS | [101340375](https://www.ncbi.nlm.nih.gov/sites/entrez?db=gene&cmd=Retrieve&dopt=full_report&list_uids=101340375) | [XM 004377468.1](https://www.ncbi.nlm.nih.gov/nuccore/XM_004377468.1) | [XP 004377525.1](https://www.ncbi.nlm.nih.gov/protein/471379455) | ✓ | ✓ |
|  |  |  |  | APAF-1 | [100127738](https://www.ncbi.nlm.nih.gov/sites/entrez?db=gene&cmd=Retrieve&dopt=full_report&list_uids=100127738) | [XM 012959843.2](https://www.ncbi.nlm.nih.gov/nuccore/XM_012959843.2) | [XP 012815297.1](https://www.ncbi.nlm.nih.gov/protein/847113621) | ✓ | ✓ |
| 210 | *Xenopus* | *tropicalis* | Amphibia | CASP9 | [100144708](https://www.ncbi.nlm.nih.gov/sites/entrez?db=gene&cmd=Retrieve&dopt=full_report&list_uids=100144708) | [NM 001123463.1](https://www.ncbi.nlm.nih.gov/nuccore/NM_001123463.1) | [NP 001116935.1](https://www.ncbi.nlm.nih.gov/protein/183986693) | ✓ | ✓ |
|  |  |  |  | CYCS | [394490](https://www.ncbi.nlm.nih.gov/sites/entrez?db=gene&cmd=Retrieve&dopt=full_report&list_uids=394490) | [NM 203564.1](https://www.ncbi.nlm.nih.gov/nuccore/NM_203564.1) | [NP 988895.1](https://www.ncbi.nlm.nih.gov/protein/45360505) | ✓ | ✓ |
|  |  |  |  | APAF-1 | [102069090](https://www.ncbi.nlm.nih.gov/sites/entrez?db=gene&cmd=Retrieve&dopt=full_report&list_uids=102069090) | [XM 005480708.2](https://www.ncbi.nlm.nih.gov/nuccore/XM_005480708.2) | [XP 005480765.1](https://www.ncbi.nlm.nih.gov/protein/542143364) | ✓ | ✓ |
| 211 | *Zonotrichia* | *albicollis* | Aves | CASP9 | [102060592](https://www.ncbi.nlm.nih.gov/sites/entrez?db=gene&cmd=Retrieve&dopt=full_report&list_uids=102060592) | [XM 014273303.1](https://www.ncbi.nlm.nih.gov/nuccore/XM_014273303.1) | [XP 014128778.1](https://www.ncbi.nlm.nih.gov/protein/929518067) | ✓ | ✓ |
|  |  |  |  | CYCS | [102069140](https://www.ncbi.nlm.nih.gov/sites/entrez?db=gene&cmd=Retrieve&dopt=full_report&list_uids=102069140) | [XM 005480872.2](https://www.ncbi.nlm.nih.gov/nuccore/XM_005480872.2) | [XP 005480929.1](https://www.ncbi.nlm.nih.gov/protein/542143711) | ✓ | ✓ |
