## Supplementary file 2-Ortho-Profile results for APAF-1 and CASP9 for "Protein coevolution and physicochemical adaptation in the APAF-1/apoptosome: structural and functional implications"

| **APAF-1 - *H. sapiens* (NP_863651.1)** |  |  |  |  |
| --- | --- | --- | --- | --- |
| **1st phase** |  |  |  |  |
| **Database: refseq_protein** |  |  |  |  |
| **Sequence against sequence- Blastp RBH** |  |  |  |  |
|  | **% coverage** | **e-value** | **% identity** | **access** |
| **Cnidaria** |  |  |  |  |
| ***A. millepora*** |  |  |  |  |
| Foward blast | 99 | 0 | 36 | AJG37574.1 |
| Reverse blast | 91 | 0 | 36 | NP_863651.1 |
| ***H. vulgaris*** |  |  |  |  |
| Foward blast | 100 | 0 | 32 | XP_012566478.1 |
| Reverse blast | 99 | 0 | 32 | NP_863651.1 |
| ***O. faveolata*** |  |  |  |  |
| Foward blast | 98 | 0 | 33 | XP_020620792.1 |
| Reverse blast | 92 | 0 | 33 | NP_863651.1 |
| **Echinodermata** |  |  |  |  |
| ***S. purpuratus*** |  |  |  |  |
| Foward blast | 99 | 0 | 35 | XP_011682983.1 |
| Reverse blast | 97 | 0 | 35 | NP_863651.1 |
| **Hemichordata** |  |  |  |  |
| ***S. kowalevskii*** |  |  |  |  |
| Foward blast | 100 | 0 | 38 | XP_006818297.1 |
| Reverse blast | 98 | 0 | 38 | NP_863651.1 |
| **Cephalochordata** |  |  |  |  |
| ***B. floridae*** |  |  |  |  |
| Foward blast | 100 | 0 | 38 | XP_002598365.1 |
| Reverse blast | 99 | 0 | 37 | NP_863651.1 |
| **APAF-1 - *H. sapiens* (NP_863651.1)** |  |  |  |  |
| **2nd phase** |  |  |  |  |
| **Database: nr** |  |  |  |  |
| **Profile against sequence - PSI - BLAST RBH** |  |  |  |  |
|  | **% coverage** | **e-value** | **% identity** | **access** |
| **Porifera** |  |  |  |  |
| ***A. queenslandica*** |  |  |  |  |
| Foward blast | 99 | 0 | 25 | XP_019855714.1 |
| Reverse blast | 91 | 0 | 25 | NP_863651.1 |
| **Protostomia** |  |  |  |  |
| ***A. transitella*** |  |  |  |  |
| Foward blast | 98 | 0 | 18 | XP_013193147.1 |
| Reverse blast | 51 | 0 | 25 | NP_863651.1 |
| ***A. rosae*** |  |  |  |  |
| Foward blast | 97 | 0 | 20 | XP_012264834.1 |
| Reverse blast | 88 | 0 | 21 | NP_863651.1 |
| ***D. magna*** |  |  |  |  |
| Foward blast | 99 | 0 | 25 | KZS20735.1 |
| Reverse blast | 99 | 0 | 25 | NP_863651.1 |
| ***D. pulex*** |  |  |  |  |
| Foward blast | 99 | 0 | 25 | EFX74325.1 |
| Reverse blast | 99 | 0 | 24 | NP_863651.1 |
| ***N. lecontei*** |  |  |  |  |
| Foward blast | 99 | 0 | 18 | XP_015511747.1 |
| Reverse blast | 87 | 0 | 19 | NP_863651.1 |
| ***O. biroi*** |  |  |  |  |
| Foward blast | 96 | 0 | 20 | XP_011342858.1 |
| Reverse blast | 78 | 0 | 21 | NP_863651.1 |
| ***O. cincta*** |  |  |  |  |
| Foward blast | 99 | 0 | 22 | ODN02648.1 |
| Reverse blast | 99 | 0 | 22 | NP_863651.1 |
| ***P. tepidariorum*** |  |  |  |  |
| Foward blast | 99 | 0 | 25 | XP_015920891.1 |
| Reverse blast | 99 | 0 | 25 | NP_863651.1 |
| ***T. cornetzi*** |  |  |  |  |
| Foward blast | 94 | 0 | 21 | XP_018369153.1 |
| Reverse blast | 91 | 0 | 20 | NP_863651.1 |
| ***T. nativa*** |  |  |  |  |
| Foward blast | 99 | 0 | 23 | KRZ60231.1* |
| Reverse blast | 80 | 0 | 22 | NP_863651.1 |
| ***T. papuae*** |  |  |  |  |
| Foward blast | 99 | 0 | 22 | KRZ71961.1 |
| Reverse blast | 81 | 0 | 23 | NP_863651.1 |
| ***T. zimbabwensis*** |  |  |  |  |
| Foward blast | 99 | 0 | 23 | KRZ19336.1* |
| Reverse blast | 81 | 0 | 23 | NP_863651.1 |
| ***T. pretiosum*** |  |  |  |  |
| Foward blast | 96 | 0 | 19 | XP_014234781.1 |
| Reverse blast | 98 | 0 | 21 | NP_863651.1 |
| **CASP9 - *H. sapiens* (NP_001220.2)** |  |  |  |  |
| **3rd phase** |  |  |  |  |
| **Database: Proteomes reference** |  |  |  |  |
| **based on profile HMM** |  |  |  |  |
|  | **ID UniProtKB** | **e-value** | **GenBank ID** |  |
| **Protostomia** |  |  |  |  |
| ***L. anatina*** |  |  |  |  |
| Foward search | A0A1S3HNC3 | 0.00059 | XP_013387550.1* |  |
| Reverse search | P55211 | 0.00016 | NP_001220.2 |  |
| **Echinodermata** |  |  |  |  |
| ***S. purpuratus*** |  |  |  |  |
| Foward search | W4YD85 | 0.00012 | XP_791872.3* |  |
| Reverse search | P55211 | 0.00013 | NP_001220.2 |  |

* The possible ortholog is different from the one detected in the search for the homologues.
