## Supplementary material for "Protein coevolution and physicochemical adaptation in the APAF-1/apoptosome: structural and functional implications": Figure 1-figure supplement 1-Non reconciliated APAF-1 phylogeny showing proteins nearby and within the human clade

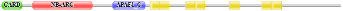

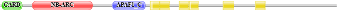

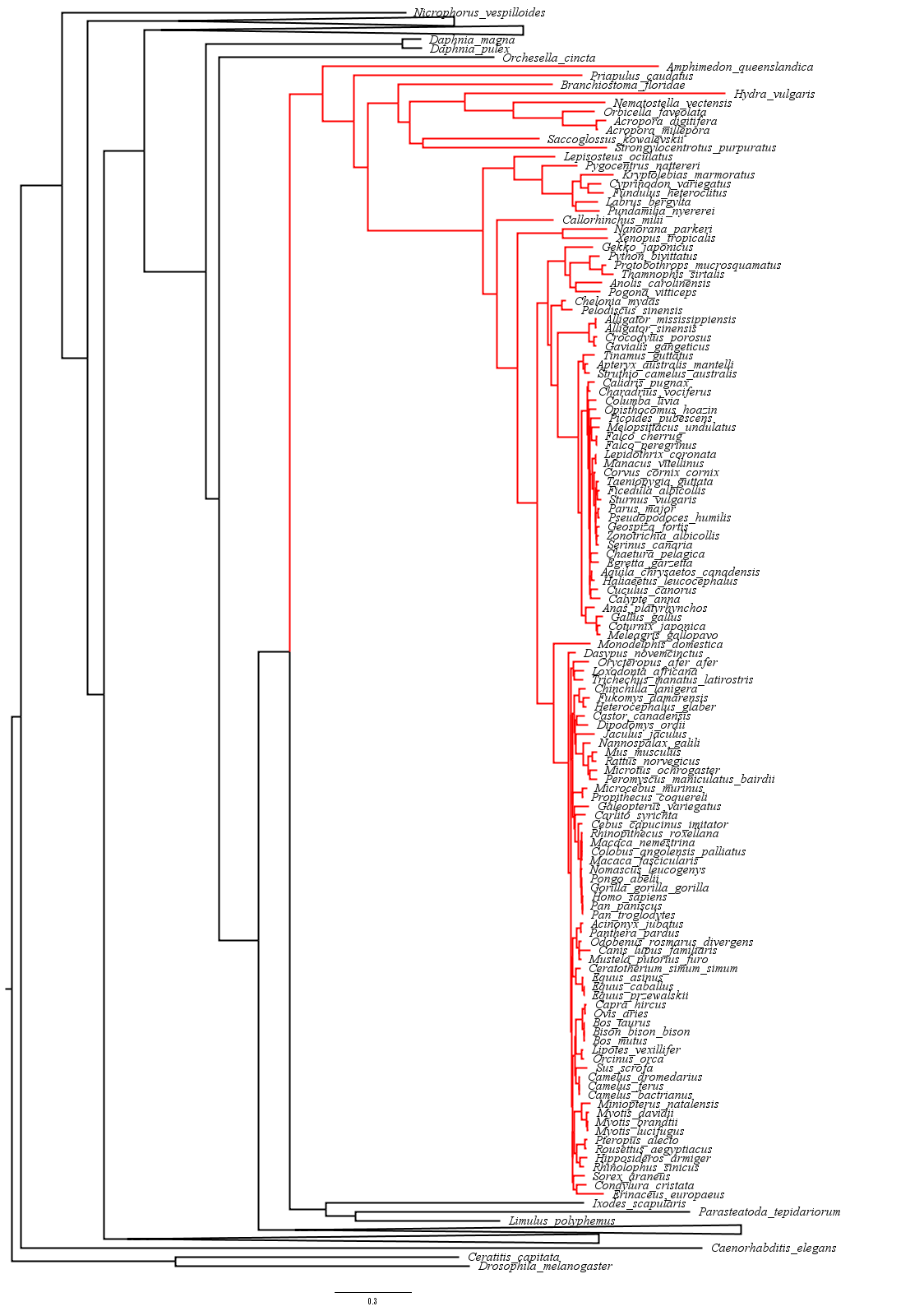

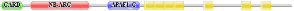

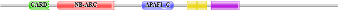

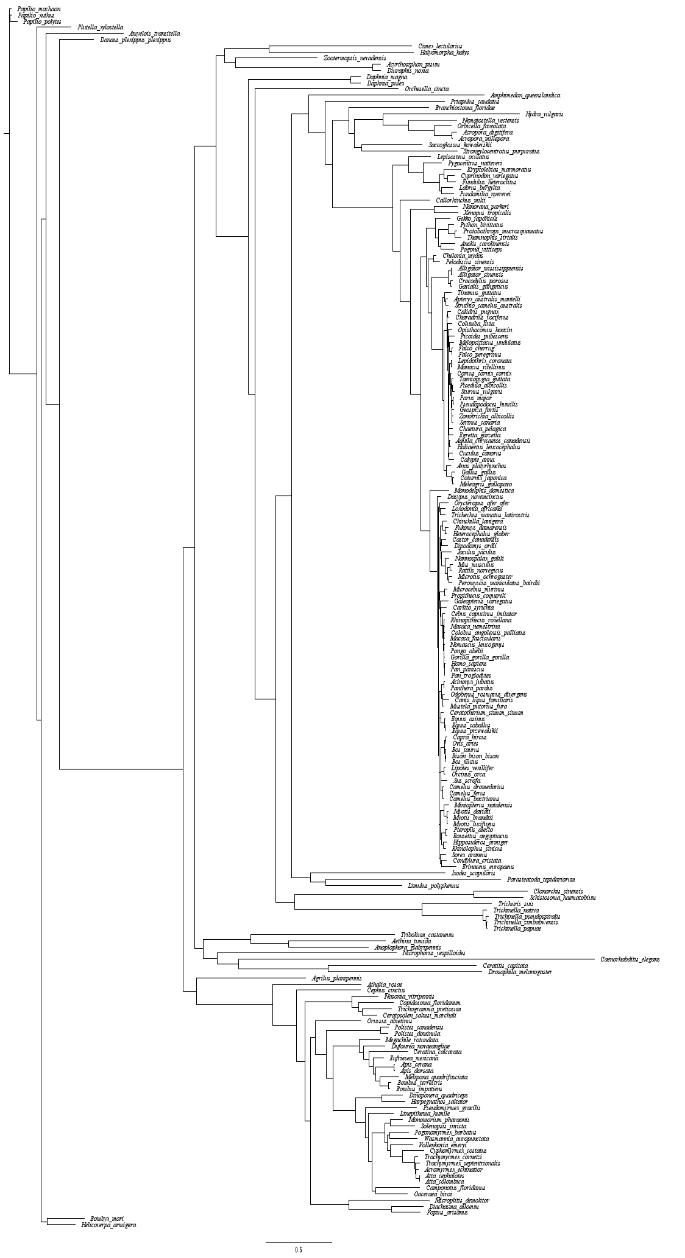

**CARD**

**NB-ARC**

**HELICAL DOMAINS**

**WD40**

**ANAPC4_WD40**

**Utp8**

**PIG-U**

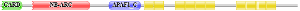

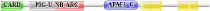

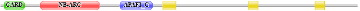

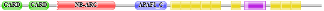

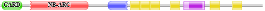

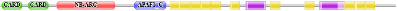

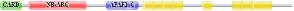

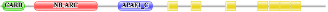

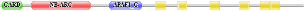

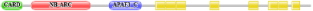

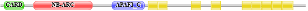

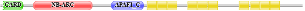

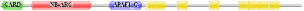

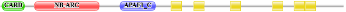

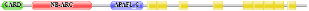

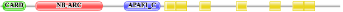

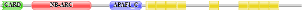

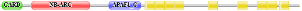

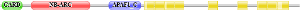

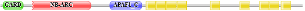

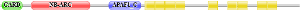

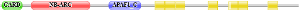

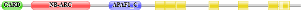

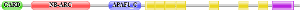

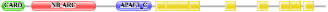

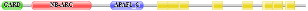

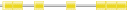

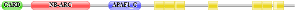

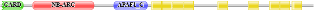

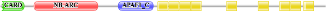

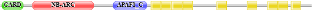

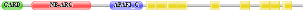

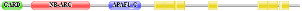

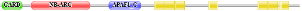

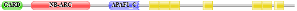

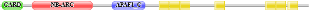

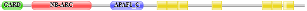

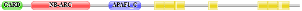

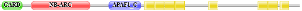

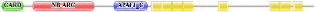

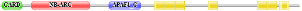

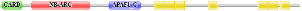

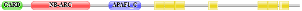

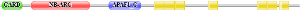

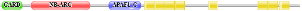

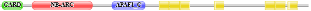

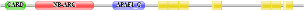

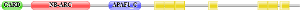

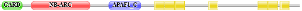

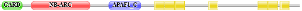

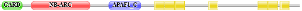

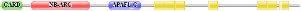

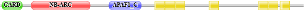

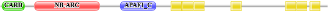

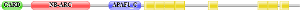

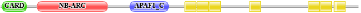

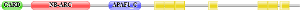

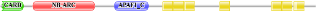

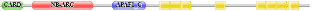

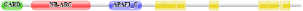

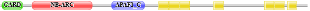

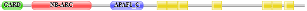

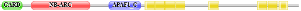

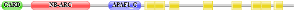

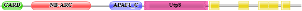

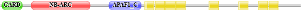

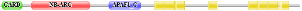

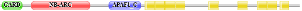

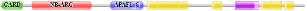

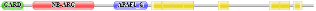

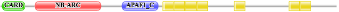

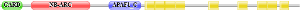

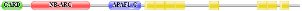

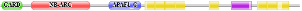

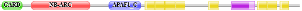

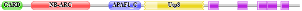

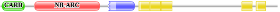

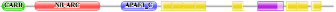

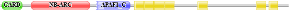

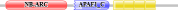

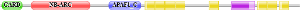

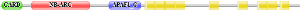

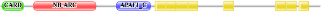

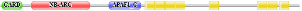

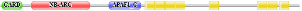

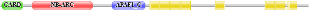

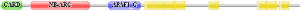

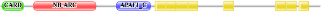

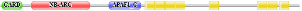

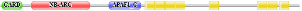

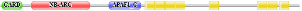

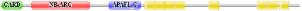

**Human**

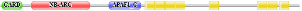

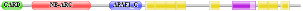

**Figure 1-figure supplement 1. Non reconciliated APAF-1 phylogeny showing proteins nearby and within the human clade.** Sequences within the human clade are in red. Arthropods except insects originate in nodes close to vertebrates. The distant nodes are collapsed. The architecture of the protein domains obtained in Pfam is exposed.
