## Supplementary material for "Protein coevolution and physicochemical adaptation in the APAF-1/apoptosome: structural and functional implications": Figure 2-figure supplement 1-Non reconciliated APAF-1 phylogeny showing the relationship between insect sequences

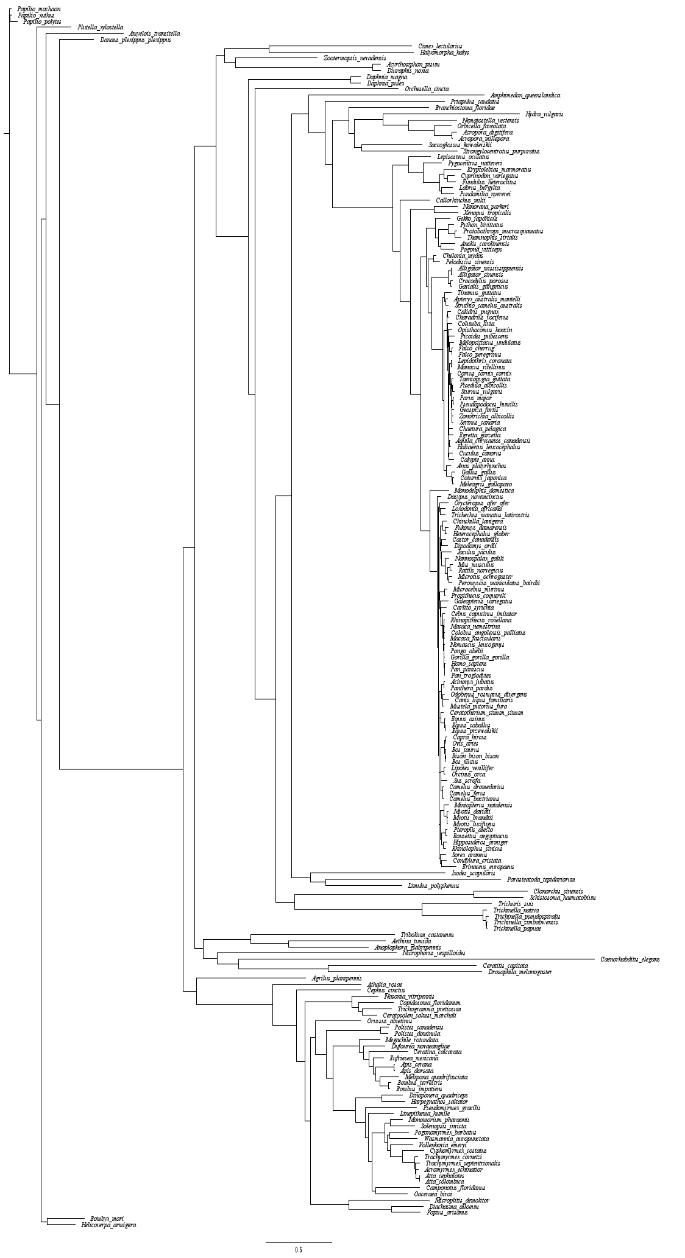

**Nup160**

**HELICAL DOMAINS**

**ANAPC4_WD40**

**WD40**

**NB-ARC**

**CARD**

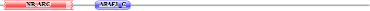

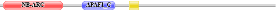

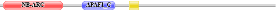

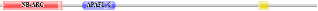

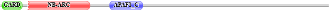

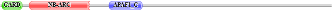

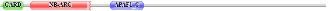

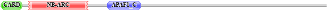

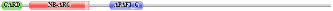

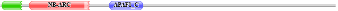

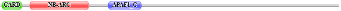

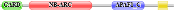

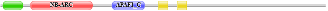

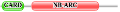

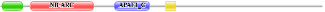

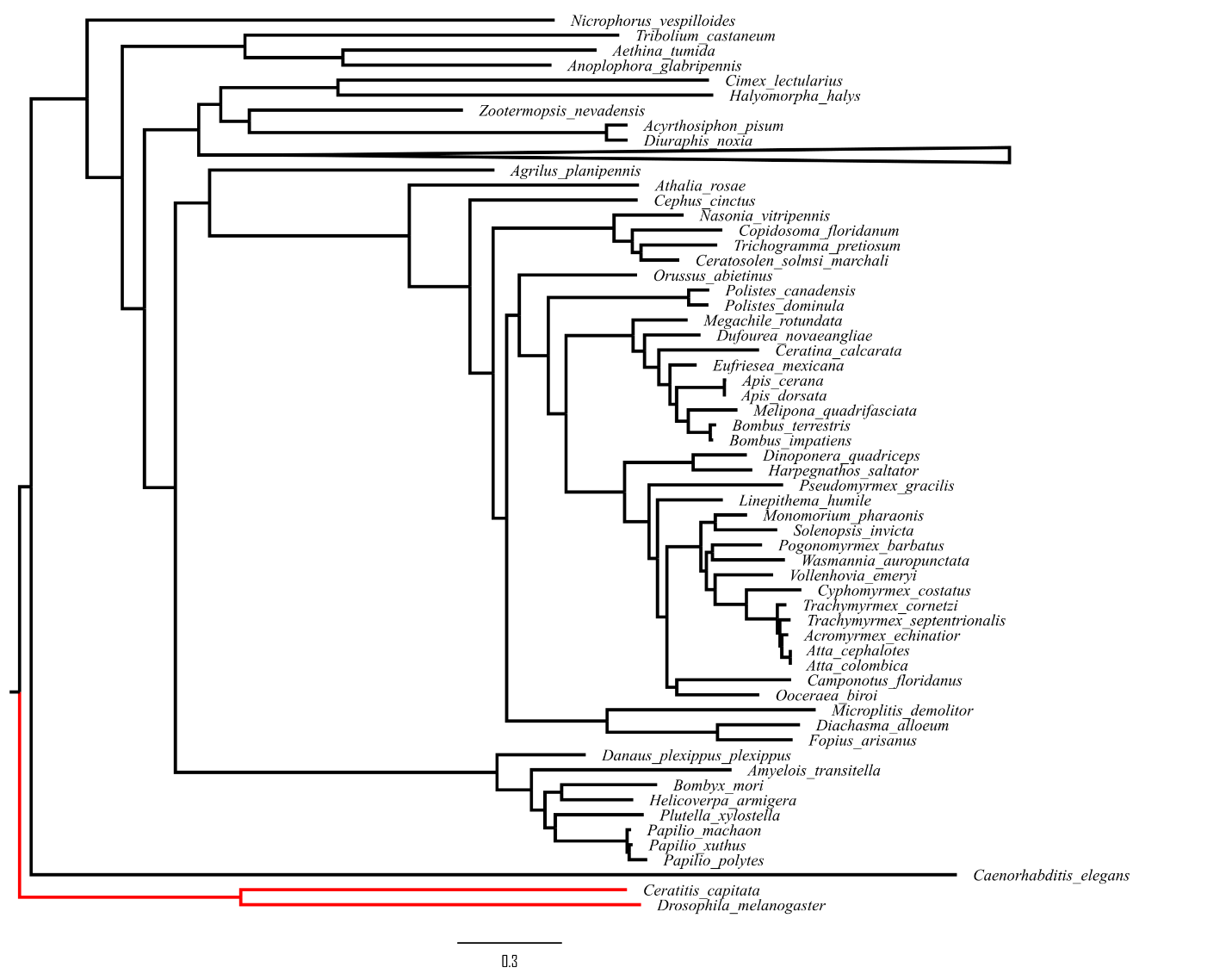

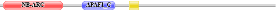

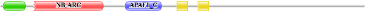

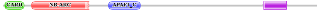

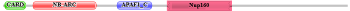

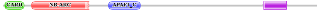

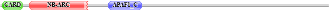

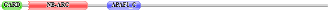

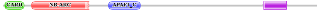

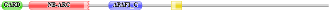

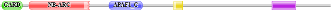

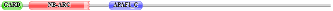

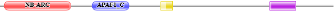

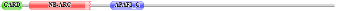

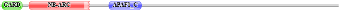

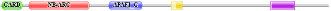

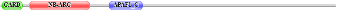

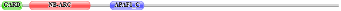

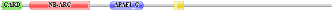

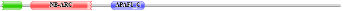

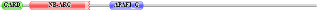

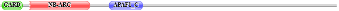

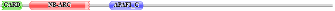

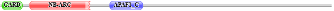

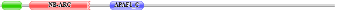

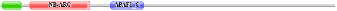

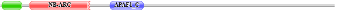

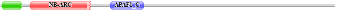

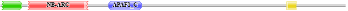

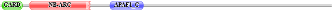

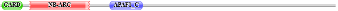

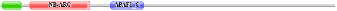

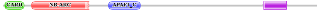

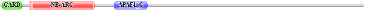

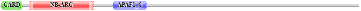

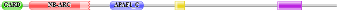

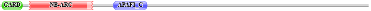

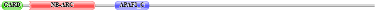

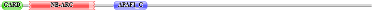

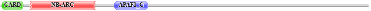

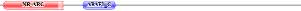

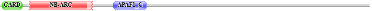

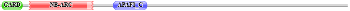

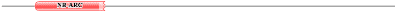

**Figure 2-figure supplement 1. Non reconciliated APAF-1 phylogeny showing the relationship between insect sequences.** The red nodes show that the Diptera sequences including *D. melanogaster* form a separate clade. The proteins of the other insects originate in different and more distant nodes. For a better visualization the nodes of the other taxa are collapsed. For each species the architecture of the protein domains obtained in Pfam is exposed.
