## Supplementary material for "Protein coevolution and physicochemical adaptation in the APAF-1/apoptosome: structural and functional implications": Figure 3-figure supplement 1-Non reconciliated APAF-1 phylogeny showing the relationship between nematode sequences

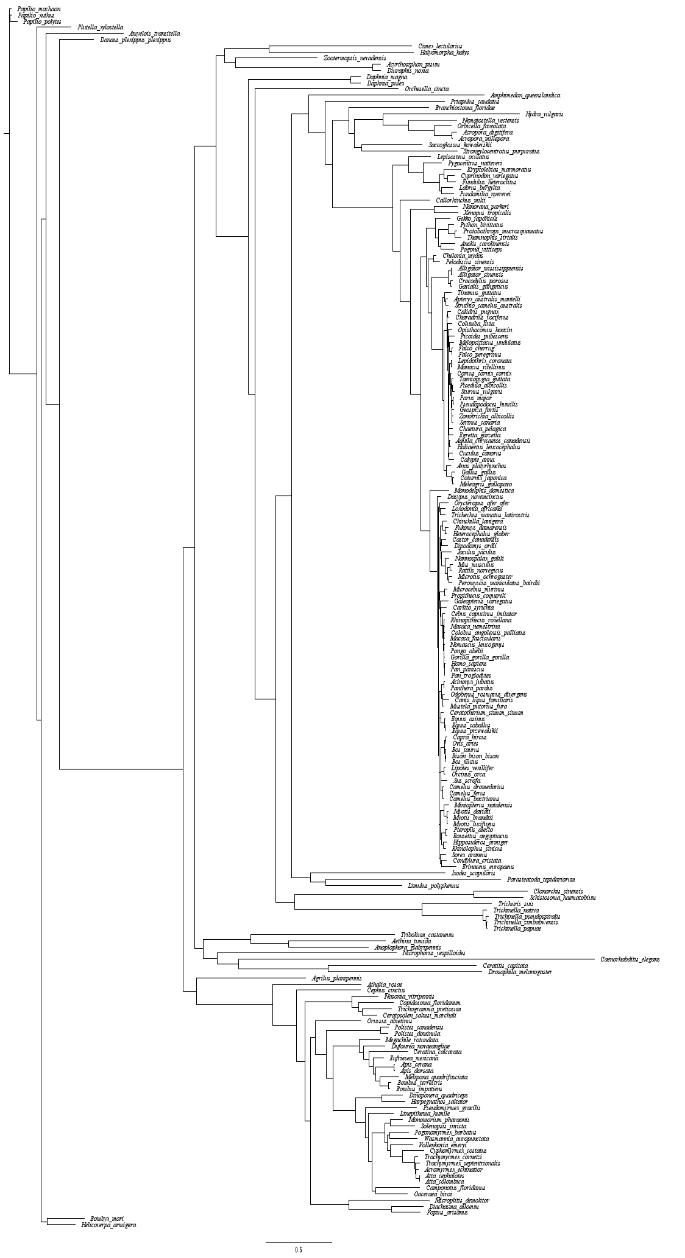

**WD40**

**NB-ARC**

**HELICAL DOMAINS**

**CARD**

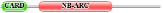

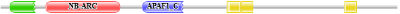

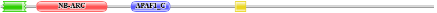

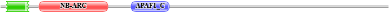

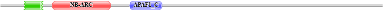

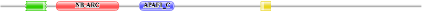

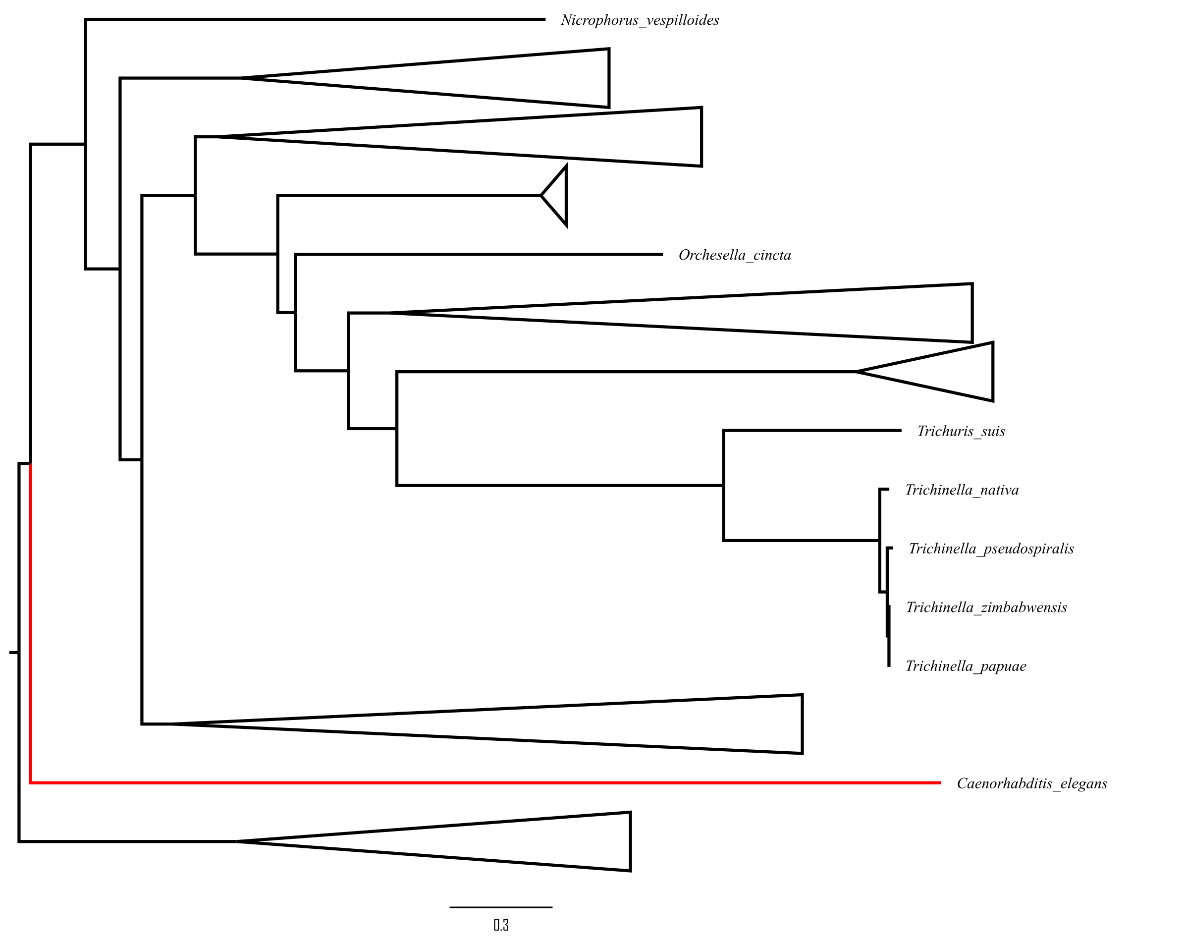

**Figure 3-figure supplement 1. Non reconciliated APAF-1 phylogeny showing the relationship between nematode sequences.** The red nodes show that the *C. elegans* sequence forms a separate clade. The proteins of the other nematodes originate in a different and more distant node. For a better visualization the nodes of the other taxa are collapsed. For each species the architecture of the protein domains is exposed. Unlike *C. elegans*, Pfam weakly detected WD40 repeats that form the β propellers at the carboxyl terminal of proteins in *T. suis*, *T. nativa* and *T. papuae*.
