## Supplementary material for "Protein coevolution and physicochemical adaptation in the APAF-1/apoptosome: structural and functional implications": Figure 3-figure supplement 2-Analysis of conserved domains obtained in the NCBI of homologues to APAF-1CED-4 in Nematoda and Plathelminthes

*C. elegans*

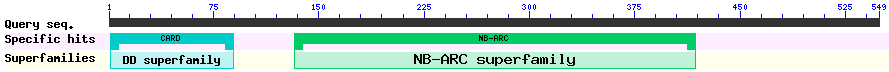

*T. suis*

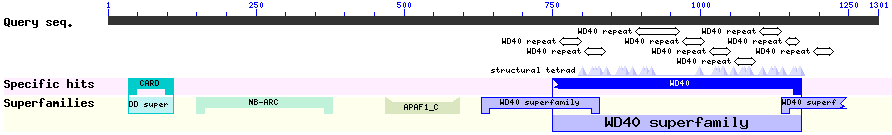

*T. nativa*

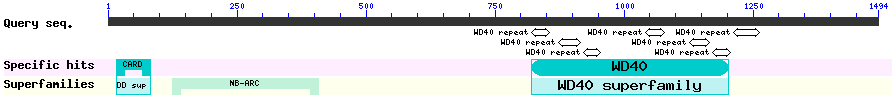

*T. papuae*

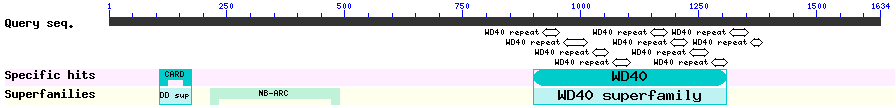

*T. pseudospiralis*

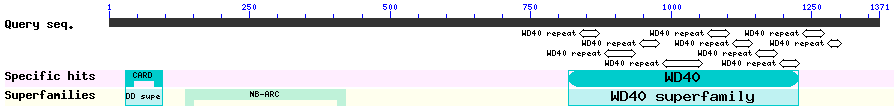

*T. zimbabwensis*

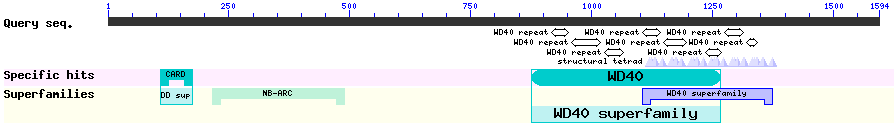

*C. sinensis*

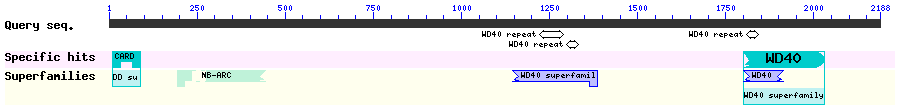

*S. haematobium*

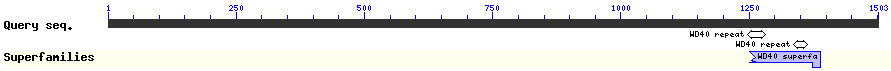

**Figure 3-figure supplement 2. Analysis of conserved domains obtained in the NCBI of homologues to APAF-1/CED-4 in Nematoda and** **Plathelminthes.** It is shown that unlike *C. elegans*, the genera *Trichinella* and *Trichuris* and the flatworms *C. sinensis* and *S. haematobium* possess the WD40 repeats that form the β propellers at the carboxyl terminal.
