## Supplementary material for "Protein coevolution and physicochemical adaptation in the APAF-1/apoptosome: structural and functional implications": Figure 4-figure supplement 1-Non reconciliated CASP9 phylogeny

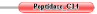

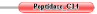

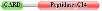

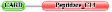

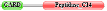

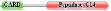

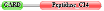

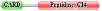

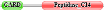

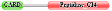

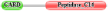

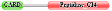

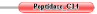

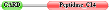

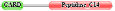

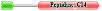

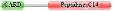

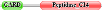

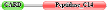

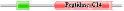

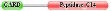

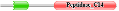

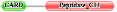

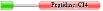

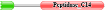

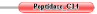

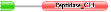

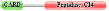

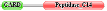

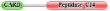

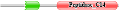

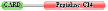

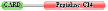

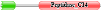

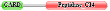

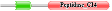

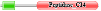

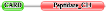

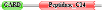

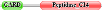

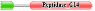

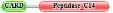

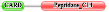

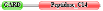

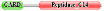

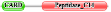

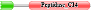

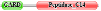

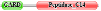

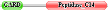

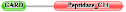

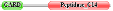

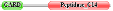

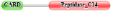

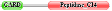

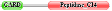

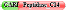

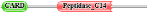

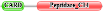

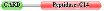

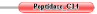

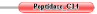

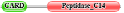

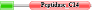

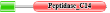

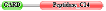

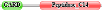

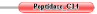

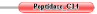

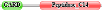

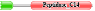

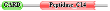

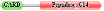

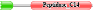

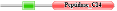

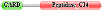

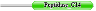

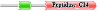

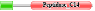

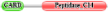

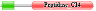

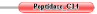

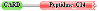

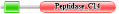

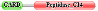

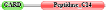

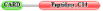

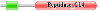

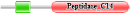

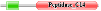

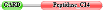

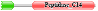

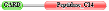

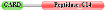

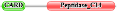

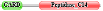

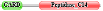

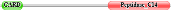

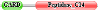

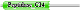

**Human**

**p20/p10**

**CARD**

**Figure 4-figure supplement 1. Non reconciliated CASP9 phylogeny.** A unique red node is observed for *Drosophila*. The other red nodes below indicate the sequences that are within the same clade as the human protein. The other taxa arise from different nodes. For each species the architecture of the protein domains obtained in Pfam is exposed.
