## Supplementary material for "Protein coevolution and physicochemical adaptation in the APAF-1/apoptosome: structural and functional implications": Suplementary file 3-Output file BIS2Analyzer with alignment and hits associated to dimensions and blocks: 8) Suplementary file 3-Output file BIS2Analyzer with alignment and hits associated to dimensions and blocks.html

Alignment and BIS clusters

|  |  |  |  |  |  |  |  |  |  |  |  |  |  |  |  |  |  |  |  |  |  |  |  |  |  |  |  |  |  |  |  |  |  |  |  |  |  |  |  |  |  |  |  |  |  |  |  |  |  |  |  |  |  |  |  |  |  |  |  |  |  |  |  |  |  |  |  |  |  |  |  |  |  |  |  |  |  |  |  |  |  |  |  |  |  |  |  |  |  |  |  |  |  |  |  |  |  |  |  |  |  |  |  |  |  |  |  |  |  |  |  |  |  |  |  |  |  |  |  |  |  |  |  |  |  |  |  |  |  |  |  |  |  |  |  |  |  |  |  |  |  |  |  |  |  |  |  |  |  |  |  |  |  |  |  |  |  |  |  |  |  |  |  |  |  |  |  |  |  |  |  |  |  |  |  |  |  |  |  |  |  |  |  |  |  |  |  |  |  |  |  |  |  |  |  |  |  |  |  |  |  |  |  |  |  |  |  |  |  |  |  |  |  |  |  |  |  |  |  |  |  |  |  |  |  |  |  |  |  |  |  |  |  |  |  |  |  |  |  |  |  |  |  |  |  |  |  |  |  |  |  |  |  |  |  |  |  |  |  |  |  |  |  |  |  |  |  |  |  |  |  |  |  |  |  |  |  |  |  |  |  |  |  |  |  |  |  |  |  |  |  |  |  |  |  |  |  |  |  |  |  |  |  |  |  |  |  |  |  |  |  |  |  |  |  |  |  |  |  |  |  |  |  |  |  |  |  |  |  |  |  |  |  |  |  |  |  |  |  |  |  |  |  |  |  |  |  |  |  |  |  |  |  |  |  |  |  |  |  |  |  |  |  |  |  |  |  |  |  |  |  |  |  |  |  |  |  |  |  |  |  |  |  |  |  |  |  |  |  |  |  |  |  |  |  |  |  |  |  |  |  |  |  |  |  |  |  |  |  |  |  |  |  |  |  |  |  |  |  |  |  |  |  |  |  |  |  |  |  |  |  |  |  |  |  |  |  |  |  |  |  |  |  |  |  |  |  |  |  |  |  |  |  |  |  |  |  |  |  |  |  |  |  |  |  |  |  |  |  |  |  |  |  |  |  |  |  |  |  |  |  |  |  |  |  |  |  |  |  |  |  |  |  |  |  |  |  |  |  |  |  |  |  |  |  |  |  |  |  |  |  |  |  |  |  |  |  |  |  |  |  |  |  |  |  |  |  |  |  |  |  |  |  |  |  |  |  |  |  |  |  |  |  |  |  |  |  |  |  |  |  |  |  |  |  |  |  |  |  |  |  |  |  |  |  |  |  |  |  |  |  |  |  |  |  |  |  |  |  |  |  |  |  |  |  |  |  |  |  |  |  |  |  |  |  |  |  |  |  |  |  |  |  |  |  |  |  |  |  |  |  |  |  |  |  |  |  |  |  |  |  |  |  |  |  |  |  |  |  |  |  |  |  |  |  |  |  |  |  |  |  |  |  |  |  |  |  |  |  |  |  |  |  |  |  |  |  |  |  |  |  |  |  |  |  |  |  |  |  |  |  |  |  |  |  |  |  |  |  |  |  |  |  |  |  |  |  |  |  |  |  |  |  |  |  |  |  |  |  |  |  |  |  |  |  |  |  |  |  |  |  |  |  |  |  |  |  |  |  |  |  |  |  |  |  |  |  |  |  |  |  |  |  |  |  |  |  |  |  |  |  |  |  |  |  |  |  |  |  |  |  |  |  |  |  |  |  |  |  |  |  |  |  |  |  |  |  |  |  |  |  |  |  |  |  |  |  |  |  |  |  |  |  |  |  |  |  |  |  |  |  |  |  |  |  |  |  |  |  |  |  |  |  |  |  |  |  |  |  |  |  |  |  |  |  |  |  |  |  |  |  |  |  |  |  |  |  |  |  |  |  |  |  |  |  |  |  |  |  |  |  |  |  |  |  |  |  |  |  |  |  |  |  |  |  |  |  |  |  |  |  |  |  |  |  |  |  |  |  |  |  |  |  |  |  |  |  |  |  |  |  |  |  |  |  |  |  |  |  |  |  |  |  |  |  |  |  |  |  |  |  |  |  |  |  |  |  |  |  |  |  |  |  |  |  |  |  |  |  |  |  |  |  |  |  |  |  |  |  |  |  |  |  |  |  |  |  |  |  |  |  |  |  |  |  |  |  |  |  |  |  |  |  |  |  |  |  |  |  |  |  |  |  |  |  |  |  |  |  |  |  |  |  |  |  |  |  |  |  |  |  |  |  |  |  |  |  |  |  |  |  |  |  |  |  |  |  |  |  |  |  |  |  |  |  |  |  |  |  |  |  |  |  |  |  |  |  |  |  |  |  |  |  |  |  |  |  |  |  |  |  |  |  |  |  |  |  |  |  |  |  |  |  |  |  |  |  |  |  |  |  |  |  |  |  |  |  |  |  |  |  |  |  |  |  |  |  |  |  |  |  |  |  |  |  |  |  |  |  |  |  |  |  |  |  |  |  |  |  |  |  |  |  |  |  |  |  |  |  |  |  |  |  |  |  |  |  |  |  |  |  |  |  |  |  |  |  |  |  |  |  |  |  |  |  |  |  |  |  |  |  |  |  |  |  |  |  |  |  |  |  |  |  |  |  |  |  |  |  |  |  |  |  |  |  |  |  |  |  |  |  |  |  |  |  |  |  |  |  |  |  |  |  |  |  |  |  |  |  |  |  |  |  |  |  |  |  |  |  |  |  |  |  |  |  |  |  |  |  |  |  |  |  |  |  |  |  |  |  |  |  |  |  |  |  |  |  |  |  |  |  |  |  |  |  |  |  |  |  |  |  |  |  |  |  |  |  |  |  |  |  |  |  |  |  |  |  |  |  |  |  |  |  |  |  |  |  |  |  |  |  |  |  |  |  |  |  |  |  |  |  |  |  |  |  |  |  |  |  |  |  |  |  |  |  |  |  |  |  |  |  |  |  |  |  |  |  |  |  |  |  |  |  |  |  |  |  |  |  |  |  |  |  |  |  |  |  |  |  |  |  |  |  |  |  |  |  |  |  |  |  |  |  |  |  |  |  |  |  |  |  |  |  |  |  |  |  |  |  |  |  |  |  |  |  |  |  |  |  |  |  |  |  |  |  |  |  |  |  |  |  |  |  |  |  |  |  |  |  |  |  |  |  |  |  |  |  |  |  |  |  |  |  |  |  |  |  |  |  |  |  |  |  |  |  |  |  |  |  |  |  |  |  |  |  |  |  |  |  |  |  |  |  |  |  |  |  |  |  |  |  |  |  |  |  |  |  |  |  |  |  |  |  |  |  |  |  |  |  |  |  |  |  |  |  |  |  |  |  |  |  |  |  |  |  |  |  |  |  |  |  |  |  |  |  |  |  |  |  |  |  |  |  |  |  |  |  |  |  |  |  |  |  |  |  |  |  |  |  |  |  |  |  |  |  |  |  |  |  |  |  |  |  |  |  |  |  |  |  |  |  |  |  |  |  |  |  |  |  |  |  |  |  |  |  |  |  |  |  |  |  |  |  |  |  |  |  |  |  |  |  |  |  |  |  |  |  |  |  |  |  |  |  |  |  |  |  |  |  |  |  |  |  |  |  |  |  |  |  |  |  |  |  |  |  |  |  |  |  |  |  |  |  |  |  |  |  |  |  |  |  |  |  |  |  |  |  |  |  |  |  |  |  |  |  |  |  |  |  |  |  |  |  |  |  |  |  |  |  |  |  |  |  |  |  |  |  |  |  |  |  |  |  |  |  |  |  |  |  |  |  |  |  |  |  |  |  |  |  |  |  |  |  |  |  |  |  |  |  |  |  |  |  |  |  |  |  |  |  |  |  |  |  |  |  |  |  |  |  |  |  |  |  |  |  |  |  |  |  |  |  |  |  |  |  |  |  |  |  |  |  |  |  |  |  |  |  |  |  |  |  |  |  |  |  |  |  |  |  |  |  |  |  |  |  |  |  |  |  |  |  |  |  |  |  |  |  |  |  |  |  |  |  |  |  |  |  |  |  |  |  |  |  |  |  |  |  |  |  |  |  |  |  |  |  |  |  |  |  |  |  |  |  |  |  |  |  |  |  |  |  |  |  |  |  |  |  |  |  |  |  |  |  |  |  |  |  |  |  |  |  |  |  |  |  |  |  |  |  |  |  |  |  |  |  |  |  |  |  |  |  |  |  |  |  |  |  |  |  |  |  |  |  |  |  |  |  |  |  |  |  |  |  |  |  |  |  |  |  |  |  |  |  |  |  |  |  |  |  |  |  |  |  |  |  |  |  |  |  |  |  |  |  |  |  |  |  |  |  |  |  |  |  |  |  |  |  |  |  |  |  |  |  |  |  |  |  |  |  |  |  |  |  |  |  |  |  |  |  |  |  |  |  |  |  |  |  |  |  |  |  |  |  |  |  |  |  |  |  |  |  |  |  |  |  |  |  |  |  |  |  |  |  |  |  |  |  |  |  |  |  |  |  |  |  |  |  |  |  |  |  |  |  |  |  |  |  |  |  |  |  |  |  |  |  |  |  |  |  |  |  |  |  |  |  |  |  |  |  |  |  |  |  |  |  |  |  |  |  |  |  |  |  |  |  |  |  |  |  |  |  |  |  |  |  |  |  |  |  |  |  |  |  |  |  |  |  |  |  |  |  |  |  |  |  |  |  |  |  |  |  |  |  |  |  |  |  |  |  |  |  |  |  |  |  |  |  |  |  |  |  |  |  |  |  |  |  |  |  |  |  |  |  |  |  |  |  |  |  |  |  |  |  |  |  |  |  |  |  |  |  |  |  |  |  |  |  |  |  |  |  |  |  |  |  |  |  |  |  |  |  |  |  |  |  |  |  |  |  |  |  |  |  |  |  |  |  |  |  |  |  |  |  |  |  |  |  |  |  |  |  |  |  |  |  |  |  |  |  |  |  |  |  |  |  |  |  |  |  |  |  |  |  |  |  |  |  |  |  |  |  |  |  |  |  |  |  |  |  |  |  |  |  |  |  |  |  |  |  |  |  |  |  |  |  |  |  |  |  |  |  |  |  |  |  |  |  |  |  |  |  |  |  |  |  |  |  |  |  |  |  |  |  |  |  |  |  |  |  |  |  |  |  |  |  |  |  |  |  |  |  |  |  |  |  |  |  |  |  |  |  |  |  |  |  |  |  |  |  |  |  |  |  |  |  |  |  |  |  |  |  |  |  |  |  |  |  |  |  |  |  |  |  |  |  |  |  |  |  |  |  |  |  |  |  |  |  |  |  |  |  |  |  |  |  |  |  |  |  |  |  |  |  |  |  |  |  |  |  |  |  |  |  |  |  |  |  |  |  |  |  |  |  |  |  |  |  |  |  |  |  |  |  |  |  |  |  |  |  |  |  |  |  |  |  |  |  |  |  |  |  |  |  |  |  |  |  |  |  |  |  |  |  |  |  |  |  |  |  |  |  |  |  |  |  |  |  |  |  |  |  |  |  |  |  |  |  |  |  |  |  |  |  |  |  |  |  |  |  |  |  |  |  |  |  |  |  |  |  |  |  |  |  |  |  |  |  |  |  |  |  |  |  |  |  |  |  |  |  |  |  |  |  |  |  |  |  |  |  |  |  |  |  |  |  |  |  |  |  |  |  |  |  |  |  |  |  |  |  |  |  |  |  |  |  |  |  |  |  |  |  |  |  |  |  |  |  |  |  |  |  |  |  |  |  |  |  |  |  |  |  |  |  |  |  |  |  |  |  |  |  |  |  |  |  |  |  |  |  |  |  |  |  |  |  |  |  |  |  |  |  |  |  |  |  |  |  |  |  |  |  |  |  |  |  |  |  |  |  |  |  |  |  |  |  |  |  |  |  |  |  |  |  |  |  |  |  |  |  |  |  |  |  |  |  |  |  |  |  |  |  |  |  |  |  |  |  |  |  |  |  |  |  |  |  |  |  |  |  |  |  |  |  |  |  |  |  |  |  |  |  |  |  |  |  |  |  |  |  |  |  |  |  |  |  |  |  |  |  |  |  |  |  |  |  |  |  |  |  |  |  |  |  |  |  |  |  |  |  |  |  |  |  |  |  |  |  |  |  |  |  |  |  |  |  |  |  |  |  |  |  |  |  |  |  |  |  |  |  |  |  |  |  |  |  |  |  |  |  |  |  |  |  |  |  |  |  |  |  |  |  |  |  |  |  |  |  |  |  |  |  |  |  |  |  |  |  |  |  |  |  |  |  |  |  |  |  |  |  |  |  |  |  |  |  |  |  |  |  |  |  |  |  |  |  |  |  |  |  |  |  |  |  |  |  |  |  |  |  |  |  |  |  |  |  |  |  |  |  |  |  |  |  |  |  |  |  |  |  |  |  |  |  |  |  |  |  |  |  |  |  |  |  |  |  |  |  |  |  |  |  |  |  |  |  |  |  |  |  |  |  |  |  |  |  |  |  |  |  |  |  |  |  |  |  |  |  |  |  |  |  |  |  |  |  |  |  |  |  |  |  |  |  |  |  |  |  |  |  |  |  |  |  |  |  |  |  |  |  |  |  |  |  |  |  |  |  |  |  |  |  |  |  |  |  |  |  |  |  |  |  |  |  |  |  |  |  |  |  |  |  |  |  |  |  |  |  |  |  |  |  |  |  |  |  |  |  |  |  |  |  |  |  |  |  |  |  |  |  |  |  |  |  |  |  |  |  |  |  |  |  |  |  |  |  |  |  |  |  |  |  |  |  |  |  |  |  |  |  |  |  |  |  |  |  |  |  |  |  |  |  |  |  |  |  |  |  |  |  |  |  |  |  |  |  |  |  |  |  |  |  |  |  |  |  |  |  |  |  |  |  |  |  |  |  |  |  |  |  |  |  |  |  |  |  |  |  |  |  |  |  |  |  |  |  |  |  |  |  |  |  |  |  |  |  |  |  |  |  |  |  |  |  |  |  |  |  |  |  |  |  |  |  |  |  |  |  |  |  |  |  |  |  |  |  |  |  |  |  |  |  |  |  |  |  |  |  |  |  |  |  |  |  |  |  |  |  |  |  |  |  |  |  |  |  |  |  |  |  |  |  |  |  |  |  |  |  |  |  |  |  |  |  |  |  |  |  |  |  |  |  |  |  |  |  |  |  |  |  |  |  |  |  |  |  |  |  |  |  |  |  |  |  |  |  |  |  |  |  |  |  |  |  |  |  |  |  |  |  |  |  |  |  |  |  |  |  |  |  |  |  |  |  |  |  |  |  |  |  |  |  |  |  |  |  |  |  |  |  |  |  |  |  |  |  |  |  |  |  |  |  |  |  |  |  |  |  |  |  |  |  |  |  |  |  |  |  |  |  |  |  |  |  |  |  |  |  |  |  |  |  |  |  |  |  |  |  |  |  |  |  |  |  |  |  |  |  |  |  |  |  |  |  |  |  |  |  |  |  |  |  |  |  |  |  |  |  |  |  |  |  |  |  |  |  |  |  |  |  |  |  |  |  |  |  |  |  |  |  |  |  |  |  |  |  |  |  |  |  |  |  |  |  |  |  |  |  |  |  |  |  |  |  |  |  |  |  |  |  |  |  |  |  |  |  |  |  |  |  |  |  |  |  |  |  |  |  |  |  |  |  |  |  |  |  |  |  |  |  |  |  |  |  |  |  |  |  |  |  |  |  |  |  |  |  |  |  |  |  |  |  |  |  |  |  |  |  |  |  |  |  |  |  |  |  |  |  |  |  |  |  |  |  |  |  |  |  |  |  |  |  |  |  |  |  |  |  |  |  |  |  |  |  |  |  |  |  |  |  |  |  |  |  |  |  |  |  |  |  |  |  |  |  |  |  |  |  |  |  |  |  |  |  |  |  |  |  |  |  |  |  |  |  |  |  |  |  |  |  |  |  |  |  |  |  |  |  |  |  |  |  |  |  |  |  |  |  |  |  |  |  |  |  |  |  |  |  |  |  |  |  |  |  |  |  |  |  |  |  |  |  |  |  |  |  |  |  |  |  |  |  |  |  |  |  |  |  |  |  |  |  |  |  |  |  |  |  |  |  |  |  |  |  |  |  |  |  |  |  |  |  |  |  |  |  |  |  |  |  |  |  |  |  |  |  |  |  |  |  |  |  |  |  |  |  |  |  |  |  |  |  |  |  |  |  |  |  |  |  |  |  |  |  |  |  |  |  |  |  |  |  |  |  |  |  |  |  |  |  |  |  |  |  |  |  |  |  |  |  |  |  |  |  |  |  |  |  |  |  |  |  |  |  |  |  |  |  |  |  |  |  |  |  |  |  |  |  |  |  |  |  |  |  |  |  |  |  |  |  |  |  |  |  |  |  |  |  |  |  |  |  |  |  |  |  |  |  |  |  |  |  |  |  |  |  |  |  |  |  |  |  |  |  |  |  |  |  |  |  |  |  |  |  |  |  |  |  |  |  |  |  |  |  |  |  |  |  |  |  |  |  |  |  |  |  |  |  |  |  |  |  |  |  |  |  |  |  |  |  |  |  |  |  |  |  |  |  |  |  |  |  |  |  |  |  |  |  |  |  |  |  |  |  |  |  |  |  |  |  |  |  |  |  |  |  |  |  |  |  |  |  |  |  |  |  |  |  |  |  |  |  |  |  |  |  |  |  |  |  |  |  |  |  |  |  |  |  |  |  |  |  |  |  |  |  |  |  |  |  |  |  |  |  |  |  |  |  |  |  |  |  |  |  |  |  |  |  |  |  |  |  |  |  |  |  |  |  |  |  |  |  |  |  |  |  |  |  |  |  |  |  |  |  |  |  |  |  |  |  |  |  |  |  |  |  |  |  |  |  |  |  |  |  |  |  |  |  |  |  |  |  |  |  |  |  |  |  |  |  |  |  |  |  |  |  |  |  |  |  |  |  |  |  |  |  |  |  |  |  |  |  |  |  |  |  |  |  |  |  |  |  |  |  |  |  |  |  |  |  |  |  |  |  |  |  |  |  |  |  |  |  |  |  |  |  |  |  |  |  |  |  |  |  |  |  |  |  |  |  |  |  |  |  |  |  |  |  |  |  |  |  |  |  |  |  |  |  |  |  |  |  |  |  |  |  |  |  |  |  |  |  |  |  |  |  |  |  |  |  |  |  |  |  |  |  |  |  |  |  |  |  |  |  |  |  |  |  |  |  |  |  |  |  |  |  |  |  |  |  |  |  |  |  |  |  |  |  |  |  |  |  |  |  |  |  |  |  |  |  |  |  |  |  |  |  |  |  |  |  |  |  |  |  |  |  |  |  |  |  |  |  |  |  |  |  |  |  |  |  |  |  |  |  |  |  |  |  |  |  |  |  |  |  |  |  |  |  |  |  |  |  |  |  |  |  |  |  |  |  |  |  |  |  |  |  |  |  |  |  |  |  |  |  |  |  |  |  |  |  |  |  |  |  |  |  |  |  |  |  |  |  |  |  |  |  |  |  |  |  |  |  |  |  |  |  |  |  |  |  |  |  |  |  |  |  |  |  |  |  |  |  |  |  |  |  |  |  |  |  |  |  |  |  |  |  |  |  |  |  |  |  |  |  |  |  |  |  |  |  |  |  |  |  |  |  |  |  |  |  |  |  |  |  |  |  |  |  |  |  |  |  |  |  |  |  |  |  |  |  |  |  |  |  |  |  |  |  |  |  |  |  |  |  |  |  |  |  |  |  |  |  |  |  |  |  |  |  |  |  |  |  |  |  |  |  |  |  |  |  |  |  |  |  |  |  |  |  |  |  |  |  |  |  |  |  |  |  |  |  |  |  |  |  |  |  |  |  |  |  |  |  |  |  |  |  |  |  |  |  |  |  |  |  |  |  |  |  |  |  |  |  |  |  |  |  |  |  |  |  |  |  |  |  |  |  |  |  |  |  |  |  |  |  |  |  |  |  |  |  |  |  |  |  |  |  |  |  |  |  |  |  |  |  |  |  |  |  |  |  |  |  |  |  |  |  |  |  |  |  |  |  |  |  |  |  |  |  |  |  |  |  |  |  |  |  |  |  |  |  |  |  |  |  |  |  |  |  |  |  |  |  |  |  |  |  |  |  |  |  |  |  |  |  |  |  |  |  |  |  |  |  |  |  |  |  |  |  |  |  |  |  |  |  |  |  |  |  |  |  |  |  |  |  |  |  |  |  |  |  |  |  |  |  |  |  |  |  |  |  |  |  |  |  |  |  |  |  |  |  |  |  |  |  |  |  |  |  |  |  |  |  |  |  |  |  |  |  |  |  |  |  |  |  |  |  |  |  |  |  |  |  |  |  |  |  |  |  |  |  |  |  |  |  |  |  |  |  |  |  |  |  |  |  |  |  |  |  |  |  |  |  |  |  |  |  |  |  |  |  |  |  |  |  |  |  |  |  |  |  |  |  |  |  |  |  |  |  |  |  |  |  |  |  |  |  |  |  |  |  |  |  |  |  |  |  |  |  |  |  |  |  |  |  |  |  |  |  |  |  |  |  |  |  |  |  |  |  |  |  |  |  |  |  |  |  |  |  |  |  |  |  |  |  |  |  |  |  |  |  |  |  |  |  |  |  |  |  |  |  |  |  |  |  |  |  |  |  |  |  |  |  |  |  |  |  |  |  |  |  |  |  |  |  |  |  |  |  |  |  |  |  |  |  |  |  |  |  |  |  |  |  |  |  |  |  |  |  |  |  |  |  |  |  |  |  |  |  |  |  |  |  |  |  |  |  |  |  |  |  |  |  |  |  |  |  |  |  |  |  |  |  |  |  |  |  |  |  |  |  |  |  |  |  |  |  |  |  |  |  |  |  |  |  |  |  |  |  |  |  |  |  |  |  |  |  |  |  |  |  |  |  |  |  |  |  |  |  |  |  |  |  |  |  |  |  |  |  |  |  |  |  |  |  |  |  |  |  |  |  |  |  |  |  |  |  |  |  |  |  |  |  |  |  |  |  |  |  |  |  |  |  |  |  |  |  |  |  |  |  |  |  |  |  |  |  |  |  |  |  |  |  |  |  |  |  |  |  |  |  |  |  |  |  |  |  |  |  |  |  |  |  |  |  |  |  |  |  |  |  |  |  |  |  |  |  |  |  |  |  |  |  |  |  |  |  |  |  |  |  |  |  |  |  |  |  |  |  |  |  |  |  |  |  |  |  |  |  |  |  |  |  |  |  |  |  |  |  |  |  |  |  |  |  |  |  |  |  |  |  |  |  |  |  |  |  |  |  |  |  |  |  |  |  |  |  |  |  |  |  |  |  |  |  |  |  |  |  |  |  |  |  |  |  |  |  |  |  |  |  |  |  |  |  |  |  |  |  |  |  |  |  |  |  |  |  |  |  |  |  |  |  |  |  |  |  |  |  |  |  |  |  |  |  |  |  |  |  |  |  |  |  |  |  |  |  |  |  |  |  |  |  |  |  |  |  |  |  |  |  |  |  |  |  |  |  |  |  |  |  |  |  |  |  |  |  |  |  |  |  |  |  |  |  |  |  |  |  |  |  |  |  |  |  |  |  |  |  |  |  |  |  |  |  |  |  |  |  |  |  |  |  |  |  |  |  |  |  |  |  |  |  |  |  |  |  |  |  |  |  |  |  |  |  |  |  |  |  |  |  |  |  |  |  |  |  |  |  |  |  |  |  |  |  |  |  |  |  |  |  |  |  |  |  |  |  |  |  |  |  |  |  |  |  |  |  |  |  |  |  |  |  |  |  |  |  |  |  |  |  |  |  |  |  |  |  |  |  |  |  |  |  |  |  |  |  |  |  |  |  |  |  |  |  |  |  |  |  |  |  |  |  |  |  |  |  |  |  |  |  |  |  |  |  |  |  |  |  |  |  |  |  |  |  |  |  |  |  |  |  |  |  |  |  |  |  |  |  |  |  |  |  |  |  |  |  |  |  |  |  |  |  |  |  |  |  |  |  |  |  |  |  |  |  |  |  |  |  |  |  |  |  |  |  |  |  |  |  |  |  |  |  |  |  |  |  |  |  |  |  |  |  |  |  |  |  |  |  |  |  |  |  |  |  |  |  |  |  |  |  |  |  |  |  |  |  |  |  |  |  |  |  |  |  |  |  |  |  |  |  |  |  |  |  |  |  |  |  |  |  |  |  |  |  |  |  |  |  |  |  |  |  |  |  |  |  |  |  |  |  |  |  |  |  |  |  |  |  |  |  |  |  |  |  |  |  |  |  |  |  |  |  |  |  |  |  |  |  |  |  |  |  |  |  |  |  |  |  |  |  |  |  |  |  |  |  |  |  |  |  |  |  |  |  |  |  |  |  |  |  |  |  |  |  |  |  |  |  |  |  |  |  |  |  |  |  |  |  |  |  |  |  |  |  |  |  |  |  |
| --- | --- | --- | --- | --- | --- | --- | --- | --- | --- | --- | --- | --- | --- | --- | --- | --- | --- | --- | --- | --- | --- | --- | --- | --- | --- | --- | --- | --- | --- | --- | --- | --- | --- | --- | --- | --- | --- | --- | --- | --- | --- | --- | --- | --- | --- | --- | --- | --- | --- | --- | --- | --- | --- | --- | --- | --- | --- | --- | --- | --- | --- | --- | --- | --- | --- | --- | --- | --- | --- | --- | --- | --- | --- | --- | --- | --- | --- | --- | --- | --- | --- | --- | --- | --- | --- | --- | --- | --- | --- | --- | --- | --- | --- | --- | --- | --- | --- | --- | --- | --- | --- | --- | --- | --- | --- | --- | --- | --- | --- | --- | --- | --- | --- | --- | --- | --- | --- | --- | --- | --- | --- | --- | --- | --- | --- | --- | --- | --- | --- | --- | --- | --- | --- | --- | --- | --- | --- | --- | --- | --- | --- | --- | --- | --- | --- | --- | --- | --- | --- | --- | --- | --- | --- | --- | --- | --- | --- | --- | --- | --- | --- | --- | --- | --- | --- | --- | --- | --- | --- | --- | --- | --- | --- | --- | --- | --- | --- | --- | --- | --- | --- | --- | --- | --- | --- | --- | --- | --- | --- | --- | --- | --- | --- | --- | --- | --- | --- | --- | --- | --- | --- | --- | --- | --- | --- | --- | --- | --- | --- | --- | --- | --- | --- | --- | --- | --- | --- | --- | --- | --- | --- | --- | --- | --- | --- | --- | --- | --- | --- | --- | --- | --- | --- | --- | --- | --- | --- | --- | --- | --- | --- | --- | --- | --- | --- | --- | --- | --- | --- | --- | --- | --- | --- | --- | --- | --- | --- | --- | --- | --- | --- | --- | --- | --- | --- | --- | --- | --- | --- | --- | --- | --- | --- | --- | --- | --- | --- | --- | --- | --- | --- | --- | --- | --- | --- | --- | --- | --- | --- | --- | --- | --- | --- | --- | --- | --- | --- | --- | --- | --- | --- | --- | --- | --- | --- | --- | --- | --- | --- | --- | --- | --- | --- | --- | --- | --- | --- | --- | --- | --- | --- | --- | --- | --- | --- | --- | --- | --- | --- | --- | --- | --- | --- | --- | --- | --- | --- | --- | --- | --- | --- | --- | --- | --- | --- | --- | --- | --- | --- | --- | --- | --- | --- | --- | --- | --- | --- | --- | --- | --- | --- | --- | --- | --- | --- | --- | --- | --- | --- | --- | --- | --- | --- | --- | --- | --- | --- | --- | --- | --- | --- | --- | --- | --- | --- | --- | --- | --- | --- | --- | --- | --- | --- | --- | --- | --- | --- | --- | --- | --- | --- | --- | --- | --- | --- | --- | --- | --- | --- | --- | --- | --- | --- | --- | --- | --- | --- | --- | --- | --- | --- | --- | --- | --- | --- | --- | --- | --- | --- | --- | --- | --- | --- | --- | --- | --- | --- | --- | --- | --- | --- | --- | --- | --- | --- | --- | --- | --- | --- | --- | --- | --- | --- | --- | --- | --- | --- | --- | --- | --- | --- | --- | --- | --- | --- | --- | --- | --- | --- | --- | --- | --- | --- | --- | --- | --- | --- | --- | --- | --- | --- | --- | --- | --- | --- | --- | --- | --- | --- | --- | --- | --- | --- | --- | --- | --- | --- | --- | --- | --- | --- | --- | --- | --- | --- | --- | --- | --- | --- | --- | --- | --- | --- | --- | --- | --- | --- | --- | --- | --- | --- | --- | --- | --- | --- | --- | --- | --- | --- | --- | --- | --- | --- | --- | --- | --- | --- | --- | --- | --- | --- | --- | --- | --- | --- | --- | --- | --- | --- | --- | --- | --- | --- | --- | --- | --- | --- | --- | --- | --- | --- | --- | --- | --- | --- | --- | --- | --- | --- | --- | --- | --- | --- | --- | --- | --- | --- | --- | --- | --- | --- | --- | --- | --- | --- | --- | --- | --- | --- | --- | --- | --- | --- | --- | --- | --- | --- | --- | --- | --- | --- | --- | --- | --- | --- | --- | --- | --- | --- | --- | --- | --- | --- | --- | --- | --- | --- | --- | --- | --- | --- | --- | --- | --- | --- | --- | --- | --- | --- | --- | --- | --- | --- | --- | --- | --- | --- | --- | --- | --- | --- | --- | --- | --- | --- | --- | --- | --- | --- | --- | --- | --- | --- | --- | --- | --- | --- | --- | --- | --- | --- | --- | --- | --- | --- | --- | --- | --- | --- | --- | --- | --- | --- | --- | --- | --- | --- | --- | --- | --- | --- | --- | --- | --- | --- | --- | --- | --- | --- | --- | --- | --- | --- | --- | --- | --- | --- | --- | --- | --- | --- | --- | --- | --- | --- | --- | --- | --- | --- | --- | --- | --- | --- | --- | --- | --- | --- | --- | --- | --- | --- | --- | --- | --- | --- | --- | --- | --- | --- | --- | --- | --- | --- | --- | --- | --- | --- | --- | --- | --- | --- | --- | --- | --- | --- | --- | --- | --- | --- | --- | --- | --- | --- | --- | --- | --- | --- | --- | --- | --- | --- | --- | --- | --- | --- | --- | --- | --- | --- | --- | --- | --- | --- | --- | --- | --- | --- | --- | --- | --- | --- | --- | --- | --- | --- | --- | --- | --- | --- | --- | --- | --- | --- | --- | --- | --- | --- | --- | --- | --- | --- | --- | --- | --- | --- | --- | --- | --- | --- | --- | --- | --- | --- | --- | --- | --- | --- | --- | --- | --- | --- | --- | --- | --- | --- | --- | --- | --- | --- | --- | --- | --- | --- | --- | --- | --- | --- | --- | --- | --- | --- | --- | --- | --- | --- | --- | --- | --- | --- | --- | --- | --- | --- | --- | --- | --- | --- | --- | --- | --- | --- | --- | --- | --- | --- | --- | --- | --- | --- | --- | --- | --- | --- | --- | --- | --- | --- | --- | --- | --- | --- | --- | --- | --- | --- | --- | --- | --- | --- | --- | --- | --- | --- | --- | --- | --- | --- | --- | --- | --- | --- | --- | --- | --- | --- | --- | --- | --- | --- | --- | --- | --- | --- | --- | --- | --- | --- | --- | --- | --- | --- | --- | --- | --- | --- | --- | --- | --- | --- | --- | --- | --- | --- | --- | --- | --- | --- | --- | --- | --- | --- | --- | --- | --- | --- | --- | --- | --- | --- | --- | --- | --- | --- | --- | --- | --- | --- | --- | --- | --- | --- | --- | --- | --- | --- | --- | --- | --- | --- | --- | --- | --- | --- | --- | --- | --- | --- | --- | --- | --- | --- | --- | --- | --- | --- | --- | --- | --- | --- | --- | --- | --- | --- | --- | --- | --- | --- | --- | --- | --- | --- | --- | --- | --- | --- | --- | --- | --- | --- | --- | --- | --- | --- | --- | --- | --- | --- | --- | --- | --- | --- | --- | --- | --- | --- | --- | --- | --- | --- | --- | --- | --- | --- | --- | --- | --- | --- | --- | --- | --- | --- | --- | --- | --- | --- | --- | --- | --- | --- | --- | --- | --- | --- | --- | --- | --- | --- | --- | --- | --- | --- | --- | --- | --- | --- | --- | --- | --- | --- | --- | --- | --- | --- | --- | --- | --- | --- | --- | --- | --- | --- | --- | --- | --- | --- | --- | --- | --- | --- | --- | --- | --- | --- | --- | --- | --- | --- | --- | --- | --- | --- | --- | --- | --- | --- | --- | --- | --- | --- | --- | --- | --- | --- | --- | --- | --- | --- | --- | --- | --- | --- | --- | --- | --- | --- | --- | --- | --- | --- | --- | --- | --- | --- | --- | --- | --- | --- | --- | --- | --- | --- | --- | --- | --- | --- | --- | --- | --- | --- | --- | --- | --- | --- | --- | --- | --- | --- | --- | --- | --- | --- | --- | --- | --- | --- | --- | --- | --- | --- | --- | --- | --- | --- | --- | --- | --- | --- | --- | --- | --- | --- | --- | --- | --- | --- | --- | --- | --- | --- | --- | --- | --- | --- | --- | --- | --- | --- | --- | --- | --- | --- | --- | --- | --- | --- | --- | --- | --- | --- | --- | --- | --- | --- | --- | --- | --- | --- | --- | --- | --- | --- | --- | --- | --- | --- | --- | --- | --- | --- | --- | --- | --- | --- | --- | --- | --- | --- | --- | --- | --- | --- | --- | --- | --- | --- | --- | --- | --- | --- | --- | --- | --- | --- | --- | --- | --- | --- | --- | --- | --- | --- | --- | --- | --- | --- | --- | --- | --- | --- | --- | --- | --- | --- | --- | --- | --- | --- | --- | --- | --- | --- | --- | --- | --- | --- | --- | --- | --- | --- | --- | --- | --- | --- | --- | --- | --- | --- | --- | --- | --- | --- | --- | --- | --- | --- | --- | --- | --- | --- | --- | --- | --- | --- | --- | --- | --- | --- | --- | --- | --- | --- | --- | --- | --- | --- | --- | --- | --- | --- | --- | --- | --- | --- | --- | --- | --- | --- | --- | --- | --- | --- | --- | --- | --- | --- | --- | --- | --- | --- | --- | --- | --- | --- | --- | --- | --- | --- | --- | --- | --- | --- | --- | --- | --- | --- | --- | --- | --- | --- | --- | --- | --- | --- | --- | --- | --- | --- | --- | --- | --- | --- | --- | --- | --- | --- | --- | --- | --- | --- | --- | --- | --- | --- | --- | --- | --- | --- | --- | --- | --- | --- | --- | --- | --- | --- | --- | --- | --- | --- | --- | --- | --- | --- | --- | --- | --- | --- | --- | --- | --- | --- | --- | --- | --- | --- | --- | --- | --- | --- | --- | --- | --- | --- | --- | --- | --- | --- | --- | --- | --- | --- | --- | --- | --- | --- | --- | --- | --- | --- | --- | --- | --- | --- | --- | --- | --- | --- | --- | --- | --- | --- | --- | --- | --- | --- | --- | --- | --- | --- | --- | --- | --- | --- | --- | --- | --- | --- | --- | --- | --- | --- | --- | --- | --- | --- | --- | --- | --- | --- | --- | --- | --- | --- | --- | --- | --- | --- | --- | --- | --- | --- | --- | --- | --- | --- | --- | --- | --- | --- | --- | --- | --- | --- | --- | --- | --- | --- | --- | --- | --- | --- | --- | --- | --- | --- | --- | --- | --- | --- | --- | --- | --- | --- | --- | --- | --- | --- | --- | --- | --- | --- | --- | --- | --- | --- | --- | --- | --- | --- | --- | --- | --- | --- | --- | --- | --- | --- | --- | --- | --- | --- | --- | --- | --- | --- | --- | --- | --- | --- | --- | --- | --- | --- | --- | --- | --- | --- | --- | --- | --- | --- | --- | --- | --- | --- | --- | --- | --- | --- | --- | --- | --- | --- | --- | --- | --- | --- | --- | --- | --- | --- | --- | --- | --- | --- | --- | --- | --- | --- | --- | --- | --- | --- | --- | --- | --- | --- | --- | --- | --- | --- | --- | --- | --- | --- | --- | --- | --- | --- | --- | --- | --- | --- | --- | --- | --- | --- | --- | --- | --- | --- | --- | --- | --- | --- | --- | --- | --- | --- | --- | --- | --- | --- | --- | --- | --- | --- | --- | --- | --- | --- | --- | --- | --- | --- | --- | --- | --- | --- | --- | --- | --- | --- | --- | --- | --- | --- | --- | --- | --- | --- | --- | --- | --- | --- | --- | --- | --- | --- | --- | --- | --- | --- | --- | --- | --- | --- | --- | --- | --- | --- | --- | --- | --- | --- | --- | --- | --- | --- | --- | --- | --- | --- | --- | --- | --- | --- | --- | --- | --- | --- | --- | --- | --- | --- | --- | --- | --- | --- | --- | --- | --- | --- | --- | --- | --- | --- | --- | --- | --- | --- | --- | --- | --- | --- | --- | --- | --- | --- | --- | --- | --- | --- | --- | --- | --- | --- | --- | --- | --- | --- | --- | --- | --- | --- | --- | --- | --- | --- | --- | --- | --- | --- | --- | --- | --- | --- | --- | --- | --- | --- | --- | --- | --- | --- | --- | --- | --- | --- | --- | --- | --- | --- | --- | --- | --- | --- | --- | --- | --- | --- | --- | --- | --- | --- | --- | --- | --- | --- | --- | --- | --- | --- | --- | --- | --- | --- | --- | --- | --- | --- | --- | --- | --- | --- | --- | --- | --- | --- | --- | --- | --- | --- | --- | --- | --- | --- | --- | --- | --- | --- | --- | --- | --- | --- | --- | --- | --- | --- | --- | --- | --- | --- | --- | --- | --- | --- | --- | --- | --- | --- | --- | --- | --- | --- | --- | --- | --- | --- | --- | --- | --- | --- | --- | --- | --- | --- | --- | --- | --- | --- | --- | --- | --- | --- | --- | --- | --- | --- | --- | --- | --- | --- | --- | --- | --- | --- | --- | --- | --- | --- | --- | --- | --- | --- | --- | --- | --- | --- | --- | --- | --- | --- | --- | --- | --- | --- | --- | --- | --- | --- | --- | --- | --- | --- | --- | --- | --- | --- | --- | --- | --- | --- | --- | --- | --- | --- | --- | --- | --- | --- | --- | --- | --- | --- | --- | --- | --- | --- | --- | --- | --- | --- | --- | --- | --- | --- | --- | --- | --- | --- | --- | --- | --- | --- | --- | --- | --- | --- | --- | --- | --- | --- | --- | --- | --- | --- | --- | --- | --- | --- | --- | --- | --- | --- | --- | --- | --- | --- | --- | --- | --- | --- | --- | --- | --- | --- | --- | --- | --- | --- | --- | --- | --- | --- | --- | --- | --- | --- | --- | --- | --- | --- | --- | --- | --- | --- | --- | --- | --- | --- | --- | --- | --- | --- | --- | --- | --- | --- | --- | --- | --- | --- | --- | --- | --- | --- | --- | --- | --- | --- | --- | --- | --- | --- | --- | --- | --- | --- | --- | --- | --- | --- | --- | --- | --- | --- | --- | --- | --- | --- | --- | --- | --- | --- | --- | --- | --- | --- | --- | --- | --- | --- | --- | --- | --- | --- | --- | --- | --- | --- | --- | --- | --- | --- | --- | --- | --- | --- | --- | --- | --- | --- | --- | --- | --- | --- | --- | --- | --- | --- | --- | --- | --- | --- | --- | --- | --- | --- | --- | --- | --- | --- | --- | --- | --- | --- | --- | --- | --- | --- | --- | --- | --- | --- | --- | --- | --- | --- | --- | --- | --- | --- | --- | --- | --- | --- | --- | --- | --- | --- | --- | --- | --- | --- | --- | --- | --- | --- | --- | --- | --- | --- | --- | --- | --- | --- | --- | --- | --- | --- | --- | --- | --- | --- | --- | --- | --- | --- | --- | --- | --- | --- | --- | --- | --- | --- | --- | --- | --- | --- | --- | --- | --- | --- | --- | --- | --- | --- | --- | --- | --- | --- | --- | --- | --- | --- | --- | --- | --- | --- | --- | --- | --- | --- | --- | --- | --- | --- | --- | --- | --- | --- | --- | --- | --- | --- | --- | --- | --- | --- | --- | --- | --- | --- | --- | --- | --- | --- | --- | --- | --- | --- | --- | --- | --- | --- | --- | --- | --- | --- | --- | --- | --- | --- | --- | --- | --- | --- | --- | --- | --- | --- | --- | --- | --- | --- | --- | --- | --- | --- | --- | --- | --- | --- | --- | --- | --- | --- | --- | --- | --- | --- | --- | --- | --- | --- | --- | --- | --- | --- | --- | --- | --- | --- | --- | --- | --- | --- | --- | --- | --- | --- | --- | --- | --- | --- | --- | --- | --- | --- | --- | --- | --- | --- | --- | --- | --- | --- | --- | --- | --- | --- | --- | --- | --- | --- | --- | --- | --- | --- | --- | --- | --- | --- | --- | --- | --- | --- | --- | --- | --- | --- | --- | --- | --- | --- | --- | --- | --- | --- | --- | --- | --- | --- | --- | --- | --- | --- | --- | --- | --- | --- | --- | --- | --- | --- | --- | --- | --- | --- | --- | --- | --- | --- | --- | --- | --- | --- | --- | --- | --- | --- | --- | --- | --- | --- | --- | --- | --- | --- | --- | --- | --- | --- | --- | --- | --- | --- | --- | --- | --- | --- | --- | --- | --- | --- | --- | --- | --- | --- | --- | --- | --- | --- | --- | --- | --- | --- | --- | --- | --- | --- | --- | --- | --- | --- | --- | --- | --- | --- | --- | --- | --- | --- | --- | --- | --- | --- | --- | --- | --- | --- | --- | --- | --- | --- | --- | --- | --- | --- | --- | --- | --- | --- | --- | --- | --- | --- | --- | --- | --- | --- | --- | --- | --- | --- | --- | --- | --- | --- | --- | --- | --- | --- | --- | --- | --- | --- | --- | --- | --- | --- | --- | --- | --- | --- | --- | --- | --- | --- | --- | --- | --- | --- | --- | --- | --- | --- | --- | --- | --- | --- | --- | --- | --- | --- | --- | --- | --- | --- | --- | --- | --- | --- | --- | --- | --- | --- | --- | --- | --- | --- | --- | --- | --- | --- | --- | --- | --- | --- | --- | --- | --- | --- | --- | --- | --- | --- | --- | --- | --- | --- | --- | --- | --- | --- | --- | --- | --- | --- | --- | --- | --- | --- | --- | --- | --- | --- | --- | --- | --- | --- | --- | --- | --- | --- | --- | --- | --- | --- | --- | --- | --- | --- | --- | --- | --- | --- | --- | --- | --- | --- | --- | --- | --- | --- | --- | --- | --- | --- | --- | --- | --- | --- | --- | --- | --- | --- | --- | --- | --- | --- | --- | --- | --- | --- | --- | --- | --- | --- | --- | --- | --- | --- | --- | --- | --- | --- | --- | --- | --- | --- | --- | --- | --- | --- | --- | --- | --- | --- | --- | --- | --- | --- | --- | --- | --- | --- | --- | --- | --- | --- | --- | --- | --- | --- | --- | --- | --- | --- | --- | --- | --- | --- | --- | --- | --- | --- | --- | --- | --- | --- | --- | --- | --- | --- | --- | --- | --- | --- | --- | --- | --- | --- | --- | --- | --- | --- | --- | --- | --- | --- | --- | --- | --- | --- | --- | --- | --- | --- | --- | --- | --- | --- | --- | --- | --- | --- | --- | --- | --- | --- | --- | --- | --- | --- | --- | --- | --- | --- | --- | --- | --- | --- | --- | --- | --- | --- | --- | --- | --- | --- | --- | --- | --- | --- | --- | --- | --- | --- | --- | --- | --- | --- | --- | --- | --- | --- | --- | --- | --- | --- | --- | --- | --- | --- | --- | --- | --- | --- | --- | --- | --- | --- | --- | --- | --- | --- | --- | --- | --- | --- | --- | --- | --- | --- | --- | --- | --- | --- | --- | --- | --- | --- | --- | --- | --- | --- | --- | --- | --- | --- | --- | --- | --- | --- | --- | --- | --- | --- | --- | --- | --- | --- | --- | --- | --- | --- | --- | --- | --- | --- | --- | --- | --- | --- | --- | --- | --- | --- | --- | --- | --- | --- | --- | --- | --- | --- | --- | --- | --- | --- | --- | --- | --- | --- | --- | --- | --- | --- | --- | --- | --- | --- | --- | --- | --- | --- | --- | --- | --- | --- | --- | --- | --- | --- | --- | --- | --- | --- | --- | --- | --- | --- | --- | --- | --- | --- | --- | --- | --- | --- | --- | --- | --- | --- | --- | --- | --- | --- | --- | --- | --- | --- | --- | --- | --- | --- | --- | --- | --- | --- | --- | --- | --- | --- | --- | --- | --- | --- | --- | --- | --- | --- | --- | --- | --- | --- | --- | --- | --- | --- | --- | --- | --- | --- | --- | --- | --- | --- | --- | --- | --- | --- | --- | --- | --- | --- | --- | --- | --- | --- | --- | --- | --- | --- | --- | --- | --- | --- | --- | --- | --- | --- | --- | --- | --- | --- | --- | --- | --- | --- | --- | --- | --- | --- | --- | --- | --- | --- | --- | --- | --- | --- | --- | --- | --- | --- | --- | --- | --- | --- | --- | --- | --- | --- | --- | --- | --- | --- | --- | --- | --- | --- | --- | --- | --- | --- | --- | --- | --- | --- | --- | --- | --- | --- | --- | --- | --- | --- | --- | --- | --- | --- | --- | --- | --- | --- | --- | --- | --- | --- | --- | --- | --- | --- | --- | --- | --- | --- | --- | --- | --- | --- | --- | --- | --- | --- | --- | --- | --- | --- | --- | --- | --- | --- | --- | --- | --- | --- | --- | --- | --- | --- | --- | --- | --- | --- | --- | --- | --- | --- | --- | --- | --- | --- | --- | --- | --- | --- | --- | --- | --- | --- | --- | --- | --- | --- | --- | --- | --- | --- | --- | --- | --- | --- | --- | --- | --- | --- | --- | --- | --- | --- | --- | --- | --- | --- | --- | --- | --- | --- | --- | --- | --- | --- | --- | --- | --- | --- | --- | --- | --- | --- | --- | --- | --- | --- | --- | --- | --- | --- | --- | --- | --- | --- | --- | --- | --- | --- | --- | --- | --- | --- | --- | --- | --- | --- | --- | --- | --- | --- | --- | --- | --- | --- | --- | --- | --- | --- | --- | --- | --- | --- | --- | --- | --- | --- | --- | --- | --- | --- | --- | --- | --- | --- | --- | --- | --- | --- | --- | --- | --- | --- | --- | --- | --- | --- | --- | --- | --- | --- | --- | --- | --- | --- | --- | --- | --- | --- | --- | --- | --- | --- | --- | --- | --- | --- | --- | --- | --- | --- | --- | --- | --- | --- | --- | --- | --- | --- | --- | --- | --- | --- | --- | --- | --- | --- | --- | --- | --- | --- | --- | --- | --- | --- | --- | --- | --- | --- | --- | --- | --- | --- | --- | --- | --- | --- | --- | --- | --- | --- | --- | --- | --- | --- | --- | --- | --- | --- | --- | --- | --- | --- | --- | --- | --- | --- | --- | --- | --- | --- | --- | --- | --- | --- | --- | --- | --- | --- | --- | --- | --- | --- | --- | --- | --- | --- | --- | --- | --- | --- | --- | --- | --- | --- | --- | --- | --- | --- | --- | --- | --- | --- | --- | --- | --- | --- | --- | --- | --- | --- | --- | --- | --- | --- | --- | --- | --- | --- | --- | --- | --- | --- | --- | --- | --- | --- | --- | --- | --- | --- | --- | --- | --- | --- | --- | --- | --- | --- | --- | --- | --- | --- | --- | --- | --- | --- | --- | --- | --- | --- | --- | --- | --- | --- | --- | --- | --- | --- | --- | --- | --- | --- | --- | --- | --- | --- | --- | --- | --- | --- | --- | --- | --- | --- | --- | --- | --- | --- | --- | --- | --- | --- | --- | --- | --- | --- | --- | --- | --- | --- | --- | --- | --- | --- | --- | --- | --- | --- | --- | --- | --- | --- | --- | --- | --- | --- | --- | --- | --- | --- | --- | --- | --- | --- | --- | --- | --- | --- | --- | --- | --- | --- | --- | --- | --- | --- | --- | --- | --- | --- | --- | --- | --- | --- | --- | --- | --- | --- | --- | --- | --- | --- | --- | --- | --- | --- | --- | --- | --- | --- | --- | --- | --- | --- | --- | --- | --- | --- | --- | --- | --- | --- | --- | --- | --- | --- | --- | --- | --- | --- | --- | --- | --- | --- | --- | --- | --- | --- | --- | --- | --- | --- | --- | --- | --- | --- | --- | --- | --- | --- | --- | --- | --- | --- | --- | --- | --- | --- | --- | --- | --- | --- | --- | --- | --- | --- | --- | --- | --- | --- | --- | --- | --- | --- | --- | --- | --- | --- | --- | --- | --- | --- | --- | --- | --- | --- | --- | --- | --- | --- | --- | --- | --- | --- | --- | --- | --- | --- | --- | --- | --- | --- | --- | --- | --- | --- | --- | --- | --- | --- | --- | --- | --- | --- | --- | --- | --- | --- | --- | --- | --- | --- | --- | --- | --- | --- | --- | --- | --- | --- | --- | --- | --- | --- | --- | --- | --- | --- | --- | --- | --- | --- | --- | --- | --- | --- | --- | --- | --- | --- | --- | --- | --- | --- | --- | --- | --- | --- | --- | --- | --- | --- | --- | --- | --- | --- | --- | --- | --- | --- | --- | --- | --- | --- | --- | --- | --- | --- | --- | --- | --- | --- | --- | --- | --- | --- | --- | --- | --- | --- | --- | --- | --- | --- | --- | --- | --- | --- | --- | --- | --- | --- | --- | --- | --- | --- | --- | --- | --- | --- | --- | --- | --- | --- | --- | --- | --- | --- | --- | --- | --- | --- | --- | --- | --- | --- | --- | --- | --- | --- | --- | --- | --- | --- | --- | --- | --- | --- | --- | --- | --- | --- | --- | --- | --- | --- | --- | --- | --- | --- | --- | --- | --- | --- | --- | --- | --- | --- | --- | --- | --- | --- | --- | --- | --- | --- | --- | --- | --- | --- | --- | --- | --- | --- | --- | --- | --- | --- | --- | --- | --- | --- | --- | --- | --- | --- | --- | --- | --- | --- | --- | --- | --- | --- | --- | --- | --- | --- | --- | --- | --- | --- | --- | --- | --- | --- | --- | --- | --- | --- | --- | --- | --- | --- | --- | --- | --- | --- | --- | --- | --- | --- | --- | --- | --- | --- | --- | --- | --- | --- | --- | --- | --- | --- | --- | --- | --- | --- | --- | --- | --- | --- | --- | --- | --- | --- | --- | --- | --- | --- | --- | --- | --- | --- | --- | --- | --- | --- | --- | --- | --- | --- | --- | --- | --- | --- | --- | --- | --- | --- | --- | --- | --- | --- | --- | --- | --- | --- | --- | --- | --- | --- | --- | --- | --- | --- | --- | --- | --- | --- | --- | --- | --- | --- | --- | --- | --- | --- | --- | --- | --- | --- | --- | --- | --- | --- | --- | --- | --- | --- | --- | --- | --- | --- | --- | --- | --- | --- | --- | --- | --- | --- | --- | --- | --- | --- | --- | --- | --- | --- | --- | --- | --- | --- | --- | --- | --- | --- | --- | --- | --- | --- | --- | --- | --- | --- | --- | --- | --- | --- | --- | --- | --- | --- | --- | --- | --- | --- | --- | --- | --- | --- | --- | --- | --- | --- | --- | --- | --- | --- | --- | --- | --- | --- | --- | --- | --- | --- | --- | --- | --- | --- | --- | --- | --- | --- | --- | --- | --- | --- | --- | --- | --- | --- | --- | --- | --- | --- | --- | --- | --- | --- | --- | --- | --- | --- | --- | --- | --- | --- | --- | --- | --- | --- | --- | --- | --- | --- | --- | --- | --- | --- | --- | --- | --- | --- | --- | --- | --- | --- | --- | --- | --- | --- | --- | --- | --- | --- | --- | --- | --- | --- | --- | --- | --- | --- | --- | --- | --- | --- | --- | --- | --- | --- | --- | --- | --- | --- | --- | --- | --- | --- | --- | --- | --- | --- | --- | --- | --- | --- | --- | --- | --- | --- | --- | --- | --- | --- | --- | --- | --- | --- | --- | --- | --- | --- | --- | --- | --- | --- | --- | --- | --- | --- | --- | --- | --- | --- | --- | --- | --- | --- | --- | --- | --- | --- | --- | --- | --- | --- | --- | --- | --- | --- | --- | --- | --- | --- | --- | --- | --- | --- | --- | --- | --- | --- | --- | --- | --- | --- | --- | --- | --- | --- | --- | --- | --- | --- | --- | --- | --- | --- | --- | --- | --- | --- | --- | --- | --- | --- | --- | --- | --- | --- | --- | --- | --- | --- | --- | --- | --- | --- | --- | --- | --- | --- | --- | --- | --- | --- | --- | --- | --- | --- | --- | --- | --- | --- | --- | --- | --- | --- | --- | --- | --- | --- | --- | --- | --- | --- | --- | --- | --- | --- | --- | --- | --- | --- | --- | --- | --- | --- | --- | --- | --- | --- | --- | --- | --- | --- | --- | --- | --- | --- | --- | --- | --- | --- | --- | --- | --- | --- | --- | --- | --- | --- | --- | --- | --- | --- | --- | --- | --- | --- | --- | --- | --- | --- | --- | --- | --- | --- | --- | --- | --- | --- | --- | --- | --- | --- | --- | --- | --- | --- | --- | --- | --- | --- | --- | --- | --- | --- | --- | --- | --- | --- | --- | --- | --- | --- | --- | --- | --- | --- | --- | --- | --- | --- | --- | --- | --- | --- | --- | --- | --- | --- | --- | --- | --- | --- | --- | --- | --- | --- | --- | --- | --- | --- | --- | --- | --- | --- | --- | --- | --- | --- | --- | --- | --- | --- | --- | --- | --- | --- | --- | --- | --- | --- | --- | --- | --- | --- | --- | --- | --- | --- | --- | --- | --- | --- | --- | --- | --- | --- | --- | --- | --- | --- | --- | --- | --- | --- | --- | --- | --- | --- | --- | --- | --- | --- | --- | --- | --- | --- | --- | --- | --- | --- | --- | --- | --- | --- | --- | --- | --- | --- | --- | --- | --- | --- | --- | --- | --- | --- | --- | --- | --- | --- | --- | --- | --- | --- | --- | --- | --- | --- | --- | --- | --- | --- | --- | --- | --- | --- | --- | --- | --- | --- | --- | --- | --- | --- | --- | --- | --- | --- | --- | --- | --- | --- | --- | --- | --- | --- | --- | --- | --- | --- | --- | --- | --- | --- | --- | --- | --- | --- | --- | --- | --- | --- | --- | --- | --- | --- | --- | --- | --- | --- | --- | --- | --- | --- | --- | --- | --- | --- | --- | --- | --- | --- | --- | --- | --- | --- | --- | --- | --- | --- | --- | --- | --- | --- | --- | --- | --- | --- | --- | --- | --- | --- | --- | --- | --- | --- | --- | --- | --- | --- | --- | --- | --- | --- | --- | --- | --- | --- | --- | --- | --- | --- | --- | --- | --- | --- | --- | --- | --- | --- | --- | --- | --- | --- | --- | --- | --- | --- | --- | --- | --- | --- | --- | --- | --- | --- | --- | --- | --- | --- | --- | --- | --- | --- | --- | --- | --- | --- | --- | --- | --- | --- | --- | --- | --- | --- | --- | --- | --- | --- | --- | --- | --- | --- | --- | --- | --- | --- | --- | --- | --- | --- | --- | --- | --- | --- | --- | --- | --- | --- | --- | --- | --- | --- | --- | --- | --- | --- | --- | --- | --- | --- | --- | --- | --- | --- | --- | --- | --- | --- | --- | --- | --- | --- | --- | --- | --- | --- | --- | --- | --- | --- | --- | --- | --- | --- | --- | --- | --- | --- | --- | --- | --- | --- | --- | --- | --- | --- | --- | --- | --- | --- | --- | --- | --- | --- | --- | --- | --- | --- | --- | --- | --- | --- | --- | --- | --- | --- | --- | --- | --- | --- | --- | --- | --- | --- | --- | --- | --- | --- | --- | --- | --- | --- | --- | --- | --- | --- | --- | --- | --- | --- | --- | --- | --- | --- | --- | --- | --- | --- | --- | --- | --- | --- | --- | --- | --- | --- | --- | --- | --- | --- | --- | --- | --- | --- | --- | --- | --- | --- | --- | --- | --- | --- | --- | --- | --- | --- | --- | --- | --- | --- | --- | --- | --- | --- | --- | --- | --- | --- | --- | --- | --- | --- | --- | --- | --- | --- | --- | --- | --- | --- | --- | --- | --- | --- | --- | --- | --- | --- | --- | --- | --- | --- | --- | --- | --- | --- | --- | --- | --- | --- | --- | --- | --- | --- | --- | --- | --- | --- | --- | --- | --- | --- | --- | --- | --- | --- | --- | --- | --- | --- | --- | --- | --- | --- | --- | --- | --- | --- | --- | --- | --- | --- | --- | --- | --- | --- | --- | --- | --- | --- | --- | --- | --- | --- | --- | --- | --- | --- | --- | --- | --- | --- | --- | --- | --- | --- | --- | --- | --- | --- | --- | --- | --- | --- | --- | --- | --- | --- | --- | --- | --- | --- | --- | --- | --- | --- | --- | --- | --- | --- | --- | --- | --- | --- | --- | --- | --- | --- | --- | --- | --- | --- | --- | --- | --- | --- | --- | --- | --- | --- | --- | --- | --- | --- | --- | --- | --- | --- | --- | --- | --- | --- | --- | --- | --- | --- | --- | --- | --- | --- | --- | --- | --- | --- | --- | --- | --- | --- | --- | --- | --- | --- | --- | --- | --- | --- | --- | --- | --- | --- | --- | --- | --- | --- | --- | --- | --- | --- | --- | --- | --- | --- | --- | --- | --- | --- | --- | --- | --- | --- | --- | --- | --- | --- | --- | --- | --- | --- | --- | --- | --- | --- | --- | --- | --- | --- | --- | --- | --- | --- | --- | --- | --- | --- | --- | --- | --- | --- | --- | --- | --- | --- | --- | --- | --- | --- | --- | --- | --- | --- | --- | --- | --- | --- | --- | --- | --- | --- | --- | --- | --- | --- | --- | --- | --- | --- | --- | --- | --- | --- | --- | --- | --- | --- | --- | --- | --- | --- | --- | --- | --- | --- | --- | --- | --- | --- | --- | --- | --- | --- | --- | --- | --- | --- | --- | --- | --- | --- | --- | --- | --- | --- | --- | --- | --- | --- | --- | --- | --- | --- | --- | --- | --- | --- | --- | --- | --- | --- | --- | --- | --- | --- | --- | --- | --- | --- | --- | --- | --- | --- | --- | --- | --- | --- | --- | --- | --- | --- | --- | --- | --- | --- | --- | --- | --- | --- | --- | --- | --- | --- | --- | --- | --- | --- | --- | --- | --- | --- | --- | --- | --- | --- | --- | --- | --- | --- | --- | --- | --- | --- | --- | --- | --- | --- | --- | --- | --- | --- | --- | --- | --- | --- | --- | --- | --- | --- | --- | --- | --- | --- | --- | --- | --- | --- | --- | --- | --- | --- | --- | --- | --- | --- | --- | --- | --- | --- | --- | --- | --- | --- | --- | --- | --- | --- | --- | --- | --- | --- | --- | --- | --- | --- | --- | --- | --- | --- | --- | --- | --- | --- | --- | --- | --- | --- | --- | --- | --- | --- | --- | --- | --- | --- | --- | --- | --- | --- | --- | --- | --- | --- | --- | --- | --- | --- | --- | --- | --- | --- | --- | --- | --- | --- | --- | --- | --- | --- | --- | --- | --- | --- | --- | --- | --- | --- | --- | --- | --- | --- | --- | --- | --- | --- | --- | --- | --- | --- | --- | --- | --- | --- | --- | --- | --- | --- | --- | --- | --- | --- | --- | --- | --- | --- | --- | --- | --- | --- | --- | --- | --- | --- | --- | --- | --- | --- | --- | --- | --- | --- | --- | --- | --- | --- | --- | --- | --- | --- | --- | --- | --- | --- | --- | --- | --- | --- | --- | --- | --- | --- | --- | --- | --- | --- | --- | --- | --- | --- | --- | --- | --- | --- | --- | --- | --- | --- | --- | --- | --- | --- | --- | --- | --- | --- | --- | --- | --- | --- | --- | --- | --- | --- | --- | --- | --- | --- | --- | --- | --- | --- | --- | --- | --- | --- | --- | --- | --- | --- | --- | --- | --- | --- | --- | --- | --- | --- | --- | --- | --- | --- | --- | --- | --- | --- | --- | --- | --- | --- | --- | --- | --- | --- | --- | --- | --- | --- | --- | --- | --- | --- | --- | --- | --- | --- | --- | --- | --- | --- | --- | --- | --- | --- | --- | --- | --- | --- | --- | --- | --- | --- | --- | --- | --- | --- | --- | --- | --- | --- | --- | --- | --- | --- | --- | --- | --- | --- | --- | --- | --- | --- | --- | --- | --- | --- | --- | --- | --- | --- | --- | --- | --- | --- | --- | --- | --- | --- | --- | --- | --- | --- | --- | --- | --- | --- | --- | --- | --- | --- | --- | --- | --- | --- | --- | --- | --- | --- | --- | --- | --- | --- | --- | --- | --- | --- | --- | --- | --- | --- | --- | --- | --- | --- | --- | --- | --- | --- | --- | --- | --- | --- | --- | --- | --- | --- | --- | --- | --- | --- | --- | --- | --- | --- | --- | --- | --- | --- | --- | --- | --- | --- | --- | --- | --- | --- | --- | --- | --- | --- | --- | --- | --- | --- | --- | --- | --- | --- | --- | --- | --- | --- | --- | --- | --- | --- | --- | --- | --- | --- | --- | --- | --- | --- | --- | --- | --- | --- | --- | --- | --- | --- | --- | --- | --- | --- | --- | --- | --- | --- | --- | --- | --- | --- | --- | --- | --- | --- | --- | --- | --- | --- | --- | --- | --- | --- | --- | --- | --- | --- | --- | --- | --- | --- | --- | --- | --- | --- | --- | --- | --- | --- | --- | --- | --- | --- | --- | --- | --- | --- | --- | --- | --- | --- | --- | --- | --- | --- | --- | --- | --- | --- | --- | --- | --- | --- | --- | --- |
|  |  |  |  |  |  |  |  | 1 | 0 |  |  |  |  |  |  |  |  | 2 | 0 |  |  |  |  |  |  |  |  | 3 | 0 |  |  |  |  |  |  |  |  | 4 | 0 |  |  |  |  |  |  |  |  | 5 | 0 |  |  |  |  |  |  |  |  | 6 | 0 |  |  |  |  |  |  |  |  | 7 | 0 |  |  |  |  |  |  |  |  | 8 | 0 |  |  |  |  |  |  |  |  | 9 | 0 |  |  |  |  |  |  |  | 1 | 0 | 0 |  |  |  |  |  |  |  | 1 | 1 | 0 |  |  |  |  |  |  |  | 1 | 2 | 0 |  |  |  |  |  |  |  | 1 | 3 | 0 |  |  |  |  |  |  |  | 1 | 4 | 0 |  |  |  |  |  |  |  | 1 | 5 | 0 |  |  |  |  |  |  |  | 1 | 6 | 0 |  |  |  |  |  |  |  | 1 | 7 | 0 |  |  |  |  |  |  |  | 1 | 8 | 0 |  |  |  |  |  |  |  | 1 | 9 | 0 |  |  |  |  |  |  |  | 2 | 0 | 0 |  |  |  |  |  |  |  | 2 | 1 | 0 |  |  |  |  |  |  |  | 2 | 2 | 0 |  |  |  |  |  |  |  | 2 | 3 | 0 |  |  |  |  |  |  |  | 2 | 4 | 0 |  |  |  |  |  |  |  | 2 | 5 | 0 |  |  |  |  |  |  |  | 2 | 6 | 0 |  |  |  |  |  |  |  | 2 | 7 | 0 |  |  |  |  |  |  |  | 2 | 8 | 0 |  |  |  |  |  |  |  | 2 | 9 | 0 |  |  |  |  |  |  |  | 3 | 0 | 0 |  |  |  |  |  |  |  | 3 | 1 | 0 |  |  |  |  |  |  |  | 3 | 2 | 0 |  |  |  |  |  |  |  | 3 | 3 | 0 |  |  |  |  |  |  |  | 3 | 4 | 0 |  |  |  |  |  |  |  | 3 | 5 | 0 |  |  |  |  |  |  |  | 3 | 6 | 0 |  |  |  |  |  |  |  | 3 | 7 | 0 |  |  |  |  |  |  |  | 3 | 8 | 0 |  |  |  |  |  |  |  | 3 | 9 | 0 |  |  |  |  |  |  |  | 4 | 0 | 0 |  |  |  |  |  |  |  | 4 | 1 | 0 |  |  |  |  |  |  |  | 4 | 2 | 0 |  |  |  |  |  |  |  | 4 | 3 | 0 |  |  |  |  |  |  |  | 4 | 4 | 0 |  |  |  |  |  |  |  | 4 | 5 | 0 |  |  |  |  |  |  |  | 4 | 6 | 0 |  |  |  |  |  |  |  | 4 | 7 | 0 |  |  |  |  |  |  |  | 4 | 8 | 0 |  |  |  |  |  |  |  | 4 | 9 | 0 |  |  |  |  |  |  |  | 5 | 0 | 0 |  |  |  |  |  |  |  | 5 | 1 | 0 |  |  |  |  |  |  |  | 5 | 2 | 0 |  |  |  |  |  |  |  | 5 | 3 | 0 |  |  |  |  |  |  |  | 5 | 4 | 0 |  |  |  |  |  |  |  | 5 | 5 | 0 |  |  |  |  |  |  |  | 5 | 6 | 0 |  |  |  |  |  |  |  | 5 | 7 | 0 |  |  |  |  |  |  |  | 5 | 8 | 0 |  |  |  |  |  |  |  | 5 | 9 | 0 |  |  |  |  |  |  |  | 6 | 0 | 0 |  |  |  |  |  |  |  | 6 | 1 | 0 |  |  |  |  |  |  |  | 6 | 2 | 0 |  |  |  |  |  |  |  | 6 | 3 | 0 |  |  |  |  |  |  |  | 6 | 4 | 0 |  |  |  |  |  |  |  | 6 | 5 | 0 |  |  |  |  |  |  |  | 6 | 6 | 0 |  |  |  |  |  |  |  | 6 | 7 | 0 |  |  |  |  |  |  |  | 6 | 8 | 0 |  |  |  |  |  |  |  | 6 | 9 | 0 |  |  |  |  |  |  |  | 7 | 0 | 0 |  |  |  |  |  |  |  | 7 | 1 | 0 |  |  |  |  |  |  |  | 7 | 2 | 0 |  |  |  |  |  |  |  | 7 | 3 | 0 |  |  |  |  |  |  |  | 7 | 4 | 0 |  |  |  |  |  |  |  | 7 | 5 | 0 |  |  |  |  |  |  |  | 7 | 6 | 0 |  |  |  |  |  |  |  | 7 | 7 | 0 |  |  |  |  |  |  |  | 7 | 8 | 0 |  |  |  |  |  |  |  | 7 | 9 | 0 |  |  |  |  |  |  |  | 8 | 0 | 0 |  |  |  |  |  |  |  | 8 | 1 | 0 |  |  |  |  |  |  |  | 8 | 2 | 0 |  |  |  |  |  |  |  | 8 | 3 | 0 |  |  |  |  |  |  |  | 8 | 4 | 0 |  |  |  |  |  |  |  | 8 | 5 | 0 |  |  |  |  |  |  |  | 8 | 6 | 0 |  |  |  |  |  |  |  | 8 | 7 | 0 |  |  |  |  |  |  |  | 8 | 8 | 0 |  |  |  |  |  |  |  | 8 | 9 | 0 |  |  |  |  |  |  |  | 9 | 0 | 0 |  |  |  |  |  |  |  | 9 | 1 | 0 |  |  |  |  |  |  |  | 9 | 2 | 0 |  |  |  |  |  |  |  | 9 | 3 | 0 |  |  |  |  |  |  |  | 9 | 4 | 0 |  |  |  |  |  |  |  | 9 | 5 | 0 |  |  |  |  |  |  |  | 9 | 6 | 0 |  |  |  |  |  |  |  | 9 | 7 | 0 |  |  |  |  |  |  |  | 9 | 8 | 0 |  |  |  |  |  |  |  | 9 | 9 | 0 |  |  |  |  |  |  | 1 | 0 | 0 | 0 |  |  |  |  |  |  | 1 | 0 | 1 | 0 |  |  |  |  |  |  | 1 | 0 | 2 | 0 |  |  |  |  |  |  | 1 | 0 | 3 | 0 |  |  |  |  |  |  | 1 | 0 | 4 | 0 |  |  |  |  |  |  | 1 | 0 | 5 | 0 |  |  |  |  |  |  | 1 | 0 | 6 | 0 |  |  |  |  |  |  | 1 | 0 | 7 | 0 |  |  |  |  |  |  | 1 | 0 | 8 | 0 |  |  |  |  |  |  | 1 | 0 | 9 | 0 |  |  |  |  |  |  | 1 | 1 | 0 | 0 |  |  |  |  |  |  | 1 | 1 | 1 | 0 |  |  |  |  |  |  | 1 | 1 | 2 | 0 |  |  |  |  |  |  | 1 | 1 | 3 | 0 |  |  |  |  |  |  | 1 | 1 | 4 | 0 |  |  |  |  |  |  | 1 | 1 | 5 | 0 |  |  |  |  |  |  | 1 | 1 | 6 | 0 |  |  |  |  |  |  | 1 | 1 | 7 | 0 |  |  |  |  |  |  | 1 | 1 | 8 | 0 |  |  |  |  |  |  | 1 | 1 | 9 | 0 |  |  |  |  |  |  | 1 | 2 | 0 | 0 |  |  |  |  |  |  | 1 | 2 | 1 | 0 |  |  |  |  |  |  | 1 | 2 | 2 | 0 |  |  |  |  |  |  | 1 | 2 | 3 | 0 |  |  |  |  |  |  | 1 | 2 | 4 | 0 |  |  |  |  |  |  | 1 | 2 | 5 | 0 |  |  |  |  |  |  | 1 | 2 | 6 | 0 |  |  |  |  |  |  | 1 | 2 | 7 | 0 |  |  |  |  |  |  | 1 | 2 | 8 | 0 |  |  |  |  |  |  | 1 | 2 | 9 | 0 |  |  |  |  |  |  | 1 | 3 | 0 | 0 |  |  |  |  |  |  | 1 | 3 | 1 | 0 |  |  |  |  |  |  | 1 | 3 | 2 | 0 |  |  |  |  |  |  | 1 | 3 | 3 | 0 |  |  |  |  |  |  | 1 | 3 | 4 | 0 |  |  |  |  |  |  | 1 | 3 | 5 | 0 |  |  |  |  |  |  | 1 | 3 | 6 | 0 |  |  |  |  |  |  | 1 | 3 | 7 | 0 |  |  |  |  |  |  | 1 | 3 | 8 | 0 |  |  |  |  |  |  | 1 | 3 | 9 | 0 |  |  |  |  |  |  | 1 | 4 | 0 | 0 |  |  |  |  |  |  | 1 | 4 | 1 | 0 |  |  |  |  |  |  | 1 | 4 | 2 | 0 |  |  |  |  |  |  | 1 | 4 | 3 | 0 |  |  |  |  |  |  | 1 | 4 | 4 | 0 |  |  |  |  |  |  | 1 | 4 | 5 | 0 |  |  |  |  |  |  | 1 | 4 | 6 | 0 |  |  |  |  |  |  | 1 | 4 | 7 | 0 |  |  |  |  |  |  | 1 | 4 | 8 | 0 |  |  |  |  |  |  | 1 | 4 | 9 | 0 |  |  |  |  |  |  | 1 | 5 | 0 | 0 |  |  |  |  |  |  | 1 | 5 | 1 | 0 |  |  |  |  |  |  | 1 | 5 | 2 | 0 |  |  |  |  |  |  | 1 | 5 | 3 | 0 |  |  |  |  |  |  | 1 | 5 | 4 | 0 |  |  |  |  |  |  | 1 | 5 | 5 | 0 |  |  |  |  |  |  | 1 | 5 | 6 | 0 |  |  |  |  |  |  | 1 | 5 | 7 | 0 |  |  |  |  |  |  | 1 | 5 | 8 | 0 |  |  |  |  |  |  | 1 | 5 | 9 | 0 |  |  |  |  |  |  | 1 | 6 | 0 | 0 |  |  |  |  |  |  | 1 | 6 | 1 | 0 |  |  |  |  |  |  | 1 | 6 | 2 | 0 |  |  |  |  |  |  | 1 | 6 | 3 | 0 |  |  |  |  |  |  | 1 | 6 | 4 | 0 |  |  |  |  |  |  | 1 | 6 | 5 | 0 |  |  |  |  |  |  | 1 | 6 | 6 | 0 |  |  |  |  |  |  | 1 | 6 | 7 | 0 |  |  |  |  |  |  | 1 | 6 | 8 | 0 |  |  |  |  |  |  | 1 | 6 | 9 | 0 |  |  |  |  |  |  | 1 | 7 | 0 | 0 |  |  |  |  |  |  | 1 | 7 | 1 | 0 |  |  |  |  |  |  | 1 | 7 | 2 | 0 |  |  |  |  |  |  | 1 | 7 | 3 | 0 |  |  |  |  |  |  | 1 | 7 | 4 | 0 |  |  |  |  |  |  | 1 | 7 | 5 | 0 |  |  |  |  |  |  | 1 | 7 | 6 | 0 |  |  |  |  |  |  | 1 | 7 | 7 | 0 |  |  |  |  |  |  | 1 | 7 | 8 | 0 |  |  |  |  |  |  | 1 | 7 | 9 | 0 |  |  |  |  |  |  | 1 | 8 | 0 | 0 |  |  |  |  |  |  | 1 | 8 | 1 | 0 |  |  |  |  |  |  | 1 | 8 | 2 | 0 |  |  |  |  |  |  | 1 | 8 | 3 | 0 |  |  |  |  |  |  | 1 | 8 | 4 | 0 |  |  |  |  |  |  | 1 | 8 | 5 | 0 |  |  |  |  |  |  | 1 | 8 | 6 | 0 |  |  |  |  |  |  | 1 | 8 | 7 | 0 |  |  |  |  |  |  | 1 | 8 | 8 | 0 |  |  |  |  |  |  | 1 | 8 | 9 | 0 |  |  |  |  |  |  | 1 | 9 | 0 | 0 |  |  |  |  |  |  | 1 | 9 | 1 | 0 |  |  |  |  |  |  | 1 | 9 | 2 | 0 |  |  |  |  |  |  | 1 | 9 | 3 | 0 |  |  |  |  |  |  | 1 | 9 | 4 | 0 |  |  |  |  |  |  | 1 | 9 | 5 | 0 |  |  |  |  |  |  | 1 | 9 | 6 | 0 |  |  |  |  |  |  | 1 | 9 | 7 | 0 |  |  |  |  |  |  | 1 | 9 | 8 | 0 |  |  |  |  |  |  | 1 | 9 | 9 | 0 |  |  |  |  |  |  | 2 | 0 | 0 | 0 |  |  |  |  |  |  | 2 | 0 | 1 | 0 |  |  |  |  |  |  | 2 | 0 | 2 | 0 |  |  |  |  |  |  | 2 | 0 | 3 | 0 |  |  |  |  |  |  | 2 | 0 | 4 | 0 |  |  |  |  |  |  | 2 | 0 | 5 | 0 |  |  |  |  |  |  | 2 | 0 | 6 | 0 |  |  |  |  |  |  | 2 | 0 | 7 | 0 |  |  |  |  |  |  | 2 | 0 | 8 | 0 |  |  |  |  |  |  | 2 | 0 | 9 | 0 |  |  |  |  |  |  | 2 | 1 | 0 | 0 |  |  |  |  |  |  | 2 | 1 | 1 | 0 |  |  |  |  |  |  | 2 | 1 | 2 | 0 |  |  |  |  |  |  | 2 | 1 | 3 | 0 |  |  |  |  |  |  | 2 | 1 | 4 | 0 |  |  |  |  |  |  | 2 | 1 | 5 | 0 |  |  |  |  |  |  | 2 | 1 | 6 | 0 |  |  |  |  |  |  | 2 | 1 | 7 | 0 |  |  |  |  |  |  | 2 | 1 | 8 | 0 |  |  |  |  |  |  | 2 | 1 | 9 | 0 |  |  |  |  |  |  | 2 | 2 | 0 | 0 |  |  |  |  |  |  | 2 | 2 | 1 | 0 |  |  |  |  |  |  | 2 | 2 | 2 | 0 |  |  |  |  |  |  | 2 | 2 | 3 | 0 |  |  |  |  |  |  | 2 | 2 | 4 | 0 |  |  |  |  |  |  | 2 | 2 | 5 | 0 |  |  |  |  |  |  | 2 | 2 | 6 | 0 |  |  |  |  |  |  | 2 | 2 | 7 | 0 |  |  |  |  |  |  | 2 | 2 | 8 | 0 |  |  |  |  |  |  | 2 | 2 | 9 | 0 |  |  |  |  |  |  | 2 | 3 | 0 | 0 |  |  |  |  |  |  | 2 | 3 | 1 | 0 |  |  |  |  |  |  | 2 | 3 | 2 | 0 |  |  |  |  |  |  | 2 | 3 | 3 | 0 |  |  |  |  |  |  | 2 | 3 | 4 | 0 |  |  |  |  |  |  | 2 | 3 | 5 | 0 |  |  |  |  |  |  | 2 | 3 | 6 | 0 |  |  |  |  |  |  | 2 | 3 | 7 | 0 |  |  |  |  |  |  | 2 | 3 | 8 | 0 |  |  |  |  |  |  | 2 | 3 | 9 | 0 |  |  |  |  |  |  | 2 | 4 | 0 | 0 |  |  |  |  |  |  | 2 | 4 | 1 | 0 |  |  |  |  |  |  | 2 | 4 | 2 | 0 |  |  |  |  |  |  | 2 | 4 | 3 | 0 |  |  |  |  |  |  | 2 | 4 | 4 | 0 |  |  |  |  |  |  | 2 | 4 | 5 | 0 |  |  |  |  |  |  | 2 | 4 | 6 | 0 |  |  |  |  |  |  | 2 | 4 | 7 | 0 |  |  |  |  |  |  | 2 | 4 | 8 | 0 |  |  |  |  |  |  | 2 | 4 | 9 | 0 |  |  |  |  |  |  | 2 | 5 | 0 | 0 |  |  |  |  |  |  | 2 | 5 | 1 | 0 |  |  |  |  |  |  | 2 | 5 | 2 | 0 |  |  |  |  |  |  | 2 | 5 | 3 | 0 |  |  |  |  |  |  | 2 | 5 | 4 | 0 |  |  |  |  |  |  | 2 | 5 | 5 | 0 |  |  |  |  |  |  | 2 | 5 | 6 | 0 |  |  |  |  |  |  | 2 | 5 | 7 | 0 |  |  |  |  |  |  | 2 | 5 | 8 | 0 |  |  |  |  |  |  | 2 | 5 | 9 | 0 |  |  |  |  |  |  | 2 | 6 | 0 | 0 |  |  |  |  |  |  | 2 | 6 | 1 | 0 |  |  |  |  |  |  | 2 | 6 | 2 | 0 |  |  |  |  |  |  | 2 | 6 | 3 | 0 |  |  |  |  |  |  | 2 | 6 | 4 | 0 |  |  |  |  |  |  | 2 | 6 | 5 | 0 |  |  |  |  |  |  | 2 | 6 | 6 | 0 |  |  |  |  |  |  | 2 | 6 | 7 | 0 |  |  |  |  |  |  | 2 | 6 | 8 | 0 |  |  |  |  |  |  | 2 | 6 | 9 | 0 |  |  |  |  |  |  | 2 | 7 | 0 | 0 |  |  |  |  |  |  | 2 | 7 | 1 | 0 |  |  |  |  |  |  | 2 | 7 | 2 | 0 |  |  |  |  |  |  | 2 | 7 | 3 | 0 |  |  |  |  |  |  | 2 | 7 | 4 | 0 |  |  |  |  |  |  | 2 | 7 | 5 | 0 |  |  |  |  |  |  | 2 | 7 | 6 | 0 |  |  |  |  |  |  | 2 | 7 | 7 | 0 |  |  |  |  |  |  | 2 | 7 | 8 | 0 |  |  |  |  |  |  | 2 | 7 | 9 | 0 |  |  |  |  |  |  | 2 | 8 | 0 | 0 |  |  |  |  |  |  | 2 | 8 | 1 | 0 |  |  |  |  |  |  | 2 | 8 | 2 | 0 |  |  |  |  |  |  | 2 | 8 | 3 | 0 |  |  |  |  |  |  | 2 | 8 | 4 | 0 |  |  |  |  |  |  | 2 | 8 | 5 | 0 |  |  |  |  |  |  | 2 | 8 | 6 | 0 |  |  |  |  |  |  | 2 | 8 | 7 | 0 |  |  |  |  |  |  | 2 | 8 | 8 | 0 |  |  |  |  |  |  | 2 | 8 | 9 | 0 |  |  |  |  |  |  | 2 | 9 | 0 | 0 |  |  |  |  |  |  | 2 | 9 | 1 | 0 |  |  |  |  |  |  | 2 | 9 | 2 | 0 |  |  |  |  |  |  | 2 | 9 | 3 | 0 |  |  |  |  |  |  | 2 | 9 | 4 | 0 |  |  |  |  |  |  | 2 | 9 | 5 | 0 |  |  |  |  |  |  | 2 | 9 | 6 | 0 |  |  |  |  |  |  | 2 | 9 | 7 | 0 |  |  |  |  |  |  | 2 | 9 | 8 | 0 |  |  |  |  |  |  | 2 | 9 | 9 | 0 |  |  |  |  |  |  | 3 | 0 | 0 | 0 |  |  |  |  |  |  | 3 | 0 | 1 | 0 |  |  |  |  |  |  | 3 | 0 | 2 | 0 |  |  |  |  |  |  | 3 | 0 | 3 | 0 |  |  |  |  |  |  | 3 | 0 | 4 | 0 |  |  |  |  |  |  | 3 | 0 | 5 | 0 |  |  |  |  |  |  | 3 | 0 | 6 | 0 |  |  |  |  |  |  | 3 | 0 | 7 | 0 |  |  |  |  |  |  | 3 | 0 | 8 | 0 |  |  |  |  |  |  | 3 | 0 | 9 | 0 |  |  |  |  |  |  | 3 | 1 | 0 | 0 |  |  |  |  |  |  | 3 | 1 | 1 | 0 |  |  |  |  |  |  | 3 | 1 | 2 | 0 |  |  |  |  |  |  | 3 | 1 | 3 | 0 |  |  |  |  |  |  | 3 | 1 | 4 | 0 |  |  |  |  |  |  | 3 | 1 | 5 | 0 |  |  |  |  |  |  | 3 | 1 | 6 | 0 |  |  |  |  |  |  | 3 | 1 | 7 | 0 |  |  |  |  |  |  | 3 | 1 | 8 | 0 |  |  |  |  |  |  | 3 | 1 | 9 | 0 |  |  |  |  |  |  | 3 | 2 | 0 | 0 |  |  |  |  |  |  | 3 | 2 | 1 | 0 |  |  |  |  |  |  | 3 | 2 | 2 | 0 |  |  |  |  |  |  | 3 | 2 | 3 | 0 |  |  |  |  |  |  | 3 | 2 | 4 | 0 |  |  |  |  |  |  | 3 | 2 | 5 | 0 |  |  |  |  |  |  | 3 | 2 | 6 | 0 |  |  |  |  |  |  | 3 | 2 | 7 | 0 |  |  |  |  |  |  | 3 | 2 | 8 | 0 |  |  |  |  |  |  | 3 | 2 | 9 | 0 |  |  |  |  |  |  | 3 | 3 | 0 | 0 |  |  |  |  |  |  | 3 | 3 | 1 | 0 |  |  |  |  |  |  | 3 | 3 | 2 | 0 |  |  |  |  |  |  | 3 | 3 | 3 | 0 |  |  |  |  |  |  | 3 | 3 | 4 | 0 |  |  |  |  |  |  | 3 | 3 | 5 | 0 |  |  |  |  |  |  | 3 | 3 | 6 | 0 |  |  |  |  |  |  | 3 | 3 | 7 | 0 |  |  |  |  |  |  | 3 | 3 | 8 | 0 |  |  |  |  |  |  | 3 | 3 | 9 | 0 |  |  |  |  |  |  | 3 | 4 | 0 | 0 |  |  |  |  |  |  | 3 | 4 | 1 | 0 |  |  |  |  |  |  | 3 | 4 | 2 | 0 |  |  |  |  |  |  | 3 | 4 | 3 | 0 |  |  |  |  |  |  | 3 | 4 | 4 | 0 |  |  |  |  |  |  | 3 | 4 | 5 | 0 |  |  |  |  |  |  | 3 | 4 | 6 | 0 |  |  |  |  |  |  | 3 | 4 | 7 | 0 |  |  |  |  |  |  | 3 | 4 | 8 | 0 |  |  |  |  |  |  | 3 | 4 | 9 | 0 |  |  |  |  |  |  | 3 | 5 | 0 | 0 |  |  |  |  |  |  | 3 | 5 | 1 | 0 |  |  |  |  |  |  | 3 | 5 | 2 | 0 |  |  |  |  |  |  | 3 | 5 | 3 | 0 |  |  |  |  |  |  | 3 | 5 | 4 | 0 |  |  |  |  |  |  | 3 | 5 | 5 | 0 |  |  |  |  |  |  | 3 | 5 | 6 | 0 |  |  |  |  |  |  | 3 | 5 | 7 | 0 |  |  |  |  |  |  | 3 | 5 | 8 | 0 |  |  |  |  |  |  | 3 | 5 | 9 | 0 |  |  |  |  |  |  | 3 | 6 | 0 | 0 |  |  |  |  |  |  | 3 | 6 | 1 | 0 |  |  |  |  |  |  | 3 | 6 | 2 | 0 |  |  |  |  |  |  | 3 | 6 | 3 | 0 |  |  |  |  |  |  | 3 | 6 | 4 | 0 |  |  |  |  |  |  | 3 | 6 | 5 | 0 |  |  |  |  |  |  | 3 | 6 | 6 | 0 |  |  |  |  |  |  | 3 | 6 | 7 | 0 |  |  |  |  |  |  | 3 | 6 | 8 | 0 |  |  |  |  |  |  | 3 | 6 | 9 | 0 |  |  |  |  |  |  | 3 | 7 | 0 | 0 |  |  |  |  |  |  | 3 | 7 | 1 | 0 |  |  |  |  |  |  | 3 | 7 | 2 | 0 |  |  |  |  |  |  | 3 | 7 | 3 | 0 |  |  |  |  |  |  | 3 | 7 | 4 | 0 |  |  |  |  |  |  | 3 | 7 | 5 | 0 |  |  |  |  |  |  | 3 | 7 | 6 | 0 |  |  |  |  |  |  | 3 | 7 | 7 | 0 |  |  |  |  |  |  | 3 | 7 | 8 | 0 |  |  |  |  |  |  | 3 | 7 | 9 | 0 |  |  |  |  |  |  | 3 | 8 | 0 | 0 |  |  |  |  |  |  | 3 | 8 | 1 | 0 |  |  |  |  |  |  | 3 | 8 | 2 | 0 |  |  |  |  |  |  | 3 | 8 | 3 | 0 |  |  |  |  |  |  | 3 | 8 | 4 | 0 |  |  |  |  |  |  | 3 | 8 | 5 | 0 |  |  |  |  |  |  | 3 | 8 | 6 | 0 |  |  |  |  |  |  | 3 | 8 | 7 | 0 |  |  |  |  |  |  | 3 | 8 | 8 | 0 |  |  |  |  |  |  | 3 | 8 | 9 | 0 |  |  |  |  |  |  | 3 | 9 | 0 | 0 |  |  |  |  |  |  | 3 | 9 | 1 | 0 |  |  |  |  |  |  | 3 | 9 | 2 | 0 |  |  |  |  |  |  | 3 | 9 | 3 | 0 |  |  |  |  |  |  | 3 | 9 | 4 | 0 |  |  |  |  |  |  | 3 | 9 | 5 | 0 |  |  |  |  |  |  | 3 | 9 | 6 | 0 |  |  |  |  |  |  | 3 | 9 | 7 | 0 |  |  |  |  |  |  | 3 | 9 | 8 | 0 |  |  |  |  |  |  | 3 | 9 | 9 | 0 |  |  |  |  |  |  | 4 | 0 | 0 | 0 |  |  |  |  |  |  | 4 | 0 | 1 | 0 |  |  |  |  |  |  | 4 | 0 | 2 | 0 |  |  |  |  |  |  | 4 | 0 | 3 | 0 |  |  |  |  |  |  | 4 | 0 | 4 | 0 |  |  |  |  |  |  | 4 | 0 | 5 | 0 |  |  |  |  |  |  | 4 | 0 | 6 | 0 |  |  |  |  |  |  | 4 | 0 | 7 | 0 |  |  |  |  |  |  | 4 | 0 | 8 | 0 |  |  |  |  |  |  | 4 | 0 | 9 | 0 |  |  |  |  |  |  | 4 | 1 | 0 | 0 |  |  |  |  |  |  | 4 | 1 | 1 | 0 |  |  |  |  |  |  | 4 | 1 | 2 | 0 |  |  |  |  |  |  | 4 | 1 | 3 | 0 |  |  |  |  |  |  | 4 | 1 | 4 | 0 |  |  |  |  |  |  | 4 | 1 | 5 | 0 |  |  |  |  |  |  | 4 | 1 | 6 | 0 |  |  |  |  |  |  | 4 | 1 | 7 | 0 |  |  |  |  |  |  | 4 | 1 | 8 | 0 |  |  |  |  |  |  | 4 | 1 | 9 | 0 |  |  |  |  |  |  | 4 | 2 | 0 | 0 |  |  |  |  |  |  | 4 | 2 | 1 | 0 |  |  |  |  |  |  | 4 | 2 | 2 | 0 |  |  |  |  |  |  | 4 | 2 | 3 | 0 |  |  |  |  |  |  | 4 | 2 | 4 | 0 |  |  |  |  |  |  | 4 | 2 | 5 | 0 |  |  |  |  |  |  | 4 | 2 | 6 | 0 |  |  |  |  |  |  | 4 | 2 | 7 | 0 |  |  |  |  |  |  | 4 | 2 | 8 | 0 |  |  |  |  |  |  | 4 | 2 | 9 | 0 |  |  |  |  |  |  | 4 | 3 | 0 | 0 |  |  |  |  |  |  | 4 | 3 | 1 | 0 |  |  |  |  |  |  | 4 | 3 | 2 | 0 |  |  |  |  |  |  | 4 | 3 | 3 | 0 |  |  |  |  |  |  | 4 | 3 | 4 | 0 |  |  |  |  |  |  | 4 | 3 | 5 | 0 |  |  |  |  |  |  | 4 | 3 | 6 | 0 |  |  |  |  |  |  | 4 | 3 | 7 | 0 |  |  |  |  |  |  | 4 | 3 | 8 | 0 |  |  |  |  |  |  | 4 | 3 | 9 | 0 |  |  |  |  |  |  | 4 | 4 | 0 | 0 |  |  |  |  |  |  | 4 | 4 | 1 | 0 |  |  |  |  |  |  | 4 | 4 | 2 | 0 |  |  |  |  |  |  | 4 | 4 | 3 | 0 |  |  |  |  |  |  | 4 | 4 | 4 | 0 |  |  |  |  |  |  | 4 | 4 | 5 | 0 |  |  |  |  |  |  | 4 | 4 | 6 | 0 |  |  |  |  |  |  | 4 | 4 | 7 | 0 |  |  |  |  |  |  | 4 | 4 | 8 | 0 |  |  |  |  |  |  | 4 | 4 | 9 | 0 |  |  |  |  |  |  | 4 | 5 | 0 | 0 |  |  |  |  |  |  | 4 | 5 | 1 | 0 |  |  |  |  |  |  | 4 | 5 | 2 | 0 |  |  |  |  |  |  | 4 | 5 | 3 | 0 |  |  |  |  |  |  | 4 | 5 | 4 | 0 |  |  |  |  |  |  | 4 | 5 | 5 | 0 |  |  |  |  |  |  | 4 | 5 | 6 | 0 |  |  |  |  |  |  | 4 | 5 | 7 | 0 |  |  |  |  |  |  | 4 | 5 | 8 | 0 |  |  |  |  |  |  | 4 | 5 | 9 | 0 |  |  |  |  |  |  | 4 | 6 | 0 | 0 |  |  |  |  |  |  | 4 | 6 | 1 | 0 |  |  |  |  |  |  | 4 | 6 | 2 | 0 |  |  |  |  |  |  | 4 | 6 | 3 | 0 |  |  |  |  |  |  | 4 | 6 | 4 | 0 |  |  |  |  |  |  | 4 | 6 | 5 | 0 |  |  |  |  |  |  | 4 | 6 | 6 | 0 |  |  |  |  |  |  | 4 | 6 | 7 | 0 |  |  |  |  |  |  | 4 | 6 | 8 | 0 |  |  |  |  |  |  | 4 | 6 | 9 | 0 |  |  |  |  |  |  | 4 | 7 | 0 | 0 |  |  |  |  |  |  | 4 | 7 | 1 | 0 |  |  |  |  |  |  | 4 | 7 | 2 | 0 |  |  |  |  |  |  | 4 | 7 | 3 | 0 |  |  |  |  |  |  | 4 | 7 | 4 | 0 |  |  |  |  |  |  | 4 | 7 | 5 | 0 |  |  |  |  |  |  | 4 | 7 | 6 | 0 |  |  |  |  |  |  | 4 | 7 | 7 | 0 |  |  |  |  |  |  | 4 | 7 | 8 | 0 |  |  |  |  |  |  | 4 | 7 | 9 | 0 |  |  |  |  |  |  | 4 | 8 | 0 | 0 |  |  |  |  |  |  | 4 | 8 | 1 | 0 |  |  |  |  |  |  | 4 | 8 | 2 | 0 |  |  |  |  |  |  | 4 | 8 | 3 | 0 |  |  |  |  |  |  | 4 | 8 | 4 | 0 |  |  |  |  |  |  | 4 | 8 | 5 | 0 |  |  |  |  |  |  | 4 | 8 | 6 | 0 |  |  |  |  |  |  | 4 | 8 | 7 | 0 |  |  |  |  |  |  | 4 | 8 | 8 | 0 |  |  |  |  |  |  | 4 | 8 | 9 | 0 |  |  |  |  |  |  | 4 | 9 | 0 | 0 |  |  |  |  |  |  | 4 | 9 | 1 | 0 |  |  |  |  |  |  | 4 | 9 | 2 | 0 |  |  |  |  |  |  | 4 | 9 | 3 | 0 |  |  |  |  |  |  | 4 | 9 | 4 | 0 |  |  |  |  |  |  | 4 | 9 | 5 | 0 |  |  |  |  |  |  | 4 | 9 | 6 | 0 |  |  |  |  |  |  | 4 | 9 | 7 | 0 |  |  |  |  |  |  | 4 | 9 | 8 | 0 |  |  |  |  |  |  | 4 | 9 | 9 | 0 |  |  |  |  |  |  | 5 | 0 | 0 | 0 |  |  |  |  |  |  | 5 | 0 | 1 | 0 |  |  |  |  |  |  | 5 | 0 | 2 | 0 |  |  |  |  |  |  | 5 | 0 | 3 | 0 |  |  |  |  |  |  | 5 | 0 | 4 | 0 |  |  |  |  |  |  | 5 | 0 | 5 | 0 |  |  |  |  |  |  | 5 | 0 | 6 | 0 |  |  |  |  |  |  | 5 | 0 | 7 | 0 |  |  |  |  |  |  | 5 | 0 | 8 | 0 |  |  |  |  |  |  | 5 | 0 | 9 | 0 |  |  |  |  |  |  | 5 | 1 | 0 | 0 |  |  |  |  |  |  | 5 | 1 | 1 | 0 |  |  |  |  |  |  | 5 | 1 | 2 | 0 |  |  |  |  |  |  | 5 | 1 | 3 | 0 |  |  |  |  |  |  | 5 | 1 | 4 | 0 |  |  |  |  |  |  | 5 | 1 | 5 | 0 |  |  |  |  |  |  | 5 | 1 | 6 | 0 |  |  |  |  |  |  | 5 | 1 | 7 | 0 |  |  |  |  |  |  | 5 | 1 | 8 | 0 |  |  |  |  |  |  | 5 | 1 | 9 | 0 |  |  |  |  |  |  | 5 | 2 | 0 | 0 |  |  |  |  |  |  | 5 | 2 | 1 | 0 |  |  |  |  |  |  | 5 | 2 | 2 | 0 |  |  |  |  |  |  | 5 | 2 | 3 | 0 |  |  |  |  |  |  | 5 | 2 | 4 | 0 |  |  |  |  |  |  | 5 | 2 | 5 | 0 |  |  |  |  |  |  | 5 | 2 | 6 | 0 |  |  |  |  |  |  | 5 | 2 | 7 | 0 |  |  |  |  |  |  | 5 | 2 | 8 | 0 |  |  |  |  |  |  | 5 | 2 | 9 | 0 |  |  |  |  |  |  | 5 | 3 | 0 | 0 |  |  |  |  |  |  | 5 | 3 | 1 | 0 |  |  |  |  |  |  | 5 | 3 | 2 | 0 |  |  |  |  |  |  | 5 | 3 | 3 | 0 |  |  |  |  |  |  | 5 | 3 | 4 | 0 |  |  |  |  |  |  | 5 | 3 | 5 | 0 |  |  |  |  |  |  | 5 | 3 | 6 | 0 |  |  |  |  |  |  | 5 | 3 | 7 | 0 |  |  |  |  |  |  | 5 | 3 | 8 | 0 |  |  |  |  |  |  | 5 | 3 | 9 | 0 |  |  |  |  |  |  | 5 | 4 | 0 | 0 |  |  |  |  |  |  | 5 | 4 | 1 | 0 |  |  |  |  |  |  | 5 | 4 | 2 | 0 |  |  |  |  |  |  |  |  |
|  |  |  |  |  |  |  |  |  | \| |  |  |  |  |  |  |  |  |  | \| |  |  |  |  |  |  |  |  |  | \| |  |  |  |  |  |  |  |  |  | \| |  |  |  |  |  |  |  |  |  | \| |  |  |  |  |  |  |  |  |  | \| |  |  |  |  |  |  |  |  |  | \| |  |  |  |  |  |  |  |  |  | \| |  |  |  |  |  |  |  |  |  | \| |  |  |  |  |  |  |  |  |  | \| |  |  |  |  |  |  |  |  |  | \| |  |  |  |  |  |  |  |  |  | \| |  |  |  |  |  |  |  |  |  | \| |  |  |  |  |  |  |  |  |  | \| |  |  |  |  |  |  |  |  |  | \| |  |  |  |  |  |  |  |  |  | \| |  |  |  |  |  |  |  |  |  | \| |  |  |  |  |  |  |  |  |  | \| |  |  |  |  |  |  |  |  |  | \| |  |  |  |  |  |  |  |  |  | \| |  |  |  |  |  |  |  |  |  | \| |  |  |  |  |  |  |  |  |  | \| |  |  |  |  |  |  |  |  |  | \| |  |  |  |  |  |  |  |  |  | \| |  |  |  |  |  |  |  |  |  | \| |  |  |  |  |  |  |  |  |  | \| |  |  |  |  |  |  |  |  |  | \| |  |  |  |  |  |  |  |  |  | \| |  |  |  |  |  |  |  |  |  | \| |  |  |  |  |  |  |  |  |  | \| |  |  |  |  |  |  |  |  |  | \| |  |  |  |  |  |  |  |  |  | \| |  |  |  |  |  |  |  |  |  | \| |  |  |  |  |  |  |  |  |  | \| |  |  |  |  |  |  |  |  |  | \| |  |  |  |  |  |  |  |  |  | \| |  |  |  |  |  |  |  |  |  | \| |  |  |  |  |  |  |  |  |  | \| |  |  |  |  |  |  |  |  |  | \| |  |  |  |  |  |  |  |  |  | \| |  |  |  |  |  |  |  |  |  | \| |  |  |  |  |  |  |  |  |  | \| |  |  |  |  |  |  |  |  |  | \| |  |  |  |  |  |  |  |  |  | \| |  |  |  |  |  |  |  |  |  | \| |  |  |  |  |  |  |  |  |  | \| |  |  |  |  |  |  |  |  |  | \| |  |  |  |  |  |  |  |  |  | \| |  |  |  |  |  |  |  |  |  | \| |  |  |  |  |  |  |  |  |  | \| |  |  |  |  |  |  |  |  |  | \| |  |  |  |  |  |  |  |  |  | \| |  |  |  |  |  |  |  |  |  | \| |  |  |  |  |  |  |  |  |  | \| |  |  |  |  |  |  |  |  |  | \| |  |  |  |  |  |  |  |  |  | \| |  |  |  |  |  |  |  |  |  | \| |  |  |  |  |  |  |  |  |  | \| |  |  |  |  |  |  |  |  |  | \| |  |  |  |  |  |  |  |  |  | \| |  |  |  |  |  |  |  |  |  | \| |  |  |  |  |  |  |  |  |  | \| |  |  |  |  |  |  |  |  |  | \| |  |  |  |  |  |  |  |  |  | \| |  |  |  |  |  |  |  |  |  | \| |  |  |  |  |  |  |  |  |  | \| |  |  |  |  |  |  |  |  |  | \| |  |  |  |  |  |  |  |  |  | \| |  |  |  |  |  |  |  |  |  | \| |  |  |  |  |  |  |  |  |  | \| |  |  |  |  |  |  |  |  |  | \| |  |  |  |  |  |  |  |  |  | \| |  |  |  |  |  |  |  |  |  | \| |  |  |  |  |  |  |  |  |  | \| |  |  |  |  |  |  |  |  |  | \| |  |  |  |  |  |  |  |  |  | \| |  |  |  |  |  |  |  |  |  | \| |  |  |  |  |  |  |  |  |  | \| |  |  |  |  |  |  |  |  |  | \| |  |  |  |  |  |  |  |  |  | \| |  |  |  |  |  |  |  |  |  | \| |  |  |  |  |  |  |  |  |  | \| |  |  |  |  |  |  |  |  |  | \| |  |  |  |  |  |  |  |  |  | \| |  |  |  |  |  |  |  |  |  | \| |  |  |  |  |  |  |  |  |  | \| |  |  |  |  |  |  |  |  |  | \| |  |  |  |  |  |  |  |  |  | \| |  |  |  |  |  |  |  |  |  | \| |  |  |  |  |  |  |  |  |  | \| |  |  |  |  |  |  |  |  |  | \| |  |  |  |  |  |  |  |  |  | \| |  |  |  |  |  |  |  |  |  | \| |  |  |  |  |  |  |  |  |  | \| |  |  |  |  |  |  |  |  |  | \| |  |  |  |  |  |  |  |  |  | \| |  |  |  |  |  |  |  |  |  | \| |  |  |  |  |  |  |  |  |  | \| |  |  |  |  |  |  |  |  |  | \| |  |  |  |  |  |  |  |  |  | \| |  |  |  |  |  |  |  |  |  | \| |  |  |  |  |  |  |  |  |  | \| |  |  |  |  |  |  |  |  |  | \| |  |  |  |  |  |  |  |  |  | \| |  |  |  |  |  |  |  |  |  | \| |  |  |  |  |  |  |  |  |  | \| |  |  |  |  |  |  |  |  |  | \| |  |  |  |  |  |  |  |  |  | \| |  |  |  |  |  |  |  |  |  | \| |  |  |  |  |  |  |  |  |  | \| |  |  |  |  |  |  |  |  |  | \| |  |  |  |  |  |  |  |  |  | \| |  |  |  |  |  |  |  |  |  | \| |  |  |  |  |  |  |  |  |  | \| |  |  |  |  |  |  |  |  |  | \| |  |  |  |  |  |  |  |  |  | \| |  |  |  |  |  |  |  |  |  | \| |  |  |  |  |  |  |  |  |  | \| |  |  |  |  |  |  |  |  |  | \| |  |  |  |  |  |  |  |  |  | \| |  |  |  |  |  |  |  |  |  | \| |  |  |  |  |  |  |  |  |  | \| |  |  |  |  |  |  |  |  |  | \| |  |  |  |  |  |  |  |  |  | \| |  |  |  |  |  |  |  |  |  | \| |  |  |  |  |  |  |  |  |  | \| |  |  |  |  |  |  |  |  |  | \| |  |  |  |  |  |  |  |  |  | \| |  |  |  |  |  |  |  |  |  | \| |  |  |  |  |  |  |  |  |  | \| |  |  |  |  |  |  |  |  |  | \| |  |  |  |  |  |  |  |  |  | \| |  |  |  |  |  |  |  |  |  | \| |  |  |  |  |  |  |  |  |  | \| |  |  |  |  |  |  |  |  |  | \| |  |  |  |  |  |  |  |  |  | \| |  |  |  |  |  |  |  |  |  | \| |  |  |  |  |  |  |  |  |  | \| |  |  |  |  |  |  |  |  |  | \| |  |  |  |  |  |  |  |  |  | \| |  |  |  |  |  |  |  |  |  | \| |  |  |  |  |  |  |  |  |  | \| |  |  |  |  |  |  |  |  |  | \| |  |  |  |  |  |  |  |  |  | \| |  |  |  |  |  |  |  |  |  | \| |  |  |  |  |  |  |  |  |  | \| |  |  |  |  |  |  |  |  |  | \| |  |  |  |  |  |  |  |  |  | \| |  |  |  |  |  |  |  |  |  | \| |  |  |  |  |  |  |  |  |  | \| |  |  |  |  |  |  |  |  |  | \| |  |  |  |  |  |  |  |  |  | \| |  |  |  |  |  |  |  |  |  | \| |  |  |  |  |  |  |  |  |  | \| |  |  |  |  |  |  |  |  |  | \| |  |  |  |  |  |  |  |  |  | \| |  |  |  |  |  |  |  |  |  | \| |  |  |  |  |  |  |  |  |  | \| |  |  |  |  |  |  |  |  |  | \| |  |  |  |  |  |  |  |  |  | \| |  |  |  |  |  |  |  |  |  | \| |  |  |  |  |  |  |  |  |  | \| |  |  |  |  |  |  |  |  |  | \| |  |  |  |  |  |  |  |  |  | \| |  |  |  |  |  |  |  |  |  | \| |  |  |  |  |  |  |  |  |  | \| |  |  |  |  |  |  |  |  |  | \| |  |  |  |  |  |  |  |  |  | \| |  |  |  |  |  |  |  |  |  | \| |  |  |  |  |  |  |  |  |  | \| |  |  |  |  |  |  |  |  |  | \| |  |  |  |  |  |  |  |  |  | \| |  |  |  |  |  |  |  |  |  | \| |  |  |  |  |  |  |  |  |  | \| |  |  |  |  |  |  |  |  |  | \| |  |  |  |  |  |  |  |  |  | \| |  |  |  |  |  |  |  |  |  | \| |  |  |  |  |  |  |  |  |  | \| |  |  |  |  |  |  |  |  |  | \| |  |  |  |  |  |  |  |  |  | \| |  |  |  |  |  |  |  |  |  | \| |  |  |  |  |  |  |  |  |  | \| |  |  |  |  |  |  |  |  |  | \| |  |  |  |  |  |  |  |  |  | \| |  |  |  |  |  |  |  |  |  | \| |  |  |  |  |  |  |  |  |  | \| |  |  |  |  |  |  |  |  |  | \| |  |  |  |  |  |  |  |  |  | \| |  |  |  |  |  |  |  |  |  | \| |  |  |  |  |  |  |  |  |  | \| |  |  |  |  |  |  |  |  |  | \| |  |  |  |  |  |  |  |  |  | \| |  |  |  |  |  |  |  |  |  | \| |  |  |  |  |  |  |  |  |  | \| |  |  |  |  |  |  |  |  |  | \| |  |  |  |  |  |  |  |  |  | \| |  |  |  |  |  |  |  |  |  | \| |  |  |  |  |  |  |  |  |  | \| |  |  |  |  |  |  |  |  |  | \| |  |  |  |  |  |  |  |  |  | \| |  |  |  |  |  |  |  |  |  | \| |  |  |  |  |  |  |  |  |  | \| |  |  |  |  |  |  |  |  |  | \| |  |  |  |  |  |  |  |  |  | \| |  |  |  |  |  |  |  |  |  | \| |  |  |  |  |  |  |  |  |  | \| |  |  |  |  |  |  |  |  |  | \| |  |  |  |  |  |  |  |  |  | \| |  |  |  |  |  |  |  |  |  | \| |  |  |  |  |  |  |  |  |  | \| |  |  |  |  |  |  |  |  |  | \| |  |  |  |  |  |  |  |  |  | \| |  |  |  |  |  |  |  |  |  | \| |  |  |  |  |  |  |  |  |  | \| |  |  |  |  |  |  |  |  |  | \| |  |  |  |  |  |  |  |  |  | \| |  |  |  |  |  |  |  |  |  | \| |  |  |  |  |  |  |  |  |  | \| |  |  |  |  |  |  |  |  |  | \| |  |  |  |  |  |  |  |  |  | \| |  |  |  |  |  |  |  |  |  | \| |  |  |  |  |  |  |  |  |  | \| |  |  |  |  |  |  |  |  |  | \| |  |  |  |  |  |  |  |  |  | \| |  |  |  |  |  |  |  |  |  | \| |  |  |  |  |  |  |  |  |  | \| |  |  |  |  |  |  |  |  |  | \| |  |  |  |  |  |  |  |  |  | \| |  |  |  |  |  |  |  |  |  | \| |  |  |  |  |  |  |  |  |  | \| |  |  |  |  |  |  |  |  |  | \| |  |  |  |  |  |  |  |  |  | \| |  |  |  |  |  |  |  |  |  | \| |  |  |  |  |  |  |  |  |  | \| |  |  |  |  |  |  |  |  |  | \| |  |  |  |  |  |  |  |  |  | \| |  |  |  |  |  |  |  |  |  | \| |  |  |  |  |  |  |  |  |  | \| |  |  |  |  |  |  |  |  |  | \| |  |  |  |  |  |  |  |  |  | \| |  |  |  |  |  |  |  |  |  | \| |  |  |  |  |  |  |  |  |  | \| |  |  |  |  |  |  |  |  |  | \| |  |  |  |  |  |  |  |  |  | \| |  |  |  |  |  |  |  |  |  | \| |  |  |  |  |  |  |  |  |  | \| |  |  |  |  |  |  |  |  |  | \| |  |  |  |  |  |  |  |  |  | \| |  |  |  |  |  |  |  |  |  | \| |  |  |  |  |  |  |  |  |  | \| |  |  |  |  |  |  |  |  |  | \| |  |  |  |  |  |  |  |  |  | \| |  |  |  |  |  |  |  |  |  | \| |  |  |  |  |  |  |  |  |  | \| |  |  |  |  |  |  |  |  |  | \| |  |  |  |  |  |  |  |  |  | \| |  |  |  |  |  |  |  |  |  | \| |  |  |  |  |  |  |  |  |  | \| |  |  |  |  |  |  |  |  |  | \| |  |  |  |  |  |  |  |  |  | \| |  |  |  |  |  |  |  |  |  | \| |  |  |  |  |  |  |  |  |  | \| |  |  |  |  |  |  |  |  |  | \| |  |  |  |  |  |  |  |  |  | \| |  |  |  |  |  |  |  |  |  | \| |  |  |  |  |  |  |  |  |  | \| |  |  |  |  |  |  |  |  |  | \| |  |  |  |  |  |  |  |  |  | \| |  |  |  |  |  |  |  |  |  | \| |  |  |  |  |  |  |  |  |  | \| |  |  |  |  |  |  |  |  |  | \| |  |  |  |  |  |  |  |  |  | \| |  |  |  |  |  |  |  |  |  | \| |  |  |  |  |  |  |  |  |  | \| |  |  |  |  |  |  |  |  |  | \| |  |  |  |  |  |  |  |  |  | \| |  |  |  |  |  |  |  |  |  | \| |  |  |  |  |  |  |  |  |  | \| |  |  |  |  |  |  |  |  |  | \| |  |  |  |  |  |  |  |  |  | \| |  |  |  |  |  |  |  |  |  | \| |  |  |  |  |  |  |  |  |  | \| |  |  |  |  |  |  |  |  |  | \| |  |  |  |  |  |  |  |  |  | \| |  |  |  |  |  |  |  |  |  | \| |  |  |  |  |  |  |  |  |  | \| |  |  |  |  |  |  |  |  |  | \| |  |  |  |  |  |  |  |  |  | \| |  |  |  |  |  |  |  |  |  | \| |  |  |  |  |  |  |  |  |  | \| |  |  |  |  |  |  |  |  |  | \| |  |  |  |  |  |  |  |  |  | \| |  |  |  |  |  |  |  |  |  | \| |  |  |  |  |  |  |  |  |  | \| |  |  |  |  |  |  |  |  |  | \| |  |  |  |  |  |  |  |  |  | \| |  |  |  |  |  |  |  |  |  | \| |  |  |  |  |  |  |  |  |  | \| |  |  |  |  |  |  |  |  |  | \| |  |  |  |  |  |  |  |  |  | \| |  |  |  |  |  |  |  |  |  | \| |  |  |  |  |  |  |  |  |  | \| |  |  |  |  |  |  |  |  |  | \| |  |  |  |  |  |  |  |  |  | \| |  |  |  |  |  |  |  |  |  | \| |  |  |  |  |  |  |  |  |  | \| |  |  |  |  |  |  |  |  |  | \| |  |  |  |  |  |  |  |  |  | \| |  |  |  |  |  |  |  |  |  | \| |  |  |  |  |  |  |  |  |  | \| |  |  |  |  |  |  |  |  |  | \| |  |  |  |  |  |  |  |  |  | \| |  |  |  |  |  |  |  |  |  | \| |  |  |  |  |  |  |  |  |  | \| |  |  |  |  |  |  |  |  |  | \| |  |  |  |  |  |  |  |  |  | \| |  |  |  |  |  |  |  |  |  | \| |  |  |  |  |  |  |  |  |  | \| |  |  |  |  |  |  |  |  |  | \| |  |  |  |  |  |  |  |  |  | \| |  |  |  |  |  |  |  |  |  | \| |  |  |  |  |  |  |  |  |  | \| |  |  |  |  |  |  |  |  |  | \| |  |  |  |  |  |  |  |  |  | \| |  |  |  |  |  |  |  |  |  | \| |  |  |  |  |  |  |  |  |  | \| |  |  |  |  |  |  |  |  |  | \| |  |  |  |  |  |  |  |  |  | \| |  |  |  |  |  |  |  |  |  | \| |  |  |  |  |  |  |  |  |  | \| |  |  |  |  |  |  |  |  |  | \| |  |  |  |  |  |  |  |  |  | \| |  |  |  |  |  |  |  |  |  | \| |  |  |  |  |  |  |  |  |  | \| |  |  |  |  |  |  |  |  |  | \| |  |  |  |  |  |  |  |  |  | \| |  |  |  |  |  |  |  |  |  | \| |  |  |  |  |  |  |  |  |  | \| |  |  |  |  |  |  |  |  |  | \| |  |  |  |  |  |  |  |  |  | \| |  |  |  |  |  |  |  |  |  | \| |  |  |  |  |  |  |  |  |  | \| |  |  |  |  |  |  |  |  |  | \| |  |  |  |  |  |  |  |  |  | \| |  |  |  |  |  |  |  |  |  | \| |  |  |  |  |  |  |  |  |  | \| |  |  |  |  |  |  |  |  |  | \| |  |  |  |  |  |  |  |  |  | \| |  |  |  |  |  |  |  |  |  | \| |  |  |  |  |  |  |  |  |  | \| |  |  |  |  |  |  |  |  |  | \| |  |  |  |  |  |  |  |  |  | \| |  |  |  |  |  |  |  |  |  | \| |  |  |  |  |  |  |  |  |  | \| |  |  |  |  |  |  |  |  |  | \| |  |  |  |  |  |  |  |  |  | \| |  |  |  |  |  |  |  |  |  | \| |  |  |  |  |  |  |  |  |  | \| |  |  |  |  |  |  |  |  |  | \| |  |  |  |  |  |  |  |  |  | \| |  |  |  |  |  |  |  |  |  | \| |  |  |  |  |  |  |  |  |  | \| |  |  |  |  |  |  |  |  |  | \| |  |  |  |  |  |  |  |  |  | \| |  |  |  |  |  |  |  |  |  | \| |  |  |  |  |  |  |  |  |  | \| |  |  |  |  |  |  |  |  |  | \| |  |  |  |  |  |  |  |  |  | \| |  |  |  |  |  |  |  |  |  | \| |  |  |  |  |  |  |  |  |  | \| |  |  |  |  |  |  |  |  |  | \| |  |  |  |  |  |  |  |  |  | \| |  |  |  |  |  |  |  |  |  | \| |  |  |  |  |  |  |  |  |  | \| |  |  |  |  |  |  |  |  |  | \| |  |  |  |  |  |  |  |  |  | \| |  |  |  |  |  |  |  |  |  | \| |  |  |  |  |  |  |  |  |  | \| |  |  |  |  |  |  |  |  |  | \| |  |  |  |  |  |  |  |  |  | \| |  |  |  |  |  |  |  |  |  | \| |  |  |  |  |  |  |  |  |  | \| |  |  |  |  |  |  |  |  |  | \| |  |  |  |  |  |  |  |  |  | \| |  |  |  |  |  |  |  |  |  | \| |  |  |  |  |  |  |  |  |  | \| |  |  |  |  |  |  |  |  |  | \| |  |  |  |  |  |  |  |  |  | \| |  |  |  |  |  |  |  |  |  | \| |  |  |  |  |  |  |  |  |  | \| |  |  |  |  |  |  |  |  |  | \| |  |  |  |  |  |  |  |  |  | \| |  |  |  |  |  |  |  |  |  | \| |  |  |  |  |  |  |  |  |  | \| |  |  |  |  |  |  |  |  |  | \| |  |  |  |  |  |  |  |  |  | \| |  |  |  |  |  |  |  |  |  | \| |  |  |  |  |  |  |  |  |  | \| |  |  |  |  |  |  |  |  |  | \| |  |  |  |  |  |  |  |  |  | \| |  |  |  |  |  |  |  |  |  | \| |  |  |  |  |  |  |  |  |  | \| |  |  |  |  |  |  |  |  |  | \| |  |  |  |  |  |  |  |  |  | \| |  |  |  |  |  |  |  |  |  | \| |  |  |  |  |  |  |  |  |  | \| |  |  |  |  |  |  |  |  |  | \| |  |  |  |  |  |  |  |  |  | \| |  |  |  |  |  |  |  |  |  | \| |  |  |  |  |  |  |  |  |  | \| |  |  |  |  |  |  |  |  |  | \| |  |  |  |  |  |  |  |  |  | \| |  |  |  |  |  |  |  |  |  | \| |  |  |  |  |  |  |  |  |  | \| |  |  |  |  |  |  |  |  |  | \| |  |  |  |  |  |  |  |  |  | \| |  |  |  |  |  |  |  |  |  | \| |  |  |  |  |  |  |  |  |  | \| |  |  |  |  |  |  |  |  |  | \| |  |  |  |  |  |  |  |  |  | \| |  |  |  |  |  |  |  |  |  | \| |  |  |  |  |  |  |  |  |  | \| |  |  |  |  |  |  |  |  |  | \| |  |  |  |  |  |  |  |  |  | \| |  |  |  |  |  |  |  |  |  | \| |  |  |  |  |  |  |  |  |  | \| |  |  |  |  |  |  |  |  |  | \| |  |  |  |  |  |  |  |  |  | \| |  |  |  |  |  |  |  |  |  | \| |  |  |  |  |  |  |  |  |  | \| |  |  |  |  |  |  |  |  |  | \| |  |  |  |  |  |  |  |  |  | \| |  |  |  |  |  |  |  |  |  | \| |  |  |  |  |  |  |  |  |  | \| |  |  |  |  |  |  |  |  |  | \| |  |  |  |  |  |  |  |  |  | \| |  |  |  |  |  |  |  |  |  | \| |  |  |  |  |  |  |  |  |  | \| |  |  |  |  |  |  |  |  |  | \| |  |  |  |  |  |  |  |  |  | \| |  |  |  |  |  |  |  |  |  | \| |  |  |  |  |  |  |  |  |  | \| |  |  |  |  |  |  |  |  |  | \| |  |  |  |  |  |  |  |  |  | \| |  |  |  |  |  |  |  |  |  | \| |  |  |  |  |  |  |  |  |  | \| |  |  |  |  |  |  |  |  |  | \| |  |  |  |  |  |  |  |  |  | \| |  |  |  |  |  |  |  |  |  | \| |  |  |  |  |  |  |  |  |  | \| |  |  |  |  |  |  |  |  |  | \| |  |  |  |  |  |  |  |  |  | \| |  |  |  |  |  |  |  |  |  | \| |  |  |  |  |  |  |  |  |  | \| |  |  |  |  |  |  |  |  |  | \| |  |  |  |  |  |  |  |  |  | \| |  |  |  |  |  |  |  |  |  | \| |  |  |  |  |  |  |  |  |  | \| |  |  |  |  |  |  |  |  |  | \| |  |  |  |  |  |  |  |  |  | \| |  |  |  |  |  |  |  |  |  | \| |  |  |  |  |  |  |  |  |  | \| |  |  |  |  |  |  |  |  |  | \| |  |  |  |  |  |  |  |  |  | \| |  |  |  |  |  |  |  |  |  | \| |  |  |  |  |  |  |  |  |  | \| |  |  |  |  |  |  |  |  |  | \| |  |  |  |  |  |  |  |  |  | \| |  |  |  |  |  |  |  |  |  | \| |  |  |  |  |  |  |  |  |  | \| |  |  |  |  |  |  |  |  |  | \| |  |  |  |  |  |  |  |  |  | \| |  |  |  |  |  |  |  |  |  | \| |  |  |  |  |  |  |  |  |  | \| |  |  |  |  |  |  |  |  |  | \| |  |  |  |  |  |  |  |  |  | \| |  |  |  |  |  |  |  |  |  | \| |  |  |  |  |  |  |  |  |  | \| |  |  |  |  |  |  |  |  |  | \| |  |  |  |  |  |  |  |  |  | \| |  |  |  |  |  |  |  |  |  | \| |  |  |  |  |  |  |  |  |  | \| |  |  |  |  |  |  |  |  |  | \| |  |  |  |  |  |  |  |  |  | \| |  |  |  |  |  |  |  |  |  | \| |  |  |  |  |  |  |  |  |  | \| |  |  |  |  |  |  |  |  |  | \| |  |  |  |  |  |  |  |  |  | \| |  |  |  |  |  |  |  |  |  | \| |  |  |  |  |  |  |  |  |  | \| |  |  |  |  |  |  |  |  |  | \| |  |  |  |  |  |  |  |  |  | \| |  |  |  |  |  |  |  |  |  | \| |  |  |  |  |  |  |  |  |  | \| |  |  |  |  |  |  |  |  |  | \| |  |  |  |  |  |  |  |  |  | \| |  |  |  |  |  |  |  |  |  | \| |  |  |  |  |  |  |  |  |  | \| |  |  |  |  |  |  |  |  |  | \| |  |  |  |  |  |  |  |  |  | \| |  |  |  |  |  |  |  |  |  | \| |  |  |  |  |  |  |  |  |  | \| |  |  |  |  |  |  |  |  |  | \| |  |  |  |  |  |  |  |  |  | \| |  |  |  |  |  |  |  |  |  | \| |  |  |  |  |  |  |  |  |  | \| |  |  |  |  |  |  |  |  |  | \| |  |  |  |  |  |  |  |  |  | \| |  |  |  |  |  |  |  |  |  | \| |  |  |  |  |  |  |  |  |  | \| |  |  |  |  |  |  |  |  |  | \| |  |  |  |  |  |  |  |  |  | \| |  |  |  |  |  |  |  |  |  | \| |  |  |  |  |  |  |  |  |  | \| |  |  |  |  |  |  |  |  |  | \| |  |  |  |  |  |  |  |  |  | \| |  |  |  |  |  |  |  |  |  | \| |  |  |  |  |  |  |  |  |  | \| |  |  |  |  |  |  |  |  |  | \| |  |  |  |  |  |  |  |  |  | \| |  |  |  |  |  |  |  |  |  | \| |  |  |  |  |  |  |  |  |  | \| |  |  |  |  |  |  |  |  |  | \| |  |  |  |  |  |  |  |  |  | \| |  |  |  |  |  |  |  |  |  | \| |  |  |  |  |  |  |  |  |  | \| |  |  |  |  |  |  |  |  |  | \| |  |  |  |  |  |  |  |  |  | \| |  |  |  |  |  |  |  |  |  | \| |  |  |  |  |  |  |  |  |  | \| |  |  |  |  |  |  |  |  |  | \| |  |  |  |  |  |  |  |  |  | \| |  |  |  |  |  |  |  |  |  | \| |  |  |  |  |  |  |  |  |  | \| |  |  |  |  |  |  |  |  |  | \| |  |  |  |  |  |  |  |  |  | \| |  |  |  |  |  |  |  |  |  | \| |  |  |  |  |  |  |  |  |  | \| |  |  |  |  |  |  |  |  |  | \| |  |  |  |  |  |  |  |  |  | \| |  |  |  |  |  |  |  |  |  | \| |  |  |  |  |  |  |  |  |  | \| |  |  |  |  |  |  |  |  |
| Acinonyx\_jubatus | M | - | - | - | - | - | - | - | - | - | - | - | - | - | - | - | - | - | - | - | - | - | - | - | - | - | - | - | - | - | - | - | - | - | - | - | - | - | - | - | - | - | - | - | - | - | - | - | - | - | - | - | - | - | - | - | - | - | - | - | - | - | - | - | - | - | - | - | - | - | - | - | - | - | - | - | - | - | - | - | - | - | - | - | - | - | - | - | - | - | - | - | - | - | - | - | - | - | D | A | K | A | - | R | N | - | - | - | - | - | - | - | - | - | - | - | - | - | - | - | - | C | L | L | - | - | - | - | - | - | - | - | - | - | - | - | - | - | - | - | - | - | - | - | - | - | - | - | - | - | - | - | - | - | - | - | - | - | - | - | - | - | - | - | - | - | - | - | - | - | - | - | - | - | - | - | - | - | - | - | - | - | - | - | - | - | - | - | - | - | - | - | - | - | - | - | - | - | - | - | - | - | - | - | - | - | - | - | - | - | - | - | - | - | - | - | - | - | - | - | - | - | - | - | - | - | - | - | - | - | - | - | - | - | - | - | - | - | - | - | - | - | - | - | - | Q | H | R | E | A | L | E | K | D | I | K | T | S | Y | - | I | M | D | R | M | I | N | D | G | V | L | T | V | S | E | E | E | K | V | K | N | E | - | - | - | - | - | - | - | - | - | - | - | - | - | - | - | - | - | P | T | Q | Q | Q | R | A | A | L | L | I | K | M | I | L | K | K | - | - | - | - | D | N | Y | S | Y | I | S | - | - | - | - | - | F | Y | N | A | L | - | I | H | E | G | - | - | - | - | - | - | Y | K | D | L | A | G | L | L | H | G | G | M | P | V | I | S | S | S | N | G | G | K | - | - | - | - | - | - | - | - | - | - | - | - | - | - | - | - | - | - | - | - | - | - | D | S | V | G | G | I | T | - | - | - | - | - | - | - | - | - | - | - | S | Y | V | R | T | V | L | C | E | G | G | - | - | - | - | - | - | - | V | P | Q | R | P | V | V | F | V | T | R | K | K | L | V | N | S | I | - | - | - | - | - | - | - | - | - | - | Q | Q | N | L | F | - | - | - | - | - | - | - | - | - | - | - | - | - | - | - | - | - | - | - | - | - | - | - | - | - | - | - | - | - | - | - | - | K | L | N | G | E | P | - | - | - | - | - | - | - | - | - | - | - | - | - | - | - | - | - | - | - | - | - | - | - | - | - | - | - | - | - | - | - | - | - | - | - | - | - | - | G | W | V | I | I | Y | G | M | A | G | C | G | K | S | V | L | A | A | E | T | V | - | R | D | H | S | F | L | D | G | C | F | P | G | G | V | H | W | V | S | V | G | - | - | - | - | - | - | - | K | - | - | - | - | - | - | Q | D | K | S | - | - | - | - | - | - | - | - | - | - | - | - | G | L | L | M | K | L | Q | N | L | C | A | R | L | D | - | - | - | - | Q | - | - | - | - | - | - | - | - | - | - | - | - | - | - | - | D | E | S | F | S | Q | R | L | P | L | N | I | - | - | - | - | - | - | - | - | - | - | - | - | - | - | - | - | E | E | A | K | D | R | L | R | I | L | M | L | R | - | - | - | - | - | - | - | - | - | - | - | - | K | H | P | R | S | L | L | - | - | I | L | D | D | V | W | D | P | W | V | L | K | A | F | D | - | N | Q | C | Q | I | L | I | T | T | R | D | K | S | V | T | D | S | V | - | M | G | P | - | - | K | Y | V | V | S | V | E | S | - | - | - | G | L | G | K | E | K | G | L | E | I | L | S | L | F | V | N | V | - | K | K | A | D | L | P | E | - | - | - | - | - | - | - | - | - | - | - | - | - | - | Q | A | H | S | I | I | K | E | C | K | - | - | - | - | - | - | - | - | - | - | - | - | - | - | - | - | - | - | - | G | - | S | P | L | V | V | S | L | I | G | A | L | L | R | D | F | P | N | - | - | - | - | - | - | - | - | - | - | R | W | D | Y | Y | L | - | R | Q | L | Q | N | K | Q | F | K | R | I | - | - | - | - | - | - | - | - | - | - | - | - | - | - | - | - | - | - | - | - | - | - | R | K | S | S | F | Y | D | - | - | - | - | Y | E | A | L | D | E | A | M | S | I | S | V | E | - | M | L | R | - | - | - | E | D | I | K | D | Y | - | Y | T | D | L | S | I | L | Q | K | D | V | K | V | P | T | K | V | L | C | I | L | W | D | - | - | - | - | M | E | T | E | E | V | E | D | I | L | Q | E | F | - | - | - | - | - | - | - | - | - | - | - | V | N | K | S | L | L | F | C | D | R | N | G | K | S | - | - | - | - | - | - | - | - | - | F | C | Y | Y | L | H | D | L | Q | V | D | F | L | I | E | K | - | - | N | R | N | Q | - | L | - | Q | D | L | H | K | K | L | I | T | Q | F | Q | R | - | H | H | - | - | - | - | - | - | Q | P | H | T | L | S | P | D | - | Q | E | - | - | - | - | - | - | - | D | C | M | Y | W | Y | N | F | L | A | Y | H | M | A | S | A | N | - | - | - | - | M | H | K | E | L | C | A | L | M | F | S | L | D | W | I | K | A | K | T | E | L | V | G | P | A | - | - | - | - | - | - | - | - | H | L | I | H | E | F | V | E | Y | R | H | I | L | D | E | K | - | - | - | - | - | - | - | D | C | A | V | C | E | N | F | Q | - | - | - | E | F | L | S | L | N | G | H | L | L | G | R | Q | P | F | - | - | - | - | - | - | - | - | - | - | - | - | - | - | - | - | - | - | - | - | - | - | - | - | - | - | - | - | - | - | - | - | - | - | - | - | - | - | - | - | - | - | - | - | - | - | - | - | - | - | - | - | - | - | - | - | - | - | - | - | - | - | - | - | - | - | - | - | - | - | - | P | N | I | V | Q | L | G | L | C | E | P | E | T | S | E | V | Y | Q | Q | A | K | L | Q | A | K | Q | E | V | D | N | G | - | - | - | - | - | - | - | - | - | - | - | - | - | - | - | - | - | - | - | - | - | - | - | - | - | - | - | - | - | M | L | Y | L | - | - | - | - | - | - | - | - | - | - | E | W | I | N | K | - | - | - | - | - | - | - | K | N | N | K | N | L | S | R | - | - | - | - | - | - | - | - | - | - | L | V | V | R | P | H | T | D | - | - | - | - | - | - | - | - | - | - | - | - | - | - | - | A | V | Y | H | A | C | - | - | - | F | S | - | E | D | G | Q | R | I | A | - | - | - | - | - | - | - | - | - | S | C | G | V | D | K | T | L | Q | V | F | K | - | - | - | - | - | A | E | T | G | E | K | L | L | E | I | - | K | A | H | E | D | E | - | - | - | V | L | C | C | A | F | S | - | A | - | - | - | - | - | - | - | - | - | - | - | - | - | D | D | R | F | I | A | T | C | S | V | D | K | K | V | K | I | - | - | - | W | N | S | M | - | - | - | - | - | - | - | - | - | - | - | - | - | - | - | - | - | - | - | - | - | - | - | - | - | - | - | - | - | - | - | - | - | - | - | - | - | - | - | - | - | - | - | - | - | - | - | - | - | - | - | - | - | - | - | - | - | - | - | - | - | - | - | - | - | - | - | - | - | - | - | - | - | - | - | - | - | - | - | - | - | - | - | - | - | - | - | - | - | - | - | - | - | - | - | - | - | - | - | - | - | - | - | - | - | - | - | - | - | - | - | - | - | - | - | - | - | - | - | - | - | - | - | - | - | - | - | - | - | - | - | - | - | - | - | - | - | - | - | - | T | G | E | L | V | H | V | Y | - | D | E | H | S | - | - | - | - | - | E | Q | V | N | - | - | - | - | - | - | - | - | - | - | - | - | - | C | C | H | - | - | - | - | - | - | - | - | - | - | - | - | - | F | T | N | N | G | - | - | - | - | - | - | - | - | - | - | - | - | - | - | - | - | H | H | - | L | L | L | A | T | A | S | S | - | - | - | - | D | C | F | L | - | - | - | - | - | - | - | - | - | - | - | - | K | L | W | D | L | - | - | - | T | Q | K | E | C | R | N | T | M | F | - | - | - | - | - | - | - | - | - | - | - | - | - | - | - | - | - | G | H | T | N | S | - | - | - | - | - | - | - | - | - | - | - | - | - | - | - | - | V | N | H | C | R | F | S | P | D | D | K | - | - | - | - | - | - | - | - | - | - | - | - | - | - | - | - | - | - | - | - | - | - | - | - | - | - | - | - | L | L | A | S | C | S | A | D | G | T | L | K | L | - | - | - | - | - | - | - | - | - | - | - | - | - | - | - | - | - | - | - | W | D | - | V | K | S | A | N | E | R | E | S | - | - | - | - | - | - | I | N | - | - | - | - | - | - | - | - | - | - | - | - | - | - | - | - | - | - | - | - | - | - | V | K | Q | F | - | - | - | - | - | - | - | - | F | L | N | L | E | E | P | Q | E | - | - | - | - | - | - | - | - | - | - | - | - | - | - | - | - | - | - | - | - | - | - | - | - | - | - | - | D | M | E | - | V | I | V | K | C | C | - | - | - | - | - | - | - | - | - | - | - | - | - | - | - | - | S | W | S | A | D | G | A | R | I | M | V | A | A | K | N | K | I | - | - | - | - | - | - | - | - | - | - | - | F | L | F | D | I | H | A | - | - | - | S | G | - | - | - | - | - | - | - | - | - | - | - | - | - | - | - | - | - | - | - | - | - | - | - | - | - | - | - | - | - | - | - | - | - | - | - | - | - | - | - | - | - | - | - | - | - | - | - | - | - | - | - | - | - | - | - | - | - | - | - | - | - | - | - | - | - | - | - | - | - | - | - | - | - | - | - | - | - | - | - | - | - | - | - | - | - | - | - | - | - | - | - | - | - | - | - | - | - | - | - | - | - | - | - | - | - | - | - | - | - | - | - | - | - | - | - | - | - | - | - | - | - | - | - | - | - | - | L | L | A | - | - | - | - | - | - | - | - | - | - | - | - | - | - | - | - | - | - | - | - | - | - | - | - | - | - | - | - | - | - | - | - | - | - | - | - | - | - | - | - | - | - | - | - | - | - | - | - | - | - | - | - | - | - | - | - | - | - | - | - | - | - | - | - | - | - | - | - | - | - | - | - | - | - | - | - | - | - | - | - | - | - | - | - | - | - | - | - | - | - | - | - | - | - | - | - | - | - | - | - | - | - | - | - | - | - | - | - | - | - | - | - | - | - | - | - | - | - | - | - | - | - | - | - | - | - | - | - | - | - | - | - | - | - | - | - | - | - | - | - | - | - | - | - | - | - | - | - | - | - | - | - | - | - | - | - | - | - | - | - | - | - | - | - | - | - | - | - | - | - | - | - | - | - | - | - | - | - | - | - | - | - | - | E | I | - | - | - | H | T | G | H | H | S | T | I | Q | Y | C | - | - | - | - | - | - | - | - | - | - | - | - | - | - | - | - | - | - | - | - | - | - | - | - | - | D | F | S | P | Q | N | R | L | A | V | V | A | L | S | Q | - | - | - | - | - | - | C | C | V | E | - | - | - | - | - | - | - | - | - | - | - | - | - | - | - | - | - | L | W | N | M | D | S | C | - | - | - | - | - | - | - | - | - | - | - | - | - | - | - | - | - | - | - | - | - | - | - | - | - | - | - | - | - | L | K | V | A | D | C | R | G | H | L | S | - | - | - | - | - | - | - | - | - | - | - | - | - | - | - | - | - | - | - | - | - | - | - | - | - | - | - | - | - | - | - | - | - | - | - | - | - | - | - | - | - | - | - | - | W | V | H | C | V | M | F | S | - | - | - | - | - | - | - | - | - | - | - | P | D | G | S | - | - | - | - | - | - | - | - | - | - | S | F | L | T | S | S | D | D | Q | - | - | - | - | - | - | - | - | - | - | - | - | - | - | T | I | R | L | - | W | - | - | - | - | - | - | - | - | - | - | - | - | - | - | - | - | - | - | - | - | - | - | - | - | - | - | - | - | - | - | - | - | - | - | - | - | - | - | E | T | K | K | V | C | E | N | S | A | I | V | L | K | Q | D | - | - | - | - | - | - | - | - | - | - | - | - | - | - | - | - | I | D | V | V | F | - | - | - | - | - | - | - | - | - | - | - | - | - | - | - | - | - | - | - | - | - | - | - | - | - | - | - | - | - | - | - | - | - | - | - | - | - | - | - | - | - | - | - | - | - | - | - | - | - | - | - | - | - | - | Q | E | N | E | V | M | - | - | - | - | - | - | - | - | - | - | - | - | - | - | - | - | - | - | - | - | - | - | - | - | - | - | - | - | - | - | - | - | - | - | - | - | V | L | A | V | D | N | V | R | H | L | Q | L | - | I | N | G | K | T | - | - | - | - | - | - | - | - | - | - | G | Q | T | D | Y | L | T | E | A | - | - | - | - | - | - | - | - | - | - | - | - | - | - | - | - | - | - | - | - | - | - | Q | - | - | - | - | - | - | - | - | - | - | - | V | S | C | C | C | L | S | P | H | L | E | - | - | - | - | - | - | - | - | - | Y | I | A | F | G | G | - | - | - | - | - | - | - | - | - | - | - | - | - | - | - | - | - | - | - | - | - | - | - | - | - | - | - | - | - | - | - | - | - | - | - | - | E | - | - | - | - | - | - | - | - | - | D | G | A | - | - | - | - | - | - | - | - | - | - | I | - | - | - | - | - | - | - | - | - | - | - | - | - | - | - | - | - | - | - | - | - | - | - | - | - | - | - | - | - | - | - | - | - | - | - | - | - | - | - | E | I | L | E | L | L | N | N | R | I | F | Q | S | R | I | G | - | - | - | - | - | H | - | - | - | - | - | - | - | - | - | - | - | - | K | K | T | V | R | H | I | Q | F | T | - | - | - | - | - | - | - | - | - | - | - | - | - | - | - | - | - | - | - | - | - | - | - | - | - | - | - | - | - | - | - | - | - | - | - | - | - | D | D | E | K | T | L | I | S | S | S | - | D | D | S | - | - | - | - | - | - | - | - | S | I | Q | - | V | - | - | - | - | - | - | - | - | - | - | - | - | - | - | - | - | - | - | - | - | - | - | - | - | - | - | - | - | - | - | - | - | - | - | - | - | - | - | - | W | N | - | - | - | - | - | - | - | - | - | - | W | Q | S | E | E | Y | V | F | - | - | - | - | - | - | - | - | - | - | - | - | - | - | - | - | - | - | - | - | - | - | - | - | - | - | - | - | - | - | - | - | - | - | - | - | - | - | - | - | - | - | - | - | - | - | - | - | - | - | L | Q | A | H | Q | - | - | - | - | - | - | - | - | - | - | - | - | - | - | - | - | - | - | - | - | - | - | - | - | - | - | E | T | V | K | D | F | R | L | L | - | - | - | - | - | - | - | - | - | - | - | - | - | - | - | - | - | - | - | - | - | - | - | K | N | - | S | R | L | L | - | S | W | - | - | - | S | F | D | G | T | V | - | - | - | - | - | - | - | - | - | - | - | - | - | - | - | - | - | - | - | - | - | - | - | - | - | - | - | - | - | - | - | - | - | - | - | K | V | W | S | I | I | T | G | R | I | - | - | - | - | - | E | K | D | F | V | C | - | - | - | - | - | - | - | H | Q | - | - | - | - | - | - | - | - | - | - | - | - | - | - | - | - | - | - | - | D | T | V | L | - | - | - | - | - | - | - | - | - | - | - | - | - | - | - | S | C | D | - | - | - | - | - | - | - | - | - | - | - | - | - | - | - | I | S | P | D | A | T | - | K | F | S | S | T | - | S | A | D | K | T | - | - | - | - | - | - | A | K | I | W | S | F | E | L | L | - | - | - | - | - | - | - | - | - | - | - | - | - | - | - | - | - | - | - | - | - | - | - | - | S | P | L | H | E | - | - | - | - | - | - | - | - | L | - | - | - | - | R | G | H | K | G | C | V | - | - | - | - | - | - | - | - | - | R | C | S | - | A | F | S | - | - | - | - | - | - | - | - | - | V | D | S | T | - | L | L | A | T | - | - | - | - | - | - | - | - | G | D | D | N | G | E | I | R | I | W | N | V | S | N | G | E | L | L | - | - | - | - | - | - | - | - | - | - | - | - | - | - | - | - | - | - | - | - | - | - | - | - | - | - | - | - | - | - | - | - | - | - | - | - | - | - | - | - | - | - | - | - | - | - | - | - | - | - | - | - | - | - | - | - | H | L | C | A | P | I | S | V | E | - | - | - | - | - | - | - | - | - | - | - | - | - | - | - | - | - | - | - | - | E | G | A | A | T | H | G | G | W | - | - | - | - | - | - | - | - | V | T | D | L | C | F | S | P | D | S | K | - | - | - | - | - | - | - | - | - | - | - | - | - | - | - | - | - | - | - | - | - | - | - | - | - | - | - | - | - | - | - | - | - | - | - | - | - | - | - | - | - | - | - | - | - | M | L | V | S | - | - | A | G | G | Y | L | K | - | - | - | - | - | - | - | - | W | W | S | V | D | T | - | - | - | - | - | - | - | - | - | - | - | - | - | - | - | - | - | - | - | - | - | - | - | - | - | - | - | - | - | - | - | - | - | - | - | - | - | - | - | - | - | - | - | - | - | - | - | - | - | - | - | - | - | - | - | - | - | - | - | - | - | - | - | - | - | - | - | - | - | - | - | - | - | - | - | - | - | - | - | - | - | - | - | - | - | - | - | - | - | - | - | - | - | - | - | - | - | - | - | - | - | - | - | - | - | - | - | - | - | - | - | - | - | - | - | - | - | - | - | - | - | - | - | - | - | - | - | - | - | - | - | - | - | - | - | - | - | - | - | - | - | - | - | - | - | - | - | - | - | - | - | - | - | - | - | - | - | - | - | - | - | - | - | - | - | - | - | - | - | - | - | - | - | - | - | - | - | - | - | - | - | - | - | - | - | - | - | - | - | - | - | - | - | - | - | - | - | - | - | - | - | - | - | - | - | - | - | - | - | - | - | - | - | - | - | - | - | - | - | - | - | - | - | - | - | - | - | - | - | - | - | - | - | - | - | - | - | - | - | - | - | - | - | - | - | - | - | - | - | - | - | - | - | - | - | - | - | - | - | - | - | - | - | - | - | - | - | G | E | S | S | Q | T | F | Y | T | - | - | - | - | - | N | G | - | - | - | T | N | L | K | K | I | - | - | - | H | V | S | P | - | - | - | - | D | F | K | T | - | - | - | - | - | - | - | - | - | - | - | - | - | - | - | - | - | - | - | - | - | - | - | - | - | - | Y | V | T | V | D | - | N | L | G | I | L | - | - | Y | I | L | Q | M | - | - | - | - | - | - | - | - | - | - | - | - | - | - | - | - | - | - | - | - | - | - | - | - | - | - | - | - | - | - | - | - | - | - | L | E | - | - | - | - | - | - | - | - | - | - | - | - | - | - | - | - | - | - | - | - | - | - | - | - | - | - | - | - | - | - | - | - | - | - | - | - | - | - | - | - | - | - | - | - | - | - | - | - | - | - | - | - | M | - | - | - | - | - | - | - | - | - | - | - | - | - | - | - | - | - | - | - | - | - | - | - | - | - | - | - | - | - | - | - | - | - | - | - | - | - | - | - | - | - | - | - | - | - | - | - | - | - | - | - | - | - | - | - | - | - | - | - | - | - | - | - | - | - | - | - | - | - | - | - | - | - | - | - | - | - | - | - | - | - | - | - | - | - | - | - | - | - | - | - | - | - | - | - | - | - | - | - | - | - | - | - | - | - | - | - | - | - | - | - | - | - | - | - | - | - | - | - | - | - | - | - | - | - | - | - | - | - | - | - | - | - | - | - | - | - | - | - | - | - | - | - | - | - | - | - | - | - | - | - | - | - | - | - | - | - | - | - | - | - | - | - | - | - | - | - | - | - | - | - | - | - | - | - | - | - | - | - | - | - | - | - | - | - | - | - | - | - | - | - | - | - | - | - | - | - | - | - | - | - | - | - | - | - | - | - | - | - | - | - | - | - | - | - | - | - | - | - | - | - | - | - | - | - | - | - | - | - | - | - | - | - | - | - | - | - | - | - | - | - | - | - | - | - | - | - | - | - | - | - | - | - | - | - | - | - | - | - | - | - | - | - | - | - | - | - | - | - | - | - | - | - | - | - | - | - | - | - | - | - | - | - | - | - | - | - | - | - | - | - | - | - | - | - | - | - | - | - | - | - | - | - | - | - | - | - | G | R | Q | A | E | - | - | - | - | K | Q | N | P | - | Q | A | V | - | - | - | - | - | - | - | - | - | - | - | - | - | - | - | - | - | - | - | - | - | - | - | - | - | - | R | P | L | D | - | - | - | - | - | - | - | - | - | - | - | - | - | - | - | L | M | P | V | - | - | - | - | - | - | - | - | - | - | - | - | - | - | - | - | - | - | - | - | - | - | - | - | - | - | - | - | - | - | - | - | - | - | - | - | - | - | - | V | V | G | P | M | G | L | K | P | E | E | L | R | - | - | R | R | Q | Y | P | L | K | P | - | T | - | P | - | - | - | - | - | - | - | - | - | - | - | - | - | - | - | - | - | - | - | - | - | - | - | - | - | - | - | - | - | - | - | - | - | - | - | - | - | - | - | - | - | - | - | - | - | - | - | - | - | - | - | E | K | L | T | P | V | - | - | - | - | - | - | - | - | - | - | - | - | - | - | - | - | - | - | - | - | - | - | - | - | - | - | V | L | - | - | - | - | - | - | - | - | - | - | - | - | - | - | - | - | - | - | - | - | - | - | - | - | - | - | - | - | - | - | - | - | - | - | - | - | - | - | - | - | - | - | - | - | - | - | - | - | - | - | - | - | - | - | - | - | - | - | - | - | - | - | - | - | - | - | - | - | G | P | E | E | L | W | P | - | - | - | - | - | - | - | - | - | - | - | - | - | - | - | - | - | - | - | - | - | - | - | - | - | - | A | K | L | R | P | - | - | - | - | - | - | E | V | L | R | P | - | E | V | - | - | - | - | - | - | - | - | - | - | - | - | - | - | - | - | - | - | P | R | P | V | D | I | G | S | G | P | F | - | - | - | - | - | - | - | - | - | - | - | - | - | S | D | I | C | - | - | - | - | - | - | - | - | - | - | A | - | Q | E | I | S | K | R | N | A | D | - | - | - | - | - | L | A | Y | V | - | - | L | N | A | D | - | P | C | G | H | C | L | I | I | N | N | M | N | F | C | L | E | S | - | - | R | L | Q | T | R | T | G | S | N | I | D | C | E | K | L | Q | R | R | F | - | F | L | L | H | F | V | V | D | V | K | H | D | Q | T | A | K | Q | M | V | Q | V | L | V | E | L | A | Q | R | D | H | - | G | A | L | D | C | C | V | V | V | I | L | S | H | G | C | Q | A | S | H | R | Q | F | P | G | A | - | V | Y | G | T | D | G | C | P | V | S | I | E | K | I | V | N | I | F | N | G | S | G | C | P | S | L | G | G | K | P | K | L | F | F | I | Q | A | C | - | - | G | G | E | R | K | D | H | G | V | E | V | A | S | T | - | - | - | S | P | E | D | - | R | T | P | G | S | D | - | - | - | - | - | - | - | S | E | P | D | A | V | P | F | Q | E | G | P | G | T | F | - | - | - | - | - | - | - | - | D | Q | P | D | A | V | - | - | - | - | - | - | - | S | S | L | - | - | - | P | T | P | S | D | I | F | V | S | Y | S | T | F | P | - | - | G | - | - | - | - | - | - | - | - | F | V | S | W | R | D | T | K | S | G | S | W | Y | V | E | T | L | D | G | V | F | E | Q | W | A | R | - | S | - | - | - | - | - | - | - | - | - | - | - | - | - | - | - | - | - | - | - | - | - | - | - | - | - | - | - | - | - | - | - | E | D | L | Q | T | L | M | - | - | - | - | - | L | R | - | - | - | - | - | - | - | - | - | - | - | - | V | A | N | A | V | - | - | - | - | - | S | Q | K | - | - | - | - | - | - | - | G | I | Y | - | K | Q | I | - | - | P | G | - | - | - | - | - | - | - | - | - | - | - | - | - | - | - | C | F | N | F | - | - | - | - | - | - | - | - | - | L | R | K | K | L | Y | F | K | M | - | - | - | - | - | - | - | - | - | - | - | - | - | - | - | - | - | - | - | - | - | - | - | - | - | - | - | - | - | - | - | - | - | - | - | - | - | - | M | - | - | - | - | - | - | - | - | - | - | - | - | - | - | - | - | - | - | - | - | - | - | - | - | - | - | - | - | - | - | - | - | - | - | - | - | - | - | - | - | - | - | - | - | - | - | - | - | - | - | - | - | G | D | V | E | K | G | K | K | I | F | V | Q | K | C | A | Q | C | H | T | V | E | K | G | G | K | H | K | T | G | P | N | L | H | G | L | F | G | R | K | T | G | Q | A | P | G | F | S | Y | T | D | A | N | K | N | K | G | I | T | W | G | E | E | T | L | M | E | Y | L | E | N | P | K | K | Y | I | P | G | T | K | M | I | F | A | G | I | K | K | A | G | E | R | A | D | L | I | A | Y | L | - | - | - | - | - | - | - | - | - | - | - | - |
| Alligator\_mississippiensis | M | - | - | - | - | - | - | - | - | - | - | - | - | - | - | - | - | - | - | - | - | - | - | - | - | - | - | - | - | - | - | - | - | - | - | - | - | - | - | - | - | - | - | - | - | - | - | - | - | - | - | - | - | - | - | - | - | - | - | - | - | - | - | - | - | - | - | - | - | - | - | - | - | - | - | - | - | - | - | - | - | - | - | - | - | - | - | - | - | - | - | - | - | - | - | - | - | - | D | V | K | S | - | R | N | - | - | - | - | - | - | - | - | - | - | - | - | - | - | - | - | H | L | L | - | - | - | - | - | - | - | - | - | - | - | - | - | - | - | - | - | - | - | - | - | - | - | - | - | - | - | - | - | - | - | - | - | - | - | - | - | - | - | - | - | - | - | - | - | - | - | - | - | - | - | - | - | - | - | - | - | - | - | - | - | - | - | - | - | - | - | - | - | - | - | - | - | - | - | - | - | - | - | - | - | - | - | - | - | - | - | - | - | - | - | - | - | - | - | - | - | - | - | - | - | - | - | - | - | - | - | - | - | - | - | - | - | - | - | - | - | - | - | - | - | L | N | R | L | A | L | E | R | D | I | K | T | S | Y | - | V | M | D | H | M | I | T | D | Q | V | L | T | L | Q | E | E | E | K | V | K | A | Q | - | - | - | - | - | - | - | - | - | - | - | - | - | - | - | - | - | A | T | Q | R | D | R | A | A | M | L | I | N | I | I | L | K | K | - | - | - | - | D | N | N | S | Y | I | S | - | - | - | - | - | F | Y | N | A | L | - | L | H | E | G | - | - | - | - | - | - | Y | K | D | L | A | A | L | L | H | D | G | I | P | S | I | T | S | Y | D | R | - | K | - | - | - | - | - | - | - | - | - | - | - | - | - | - | - | - | - | - | - | - | - | - | D | S | T | D | G | V | A | - | - | - | - | - | - | - | - | - | - | - | F | C | V | R | T | I | L | C | E | G | G | - | - | - | - | - | - | - | V | P | Q | R | P | V | V | F | V | T | R | P | E | L | V | N | E | I | - | - | - | - | - | - | - | - | - | - | R | Q | K | L | Y | - | - | - | - | - | - | - | - | - | - | - | - | - | - | - | - | - | - | - | - | - | - | - | - | - | - | - | - | - | - | - | - | N | L | R | N | D | P | - | - | - | - | - | - | - | - | - | - | - | - | - | - | - | - | - | - | - | - | - | - | - | - | - | - | - | - | - | - | - | - | - | - | - | - | - | - | G | W | V | T | V | Y | G | M | A | G | S | G | K | S | V | L | T | A | E | A | L | - | R | D | Q | S | L | L | E | E | C | F | P | G | G | V | H | W | I | S | V | G | - | - | - | - | - | - | - | K | - | - | - | - | - | - | Q | D | N | A | - | - | - | - | - | - | - | - | - | - | - | - | G | L | L | M | K | L | H | N | L | C | S | R | L | D | - | - | - | - | L | - | - | - | - | - | - | - | - | - | - | - | - | - | - | - | D | F | T | Y | A | Q | R | P | P | L | N | I | - | - | - | - | - | - | - | - | - | - | - | - | - | - | - | - | G | E | A | K | D | R | L | R | L | L | M | L | H | - | - | - | - | - | - | - | - | - | - | - | - | K | Y | P | R | S | L | L | - | - | V | L | D | D | I | W | D | S | Q | T | L | K | A | F | D | - | N | H | C | K | V | L | I | T | S | R | D | R | S | V | T | D | H | V | - | T | G | R | - | - | K | Y | E | V | H | M | E | S | - | - | - | G | L | N | D | E | K | G | L | E | I | L | S | L | F | V | D | M | - | K | I | K | D | M | P | A | - | - | - | - | - | - | - | - | - | - | - | - | - | - | Q | A | D | K | I | V | R | E | C | K | - | - | - | - | - | - | - | - | - | - | - | - | - | - | - | - | - | - | - | G | - | S | P | L | V | L | S | L | I | G | A | L | L | R | D | F | P | N | - | - | - | - | - | - | - | - | - | - | R | W | E | Y | Y | L | - | R | Q | L | Q | N | K | Q | F | R | R | I | - | - | - | - | - | - | - | - | - | - | - | - | - | - | - | - | - | - | - | - | - | - | K | K | S | S | S | Y | D | - | - | - | - | Y | E | A | L | D | E | A | M | S | I | S | V | E | - | M | L | N | - | - | - | E | N | F | K | D | F | - | Y | K | D | L | S | I | L | P | K | D | V | K | V | P | T | K | V | F | C | I | L | W | D | - | - | - | - | M | E | P | E | E | V | E | D | M | Q | Q | E | F | - | - | - | - | - | - | - | - | - | - | - | V | N | K | S | L | L | F | C | D | R | N | G | K | S | - | - | - | - | - | - | - | - | - | F | H | Y | Y | L | H | D | L | Q | L | D | F | L | I | E | K | - | - | N | R | E | Q | - | L | - | Q | E | L | H | E | K | I | V | K | Q | Y | K | K | - | H | Y | - | - | - | - | - | - | R | L | C | I | P | T | P | G | - | E | E | - | - | - | - | - | - | - | D | C | M | Y | W | Y | N | F | L | A | F | H | M | A | S | A | K | - | - | - | - | M | Q | E | E | L | C | D | L | M | F | S | L | D | W | I | K | A | K | T | E | L | V | G | P | A | - | - | - | - | - | - | - | - | H | L | I | H | E | Y | V | E | Y | S | N | I | L | D | Q | K | - | - | - | - | - | - | - | N | S | L | A | R | E | N | F | Q | - | - | - | K | F | L | S | L | N | G | H | L | L | G | R | Q | P | F | - | - | - | - | - | - | - | - | - | - | - | - | - | - | - | - | - | - | - | - | - | - | - | - | - | - | - | - | - | - | - | - | - | - | - | - | - | - | - | - | - | - | - | - | - | - | - | - | - | - | - | - | - | - | - | - | - | - | - | - | - | - | - | - | - | - | - | - | - | - | - | P | D | I | V | Q | L | G | L | C | Q | P | E | T | S | E | V | Y | Q | Q | A | K | L | K | A | H | Q | E | V | D | K | G | - | - | - | - | - | - | - | - | - | - | - | - | - | - | - | - | - | - | - | - | - | - | - | - | - | - | - | - | - | A | F | Y | V | - | - | - | - | - | - | - | - | - | - | E | W | I | N | K | - | - | - | - | - | - | - | - | G | M | K | N | L | S | R | - | - | - | - | - | - | - | - | - | - | L | V | V | R | P | H | T | D | - | - | - | - | - | - | - | - | - | - | - | - | - | - | - | A | V | Y | H | A | C | - | - | - | F | S | - | Q | D | G | Q | K | I | A | - | - | - | - | - | - | - | - | - | S | C | G | A | D | R | T | L | Q | V | F | K | - | - | - | - | - | A | E | T | G | E | K | L | Q | N | V | - | E | A | H | D | D | E | - | - | - | V | L | Y | C | A | F | S | - | P | - | - | - | - | - | - | - | - | - | - | - | - | - | D | G | R | F | I | A | T | C | S | A | D | N | K | A | K | I | - | - | - | W | C | L | R | - | - | - | - | - | - | - | - | - | - | - | - | - | - | - | - | - | - | - | - | - | - | - | - | - | - | - | - | - | - | - | - | - | - | - | - | - | - | - | - | - | - | - | - | - | - | - | - | - | - | - | - | - | - | - | - | - | - | - | - | - | - | - | - | - | - | - | - | - | - | - | - | - | - | - | - | - | - | - | - | - | - | - | - | - | - | - | - | - | - | - | - | - | - | - | - | - | - | - | - | - | - | - | - | - | - | - | - | - | - | - | - | - | - | - | - | - | - | - | - | - | - | - | - | - | - | - | - | - | - | - | - | - | - | - | - | - | - | - | - | T | G | K | R | I | F | D | C | - | V | E | H | T | - | - | - | - | - | E | Q | V | N | - | - | - | - | - | - | - | - | - | - | - | - | - | C | C | Q | - | - | - | - | - | - | - | - | - | - | - | - | - | F | N | S | G | N | - | - | - | - | - | - | - | - | - | - | - | - | - | - | - | - | N | E | - | L | L | L | T | T | C | S | N | - | - | - | - | D | S | L | L | - | - | - | - | - | - | - | - | - | - | - | - | K | L | W | D | I | - | - | - | R | N | K | R | C | Q | N | T | M | F | - | - | - | - | - | - | - | - | - | - | - | - | - | - | - | - | - | G | H | T | S | S | - | - | - | - | - | - | - | - | - | - | - | - | - | - | - | - | V | N | H | C | R | F | S | P | D | D | K | - | - | - | - | - | - | - | - | - | - | - | - | - | - | - | - | - | - | - | - | - | - | - | - | - | - | - | - | Y | V | A | S | C | S | T | D | G | T | V | K | L | - | - | - | - | - | - | - | - | - | - | - | - | - | - | - | - | - | - | - | W | E | - | T | G | S | G | N | E | L | K | C | - | - | - | - | - | - | I | K | - | - | - | - | - | - | - | - | - | - | - | - | - | - | - | - | - | - | - | - | - | - | I | K | D | F | - | - | - | - | - | - | - | - | F | R | N | A | D | E | R | - | - | - | - | - | - | - | - | - | - | - | - | - | - | - | - | - | - | - | - | - | - | - | - | - | - | - | - | - | E | V | E | - | A | L | P | K | C | C | - | - | - | - | - | - | - | - | - | - | - | - | - | - | - | - | S | W | S | T | D | G | A | T | V | M | V | A | A | K | N | K | L | - | - | - | - | - | - | - | - | - | - | - | L | L | L | D | V | E | T | - | - | - | S | D | - | - | - | - | - | - | - | - | - | - | - | - | - | - | - | - | - | - | - | - | - | - | - | - | - | - | - | - | - | - | - | - | - | - | - | - | - | - | - | - | - | - | - | - | - | - | - | - | - | - | - | - | - | - | - | - | - | - | - | - | - | - | - | - | - | - | - | - | - | - | - | - | - | - | - | - | - | - | - | - | - | - | - | - | - | - | - | - | - | - | - | - | - | - | - | - | - | - | - | - | - | - | - | - | - | - | - | - | - | - | - | - | - | - | - | - | - | - | - | - | - | - | - | - | - | - | L | L | E | - | - | - | - | - | - | - | - | - | - | - | - | - | - | - | - | - | - | - | - | - | - | - | - | - | - | - | - | - | - | - | - | - | - | - | - | - | - | - | - | - | - | - | - | - | - | - | - | - | - | - | - | - | - | - | - | - | - | - | - | - | - | - | - | - | - | - | - | - | - | - | - | - | - | - | - | - | - | - | - | - | - | - | - | - | - | - | - | - | - | - | - | - | - | - | - | - | - | - | - | - | - | - | - | - | - | - | - | - | - | - | - | - | - | - | - | - | - | - | - | - | - | - | - | - | - | - | - | - | - | - | - | - | - | - | - | - | - | - | - | - | - | - | - | - | - | - | - | - | - | - | - | - | - | - | - | - | - | - | - | - | - | - | - | - | - | - | - | - | - | - | - | - | - | - | - | - | - | - | - | - | - | - | E | V | - | - | - | L | - | S | Y | Q | N | T | I | Q | Y | Y | - | - | - | - | - | - | - | - | - | - | - | - | - | - | - | - | - | - | - | - | - | - | - | - | - | D | F | C | P | S | N | Q | L | V | A | V | A | F | S | H | - | - | - | - | - | - | Y | S | V | E | - | - | - | - | - | - | - | - | - | - | - | - | - | - | - | - | - | L | W | N | T | K | T | K | - | - | - | - | - | - | - | - | - | - | - | - | - | - | - | - | - | - | - | - | - | - | - | - | - | - | - | - | - | T | K | V | A | D | C | R | E | H | M | S | - | - | - | - | - | - | - | - | - | - | - | - | - | - | - | - | - | - | - | - | - | - | - | - | - | - | - | - | - | - | - | - | - | - | - | - | - | - | - | - | - | - | - | - | W | V | H | C | V | T | F | S | - | - | - | - | - | - | - | - | - | - | - | P | D | G | S | - | - | - | - | - | - | - | - | - | - | L | F | L | T | S | S | D | D | Q | - | - | - | - | - | - | - | - | - | - | - | - | - | - | T | I | R | I | - | W | - | - | - | - | - | - | - | - | - | - | - | - | - | - | - | - | - | - | - | - | - | - | - | - | - | - | - | - | - | - | - | - | - | - | - | - | - | - | E | T | S | K | V | C | K | P | P | E | I | A | L | K | N | E | - | - | - | - | - | - | - | - | - | - | - | - | - | - | - | - | L | D | V | A | F | - | - | - | - | - | - | - | - | - | - | - | - | - | - | - | - | - | - | - | - | - | - | - | - | - | - | - | - | - | - | - | - | - | - | - | - | - | - | - | - | - | - | - | - | - | - | - | - | - | - | - | - | - | - | K | G | D | E | M | M | - | - | - | - | - | - | - | - | - | - | - | - | - | - | - | - | - | - | - | - | - | - | - | - | - | - | - | - | - | - | - | - | - | - | - | - | I | L | A | I | L | N | H | T | H | L | Q | L | - | I | N | G | N | T | - | - | - | - | - | - | - | - | - | - | G | H | T | V | S | L | T | D | V | - | - | - | - | - | - | - | - | - | - | - | - | - | - | - | - | - | - | - | - | - | - | E | D | S | - | - | - | - | - | - | - | - | - | I | S | C | C | C | L | S | E | D | L | Q | - | - | - | - | - | - | - | - | - | F | A | A | F | G | Q | - | - | - | - | - | - | - | - | - | - | - | - | - | - | - | - | - | - | - | - | - | - | - | - | - | - | - | - | - | - | - | - | - | - | - | - | K | - | - | - | - | - | - | - | - | - | N | G | T | - | - | - | - | - | - | - | - | - | - | V | - | - | - | - | - | - | - | - | - | - | - | - | - | - | - | - | - | - | - | - | - | - | - | - | - | - | - | - | - | - | - | - | - | - | - | - | - | - | - | K | V | L | E | V | S | G | G | R | V | L | K | S | R | H | G | - | - | - | - | - | H | - | - | - | - | - | - | - | - | - | - | - | - | K | T | S | V | Q | H | C | L | F | T | - | - | - | - | - | - | - | - | - | - | - | - | - | - | - | - | - | - | - | - | - | - | - | - | - | - | - | - | - | - | - | - | - | - | - | - | - | S | D | G | Q | T | L | I | S | S | D | - | D | D | P | - | - | - | - | - | - | - | - | I | I | Q | - | V | - | - | - | - | - | - | - | - | - | - | - | - | - | - | - | - | - | - | - | - | - | - | - | - | - | - | - | - | - | - | - | - | - | - | - | - | - | - | - | W | N | - | - | - | - | - | - | - | - | - | - | W | E | L | E | E | C | L | F | - | - | - | - | - | - | - | - | - | - | - | - | - | - | - | - | - | - | - | - | - | - | - | - | - | - | - | - | - | - | - | - | - | - | - | - | - | - | - | - | - | - | - | - | - | - | - | - | - | - | L | E | G | H | N | - | - | - | - | - | - | - | - | - | - | - | - | - | - | - | - | - | - | - | - | - | - | - | - | - | - | E | T | I | K | N | F | R | L | L | - | - | - | - | - | - | - | - | - | - | - | - | - | - | - | - | - | - | - | - | - | - | - | E | N | - | S | R | L | L | - | S | W | - | - | - | S | F | D | G | T | V | - | - | - | - | - | - | - | - | - | - | - | - | - | - | - | - | - | - | - | - | - | - | - | - | - | - | - | - | - | - | - | - | - | - | - | K | V | W | N | I | V | H | G | K | I | - | - | - | - | - | E | V | N | F | A | C | - | - | - | - | - | - | - | H | K | - | - | - | - | - | - | - | - | - | - | - | - | - | - | - | - | - | - | - | D | T | V | L | - | - | - | - | - | - | - | - | - | - | - | - | - | - | - | S | C | A | - | - | - | - | - | - | - | - | - | - | - | - | - | - | - | V | S | P | D | A | T | - | K | F | S | S | T | - | S | V | D | K | T | - | - | - | - | - | - | A | K | I | W | S | F | K | S | S | - | - | - | - | - | - | - | - | - | - | - | - | - | - | - | - | - | - | - | - | - | - | - | - | S | I | L | H | E | - | - | - | - | - | - | - | - | L | - | - | - | - | K | G | H | E | A | C | V | - | - | - | - | - | - | - | - | - | R | C | C | - | T | F | S | - | - | - | - | - | - | - | - | - | F | D | N | K | - | L | L | A | T | - | - | - | - | - | - | - | - | G | D | D | K | R | E | I | R | I | W | D | V | S | S | G | E | L | L | - | - | - | - | - | - | - | - | - | - | - | - | - | - | - | - | - | - | - | - | - | - | - | - | - | - | - | - | - | - | - | - | - | - | - | - | - | - | - | - | - | - | - | - | - | - | - | - | - | - | - | - | - | - | - | - | H | L | C | S | P | V | I | I | D | - | - | - | - | - | - | - | - | - | - | - | - | - | - | - | - | - | - | - | - | E | G | G | L | T | H | D | Y | W | - | - | - | - | - | - | - | - | V | T | D | L | C | F | S | P | D | S | K | - | - | - | - | - | - | - | - | - | - | - | - | - | - | - | - | - | - | - | - | - | - | - | - | - | - | - | - | - | - | - | - | - | - | - | - | - | - | - | - | - | - | - | - | - | M | L | V | S | - | - | S | G | G | Y | L | K | - | - | - | - | - | - | - | - | W | W | N | A | V | T | - | - | - | - | - | - | - | - | - | - | - | - | - | - | - | - | - | - | - | - | - | - | - | - | - | - | - | - | - | - | - | - | - | - | - | - | - | - | - | - | - | - | - | - | - | - | - | - | - | - | - | - | - | - | - | - | - | - | - | - | - | - | - | - | - | - | - | - | - | - | - | - | - | - | - | - | - | - | - | - | - | - | - | - | - | - | - | - | - | - | - | - | - | - | - | - | - | - | - | - | - | - | - | - | - | - | - | - | - | - | - | - | - | - | - | - | - | - | - | - | - | - | - | - | - | - | - | - | - | - | - | - | - | - | - | - | - | - | - | - | - | - | - | - | - | - | - | - | - | - | - | - | - | - | - | - | - | - | - | - | - | - | - | - | - | - | - | - | - | - | - | - | - | - | - | - | - | - | - | - | - | - | - | - | - | - | - | - | - | - | - | - | - | - | - | - | - | - | - | - | - | - | - | - | - | - | - | - | - | - | - | - | - | - | - | - | - | - | - | - | - | - | - | - | - | - | - | - | - | - | - | - | - | - | - | - | - | - | - | - | - | - | - | - | - | - | - | - | - | - | - | - | - | - | - | - | - | - | - | - | - | - | - | - | - | - | - | G | E | A | L | Q | T | F | Y | T | - | - | - | - | - | N | G | - | - | - | T | D | L | Q | S | I | - | - | - | Y | V | S | S | - | - | - | - | N | F | K | I | - | - | - | - | - | - | - | - | - | - | - | - | - | - | - | - | - | - | - | - | - | - | - | - | - | - | Y | V | T | V | D | - | N | L | G | I | L | - | - | Y | V | L | Q | M | - | - | - | - | - | - | - | - | - | - | - | - | - | - | - | - | - | - | - | - | - | - | - | - | - | - | - | - | - | - | - | - | - | - | M | E | - | - | - | - | - | - | - | - | - | - | - | - | - | - | - | - | - | - | - | - | - | - | - | - | - | - | - | - | - | - | - | - | - | - | - | - | - | - | - | - | - | - | - | - | - | - | - | - | - | - | - | - | M | - | - | - | - | - | - | - | - | - | - | - | - | - | - | - | - | - | - | - | - | - | - | - | - | - | - | - | - | - | - | - | - | - | - | - | - | - | - | - | - | - | - | - | - | - | - | - | - | - | - | - | - | - | - | E | A | E | Q | R | Q | - | - | - | - | - | - | - | - | - | - | - | - | - | - | - | - | - | - | - | - | R | L | Q | R | - | - | - | - | - | - | - | - | - | - | - | - | - | - | - | - | - | - | - | - | - | - | - | - | - | - | - | - | - | - | G | R | - | - | - | - | - | - | - | - | V | R | L | - | V | R | G | L | Q | L | E | A | L | W | - | - | - | - | - | - | - | - | - | - | - | - | - | - | - | - | - | - | - | - | - | - | - | - | - | - | - | - | - | - | - | - | - | - | - | - | - | - | - | - | - | - | - | - | - | G | P | L | V | R | D | G | V | - | - | - | - | - | - | - | - | - | - | - | - | - | - | - | - | - | - | - | - | - | - | - | - | F | S | R | D | M | L | D | D | - | - | - | - | - | - | - | - | - | - | - | - | - | - | - | - | - | - | - | - | - | - | - | - | - | I | Q | R | A | - | - | - | - | - | - | - | - | - | - | G | S | P | R | D | Q | - | A | R | Q | L | L | I | D | L | E | T | R | G | - | K | Q | A | F | P | T | F | V | S | I | L | R | E | T | G | Q | K | D | L | A | D | F | L | S | D | G | C | A | F | P | - | - | - | - | R | P | S | S | - | - | - | - | - | - | - | - | - | - | - | - | - | - | - | - | - | - | - | - | - | - | - | - | - | - | - | - | - | - | - | - | I | D | - | - | - | - | - | - | - | - | - | - | - | - | - | - | - | I | K | P | I | - | - | - | - | - | - | - | - | - | - | - | - | - | - | - | - | - | - | - | - | - | - | - | - | - | - | - | - | - | - | - | - | - | - | - | - | - | - | - | E | I | G | P | R | G | G | R | G | R | A | G | G | - | - | - | - | - | - | - | - | - | - | - | R | - | - | - | - | - | - | - | - | - | - | - | - | - | - | - | - | - | - | - | - | - | - | - | - | - | - | - | - | - | - | - | - | - | - | - | - | - | - | - | - | - | - | - | - | - | - | - | - | - | - | - | - | - | - | - | - | - | - | - | - | - | - | - | - | - | - | - | - | - | - | - | - | - | - | - | - | - | - | - | - | - | - | - | - | - | - | - | - | - | - | - | - | - | - | - | - | - | - | - | - | - | - | - | - | - | - | - | - | - | - | - | - | - | - | - | - | - | - | - | - | - | - | - | - | - | - | - | - | - | - | - | - | - | - | - | - | - | - | - | - | - | - | - | - | - | - | - | - | - | - | - | - | - | - | - | - | D | S | E | Y | L | P | - | - | - | - | - | - | - | - | - | - | - | - | - | - | - | - | - | - | - | - | - | - | - | - | - | - | L | P | V | Q | A | - | - | - | - | - | - | R | T | E | R | D | - | Q | E | L | D | - | - | - | - | - | - | - | - | - | - | - | - | - | - | - | H | P | G | T | V | A | H | S | - | - | - | - | - | - | - | - | - | - | - | - | - | - | - | L | C | V | A | G | A | - | - | - | - | - | - | - | - | - | - | A | - | A | D | R | F | R | Q | N | S | D | - | - | - | - | - | M | V | Y | V | - | - | L | K | S | R | - | P | C | G | Y | C | L | I | I | N | N | V | H | F | S | P | E | S | - | - | G | L | S | L | R | Q | G | S | D | L | D | Y | E | K | L | K | R | R | F | - | S | L | F | H | F | E | V | L | T | R | Q | D | L | K | A | K | E | M | Y | S | E | L | W | A | L | A | K | K | D | H | - | S | A | M | D | C | C | L | V | V | I | L | S | H | G | R | Q | R | S | H | I | Q | F | P | G | E | - | V | C | G | T | D | G | N | A | I | P | V | E | K | I | V | N | C | F | N | G | L | N | C | W | S | L | R | G | K | P | K | L | F | I | I | Q | A | C | - | - | G | G | E | E | K | D | Y | G | F | E | L | D | S | D | - | - | - | S | S | G | N | - | Q | G | Y | G | A | S | - | - | - | - | - | - | - | L | E | S | D | A | S | P | L | K | P | S | S | G | N | L | - | - | - | - | - | - | - | - | D | E | P | D | A | V | - | - | - | - | - | - | - | A | S | V | - | - | - | P | K | A | S | D | I | L | V | S | Y | S | T | F | P | - | - | G | - | - | - | - | - | - | - | - | F | V | S | W | R | D | R | K | S | G | S | W | F | V | E | T | L | D | R | I | L | E | Q | Y | A | D | - | S | - | - | - | - | - | - | - | - | - | - | - | - | - | - | - | - | - | - | - | - | - | - | - | - | - | - | - | - | - | - | - | E | D | L | Q | K | L | L | - | - | - | - | - | L | R | - | - | - | - | - | - | - | - | - | - | - | - | V | A | N | E | V | - | - | - | - | - | S | T | K | - | - | - | - | - | - | - | E | K | Y | - | K | Q | M | - | - | P | G | - | - | - | - | - | - | - | - | - | - | - | - | - | - | - | C | Y | H | F | - | - | - | - | - | - | - | - | - | L | R | K | W | F | F | F | M | T | K | - | - | - | - | - | - | - | - | - | - | - | - | - | - | - | - | - | - | - | - | - | - | - | - | - | - | - | - | - | - | - | - | - | - | - | - | - | M | - | - | - | - | - | - | - | - | - | - | - | - | - | - | - | - | - | - | - | - | - | - | - | - | - | - | - | - | - | - | - | - | - | - | - | - | - | - | - | - | - | - | - | - | - | - | - | - | - | - | - | - | G | D | V | E | K | G | K | K | I | F | V | Q | K | C | A | Q | C | H | T | V | E | K | G | G | K | H | K | T | G | P | N | L | H | G | L | I | G | R | K | T | G | Q | A | P | G | F | S | Y | T | E | A | N | K | N | K | G | I | T | W | G | E | E | T | L | M | E | Y | L | E | N | P | K | K | Y | I | P | G | T | K | M | I | F | A | G | I | K | K | K | P | E | R | A | D | L | I | A | Y | L | K | E | A | T | A | N | - | - | - | - | - | - |
| Alligator\_sinensis | M | - | - | - | - | - | - | - | - | - | - | - | - | - | - | - | - | - | - | - | - | - | - | - | - | - | - | - | - | - | - | - | - | - | - | - | - | - | - | - | - | - | - | - | - | - | - | - | - | - | - | - | - | - | - | - | - | - | - | - | - | - | - | - | - | - | - | - | - | - | - | - | - | - | - | - | - | - | - | - | - | - | - | - | - | - | - | - | - | - | - | - | - | - | - | - | - | - | D | V | K | S | - | R | N | - | - | - | - | - | - | - | - | - | - | - | - | - | - | - | - | H | L | L | - | - | - | - | - | - | - | - | - | - | - | - | - | - | - | - | - | - | - | - | - | - | - | - | - | - | - | - | - | - | - | - | - | - | - | - | - | - | - | - | - | - | - | - | - | - | - | - | - | - | - | - | - | - | - | - | - | - | - | - | - | - | - | - | - | - | - | - | - | - | - | - | - | - | - | - | - | - | - | - | - | - | - | - | - | - | - | - | - | - | - | - | - | - | - | - | - | - | - | - | - | - | - | - | - | - | - | - | - | - | - | - | - | - | - | - | - | - | - | - | - | L | N | R | L | A | L | E | R | D | I | K | T | S | Y | - | V | M | D | H | M | I | T | D | Q | V | L | T | L | Q | E | E | E | K | V | K | A | Q | - | - | - | - | - | - | - | - | - | - | - | - | - | - | - | - | - | A | T | Q | R | D | R | A | A | M | L | I | N | I | I | L | K | K | - | - | - | - | D | N | N | S | Y | I | S | - | - | - | - | - | F | Y | N | A | L | - | L | H | E | G | - | - | - | - | - | - | Y | K | D | L | A | A | L | L | H | D | G | I | P | S | I | T | S | Y | D | R | - | K | - | - | - | - | - | - | - | - | - | - | - | - | - | - | - | - | - | - | - | - | - | - | D | S | T | D | G | V | A | - | - | - | - | - | - | - | - | - | - | - | F | C | V | R | T | I | L | C | E | G | G | - | - | - | - | - | - | - | V | P | Q | R | P | V | V | F | V | T | R | P | E | L | V | N | E | I | - | - | - | - | - | - | - | - | - | - | R | Q | K | L | Y | - | - | - | - | - | - | - | - | - | - | - | - | - | - | - | - | - | - | - | - | - | - | - | - | - | - | - | - | - | - | - | - | N | L | R | N | D | P | - | - | - | - | - | - | - | - | - | - | - | - | - | - | - | - | - | - | - | - | - | - | - | - | - | - | - | - | - | - | - | - | - | - | - | - | - | - | G | W | V | T | V | Y | G | M | A | G | S | G | K | S | V | L | T | A | E | A | L | - | R | D | Q | S | L | L | E | E | C | F | P | G | G | V | H | W | I | S | V | G | - | - | - | - | - | - | - | K | - | - | - | - | - | - | Q | D | N | A | - | - | - | - | - | - | - | - | - | - | - | - | G | L | L | M | K | L | H | N | L | C | S | R | L | D | - | - | - | - | L | - | - | - | - | - | - | - | - | - | - | - | - | - | - | - | D | F | T | Y | A | Q | R | P | P | L | N | I | - | - | - | - | - | - | - | - | - | - | - | - | - | - | - | - | G | E | A | K | D | R | L | R | L | L | M | L | H | - | - | - | - | - | - | - | - | - | - | - | - | K | Y | P | R | S | L | L | - | - | V | L | D | D | I | W | D | S | Q | T | L | K | A | F | D | - | N | H | C | K | V | L | I | T | S | R | D | R | S | V | T | D | H | V | - | T | G | R | - | - | K | Y | E | V | H | M | E | S | - | - | - | G | L | N | D | E | K | G | L | E | I | L | S | L | F | V | D | M | - | K | I | K | D | M | P | A | - | - | - | - | - | - | - | - | - | - | - | - | - | - | Q | A | D | K | I | V | R | E | C | K | - | - | - | - | - | - | - | - | - | - | - | - | - | - | - | - | - | - | - | G | - | S | P | L | V | L | S | L | I | G | A | L | L | R | D | F | P | N | - | - | - | - | - | - | - | - | - | - | R | W | E | Y | Y | L | - | R | Q | L | Q | N | K | Q | F | R | R | I | - | - | - | - | - | - | - | - | - | - | - | - | - | - | - | - | - | - | - | - | - | - | K | K | S | S | S | Y | D | - | - | - | - | Y | E | A | L | D | E | A | M | S | I | S | V | E | - | M | L | N | - | - | - | E | N | F | K | D | F | - | Y | K | D | L | S | I | L | P | K | D | V | K | V | P | T | K | V | F | C | I | L | W | D | - | - | - | - | M | E | P | E | E | V | E | D | M | Q | Q | E | F | - | - | - | - | - | - | - | - | - | - | - | V | N | K | S | L | L | F | C | D | R | N | G | K | S | - | - | - | - | - | - | - | - | - | F | H | Y | Y | L | H | D | L | Q | L | D | F | L | I | E | K | - | - | N | R | E | Q | - | L | - | Q | E | L | H | E | K | I | V | K | Q | Y | K | K | - | H | Y | - | - | - | - | - | - | R | L | C | I | P | T | P | G | - | E | E | - | - | - | - | - | - | - | D | C | M | Y | W | Y | N | F | L | V | F | H | M | A | S | A | K | - | - | - | - | M | Q | E | E | L | R | D | L | M | F | S | L | D | W | I | K | A | K | T | E | L | V | G | P | A | - | - | - | - | - | - | - | - | H | L | I | H | E | Y | V | E | Y | S | N | I | L | D | Q | K | - | - | - | - | - | - | - | N | S | L | A | R | E | N | F | Q | - | - | - | K | F | L | S | L | N | G | H | L | L | G | R | Q | P | F | - | - | - | - | - | - | - | - | - | - | - | - | - | - | - | - | - | - | - | - | - | - | - | - | - | - | - | - | - | - | - | - | - | - | - | - | - | - | - | - | - | - | - | - | - | - | - | - | - | - | - | - | - | - | - | - | - | - | - | - | - | - | - | - | - | - | - | - | - | - | - | P | D | I | V | Q | L | G | L | C | Q | P | E | T | S | E | V | Y | Q | Q | A | K | L | K | A | H | Q | E | V | D | K | G | - | - | - | - | - | - | - | - | - | - | - | - | - | - | - | - | - | - | - | - | - | - | - | - | - | - | - | - | - | A | F | Y | V | - | - | - | - | - | - | - | - | - | - | E | W | I | N | K | - | - | - | - | - | - | - | - | G | M | K | N | L | S | R | - | - | - | - | - | - | - | - | - | - | L | V | V | R | P | H | T | D | - | - | - | - | - | - | - | - | - | - | - | - | - | - | - | A | V | Y | H | A | C | - | - | - | F | S | - | Q | D | G | Q | K | I | A | - | - | - | - | - | - | - | - | - | S | C | G | A | D | R | T | L | Q | V | F | K | - | - | - | - | - | A | E | T | G | E | K | L | Q | N | I | - | E | A | H | D | D | E | - | - | - | V | L | Y | C | A | F | S | - | P | - | - | - | - | - | - | - | - | - | - | - | - | - | D | G | R | F | I | A | T | C | S | A | D | N | K | A | K | I | - | - | - | W | C | L | R | - | - | - | - | - | - | - | - | - | - | - | - | - | - | - | - | - | - | - | - | - | - | - | - | - | - | - | - | - | - | - | - | - | - | - | - | - | - | - | - | - | - | - | - | - | - | - | - | - | - | - | - | - | - | - | - | - | - | - | - | - | - | - | - | - | - | - | - | - | - | - | - | - | - | - | - | - | - | - | - | - | - | - | - | - | - | - | - | - | - | - | - | - | - | - | - | - | - | - | - | - | - | - | - | - | - | - | - | - | - | - | - | - | - | - | - | - | - | - | - | - | - | - | - | - | - | - | - | - | - | - | - | - | - | - | - | - | - | - | - | T | G | K | R | I | F | D | C | - | G | E | H | T | - | - | - | - | - | E | Q | V | N | - | - | - | - | - | - | - | - | - | - | - | - | - | C | C | Q | - | - | - | - | - | - | - | - | - | - | - | - | - | F | N | S | G | N | - | - | - | - | - | - | - | - | - | - | - | - | - | - | - | - | N | E | - | L | L | L | T | T | C | S | N | - | - | - | - | D | S | L | L | - | - | - | - | - | - | - | - | - | - | - | - | K | L | W | D | V | - | - | - | R | N | K | R | C | Q | N | T | M | F | - | - | - | - | - | - | - | - | - | - | - | - | - | - | - | - | - | G | H | T | S | S | - | - | - | - | - | - | - | - | - | - | - | - | - | - | - | - | V | N | H | C | R | F | S | P | D | D | K | - | - | - | - | - | - | - | - | - | - | - | - | - | - | - | - | - | - | - | - | - | - | - | - | - | - | - | - | Y | V | A | S | C | S | A | D | G | T | V | K | L | - | - | - | - | - | - | - | - | - | - | - | - | - | - | - | - | - | - | - | W | E | - | T | G | S | G | N | E | L | K | C | - | - | - | - | - | - | I | R | - | - | - | - | - | - | - | - | - | - | - | - | - | - | - | - | - | - | - | - | - | - | I | K | D | F | - | - | - | - | - | - | - | - | F | R | N | A | D | E | Q | - | - | - | - | - | - | - | - | - | - | - | - | - | - | - | - | - | - | - | - | - | - | - | - | - | - | - | - | - | E | V | E | - | A | L | P | K | C | C | - | - | - | - | - | - | - | - | - | - | - | - | - | - | - | - | S | W | S | T | D | G | A | T | V | M | V | A | A | K | N | K | L | - | - | - | - | - | - | - | - | - | - | - | L | L | L | D | V | E | T | - | - | - | S | D | - | - | - | - | - | - | - | - | - | - | - | - | - | - | - | - | - | - | - | - | - | - | - | - | - | - | - | - | - | - | - | - | - | - | - | - | - | - | - | - | - | - | - | - | - | - | - | - | - | - | - | - | - | - | - | - | - | - | - | - | - | - | - | - | - | - | - | - | - | - | - | - | - | - | - | - | - | - | - | - | - | - | - | - | - | - | - | - | - | - | - | - | - | - | - | - | - | - | - | - | - | - | - | - | - | - | - | - | - | - | - | - | - | - | - | - | - | - | - | - | - | - | - | - | - | - | L | L | E | - | - | - | - | - | - | - | - | - | - | - | - | - | - | - | - | - | - | - | - | - | - | - | - | - | - | - | - | - | - | - | - | - | - | - | - | - | - | - | - | - | - | - | - | - | - | - | - | - | - | - | - | - | - | - | - | - | - | - | - | - | - | - | - | - | - | - | - | - | - | - | - | - | - | - | - | - | - | - | - | - | - | - | - | - | - | - | - | - | - | - | - | - | - | - | - | - | - | - | - | - | - | - | - | - | - | - | - | - | - | - | - | - | - | - | - | - | - | - | - | - | - | - | - | - | - | - | - | - | - | - | - | - | - | - | - | - | - | - | - | - | - | - | - | - | - | - | - | - | - | - | - | - | - | - | - | - | - | - | - | - | - | - | - | - | - | - | - | - | - | - | - | - | - | - | - | - | - | - | - | - | - | - | E | V | - | - | - | L | - | S | Y | Q | N | T | I | Q | Y | Y | - | - | - | - | - | - | - | - | - | - | - | - | - | - | - | - | - | - | - | - | - | - | - | - | - | D | F | C | P | S | N | Q | L | V | A | V | A | F | S | H | - | - | - | - | - | - | Y | S | V | E | - | - | - | - | - | - | - | - | - | - | - | - | - | - | - | - | - | L | W | N | T | K | T | K | - | - | - | - | - | - | - | - | - | - | - | - | - | - | - | - | - | - | - | - | - | - | - | - | - | - | - | - | - | T | K | V | A | D | C | R | E | H | M | S | - | - | - | - | - | - | - | - | - | - | - | - | - | - | - | - | - | - | - | - | - | - | - | - | - | - | - | - | - | - | - | - | - | - | - | - | - | - | - | - | - | - | - | - | W | V | H | C | V | T | F | S | - | - | - | - | - | - | - | - | - | - | - | P | D | G | S | - | - | - | - | - | - | - | - | - | - | L | F | L | T | S | S | N | D | Q | - | - | - | - | - | - | - | - | - | - | - | - | - | - | T | I | R | I | - | W | - | - | - | - | - | - | - | - | - | - | - | - | - | - | - | - | - | - | - | - | - | - | - | - | - | - | - | - | - | - | - | - | - | - | - | - | - | - | E | T | S | K | V | C | K | P | P | E | I | A | L | K | N | E | - | - | - | - | - | - | - | - | - | - | - | - | - | - | - | - | L | D | V | A | F | - | - | - | - | - | - | - | - | - | - | - | - | - | - | - | - | - | - | - | - | - | - | - | - | - | - | - | - | - | - | - | - | - | - | - | - | - | - | - | - | - | - | - | - | - | - | - | - | - | - | - | - | - | - | K | G | D | E | M | M | - | - | - | - | - | - | - | - | - | - | - | - | - | - | - | - | - | - | - | - | - | - | - | - | - | - | - | - | - | - | - | - | - | - | - | - | I | L | A | I | L | N | H | T | H | L | Q | L | - | I | N | G | N | T | - | - | - | - | - | - | - | - | - | - | G | H | T | V | S | L | T | D | V | - | - | - | - | - | - | - | - | - | - | - | - | - | - | - | - | - | - | - | - | - | - | E | D | S | - | - | - | - | - | - | - | - | - | I | S | C | C | C | L | S | E | D | L | Q | - | - | - | - | - | - | - | - | - | F | A | A | F | G | Q | - | - | - | - | - | - | - | - | - | - | - | - | - | - | - | - | - | - | - | - | - | - | - | - | - | - | - | - | - | - | - | - | - | - | - | - | K | - | - | - | - | - | - | - | - | - | N | G | T | - | - | - | - | - | - | - | - | - | - | V | - | - | - | - | - | - | - | - | - | - | - | - | - | - | - | - | - | - | - | - | - | - | - | - | - | - | - | - | - | - | - | - | - | - | - | - | - | - | - | K | V | L | E | V | S | G | G | R | V | L | K | S | R | R | G | - | - | - | - | - | H | - | - | - | - | - | - | - | - | - | - | - | - | K | T | S | V | Q | H | C | L | F | T | - | - | - | - | - | - | - | - | - | - | - | - | - | - | - | - | - | - | - | - | - | - | - | - | - | - | - | - | - | - | - | - | - | - | - | - | - | S | D | G | Q | T | L | I | S | S | D | - | D | D | P | - | - | - | - | - | - | - | - | I | I | Q | - | V | - | - | - | - | - | - | - | - | - | - | - | - | - | - | - | - | - | - | - | - | - | - | - | - | - | - | - | - | - | - | - | - | - | - | - | - | - | - | - | W | N | - | - | - | - | - | - | - | - | - | - | W | E | L | E | E | C | L | F | - | - | - | - | - | - | - | - | - | - | - | - | - | - | - | - | - | - | - | - | - | - | - | - | - | - | - | - | - | - | - | - | - | - | - | - | - | - | - | - | - | - | - | - | - | - | - | - | - | - | L | E | G | H | N | - | - | - | - | - | - | - | - | - | - | - | - | - | - | - | - | - | - | - | - | - | - | - | - | - | - | E | T | I | K | N | F | R | L | L | - | - | - | - | - | - | - | - | - | - | - | - | - | - | - | - | - | - | - | - | - | - | - | E | N | - | S | R | L | L | - | S | W | - | - | - | S | F | D | G | T | V | - | - | - | - | - | - | - | - | - | - | - | - | - | - | - | - | - | - | - | - | - | - | - | - | - | - | - | - | - | - | - | - | - | - | - | K | V | W | N | I | I | H | G | K | I | - | - | - | - | - | E | V | N | F | A | C | - | - | - | - | - | - | - | H | K | - | - | - | - | - | - | - | - | - | - | - | - | - | - | - | - | - | - | - | D | T | V | L | - | - | - | - | - | - | - | - | - | - | - | - | - | - | - | S | C | A | - | - | - | - | - | - | - | - | - | - | - | - | - | - | - | V | S | P | D | A | T | - | K | F | S | S | T | - | S | V | D | K | T | - | - | - | - | - | - | A | K | I | W | S | F | K | S | S | - | - | - | - | - | - | - | - | - | - | - | - | - | - | - | - | - | - | - | - | - | - | - | - | S | I | L | H | E | - | - | - | - | - | - | - | - | L | - | - | - | - | K | G | H | E | A | C | V | - | - | - | - | - | - | - | - | - | R | C | C | - | T | F | S | - | - | - | - | - | - | - | - | - | F | D | N | K | - | L | L | A | T | - | - | - | - | - | - | - | - | G | D | D | K | R | E | I | R | I | W | D | V | S | S | G | E | L | L | - | - | - | - | - | - | - | - | - | - | - | - | - | - | - | - | - | - | - | - | - | - | - | - | - | - | - | - | - | - | - | - | - | - | - | - | - | - | - | - | - | - | - | - | - | - | - | - | - | - | - | - | - | - | - | - | H | L | C | S | P | V | I | I | D | - | - | - | - | - | - | - | - | - | - | - | - | - | - | - | - | - | - | - | - | E | G | G | L | T | H | D | Y | W | - | - | - | - | - | - | - | - | V | T | D | L | C | F | S | P | D | S | K | - | - | - | - | - | - | - | - | - | - | - | - | - | - | - | - | - | - | - | - | - | - | - | - | - | - | - | - | - | - | - | - | - | - | - | - | - | - | - | - | - | - | - | - | - | M | L | V | S | - | - | S | G | G | Y | L | K | - | - | - | - | - | - | - | - | W | W | N | A | V | T | - | - | - | - | - | - | - | - | - | - | - | - | - | - | - | - | - | - | - | - | - | - | - | - | - | - | - | - | - | - | - | - | - | - | - | - | - | - | - | - | - | - | - | - | - | - | - | - | - | - | - | - | - | - | - | - | - | - | - | - | - | - | - | - | - | - | - | - | - | - | - | - | - | - | - | - | - | - | - | - | - | - | - | - | - | - | - | - | - | - | - | - | - | - | - | - | - | - | - | - | - | - | - | - | - | - | - | - | - | - | - | - | - | - | - | - | - | - | - | - | - | - | - | - | - | - | - | - | - | - | - | - | - | - | - | - | - | - | - | - | - | - | - | - | - | - | - | - | - | - | - | - | - | - | - | - | - | - | - | - | - | - | - | - | - | - | - | - | - | - | - | - | - | - | - | - | - | - | - | - | - | - | - | - | - | - | - | - | - | - | - | - | - | - | - | - | - | - | - | - | - | - | - | - | - | - | - | - | - | - | - | - | - | - | - | - | - | - | - | - | - | - | - | - | - | - | - | - | - | - | - | - | - | - | - | - | - | - | - | - | - | - | - | - | - | - | - | - | - | - | - | - | - | - | - | - | - | - | - | - | - | - | - | - | - | - | - | G | E | A | L | Q | T | F | Y | T | - | - | - | - | - | N | G | - | - | - | T | D | L | Q | S | I | - | - | - | Y | V | S | S | - | - | - | - | N | F | K | I | - | - | - | - | - | - | - | - | - | - | - | - | - | - | - | - | - | - | - | - | - | - | - | - | - | - | Y | V | T | V | D | - | N | L | G | I | L | - | - | Y | V | L | Q | M | - | - | - | - | - | - | - | - | - | - | - | - | - | - | - | - | - | - | - | - | - | - | - | - | - | - | - | - | - | - | - | - | - | - | M | E | - | - | - | - | - | - | - | - | - | - | - | - | - | - | - | - | - | - | - | - | - | - | - | - | - | - | - | - | - | - | - | - | - | - | - | - | - | - | - | - | - | - | - | - | - | - | - | - | - | - | - | - | M | - | - | - | - | - | - | - | - | - | - | - | - | - | - | - | - | - | - | - | - | - | - | - | - | - | - | - | - | - | - | - | - | - | - | - | - | - | - | - | - | - | - | - | - | - | - | - | - | - | - | - | - | - | - | E | A | E | Q | R | Q | - | - | - | - | - | - | - | - | - | - | - | - | - | - | - | - | - | - | - | - | R | L | Q | R | - | - | - | - | - | - | - | - | - | - | - | - | - | - | - | - | - | - | - | - | - | - | - | - | - | - | - | - | - | - | G | R | - | - | - | - | - | - | - | - | V | R | L | - | V | R | G | L | Q | V | E | A | L | W | - | - | - | - | - | - | - | - | - | - | - | - | - | - | - | - | - | - | - | - | - | - | - | - | - | - | - | - | - | - | - | - | - | - | - | - | - | - | - | - | - | - | - | - | - | G | P | L | V | R | D | G | V | - | - | - | - | - | - | - | - | - | - | - | - | - | - | - | - | - | - | - | - | - | - | - | - | F | S | R | D | M | L | D | D | - | - | - | - | - | - | - | - | - | - | - | - | - | - | - | - | - | - | - | - | - | - | - | - | - | I | Q | R | A | - | - | - | - | - | - | - | - | - | - | G | T | L | R | D | Q | - | A | R | Q | L | L | I | D | L | E | T | R | G | - | K | Q | A | F | P | T | F | V | S | I | L | H | E | T | G | Q | K | D | L | A | D | F | L | S | E | G | C | A | F | P | - | - | - | - | R | P | S | S | - | - | - | - | - | - | - | - | - | - | - | - | - | - | - | - | - | - | - | - | - | - | - | - | - | - | - | - | - | - | - | - | I | D | - | - | - | - | - | - | - | - | - | - | - | - | - | - | - | I | K | P | I | - | - | - | - | - | - | - | - | - | - | - | - | - | - | - | - | - | - | - | - | - | - | - | - | - | - | - | - | - | - | - | - | - | - | - | - | - | - | - | E | I | G | L | R | G | G | K | G | R | A | G | G | - | - | - | - | - | - | - | - | - | - | - | R | - | - | - | - | - | - | - | - | - | - | - | - | - | - | - | - | - | - | - | - | - | - | - | - | - | - | - | - | - | - | - | - | - | - | - | - | - | - | - | - | - | - | - | - | - | - | - | - | - | - | - | - | - | - | - | - | - | - | - | - | - | - | - | - | - | - | - | - | - | - | - | - | - | - | - | - | - | - | - | - | - | - | - | - | - | - | - | - | - | - | - | - | - | - | - | - | - | - | - | - | - | - | - | - | - | - | - | - | - | - | - | - | - | - | - | - | - | - | - | - | - | - | - | - | - | - | - | - | - | - | - | - | - | - | - | - | - | - | - | - | - | - | - | - | - | - | - | - | - | - | - | - | - | - | - | - | D | S | E | Y | L | P | - | - | - | - | - | - | - | - | - | - | - | - | - | - | - | - | - | - | - | - | - | - | - | - | - | - | L | P | V | Q | A | - | - | - | - | - | - | R | T | E | R | D | - | Q | E | L | D | - | - | - | - | - | - | - | - | - | - | - | - | - | - | - | H | P | G | A | V | A | H | S | - | - | - | - | - | - | - | - | - | - | - | - | - | - | - | - | - | - | - | - | A | - | - | - | - | - | - | - | - | - | - | A | - | A | D | R | S | R | Q | N | S | D | - | - | - | - | - | M | V | Y | V | - | - | L | K | S | R | - | P | C | G | Y | C | L | I | I | N | N | V | H | F | S | P | E | S | - | - | G | L | S | L | R | Q | G | S | D | L | D | Y | E | K | L | K | R | R | F | - | S | L | F | H | F | E | V | L | T | R | Q | D | L | K | A | K | E | M | Y | S | E | L | W | A | L | A | Q | K | D | H | - | S | A | M | D | C | C | L | V | V | I | L | S | H | G | R | Q | R | S | H | I | Q | F | P | G | E | - | V | C | G | T | D | G | N | A | I | P | V | E | R | I | V | N | C | F | N | G | L | N | C | W | S | L | R | G | K | P | K | L | F | I | I | Q | A | C | - | - | G | G | E | E | R | D | Y | G | F | E | L | D | S | D | - | - | - | S | S | G | N | - | Q | G | Y | G | A | S | - | - | - | - | - | - | - | L | Q | S | D | A | S | P | L | K | P | S | S | G | N | L | - | - | - | - | - | - | - | - | D | E | P | D | A | V | - | - | - | - | - | - | - | T | S | L | - | - | - | P | T | A | S | D | I | L | V | S | Y | S | T | F | P | - | - | G | - | - | - | - | - | - | - | - | F | V | S | W | R | D | R | K | S | G | S | W | F | V | E | T | L | D | R | I | L | E | Q | Y | A | D | - | S | - | - | - | - | - | - | - | - | - | - | - | - | - | - | - | - | - | - | - | - | - | - | - | - | - | - | - | - | - | - | - | E | D | L | Q | K | L | L | - | - | - | - | - | L | R | - | - | - | - | - | - | - | - | - | - | - | - | V | A | N | E | V | - | - | - | - | - | S | S | K | - | - | - | - | - | - | - | E | K | Y | - | K | Q | M | - | - | P | G | - | - | - | - | - | - | - | - | - | - | - | - | - | - | - | C | Y | H | F | - | - | - | - | - | - | - | - | - | L | R | K | W | F | F | F | M | T | K | - | - | - | - | - | - | - | - | - | - | - | - | - | - | - | - | - | - | - | - | - | - | - | - | - | - | - | - | - | - | - | - | - | - | - | - | - | M | - | - | - | - | - | - | - | - | - | - | - | - | - | - | - | - | - | - | - | - | - | - | - | - | - | - | - | - | - | - | - | - | - | - | - | - | - | - | - | - | - | - | - | - | - | - | - | - | - | - | - | - | G | D | V | E | K | G | K | K | I | F | V | Q | K | C | A | Q | C | H | T | V | E | K | G | G | K | H | K | T | G | P | N | L | H | G | L | I | G | R | K | T | G | Q | A | P | G | F | S | Y | T | E | A | N | K | N | K | G | I | T | W | G | E | E | T | L | M | E | Y | L | E | N | P | K | K | Y | I | P | G | T | K | M | I | F | A | G | I | K | K | K | P | E | R | A | D | L | I | A | Y | L | K | E | A | T | A | N | - | - | - | - | - | - |
| Anas\_platyrhynchos | M | - | - | - | - | - | - | - | - | - | - | - | - | - | - | - | - | - | - | - | - | - | - | - | - | - | - | - | - | - | - | - | - | - | - | - | - | - | - | - | - | - | - | - | - | - | - | - | - | - | - | - | - | - | - | - | - | - | - | - | - | - | - | - | - | - | - | - | - | - | - | - | - | - | - | - | - | - | - | - | - | - | - | - | - | - | - | - | - | - | - | - | - | - | - | - | - | - | D | V | K | S | - | R | N | - | - | - | - | - | - | - | - | - | - | - | - | - | - | - | - | Y | L | L | - | - | - | - | - | - | - | - | - | - | - | - | - | - | - | - | - | - | - | - | - | - | - | - | - | - | - | - | - | - | - | - | - | - | - | - | - | - | - | - | - | - | - | - | - | - | - | - | - | - | - | - | - | - | - | - | - | - | - | - | - | - | - | - | - | - | - | - | - | - | - | - | - | - | - | - | - | - | - | - | - | - | - | - | - | - | - | - | - | - | - | - | - | - | - | - | - | - | - | - | - | - | - | - | - | - | - | - | - | - | - | - | - | - | - | - | - | - | - | - | - | M | N | R | Q | A | L | E | R | D | I | K | T | C | Y | - | I | M | D | H | M | I | S | D | E | V | L | T | T | E | E | E | E | K | V | K | Q | Q | - | - | - | - | - | - | - | - | - | - | - | - | - | - | - | - | - | N | T | Q | K | E | R | A | A | M | L | I | N | I | I | L | T | K | - | - | - | - | D | N | N | S | Y | R | S | - | - | - | - | - | F | Y | N | A | L | - | L | H | E | G | - | - | - | - | - | - | Y | R | D | L | A | A | L | L | Q | D | G | I | P | V | F | S | - | G | N | G | - | K | - | - | - | - | - | - | - | - | - | - | - | - | - | - | - | - | - | - | - | - | - | - | S | S | V | D | G | M | T | - | - | - | - | - | - | - | - | - | - | - | S | Y | V | K | T | V | L | C | E | G | G | - | - | - | - | - | - | - | V | P | Q | R | P | V | V | F | V | T | R | P | E | L | V | D | A | I | - | - | - | - | - | - | - | - | - | - | K | Q | K | L | R | - | - | - | - | - | - | - | - | - | - | - | - | - | - | - | - | - | - | - | - | - | - | - | - | - | - | - | - | - | - | - | - | S | L | G | S | D | P | - | - | - | - | - | - | - | - | - | - | - | - | - | - | - | - | - | - | - | - | - | - | - | - | - | - | - | - | - | - | - | - | - | - | - | - | - | - | G | W | V | T | V | Y | G | M | A | G | C | G | K | T | V | L | T | A | E | A | L | - | R | D | H | Q | L | L | E | D | Y | F | P | G | G | V | H | W | I | S | V | G | - | - | - | - | - | - | - | K | - | - | - | - | - | - | Q | D | K | A | - | - | - | - | - | - | - | - | - | - | - | - | G | L | L | I | K | L | Q | N | L | C | S | R | L | E | - | - | - | - | H | - | - | - | - | - | - | - | - | - | - | - | - | - | - | - | D | S | T | L | S | Q | R | P | P | L | N | I | - | - | - | - | - | - | - | - | - | - | - | - | - | - | - | - | E | E | A | K | D | R | L | R | L | L | M | L | R | - | - | - | - | - | - | - | - | - | - | - | - | K | Y | P | R | S | L | L | - | - | V | L | D | D | I | W | D | S | W | V | L | K | A | F | D | - | N | Q | C | Q | V | L | I | T | S | R | D | R | S | V | T | D | A | V | - | S | G | N | - | - | K | Y | E | I | H | V | E | S | - | - | - | G | L | A | H | E | K | G | L | E | V | L | S | L | F | V | N | M | - | K | I | P | E | L | P | E | - | - | - | - | - | - | - | - | - | - | - | - | - | - | Q | A | N | C | I | I | R | E | C | K | - | - | - | - | - | - | - | - | - | - | - | - | - | - | - | - | - | - | - | G | - | S | P | L | V | I | S | L | I | G | A | L | L | R | D | F | P | S | - | - | - | - | - | - | - | - | - | - | R | W | E | Y | Y | L | - | K | Q | L | Q | N | K | Q | F | K | R | I | - | - | - | - | - | - | - | - | - | - | - | - | - | - | - | - | - | - | - | - | - | - | R | K | S | S | S | Y | D | - | - | - | - | Y | E | A | L | D | E | A | M | S | I | S | V | E | - | Q | L | D | - | - | - | N | D | F | K | D | Y | - | Y | K | D | L | S | I | L | P | K | D | V | K | V | P | T | K | V | L | C | I | L | W | D | - | - | - | - | M | E | T | E | E | V | E | D | I | L | Q | E | F | - | - | - | - | - | - | - | - | - | - | - | V | N | K | S | L | L | F | C | D | R | N | G | K | S | - | - | - | - | - | - | - | - | - | F | H | Y | Y | L | H | D | L | Q | L | D | F | L | T | E | K | - | - | N | R | N | Q | - | L | - | Q | E | L | H | K | N | I | V | N | Q | Y | K | K | - | H | Y | - | - | - | - | - | - | K | L | N | M | P | I | P | S | - | Q | E | - | - | - | - | - | - | - | D | C | M | Y | W | Y | N | F | L | A | Y | H | M | A | G | A | N | - | - | - | - | M | Q | K | E | L | R | E | L | M | F | S | L | D | W | I | K | A | K | T | E | L | V | G | P | A | - | - | - | - | - | - | - | - | H | L | I | H | E | Y | V | E | Y | S | S | V | L | D | Q | K | - | - | - | - | - | - | - | D | S | I | V | R | E | N | F | Q | - | - | - | E | F | L | S | L | N | G | H | L | L | G | R | Q | P | F | - | - | - | - | - | - | - | - | - | - | - | - | - | - | - | - | - | - | - | - | - | - | - | - | - | - | - | - | - | - | - | - | - | - | - | - | - | - | - | - | - | - | - | - | - | - | - | - | - | - | - | - | - | - | - | - | - | - | - | - | - | - | - | - | - | - | - | - | - | - | - | P | D | I | I | Q | L | G | L | C | Q | P | D | T | S | E | V | Y | Q | Q | A | K | L | R | A | Q | S | E | - | - | T | G | - | - | - | - | - | - | - | - | - | - | - | - | - | - | - | - | - | - | - | - | - | - | - | - | - | - | - | - | - | A | F | Y | L | - | - | - | - | - | - | - | - | - | - | E | W | I | N | K | - | - | - | - | - | - | - | K | S | L | K | N | L | S | R | - | - | - | - | - | - | - | - | - | - | L | V | V | R | P | H | K | D | - | - | - | - | - | - | - | - | - | - | - | - | - | - | - | A | V | Y | H | A | C | - | - | - | F | S | - | Q | D | R | Q | R | I | A | - | - | - | - | - | - | - | - | - | S | C | G | A | D | K | T | L | Q | V | F | K | - | - | - | - | - | T | E | S | G | E | M | L | L | E | I | - | T | A | H | E | D | E | - | - | - | I | L | C | C | A | F | S | - | A | - | - | - | - | - | - | - | - | - | - | - | - | - | D | D | K | Y | V | A | T | C | S | A | D | K | K | V | K | I | - | - | - | W | N | S | R | - | - | - | - | - | - | - | - | - | - | - | - | - | - | - | - | - | - | - | - | - | - | - | - | - | - | - | - | - | - | - | - | - | - | - | - | - | - | - | - | - | - | - | - | - | - | - | - | - | - | - | - | - | - | - | - | - | - | - | - | - | - | - | - | - | - | - | - | - | - | - | - | - | - | - | - | - | - | - | - | - | - | - | - | - | - | - | - | - | - | - | - | - | - | - | - | - | - | - | - | - | - | - | - | - | - | - | - | - | - | - | - | - | - | - | - | - | - | - | - | - | - | - | - | - | - | - | - | - | - | - | - | - | - | - | - | - | - | - | - | T | G | K | C | M | H | I | F | - | E | E | H | T | - | - | - | - | - | E | Q | V | N | - | - | - | - | - | - | - | - | - | - | - | - | - | C | C | Q | - | - | - | - | - | - | - | - | - | - | - | - | - | F | N | N | R | S | - | - | - | - | - | - | - | - | - | - | - | - | - | - | - | - | G | Q | - | Y | L | L | A | T | C | S | N | - | - | - | - | D | T | F | I | - | - | - | - | - | - | - | - | - | - | - | - | K | L | W | D | L | - | - | - | K | G | K | Y | C | R | N | T | M | F | - | - | - | - | - | - | - | - | - | - | - | - | - | - | - | - | - | G | H | V | N | S | - | - | - | - | - | - | - | - | - | - | - | - | - | - | - | - | V | N | H | C | R | F | S | P | N | D | E | - | - | - | - | - | - | - | - | - | - | - | - | - | - | - | - | - | - | - | - | - | - | - | - | - | - | - | - | Y | V | A | S | C | S | T | D | G | T | V | K | L | - | - | - | - | - | - | - | - | - | - | - | - | - | - | - | - | - | - | - | W | E | - | A | R | S | A | N | E | L | K | T | - | - | - | - | - | - | I | E | - | - | - | - | - | - | - | - | - | - | - | - | - | - | - | - | - | - | - | - | - | - | I | K | D | F | - | - | - | - | - | - | - | - | F | Q | N | A | D | E | Q | - | D | - | - | - | - | - | - | - | - | - | - | - | - | - | - | - | - | - | - | - | - | - | - | - | - | - | - | - | D | V | E | - | V | L | V | K | C | C | - | - | - | - | - | - | - | - | - | - | - | - | - | - | - | - | S | W | S | R | N | S | D | M | I | L | A | V | A | K | N | K | L | - | - | - | - | - | - | - | - | - | - | - | L | L | F | D | V | K | T | - | - | - | S | D | - | - | - | - | - | - | - | - | - | - | - | - | - | - | - | - | - | - | - | - | - | - | - | - | - | - | - | - | - | - | - | - | - | - | - | - | - | - | - | - | - | - | - | - | - | - | - | - | - | - | - | - | - | - | - | - | - | - | - | - | - | - | - | - | - | - | - | - | - | - | - | - | - | - | - | - | - | - | - | - | - | - | - | - | - | - | - | - | - | - | - | - | - | - | - | - | - | - | - | - | - | - | - | - | - | - | - | - | - | - | - | - | - | - | - | - | - | - | - | - | - | - | - | - | - | - | L | L | T | - | - | - | - | - | - | - | - | - | - | - | - | - | - | - | - | - | - | - | - | - | - | - | - | - | - | - | - | - | - | - | - | - | - | - | - | - | - | - | - | - | - | - | - | - | - | - | - | - | - | - | - | - | - | - | - | - | - | - | - | - | - | - | - | - | - | - | - | - | - | - | - | - | - | - | - | - | - | - | - | - | - | - | - | - | - | - | - | - | - | - | - | - | - | - | - | - | - | - | - | - | - | - | - | - | - | - | - | - | - | - | - | - | - | - | - | - | - | - | - | - | - | - | - | - | - | - | - | - | - | - | - | - | - | - | - | - | - | - | - | - | - | - | - | - | - | - | - | - | - | - | - | - | - | - | - | - | - | - | - | - | - | - | - | - | - | - | - | - | - | - | - | - | - | - | - | - | - | - | - | - | - | - | Q | V | - | - | - | I | V | S | H | H | S | T | I | Q | Y | C | - | - | - | - | - | - | - | - | - | - | - | - | - | - | - | - | - | - | - | - | - | - | - | - | - | D | F | C | A | G | D | E | L | V | A | V | A | L | S | H | - | - | - | - | - | - | C | S | V | E | - | - | - | - | - | - | - | - | - | - | - | - | - | - | - | - | - | L | W | N | I | K | S | L | - | - | - | - | - | - | - | - | - | - | - | - | - | - | - | - | - | - | - | - | - | - | - | - | - | - | - | - | - | S | K | V | A | D | C | R | G | H | M | S | - | - | - | - | - | - | - | - | - | - | - | - | - | - | - | - | - | - | - | - | - | - | - | - | - | - | - | - | - | - | - | - | - | - | - | - | - | - | - | - | - | - | - | - | W | V | H | C | V | T | F | S | - | - | - | - | - | - | - | - | - | - | - | P | D | G | S | - | - | - | - | - | - | - | - | - | - | L | F | L | T | S | S | D | D | Q | - | - | - | - | - | - | - | - | - | - | - | - | - | - | T | I | R | I | - | W | - | - | - | - | - | - | - | - | - | - | - | - | - | - | - | - | - | - | - | - | - | - | - | - | - | - | - | - | - | - | - | - | - | - | - | - | - | - | E | T | N | K | V | C | K | S | S | D | A | E | L | K | S | E | - | - | - | - | - | - | - | - | - | - | - | - | - | - | - | - | L | D | A | V | F | - | - | - | - | - | - | - | - | - | - | - | - | - | - | - | - | - | - | - | - | - | - | - | - | - | - | - | - | - | - | - | - | - | - | - | - | - | - | - | - | - | - | - | - | - | - | - | - | - | - | - | - | - | - | H | N | G | E | V | M | - | - | - | - | - | - | - | - | - | - | - | - | - | - | - | - | - | - | - | - | - | - | - | - | - | - | - | - | - | - | - | - | - | - | - | - | I | L | A | I | Y | N | Q | K | H | I | Q | L | - | I | N | G | N | T | - | - | - | - | - | - | - | - | - | - | D | N | I | I | M | Q | T | E | P | - | - | - | - | - | - | - | - | - | - | - | - | - | - | - | - | - | - | - | - | - | - | Q | E | S | - | - | - | - | - | - | - | - | L | I | C | C | C | C | L | S | E | D | L | K | - | - | - | - | - | - | - | - | - | F | A | A | V | G | Q | - | - | - | - | - | - | - | - | - | - | - | - | - | - | - | - | - | - | - | - | - | - | - | - | - | - | - | - | - | - | - | - | - | - | - | - | E | - | - | - | - | - | - | - | - | - | N | G | T | - | - | - | - | - | - | - | - | - | - | V | - | - | - | - | - | - | - | - | - | - | - | - | - | - | - | - | - | - | - | - | - | - | - | - | - | - | - | - | - | - | - | - | - | - | - | - | - | - | - | K | V | L | Q | L | L | D | G | K | V | L | K | C | R | A | A | - | - | - | - | - | Y | - | - | - | - | - | - | - | - | - | - | - | - | K | T | S | V | Q | H | C | R | F | T | - | - | - | - | - | - | - | - | - | - | - | - | - | - | - | - | - | - | - | - | - | - | - | - | - | - | - | - | - | - | - | - | - | - | - | - | - | S | D | C | Q | T | L | I | S | S | A | - | H | D | A | - | - | - | - | - | - | - | - | V | I | Q | - | V | - | - | - | - | - | - | - | - | - | - | - | - | - | - | - | - | - | - | - | - | - | - | - | - | - | - | - | - | - | - | - | - | - | - | - | - | - | - | - | W | N | - | - | - | - | - | - | - | - | - | - | W | Q | L | N | E | C | V | F | - | - | - | - | - | - | - | - | - | - | - | - | - | - | - | - | - | - | - | - | - | - | - | - | - | - | - | - | - | - | - | - | - | - | - | - | - | - | - | - | - | - | - | - | - | - | - | - | - | - | L | R | G | H | K | - | - | - | - | - | - | - | - | - | - | - | - | - | - | - | - | - | - | - | - | - | - | - | - | - | - | E | A | I | K | D | F | R | L | L | - | - | - | - | - | - | - | - | - | - | - | - | - | - | - | - | - | - | - | - | - | - | - | E | N | - | S | K | L | L | - | S | W | - | - | - | S | F | D | G | T | V | - | - | - | - | - | - | - | - | - | - | - | - | - | - | - | - | - | - | - | - | - | - | - | - | - | - | - | - | - | - | - | - | - | - | - | K | V | W | N | I | I | T | G | K | T | - | - | - | - | - | E | K | D | F | A | C | - | - | - | - | - | - | - | H | G | - | - | - | - | - | - | - | - | - | - | - | - | - | - | - | - | - | - | - | G | A | V | L | - | - | - | - | - | - | - | - | - | - | - | - | - | - | - | S | C | A | - | - | - | - | - | - | - | - | - | - | - | - | - | - | - | V | S | P | D | G | S | - | K | F | A | S | T | - | S | A | D | K | T | - | - | - | - | - | - | A | K | I | W | S | F | E | S | S | - | - | - | - | - | - | - | - | - | - | - | - | - | - | - | - | - | - | - | - | - | - | - | - | C | V | L | H | E | - | - | - | - | - | - | - | - | L | - | - | - | - | K | G | H | E | A | C | V | - | - | - | - | - | - | - | - | - | R | C | C | - | T | F | S | - | - | - | - | - | - | - | - | - | P | N | N | K | - | L | L | A | T | - | - | - | - | - | - | - | - | G | D | D | K | G | D | I | R | I | W | D | I | L | T | G | G | L | L | - | - | - | - | - | - | - | - | - | - | - | - | - | - | - | - | - | - | - | - | - | - | - | - | - | - | - | - | - | - | - | - | - | - | - | - | - | - | - | - | - | - | - | - | - | - | - | - | - | - | - | - | - | - | - | - | H | F | C | S | L | V | T | V | D | - | - | - | - | - | - | - | - | - | - | - | - | - | - | - | - | - | - | - | - | E | G | E | P | T | H | G | G | W | - | - | - | - | - | - | - | - | V | T | D | L | S | F | S | S | D | S | K | - | - | - | - | - | - | - | - | - | - | - | - | - | - | - | - | - | - | - | - | - | - | - | - | - | - | - | - | - | - | - | - | - | - | - | - | - | - | - | - | - | - | - | - | - | M | L | V | S | - | - | S | G | G | Y | L | K | - | - | - | - | - | - | - | - | W | W | N | V | T | T | - | - | - | - | - | - | - | - | - | - | - | - | - | - | - | - | - | - | - | - | - | - | - | - | - | - | - | - | - | - | - | - | - | - | - | - | - | - | - | - | - | - | - | - | - | - | - | - | - | - | - | - | - | - | - | - | - | - | - | - | - | - | - | - | - | - | - | - | - | - | - | - | - | - | - | - | - | - | - | - | - | - | - | - | - | - | - | - | - | - | - | - | - | - | - | - | - | - | - | - | - | - | - | - | - | - | - | - | - | - | - | - | - | - | - | - | - | - | - | - | - | - | - | - | - | - | - | - | - | - | - | - | - | - | - | - | - | - | - | - | - | - | - | - | - | - | - | - | - | - | - | - | - | - | - | - | - | - | - | - | - | - | - | - | - | - | - | - | - | - | - | - | - | - | - | - | - | - | - | - | - | - | - | - | - | - | - | - | - | - | - | - | - | - | - | - | - | - | - | - | - | - | - | - | - | - | - | - | - | - | - | - | - | - | - | - | - | - | - | - | - | - | - | - | - | - | - | - | - | - | - | - | - | - | - | - | - | - | - | - | - | - | - | - | - | - | - | - | - | - | - | - | - | - | - | - | - | - | - | - | - | - | - | - | - | - | - | G | E | S | L | Q | T | F | Y | T | - | - | - | - | - | N | G | - | - | - | T | N | L | K | S | I | - | - | - | H | V | S | P | - | - | - | - | D | F | K | V | - | - | - | - | - | - | - | - | - | - | - | - | - | - | - | - | - | - | - | - | - | - | - | - | - | - | Y | V | T | V | D | - | N | L | G | I | L | - | - | Y | V | L | Q | K | - | - | - | - | - | - | - | - | - | - | - | - | - | - | - | - | - | - | - | - | - | - | - | - | - | - | - | - | - | - | - | - | - | - | L | - | - | - | - | - | - | - | - | - | - | - | - | - | - | - | - | - | - | - | - | - | - | - | - | - | - | - | - | - | - | - | - | - | - | - | - | - | - | - | - | - | - | - | - | - | - | - | - | - | - | - | - | - | M | G | P | Y | S | C | V | W | - | - | - | - | - | - | - | - | - | - | - | - | - | - | - | - | - | - | - | - | - | - | - | - | - | - | - | - | - | - | - | - | - | - | - | - | - | - | - | - | - | - | - | - | - | - | V | P | T | A | T | Y | W | - | - | - | - | - | - | - | - | - | - | D | P | - | - | - | - | - | - | - | - | - | - | - | - | - | - | - | - | - | - | - | - | - | T | A | - | - | - | - | - | - | - | - | - | - | - | - | - | - | - | - | - | - | - | G | Y | - | - | - | - | - | - | - | G | - | - | - | - | A | L | Q | L | C | M | G | P | Y | S | - | - | - | - | - | - | - | - | - | - | - | - | - | - | - | - | - | - | - | - | - | - | - | - | - | - | - | - | - | - | - | - | - | - | - | - | - | - | - | - | - | - | - | - | - | H | L | L | G | P | Y | D | W | - | - | - | - | - | - | - | - | - | - | - | - | - | - | - | - | - | - | - | - | - | - | - | - | A | S | S | C | F | F | L | P | - | - | - | - | - | - | - | - | - | - | - | - | - | - | - | - | - | - | - | - | - | - | - | - | - | P | Q | S | L | - | - | - | - | - | - | - | - | - | - | R | T | R | R | E | Q | - | V | R | Q | L | V | I | D | L | E | S | R | G | - | Q | Q | A | F | P | V | F | L | S | I | L | R | D | T | G | Q | G | E | L | A | D | A | L | S | E | A | C | G | S | L | - | - | - | - | P | S | Q | P | - | - | - | - | - | - | - | - | - | - | - | - | - | - | - | - | - | - | - | - | - | - | - | - | - | - | - | - | - | - | - | - | I | N | - | - | - | - | - | - | - | - | - | - | - | - | - | - | - | L | R | P | V | - | - | - | - | - | - | - | - | - | - | - | - | - | - | - | - | - | - | - | - | - | - | - | - | - | - | - | - | - | - | - | - | - | - | - | - | - | - | - | Q | L | E | L | S | G | E | K | R | G | K | S | E | - | - | - | - | - | - | - | - | - | - | - | S | - | - | - | - | - | - | - | - | - | - | - | - | - | - | - | - | - | - | - | - | - | - | - | - | - | - | - | - | - | - | - | - | - | - | - | - | - | - | - | - | - | - | - | - | - | - | - | - | - | - | - | - | - | - | - | - | - | - | - | - | - | - | - | - | - | - | - | - | - | - | - | - | - | - | - | - | - | - | - | - | - | - | - | - | - | - | - | - | - | - | - | - | - | - | - | - | - | - | - | - | - | - | - | - | - | - | - | - | - | - | - | - | - | - | - | - | - | - | - | - | - | - | - | - | - | - | - | - | - | - | - | - | - | - | - | - | - | - | - | - | - | - | - | - | - | - | - | - | - | - | - | - | - | - | - | - | T | V | E | R | L | S | - | - | - | - | - | - | - | - | - | - | - | - | - | - | - | - | - | - | - | - | - | - | - | - | - | - | I | P | V | Q | D | - | - | - | - | - | - | K | N | E | Q | F | - | R | A | - | - | - | - | - | - | - | - | - | - | - | - | - | - | - | - | - | - | P | P | A | P | A | K | G | - | - | - | - | - | - | - | - | - | - | - | - | - | - | - | - | - | - | - | - | S | - | - | - | - | - | - | - | - | - | - | A | - | V | D | - | - | K | S | D | Q | N | - | - | - | - | - | L | V | Y | Q | - | - | L | K | A | D | - | P | C | G | H | C | L | I | I | N | N | I | T | F | S | P | D | S | - | - | G | L | S | T | R | T | G | S | D | V | D | C | E | R | L | E | K | R | F | - | K | S | L | C | F | R | V | Q | T | L | R | N | L | K | A | Q | E | I | N | A | K | L | R | S | L | A | Q | Q | D | H | - | S | A | L | D | C | C | L | V | V | I | L | S | H | G | C | Q | T | S | H | N | Q | F | P | G | G | - | I | Y | G | T | D | G | I | S | I | P | I | E | K | I | V | K | H | F | D | G | S | N | C | P | S | L | R | G | K | P | K | L | F | F | I | Q | A | C | - | - | G | G | D | Q | K | D | Q | G | F | E | V | D | C | D | - | - | - | S | P | E | D | - | E | T | C | R | S | S | - | - | - | - | - | - | - | V | E | S | D | A | T | P | F | Q | T | P | S | G | N | L | - | - | - | - | - | - | - | - | D | E | P | D | A | V | - | - | - | - | - | - | - | A | S | L | - | - | - | P | T | P | G | D | I | L | V | S | Y | S | T | F | P | - | - | G | - | - | - | - | - | - | - | - | F | V | S | W | R | D | K | S | S | G | S | W | Y | V | E | T | L | D | S | M | L | G | Q | H | A | H | - | S | - | - | - | - | - | - | - | - | - | - | - | - | - | - | - | - | - | - | - | - | - | - | - | - | - | - | - | - | - | - | - | E | D | L | L | T | V | L | - | - | - | - | - | V | R | - | - | - | - | - | - | - | - | - | - | - | - | V | A | N | A | V | - | - | - | - | - | S | A | K | - | - | - | - | - | - | - | G | K | F | - | K | Q | I | - | - | P | G | - | - | - | - | - | - | - | - | - | - | - | - | - | - | - | C | F | N | F | - | - | - | - | - | - | - | - | - | L | R | K | K | F | F | F | M | C | K | - | - | - | - | - | - | - | - | - | - | - | - | - | - | - | - | - | - | - | - | - | - | - | - | - | - | - | - | - | - | - | - | - | - | - | - | - | M | - | - | - | - | - | - | - | - | - | - | - | - | - | - | - | - | - | - | - | - | - | - | - | - | - | - | - | - | - | - | - | - | - | - | - | - | - | - | - | - | - | - | - | - | - | - | - | - | - | - | - | - | G | D | V | E | K | G | K | K | I | F | V | Q | K | C | S | Q | C | H | T | V | E | K | G | G | K | H | K | T | G | P | N | L | H | G | L | F | G | R | K | T | G | Q | A | E | G | F | S | Y | T | D | A | N | K | N | K | G | I | T | W | G | E | D | T | L | M | E | Y | L | E | N | P | K | K | Y | I | P | G | T | K | M | I | F | A | G | I | K | K | K | S | E | R | A | D | L | I | A | Y | L | K | D | A | T | A | K | - | - | - | - | - | - |
| Anolis\_carolinensis | M | - | - | - | - | - | - | - | - | - | - | - | - | - | - | - | - | - | - | - | - | - | - | - | - | - | - | - | - | - | - | - | - | - | - | - | - | - | - | - | - | - | - | - | - | - | - | - | - | - | - | - | - | - | - | - | - | - | - | - | - | - | - | - | - | - | - | - | - | - | - | - | - | - | - | - | - | - | - | - | - | - | - | - | - | - | - | - | - | - | - | - | - | - | - | - | - | - | D | L | T | A | - | R | S | - | - | - | - | - | - | - | - | - | - | - | - | - | - | - | - | H | L | L | - | - | - | - | - | - | - | - | - | - | - | - | - | - | - | - | - | - | - | - | - | - | - | - | - | - | - | - | - | - | - | - | - | - | - | - | - | - | - | - | - | - | - | - | - | - | - | - | - | - | - | - | - | - | - | - | - | - | - | - | - | - | - | - | - | - | - | - | - | - | - | - | - | - | - | - | - | - | - | - | - | - | - | - | - | - | - | - | - | - | - | - | - | - | - | - | - | - | - | - | - | - | - | - | - | - | - | - | - | - | - | - | - | - | - | - | - | - | - | - | - | Q | N | R | E | A | L | E | R | D | I | K | T | S | Y | - | V | I | D | H | M | I | A | D | G | A | L | T | P | Q | E | E | A | E | V | K | A | V | - | - | - | - | - | - | - | - | - | - | - | - | - | - | - | - | - | T | T | Q | R | D | R | A | A | K | L | I | K | T | I | L | K | K | - | - | - | - | D | D | H | A | Y | I | S | - | - | - | - | - | F | Y | N | A | L | - | V | H | E | G | - | - | - | - | - | - | Y | K | D | L | A | A | L | L | Q | D | G | I | P | I | I | S | P | R | N | G | - | N | - | - | - | - | - | - | - | - | - | - | - | - | - | - | - | - | - | - | - | - | - | - | K | S | T | D | G | V | T | - | - | - | - | - | - | - | - | - | - | - | S | Y | V | K | T | I | L | C | E | G | G | - | - | - | - | - | - | - | V | P | Q | R | P | V | V | F | A | T | R | P | E | L | T | K | R | I | - | - | - | - | - | - | - | - | - | - | Q | R | K | L | Y | - | - | - | - | - | - | - | - | - | - | - | - | - | - | - | - | - | - | - | - | - | - | - | - | - | - | - | - | - | - | - | - | K | L | K | S | D | P | - | - | - | - | - | - | - | - | - | - | - | - | - | - | - | - | - | - | - | - | - | - | - | - | - | - | - | - | - | - | - | - | - | - | - | - | - | - | G | W | V | V | I | Y | G | M | A | G | C | G | K | T | V | L | A | A | D | A | L | - | R | D | H | N | L | L | K | D | C | F | P | G | G | V | H | W | V | S | I | G | - | - | - | - | - | - | - | K | - | - | - | - | - | - | Q | D | K | S | - | - | - | - | - | - | - | - | - | - | - | - | G | L | L | I | K | L | Q | N | L | C | L | R | L | D | - | - | - | - | Q | - | - | - | - | - | - | - | - | - | - | - | - | - | - | - | D | V | T | Y | L | Q | R | P | P | L | N | I | - | - | - | - | - | - | - | - | - | - | - | - | - | - | - | - | E | E | A | K | D | R | L | R | V | L | L | F | R | - | - | - | - | - | - | - | - | - | - | - | - | K | Y | P | R | S | L | L | - | - | V | L | D | D | I | W | D | S | W | V | L | K | A | F | D | - | S | H | F | Q | I | L | L | T | S | R | D | R | S | V | T | D | S | V | - | T | G | K | - | - | K | Y | E | V | P | V | E | S | - | - | - | G | L | S | H | E | K | A | L | E | I | L | S | L | F | V | N | M | - | E | I | T | E | L | P | E | - | - | - | - | - | - | - | - | - | - | - | - | - | - | K | A | D | W | I | V | K | E | C | K | - | - | - | - | - | - | - | - | - | - | - | - | - | - | - | - | - | - | - | G | - | S | P | L | V | V | S | L | I | G | A | L | L | R | D | F | P | N | - | - | - | - | - | - | - | - | - | - | R | W | E | Y | Y | L | - | R | Q | L | Q | N | K | Q | F | R | R | I | - | - | - | - | - | - | - | - | - | - | - | - | - | - | - | - | - | - | - | - | - | - | K | K | S | S | S | Y | D | - | - | - | - | Y | E | A | L | D | E | A | M | S | I | S | V | E | - | M | L | P | - | - | - | E | N | S | K | D | Y | - | Y | R | D | L | S | V | L | Q | K | D | V | K | V | P | S | K | V | F | C | V | L | W | D | - | - | - | - | I | E | T | E | E | A | E | D | I | L | Q | G | F | - | - | - | - | - | - | - | - | - | - | - | V | N | K | S | L | L | F | C | D | R | N | G | K | S | - | - | - | - | - | - | - | - | - | F | H | Y | Y | L | H | D | L | Q | L | D | F | L | T | E | K | - | - | N | R | D | Q | - | L | - | Q | A | L | H | K | K | V | V | H | Q | Y | K | N | - | H | Y | - | - | - | - | - | - | A | Q | Y | M | P | N | S | T | - | Q | E | - | - | - | - | - | - | - | D | W | M | Y | W | Y | N | F | L | A | Y | H | M | S | S | A | S | - | - | - | - | M | H | D | E | L | C | A | L | M | F | S | L | E | W | I | K | T | K | T | E | L | V | G | P | A | - | - | - | - | - | - | - | - | H | L | I | H | E | Y | V | E | Y | S | E | I | L | N | E | K | - | - | - | - | - | - | - | D | P | I | A | R | E | N | F | Q | - | - | - | E | F | L | S | L | N | G | H | L | L | G | R | Q | P | S | - | - | - | - | - | - | - | - | - | - | - | - | - | - | - | - | - | - | - | - | - | - | - | - | - | - | - | - | - | - | - | - | - | - | - | - | - | - | - | - | - | - | - | - | - | - | - | - | - | - | - | - | - | - | - | - | - | - | - | - | - | - | - | - | - | - | - | - | - | - | - | L | D | I | V | Q | L | G | L | C | Q | P | E | S | S | K | V | Y | Q | Q | A | K | L | Q | A | Q | Q | G | L | D | K | G | - | - | - | - | - | - | - | - | - | - | - | - | - | - | - | - | - | - | - | - | - | - | - | - | - | - | - | - | - | S | F | Y | V | - | - | - | - | - | - | - | - | - | - | E | W | M | N | K | - | - | - | - | - | - | - | Q | S | I | K | N | T | Y | R | - | - | - | - | - | - | - | - | - | - | L | V | V | R | P | H | T | D | - | - | - | - | - | - | - | - | - | - | - | - | - | - | - | A | V | Y | H | A | C | - | - | - | F | S | - | Q | D | R | H | R | I | A | - | - | - | - | - | - | - | - | - | S | C | G | A | D | K | T | L | Q | I | F | K | - | - | - | - | - | A | E | S | G | E | K | L | L | E | V | - | K | A | H | D | D | E | - | - | - | V | L | C | C | S | F | S | - | A | - | - | - | - | - | - | - | - | - | - | - | - | - | D | D | R | F | V | A | T | C | S | A | D | K | K | V | K | V | - | - | - | W | N | S | R | - | - | - | - | - | - | - | - | - | - | - | - | - | - | - | - | - | - | - | - | - | - | - | - | - | - | - | - | - | - | - | - | - | - | - | - | - | - | - | - | - | - | - | - | - | - | - | - | - | - | - | - | - | - | - | - | - | - | - | - | - | - | - | - | - | - | - | - | - | - | - | - | - | - | - | - | - | - | - | - | - | - | - | - | - | - | - | - | - | - | - | - | - | - | - | - | - | - | - | - | - | - | - | - | - | - | - | - | - | - | - | - | - | - | - | - | - | - | - | - | - | - | - | - | - | - | - | - | - | - | - | - | - | - | - | - | - | - | - | - | T | G | Q | P | V | C | I | F | - | E | E | H | T | - | - | - | - | - | E | Q | V | N | - | - | - | - | - | - | - | - | - | - | - | - | - | C | C | Q | - | - | - | - | - | - | - | - | - | - | - | - | - | F | S | N | T | N | - | - | - | - | - | - | - | - | - | - | - | - | - | - | - | - | H | N | Q | F | L | L | A | T | C | S | N | - | - | - | - | D | S | F | I | - | - | - | - | - | - | - | - | - | - | - | - | K | L | W | D | L | - | - | - | N | K | K | Y | C | R | N | T | L | F | - | - | - | - | - | - | - | - | - | - | - | - | - | - | - | - | - | D | H | S | S | S | - | - | - | - | - | - | - | - | - | - | - | - | - | - | - | - | V | T | H | C | R | F | S | P | N | D | K | - | - | - | - | - | - | - | - | - | - | - | - | - | - | - | - | - | - | - | - | - | - | - | - | - | - | - | - | Y | I | A | S | C | S | A | D | G | T | L | M | L | - | - | - | - | - | - | - | - | - | - | - | - | - | - | - | - | - | - | - | W | Y | - | A | Q | S | A | N | K | R | K | T | - | - | - | - | - | - | V | D | - | - | - | - | - | - | - | - | - | - | - | - | - | - | - | - | - | - | - | - | - | - | I | R | G | N | - | - | - | - | - | - | - | - | F | R | N | V | D | E | Y | Q | D | - | - | - | - | - | - | - | - | - | - | - | - | - | - | - | - | - | - | - | - | - | - | - | - | - | - | - | D | M | E | - | V | L | V | K | C | C | - | - | - | - | - | - | - | - | - | - | - | - | - | - | - | - | S | W | S | S | D | S | T | T | I | V | V | A | A | K | N | K | L | - | - | - | - | - | - | - | - | - | - | - | F | R | T | D | T | K | I | - | - | - | S | T | - | - | - | - | - | - | - | - | - | - | - | - | - | - | - | - | - | - | - | - | - | - | - | - | - | - | - | - | - | - | - | - | - | - | - | - | - | - | - | - | - | - | - | - | - | - | - | - | - | - | - | - | - | - | - | - | - | - | - | - | - | - | - | - | - | - | - | - | - | - | - | - | - | - | - | - | - | - | - | - | - | - | - | - | - | - | - | - | - | - | - | - | - | - | - | - | - | - | - | - | - | - | - | - | - | - | - | - | - | - | - | - | - | - | - | - | - | - | - | - | - | - | - | - | - | - | S | L | P | - | - | - | - | - | - | - | - | - | - | - | - | - | - | - | - | - | - | - | - | - | - | - | - | - | - | - | - | - | - | - | - | - | - | - | - | - | - | - | - | - | - | - | - | - | - | - | - | - | - | - | - | - | - | - | - | - | - | - | - | - | - | - | - | - | - | - | - | - | - | - | - | - | - | - | - | - | - | - | - | - | - | - | - | - | - | - | - | - | - | - | - | - | - | - | - | - | - | - | - | - | - | - | - | - | - | - | - | - | - | - | - | - | - | - | - | - | - | - | - | - | - | - | - | - | - | - | - | - | - | - | - | - | - | - | - | - | - | - | - | - | - | - | - | - | - | - | - | - | - | - | - | - | - | - | - | - | - | - | - | - | - | - | - | - | - | - | - | - | - | - | - | - | - | - | - | - | - | - | - | - | - | - | E | V | - | - | - | L | I | N | H | H | Y | T | I | Q | Y | C | - | - | - | - | - | - | - | - | - | - | - | - | - | - | - | - | - | - | - | - | - | - | - | - | - | D | F | C | P | N | S | Q | M | V | A | V | A | L | S | N | - | - | - | - | - | - | Y | S | V | E | - | - | - | - | - | - | - | - | - | - | - | - | - | - | - | - | - | I | W | N | I | E | S | C | - | - | - | - | - | - | - | - | - | - | - | - | - | - | - | - | - | - | - | - | - | - | - | - | - | - | - | - | - | S | K | I | A | D | C | R | G | H | M | S | - | - | - | - | - | - | - | - | - | - | - | - | - | - | - | - | - | - | - | - | - | - | - | - | - | - | - | - | - | - | - | - | - | - | - | - | - | - | - | - | - | - | - | - | W | V | H | S | V | T | F | S | - | - | - | - | - | - | - | - | - | - | - | F | D | G | S | - | - | - | - | - | - | - | - | - | - | L | F | L | T | S | S | D | D | Q | - | - | - | - | - | - | - | - | - | - | - | - | - | - | T | I | R | I | - | W | - | - | - | - | - | - | - | - | - | - | - | - | - | - | - | - | - | - | - | - | - | - | - | - | - | - | - | - | - | - | - | - | - | - | - | - | - | - | E | T | S | K | V | C | N | F | S | A | I | V | L | N | R | E | - | - | - | - | - | - | - | - | - | - | - | - | - | - | - | - | I | D | V | L | F | - | - | - | - | - | - | - | - | - | - | - | - | - | - | - | - | - | - | - | - | - | - | - | - | - | - | - | - | - | - | - | - | - | - | - | - | - | - | - | - | - | - | - | - | - | - | - | - | - | - | - | - | - | - | L | D | S | D | V | I | - | - | - | - | - | - | - | - | - | - | - | - | - | - | - | - | - | - | - | - | - | - | - | - | - | - | - | - | - | - | - | - | - | - | - | - | I | L | A | I | D | S | S | K | R | L | Q | L | - | I | S | G | Y | T | - | - | - | - | - | - | - | - | - | - | G | N | T | I | Y | L | T | K | E | - | - | - | - | - | - | - | - | - | - | - | - | - | - | - | - | - | - | - | - | - | - | Q | D | S | - | - | - | - | - | - | - | - | L | I | C | C | C | C | L | S | H | D | R | K | - | - | - | - | - | - | - | - | - | V | A | A | F | G | E | - | - | - | - | - | - | - | - | - | - | - | - | - | - | - | - | - | - | - | - | - | - | - | - | - | - | - | - | - | - | - | - | - | - | - | - | N | - | - | - | - | - | - | - | - | - | N | G | T | - | - | - | - | - | - | - | - | - | - | V | - | - | - | - | - | - | - | - | - | - | - | - | - | - | - | - | - | - | - | - | - | - | - | - | - | - | - | - | - | - | - | - | - | - | - | - | - | - | - | K | V | L | S | L | Q | N | G | N | T | L | N | S | R | T | G | - | - | - | - | - | H | - | - | - | - | - | - | - | - | - | - | - | - | R | E | A | V | Q | H | C | Q | F | T | - | - | - | - | - | - | - | - | - | - | - | - | - | - | - | - | - | - | - | - | - | - | - | - | - | - | - | - | - | - | - | - | - | - | - | - | - | S | D | G | K | T | L | I | S | S | S | - | D | D | S | - | - | - | - | - | - | - | - | T | I | Q | - | V | - | - | - | - | - | - | - | - | - | - | - | - | - | - | - | - | - | - | - | - | - | - | - | - | - | - | - | - | - | - | - | - | - | - | - | - | - | - | - | W | N | - | - | - | - | - | - | - | - | - | - | W | Q | S | E | E | C | I | L | - | - | - | - | - | - | - | - | - | - | - | - | - | - | - | - | - | - | - | - | - | - | - | - | - | - | - | - | - | - | - | - | - | - | - | - | - | - | - | - | - | - | - | - | - | - | - | - | - | - | L | K | G | H | N | - | - | - | - | - | - | - | - | - | - | - | - | - | - | - | - | - | - | - | - | - | - | - | - | - | - | E | R | V | K | K | F | L | L | L | - | - | - | - | - | - | - | - | - | - | - | - | - | - | - | - | - | - | - | - | - | - | - | K | D | - | S | R | L | L | - | S | W | - | - | - | S | F | D | G | T | V | - | - | - | - | - | - | - | - | - | - | - | - | - | - | - | - | - | - | - | - | - | - | - | - | - | - | - | - | - | - | - | - | - | - | - | K | L | W | N | I | V | T | G | E | I | - | - | - | - | - | E | K | D | V | I | C | - | - | - | - | - | - | - | H | E | - | - | - | - | - | - | - | - | - | - | - | - | - | - | - | - | - | - | - | N | A | I | L | - | - | - | - | - | - | - | - | - | - | - | - | - | - | - | S | C | A | - | - | - | - | - | - | - | - | - | - | - | - | - | - | - | I | S | P | D | A | T | - | M | F | S | T | T | - | S | S | D | K | T | - | - | - | - | - | - | A | K | I | W | S | F | E | T | S | - | - | - | - | - | - | - | - | - | - | - | - | - | - | - | - | - | - | - | - | - | - | - | - | S | M | L | H | K | - | - | - | - | - | - | - | - | L | - | - | - | - | N | S | H | E | S | C | V | - | - | - | - | - | - | - | - | - | R | C | C | - | A | F | S | - | - | - | - | - | - | - | - | - | L | N | N | E | - | L | L | A | T | - | - | - | - | - | - | - | - | G | D | D | N | R | E | I | R | V | W | D | V | S | R | G | E | L | L | - | - | - | - | - | - | - | - | - | - | - | - | - | - | - | - | - | - | - | - | - | - | - | - | - | - | - | - | - | - | - | - | - | - | - | - | - | - | - | - | - | - | - | - | - | - | - | - | - | - | - | - | - | - | - | - | H | L | C | S | R | A | S | I | D | - | - | - | - | - | - | - | - | - | - | - | - | - | - | - | - | - | - | - | - | E | G | E | S | A | H | E | G | W | - | - | - | - | - | - | - | - | V | T | D | L | C | F | S | P | D | N | K | - | - | - | - | - | - | - | - | - | - | - | - | - | - | - | - | - | - | - | - | - | - | - | - | - | - | - | - | - | - | - | - | - | - | - | - | - | - | - | - | - | - | - | - | - | I | L | V | S | - | - | S | A | G | Y | I | K | - | - | - | - | - | - | - | - | W | W | N | V | D | T | - | - | - | - | - | - | - | - | - | - | - | - | - | - | - | - | - | - | - | - | - | - | - | - | - | - | - | - | - | - | - | - | - | - | - | - | - | - | - | - | - | - | - | - | - | - | - | - | - | - | - | - | - | - | - | - | - | - | - | - | - | - | - | - | - | - | - | - | - | - | - | - | - | - | - | - | - | - | - | - | - | - | - | - | - | - | - | - | - | - | - | - | - | - | - | - | - | - | - | - | - | - | - | - | - | - | - | - | - | - | - | - | - | - | - | - | - | - | - | - | - | - | - | - | - | - | - | - | - | - | - | - | - | - | - | - | - | - | - | - | - | - | - | - | - | - | - | - | - | - | - | - | - | - | - | - | - | - | - | - | - | - | - | - | - | - | - | - | - | - | - | - | - | - | - | - | - | - | - | - | - | - | - | - | - | - | - | - | - | - | - | - | - | - | - | - | - | - | - | - | - | - | - | - | - | - | - | - | - | - | - | - | - | - | - | - | - | - | - | - | - | - | - | - | - | - | - | - | - | - | - | - | - | - | - | - | - | - | - | - | - | - | - | - | - | - | - | - | - | - | - | - | - | - | - | - | - | - | - | - | - | - | - | - | - | - | - | G | Q | P | L | Q | T | F | Y | T | - | - | - | - | - | N | G | - | - | - | T | D | L | K | S | L | - | - | - | H | V | S | P | - | - | - | - | D | F | K | V | - | - | - | - | - | - | - | - | - | - | - | - | - | - | - | - | - | - | - | - | - | - | - | - | - | - | C | V | T | V | D | - | N | L | G | I | L | - | - | Y | V | L | R | I | - | - | - | - | - | - | - | - | - | - | - | - | - | - | - | - | - | - | - | - | - | - | - | - | - | - | - | - | - | - | - | - | - | - | M | E | - | - | - | - | - | - | - | - | - | - | - | - | - | - | - | - | - | - | - | - | - | - | - | - | - | - | - | - | - | - | - | - | - | - | - | - | - | - | - | - | - | - | - | - | - | - | - | - | - | - | - | - | M | - | - | - | - | - | - | - | - | - | - | - | - | - | - | - | - | - | - | - | - | - | - | - | - | - | - | - | - | - | - | - | - | - | - | - | - | - | - | - | - | - | - | - | - | - | - | - | - | - | - | - | - | - | - | E | E | S | H | R | A | - | - | - | - | - | - | - | - | - | - | - | - | - | - | - | - | - | - | - | - | L | L | R | R | - | - | - | - | - | - | - | - | - | - | - | - | - | - | - | - | - | - | - | - | - | - | - | - | - | - | - | - | - | - | N | R | - | - | - | - | - | - | - | - | V | R | L | - | V | K | E | L | R | V | D | P | L | W | - | - | - | - | - | - | - | - | - | - | - | - | - | - | - | - | - | - | - | - | - | - | - | - | - | - | - | - | - | - | - | - | - | - | - | - | - | - | - | - | - | - | - | - | - | D | L | L | L | Q | E | G | I | - | - | - | - | - | - | - | - | - | - | - | - | - | - | - | - | - | - | - | - | - | - | - | - | F | S | Q | D | M | I | E | E | - | - | - | - | - | - | - | - | - | - | - | - | - | - | - | - | - | - | - | - | - | - | - | - | - | I | Q | G | S | - | - | - | - | - | - | - | - | - | - | G | A | R | S | A | Q | - | A | R | Q | L | C | I | D | L | E | T | R | G | - | K | R | A | L | P | V | F | V | H | C | L | Q | E | T | G | Q | G | A | L | G | D | L | L | G | R | G | A | N | Q | V | - | - | - | - | P | E | S | L | - | - | - | - | - | - | - | - | - | - | - | - | - | - | - | - | - | - | - | - | - | - | - | - | - | - | - | - | - | - | - | - | S | D | H | - | - | - | - | - | - | - | - | - | - | - | - | - | - | I | R | P | V | - | - | - | - | - | - | - | - | - | - | - | - | - | - | - | - | - | - | - | - | - | - | - | - | - | - | - | - | - | - | - | - | - | - | - | - | - | - | - | - | - | - | - | - | - | - | - | - | - | - | - | - | - | - | - | - | - | - | - | - | - | - | - | - | - | - | - | - | - | - | - | - | - | - | - | - | - | - | - | - | - | - | - | - | - | - | - | - | - | - | - | - | - | - | - | - | - | - | - | - | - | - | - | - | - | - | - | - | - | - | - | - | - | - | - | - | - | - | - | - | - | - | - | - | - | - | - | - | - | - | - | - | - | - | - | - | - | - | - | - | - | - | - | - | - | - | - | - | - | - | - | - | - | - | - | - | - | - | - | - | - | - | - | - | - | - | - | - | - | - | - | - | - | - | - | - | - | - | - | - | - | - | - | - | - | - | - | - | - | - | - | - | - | - | - | - | - | - | - | - | - | - | - | - | - | - | - | - | - | - | - | - | - | - | - | - | - | - | - | - | - | - | - | - | - | P | - | - | - | - | - | - | - | - | - | - | - | - | - | - | - | - | - | - | - | - | - | - | - | - | - | - | I | R | T | D | T | - | - | - | - | - | - | V | A | E | R | P | - | Q | E | P | R | - | - | - | - | - | - | - | - | - | - | - | - | - | - | - | P | E | I | T | P | I | H | G | - | - | - | - | - | - | - | - | - | - | - | - | - | - | - | - | - | - | - | - | S | - | - | - | - | - | - | - | - | - | - | G | - | G | E | K | S | R | R | N | S | E | - | - | - | - | - | T | A | Y | V | - | - | L | T | H | D | - | P | C | G | Y | C | L | I | I | N | N | V | N | F | S | E | A | S | - | - | Q | L | G | R | R | S | G | S | D | L | D | C | E | R | M | E | K | R | F | - | K | S | L | R | F | E | V | L | T | R | R | D | L | P | A | Q | A | I | A | E | E | L | Q | S | L | A | R | R | D | H | - | S | S | M | D | C | C | L | V | V | I | L | S | H | G | C | Q | T | S | H | V | Q | F | P | G | G | - | I | F | G | T | D | G | R | R | L | A | V | E | K | M | V | S | L | F | D | G | S | H | C | P | S | L | R | G | K | P | K | I | F | F | I | Q | A | C | - | - | G | G | E | E | K | D | R | G | F | Q | V | D | L | D | - | - | - | P | P | F | Y | - | - | - | N | P | A | S | - | - | - | - | - | - | - | I | E | S | D | A | I | A | F | Q | T | Q | P | G | N | T | - | - | - | - | - | - | - | - | D | E | P | D | A | I | - | - | - | - | - | - | - | A | S | L | - | - | - | P | T | P | S | D | I | L | V | S | Y | S | T | F | P | - | - | G | - | - | - | - | - | - | - | - | F | V | S | W | R | D | K | L | T | G | S | W | Y | I | E | S | L | D | Q | V | L | E | Q | F | V | R | - | S | - | - | - | - | - | - | - | - | - | - | - | - | - | - | - | - | - | - | - | - | - | - | - | - | - | - | - | - | - | - | - | E | D | L | L | D | M | L | - | - | - | - | - | L | R | - | - | - | - | - | - | - | - | - | - | - | - | V | A | N | A | V | - | - | - | - | - | S | A | K | - | - | - | - | - | - | - | G | T | F | - | K | Q | M | - | - | P | G | - | - | - | - | - | - | - | - | - | - | - | - | - | - | - | C | F | N | F | - | - | - | - | - | - | - | - | - | L | R | K | R | F | F | F | M | V | R | - | - | - | - | - | - | - | - | - | - | - | - | - | - | - | - | - | - | - | - | - | - | - | - | - | - | - | - | - | - | - | - | - | - | - | - | - | M | - | - | - | - | - | - | - | - | - | - | - | - | - | - | - | - | - | - | - | - | - | - | - | - | - | - | - | - | - | - | - | - | - | - | - | - | - | - | - | - | - | - | - | - | - | - | - | - | - | - | - | - | G | D | I | E | K | G | K | K | I | F | V | Q | K | C | S | Q | C | H | T | V | E | K | G | G | K | H | K | T | G | P | N | L | H | G | L | F | G | R | K | T | G | Q | A | D | G | F | S | Y | T | E | A | N | K | S | K | G | I | I | W | G | E | D | T | L | M | E | Y | L | E | N | P | K | K | Y | I | P | G | T | K | M | I | F | A | G | I | K | K | K | G | E | R | A | D | L | I | A | Y | L | K | D | S | T | S | K | - | - | - | - | - | - |
| Apteryx\_australis\_mantelli | M | - | - | - | - | - | - | - | - | - | - | - | - | - | - | - | - | - | - | - | - | - | - | - | - | - | - | - | - | - | - | - | - | - | - | - | - | - | - | - | - | - | - | - | - | - | - | - | - | - | - | - | - | - | - | - | - | - | - | - | - | - | - | - | - | - | - | - | - | - | - | - | - | - | - | - | - | - | - | - | - | - | - | - | - | - | - | - | - | - | - | - | - | - | - | - | - | - | D | V | K | S | - | R | N | - | - | - | - | - | - | - | - | - | - | - | - | - | - | - | - | Y | L | L | - | - | - | - | - | - | - | - | - | - | - | - | - | - | - | - | - | - | - | - | - | - | - | - | - | - | - | - | - | - | - | - | - | - | - | - | - | - | - | - | - | - | - | - | - | - | - | - | - | - | - | - | - | - | - | - | - | - | - | - | - | - | - | - | - | - | - | - | - | - | - | - | - | - | - | - | - | - | - | - | - | - | - | - | - | - | - | - | - | - | - | - | - | - | - | - | - | - | - | - | - | - | - | - | - | - | - | - | - | - | - | - | - | - | - | - | - | - | - | - | - | L | N | R | Q | A | L | E | K | D | I | K | T | S | Y | - | I | M | D | H | M | I | A | D | Q | V | L | T | L | Q | E | E | E | K | V | R | Q | Q | - | - | - | - | - | - | - | - | - | - | - | - | - | - | - | - | - | N | T | Q | K | E | R | A | A | M | L | I | N | I | L | L | K | K | - | - | - | - | D | N | N | S | Y | R | S | - | - | - | - | - | F | Y | N | A | L | - | L | H | E | G | - | - | - | - | - | - | Y | R | D | L | A | A | L | L | Q | D | G | I | P | T | I | S | S | G | S | V | - | K | - | - | - | - | - | - | - | - | - | - | - | - | - | - | - | - | - | - | - | - | - | - | S | S | L | D | G | M | T | - | - | - | - | - | - | - | - | - | - | - | S | Y | V | K | T | I | L | C | E | G | G | - | - | - | - | - | - | - | V | P | Q | R | P | V | V | F | V | T | R | P | K | L | V | D | A | I | - | - | - | - | - | - | - | - | - | - | K | E | K | L | Y | - | - | - | - | - | - | - | - | - | - | - | - | - | - | - | - | - | - | - | - | - | - | - | - | - | - | - | - | - | - | - | - | C | L | Q | N | D | P | - | - | - | - | - | - | - | - | - | - | - | - | - | - | - | - | - | - | - | - | - | - | - | - | - | - | - | - | - | - | - | - | - | - | - | - | - | - | G | W | V | T | V | Y | G | M | A | G | C | G | K | T | V | L | T | A | E | A | L | - | R | D | H | Q | L | L | E | G | Y | F | P | G | G | V | H | W | I | S | V | G | - | - | - | - | - | - | - | K | - | - | - | - | - | - | Q | D | K | A | - | - | - | - | - | - | - | - | - | - | - | - | G | L | L | I | K | L | Q | N | L | C | S | R | L | D | - | - | - | - | H | - | - | - | - | - | - | - | - | - | - | - | - | - | - | - | D | C | T | L | S | Q | R | P | P | L | N | I | - | - | - | - | - | - | - | - | - | - | - | - | - | - | - | - | E | E | A | K | D | R | L | R | L | L | M | L | R | - | - | - | - | - | - | - | - | - | - | - | - | K | Y | P | R | S | L | L | - | - | V | L | D | D | I | W | D | S | W | V | L | K | A | F | D | - | N | Q | C | Q | V | L | I | T | S | R | D | R | S | V | T | D | A | V | - | A | G | K | - | - | K | Y | E | I | H | V | E | S | - | - | - | G | L | A | H | E | K | G | L | E | V | L | S | L | F | V | N | K | - | K | I | T | E | L | P | E | - | - | - | - | - | - | - | - | - | - | - | - | - | - | Q | A | N | C | I | V | R | E | C | K | - | - | - | - | - | - | - | - | - | - | - | - | - | - | - | - | - | - | - | G | - | S | P | L | V | I | S | L | I | G | A | L | L | R | D | F | P | G | - | - | - | - | - | - | - | - | - | - | R | W | E | Y | Y | L | - | K | Q | L | Q | N | K | Q | F | K | R | I | - | - | - | - | - | - | - | - | - | - | - | - | - | - | - | - | - | - | - | - | - | - | R | K | S | S | S | Y | D | - | - | - | - | Y | E | A | L | D | E | A | M | S | I | S | V | E | - | Q | L | N | - | - | - | E | N | F | K | D | Y | - | Y | K | D | L | S | I | L | P | K | D | V | K | V | P | T | K | V | L | C | I | L | W | D | - | - | - | - | M | E | T | E | E | V | E | D | I | L | Q | E | F | - | - | - | - | - | - | - | - | - | - | - | V | N | K | S | L | L | F | C | D | R | N | G | K | S | - | - | - | - | - | - | - | - | - | F | H | Y | Y | L | H | D | L | Q | L | D | F | L | T | E | K | - | - | N | R | N | Q | - | L | - | Q | E | L | H | K | N | I | V | N | Q | Y | K | K | - | Y | Y | - | - | - | - | - | - | K | L | N | M | P | I | P | S | - | Q | E | - | - | - | - | - | - | - | D | C | M | Y | W | Y | N | F | L | A | Y | H | M | A | G | A | N | - | - | - | - | M | Q | K | E | L | C | D | L | M | F | S | L | D | W | I | K | A | K | T | E | L | V | G | P | A | - | - | - | - | - | - | - | - | H | L | I | H | E | Y | V | E | Y | S | S | V | L | D | Q | K | - | - | - | - | - | - | - | G | S | I | V | R | E | N | F | Q | - | - | - | E | F | L | S | L | N | G | H | L | L | G | R | Q | P | F | - | - | - | - | - | - | - | - | - | - | - | - | - | - | - | - | - | - | - | - | - | - | - | - | - | - | - | - | - | - | - | - | - | - | - | - | - | - | - | - | - | - | - | - | - | - | - | - | - | - | - | - | - | - | - | - | - | - | - | - | - | - | - | - | - | - | - | - | - | - | - | P | D | I | I | Q | L | G | L | C | Q | P | E | T | S | E | V | Y | Q | Q | A | K | L | L | A | Q | Q | E | - | - | A | G | - | - | - | - | - | - | - | - | - | - | - | - | - | - | - | - | - | - | - | - | - | - | - | - | - | - | - | - | - | V | F | Y | L | - | - | - | - | - | - | - | - | - | - | E | W | I | N | K | - | - | - | - | - | - | - | K | S | M | K | N | L | S | R | - | - | - | - | - | - | - | - | - | - | L | V | V | R | P | H | R | D | - | - | - | - | - | - | - | - | - | - | - | - | - | - | - | A | V | Y | H | A | C | - | - | - | F | S | - | Q | D | R | Q | R | I | A | - | - | - | - | - | - | - | - | - | S | C | G | A | D | K | T | L | Q | V | F | K | - | - | - | - | - | A | E | S | G | E | R | L | L | E | I | - | S | A | H | D | D | E | - | - | - | I | L | C | C | S | F | S | - | A | - | - | - | - | - | - | - | - | - | - | - | - | - | D | G | T | F | V | A | T | C | S | A | D | K | K | V | K | I | - | - | - | W | N | S | R | - | - | - | - | - | - | - | - | - | - | - | - | - | - | - | - | - | - | - | - | - | - | - | - | - | - | - | - | - | - | - | - | - | - | - | - | - | - | - | - | - | - | - | - | - | - | - | - | - | - | - | - | - | - | - | - | - | - | - | - | - | - | - | - | - | - | - | - | - | - | - | - | - | - | - | - | - | - | - | - | - | - | - | - | - | - | - | - | - | - | - | - | - | - | - | - | - | - | - | - | - | - | - | - | - | - | - | - | - | - | - | - | - | - | - | - | - | - | - | - | - | - | - | - | - | - | - | - | - | - | - | - | - | - | - | - | - | - | - | - | T | G | Q | C | M | H | V | Y | - | E | E | H | S | - | - | - | - | - | E | Q | V | N | - | - | - | - | - | - | - | - | - | - | - | - | - | C | C | Q | - | - | - | - | - | - | - | - | - | - | - | - | - | F | N | N | R | S | - | - | - | - | - | - | - | - | - | - | - | - | - | - | - | - | G | Q | - | Y | L | L | A | T | C | S | N | - | - | - | - | D | T | F | I | - | - | - | - | - | - | - | - | - | - | - | - | K | L | W | D | L | - | - | - | N | R | N | Y | C | R | N | T | M | F | - | - | - | - | - | - | - | - | - | - | - | - | - | - | - | - | - | G | H | V | N | S | - | - | - | - | - | - | - | - | - | - | - | - | - | - | - | - | V | N | H | C | K | F | S | P | N | D | E | - | - | - | - | - | - | - | - | - | - | - | - | - | - | - | - | - | - | - | - | - | - | - | - | - | - | - | - | Y | V | A | S | C | S | T | D | G | T | V | K | L | - | - | - | - | - | - | - | - | - | - | - | - | - | - | - | - | - | - | - | W | E | - | V | L | S | A | N | E | L | K | T | - | - | - | - | - | - | I | E | - | - | - | - | - | - | - | - | - | - | - | - | - | - | - | - | - | - | - | - | - | - | I | K | D | F | - | - | - | - | - | - | - | - | F | K | T | A | D | E | Q | P | D | - | - | - | - | - | - | - | - | - | - | - | - | - | - | - | - | - | - | - | - | - | - | - | - | - | - | - | D | V | E | - | I | L | V | K | C | C | - | - | - | - | - | - | - | - | - | - | - | - | - | - | - | - | S | W | S | G | N | S | A | T | V | L | V | V | A | K | N | K | L | - | - | - | - | - | - | - | - | - | - | - | L | L | F | D | V | K | T | - | - | - | S | D | - | - | - | - | - | - | - | - | - | - | - | - | - | - | - | - | - | - | - | - | - | - | - | - | - | - | - | - | - | - | - | - | - | - | - | - | - | - | - | - | - | - | - | - | - | - | - | - | - | - | - | - | - | - | - | - | - | - | - | - | - | - | - | - | - | - | - | - | - | - | - | - | - | - | - | - | - | - | - | - | - | - | - | - | - | - | - | - | - | - | - | - | - | - | - | - | - | - | - | - | - | - | - | - | - | - | - | - | - | - | - | - | - | - | - | - | - | - | - | - | - | - | - | - | - | - | L | L | K | - | - | - | - | - | - | - | - | - | - | - | - | - | - | - | - | - | - | - | - | - | - | - | - | - | - | - | - | - | - | - | - | - | - | - | - | - | - | - | - | - | - | - | - | - | - | - | - | - | - | - | - | - | - | - | - | - | - | - | - | - | - | - | - | - | - | - | - | - | - | - | - | - | - | - | - | - | - | - | - | - | - | - | - | - | - | - | - | - | - | - | - | - | - | - | - | - | - | - | - | - | - | - | - | - | - | - | - | - | - | - | - | - | - | - | - | - | - | - | - | - | - | - | - | - | - | - | - | - | - | - | - | - | - | - | - | - | - | - | - | - | - | - | - | - | - | - | - | - | - | - | - | - | - | - | - | - | - | - | - | - | - | - | - | - | - | - | - | - | - | - | - | - | - | - | - | - | - | - | - | - | - | - | Q | I | - | - | - | I | V | S | H | H | S | T | I | Q | Y | C | - | - | - | - | - | - | - | - | - | - | - | - | - | - | - | - | - | - | - | - | - | - | - | - | - | D | F | C | P | G | D | E | L | V | A | V | A | L | S | H | - | - | - | - | - | - | C | S | V | E | - | - | - | - | - | - | - | - | - | - | - | - | - | - | - | - | - | L | W | N | I | K | S | L | - | - | - | - | - | - | - | - | - | - | - | - | - | - | - | - | - | - | - | - | - | - | - | - | - | - | - | - | - | S | K | V | A | D | C | R | G | H | M | S | - | - | - | - | - | - | - | - | - | - | - | - | - | - | - | - | - | - | - | - | - | - | - | - | - | - | - | - | - | - | - | - | - | - | - | - | - | - | - | - | - | - | - | - | W | V | H | C | V | T | F | S | - | - | - | - | - | - | - | - | - | - | - | P | D | G | S | - | - | - | - | - | - | - | - | - | - | L | F | M | T | S | S | D | D | Q | - | - | - | - | - | - | - | - | - | - | - | - | - | - | T | I | R | I | - | W | - | - | - | - | - | - | - | - | - | - | - | - | - | - | - | - | - | - | - | - | - | - | - | - | - | - | - | - | - | - | - | - | - | - | - | - | - | - | E | T | N | K | V | C | K | S | S | D | A | V | L | K | S | E | - | - | - | - | - | - | - | - | - | - | - | - | - | - | - | - | L | D | V | V | F | - | - | - | - | - | - | - | - | - | - | - | - | - | - | - | - | - | - | - | - | - | - | - | - | - | - | - | - | - | - | - | - | - | - | - | - | - | - | - | - | - | - | - | - | - | - | - | - | - | - | - | - | - | - | H | N | G | E | V | M | - | - | - | - | - | - | - | - | - | - | - | - | - | - | - | - | - | - | - | - | - | - | - | - | - | - | - | - | - | - | - | - | - | - | - | - | I | L | A | I | Y | N | Q | Q | H | L | Q | L | - | I | N | G | N | T | - | - | - | - | - | - | - | - | - | - | D | S | I | I | M | Q | T | E | A | - | - | - | - | - | - | - | - | - | - | - | - | - | - | - | - | - | - | - | - | - | - | Q | E | S | - | - | - | - | - | - | - | - | L | I | C | C | C | C | L | S | E | D | L | K | - | - | - | - | - | - | - | - | - | F | A | A | F | G | Q | - | - | - | - | - | - | - | - | - | - | - | - | - | - | - | - | - | - | - | - | - | - | - | - | - | - | - | - | - | - | - | - | - | - | - | - | E | - | - | - | - | - | - | - | - | - | N | G | T | - | - | - | - | - | - | - | - | - | - | I | - | - | - | - | - | - | - | - | - | - | - | - | - | - | - | - | - | - | - | - | - | - | - | - | - | - | - | - | - | - | - | - | - | - | - | - | - | - | - | K | I | L | Q | L | S | D | G | K | V | L | K | T | L | E | A | - | - | - | - | - | Y | - | - | - | - | - | - | - | - | - | - | - | - | R | T | S | V | Q | H | C | Q | F | T | - | - | - | - | - | - | - | - | - | - | - | - | - | - | - | - | - | - | - | - | - | - | - | - | - | - | - | - | - | - | - | - | - | - | - | - | - | S | D | C | Q | T | L | I | S | S | A | - | H | D | S | - | - | - | - | - | - | - | - | V | I | Q | - | V | - | - | - | - | - | - | - | - | - | - | - | - | - | - | - | - | - | - | - | - | - | - | - | - | - | - | - | - | - | - | - | - | - | - | - | - | - | - | - | W | S | - | - | - | - | - | - | - | - | - | - | W | Q | L | K | E | C | V | F | - | - | - | - | - | - | - | - | - | - | - | - | - | - | - | - | - | - | - | - | - | - | - | - | - | - | - | - | - | - | - | - | - | - | - | - | - | - | - | - | - | - | - | - | - | - | - | - | - | - | L | R | G | H | K | - | - | - | - | - | - | - | - | - | - | - | - | - | - | - | - | - | - | - | - | - | - | - | - | - | - | E | A | V | K | D | F | R | L | L | - | - | - | - | - | - | - | - | - | - | - | - | - | - | - | - | - | - | - | - | - | - | - | E | N | - | S | K | L | L | - | S | W | - | - | - | S | F | D | G | T | V | - | - | - | - | - | - | - | - | - | - | - | - | - | - | - | - | - | - | - | - | - | - | - | - | - | - | - | - | - | - | - | - | - | - | - | K | V | W | N | I | I | T | G | K | A | - | - | - | - | - | E | K | D | F | A | C | - | - | - | - | - | - | - | H | G | - | - | - | - | - | - | - | - | - | - | - | - | - | - | - | - | - | - | - | D | T | V | L | - | - | - | - | - | - | - | - | - | - | - | - | - | - | - | S | C | A | - | - | - | - | - | - | - | - | - | - | - | - | - | - | - | V | S | P | D | G | S | - | K | F | S | S | A | - | S | A | D | K | T | - | - | - | - | - | - | A | K | I | W | S | F | E | S | S | - | - | - | - | - | - | - | - | - | - | - | - | - | - | - | - | - | - | - | - | - | - | - | - | S | V | L | H | E | - | - | - | - | - | - | - | - | L | - | - | - | - | N | G | H | E | A | C | V | - | - | - | - | - | - | - | - | - | R | C | C | - | T | F | S | - | - | - | - | - | - | - | - | - | S | N | N | K | - | L | L | A | T | - | - | - | - | - | - | - | - | G | D | D | K | G | E | I | R | I | W | Y | T | L | T | G | E | L | L | - | - | - | - | - | - | - | - | - | - | - | - | - | - | - | - | - | - | - | - | - | - | - | - | - | - | - | - | - | - | - | - | - | - | - | - | - | - | - | - | - | - | - | - | - | - | - | - | - | - | - | - | - | - | - | - | H | F | C | S | P | V | T | V | D | - | - | - | - | - | - | - | - | - | - | - | - | - | - | - | - | - | - | - | - | E | G | E | P | T | H | G | G | W | - | - | - | - | - | - | - | - | V | T | D | L | S | F | S | P | D | S | K | - | - | - | - | - | - | - | - | - | - | - | - | - | - | - | - | - | - | - | - | - | - | - | - | - | - | - | - | - | - | - | - | - | - | - | - | - | - | - | - | - | - | - | - | - | V | L | V | S | - | - | S | G | G | Y | L | K | - | - | - | - | - | - | - | - | W | W | N | V | T | T | - | - | - | - | - | - | - | - | - | - | - | - | - | - | - | - | - | - | - | - | - | - | - | - | - | - | - | - | - | - | - | - | - | - | - | - | - | - | - | - | - | - | - | - | - | - | - | - | - | - | - | - | - | - | - | - | - | - | - | - | - | - | - | - | - | - | - | - | - | - | - | - | - | - | - | - | - | - | - | - | - | - | - | - | - | - | - | - | - | - | - | - | - | - | - | - | - | - | - | - | - | - | - | - | - | - | - | - | - | - | - | - | - | - | - | - | - | - | - | - | - | - | - | - | - | - | - | - | - | - | - | - | - | - | - | - | - | - | - | - | - | - | - | - | - | - | - | - | - | - | - | - | - | - | - | - | - | - | - | - | - | - | - | - | - | - | - | - | - | - | - | - | - | - | - | - | - | - | - | - | - | - | - | - | - | - | - | - | - | - | - | - | - | - | - | - | - | - | - | - | - | - | - | - | - | - | - | - | - | - | - | - | - | - | - | - | - | - | - | - | - | - | - | - | - | - | - | - | - | - | - | - | - | - | - | - | - | - | - | - | - | - | - | - | - | - | - | - | - | - | - | - | - | - | - | - | - | - | - | - | - | - | - | - | - | - | - | G | E | S | L | Q | T | F | Y | T | - | - | - | - | - | N | G | - | - | - | T | N | L | K | S | I | - | - | - | H | V | S | P | - | - | - | - | N | F | K | V | - | - | - | - | - | - | - | - | - | - | - | - | - | - | - | - | - | - | - | - | - | - | - | - | - | - | Y | V | T | V | D | - | N | L | G | I | L | - | - | Y | V | L | Q | K | - | - | - | - | - | - | - | - | - | - | - | - | - | - | - | - | - | - | - | - | - | - | - | - | - | - | - | - | - | - | - | - | - | - | M | - | - | - | - | - | - | - | - | - | - | - | - | - | - | - | - | - | - | - | - | - | - | - | - | - | - | - | - | - | - | - | - | - | - | - | - | - | - | - | - | - | - | - | - | - | - | - | - | - | - | - | - | - | M | - | - | - | - | - | - | - | - | - | - | - | - | - | - | - | - | - | - | - | - | - | - | - | - | - | - | - | - | - | - | - | - | - | - | - | - | - | - | - | - | - | - | - | - | - | - | - | - | - | - | - | - | - | - | Q | T | - | - | - | - | - | - | - | - | - | - | - | - | - | - | - | - | - | - | - | - | - | - | - | - | - | - | - | - | - | - | - | - | - | - | - | - | - | - | - | - | - | - | - | - | - | - | - | - | - | - | - | - | - | - | - | - | - | - | - | - | - | - | - | - | - | - | - | - | - | - | - | - | - | - | - | - | - | - | - | - | - | - | - | - | - | - | - | - | - | - | - | - | - | - | - | - | - | - | - | - | - | - | - | - | - | - | - | - | - | - | - | - | - | - | - | - | - | - | - | - | - | - | - | - | - | - | - | - | - | - | - | - | - | - | - | - | - | - | - | - | - | - | - | - | - | - | - | - | - | - | - | - | - | - | - | - | - | - | - | - | - | - | - | - | - | - | - | - | - | - | - | - | - | - | - | - | - | - | - | - | - | - | - | - | - | - | - | - | - | - | - | - | - | - | S | A | - | - | - | - | - | - | - | - | - | - | G | T | P | R | D | Q | - | A | R | Q | L | V | I | D | L | E | T | R | G | - | K | Q | A | F | P | I | F | L | S | I | L | R | D | T | G | Q | G | D | L | A | E | L | L | S | E | A | C | E | G | S | - | - | - | - | P | S | Q | P | - | - | - | - | - | - | - | - | - | - | - | - | - | - | - | - | - | - | - | - | - | - | - | - | - | - | - | - | - | - | - | - | V | Q | - | - | - | - | - | - | - | - | - | - | - | - | - | - | - | V | K | P | L | - | - | - | - | - | - | - | - | - | - | - | - | - | - | - | - | - | - | - | - | - | - | - | - | - | - | - | - | - | - | - | - | - | - | - | - | - | - | - | E | L | H | L | G | G | E | T | R | D | K | S | A | - | - | - | - | - | - | - | - | - | - | - | S | - | - | - | - | - | - | - | - | - | - | - | - | - | - | - | - | - | - | - | - | - | - | - | - | - | - | - | - | - | - | - | - | - | - | - | - | - | - | - | - | - | - | - | - | - | - | - | - | - | - | - | - | - | - | - | - | - | - | - | - | - | - | - | - | - | - | - | - | - | - | - | - | - | - | - | - | - | - | - | - | - | - | - | - | - | - | - | - | - | - | - | - | - | - | - | - | - | - | - | - | - | - | - | - | - | - | - | - | - | - | - | - | - | - | - | - | - | - | - | - | - | - | - | - | - | - | - | - | - | - | - | - | - | - | - | - | - | - | - | - | - | - | - | - | - | - | - | - | - | - | - | - | - | - | - | - | A | L | E | R | V | P | - | - | - | - | - | - | - | - | - | - | - | - | - | - | - | - | - | - | - | - | - | - | - | - | - | - | V | P | V | Q | D | - | - | - | - | - | - | M | S | D | A | F | - | R | T | - | - | - | - | - | - | - | - | - | - | - | - | - | - | - | - | - | - | P | P | A | A | P | R | G | - | - | - | - | - | - | - | - | - | - | - | - | - | - | - | - | - | - | - | - | S | - | - | - | - | - | - | - | - | - | - | A | - | V | D | S | L | K | R | N | S | D | - | - | - | - | - | L | V | Y | D | - | - | L | K | A | D | - | P | C | G | Y | C | L | I | L | N | N | V | N | F | S | R | D | S | - | - | D | L | S | T | R | A | G | S | N | I | D | C | E | K | L | E | K | R | F | - | K | S | L | H | F | S | V | L | I | R | Q | D | L | K | A | R | E | I | V | R | E | L | Q | E | L | A | K | L | D | H | - | S | A | L | D | C | C | L | V | V | I | L | S | H | G | C | Q | T | S | H | I | Q | F | P | G | G | - | I | Y | G | T | D | G | K | A | I | P | V | E | K | I | V | N | Y | F | N | G | S | H | C | P | S | L | R | G | K | P | K | L | F | F | I | Q | A | C | - | - | G | G | E | Q | K | D | Q | G | F | E | V | E | S | D | - | - | - | S | P | G | D | - | E | A | R | R | V | S | - | - | - | - | - | - | - | V | E | S | D | A | T | P | F | Q | A | P | S | G | A | L | - | - | - | - | - | - | - | - | D | E | P | D | A | V | - | - | - | - | - | - | - | A | S | L | - | - | - | P | T | P | S | D | I | L | V | S | Y | S | T | F | P | - | - | G | - | - | - | - | - | - | - | - | E | V | S | C | A | N | R | L | P | G | K | E | - | - | - | - | - | - | - | - | - | - | - | - | - | - | - | - | - | - | - | - | - | - | - | - | - | - | - | - | - | - | - | - | - | - | - | - | - | - | - | - | - | - | - | - | - | - | - | - | - | - | - | - | - | - | - | - | - | - | - | - | - | - | - | - | - | - | - | - | - | - | - | - | - | - | - | - | - | - | - | - | - | - | - | - | - | - | - | - | - | - | - | - | - | - | - | - | - | - | - | - | - | - | - | - | - | - | - | - | - | - | - | - | - | - | - | - | - | - | - | - | - | - | - | - | - | - | - | - | - | - | - | - | - | - | - | - | - | - | - | - | - | - | - | - | - | - | - | - | - | - | - | - | - | - | - | - | - | - | - | - | - | - | - | - | - | - | - | - | - | - | - | - | - | - | - | - | - | - | - | M | - | - | - | - | - | - | - | - | - | - | - | - | - | - | - | - | - | - | - | - | - | - | - | - | - | - | - | - | - | - | - | - | - | - | - | - | - | - | - | - | - | - | - | - | - | - | - | - | - | - | - | - | G | D | I | E | K | G | K | K | I | F | V | Q | K | C | S | Q | C | H | T | V | E | K | G | G | K | H | K | T | G | P | N | L | N | G | L | F | G | R | K | T | G | Q | A | E | G | F | S | Y | T | D | A | N | K | N | K | G | I | T | W | G | E | D | T | L | M | E | Y | L | E | N | P | K | K | Y | I | P | G | T | K | M | I | F | A | G | I | K | K | K | S | E | R | A | D | L | I | A | Y | L | K | D | A | T | S | K | - | - | - | - | - | - |
| Aquila\_chrysaetos\_canadensis | M | - | - | - | - | - | - | - | - | - | - | - | - | - | - | - | - | - | - | - | - | - | - | - | - | - | - | - | - | - | - | - | - | - | - | - | - | - | - | - | - | - | - | - | - | - | - | - | - | - | - | - | - | - | - | - | - | - | - | - | - | - | - | - | - | - | - | - | - | - | - | - | - | - | - | - | - | - | - | - | - | - | - | - | - | - | - | - | - | - | - | - | - | - | - | - | - | - | D | I | K | S | - | R | N | - | - | - | - | - | - | - | - | - | - | - | - | - | - | - | - | Y | L | L | - | - | - | - | - | - | - | - | - | - | - | - | - | - | - | - | - | - | - | - | - | - | - | - | - | - | - | - | - | - | - | - | - | - | - | - | - | - | - | - | - | - | - | - | - | - | - | - | - | - | - | - | - | - | - | - | - | - | - | - | - | - | - | - | - | - | - | - | - | - | - | - | - | - | - | - | - | - | - | - | - | - | - | - | - | - | - | - | - | - | - | - | - | - | - | - | - | - | - | - | - | - | - | - | - | - | - | - | - | - | - | - | - | - | - | - | - | - | - | - | - | M | N | R | Q | A | L | E | K | D | I | K | T | S | Y | - | I | M | D | R | M | I | S | D | E | V | L | T | L | Q | E | E | E | R | V | K | Q | Q | - | - | - | - | - | - | - | - | - | - | - | - | - | - | - | - | - | N | T | Q | K | E | R | A | A | M | L | I | N | I | I | L | T | K | - | - | - | - | D | N | N | S | Y | R | S | - | - | - | - | - | F | Y | N | A | L | - | L | H | E | G | - | - | - | - | - | - | Y | R | D | L | A | A | L | L | Q | D | G | I | P | A | I | S | S | G | N | G | - | K | - | - | - | - | - | - | - | - | - | - | - | - | - | - | - | - | - | - | - | - | - | - | S | S | M | D | G | M | T | - | - | - | - | - | - | - | - | - | - | - | S | Y | V | K | T | V | L | C | E | G | G | - | - | - | - | - | - | - | V | P | Q | R | P | V | V | F | V | T | R | P | K | L | V | D | A | I | - | - | - | - | - | - | - | - | - | - | K | Q | K | L | C | - | - | - | - | - | - | - | - | - | - | - | - | - | - | - | - | - | - | - | - | - | - | - | - | - | - | - | - | - | - | - | - | C | L | G | S | D | P | - | - | - | - | - | - | - | - | - | - | - | - | - | - | - | - | - | - | - | - | - | - | - | - | - | - | - | - | - | - | - | - | - | - | - | - | - | - | G | W | V | T | V | Y | G | M | A | G | C | G | K | T | V | L | T | A | E | A | L | - | R | D | H | Q | L | L | E | D | Y | F | P | G | G | V | H | W | I | S | V | G | - | - | - | - | - | - | - | K | - | - | - | - | - | - | Q | D | K | A | - | - | - | - | - | - | - | - | - | - | - | - | G | L | L | I | K | L | Q | N | L | C | S | R | L | E | - | - | - | - | H | - | - | - | - | - | - | - | - | - | - | - | - | - | - | - | D | S | S | L | S | Q | R | P | P | L | N | I | - | - | - | - | - | - | - | - | - | - | - | - | - | - | - | - | E | E | A | K | D | R | L | R | L | L | M | L | R | - | - | - | - | - | - | - | - | - | - | - | - | K | Y | P | R | S | L | L | - | - | I | L | D | D | I | W | D | S | W | V | L | K | A | F | D | - | N | Q | C | Q | V | L | I | T | S | R | D | R | S | V | T | D | A | V | - | A | G | N | - | - | K | Y | E | V | H | V | E | S | - | - | - | G | L | A | H | E | K | G | L | E | I | L | S | L | F | V | N | M | - | K | I | S | E | L | P | E | - | - | - | - | - | - | - | - | - | - | - | - | - | - | Q | A | N | C | L | V | R | E | C | K | - | - | - | - | - | - | - | - | - | - | - | - | - | - | - | - | - | - | - | G | - | S | P | L | V | I | S | L | I | G | A | L | L | R | D | F | P | S | - | - | - | - | - | - | - | - | - | - | R | W | E | Y | Y | L | - | K | Q | L | Q | N | K | Q | F | K | R | I | - | - | - | - | - | - | - | - | - | - | - | - | - | - | - | - | - | - | - | - | - | - | R | K | S | S | S | Y | D | - | - | - | - | Y | E | A | L | D | E | A | M | S | I | S | V | E | - | Q | L | N | - | - | - | D | N | Y | K | D | Y | - | Y | K | D | L | S | V | L | P | K | D | V | K | V | P | T | K | V | L | C | I | L | W | D | - | - | - | - | M | E | T | E | E | V | E | D | I | L | Q | E | F | - | - | - | - | - | - | - | - | - | - | - | V | N | K | S | L | L | F | C | D | R | N | G | K | S | - | - | - | - | - | - | - | - | - | F | H | Y | Y | L | H | D | L | Q | L | D | F | L | T | E | K | - | - | N | R | N | Q | - | L | - | Q | E | L | H | K | N | I | V | N | Q | Y | K | K | - | Y | Y | - | - | - | - | - | - | K | L | N | T | P | V | P | S | - | Q | E | - | - | - | - | - | - | - | D | C | M | Y | W | Y | N | F | L | A | Y | H | M | A | G | A | N | - | - | - | - | M | Q | K | E | L | R | D | L | M | F | S | L | D | W | I | K | A | K | T | E | L | V | G | P | A | - | - | - | - | - | - | - | - | H | L | I | H | E | Y | V | E | Y | S | S | I | L | D | Q | K | - | - | - | - | - | - | - | D | S | T | V | R | E | N | F | Q | - | - | - | E | F | L | S | L | N | G | H | L | L | G | R | Q | P | F | - | - | - | - | - | - | - | - | - | - | - | - | - | - | - | - | - | - | - | - | - | - | - | - | - | - | - | - | - | - | - | - | - | - | - | - | - | - | - | - | - | - | - | - | - | - | - | - | - | - | - | - | - | - | - | - | - | - | - | - | - | - | - | - | - | - | - | - | - | - | - | P | D | I | I | Q | L | G | L | C | Q | P | E | T | S | E | V | Y | Q | Q | A | K | L | Y | A | Q | R | Q | - | - | T | G | - | - | - | - | - | - | - | - | - | - | - | - | - | - | - | - | - | - | - | - | - | - | - | - | - | - | - | - | - | T | F | Y | L | - | - | - | - | - | - | - | - | - | - | E | W | I | N | K | - | - | - | - | - | - | - | K | S | L | K | N | L | Y | R | - | - | - | - | - | - | - | - | - | - | L | V | V | R | P | H | R | D | - | - | - | - | - | - | - | - | - | - | - | - | - | - | - | A | V | Y | H | A | C | - | - | - | F | S | - | K | D | R | Q | R | I | A | - | - | - | - | - | - | - | - | - | S | C | G | A | D | K | T | L | Q | V | F | K | - | - | - | - | - | A | E | S | G | E | R | L | L | E | I | - | S | A | H | D | D | E | - | - | - | I | L | C | C | A | F | S | - | A | - | - | - | - | - | - | - | - | - | - | - | - | - | D | G | E | F | V | A | T | C | S | A | D | K | K | V | K | I | - | - | - | W | N | S | R | - | - | - | - | - | - | - | - | - | - | - | - | - | - | - | - | - | - | - | - | - | - | - | - | - | - | - | - | - | - | - | - | - | - | - | - | - | - | - | - | - | - | - | - | - | - | - | - | - | - | - | - | - | - | - | - | - | - | - | - | - | - | - | - | - | - | - | - | - | - | - | - | - | - | - | - | - | - | - | - | - | - | - | - | - | - | - | - | - | - | - | - | - | - | - | - | - | - | - | - | - | - | - | - | - | - | - | - | - | - | - | - | - | - | - | - | - | - | - | - | - | - | - | - | - | - | - | - | - | - | - | - | - | - | - | - | - | - | - | - | T | G | Q | R | K | C | V | Y | - | E | E | H | S | - | - | - | - | - | E | Q | V | N | - | - | - | - | - | - | - | - | - | - | - | - | - | C | C | Q | - | - | - | - | - | - | - | - | - | - | - | - | - | F | N | N | R | S | - | - | - | - | - | - | - | - | - | - | - | - | - | - | - | - | G | Q | - | Y | L | L | V | T | C | S | N | - | - | - | - | D | T | F | I | - | - | - | - | - | - | - | - | - | - | - | - | K | L | W | D | L | - | - | - | N | K | K | Y | C | R | N | T | M | M | - | - | - | - | - | - | - | - | - | - | - | - | - | - | - | - | - | G | H | E | N | S | - | - | - | - | - | - | - | - | - | - | - | - | - | - | - | - | V | N | H | C | R | F | S | P | N | D | E | - | - | - | - | - | - | - | - | - | - | - | - | - | - | - | - | - | - | - | - | - | - | - | - | - | - | - | - | Y | V | A | S | C | S | T | D | G | T | V | K | L | - | - | - | - | - | - | - | - | - | - | - | - | - | - | - | - | - | - | - | W | E | - | A | R | S | G | N | E | L | K | S | - | - | - | - | - | - | I | E | - | - | - | - | - | - | - | - | - | - | - | - | - | - | - | - | - | - | - | - | - | - | I | K | D | F | - | - | - | - | - | - | - | - | F | K | N | A | D | E | Q | P | D | - | - | - | - | - | - | - | - | - | - | - | - | - | - | - | - | - | - | - | - | - | - | - | - | - | - | - | D | V | E | - | V | L | V | K | C | C | - | - | - | - | - | - | - | - | - | - | - | - | - | - | - | - | S | W | S | R | N | G | D | M | I | L | V | V | A | K | N | K | L | - | - | - | - | - | - | - | - | - | - | - | L | L | F | D | V | K | T | - | - | - | C | G | - | - | - | - | - | - | - | - | - | - | - | - | - | - | - | - | - | - | - | - | - | - | - | - | - | - | - | - | - | - | - | - | - | - | - | - | - | - | - | - | - | - | - | - | - | - | - | - | - | - | - | - | - | - | - | - | - | - | - | - | - | - | - | - | - | - | - | - | - | - | - | - | - | - | - | - | - | - | - | - | - | - | - | - | - | - | - | - | - | - | - | - | - | - | - | - | - | - | - | - | - | - | - | - | - | - | - | - | - | - | - | - | - | - | - | - | - | - | - | - | - | - | - | - | - | - | L | L | S | - | - | - | - | - | - | - | - | - | - | - | - | - | - | - | - | - | - | - | - | - | - | - | - | - | - | - | - | - | - | - | - | - | - | - | - | - | - | - | - | - | - | - | - | - | - | - | - | - | - | - | - | - | - | - | - | - | - | - | - | - | - | - | - | - | - | - | - | - | - | - | - | - | - | - | - | - | - | - | - | - | - | - | - | - | - | - | - | - | - | - | - | - | - | - | - | - | - | - | - | - | - | - | - | - | - | - | - | - | - | - | - | - | - | - | - | - | - | - | - | - | - | - | - | - | - | - | - | - | - | - | - | - | - | - | - | - | - | - | - | - | - | - | - | - | - | - | - | - | - | - | - | - | - | - | - | - | - | - | - | - | - | - | - | - | - | - | - | - | - | - | - | - | - | - | - | - | - | - | - | - | - | - | Q | V | - | - | - | I | V | S | H | H | S | T | I | Q | Y | C | - | - | - | - | - | - | - | - | - | - | - | - | - | - | - | - | - | - | - | - | - | - | - | - | - | D | F | C | P | G | D | E | L | V | A | V | A | L | S | H | - | - | - | - | - | - | C | S | V | E | - | - | - | - | - | - | - | - | - | - | - | - | - | - | - | - | - | L | W | N | I | K | S | L | - | - | - | - | - | - | - | - | - | - | - | - | - | - | - | - | - | - | - | - | - | - | - | - | - | - | - | - | - | S | K | V | A | Y | C | R | G | H | M | S | - | - | - | - | - | - | - | - | - | - | - | - | - | - | - | - | - | - | - | - | - | - | - | - | - | - | - | - | - | - | - | - | - | - | - | - | - | - | - | - | - | - | - | - | W | V | H | C | V | T | F | S | - | - | - | - | - | - | - | - | - | - | - | P | D | G | S | - | - | - | - | - | - | - | - | - | - | L | F | L | T | S | S | D | D | Q | - | - | - | - | - | - | - | - | - | - | - | - | - | - | T | I | R | I | - | W | - | - | - | - | - | - | - | - | - | - | - | - | - | - | - | - | - | - | - | - | - | - | - | - | - | - | - | - | - | - | - | - | - | - | - | - | - | - | E | T | K | K | V | C | K | S | S | D | A | V | L | K | S | E | - | - | - | - | - | - | - | - | - | - | - | - | - | - | - | - | L | D | V | V | F | - | - | - | - | - | - | - | - | - | - | - | - | - | - | - | - | - | - | - | - | - | - | - | - | - | - | - | - | - | - | - | - | - | - | - | - | - | - | - | - | - | - | - | - | - | - | - | - | - | - | - | - | - | - | H | N | G | E | V | M | - | - | - | - | - | - | - | - | - | - | - | - | - | - | - | - | - | - | - | - | - | - | - | - | - | - | - | - | - | - | - | - | - | - | - | - | I | L | A | I | Y | N | Q | K | H | L | Q | L | - | I | N | G | N | T | - | - | - | - | - | - | - | - | - | - | D | S | V | I | M | Q | T | A | A | - | - | - | - | - | - | - | - | - | - | - | - | - | - | - | - | - | - | - | - | - | - | P | E | S | - | - | - | - | - | - | - | - | P | I | S | C | C | C | L | S | E | D | L | K | - | - | - | - | - | - | - | - | - | F | A | A | F | G | Q | - | - | - | - | - | - | - | - | - | - | - | - | - | - | - | - | - | - | - | - | - | - | - | - | - | - | - | - | - | - | - | - | - | - | - | - | E | - | - | - | - | - | - | - | - | - | S | G | T | - | - | - | - | - | - | - | - | - | - | I | - | - | - | - | - | - | - | - | - | - | - | - | - | - | - | - | - | - | - | - | - | - | - | - | - | - | - | - | - | - | - | - | - | - | - | - | - | - | - | K | V | L | Q | L | L | D | G | K | V | L | K | S | Q | E | A | - | - | - | - | - | Y | - | - | - | - | - | - | - | - | - | - | - | - | R | T | S | V | Q | H | C | R | F | T | - | - | - | - | - | - | - | - | - | - | - | - | - | - | - | - | - | - | - | - | - | - | - | - | - | - | - | - | - | - | - | - | - | - | - | - | - | S | D | C | Q | T | L | I | S | S | A | - | H | D | S | - | - | - | - | - | - | - | - | V | I | Q | - | V | - | - | - | - | - | - | - | - | - | - | - | - | - | - | - | - | - | - | - | - | - | - | - | - | - | - | - | - | - | - | - | - | - | - | - | - | - | - | - | W | N | - | - | - | - | - | - | - | - | - | - | W | Q | L | S | E | C | V | L | - | - | - | - | - | - | - | - | - | - | - | - | - | - | - | - | - | - | - | - | - | - | - | - | - | - | - | - | - | - | - | - | - | - | - | - | - | - | - | - | - | - | - | - | - | - | - | - | - | - | L | R | G | H | K | - | - | - | - | - | - | - | - | - | - | - | - | - | - | - | - | - | - | - | - | - | - | - | - | - | - | E | A | I | K | D | F | R | L | L | - | - | - | - | - | - | - | - | - | - | - | - | - | - | - | - | - | - | - | - | - | - | - | E | N | - | S | K | L | L | - | S | W | - | - | - | S | F | D | G | T | V | - | - | - | - | - | - | - | - | - | - | - | - | - | - | - | - | - | - | - | - | - | - | - | - | - | - | - | - | - | - | - | - | - | - | - | K | V | W | N | I | T | T | G | N | T | - | - | - | - | - | E | K | D | F | A | C | - | - | - | - | - | - | - | H | G | - | - | - | - | - | - | - | - | - | - | - | - | - | - | - | - | - | - | - | G | A | V | L | - | - | - | - | - | - | - | - | - | - | - | - | - | - | - | S | C | A | - | - | - | - | - | - | - | - | - | - | - | - | - | - | - | V | S | P | D | G | S | - | K | F | S | S | T | - | S | A | D | K | T | - | - | - | - | - | - | A | K | I | W | S | F | E | S | S | - | - | - | - | - | - | - | - | - | - | - | - | - | - | - | - | - | - | - | - | - | - | - | - | S | V | L | H | E | - | - | - | - | - | - | - | - | L | - | - | - | - | K | S | H | E | A | C | V | - | - | - | - | - | - | - | - | - | R | C | C | - | T | F | S | - | - | - | - | - | - | - | - | - | P | N | N | K | - | L | L | A | T | - | - | - | - | - | - | - | - | G | D | D | K | G | E | I | R | I | W | D | I | L | T | G | A | L | L | - | - | - | - | - | - | - | - | - | - | - | - | - | - | - | - | - | - | - | - | - | - | - | - | - | - | - | - | - | - | - | - | - | - | - | - | - | - | - | - | - | - | - | - | - | - | - | - | - | - | - | - | - | - | - | - | H | F | C | S | P | V | T | V | D | - | - | - | - | - | - | - | - | - | - | - | - | - | - | - | - | - | - | - | - | E | G | E | P | T | H | G | G | W | - | - | - | - | - | - | - | - | V | T | D | L | S | F | S | P | D | S | K | - | - | - | - | - | - | - | - | - | - | - | - | - | - | - | - | - | - | - | - | - | - | - | - | - | - | - | - | - | - | - | - | - | - | - | - | - | - | - | - | - | - | - | - | - | M | L | V | S | - | - | S | G | G | F | L | K | - | - | - | - | - | - | - | - | W | W | N | V | T | T | - | - | - | - | - | - | - | - | - | - | - | - | - | - | - | - | - | - | - | - | - | - | - | - | - | - | - | - | - | - | - | - | - | - | - | - | - | - | - | - | - | - | - | - | - | - | - | - | - | - | - | - | - | - | - | - | - | - | - | - | - | - | - | - | - | - | - | - | - | - | - | - | - | - | - | - | - | - | - | - | - | - | - | - | - | - | - | - | - | - | - | - | - | - | - | - | - | - | - | - | - | - | - | - | - | - | - | - | - | - | - | - | - | - | - | - | - | - | - | - | - | - | - | - | - | - | - | - | - | - | - | - | - | - | - | - | - | - | - | - | - | - | - | - | - | - | - | - | - | - | - | - | - | - | - | - | - | - | - | - | - | - | - | - | - | - | - | - | - | - | - | - | - | - | - | - | - | - | - | - | - | - | - | - | - | - | - | - | - | - | - | - | - | - | - | - | - | - | - | - | - | - | - | - | - | - | - | - | - | - | - | - | - | - | - | - | - | - | - | - | - | - | - | - | - | - | - | - | - | - | - | - | - | - | - | - | - | - | - | - | - | - | - | - | - | - | - | - | - | - | - | - | - | - | - | - | - | - | - | - | - | - | - | - | - | - | - | G | E | S | L | Q | T | F | Y | T | - | - | - | - | - | N | G | - | - | - | T | N | L | K | S | I | - | - | - | H | V | S | P | - | - | - | - | N | F | R | V | - | - | - | - | - | - | - | - | - | - | - | - | - | - | - | - | - | - | - | - | - | - | - | - | - | - | - | - | - | - | - | - | - | - | - | - | - | - | - | - | - | - | - | - | - | - | - | - | - | - | - | - | - | - | - | - | - | - | - | - | - | - | - | - | - | - | - | - | - | - | - | - | - | - | - | - | - | - | - | - | - | - | - | - | - | - | - | - | - | - | - | - | - | - | - | - | - | - | - | - | - | - | - | - | - | - | - | - | - | - | - | - | - | - | - | - | - | - | - | - | - | - | - | - | - | - | - | - | - | - | - | - | M | - | - | - | - | - | - | - | - | - | - | - | - | - | - | - | - | - | - | - | - | - | - | - | - | - | - | - | - | - | - | - | - | - | - | - | - | - | - | - | - | - | - | - | - | - | - | - | - | - | - | - | - | - | - | E | E | A | Q | R | R | - | - | - | - | - | - | - | - | - | - | - | - | - | - | - | - | - | - | - | - | A | L | R | R | - | - | - | - | - | - | - | - | - | - | - | - | - | - | - | - | - | - | - | - | - | - | - | - | - | - | - | - | - | - | G | R | - | - | - | - | - | - | - | - | A | R | L | - | V | A | A | L | R | V | A | P | L | W | - | - | - | - | - | - | - | - | - | - | - | - | - | - | - | - | - | - | - | - | - | - | - | - | - | - | - | - | - | - | - | - | - | - | - | - | - | - | - | - | - | - | - | - | - | D | P | L | E | D | R | G | L | - | - | - | - | - | - | - | - | - | - | - | - | - | - | - | - | - | - | - | - | - | - | - | - | F | T | R | P | M | V | E | E | - | - | - | - | - | - | - | - | - | - | - | - | - | - | - | - | - | - | - | - | - | - | - | - | - | L | Q | S | A | - | - | - | - | - | - | - | - | - | - | G | S | R | G | E | Q | - | A | R | Q | L | V | I | D | L | E | T | R | G | - | K | R | A | F | P | V | F | L | S | I | L | R | D | T | G | Q | G | D | L | A | D | M | L | I | E | E | C | E | C | P | - | - | - | - | P | A | P | P | - | Q | - | - | - | - | - | - | - | - | - | - | - | - | - | - | - | - | - | - | - | - | - | - | - | - | - | - | - | - | - | L | A | D | - | - | - | - | - | - | - | - | - | - | - | - | - | - | - | P | R | P | V | - | - | - | - | - | - | - | - | - | - | - | - | - | - | - | - | - | - | - | - | - | - | - | - | - | - | - | - | - | - | - | - | - | - | - | - | - | - | - | E | L | D | L | G | G | E | K | H | R | K | N | V | - | - | - | - | - | - | - | - | - | - | - | S | - | - | - | - | - | - | - | - | - | - | - | - | - | - | - | - | - | - | - | - | - | - | - | - | - | - | - | - | - | - | - | - | - | - | - | - | - | - | - | - | - | - | - | - | - | - | - | - | - | - | - | - | - | - | - | - | - | - | - | - | - | - | - | - | - | - | - | - | - | - | - | - | - | - | - | - | - | - | - | - | - | - | - | - | - | - | - | - | - | - | - | - | - | - | - | - | - | - | - | - | - | - | - | - | - | - | - | - | - | - | - | - | - | - | - | - | - | - | - | - | - | - | - | - | - | - | - | - | - | - | - | - | - | - | - | - | - | - | - | - | - | - | - | - | - | - | - | - | - | - | - | - | - | - | - | - | F | P | E | R | L | S | - | - | - | - | - | - | - | - | - | - | - | - | - | - | - | - | - | - | - | - | - | - | - | - | - | - | I | P | V | Q | A | - | - | - | - | - | - | E | S | E | R | P | - | Q | T | - | - | - | - | - | - | - | - | - | - | - | - | - | - | - | - | - | - | P | P | A | P | A | R | G | - | - | - | - | - | - | - | - | - | - | - | - | - | - | - | - | - | - | - | - | S | - | - | - | - | - | - | - | - | - | - | A | - | V | D | - | - | K | R | N | H | D | - | - | - | - | - | L | V | Y | E | - | - | L | K | A | D | - | P | C | G | H | C | L | I | I | N | N | V | N | F | S | R | D | S | - | - | N | L | L | T | R | D | G | S | D | V | D | C | R | K | L | E | K | R | F | - | K | A | L | R | F | D | V | L | T | R | R | D | L | E | A | K | A | M | V | S | E | L | R | K | L | A | R | Q | D | H | - | R | A | L | D | C | C | V | V | V | I | L | S | H | G | C | Q | M | S | H | I | Q | F | P | G | G | - | I | Y | G | T | D | G | K | P | V | P | V | E | E | I | V | N | Y | F | N | G | S | N | C | P | S | L | R | G | K | P | K | L | F | F | I | Q | A | C | - | - | G | G | E | Q | R | D | R | G | F | V | V | D | C | D | - | - | - | S | P | G | D | - | D | A | P | G | G | S | - | - | - | - | - | - | - | L | E | S | D | A | T | P | F | Q | A | P | S | G | N | V | - | - | - | - | - | - | - | - | D | E | P | D | A | V | - | - | - | - | - | - | - | A | S | L | - | - | - | P | T | P | S | D | I | L | V | S | Y | S | T | F | P | - | - | G | - | - | - | - | - | - | - | - | F | V | S | W | R | E | T | A | R | G | S | W | Y | V | E | M | L | D | S | V | L | E | Q | Y | A | H | - | S | - | - | - | - | - | - | - | - | - | - | - | - | - | - | - | - | - | - | - | - | - | - | - | - | - | - | - | - | - | - | - | E | H | L | L | N | M | L | - | - | - | - | - | L | R | - | - | - | - | - | - | - | - | - | - | - | - | V | A | D | A | V | - | - | - | - | - | S | A | K | - | - | - | - | - | - | - | G | Q | Y | - | K | Q | M | - | - | P | G | - | - | - | - | - | - | - | - | - | - | - | - | - | - | - | C | F | N | F | - | - | - | - | - | - | - | - | - | L | R | K | K | F | F | F | A | Y | K | - | - | - | - | - | - | - | - | - | - | - | - | - | - | - | - | - | - | - | - | - | - | - | - | - | - | - | - | - | - | - | - | - | - | - | - | - | M | - | - | - | - | - | - | - | - | - | - | - | - | - | - | - | - | - | - | - | - | - | - | - | - | - | - | - | - | - | - | - | - | - | - | - | - | - | - | - | - | - | - | - | - | - | - | - | - | - | - | - | - | G | D | I | E | K | G | R | K | I | F | V | Q | K | C | S | Q | C | H | T | V | E | K | G | G | K | H | K | T | G | P | N | L | N | G | L | F | G | R | K | T | G | Q | A | E | G | F | S | Y | T | D | A | N | K | N | K | G | I | T | W | G | E | D | T | L | M | E | Y | L | E | N | P | K | K | Y | I | P | G | T | K | M | I | F | A | G | I | K | K | K | S | E | R | A | D | L | I | A | Y | L | K | D | A | T | S | K | - | - | - | - | - | - |
| Bison\_bison\_bison | M | - | - | - | - | - | - | - | - | - | - | - | - | - | - | - | - | - | - | - | - | - | - | - | - | - | - | - | - | - | - | - | - | - | - | - | - | - | - | - | - | - | - | - | - | - | - | - | - | - | - | - | - | - | - | - | - | - | - | - | - | - | - | - | - | - | - | - | - | - | - | - | - | - | - | - | - | - | - | - | - | - | - | - | - | - | - | - | - | - | - | - | - | - | - | - | - | - | D | A | K | A | - | R | N | - | - | - | - | - | - | - | - | - | - | - | - | - | - | - | - | C | L | L | - | - | - | - | - | - | - | - | - | - | - | - | - | - | - | - | - | - | - | - | - | - | - | - | - | - | - | - | - | - | - | - | - | - | - | - | - | - | - | - | - | - | - | - | - | - | - | - | - | - | - | - | - | - | - | - | - | - | - | - | - | - | - | - | - | - | - | - | - | - | - | - | - | - | - | - | - | - | - | - | - | - | - | - | - | - | - | - | - | - | - | - | - | - | - | - | - | - | - | - | - | - | - | - | - | - | - | - | - | - | - | - | - | - | - | - | - | - | - | - | - | Q | H | R | E | A | L | E | R | D | I | K | T | S | Y | - | I | M | D | H | M | I | S | N | G | V | L | T | V | S | E | E | E | K | V | K | N | E | - | - | - | - | - | - | - | - | - | - | - | - | - | - | - | - | - | P | T | Q | R | Q | R | A | A | M | L | I | K | M | I | L | K | K | - | - | - | - | D | N | H | S | Y | I | S | - | - | - | - | - | F | Y | N | A | L | - | L | H | E | G | - | - | - | - | - | - | Y | K | D | L | A | A | L | L | H | G | G | I | P | V | V | S | S | S | D | G | - | K | - | - | - | - | - | - | - | - | - | - | - | - | - | - | - | - | - | - | - | - | - | - | D | S | V | S | G | I | T | - | - | - | - | - | - | - | - | - | - | - | S | Y | V | R | T | V | L | C | E | G | G | - | - | - | - | - | - | - | V | P | Q | R | P | V | V | F | V | T | R | K | K | L | V | N | A | I | - | - | - | - | - | - | - | - | - | - | Q | Q | K | L | S | - | - | - | - | - | - | - | - | - | - | - | - | - | - | - | - | - | - | - | - | - | - | - | - | - | - | - | - | - | - | - | - | K | L | N | G | E | P | - | - | - | - | - | - | - | - | - | - | - | - | - | - | - | - | - | - | - | - | - | - | - | - | - | - | - | - | - | - | - | - | - | - | - | - | - | - | G | W | V | T | I | Y | G | M | A | G | C | G | K | S | V | L | A | A | E | A | V | - | R | D | H | S | F | L | E | D | C | F | P | G | G | V | H | W | V | S | V | G | - | - | - | - | - | - | - | K | - | - | - | - | - | - | Q | D | K | S | - | - | - | - | - | - | - | - | - | - | - | - | G | L | L | M | K | L | Q | N | L | C | A | R | L | D | - | - | - | - | Q | - | - | - | - | - | - | - | - | - | - | - | - | - | - | - | D | E | S | F | S | Q | R | L | P | L | N | I | - | - | - | - | - | - | - | - | - | - | - | - | - | - | - | - | E | E | A | K | D | R | L | R | I | L | M | L | R | - | - | - | - | - | - | - | - | - | - | - | - | K | H | P | R | S | L | L | - | - | I | L | D | D | I | W | D | P | W | V | L | K | A | F | D | - | N | Q | C | Q | I | L | L | T | T | R | D | K | S | V | T | D | S | V | - | M | G | P | - | - | K | Y | V | V | P | V | E | S | - | - | - | G | L | G | K | E | K | G | L | E | I | L | S | L | F | V | N | M | - | K | K | A | D | L | P | E | - | - | - | - | - | - | - | - | - | - | - | - | - | - | Q | A | H | S | I | I | Q | E | C | K | - | - | - | - | - | - | - | - | - | - | - | - | - | - | - | - | - | - | - | G | - | S | P | L | V | V | S | L | I | G | A | L | L | R | D | F | P | N | - | - | - | - | - | - | - | - | - | - | R | W | E | Y | Y | L | - | R | Q | L | Q | N | K | Q | F | K | R | I | - | - | - | - | - | - | - | - | - | - | - | - | - | - | - | - | - | - | - | - | - | - | R | K | S | S | S | Y | D | - | - | - | - | Y | E | A | L | D | E | A | M | S | I | S | V | E | - | M | L | R | - | - | - | E | D | I | K | D | Y | - | Y | T | D | L | S | I | L | Q | K | D | V | K | V | P | T | K | V | L | C | I | L | W | D | - | - | - | - | L | E | T | E | E | V | E | D | I | L | Q | E | F | - | - | - | - | - | - | - | - | - | - | - | V | N | K | S | L | L | F | C | D | R | N | G | K | S | - | - | - | - | - | - | - | - | - | F | L | Y | Y | L | H | D | L | Q | V | D | F | L | A | E | K | - | - | N | C | S | Q | - | L | - | Q | D | L | H | K | K | M | V | T | Q | F | Q | R | - | H | H | - | - | - | - | - | - | Q | L | Y | S | L | S | P | D | - | Q | E | - | - | - | - | - | - | - | D | C | M | Y | W | Y | S | F | L | A | Y | H | M | A | S | A | N | - | - | - | - | M | H | K | E | L | C | A | L | M | F | S | L | D | W | I | K | A | K | T | E | L | V | G | P | A | - | - | - | - | - | - | - | - | H | L | I | H | E | F | V | E | Y | R | H | I | L | D | E | K | - | - | - | - | - | - | - | D | G | I | D | C | E | N | F | Q | - | - | - | E | F | L | S | L | N | G | H | L | L | G | R | Q | P | F | - | - | - | - | - | - | - | - | - | - | - | - | - | - | - | - | - | - | - | - | - | - | - | - | - | - | - | - | - | - | - | - | - | - | - | - | - | - | - | - | - | - | - | - | - | - | - | - | - | - | - | - | - | - | - | - | - | - | - | - | - | - | - | - | - | - | - | - | - | - | - | P | N | I | V | Q | L | G | L | C | E | L | E | T | S | E | V | Y | R | Q | A | K | L | Q | A | K | Q | E | V | D | N | G | - | - | - | - | - | - | - | - | - | - | - | - | - | - | - | - | - | - | - | - | - | - | - | - | - | - | - | - | - | M | L | Y | L | - | - | - | - | - | - | - | - | - | - | E | W | V | N | K | - | - | - | - | - | - | - | K | N | M | K | N | L | S | R | - | - | - | - | - | - | - | - | - | - | L | V | V | R | P | H | T | D | - | - | - | - | - | - | - | - | - | - | - | - | - | - | - | A | V | Y | H | A | C | - | - | - | F | S | - | E | D | G | Q | R | I | A | - | - | - | - | - | - | - | - | - | S | C | G | A | D | K | T | L | Q | V | F | K | - | - | - | - | - | A | E | T | G | E | K | L | L | E | I | - | K | A | H | E | D | E | - | - | - | V | L | C | C | A | F | S | - | A | - | - | - | - | - | - | - | - | - | - | - | - | - | D | D | K | F | I | A | T | C | S | V | D | K | K | V | K | I | - | - | - | W | N | S | M | - | - | - | - | - | - | - | - | - | - | - | - | - | - | - | - | - | - | - | - | - | - | - | - | - | - | - | - | - | - | - | - | - | - | - | - | - | - | - | - | - | - | - | - | - | - | - | - | - | - | - | - | - | - | - | - | - | - | - | - | - | - | - | - | - | - | - | - | - | - | - | - | - | - | - | - | - | - | - | - | - | - | - | - | - | - | - | - | - | - | - | - | - | - | - | - | - | - | - | - | - | - | - | - | - | - | - | - | - | - | - | - | - | - | - | - | - | - | - | - | - | - | - | - | - | - | - | - | - | - | - | - | - | - | - | - | - | - | - | - | T | G | E | L | V | R | I | Y | - | D | E | H | S | - | - | - | - | - | E | Q | V | N | - | - | - | - | - | - | - | - | - | - | - | - | - | C | C | H | - | - | - | - | - | - | - | - | - | - | - | - | - | F | T | N | N | S | - | - | - | - | - | - | - | - | - | - | - | - | - | - | - | - | N | H | - | L | L | L | A | T | G | S | N | - | - | - | - | D | Y | F | L | - | - | - | - | - | - | - | - | - | - | - | - | K | L | W | D | L | - | - | - | N | Q | E | E | C | R | N | T | M | F | - | - | - | - | - | - | - | - | - | - | - | - | - | - | - | - | - | G | H | T | N | S | - | - | - | - | - | - | - | - | - | - | - | - | - | - | - | - | V | S | H | C | R | F | S | P | D | D | Q | - | - | - | - | - | - | - | - | - | - | - | - | - | - | - | - | - | - | - | - | - | - | - | - | - | - | - | - | V | L | A | S | C | S | A | D | G | T | L | K | L | - | - | - | - | - | - | - | - | - | - | - | - | - | - | - | - | - | - | - | W | D | - | V | K | S | A | N | E | R | K | S | - | - | - | - | - | - | I | N | - | - | - | - | - | - | - | - | - | - | - | - | - | - | - | - | - | - | - | - | - | - | V | K | Q | F | - | - | - | - | - | - | - | - | F | L | N | S | E | E | P | Q | E | - | - | - | - | - | - | - | - | - | - | - | - | - | - | - | - | - | - | - | - | - | - | - | - | - | - | - | D | L | E | - | V | I | V | K | C | C | - | - | - | - | - | - | - | - | - | - | - | - | - | - | - | - | S | W | S | A | D | G | A | K | I | M | V | A | A | K | N | K | V | - | - | - | - | - | - | - | - | - | - | - | F | L | F | D | I | H | S | - | - | - | S | G | - | - | - | - | - | - | - | - | - | - | - | - | - | - | - | - | - | - | - | - | - | - | - | - | - | - | - | - | - | - | - | - | - | - | - | - | - | - | - | - | - | - | - | - | - | - | - | - | - | - | - | - | - | - | - | - | - | - | - | - | - | - | - | - | - | - | - | - | - | - | - | - | - | - | - | - | - | - | - | - | - | - | - | - | - | - | - | - | - | - | - | - | - | - | - | - | - | - | - | - | - | - | - | - | - | - | - | - | - | - | - | - | - | - | - | - | - | - | - | - | - | - | - | - | - | - | L | L | A | - | - | - | - | - | - | - | - | - | - | - | - | - | - | - | - | - | - | - | - | - | - | - | - | - | - | - | - | - | - | - | - | - | - | - | - | - | - | - | - | - | - | - | - | - | - | - | - | - | - | - | - | - | - | - | - | - | - | - | - | - | - | - | - | - | - | - | - | - | - | - | - | - | - | - | - | - | - | - | - | - | - | - | - | - | - | - | - | - | - | - | - | - | - | - | - | - | - | - | - | - | - | - | - | - | - | - | - | - | - | - | - | - | - | - | - | - | - | - | - | - | - | - | - | - | - | - | - | - | - | - | - | - | - | - | - | - | - | - | - | - | - | - | - | - | - | - | - | - | - | - | - | - | - | - | - | - | - | - | - | - | - | - | - | - | - | - | - | - | - | - | - | - | - | - | - | - | - | - | - | - | - | - | E | M | - | - | - | H | M | G | H | H | S | A | V | Q | Y | C | - | - | - | - | - | - | - | - | - | - | - | - | - | - | - | - | - | - | - | - | - | - | - | - | - | D | F | S | P | H | N | H | L | A | V | V | A | L | S | Q | - | - | - | - | - | - | Y | C | V | E | - | - | - | - | - | - | - | - | - | - | - | - | - | - | - | - | - | L | W | S | M | D | S | G | - | - | - | - | - | - | - | - | - | - | - | - | - | - | - | - | - | - | - | - | - | - | - | - | - | - | - | - | - | L | K | V | A | D | C | R | G | H | L | S | - | - | - | - | - | - | - | - | - | - | - | - | - | - | - | - | - | - | - | - | - | - | - | - | - | - | - | - | - | - | - | - | - | - | - | - | - | - | - | - | - | - | - | - | W | V | H | C | V | M | F | S | - | - | - | - | - | - | - | - | - | - | - | P | D | G | S | - | - | - | - | - | - | - | - | - | - | S | F | L | T | S | S | D | D | Q | - | - | - | - | - | - | - | - | - | - | - | - | - | - | T | I | R | L | - | W | - | - | - | - | - | - | - | - | - | - | - | - | - | - | - | - | - | - | - | - | - | - | - | - | - | - | - | - | - | - | - | - | - | - | - | - | - | - | E | T | K | K | V | C | K | N | S | A | T | V | L | K | Q | E | - | - | - | - | - | - | - | - | - | - | - | - | - | - | - | - | I | D | V | V | F | - | - | - | - | - | - | - | - | - | - | - | - | - | - | - | - | - | - | - | - | - | - | - | - | - | - | - | - | - | - | - | - | - | - | - | - | - | - | - | - | - | - | - | - | - | - | - | - | - | - | - | - | - | - | Q | E | N | E | V | T | - | - | - | - | - | - | - | - | - | - | - | - | - | - | - | - | - | - | - | - | - | - | - | - | - | - | - | - | - | - | - | - | - | - | - | - | V | L | A | V | D | N | I | R | R | L | Q | L | - | I | N | G | N | T | - | - | - | - | - | - | - | - | - | - | G | Q | I | D | Y | L | T | E | A | - | - | - | - | - | - | - | - | - | - | - | - | - | - | - | - | - | - | - | - | - | - | Q | - | - | - | - | - | - | - | - | - | - | - | V | S | C | C | C | L | S | S | H | L | Q | - | - | - | - | - | - | - | - | - | Y | L | A | F | G | G | - | - | - | - | - | - | - | - | - | - | - | - | - | - | - | - | - | - | - | - | - | - | - | - | - | - | - | - | - | - | - | - | - | - | - | - | E | - | - | - | - | - | - | - | - | - | D | G | A | - | - | - | - | - | - | - | - | - | - | I | - | - | - | - | - | - | - | - | - | - | - | - | - | - | - | - | - | - | - | - | - | - | - | - | - | - | - | - | - | - | - | - | - | - | - | - | - | - | - | Q | I | L | E | L | L | N | N | R | I | F | Q | S | R | N | G | - | - | - | - | - | H | - | - | - | - | - | - | - | - | - | - | - | - | K | T | T | V | R | H | I | Q | F | T | - | - | - | - | - | - | - | - | - | - | - | - | - | - | - | - | - | - | - | - | - | - | - | - | - | - | - | - | - | - | - | - | - | - | - | - | - | A | D | G | K | T | L | I | S | S | S | - | D | D | S | - | - | - | - | - | - | - | - | A | V | Q | - | I | - | - | - | - | - | - | - | - | - | - | - | - | - | - | - | - | - | - | - | - | - | - | - | - | - | - | - | - | - | - | - | - | - | - | - | - | - | - | - | W | N | - | - | - | - | - | - | - | - | - | - | W | Q | S | E | D | P | F | F | - | - | - | - | - | - | - | - | - | - | - | - | - | - | - | - | - | - | - | - | - | - | - | - | - | - | - | - | - | - | - | - | - | - | - | - | - | - | - | - | - | - | - | - | - | - | - | - | - | - | L | Q | A | H | Q | - | - | - | - | - | - | - | - | - | - | - | - | - | - | - | - | - | - | - | - | - | - | - | - | - | - | E | T | V | K | D | F | R | L | L | - | - | - | - | - | - | - | - | - | - | - | - | - | - | - | - | - | - | - | - | - | - | - | K | N | - | S | R | L | L | - | S | W | - | - | - | S | F | D | G | T | V | - | - | - | - | - | - | - | - | - | - | - | - | - | - | - | - | - | - | - | - | - | - | - | - | - | - | - | - | - | - | - | - | - | - | - | K | V | W | N | I | I | T | R | R | I | - | - | - | - | - | E | K | D | F | V | C | - | - | - | - | - | - | - | H | Q | - | - | - | - | - | - | - | - | - | - | - | - | - | - | - | - | - | - | - | D | T | V | L | - | - | - | - | - | - | - | - | - | - | - | - | - | - | - | S | C | D | - | - | - | - | - | - | - | - | - | - | - | - | - | - | - | I | S | S | D | A | T | - | K | F | S | S | T | - | S | A | D | K | T | - | - | - | - | - | - | A | K | I | W | S | F | E | L | L | - | - | - | - | - | - | - | - | - | - | - | - | - | - | - | - | - | - | - | - | - | - | - | - | S | P | L | H | E | - | - | - | - | - | - | - | - | L | - | - | - | - | R | G | H | K | G | C | V | - | - | - | - | - | - | - | - | - | R | C | S | - | S | F | S | - | - | - | - | - | - | - | - | - | V | N | N | S | - | L | L | A | T | - | - | - | - | - | - | - | - | G | D | D | N | G | E | I | R | I | W | N | V | S | N | G | E | L | L | - | - | - | - | - | - | - | - | - | - | - | - | - | - | - | - | - | - | - | - | - | - | - | - | - | - | - | - | - | - | - | - | - | - | - | - | - | - | - | - | - | - | - | - | - | - | - | - | - | - | - | - | - | - | - | - | H | L | C | A | P | I | S | V | E | - | - | - | - | - | - | - | - | - | - | - | - | - | - | - | - | - | - | - | - | E | G | A | A | T | H | G | G | W | - | - | - | - | - | - | - | - | V | T | D | L | C | F | S | P | D | T | K | - | - | - | - | - | - | - | - | - | - | - | - | - | - | - | - | - | - | - | - | - | - | - | - | - | - | - | - | - | - | - | - | - | - | - | - | - | - | - | - | - | - | - | - | - | M | L | V | S | - | - | A | G | G | Y | L | K | - | - | - | - | - | - | - | - | W | W | N | V | V | T | - | - | - | - | - | - | - | - | - | - | - | - | - | - | - | - | - | - | - | - | - | - | - | - | - | - | - | - | - | - | - | - | - | - | - | - | - | - | - | - | - | - | - | - | - | - | - | - | - | - | - | - | - | - | - | - | - | - | - | - | - | - | - | - | - | - | - | - | - | - | - | - | - | - | - | - | - | - | - | - | - | - | - | - | - | - | - | - | - | - | - | - | - | - | - | - | - | - | - | - | - | - | - | - | - | - | - | - | - | - | - | - | - | - | - | - | - | - | - | - | - | - | - | - | - | - | - | - | - | - | - | - | - | - | - | - | - | - | - | - | - | - | - | - | - | - | - | - | - | - | - | - | - | - | - | - | - | - | - | - | - | - | - | - | - | - | - | - | - | - | - | - | - | - | - | - | - | - | - | - | - | - | - | - | - | - | - | - | - | - | - | - | - | - | - | - | - | - | - | - | - | - | - | - | - | - | - | - | - | - | - | - | - | - | - | - | - | - | - | - | - | - | - | - | - | - | - | - | - | - | - | - | - | - | - | - | - | - | - | - | - | - | - | - | - | - | - | - | - | - | - | - | - | - | - | - | - | - | - | - | - | - | - | - | - | - | - | G | E | S | S | Q | T | F | Y | T | - | - | - | - | - | N | G | - | - | - | T | N | L | K | K | I | - | - | - | H | V | S | P | - | - | - | - | D | F | K | T | - | - | - | - | - | - | - | - | - | - | - | - | - | - | - | - | - | - | - | - | - | - | - | - | - | - | Y | V | T | V | D | - | N | L | G | I | L | - | - | Y | I | L | K | I | - | - | - | - | - | - | - | - | - | - | - | - | - | - | - | - | - | - | - | - | - | - | - | - | - | - | - | - | - | - | - | - | - | - | L | E | - | - | - | - | - | - | - | - | - | - | - | - | - | - | - | - | - | - | - | - | - | - | - | - | - | - | - | - | - | - | - | - | - | - | - | - | - | - | - | - | - | - | - | - | - | - | - | - | - | - | - | - | M | - | - | - | - | - | - | - | - | - | - | - | - | - | - | - | - | - | - | - | - | - | - | - | - | - | - | - | - | - | - | - | - | - | - | - | - | - | - | - | - | - | - | - | - | - | - | - | - | - | - | - | - | - | - | - | - | - | - | - | - | - | - | - | - | - | - | - | - | - | - | - | - | - | - | - | - | - | - | - | - | - | - | - | - | - | - | - | - | - | - | - | - | - | - | - | - | - | - | - | - | - | - | - | - | - | - | - | - | - | - | - | - | - | - | - | - | - | - | - | - | - | - | - | - | - | - | - | - | - | - | - | - | - | - | - | - | - | - | - | - | - | - | - | - | - | - | - | - | - | - | - | - | - | - | - | - | - | - | - | - | - | - | - | - | - | - | - | - | - | - | - | - | - | - | - | - | - | - | - | - | - | - | - | - | - | - | - | - | - | - | - | - | - | - | - | - | - | - | - | - | - | - | - | - | - | - | - | - | - | - | - | - | - | - | - | - | - | - | - | - | I | E | D | - | - | - | - | - | - | - | - | - | - | - | - | - | - | - | - | - | - | - | - | - | - | - | - | - | I | Q | R | A | G | - | - | - | - | - | - | - | - | S | G | S | R | R | D | Q | - | A | R | Q | L | I | L | D | L | E | T | R | G | - | S | Q | A | L | P | L | F | I | S | C | L | E | D | T | G | Q | D | T | L | A | S | L | L | K | T | S | R | Q | A | A | - | - | - | - | K | Q | D | V | - | E | A | I | - | - | - | - | - | - | - | - | - | - | - | - | - | - | - | - | - | - | - | - | - | - | - | - | - | - | R | P | L | D | - | - | - | - | - | - | - | - | - | - | - | - | - | - | - | L | K | P | V | - | - | - | - | - | - | - | - | - | - | - | - | - | - | - | - | - | - | - | - | - | - | - | - | - | - | - | - | - | - | - | - | - | - | - | - | - | - | - | V | L | G | P | E | G | L | K | P | G | E | L | R | - | V | M | K | Q | D | P | S | K | L | - | S | - | Q | - | - | - | - | - | - | - | - | - | - | - | - | - | - | - | - | - | - | - | - | - | - | - | - | - | - | - | - | - | - | - | - | - | - | - | - | - | - | - | - | - | - | - | - | - | - | - | - | - | - | - | G | K | L | A | P | V | - | - | - | - | - | - | - | - | - | - | - | - | - | - | - | - | - | - | - | - | - | - | - | - | - | - | V | L | - | - | - | - | - | - | - | - | - | - | - | - | - | - | - | - | - | - | - | - | - | - | - | - | - | - | - | - | - | - | - | - | - | - | - | - | - | - | - | - | - | - | - | - | - | - | - | - | - | - | - | - | - | - | - | - | - | - | - | - | - | - | - | - | - | - | - | - | G | P | E | E | L | W | P | - | - | - | - | - | - | - | - | - | - | - | - | - | - | - | - | - | - | - | - | - | - | - | - | - | - | P | K | L | R | P | - | - | - | - | - | - | E | V | L | R | P | - | E | V | - | - | - | - | - | - | - | - | - | - | - | - | - | - | - | - | - | - | P | R | A | V | D | A | G | S | G | G | F | - | - | - | - | - | - | - | - | - | - | - | - | - | T | D | V | C | - | - | - | - | - | - | - | - | - | - | P | - | Q | D | R | A | K | G | N | A | D | - | - | - | - | - | L | A | Y | V | - | - | L | N | A | D | - | P | C | G | H | C | L | I | I | N | N | V | N | F | C | R | E | S | - | - | G | L | R | A | R | T | G | S | N | I | D | C | E | R | M | W | R | R | F | - | H | L | L | Q | F | V | V | E | V | K | C | D | L | T | A | K | Q | M | V | Q | A | L | M | Q | L | A | R | Q | D | H | - | S | A | L | D | C | C | V | V | V | I | L | S | H | G | C | Q | A | S | H | L | Q | F | P | G | A | - | V | Y | G | T | D | G | C | P | V | S | V | E | R | I | V | N | T | F | N | G | T | G | C | P | S | L | R | G | K | P | K | L | F | F | I | Q | A | C | - | - | G | G | E | Q | K | D | H | G | F | E | V | E | S | T | - | - | - | S | P | E | D | - | K | T | P | S | S | D | - | - | - | - | - | - | - | P | E | A | D | A | T | P | F | Q | E | G | P | R | S | I | - | - | - | - | - | - | - | - | D | E | P | D | A | V | - | - | - | - | - | - | - | S | S | L | - | - | - | P | T | P | S | D | I | L | V | S | Y | S | T | F | P | - | - | G | - | - | - | - | - | - | - | - | F | V | S | W | R | D | P | K | S | G | S | W | Y | I | E | T | L | D | S | I | F | E | Q | W | A | H | - | C | - | - | - | - | - | - | - | - | - | - | - | - | - | - | - | - | - | - | - | - | - | - | - | - | - | - | - | - | - | - | - | E | D | L | Q | T | L | L | - | - | - | - | - | L | R | - | - | - | - | - | - | - | - | - | - | - | - | V | A | N | A | V | - | - | - | - | - | S | V | K | - | - | - | - | - | - | - | G | I | Y | - | K | Q | M | - | - | P | G | - | - | - | - | - | - | - | - | - | - | - | - | - | - | - | C | F | N | F | - | - | - | - | - | - | - | - | - | L | R | K | K | L | F | F | K | T | - | - | - | - | - | - | - | - | - | - | - | - | - | - | - | - | - | - | - | - | - | - | - | - | - | - | - | - | - | - | - | - | - | - | - | - | - | - | M | - | - | - | - | - | - | - | - | - | - | - | - | - | - | - | - | - | - | - | - | - | - | - | - | - | - | - | - | - | - | - | - | - | - | - | - | - | - | - | - | - | - | - | - | - | - | - | - | - | - | - | - | G | D | V | E | K | G | K | K | I | F | V | Q | K | C | A | Q | C | H | T | V | E | K | G | G | K | H | K | T | G | P | N | L | H | G | L | F | G | R | K | T | G | Q | A | P | G | F | S | Y | T | D | A | N | K | N | K | G | I | T | W | G | E | E | T | L | M | E | Y | L | E | N | P | K | K | Y | I | P | G | T | K | M | I | F | A | G | I | K | K | K | G | E | R | E | D | L | I | A | Y | L | K | K | A | T | N | E | - | - | - | - | - | - |
| Bos\_mutus | M | - | - | - | - | - | - | - | - | - | - | - | - | - | - | - | - | - | - | - | - | - | - | - | - | - | - | - | - | - | - | - | - | - | - | - | - | - | - | - | - | - | - | - | - | - | - | - | - | - | - | - | - | - | - | - | - | - | - | - | - | - | - | - | - | - | - | - | - | - | - | - | - | - | - | - | - | - | - | - | - | - | - | - | - | - | - | - | - | - | - | - | - | - | - | - | - | - | D | A | K | A | - | R | N | - | - | - | - | - | - | - | - | - | - | - | - | - | - | - | - | C | L | L | - | - | - | - | - | - | - | - | - | - | - | - | - | - | - | - | - | - | - | - | - | - | - | - | - | - | - | - | - | - | - | - | - | - | - | - | - | - | - | - | - | - | - | - | - | - | - | - | - | - | - | - | - | - | - | - | - | - | - | - | - | - | - | - | - | - | - | - | - | - | - | - | - | - | - | - | - | - | - | - | - | - | - | - | - | - | - | - | - | - | - | - | - | - | - | - | - | - | - | - | - | - | - | - | - | - | - | - | - | - | - | - | - | - | - | - | - | - | - | - | - | Q | H | R | E | A | L | E | R | D | I | K | T | S | Y | - | I | M | D | H | M | I | S | N | G | V | L | T | V | S | E | E | E | K | V | K | N | E | - | - | - | - | - | - | - | - | - | - | - | - | - | - | - | - | - | P | T | Q | R | Q | R | A | A | M | L | I | K | M | I | L | K | K | - | - | - | - | D | N | H | S | Y | I | S | - | - | - | - | - | F | Y | N | A | L | - | L | H | E | G | - | - | - | - | - | - | Y | K | D | L | A | A | L | L | H | G | G | V | P | V | V | S | S | S | D | G | - | K | - | - | - | - | - | - | - | - | - | - | - | - | - | - | - | - | - | - | - | - | - | - | D | S | V | S | G | I | T | - | - | - | - | - | - | - | - | - | - | - | S | Y | V | R | T | V | L | C | E | G | G | - | - | - | - | - | - | - | V | P | Q | R | P | V | V | F | V | T | R | K | K | L | V | N | A | I | - | - | - | - | - | - | - | - | - | - | Q | Q | K | L | S | - | - | - | - | - | - | - | - | - | - | - | - | - | - | - | - | - | - | - | - | - | - | - | - | - | - | - | - | - | - | - | - | K | L | N | G | E | P | - | - | - | - | - | - | - | - | - | - | - | - | - | - | - | - | - | - | - | - | - | - | - | - | - | - | - | - | - | - | - | - | - | - | - | - | - | - | G | W | V | T | I | Y | G | M | A | G | C | G | K | S | V | L | A | A | E | A | V | - | R | D | H | S | F | L | E | D | C | F | P | G | G | V | H | W | V | S | V | G | - | - | - | - | - | - | - | K | - | - | - | - | - | - | Q | D | K | S | - | - | - | - | - | - | - | - | - | - | - | - | G | L | L | M | K | L | Q | N | L | C | A | R | L | D | - | - | - | - | Q | - | - | - | - | - | - | - | - | - | - | - | - | - | - | - | D | E | S | F | S | Q | R | L | P | L | N | I | - | - | - | - | - | - | - | - | - | - | - | - | - | - | - | - | E | E | A | K | D | R | L | R | I | L | M | L | R | - | - | - | - | - | - | - | - | - | - | - | - | K | H | P | R | S | L | L | - | - | I | L | D | D | I | W | D | P | W | V | L | K | A | F | D | - | N | Q | C | Q | I | L | L | T | T | R | D | K | S | V | T | D | S | V | - | M | G | P | - | - | K | Y | V | V | P | V | E | S | - | - | - | G | L | G | K | E | K | G | L | E | I | L | S | L | F | V | N | M | - | K | K | A | D | L | P | E | - | - | - | - | - | - | - | - | - | - | - | - | - | - | Q | A | H | S | I | I | Q | E | C | K | - | - | - | - | - | - | - | - | - | - | - | - | - | - | - | - | - | - | - | G | - | S | P | L | V | V | S | L | I | G | A | L | L | R | D | F | P | N | - | - | - | - | - | - | - | - | - | - | R | W | E | Y | Y | L | - | R | Q | L | Q | N | K | Q | F | K | R | I | - | - | - | - | - | - | - | - | - | - | - | - | - | - | - | - | - | - | - | - | - | - | R | K | S | S | S | Y | D | - | - | - | - | Y | E | A | L | D | E | A | M | S | I | S | V | E | - | M | L | R | - | - | - | E | D | I | K | D | Y | - | Y | T | D | L | S | I | L | Q | K | D | V | K | V | P | T | K | V | L | C | I | L | W | D | - | - | - | - | L | E | T | E | E | V | E | D | I | L | Q | E | F | - | - | - | - | - | - | - | - | - | - | - | V | N | K | S | L | L | F | C | D | R | N | G | K | S | - | - | - | - | - | - | - | - | - | F | L | Y | Y | L | H | D | L | Q | V | D | F | L | A | E | K | - | - | N | R | S | Q | - | L | - | Q | D | L | H | K | K | M | V | T | Q | F | Q | R | - | H | H | - | - | - | - | - | - | Q | L | Y | S | L | S | P | D | - | Q | E | - | - | - | - | - | - | - | D | C | M | Y | W | Y | S | F | L | A | Y | H | M | A | S | A | N | - | - | - | - | M | H | K | E | L | C | A | L | M | F | S | L | D | W | I | K | A | K | T | E | L | V | G | P | A | - | - | - | - | - | - | - | - | H | L | I | H | E | F | V | E | Y | R | H | I | L | D | E | K | - | - | - | - | - | - | - | D | G | I | D | C | E | N | F | Q | - | - | - | E | F | L | S | L | N | G | H | L | L | G | R | Q | P | F | - | - | - | - | - | - | - | - | - | - | - | - | - | - | - | - | - | - | - | - | - | - | - | - | - | - | - | - | - | - | - | - | - | - | - | - | - | - | - | - | - | - | - | - | - | - | - | - | - | - | - | - | - | - | - | - | - | - | - | - | - | - | - | - | - | - | - | - | - | - | - | P | N | I | V | Q | L | G | L | C | E | L | E | T | S | E | V | Y | R | Q | A | K | L | Q | A | K | Q | E | V | D | N | G | - | - | - | - | - | - | - | - | - | - | - | - | - | - | - | - | - | - | - | - | - | - | - | - | - | - | - | - | - | M | L | Y | L | - | - | - | - | - | - | - | - | - | - | E | W | V | N | K | - | - | - | - | - | - | - | K | N | M | K | N | L | S | R | - | - | - | - | - | - | - | - | - | - | L | V | V | R | P | H | T | D | - | - | - | - | - | - | - | - | - | - | - | - | - | - | - | A | V | Y | H | A | C | - | - | - | F | S | - | E | D | G | Q | R | I | A | - | - | - | - | - | - | - | - | - | S | C | G | A | D | K | T | L | Q | V | F | K | - | - | - | - | - | A | E | T | G | E | K | L | L | E | I | - | K | A | H | E | D | E | - | - | - | V | L | C | C | A | F | S | - | A | - | - | - | - | - | - | - | - | - | - | - | - | - | D | D | K | F | I | A | T | C | S | V | D | K | K | V | K | I | - | - | - | W | N | S | M | - | - | - | - | - | - | - | - | - | - | - | - | - | - | - | - | - | - | - | - | - | - | - | - | - | - | - | - | - | - | - | - | - | - | - | - | - | - | - | - | - | - | - | - | - | - | - | - | - | - | - | - | - | - | - | - | - | - | - | - | - | - | - | - | - | - | - | - | - | - | - | - | - | - | - | - | - | - | - | - | - | - | - | - | - | - | - | - | - | - | - | - | - | - | - | - | - | - | - | - | - | - | - | - | - | - | - | - | - | - | - | - | - | - | - | - | - | - | - | - | - | - | - | - | - | - | - | - | - | - | - | - | - | - | - | - | - | - | - | - | T | G | E | L | V | R | I | Y | - | D | E | H | S | - | - | - | - | - | E | Q | V | N | - | - | - | - | - | - | - | - | - | - | - | - | - | C | C | H | - | - | - | - | - | - | - | - | - | - | - | - | - | F | T | N | N | S | - | - | - | - | - | - | - | - | - | - | - | - | - | - | - | - | N | H | - | L | L | L | A | T | G | S | N | - | - | - | - | D | Y | F | L | - | - | - | - | - | - | - | - | - | - | - | - | K | L | W | D | L | - | - | - | N | Q | E | E | C | R | N | T | M | F | - | - | - | - | - | - | - | - | - | - | - | - | - | - | - | - | - | G | H | T | N | S | - | - | - | - | - | - | - | - | - | - | - | - | - | - | - | - | V | S | H | C | R | F | S | P | D | D | Q | - | - | - | - | - | - | - | - | - | - | - | - | - | - | - | - | - | - | - | - | - | - | - | - | - | - | - | - | V | L | A | S | C | S | A | D | G | T | L | K | L | - | - | - | - | - | - | - | - | - | - | - | - | - | - | - | - | - | - | - | W | D | - | V | K | S | A | N | E | R | K | S | - | - | - | - | - | - | I | N | - | - | - | - | - | - | - | - | - | - | - | - | - | - | - | - | - | - | - | - | - | - | V | K | Q | F | - | - | - | - | - | - | - | - | F | L | N | S | E | E | P | Q | E | - | - | - | - | - | - | - | - | - | - | - | - | - | - | - | - | - | - | - | - | - | - | - | - | - | - | - | D | L | E | - | V | I | V | K | C | C | - | - | - | - | - | - | - | - | - | - | - | - | - | - | - | - | S | W | S | A | D | G | A | K | I | M | V | A | A | K | N | K | V | - | - | - | - | - | - | - | - | - | - | - | F | L | F | D | I | H | S | - | - | - | S | G | - | - | - | - | - | - | - | - | - | - | - | - | - | - | - | - | - | - | - | - | - | - | - | - | - | - | - | - | - | - | - | - | - | - | - | - | - | - | - | - | - | - | - | - | - | - | - | - | - | - | - | - | - | - | - | - | - | - | - | - | - | - | - | - | - | - | - | - | - | - | - | - | - | - | - | - | - | - | - | - | - | - | - | - | - | - | - | - | - | - | - | - | - | - | - | - | - | - | - | - | - | - | - | - | - | - | - | - | - | - | - | - | - | - | - | - | - | - | - | - | - | - | - | - | - | - | L | L | A | - | - | - | - | - | - | - | - | - | - | - | - | - | - | - | - | - | - | - | - | - | - | - | - | - | - | - | - | - | - | - | - | - | - | - | - | - | - | - | - | - | - | - | - | - | - | - | - | - | - | - | - | - | - | - | - | - | - | - | - | - | - | - | - | - | - | - | - | - | - | - | - | - | - | - | - | - | - | - | - | - | - | - | - | - | - | - | - | - | - | - | - | - | - | - | - | - | - | - | - | - | - | - | - | - | - | - | - | - | - | - | - | - | - | - | - | - | - | - | - | - | - | - | - | - | - | - | - | - | - | - | - | - | - | - | - | - | - | - | - | - | - | - | - | - | - | - | - | - | - | - | - | - | - | - | - | - | - | - | - | - | - | - | - | - | - | - | - | - | - | - | - | - | - | - | - | - | - | - | - | - | - | - | E | M | - | - | - | H | M | G | H | H | S | A | V | Q | Y | C | - | - | - | - | - | - | - | - | - | - | - | - | - | - | - | - | - | - | - | - | - | - | - | - | - | D | F | S | P | H | N | H | L | A | V | V | A | L | S | Q | - | - | - | - | - | - | Y | C | V | E | - | - | - | - | - | - | - | - | - | - | - | - | - | - | - | - | - | L | W | S | M | D | S | G | - | - | - | - | - | - | - | - | - | - | - | - | - | - | - | - | - | - | - | - | - | - | - | - | - | - | - | - | - | L | K | V | A | D | C | R | G | H | L | S | - | - | - | - | - | - | - | - | - | - | - | - | - | - | - | - | - | - | - | - | - | - | - | - | - | - | - | - | - | - | - | - | - | - | - | - | - | - | - | - | - | - | - | - | W | V | H | C | V | M | F | S | - | - | - | - | - | - | - | - | - | - | - | P | D | G | S | - | - | - | - | - | - | - | - | - | - | S | F | L | T | S | S | D | D | Q | - | - | - | - | - | - | - | - | - | - | - | - | - | - | T | I | R | L | - | W | - | - | - | - | - | - | - | - | - | - | - | - | - | - | - | - | - | - | - | - | - | - | - | - | - | - | - | - | - | - | - | - | - | - | - | - | - | - | E | T | K | K | V | C | K | N | S | A | T | V | L | K | Q | E | - | - | - | - | - | - | - | - | - | - | - | - | - | - | - | - | I | D | V | V | F | - | - | - | - | - | - | - | - | - | - | - | - | - | - | - | - | - | - | - | - | - | - | - | - | - | - | - | - | - | - | - | - | - | - | - | - | - | - | - | - | - | - | - | - | - | - | - | - | - | - | - | - | - | - | Q | E | N | E | V | T | - | - | - | - | - | - | - | - | - | - | - | - | - | - | - | - | - | - | - | - | - | - | - | - | - | - | - | - | - | - | - | - | - | - | - | - | V | L | A | V | D | N | I | R | R | L | Q | L | - | I | N | G | N | T | - | - | - | - | - | - | - | - | - | - | G | Q | I | D | Y | L | T | E | A | - | - | - | - | - | - | - | - | - | - | - | - | - | - | - | - | - | - | - | - | - | - | Q | - | - | - | - | - | - | - | - | - | - | - | V | S | C | C | C | L | S | P | H | L | Q | - | - | - | - | - | - | - | - | - | Y | L | A | F | G | G | - | - | - | - | - | - | - | - | - | - | - | - | - | - | - | - | - | - | - | - | - | - | - | - | - | - | - | - | - | - | - | - | - | - | - | - | E | - | - | - | - | - | - | - | - | - | D | G | A | - | - | - | - | - | - | - | - | - | - | I | - | - | - | - | - | - | - | - | - | - | - | - | - | - | - | - | - | - | - | - | - | - | - | - | - | - | - | - | - | - | - | - | - | - | - | - | - | - | - | Q | I | L | E | L | L | N | N | R | I | F | Q | S | R | N | G | - | - | - | - | - | H | - | - | - | - | - | - | - | - | - | - | - | - | K | T | T | V | R | H | I | Q | F | T | - | - | - | - | - | - | - | - | - | - | - | - | - | - | - | - | - | - | - | - | - | - | - | - | - | - | - | - | - | - | - | - | - | - | - | - | - | A | D | G | K | T | L | I | S | S | S | - | D | D | S | - | - | - | - | - | - | - | - | A | V | Q | - | I | - | - | - | - | - | - | - | - | - | - | - | - | - | - | - | - | - | - | - | - | - | - | - | - | - | - | - | - | - | - | - | - | - | - | - | - | - | - | - | W | N | - | - | - | - | - | - | - | - | - | - | W | Q | S | E | D | P | F | F | - | - | - | - | - | - | - | - | - | - | - | - | - | - | - | - | - | - | - | - | - | - | - | - | - | - | - | - | - | - | - | - | - | - | - | - | - | - | - | - | - | - | - | - | - | - | - | - | - | - | L | Q | A | H | Q | - | - | - | - | - | - | - | - | - | - | - | - | - | - | - | - | - | - | - | - | - | - | - | - | - | - | E | T | V | K | D | F | R | L | L | - | - | - | - | - | - | - | - | - | - | - | - | - | - | - | - | - | - | - | - | - | - | - | K | N | - | S | R | L | L | - | S | W | - | - | - | S | F | D | G | T | V | - | - | - | - | - | - | - | - | - | - | - | - | - | - | - | - | - | - | - | - | - | - | - | - | - | - | - | - | - | - | - | - | - | - | - | K | V | W | N | I | I | T | R | R | I | - | - | - | - | - | E | K | D | F | V | C | - | - | - | - | - | - | - | H | Q | - | - | - | - | - | - | - | - | - | - | - | - | - | - | - | - | - | - | - | D | T | V | L | - | - | - | - | - | - | - | - | - | - | - | - | - | - | - | S | C | D | - | - | - | - | - | - | - | - | - | - | - | - | - | - | - | I | S | S | D | A | T | - | K | F | S | S | T | - | S | A | D | K | T | - | - | - | - | - | - | A | K | I | W | S | F | E | L | L | - | - | - | - | - | - | - | - | - | - | - | - | - | - | - | - | - | - | - | - | - | - | - | - | S | P | L | H | E | - | - | - | - | - | - | - | - | L | - | - | - | - | R | G | H | K | G | C | V | - | - | - | - | - | - | - | - | - | R | C | S | - | S | F | S | - | - | - | - | - | - | - | - | - | V | N | N | S | - | L | L | A | T | - | - | - | - | - | - | - | - | G | D | D | N | G | E | I | R | I | W | N | V | S | N | G | E | L | L | - | - | - | - | - | - | - | - | - | - | - | - | - | - | - | - | - | - | - | - | - | - | - | - | - | - | - | - | - | - | - | - | - | - | - | - | - | - | - | - | - | - | - | - | - | - | - | - | - | - | - | - | - | - | - | - | H | L | C | A | P | I | S | V | E | - | - | - | - | - | - | - | - | - | - | - | - | - | - | - | - | - | - | - | - | E | G | A | A | T | H | G | G | W | - | - | - | - | - | - | - | - | V | T | D | L | C | F | S | P | D | T | K | - | - | - | - | - | - | - | - | - | - | - | - | - | - | - | - | - | - | - | - | - | - | - | - | - | - | - | - | - | - | - | - | - | - | - | - | - | - | - | - | - | - | - | - | - | M | L | V | S | - | - | A | G | G | Y | L | K | - | - | - | - | - | - | - | - | W | W | N | V | V | T | - | - | - | - | - | - | - | - | - | - | - | - | - | - | - | - | - | - | - | - | - | - | - | - | - | - | - | - | - | - | - | - | - | - | - | - | - | - | - | - | - | - | - | - | - | - | - | - | - | - | - | - | - | - | - | - | - | - | - | - | - | - | - | - | - | - | - | - | - | - | - | - | - | - | - | - | - | - | - | - | - | - | - | - | - | - | - | - | - | - | - | - | - | - | - | - | - | - | - | - | - | - | - | - | - | - | - | - | - | - | - | - | - | - | - | - | - | - | - | - | - | - | - | - | - | - | - | - | - | - | - | - | - | - | - | - | - | - | - | - | - | - | - | - | - | - | - | - | - | - | - | - | - | - | - | - | - | - | - | - | - | - | - | - | - | - | - | - | - | - | - | - | - | - | - | - | - | - | - | - | - | - | - | - | - | - | - | - | - | - | - | - | - | - | - | - | - | - | - | - | - | - | - | - | - | - | - | - | - | - | - | - | - | - | - | - | - | - | - | - | - | - | - | - | - | - | - | - | - | - | - | - | - | - | - | - | - | - | - | - | - | - | - | - | - | - | - | - | - | - | - | - | - | - | - | - | - | - | - | - | - | - | - | - | - | - | - | G | E | S | S | Q | T | F | Y | T | - | - | - | - | - | N | G | - | - | - | T | N | L | K | K | I | - | - | - | H | V | S | P | - | - | - | - | D | F | K | T | - | - | - | - | - | - | - | - | - | - | - | - | - | - | - | - | - | - | - | - | - | - | - | - | - | - | Y | V | T | V | D | - | N | L | G | I | L | - | - | Y | I | L | K | I | - | - | - | - | - | - | - | - | - | - | - | - | - | - | - | - | - | - | - | - | - | - | - | - | - | - | - | - | - | - | - | - | - | - | L | E | - | - | - | - | - | - | - | - | - | - | - | - | - | - | - | - | - | - | - | - | - | - | - | - | - | - | - | - | - | - | - | - | - | - | - | - | - | - | - | - | - | - | - | - | - | - | - | - | - | - | - | - | M | L | K | D | T | - | - | Q | F | H | C | G | - | - | - | - | - | - | - | - | Q | G | T | E | L | E | G | - | - | Y | - | - | - | - | - | - | - | - | - | - | - | - | - | - | - | - | - | - | - | - | - | - | - | - | - | D | A | L | E | R | V | E | L | - | - | - | - | - | G | G | T | D | Q | - | - | - | G | S | R | H | D | I | L | S | R | N | M | T | I | - | - | - | - | - | - | - | - | - | - | - | - | - | - | - | - | - | - | - | - | - | - | - | - | - | P | H | P | - | - | - | - | - | - | - | - | A | P | R | - | V | Y | E | - | - | - | - | - | - | - | - | - | - | - | - | - | - | - | - | - | - | - | - | - | - | - | - | - | - | - | - | - | - | - | - | - | - | - | - | - | - | - | - | - | - | - | - | - | - | - | - | - | - | - | - | - | - | - | - | C | R | E | L | - | - | - | - | - | - | - | - | - | - | - | - | - | - | - | - | - | - | - | - | - | - | - | - | F | T | P | D | M | I | E | D | - | - | - | - | - | - | - | - | - | - | - | - | - | - | - | - | - | - | - | - | - | - | - | - | - | I | Q | R | A | G | - | - | - | - | - | - | - | - | S | G | S | R | R | D | Q | - | A | R | Q | L | I | L | D | L | E | T | R | G | - | S | Q | A | L | P | L | F | I | S | C | L | E | D | T | G | Q | D | T | L | A | S | L | L | K | T | S | R | Q | A | A | - | - | - | - | K | Q | D | V | - | E | A | I | - | - | - | - | - | - | - | - | - | - | - | - | - | - | - | - | - | - | - | - | - | - | - | - | - | - | R | P | L | D | - | - | - | - | - | - | - | - | - | - | - | - | - | - | - | L | K | P | V | - | - | - | - | - | - | - | - | - | - | - | - | - | - | - | - | - | - | - | - | - | - | - | - | - | - | - | - | - | - | - | - | - | - | - | - | - | - | - | V | L | G | P | E | G | L | K | P | G | E | L | R | - | V | M | K | Q | D | P | S | K | L | - | S | - | Q | - | - | - | - | - | - | - | - | - | - | - | - | - | - | - | - | - | - | - | - | - | - | - | - | - | - | - | - | - | - | - | - | - | - | - | - | - | - | - | - | - | - | - | - | - | - | - | - | - | - | - | G | K | L | A | P | V | - | - | - | - | - | - | - | - | - | - | - | - | - | - | - | - | - | - | - | - | - | - | - | - | - | - | V | L | - | - | - | - | - | - | - | - | - | - | - | - | - | - | - | - | - | - | - | - | - | - | - | - | - | - | - | - | - | - | - | - | - | - | - | - | - | - | - | - | - | - | - | - | - | - | - | - | - | - | - | - | - | - | - | - | - | - | - | - | - | - | - | - | - | - | - | - | G | P | E | E | L | W | P | - | - | - | - | - | - | - | - | - | - | - | - | - | - | - | - | - | - | - | - | - | - | - | - | - | - | P | K | L | R | P | - | - | - | - | - | - | E | V | L | R | P | - | E | V | - | - | - | - | - | - | - | - | - | - | - | - | - | - | - | - | - | - | P | R | A | V | D | A | G | S | G | G | F | - | - | - | - | - | - | - | - | - | - | - | - | - | T | D | V | C | - | - | - | - | - | - | - | - | - | - | P | - | Q | D | R | A | K | G | N | A | D | - | - | - | - | - | L | A | Y | V | - | - | L | N | A | D | - | P | C | G | H | C | L | I | I | N | N | V | N | F | C | R | E | S | - | - | G | L | R | A | R | T | G | S | N | I | D | C | E | R | M | R | R | R | F | - | H | L | L | Q | F | V | V | E | V | K | C | D | L | T | A | K | Q | M | V | Q | A | L | M | Q | L | A | R | Q | D | H | - | S | A | V | D | C | C | V | V | V | I | L | S | H | G | C | Q | A | S | H | L | R | F | P | G | A | - | V | Y | G | T | D | G | C | P | V | S | V | E | R | I | V | N | T | F | N | G | T | G | C | P | S | L | R | G | K | P | K | L | F | F | I | Q | A | C | - | - | G | G | E | Q | K | D | H | G | F | E | V | E | S | T | - | - | - | S | P | E | D | - | K | T | P | S | S | D | - | - | - | - | - | - | - | P | E | A | D | A | T | P | F | Q | E | G | P | R | S | I | - | - | - | - | - | - | - | - | D | E | P | D | A | V | - | - | - | - | - | - | - | S | S | L | - | - | - | P | T | P | S | D | I | L | V | S | Y | S | T | F | P | - | - | G | - | - | - | - | - | - | - | - | F | V | S | W | R | D | P | K | S | G | S | W | Y | I | E | T | L | D | S | I | F | E | Q | W | A | H | - | C | - | - | - | - | - | - | - | - | - | - | - | - | - | - | - | - | - | - | - | - | - | - | - | - | - | - | - | - | - | - | - | E | D | L | Q | T | L | L | - | - | - | - | - | L | R | - | - | - | - | - | - | - | - | - | - | - | - | V | A | N | A | V | - | - | - | - | - | S | V | K | - | - | - | - | - | - | - | G | I | Y | - | K | Q | M | - | - | P | G | - | - | - | - | - | - | - | - | - | - | - | - | - | - | - | C | F | N | F | - | - | - | - | - | - | - | - | - | L | R | K | K | L | F | F | K | T | - | - | - | - | - | - | - | - | - | - | - | - | - | - | - | - | - | - | - | - | - | - | - | - | - | - | - | - | - | - | - | - | - | - | - | - | - | - | M | - | - | - | - | - | - | - | - | - | - | - | - | - | - | - | - | - | - | - | - | - | - | - | - | - | - | - | - | - | - | - | - | - | - | - | - | - | - | - | - | - | - | - | - | - | - | - | - | - | - | - | - | G | D | V | E | K | G | K | K | I | F | V | Q | K | C | A | Q | C | H | T | V | E | K | G | G | K | H | K | T | G | P | N | L | H | G | L | F | G | R | K | S | G | Q | A | P | G | F | S | Y | T | D | A | N | K | N | K | G | I | T | W | G | E | E | T | L | M | E | Y | L | E | N | P | K | K | Y | I | P | G | T | K | M | I | F | A | G | I | K | K | K | G | E | R | E | D | L | I | A | Y | L | K | K | A | T | N | E | - | - | - | - | - | - |
| Bos\_taurus | M | - | - | - | - | - | - | - | - | - | - | - | - | - | - | - | - | - | - | - | - | - | - | - | - | - | - | - | - | - | - | - | - | - | - | - | - | - | - | - | - | - | - | - | - | - | - | - | - | - | - | - | - | - | - | - | - | - | - | - | - | - | - | - | - | - | - | - | - | - | - | - | - | - | - | - | - | - | - | - | - | - | - | - | - | - | - | - | - | - | - | - | - | - | - | - | - | - | D | A | K | A | - | R | N | - | - | - | - | - | - | - | - | - | - | - | - | - | - | - | - | C | L | L | - | - | - | - | - | - | - | - | - | - | - | - | - | - | - | - | - | - | - | - | - | - | - | - | - | - | - | - | - | - | - | - | - | - | - | - | - | - | - | - | - | - | - | - | - | - | - | - | - | - | - | - | - | - | - | - | - | - | - | - | - | - | - | - | - | - | - | - | - | - | - | - | - | - | - | - | - | - | - | - | - | - | - | - | - | - | - | - | - | - | - | - | - | - | - | - | - | - | - | - | - | - | - | - | - | - | - | - | - | - | - | - | - | - | - | - | - | - | - | - | - | Q | H | R | E | A | L | E | R | D | I | K | T | S | Y | - | I | M | D | H | M | I | S | N | G | V | L | T | V | S | E | E | E | K | V | K | N | E | - | - | - | - | - | - | - | - | - | - | - | - | - | - | - | - | - | P | T | Q | R | Q | R | A | A | M | L | I | K | M | I | L | K | K | - | - | - | - | D | N | H | S | Y | I | S | - | - | - | - | - | F | Y | N | A | L | - | L | H | E | G | - | - | - | - | - | - | Y | K | D | L | A | A | L | L | H | G | G | I | P | V | I | S | S | S | D | - | - | - | - | - | - | - | - | - | - | - | - | - | - | - | - | - | - | - | - | - | - | - | - | - | - | - | - | - | - | - | - | - | - | - | - | - | - | - | - | - | - | - | - | - | V | R | T | V | L | C | E | G | G | - | - | - | - | - | - | - | V | P | Q | R | P | V | V | F | V | T | R | K | K | L | V | N | A | I | - | - | - | - | - | - | - | - | - | - | Q | Q | K | L | S | - | - | - | - | - | - | - | - | - | - | - | - | - | - | - | - | - | - | - | - | - | - | - | - | - | - | - | - | - | - | - | - | K | L | N | G | E | P | - | - | - | - | - | - | - | - | - | - | - | - | - | - | - | - | - | - | - | - | - | - | - | - | - | - | - | - | - | - | - | - | - | - | - | - | - | - | G | W | V | T | I | Y | G | M | A | G | C | G | K | S | V | L | A | A | E | A | V | - | R | D | H | S | F | L | E | D | C | F | P | G | G | V | H | W | V | S | V | G | - | - | - | - | - | - | - | K | - | - | - | - | - | - | Q | D | K | S | - | - | - | - | - | - | - | - | - | - | - | - | G | L | L | M | K | L | Q | N | L | C | A | R | L | D | - | - | - | - | Q | - | - | - | - | - | - | - | - | - | - | - | - | - | - | - | D | E | S | F | S | Q | R | L | P | L | N | I | - | - | - | - | - | - | - | - | - | - | - | - | - | - | - | - | E | E | A | K | D | R | L | R | I | L | M | L | R | - | - | - | - | - | - | - | - | - | - | - | - | K | H | P | R | S | L | L | - | - | I | L | D | D | I | W | D | P | W | V | L | K | A | F | D | - | N | Q | C | Q | I | L | L | T | T | R | D | K | S | V | T | D | S | V | - | M | G | P | - | - | K | Y | V | V | P | V | E | S | - | - | - | G | L | G | K | E | K | G | L | E | I | L | S | L | F | V | N | M | - | K | K | A | D | L | P | E | - | - | - | - | - | - | - | - | - | - | - | - | - | - | Q | A | H | S | I | I | Q | E | C | K | - | - | - | - | - | - | - | - | - | - | - | - | - | - | - | - | - | - | - | G | - | S | P | L | V | V | S | L | I | G | A | L | L | R | D | F | P | N | - | - | - | - | - | - | - | - | - | - | R | W | E | Y | Y | L | - | R | Q | L | Q | N | K | Q | F | K | R | I | - | - | - | - | - | - | - | - | - | - | - | - | - | - | - | - | - | - | - | - | - | - | R | K | S | S | S | Y | D | - | - | - | - | Y | E | A | L | D | E | A | M | S | I | S | V | E | - | M | L | R | - | - | - | E | D | I | K | D | Y | - | Y | T | D | L | S | I | L | Q | K | D | V | K | V | P | T | K | V | L | C | I | L | W | D | - | - | - | - | L | E | T | E | E | V | E | D | I | L | Q | E | F | - | - | - | - | - | - | - | - | - | - | - | V | N | K | S | L | L | F | C | D | R | N | G | K | S | - | - | - | - | - | - | - | - | - | F | L | Y | Y | L | H | D | L | Q | V | D | F | L | A | E | K | - | - | N | R | S | Q | - | L | - | Q | D | L | H | K | K | M | V | T | Q | F | Q | R | - | H | H | - | - | - | - | - | - | Q | L | Y | S | L | S | P | D | - | Q | E | - | - | - | - | - | - | - | D | C | M | Y | W | Y | S | F | L | A | Y | H | M | A | S | A | N | - | - | - | - | M | H | K | E | L | C | A | L | M | F | S | L | D | W | I | K | A | K | T | E | L | V | G | P | A | - | - | - | - | - | - | - | - | H | L | I | H | E | F | V | E | Y | R | H | I | L | D | E | K | - | - | - | - | - | - | - | D | G | I | D | C | E | N | F | Q | - | - | - | E | F | L | S | L | N | G | H | L | L | G | R | Q | P | F | - | - | - | - | - | - | - | - | - | - | - | - | - | - | - | - | - | - | - | - | - | - | - | - | - | - | - | - | - | - | - | - | - | - | - | - | - | - | - | - | - | - | - | - | - | - | - | - | - | - | - | - | - | - | - | - | - | - | - | - | - | - | - | - | - | - | - | - | - | - | - | P | N | I | V | Q | L | G | L | C | E | L | E | T | S | E | V | Y | R | Q | A | K | L | Q | A | K | Q | E | V | D | N | G | - | - | - | - | - | - | - | - | - | - | - | - | - | - | - | - | - | - | - | - | - | - | - | - | - | - | - | - | - | M | L | Y | L | - | - | - | - | - | - | - | - | - | - | E | W | V | N | K | - | - | - | - | - | - | - | K | N | M | K | N | L | S | R | - | - | - | - | - | - | - | - | - | - | L | V | V | R | P | H | T | D | - | - | - | - | - | - | - | - | - | - | - | - | - | - | - | A | V | Y | H | A | C | - | - | - | F | S | - | E | D | G | Q | R | I | A | - | - | - | - | - | - | - | - | - | S | C | G | A | D | K | T | L | Q | V | F | K | - | - | - | - | - | A | E | T | G | E | K | L | L | E | I | - | K | A | H | E | D | E | - | - | - | V | L | C | C | A | F | S | - | A | - | - | - | - | - | - | - | - | - | - | - | - | - | D | D | K | F | I | A | T | C | S | V | D | K | K | V | K | I | - | - | - | W | N | S | M | - | - | - | - | - | - | - | - | - | - | - | - | - | - | - | - | - | - | - | - | - | - | - | - | - | - | - | - | - | - | - | - | - | - | - | - | - | - | - | - | - | - | - | - | - | - | - | - | - | - | - | - | - | - | - | - | - | - | - | - | - | - | - | - | - | - | - | - | - | - | - | - | - | - | - | - | - | - | - | - | - | - | - | - | - | - | - | - | - | - | - | - | - | - | - | - | - | - | - | - | - | - | - | - | - | - | - | - | - | - | - | - | - | - | - | - | - | - | - | - | - | - | - | - | - | - | - | - | - | - | - | - | - | - | - | - | - | - | - | - | T | G | E | L | V | R | I | Y | - | D | E | H | S | - | - | - | - | - | E | Q | V | N | - | - | - | - | - | - | - | - | - | - | - | - | - | C | C | H | - | - | - | - | - | - | - | - | - | - | - | - | - | F | T | N | N | S | - | - | - | - | - | - | - | - | - | - | - | - | - | - | - | - | N | H | - | L | L | L | A | T | G | S | N | - | - | - | - | D | Y | F | L | - | - | - | - | - | - | - | - | - | - | - | - | K | L | W | D | L | - | - | - | N | Q | E | E | C | R | N | T | M | F | - | - | - | - | - | - | - | - | - | - | - | - | - | - | - | - | - | G | H | T | N | S | - | - | - | - | - | - | - | - | - | - | - | - | - | - | - | - | V | S | H | C | R | F | S | P | D | D | Q | - | - | - | - | - | - | - | - | - | - | - | - | - | - | - | - | - | - | - | - | - | - | - | - | - | - | - | - | V | L | A | S | C | S | A | D | G | T | L | K | L | - | - | - | - | - | - | - | - | - | - | - | - | - | - | - | - | - | - | - | W | D | - | V | K | S | A | N | E | R | K | S | - | - | - | - | - | - | I | N | - | - | - | - | - | - | - | - | - | - | - | - | - | - | - | - | - | - | - | - | - | - | V | K | Q | F | - | - | - | - | - | - | - | - | F | L | N | S | E | E | P | Q | E | - | - | - | - | - | - | - | - | - | - | - | - | - | - | - | - | - | - | - | - | - | - | - | - | - | - | - | D | L | E | - | V | I | V | K | C | C | - | - | - | - | - | - | - | - | - | - | - | - | - | - | - | - | S | W | S | A | D | G | A | K | I | M | V | A | A | K | N | K | V | - | - | - | - | - | - | - | - | - | - | - | F | L | F | D | I | H | S | - | - | - | S | G | - | - | - | - | - | - | - | - | - | - | - | - | - | - | - | - | - | - | - | - | - | - | - | - | - | - | - | - | - | - | - | - | - | - | - | - | - | - | - | - | - | - | - | - | - | - | - | - | - | - | - | - | - | - | - | - | - | - | - | - | - | - | - | - | - | - | - | - | - | - | - | - | - | - | - | - | - | - | - | - | - | - | - | - | - | - | - | - | - | - | - | - | - | - | - | - | - | - | - | - | - | - | - | - | - | - | - | - | - | - | - | - | - | - | - | - | - | - | - | - | - | - | - | - | - | - | L | L | A | - | - | - | - | - | - | - | - | - | - | - | - | - | - | - | - | - | - | - | - | - | - | - | - | - | - | - | - | - | - | - | - | - | - | - | - | - | - | - | - | - | - | - | - | - | - | - | - | - | - | - | - | - | - | - | - | - | - | - | - | - | - | - | - | - | - | - | - | - | - | - | - | - | - | - | - | - | - | - | - | - | - | - | - | - | - | - | - | - | - | - | - | - | - | - | - | - | - | - | - | - | - | - | - | - | - | - | - | - | - | - | - | - | - | - | - | - | - | - | - | - | - | - | - | - | - | - | - | - | - | - | - | - | - | - | - | - | - | - | - | - | - | - | - | - | - | - | - | - | - | - | - | - | - | - | - | - | - | - | - | - | - | - | - | - | - | - | - | - | - | - | - | - | - | - | - | - | - | - | - | - | - | - | E | M | - | - | - | H | M | G | H | H | S | A | V | Q | Y | C | - | - | - | - | - | - | - | - | - | - | - | - | - | - | - | - | - | - | - | - | - | - | - | - | - | D | F | S | P | H | N | H | L | A | V | V | A | L | S | Q | - | - | - | - | - | - | Y | C | V | E | - | - | - | - | - | - | - | - | - | - | - | - | - | - | - | - | - | L | W | S | M | D | S | G | - | - | - | - | - | - | - | - | - | - | - | - | - | - | - | - | - | - | - | - | - | - | - | - | - | - | - | - | - | L | K | V | A | D | C | R | G | H | L | S | - | - | - | - | - | - | - | - | - | - | - | - | - | - | - | - | - | - | - | - | - | - | - | - | - | - | - | - | - | - | - | - | - | - | - | - | - | - | - | - | - | - | - | - | W | V | H | C | V | M | F | S | - | - | - | - | - | - | - | - | - | - | - | P | D | G | S | - | - | - | - | - | - | - | - | - | - | S | F | L | T | S | S | D | D | Q | - | - | - | - | - | - | - | - | - | - | - | - | - | - | T | I | R | L | - | W | - | - | - | - | - | - | - | - | - | - | - | - | - | - | - | - | - | - | - | - | - | - | - | - | - | - | - | - | - | - | - | - | - | - | - | - | - | - | E | T | K | K | V | C | K | N | S | A | T | V | L | K | Q | E | - | - | - | - | - | - | - | - | - | - | - | - | - | - | - | - | I | D | V | V | F | - | - | - | - | - | - | - | - | - | - | - | - | - | - | - | - | - | - | - | - | - | - | - | - | - | - | - | - | - | - | - | - | - | - | - | - | - | - | - | - | - | - | - | - | - | - | - | - | - | - | - | - | - | - | Q | E | N | E | V | T | - | - | - | - | - | - | - | - | - | - | - | - | - | - | - | - | - | - | - | - | - | - | - | - | - | - | - | - | - | - | - | - | - | - | - | - | V | L | A | V | D | N | I | R | R | L | Q | L | - | I | N | G | N | T | - | - | - | - | - | - | - | - | - | - | G | Q | I | D | Y | L | T | E | A | - | - | - | - | - | - | - | - | - | - | - | - | - | - | - | - | - | - | - | - | - | - | Q | - | - | - | - | - | - | - | - | - | - | - | V | S | C | C | C | L | S | P | H | L | Q | - | - | - | - | - | - | - | - | - | Y | L | A | F | G | G | - | - | - | - | - | - | - | - | - | - | - | - | - | - | - | - | - | - | - | - | - | - | - | - | - | - | - | - | - | - | - | - | - | - | - | - | E | - | - | - | - | - | - | - | - | - | D | G | A | - | - | - | - | - | - | - | - | - | - | I | - | - | - | - | - | - | - | - | - | - | - | - | - | - | - | - | - | - | - | - | - | - | - | - | - | - | - | - | - | - | - | - | - | - | - | - | - | - | - | Q | I | L | E | L | L | N | N | R | I | F | Q | S | R | N | G | - | - | - | - | - | H | - | - | - | - | - | - | - | - | - | - | - | - | K | T | T | V | R | H | I | Q | F | T | - | - | - | - | - | - | - | - | - | - | - | - | - | - | - | - | - | - | - | - | - | - | - | - | - | - | - | - | - | - | - | - | - | - | - | - | - | A | D | G | K | T | L | I | S | S | S | - | D | D | S | - | - | - | - | - | - | - | - | A | V | Q | - | I | - | - | - | - | - | - | - | - | - | - | - | - | - | - | - | - | - | - | - | - | - | - | - | - | - | - | - | - | - | - | - | - | - | - | - | - | - | - | - | W | N | - | - | - | - | - | - | - | - | - | - | W | Q | S | E | D | P | F | F | - | - | - | - | - | - | - | - | - | - | - | - | - | - | - | - | - | - | - | - | - | - | - | - | - | - | - | - | - | - | - | - | - | - | - | - | - | - | - | - | - | - | - | - | - | - | - | - | - | - | L | Q | A | H | Q | - | - | - | - | - | - | - | - | - | - | - | - | - | - | - | - | - | - | - | - | - | - | - | - | - | - | E | T | V | K | D | F | R | L | L | - | - | - | - | - | - | - | - | - | - | - | - | - | - | - | - | - | - | - | - | - | - | - | K | N | - | S | R | L | L | - | S | W | - | - | - | S | F | D | G | T | V | - | - | - | - | - | - | - | - | - | - | - | - | - | - | - | - | - | - | - | - | - | - | - | - | - | - | - | - | - | - | - | - | - | - | - | K | V | W | N | I | I | T | R | R | I | - | - | - | - | - | E | K | D | F | V | C | - | - | - | - | - | - | - | H | Q | - | - | - | - | - | - | - | - | - | - | - | - | - | - | - | - | - | - | - | D | T | V | L | - | - | - | - | - | - | - | - | - | - | - | - | - | - | - | S | C | D | - | - | - | - | - | - | - | - | - | - | - | - | - | - | - | I | S | S | D | A | T | - | K | F | S | S | T | - | S | A | D | K | T | - | - | - | - | - | - | A | K | I | W | S | F | E | L | L | - | - | - | - | - | - | - | - | - | - | - | - | - | - | - | - | - | - | - | - | - | - | - | - | S | P | L | H | E | - | - | - | - | - | - | - | - | L | - | - | - | - | R | G | H | K | G | C | V | - | - | - | - | - | - | - | - | - | R | C | S | - | S | F | S | - | - | - | - | - | - | - | - | - | V | N | N | T | - | L | L | A | T | - | - | - | - | - | - | - | - | G | D | D | N | G | E | I | R | I | W | N | V | S | N | G | E | L | L | - | - | - | - | - | - | - | - | - | - | - | - | - | - | - | - | - | - | - | - | - | - | - | - | - | - | - | - | - | - | - | - | - | - | - | - | - | - | - | - | - | - | - | - | - | - | - | - | - | - | - | - | - | - | - | - | H | L | C | A | P | I | S | V | E | - | - | - | - | - | - | - | - | - | - | - | - | - | - | - | - | - | - | - | - | E | G | A | A | T | H | G | G | W | - | - | - | - | - | - | - | - | V | T | D | L | C | F | S | P | D | T | K | - | - | - | - | - | - | - | - | - | - | - | - | - | - | - | - | - | - | - | - | - | - | - | - | - | - | - | - | - | - | - | - | - | - | - | - | - | - | - | - | - | - | - | - | - | M | L | V | S | - | - | A | G | G | Y | L | K | - | - | - | - | - | - | - | - | W | W | N | V | V | T | - | - | - | - | - | - | - | - | - | - | - | - | - | - | - | - | - | - | - | - | - | - | - | - | - | - | - | - | - | - | - | - | - | - | - | - | - | - | - | - | - | - | - | - | - | - | - | - | - | - | - | - | - | - | - | - | - | - | - | - | - | - | - | - | - | - | - | - | - | - | - | - | - | - | - | - | - | - | - | - | - | - | - | - | - | - | - | - | - | - | - | - | - | - | - | - | - | - | - | - | - | - | - | - | - | - | - | - | - | - | - | - | - | - | - | - | - | - | - | - | - | - | - | - | - | - | - | - | - | - | - | - | - | - | - | - | - | - | - | - | - | - | - | - | - | - | - | - | - | - | - | - | - | - | - | - | - | - | - | - | - | - | - | - | - | - | - | - | - | - | - | - | - | - | - | - | - | - | - | - | - | - | - | - | - | - | - | - | - | - | - | - | - | - | - | - | - | - | - | - | - | - | - | - | - | - | - | - | - | - | - | - | - | - | - | - | - | - | - | - | - | - | - | - | - | - | - | - | - | - | - | - | - | - | - | - | - | - | - | - | - | - | - | - | - | - | - | - | - | - | - | - | - | - | - | - | - | - | - | - | - | - | - | - | - | - | - | G | E | S | S | Q | T | F | Y | T | - | - | - | - | - | N | G | - | - | - | T | N | L | K | K | I | - | - | - | H | V | S | P | - | - | - | - | D | F | K | T | - | - | - | - | - | - | - | - | - | - | - | - | - | - | - | - | - | - | - | - | - | - | - | - | - | - | Y | V | T | V | D | - | N | L | G | I | L | - | - | Y | I | L | K | I | - | - | - | - | - | - | - | - | - | - | - | - | - | - | - | - | - | - | - | - | - | - | - | - | - | - | - | - | - | - | - | - | - | - | L | E | - | - | - | - | - | - | - | - | - | - | - | - | - | - | - | - | - | - | - | - | - | - | - | - | - | - | - | - | - | - | - | - | - | - | - | - | - | - | - | - | - | - | - | - | - | - | - | - | - | - | - | - | M | - | - | - | - | - | - | - | - | - | - | - | - | - | - | - | - | - | - | - | - | - | - | - | - | - | - | - | - | - | - | - | - | - | - | - | - | - | - | - | - | - | - | - | - | - | - | - | - | - | - | - | - | - | - | N | E | A | D | R | L | - | - | - | - | - | - | - | - | - | - | - | - | - | - | - | - | - | - | - | - | L | L | R | R | - | - | - | - | - | - | - | - | - | - | - | - | - | - | - | - | - | - | - | - | - | - | - | - | - | - | - | - | - | - | Y | R | - | - | - | - | - | - | - | - | V | R | L | - | V | G | E | L | Q | V | A | S | L | W | - | - | - | - | - | - | - | - | - | - | - | - | - | - | - | - | - | - | - | - | - | - | - | - | - | - | - | - | - | - | - | - | - | - | - | - | - | - | - | - | - | - | - | - | - | D | A | L | L | S | R | E | L | - | - | - | - | - | - | - | - | - | - | - | - | - | - | - | - | - | - | - | - | - | - | - | - | F | T | P | D | M | I | E | D | - | - | - | - | - | - | - | - | - | - | - | - | - | - | - | - | - | - | - | - | - | - | - | - | - | I | Q | R | A | G | - | - | - | - | - | - | - | - | S | G | S | R | R | D | Q | - | A | R | Q | L | I | L | D | L | E | T | R | G | - | S | Q | A | L | P | L | F | I | S | C | L | E | D | T | G | Q | D | T | L | A | S | L | L | K | T | S | R | Q | A | A | - | - | - | - | K | Q | D | - | - | - | - | - | - | - | - | - | - | - | - | - | - | - | - | - | - | - | - | - | - | - | - | - | - | - | - | - | - | - | - | - | - | - | - | - | - | - | - | - | - | - | - | - | - | - | - | - | - | - | - | - | - | - | - | - | - | - | - | - | - | - | - | - | - | - | - | - | - | - | - | - | - | - | - | - | - | - | - | - | - | - | - | - | - | - | - | - | - | - | - | - | - | - | - | - | - | - | - | - | - | - | - | - | - | - | - | - | - | - | D | P | S | K | P | - | S | - | Q | - | - | - | - | - | - | - | - | - | - | - | - | - | - | - | - | - | - | - | - | - | - | - | - | - | - | - | - | - | - | - | - | - | - | - | - | - | - | - | - | - | - | - | - | - | - | - | - | - | - | - | G | K | L | A | P | V | - | - | - | - | - | - | - | - | - | - | - | - | - | - | - | - | - | - | - | - | - | - | - | - | - | - | V | L | - | - | - | - | - | - | - | - | - | - | - | - | - | - | - | - | - | - | - | - | - | - | - | - | - | - | - | - | - | - | - | - | - | - | - | - | - | - | - | - | - | - | - | - | - | - | - | - | - | - | - | - | - | - | - | - | - | - | - | - | - | - | - | - | - | - | - | - | G | P | E | E | L | W | P | - | - | - | - | - | - | - | - | - | - | - | - | - | - | - | - | - | - | - | - | - | - | - | - | - | - | A | K | L | R | P | - | - | - | - | - | - | E | V | L | R | P | - | E | V | - | - | - | - | - | - | - | - | - | - | - | - | - | - | - | - | - | - | P | R | A | V | D | A | G | S | G | G | F | - | - | - | - | - | - | - | - | - | - | - | - | - | T | D | V | C | - | - | - | - | - | - | - | - | - | - | P | - | Q | D | R | A | K | G | N | A | D | - | - | - | - | - | L | A | Y | V | - | - | L | N | A | D | - | P | C | G | H | C | L | I | I | N | N | V | N | F | C | R | E | S | - | - | G | L | R | A | R | T | G | S | N | I | D | C | E | R | M | R | Q | R | F | - | H | L | L | Q | F | V | V | E | V | K | C | D | L | T | A | K | Q | M | V | Q | A | L | M | Q | L | A | R | Q | D | H | - | S | A | L | D | C | C | V | V | V | I | L | S | H | G | C | Q | A | S | H | L | Q | F | P | G | A | - | V | Y | G | T | D | G | C | P | V | S | V | E | R | I | V | N | T | F | N | G | T | G | C | P | S | L | R | G | K | P | K | L | F | F | I | Q | A | C | - | - | G | G | E | Q | K | D | H | G | F | E | V | E | S | T | - | - | - | S | P | E | D | - | K | T | P | S | S | D | - | - | - | - | - | - | - | P | E | A | D | A | T | P | F | Q | E | G | P | R | S | I | - | - | - | - | - | - | - | - | D | E | P | D | A | V | - | - | - | - | - | - | - | S | S | L | - | - | - | P | T | P | S | D | I | L | V | S | Y | S | T | F | P | - | - | G | - | - | - | - | - | - | - | - | F | V | S | W | R | D | P | K | S | G | S | W | Y | I | E | T | L | D | S | I | F | E | Q | W | A | H | - | C | - | - | - | - | - | - | - | - | - | - | - | - | - | - | - | - | - | - | - | - | - | - | - | - | - | - | - | - | - | - | - | E | D | L | Q | T | L | L | - | - | - | - | - | L | R | - | - | - | - | - | - | - | - | - | - | - | - | V | A | N | A | V | - | - | - | - | - | S | V | K | - | - | - | - | - | - | - | G | I | Y | - | K | Q | M | - | - | P | G | - | - | - | - | - | - | - | - | - | - | - | - | - | - | - | C | F | N | F | - | - | - | - | - | - | - | - | - | L | R | K | K | L | F | F | K | T | - | - | - | - | - | - | - | - | - | - | - | - | - | - | - | - | - | - | - | - | - | - | - | - | - | - | - | - | - | - | - | - | - | - | - | - | - | - | M | - | - | - | - | - | - | - | - | - | - | - | - | - | - | - | - | - | - | - | - | - | - | - | - | - | - | - | - | - | - | - | - | - | - | - | - | - | - | - | - | - | - | - | - | - | - | - | - | - | - | - | - | G | D | V | E | K | G | K | K | I | F | V | Q | K | C | A | Q | C | H | T | V | E | K | G | G | K | H | K | T | G | P | N | L | H | G | L | F | G | R | K | T | G | Q | A | P | G | F | S | Y | T | D | A | N | K | N | K | G | I | T | W | G | E | E | T | L | M | E | Y | L | E | N | P | K | K | Y | I | P | G | T | K | M | I | F | A | G | I | K | K | K | G | E | R | E | D | L | I | A | Y | L | K | K | A | T | N | E | - | - | - | - | - | - |
| Calidris\_pugnax | M | - | - | - | - | - | - | - | - | - | - | - | - | - | - | - | - | - | - | - | - | - | - | - | - | - | - | - | - | - | - | - | - | - | - | - | - | - | - | - | - | - | - | - | - | - | - | - | - | - | - | - | - | - | - | - | - | - | - | - | - | - | - | - | - | - | - | - | - | - | - | - | - | - | - | - | - | - | - | - | - | - | - | - | - | - | - | - | - | - | - | - | - | - | - | - | - | - | D | V | K | S | - | R | N | - | - | - | - | - | - | - | - | - | - | - | - | - | - | - | - | Y | L | L | - | - | - | - | - | - | - | - | - | - | - | - | - | - | - | - | - | - | - | - | - | - | - | - | - | - | - | - | - | - | - | - | - | - | - | - | - | - | - | - | - | - | - | - | - | - | - | - | - | - | - | - | - | - | - | - | - | - | - | - | - | - | - | - | - | - | - | - | - | - | - | - | - | - | - | - | - | - | - | - | - | - | - | - | - | - | - | - | - | - | - | - | - | - | - | - | - | - | - | - | - | - | - | - | - | - | - | - | - | - | - | - | - | - | - | - | - | - | - | - | - | M | N | R | Q | A | L | E | K | D | I | K | T | S | Y | - | I | M | D | R | M | I | S | D | E | V | L | T | L | Q | E | E | E | K | V | R | Q | Q | - | - | - | - | - | - | - | - | - | - | - | - | - | - | - | - | - | S | T | R | Q | E | R | A | A | M | L | I | N | I | I | L | T | K | - | - | - | - | D | N | N | S | Y | R | S | - | - | - | - | - | F | Y | N | A | L | - | L | H | E | G | - | - | - | - | - | - | Y | R | D | L | A | A | L | L | Q | D | G | I | P | S | I | S | P | G | N | G | - | K | - | - | - | - | - | - | - | - | - | - | - | - | - | - | - | - | - | - | - | - | - | - | S | S | M | D | G | M | T | - | - | - | - | - | - | - | - | - | - | - | S | Y | V | K | T | I | L | C | E | G | G | - | - | - | - | - | - | - | V | P | Q | R | P | V | V | F | V | T | R | P | K | L | V | D | A | I | - | - | - | - | - | - | - | - | - | - | K | Q | K | L | C | - | - | - | - | - | - | - | - | - | - | - | - | - | - | - | - | - | - | - | - | - | - | - | - | - | - | - | - | - | - | - | - | C | L | G | S | D | P | - | - | - | - | - | - | - | - | - | - | - | - | - | - | - | - | - | - | - | - | - | - | - | - | - | - | - | - | - | - | - | - | - | - | - | - | - | - | G | W | V | T | V | Y | G | M | A | G | C | G | K | T | V | L | T | A | E | A | L | - | R | D | H | Q | L | L | E | D | Y | F | P | G | G | V | H | W | I | S | V | G | - | - | - | - | - | - | - | K | - | - | - | - | - | - | Q | D | K | A | - | - | - | - | - | - | - | - | - | - | - | - | G | L | L | I | K | L | Q | N | L | C | S | R | L | E | - | - | - | - | Q | - | - | - | - | - | - | - | - | - | - | - | - | - | - | - | D | S | T | L | S | Q | R | P | P | L | N | I | - | - | - | - | - | - | - | - | - | - | - | - | - | - | - | - | E | E | A | K | D | R | L | R | L | L | M | L | R | - | - | - | - | - | - | - | - | - | - | - | - | K | Y | P | R | S | L | L | - | - | I | L | D | D | I | W | D | S | W | V | L | K | A | F | D | - | N | Q | C | Q | V | L | I | T | S | R | D | R | S | V | T | D | A | V | - | A | G | N | - | - | K | Y | E | V | H | V | E | S | - | - | - | G | L | A | H | E | K | G | L | E | I | L | S | L | F | V | N | T | - | K | I | S | E | L | P | E | - | - | - | - | - | - | - | - | - | - | - | - | - | - | Q | A | N | C | L | V | R | E | C | K | - | - | - | - | - | - | - | - | - | - | - | - | - | - | - | - | - | - | - | G | - | S | P | L | V | I | S | L | I | G | A | L | L | R | D | F | P | S | - | - | - | - | - | - | - | - | - | - | R | W | E | Y | Y | L | - | K | Q | L | Q | N | K | Q | F | K | R | I | - | - | - | - | - | - | - | - | - | - | - | - | - | - | - | - | - | - | - | - | - | - | R | K | S | S | S | Y | D | - | - | - | - | Y | E | A | L | D | E | A | M | S | I | S | V | E | - | Q | L | N | - | - | - | D | N | Y | K | D | Y | - | Y | K | D | L | S | I | L | P | K | D | V | K | V | P | T | K | V | L | C | I | L | W | D | - | - | - | - | M | E | T | E | E | V | E | D | I | L | Q | E | F | - | - | - | - | - | - | - | - | - | - | - | V | N | K | S | L | L | F | C | D | R | N | G | K | S | - | - | - | - | - | - | - | - | - | F | H | Y | Y | L | H | D | L | Q | L | D | F | L | T | E | K | - | - | N | R | N | Q | - | L | - | Q | E | L | H | K | N | I | V | N | Q | Y | K | K | - | Y | Y | - | - | - | - | - | - | K | L | N | T | P | V | P | S | - | Q | E | - | - | - | - | - | - | - | D | C | M | Y | W | Y | N | F | L | A | Y | H | M | A | G | A | N | - | - | - | - | M | Q | K | E | L | C | D | L | M | F | S | L | D | W | I | K | A | K | T | E | L | V | G | P | A | - | - | - | - | - | - | - | - | H | L | I | H | E | Y | V | E | Y | S | S | I | L | D | Q | K | - | - | - | - | - | - | - | D | S | T | V | R | E | N | F | Q | - | - | - | E | F | L | S | L | N | G | H | L | L | G | R | Q | P | F | - | - | - | - | - | - | - | - | - | - | - | - | - | - | - | - | - | - | - | - | - | - | - | - | - | - | - | - | - | - | - | - | - | - | - | - | - | - | - | - | - | - | - | - | - | - | - | - | - | - | - | - | - | - | - | - | - | - | - | - | - | - | - | - | - | - | - | - | - | - | - | P | D | I | V | Q | L | G | L | C | Q | P | E | T | S | E | V | Y | Q | Q | A | K | L | H | A | Q | R | E | - | - | R | G | - | - | - | - | - | - | - | - | - | - | - | - | - | - | - | - | - | - | - | - | - | - | - | - | - | - | - | - | - | A | F | Y | L | - | - | - | - | - | - | - | - | - | - | E | W | I | N | K | - | - | - | - | - | - | - | K | S | L | K | N | L | Y | R | - | - | - | - | - | - | - | - | - | - | L | V | V | R | P | H | R | D | - | - | - | - | - | - | - | - | - | - | - | - | - | - | - | A | V | Y | H | A | C | - | - | - | F | S | - | K | D | R | Q | R | I | A | - | - | - | - | - | - | - | - | - | S | C | G | A | D | K | T | L | Q | V | F | K | - | - | - | - | - | T | E | S | G | E | R | L | L | E | I | - | S | A | H | D | D | E | - | - | - | I | L | C | C | A | F | S | - | A | - | - | - | - | - | - | - | - | - | - | - | - | - | D | G | E | F | V | A | T | C | S | A | D | K | K | V | K | I | - | - | - | W | N | S | R | - | - | - | - | - | - | - | - | - | - | - | - | - | - | - | - | - | - | - | - | - | - | - | - | - | - | - | - | - | - | - | - | - | - | - | - | - | - | - | - | - | - | - | - | - | - | - | - | - | - | - | - | - | - | - | - | - | - | - | - | - | - | - | - | - | - | - | - | - | - | - | - | - | - | - | - | - | - | - | - | - | - | - | - | - | - | - | - | - | - | - | - | - | - | - | - | - | - | - | - | - | - | - | - | - | - | - | - | - | - | - | - | - | - | - | - | - | - | - | - | - | - | - | - | - | - | - | - | - | - | - | - | - | - | - | - | - | - | - | - | T | G | Q | C | K | C | V | Y | - | E | E | H | S | - | - | - | - | - | E | Q | V | N | - | - | - | - | - | - | - | - | - | - | - | - | - | C | C | Q | - | - | - | - | - | - | - | - | - | - | - | - | - | F | N | N | G | S | - | - | - | - | - | - | - | - | - | - | - | - | - | - | - | - | G | Q | - | Y | L | L | A | T | C | S | N | - | - | - | - | D | T | F | I | - | - | - | - | - | - | - | - | - | - | - | - | K | L | W | D | L | - | - | - | N | K | K | Y | C | R | N | T | M | I | - | - | - | - | - | - | - | - | - | - | - | - | - | - | - | - | - | G | H | I | N | S | - | - | - | - | - | - | - | - | - | - | - | - | - | - | - | - | V | S | H | C | R | F | S | P | N | D | E | - | - | - | - | - | - | - | - | - | - | - | - | - | - | - | - | - | - | - | - | - | - | - | - | - | - | - | - | Y | I | A | S | C | S | T | D | G | T | V | K | L | - | - | - | - | - | - | - | - | - | - | - | - | - | - | - | - | - | - | - | W | E | - | T | R | S | A | N | E | L | K | S | - | - | - | - | - | - | I | E | - | - | - | - | - | - | - | - | - | - | - | - | - | - | - | - | - | - | - | - | - | - | I | K | E | F | - | - | - | - | - | - | - | - | F | E | N | A | D | E | Q | S | D | - | - | - | - | - | - | - | - | - | - | - | - | - | - | - | - | - | - | - | - | - | - | - | - | - | - | - | D | V | E | - | V | L | V | K | C | C | - | - | - | - | - | - | - | - | - | - | - | - | - | - | - | - | S | W | S | K | N | G | N | M | I | L | V | V | A | K | N | K | V | - | - | - | - | - | - | - | - | - | - | - | L | L | F | D | V | K | T | - | - | - | C | D | - | - | - | - | - | - | - | - | - | - | - | - | - | - | - | - | - | - | - | - | - | - | - | - | - | - | - | - | - | - | - | - | - | - | - | - | - | - | - | - | - | - | - | - | - | - | - | - | - | - | - | - | - | - | - | - | - | - | - | - | - | - | - | - | - | - | - | - | - | - | - | - | - | - | - | - | - | - | - | - | - | - | - | - | - | - | - | - | - | - | - | - | - | - | - | - | - | - | - | - | - | - | - | - | - | - | - | - | - | - | - | - | - | - | - | - | - | - | - | - | - | - | - | - | - | - | L | L | T | - | - | - | - | - | - | - | - | - | - | - | - | - | - | - | - | - | - | - | - | - | - | - | - | - | - | - | - | - | - | - | - | - | - | - | - | - | - | - | - | - | - | - | - | - | - | - | - | - | - | - | - | - | - | - | - | - | - | - | - | - | - | - | - | - | - | - | - | - | - | - | - | - | - | - | - | - | - | - | - | - | - | - | - | - | - | - | - | - | - | - | - | - | - | - | - | - | - | - | - | - | - | - | - | - | - | - | - | - | - | - | - | - | - | - | - | - | - | - | - | - | - | - | - | - | - | - | - | - | - | - | - | - | - | - | - | - | - | - | - | - | - | - | - | - | - | - | - | - | - | - | - | - | - | - | - | - | - | - | - | - | - | - | - | - | - | - | - | - | - | - | - | - | - | - | - | - | - | - | - | - | - | - | Q | V | - | - | - | I | I | S | H | H | S | T | I | Q | Y | C | - | - | - | - | - | - | - | - | - | - | - | - | - | - | - | - | - | - | - | - | - | - | - | - | - | D | F | C | P | G | D | E | L | V | A | V | A | L | S | H | - | - | - | - | - | - | C | S | V | E | - | - | - | - | - | - | - | - | - | - | - | - | - | - | - | - | - | L | W | N | I | K | S | L | - | - | - | - | - | - | - | - | - | - | - | - | - | - | - | - | - | - | - | - | - | - | - | - | - | - | - | - | - | S | K | V | A | Y | C | R | G | H | M | S | - | - | - | - | - | - | - | - | - | - | - | - | - | - | - | - | - | - | - | - | - | - | - | - | - | - | - | - | - | - | - | - | - | - | - | - | - | - | - | - | - | - | - | - | W | V | H | C | V | T | F | S | - | - | - | - | - | - | - | - | - | - | - | P | D | G | S | - | - | - | - | - | - | - | - | - | - | L | F | L | T | S | S | D | D | Q | - | - | - | - | - | - | - | - | - | - | - | - | - | - | T | V | R | I | - | W | - | - | - | - | - | - | - | - | - | - | - | - | - | - | - | - | - | - | - | - | - | - | - | - | - | - | - | - | - | - | - | - | - | - | - | - | - | - | E | T | R | K | V | C | Q | S | S | D | A | V | L | K | S | E | - | - | - | - | - | - | - | - | - | - | - | - | - | - | - | - | L | D | V | V | F | - | - | - | - | - | - | - | - | - | - | - | - | - | - | - | - | - | - | - | - | - | - | - | - | - | - | - | - | - | - | - | - | - | - | - | - | - | - | - | - | - | - | - | - | - | - | - | - | - | - | - | - | - | - | H | N | G | E | V | M | - | - | - | - | - | - | - | - | - | - | - | - | - | - | - | - | - | - | - | - | - | - | - | - | - | - | - | - | - | - | - | - | - | - | - | - | I | L | A | I | Y | N | Q | K | H | L | Q | L | - | I | N | G | N | T | - | - | - | - | - | - | - | - | - | - | D | S | V | I | M | Q | T | A | A | - | - | - | - | - | - | - | - | - | - | - | - | - | - | - | - | - | - | - | - | - | - | Q | E | S | - | - | - | - | - | - | - | - | P | I | S | C | C | C | L | S | E | D | L | K | - | - | - | - | - | - | - | - | - | F | A | A | F | G | Q | - | - | - | - | - | - | - | - | - | - | - | - | - | - | - | - | - | - | - | - | - | - | - | - | - | - | - | - | - | - | - | - | - | - | - | - | E | - | - | - | - | - | - | - | - | - | S | G | T | - | - | - | - | - | - | - | - | - | - | I | - | - | - | - | - | - | - | - | - | - | - | - | - | - | - | - | - | - | - | - | - | - | - | - | - | - | - | - | - | - | - | - | - | - | - | - | - | - | - | K | V | L | Q | V | L | D | G | K | V | L | N | S | R | E | A | - | - | - | - | - | F | - | - | - | - | - | - | - | - | - | - | - | - | R | T | S | V | Q | H | C | R | F | T | - | - | - | - | - | - | - | - | - | - | - | - | - | - | - | - | - | - | - | - | - | - | - | - | - | - | - | - | - | - | - | - | - | - | - | - | - | S | D | C | Q | T | L | I | S | S | A | - | H | D | S | - | - | - | - | - | - | - | - | V | I | Q | - | V | - | - | - | - | - | - | - | - | - | - | - | - | - | - | - | - | - | - | - | - | - | - | - | - | - | - | - | - | - | - | - | - | - | - | - | - | - | - | - | W | N | - | - | - | - | - | - | - | - | - | - | W | L | L | S | E | C | V | F | - | - | - | - | - | - | - | - | - | - | - | - | - | - | - | - | - | - | - | - | - | - | - | - | - | - | - | - | - | - | - | - | - | - | - | - | - | - | - | - | - | - | - | - | - | - | - | - | - | - | L | R | G | H | K | - | - | - | - | - | - | - | - | - | - | - | - | - | - | - | - | - | - | - | - | - | - | - | - | - | - | E | A | I | K | D | F | R | L | L | - | - | - | - | - | - | - | - | - | - | - | - | - | - | - | - | - | - | - | - | - | - | - | G | N | - | S | K | L | L | - | S | W | - | - | - | S | F | D | G | T | V | - | - | - | - | - | - | - | - | - | - | - | - | - | - | - | - | - | - | - | - | - | - | - | - | - | - | - | - | - | - | - | - | - | - | - | K | V | W | N | I | T | T | G | N | T | - | - | - | - | - | E | K | D | F | A | C | - | - | - | - | - | - | - | H | G | - | - | - | - | - | - | - | - | - | - | - | - | - | - | - | - | - | - | - | G | A | V | L | - | - | - | - | - | - | - | - | - | - | - | - | - | - | - | S | C | A | - | - | - | - | - | - | - | - | - | - | - | - | - | - | - | V | S | P | D | G | S | - | K | F | S | S | T | - | S | A | D | K | T | - | - | - | - | - | - | A | K | I | W | S | F | E | S | S | - | - | - | - | - | - | - | - | - | - | - | - | - | - | - | - | - | - | - | - | - | - | - | - | S | V | L | H | E | - | - | - | - | - | - | - | - | L | - | - | - | - | K | G | H | E | A | C | V | - | - | - | - | - | - | - | - | - | R | C | C | - | T | F | S | - | - | - | - | - | - | - | - | - | P | N | N | K | - | L | L | A | T | - | - | - | - | - | - | - | - | G | D | D | K | G | E | I | R | I | W | D | I | L | T | G | S | L | L | - | - | - | - | - | - | - | - | - | - | - | - | - | - | - | - | - | - | - | - | - | - | - | - | - | - | - | - | - | - | - | - | - | - | - | - | - | - | - | - | - | - | - | - | - | - | - | - | - | - | - | - | - | - | - | - | H | F | C | S | P | V | T | V | D | - | - | - | - | - | - | - | - | - | - | - | - | - | - | - | - | - | - | - | - | E | G | E | P | T | H | G | G | W | - | - | - | - | - | - | - | - | V | T | D | L | S | F | S | P | D | S | K | - | - | - | - | - | - | - | - | - | - | - | - | - | - | - | - | - | - | - | - | - | - | - | - | - | - | - | - | - | - | - | - | - | - | - | - | - | - | - | - | - | - | - | - | - | M | L | V | S | - | - | S | G | G | Y | L | K | - | - | - | - | - | - | - | - | W | W | N | V | T | T | - | - | - | - | - | - | - | - | - | - | - | - | - | - | - | - | - | - | - | - | - | - | - | - | - | - | - | - | - | - | - | - | - | - | - | - | - | - | - | - | - | - | - | - | - | - | - | - | - | - | - | - | - | - | - | - | - | - | - | - | - | - | - | - | - | - | - | - | - | - | - | - | - | - | - | - | - | - | - | - | - | - | - | - | - | - | - | - | - | - | - | - | - | - | - | - | - | - | - | - | - | - | - | - | - | - | - | - | - | - | - | - | - | - | - | - | - | - | - | - | - | - | - | - | - | - | - | - | - | - | - | - | - | - | - | - | - | - | - | - | - | - | - | - | - | - | - | - | - | - | - | - | - | - | - | - | - | - | - | - | - | - | - | - | - | - | - | - | - | - | - | - | - | - | - | - | - | - | - | - | - | - | - | - | - | - | - | - | - | - | - | - | - | - | - | - | - | - | - | - | - | - | - | - | - | - | - | - | - | - | - | - | - | - | - | - | - | - | - | - | - | - | - | - | - | - | - | - | - | - | - | - | - | - | - | - | - | - | - | - | - | - | - | - | - | - | - | - | - | - | - | - | - | - | - | - | - | - | - | - | - | - | - | - | - | - | - | G | E | S | L | Q | T | F | Y | T | - | - | - | - | - | N | G | - | - | - | T | N | L | K | S | I | - | - | - | H | V | S | P | - | - | - | - | N | F | K | V | - | - | - | - | - | - | - | - | - | - | - | - | - | - | - | - | - | - | - | - | - | - | - | - | - | - | Y | V | T | V | D | - | N | L | G | I | L | - | - | Y | V | L | Q | K | - | - | - | - | - | - | - | - | - | - | - | - | - | - | - | - | - | - | - | - | - | - | - | - | - | - | - | - | - | - | - | - | - | - | L | - | - | - | - | - | - | - | - | - | - | - | - | - | - | - | - | - | - | - | - | - | - | - | - | - | - | - | - | - | - | - | - | - | - | - | - | - | - | - | - | - | - | - | - | - | - | - | - | - | - | - | - | - | M | - | - | - | - | - | - | - | - | - | - | - | - | - | - | - | - | - | - | - | - | - | - | - | - | - | - | - | - | - | - | - | - | - | - | - | - | - | - | - | - | - | - | - | - | - | - | - | - | - | - | - | - | - | - | E | E | T | Q | R | R | - | - | - | - | - | - | - | - | - | - | - | - | - | - | - | - | - | - | - | - | A | L | R | R | - | - | - | - | - | - | - | - | - | - | - | - | - | - | - | - | - | - | - | - | - | - | - | - | - | - | - | - | - | - | G | R | - | - | - | - | - | - | - | - | A | R | L | - | V | A | E | L | R | V | A | P | L | W | - | - | - | - | - | - | - | - | - | - | - | - | - | - | - | - | - | - | - | - | - | - | - | - | - | - | - | - | - | - | - | - | - | - | - | - | - | - | - | - | - | - | - | - | - | D | P | L | E | D | R | G | L | - | - | - | - | - | - | - | - | - | - | - | - | - | - | - | - | - | - | - | - | - | - | - | - | F | T | R | T | M | V | E | E | - | - | - | - | - | - | - | - | - | - | - | - | - | - | - | - | - | - | - | - | - | - | - | - | - | L | Q | S | A | - | - | - | - | - | - | - | - | - | - | G | S | R | G | D | Q | - | A | R | Q | L | V | I | D | L | E | T | R | G | - | K | Q | A | F | P | L | F | L | S | I | L | R | D | T | G | Q | G | D | L | A | D | M | L | M | K | E | C | Q | C | P | - | - | - | - | L | V | P | P | - | Q | - | - | - | - | - | - | - | - | - | - | - | - | - | - | - | - | - | - | - | - | - | - | - | - | - | - | - | - | - | L | A | E | - | - | - | - | - | - | - | - | - | - | - | - | - | - | - | L | R | P | V | - | - | - | - | - | - | - | - | - | - | - | - | - | - | - | - | - | - | - | - | - | - | - | - | - | - | - | - | - | - | - | - | - | - | - | - | - | - | - | E | L | E | L | R | G | K | K | H | N | K | N | V | - | - | - | - | - | - | - | - | - | - | - | N | - | - | - | - | - | - | - | - | - | - | - | - | - | - | - | - | - | - | - | - | - | - | - | - | - | - | - | - | - | - | - | - | - | - | - | - | - | - | - | - | - | - | - | - | - | - | - | - | - | - | - | - | - | - | - | - | - | - | - | - | - | - | - | - | - | - | - | - | - | - | - | - | - | - | - | - | - | - | - | - | - | - | - | - | - | - | - | - | - | - | - | - | - | - | - | - | - | - | - | - | - | - | - | - | - | - | - | - | - | - | - | - | - | - | - | - | - | - | - | - | - | - | - | - | - | - | - | - | - | - | - | - | - | - | - | - | - | - | - | - | - | - | - | - | - | - | - | - | - | - | - | - | - | - | - | - | L | P | E | R | L | S | - | - | - | - | - | - | - | - | - | - | - | - | - | - | - | - | - | - | - | - | - | - | - | - | - | - | I | P | V | Q | V | - | - | - | - | - | - | E | S | E | R | P | - | R | T | - | - | - | - | - | - | - | - | - | - | - | - | - | - | - | - | - | - | P | P | A | P | A | L | G | - | - | - | - | - | - | - | - | - | - | - | - | - | - | - | - | - | - | - | - | L | - | - | - | - | - | - | - | - | - | - | A | - | V | E | - | - | K | R | N | R | D | - | - | - | - | - | L | V | Y | E | - | - | L | K | A | D | - | P | C | G | H | C | L | I | L | N | N | V | D | F | S | R | D | S | - | - | D | L | S | T | R | V | G | S | N | V | D | C | K | K | L | E | K | R | F | - | K | S | L | S | F | D | V | L | T | R | Q | N | L | K | A | Q | E | M | V | S | E | L | Q | K | L | A | Q | Q | D | H | - | G | A | L | D | C | C | I | V | V | I | L | S | H | G | C | Q | T | S | H | N | Q | F | P | G | G | - | V | Y | G | T | D | G | K | P | I | P | I | E | K | I | V | N | Y | F | N | G | S | N | C | P | S | L | R | G | K | P | K | L | F | F | I | Q | A | C | - | - | G | G | E | Q | R | D | R | G | F | V | V | D | C | D | - | - | - | S | P | G | D | - | E | A | P | G | G | A | - | - | - | - | - | - | - | L | E | S | D | A | T | P | F | R | A | P | V | G | G | T | - | - | - | - | - | - | - | - | D | E | P | D | A | V | - | - | - | - | - | - | - | A | S | L | - | - | - | P | T | P | S | D | I | L | V | S | Y | S | T | F | P | - | - | G | - | - | - | - | - | - | - | - | F | V | S | W | R | D | A | S | S | G | S | W | Y | V | E | T | L | D | R | V | L | E | Q | Y | A | H | - | T | - | - | - | - | - | - | - | - | - | - | - | - | - | - | - | - | - | - | - | - | - | - | - | - | - | - | - | - | - | - | - | E | D | L | L | S | M | L | - | - | - | - | - | L | R | - | - | - | - | - | - | - | - | - | - | - | - | V | A | D | A | V | - | - | - | - | - | S | A | K | - | - | - | - | - | - | - | G | R | Y | - | K | Q | I | - | - | P | G | - | - | - | - | - | - | - | - | - | - | - | - | - | - | - | C | F | N | F | - | - | - | - | - | - | - | - | - | L | R | K | K | F | F | F | T | C | K | - | - | - | - | - | - | - | - | - | - | - | - | - | - | - | - | - | - | - | - | - | - | - | - | - | - | - | - | - | - | - | - | - | - | - | - | - | M | - | - | - | - | - | - | - | - | - | - | - | - | - | - | - | - | - | - | - | - | - | - | - | - | - | - | - | - | - | - | - | - | - | - | - | - | - | - | - | - | - | - | - | - | - | - | - | - | - | - | - | - | G | D | I | E | K | G | K | K | I | F | V | Q | K | C | S | Q | C | H | T | V | E | K | G | G | K | H | K | T | G | P | N | L | N | G | L | F | G | R | K | T | G | Q | A | A | G | F | S | Y | T | D | A | N | K | N | K | G | I | T | W | G | E | D | T | L | M | E | Y | L | E | N | P | K | K | Y | I | P | G | T | K | M | I | F | A | G | I | K | K | K | S | E | R | A | D | L | I | A | Y | L | K | D | A | T | S | K | - | - | - | - | - | - |
| Callorhinchus\_milii | M | - | - | - | - | - | - | - | - | - | - | - | - | - | - | - | - | - | - | - | - | - | - | - | - | - | - | - | - | - | - | - | - | - | - | - | - | - | - | - | - | - | - | - | - | - | - | - | - | - | - | - | - | - | - | - | - | - | - | - | - | - | - | - | - | - | - | - | - | - | - | - | - | - | - | - | - | - | - | - | - | - | - | - | - | - | - | - | - | - | - | - | - | - | - | - | - | - | E | E | R | T | - | R | S | - | - | - | - | - | - | - | - | - | - | - | - | - | - | - | - | C | L | L | - | - | - | - | - | - | - | - | - | - | - | - | - | - | - | - | - | - | - | - | - | - | - | - | - | - | - | - | - | - | - | - | - | - | - | - | - | - | - | - | - | - | - | - | - | - | - | - | - | - | - | - | - | - | - | - | - | - | - | - | - | - | - | - | - | - | - | - | - | - | - | - | - | - | - | - | - | - | - | - | - | - | - | - | - | - | - | - | - | - | - | - | - | - | - | - | - | - | - | - | - | - | - | - | - | - | - | - | - | - | - | - | - | - | - | - | - | - | - | - | - | R | H | R | S | A | L | E | Q | D | I | K | T | S | Y | - | I | M | D | H | L | I | S | E | D | V | L | T | V | V | D | E | Q | K | V | K | A | K | - | - | - | - | - | - | - | - | - | - | - | - | - | - | - | - | - | T | T | Q | R | E | R | A | A | L | L | L | D | M | V | L | E | K | - | - | - | - | D | N | Y | G | F | M | S | - | - | - | - | - | F | Y | N | A | L | - | L | N | E | G | - | - | - | - | - | - | Y | K | D | L | A | A | L | L | Q | D | D | I | P | V | T | S | P | P | T | I | - | K | - | - | - | - | - | - | - | - | - | - | - | - | - | - | - | - | - | - | - | - | - | - | S | F | V | D | G | V | T | - | - | - | - | - | - | - | - | - | - | - | P | Y | V | Q | T | M | L | C | E | G | G | - | - | - | - | - | - | - | V | P | Q | R | P | V | V | F | V | D | R | P | K | L | V | Q | T | I | - | - | - | - | - | - | - | - | - | - | R | K | E | L | I | - | - | - | - | - | - | - | - | - | - | - | - | - | - | - | - | - | - | - | - | - | - | - | - | - | - | - | - | - | - | - | - | K | L | K | Q | G | P | - | - | - | - | - | - | - | - | - | - | - | - | - | - | - | - | - | - | - | - | - | - | - | - | - | - | - | - | - | - | - | - | - | - | - | - | - | - | G | W | I | T | I | H | G | M | A | G | S | G | K | S | V | L | A | A | E | A | L | - | R | N | H | S | V | I | D | D | C | F | P | G | G | V | H | W | V | S | I | G | - | - | - | - | - | - | - | K | - | - | - | - | - | - | Q | D | R | V | - | - | - | - | - | - | - | - | - | - | - | - | G | L | L | V | K | L | Q | N | L | C | S | R | L | D | - | - | - | - | Q | - | - | - | - | - | - | - | - | - | - | - | - | - | - | - | E | S | Q | I | Y | P | N | P | P | L | N | M | - | - | - | - | - | - | - | - | - | - | - | - | - | - | - | - | D | E | A | R | D | R | L | R | F | L | M | A | R | - | - | - | - | - | - | - | - | - | - | - | - | K | H | P | R | S | L | L | - | - | I | L | D | D | V | W | E | S | W | V | L | K | C | F | D | - | I | H | S | R | V | L | I | T | S | R | D | K | S | V | T | D | T | V | - | A | G | N | - | - | K | S | L | V | P | V | Q | S | - | - | - | G | L | T | G | D | K | G | L | E | V | L | A | R | F | V | N | L | - | E | V | S | S | L | P | Q | - | - | - | - | - | - | - | - | - | - | - | - | - | - | E | A | Y | N | I | V | Q | E | C | K | - | - | - | - | - | - | - | - | - | - | - | - | - | - | - | - | - | - | - | G | - | S | P | L | V | V | S | L | I | G | A | L | L | R | E | F | P | N | - | - | - | - | - | - | - | - | - | - | R | W | E | Y | Y | L | - | K | Q | L | Q | N | K | Q | F | K | R | I | - | - | - | - | - | - | - | - | - | - | - | - | - | - | - | - | - | - | - | - | - | - | R | K | S | S | S | Y | D | - | - | - | - | Y | E | A | L | D | E | A | M | A | I | S | I | D | - | V | L | E | - | - | - | E | Y | Q | K | E | H | - | Y | K | D | L | S | V | L | K | K | D | V | K | V | P | T | K | V | L | C | V | L | W | E | - | - | - | - | K | D | T | E | E | V | E | D | I | L | Q | L | F | - | - | - | - | - | - | - | - | - | - | - | V | N | K | S | L | L | F | C | D | R | N | G | K | S | - | - | - | - | - | - | - | - | - | F | Q | Y | Y | L | H | D | L | Q | L | D | F | L | T | E | Q | - | - | N | R | N | R | - | L | - | P | E | L | H | A | K | M | V | R | Q | Y | Q | K | - | V | H | - | - | - | - | - | - | E | K | H | F | P | L | P | S | - | D | E | - | - | - | - | - | - | - | D | C | M | Y | W | Y | S | F | L | I | G | H | M | A | E | A | L | - | - | - | - | M | H | K | E | L | Y | L | L | M | F | S | M | D | W | I | K | A | K | T | E | L | M | G | P | A | - | - | - | - | - | - | - | - | P | L | I | N | D | F | V | E | Y | R | H | I | L | D | K | E | - | - | - | - | - | - | - | N | S | T | V | R | D | N | F | Q | - | - | - | E | F | L | S | L | K | G | H | L | L | V | Q | K | P | F | - | - | - | - | - | - | - | - | - | - | - | - | - | - | - | - | - | - | - | - | - | - | - | - | - | - | - | - | - | - | - | - | - | - | - | - | - | - | - | - | - | - | - | - | - | - | - | - | - | - | - | - | - | - | - | - | - | - | - | - | - | - | - | - | - | - | - | - | - | - | - | P | D | I | V | Q | L | G | L | C | Q | P | D | C | S | E | V | Y | R | Q | A | R | L | E | A | G | K | T | T | K | D | R | - | - | - | - | - | - | - | - | - | - | - | - | - | - | - | - | - | - | - | - | - | - | - | - | - | - | - | - | - | V | V | Y | L | - | - | - | - | - | - | - | - | - | - | E | W | M | N | K | - | - | - | - | - | - | - | - | N | L | Q | S | L | T | R | - | - | - | - | - | - | - | - | - | - | L | V | V | R | P | H | S | D | - | - | - | - | - | - | - | - | - | - | - | - | - | - | - | A | V | L | H | G | C | - | - | - | F | S | - | P | D | G | Q | R | I | A | - | - | - | - | - | - | - | - | - | T | C | G | V | D | K | N | F | Q | V | F | K | - | - | - | - | - | S | T | T | G | E | K | V | L | E | V | - | A | A | H | D | D | D | - | - | - | V | L | C | C | V | F | S | - | P | - | - | - | - | - | - | - | - | - | - | - | - | - | D | N | K | F | I | A | T | C | S | S | D | T | K | V | K | V | - | - | - | W | N | S | V | - | - | - | - | - | - | - | - | - | - | - | - | - | - | - | - | - | - | - | - | - | - | - | - | - | - | - | - | - | - | - | - | - | - | - | - | - | - | - | - | - | - | - | - | - | - | - | - | - | - | - | - | - | - | - | - | - | - | - | - | - | - | - | - | - | - | - | - | - | - | - | - | - | - | - | - | - | - | - | - | - | - | - | - | - | - | - | - | - | - | - | - | - | - | - | - | - | - | - | - | - | - | - | - | - | - | - | - | - | - | - | - | - | - | - | - | - | - | - | - | - | - | - | - | - | - | - | - | - | - | - | - | - | - | - | - | - | - | - | - | T | G | E | L | L | Q | T | Y | - | S | D | H | T | - | - | - | - | - | E | Q | V | N | - | - | - | - | - | - | - | - | - | - | - | - | - | H | C | Q | - | - | - | - | - | - | - | - | - | - | - | - | - | F | S | N | L | V | - | - | - | - | - | - | - | - | - | - | - | - | - | - | - | - | Q | T | - | N | L | L | A | T | C | S | N | - | - | - | - | D | T | Y | L | - | - | - | - | - | - | - | - | - | - | - | - | K | T | W | D | L | - | - | - | K | G | K | S | C | K | N | T | L | F | - | - | - | - | - | - | - | - | - | - | - | - | - | - | - | - | - | G | H | D | N | S | - | - | - | - | - | - | - | - | - | - | - | - | - | - | - | - | V | N | H | C | N | F | S | P | D | D | K | - | - | - | - | - | - | - | - | - | - | - | - | - | - | - | - | - | - | - | - | - | - | - | - | - | - | - | - | Y | L | A | S | C | S | N | D | G | T | V | K | L | - | - | - | - | - | - | - | - | - | - | - | - | - | - | - | - | - | - | - | W | D | - | V | S | S | A | N | E | W | K | F | - | - | - | - | - | - | F | D | - | - | - | - | - | - | - | - | - | - | - | - | - | - | - | - | - | - | - | - | - | - | V | K | N | E | - | - | - | - | - | - | - | - | F | M | N | T | D | D | N | Q | E | - | - | - | - | - | - | - | - | - | - | - | - | - | - | - | - | - | - | - | - | - | - | - | - | - | - | - | K | E | E | - | V | L | V | K | C | C | - | - | - | - | - | - | - | - | - | - | - | - | - | - | - | - | T | W | S | S | D | G | S | Q | L | I | A | A | T | K | N | I | L | - | - | - | - | - | - | - | - | - | - | - | F | I | F | N | V | A | T | - | - | - | G | D | - | - | - | - | - | - | - | - | - | - | - | - | - | - | - | - | - | - | - | - | - | - | - | - | - | - | - | - | - | - | - | - | - | - | - | - | - | - | - | - | - | - | - | - | - | - | - | - | - | - | - | - | - | - | - | - | - | - | - | - | - | - | - | - | - | - | - | - | - | - | - | - | - | - | - | - | - | - | - | - | - | - | - | - | - | - | - | - | - | - | - | - | - | - | - | - | - | - | - | - | - | - | - | - | - | - | - | - | - | - | - | - | - | - | - | - | - | - | - | - | - | - | - | - | - | - | I | E | A | - | - | - | - | - | - | - | - | - | - | - | - | - | - | - | - | - | - | - | - | - | - | - | - | - | - | - | - | - | - | - | - | - | - | - | - | - | - | - | - | - | - | - | - | - | - | - | - | - | - | - | - | - | - | - | - | - | - | - | - | - | - | - | - | - | - | - | - | - | - | - | - | - | - | - | - | - | - | - | - | - | - | - | - | - | - | - | - | - | - | - | - | - | - | - | - | - | - | - | - | - | - | - | - | - | - | - | - | - | - | - | - | - | - | - | - | - | - | - | - | - | - | - | - | - | - | - | - | - | - | - | - | - | - | - | - | - | - | - | - | - | - | - | - | - | - | - | - | - | - | - | - | - | - | - | - | - | - | - | - | - | - | - | - | - | - | - | - | - | - | - | - | - | - | - | - | - | - | - | - | - | - | - | K | L | - | - | - | E | S | R | C | H | S | T | I | Q | Y | C | - | - | - | - | - | - | - | - | - | - | - | - | - | - | - | - | - | - | - | - | - | - | - | - | - | H | A | S | P | I | S | S | L | V | A | V | A | L | S | H | - | - | - | - | - | - | Y | S | V | E | - | - | - | - | - | - | - | - | - | - | - | - | - | - | - | - | - | L | W | D | I | K | L | Q | - | - | - | - | - | - | - | - | - | - | - | - | - | - | - | - | - | - | - | - | - | - | - | - | - | - | - | - | - | K | R | V | A | D | C | S | G | H | L | S | - | - | - | - | - | - | - | - | - | - | - | - | - | - | - | - | - | - | - | - | - | - | - | - | - | - | - | - | - | - | - | - | - | - | - | - | - | - | - | - | - | - | - | - | W | V | H | C | V | Q | F | S | - | - | - | - | - | - | - | - | - | - | - | P | D | G | S | - | - | - | - | - | - | - | - | - | - | M | L | L | S | S | S | D | D | Q | - | - | - | - | - | - | - | - | - | - | - | - | - | - | T | V | R | L | - | W | - | - | - | - | - | - | - | - | - | - | - | - | - | - | - | - | - | - | - | - | - | - | - | - | - | - | - | - | - | - | - | - | - | - | - | - | - | - | E | T | N | K | V | H | H | T | S | S | I | S | L | M | Q | V | - | - | - | - | - | - | - | - | - | - | - | - | - | - | - | - | M | D | V | V | F | - | - | - | - | - | - | - | - | - | - | - | - | - | - | - | - | - | - | - | - | - | - | - | - | - | - | - | - | - | - | - | - | - | - | - | - | - | - | - | - | - | - | - | - | - | - | - | - | - | - | - | - | - | - | E | Q | N | E | V | V | - | - | - | - | - | - | - | - | - | - | - | - | - | - | - | - | - | - | - | - | - | - | - | - | - | - | - | - | - | - | - | - | - | - | - | - | V | L | V | A | D | S | K | N | C | L | Q | L | - | I | N | G | K | T | - | - | - | - | - | - | - | - | - | - | G | D | V | M | C | Q | T | E | V | - | - | - | - | - | - | - | - | - | - | - | - | - | - | - | - | - | - | - | - | - | - | Q | E | S | - | - | - | - | - | - | - | - | R | I | C | C | C | C | L | S | L | D | L | N | - | - | - | - | - | - | - | - | - | L | A | A | F | G | R | G | D | G | I | L | K | A | G | V | K | S | A | K | L | I | L | Q | S | F | G | W | K | T | Q | P | S | Q | Q | S | P | L | V | A | I | G | L | E | - | - | - | - | - | - | - | - | - | N | G | T | - | - | - | - | - | - | - | - | - | - | V | - | - | - | - | - | - | - | - | - | - | - | - | - | - | - | - | - | - | - | - | - | - | - | - | - | - | - | - | - | - | - | - | - | - | - | - | - | - | - | K | V | L | D | R | H | S | G | R | V | L | H | M | L | N | K | - | - | - | - | - | H | - | - | - | - | - | - | - | - | - | - | - | - | T | K | C | V | Q | R | C | Q | F | T | - | - | - | - | - | - | - | - | - | - | - | - | - | - | - | - | - | - | - | - | - | - | - | - | - | - | - | - | - | - | - | - | - | - | - | - | - | S | D | G | E | T | L | I | S | S | S | - | D | D | T | - | - | - | - | - | - | - | - | T | L | L | - | V | - | - | - | - | - | - | - | - | - | - | - | - | - | - | - | - | - | - | - | - | - | - | - | - | - | - | - | - | - | - | - | - | - | - | - | - | - | - | - | W | N | - | - | - | - | - | - | - | - | - | - | W | R | A | G | E | Y | K | V | - | - | - | - | - | - | - | - | - | - | - | - | - | - | - | - | - | - | - | - | - | - | - | - | - | - | - | - | - | - | - | - | - | - | - | - | - | - | - | - | - | - | - | - | - | - | - | - | - | - | L | K | G | H | I | - | - | - | - | - | - | - | - | - | - | - | - | - | - | - | - | - | - | - | - | - | - | - | - | - | - | E | Q | V | K | N | F | K | V | L | - | - | - | - | - | - | - | - | - | - | - | - | - | - | - | - | - | - | - | - | - | - | - | D | D | - | S | Q | L | L | - | S | W | - | - | - | S | F | D | G | T | M | - | - | - | - | - | - | - | - | - | - | - | - | - | - | - | - | - | - | - | - | - | - | - | - | - | - | - | - | - | - | - | - | - | - | - | K | V | W | N | F | I | S | G | V | M | - | - | - | - | - | L | K | S | Y | G | C | - | - | - | - | - | - | - | H | N | - | - | - | - | - | - | - | - | - | - | - | - | - | - | - | - | - | - | - | G | A | I | L | - | - | - | - | - | - | - | - | - | - | - | - | - | - | - | S | C | D | - | - | - | - | - | - | - | - | - | - | - | - | - | - | - | V | S | P | D | G | S | - | K | L | A | S | A | - | S | A | D | K | Y | - | - | - | - | - | - | A | K | I | W | N | F | S | R | T | - | - | - | - | - | - | - | - | - | - | - | - | - | - | - | - | - | - | - | - | - | - | - | - | A | P | L | Y | V | - | - | - | - | - | - | - | - | L | - | - | - | - | T | G | H | K | A | C | V | - | - | - | - | - | - | - | - | - | R | S | C | - | R | F | S | - | - | - | - | - | - | - | - | - | K | D | N | E | - | Y | L | A | T | - | - | - | - | - | - | - | - | G | D | D | N | G | E | I | R | I | W | R | V | R | D | G | C | L | L | - | - | - | - | - | - | - | - | - | - | - | - | - | - | - | - | - | - | - | - | - | - | - | - | - | - | - | - | - | - | - | - | - | - | - | - | - | - | - | - | - | - | - | - | - | - | - | - | - | - | - | - | - | - | - | - | H | V | C | P | K | - | - | - | E | - | - | - | - | - | - | - | - | - | - | - | - | - | - | - | - | - | - | - | - | S | E | D | S | K | H | G | G | W | - | - | - | - | - | - | - | - | V | T | D | L | H | F | S | P | N | S | K | - | - | - | - | - | - | - | - | - | - | - | - | - | - | - | - | - | - | - | - | - | - | - | - | - | - | - | - | - | - | - | - | - | - | - | - | - | - | - | - | - | - | - | - | - | M | L | I | S | - | - | T | G | G | Y | I | K | - | - | - | - | - | - | - | - | W | W | D | L | E | N | - | - | - | - | - | - | - | - | - | - | - | - | - | - | - | - | - | - | - | - | - | - | - | - | - | - | - | - | - | - | - | - | - | - | - | - | - | - | - | - | - | - | - | - | - | - | - | - | - | - | - | - | - | - | - | - | - | - | - | - | - | - | - | - | - | - | - | - | - | - | - | - | - | - | - | - | - | - | - | - | - | - | - | - | - | - | - | - | - | - | - | - | - | - | - | - | - | - | - | - | - | - | - | - | - | - | - | - | - | - | - | - | - | - | - | - | - | - | - | - | - | - | - | - | - | - | - | - | - | - | - | - | - | - | - | - | - | - | - | - | - | - | - | - | - | - | - | - | - | - | - | - | - | - | - | - | - | - | - | - | - | - | - | - | - | - | - | - | - | - | - | - | - | - | - | - | - | - | - | - | - | - | - | - | - | - | - | - | - | - | - | - | - | - | - | - | - | - | - | - | - | - | - | - | - | - | - | - | - | - | - | - | - | - | - | - | - | - | - | - | - | - | - | - | - | - | - | - | - | - | - | - | - | - | - | - | - | - | - | - | - | - | - | - | - | - | - | - | - | - | - | - | - | - | - | - | - | - | - | - | - | - | - | - | - | - | - | G | E | F | L | Q | T | F | Y | T | - | - | - | - | - | N | G | - | - | - | T | N | L | K | V | I | - | - | - | Y | V | S | A | - | - | - | - | D | F | Q | T | - | - | - | - | - | - | - | - | - | - | - | - | - | - | - | - | - | - | - | - | - | - | - | - | - | - | F | V | T | I | D | - | N | I | G | T | L | - | - | Y | V | L | K | M | - | - | - | - | - | - | - | - | - | - | - | - | - | - | - | - | - | - | - | - | - | - | - | - | - | - | - | - | - | - | - | - | - | - | I | E | R | W | G | G | K | K | D | S | C | D | L | - | - | - | - | - | - | - | - | - | - | - | - | - | - | - | - | - | - | - | - | - | - | - | - | - | - | - | - | - | - | - | - | - | - | - | - | Q | M | - | - | - | M | - | - | - | - | - | - | - | - | - | - | - | - | - | - | - | - | - | - | - | - | - | - | - | - | - | - | - | - | - | - | - | - | - | - | - | - | - | - | - | - | - | - | - | - | - | - | - | - | - | - | - | - | - | - | E | E | G | H | R | Q | - | - | - | - | - | - | - | - | - | - | - | - | - | - | - | - | - | - | - | - | L | L | R | R | - | - | - | - | - | - | - | - | - | - | - | - | - | - | - | - | - | - | - | - | - | - | - | - | - | - | - | - | - | - | R | R | - | - | - | - | - | - | - | - | V | E | L | - | V | T | E | L | Q | P | E | L | L | L | - | - | - | - | - | - | - | - | - | - | - | - | - | - | - | - | - | - | - | - | - | - | - | - | - | - | - | - | - | - | - | - | - | - | - | - | - | - | - | - | - | - | - | - | - | D | P | L | L | A | K | G | I | - | - | - | - | - | - | - | - | - | - | - | - | - | - | - | - | - | - | - | - | - | - | - | - | F | S | Q | D | M | L | E | E | - | - | - | - | - | - | - | - | - | - | - | - | - | - | - | - | - | - | - | - | - | - | - | - | - | I | Q | K | S | - | - | - | - | - | - | - | - | - | - | G | T | R | R | D | Q | - | A | R | Q | L | V | I | D | L | Q | T | R | G | - | R | R | A | F | P | A | F | L | E | S | L | E | E | T | G | Q | H | E | L | V | K | L | L | Q | E | G | L | T | K | H | - | - | - | - | P | G | R | P | - | K | T | S | - | - | - | - | - | - | - | - | - | - | - | - | - | - | - | - | - | - | - | - | - | - | - | - | - | - | W | P | E | - | - | - | - | - | - | - | - | - | - | - | - | - | - | - | - | Y | T | P | V | - | - | - | - | - | - | - | - | - | - | - | - | - | - | - | - | - | - | - | - | - | - | - | - | - | - | - | - | - | - | - | - | - | - | - | - | - | - | - | - | - | - | - | - | - | - | - | - | - | - | - | - | - | - | - | - | - | - | - | - | - | - | - | - | - | - | - | - | - | - | - | - | - | - | - | - | - | - | - | - | - | - | - | - | - | - | - | - | - | - | - | - | - | - | - | - | - | - | - | - | - | - | - | - | - | - | - | - | - | - | - | - | - | - | - | - | - | - | - | - | - | - | - | - | - | - | - | - | - | - | - | - | - | - | - | - | - | - | - | - | - | - | - | - | - | - | - | - | - | - | - | - | - | - | - | - | - | - | - | - | - | - | - | - | - | - | - | - | - | - | - | - | - | - | - | - | - | - | - | - | - | - | - | - | - | - | - | - | - | - | - | - | - | - | - | - | - | - | - | - | - | - | - | - | - | - | - | - | - | - | - | - | - | - | - | - | - | - | - | - | - | - | - | - | - | - | - | - | - | - | - | - | - | - | - | - | - | - | - | - | - | - | - | - | - | - | - | - | - | - | - | - | - | - | - | - | - | - | - | - | - | - | - | - | - | - | R | P | - | - | - | - | - | - | - | - | - | - | - | - | - | - | - | - | - | - | - | - | - | P | A | P | V | Q | C | S | D | T | D | L | - | - | - | - | - | - | - | - | - | - | - | - | - | N | I | C | R | - | - | - | - | - | - | - | - | - | K | P | - | K | L | G | R | N | W | N | D | E | - | - | - | - | - | M | I | Y | K | - | - | M | D | V | V | - | P | R | G | Y | C | L | I | I | N | N | V | D | F | S | K | S | S | - | - | R | L | G | R | R | E | G | S | D | I | D | R | D | N | V | K | R | L | F | - | K | K | L | Q | F | E | V | I | A | K | Q | N | L | T | K | Q | E | L | E | K | E | L | R | E | L | A | H | K | D | H | - | T | R | F | D | C | C | V | V | V | L | L | S | H | G | N | E | A | Q | H | S | Q | F | P | G | A | - | I | Y | G | V | D | G | N | P | I | S | V | Q | Q | I | V | R | H | F | D | G | S | N | C | P | T | L | R | K | K | P | K | L | F | F | I | Q | A | C | - | - | G | G | G | M | K | D | L | G | F | K | T | A | D | E | I | D | G | A | P | G | L | - | L | S | S | G | E | S | - | - | - | - | - | - | - | L | Q | M | D | A | V | P | C | Q | - | - | - | - | T | L | - | - | - | - | - | - | - | - | D | E | I | D | A | T | - | - | - | - | - | - | - | S | S | L | - | - | - | P | T | P | S | D | I | L | V | S | Y | S | T | F | P | - | - | G | - | - | - | - | - | - | - | - | Y | V | S | W | R | D | K | N | E | G | S | W | Y | M | K | T | L | V | E | V | F | S | E | Y | A | H | - | L | - | - | - | - | - | - | - | - | - | - | - | - | - | - | - | - | - | - | - | - | - | - | - | - | - | - | - | - | - | - | - | E | D | L | Q | R | L | L | - | - | - | - | - | T | M | - | - | - | - | - | - | - | - | - | - | - | - | V | T | K | R | I | S | E | D | - | E | E | A | K | - | - | - | - | - | - | - | G | I | Y | - | K | Q | I | - | - | P | G | - | - | - | - | - | - | - | - | - | - | - | - | - | - | - | S | F | N | F | - | - | - | - | - | - | - | - | - | L | R | K | Q | L | Y | F | I | K | P | L | D | Q | - | - | - | - | - | - | - | - | - | - | - | - | - | - | - | - | - | - | - | - | - | - | - | - | - | - | - | - | - | - | - | - | - | - | M | - | - | - | - | - | - | - | - | - | - | - | - | - | - | - | - | - | - | - | - | - | - | - | - | - | - | - | - | - | - | - | - | - | - | - | - | - | - | - | - | - | - | - | - | - | - | - | - | - | - | - | - | G | D | I | N | K | G | K | K | I | F | V | Q | K | C | A | Q | C | H | T | V | E | Q | G | G | K | H | K | T | G | P | N | L | S | G | L | I | G | R | K | T | G | Q | S | P | G | F | S | Y | T | E | A | N | K | N | K | G | I | V | W | G | E | E | T | L | M | E | Y | L | A | N | P | K | K | Y | I | P | G | T | K | M | I | F | A | G | I | K | K | A | N | E | R | T | D | L | I | A | Y | L | K | S | A | T | S | Q | - | - | - | - | - | - |
| Calypte\_anna | M | - | - | - | - | - | - | - | - | - | - | - | - | - | - | - | - | - | - | - | - | - | - | - | - | - | - | - | - | - | - | - | - | - | - | - | - | - | - | - | - | - | - | - | - | - | - | - | - | - | - | - | - | - | - | - | - | - | - | - | - | - | - | - | - | - | - | - | - | - | - | - | - | - | - | - | - | - | - | - | - | - | - | - | - | - | - | - | - | - | - | - | - | - | - | - | - | - | D | V | K | S | - | R | N | - | - | - | - | - | - | - | - | - | - | - | - | - | - | - | - | Y | L | L | - | - | - | - | - | - | - | - | - | - | - | - | - | - | - | - | - | - | - | - | - | - | - | - | - | - | - | - | - | - | - | - | - | - | - | - | - | - | - | - | - | - | - | - | - | - | - | - | - | - | - | - | - | - | - | - | - | - | - | - | - | - | - | - | - | - | - | - | - | - | - | - | - | - | - | - | - | - | - | - | - | - | - | - | - | - | - | - | - | - | - | - | - | - | - | - | - | - | - | - | - | - | - | - | - | - | - | - | - | - | - | - | - | - | - | - | - | - | - | - | - | M | N | R | Q | A | L | E | R | D | I | K | T | S | Y | - | I | M | D | R | M | I | S | D | E | V | L | T | L | Q | E | E | E | R | V | K | Q | Q | - | - | - | - | - | - | - | - | - | - | - | - | - | - | - | - | - | N | T | Q | K | E | R | A | A | M | L | I | N | I | I | L | T | K | - | - | - | - | D | N | N | S | Y | R | S | - | - | - | - | - | F | Y | N | A | L | - | L | H | E | G | - | - | - | - | - | - | Y | R | D | L | A | A | L | L | Q | D | G | I | P | A | I | S | S | G | N | G | - | K | - | - | - | - | - | - | - | - | - | - | - | - | - | - | - | - | - | - | - | - | - | - | S | S | V | D | G | M | T | - | - | - | - | - | - | - | - | - | - | - | S | Y | V | K | T | V | L | R | E | G | G | - | - | - | - | - | - | - | V | P | Q | R | P | V | V | F | V | T | R | P | K | L | V | D | A | I | - | - | - | - | - | - | - | - | - | - | K | K | N | L | C | - | - | - | - | - | - | - | - | - | - | - | - | - | - | - | - | - | - | - | - | - | - | - | - | - | - | - | - | - | - | - | - | C | L | G | S | D | P | - | - | - | - | - | - | - | - | - | - | - | - | - | - | - | - | - | - | - | - | - | - | - | - | - | - | - | - | - | - | - | - | - | - | - | - | - | - | G | W | V | T | V | Y | G | M | A | G | C | G | K | T | V | L | T | A | E | A | L | - | R | D | H | Q | L | L | E | D | Y | F | P | G | G | V | H | W | I | S | V | G | - | - | - | - | - | - | - | K | - | - | - | - | - | - | Q | D | K | A | - | - | - | - | - | - | - | - | - | - | - | - | G | L | L | I | K | L | Q | N | L | C | S | R | L | E | - | - | - | - | H | - | - | - | - | - | - | - | - | - | - | - | - | - | - | - | D | S | T | L | S | Q | R | P | P | L | N | I | - | - | - | - | - | - | - | - | - | - | - | - | - | - | - | - | E | E | A | K | D | R | L | R | L | L | M | L | R | - | - | - | - | - | - | - | - | - | - | - | - | K | Y | P | R | S | L | L | - | - | I | L | D | D | V | W | D | S | W | V | L | K | A | F | D | - | N | Q | C | Q | I | L | I | T | S | R | D | R | S | V | T | D | A | V | - | A | G | N | - | - | K | Y | E | V | H | V | E | S | - | - | - | G | L | A | H | E | K | G | L | E | I | L | S | L | F | V | N | M | - | K | I | S | E | L | P | E | - | - | - | - | - | - | - | - | - | - | - | - | - | - | Q | A | N | C | L | V | R | E | C | K | - | - | - | - | - | - | - | - | - | - | - | - | - | - | - | - | - | - | - | G | - | S | P | L | V | I | S | L | I | G | A | L | L | R | D | F | P | N | - | - | - | - | - | - | - | - | - | - | R | W | E | Y | Y | L | - | K | Q | L | Q | N | K | Q | F | K | R | I | - | - | - | - | - | - | - | - | - | - | - | - | - | - | - | - | - | - | - | - | - | - | R | K | S | S | S | Y | D | - | - | - | - | Y | E | A | L | D | E | A | M | S | I | S | V | E | - | Q | L | N | - | - | - | D | N | Y | K | D | Y | - | Y | K | D | L | S | I | F | P | K | D | V | R | V | P | T | K | V | L | C | I | L | W | D | - | - | - | - | M | E | T | E | E | V | E | D | I | L | Q | E | F | - | - | - | - | - | - | - | - | - | - | - | V | N | K | S | L | L | F | C | D | R | N | G | K | S | - | - | - | - | - | - | - | - | - | F | H | Y | Y | L | H | D | L | Q | L | D | F | L | T | E | K | - | - | N | R | N | Q | - | L | - | Q | E | L | H | K | N | I | V | N | Q | Y | K | K | - | Y | Y | - | - | - | - | - | - | K | V | N | T | P | V | P | S | - | Q | E | - | - | - | - | - | - | - | D | C | M | Y | W | Y | N | F | L | A | Y | H | M | A | G | A | N | - | - | - | - | M | Q | K | E | L | C | D | L | M | F | S | L | D | W | I | K | A | K | T | E | L | V | G | P | A | - | - | - | - | - | - | - | - | H | L | I | H | E | Y | V | E | Y | S | S | V | L | D | Q | K | - | - | - | - | - | - | - | G | S | T | V | R | E | N | F | Q | - | - | - | E | F | L | S | L | N | G | H | L | L | G | R | Q | P | F | - | - | - | - | - | - | - | - | - | - | - | - | - | - | - | - | - | - | - | - | - | - | - | - | - | - | - | - | - | - | - | - | - | - | - | - | - | - | - | - | - | - | - | - | - | - | - | - | - | - | - | - | - | - | - | - | - | - | - | - | - | - | - | - | - | - | - | - | - | - | - | P | D | I | I | Q | L | G | L | C | Q | P | E | A | S | E | V | Y | Q | Q | A | K | L | H | A | Q | R | E | - | - | T | G | - | - | - | - | - | - | - | - | - | - | - | - | - | - | - | - | - | - | - | - | - | - | - | - | - | - | - | - | - | A | F | Y | L | - | - | - | - | - | - | - | - | - | - | E | W | I | N | K | - | - | - | - | - | - | - | K | F | L | K | N | L | Y | R | - | - | - | - | - | - | - | - | - | - | L | V | V | R | P | H | R | D | - | - | - | - | - | - | - | - | - | - | - | - | - | - | - | A | V | Y | H | A | C | - | - | - | F | S | - | K | D | R | Q | R | I | A | - | - | - | - | - | - | - | - | - | S | C | G | A | D | K | T | L | Q | V | F | K | - | - | - | - | - | A | E | S | G | E | R | L | L | S | I | - | S | A | H | D | D | E | - | - | - | I | L | C | C | T | F | S | - | A | - | - | - | - | - | - | - | - | - | - | - | - | - | D | G | E | F | V | A | T | C | S | A | D | K | K | V | K | I | - | - | - | W | N | S | R | - | - | - | - | - | - | - | - | - | - | - | - | - | - | - | - | - | - | - | - | - | - | - | - | - | - | - | - | - | - | - | - | - | - | - | - | - | - | - | - | - | - | - | - | - | - | - | - | - | - | - | - | - | - | - | - | - | - | - | - | - | - | - | - | - | - | - | - | - | - | - | - | - | - | - | - | - | - | - | - | - | - | - | - | - | - | - | - | - | - | - | - | - | - | - | - | - | - | - | - | - | - | - | - | - | - | - | - | - | - | - | - | - | - | - | - | - | - | - | - | - | - | - | - | - | - | - | - | - | - | - | - | - | - | - | - | - | - | - | - | T | G | Q | C | R | C | V | Y | - | E | E | H | T | - | - | - | - | - | E | Q | V | N | - | - | - | - | - | - | - | - | - | - | - | - | - | C | C | Q | - | - | - | - | - | - | - | - | - | - | - | - | - | F | S | N | R | S | - | - | - | - | - | - | - | - | - | - | - | - | - | - | - | - | G | Q | - | Y | V | L | A | T | C | S | S | - | - | - | - | D | T | F | I | - | - | - | - | - | - | - | - | - | - | - | - | K | L | W | D | L | - | - | - | N | K | K | Y | C | R | N | T | M | I | - | - | - | - | - | - | - | - | - | - | - | - | - | - | - | - | - | G | H | E | N | S | - | - | - | - | - | - | - | - | - | - | - | - | - | - | - | - | V | I | H | C | R | F | S | P | N | D | E | - | - | - | - | - | - | - | - | - | - | - | - | - | - | - | - | - | - | - | - | - | - | - | - | - | - | - | - | C | V | A | S | C | S | T | D | G | T | V | K | L | - | - | - | - | - | - | - | - | - | - | - | - | - | - | - | - | - | - | - | W | E | - | S | R | S | G | N | E | L | K | S | - | - | - | - | - | - | I | E | - | - | - | - | - | - | - | - | - | - | - | - | - | - | - | - | - | - | - | - | - | - | I | K | D | F | - | - | - | - | - | - | - | - | F | K | N | A | D | E | Q | A | D | - | - | - | - | - | - | - | - | - | - | - | - | - | - | - | - | - | - | - | - | - | - | - | - | - | - | - | E | V | D | - | I | F | V | K | C | C | - | - | - | - | - | - | - | - | - | - | - | - | - | - | - | - | S | W | S | R | N | G | D | M | I | L | V | V | A | K | S | K | L | - | - | - | - | - | - | - | - | - | - | - | L | L | F | D | V | K | T | - | - | - | C | D | - | - | - | - | - | - | - | - | - | - | - | - | - | - | - | - | - | - | - | - | - | - | - | - | - | - | - | - | - | - | - | - | - | - | - | - | - | - | - | - | - | - | - | - | - | - | - | - | - | - | - | - | - | - | - | - | - | - | - | - | - | - | - | - | - | - | - | - | - | - | - | - | - | - | - | - | - | - | - | - | - | - | - | - | - | - | - | - | - | - | - | - | - | - | - | - | - | - | - | - | - | - | - | - | - | - | - | - | - | - | - | - | - | - | - | - | - | - | - | - | - | - | - | - | - | - | L | L | T | - | - | - | - | - | - | - | - | - | - | - | - | - | - | - | - | - | - | - | - | - | - | - | - | - | - | - | - | - | - | - | - | - | - | - | - | - | - | - | - | - | - | - | - | - | - | - | - | - | - | - | - | - | - | - | - | - | - | - | - | - | - | - | - | - | - | - | - | - | - | - | - | - | - | - | - | - | - | - | - | - | - | - | - | - | - | - | - | - | - | - | - | - | - | - | - | - | - | - | - | - | - | - | - | - | - | - | - | - | - | - | - | - | - | - | - | - | - | - | - | - | - | - | - | - | - | - | - | - | - | - | - | - | - | - | - | - | - | - | - | - | - | - | - | - | - | - | - | - | - | - | - | - | - | - | - | - | - | - | - | - | - | - | - | - | - | - | - | - | - | - | - | - | - | - | - | - | - | - | - | - | - | - | Q | V | - | - | - | I | - | N | H | H | S | T | I | Q | Y | C | - | - | - | - | - | - | - | - | - | - | - | - | - | - | - | - | - | - | - | - | - | - | - | - | - | D | F | C | P | G | D | E | L | V | A | V | A | L | S | H | - | - | - | - | - | - | C | S | V | E | - | - | - | - | - | - | - | - | - | - | - | - | - | - | - | - | - | L | W | N | I | K | S | L | - | - | - | - | - | - | - | - | - | - | - | - | - | - | - | - | - | - | - | - | - | - | - | - | - | - | - | - | - | S | K | V | A | D | C | R | G | H | M | S | - | - | - | - | - | - | - | - | - | - | - | - | - | - | - | - | - | - | - | - | - | - | - | - | - | - | - | - | - | - | - | - | - | - | - | - | - | - | - | - | - | - | - | - | W | V | H | C | V | T | F | S | - | - | - | - | - | - | - | - | - | - | - | S | D | G | S | - | - | - | - | - | - | - | - | - | - | L | F | L | T | S | S | D | D | Q | - | - | - | - | - | - | - | - | - | - | - | - | - | - | T | I | R | I | - | W | - | - | - | - | - | - | - | - | - | - | - | - | - | - | - | - | - | - | - | - | - | - | - | - | - | - | - | - | - | - | - | - | - | - | - | - | - | - | E | T | N | K | V | C | K | S | S | N | D | V | L | K | S | E | - | - | - | - | - | - | - | - | - | - | - | - | - | - | - | - | L | D | V | V | F | - | - | - | - | - | - | - | - | - | - | - | - | - | - | - | - | - | - | - | - | - | - | - | - | - | - | - | - | - | - | - | - | - | - | - | - | - | - | - | - | - | - | - | - | - | - | - | - | - | - | - | - | - | - | H | N | G | E | V | M | - | - | - | - | - | - | - | - | - | - | - | - | - | - | - | - | - | - | - | - | - | - | - | - | - | - | - | - | - | - | - | - | - | - | - | - | I | L | A | I | Y | N | Q | K | H | L | Q | L | - | I | S | G | N | T | - | - | - | - | - | - | - | - | - | - | D | S | V | I | M | Q | T | A | A | - | - | - | - | - | - | - | - | - | - | - | - | - | - | - | - | - | - | - | - | - | - | Q | E | S | - | - | - | - | - | - | - | - | P | I | S | C | C | C | L | S | E | D | L | K | - | - | - | - | - | - | - | - | - | F | A | A | F | G | Q | - | - | - | - | - | - | - | - | - | - | - | - | - | - | - | - | - | - | - | - | - | - | - | - | - | - | - | - | - | - | - | - | - | - | - | - | E | - | - | - | - | - | - | - | - | - | S | G | T | - | - | - | - | - | - | - | - | - | - | L | - | - | - | - | - | - | - | - | - | - | - | - | - | - | - | - | - | - | - | - | - | - | - | - | - | - | - | - | - | - | - | - | - | - | - | - | - | - | - | K | V | L | Q | L | L | D | G | K | I | L | K | T | Q | E | A | - | - | - | - | - | Y | - | - | - | - | - | - | - | - | - | - | - | - | R | T | S | V | Q | H | C | R | F | T | - | - | - | - | - | - | - | - | - | - | - | - | - | - | - | - | - | - | - | - | - | - | - | - | - | - | - | - | - | - | - | - | - | - | - | - | - | S | D | C | Q | S | L | I | S | S | A | - | H | D | S | - | - | - | - | - | - | - | - | V | I | Q | - | V | - | - | - | - | - | - | - | - | - | - | - | - | - | - | - | - | - | - | - | - | - | - | - | - | - | - | - | - | - | - | - | - | - | - | - | - | - | - | - | W | N | - | - | - | - | - | - | - | - | - | - | W | Q | F | S | Q | C | V | F | - | - | - | - | - | - | - | - | - | - | - | - | - | - | - | - | - | - | - | - | - | - | - | - | - | - | - | - | - | - | - | - | - | - | - | - | - | - | - | - | - | - | - | - | - | - | - | - | - | - | L | R | G | H | K | - | - | - | - | - | - | - | - | - | - | - | - | - | - | - | - | - | - | - | - | - | - | - | - | - | - | E | A | I | K | D | F | R | L | L | - | - | - | - | - | - | - | - | - | - | - | - | - | - | - | - | - | - | - | - | - | - | - | E | N | - | S | K | L | L | - | S | W | - | - | - | S | F | D | G | T | V | - | - | - | - | - | - | - | - | - | - | - | - | - | - | - | - | - | - | - | - | - | - | - | - | - | - | - | - | - | - | - | - | - | - | - | K | V | W | N | I | T | T | G | N | T | - | - | - | - | - | E | E | D | F | A | C | - | - | - | - | - | - | - | H | G | - | - | - | - | - | - | - | - | - | - | - | - | - | - | - | - | - | - | - | G | A | V | L | - | - | - | - | - | - | - | - | - | - | - | - | - | - | - | S | C | A | - | - | - | - | - | - | - | - | - | - | - | - | - | - | - | V | S | P | D | G | T | - | K | F | S | S | T | - | S | A | D | K | T | - | - | - | - | - | - | A | K | I | W | S | F | E | S | S | - | - | - | - | - | - | - | - | - | - | - | - | - | - | - | - | - | - | - | - | - | - | - | - | S | V | L | H | E | - | - | - | - | - | - | - | - | L | - | - | - | - | K | G | H | E | A | C | V | - | - | - | - | - | - | - | - | - | R | C | C | - | A | F | S | - | - | - | - | - | - | - | - | - | P | N | N | K | - | V | L | A | T | - | - | - | - | - | - | - | - | G | D | D | K | G | E | I | R | I | W | D | I | L | T | G | A | F | L | - | - | - | - | - | - | - | - | - | - | - | - | - | - | - | - | - | - | - | - | - | - | - | - | - | - | - | - | - | - | - | - | - | - | - | - | - | - | - | - | - | - | - | - | - | - | - | - | - | - | - | - | - | - | - | - | H | F | C | S | P | V | T | V | D | - | - | - | - | - | - | - | - | - | - | - | - | - | - | - | - | - | - | - | - | E | G | E | L | T | H | G | G | W | - | - | - | - | - | - | - | - | V | T | D | L | S | F | S | S | D | S | K | - | - | - | - | - | - | - | - | - | - | - | - | - | - | - | - | - | - | - | - | - | - | - | - | - | - | - | - | - | - | - | - | - | - | - | - | - | - | - | - | - | - | - | - | - | M | L | V | S | - | - | S | G | G | Y | I | K | - | - | - | - | - | - | - | - | W | W | N | V | T | T | - | - | - | - | - | - | - | - | - | - | - | - | - | - | - | - | - | - | - | - | - | - | - | - | - | - | - | - | - | - | - | - | - | - | - | - | - | - | - | - | - | - | - | - | - | - | - | - | - | - | - | - | - | - | - | - | - | - | - | - | - | - | - | - | - | - | - | - | - | - | - | - | - | - | - | - | - | - | - | - | - | - | - | - | - | - | - | - | - | - | - | - | - | - | - | - | - | - | - | - | - | - | - | - | - | - | - | - | - | - | - | - | - | - | - | - | - | - | - | - | - | - | - | - | - | - | - | - | - | - | - | - | - | - | - | - | - | - | - | - | - | - | - | - | - | - | - | - | - | - | - | - | - | - | - | - | - | - | - | - | - | - | - | - | - | - | - | - | - | - | - | - | - | - | - | - | - | - | - | - | - | - | - | - | - | - | - | - | - | - | - | - | - | - | - | - | - | - | - | - | - | - | - | - | - | - | - | - | - | - | - | - | - | - | - | - | - | - | - | - | - | - | - | - | - | - | - | - | - | - | - | - | - | - | - | - | - | - | - | - | - | - | - | - | - | - | - | - | - | - | - | - | - | - | - | - | - | - | - | - | - | - | - | - | - | - | - | G | E | S | L | Q | T | F | Y | T | - | - | - | - | - | N | G | - | - | - | T | N | L | K | S | I | - | - | - | H | V | S | P | - | - | - | - | D | F | K | V | - | - | - | - | - | - | - | - | - | - | - | - | - | - | - | - | - | - | - | - | - | - | - | - | - | - | Y | V | T | V | D | - | N | L | G | I | L | - | - | Y | I | L | Q | K | - | - | - | - | - | - | - | - | - | - | - | - | - | - | - | - | - | - | - | - | - | - | - | - | - | - | - | - | - | - | - | - | - | - | L | - | - | - | - | - | - | - | - | - | - | - | - | - | - | - | - | - | - | - | - | - | - | - | - | - | - | - | - | - | - | - | - | - | - | - | - | - | - | - | - | - | - | - | - | - | - | - | - | - | - | - | - | - | M | - | - | - | - | - | - | - | - | - | - | - | - | - | - | - | - | - | - | - | - | - | - | - | - | - | - | - | - | - | - | - | - | - | - | - | - | - | - | - | - | - | - | - | - | - | - | - | - | - | - | - | - | - | - | E | E | A | Q | R | R | - | - | - | - | - | - | - | - | - | - | - | - | - | - | - | - | - | - | - | - | A | L | R | R | - | - | - | - | - | - | - | - | - | - | - | - | - | - | - | - | - | - | - | - | - | - | - | - | - | - | - | - | - | - | A | R | - | - | - | - | - | - | - | - | P | R | A | - | V | A | E | L | R | V | E | P | L | W | - | - | - | - | - | - | - | - | - | - | - | - | - | - | - | - | - | - | - | - | - | - | - | - | - | - | - | - | - | - | - | - | - | - | - | - | - | - | - | - | - | - | - | - | - | G | L | L | E | E | R | G | L | - | - | - | - | - | - | - | - | - | - | - | - | - | - | - | - | - | - | - | - | - | - | - | - | F | T | R | P | M | I | E | D | - | - | - | - | - | - | - | - | - | - | - | - | - | - | - | - | - | - | - | - | - | - | - | - | - | L | Q | S | A | - | - | - | - | - | - | - | - | - | - | G | S | R | A | D | Q | - | A | R | Q | L | L | I | D | L | E | T | R | G | - | K | Q | A | F | P | L | F | L | S | I | L | R | D | T | E | Q | C | S | L | A | D | V | L | L | E | A | S | E | H | P | S | - | - | A | V | A | P | S | - | Q | - | - | - | - | - | - | - | - | - | - | - | - | - | - | - | - | - | - | - | - | - | - | - | - | - | - | - | - | - | L | Q | H | - | - | - | - | - | - | - | - | - | - | - | - | - | - | - | L | K | P | L | - | - | - | - | - | - | - | - | - | - | - | - | - | - | - | - | - | - | - | - | - | - | - | - | - | - | - | - | - | - | - | - | - | - | - | - | - | - | - | Q | L | E | L | P | G | E | K | H | K | K | - | - | - | - | - | - | - | - | - | - | - | - | - | - | - | - | - | - | - | - | - | - | - | - | - | - | - | - | - | - | - | - | - | - | - | - | - | - | - | - | - | - | - | - | - | - | - | - | - | - | - | - | - | - | - | - | - | - | - | - | - | - | - | - | - | - | - | - | - | - | - | - | - | - | - | - | - | - | - | - | - | - | - | - | - | - | - | - | - | - | - | - | - | - | - | - | - | - | - | - | - | - | - | - | - | - | - | - | - | - | - | - | - | - | - | - | - | - | - | - | - | - | - | - | - | - | - | - | - | - | - | - | - | - | - | - | - | - | - | - | - | - | - | - | - | - | - | - | - | - | - | - | - | - | - | - | - | - | - | - | - | - | - | - | - | - | - | - | - | - | - | - | - | - | - | - | - | - | - | - | - | - | - | - | - | - | - | - | - | - | - | - | - | - | - | - | - | - | - | - | - | - | - | - | - | - | - | - | - | - | - | - | - | - | - | - | - | - | - | - | - | - | - | - | - | - | - | - | - | - | - | - | - | - | - | - | - | - | - | - | - | - | - | - | - | G | - | - | - | - | - | - | - | - | - | - | - | - | - | - | - | - | - | - | - | - | L | - | - | - | - | - | - | - | - | - | - | A | - | A | G | - | - | K | R | S | H | D | - | - | - | - | - | L | V | Y | E | - | - | L | R | G | D | - | P | C | G | Y | C | L | I | L | N | N | V | T | F | R | Q | G | S | - | - | G | L | S | T | R | V | G | S | D | V | D | C | Q | K | L | Q | K | R | F | - | Q | A | L | C | F | Q | V | L | T | R | R | D | L | T | A | Q | E | I | V | L | E | L | Q | E | L | A | R | Q | D | H | - | S | A | L | D | C | C | V | V | V | I | L | S | H | G | C | Q | T | S | H | I | Q | F | P | G | G | - | I | Y | G | T | D | G | K | P | I | P | I | E | K | I | V | N | Y | F | N | G | S | Q | C | P | S | L | R | G | K | P | K | L | F | F | I | Q | A | C | - | - | G | G | E | E | R | D | R | G | F | V | V | D | C | D | - | - | - | S | P | P | D | - | N | A | P | G | G | S | - | - | - | - | - | - | - | L | E | S | D | A | T | P | F | Q | A | P | P | G | N | V | - | - | - | - | - | - | - | - | D | E | L | D | A | V | - | - | - | - | - | - | - | A | S | L | - | - | - | P | T | P | S | D | I | L | V | S | Y | S | T | F | P | - | - | G | - | - | - | - | - | - | - | - | F | V | S | W | R | E | A | L | S | G | S | W | Y | V | E | T | L | D | S | V | L | E | Q | Y | H | P | - | A | - | - | - | - | - | - | - | - | - | - | - | - | - | - | - | - | - | - | - | - | - | - | - | - | - | - | - | - | - | - | - | P | Q | L | L | S | L | S | - | - | - | - | - | S | Q | - | - | - | - | - | - | - | - | - | - | - | - | V | A | H | I | V | - | - | - | - | - | S | A | K | - | - | - | - | - | - | - | G | T | F | - | K | Q | I | - | - | P | G | - | - | - | - | - | - | - | - | - | - | - | - | - | - | - | C | F | N | F | - | - | - | - | - | - | - | - | - | L | R | K | K | F | F | F | M | S | K | - | - | - | - | - | - | - | - | - | - | - | - | - | - | - | - | - | - | - | - | - | - | - | - | - | - | - | - | - | - | - | - | - | - | - | - | - | M | - | - | - | - | - | - | - | - | - | - | - | - | - | - | - | - | - | - | - | - | - | - | - | - | - | - | - | - | - | - | - | - | - | - | - | - | - | - | - | - | - | - | - | - | - | - | - | - | - | - | - | - | G | D | I | E | K | G | K | K | I | F | V | Q | K | C | S | Q | C | H | T | V | E | K | G | G | K | H | K | T | G | P | N | L | N | G | I | F | G | R | K | T | G | Q | A | E | G | F | S | Y | T | D | A | N | K | N | K | G | I | T | W | G | E | D | T | L | M | E | Y | L | E | N | P | K | K | Y | I | P | G | T | K | M | I | F | A | G | I | K | K | K | S | E | R | V | D | L | I | A | Y | L | K | S | A | T | A | K | - | - | - | - | - | - |
| Camelus\_bactrianus | M | - | - | - | - | - | - | - | - | - | - | - | - | - | - | - | - | - | - | - | - | - | - | - | - | - | - | - | - | - | - | - | - | - | - | - | - | - | - | - | - | - | - | - | - | - | - | - | - | - | - | - | - | - | - | - | - | - | - | - | - | - | - | - | - | - | - | - | - | - | - | - | - | - | - | - | - | - | - | - | - | - | - | - | - | - | - | - | - | - | - | - | - | - | - | - | - | - | D | A | K | A | - | R | N | - | - | - | - | - | - | - | - | - | - | - | - | - | - | - | - | C | L | L | - | - | - | - | - | - | - | - | - | - | - | - | - | - | - | - | - | - | - | - | - | - | - | - | - | - | - | - | - | - | - | - | - | - | - | - | - | - | - | - | - | - | - | - | - | - | - | - | - | - | - | - | - | - | - | - | - | - | - | - | - | - | - | - | - | - | - | - | - | - | - | - | - | - | - | - | - | - | - | - | - | - | - | - | - | - | - | - | - | - | - | - | - | - | - | - | - | - | - | - | - | - | - | - | - | - | - | - | - | - | - | - | - | - | - | - | - | - | - | - | - | Q | H | R | E | A | L | E | R | D | I | K | T | S | Y | - | I | M | D | H | M | I | S | N | G | V | L | T | V | S | E | E | E | K | V | K | N | E | - | - | - | - | - | - | - | - | - | - | - | - | - | - | - | - | - | P | T | Q | Q | Q | R | A | A | M | L | I | K | M | I | L | K | K | - | - | - | - | D | N | Y | S | Y | V | S | - | - | - | - | - | F | Y | N | A | L | - | L | H | E | G | - | - | - | - | - | - | Y | K | D | F | A | A | L | L | H | A | G | I | P | V | I | S | S | S | N | G | - | K | - | - | - | - | - | - | - | - | - | - | - | - | - | - | - | - | - | - | - | - | - | - | D | S | V | G | G | I | T | - | - | - | - | - | - | - | - | - | - | - | S | Y | V | R | T | V | L | C | E | G | G | - | - | - | - | - | - | - | V | P | Q | R | P | V | V | F | V | T | R | K | K | L | V | H | A | I | - | - | - | - | - | - | - | - | - | - | Q | Q | K | L | F | - | - | - | - | - | - | - | - | - | - | - | - | - | - | - | - | - | - | - | - | - | - | - | - | - | - | - | - | - | - | - | - | K | L | S | G | E | P | - | - | - | - | - | - | - | - | - | - | - | - | - | - | - | - | - | - | - | - | - | - | - | - | - | - | - | - | - | - | - | - | - | - | - | - | - | - | G | W | V | T | I | Y | G | M | A | G | C | G | K | S | V | L | A | A | E | A | V | - | R | D | H | S | L | L | E | V | C | F | P | G | G | V | H | W | V | S | V | G | - | - | - | - | - | - | - | K | - | - | - | - | - | - | Q | D | K | S | - | - | - | - | - | - | - | - | - | - | - | - | G | L | L | M | K | L | Q | N | L | C | T | R | L | D | - | - | - | - | Q | - | - | - | - | - | - | - | - | - | - | - | - | - | - | - | D | E | S | F | S | Q | R | L | P | L | N | I | - | - | - | - | - | - | - | - | - | - | - | - | - | - | - | - | E | E | A | K | D | R | L | R | I | L | M | L | R | - | - | - | - | - | - | - | - | - | - | - | - | K | H | P | R | S | L | L | - | - | I | L | D | D | I | W | D | P | W | V | L | K | A | F | D | - | N | Q | C | Q | I | L | L | T | T | R | D | K | S | V | T | D | S | V | - | M | G | P | - | - | K | Y | V | V | P | V | E | S | - | - | - | G | L | G | K | E | K | G | L | E | I | L | S | L | F | V | N | M | - | K | K | A | D | L | P | E | - | - | - | - | - | - | - | - | - | - | - | - | - | - | Q | A | H | S | I | I | K | E | C | K | - | - | - | - | - | - | - | - | - | - | - | - | - | - | - | - | - | - | - | G | - | S | P | L | V | V | S | L | I | G | A | L | L | R | D | F | P | N | - | - | - | - | - | - | - | - | - | - | R | W | E | Y | Y | L | - | K | Q | L | Q | N | K | Q | F | K | R | I | - | - | - | - | - | - | - | - | - | - | - | - | - | - | - | - | - | - | - | - | - | - | R | K | S | S | S | Y | D | - | - | - | - | Y | E | A | L | D | E | A | M | S | I | S | V | E | - | M | L | R | - | - | - | E | D | I | K | D | Y | - | Y | T | D | L | S | I | L | Q | K | D | V | K | V | P | T | K | V | L | C | I | L | W | D | - | - | - | - | M | E | T | E | E | V | E | D | I | L | Q | E | F | - | - | - | - | - | - | - | - | - | - | - | V | N | K | S | L | L | F | C | D | R | N | G | K | S | - | - | - | - | - | - | - | - | - | F | L | Y | Y | L | H | D | L | Q | V | D | F | L | T | E | K | - | - | N | R | S | Q | - | L | - | Q | D | L | H | K | K | I | I | T | Q | F | Q | R | - | H | R | - | - | - | - | - | - | R | L | H | T | L | S | P | D | - | Q | E | - | - | - | - | - | - | - | D | C | M | Y | W | Y | N | F | L | A | Y | H | M | A | S | A | N | - | - | - | - | M | H | K | E | L | C | A | L | M | F | S | L | D | W | I | K | A | K | T | E | L | V | G | P | A | - | - | - | - | - | - | - | - | H | L | I | H | E | F | V | E | Y | R | H | M | L | D | E | K | - | - | - | - | - | - | - | D | G | T | V | C | E | N | F | Q | - | - | - | E | F | L | S | L | N | G | H | L | L | G | R | Q | P | F | - | - | - | - | - | - | - | - | - | - | - | - | - | - | - | - | - | - | - | - | - | - | - | - | - | - | - | - | - | - | - | - | - | - | - | - | - | - | - | - | - | - | - | - | - | - | - | - | - | - | - | - | - | - | - | - | - | - | - | - | - | - | - | - | - | - | - | - | - | - | - | P | N | I | V | Q | L | G | L | C | E | L | E | T | S | E | V | Y | Q | Q | A | K | L | Q | A | K | Q | E | V | D | N | G | - | - | - | - | - | - | - | - | - | - | - | - | - | - | - | - | - | - | - | - | - | - | - | - | - | - | - | - | - | M | L | Y | L | - | - | - | - | - | - | - | - | - | - | E | W | I | N | K | - | - | - | - | - | - | - | K | N | I | K | N | L | S | R | - | - | - | - | - | - | - | - | - | - | L | V | V | C | P | H | T | D | - | - | - | - | - | - | - | - | - | - | - | - | - | - | - | A | V | Y | H | A | C | - | - | - | F | S | - | E | D | G | Q | R | I | A | - | - | - | - | - | - | - | - | - | S | C | G | A | D | K | T | L | Q | V | F | K | - | - | - | - | - | A | E | T | G | E | K | L | L | E | I | - | K | A | H | E | D | E | - | - | - | V | L | C | C | A | F | S | - | A | - | - | - | - | - | - | - | - | - | - | - | - | - | D | D | K | F | I | A | T | C | S | V | D | K | K | V | K | I | - | - | - | W | N | S | M | - | - | - | - | - | - | - | - | - | - | - | - | - | - | - | - | - | - | - | - | - | - | - | - | - | - | - | - | - | - | - | - | - | - | - | - | - | - | - | - | - | - | - | - | - | - | - | - | - | - | - | - | - | - | - | - | - | - | - | - | - | - | - | - | - | - | - | - | - | - | - | - | - | - | - | - | - | - | - | - | - | - | - | - | - | - | - | - | - | - | - | - | - | - | - | - | - | - | - | - | - | - | - | - | - | - | - | - | - | - | - | - | - | - | - | - | - | - | - | - | - | - | - | - | - | - | - | - | - | - | - | - | - | - | - | - | - | - | - | - | T | G | E | L | V | H | T | Y | - | D | E | H | S | - | - | - | - | - | E | Q | V | N | - | - | - | - | - | - | - | - | - | - | - | - | - | C | C | H | - | - | - | - | - | - | - | - | - | - | - | - | - | F | T | N | N | S | - | - | - | - | - | - | - | - | - | - | - | - | - | - | - | - | N | H | - | L | L | L | A | T | G | S | S | - | - | - | - | D | C | F | L | - | - | - | - | - | - | - | - | - | - | - | - | K | L | W | D | V | - | - | - | N | Q | K | E | C | R | N | T | M | F | - | - | - | - | - | - | - | - | - | - | - | - | - | - | - | - | - | G | H | T | N | S | - | - | - | - | - | - | - | - | - | - | - | - | - | - | - | - | V | N | H | C | R | F | S | P | D | D | K | - | - | - | - | - | - | - | - | - | - | - | - | - | - | - | - | - | - | - | - | - | - | - | - | - | - | - | - | L | L | A | S | C | S | A | D | G | T | L | K | L | - | - | - | - | - | - | - | - | - | - | - | - | - | - | - | - | - | - | - | W | D | - | V | K | S | A | N | E | R | K | S | - | - | - | - | - | - | I | N | - | - | - | - | - | - | - | - | - | - | - | - | - | - | - | - | - | - | - | - | - | - | V | K | H | F | - | - | - | - | - | - | - | - | F | L | N | S | E | E | P | Q | E | - | - | - | - | - | - | - | - | - | - | - | - | - | - | - | - | - | - | - | - | - | - | - | - | - | - | - | D | M | E | - | V | I | V | K | C | C | - | - | - | - | - | - | - | - | - | - | - | - | - | - | - | - | S | W | S | A | D | G | A | R | I | M | V | A | A | K | N | K | V | - | - | - | - | - | - | - | - | - | - | - | F | L | F | D | I | H | T | - | - | - | S | G | - | - | - | - | - | - | - | - | - | - | - | - | - | - | - | - | - | - | - | - | - | - | - | - | - | - | - | - | - | - | - | - | - | - | - | - | - | - | - | - | - | - | - | - | - | - | - | - | - | - | - | - | - | - | - | - | - | - | - | - | - | - | - | - | - | - | - | - | - | - | - | - | - | - | - | - | - | - | - | - | - | - | - | - | - | - | - | - | - | - | - | - | - | - | - | - | - | - | - | - | - | - | - | - | - | - | - | - | - | - | - | - | - | - | - | - | - | - | - | - | - | - | - | - | - | - | L | L | A | - | - | - | - | - | - | - | - | - | - | - | - | - | - | - | - | - | - | - | - | - | - | - | - | - | - | - | - | - | - | - | - | - | - | - | - | - | - | - | - | - | - | - | - | - | - | - | - | - | - | - | - | - | - | - | - | - | - | - | - | - | - | - | - | - | - | - | - | - | - | - | - | - | - | - | - | - | - | - | - | - | - | - | - | - | - | - | - | - | - | - | - | - | - | - | - | - | - | - | - | - | - | - | - | - | - | - | - | - | - | - | - | - | - | - | - | - | - | - | - | - | - | - | - | - | - | - | - | - | - | - | - | - | - | - | - | - | - | - | - | - | - | - | - | - | - | - | - | - | - | - | - | - | - | - | - | - | - | - | - | - | - | - | - | - | - | - | - | - | - | - | - | - | - | - | - | - | - | - | - | - | - | - | E | I | - | - | - | H | T | G | H | H | S | T | V | Q | Y | C | - | - | - | - | - | - | - | - | - | - | - | - | - | - | - | - | - | - | - | - | - | - | - | - | - | D | F | S | P | H | N | H | L | A | L | V | A | L | S | Q | - | - | - | - | - | - | Y | C | V | E | - | - | - | - | - | - | - | - | - | - | - | - | - | - | - | - | - | L | W | N | V | D | S | C | - | - | - | - | - | - | - | - | - | - | - | - | - | - | - | - | - | - | - | - | - | - | - | - | - | - | - | - | - | L | K | V | A | D | C | R | G | H | L | S | - | - | - | - | - | - | - | - | - | - | - | - | - | - | - | - | - | - | - | - | - | - | - | - | - | - | - | - | - | - | - | - | - | - | - | - | - | - | - | - | - | - | - | - | W | V | H | C | V | M | F | S | - | - | - | - | - | - | - | - | - | - | - | P | D | G | S | - | - | - | - | - | - | - | - | - | - | S | F | L | T | S | S | D | D | Q | - | - | - | - | - | - | - | - | - | - | - | - | - | - | T | I | R | L | - | W | - | - | - | - | - | - | - | - | - | - | - | - | - | - | - | - | - | - | - | - | - | - | - | - | - | - | - | - | - | - | - | - | - | - | - | - | - | - | E | T | K | K | V | C | K | N | S | A | I | V | L | K | Q | E | - | - | - | - | - | - | - | - | - | - | - | - | - | - | - | - | I | D | V | V | F | - | - | - | - | - | - | - | - | - | - | - | - | - | - | - | - | - | - | - | - | - | - | - | - | - | - | - | - | - | - | - | - | - | - | - | - | - | - | - | - | - | - | - | - | - | - | - | - | - | - | - | - | - | - | Q | E | N | E | V | M | - | - | - | - | - | - | - | - | - | - | - | - | - | - | - | - | - | - | - | - | - | - | - | - | - | - | - | - | - | - | - | - | - | - | - | - | V | L | A | V | D | N | I | R | R | L | Q | L | - | I | N | G | K | T | - | - | - | - | - | - | - | - | - | - | G | Q | I | D | Y | L | T | E | A | - | - | - | - | - | - | - | - | - | - | - | - | - | - | - | - | - | - | - | - | - | - | H | - | - | - | - | - | - | - | - | - | - | - | V | S | C | C | C | L | S | P | H | L | Q | - | - | - | - | - | - | - | - | - | Y | V | A | F | G | G | - | - | - | - | - | - | - | - | - | - | - | - | - | - | - | - | - | - | - | - | - | - | - | - | - | - | - | - | - | - | - | - | - | - | - | - | E | - | - | - | - | - | - | - | - | - | D | G | A | - | - | - | - | - | - | - | - | - | - | I | - | - | - | - | - | - | - | - | - | - | - | - | - | - | - | - | - | - | - | - | - | - | - | - | - | - | - | - | - | - | - | - | - | - | - | - | - | - | - | Q | I | L | E | L | L | N | N | R | V | F | Q | S | K | I | G | - | - | - | - | - | H | - | - | - | - | - | - | - | - | - | - | - | - | K | N | T | V | R | H | I | Q | F | T | - | - | - | - | - | - | - | - | - | - | - | - | - | - | - | - | - | - | - | - | - | - | - | - | - | - | - | - | - | - | - | - | - | - | - | - | - | A | D | G | K | N | L | I | S | S | S | - | D | D | C | - | - | - | - | - | - | - | - | T | I | Q | - | I | - | - | - | - | - | - | - | - | - | - | - | - | - | - | - | - | - | - | - | - | - | - | - | - | - | - | - | - | - | - | - | - | - | - | - | - | - | - | - | W | N | - | - | - | - | - | - | - | - | - | - | W | Q | S | E | E | Y | V | F | - | - | - | - | - | - | - | - | - | - | - | - | - | - | - | - | - | - | - | - | - | - | - | - | - | - | - | - | - | - | - | - | - | - | - | - | - | - | - | - | - | - | - | - | - | - | - | - | - | - | L | Q | A | H | Q | - | - | - | - | - | - | - | - | - | - | - | - | - | - | - | - | - | - | - | - | - | - | - | - | - | - | E | A | V | K | D | F | K | L | L | - | - | - | - | - | - | - | - | - | - | - | - | - | - | - | - | - | - | - | - | - | - | - | K | N | - | S | R | L | L | - | S | W | - | - | - | S | F | D | G | T | V | - | - | - | - | - | - | - | - | - | - | - | - | - | - | - | - | - | - | - | - | - | - | - | - | - | - | - | - | - | - | - | - | - | - | - | K | V | W | N | F | I | T | G | R | I | - | - | - | - | - | E | K | D | F | V | C | - | - | - | - | - | - | - | H | Q | - | - | - | - | - | - | - | - | - | - | - | - | - | - | - | - | - | - | - | D | T | V | L | - | - | - | - | - | - | - | - | - | - | - | - | - | - | - | S | C | D | - | - | - | - | - | - | - | - | - | - | - | - | - | - | - | I | S | P | D | A | T | - | K | F | S | S | T | - | S | A | D | K | T | - | - | - | - | - | - | A | K | I | W | S | F | E | L | I | - | - | - | - | - | - | - | - | - | - | - | - | - | - | - | - | - | - | - | - | - | - | - | - | S | P | L | H | E | - | - | - | - | - | - | - | - | L | - | - | - | - | R | G | H | K | G | C | V | - | - | - | - | - | - | - | - | - | R | C | S | - | A | F | S | - | - | - | - | - | - | - | - | - | V | D | S | T | - | L | L | A | T | - | - | - | - | - | - | - | - | G | D | D | N | G | E | I | R | I | W | N | V | S | N | G | E | L | L | - | - | - | - | - | - | - | - | - | - | - | - | - | - | - | - | - | - | - | - | - | - | - | - | - | - | - | - | - | - | - | - | - | - | - | - | - | - | - | - | - | - | - | - | - | - | - | - | - | - | - | - | - | - | - | - | H | L | C | A | P | I | S | V | E | - | - | - | - | - | - | - | - | - | - | - | - | - | - | - | - | - | - | - | - | E | G | A | A | T | H | G | G | W | - | - | - | - | - | - | - | - | V | T | D | L | C | F | S | P | N | S | K | - | - | - | - | - | - | - | - | - | - | - | - | - | - | - | - | - | - | - | - | - | - | - | - | - | - | - | - | - | - | - | - | - | - | - | - | - | - | - | - | - | - | - | - | - | M | L | V | S | - | - | A | G | G | Y | L | K | - | - | - | - | - | - | - | - | W | W | N | V | V | T | - | - | - | - | - | - | - | - | - | - | - | - | - | - | - | - | - | - | - | - | - | - | - | - | - | - | - | - | - | - | - | - | - | - | - | - | - | - | - | - | - | - | - | - | - | - | - | - | - | - | - | - | - | - | - | - | - | - | - | - | - | - | - | - | - | - | - | - | - | - | - | - | - | - | - | - | - | - | - | - | - | - | - | - | - | - | - | - | - | - | - | - | - | - | - | - | - | - | - | - | - | - | - | - | - | - | - | - | - | - | - | - | - | - | - | - | - | - | - | - | - | - | - | - | - | - | - | - | - | - | - | - | - | - | - | - | - | - | - | - | - | - | - | - | - | - | - | - | - | - | - | - | - | - | - | - | - | - | - | - | - | - | - | - | - | - | - | - | - | - | - | - | - | - | - | - | - | - | - | - | - | - | - | - | - | - | - | - | - | - | - | - | - | - | - | - | - | - | - | - | - | - | - | - | - | - | - | - | - | - | - | - | - | - | - | - | - | - | - | - | - | - | - | - | - | - | - | - | - | - | - | - | - | - | - | - | - | - | - | - | - | - | - | - | - | - | - | - | - | - | - | - | - | - | - | - | - | - | - | - | - | - | - | - | - | - | - | G | E | S | S | Q | T | F | Y | T | - | - | - | - | - | N | G | - | - | - | T | N | L | K | K | I | - | - | - | H | V | S | P | - | - | - | - | D | F | K | T | - | - | - | - | - | - | - | - | - | - | - | - | - | - | - | - | - | - | - | - | - | - | - | - | - | - | Y | V | T | V | D | - | N | L | G | I | L | - | - | Y | I | L | K | I | - | - | - | - | - | - | - | - | - | - | - | - | - | - | - | - | - | - | - | - | - | - | - | - | - | - | - | - | - | - | - | - | - | - | L | E | - | - | - | - | - | - | - | - | - | - | - | - | - | - | - | - | - | - | - | - | - | - | - | - | - | - | - | - | - | - | - | - | - | - | - | - | - | - | - | - | - | - | - | - | - | - | - | - | - | - | - | - | M | - | - | - | - | - | - | - | - | - | - | - | - | - | - | - | - | - | - | - | - | - | - | - | - | - | - | - | - | - | - | - | - | - | - | - | - | - | - | - | - | - | - | - | - | - | - | - | - | - | - | - | - | - | - | - | - | - | - | - | - | - | - | - | - | - | - | - | - | - | - | - | - | - | - | - | - | - | - | - | - | - | - | - | - | - | - | - | - | - | - | - | - | - | - | - | - | - | - | - | - | - | - | - | - | - | - | - | - | - | - | - | - | - | - | - | - | - | - | - | - | - | - | - | - | - | - | - | - | - | - | - | - | - | - | - | - | - | - | - | - | - | - | - | - | - | - | - | - | - | - | - | - | - | - | - | - | - | - | - | - | - | - | - | - | - | - | - | - | - | - | - | - | - | - | - | - | - | - | - | - | - | - | - | - | - | - | - | - | - | - | - | - | - | - | - | - | - | - | - | - | - | - | - | - | - | - | - | - | - | - | - | - | - | - | - | - | - | - | - | - | - | - | - | - | - | - | - | - | - | - | - | - | - | - | - | - | - | - | - | - | - | - | - | - | - | - | - | - | - | - | - | - | - | - | - | - | - | - | - | - | - | - | - | - | - | - | - | - | - | - | - | - | - | - | - | - | - | - | - | - | - | - | - | - | - | - | - | - | - | - | - | - | - | - | - | - | - | - | - | - | L | A | S | L | L | R | T | S | S | Q | A | A | - | - | - | - | R | Q | H | L | - | E | A | I | - | - | - | - | - | - | - | - | - | - | - | - | - | - | - | - | - | - | - | - | - | - | - | - | - | - | R | P | L | D | - | - | - | - | - | - | - | - | - | - | - | - | - | - | - | L | K | P | V | - | - | - | - | - | - | - | - | - | - | - | - | - | - | - | - | - | - | - | - | - | - | - | - | - | - | - | - | - | - | - | - | - | - | - | - | - | - | - | V | L | R | P | E | G | L | K | P | E | S | L | G | - | V | G | K | Q | D | P | L | M | P | - | T | - | P | - | - | - | - | - | - | - | - | - | - | - | - | - | - | - | - | - | - | - | - | - | - | - | - | - | - | - | - | - | - | - | - | - | - | - | - | - | - | - | - | - | - | - | - | - | - | - | - | - | - | - | G | K | L | V | P | V | - | - | - | - | - | - | - | - | - | - | - | - | - | - | - | - | - | - | - | - | - | - | - | - | - | - | V | L | - | - | - | - | - | - | - | - | - | - | - | - | - | - | - | - | - | - | - | - | - | - | - | - | - | - | - | - | - | - | - | - | - | - | - | - | - | - | - | - | - | - | - | - | - | - | - | - | - | - | - | - | - | - | - | - | - | - | - | - | - | - | - | - | - | - | - | - | G | P | K | E | L | W | P | - | - | - | - | - | - | - | - | - | - | - | - | - | - | - | - | - | - | - | - | - | - | - | - | - | - | A | K | L | R | P | - | - | - | - | - | - | E | V | L | R | P | - | E | V | - | - | - | - | - | - | - | - | - | - | - | - | - | - | - | - | - | - | P | R | P | V | D | I | G | S | G | G | L | - | - | - | - | - | - | - | - | - | - | - | - | - | S | D | V | C | - | - | - | - | - | - | - | - | - | - | A | - | Q | D | R | A | K | G | N | A | N | - | - | - | - | - | L | A | Y | V | - | - | L | S | A | D | - | P | C | G | R | C | L | I | V | N | N | V | H | F | R | R | E | S | - | - | G | L | R | T | R | T | G | S | S | V | D | C | E | R | L | Q | R | R | F | - | H | L | L | H | F | V | V | E | V | E | C | D | L | T | A | K | Q | M | V | Q | A | L | V | E | L | A | R | R | D | H | - | S | A | L | D | C | C | V | V | V | I | L | S | H | G | C | Q | A | S | H | L | Q | F | P | G | A | - | V | Y | G | T | D | G | C | P | V | S | V | E | R | I | V | N | I | F | N | G | T | G | C | P | S | L | R | G | K | P | K | L | F | F | I | Q | A | C | - | - | G | G | E | Q | K | D | H | G | F | E | V | A | S | T | - | - | - | F | P | E | G | - | Q | T | P | D | S | D | - | - | - | - | - | - | - | P | E | P | D | A | T | P | F | Q | E | G | P | G | T | V | - | - | - | - | - | - | - | - | D | Q | P | D | A | V | - | - | - | - | - | - | - | S | S | L | - | - | - | P | T | P | S | D | I | L | V | S | Y | S | T | F | P | - | - | G | - | - | - | - | - | - | - | - | F | V | S | W | R | D | T | K | S | G | S | W | Y | I | E | T | L | D | S | V | F | E | Q | W | A | H | - | C | - | - | - | - | - | - | - | - | - | - | - | - | - | - | - | - | - | - | - | - | - | - | - | - | - | - | - | - | - | - | - | E | D | L | Q | T | L | L | - | - | - | - | - | L | R | - | - | - | - | - | - | - | - | - | - | - | - | V | A | N | A | V | - | - | - | - | - | S | V | K | - | - | - | - | - | - | - | G | I | Y | - | K | Q | I | - | - | P | G | - | - | - | - | - | - | - | - | - | - | - | - | - | - | - | C | F | N | F | - | - | - | - | - | - | - | - | - | L | R | K | K | L | F | F | K | T | - | - | - | - | - | - | - | - | - | - | - | - | - | - | - | - | - | - | - | - | - | - | - | - | - | - | - | - | - | - | - | - | - | - | - | - | - | - | M | - | - | - | - | - | - | - | - | - | - | - | - | - | - | - | - | - | - | - | - | - | - | - | - | - | - | - | - | - | - | - | - | - | - | - | - | - | - | - | - | - | - | - | - | - | - | - | - | - | - | - | - | G | D | V | E | K | G | K | K | I | F | V | Q | K | C | A | Q | C | H | T | V | E | K | G | G | K | H | K | T | G | P | N | L | H | G | L | F | G | R | K | T | G | Q | A | V | G | F | S | Y | T | D | A | N | K | N | K | G | I | T | W | G | E | E | T | L | M | E | Y | L | E | N | P | K | K | Y | I | P | G | T | K | M | I | F | A | G | I | K | K | K | G | E | R | A | D | L | I | A | Y | L | K | K | A | T | N | E | - | - | - | - | - | - |
| Camelus\_ferus | M | - | - | - | - | - | - | - | - | - | - | - | - | - | - | - | - | - | - | - | - | - | - | - | - | - | - | - | - | - | - | - | - | - | - | - | - | - | - | - | - | - | - | - | - | - | - | - | - | - | - | - | - | - | - | - | - | - | - | - | - | - | - | - | - | - | - | - | - | - | - | - | - | - | - | - | - | - | - | - | - | - | - | - | - | - | - | - | - | - | - | - | - | - | - | - | - | - | D | A | K | A | - | R | N | - | - | - | - | - | - | - | - | - | - | - | - | - | - | - | - | C | L | L | - | - | - | - | - | - | - | - | - | - | - | - | - | - | - | - | - | - | - | - | - | - | - | - | - | - | - | - | - | - | - | - | - | - | - | - | - | - | - | - | - | - | - | - | - | - | - | - | - | - | - | - | - | - | - | - | - | - | - | - | - | - | - | - | - | - | - | - | - | - | - | - | - | - | - | - | - | - | - | - | - | - | - | - | - | - | - | - | - | - | - | - | - | - | - | - | - | - | - | - | - | - | - | - | - | - | - | - | - | - | - | - | - | - | - | - | - | - | - | - | - | Q | H | R | E | A | L | E | R | D | I | K | T | S | Y | - | I | M | D | H | M | I | S | N | G | V | L | T | V | S | E | E | E | K | V | K | N | E | - | - | - | - | - | - | - | - | - | - | - | - | - | - | - | - | - | P | T | Q | Q | Q | R | A | A | M | L | I | K | M | I | L | K | K | - | - | - | - | D | N | Y | S | Y | V | S | - | - | - | - | - | F | Y | N | A | L | - | L | H | E | G | - | - | - | - | - | - | Y | K | D | L | A | A | L | L | H | A | G | I | P | V | I | S | S | S | N | G | - | K | - | - | - | - | - | - | - | - | - | - | - | - | - | - | - | - | - | - | - | - | - | - | D | S | V | G | G | I | T | - | - | - | - | - | - | - | - | - | - | - | S | Y | V | R | T | V | L | C | E | G | G | - | - | - | - | - | - | - | V | P | Q | R | P | V | V | F | V | T | R | K | K | L | V | H | A | I | - | - | - | - | - | - | - | - | - | - | Q | Q | K | L | F | - | - | - | - | - | - | - | - | - | - | - | - | - | - | - | - | - | - | - | - | - | - | - | - | - | - | - | - | - | - | - | - | K | L | S | G | E | P | - | - | - | - | - | - | - | - | - | - | - | - | - | - | - | - | - | - | - | - | - | - | - | - | - | - | - | - | - | - | - | - | - | - | - | - | - | - | G | W | V | T | I | Y | G | M | A | G | C | G | K | S | V | L | A | A | E | A | V | - | R | D | H | S | L | L | E | V | C | F | P | G | G | V | H | W | V | S | V | G | - | - | - | - | - | - | - | K | - | - | - | - | - | - | Q | D | K | S | - | - | - | - | - | - | - | - | - | - | - | - | G | L | L | M | K | L | Q | N | L | C | T | R | L | D | - | - | - | - | Q | - | - | - | - | - | - | - | - | - | - | - | - | - | - | - | D | E | S | F | S | Q | R | L | P | L | N | I | - | - | - | - | - | - | - | - | - | - | - | - | - | - | - | - | E | E | A | K | D | R | L | R | I | L | M | L | R | - | - | - | - | - | - | - | - | - | - | - | - | K | H | P | R | S | L | L | - | - | I | L | D | D | I | W | D | P | W | V | L | K | A | F | D | - | N | Q | C | Q | I | L | L | T | T | R | D | K | S | V | T | D | S | V | - | M | G | P | - | - | K | Y | V | V | P | V | E | S | - | - | - | G | L | G | K | E | K | G | L | E | I | L | S | L | F | V | N | M | - | K | K | A | D | L | P | E | - | - | - | - | - | - | - | - | - | - | - | - | - | - | Q | A | H | S | I | I | K | E | C | K | - | - | - | - | - | - | - | - | - | - | - | - | - | - | - | - | - | - | - | G | - | S | P | L | V | V | S | L | I | G | A | L | L | R | D | F | P | N | - | - | - | - | - | - | - | - | - | - | R | W | E | Y | Y | L | - | K | Q | L | Q | N | K | Q | F | K | R | I | - | - | - | - | - | - | - | - | - | - | - | - | - | - | - | - | - | - | - | - | - | - | R | K | S | S | S | Y | D | - | - | - | - | Y | E | A | L | D | E | A | M | S | I | S | V | E | - | M | L | R | - | - | - | E | D | I | K | D | Y | - | Y | T | D | L | S | I | L | Q | K | D | V | K | V | P | T | K | V | L | C | I | L | W | D | - | - | - | - | M | E | T | E | E | V | E | D | I | L | Q | E | F | - | - | - | - | - | - | - | - | - | - | - | V | N | K | S | L | L | F | C | D | R | N | G | K | S | - | - | - | - | - | - | - | - | - | F | L | Y | Y | L | H | D | L | Q | V | D | F | L | T | E | K | - | - | N | R | S | Q | - | L | - | Q | D | L | H | K | K | I | I | T | Q | F | Q | R | - | H | R | - | - | - | - | - | - | R | L | H | T | L | S | P | D | - | Q | E | - | - | - | - | - | - | - | D | C | M | Y | W | Y | N | F | L | A | Y | H | M | A | S | A | N | - | - | - | - | M | H | K | E | L | C | A | L | M | F | S | L | D | W | I | K | A | K | T | E | L | V | G | P | A | - | - | - | - | - | - | - | - | H | L | I | H | E | F | V | E | Y | R | H | M | L | D | E | K | - | - | - | - | - | - | - | D | G | T | V | C | E | N | F | Q | - | - | - | E | F | L | S | L | N | G | H | L | L | G | R | Q | P | F | - | - | - | - | - | - | - | - | - | - | - | - | - | - | - | - | - | - | - | - | - | - | - | - | - | - | - | - | - | - | - | - | - | - | - | - | - | - | - | - | - | - | - | - | - | - | - | - | - | - | - | - | - | - | - | - | - | - | - | - | - | - | - | - | - | - | - | - | - | - | - | P | N | I | V | Q | L | G | L | C | E | L | E | T | S | E | V | Y | Q | Q | A | K | L | Q | A | K | Q | E | V | D | N | G | - | - | - | - | - | - | - | - | - | - | - | - | - | - | - | - | - | - | - | - | - | - | - | - | - | - | - | - | - | M | L | Y | L | - | - | - | - | - | - | - | - | - | - | E | W | I | N | K | - | - | - | - | - | - | - | K | N | I | K | N | L | S | R | - | - | - | - | - | - | - | - | - | - | L | V | V | C | P | H | T | D | - | - | - | - | - | - | - | - | - | - | - | - | - | - | - | A | V | Y | H | A | C | - | - | - | F | S | - | E | D | G | Q | R | I | A | - | - | - | - | - | - | - | - | - | S | C | G | A | D | K | T | L | Q | V | F | K | - | - | - | - | - | A | E | T | G | E | K | L | L | E | I | - | K | A | H | E | D | E | - | - | - | V | L | C | C | A | F | S | - | A | - | - | - | - | - | - | - | - | - | - | - | - | - | D | D | K | F | I | A | T | C | S | V | D | K | K | V | K | I | - | - | - | W | N | S | M | - | - | - | - | - | - | - | - | - | - | - | - | - | - | - | - | - | - | - | - | - | - | - | - | - | - | - | - | - | - | - | - | - | - | - | - | - | - | - | - | - | - | - | - | - | - | - | - | - | - | - | - | - | - | - | - | - | - | - | - | - | - | - | - | - | - | - | - | - | - | - | - | - | - | - | - | - | - | - | - | - | - | - | - | - | - | - | - | - | - | - | - | - | - | - | - | - | - | - | - | - | - | - | - | - | - | - | - | - | - | - | - | - | - | - | - | - | - | - | - | - | - | - | - | - | - | - | - | - | - | - | - | - | - | - | - | - | - | - | - | T | G | E | L | V | H | T | Y | - | D | E | H | S | - | - | - | - | - | E | Q | V | N | - | - | - | - | - | - | - | - | - | - | - | - | - | C | C | H | - | - | - | - | - | - | - | - | - | - | - | - | - | F | T | N | N | S | - | - | - | - | - | - | - | - | - | - | - | - | - | - | - | - | N | H | - | L | L | L | A | T | G | S | S | - | - | - | - | D | C | F | L | - | - | - | - | - | - | - | - | - | - | - | - | K | L | W | D | V | - | - | - | N | Q | K | E | C | R | N | T | M | F | - | - | - | - | - | - | - | - | - | - | - | - | - | - | - | - | - | G | H | T | N | S | - | - | - | - | - | - | - | - | - | - | - | - | - | - | - | - | V | N | H | C | R | F | S | P | D | D | K | - | - | - | - | - | - | - | - | - | - | - | - | - | - | - | - | - | - | - | - | - | - | - | - | - | - | - | - | L | L | A | S | C | S | A | D | G | T | L | K | L | - | - | - | - | - | - | - | - | - | - | - | - | - | - | - | - | - | - | - | W | D | - | V | K | S | A | N | E | R | K | S | - | - | - | - | - | - | I | N | - | - | - | - | - | - | - | - | - | - | - | - | - | - | - | - | - | - | - | - | - | - | V | K | H | F | - | - | - | - | - | - | - | - | F | L | N | S | E | E | P | Q | E | - | - | - | - | - | - | - | - | - | - | - | - | - | - | - | - | - | - | - | - | - | - | - | - | - | - | - | D | M | E | - | V | I | V | K | C | C | - | - | - | - | - | - | - | - | - | - | - | - | - | - | - | - | S | W | S | A | D | G | A | R | I | M | V | A | A | K | N | K | V | - | - | - | - | - | - | - | - | - | - | - | F | L | F | D | I | H | T | - | - | - | S | G | - | - | - | - | - | - | - | - | - | - | - | - | - | - | - | - | - | - | - | - | - | - | - | - | - | - | - | - | - | - | - | - | - | - | - | - | - | - | - | - | - | - | - | - | - | - | - | - | - | - | - | - | - | - | - | - | - | - | - | - | - | - | - | - | - | - | - | - | - | - | - | - | - | - | - | - | - | - | - | - | - | - | - | - | - | - | - | - | - | - | - | - | - | - | - | - | - | - | - | - | - | - | - | - | - | - | - | - | - | - | - | - | - | - | - | - | - | - | - | - | - | - | - | - | - | - | L | L | A | - | - | - | - | - | - | - | - | - | - | - | - | - | - | - | - | - | - | - | - | - | - | - | - | - | - | - | - | - | - | - | - | - | - | - | - | - | - | - | - | - | - | - | - | - | - | - | - | - | - | - | - | - | - | - | - | - | - | - | - | - | - | - | - | - | - | - | - | - | - | - | - | - | - | - | - | - | - | - | - | - | - | - | - | - | - | - | - | - | - | - | - | - | - | - | - | - | - | - | - | - | - | - | - | - | - | - | - | - | - | - | - | - | - | - | - | - | - | - | - | - | - | - | - | - | - | - | - | - | - | - | - | - | - | - | - | - | - | - | - | - | - | - | - | - | - | - | - | - | - | - | - | - | - | - | - | - | - | - | - | - | - | - | - | - | - | - | - | - | - | - | - | - | - | - | - | - | - | - | - | - | - | - | E | I | - | - | - | H | T | G | H | H | S | T | V | Q | Y | C | - | - | - | - | - | - | - | - | - | - | - | - | - | - | - | - | - | - | - | - | - | - | - | - | - | D | F | S | P | H | N | H | L | A | L | V | A | L | S | Q | - | - | - | - | - | - | Y | C | V | E | - | - | - | - | - | - | - | - | - | - | - | - | - | - | - | - | - | L | W | N | V | D | S | C | - | - | - | - | - | - | - | - | - | - | - | - | - | - | - | - | - | - | - | - | - | - | - | - | - | - | - | - | - | L | K | V | A | D | C | R | G | H | L | S | - | - | - | - | - | - | - | - | - | - | - | - | - | - | - | - | - | - | - | - | - | - | - | - | - | - | - | - | - | - | - | - | - | - | - | - | - | - | - | - | - | - | - | - | W | V | H | C | V | M | F | S | - | - | - | - | - | - | - | - | - | - | - | P | D | G | S | - | - | - | - | - | - | - | - | - | - | S | F | L | T | S | S | D | D | Q | - | - | - | - | - | - | - | - | - | - | - | - | - | - | T | I | R | L | - | W | - | - | - | - | - | - | - | - | - | - | - | - | - | - | - | - | - | - | - | - | - | - | - | - | - | - | - | - | - | - | - | - | - | - | - | - | - | - | E | T | K | K | V | C | K | N | S | A | I | V | L | K | Q | E | - | - | - | - | - | - | - | - | - | - | - | - | - | - | - | - | I | D | V | V | F | - | - | - | - | - | - | - | - | - | - | - | - | - | - | - | - | - | - | - | - | - | - | - | - | - | - | - | - | - | - | - | - | - | - | - | - | - | - | - | - | - | - | - | - | - | - | - | - | - | - | - | - | - | - | Q | E | N | E | V | M | - | - | - | - | - | - | - | - | - | - | - | - | - | - | - | - | - | - | - | - | - | - | - | - | - | - | - | - | - | - | - | - | - | - | - | - | V | L | A | V | D | N | I | R | R | L | Q | L | - | I | N | G | K | T | - | - | - | - | - | - | - | - | - | - | G | Q | I | D | Y | L | T | E | A | - | - | - | - | - | - | - | - | - | - | - | - | - | - | - | - | - | - | - | - | - | - | H | - | - | - | - | - | - | - | - | - | - | - | V | S | C | C | C | L | S | P | H | L | Q | - | - | - | - | - | - | - | - | - | Y | V | A | F | G | G | - | - | - | - | - | - | - | - | - | - | - | - | - | - | - | - | - | - | - | - | - | - | - | - | - | - | - | - | - | - | - | - | - | - | - | - | E | - | - | - | - | - | - | - | - | - | D | G | A | - | - | - | - | - | - | - | - | - | - | I | - | - | - | - | - | - | - | - | - | - | - | - | - | - | - | - | - | - | - | - | - | - | - | - | - | - | - | - | - | - | - | - | - | - | - | - | - | - | - | Q | I | L | E | L | L | N | N | R | V | F | Q | S | K | I | G | - | - | - | - | - | H | - | - | - | - | - | - | - | - | - | - | - | - | K | N | T | V | R | H | I | Q | F | T | - | - | - | - | - | - | - | - | - | - | - | - | - | - | - | - | - | - | - | - | - | - | - | - | - | - | - | - | - | - | - | - | - | - | - | - | - | A | D | G | K | N | L | I | S | S | S | - | D | D | C | - | - | - | - | - | - | - | - | T | I | Q | - | I | - | - | - | - | - | - | - | - | - | - | - | - | - | - | - | - | - | - | - | - | - | - | - | - | - | - | - | - | - | - | - | - | - | - | - | - | - | - | - | W | N | - | - | - | - | - | - | - | - | - | - | W | Q | S | E | E | Y | V | F | - | - | - | - | - | - | - | - | - | - | - | - | - | - | - | - | - | - | - | - | - | - | - | - | - | - | - | - | - | - | - | - | - | - | - | - | - | - | - | - | - | - | - | - | - | - | - | - | - | - | L | Q | A | H | Q | - | - | - | - | - | - | - | - | - | - | - | - | - | - | - | - | - | - | - | - | - | - | - | - | - | - | E | A | V | K | D | F | K | L | L | - | - | - | - | - | - | - | - | - | - | - | - | - | - | - | - | - | - | - | - | - | - | - | K | N | - | S | R | L | L | - | S | W | - | - | - | S | F | D | G | T | V | - | - | - | - | - | - | - | - | - | - | - | - | - | - | - | - | - | - | - | - | - | - | - | - | - | - | - | - | - | - | - | - | - | - | - | K | V | W | N | F | I | T | G | R | I | - | - | - | - | - | E | K | D | F | V | C | - | - | - | - | - | - | - | H | Q | - | - | - | - | - | - | - | - | - | - | - | - | - | - | - | - | - | - | - | D | T | V | L | - | - | - | - | - | - | - | - | - | - | - | - | - | - | - | S | C | D | - | - | - | - | - | - | - | - | - | - | - | - | - | - | - | I | S | P | D | A | T | - | K | F | S | S | T | - | S | A | D | K | T | - | - | - | - | - | - | A | K | I | W | S | F | E | L | I | - | - | - | - | - | - | - | - | - | - | - | - | - | - | - | - | - | - | - | - | - | - | - | - | S | P | L | H | E | - | - | - | - | - | - | - | - | L | - | - | - | - | R | G | H | K | G | C | V | - | - | - | - | - | - | - | - | - | R | C | S | - | A | F | S | - | - | - | - | - | - | - | - | - | V | D | S | T | - | L | L | A | T | - | - | - | - | - | - | - | - | G | D | D | N | G | E | I | R | I | W | N | V | S | N | G | E | L | L | - | - | - | - | - | - | - | - | - | - | - | - | - | - | - | - | - | - | - | - | - | - | - | - | - | - | - | - | - | - | - | - | - | - | - | - | - | - | - | - | - | - | - | - | - | - | - | - | - | - | - | - | - | - | - | - | H | L | C | A | P | I | S | V | E | - | - | - | - | - | - | - | - | - | - | - | - | - | - | - | - | - | - | - | - | E | G | A | A | T | H | G | G | W | - | - | - | - | - | - | - | - | V | T | D | L | C | F | S | P | N | S | K | - | - | - | - | - | - | - | - | - | - | - | - | - | - | - | - | - | - | - | - | - | - | - | - | - | - | - | - | - | - | - | - | - | - | - | - | - | - | - | - | - | - | - | - | - | M | L | V | S | - | - | A | G | G | Y | L | K | - | - | - | - | - | - | - | - | W | W | N | V | V | T | - | - | - | - | - | - | - | - | - | - | - | - | - | - | - | - | - | - | - | - | - | - | - | - | - | - | - | - | - | - | - | - | - | - | - | - | - | - | - | - | - | - | - | - | - | - | - | - | - | - | - | - | - | - | - | - | - | - | - | - | - | - | - | - | - | - | - | - | - | - | - | - | - | - | - | - | - | - | - | - | - | - | - | - | - | - | - | - | - | - | - | - | - | - | - | - | - | - | - | - | - | - | - | - | - | - | - | - | - | - | - | - | - | - | - | - | - | - | - | - | - | - | - | - | - | - | - | - | - | - | - | - | - | - | - | - | - | - | - | - | - | - | - | - | - | - | - | - | - | - | - | - | - | - | - | - | - | - | - | - | - | - | - | - | - | - | - | - | - | - | - | - | - | - | - | - | - | - | - | - | - | - | - | - | - | - | - | - | - | - | - | - | - | - | - | - | - | - | - | - | - | - | - | - | - | - | - | - | - | - | - | - | - | - | - | - | - | - | - | - | - | - | - | - | - | - | - | - | - | - | - | - | - | - | - | - | - | - | - | - | - | - | - | - | - | - | - | - | - | - | - | - | - | - | - | - | - | - | - | - | - | - | - | - | - | - | - | G | E | S | S | Q | T | F | Y | T | - | - | - | - | - | N | G | - | - | - | T | N | L | K | K | I | - | - | - | H | V | S | P | - | - | - | - | D | F | K | T | - | - | - | - | - | - | - | - | - | - | - | - | - | - | - | - | - | - | - | - | - | - | - | - | - | - | Y | V | T | V | D | - | N | L | G | I | L | - | - | Y | I | L | K | I | - | - | - | - | - | - | - | - | - | - | - | - | - | - | - | - | - | - | - | - | - | - | - | - | - | - | - | - | - | - | - | - | - | - | L | E | - | - | - | - | - | - | - | - | - | - | - | - | - | - | - | - | - | - | - | - | - | - | - | - | - | - | - | - | - | - | - | - | - | - | - | - | - | - | - | - | - | - | - | - | - | - | - | - | - | - | - | - | M | - | - | - | - | - | - | - | - | - | - | - | - | - | - | - | - | - | - | - | - | - | - | - | - | - | - | - | - | - | - | - | - | - | - | - | - | - | - | - | - | - | - | - | - | - | - | - | - | - | - | - | - | - | - | - | - | - | - | - | - | - | - | - | - | - | - | - | - | - | - | - | - | - | - | - | - | - | - | - | - | - | - | - | - | - | - | - | - | - | - | - | - | - | - | - | - | - | - | - | - | - | - | - | - | - | - | - | - | - | - | - | - | - | - | - | - | - | - | - | - | - | - | - | - | - | - | - | - | - | - | - | - | - | - | - | - | - | - | - | - | - | - | - | - | - | - | - | - | - | - | - | - | - | - | - | - | - | - | - | - | - | - | - | - | - | - | - | - | - | - | - | - | - | - | - | - | - | - | - | - | - | - | - | - | - | - | - | - | - | - | - | - | - | - | - | - | - | - | - | - | - | - | - | - | - | - | - | - | - | - | - | - | - | - | - | - | - | - | - | - | I | E | E | - | - | - | - | - | - | - | - | - | - | - | - | - | - | - | - | - | - | - | - | - | - | - | - | - | I | Q | R | A | G | - | - | - | - | - | - | - | - | S | G | S | R | R | D | Q | - | A | R | Q | L | I | T | D | L | E | T | R | G | - | S | Q | A | L | S | L | F | I | S | C | L | E | D | T | G | Q | N | M | L | A | S | L | L | R | T | S | S | Q | A | A | - | - | - | - | R | Q | H | P | - | E | A | I | - | - | - | - | - | - | - | - | - | - | - | - | - | - | - | - | - | - | - | - | - | - | - | - | - | - | R | P | L | D | - | - | - | - | - | - | - | - | - | - | - | - | - | - | - | L | K | P | V | - | - | - | - | - | - | - | - | - | - | - | - | - | - | - | - | - | - | - | - | - | - | - | - | - | - | - | - | - | - | - | - | - | - | - | - | - | - | - | V | L | R | P | E | G | L | K | P | E | S | L | G | - | V | G | K | Q | D | P | L | M | P | - | T | - | P | - | - | - | - | - | - | - | - | - | - | - | - | - | - | - | - | - | - | - | - | - | - | - | - | - | - | - | - | - | - | - | - | - | - | - | - | - | - | - | - | - | - | - | - | - | - | - | - | - | - | - | G | K | L | V | P | V | - | - | - | - | - | - | - | - | - | - | - | - | - | - | - | - | - | - | - | - | - | - | - | - | - | - | V | L | - | - | - | - | - | - | - | - | - | - | - | - | - | - | - | - | - | - | - | - | - | - | - | - | - | - | - | - | - | - | - | - | - | - | - | - | - | - | - | - | - | - | - | - | - | - | - | - | - | - | - | - | - | - | - | - | - | - | - | - | - | - | - | - | - | - | - | - | G | P | K | E | L | W | P | - | - | - | - | - | - | - | - | - | - | - | - | - | - | - | - | - | - | - | - | - | - | - | - | - | - | A | K | L | R | P | - | - | - | - | - | - | E | V | L | R | P | - | E | V | - | - | - | - | - | - | - | - | - | - | - | - | - | - | - | - | - | - | P | R | P | V | D | I | G | S | G | G | L | - | - | - | - | - | - | - | - | - | - | - | - | - | S | D | V | C | - | - | - | - | - | - | - | - | - | - | A | - | Q | D | R | A | K | G | N | A | N | - | - | - | - | - | L | A | Y | V | - | - | L | S | A | D | - | P | C | G | R | C | L | I | V | N | N | V | H | F | R | R | E | S | - | - | G | L | R | T | R | T | G | S | S | V | D | C | E | R | L | Q | R | R | F | - | H | L | L | H | F | V | V | E | V | E | C | D | L | T | A | K | Q | M | V | Q | A | L | V | E | L | A | R | R | D | H | - | S | A | L | D | C | C | V | V | V | I | L | S | H | G | C | Q | A | S | H | L | Q | F | P | G | A | - | V | Y | G | T | D | G | C | P | V | S | V | E | R | I | V | N | I | F | N | G | T | G | C | P | S | L | R | G | K | P | K | L | F | F | I | Q | A | C | - | - | G | G | E | Q | K | D | H | G | F | E | V | A | S | T | - | - | - | F | P | E | G | - | Q | T | P | D | S | D | - | - | - | - | - | - | - | P | E | P | D | A | T | P | F | Q | E | G | P | G | T | V | - | - | - | - | - | - | - | - | D | Q | P | D | A | V | - | - | - | - | - | - | - | S | S | L | - | - | - | P | T | P | S | D | I | L | V | S | Y | S | T | F | P | - | - | G | - | - | - | - | - | - | - | - | F | V | S | W | R | D | T | K | S | G | S | W | Y | I | E | T | L | D | S | V | F | E | Q | W | A | H | - | C | - | - | - | - | - | - | - | - | - | - | - | - | - | - | - | - | - | - | - | - | - | - | - | - | - | - | - | - | - | - | - | E | D | L | Q | T | L | L | - | - | - | - | - | L | R | - | - | - | - | - | - | - | - | - | - | - | - | V | A | N | A | V | - | - | - | - | - | S | V | K | - | - | - | - | - | - | - | G | I | Y | - | K | Q | I | - | - | P | G | - | - | - | - | - | - | - | - | - | - | - | - | - | - | - | C | F | N | F | - | - | - | - | - | - | - | - | - | L | R | K | K | L | F | F | K | T | - | - | - | - | - | - | - | - | - | - | - | - | - | - | - | - | - | - | - | - | - | - | - | - | - | - | - | - | - | - | - | - | - | - | - | - | - | - | M | - | - | - | - | - | - | - | - | - | - | - | - | - | - | - | - | - | - | - | - | - | - | - | - | - | - | - | - | - | - | - | - | - | - | - | - | - | - | - | - | - | - | - | - | - | - | - | - | - | - | - | - | G | D | V | E | K | G | K | K | I | F | V | Q | K | C | A | Q | C | H | T | V | E | K | G | G | K | H | K | T | G | P | N | L | H | G | L | F | G | R | K | T | G | Q | A | V | G | F | S | Y | T | D | A | N | K | N | K | G | I | T | W | G | E | E | T | L | M | E | Y | L | E | N | P | K | K | Y | I | P | G | T | K | M | I | F | A | G | I | K | K | K | G | E | R | A | D | L | I | A | Y | L | K | K | A | T | N | E | - | - | - | - | - | - |
| Camelus\_dromedarius | M | - | - | - | - | - | - | - | - | - | - | - | - | - | - | - | - | - | - | - | - | - | - | - | - | - | - | - | - | - | - | - | - | - | - | - | - | - | - | - | - | - | - | - | - | - | - | - | - | - | - | - | - | - | - | - | - | - | - | - | - | - | - | - | - | - | - | - | - | - | - | - | - | - | - | - | - | - | - | - | - | - | - | - | - | - | - | - | - | - | - | - | - | - | - | - | - | - | D | A | K | A | - | R | N | - | - | - | - | - | - | - | - | - | - | - | - | - | - | - | - | C | L | L | - | - | - | - | - | - | - | - | - | - | - | - | - | - | - | - | - | - | - | - | - | - | - | - | - | - | - | - | - | - | - | - | - | - | - | - | - | - | - | - | - | - | - | - | - | - | - | - | - | - | - | - | - | - | - | - | - | - | - | - | - | - | - | - | - | - | - | - | - | - | - | - | - | - | - | - | - | - | - | - | - | - | - | - | - | - | - | - | - | - | - | - | - | - | - | - | - | - | - | - | - | - | - | - | - | - | - | - | - | - | - | - | - | - | - | - | - | - | - | - | - | Q | H | R | E | A | L | E | R | D | I | K | T | S | Y | - | I | M | D | H | M | I | S | N | G | V | L | T | V | S | E | E | E | K | V | K | N | E | - | - | - | - | - | - | - | - | - | - | - | - | - | - | - | - | - | P | T | Q | Q | Q | R | A | A | M | L | I | K | M | I | L | K | K | - | - | - | - | D | N | Y | S | Y | V | S | - | - | - | - | - | F | Y | N | A | L | - | L | H | E | G | - | - | - | - | - | - | Y | K | D | L | A | A | L | L | H | A | G | I | P | V | I | S | S | S | N | G | - | K | - | - | - | - | - | - | - | - | - | - | - | - | - | - | - | - | - | - | - | - | - | - | D | S | V | G | G | I | T | - | - | - | - | - | - | - | - | - | - | - | S | Y | V | R | T | V | L | C | E | G | G | - | - | - | - | - | - | - | V | P | Q | R | P | V | V | F | V | T | R | K | K | L | V | H | A | I | - | - | - | - | - | - | - | - | - | - | Q | Q | K | L | F | - | - | - | - | - | - | - | - | - | - | - | - | - | - | - | - | - | - | - | - | - | - | - | - | - | - | - | - | - | - | - | - | K | L | S | G | E | P | - | - | - | - | - | - | - | - | - | - | - | - | - | - | - | - | - | - | - | - | - | - | - | - | - | - | - | - | - | - | - | - | - | - | - | - | - | - | G | W | V | T | I | Y | G | M | A | G | C | G | K | S | V | L | A | A | E | A | V | - | R | D | H | S | L | L | E | V | C | F | P | G | G | V | H | W | V | S | V | G | - | - | - | - | - | - | - | K | - | - | - | - | - | - | Q | D | K | S | - | - | - | - | - | - | - | - | - | - | - | - | G | L | L | M | K | L | Q | N | L | C | T | R | L | D | - | - | - | - | Q | - | - | - | - | - | - | - | - | - | - | - | - | - | - | - | D | E | S | F | S | Q | R | L | P | L | N | I | - | - | - | - | - | - | - | - | - | - | - | - | - | - | - | - | E | E | A | K | D | R | L | R | I | L | M | L | R | - | - | - | - | - | - | - | - | - | - | - | - | K | H | P | R | S | L | L | - | - | I | L | D | D | I | W | D | P | W | V | L | K | A | F | D | - | N | Q | C | Q | I | L | L | T | T | R | D | K | S | V | T | D | S | V | - | M | G | P | - | - | K | Y | V | V | P | V | E | S | - | - | - | G | L | G | K | E | K | G | L | E | I | L | S | L | F | V | N | M | - | K | K | A | D | L | P | E | - | - | - | - | - | - | - | - | - | - | - | - | - | - | Q | A | H | S | I | I | K | E | C | K | - | - | - | - | - | - | - | - | - | - | - | - | - | - | - | - | - | - | - | G | - | S | P | L | V | V | S | L | I | G | A | L | L | R | D | F | P | N | - | - | - | - | - | - | - | - | - | - | R | W | E | Y | Y | L | - | K | Q | L | Q | N | K | Q | F | K | R | I | - | - | - | - | - | - | - | - | - | - | - | - | - | - | - | - | - | - | - | - | - | - | R | K | S | S | S | Y | D | - | - | - | - | Y | E | A | L | D | E | A | M | S | I | S | V | E | - | M | L | R | - | - | - | E | D | I | K | D | Y | - | Y | T | D | L | S | I | L | Q | K | D | V | K | V | P | T | K | V | L | C | I | L | W | D | - | - | - | - | M | E | T | E | E | V | E | D | I | L | Q | E | F | - | - | - | - | - | - | - | - | - | - | - | V | N | K | S | L | L | F | C | D | R | N | G | K | S | - | - | - | - | - | - | - | - | - | F | L | Y | Y | L | H | D | L | Q | V | D | F | L | T | E | K | - | - | N | R | S | Q | - | L | - | Q | D | L | H | K | K | I | I | T | Q | F | Q | R | - | H | R | - | - | - | - | - | - | R | L | H | T | L | S | P | D | - | Q | E | - | - | - | - | - | - | - | D | C | M | Y | W | Y | N | F | L | A | Y | H | M | A | S | A | N | - | - | - | - | M | H | K | E | L | C | A | L | M | F | S | L | D | W | I | K | A | K | T | E | L | V | G | P | A | - | - | - | - | - | - | - | - | H | L | I | H | E | F | V | E | Y | R | H | M | L | D | E | K | - | - | - | - | - | - | - | D | G | T | V | C | E | N | F | Q | - | - | - | E | F | L | S | L | N | G | H | L | L | G | R | Q | P | F | - | - | - | - | - | - | - | - | - | - | - | - | - | - | - | - | - | - | - | - | - | - | - | - | - | - | - | - | - | - | - | - | - | - | - | - | - | - | - | - | - | - | - | - | - | - | - | - | - | - | - | - | - | - | - | - | - | - | - | - | - | - | - | - | - | - | - | - | - | - | - | P | N | I | V | Q | L | G | L | C | E | L | E | T | S | E | V | Y | Q | Q | A | K | L | Q | A | K | Q | E | V | D | N | G | - | - | - | - | - | - | - | - | - | - | - | - | - | - | - | - | - | - | - | - | - | - | - | - | - | - | - | - | - | M | L | Y | L | - | - | - | - | - | - | - | - | - | - | E | W | I | N | K | - | - | - | - | - | - | - | K | N | I | K | N | L | S | R | - | - | - | - | - | - | - | - | - | - | L | V | V | C | P | H | T | D | - | - | - | - | - | - | - | - | - | - | - | - | - | - | - | A | V | Y | H | A | C | - | - | - | F | S | - | E | D | G | Q | R | I | A | - | - | - | - | - | - | - | - | - | S | C | G | A | D | K | T | L | Q | V | F | K | - | - | - | - | - | A | E | T | G | E | K | L | L | E | I | - | K | A | H | E | D | E | - | - | - | V | L | C | C | A | F | S | - | A | - | - | - | - | - | - | - | - | - | - | - | - | - | D | D | K | F | I | A | T | C | S | V | D | K | K | V | K | I | - | - | - | W | N | S | M | - | - | - | - | - | - | - | - | - | - | - | - | - | - | - | - | - | - | - | - | - | - | - | - | - | - | - | - | - | - | - | - | - | - | - | - | - | - | - | - | - | - | - | - | - | - | - | - | - | - | - | - | - | - | - | - | - | - | - | - | - | - | - | - | - | - | - | - | - | - | - | - | - | - | - | - | - | - | - | - | - | - | - | - | - | - | - | - | - | - | - | - | - | - | - | - | - | - | - | - | - | - | - | - | - | - | - | - | - | - | - | - | - | - | - | - | - | - | - | - | - | - | - | - | - | - | - | - | - | - | - | - | - | - | - | - | - | - | - | - | T | G | E | L | V | H | T | Y | - | D | E | H | S | - | - | - | - | - | E | Q | V | N | - | - | - | - | - | - | - | - | - | - | - | - | - | C | C | H | - | - | - | - | - | - | - | - | - | - | - | - | - | F | T | N | N | S | - | - | - | - | - | - | - | - | - | - | - | - | - | - | - | - | N | H | - | L | L | L | A | T | G | S | S | - | - | - | - | D | C | F | L | - | - | - | - | - | - | - | - | - | - | - | - | K | L | W | D | L | - | - | - | N | Q | K | E | C | R | N | T | M | F | - | - | - | - | - | - | - | - | - | - | - | - | - | - | - | - | - | G | H | T | N | S | - | - | - | - | - | - | - | - | - | - | - | - | - | - | - | - | V | N | H | C | R | F | S | P | D | D | K | - | - | - | - | - | - | - | - | - | - | - | - | - | - | - | - | - | - | - | - | - | - | - | - | - | - | - | - | L | L | A | S | C | S | A | D | G | T | L | K | L | - | - | - | - | - | - | - | - | - | - | - | - | - | - | - | - | - | - | - | W | D | - | V | K | S | A | N | E | R | K | S | - | - | - | - | - | - | I | N | - | - | - | - | - | - | - | - | - | - | - | - | - | - | - | - | - | - | - | - | - | - | V | K | H | F | - | - | - | - | - | - | - | - | F | L | N | S | E | E | P | Q | E | - | - | - | - | - | - | - | - | - | - | - | - | - | - | - | - | - | - | - | - | - | - | - | - | - | - | - | D | M | E | - | V | I | V | K | C | C | - | - | - | - | - | - | - | - | - | - | - | - | - | - | - | - | S | W | S | A | D | G | A | R | I | M | V | A | A | K | N | K | V | - | - | - | - | - | - | - | - | - | - | - | F | L | F | D | I | H | T | - | - | - | S | G | - | - | - | - | - | - | - | - | - | - | - | - | - | - | - | - | - | - | - | - | - | - | - | - | - | - | - | - | - | - | - | - | - | - | - | - | - | - | - | - | - | - | - | - | - | - | - | - | - | - | - | - | - | - | - | - | - | - | - | - | - | - | - | - | - | - | - | - | - | - | - | - | - | - | - | - | - | - | - | - | - | - | - | - | - | - | - | - | - | - | - | - | - | - | - | - | - | - | - | - | - | - | - | - | - | - | - | - | - | - | - | - | - | - | - | - | - | - | - | - | - | - | - | - | - | - | L | L | A | - | - | - | - | - | - | - | - | - | - | - | - | - | - | - | - | - | - | - | - | - | - | - | - | - | - | - | - | - | - | - | - | - | - | - | - | - | - | - | - | - | - | - | - | - | - | - | - | - | - | - | - | - | - | - | - | - | - | - | - | - | - | - | - | - | - | - | - | - | - | - | - | - | - | - | - | - | - | - | - | - | - | - | - | - | - | - | - | - | - | - | - | - | - | - | - | - | - | - | - | - | - | - | - | - | - | - | - | - | - | - | - | - | - | - | - | - | - | - | - | - | - | - | - | - | - | - | - | - | - | - | - | - | - | - | - | - | - | - | - | - | - | - | - | - | - | - | - | - | - | - | - | - | - | - | - | - | - | - | - | - | - | - | - | - | - | - | - | - | - | - | - | - | - | - | - | - | - | - | - | - | - | - | E | I | - | - | - | H | T | G | H | H | S | T | V | Q | Y | C | - | - | - | - | - | - | - | - | - | - | - | - | - | - | - | - | - | - | - | - | - | - | - | - | - | D | F | S | P | H | N | H | L | A | L | V | A | L | S | Q | - | - | - | - | - | - | Y | C | V | E | - | - | - | - | - | - | - | - | - | - | - | - | - | - | - | - | - | L | W | N | V | D | S | C | - | - | - | - | - | - | - | - | - | - | - | - | - | - | - | - | - | - | - | - | - | - | - | - | - | - | - | - | - | L | K | V | A | D | C | R | G | H | L | S | - | - | - | - | - | - | - | - | - | - | - | - | - | - | - | - | - | - | - | - | - | - | - | - | - | - | - | - | - | - | - | - | - | - | - | - | - | - | - | - | - | - | - | - | W | V | H | C | V | T | F | S | - | - | - | - | - | - | - | - | - | - | - | P | D | G | S | - | - | - | - | - | - | - | - | - | - | S | F | L | T | S | S | D | D | Q | - | - | - | - | - | - | - | - | - | - | - | - | - | - | T | I | R | L | - | W | - | - | - | - | - | - | - | - | - | - | - | - | - | - | - | - | - | - | - | - | - | - | - | - | - | - | - | - | - | - | - | - | - | - | - | - | - | - | E | T | K | K | V | C | K | N | S | A | I | V | L | K | Q | E | - | - | - | - | - | - | - | - | - | - | - | - | - | - | - | - | I | D | V | V | F | - | - | - | - | - | - | - | - | - | - | - | - | - | - | - | - | - | - | - | - | - | - | - | - | - | - | - | - | - | - | - | - | - | - | - | - | - | - | - | - | - | - | - | - | - | - | - | - | - | - | - | - | - | - | Q | E | N | E | V | M | - | - | - | - | - | - | - | - | - | - | - | - | - | - | - | - | - | - | - | - | - | - | - | - | - | - | - | - | - | - | - | - | - | - | - | - | V | L | A | V | D | N | I | R | R | L | Q | L | - | I | N | G | K | T | - | - | - | - | - | - | - | - | - | - | G | Q | I | D | Y | L | T | E | A | - | - | - | - | - | - | - | - | - | - | - | - | - | - | - | - | - | - | - | - | - | - | H | - | - | - | - | - | - | - | - | - | - | - | V | S | C | C | C | L | S | P | H | L | Q | - | - | - | - | - | - | - | - | - | Y | V | A | F | G | G | - | - | - | - | - | - | - | - | - | - | - | - | - | - | - | - | - | - | - | - | - | - | - | - | - | - | - | - | - | - | - | - | - | - | - | - | E | - | - | - | - | - | - | - | - | - | D | G | A | - | - | - | - | - | - | - | - | - | - | I | - | - | - | - | - | - | - | - | - | - | - | - | - | - | - | - | - | - | - | - | - | - | - | - | - | - | - | - | - | - | - | - | - | - | - | - | - | - | - | Q | I | L | E | L | L | N | N | R | V | F | Q | S | K | I | G | - | - | - | - | - | H | - | - | - | - | - | - | - | - | - | - | - | - | K | N | T | V | R | H | I | Q | F | T | - | - | - | - | - | - | - | - | - | - | - | - | - | - | - | - | - | - | - | - | - | - | - | - | - | - | - | - | - | - | - | - | - | - | - | - | - | A | D | G | K | N | L | I | S | S | S | - | D | D | C | - | - | - | - | - | - | - | - | T | I | Q | - | I | - | - | - | - | - | - | - | - | - | - | - | - | - | - | - | - | - | - | - | - | - | - | - | - | - | - | - | - | - | - | - | - | - | - | - | - | - | - | - | W | N | - | - | - | - | - | - | - | - | - | - | W | Q | S | E | E | Y | V | F | - | - | - | - | - | - | - | - | - | - | - | - | - | - | - | - | - | - | - | - | - | - | - | - | - | - | - | - | - | - | - | - | - | - | - | - | - | - | - | - | - | - | - | - | - | - | - | - | - | - | L | Q | A | H | Q | - | - | - | - | - | - | - | - | - | - | - | - | - | - | - | - | - | - | - | - | - | - | - | - | - | - | E | A | V | K | D | F | K | L | L | - | - | - | - | - | - | - | - | - | - | - | - | - | - | - | - | - | - | - | - | - | - | - | K | N | - | S | R | L | L | - | S | W | - | - | - | S | F | D | G | T | V | - | - | - | - | - | - | - | - | - | - | - | - | - | - | - | - | - | - | - | - | - | - | - | - | - | - | - | - | - | - | - | - | - | - | - | K | V | W | N | F | I | T | G | R | I | - | - | - | - | - | E | K | D | F | V | C | - | - | - | - | - | - | - | H | Q | - | - | - | - | - | - | - | - | - | - | - | - | - | - | - | - | - | - | - | D | T | V | L | - | - | - | - | - | - | - | - | - | - | - | - | - | - | - | S | C | D | - | - | - | - | - | - | - | - | - | - | - | - | - | - | - | I | S | P | D | A | T | - | K | F | S | S | T | - | S | A | D | K | T | - | - | - | - | - | - | A | K | I | W | S | F | E | L | I | - | - | - | - | - | - | - | - | - | - | - | - | - | - | - | - | - | - | - | - | - | - | - | - | S | P | L | H | E | - | - | - | - | - | - | - | - | L | - | - | - | - | R | G | H | K | G | C | V | - | - | - | - | - | - | - | - | - | R | C | S | - | A | F | S | - | - | - | - | - | - | - | - | - | V | D | S | T | - | L | L | A | T | - | - | - | - | - | - | - | - | G | D | D | N | G | E | I | R | I | W | N | V | S | N | G | E | L | L | - | - | - | - | - | - | - | - | - | - | - | - | - | - | - | - | - | - | - | - | - | - | - | - | - | - | - | - | - | - | - | - | - | - | - | - | - | - | - | - | - | - | - | - | - | - | - | - | - | - | - | - | - | - | - | - | H | L | C | A | P | I | S | V | E | - | - | - | - | - | - | - | - | - | - | - | - | - | - | - | - | - | - | - | - | E | G | A | A | T | H | G | G | W | - | - | - | - | - | - | - | - | V | T | D | L | C | F | S | P | N | S | K | - | - | - | - | - | - | - | - | - | - | - | - | - | - | - | - | - | - | - | - | - | - | - | - | - | - | - | - | - | - | - | - | - | - | - | - | - | - | - | - | - | - | - | - | - | M | L | V | S | - | - | A | G | G | Y | L | K | - | - | - | - | - | - | - | - | W | W | N | V | V | T | - | - | - | - | - | - | - | - | - | - | - | - | - | - | - | - | - | - | - | - | - | - | - | - | - | - | - | - | - | - | - | - | - | - | - | - | - | - | - | - | - | - | - | - | - | - | - | - | - | - | - | - | - | - | - | - | - | - | - | - | - | - | - | - | - | - | - | - | - | - | - | - | - | - | - | - | - | - | - | - | - | - | - | - | - | - | - | - | - | - | - | - | - | - | - | - | - | - | - | - | - | - | - | - | - | - | - | - | - | - | - | - | - | - | - | - | - | - | - | - | - | - | - | - | - | - | - | - | - | - | - | - | - | - | - | - | - | - | - | - | - | - | - | - | - | - | - | - | - | - | - | - | - | - | - | - | - | - | - | - | - | - | - | - | - | - | - | - | - | - | - | - | - | - | - | - | - | - | - | - | - | - | - | - | - | - | - | - | - | - | - | - | - | - | - | - | - | - | - | - | - | - | - | - | - | - | - | - | - | - | - | - | - | - | - | - | - | - | - | - | - | - | - | - | - | - | - | - | - | - | - | - | - | - | - | - | - | - | - | - | - | - | - | - | - | - | - | - | - | - | - | - | - | - | - | - | - | - | - | - | - | - | - | - | - | - | - | G | E | S | S | Q | T | F | Y | T | - | - | - | - | - | N | G | - | - | - | T | N | L | K | K | I | - | - | - | H | V | S | P | - | - | - | - | D | F | K | T | - | - | - | - | - | - | - | - | - | - | - | - | - | - | - | - | - | - | - | - | - | - | - | - | - | - | Y | V | T | V | D | - | N | L | G | I | L | - | - | Y | I | L | K | I | - | - | - | - | - | - | - | - | - | - | - | - | - | - | - | - | - | - | - | - | - | - | - | - | - | - | - | - | - | - | - | - | - | - | L | E | - | - | - | - | - | - | - | - | - | - | - | - | - | - | - | - | - | - | - | - | - | - | - | - | - | - | - | - | - | - | - | - | - | - | - | - | - | - | - | - | - | - | - | - | - | - | - | - | - | - | - | - | M | - | - | - | - | - | - | - | - | - | - | - | - | - | - | - | - | - | - | - | - | - | - | - | - | - | - | - | - | - | - | - | - | - | - | - | - | - | - | - | - | - | - | - | - | - | - | - | - | - | - | - | - | - | - | - | - | - | - | - | - | - | - | - | - | - | - | - | - | - | - | - | - | - | - | - | - | - | - | - | - | - | - | - | - | - | - | - | - | - | - | - | - | - | - | - | - | - | - | - | - | - | - | - | - | - | - | - | - | - | - | - | - | - | - | - | - | - | - | - | - | - | - | - | - | - | - | - | - | - | - | - | - | - | - | - | - | - | - | - | - | - | - | - | - | - | - | - | - | - | - | - | - | - | - | - | - | - | - | - | - | - | - | - | - | - | - | - | - | - | - | - | - | - | - | - | - | - | - | - | - | - | - | - | - | - | - | - | - | - | - | - | - | - | - | - | - | - | - | - | - | - | - | - | - | - | - | - | - | - | - | - | - | - | - | - | - | - | - | - | - | I | E | E | - | - | - | - | - | - | - | - | - | - | - | - | - | - | - | - | - | - | - | - | - | - | - | - | - | I | Q | R | A | G | - | - | - | - | - | - | - | - | S | G | S | R | R | D | Q | - | A | R | Q | L | I | T | D | L | E | T | R | G | - | S | Q | A | L | S | L | F | I | S | C | L | E | D | T | G | Q | N | M | L | A | S | L | L | R | T | S | S | Q | A | A | - | - | - | - | R | Q | H | P | - | E | A | I | - | - | - | - | - | - | - | - | - | - | - | - | - | - | - | - | - | - | - | - | - | - | - | - | - | - | R | P | L | D | - | - | - | - | - | - | - | - | - | - | - | - | - | - | - | L | K | P | V | - | - | - | - | - | - | - | - | - | - | - | - | - | - | - | - | - | - | - | - | - | - | - | - | - | - | - | - | - | - | - | - | - | - | - | - | - | - | - | V | L | R | P | E | G | L | K | P | E | S | L | G | - | V | G | K | Q | D | P | L | M | P | - | T | - | P | - | - | - | - | - | - | - | - | - | - | - | - | - | - | - | - | - | - | - | - | - | - | - | - | - | - | - | - | - | - | - | - | - | - | - | - | - | - | - | - | - | - | - | - | - | - | - | - | - | - | - | G | K | L | V | P | V | - | - | - | - | - | - | - | - | - | - | - | - | - | - | - | - | - | - | - | - | - | - | - | - | - | - | V | L | - | - | - | - | - | - | - | - | - | - | - | - | - | - | - | - | - | - | - | - | - | - | - | - | - | - | - | - | - | - | - | - | - | - | - | - | - | - | - | - | - | - | - | - | - | - | - | - | - | - | - | - | - | - | - | - | - | - | - | - | - | - | - | - | - | - | - | - | G | P | K | E | L | W | P | - | - | - | - | - | - | - | - | - | - | - | - | - | - | - | - | - | - | - | - | - | - | - | - | - | - | A | K | L | R | P | - | - | - | - | - | - | E | V | L | R | P | - | E | V | - | - | - | - | - | - | - | - | - | - | - | - | - | - | - | - | - | - | P | R | P | V | D | I | G | S | G | G | L | - | - | - | - | - | - | - | - | - | - | - | - | - | S | D | V | C | - | - | - | - | - | - | - | - | - | - | A | - | Q | D | R | A | K | G | N | A | N | - | - | - | - | - | L | A | Y | V | - | - | L | S | A | D | - | P | C | G | R | C | L | I | V | N | N | V | H | F | R | R | E | S | - | - | G | L | R | T | R | T | G | S | S | V | D | C | E | R | L | Q | R | R | F | - | H | L | L | H | F | V | V | E | V | E | C | D | L | T | A | K | Q | M | V | Q | A | L | V | E | L | A | Q | R | D | H | - | S | A | L | D | C | C | V | V | V | I | L | S | H | G | C | Q | A | S | H | L | Q | F | P | G | A | - | V | Y | G | T | D | G | C | P | V | S | V | E | R | I | V | N | I | F | N | G | T | G | C | P | S | L | R | G | K | P | K | L | F | F | I | Q | A | C | - | - | G | G | E | Q | K | D | H | G | F | E | V | A | S | T | - | - | - | F | P | E | G | - | Q | T | P | D | S | D | - | - | - | - | - | - | - | P | E | P | D | A | T | P | F | Q | E | G | P | G | T | V | - | - | - | - | - | - | - | - | D | Q | P | D | A | M | - | - | - | - | - | - | - | S | S | L | - | - | - | P | T | P | S | D | I | L | V | S | Y | S | T | F | P | - | - | G | - | - | - | - | - | - | - | - | F | V | S | W | R | D | T | K | S | G | S | W | Y | I | E | T | L | D | S | V | F | E | Q | W | A | H | - | C | - | - | - | - | - | - | - | - | - | - | - | - | - | - | - | - | - | - | - | - | - | - | - | - | - | - | - | - | - | - | - | E | D | L | Q | T | L | L | - | - | - | - | - | L | R | - | - | - | - | - | - | - | - | - | - | - | - | V | A | N | A | V | - | - | - | - | - | S | V | K | - | - | - | - | - | - | - | G | I | Y | - | K | Q | I | - | - | P | G | - | - | - | - | - | - | - | - | - | - | - | - | - | - | - | C | F | N | F | - | - | - | - | - | - | - | - | - | L | R | K | K | L | F | F | K | T | - | - | - | - | - | - | - | - | - | - | - | - | - | - | - | - | - | - | - | - | - | - | - | - | - | - | - | - | - | - | - | - | - | - | - | - | - | - | M | - | - | - | - | - | - | - | - | - | - | - | - | - | - | - | - | - | - | - | - | - | - | - | - | - | - | - | - | - | - | - | - | - | - | - | - | - | - | - | - | - | - | - | - | - | - | - | - | - | - | - | - | G | D | V | E | K | G | K | K | I | F | V | Q | K | C | A | Q | C | H | T | V | E | K | G | G | K | H | K | T | G | P | N | L | H | G | L | F | G | R | K | T | G | Q | A | V | G | F | S | Y | T | D | A | N | K | N | K | G | I | T | W | G | E | E | T | L | M | E | Y | L | E | N | P | K | K | Y | I | P | G | T | K | M | I | F | A | G | I | K | K | K | G | E | R | A | D | L | I | A | Y | L | K | K | A | T | N | E | - | - | - | - | - | - |
| Canis\_lupus\_familiaris | M | - | - | - | - | - | - | - | - | - | - | - | - | - | - | - | - | - | - | - | - | - | - | - | - | - | - | - | - | - | - | - | - | - | - | - | - | - | - | - | - | - | - | - | - | - | - | - | - | - | - | - | - | - | - | - | - | - | - | - | - | - | - | - | - | - | - | - | - | - | - | - | - | - | - | - | - | - | - | - | - | - | - | - | - | - | - | - | - | - | - | - | - | - | - | - | - | - | D | A | K | A | - | R | N | - | - | - | - | - | - | - | - | - | - | - | - | - | - | - | - | C | L | L | - | - | - | - | - | - | - | - | - | - | - | - | - | - | - | - | - | - | - | - | - | - | - | - | - | - | - | - | - | - | - | - | - | - | - | - | - | - | - | - | - | - | - | - | - | - | - | - | - | - | - | - | - | - | - | - | - | - | - | - | - | - | - | - | - | - | - | - | - | - | - | - | - | - | - | - | - | - | - | - | - | - | - | - | - | - | - | - | - | - | - | - | - | - | - | - | - | - | - | - | - | - | - | - | - | - | - | - | - | - | - | - | - | - | - | - | - | - | - | - | - | Q | Y | R | E | A | L | E | K | D | I | K | T | S | Y | - | I | M | D | R | M | I | N | D | G | V | L | T | I | S | E | E | E | K | V | K | N | E | - | - | - | - | - | - | - | - | - | - | - | - | - | - | - | - | - | P | T | Q | Q | Q | R | A | A | L | L | I | K | T | I | L | K | K | - | - | - | - | D | N | Y | S | Y | I | S | - | - | - | - | - | F | Y | N | A | L | - | I | H | E | G | - | - | - | - | - | - | Y | K | D | L | A | A | L | L | H | S | G | I | P | V | I | S | S | S | N | G | G | K | - | - | - | - | - | - | - | - | - | - | - | - | - | - | - | - | - | - | - | - | - | - | D | S | V | G | G | I | T | - | - | - | - | - | - | - | - | - | - | - | S | H | V | K | T | V | L | C | E | G | G | - | - | - | - | - | - | - | V | P | Q | R | P | V | V | F | V | T | R | K | K | L | V | N | A | I | - | - | - | - | - | - | - | - | - | - | R | Q | N | L | F | - | - | - | - | - | - | - | - | - | - | - | - | - | - | - | - | - | - | - | - | - | - | - | - | - | - | - | - | - | - | - | - | K | L | S | D | E | P | - | - | - | - | - | - | - | - | - | - | - | - | - | - | - | - | - | - | - | - | - | - | - | - | - | - | - | - | - | - | - | - | - | - | - | - | - | - | G | W | V | V | I | Y | G | M | A | G | C | G | K | S | V | L | A | A | E | A | V | - | R | D | H | F | F | L | D | G | C | F | P | G | G | V | H | W | V | S | V | G | - | - | - | - | - | - | - | K | - | - | - | - | - | - | Q | D | K | A | - | - | - | - | - | - | - | - | - | - | - | - | G | L | L | M | K | L | Q | N | L | C | T | R | L | D | - | - | - | - | Q | - | - | - | - | - | - | - | - | - | - | - | - | - | - | - | D | E | N | F | S | Q | R | P | P | L | N | I | - | - | - | - | - | - | - | - | - | - | - | - | - | - | - | - | E | E | A | K | D | R | L | R | L | L | M | L | R | - | - | - | - | - | - | - | - | - | - | - | - | K | H | P | R | S | L | L | - | - | I | L | D | D | I | W | D | S | W | I | L | K | A | F | D | - | N | Q | C | Q | I | L | L | T | T | R | D | K | S | V | T | D | S | V | - | M | G | P | - | - | K | Y | I | V | A | V | E | S | - | - | - | D | L | G | K | E | K | G | L | E | I | L | S | L | F | V | N | M | - | K | K | A | D | L | P | E | - | - | - | - | - | - | - | - | - | - | - | - | - | - | Q | A | H | S | I | I | K | E | C | K | - | - | - | - | - | - | - | - | - | - | - | - | - | - | - | - | - | - | - | G | - | S | P | L | V | V | S | L | I | G | A | L | L | R | D | F | P | N | - | - | - | - | - | - | - | - | - | - | R | W | D | Y | Y | L | - | R | Q | L | Q | N | K | Q | F | K | R | I | - | - | - | - | - | - | - | - | - | - | - | - | - | - | - | - | - | - | - | - | - | - | R | K | S | S | F | Y | D | - | - | - | - | Y | E | A | L | D | E | A | M | S | I | S | V | E | - | M | L | R | - | - | - | E | D | I | K | D | Y | - | Y | T | D | L | S | I | L | Q | K | D | V | K | V | P | T | K | V | L | C | I | L | W | D | - | - | - | - | M | E | T | E | E | V | E | D | I | L | Q | E | F | - | - | - | - | - | - | - | - | - | - | - | V | N | K | S | L | L | F | C | D | R | N | G | K | S | - | - | - | - | - | - | - | - | - | F | R | Y | Y | L | H | D | L | Q | V | D | F | L | I | E | K | - | - | N | R | N | Q | - | L | - | Q | D | L | H | K | K | I | V | T | Q | F | Q | K | - | H | Y | - | - | - | - | - | - | Q | P | H | T | L | S | P | D | - | Q | E | - | - | - | - | - | - | - | D | C | M | Y | W | Y | N | F | L | A | Y | H | M | A | S | A | S | - | - | - | - | M | H | K | E | L | C | A | L | M | F | S | L | D | W | I | K | A | K | T | E | L | V | G | P | A | - | - | - | - | - | - | - | - | H | L | I | H | E | F | V | E | Y | R | H | I | L | D | E | K | - | - | - | - | - | - | - | D | C | A | V | C | E | N | F | Q | - | - | - | E | F | L | S | L | N | G | H | L | L | G | R | Q | P | F | - | - | - | - | - | - | - | - | - | - | - | - | - | - | - | - | - | - | - | - | - | - | - | - | - | - | - | - | - | - | - | - | - | - | - | - | - | - | - | - | - | - | - | - | - | - | - | - | - | - | - | - | - | - | - | - | - | - | - | - | - | - | - | - | - | - | - | - | - | - | - | P | N | I | V | Q | L | G | L | C | E | P | E | T | S | E | V | Y | Q | Q | A | K | L | Q | A | K | Q | E | V | D | H | G | - | - | - | - | - | - | - | - | - | - | - | - | - | - | - | - | - | - | - | - | - | - | - | - | - | - | - | - | - | M | L | Y | L | - | - | - | - | - | - | - | - | - | - | E | W | I | N | K | - | - | - | - | - | - | - | K | N | I | K | N | L | S | R | - | - | - | - | - | - | - | - | - | - | L | V | V | R | P | H | T | D | - | - | - | - | - | - | - | - | - | - | - | - | - | - | - | A | V | Y | H | A | C | - | - | - | F | S | - | E | D | G | Q | R | I | A | - | - | - | - | - | - | - | - | - | S | C | G | V | D | K | T | L | Q | V | F | K | - | - | - | - | - | A | E | T | G | E | K | L | L | E | I | - | K | A | H | E | D | E | - | - | - | V | L | C | C | A | F | S | - | T | - | - | - | - | - | - | - | - | - | - | - | - | - | D | D | R | F | I | A | T | C | S | V | D | K | K | V | K | I | - | - | - | W | N | S | M | - | - | - | - | - | - | - | - | - | - | - | - | - | - | - | - | - | - | - | - | - | - | - | - | - | - | - | - | - | - | - | - | - | - | - | - | - | - | - | - | - | - | - | - | - | - | - | - | - | - | - | - | - | - | - | - | - | - | - | - | - | - | - | - | - | - | - | - | - | - | - | - | - | - | - | - | - | - | - | - | - | - | - | - | - | - | - | - | - | - | - | - | - | - | - | - | - | - | - | - | - | - | - | - | - | - | - | - | - | - | - | - | - | - | - | - | - | - | - | - | - | - | - | - | - | - | - | - | - | - | - | - | - | - | - | - | - | - | - | - | T | G | G | L | V | H | V | Y | - | D | E | H | S | - | - | - | - | - | E | Q | V | N | - | - | - | - | - | - | - | - | - | - | - | - | - | C | C | H | - | - | - | - | - | - | - | - | - | - | - | - | - | F | T | N | N | S | - | - | - | - | - | - | - | - | - | - | - | - | - | - | - | - | H | Y | - | L | L | L | A | T | A | S | S | - | - | - | - | D | C | F | L | - | - | - | - | - | - | - | - | - | - | - | - | K | L | W | D | L | - | - | - | N | Q | K | E | C | R | N | T | M | F | - | - | - | - | - | - | - | - | - | - | - | - | - | - | - | - | - | G | H | T | N | S | - | - | - | - | - | - | - | - | - | - | - | - | - | - | - | - | V | N | H | C | R | F | S | P | D | D | K | - | - | - | - | - | - | - | - | - | - | - | - | - | - | - | - | - | - | - | - | - | - | - | - | - | - | - | - | L | L | A | S | C | S | A | D | G | T | L | K | L | - | - | - | - | - | - | - | - | - | - | - | - | - | - | - | - | - | - | - | W | D | - | V | K | S | A | N | E | R | K | S | - | - | - | - | - | - | I | N | - | - | - | - | - | - | - | - | - | - | - | - | - | - | - | - | - | - | - | - | - | - | V | K | Q | F | - | - | - | - | - | - | - | - | F | L | N | S | E | E | P | Q | E | - | - | - | - | - | - | - | - | - | - | - | - | - | - | - | - | - | - | - | - | - | - | - | - | - | - | - | D | M | E | - | V | I | V | K | C | C | - | - | - | - | - | - | - | - | - | - | - | - | - | - | - | - | S | W | S | A | D | G | T | R | I | M | V | A | A | K | N | K | I | - | - | - | - | - | - | - | - | - | - | - | F | L | F | D | I | H | T | - | - | - | G | G | - | - | - | - | - | - | - | - | - | - | - | - | - | - | - | - | - | - | - | - | - | - | - | - | - | - | - | - | - | - | - | - | - | - | - | - | - | - | - | - | - | - | - | - | - | - | - | - | - | - | - | - | - | - | - | - | - | - | - | - | - | - | - | - | - | - | - | - | - | - | - | - | - | - | - | - | - | - | - | - | - | - | - | - | - | - | - | - | - | - | - | - | - | - | - | - | - | - | - | - | - | - | - | - | - | - | - | - | - | - | - | - | - | - | - | - | - | - | - | - | - | - | - | - | - | - | L | L | A | - | - | - | - | - | - | - | - | - | - | - | - | - | - | - | - | - | - | - | - | - | - | - | - | - | - | - | - | - | - | - | - | - | - | - | - | - | - | - | - | - | - | - | - | - | - | - | - | - | - | - | - | - | - | - | - | - | - | - | - | - | - | - | - | - | - | - | - | - | - | - | - | - | - | - | - | - | - | - | - | - | - | - | - | - | - | - | - | - | - | - | - | - | - | - | - | - | - | - | - | - | - | - | - | - | - | - | - | - | - | - | - | - | - | - | - | - | - | - | - | - | - | - | - | - | - | - | - | - | - | - | - | - | - | - | - | - | - | - | - | - | - | - | - | - | - | - | - | - | - | - | - | - | - | - | - | - | - | - | - | - | - | - | - | - | - | - | - | - | - | - | - | - | - | - | - | - | - | - | - | - | - | - | E | I | - | - | - | H | T | G | H | H | S | T | I | Q | Y | C | - | - | - | - | - | - | - | - | - | - | - | - | - | - | - | - | - | - | - | - | - | - | - | - | - | D | F | S | P | Q | N | H | L | A | V | V | A | L | S | Q | - | - | - | - | - | - | C | C | V | E | - | - | - | - | - | - | - | - | - | - | - | - | - | - | - | - | - | L | W | N | M | D | S | C | - | - | - | - | - | - | - | - | - | - | - | - | - | - | - | - | - | - | - | - | - | - | - | - | - | - | - | - | - | L | K | V | A | D | C | R | G | H | L | S | - | - | - | - | - | - | - | - | - | - | - | - | - | - | - | - | - | - | - | - | - | - | - | - | - | - | - | - | - | - | - | - | - | - | - | - | - | - | - | - | - | - | - | - | W | V | H | C | V | M | F | S | - | - | - | - | - | - | - | - | - | - | - | P | D | G | S | - | - | - | - | - | - | - | - | - | - | S | F | L | T | S | S | D | D | Q | - | - | - | - | - | - | - | - | - | - | - | - | - | - | T | I | R | L | - | W | - | - | - | - | - | - | - | - | - | - | - | - | - | - | - | - | - | - | - | - | - | - | - | - | - | - | - | - | - | - | - | - | - | - | - | - | - | - | E | T | K | K | V | C | K | N | S | A | I | V | L | K | Q | E | - | - | - | - | - | - | - | - | - | - | - | - | - | - | - | - | I | D | V | V | F | - | - | - | - | - | - | - | - | - | - | - | - | - | - | - | - | - | - | - | - | - | - | - | - | - | - | - | - | - | - | - | - | - | - | - | - | - | - | - | - | - | - | - | - | - | - | - | - | - | - | - | - | - | - | Q | E | N | E | V | M | - | - | - | - | - | - | - | - | - | - | - | - | - | - | - | - | - | - | - | - | - | - | - | - | - | - | - | - | - | - | - | - | - | - | - | - | V | L | A | V | D | N | I | K | R | L | Q | L | - | I | N | G | K | T | - | - | - | - | - | - | - | - | - | - | G | Q | I | D | Y | L | I | E | A | - | - | - | - | - | - | - | - | - | - | - | - | - | - | - | - | - | - | - | - | - | - | Q | - | - | - | - | - | - | - | - | - | - | - | V | S | C | C | C | L | S | P | H | L | Q | - | - | - | - | - | - | - | - | - | Y | I | A | F | G | G | - | - | - | - | - | - | - | - | - | - | - | - | - | - | - | - | - | - | - | - | - | - | - | - | - | - | - | - | - | - | - | - | - | - | - | - | E | - | - | - | - | - | - | - | - | - | D | G | A | - | - | - | - | - | - | - | - | - | - | V | - | - | - | - | - | - | - | - | - | - | - | - | - | - | - | - | - | - | - | - | - | - | - | - | - | - | - | - | - | - | - | - | - | - | - | - | - | - | - | E | I | L | E | L | L | N | N | R | I | F | Q | S | R | I | G | - | - | - | - | - | H | - | - | - | - | - | - | - | - | - | - | - | - | K | K | T | V | R | H | I | Q | F | T | - | - | - | - | - | - | - | - | - | - | - | - | - | - | - | - | - | - | - | - | - | - | - | - | - | - | - | - | - | - | - | - | - | - | - | - | - | D | D | G | K | T | L | I | S | S | S | - | D | D | S | - | - | - | - | - | - | - | - | S | I | Q | - | V | - | - | - | - | - | - | - | - | - | - | - | - | - | - | - | - | - | - | - | - | - | - | - | - | - | - | - | - | - | - | - | - | - | - | - | - | - | - | - | W | N | - | - | - | - | - | - | - | - | - | - | W | Q | S | E | E | Y | V | F | - | - | - | - | - | - | - | - | - | - | - | - | - | - | - | - | - | - | - | - | - | - | - | - | - | - | - | - | - | - | - | - | - | - | - | - | - | - | - | - | - | - | - | - | - | - | - | - | - | - | L | Q | A | H | Q | - | - | - | - | - | - | - | - | - | - | - | - | - | - | - | - | - | - | - | - | - | - | - | - | - | - | E | T | V | K | D | F | K | L | L | - | - | - | - | - | - | - | - | - | - | - | - | - | - | - | - | - | - | - | - | - | - | - | K | N | - | S | K | L | L | - | S | W | - | - | - | S | F | D | G | T | V | - | - | - | - | - | - | - | - | - | - | - | - | - | - | - | - | - | - | - | - | - | - | - | - | - | - | - | - | - | - | - | - | - | - | - | K | I | W | S | I | I | T | G | R | I | - | - | - | - | - | E | K | D | F | V | C | - | - | - | - | - | - | - | H | Q | - | - | - | - | - | - | - | - | - | - | - | - | - | - | - | - | - | - | - | D | T | V | L | - | - | - | - | - | - | - | - | - | - | - | - | - | - | - | S | C | D | - | - | - | - | - | - | - | - | - | - | - | - | - | - | - | I | S | P | D | A | T | - | K | F | S | S | T | - | S | A | D | K | T | - | - | - | - | - | - | A | K | I | W | S | F | E | L | L | - | - | - | - | - | - | - | - | - | - | - | - | - | - | - | - | - | - | - | - | - | - | - | - | S | P | L | H | E | - | - | - | - | - | - | - | - | L | - | - | - | - | R | G | H | K | G | C | V | - | - | - | - | - | - | - | - | - | R | C | S | - | V | F | S | - | - | - | - | - | - | - | - | - | V | D | S | T | - | L | L | A | T | - | - | - | - | - | - | - | - | G | D | D | N | G | E | I | R | I | W | N | V | S | N | G | E | L | L | - | - | - | - | - | - | - | - | - | - | - | - | - | - | - | - | - | - | - | - | - | - | - | - | - | - | - | - | - | - | - | - | - | - | - | - | - | - | - | - | - | - | - | - | - | - | - | - | - | - | - | - | - | - | - | - | H | L | C | A | P | I | L | V | E | - | - | - | - | - | - | - | - | - | - | - | - | - | - | - | - | - | - | - | - | E | G | A | A | T | H | G | G | W | - | - | - | - | - | - | - | - | V | T | D | L | C | F | S | P | D | S | K | - | - | - | - | - | - | - | - | - | - | - | - | - | - | - | - | - | - | - | - | - | - | - | - | - | - | - | - | - | - | - | - | - | - | - | - | - | - | - | - | - | - | - | - | - | M | L | V | S | - | - | A | G | G | Y | L | K | A | I | T | S | P | G | T | P | F | H | P | S | V | S | - | - | - | - | - | - | - | - | - | - | - | - | - | - | - | - | - | - | - | - | - | - | - | - | - | - | - | - | - | - | - | - | - | - | - | - | - | - | - | - | - | - | - | - | - | - | - | - | - | - | - | - | - | - | - | - | - | - | - | - | - | - | - | - | - | - | - | - | - | - | - | - | - | - | - | - | - | - | - | - | - | - | - | - | - | - | - | - | - | - | - | - | - | - | - | - | - | - | - | - | - | - | - | - | - | - | - | - | - | - | - | - | - | - | - | - | - | - | - | - | - | - | - | - | - | - | - | - | - | - | - | - | - | - | - | - | - | - | - | - | - | - | - | - | - | - | - | - | - | - | - | - | - | - | - | - | - | - | - | - | - | - | - | - | - | - | - | - | - | - | - | - | - | - | - | - | - | - | - | - | - | - | - | - | - | - | - | - | - | - | - | - | - | - | - | - | - | - | - | - | - | - | - | - | - | - | - | - | - | - | - | - | - | - | - | - | - | - | - | - | - | - | - | - | - | - | - | - | - | - | - | - | - | - | - | - | - | - | - | - | - | - | - | - | - | - | - | - | - | - | - | - | - | - | - | - | - | - | - | - | - | - | - | - | - | - | - | L | H | A | A | N | T | Y | V | T | K | S | - | - | W | N | G | C | S | R | S | H | K | K | R | E | - | - | - | H | - | - | - | - | - | - | - | - | - | - | - | - | - | - | - | - | - | - | - | - | - | - | - | - | - | - | - | - | - | - | - | - | - | - | - | - | - | H | A | M | L | K | - | K | A | G | Q | I | - | - | Q | Q | E | E | E | - | - | - | - | - | - | - | - | - | - | - | - | - | - | - | - | - | - | - | - | - | - | - | - | - | - | - | - | - | - | - | - | - | - | L | P | A | R | E | L | A | - | - | - | - | - | - | - | - | - | - | - | - | - | - | - | - | - | - | - | - | - | - | - | - | - | - | - | - | - | - | - | - | - | - | - | - | - | - | - | - | - | - | - | - | - | - | - | M | - | - | - | - | - | - | - | - | - | - | - | - | - | - | - | - | - | - | - | - | - | - | - | - | - | - | - | - | - | - | - | - | - | - | - | - | - | - | - | - | - | - | - | - | - | - | - | - | - | - | - | - | - | - | D | E | A | E | R | R | - | - | - | - | - | - | - | - | - | - | - | - | - | - | - | - | - | - | - | - | A | L | R | R | - | - | - | - | - | - | - | - | - | - | - | - | - | - | - | - | - | - | - | - | - | - | - | - | - | - | - | - | - | - | C | R | - | - | - | - | - | - | - | - | V | R | L | - | V | R | E | L | Q | V | A | A | L | W | - | - | - | - | - | - | - | - | - | - | - | - | - | - | - | - | - | - | - | - | - | - | - | - | - | - | - | - | - | - | - | - | - | - | - | - | - | - | - | - | - | - | - | - | - | D | A | L | L | S | R | E | L | - | - | - | - | - | - | - | - | - | - | - | - | - | - | - | - | - | - | - | - | - | - | - | - | F | T | A | A | M | V | E | D | - | - | - | - | - | - | - | - | - | - | - | - | - | - | - | - | - | - | - | - | - | - | - | - | - | L | Q | H | A | G | - | - | - | - | - | - | - | - | S | G | S | R | S | D | Q | - | A | R | Q | L | V | I | D | L | E | T | R | G | - | S | Q | A | L | P | L | F | I | S | C | L | E | D | T | G | Q | D | A | L | A | S | C | L | R | M | S | R | Q | A | V | - | - | - | - | K | Q | N | P | - | E | A | T | - | - | - | - | - | - | - | - | - | - | - | - | - | - | - | - | - | - | - | - | - | - | - | - | - | - | R | P | L | D | - | - | - | - | - | - | - | - | - | - | - | - | - | - | - | L | M | P | V | - | - | - | - | - | - | - | - | - | - | - | - | - | - | - | - | - | - | - | - | - | - | - | - | - | - | - | - | - | - | - | - | - | - | - | - | - | - | - | V | L | G | P | T | R | L | T | P | E | D | L | R | - | - | - | G | K | D | P | S | K | L | - | T | - | P | - | - | - | - | - | - | - | - | - | - | - | - | - | - | - | - | - | - | - | - | - | - | - | - | - | - | - | - | - | - | - | - | - | - | - | - | - | - | - | - | - | - | - | - | - | - | - | - | - | - | - | G | K | L | A | P | V | - | - | - | - | - | - | - | - | - | - | - | - | - | - | - | - | - | - | - | - | - | - | - | - | - | - | V | L | - | - | - | - | - | - | - | - | - | - | - | - | - | - | - | - | - | - | - | - | - | - | - | - | - | - | - | - | - | - | - | - | - | - | - | - | - | - | - | - | - | - | - | - | - | - | - | - | - | - | - | - | - | - | - | - | - | - | - | - | - | - | - | - | - | - | - | - | G | - | - | - | - | - | P | - | - | - | - | - | - | - | - | - | - | - | - | - | - | - | - | - | - | - | - | - | - | - | - | - | - | A | E | L | Q | P | - | - | - | - | - | - | Q | V | V | R | P | - | S | V | - | - | - | - | - | - | - | - | - | - | - | - | - | - | - | - | - | - | P | R | P | T | D | N | G | S | G | R | F | - | - | - | - | - | - | - | - | - | - | - | - | - | S | D | V | C | - | - | - | - | - | - | - | - | - | - | V | - | Q | E | I | S | K | G | N | G | D | - | - | - | - | - | L | A | Y | A | - | - | L | N | A | D | - | P | C | G | Y | C | L | I | I | N | N | V | N | F | C | P | E | S | - | - | R | L | T | A | R | G | G | S | N | I | D | C | E | K | L | Q | R | R | F | - | C | L | L | R | F | T | V | E | V | K | W | D | L | T | A | K | Q | M | V | Q | A | L | V | E | L | A | R | R | D | H | - | S | A | L | D | C | C | V | V | V | V | L | S | H | G | C | E | A | S | H | R | Q | F | P | G | A | - | V | Y | G | T | D | G | C | P | V | S | I | E | K | I | V | N | I | F | N | G | T | G | C | P | S | L | G | G | K | P | K | L | F | F | I | Q | A | C | - | - | G | G | E | Q | K | D | H | G | F | A | V | A | S | A | - | - | - | S | P | E | D | - | R | S | P | G | S | D | - | - | - | - | - | - | - | S | E | P | D | A | V | P | F | Q | E | G | P | G | P | F | - | - | - | - | - | - | - | - | D | Q | L | D | A | V | - | - | - | - | - | - | - | S | S | L | - | - | - | P | T | P | S | D | I | F | V | S | Y | S | T | F | P | - | - | G | - | - | - | - | - | - | - | - | F | V | S | W | R | N | T | K | S | G | S | W | Y | V | E | T | L | D | G | V | F | E | Q | W | A | H | - | A | - | - | - | - | - | - | - | - | - | - | - | - | - | - | - | - | - | - | - | - | - | - | - | - | - | - | - | - | - | - | - | E | D | L | Q | T | L | L | - | - | - | - | - | L | R | - | - | - | - | - | - | - | - | - | - | - | - | V | A | N | A | V | - | - | - | - | - | S | Q | K | - | - | - | - | - | - | - | G | V | Y | - | K | Q | I | - | - | P | G | - | - | - | - | - | - | - | - | - | - | - | - | - | - | - | C | F | N | F | - | - | - | - | - | - | - | - | - | L | R | K | K | L | Y | F | K | M | - | - | - | - | - | - | - | - | - | - | - | - | - | - | - | - | - | - | - | - | - | - | - | - | - | - | - | - | - | - | - | - | - | - | - | - | - | - | M | - | - | - | - | - | - | - | - | - | - | - | - | - | - | - | - | - | - | - | - | - | - | - | - | - | - | - | - | - | - | - | - | - | - | - | - | - | - | - | - | - | - | - | - | - | - | - | - | - | - | - | - | G | D | V | E | K | G | K | K | I | F | V | Q | K | C | A | Q | C | H | T | V | E | K | G | G | K | H | K | T | G | P | N | L | H | G | L | F | G | R | K | T | G | Q | A | P | G | F | S | Y | T | D | A | N | K | N | K | G | I | T | W | G | E | E | T | L | M | E | Y | L | E | N | P | K | K | Y | I | P | G | T | K | M | I | F | A | G | I | K | K | T | G | E | R | A | D | L | I | A | Y | L | K | K | A | T | K | E | - | - | - | - | - | - |
| Capra\_hircus | M | - | - | - | - | - | - | - | - | - | - | - | - | - | - | - | - | - | - | - | - | - | - | - | - | - | - | - | - | - | - | - | - | - | - | - | - | - | - | - | - | - | - | - | - | - | - | - | - | - | - | - | - | - | - | - | - | - | - | - | - | - | - | - | - | - | - | - | - | - | - | - | - | - | - | - | - | - | - | - | - | - | - | - | - | - | - | - | - | - | - | - | - | - | - | - | - | - | D | A | K | A | - | R | N | - | - | - | - | - | - | - | - | - | - | - | - | - | - | - | - | C | L | L | - | - | - | - | - | - | - | - | - | - | - | - | - | - | - | - | - | - | - | - | - | - | - | - | - | - | - | - | - | - | - | - | - | - | - | - | - | - | - | - | - | - | - | - | - | - | - | - | - | - | - | - | - | - | - | - | - | - | - | - | - | - | - | - | - | - | - | - | - | - | - | - | - | - | - | - | - | - | - | - | - | - | - | - | - | - | - | - | - | - | - | - | - | - | - | - | - | - | - | - | - | - | - | - | - | - | - | - | - | - | - | - | - | - | - | - | - | - | - | - | - | Q | H | R | E | A | L | E | R | D | I | K | T | S | Y | - | I | M | D | H | M | I | S | N | G | V | L | T | V | S | E | E | E | K | V | K | N | E | - | - | - | - | - | - | - | - | - | - | - | - | - | - | - | - | - | P | T | Q | R | Q | R | A | A | M | L | I | K | M | I | L | K | K | - | - | - | - | D | N | H | S | Y | I | S | - | - | - | - | - | F | Y | N | A | L | - | L | H | E | G | - | - | - | - | - | - | Y | R | D | L | A | A | L | L | H | G | G | I | P | V | V | S | S | S | D | G | - | K | - | - | - | - | - | - | - | - | - | - | - | - | - | - | - | - | - | - | - | - | - | - | D | S | V | S | G | I | T | - | - | - | - | - | - | - | - | - | - | - | S | Y | V | R | T | V | L | C | E | G | G | - | - | - | - | - | - | - | V | P | Q | R | P | V | V | F | V | T | R | K | K | L | V | N | A | I | - | - | - | - | - | - | - | - | - | - | Q | Q | K | L | S | - | - | - | - | - | - | - | - | - | - | - | - | - | - | - | - | - | - | - | - | - | - | - | - | - | - | - | - | - | - | - | - | K | L | N | G | E | P | - | - | - | - | - | - | - | - | - | - | - | - | - | - | - | - | - | - | - | - | - | - | - | - | - | - | - | - | - | - | - | - | - | - | - | - | - | - | G | W | V | T | I | Y | G | M | A | G | C | G | K | S | V | L | A | A | E | A | V | - | R | D | H | S | F | L | E | D | C | F | P | G | G | V | H | W | V | S | V | G | - | - | - | - | - | - | - | K | - | - | - | - | - | - | Q | D | K | S | - | - | - | - | - | - | - | - | - | - | - | - | G | L | L | M | K | L | Q | N | L | C | A | R | L | D | - | - | - | - | Q | - | - | - | - | - | - | - | - | - | - | - | - | - | - | - | D | E | S | F | S | Q | R | L | P | L | N | I | - | - | - | - | - | - | - | - | - | - | - | - | - | - | - | - | E | E | A | K | D | R | L | R | I | L | M | L | R | - | - | - | - | - | - | - | - | - | - | - | - | K | H | P | R | S | L | L | - | - | I | L | D | D | I | W | D | P | W | V | L | K | A | F | D | - | N | Q | C | R | I | L | L | T | T | R | D | K | S | V | T | D | S | V | - | M | G | P | - | - | K | Y | V | V | P | V | E | S | - | - | - | G | L | G | K | E | K | A | L | E | I | L | S | L | F | V | N | M | - | K | K | A | D | L | P | E | - | - | - | - | - | - | - | - | - | - | - | - | - | - | Q | A | H | S | I | I | Q | E | C | K | - | - | - | - | - | - | - | - | - | - | - | - | - | - | - | - | - | - | - | G | - | S | P | L | V | V | S | L | I | G | A | L | L | R | D | F | P | N | - | - | - | - | - | - | - | - | - | - | R | W | E | Y | Y | L | - | R | Q | L | Q | N | K | Q | F | K | R | I | - | - | - | - | - | - | - | - | - | - | - | - | - | - | - | - | - | - | - | - | - | - | R | K | S | S | S | Y | D | - | - | - | - | Y | E | A | L | D | E | A | M | S | I | S | V | E | - | M | L | R | - | - | - | E | D | I | K | D | Y | - | Y | T | D | L | S | I | V | Q | K | D | V | K | V | P | T | K | V | L | C | I | L | W | D | - | - | - | - | L | E | T | E | E | V | E | D | I | L | Q | E | F | - | - | - | - | - | - | - | - | - | - | - | V | N | K | S | L | L | F | C | D | R | N | G | K | S | - | - | - | - | - | - | - | - | - | F | L | Y | Y | L | H | D | L | Q | V | D | F | L | T | E | K | - | - | N | R | S | Q | - | L | - | Q | D | L | H | K | K | M | V | T | Q | F | Q | R | - | H | H | - | - | - | - | - | - | Q | L | Y | S | L | T | P | D | - | Q | E | - | - | - | - | - | - | - | D | C | M | Y | W | Y | N | F | L | A | Y | H | M | A | S | A | N | - | - | - | - | M | H | K | E | L | C | A | L | M | F | S | L | D | W | I | K | A | K | T | E | L | V | G | P | A | - | - | - | - | - | - | - | - | H | L | I | H | E | F | V | E | Y | R | H | I | L | D | E | K | - | - | - | - | - | - | - | D | G | T | D | C | E | N | F | Q | - | - | - | E | F | L | S | L | N | G | H | L | L | G | R | Q | P | F | - | - | - | - | - | - | - | - | - | - | - | - | - | - | - | - | - | - | - | - | - | - | - | - | - | - | - | - | - | - | - | - | - | - | - | - | - | - | - | - | - | - | - | - | - | - | - | - | - | - | - | - | - | - | - | - | - | - | - | - | - | - | - | - | - | - | - | - | - | - | - | P | N | I | V | Q | L | G | L | C | E | L | E | T | S | E | V | Y | R | Q | A | K | L | Q | A | K | Q | E | V | D | N | G | - | - | - | - | - | - | - | - | - | - | - | - | - | - | - | - | - | - | - | - | - | - | - | - | - | - | - | - | - | M | L | Y | L | - | - | - | - | - | - | - | - | - | - | E | W | V | N | K | - | - | - | - | - | - | - | K | N | M | K | N | L | S | R | - | - | - | - | - | - | - | - | - | - | L | V | V | R | P | H | T | D | - | - | - | - | - | - | - | - | - | - | - | - | - | - | - | A | V | Y | H | A | C | - | - | - | F | S | - | E | D | G | Q | R | I | A | - | - | - | - | - | - | - | - | - | S | C | G | A | D | K | T | L | Q | V | F | K | - | - | - | - | - | A | E | T | G | E | K | L | L | E | I | - | K | A | H | E | D | E | - | - | - | V | L | C | C | A | F | S | - | A | - | - | - | - | - | - | - | - | - | - | - | - | - | D | D | R | F | I | A | T | C | S | V | D | K | K | V | K | I | - | - | - | W | N | S | M | - | - | - | - | - | - | - | - | - | - | - | - | - | - | - | - | - | - | - | - | - | - | - | - | - | - | - | - | - | - | - | - | - | - | - | - | - | - | - | - | - | - | - | - | - | - | - | - | - | - | - | - | - | - | - | - | - | - | - | - | - | - | - | - | - | - | - | - | - | - | - | - | - | - | - | - | - | - | - | - | - | - | - | - | - | - | - | - | - | - | - | - | - | - | - | - | - | - | - | - | - | - | - | - | - | - | - | - | - | - | - | - | - | - | - | - | - | - | - | - | - | - | - | - | - | - | - | - | - | - | - | - | - | - | - | - | - | - | - | - | T | G | E | L | V | R | I | Y | - | D | E | H | S | - | - | - | - | - | E | Q | V | N | - | - | - | - | - | - | - | - | - | - | - | - | - | C | C | H | - | - | - | - | - | - | - | - | - | - | - | - | - | F | T | N | N | S | - | - | - | - | - | - | - | - | - | - | - | - | - | - | - | - | S | H | - | L | L | L | A | T | G | S | N | - | - | - | - | D | Y | F | L | - | - | - | - | - | - | - | - | - | - | - | - | K | L | W | D | L | - | - | - | N | Q | E | E | C | R | N | T | M | F | - | - | - | - | - | - | - | - | - | - | - | - | - | - | - | - | - | G | H | T | N | S | - | - | - | - | - | - | - | - | - | - | - | - | - | - | - | - | V | S | H | C | R | F | S | P | D | D | Q | - | - | - | - | - | - | - | - | - | - | - | - | - | - | - | - | - | - | - | - | - | - | - | - | - | - | - | - | V | L | A | S | C | S | A | D | G | T | L | K | L | - | - | - | - | - | - | - | - | - | - | - | - | - | - | - | - | - | - | - | W | D | - | V | K | S | A | N | E | R | K | S | - | - | - | - | - | - | I | N | - | - | - | - | - | - | - | - | - | - | - | - | - | - | - | - | - | - | - | - | - | - | V | K | Q | F | - | - | - | - | - | - | - | - | F | L | N | S | E | E | P | Q | E | - | - | - | - | - | - | - | - | - | - | - | - | - | - | - | - | - | - | - | - | - | - | - | - | - | - | - | D | L | E | - | V | I | V | K | C | C | - | - | - | - | - | - | - | - | - | - | - | - | - | - | - | - | S | W | S | A | D | G | A | K | I | M | V | A | A | K | N | K | V | - | - | - | - | - | - | - | - | - | - | - | F | L | F | D | I | H | T | - | - | - | S | G | - | - | - | - | - | - | - | - | - | - | - | - | - | - | - | - | - | - | - | - | - | - | - | - | - | - | - | - | - | - | - | - | - | - | - | - | - | - | - | - | - | - | - | - | - | - | - | - | - | - | - | - | - | - | - | - | - | - | - | - | - | - | - | - | - | - | - | - | - | - | - | - | - | - | - | - | - | - | - | - | - | - | - | - | - | - | - | - | - | - | - | - | - | - | - | - | - | - | - | - | - | - | - | - | - | - | - | - | - | - | - | - | - | - | - | - | - | - | - | - | - | - | - | - | - | - | L | L | A | - | - | - | - | - | - | - | - | - | - | - | - | - | - | - | - | - | - | - | - | - | - | - | - | - | - | - | - | - | - | - | - | - | - | - | - | - | - | - | - | - | - | - | - | - | - | - | - | - | - | - | - | - | - | - | - | - | - | - | - | - | - | - | - | - | - | - | - | - | - | - | - | - | - | - | - | - | - | - | - | - | - | - | - | - | - | - | - | - | - | - | - | - | - | - | - | - | - | - | - | - | - | - | - | - | - | - | - | - | - | - | - | - | - | - | - | - | - | - | - | - | - | - | - | - | - | - | - | - | - | - | - | - | - | - | - | - | - | - | - | - | - | - | - | - | - | - | - | - | - | - | - | - | - | - | - | - | - | - | - | - | - | - | - | - | - | - | - | - | - | - | - | - | - | - | - | - | - | - | - | - | - | - | E | I | - | - | - | H | M | G | H | H | S | T | V | Q | Y | C | - | - | - | - | - | - | - | - | - | - | - | - | - | - | - | - | - | - | - | - | - | - | - | - | - | D | F | S | P | H | N | H | L | A | V | V | A | L | S | Q | - | - | - | - | - | - | Y | C | V | E | - | - | - | - | - | - | - | - | - | - | - | - | - | - | - | - | - | L | W | S | M | D | S | G | - | - | - | - | - | - | - | - | - | - | - | - | - | - | - | - | - | - | - | - | - | - | - | - | - | - | - | - | - | L | K | V | A | D | C | R | G | H | L | S | - | - | - | - | - | - | - | - | - | - | - | - | - | - | - | - | - | - | - | - | - | - | - | - | - | - | - | - | - | - | - | - | - | - | - | - | - | - | - | - | - | - | - | - | W | V | H | C | V | M | F | S | - | - | - | - | - | - | - | - | - | - | - | P | D | G | S | - | - | - | - | - | - | - | - | - | - | S | F | L | T | S | S | D | D | Q | - | - | - | - | - | - | - | - | - | - | - | - | - | - | T | V | R | L | - | W | - | - | - | - | - | - | - | - | - | - | - | - | - | - | - | - | - | - | - | - | - | - | - | - | - | - | - | - | - | - | - | - | - | - | - | - | - | - | E | T | K | K | V | C | K | N | S | A | T | V | L | K | Q | E | - | - | - | - | - | - | - | - | - | - | - | - | - | - | - | - | I | D | V | V | F | - | - | - | - | - | - | - | - | - | - | - | - | - | - | - | - | - | - | - | - | - | - | - | - | - | - | - | - | - | - | - | - | - | - | - | - | - | - | - | - | - | - | - | - | - | - | - | - | - | - | - | - | - | - | Q | E | K | E | V | T | - | - | - | - | - | - | - | - | - | - | - | - | - | - | - | - | - | - | - | - | - | - | - | - | - | - | - | - | - | - | - | - | - | - | - | - | V | L | A | V | D | N | I | R | R | L | Q | L | - | I | N | G | N | T | - | - | - | - | - | - | - | - | - | - | G | Q | I | D | Y | L | T | E | A | - | - | - | - | - | - | - | - | - | - | - | - | - | - | - | - | - | - | - | - | - | - | Q | - | - | - | - | - | - | - | - | - | - | - | V | S | C | C | C | L | S | P | H | L | Q | - | - | - | - | - | - | - | - | - | Y | L | A | F | G | A | - | - | - | - | - | - | - | - | - | - | - | - | - | - | - | - | - | - | - | - | - | - | - | - | - | - | - | - | - | - | - | - | - | - | - | - | E | - | - | - | - | - | - | - | - | - | D | G | A | - | - | - | - | - | - | - | - | - | - | I | - | - | - | - | - | - | - | - | - | - | - | - | - | - | - | - | - | - | - | - | - | - | - | - | - | - | - | - | - | - | - | - | - | - | - | - | - | - | - | Q | I | L | E | L | L | N | N | R | I | F | Q | S | R | N | A | - | - | - | - | - | H | - | - | - | - | - | - | - | - | - | - | - | - | K | T | T | V | R | H | I | Q | F | T | - | - | - | - | - | - | - | - | - | - | - | - | - | - | - | - | - | - | - | - | - | - | - | - | - | - | - | - | - | - | - | - | - | - | - | - | - | A | D | G | K | T | L | I | S | S | S | - | D | D | S | - | - | - | - | - | - | - | - | A | I | Q | - | I | - | - | - | - | - | - | - | - | - | - | - | - | - | - | - | - | - | - | - | - | - | - | - | - | - | - | - | - | - | - | - | - | - | - | - | - | - | - | - | W | N | - | - | - | - | - | - | - | - | - | - | W | Q | S | E | D | P | F | F | - | - | - | - | - | - | - | - | - | - | - | - | - | - | - | - | - | - | - | - | - | - | - | - | - | - | - | - | - | - | - | - | - | - | - | - | - | - | - | - | - | - | - | - | - | - | - | - | - | - | L | Q | A | H | Q | - | - | - | - | - | - | - | - | - | - | - | - | - | - | - | - | - | - | - | - | - | - | - | - | - | - | E | T | V | K | D | F | K | L | L | - | - | - | - | - | - | - | - | - | - | - | - | - | - | - | - | - | - | - | - | - | - | - | K | N | - | S | R | L | L | - | S | W | - | - | - | S | F | D | G | T | V | - | - | - | - | - | - | - | - | - | - | - | - | - | - | - | - | - | - | - | - | - | - | - | - | - | - | - | - | - | - | - | - | - | - | - | K | V | W | N | I | T | T | R | R | I | - | - | - | - | - | E | K | D | F | V | C | - | - | - | - | - | - | - | H | Q | - | - | - | - | - | - | - | - | - | - | - | - | - | - | - | - | - | - | - | D | T | V | L | - | - | - | - | - | - | - | - | - | - | - | - | - | - | - | S | C | D | - | - | - | - | - | - | - | - | - | - | - | - | - | - | - | I | S | S | D | G | A | - | K | F | S | S | T | - | S | A | D | K | T | - | - | - | - | - | - | A | K | I | W | S | F | E | L | L | - | - | - | - | - | - | - | - | - | - | - | - | - | - | - | - | - | - | - | - | - | - | - | - | S | P | L | H | E | - | - | - | - | - | - | - | - | L | - | - | - | - | R | G | H | K | G | C | V | - | - | - | - | - | - | - | - | - | R | C | S | - | S | F | S | - | - | - | - | - | - | - | - | - | V | D | N | T | - | M | L | A | T | - | - | - | - | - | - | - | - | G | D | D | N | G | E | I | R | M | W | N | V | S | N | G | E | L | L | - | - | - | - | - | - | - | - | - | - | - | - | - | - | - | - | - | - | - | - | - | - | - | - | - | - | - | - | - | - | - | - | - | - | - | - | - | - | - | - | - | - | - | - | - | - | - | - | - | - | - | - | - | - | - | - | H | L | C | A | P | I | S | V | E | - | - | - | - | - | - | - | - | - | - | - | - | - | - | - | - | - | - | - | - | E | G | A | A | T | H | G | G | W | - | - | - | - | - | - | - | - | V | T | D | L | C | F | S | P | D | N | K | - | - | - | - | - | - | - | - | - | - | - | - | - | - | - | - | - | - | - | - | - | - | - | - | - | - | - | - | - | - | - | - | - | - | - | - | - | - | - | - | - | - | - | - | - | M | L | V | S | - | - | A | G | G | Y | L | K | - | - | - | - | - | - | - | - | W | W | N | V | V | T | - | - | - | - | - | - | - | - | - | - | - | - | - | - | - | - | - | - | - | - | - | - | - | - | - | - | - | - | - | - | - | - | - | - | - | - | - | - | - | - | - | - | - | - | - | - | - | - | - | - | - | - | - | - | - | - | - | - | - | - | - | - | - | - | - | - | - | - | - | - | - | - | - | - | - | - | - | - | - | - | - | - | - | - | - | - | - | - | - | - | - | - | - | - | - | - | - | - | - | - | - | - | - | - | - | - | - | - | - | - | - | - | - | - | - | - | - | - | - | - | - | - | - | - | - | - | - | - | - | - | - | - | - | - | - | - | - | - | - | - | - | - | - | - | - | - | - | - | - | - | - | - | - | - | - | - | - | - | - | - | - | - | - | - | - | - | - | - | - | - | - | - | - | - | - | - | - | - | - | - | - | - | - | - | - | - | - | - | - | - | - | - | - | - | - | - | - | - | - | - | - | - | - | - | - | - | - | - | - | - | - | - | - | - | - | - | - | - | - | - | - | - | - | - | - | - | - | - | - | - | - | - | - | - | - | - | - | - | - | - | - | - | - | - | - | - | - | - | - | - | - | - | - | - | - | - | - | - | - | - | - | - | - | - | - | - | - | G | E | S | S | Q | T | F | Y | T | - | - | - | - | - | N | G | - | - | - | T | N | L | K | K | I | - | - | - | H | V | S | P | - | - | - | - | D | F | K | T | - | - | - | - | - | - | - | - | - | - | - | - | - | - | - | - | - | - | - | - | - | - | - | - | - | - | Y | V | T | V | D | - | N | L | G | I | L | - | - | Y | I | L | K | I | - | - | - | - | - | - | - | - | - | - | - | - | - | - | - | - | - | - | - | - | - | - | - | - | - | - | - | - | - | - | - | - | - | - | L | E | - | - | - | - | - | - | - | - | - | - | - | - | - | - | - | - | - | - | - | - | - | - | - | - | - | - | - | - | - | - | - | - | - | - | - | - | - | - | - | - | - | - | - | - | - | - | - | - | - | - | - | - | M | - | - | - | - | - | - | - | - | - | - | - | - | - | - | - | - | - | - | - | - | - | - | - | - | - | - | - | - | - | - | - | - | - | - | - | - | - | - | - | - | - | - | - | - | - | - | - | - | - | - | - | - | - | - | D | E | A | D | R | L | - | - | - | - | - | - | - | - | - | - | - | - | - | - | - | - | - | - | - | - | L | L | R | R | - | - | - | - | - | - | - | - | - | - | - | - | - | - | - | - | - | - | - | - | - | - | - | - | - | - | - | - | - | - | Y | R | - | - | - | - | - | - | - | - | L | P | L | - | V | G | E | L | Q | V | A | S | L | W | - | - | - | - | - | - | - | - | - | - | - | - | - | - | - | - | - | - | - | - | - | - | - | - | - | - | - | - | - | - | - | - | - | - | - | - | - | - | - | - | - | - | - | - | - | D | A | L | L | S | R | E | L | - | - | - | - | - | - | - | - | - | - | - | - | - | - | - | - | - | - | - | - | - | - | - | - | F | T | P | D | M | I | E | D | - | - | - | - | - | - | - | - | - | - | - | - | - | - | - | - | - | - | - | - | - | - | - | - | - | I | Q | R | A | G | - | - | - | - | - | - | - | - | S | G | S | R | R | D | Q | - | A | R | Q | L | I | L | D | L | E | T | R | G | - | S | Q | A | L | P | L | F | I | S | C | L | E | D | T | G | Q | D | T | L | A | S | L | L | K | T | S | R | Q | A | A | - | - | - | - | K | Q | D | V | - | E | A | I | - | - | - | - | - | - | - | - | - | - | - | - | - | - | - | - | - | - | - | - | - | - | - | - | - | - | R | P | L | D | - | - | - | - | - | - | - | - | - | - | - | - | - | - | - | L | K | P | V | - | - | - | - | - | - | - | - | - | - | - | - | - | - | - | - | - | - | - | - | - | - | - | - | - | - | - | - | - | - | - | - | - | - | - | - | - | - | - | V | L | G | P | E | G | L | K | P | E | E | L | R | - | V | M | K | Q | D | P | S | K | P | - | S | - | Q | - | - | - | - | - | - | - | - | - | - | - | - | - | - | - | - | - | - | - | - | - | - | - | - | - | - | - | - | - | - | - | - | - | - | - | - | - | - | - | - | - | - | - | - | - | - | - | - | - | - | - | G | K | L | A | P | V | - | - | - | - | - | - | - | - | - | - | - | - | - | - | - | - | - | - | - | - | - | - | - | - | - | - | V | L | - | - | - | - | - | - | - | - | - | - | - | - | - | - | - | - | - | - | - | - | - | - | - | - | - | - | - | - | - | - | - | - | - | - | - | - | - | - | - | - | - | - | - | - | - | - | - | - | - | - | - | - | - | - | - | - | - | - | - | - | - | - | - | - | - | - | - | - | G | P | E | E | L | W | P | - | - | - | - | - | - | - | - | - | - | - | - | - | - | - | - | - | - | - | - | - | - | - | - | - | - | A | K | L | R | P | - | - | - | - | - | - | E | V | L | R | P | - | E | V | - | - | - | - | - | - | - | - | - | - | - | - | - | - | - | - | - | - | P | R | P | G | N | I | G | S | G | G | F | - | - | - | - | - | - | - | - | - | - | - | - | - | T | D | V | C | - | - | - | - | - | - | - | - | - | - | P | - | Q | D | R | A | K | G | N | A | D | - | - | - | - | - | L | A | Y | V | - | - | L | N | A | D | - | P | C | G | H | C | L | I | I | N | N | V | N | F | C | R | E | S | - | - | G | L | M | T | R | T | G | S | N | I | D | C | E | R | M | Q | R | R | F | - | H | L | L | Q | F | A | V | E | V | K | C | D | L | T | A | K | Q | M | V | Q | A | L | M | Q | L | A | R | R | D | H | - | G | A | L | D | C | C | V | V | V | I | L | S | H | G | C | Q | A | S | H | R | Q | F | P | G | A | - | V | Y | G | T | D | G | C | P | V | S | V | E | R | I | V | N | T | F | N | G | T | G | C | P | S | L | G | G | K | P | K | L | F | F | I | Q | A | C | - | - | G | G | E | Q | K | D | H | G | F | E | V | A | S | T | - | - | - | S | P | E | D | - | E | T | P | N | S | N | - | - | - | - | - | - | - | P | E | T | D | A | T | P | F | Q | E | G | P | R | S | T | - | - | - | - | - | - | - | - | D | E | P | D | A | V | - | - | - | - | - | - | - | S | S | L | - | - | - | P | T | P | S | D | I | L | V | S | Y | S | T | F | P | - | - | G | - | - | - | - | - | - | - | - | F | V | S | W | R | D | P | K | S | G | S | W | Y | I | E | T | L | D | S | I | F | E | Q | W | A | H | - | C | - | - | - | - | - | - | - | - | - | - | - | - | - | - | - | - | - | - | - | - | - | - | - | - | - | - | - | - | - | - | - | E | D | L | Q | T | L | L | - | - | - | - | - | L | R | - | - | - | - | - | - | - | - | - | - | - | - | V | A | N | A | V | - | - | - | - | - | S | V | K | - | - | - | - | - | - | - | G | I | Y | - | K | Q | I | - | - | P | G | - | - | - | - | - | - | - | - | - | - | - | - | - | - | - | C | F | N | F | - | - | - | - | - | - | - | - | - | L | R | K | K | L | F | F | K | T | - | - | - | - | - | - | - | - | - | - | - | - | - | - | - | - | - | - | - | - | - | - | - | - | - | - | - | - | - | - | - | - | - | - | - | - | - | - | M | - | - | - | - | - | - | - | - | - | - | - | - | - | - | - | - | - | - | - | - | - | - | - | - | - | - | - | - | - | - | - | - | - | - | - | - | - | - | - | - | - | - | - | - | - | - | - | - | - | - | - | - | G | D | V | E | K | G | K | K | I | F | V | Q | K | C | A | Q | C | H | T | V | E | K | G | G | K | H | K | T | G | P | N | L | H | G | L | F | G | R | K | T | G | Q | A | P | G | F | S | Y | T | D | A | N | K | N | K | G | I | T | W | G | E | E | T | L | M | E | Y | L | E | N | P | K | K | Y | I | P | G | T | K | M | I | F | A | G | I | K | K | K | G | E | R | E | D | L | I | A | Y | L | K | K | A | T | N | E | - | - | - | - | - | - |
| Castor\_canadensis | M | - | - | - | - | - | - | - | - | - | - | - | - | - | - | - | - | - | - | - | - | - | - | - | - | - | - | - | - | - | - | - | - | - | - | - | - | - | - | - | - | - | - | - | - | - | - | - | - | - | - | - | - | - | - | - | - | - | - | - | - | - | - | - | - | - | - | - | - | - | - | - | - | - | - | - | - | - | - | - | - | - | - | - | - | - | - | - | - | - | - | - | - | - | - | - | - | - | D | A | K | A | - | R | N | - | - | - | - | - | - | - | - | - | - | - | - | - | - | - | - | C | L | L | - | - | - | - | - | - | - | - | - | - | - | - | - | - | - | - | - | - | - | - | - | - | - | - | - | - | - | - | - | - | - | - | - | - | - | - | - | - | - | - | - | - | - | - | - | - | - | - | - | - | - | - | - | - | - | - | - | - | - | - | - | - | - | - | - | - | - | - | - | - | - | - | - | - | - | - | - | - | - | - | - | - | - | - | - | - | - | - | - | - | - | - | - | - | - | - | - | - | - | - | - | - | - | - | - | - | - | - | - | - | - | - | - | - | - | - | - | - | - | - | - | Q | Y | R | E | A | L | E | K | D | I | K | T | S | Y | - | I | M | D | H | M | I | S | D | G | V | L | T | I | S | E | E | E | K | V | K | N | E | - | - | - | - | - | - | - | - | - | - | - | - | - | - | - | - | - | P | T | K | H | Q | R | A | A | I | L | I | K | M | I | L | K | K | - | - | - | - | D | N | R | A | Y | I | S | - | - | - | - | - | F | Y | N | A | L | - | L | H | E | G | - | - | - | - | - | - | Y | K | D | L | A | A | L | L | H | S | G | I | P | V | I | S | S | S | S | G | - | K | - | - | - | - | - | - | - | - | - | - | - | - | - | - | - | - | - | - | - | - | - | - | D | S | G | G | G | I | T | - | - | - | - | - | - | - | - | - | - | - | S | Y | V | R | T | V | L | C | E | G | G | - | - | - | - | - | - | - | V | P | Q | R | P | V | V | F | V | T | R | K | K | L | V | N | A | I | - | - | - | - | - | - | - | - | - | - | Q | Q | K | L | F | - | - | - | - | - | - | - | - | - | - | - | - | - | - | - | - | - | - | - | - | - | - | - | - | - | - | - | - | - | - | - | - | K | L | N | G | E | P | - | - | - | - | - | - | - | - | - | - | - | - | - | - | - | - | - | - | - | - | - | - | - | - | - | - | - | - | - | - | - | - | - | - | - | - | - | - | G | W | V | T | I | Y | G | M | A | G | C | G | K | S | V | L | A | A | E | A | V | - | R | D | H | S | L | L | E | G | C | F | P | G | G | V | H | W | V | S | V | G | - | - | - | - | - | - | - | K | - | - | - | - | - | - | Q | D | K | S | - | - | - | - | - | - | - | - | - | - | - | - | G | L | L | M | K | L | Q | N | L | C | A | R | L | D | - | - | - | - | Q | - | - | - | - | - | - | - | - | - | - | - | - | - | - | - | D | E | S | F | S | Q | R | L | P | L | N | I | - | - | - | - | - | - | - | - | - | - | - | - | - | - | - | - | E | E | A | K | D | R | L | R | I | L | M | L | R | - | - | - | - | - | - | - | - | - | - | - | - | K | H | P | R | S | L | L | - | - | I | L | D | D | V | W | D | P | W | V | L | K | A | F | D | - | N | Q | C | Q | I | L | L | T | T | R | D | K | S | V | T | D | S | V | - | M | G | P | - | - | K | Y | E | V | P | V | E | S | - | - | - | G | L | G | K | E | K | G | L | E | I | L | S | L | F | V | N | M | - | K | K | A | D | L | P | E | - | - | - | - | - | - | - | - | - | - | - | - | - | - | Q | A | H | S | I | I | K | E | C | K | - | - | - | - | - | - | - | - | - | - | - | - | - | - | - | - | - | - | - | G | - | S | P | L | V | V | S | L | I | G | A | L | L | R | D | F | P | N | - | - | - | - | - | - | - | - | - | - | R | W | E | Y | Y | L | - | R | Q | L | Q | N | K | Q | F | K | R | I | - | - | - | - | - | - | - | - | - | - | - | - | - | - | - | - | - | - | - | - | - | - | R | K | S | S | S | Y | D | - | - | - | - | Y | E | A | L | D | E | A | M | S | I | S | V | E | - | M | L | R | - | - | - | E | N | I | K | D | Y | - | Y | T | D | L | S | I | L | Q | K | D | V | K | V | P | T | K | V | L | C | V | L | W | D | - | - | - | - | M | E | T | E | D | V | E | D | I | L | Q | E | F | - | - | - | - | - | - | - | - | - | - | - | V | N | K | S | L | L | F | C | D | R | N | G | K | S | - | - | - | - | - | - | - | - | - | F | C | Y | Y | L | H | D | L | Q | V | D | F | L | T | E | K | - | - | N | R | S | Q | - | L | - | Q | D | L | H | K | K | M | V | T | Q | F | Q | R | - | Y | Y | - | - | - | - | - | - | Q | P | H | A | L | S | P | D | - | Q | E | - | - | - | - | - | - | - | D | G | M | Y | W | Y | S | F | L | A | Y | H | M | A | S | A | N | - | - | - | - | M | H | K | E | L | C | A | L | M | F | S | L | D | W | I | K | A | K | T | E | L | V | G | P | A | - | - | - | - | - | - | - | - | H | L | I | H | E | F | V | E | Y | S | H | I | L | D | E | K | - | - | - | - | - | - | - | D | C | A | V | C | E | N | F | Q | - | - | - | E | F | L | S | L | N | G | H | L | L | G | R | Q | P | F | - | - | - | - | - | - | - | - | - | - | - | - | - | - | - | - | - | - | - | - | - | - | - | - | - | - | - | - | - | - | - | - | - | - | - | - | - | - | - | - | - | - | - | - | - | - | - | - | - | - | - | - | - | - | - | - | - | - | - | - | - | - | - | - | - | - | - | - | - | - | - | P | N | I | V | Q | L | G | L | C | E | P | E | T | S | E | V | Y | Q | Q | A | K | L | Q | A | K | Q | E | V | D | N | G | - | - | - | - | - | - | - | - | - | - | - | - | - | - | - | - | - | - | - | - | - | - | - | - | - | - | - | - | - | K | L | Y | L | - | - | - | - | - | - | - | - | - | - | E | W | T | N | K | - | - | - | - | - | - | - | K | T | I | K | N | L | S | R | - | - | - | - | - | - | - | - | - | - | L | V | V | R | P | H | T | D | - | - | - | - | - | - | - | - | - | - | - | - | - | - | - | A | V | Y | H | A | C | - | - | - | F | S | - | E | N | G | Q | R | I | A | - | - | - | - | - | - | - | - | - | S | C | G | A | D | K | T | L | Q | V | F | K | - | - | - | - | - | A | E | T | G | E | K | L | L | E | I | - | K | A | H | E | D | E | - | - | - | V | L | C | C | A | F | S | - | A | - | - | - | - | - | - | - | - | - | - | - | - | - | D | D | R | F | I | A | T | C | S | V | D | K | K | V | K | I | - | - | - | W | N | S | V | - | - | - | - | - | - | - | - | - | - | - | - | - | - | - | - | - | - | - | - | - | - | - | - | - | - | - | - | - | - | - | - | - | - | - | - | - | - | - | - | - | - | - | - | - | - | - | - | - | - | - | - | - | - | - | - | - | - | - | - | - | - | - | - | - | - | - | - | - | - | - | - | - | - | - | - | - | - | - | - | - | - | - | - | - | - | - | - | - | - | - | - | - | - | - | - | - | - | - | - | - | - | - | - | - | - | - | - | - | - | - | - | - | - | - | - | - | - | - | - | - | - | - | - | - | - | - | - | - | - | - | - | - | - | - | - | - | - | - | - | T | G | E | L | V | H | T | Y | - | E | E | H | S | - | - | - | - | - | E | Q | V | N | - | - | - | - | - | - | - | - | - | - | - | - | - | C | C | H | - | - | - | - | - | - | - | - | - | - | - | - | - | F | T | N | N | S | - | - | - | - | - | - | - | - | - | - | - | - | - | - | - | - | N | H | - | L | L | L | A | T | G | S | S | - | - | - | - | D | F | F | L | - | - | - | - | - | - | - | - | - | - | - | - | K | L | W | D | L | - | - | - | N | Q | K | E | C | R | N | T | M | F | - | - | - | - | - | - | - | - | - | - | - | - | - | - | - | - | - | G | H | T | N | S | - | - | - | - | - | - | - | - | - | - | - | - | - | - | - | - | I | N | H | C | R | F | S | P | D | D | E | - | - | - | - | - | - | - | - | - | - | - | - | - | - | - | - | - | - | - | - | - | - | - | - | - | - | - | - | L | L | A | S | C | S | A | D | G | T | L | K | L | - | - | - | - | - | - | - | - | - | - | - | - | - | - | - | - | - | - | - | W | D | - | V | R | S | A | N | E | R | K | S | - | - | - | - | - | - | I | N | - | - | - | - | - | - | - | - | - | - | - | - | - | - | - | - | - | - | - | - | - | - | V | K | Q | F | - | - | - | - | - | - | - | - | F | Q | S | S | E | D | P | Q | E | - | - | - | - | - | - | - | - | - | - | - | - | - | - | - | - | - | - | - | - | - | - | - | - | - | - | - | D | V | E | - | V | I | V | K | C | C | - | - | - | - | - | - | - | - | - | - | - | - | - | - | - | - | S | W | S | A | D | G | A | R | I | L | V | A | A | K | T | K | I | - | - | - | - | - | - | - | - | - | - | - | F | L | F | D | I | H | S | - | - | - | S | G | - | - | - | - | - | - | - | - | - | - | - | - | - | - | - | - | - | - | - | - | - | - | - | - | - | - | - | - | - | - | - | - | - | - | - | - | - | - | - | - | - | - | - | - | - | - | - | - | - | - | - | - | - | - | - | - | - | - | - | - | - | - | - | - | - | - | - | - | - | - | - | - | - | - | - | - | - | - | - | - | - | - | - | - | - | - | - | - | - | - | - | - | - | - | - | - | - | - | - | - | - | - | - | - | - | - | - | - | - | - | - | - | - | - | - | - | - | - | - | - | - | - | - | - | - | - | L | L | A | - | - | - | - | - | - | - | - | - | - | - | - | - | - | - | - | - | - | - | - | - | - | - | - | - | - | - | - | - | - | - | - | - | - | - | - | - | - | - | - | - | - | - | - | - | - | - | - | - | - | - | - | - | - | - | - | - | - | - | - | - | - | - | - | - | - | - | - | - | - | - | - | - | - | - | - | - | - | - | - | - | - | - | - | - | - | - | - | - | - | - | - | - | - | - | - | - | - | - | - | - | - | - | - | - | - | - | - | - | - | - | - | - | - | - | - | - | - | - | - | - | - | - | - | - | - | - | - | - | - | - | - | - | - | - | - | - | - | - | - | - | - | - | - | - | - | - | - | - | - | - | - | - | - | - | - | - | - | - | - | - | - | - | - | - | - | - | - | - | - | - | - | - | - | - | - | - | - | - | - | - | - | - | E | I | - | - | - | H | T | G | H | H | S | T | I | Q | Y | C | - | - | - | - | - | - | - | - | - | - | - | - | - | - | - | - | - | - | - | - | - | - | - | - | - | D | F | S | P | C | N | H | L | A | V | V | A | L | S | Q | - | - | - | - | - | - | Y | C | V | E | - | - | - | - | - | - | - | - | - | - | - | - | - | - | - | - | - | L | W | N | I | D | S | C | - | - | - | - | - | - | - | - | - | - | - | - | - | - | - | - | - | - | - | - | - | - | - | - | - | - | - | - | - | L | K | V | A | D | C | R | G | H | L | S | - | - | - | - | - | - | - | - | - | - | - | - | - | - | - | - | - | - | - | - | - | - | - | - | - | - | - | - | - | - | - | - | - | - | - | - | - | - | - | - | - | - | - | - | W | I | H | S | V | M | F | S | - | - | - | - | - | - | - | - | - | - | - | P | D | G | S | - | - | - | - | - | - | - | - | - | - | S | F | L | T | S | S | D | D | Q | - | - | - | - | - | - | - | - | - | - | - | - | - | - | T | I | R | V | - | W | - | - | - | - | - | - | - | - | - | - | - | - | - | - | - | - | - | - | - | - | - | - | - | - | - | - | - | - | - | - | - | - | - | - | - | - | - | - | E | T | K | K | V | C | K | N | S | A | I | V | L | K | Q | E | - | - | - | - | - | - | - | - | - | - | - | - | - | - | - | - | I | D | V | V | F | - | - | - | - | - | - | - | - | - | - | - | - | - | - | - | - | - | - | - | - | - | - | - | - | - | - | - | - | - | - | - | - | - | - | - | - | - | - | - | - | - | - | - | - | - | - | - | - | - | - | - | - | - | - | Q | E | N | E | V | T | - | - | - | - | - | - | - | - | - | - | - | - | - | - | - | - | - | - | - | - | - | - | - | - | - | - | - | - | - | - | - | - | - | - | - | - | V | L | A | V | D | N | I | R | R | L | Q | V | - | I | N | G | K | T | - | - | - | - | - | - | - | - | - | - | G | H | I | D | Y | L | T | E | A | - | - | - | - | - | - | - | - | - | - | - | - | - | - | - | - | - | - | - | - | - | - | Q | - | - | - | - | - | - | - | - | - | - | - | I | S | C | C | C | L | S | P | H | L | Q | - | - | - | - | - | - | - | - | - | Y | I | A | Y | G | K | - | - | - | - | - | - | - | - | - | - | - | - | - | - | - | - | - | - | - | - | - | - | - | - | - | - | - | - | - | - | - | - | - | - | - | - | E | - | - | - | - | - | - | - | - | - | D | G | A | - | - | - | - | - | - | - | - | - | - | I | - | - | - | - | - | - | - | - | - | - | - | - | - | - | - | - | - | - | - | - | - | - | - | - | - | - | - | - | - | - | - | - | - | - | - | - | - | - | - | E | I | L | E | L | L | N | N | R | V | F | K | S | K | N | R | - | - | - | - | - | H | - | - | - | - | - | - | - | - | - | - | - | - | K | K | P | V | R | H | I | Q | F | T | - | - | - | - | - | - | - | - | - | - | - | - | - | - | - | - | - | - | - | - | - | - | - | - | - | - | - | - | - | - | - | - | - | - | - | - | - | T | D | G | R | T | L | I | S | S | S | - | D | D | S | - | - | - | - | - | - | - | - | A | I | Q | - | V | - | - | - | - | - | - | - | - | - | - | - | - | - | - | - | - | - | - | - | - | - | - | - | - | - | - | - | - | - | - | - | - | - | - | - | - | - | - | - | W | N | - | - | - | - | - | - | - | - | - | - | W | Q | S | E | E | Y | V | S | - | - | - | - | - | - | - | - | - | - | - | - | - | - | - | - | - | - | - | - | - | - | - | - | - | - | - | - | - | - | - | - | - | - | - | - | - | - | - | - | - | - | - | - | - | - | - | - | - | - | L | Q | A | H | Q | - | - | - | - | - | - | - | - | - | - | - | - | - | - | - | - | - | - | - | - | - | - | - | - | - | - | E | T | V | K | D | F | R | L | L | - | - | - | - | - | - | - | - | - | - | - | - | - | - | - | - | - | - | - | - | - | - | - | K | N | - | S | R | L | L | - | S | W | - | - | - | S | F | D | G | T | V | - | - | - | - | - | - | - | - | - | - | - | - | - | - | - | - | - | - | - | - | - | - | - | - | - | - | - | - | - | - | - | - | - | - | - | K | V | W | S | V | I | T | G | R | I | - | - | - | - | - | E | K | D | F | A | C | - | - | - | - | - | - | - | H | Q | - | - | - | - | - | - | - | - | - | - | - | - | - | - | - | - | - | - | - | G | T | V | L | - | - | - | - | - | - | - | - | - | - | - | - | - | - | - | S | C | D | - | - | - | - | - | - | - | - | - | - | - | - | - | - | - | I | S | S | D | A | T | - | K | F | S | S | T | - | S | A | D | K | T | - | - | - | - | - | - | A | K | I | W | S | F | E | H | L | - | - | - | - | - | - | - | - | - | - | - | - | - | - | - | - | - | - | - | - | - | - | - | - | S | P | L | H | E | - | - | - | - | - | - | - | - | L | - | - | - | - | K | G | H | N | G | C | V | - | - | - | - | - | - | - | - | - | R | C | S | - | A | F | S | - | - | - | - | - | - | - | - | - | V | D | S | T | - | L | L | A | T | - | - | - | - | - | - | - | - | G | D | D | N | G | E | I | R | I | W | N | V | S | N | G | E | L | L | - | - | - | - | - | - | - | - | - | - | - | - | - | - | - | - | - | - | - | - | - | - | - | - | - | - | - | - | - | - | - | - | - | - | - | - | - | - | - | - | - | - | - | - | - | - | - | - | - | - | - | - | - | - | - | - | H | L | C | A | P | I | S | V | E | - | - | - | - | - | - | - | - | - | - | - | - | - | - | - | - | - | - | - | - | E | G | A | A | T | H | G | G | W | - | - | - | - | - | - | - | - | V | T | D | L | C | F | S | P | D | S | K | - | - | - | - | - | - | - | - | - | - | - | - | - | - | - | - | - | - | - | - | - | - | - | - | - | - | - | - | - | - | - | - | - | - | - | - | - | - | - | - | - | - | - | - | - | M | L | I | S | - | - | A | G | G | Y | L | K | - | - | - | - | - | - | - | - | W | W | N | A | V | T | - | - | - | - | - | - | - | - | - | - | - | - | - | - | - | - | - | - | - | - | - | - | - | - | - | - | - | - | - | - | - | - | - | - | - | - | - | - | - | - | - | - | - | - | - | - | - | - | - | - | - | - | - | - | - | - | - | - | - | - | - | - | - | - | - | - | - | - | - | - | - | - | - | - | - | - | - | - | - | - | - | - | - | - | - | - | - | - | - | - | - | - | - | - | - | - | - | - | - | - | - | - | - | - | - | - | - | - | - | - | - | - | - | - | - | - | - | - | - | - | - | - | - | - | - | - | - | - | - | - | - | - | - | - | - | - | - | - | - | - | - | - | - | - | - | - | - | - | - | - | - | - | - | - | - | - | - | - | - | - | - | - | - | - | - | - | - | - | - | - | - | - | - | - | - | - | - | - | - | - | - | - | - | - | - | - | - | - | - | - | - | - | - | - | - | - | - | - | - | - | - | - | - | - | - | - | - | - | - | - | - | - | - | - | - | - | - | - | - | - | - | - | - | - | - | - | - | - | - | - | - | - | - | - | - | - | - | - | - | - | - | - | - | - | - | - | - | - | - | - | - | - | - | - | - | - | - | - | - | - | - | - | - | - | - | - | - | G | E | S | L | Q | T | F | Y | T | - | - | - | - | - | N | G | - | - | - | T | N | L | K | K | I | - | - | - | H | V | S | P | - | - | - | - | D | F | R | T | - | - | - | - | - | - | - | - | - | - | - | - | - | - | - | - | - | - | - | - | - | - | - | - | - | - | Y | V | T | V | D | - | N | L | G | I | L | - | - | Y | I | L | Q | I | - | - | - | - | - | - | - | - | - | - | - | - | - | - | - | - | - | - | - | - | - | - | - | - | - | - | - | - | - | - | - | - | - | - | L | E | - | - | - | - | - | - | - | - | - | - | - | - | - | - | - | - | - | - | - | - | - | - | - | - | - | - | - | - | - | - | - | - | - | - | - | - | - | - | - | - | - | - | - | - | - | - | - | - | - | - | - | - | M | - | - | - | - | - | - | - | - | - | - | - | - | - | - | - | - | - | - | - | - | - | - | - | - | - | - | - | - | - | - | - | - | - | - | - | - | - | - | - | - | - | - | - | - | - | - | - | - | - | - | - | - | - | - | D | D | A | D | R | H | - | - | - | - | - | - | - | - | - | - | - | - | - | - | - | - | - | - | - | - | L | L | R | R | - | - | - | - | - | - | - | - | - | - | - | - | - | - | - | - | - | - | - | - | - | - | - | - | - | - | - | - | - | - | C | R | - | - | - | - | - | - | - | - | M | R | L | - | V | G | E | L | Q | V | A | S | L | W | - | - | - | - | - | - | - | - | - | - | - | - | - | - | - | - | - | - | - | - | - | - | - | - | - | - | - | - | - | - | - | - | - | - | - | - | - | - | - | - | - | - | - | - | - | D | A | L | L | N | R | E | L | - | - | - | - | - | - | - | - | - | - | - | - | - | - | - | - | - | - | - | - | - | - | - | - | F | T | P | D | M | I | E | D | - | - | - | - | - | - | - | - | - | - | - | - | - | - | - | - | - | - | - | - | - | - | - | - | - | I | Q | R | A | G | - | - | - | - | - | - | - | - | S | G | S | R | R | D | Q | - | A | R | Q | L | V | T | D | L | E | T | R | G | - | R | Q | A | L | P | L | F | I | S | C | L | E | D | T | G | Q | D | T | L | A | S | F | L | R | T | S | R | Q | A | G | - | - | - | - | K | Q | D | P | - | E | A | I | - | - | - | - | - | - | - | - | - | - | - | - | - | - | - | - | - | - | - | - | - | - | - | - | - | - | R | P | L | D | - | - | - | - | - | - | - | - | - | - | - | - | - | - | - | L | V | P | V | - | - | - | - | - | - | - | - | - | - | - | - | - | - | - | - | - | - | - | - | - | - | - | - | - | - | - | - | - | - | - | - | - | - | - | - | - | - | - | D | L | G | P | I | G | L | K | P | E | E | P | R | - | G | M | K | L | D | P | S | P | P | - | A | - | P | - | - | - | - | - | - | - | - | - | - | - | - | - | - | - | - | - | - | - | - | - | - | - | - | - | - | - | - | - | - | - | - | - | - | - | - | - | - | - | - | - | - | - | - | - | - | - | - | - | - | - | G | N | L | T | P | V | - | - | - | - | - | - | - | - | - | - | - | - | - | - | - | - | - | - | - | - | - | - | - | - | - | - | V | L | - | - | - | - | - | - | - | - | - | - | - | - | - | - | - | - | - | - | - | - | - | - | - | - | - | - | - | - | - | - | - | - | - | - | - | - | - | - | - | - | - | - | - | - | - | - | - | - | - | - | - | - | - | - | - | - | - | - | - | - | - | - | - | - | - | - | - | - | G | P | E | E | L | W | P | - | - | - | - | - | - | - | - | - | - | - | - | - | - | - | - | - | - | - | - | - | - | - | - | - | - | A | Q | L | R | P | - | - | - | - | - | - | E | V | L | R | P | - | E | T | - | - | - | - | - | - | - | - | - | - | - | - | - | - | - | - | - | - | P | R | P | V | D | I | G | S | G | E | S | - | - | - | - | - | - | - | - | - | - | - | - | - | R | D | A | C | - | - | - | - | - | - | - | - | - | - | P | - | Q | E | K | L | R | G | N | A | D | - | - | - | - | - | L | A | Y | F | - | - | L | D | A | D | - | P | C | G | H | C | L | I | I | N | N | V | N | F | C | H | A | S | - | - | G | L | S | M | R | T | G | S | N | V | D | C | E | K | L | Q | H | R | F | - | R | W | L | H | F | E | V | E | V | K | C | D | L | T | A | K | K | M | V | Q | A | L | V | E | L | A | Q | R | D | H | - | R | A | L | D | C | C | V | V | V | I | L | S | H | G | C | Q | A | S | H | L | Q | F | P | G | A | - | V | Y | G | T | D | G | Y | A | L | S | V | E | R | I | V | N | I | F | N | G | T | G | C | P | S | L | G | G | K | P | K | L | F | F | I | Q | A | C | - | - | G | G | E | Q | K | D | H | G | F | E | V | S | Q | A | - | - | - | S | P | E | D | - | R | A | S | D | S | N | - | - | - | - | - | - | - | S | E | P | D | A | I | P | F | Q | E | G | L | R | T | F | - | - | - | - | - | - | - | - | D | Q | L | D | A | V | - | - | - | - | - | - | - | S | S | L | - | - | - | P | T | P | S | D | I | L | V | S | Y | S | T | F | P | - | - | G | - | - | - | - | - | - | - | - | F | V | S | W | R | D | T | K | S | G | S | W | Y | I | E | T | L | D | G | V | L | E | Q | W | A | H | - | S | - | - | - | - | - | - | - | - | - | - | - | - | - | - | - | - | - | - | - | - | - | - | - | - | - | - | - | - | - | - | - | E | D | L | Q | S | L | L | - | - | - | - | - | L | R | - | - | - | - | - | - | - | - | - | - | - | - | V | A | N | A | V | - | - | - | - | - | S | E | K | - | - | - | - | - | - | - | G | I | Y | - | K | Q | I | - | - | P | G | - | - | - | - | - | - | - | - | - | - | - | - | - | - | - | C | F | N | F | - | - | - | - | - | - | - | - | - | L | R | K | K | L | F | F | K | T | S | - | - | - | - | - | - | - | - | - | - | - | - | - | - | - | - | - | - | - | - | - | - | - | - | - | - | - | - | - | - | - | - | - | - | - | - | - | M | - | - | - | - | - | - | - | - | - | - | - | - | - | - | - | - | - | - | - | - | - | - | - | - | - | - | - | - | - | - | - | - | - | - | - | - | - | - | - | - | - | - | - | - | - | - | - | - | - | - | - | - | G | D | V | E | K | G | K | K | I | F | V | Q | K | C | A | Q | C | H | T | V | E | K | G | G | K | H | K | T | G | P | N | L | H | G | L | F | G | R | K | T | G | Q | A | P | G | F | S | Y | T | D | A | N | K | N | K | G | I | T | W | G | E | D | T | L | M | E | Y | L | E | N | P | K | K | Y | I | P | G | T | K | M | I | F | A | G | I | K | K | K | G | E | R | A | D | L | I | A | Y | L | K | K | A | T | N | E | - | - | - | - | - | - |
| Cebus\_capucinus\_imitator | M | - | - | - | - | - | - | - | - | - | - | - | - | - | - | - | - | - | - | - | - | - | - | - | - | - | - | - | - | - | - | - | - | - | - | - | - | - | - | - | - | - | - | - | - | - | - | - | - | - | - | - | - | - | - | - | - | - | - | - | - | - | - | - | - | - | - | - | - | - | - | - | - | - | - | - | - | - | - | - | - | - | - | - | - | - | - | - | - | - | - | - | - | - | - | - | - | - | D | A | K | A | - | R | N | - | - | - | - | - | - | - | - | - | - | - | - | - | - | - | - | C | L | L | - | - | - | - | - | - | - | - | - | - | - | - | - | - | - | - | - | - | - | - | - | - | - | - | - | - | - | - | - | - | - | - | - | - | - | - | - | - | - | - | - | - | - | - | - | - | - | - | - | - | - | - | - | - | - | - | - | - | - | - | - | - | - | - | - | - | - | - | - | - | - | - | - | - | - | - | - | - | - | - | - | - | - | - | - | - | - | - | - | - | - | - | - | - | - | - | - | - | - | - | - | - | - | - | - | - | - | - | - | - | - | - | - | - | - | - | - | - | - | - | - | Q | H | R | E | A | L | E | K | D | I | K | T | S | Y | - | I | M | D | H | M | I | S | D | G | F | L | T | I | L | E | E | E | K | V | R | N | E | - | - | - | - | - | - | - | - | - | - | - | - | - | - | - | - | - | P | T | Q | Q | Q | R | A | A | M | L | I | K | M | I | L | K | K | - | - | - | - | D | N | G | S | Y | I | S | - | - | - | - | - | F | Y | N | A | L | - | L | H | E | G | - | - | - | - | - | - | Y | K | D | L | A | A | L | L | H | D | G | I | P | V | V | S | S | S | S | G | - | K | - | - | - | - | - | - | - | - | - | - | - | - | - | - | - | - | - | - | - | - | - | - | D | S | V | S | G | I | T | - | - | - | - | - | - | - | - | - | - | - | S | Y | V | R | T | V | L | C | E | G | G | - | - | - | - | - | - | - | V | P | Q | R | P | V | V | F | V | A | R | K | K | L | V | N | A | I | - | - | - | - | - | - | - | - | - | - | Q | Q | K | L | S | - | - | - | - | - | - | - | - | - | - | - | - | - | - | - | - | - | - | - | - | - | - | - | - | - | - | - | - | - | - | - | - | K | L | N | G | E | P | - | - | - | - | - | - | - | - | - | - | - | - | - | - | - | - | - | - | - | - | - | - | - | - | - | - | - | - | - | - | - | - | - | - | - | - | - | - | G | W | V | T | I | Y | G | M | A | G | C | G | K | S | V | L | A | A | E | A | V | - | R | D | H | S | L | L | K | R | C | F | P | G | G | V | H | W | V | S | V | G | - | - | - | - | - | - | - | K | - | - | - | - | - | - | Q | D | K | S | - | - | - | - | - | - | - | - | - | - | - | - | G | L | L | M | K | L | Q | N | L | C | T | R | L | D | - | - | - | - | Q | - | - | - | - | - | - | - | - | - | - | - | - | - | - | - | D | E | S | F | S | Q | R | L | P | L | N | I | - | - | - | - | - | - | - | - | - | - | - | - | - | - | - | - | E | E | A | K | D | R | L | R | I | L | L | L | R | - | - | - | - | - | - | - | - | - | - | - | - | K | H | P | S | S | L | L | - | - | I | L | D | D | I | W | D | P | W | V | L | K | A | F | D | - | N | Q | C | Q | I | L | L | T | T | R | D | K | S | V | T | D | S | V | - | M | G | P | - | - | N | Y | V | V | P | V | E | S | - | - | - | S | L | G | K | E | K | G | L | E | I | L | S | L | F | V | N | M | - | K | K | A | D | L | P | E | - | - | - | - | - | - | - | - | - | - | - | - | - | - | Q | A | H | S | I | I | K | E | C | K | - | - | - | - | - | - | - | - | - | - | - | - | - | - | - | - | - | - | - | G | - | S | P | L | V | V | S | L | I | G | A | L | L | R | D | F | P | N | - | - | - | - | - | - | - | - | - | - | R | W | E | Y | Y | L | - | R | Q | L | Q | N | K | Q | F | K | R | I | - | - | - | - | - | - | - | - | - | - | - | - | - | - | - | - | - | - | - | - | - | - | R | K | S | S | S | Y | D | - | - | - | - | Y | E | A | L | D | E | A | M | S | I | S | V | E | - | M | L | R | - | - | - | E | D | I | K | N | Y | - | Y | T | D | L | S | I | L | P | K | D | V | K | V | P | T | K | V | L | C | I | L | W | D | - | - | - | - | M | E | T | E | E | V | E | D | I | L | Q | E | F | - | - | - | - | - | - | - | - | - | - | - | V | N | K | S | L | L | F | C | D | R | N | G | K | S | - | - | - | - | - | - | - | - | - | F | R | Y | Y | L | H | D | L | Q | V | D | F | L | T | E | K | - | - | N | C | S | Q | - | L | - | Q | D | L | H | K | K | I | I | T | Q | F | Q | R | - | Y | Y | - | - | - | - | - | - | Q | L | P | S | L | L | P | D | - | Q | E | - | - | - | - | - | - | - | D | C | M | Y | W | Y | N | F | L | A | Y | H | M | A | S | A | K | - | - | - | - | M | H | K | E | L | C | A | L | M | F | S | L | D | W | I | K | A | K | T | E | L | V | G | P | A | - | - | - | - | - | - | - | - | H | L | I | H | E | F | V | E | Y | R | H | I | L | D | E | K | - | - | - | - | - | - | - | D | C | A | V | S | E | N | F | Q | - | - | - | E | F | L | S | L | N | G | H | L | L | G | R | Q | P | F | - | - | - | - | - | - | - | - | - | - | - | - | - | - | - | - | - | - | - | - | - | - | - | - | - | - | - | - | - | - | - | - | - | - | - | - | - | - | - | - | - | - | - | - | - | - | - | - | - | - | - | - | - | - | - | - | - | - | - | - | - | - | - | - | - | - | - | - | - | - | - | P | N | I | V | Q | L | G | L | C | E | P | E | T | S | E | V | Y | Q | Q | A | K | L | Q | A | K | Q | E | V | D | N | G | - | - | - | - | - | - | - | - | - | - | - | - | - | - | - | - | - | - | - | - | - | - | - | - | - | - | - | - | - | T | L | Y | L | - | - | - | - | - | - | - | - | - | - | E | W | T | N | K | - | - | - | - | - | - | - | K | T | I | M | N | L | S | R | - | - | - | - | - | - | - | - | - | - | L | V | V | R | P | H | T | D | - | - | - | - | - | - | - | - | - | - | - | - | - | - | - | A | V | Y | H | A | C | - | - | - | F | S | - | E | D | G | Q | R | I | A | - | - | - | - | - | - | - | - | - | S | C | G | A | D | K | T | L | Q | V | F | K | - | - | - | - | - | A | E | T | G | E | K | L | L | E | I | - | K | A | H | E | D | E | - | - | - | V | L | C | C | A | F | S | - | T | - | - | - | - | - | - | - | - | - | - | - | - | - | D | D | R | F | I | A | T | C | S | V | D | K | K | V | K | I | - | - | - | W | N | S | M | - | - | - | - | - | - | - | - | - | - | - | - | - | - | - | - | - | - | - | - | - | - | - | - | - | - | - | - | - | - | - | - | - | - | - | - | - | - | - | - | - | - | - | - | - | - | - | - | - | - | - | - | - | - | - | - | - | - | - | - | - | - | - | - | - | - | - | - | - | - | - | - | - | - | - | - | - | - | - | - | - | - | - | - | - | - | - | - | - | - | - | - | - | - | - | - | - | - | - | - | - | - | - | - | - | - | - | - | - | - | - | - | - | - | - | - | - | - | - | - | - | - | - | - | - | - | - | - | - | - | - | - | - | - | - | - | - | - | - | - | T | G | E | L | V | H | N | Y | - | D | E | H | S | - | - | - | - | - | E | Q | V | N | - | - | - | - | - | - | - | - | - | - | - | - | - | C | C | H | - | - | - | - | - | - | - | - | - | - | - | - | - | F | T | N | S | S | - | - | - | - | - | - | - | - | - | - | - | - | - | - | - | - | H | H | - | F | L | L | A | T | G | S | S | - | - | - | - | D | C | V | L | - | - | - | - | - | - | - | - | - | - | - | - | K | L | W | D | L | - | - | - | N | Q | K | P | C | R | N | T | L | F | - | - | - | - | - | - | - | - | - | - | - | - | - | - | - | - | - | G | H | I | N | S | - | - | - | - | - | - | - | - | - | - | - | - | - | - | - | - | V | N | H | C | R | F | S | P | D | D | K | - | - | - | - | - | - | - | - | - | - | - | - | - | - | - | - | - | - | - | - | - | - | - | - | - | - | - | - | L | L | A | S | C | S | A | D | G | T | L | K | L | - | - | - | - | - | - | - | - | - | - | - | - | - | - | - | - | - | - | - | W | D | - | V | T | S | A | N | E | R | K | S | - | - | - | - | - | - | I | N | - | - | - | - | - | - | - | - | - | - | - | - | - | - | - | - | - | - | - | - | - | - | V | K | Q | F | - | - | - | - | - | - | - | - | F | L | N | S | E | D | P | Q | E | - | - | - | - | - | - | - | - | - | - | - | - | - | - | - | - | - | - | - | - | - | - | - | - | - | - | - | D | M | E | - | V | I | V | K | C | C | - | - | - | - | - | - | - | - | - | - | - | - | - | - | - | - | S | W | S | A | D | G | A | R | I | M | V | A | A | K | N | K | I | - | - | - | - | - | - | - | - | - | - | - | F | L | F | D | I | H | T | - | - | - | N | D | - | - | - | - | - | - | - | - | - | - | - | - | - | - | - | - | - | - | - | - | - | - | - | - | - | - | - | - | - | - | - | - | - | - | - | - | - | - | - | - | - | - | - | - | - | - | - | - | - | - | - | - | - | - | - | - | - | - | - | - | - | - | - | - | - | - | - | - | - | - | - | - | - | - | - | - | - | - | - | - | - | - | - | - | - | - | - | - | - | - | - | - | - | - | - | - | - | - | - | - | - | - | - | - | - | - | - | - | - | - | - | - | - | - | - | - | - | - | - | - | - | - | - | - | - | - | L | L | G | - | - | - | - | - | - | - | - | - | - | - | - | - | - | - | - | - | - | - | - | - | - | - | - | - | - | - | - | - | - | - | - | - | - | - | - | - | - | - | - | - | - | - | - | - | - | - | - | - | - | - | - | - | - | - | - | - | - | - | - | - | - | - | - | - | - | - | - | - | - | - | - | - | - | - | - | - | - | - | - | - | - | - | - | - | - | - | - | - | - | - | - | - | - | - | - | - | - | - | - | - | - | - | - | - | - | - | - | - | - | - | - | - | - | - | - | - | - | - | - | - | - | - | - | - | - | - | - | - | - | - | - | - | - | - | - | - | - | - | - | - | - | - | - | - | - | - | - | - | - | - | - | - | - | - | - | - | - | - | - | - | - | - | - | - | - | - | - | - | - | - | - | - | - | - | - | - | - | - | - | - | - | - | E | I | - | - | - | H | M | G | H | H | S | T | I | Q | Y | C | - | - | - | - | - | - | - | - | - | - | - | - | - | - | - | - | - | - | - | - | - | - | - | - | - | D | F | S | P | Q | N | H | L | A | V | V | A | L | S | Q | - | - | - | - | - | - | Y | C | V | E | - | - | - | - | - | - | - | - | - | - | - | - | - | - | - | - | - | L | W | N | I | D | S | C | - | - | - | - | - | - | - | - | - | - | - | - | - | - | - | - | - | - | - | - | - | - | - | - | - | - | - | - | - | S | K | V | A | D | C | R | G | H | L | S | - | - | - | - | - | - | - | - | - | - | - | - | - | - | - | - | - | - | - | - | - | - | - | - | - | - | - | - | - | - | - | - | - | - | - | - | - | - | - | - | - | - | - | - | W | V | H | G | V | M | F | S | - | - | - | - | - | - | - | - | - | - | - | P | D | G | S | - | - | - | - | - | - | - | - | - | - | S | F | L | T | S | S | D | D | Q | - | - | - | - | - | - | - | - | - | - | - | - | - | - | T | I | R | L | - | W | - | - | - | - | - | - | - | - | - | - | - | - | - | - | - | - | - | - | - | - | - | - | - | - | - | - | - | - | - | - | - | - | - | - | - | - | - | - | E | T | K | K | V | C | K | N | S | A | I | M | L | K | Q | E | - | - | - | - | - | - | - | - | - | - | - | - | - | - | - | - | V | D | V | V | F | - | - | - | - | - | - | - | - | - | - | - | - | - | - | - | - | - | - | - | - | - | - | - | - | - | - | - | - | - | - | - | - | - | - | - | - | - | - | - | - | - | - | - | - | - | - | - | - | - | - | - | - | - | - | Q | E | N | E | V | M | - | - | - | - | - | - | - | - | - | - | - | - | - | - | - | - | - | - | - | - | - | - | - | - | - | - | - | - | - | - | - | - | - | - | - | - | V | L | A | V | D | H | I | K | R | L | Q | L | - | I | N | G | K | T | - | - | - | - | - | - | - | - | - | - | G | Q | I | D | Y | L | T | E | A | - | - | - | - | - | - | - | - | - | - | - | - | - | - | - | - | - | - | - | - | - | - | Q | - | - | - | - | - | - | - | - | - | - | - | V | S | C | C | C | L | S | P | H | L | Q | - | - | - | - | - | - | - | - | - | Y | I | A | F | G | D | - | - | - | - | - | - | - | - | - | - | - | - | - | - | - | - | - | - | - | - | - | - | - | - | - | - | - | - | - | - | - | - | - | - | - | - | E | - | - | - | - | - | - | - | - | - | N | G | A | - | - | - | - | - | - | - | - | - | - | I | - | - | - | - | - | - | - | - | - | - | - | - | - | - | - | - | - | - | - | - | - | - | - | - | - | - | - | - | - | - | - | - | - | - | - | - | - | - | - | E | I | L | E | P | V | N | N | R | I | F | Q | S | R | I | R | - | - | - | - | - | H | - | - | - | - | - | - | - | - | - | - | - | - | K | K | T | V | R | H | I | Q | F | T | - | - | - | - | - | - | - | - | - | - | - | - | - | - | - | - | - | - | - | - | - | - | - | - | - | - | - | - | - | - | - | - | - | - | - | - | - | A | D | E | K | T | L | I | S | S | S | - | D | D | S | - | - | - | - | - | - | - | - | A | I | Q | - | V | - | - | - | - | - | - | - | - | - | - | - | - | - | - | - | - | - | - | - | - | - | - | - | - | - | - | - | - | - | - | - | - | - | - | - | - | - | - | - | W | N | - | - | - | - | - | - | - | - | - | - | W | Q | L | E | E | Y | D | V | - | - | - | - | - | - | - | - | - | - | - | - | - | - | - | - | - | - | - | - | - | - | - | - | - | - | - | - | - | - | - | - | - | - | - | - | - | - | - | - | - | - | - | - | - | - | - | - | - | - | L | Q | A | H | Q | - | - | - | - | - | - | - | - | - | - | - | - | - | - | - | - | - | - | - | - | - | - | - | - | - | - | E | T | V | K | D | F | R | L | L | - | - | - | - | - | - | - | - | - | - | - | - | - | - | - | - | - | - | - | - | - | - | - | K | N | - | S | R | L | L | - | S | W | - | - | - | S | F | D | G | T | V | - | - | - | - | - | - | - | - | - | - | - | - | - | - | - | - | - | - | - | - | - | - | - | - | - | - | - | - | - | - | - | - | - | - | - | K | V | W | N | I | I | T | G | K | V | - | - | - | - | - | E | K | D | F | V | C | - | - | - | - | - | - | - | H | Q | - | - | - | - | - | - | - | - | - | - | - | - | - | - | - | - | - | - | - | G | T | V | L | - | - | - | - | - | - | - | - | - | - | - | - | - | - | - | S | C | D | - | - | - | - | - | - | - | - | - | - | - | - | - | - | - | I | S | H | D | A | T | - | K | F | S | S | T | - | S | A | D | K | T | - | - | - | - | - | - | A | K | I | W | S | F | D | L | L | - | - | - | - | - | - | - | - | - | - | - | - | - | - | - | - | - | - | - | - | - | - | - | - | S | P | L | H | E | - | - | - | - | - | - | - | - | L | - | - | - | - | R | G | H | N | G | C | V | - | - | - | - | - | - | - | - | - | R | C | S | - | A | F | S | - | - | - | - | - | - | - | - | - | V | D | S | T | - | L | L | A | T | - | - | - | - | - | - | - | - | G | D | D | N | G | E | I | R | I | W | N | V | S | N | G | E | L | L | - | - | - | - | - | - | - | - | - | - | - | - | - | - | - | - | - | - | - | - | - | - | - | - | - | - | - | - | - | - | - | - | - | - | - | - | - | - | - | - | - | - | - | - | - | - | - | - | - | - | - | - | - | - | - | - | H | L | C | A | P | L | S | - | E | - | - | - | - | - | - | - | - | - | - | - | - | - | - | - | - | - | - | - | - | E | G | A | A | T | H | G | G | W | - | - | - | - | - | - | - | - | V | T | D | L | C | F | S | P | D | G | K | - | - | - | - | - | - | - | - | - | - | - | - | - | - | - | - | - | - | - | - | - | - | - | - | - | - | - | - | - | - | - | - | - | - | - | - | - | - | - | - | - | - | - | - | - | M | L | I | S | - | - | A | G | G | Y | L | K | - | - | - | - | - | - | - | - | W | W | N | V | V | T | - | - | - | - | - | - | - | - | - | - | - | - | - | - | - | - | - | - | - | - | - | - | - | - | - | - | - | - | - | - | - | - | - | - | - | - | - | - | - | - | - | - | - | - | - | - | - | - | - | - | - | - | - | - | - | - | - | - | - | - | - | - | - | - | - | - | - | - | - | - | - | - | - | - | - | - | - | - | - | - | - | - | - | - | - | - | - | - | - | - | - | - | - | - | - | - | - | - | - | - | - | - | - | - | - | - | - | - | - | - | - | - | - | - | - | - | - | - | - | - | - | - | - | - | - | - | - | - | - | - | - | - | - | - | - | - | - | - | - | - | - | - | - | - | - | - | - | - | - | - | - | - | - | - | - | - | - | - | - | - | - | - | - | - | - | - | - | - | - | - | - | - | - | - | - | - | - | - | - | - | - | - | - | - | - | - | - | - | - | - | - | - | - | - | - | - | - | - | - | - | - | - | - | - | - | - | - | - | - | - | - | - | - | - | - | - | - | - | - | - | - | - | - | - | - | - | - | - | - | - | - | - | - | - | - | - | - | - | - | - | - | - | - | - | - | - | - | - | - | - | - | - | - | - | - | - | - | - | - | - | - | - | - | - | - | - | - | G | Q | S | L | Q | T | F | Y | T | - | - | - | - | - | N | G | - | - | - | T | N | L | K | K | I | - | - | - | H | V | S | P | - | - | - | - | D | F | K | T | - | - | - | - | - | - | - | - | - | - | - | - | - | - | - | - | - | - | - | - | - | - | - | - | - | - | Y | V | T | V | D | - | N | L | G | I | L | - | - | Y | I | L | Q | T | - | - | - | - | - | - | - | - | - | - | - | - | - | - | - | - | - | - | - | - | - | - | - | - | - | - | - | - | - | - | - | - | - | - | L | E | - | - | - | - | - | - | - | - | - | - | - | - | - | - | - | - | - | - | - | - | - | - | - | - | - | - | - | - | - | - | - | - | - | - | - | - | - | - | - | - | - | - | - | - | - | - | - | - | - | - | - | - | M | - | - | - | - | - | - | - | - | - | - | - | - | - | - | - | - | - | - | - | - | - | - | - | - | - | - | - | - | - | - | - | - | - | - | - | - | - | - | - | - | - | - | - | - | - | - | - | - | - | - | - | - | - | - | D | E | A | D | R | L | - | - | - | - | - | - | - | - | - | - | - | - | - | - | - | - | - | - | - | - | L | L | R | R | - | - | - | - | - | - | - | - | - | - | - | - | - | - | - | - | - | - | - | - | - | - | - | - | - | - | - | - | - | - | H | R | - | - | - | - | - | - | - | - | R | L | L | - | V | E | K | L | Q | V | D | Q | L | W | - | - | - | - | - | - | - | - | - | - | - | - | - | - | - | - | - | - | - | - | - | - | - | - | - | - | - | - | - | - | - | - | - | - | - | - | - | - | - | - | - | - | - | - | - | D | A | L | L | S | R | E | L | - | - | - | - | - | - | - | - | - | - | - | - | - | - | - | - | - | - | - | - | - | - | - | - | F | K | P | H | M | I | E | D | - | - | - | - | - | - | - | - | - | - | - | - | - | - | - | - | - | - | - | - | - | - | - | - | - | I | Q | R | A | G | - | - | - | - | - | - | - | - | C | G | S | R | R | D | Q | - | A | R | Q | L | I | T | D | L | E | S | R | G | - | S | Q | A | F | P | L | F | I | S | C | L | E | E | T | G | Q | C | M | L | A | S | I | L | Q | T | N | R | Q | T | E | - | - | - | - | K | F | - | - | - | - | - | - | - | - | - | - | - | - | - | - | - | - | - | - | - | - | - | - | - | - | - | - | - | - | - | - | - | - | - | - | - | - | - | - | - | - | - | - | - | - | - | - | - | - | - | - | - | L | I | P | - | - | - | - | - | - | - | - | - | - | - | - | - | - | - | - | - | - | - | - | - | - | - | - | - | - | - | - | - | - | - | - | - | - | - | - | - | - | - | - | - | - | - | - | - | - | - | - | - | - | - | - | - | - | - | - | - | - | - | - | - | - | - | - | - | - | - | - | - | - | - | - | - | - | - | - | - | - | - | - | - | - | - | - | - | - | - | - | - | - | - | - | - | - | - | - | - | - | - | - | - | - | - | - | - | - | - | - | - | - | - | - | - | - | - | - | - | - | - | - | - | - | - | - | - | - | - | - | - | - | - | - | - | - | - | - | - | - | - | - | - | - | - | - | - | - | - | - | - | - | - | - | - | - | - | - | - | - | - | - | - | - | - | - | - | - | - | - | - | - | - | - | - | - | - | - | - | - | - | - | - | - | - | - | - | - | - | - | - | - | - | - | - | - | - | - | - | - | - | - | - | - | - | - | - | - | - | - | - | - | - | - | - | - | - | - | - | - | - | - | - | - | - | - | - | - | - | - | - | - | - | - | - | - | - | - | - | - | - | - | - | - | - | - | - | - | - | - | - | - | - | - | - | - | - | - | - | - | - | - | - | - | - | - | - | - | - | K | P | - | K | D | - | - | - | - | - | - | - | - | - | - | - | - | - | - | - | - | - | - | P | R | P | V | D | I | G | S | G | G | S | - | - | - | - | - | - | - | - | - | - | - | - | - | G | D | G | G | - | - | - | - | - | - | - | - | - | - | A | - | P | E | I | L | R | Q | N | T | D | - | - | - | - | - | L | A | Y | V | - | - | M | S | T | E | - | P | C | G | H | C | L | I | I | N | N | V | N | F | S | P | E | S | - | - | R | L | S | T | R | T | G | S | D | I | D | C | G | K | L | Q | L | R | F | - | S | S | L | H | F | V | V | E | V | K | R | D | L | T | A | K | E | M | G | L | A | L | L | E | L | A | R | R | D | H | - | G | A | L | D | C | C | V | V | V | I | L | S | H | G | C | Q | A | S | H | L | Q | F | P | G | A | - | V | Y | G | T | D | G | Y | P | V | S | L | E | K | I | V | N | I | F | N | G | T | S | C | P | S | L | G | G | K | P | K | L | F | F | I | Q | A | C | - | - | G | G | E | Q | K | D | H | G | F | E | V | A | S | T | - | - | - | S | P | E | E | - | E | S | P | G | S | N | - | - | - | - | - | - | - | P | E | S | D | A | T | P | F | Q | E | D | A | W | T | F | - | - | - | - | - | - | - | - | D | Q | V | D | A | V | - | - | - | - | - | - | - | S | S | L | - | - | - | P | T | P | S | D | I | F | V | S | Y | S | T | F | P | - | - | G | - | - | - | - | - | - | - | - | F | V | S | W | R | E | P | K | R | G | S | W | Y | V | E | T | L | D | D | I | F | E | H | W | A | H | - | S | - | - | - | - | - | - | - | - | - | - | - | - | - | - | - | - | - | - | - | - | - | - | - | - | - | - | - | - | - | - | - | E | D | L | Q | S | L | L | - | - | - | - | - | L | R | - | - | - | - | - | - | - | - | - | - | - | - | V | A | N | A | V | - | - | - | - | - | S | G | K | - | - | - | - | - | - | - | G | I | Y | - | K | Q | M | - | - | P | G | - | - | - | - | - | - | - | - | - | - | - | - | - | - | - | F | F | N | F | - | - | - | - | - | - | - | - | - | L | R | K | K | L | F | F | K | T | S | - | - | - | - | - | - | - | - | - | - | - | - | - | - | - | - | - | - | - | - | - | - | - | - | - | - | - | - | - | - | - | - | - | - | - | - | - | M | - | - | - | - | - | - | - | - | - | - | - | - | - | - | - | - | - | - | - | - | - | - | - | - | - | - | - | - | - | - | - | - | - | - | - | - | - | - | - | - | - | - | - | - | - | - | - | - | - | - | - | - | G | D | V | E | K | G | K | R | I | F | I | Q | K | C | S | Q | C | H | T | V | E | K | G | G | K | H | K | T | G | P | N | L | H | G | I | F | G | R | K | T | G | Q | A | S | G | F | T | Y | T | E | A | N | K | N | K | G | I | I | W | G | E | D | T | L | M | E | Y | L | E | N | P | K | K | Y | I | P | G | T | K | M | I | F | V | G | I | K | K | K | G | E | R | E | D | L | I | A | Y | L | K | K | A | T | N | E | - | - | - | - | - | - |
| Ceratotherium\_simum\_simum | M | - | - | - | - | - | - | - | - | - | - | - | - | - | - | - | - | - | - | - | - | - | - | - | - | - | - | - | - | - | - | - | - | - | - | - | - | - | - | - | - | - | - | - | - | - | - | - | - | - | - | - | - | - | - | - | - | - | - | - | - | - | - | - | - | - | - | - | - | - | - | - | - | - | - | - | - | - | - | - | - | - | - | - | - | - | - | - | - | - | - | - | - | - | - | - | - | - | D | A | K | A | - | R | N | - | - | - | - | - | - | - | - | - | - | - | - | - | - | - | - | C | L | L | - | - | - | - | - | - | - | - | - | - | - | - | - | - | - | - | - | - | - | - | - | - | - | - | - | - | - | - | - | - | - | - | - | - | - | - | - | - | - | - | - | - | - | - | - | - | - | - | - | - | - | - | - | - | - | - | - | - | - | - | - | - | - | - | - | - | - | - | - | - | - | - | - | - | - | - | - | - | - | - | - | - | - | - | - | - | - | - | - | - | - | - | - | - | - | - | - | - | - | - | - | - | - | - | - | - | - | - | - | - | - | - | - | - | - | - | - | - | - | - | - | Q | H | R | E | A | L | E | K | D | I | K | T | S | Y | - | I | M | D | H | M | I | S | D | G | V | L | T | V | L | E | E | E | K | V | K | N | E | - | - | - | - | - | - | - | - | - | - | - | - | - | - | - | - | - | P | T | R | Q | Q | K | A | A | M | L | I | K | L | I | L | K | K | - | - | - | - | D | N | Y | S | Y | I | S | - | - | - | - | - | F | Y | K | A | L | - | L | H | E | G | - | - | - | - | - | - | Y | K | D | L | A | A | R | L | H | G | G | I | P | V | I | S | S | S | N | G | - | K | - | - | - | - | - | - | - | - | - | - | - | - | - | - | - | - | - | - | - | - | - | - | D | S | V | G | G | I | T | - | - | - | - | - | - | - | - | - | - | - | S | Y | V | K | T | V | L | C | E | G | G | - | - | - | - | - | - | - | V | P | Q | R | P | V | V | F | V | T | R | K | K | L | V | N | A | I | - | - | - | - | - | - | - | - | - | - | Q | Q | K | L | F | - | - | - | - | - | - | - | - | - | - | - | - | - | - | - | - | - | - | - | - | - | - | - | - | - | - | - | - | - | - | - | - | K | L | G | G | E | P | - | - | - | - | - | - | - | - | - | - | - | - | - | - | - | - | - | - | - | - | - | - | - | - | - | - | - | - | - | - | - | - | - | - | - | - | - | - | G | W | V | T | I | Y | G | M | A | G | C | G | K | S | V | L | A | A | E | A | V | - | R | D | H | S | F | L | E | G | C | F | P | G | G | V | H | W | V | S | V | G | - | - | - | - | - | - | - | K | - | - | - | - | - | - | Q | D | K | S | - | - | - | - | - | - | - | - | - | - | - | - | G | L | L | M | K | L | Q | N | L | C | T | R | L | D | - | - | - | - | Q | - | - | - | - | - | - | - | - | - | - | - | - | - | - | - | D | E | S | F | S | Q | R | L | P | L | N | I | - | - | - | - | - | - | - | - | - | - | - | - | - | - | - | - | E | E | A | K | D | R | L | R | I | L | M | L | R | - | - | - | - | - | - | - | - | - | - | - | - | K | H | P | R | S | L | L | - | - | I | L | D | D | I | W | D | P | W | V | L | K | A | F | D | - | N | Q | C | Q | I | L | L | T | T | R | D | K | S | V | T | D | S | V | - | M | G | L | - | - | K | Y | L | V | P | V | E | S | - | - | - | G | L | G | K | E | K | G | L | E | I | L | S | L | F | V | N | M | - | K | K | A | D | L | P | E | - | - | - | - | - | - | - | - | - | - | - | - | - | - | Q | A | H | S | I | I | K | E | C | K | - | - | - | - | - | - | - | - | - | - | - | - | - | - | - | - | - | - | - | G | - | S | P | L | V | V | S | L | I | G | A | L | L | R | D | F | P | N | - | - | - | - | - | - | - | - | - | - | R | W | E | Y | Y | L | - | R | Q | L | Q | N | K | Q | F | K | R | I | - | - | - | - | - | - | - | - | - | - | - | - | - | - | - | - | - | - | - | - | - | - | R | K | S | S | S | Y | D | - | - | - | - | Y | E | A | L | D | E | A | M | S | I | S | V | E | - | M | L | R | - | - | - | E | D | I | K | D | Y | - | Y | T | D | L | S | I | L | Q | K | D | I | K | V | P | T | K | V | L | C | I | L | W | D | - | - | - | - | M | E | T | E | E | V | E | D | I | L | Q | E | F | - | - | - | - | - | - | - | - | - | - | - | V | N | K | S | L | L | F | C | D | R | N | G | K | S | - | - | - | - | - | - | - | - | - | F | R | Y | Y | L | H | D | L | Q | I | D | F | L | T | E | K | - | - | N | R | N | Q | - | L | - | Q | D | L | H | K | K | I | V | T | Q | F | Q | R | - | H | H | - | - | - | - | - | - | K | L | H | T | L | S | P | D | - | Q | E | - | - | - | - | - | - | - | D | C | M | Y | W | Y | S | F | L | A | Y | H | M | A | S | A | N | - | - | - | - | M | H | K | E | L | C | A | L | M | F | S | L | D | W | I | K | A | K | T | E | L | V | G | P | A | - | - | - | - | - | - | - | - | H | L | I | H | E | F | M | E | Y | R | H | I | L | D | T | K | E | I | S | V | C | S | Q | D | C | S | V | C | E | N | F | Q | - | - | - | E | F | L | S | L | N | G | H | L | L | G | R | Q | P | F | - | - | - | - | - | - | - | - | - | - | - | - | - | - | - | - | - | - | - | - | - | - | - | - | - | - | - | - | - | - | - | - | - | - | - | - | - | - | - | - | - | - | - | - | - | - | - | - | - | - | - | - | - | - | - | - | - | - | - | - | - | - | - | - | - | - | - | - | - | - | - | P | N | I | V | Q | L | G | L | C | E | P | E | T | S | E | V | Y | R | Q | A | K | L | Q | A | K | Q | E | V | D | N | G | - | - | - | - | - | - | - | - | - | - | - | - | - | - | - | - | - | - | - | - | - | - | - | - | - | - | - | - | - | M | L | Y | L | - | - | - | - | - | - | - | - | - | - | E | W | V | N | K | - | - | - | - | - | - | - | K | N | I | K | N | L | S | R | - | - | - | - | - | - | - | - | - | - | L | V | V | R | P | H | T | D | - | - | - | - | - | - | - | - | - | - | - | - | - | - | - | A | V | Y | H | A | C | - | - | - | F | S | - | E | D | G | Q | R | I | A | - | - | - | - | - | - | - | - | - | S | C | G | A | D | K | T | L | Q | V | F | K | - | - | - | - | - | A | E | T | G | E | K | L | L | E | I | - | K | A | H | E | D | E | - | - | - | V | L | C | C | A | F | S | - | T | - | - | - | - | - | - | - | - | - | - | - | - | - | D | D | R | F | I | A | T | C | A | V | D | K | K | V | K | I | - | - | - | W | N | S | V | - | - | - | - | - | - | - | - | - | - | - | - | - | - | - | - | - | - | - | - | - | - | - | - | - | - | - | - | - | - | - | - | - | - | - | - | - | - | - | - | - | - | - | - | - | - | - | - | - | - | - | - | - | - | - | - | - | - | - | - | - | - | - | - | - | - | - | - | - | - | - | - | - | - | - | - | - | - | - | - | - | - | - | - | - | - | - | - | - | - | - | - | - | - | - | - | - | - | - | - | - | - | - | - | - | - | - | - | - | - | - | - | - | - | - | - | - | - | - | - | - | - | - | - | - | - | - | - | - | - | - | - | - | - | - | - | - | - | - | - | T | G | E | L | V | H | T | Y | - | D | E | H | S | - | - | - | - | - | E | Q | V | N | - | - | - | - | - | - | - | - | - | - | - | - | - | C | C | H | - | - | - | - | - | - | - | - | - | - | - | - | - | F | T | N | N | S | - | - | - | - | - | - | - | - | - | - | - | - | - | - | - | - | H | H | - | L | L | L | A | T | G | S | S | - | - | - | - | D | C | F | L | - | - | - | - | - | - | - | - | - | - | - | - | K | L | W | D | L | - | - | - | N | Q | K | D | C | R | N | T | M | F | - | - | - | - | - | - | - | - | - | - | - | - | - | - | - | - | - | G | H | T | N | S | - | - | - | - | - | - | - | - | - | - | - | - | - | - | - | - | V | N | H | C | R | F | S | P | D | D | E | - | - | - | - | - | - | - | - | - | - | - | - | - | - | - | - | - | - | - | - | - | - | - | - | - | - | - | - | L | L | A | S | C | S | A | D | G | S | L | K | L | - | - | - | - | - | - | - | - | - | - | - | - | - | - | - | - | - | - | - | W | D | - | V | K | S | A | N | E | R | K | S | - | - | - | - | - | - | I | N | - | - | - | - | - | - | - | - | - | - | - | - | - | - | - | - | - | - | - | - | - | - | V | K | Q | F | - | - | - | - | - | - | - | - | F | L | N | S | E | E | P | Q | E | - | - | - | - | - | - | - | - | - | - | - | - | - | - | - | - | - | - | - | - | - | - | - | - | - | - | - | D | M | E | - | V | I | V | K | C | C | - | - | - | - | - | - | - | - | - | - | - | - | - | - | - | - | S | W | S | S | D | G | A | R | I | M | V | A | A | K | N | K | I | - | - | - | - | - | - | - | - | - | - | - | F | L | F | D | I | H | T | - | - | - | S | G | - | - | - | - | - | - | - | - | - | - | - | - | - | - | - | - | - | - | - | - | - | - | - | - | - | - | - | - | - | - | - | - | - | - | - | - | - | - | - | - | - | - | - | - | - | - | - | - | - | - | - | - | - | - | - | - | - | - | - | - | - | - | - | - | - | - | - | - | - | - | - | - | - | - | - | - | - | - | - | - | - | - | - | - | - | - | - | - | - | - | - | - | - | - | - | - | - | - | - | - | - | - | - | - | - | - | - | - | - | - | - | - | - | - | - | - | - | - | - | - | - | - | - | - | - | - | L | L | A | - | - | - | - | - | - | - | - | - | - | - | - | - | - | - | - | - | - | - | - | - | - | - | - | - | - | - | - | - | - | - | - | - | - | - | - | - | - | - | - | - | - | - | - | - | - | - | - | - | - | - | - | - | - | - | - | - | - | - | - | - | - | - | - | - | - | - | - | - | - | - | - | - | - | - | - | - | - | - | - | - | - | - | - | - | - | - | - | - | - | - | - | - | - | - | - | - | - | - | - | - | - | - | - | - | - | - | - | - | - | - | - | - | - | - | - | - | - | - | - | - | - | - | - | - | - | - | - | - | - | - | - | - | - | - | - | - | - | - | - | - | - | - | - | - | - | - | - | - | - | - | - | - | - | - | - | - | - | - | - | - | - | - | - | - | - | - | - | - | - | - | - | - | - | - | - | - | - | - | - | - | - | - | E | I | - | - | - | H | T | G | H | H | S | T | I | Q | Y | C | - | - | - | - | - | - | - | - | - | - | - | - | - | - | - | - | - | - | - | - | - | - | - | - | - | D | F | S | S | N | N | H | L | A | L | V | A | L | S | Q | - | - | - | - | - | - | Y | C | V | E | - | - | - | - | - | - | - | - | - | - | - | - | - | - | - | - | - | L | W | N | M | D | S | C | - | - | - | - | - | - | - | - | - | - | - | - | - | - | - | - | - | - | - | - | - | - | - | - | - | - | - | - | - | S | K | V | A | D | C | R | G | H | L | S | - | - | - | - | - | - | - | - | - | - | - | - | - | - | - | - | - | - | - | - | - | - | - | - | - | - | - | - | - | - | - | - | - | - | - | - | - | - | - | - | - | - | - | - | W | V | H | C | A | M | F | S | - | - | - | - | - | - | - | - | - | - | - | P | D | G | S | - | - | - | - | - | - | - | - | - | - | S | F | L | T | S | S | D | D | E | - | - | - | - | - | - | - | - | - | - | - | - | - | - | T | I | R | L | - | W | - | - | - | - | - | - | - | - | - | - | - | - | - | - | - | - | - | - | - | - | - | - | - | - | - | - | - | - | - | - | - | - | - | - | - | - | - | - | E | T | K | K | V | C | K | N | S | A | I | V | L | K | Q | E | - | - | - | - | - | - | - | - | - | - | - | - | - | - | - | - | I | D | V | V | F | - | - | - | - | - | - | - | - | - | - | - | - | - | - | - | - | - | - | - | - | - | - | - | - | - | - | - | - | - | - | - | - | - | - | - | - | - | - | - | - | - | - | - | - | - | - | - | - | - | - | - | - | - | - | Q | E | N | E | V | M | - | - | - | - | - | - | - | - | - | - | - | - | - | - | - | - | - | - | - | - | - | - | - | - | - | - | - | - | - | - | - | - | - | - | - | - | I | L | A | V | D | N | M | K | R | L | Q | L | - | I | N | G | K | T | - | - | - | - | - | - | - | - | - | - | G | Q | I | D | Y | L | T | E | A | - | - | - | - | - | - | - | - | - | - | - | - | - | - | - | - | - | - | - | - | - | - | Q | - | - | - | - | - | - | - | - | - | - | - | V | S | C | C | C | L | S | P | R | L | Q | - | - | - | - | - | - | - | - | - | Y | V | A | L | G | G | - | - | - | - | - | - | - | - | - | - | - | - | - | - | - | - | - | - | - | - | - | - | - | - | - | - | - | - | - | - | - | - | - | - | - | - | E | - | - | - | - | - | - | - | - | - | D | G | A | - | - | - | - | - | - | - | - | - | - | I | - | - | - | - | - | - | - | - | - | - | - | - | - | - | - | - | - | - | - | - | - | - | - | - | - | - | - | - | - | - | - | - | - | - | - | - | - | - | - | E | I | L | E | L | L | N | N | R | I | F | Q | S | R | I | G | - | - | - | - | - | H | - | - | - | - | - | - | - | - | - | - | - | - | K | S | T | V | R | H | I | Q | F | T | - | - | - | - | - | - | - | - | - | - | - | - | - | - | - | - | - | - | - | - | - | - | - | - | - | - | - | - | - | - | - | - | - | - | - | - | - | T | D | G | K | T | L | I | S | S | S | - | D | D | S | - | - | - | - | - | - | - | - | A | I | Q | - | V | - | - | - | - | - | - | - | - | - | - | - | - | - | - | - | - | - | - | - | - | - | - | - | - | - | - | - | - | - | - | - | - | - | - | - | - | - | - | - | W | N | - | - | - | - | - | - | - | - | - | - | W | Q | S | E | E | Y | V | F | - | - | - | - | - | - | - | - | - | - | - | - | - | - | - | - | - | - | - | - | - | - | - | - | - | - | - | - | - | - | - | - | - | - | - | - | - | - | - | - | - | - | - | - | - | - | - | - | - | - | L | Q | A | H | Q | - | - | - | - | - | - | - | - | - | - | - | - | - | - | - | - | - | - | - | - | - | - | - | - | - | - | E | T | V | K | D | F | R | L | L | - | - | - | - | - | - | - | - | - | - | - | - | - | - | - | - | - | - | - | - | - | - | - | K | N | - | S | R | L | L | - | S | W | - | - | - | S | F | D | G | T | V | - | - | - | - | - | - | - | - | - | - | - | - | - | - | - | - | - | - | - | - | - | - | - | - | - | - | - | - | - | - | - | - | - | - | - | K | V | W | N | I | I | T | G | R | I | - | - | - | - | - | E | K | D | F | V | C | - | - | - | - | - | - | - | H | Q | - | - | - | - | - | - | - | - | - | - | - | - | - | - | - | - | - | - | - | D | T | V | L | - | - | - | - | - | - | - | - | - | - | - | - | - | - | - | S | C | D | - | - | - | - | - | - | - | - | - | - | - | - | - | - | - | I | S | P | D | G | T | - | K | F | S | S | T | - | S | A | D | K | T | - | - | - | - | - | - | A | K | I | W | S | F | E | L | F | - | - | - | - | - | - | - | - | - | - | - | - | - | - | - | - | - | - | - | - | - | - | - | - | S | P | L | H | E | - | - | - | - | - | - | - | - | L | - | - | - | - | R | G | H | K | G | C | V | - | - | - | - | - | - | - | - | - | R | C | S | - | V | F | S | - | - | - | - | - | - | - | - | - | V | D | S | T | - | L | L | A | T | - | - | - | - | - | - | - | - | G | D | D | N | G | E | I | R | I | W | N | V | S | N | G | E | L | L | - | - | - | - | - | - | - | - | - | - | - | - | - | - | - | - | - | - | - | - | - | - | - | - | - | - | - | - | - | - | - | - | - | - | - | - | - | - | - | - | - | - | - | - | - | - | - | - | - | - | - | - | - | - | - | - | H | L | C | A | P | I | S | V | E | - | - | - | - | - | - | - | - | - | - | - | - | - | - | - | - | - | - | - | - | E | G | A | A | T | H | G | G | W | - | - | - | - | - | - | - | - | V | T | D | L | C | F | S | P | D | S | K | - | - | - | - | - | - | - | - | - | - | - | - | - | - | - | - | - | - | - | - | - | - | - | - | - | - | - | - | - | - | - | - | - | - | - | - | - | - | - | - | - | - | - | - | - | M | L | V | S | - | - | A | E | G | Y | L | K | - | - | - | - | - | - | - | - | W | W | N | V | V | T | - | - | - | - | - | - | - | - | - | - | - | - | - | - | - | - | - | - | - | - | - | - | - | - | - | - | - | - | - | - | - | - | - | - | - | - | - | - | - | - | - | - | - | - | - | - | - | - | - | - | - | - | - | - | - | - | - | - | - | - | - | - | - | - | - | - | - | - | - | - | - | - | - | - | - | - | - | - | - | - | - | - | - | - | - | - | - | - | - | - | - | - | - | - | - | - | - | - | - | - | - | - | - | - | - | - | - | - | - | - | - | - | - | - | - | - | - | - | - | - | - | - | - | - | - | - | - | - | - | - | - | - | - | - | - | - | - | - | - | - | - | - | - | - | - | - | - | - | - | - | - | - | - | - | - | - | - | - | - | - | - | - | - | - | - | - | - | - | - | - | - | - | - | - | - | - | - | - | - | - | - | - | - | - | - | - | - | - | - | - | - | - | - | - | - | - | - | - | - | - | - | - | - | - | - | - | - | - | - | - | - | - | - | - | - | - | - | - | - | - | - | - | - | - | - | - | - | - | - | - | - | - | - | - | - | - | - | - | - | - | - | - | - | - | - | - | - | - | - | - | - | - | - | - | - | - | - | - | - | - | - | - | - | - | - | - | - | G | E | S | S | Q | T | F | Y | T | - | - | - | - | - | N | G | - | - | - | T | N | L | K | K | I | - | - | - | H | V | S | P | - | - | - | - | D | F | K | T | - | - | - | - | - | - | - | - | - | - | - | - | - | - | - | - | - | - | - | - | - | - | - | - | - | - | Y | V | T | V | D | - | N | L | G | I | L | - | - | Y | I | L | Q | V | - | - | - | - | - | - | - | - | - | - | - | - | - | - | - | - | - | - | - | - | - | - | - | - | - | - | - | - | - | - | - | - | - | - | L | E | - | - | - | - | - | - | - | - | - | - | - | - | - | - | - | - | - | - | - | - | - | - | - | - | - | - | - | - | - | - | - | - | - | - | - | - | - | - | - | - | - | - | - | - | - | - | - | - | - | - | - | - | M | - | - | - | - | - | - | - | - | - | - | - | - | - | - | - | - | - | - | - | - | - | - | - | - | - | - | - | - | - | - | - | - | - | - | - | - | - | - | - | - | - | - | - | - | - | - | - | - | - | - | - | - | - | - | D | E | A | D | R | Q | - | - | - | - | - | - | - | - | - | - | - | - | - | - | - | - | - | - | - | - | L | L | R | R | - | - | - | - | - | - | - | - | - | - | - | - | - | - | - | - | - | - | - | - | - | - | - | - | - | - | - | - | - | - | C | R | - | - | - | - | - | - | - | - | V | R | L | - | V | G | E | L | Q | V | D | P | L | W | - | - | - | - | - | - | - | - | - | - | - | - | - | - | - | - | - | - | - | - | - | - | - | - | - | - | - | - | - | - | - | - | - | - | - | - | - | - | - | - | - | - | - | - | - | D | A | L | L | T | R | E | L | - | - | - | - | - | - | - | - | - | - | - | - | - | - | - | - | - | - | - | - | - | - | - | - | F | T | P | D | M | I | E | D | - | - | - | - | - | - | - | - | - | - | - | - | - | - | - | - | - | - | - | - | - | - | - | - | - | I | Q | R | A | G | - | - | - | - | - | - | - | - | S | G | S | R | R | D | Q | - | A | R | Q | L | I | I | D | L | E | T | R | G | - | S | Q | A | L | P | L | F | I | S | C | L | E | D | T | G | Q | D | T | L | A | S | F | L | R | T | S | S | H | A | A | - | - | - | - | K | W | N | P | - | E | A | V | - | - | - | - | - | - | - | - | - | - | - | - | - | - | - | - | - | - | - | - | - | - | - | - | - | - | R | T | L | N | - | - | - | - | - | - | - | - | - | - | - | - | - | - | - | L | M | P | V | - | - | - | - | - | - | - | - | - | - | - | - | - | - | - | - | - | - | - | - | - | - | - | - | - | - | - | - | - | - | - | - | - | - | - | - | - | - | - | V | L | G | P | - | - | - | - | - | E | L | L | R | - | E | V | K | Q | D | P | L | K | P | - | T | - | P | - | - | - | - | - | - | - | - | - | - | - | - | - | - | - | - | - | - | - | - | - | - | - | - | - | - | - | - | - | - | - | - | - | - | - | - | - | - | - | - | - | - | - | - | - | - | - | - | - | - | - | G | K | L | T | P | V | - | - | - | - | - | - | - | - | - | - | - | - | - | - | - | - | - | - | - | - | - | - | - | - | - | - | V | V | - | - | - | - | - | - | - | - | - | - | - | - | - | - | - | - | - | - | - | - | - | - | - | - | - | - | - | - | - | - | - | - | - | - | - | - | - | - | - | - | - | - | - | - | - | - | - | - | - | - | - | - | - | - | - | - | - | - | - | - | - | - | - | - | - | - | - | - | G | P | E | E | L | W | P | - | - | - | - | - | - | - | - | - | - | - | - | - | - | - | - | - | - | - | - | - | - | - | - | - | - | A | K | L | R | P | - | - | - | - | - | - | E | V | L | R | P | - | E | A | - | - | - | - | - | - | - | - | - | - | - | - | - | - | - | - | - | - | P | R | P | G | D | I | G | S | G | G | F | - | - | - | - | - | - | - | - | - | - | - | - | - | S | D | V | R | - | - | - | - | - | - | - | - | - | - | A | - | Q | E | R | W | R | G | N | A | N | - | - | - | - | - | L | A | Y | V | - | - | L | D | A | D | - | P | C | G | H | C | L | I | I | N | N | V | N | F | C | L | E | S | - | - | G | L | S | T | R | T | G | S | N | I | D | C | E | K | L | Q | R | R | F | - | R | L | L | H | F | V | V | D | V | K | G | D | L | T | A | K | Q | M | V | Q | A | L | T | E | L | A | R | R | D | H | - | G | A | L | D | C | C | V | V | V | I | L | S | H | G | C | Q | A | S | H | L | Q | L | P | G | A | - | V | Y | G | T | D | G | C | P | V | S | V | E | R | I | V | N | I | F | N | G | T | G | C | P | S | L | G | G | K | P | K | L | F | F | I | Q | A | C | - | - | G | G | D | Q | K | D | Y | G | F | E | V | V | S | T | - | - | - | S | P | E | D | - | R | T | P | G | S | D | - | - | - | - | - | - | - | S | E | P | D | A | A | P | F | Q | E | G | P | G | A | F | - | - | - | - | - | - | - | - | D | Q | P | D | A | V | - | - | - | - | - | - | - | S | T | L | - | - | - | P | T | A | S | D | I | F | V | S | Y | S | T | F | P | - | - | G | - | - | - | - | - | - | - | - | F | V | S | W | R | D | T | K | S | G | S | W | Y | V | E | T | L | D | G | V | F | E | Q | W | A | H | - | S | - | - | - | - | - | - | - | - | - | - | - | - | - | - | - | - | - | - | - | - | - | - | - | - | - | - | - | - | - | - | - | E | D | L | Q | T | L | L | - | - | - | - | - | L | R | - | - | - | - | - | - | - | - | - | - | - | - | V | A | N | A | V | - | - | - | - | - | S | V | K | - | - | - | - | - | - | - | G | I | Y | - | K | Q | S | - | - | P | G | - | - | - | - | - | - | - | - | - | - | - | - | - | - | - | C | L | N | F | - | - | - | - | - | - | - | - | - | L | R | K | K | L | F | F | K | T | - | - | - | - | - | - | - | - | - | - | - | - | - | - | - | - | - | - | - | - | - | - | - | - | - | - | - | - | - | - | - | - | - | - | - | - | - | - | M | - | - | - | - | - | - | - | - | - | - | - | - | - | - | - | - | - | - | - | - | - | - | - | - | - | - | - | - | - | - | - | - | - | - | - | - | - | - | - | - | - | - | - | - | - | - | - | - | - | - | - | - | G | D | V | E | K | G | K | K | I | F | V | Q | K | C | A | Q | C | H | T | V | E | K | G | G | K | H | K | T | G | P | N | L | H | G | L | F | G | R | K | T | G | Q | A | P | G | F | S | Y | T | D | A | N | K | N | K | G | I | T | W | K | E | E | T | L | M | E | Y | L | E | N | P | K | K | Y | I | P | G | T | K | M | I | F | A | G | I | K | K | K | A | E | R | E | D | L | I | A | Y | L | K | K | A | T | N | E | - | - | - | - | - | - |
| Chaetura\_pelagica | M | - | - | - | - | - | - | - | - | - | - | - | - | - | - | - | - | - | - | - | - | - | - | - | - | - | - | - | - | - | - | - | - | - | - | - | - | - | - | - | - | - | - | - | - | - | - | - | - | - | - | - | - | - | - | - | - | - | - | - | - | - | - | - | - | - | - | - | - | - | - | - | - | - | - | - | - | - | - | - | - | - | - | - | - | - | - | - | - | - | - | - | - | - | - | - | - | - | D | V | K | S | - | R | N | - | - | - | - | - | - | - | - | - | - | - | - | - | - | - | - | Y | L | L | - | - | - | - | - | - | - | - | - | - | - | - | - | - | - | - | - | - | - | - | - | - | - | - | - | - | - | - | - | - | - | - | - | - | - | - | - | - | - | - | - | - | - | - | - | - | - | - | - | - | - | - | - | - | - | - | - | - | - | - | - | - | - | - | - | - | - | - | - | - | - | - | - | - | - | - | - | - | - | - | - | - | - | - | - | - | - | - | - | - | - | - | - | - | - | - | - | - | - | - | - | - | - | - | - | - | - | - | - | - | - | - | - | - | - | - | - | - | - | - | - | M | N | R | Q | A | L | E | K | D | I | K | T | S | Y | - | I | M | D | R | M | I | S | D | E | V | L | T | L | Q | E | E | E | R | V | K | Q | Q | - | - | - | - | - | - | - | - | - | - | - | - | - | - | - | - | - | N | T | R | E | E | R | A | A | M | L | I | N | I | I | L | T | K | - | - | - | - | D | N | N | S | Y | R | S | - | - | - | - | - | F | Y | N | A | L | - | L | H | E | G | - | - | - | - | - | - | Y | R | E | L | A | A | L | L | Q | D | G | I | P | A | I | S | S | G | N | R | - | K | - | - | - | - | - | - | - | - | - | - | - | - | - | - | - | - | - | - | - | - | - | - | S | S | V | D | G | M | T | - | - | - | - | - | - | - | - | - | - | - | S | Y | V | K | M | I | L | C | E | G | G | - | - | - | - | - | - | - | V | P | Q | R | P | V | V | F | V | T | R | P | K | L | V | D | A | I | - | - | - | - | - | - | - | - | - | - | K | K | K | L | C | - | - | - | - | - | - | - | - | - | - | - | - | - | - | - | - | - | - | - | - | - | - | - | - | - | - | - | - | - | - | - | - | C | L | G | S | E | P | - | - | - | - | - | - | - | - | - | - | - | - | - | - | - | - | - | - | - | - | - | - | - | - | - | - | - | - | - | - | - | - | - | - | - | - | - | - | A | W | V | T | V | Y | G | M | A | G | C | G | K | T | V | L | T | A | E | A | L | - | R | D | H | Q | L | L | E | D | Y | F | P | G | G | V | H | W | V | S | V | G | - | - | - | - | - | - | - | K | - | - | - | - | - | - | Q | D | K | A | - | - | - | - | - | - | - | - | - | - | - | - | G | L | L | I | K | L | Q | N | L | C | S | R | L | E | - | - | - | - | H | - | - | - | - | - | - | - | - | - | - | - | - | - | - | - | D | S | A | L | S | Q | R | P | P | L | N | I | - | - | - | - | - | - | - | - | - | - | - | - | - | - | - | - | E | E | A | K | D | R | L | R | L | L | M | L | R | - | - | - | - | - | - | - | - | - | - | - | - | K | Y | P | R | S | L | L | - | - | I | L | D | D | V | W | D | S | W | V | L | K | A | F | D | - | N | Q | C | Q | V | L | I | T | S | R | D | R | S | V | T | D | A | V | - | A | G | K | - | - | K | Y | E | V | H | V | E | S | - | - | - | G | L | A | H | E | K | G | L | E | I | L | S | L | F | V | N | M | - | K | I | S | E | L | P | E | - | - | - | - | - | - | - | - | - | - | - | - | - | - | Q | A | N | F | L | V | R | E | C | K | - | - | - | - | - | - | - | - | - | - | - | - | - | - | - | - | - | - | - | G | - | S | P | L | V | I | S | L | I | G | A | L | L | R | D | F | P | N | - | - | - | - | - | - | - | - | - | - | R | W | E | Y | Y | L | - | K | Q | L | Q | N | K | Q | F | K | R | I | - | - | - | - | - | - | - | - | - | - | - | - | - | - | - | - | - | - | - | - | - | - | R | K | S | S | S | Y | D | - | - | - | - | Y | E | A | L | D | E | A | M | S | I | S | V | E | - | Q | L | N | - | - | - | D | N | C | K | D | Y | - | Y | K | D | L | S | I | L | P | K | D | V | R | V | P | T | K | V | L | C | I | L | W | D | - | - | - | - | M | E | T | E | E | V | E | D | I | L | Q | E | F | - | - | - | - | - | - | - | - | - | - | - | V | N | K | S | L | L | F | C | D | R | N | G | N | S | - | - | - | - | - | - | - | - | - | F | Q | Y | Y | L | H | D | L | Q | L | D | F | L | T | E | K | - | - | N | R | N | Q | - | L | - | Q | E | L | H | K | N | I | V | N | Q | Y | K | K | - | Y | Y | - | - | - | - | - | - | K | L | N | T | P | V | P | S | - | Q | E | - | - | - | - | - | - | - | D | C | M | Y | W | Y | N | F | L | A | Y | H | M | A | G | A | N | - | - | - | - | M | Q | K | E | L | H | D | L | M | F | S | L | D | W | I | K | A | K | T | E | L | V | G | P | A | - | - | - | - | - | - | - | - | H | L | I | H | E | Y | V | E | Y | S | S | I | L | D | Q | K | - | - | - | - | - | - | - | G | S | T | V | R | E | N | F | Q | - | - | - | E | F | L | S | L | N | G | H | L | L | G | R | Q | P | F | - | - | - | - | - | - | - | - | - | - | - | - | - | - | - | - | - | - | - | - | - | - | - | - | - | - | - | - | - | - | - | - | - | - | - | - | - | - | - | - | - | - | - | - | - | - | - | - | - | - | - | - | - | - | - | - | - | - | - | - | - | - | - | - | - | - | - | - | - | - | - | P | D | I | I | Q | L | G | L | C | Q | P | E | T | S | E | V | Y | Q | Q | A | K | L | H | A | Q | R | E | - | - | T | G | - | - | - | - | - | - | - | - | - | - | - | - | - | - | - | - | - | - | - | - | - | - | - | - | - | - | - | - | - | A | F | Y | L | - | - | - | - | - | - | - | - | - | - | E | W | I | N | K | - | - | - | - | - | - | - | K | S | L | K | N | L | Y | R | - | - | - | - | - | - | - | - | - | - | L | V | V | R | P | H | R | D | - | - | - | - | - | - | - | - | - | - | - | - | - | - | - | A | V | Y | H | A | C | - | - | - | F | S | - | K | D | R | K | R | I | A | - | - | - | - | - | - | - | - | - | S | C | G | A | D | K | T | L | Q | V | F | K | - | - | - | - | - | A | E | S | G | E | R | L | L | E | I | - | S | A | H | D | D | E | - | - | - | I | F | C | C | A | F | S | - | A | - | - | - | - | - | - | - | - | - | - | - | - | - | D | G | E | F | V | A | T | C | S | A | D | K | K | V | K | I | - | - | - | W | N | S | R | - | - | - | - | - | - | - | - | - | - | - | - | - | - | - | - | - | - | - | - | - | - | - | - | - | - | - | - | - | - | - | - | - | - | - | - | - | - | - | - | - | - | - | - | - | - | - | - | - | - | - | - | - | - | - | - | - | - | - | - | - | - | - | - | - | - | - | - | - | - | - | - | - | - | - | - | - | - | - | - | - | - | - | - | - | - | - | - | - | - | - | - | - | - | - | - | - | - | - | - | - | - | - | - | - | - | - | - | - | - | - | - | - | - | - | - | - | - | - | - | - | - | - | - | - | - | - | - | - | - | - | - | - | - | - | - | - | - | - | - | T | G | Q | C | R | C | V | Y | - | E | E | H | S | - | - | - | - | - | E | Q | V | N | - | - | - | - | - | - | - | - | - | - | - | - | - | C | C | Q | - | - | - | - | - | - | - | - | - | - | - | - | - | F | N | N | R | G | - | - | - | - | - | - | - | - | - | - | - | - | - | - | - | - | G | Q | - | Y | L | L | A | T | C | S | S | - | - | - | - | D | T | F | I | - | - | - | - | - | - | - | - | - | - | - | - | K | L | W | D | L | - | - | - | N | K | K | Y | C | R | N | T | M | I | - | - | - | - | - | - | - | - | - | - | - | - | - | - | - | - | - | G | H | V | N | S | - | - | - | - | - | - | - | - | - | - | - | - | - | - | - | - | V | I | H | C | R | F | S | P | N | D | E | - | - | - | - | - | - | - | - | - | - | - | - | - | - | - | - | - | - | - | - | - | - | - | - | - | - | - | - | Y | V | A | S | C | S | T | D | G | T | V | K | L | - | - | - | - | - | - | - | - | - | - | - | - | - | - | - | - | - | - | - | W | E | - | A | R | S | G | N | E | L | K | S | - | - | - | - | - | - | I | E | - | - | - | - | - | - | - | - | - | - | - | - | - | - | - | - | - | - | - | - | - | - | I | K | D | F | - | - | - | - | - | - | - | - | F | K | N | A | D | E | Q | P | D | - | - | - | - | - | - | - | - | - | - | - | - | - | - | - | - | - | - | - | - | - | - | - | - | - | - | - | D | V | E | - | A | L | V | K | C | C | - | - | - | - | - | - | - | - | - | - | - | - | - | - | - | - | S | W | S | R | N | G | E | M | I | L | V | A | A | K | N | K | L | - | - | - | - | - | - | - | - | - | - | - | L | L | F | D | V | K | T | - | - | - | C | D | - | - | - | - | - | - | - | - | - | - | - | - | - | - | - | - | - | - | - | - | - | - | - | - | - | - | - | - | - | - | - | - | - | - | - | - | - | - | - | - | - | - | - | - | - | - | - | - | - | - | - | - | - | - | - | - | - | - | - | - | - | - | - | - | - | - | - | - | - | - | - | - | - | - | - | - | - | - | - | - | - | - | - | - | - | - | - | - | - | - | - | - | - | - | - | - | - | - | - | - | - | - | - | - | - | - | - | - | - | - | - | - | - | - | - | - | - | - | - | - | - | - | - | - | - | - | L | L | T | - | - | - | - | - | - | - | - | - | - | - | - | - | - | - | - | - | - | - | - | - | - | - | - | - | - | - | - | - | - | - | - | - | - | - | - | - | - | - | - | - | - | - | - | - | - | - | - | - | - | - | - | - | - | - | - | - | - | - | - | - | - | - | - | - | - | - | - | - | - | - | - | - | - | - | - | - | - | - | - | - | - | - | - | - | - | - | - | - | - | - | - | - | - | - | - | - | - | - | - | - | - | - | - | - | - | - | - | - | - | - | - | - | - | - | - | - | - | - | - | - | - | - | - | - | - | - | - | - | - | - | - | - | - | - | - | - | - | - | - | - | - | - | - | - | - | - | - | - | - | - | - | - | - | - | - | - | - | - | - | - | - | - | - | - | - | - | - | - | - | - | - | - | - | - | - | - | - | - | - | - | - | - | Q | V | - | - | - | I | V | S | H | H | S | T | I | Q | Y | C | - | - | - | - | - | - | - | - | - | - | - | - | - | - | - | - | - | - | - | - | - | - | - | - | - | D | F | C | P | G | D | E | L | V | A | V | A | L | S | H | - | - | - | - | - | - | C | S | V | E | - | - | - | - | - | - | - | - | - | - | - | - | - | - | - | - | - | L | W | N | I | K | S | L | - | - | - | - | - | - | - | - | - | - | - | - | - | - | - | - | - | - | - | - | - | - | - | - | - | - | - | - | - | S | K | V | A | Y | C | R | G | H | M | S | - | - | - | - | - | - | - | - | - | - | - | - | - | - | - | - | - | - | - | - | - | - | - | - | - | - | - | - | - | - | - | - | - | - | - | - | - | - | - | - | - | - | - | - | W | V | H | C | V | T | F | S | - | - | - | - | - | - | - | - | - | - | - | P | D | G | S | - | - | - | - | - | - | - | - | - | - | L | F | L | T | S | S | D | D | Q | - | - | - | - | - | - | - | - | - | - | - | - | - | - | T | I | R | I | - | W | - | - | - | - | - | - | - | - | - | - | - | - | - | - | - | - | - | - | - | - | - | - | - | - | - | - | - | - | - | - | - | - | - | - | - | - | - | - | E | T | K | K | V | C | K | S | S | D | A | V | L | K | S | E | - | - | - | - | - | - | - | - | - | - | - | - | - | - | - | - | L | D | V | V | F | - | - | - | - | - | - | - | - | - | - | - | - | - | - | - | - | - | - | - | - | - | - | - | - | - | - | - | - | - | - | - | - | - | - | - | - | - | - | - | - | - | - | - | - | - | - | - | - | - | - | - | - | - | - | H | N | D | E | V | M | - | - | - | - | - | - | - | - | - | - | - | - | - | - | - | - | - | - | - | - | - | - | - | - | - | - | - | - | - | - | - | - | - | - | - | - | I | L | A | I | Y | N | Q | K | H | L | Q | L | - | I | N | G | N | T | - | - | - | - | - | - | - | - | - | - | D | S | V | V | V | Q | T | A | A | - | - | - | - | - | - | - | - | - | - | - | - | - | - | - | - | - | - | - | - | - | - | Q | E | S | - | - | - | - | - | - | - | - | P | I | S | C | C | C | L | S | G | D | L | R | - | - | - | - | - | - | - | - | - | F | A | A | F | G | Q | - | - | - | - | - | - | - | - | - | - | - | - | - | - | - | - | - | - | - | - | - | - | - | - | - | - | - | - | - | - | - | - | - | - | - | - | E | - | - | - | - | - | - | - | - | - | S | G | T | - | - | - | - | - | - | - | - | - | - | I | - | - | - | - | - | - | - | - | - | - | - | - | - | - | - | - | - | - | - | - | - | - | - | - | - | - | - | - | - | - | - | - | - | - | - | - | - | - | - | K | V | L | Q | L | L | D | G | K | I | L | N | S | W | E | A | - | - | - | - | - | Y | - | - | - | - | - | - | - | - | - | - | - | - | R | T | S | V | Q | H | C | R | F | T | - | - | - | - | - | - | - | - | - | - | - | - | - | - | - | - | - | - | - | - | - | - | - | - | - | - | - | - | - | - | - | - | - | - | - | - | - | S | D | C | Q | T | L | I | S | S | A | - | H | D | S | - | - | - | - | - | - | - | - | T | I | Q | - | V | - | - | - | - | - | - | - | - | - | - | - | - | - | - | - | - | - | - | - | - | - | - | - | - | - | - | - | - | - | - | - | - | - | - | - | - | - | - | - | W | N | - | - | - | - | - | - | - | - | - | - | W | Q | L | N | E | C | V | F | - | - | - | - | - | - | - | - | - | - | - | - | - | - | - | - | - | - | - | - | - | - | - | - | - | - | - | - | - | - | - | - | - | - | - | - | - | - | - | - | - | - | - | - | - | - | - | - | - | - | L | R | G | H | K | - | - | - | - | - | - | - | - | - | - | - | - | - | - | - | - | - | - | - | - | - | - | - | - | - | - | E | A | I | K | D | F | R | L | L | - | - | - | - | - | - | - | - | - | - | - | - | - | - | - | - | - | - | - | - | - | - | - | E | N | - | S | K | L | L | - | S | W | - | - | - | S | F | D | G | T | V | - | - | - | - | - | - | - | - | - | - | - | - | - | - | - | - | - | - | - | - | - | - | - | - | - | - | - | - | - | - | - | - | - | - | - | K | V | W | N | I | T | T | G | N | T | - | - | - | - | - | E | K | E | F | A | C | - | - | - | - | - | - | - | H | G | - | - | - | - | - | - | - | - | - | - | - | - | - | - | - | - | - | - | - | G | A | V | L | - | - | - | - | - | - | - | - | - | - | - | - | - | - | - | S | C | A | - | - | - | - | - | - | - | - | - | - | - | - | - | - | - | V | S | P | D | N | S | - | T | F | S | S | T | - | S | A | D | K | T | - | - | - | - | - | - | A | K | I | W | S | F | E | S | S | - | - | - | - | - | - | - | - | - | - | - | - | - | - | - | - | - | - | - | - | - | - | - | - | S | V | L | H | E | - | - | - | - | - | - | - | - | L | - | - | - | - | K | G | H | E | A | C | V | - | - | - | - | - | - | - | - | - | R | C | C | - | T | F | S | - | - | - | - | - | - | - | - | - | P | N | N | K | - | L | L | A | T | - | - | - | - | - | - | - | - | G | D | D | K | G | E | I | R | I | W | D | T | L | T | G | A | L | L | - | - | - | - | - | - | - | - | - | - | - | - | - | - | - | - | - | - | - | - | - | - | - | - | - | - | - | - | - | - | - | - | - | - | - | - | - | - | - | - | - | - | - | - | - | - | - | - | - | - | - | - | - | - | - | - | H | F | C | S | P | V | T | V | D | - | - | - | - | - | - | - | - | - | - | - | - | - | - | - | - | - | - | - | - | E | G | E | P | A | H | G | G | W | - | - | - | - | - | - | - | - | V | T | D | L | S | F | S | P | D | S | K | - | - | - | - | - | - | - | - | - | - | - | - | - | - | - | - | - | - | - | - | - | - | - | - | - | - | - | - | - | - | - | - | - | - | - | - | - | - | - | - | - | - | - | - | - | M | L | V | S | - | - | S | G | G | Y | L | K | - | - | - | - | - | - | - | - | W | W | N | V | T | T | - | - | - | - | - | - | - | - | - | - | - | - | - | - | - | - | - | - | - | - | - | - | - | - | - | - | - | - | - | - | - | - | - | - | - | - | - | - | - | - | - | - | - | - | - | - | - | - | - | - | - | - | - | - | - | - | - | - | - | - | - | - | - | - | - | - | - | - | - | - | - | - | - | - | - | - | - | - | - | - | - | - | - | - | - | - | - | - | - | - | - | - | - | - | - | - | - | - | - | - | - | - | - | - | - | - | - | - | - | - | - | - | - | - | - | - | - | - | - | - | - | - | - | - | - | - | - | - | - | - | - | - | - | - | - | - | - | - | - | - | - | - | - | - | - | - | - | - | - | - | - | - | - | - | - | - | - | - | - | - | - | - | - | - | - | - | - | - | - | - | - | - | - | - | - | - | - | - | - | - | - | - | - | - | - | - | - | - | - | - | - | - | - | - | - | - | - | - | - | - | - | - | - | - | - | - | - | - | - | - | - | - | - | - | - | - | - | - | - | - | - | - | - | - | - | - | - | - | - | - | - | - | - | - | - | - | - | - | - | - | - | - | - | - | - | - | - | - | - | - | - | - | - | - | - | - | - | - | - | - | - | - | - | - | - | - | - | G | E | S | L | Q | T | F | Y | T | - | - | - | - | - | N | G | - | - | - | T | N | L | K | S | I | - | - | - | H | V | S | P | - | - | - | - | N | F | K | V | - | - | - | - | - | - | - | - | - | - | - | - | - | - | - | - | - | - | - | - | - | - | - | - | - | - | Y | V | T | V | D | - | N | L | G | I | L | - | - | Y | V | L | Q | K | - | - | - | - | - | - | - | - | - | - | - | - | - | - | - | - | - | - | - | - | - | - | - | - | - | - | - | - | - | - | - | - | - | - | L | - | - | - | - | - | - | - | - | - | - | - | - | - | - | - | - | - | - | - | - | - | - | - | - | - | - | - | - | - | - | - | - | - | - | - | - | - | - | - | - | - | - | - | - | - | - | - | - | - | - | - | - | - | M | - | - | - | - | - | - | - | - | - | - | - | - | - | - | - | - | - | - | - | - | - | - | - | - | - | - | - | - | - | - | - | - | - | - | - | - | - | - | - | - | - | - | - | - | - | - | - | - | - | - | - | - | - | - | - | - | - | - | - | - | - | - | - | - | - | - | - | - | - | - | - | - | - | - | - | - | - | - | - | - | - | - | - | - | - | - | - | - | - | - | - | - | - | - | - | - | - | - | - | - | - | - | - | - | - | - | - | - | - | - | - | - | - | - | S | R | - | - | - | - | - | - | - | - | - | - | - | - | - | - | - | - | - | - | - | - | - | - | - | - | - | - | - | - | - | - | - | - | - | - | - | - | - | - | - | - | - | - | - | - | - | - | - | - | - | - | - | - | - | - | - | - | - | - | - | - | - | - | - | - | - | - | - | - | - | - | - | - | - | - | - | - | - | - | - | - | - | - | - | - | - | - | - | - | - | - | - | - | - | - | - | - | - | - | - | R | S | P | S | A | Q | L | Q | L | - | - | - | - | - | - | - | - | - | - | - | - | - | - | - | - | - | - | - | - | - | - | - | P | A | E | S | A | - | - | - | - | - | - | - | - | - | - | P | S | R | G | E | Q | - | A | R | Q | L | L | I | D | L | E | T | R | G | - | N | Q | A | F | P | I | F | L | S | I | L | R | D | T | G | Q | G | E | L | A | D | L | L | L | E | E | C | E | L | P | - | - | - | - | L | V | P | P | - | E | - | - | - | - | - | - | - | - | - | - | - | - | - | - | - | - | - | - | - | - | - | - | - | - | - | - | - | - | - | P | R | D | - | - | - | - | - | - | - | - | - | - | - | - | - | - | - | L | R | P | V | - | - | - | - | - | - | - | - | - | - | - | - | - | - | - | - | - | - | - | - | - | - | - | - | - | - | - | - | - | - | - | - | - | - | - | - | - | - | - | P | L | D | L | P | G | E | K | H | D | R | G | G | - | - | - | - | - | - | - | - | - | - | - | T | - | - | - | - | - | - | - | - | - | - | - | - | - | - | - | - | - | - | - | - | - | - | - | - | - | - | - | - | - | - | - | - | - | - | - | - | - | - | - | - | - | - | - | - | - | - | - | - | - | - | - | - | - | - | - | - | - | - | - | - | - | - | - | - | - | - | - | - | - | - | - | - | - | - | - | - | - | - | - | - | - | - | - | - | - | - | - | - | - | - | - | - | - | - | - | - | - | - | - | - | - | - | - | - | - | - | - | - | - | - | - | - | - | - | - | - | - | - | - | - | - | - | - | - | - | - | - | - | - | - | - | - | - | - | - | - | - | - | - | - | - | - | - | - | - | - | - | - | - | - | - | - | - | - | - | - | F | P | E | R | V | S | - | - | - | - | - | - | - | - | - | - | - | - | - | - | - | - | - | - | - | - | - | - | - | - | - | - | I | P | V | Q | A | - | - | - | - | - | - | E | S | E | R | P | - | P | R | - | - | - | - | - | - | - | - | - | - | - | - | - | - | - | - | - | - | - | - | - | - | T | Q | G | - | - | - | - | - | - | - | - | - | - | - | - | - | - | - | - | - | - | - | - | L | - | - | - | - | - | - | - | - | - | - | A | - | V | D | - | - | K | R | N | E | D | - | - | - | - | - | L | V | Y | Q | - | - | L | R | A | N | - | P | C | G | Y | C | L | I | L | N | N | V | T | F | S | R | A | S | - | - | G | L | S | R | R | D | G | S | D | L | D | S | Q | K | L | E | K | R | F | - | R | S | L | G | F | H | I | L | P | R | R | D | L | T | A | Q | E | M | V | S | E | L | Q | K | L | A | Q | Q | D | H | - | T | T | L | D | C | C | V | V | V | I | L | S | H | G | C | Q | T | S | H | I | Q | F | P | G | G | - | V | F | G | T | D | G | K | P | I | P | I | E | K | I | V | N | Y | F | N | G | S | H | C | P | S | L | R | G | K | P | K | L | F | F | I | Q | A | C | - | - | G | G | E | Q | R | D | R | G | F | V | V | E | S | D | - | - | - | S | P | G | D | - | E | A | P | G | G | S | - | - | - | - | - | - | - | S | E | S | D | A | T | P | F | Q | P | P | P | G | N | V | - | - | - | - | - | - | - | - | D | E | P | D | A | V | - | - | - | - | - | - | - | A | S | L | - | - | - | P | T | P | S | D | I | L | V | S | Y | S | T | F | P | - | - | G | - | - | - | - | - | - | - | - | F | V | S | W | R | E | A | S | S | G | S | W | Y | V | E | T | L | D | S | V | L | E | Q | Y | A | P | - | S | - | - | - | - | - | - | - | - | - | - | - | - | - | - | - | - | - | - | - | - | - | - | - | - | - | - | - | - | - | - | - | E | D | L | L | T | L | S | - | - | - | - | - | S | Q | - | - | - | - | - | - | - | - | - | - | - | - | V | A | H | A | V | - | - | - | - | - | S | A | K | - | - | - | - | - | - | - | G | K | Y | - | K | Q | I | - | - | P | G | - | - | - | - | - | - | - | - | - | - | - | - | - | - | - | C | F | N | F | - | - | - | - | - | - | - | - | - | L | R | K | K | F | F | F | R | C | K | - | - | - | - | - | - | - | - | - | - | - | - | - | - | - | - | - | - | - | - | - | - | - | - | - | - | - | - | - | - | - | - | - | - | - | - | - | M | - | - | - | - | - | - | - | - | - | - | - | - | - | - | - | - | - | - | - | - | - | - | - | - | - | - | - | - | - | - | - | - | - | - | - | - | - | - | - | - | - | - | - | - | - | - | - | - | - | - | - | - | G | D | I | E | K | G | K | K | I | F | V | Q | K | C | S | Q | C | H | T | V | E | K | G | G | K | H | K | T | G | P | N | L | N | G | L | F | G | R | K | T | G | Q | A | E | G | F | S | Y | T | D | A | N | K | N | K | G | I | T | W | G | E | D | T | L | M | E | Y | L | E | N | P | K | K | Y | I | P | G | T | K | M | I | F | A | G | I | K | K | K | T | E | R | V | D | L | I | A | Y | L | K | D | A | T | A | K | - | - | - | - | - | - |
| Charadrius\_vociferus | M | - | - | - | - | - | - | - | - | - | - | - | - | - | - | - | - | - | - | - | - | - | - | - | - | - | - | - | - | - | - | - | - | - | - | - | - | - | - | - | - | - | - | - | - | - | - | - | - | - | - | - | - | - | - | - | - | - | - | - | - | - | - | - | - | - | - | - | - | - | - | - | - | - | - | - | - | - | - | - | - | - | - | - | - | - | - | - | - | - | - | - | - | - | - | - | - | - | D | A | K | S | - | R | N | - | - | - | - | - | - | - | - | - | - | - | - | - | - | - | - | Y | L | L | - | - | - | - | - | - | - | - | - | - | - | - | - | - | - | - | - | - | - | - | - | - | - | - | - | - | - | - | - | - | - | - | - | - | - | - | - | - | - | - | - | - | - | - | - | - | - | - | - | - | - | - | - | - | - | - | - | - | - | - | - | - | - | - | - | - | - | - | - | - | - | - | - | - | - | - | - | - | - | - | - | - | - | - | - | - | - | - | - | - | - | - | - | - | - | - | - | - | - | - | - | - | - | - | - | - | - | - | - | - | - | - | - | - | - | - | - | - | - | - | - | R | N | R | Q | A | L | E | K | D | I | K | T | S | Y | - | I | M | D | R | M | I | S | D | E | V | L | T | L | Q | E | E | E | K | V | K | Q | Q | - | - | - | - | - | - | - | - | - | - | - | - | - | - | - | - | - | N | T | Q | K | E | R | A | A | M | L | I | N | I | I | L | T | K | - | - | - | - | D | N | N | S | Y | R | S | - | - | - | - | - | F | Y | N | A | L | - | L | H | E | G | - | - | - | - | - | - | Y | R | D | L | A | A | L | L | Q | D | G | I | P | A | I | S | S | G | N | G | - | K | - | - | - | - | - | - | - | - | - | - | - | - | - | - | - | - | - | - | - | - | - | - | S | S | M | D | G | M | T | - | - | - | - | - | - | - | - | - | - | - | S | Y | V | K | T | I | L | C | E | G | G | - | - | - | - | - | - | - | V | P | Q | R | P | V | V | F | V | T | R | P | K | L | V | D | A | I | - | - | - | - | - | - | - | - | - | - | K | Q | K | L | C | - | - | - | - | - | - | - | - | - | - | - | - | - | - | - | - | - | - | - | - | - | - | - | - | - | - | - | - | - | - | - | - | C | L | G | S | D | P | - | - | - | - | - | - | - | - | - | - | - | - | - | - | - | - | - | - | - | - | - | - | - | - | - | - | - | - | - | - | - | - | - | - | - | - | - | - | G | W | V | T | V | Y | G | M | A | G | C | G | K | T | V | L | T | A | E | A | L | - | R | D | H | Q | L | L | E | D | Y | F | P | G | G | V | H | W | I | S | V | G | - | - | - | - | - | - | - | K | - | - | - | - | - | - | Q | D | K | A | - | - | - | - | - | - | - | - | - | - | - | - | G | L | L | I | K | L | Q | N | L | C | S | R | L | E | - | - | - | - | H | - | - | - | - | - | - | - | - | - | - | - | - | - | - | - | D | S | T | L | S | Q | R | P | P | L | N | I | - | - | - | - | - | - | - | - | - | - | - | - | - | - | - | - | E | E | A | K | D | R | L | R | L | L | M | L | R | - | - | - | - | - | - | - | - | - | - | - | - | K | Y | P | R | S | L | L | - | - | I | L | D | D | I | W | D | S | W | V | L | K | A | F | D | - | N | Q | C | Q | V | L | I | T | S | R | D | R | S | V | T | D | A | V | - | A | G | N | - | - | K | Y | E | V | H | V | E | S | - | - | - | G | L | A | H | E | K | G | L | E | I | L | S | L | F | V | N | M | - | K | I | S | E | L | P | E | - | - | - | - | - | - | - | - | - | - | - | - | - | - | Q | A | N | C | L | V | R | E | C | K | - | - | - | - | - | - | - | - | - | - | - | - | - | - | - | - | - | - | - | G | - | S | P | L | V | I | S | L | I | G | A | L | L | R | D | F | P | S | - | - | - | - | - | - | - | - | - | - | R | W | E | Y | Y | L | - | K | Q | L | Q | H | K | Q | F | K | R | I | - | - | - | - | - | - | - | - | - | - | - | - | - | - | - | - | - | - | - | - | - | - | R | K | S | S | S | Y | D | - | - | - | - | Y | E | A | L | D | E | A | M | S | I | S | V | E | - | Q | L | N | - | - | - | D | N | Y | K | D | Y | - | Y | K | D | L | S | I | L | P | K | D | V | K | V | P | T | K | V | L | C | I | L | W | D | - | - | - | - | M | E | T | E | E | V | E | D | I | L | Q | E | F | - | - | - | - | - | - | - | - | - | - | - | V | N | K | S | L | L | F | C | D | R | N | G | K | S | - | - | - | - | - | - | - | - | - | F | H | Y | Y | L | H | D | L | Q | L | D | F | L | T | E | K | - | - | N | R | N | Q | - | L | - | Q | E | L | H | K | N | I | V | N | Q | Y | K | K | - | Y | Y | - | - | - | - | - | - | K | L | N | T | P | V | P | S | - | Q | E | - | - | - | - | - | - | - | D | C | M | Y | W | Y | N | F | L | A | Y | H | M | A | G | A | N | - | - | - | - | M | Q | K | E | L | H | D | L | M | F | S | L | D | W | I | K | A | K | T | E | L | V | G | P | A | - | - | - | - | - | - | - | - | H | L | I | H | E | Y | V | E | Y | S | S | I | L | D | Q | K | - | - | - | - | - | - | - | D | S | T | V | R | E | N | F | Q | - | - | - | E | F | L | S | L | N | G | H | L | L | G | R | Q | P | F | - | - | - | - | - | - | - | - | - | - | - | - | - | - | - | - | - | - | - | - | - | - | - | - | - | - | - | - | - | - | - | - | - | - | - | - | - | - | - | - | - | - | - | - | - | - | - | - | - | - | - | - | - | - | - | - | - | - | - | - | - | - | - | - | - | - | - | - | - | - | - | P | D | I | V | Q | L | G | L | C | Q | P | E | T | - | - | - | - | - | - | - | - | - | - | - | - | - | - | - | - | - | - | - | - | - | - | - | - | - | - | - | - | - | - | - | - | - | - | - | - | - | - | - | - | - | - | - | - | - | - | - | - | - | - | - | - | - | - | - | - | - | - | - | - | - | - | - | - | N | K | - | - | - | - | - | - | - | K | S | L | K | N | L | Y | R | - | - | - | - | - | - | - | - | - | - | L | V | V | R | P | H | R | D | - | - | - | - | - | - | - | - | - | - | - | - | - | - | - | A | V | Y | H | A | C | - | - | - | F | S | - | K | D | R | Q | R | I | A | - | - | - | - | - | - | - | - | - | S | C | G | A | D | K | T | L | Q | V | F | K | - | - | - | - | - | A | E | S | G | E | R | L | L | E | I | - | S | A | H | D | D | E | - | - | - | I | L | C | C | A | F | S | - | A | - | - | - | - | - | - | - | - | - | - | - | - | - | D | G | E | F | V | A | T | C | S | A | D | K | K | V | K | I | - | - | - | W | N | S | R | - | - | - | - | - | - | - | - | - | - | - | - | - | - | - | - | - | - | - | - | - | - | - | - | - | - | - | - | - | - | - | - | - | - | - | - | - | - | - | - | - | - | - | - | - | - | - | - | - | - | - | - | - | - | - | - | - | - | - | - | - | - | - | - | - | - | - | - | - | - | - | - | - | - | - | - | - | - | - | - | - | - | - | - | - | - | - | - | - | - | - | - | - | - | - | - | - | - | - | - | - | - | - | - | - | - | - | - | - | - | - | - | - | - | - | - | - | - | - | - | - | - | - | - | - | - | - | - | - | - | - | - | - | - | - | - | - | - | - | - | T | G | Q | C | K | C | V | Y | - | E | E | H | A | - | - | - | - | - | E | Q | V | N | - | - | - | - | - | - | - | - | - | - | - | - | - | C | C | Q | - | - | - | - | - | - | - | - | - | - | - | - | - | F | N | N | G | T | - | - | - | - | - | - | - | - | - | - | - | - | - | - | - | - | G | Q | - | Y | L | L | A | T | C | S | N | - | - | - | - | D | T | F | I | - | - | - | - | - | - | - | - | - | - | - | - | K | L | W | D | L | - | - | - | N | K | K | Y | C | R | N | T | M | I | - | - | - | - | - | - | - | - | - | - | - | - | - | - | - | - | - | G | H | V | N | S | - | - | - | - | - | - | - | - | - | - | - | - | - | - | - | - | V | I | H | C | R | F | S | P | N | D | E | - | - | - | - | - | - | - | - | - | - | - | - | - | - | - | - | - | - | - | - | - | - | - | - | - | - | - | - | Y | V | A | S | C | S | A | D | G | T | V | K | L | - | - | - | - | - | - | - | - | - | - | - | - | - | - | - | - | - | - | - | W | E | - | A | R | S | A | N | E | L | K | S | - | - | - | - | - | - | I | E | - | - | - | - | - | - | - | - | - | - | - | - | - | - | - | - | - | - | - | - | - | - | I | K | D | F | - | - | - | - | - | - | - | - | F | R | N | A | D | E | Q | P | D | - | - | - | - | - | - | - | - | - | - | - | - | - | - | - | - | - | - | - | - | - | - | - | - | - | - | - | D | V | E | - | V | L | V | K | C | C | - | - | - | - | - | - | - | - | - | - | - | - | - | - | - | - | S | W | S | R | N | G | D | M | L | L | V | V | A | K | N | K | L | - | - | - | - | - | - | - | - | - | - | - | L | L | F | D | V | K | T | - | - | - | C | D | - | - | - | - | - | - | - | - | - | - | - | - | - | - | - | - | - | - | - | - | - | - | - | - | - | - | - | - | - | - | - | - | - | - | - | - | - | - | - | - | - | - | - | - | - | - | - | - | - | - | - | - | - | - | - | - | - | - | - | - | - | - | - | - | - | - | - | - | - | - | - | - | - | - | - | - | - | - | - | - | - | - | - | - | - | - | - | - | - | - | - | - | - | - | - | - | - | - | - | - | - | - | - | - | - | - | - | - | - | - | - | - | - | - | - | - | - | - | - | - | - | - | - | - | - | - | L | L | T | - | - | - | - | - | - | - | - | - | - | - | - | - | - | - | - | - | - | - | - | - | - | - | - | - | - | - | - | - | - | - | - | - | - | - | - | - | - | - | - | - | - | - | - | - | - | - | - | - | - | - | - | - | - | - | - | - | - | - | - | - | - | - | - | - | - | - | - | - | - | - | - | - | - | - | - | - | - | - | - | - | - | - | - | - | - | - | - | - | - | - | - | - | - | - | - | - | - | - | - | - | - | - | - | - | - | - | - | - | - | - | - | - | - | - | - | - | - | - | - | - | - | - | - | - | - | - | - | - | - | - | - | - | - | - | - | - | - | - | - | - | - | - | - | - | - | - | - | - | - | - | - | - | - | - | - | - | - | - | - | - | - | - | - | - | - | - | - | - | - | - | - | - | - | - | - | - | - | - | - | - | - | - | Q | V | - | - | - | I | V | S | H | H | S | T | I | Q | Y | C | - | - | - | - | - | - | - | - | - | - | - | - | - | - | - | - | - | - | - | - | - | - | - | - | - | D | F | C | P | G | D | E | L | V | A | V | A | L | S | H | - | - | - | - | - | - | C | S | V | E | - | - | - | - | - | - | - | - | - | - | - | - | - | - | - | - | - | L | W | N | I | K | S | L | - | - | - | - | - | - | - | - | - | - | - | - | - | - | - | - | - | - | - | - | - | - | - | - | - | - | - | - | - | S | K | V | A | Y | C | R | G | H | M | S | - | - | - | - | - | - | - | - | - | - | - | - | - | - | - | - | - | - | - | - | - | - | - | - | - | - | - | - | - | - | - | - | - | - | - | - | - | - | - | - | - | - | - | - | W | V | H | C | V | T | F | S | - | - | - | - | - | - | - | - | - | - | - | P | D | G | S | - | - | - | - | - | - | - | - | - | - | L | F | L | T | S | S | D | D | Q | - | - | - | - | - | - | - | - | - | - | - | - | - | - | T | I | R | I | - | W | - | - | - | - | - | - | - | - | - | - | - | - | - | - | - | - | - | - | - | - | - | - | - | - | - | - | - | - | - | - | - | - | - | - | - | - | - | - | E | T | K | K | V | C | K | S | S | D | A | V | L | K | S | E | - | - | - | - | - | - | - | - | - | - | - | - | - | - | - | - | L | D | V | V | F | - | - | - | - | - | - | - | - | - | - | - | - | - | - | - | - | - | - | - | - | - | - | - | - | - | - | - | - | - | - | - | - | - | - | - | - | - | - | - | - | - | - | - | - | - | - | - | - | - | - | - | - | - | - | H | S | D | E | V | M | - | - | - | - | - | - | - | - | - | - | - | - | - | - | - | - | - | - | - | - | - | - | - | - | - | - | - | - | - | - | - | - | - | - | - | - | I | L | A | I | Y | N | Q | K | H | L | Q | L | - | I | N | G | N | T | - | - | - | - | - | - | - | - | - | - | D | S | V | V | M | H | T | A | A | - | - | - | - | - | - | - | - | - | - | - | - | - | - | - | - | - | - | - | - | - | - | Q | E | S | - | - | - | - | - | - | - | - | P | I | S | C | C | C | L | S | E | D | L | K | - | - | - | - | - | - | - | - | - | F | A | A | F | G | Q | - | - | - | - | - | - | - | - | - | - | - | - | - | - | - | - | - | - | - | - | - | - | - | - | - | - | - | - | - | - | - | - | - | - | - | - | E | - | - | - | - | - | - | - | - | - | S | G | T | - | - | - | - | - | - | - | - | - | - | I | - | - | - | - | - | - | - | - | - | - | - | - | - | - | - | - | - | - | - | - | - | - | - | - | - | - | - | - | - | - | - | - | - | - | - | - | - | - | - | K | V | L | Q | L | L | D | G | K | V | L | K | S | R | E | A | - | - | - | - | - | Y | - | - | - | - | - | - | - | - | - | - | - | - | R | T | S | V | Q | H | C | R | F | T | - | - | - | - | - | - | - | - | - | - | - | - | - | - | - | - | - | - | - | - | - | - | - | - | - | - | - | - | - | - | - | - | - | - | - | - | - | S | D | C | Q | T | L | I | S | S | A | - | H | D | S | - | - | - | - | - | - | - | - | V | I | Q | - | V | - | - | - | - | - | - | - | - | - | - | - | - | - | - | - | - | - | - | - | - | - | - | - | - | - | - | - | - | - | - | - | - | - | - | - | - | - | - | - | W | N | - | - | - | - | - | - | - | - | - | - | W | Q | L | S | E | C | V | F | - | - | - | - | - | - | - | - | - | - | - | - | - | - | - | - | - | - | - | - | - | - | - | - | - | - | - | - | - | - | - | - | - | - | - | - | - | - | - | - | - | - | - | - | - | - | - | - | - | - | L | R | G | H | K | - | - | - | - | - | - | - | - | - | - | - | - | - | - | - | - | - | - | - | - | - | - | - | - | - | - | E | A | I | K | D | F | R | L | L | - | - | - | - | - | - | - | - | - | - | - | - | - | - | - | - | - | - | - | - | - | - | - | E | N | - | S | K | L | L | - | S | W | - | - | - | S | F | D | G | T | V | - | - | - | - | - | - | - | - | - | - | - | - | - | - | - | - | - | - | - | - | - | - | - | - | - | - | - | - | - | - | - | - | - | - | - | K | V | W | N | I | T | T | G | N | T | - | - | - | - | - | E | K | D | F | A | C | - | - | - | - | - | - | - | H | G | - | - | - | - | - | - | - | - | - | - | - | - | - | - | - | - | - | - | - | G | A | V | L | - | - | - | - | - | - | - | - | - | - | - | - | - | - | - | S | C | A | - | - | - | - | - | - | - | - | - | - | - | - | - | - | - | V | S | P | D | G | S | - | K | F | S | S | T | - | S | A | D | K | T | - | - | - | - | - | - | A | K | I | W | S | F | E | S | S | - | - | - | - | - | - | - | - | - | - | - | - | - | - | - | - | - | - | - | - | - | - | - | - | S | V | L | H | E | - | - | - | - | - | - | - | - | L | - | - | - | - | K | G | H | E | A | C | V | - | - | - | - | - | - | - | - | - | R | C | C | - | T | F | S | - | - | - | - | - | - | - | - | - | P | N | N | K | - | L | L | A | T | - | - | - | - | - | - | - | - | G | D | D | K | G | E | I | R | I | W | D | I | L | T | G | A | L | L | - | - | - | - | - | - | - | - | - | - | - | - | - | - | - | - | - | - | - | - | - | - | - | - | - | - | - | - | - | - | - | - | - | - | - | - | - | - | - | - | - | - | - | - | - | - | - | - | - | - | - | - | - | - | - | - | H | F | C | S | P | V | T | V | D | - | - | - | - | - | - | - | - | - | - | - | - | - | - | - | - | - | - | - | - | E | G | E | P | T | H | G | G | W | - | - | - | - | - | - | - | - | V | T | D | L | S | F | S | P | D | S | K | - | - | - | - | - | - | - | - | - | - | - | - | - | - | - | - | - | - | - | - | - | - | - | - | - | - | - | - | - | - | - | - | - | - | - | - | - | - | - | - | - | - | - | - | - | M | L | V | S | - | - | S | G | G | Y | L | K | - | - | - | - | - | - | - | - | W | W | N | V | I | T | - | - | - | - | - | - | - | - | - | - | - | - | - | - | - | - | - | - | - | - | - | - | - | - | - | - | - | - | - | - | - | - | - | - | - | - | - | - | - | - | - | - | - | - | - | - | - | - | - | - | - | - | - | - | - | - | - | - | - | - | - | - | - | - | - | - | - | - | - | - | - | - | - | - | - | - | - | - | - | - | - | - | - | - | - | - | - | - | - | - | - | - | - | - | - | - | - | - | - | - | - | - | - | - | - | - | - | - | - | - | - | - | - | - | - | - | - | - | - | - | - | - | - | - | - | - | - | - | - | - | - | - | - | - | - | - | - | - | - | - | - | - | - | - | - | - | - | - | - | - | - | - | - | - | - | - | - | - | - | - | - | - | - | - | - | - | - | - | - | - | - | - | - | - | - | - | - | - | - | - | - | - | - | - | - | - | - | - | - | - | - | - | - | - | - | - | - | - | - | - | - | - | - | - | - | - | - | - | - | - | - | - | - | - | - | - | - | - | - | - | - | - | - | - | - | - | - | - | - | - | - | - | - | - | - | - | - | - | - | - | - | - | - | - | - | - | - | - | - | - | - | - | - | - | - | - | - | - | - | - | - | - | - | - | - | - | - | G | E | S | L | Q | T | F | Y | T | - | - | - | - | - | N | G | - | - | - | T | N | L | K | S | I | - | - | - | H | V | C | P | - | - | - | - | N | F | E | V | - | - | - | - | - | - | - | - | - | - | - | - | - | - | - | - | - | - | - | - | - | - | - | - | - | - | Y | V | T | V | D | - | N | L | G | I | L | - | - | Y | V | L | Q | K | - | - | - | - | - | - | - | - | - | - | - | - | - | - | - | - | - | - | - | - | - | - | - | - | - | - | - | - | - | - | - | - | - | - | L | - | - | - | - | - | - | - | - | - | - | - | - | - | - | - | - | - | - | - | - | - | - | - | - | - | - | - | - | - | - | - | - | - | - | - | - | - | - | - | - | - | - | - | - | - | - | - | - | - | - | - | - | - | M | - | - | - | - | - | - | - | - | - | - | - | - | - | - | - | - | - | - | - | - | - | - | - | - | - | - | - | - | - | - | - | - | - | - | - | - | - | - | - | - | - | - | - | - | - | - | - | - | - | - | - | - | - | - | E | - | - | - | - | - | - | - | - | - | - | - | - | - | - | - | - | - | - | - | - | - | - | - | - | - | - | L | C | L | - | - | - | - | - | - | - | - | - | - | - | - | - | - | - | - | - | - | - | - | - | - | - | - | - | - | - | - | - | - | S | Q | - | - | - | - | - | - | - | - | P | Q | G | - | - | - | - | - | - | - | - | - | - | - | - | - | - | - | - | - | - | - | - | - | - | - | - | - | - | - | - | - | - | - | - | - | - | - | - | - | - | - | - | - | - | - | - | - | - | - | - | - | - | - | - | - | - | - | - | - | - | - | - | - | - | - | - | - | - | - | - | - | - | - | - | - | - | - | - | - | - | - | - | - | - | - | - | - | - | - | - | - | Q | P | P | V | C | P | S | A | - | - | - | - | - | - | - | - | - | - | - | - | - | - | - | - | - | - | - | - | - | - | - | F | A | W | S | A | - | - | - | - | - | - | - | - | - | - | G | S | R | G | E | Q | - | A | R | Q | L | V | I | D | L | E | T | R | G | - | K | Q | A | F | P | V | F | L | S | I | L | R | D | T | G | Q | G | D | L | A | D | M | L | I | E | E | C | E | C | L | - | - | - | - | P | V | S | S | - | Q | - | - | - | - | - | - | - | - | - | - | - | - | - | - | - | - | - | - | - | - | - | - | - | - | - | - | - | - | - | M | V | D | - | - | - | - | - | - | - | - | - | - | - | - | - | - | - | L | R | P | V | - | - | - | - | - | - | - | - | - | - | - | - | - | - | - | - | - | - | - | - | - | - | - | - | - | - | - | - | - | - | - | - | - | - | - | - | - | - | - | E | L | D | L | R | G | E | K | H | H | K | N | V | - | - | - | - | - | - | - | - | - | - | - | N | - | - | - | - | - | - | - | - | - | - | - | - | - | - | - | - | - | - | - | - | - | - | - | - | - | - | - | - | - | - | - | - | - | - | - | - | - | - | - | - | - | - | - | - | - | - | - | - | - | - | - | - | - | - | - | - | - | - | - | - | - | - | - | - | - | - | - | - | - | - | - | - | - | - | - | - | - | - | - | - | - | - | - | - | - | - | - | - | - | - | - | - | - | - | - | - | - | - | - | - | - | - | - | - | - | - | - | - | - | - | - | - | - | - | - | - | - | - | - | - | - | - | - | - | - | - | - | - | - | - | - | - | - | - | - | - | - | - | - | - | - | - | - | - | - | - | - | - | - | - | - | - | - | - | - | - | F | P | E | R | L | S | - | - | - | - | - | - | - | - | - | - | - | - | - | - | - | - | - | - | - | - | - | - | - | - | - | - | I | P | V | Q | A | - | - | - | - | - | - | E | S | E | R | P | - | R | M | - | - | - | - | - | - | - | - | - | - | - | - | - | - | - | - | - | - | - | - | - | - | - | - | G | - | - | - | - | - | - | - | - | - | - | - | - | - | - | - | - | - | - | - | - | S | - | - | - | - | - | - | - | - | - | - | A | - | V | D | - | - | K | M | N | R | T | E | - | - | S | C | V | V | C | E | - | - | L | K | A | D | - | P | C | G | H | C | L | I | V | N | N | V | N | F | S | S | D | S | - | - | N | L | S | T | R | E | G | S | D | V | D | C | K | K | L | K | Q | R | F | - | K | A | L | H | F | N | V | L | T | R | Q | N | L | K | A | Q | D | M | V | S | E | L | K | K | L | A | Q | Q | D | H | - | R | A | L | D | C | C | V | V | V | I | L | S | H | G | C | Q | T | S | H | I | Q | F | P | G | S | - | I | Y | G | T | D | G | E | P | I | A | I | E | K | I | V | N | C | F | N | G | S | N | C | P | S | L | R | G | K | P | K | L | F | F | I | Q | A | C | - | - | G | G | E | Q | R | D | Q | G | C | V | V | D | C | D | - | - | - | S | P | G | D | - | E | A | P | G | G | S | - | - | - | - | - | - | - | L | E | S | D | A | T | P | F | R | T | L | V | G | S | T | - | - | - | - | - | - | - | - | D | E | P | D | A | V | - | - | - | - | - | - | - | A | S | L | - | - | - | P | T | P | S | D | I | L | V | S | Y | S | T | F | P | - | - | G | - | - | - | - | - | - | - | - | F | V | S | W | R | D | V | S | R | G | S | W | Y | V | E | M | L | D | S | V | L | E | Q | Y | A | H | F | S | - | - | - | - | - | - | - | - | - | - | - | - | - | - | - | - | - | - | - | - | - | - | - | - | - | - | - | - | - | - | - | P | Q | L | L | S | F | C | - | - | - | - | - | S | Q | - | - | - | - | - | - | - | - | - | - | - | - | V | A | N | A | V | - | - | - | - | - | A | D | K | - | - | - | - | - | - | - | G | K | Y | - | K | Q | I | - | - | P | G | - | - | - | - | - | - | - | - | - | - | - | - | - | - | - | C | F | N | F | - | - | - | - | - | - | - | - | - | L | R | K | K | F | F | F | M | C | K | - | - | - | - | - | - | - | - | - | - | - | - | - | - | - | - | - | - | - | - | - | - | - | - | - | - | - | - | - | - | - | - | - | - | - | - | - | M | - | - | - | - | - | - | - | - | - | - | - | - | - | - | - | - | - | - | - | - | - | - | - | - | - | - | - | - | - | - | - | - | - | - | - | - | - | - | - | - | - | - | - | - | - | - | - | - | - | - | - | - | G | D | I | E | K | G | K | K | I | F | V | Q | K | C | S | Q | C | H | T | V | E | K | G | G | K | H | K | T | G | P | N | L | N | G | L | F | G | R | K | T | G | Q | A | E | G | F | S | Y | T | D | A | N | K | N | K | G | I | T | W | G | E | D | T | L | M | E | Y | L | E | N | P | K | K | Y | I | P | G | T | K | M | I | F | A | G | I | K | K | K | S | E | R | A | D | L | I | A | Y | L | K | D | A | T | S | K | - | - | - | - | - | - |
| Chelonia\_mydas | M | - | - | - | - | - | - | - | - | - | - | - | - | - | - | - | - | - | - | - | - | - | - | - | - | - | - | - | - | - | - | - | - | - | - | - | - | - | - | - | - | - | - | - | - | - | - | - | - | - | - | - | - | - | - | - | - | - | - | - | - | - | - | - | - | - | - | - | - | - | - | - | - | - | - | - | - | - | - | - | - | - | - | - | - | - | - | - | - | - | - | - | - | - | - | - | - | - | D | V | K | S | - | R | N | - | - | - | - | - | - | - | - | - | - | - | - | - | - | - | - | H | L | L | - | - | - | - | - | - | - | - | - | - | - | - | - | - | - | - | - | - | - | - | - | - | - | - | - | - | - | - | - | - | - | - | - | - | - | - | - | - | - | - | - | - | - | - | - | - | - | - | - | - | - | - | - | - | - | - | - | - | - | - | - | - | - | - | - | - | - | - | - | - | - | - | - | - | - | - | - | - | - | - | - | - | - | - | - | - | - | - | - | - | - | - | - | - | - | - | - | - | - | - | - | - | - | - | - | - | - | - | - | - | - | - | - | - | - | - | - | - | - | - | - | L | N | H | E | A | L | E | R | D | I | K | T | S | Y | - | I | M | D | H | M | V | A | D | E | V | L | T | L | Q | E | E | E | K | V | K | A | Q | - | - | - | - | - | - | - | - | - | - | - | - | - | - | - | - | - | T | T | Q | K | E | R | A | A | M | L | I | K | I | I | L | A | K | - | - | - | - | D | N | Y | S | Y | I | S | - | - | - | - | - | F | Y | N | A | L | - | L | H | E | G | - | - | - | - | - | - | Y | K | D | L | A | A | L | L | Q | D | G | I | P | I | I | S | S | - | - | - | - | - | - | - | - | - | - | - | - | - | - | - | - | - | - | - | - | - | - | - | - | - | - | - | - | - | - | - | - | - | - | - | - | - | - | - | - | - | - | - | - | - | S | T | V | R | T | V | L | C | E | G | G | - | - | - | - | - | - | - | V | P | Q | R | P | V | V | F | V | I | R | P | K | L | V | K | A | I | - | - | - | - | - | - | - | - | - | - | Q | Q | K | L | Y | - | - | - | - | - | - | - | - | - | - | - | - | - | - | - | - | - | - | - | - | - | - | - | - | - | - | - | - | - | - | - | - | N | L | R | S | D | P | - | - | - | - | - | - | - | - | - | - | - | - | - | - | - | - | - | - | - | - | - | - | - | - | - | - | - | - | - | - | - | - | - | - | - | - | - | - | G | W | V | T | V | Y | G | M | A | G | C | G | K | S | V | L | T | A | E | A | L | - | R | D | H | H | L | L | E | D | C | F | P | G | G | V | H | W | I | S | V | G | - | - | - | - | - | - | - | K | - | - | - | - | - | - | Q | D | K | A | - | - | - | - | - | - | - | - | - | - | - | - | G | L | L | M | K | L | Q | N | L | C | S | R | L | D | - | - | - | - | Q | - | - | - | - | - | - | - | - | - | - | - | - | - | - | - | D | F | S | L | S | Q | R | P | P | L | N | I | - | - | - | - | - | - | - | - | - | - | - | - | - | - | - | - | E | E | A | K | D | R | L | R | L | L | I | L | R | - | - | - | - | - | - | - | - | - | - | - | - | K | Y | P | R | C | L | M | - | - | V | L | D | D | I | W | D | S | W | V | L | K | A | F | D | - | N | Q | C | Q | V | L | I | T | S | R | D | R | S | V | T | D | S | V | - | T | G | N | - | - | K | Y | E | V | P | V | E | S | - | - | - | G | L | A | H | D | K | G | L | E | I | L | S | L | F | V | N | M | - | K | T | T | D | L | P | E | - | - | - | - | - | - | - | - | - | - | - | - | - | - | E | A | D | S | I | V | R | E | C | K | - | - | - | - | - | - | - | - | - | - | - | - | - | - | - | - | - | - | - | G | - | S | P | L | V | V | S | L | I | G | A | L | L | R | D | F | P | N | - | - | - | - | - | - | - | - | - | - | R | W | E | Y | Y | L | - | R | Q | L | Q | N | K | Q | F | K | R | I | - | - | - | - | - | - | - | - | - | - | - | - | - | - | - | - | - | - | - | - | - | - | R | K | S | S | S | Y | D | - | - | - | - | Y | E | A | L | D | E | A | M | S | M | S | V | E | - | M | L | N | - | - | - | E | N | F | K | D | Y | - | Y | K | D | L | S | V | L | Q | K | D | V | K | V | P | T | K | V | L | C | I | L | W | D | - | - | - | - | M | E | T | E | E | V | E | D | I | L | Q | E | F | - | - | - | - | - | - | - | - | - | - | - | V | N | K | S | L | L | F | C | D | R | N | G | K | S | - | - | - | - | - | - | - | - | - | F | H | Y | Y | L | H | D | L | Q | L | D | F | L | T | E | K | - | - | N | C | N | Q | - | L | - | Q | E | L | H | K | K | I | V | N | Q | Y | M | K | - | H | Y | - | - | - | - | - | - | Q | L | H | I | P | T | P | A | - | Q | E | - | - | - | - | - | - | - | D | C | M | Y | W | Y | N | F | L | A | Y | H | M | A | S | A | N | - | - | - | - | M | Q | N | E | L | C | A | L | M | F | S | L | D | W | I | K | A | K | T | E | L | V | G | P | A | - | - | - | - | - | - | - | - | H | L | I | H | E | Y | V | E | Y | S | H | I | L | D | Q | K | - | - | - | - | - | - | - | D | S | I | V | R | E | N | F | Q | - | - | - | E | F | L | S | L | N | G | H | L | L | G | R | Q | P | F | - | - | - | - | - | - | - | - | - | - | - | - | - | - | - | - | - | - | - | - | - | - | - | - | - | - | - | - | - | - | - | - | - | - | - | - | - | - | - | - | - | - | - | - | - | - | - | - | - | - | - | - | - | - | - | - | - | - | - | - | - | - | - | - | - | - | - | - | - | - | - | P | N | I | I | Q | L | G | L | C | Q | P | E | T | S | E | V | Y | Q | Q | A | K | L | Q | A | Q | Q | E | I | V | K | G | - | - | - | - | - | - | - | - | - | - | - | - | - | - | - | - | - | - | - | - | - | - | - | - | - | - | - | - | - | A | F | Y | L | - | - | - | - | - | - | - | - | - | - | E | W | I | N | K | - | - | - | - | - | - | - | K | S | M | K | N | L | S | R | - | - | - | - | - | - | - | - | - | - | L | V | V | R | P | H | T | D | - | - | - | - | - | - | - | - | - | - | - | - | - | - | - | A | V | Y | H | A | C | - | - | - | F | S | - | K | D | G | Q | R | I | A | - | - | - | - | - | - | - | - | - | S | C | G | A | D | K | T | L | Q | V | F | K | - | - | - | - | - | A | K | S | G | E | K | L | L | E | I | - | K | A | H | D | D | E | - | - | - | V | L | C | C | T | F | S | - | V | - | - | - | - | - | - | - | - | - | - | - | - | - | D | D | R | F | V | A | T | C | S | V | D | K | K | V | K | I | - | - | - | W | N | S | R | - | - | - | - | - | - | - | - | - | - | - | - | - | - | - | - | - | - | - | - | - | - | - | - | - | - | - | - | - | - | - | - | - | - | - | - | - | - | - | - | - | - | - | - | - | - | - | - | - | - | - | - | - | - | - | - | - | - | - | - | - | - | - | - | - | - | - | - | - | - | - | - | - | - | - | - | - | - | - | - | - | - | - | - | - | - | - | - | - | - | - | - | - | - | - | - | - | - | - | - | - | - | - | - | - | - | - | - | - | - | - | - | - | - | - | - | - | - | - | - | - | - | - | - | - | - | - | - | - | - | - | - | - | - | - | - | - | - | - | - | T | G | Q | L | V | Q | V | Y | - | E | E | H | V | - | - | - | - | - | E | Q | V | N | - | - | - | - | - | - | - | - | - | - | - | - | - | C | C | Q | - | - | - | - | - | - | - | - | - | - | - | - | - | F | N | N | T | N | - | - | - | - | - | - | - | - | - | - | - | - | - | - | - | - | D | Q | - | L | L | L | A | T | C | S | N | - | - | - | - | D | T | Y | V | - | - | - | - | - | - | - | - | - | - | - | - | K | L | W | N | L | - | - | - | N | K | K | H | C | R | N | T | M | F | - | - | - | - | - | - | - | - | - | - | - | - | - | - | - | - | - | G | H | L | N | S | - | - | - | - | - | - | - | - | - | - | - | - | - | - | - | - | V | S | H | C | K | F | S | P | D | D | K | - | - | - | - | - | - | - | - | - | - | - | - | - | - | - | - | - | - | - | - | - | - | - | - | - | - | - | - | Y | V | A | S | C | S | A | D | G | T | L | K | L | - | - | - | - | - | - | - | - | - | - | - | - | - | - | - | - | - | - | - | W | E | - | V | C | S | A | N | E | L | K | T | - | - | - | - | - | - | I | D | - | - | - | - | - | - | - | - | - | - | - | - | - | - | - | - | - | - | - | - | - | - | I | K | D | F | - | - | - | - | - | - | - | - | F | R | N | A | D | E | H | Q | D | - | - | - | - | - | - | - | - | - | - | - | - | - | - | - | - | - | - | - | - | - | - | - | - | - | - | - | D | V | E | - | V | L | V | K | C | C | - | - | - | - | - | - | - | - | - | - | - | - | - | - | - | - | S | W | S | A | N | C | A | T | I | M | V | A | A | K | N | R | L | - | - | - | - | - | - | - | - | - | - | - | L | L | L | D | V | I | T | - | - | - | S | D | - | - | - | - | - | - | - | - | - | - | - | - | - | - | - | - | - | - | - | - | - | - | - | - | - | - | - | - | - | - | - | - | - | - | - | - | - | - | - | - | - | - | - | - | - | - | - | - | - | - | - | - | - | - | - | - | - | - | - | - | - | - | - | - | - | - | - | - | - | - | - | - | - | - | - | - | - | - | - | - | - | - | - | - | - | - | - | - | - | - | - | - | - | - | - | - | - | - | - | - | - | - | - | - | - | - | - | - | - | - | - | - | - | - | - | - | - | - | - | - | - | - | - | - | - | - | L | L | A | - | - | - | - | - | - | - | - | - | - | - | - | - | - | - | - | - | - | - | - | - | - | - | - | - | - | - | - | - | - | - | - | - | - | - | - | - | - | - | - | - | - | - | - | - | - | - | - | - | - | - | - | - | - | - | - | - | - | - | - | - | - | - | - | - | - | - | - | - | - | - | - | - | - | - | - | - | - | - | - | - | - | - | - | - | - | - | - | - | - | - | - | - | - | - | - | - | - | - | - | - | - | - | - | - | - | - | - | - | - | - | - | - | - | - | - | - | - | - | - | - | - | - | - | - | - | - | - | - | - | - | - | - | - | - | - | - | - | - | - | - | - | - | - | - | - | - | - | - | - | - | - | - | - | - | - | - | - | - | - | - | - | - | - | - | - | - | - | - | - | - | - | - | - | - | - | - | - | - | - | - | - | - | E | V | - | - | - | L | I | S | H | H | S | T | I | Q | Y | C | - | - | - | - | - | - | - | - | - | - | - | - | - | - | - | - | - | - | - | - | - | - | - | - | - | D | F | C | P | G | N | Q | L | V | A | V | A | L | S | H | - | - | - | - | - | - | Y | S | V | E | - | - | - | - | - | - | - | - | - | - | - | - | - | - | - | - | - | L | W | N | T | E | S | F | - | - | - | - | - | - | - | - | - | - | - | - | - | - | - | - | - | - | - | - | - | - | - | - | - | - | - | - | - | S | K | V | A | D | C | R | G | H | L | S | - | - | - | - | - | - | - | - | - | - | - | - | - | - | - | - | - | - | - | - | - | - | - | - | - | - | - | - | - | - | - | - | - | - | - | - | - | - | - | - | - | - | - | - | W | V | H | C | V | T | F | S | - | - | - | - | - | - | - | - | - | - | - | P | D | G | S | - | - | - | - | - | - | - | - | - | - | L | F | L | T | S | S | D | D | Q | - | - | - | - | - | - | - | - | - | - | - | - | - | - | T | I | R | I | - | W | - | - | - | - | - | - | - | - | - | - | - | - | - | - | - | - | - | - | - | - | - | - | - | - | - | - | - | - | - | - | - | - | - | - | - | - | - | - | E | S | S | K | V | C | K | S | S | A | V | V | L | K | R | E | - | - | - | - | - | - | - | - | - | - | - | - | - | - | - | - | L | D | V | V | F | - | - | - | - | - | - | - | - | - | - | - | - | - | - | - | - | - | - | - | - | - | - | - | - | - | - | - | - | - | - | - | - | - | - | - | - | - | - | - | - | - | - | - | - | - | - | - | - | - | - | - | - | - | - | Q | D | D | E | V | M | - | - | - | - | - | - | - | - | - | - | - | - | - | - | - | - | - | - | - | - | - | - | - | - | - | - | - | - | - | - | - | - | - | - | - | - | V | L | A | I | L | N | Q | K | H | L | Q | L | - | I | N | G | N | T | - | - | - | - | - | - | - | - | - | - | G | H | T | D | Y | L | T | E | A | - | - | - | - | - | - | - | - | - | - | - | - | - | - | - | - | - | - | - | - | - | - | Q | E | S | - | - | - | - | - | - | - | - | L | I | C | C | C | C | L | S | E | D | L | Q | - | - | - | - | - | - | - | - | - | Y | A | A | F | G | E | - | - | - | - | - | - | - | - | - | - | - | - | - | - | - | - | - | - | - | - | - | - | - | - | - | - | - | - | - | - | - | - | - | - | - | - | E | - | - | - | - | - | - | - | - | - | N | G | T | - | - | - | - | - | - | - | - | - | - | V | - | - | - | - | - | - | - | - | - | - | - | - | - | - | - | - | - | - | - | - | - | - | - | - | - | - | - | - | - | - | - | - | - | - | - | - | - | - | - | K | V | L | K | L | S | N | R | R | V | F | K | S | R | T | G | - | - | - | - | - | H | - | - | - | - | - | - | - | - | - | - | - | - | E | K | S | V | Q | H | C | Q | F | T | - | - | - | - | - | - | - | - | - | - | - | - | - | - | - | - | - | - | - | - | - | - | - | - | - | - | - | - | - | - | - | - | - | - | - | - | - | S | D | G | Q | T | L | I | S | S | S | - | D | D | S | - | - | - | - | - | - | - | - | T | I | Q | - | V | - | - | - | - | - | - | - | - | - | - | - | - | - | - | - | - | - | - | - | - | - | - | - | - | - | - | - | - | - | - | - | - | - | - | - | - | - | - | - | W | N | - | - | - | - | - | - | - | - | - | - | W | Q | S | E | E | Y | I | F | - | - | - | - | - | - | - | - | - | - | - | - | - | - | - | - | - | - | - | - | - | - | - | - | - | - | - | - | - | - | - | - | - | - | - | - | - | - | - | - | - | - | - | - | - | - | - | - | - | - | L | K | G | H | K | - | - | - | - | - | - | - | - | - | - | - | - | - | - | - | - | - | - | - | - | - | - | - | - | - | - | E | A | V | K | N | F | R | L | L | - | - | - | - | - | - | - | - | - | - | - | - | - | - | - | - | - | - | - | - | - | - | - | E | N | - | S | R | L | L | - | S | W | - | - | - | S | F | D | G | T | V | - | - | - | - | - | - | - | - | - | - | - | - | - | - | - | - | - | - | - | - | - | - | - | - | - | - | - | - | - | - | - | - | - | - | - | K | V | W | N | I | C | T | G | K | I | - | - | - | - | - | E | D | D | F | V | C | - | - | - | - | - | - | - | H | E | - | - | - | - | - | - | - | - | - | - | - | - | - | - | - | - | - | - | - | D | A | V | L | - | - | - | - | - | - | - | - | - | - | - | - | - | - | - | S | C | A | - | - | - | - | - | - | - | - | - | - | - | - | - | - | - | I | S | P | D | A | T | - | K | F | S | S | T | - | S | A | D | K | T | - | - | - | - | - | - | A | K | I | W | S | F | E | S | S | - | - | - | - | - | - | - | - | - | - | - | - | - | - | - | - | - | - | - | - | - | - | - | - | S | V | L | H | E | - | - | - | - | - | - | - | - | L | - | - | - | - | N | G | H | K | G | C | V | - | - | - | - | - | - | - | - | - | R | C | C | - | T | F | S | - | - | - | - | - | - | - | - | - | F | D | D | K | - | F | L | A | T | - | - | - | - | - | - | - | - | G | D | D | N | G | E | I | R | I | W | D | V | S | R | G | E | L | L | - | - | - | - | - | - | - | - | - | - | - | - | - | - | - | - | - | - | - | - | - | - | - | - | - | - | - | - | - | - | - | - | - | - | - | - | - | - | - | - | - | - | - | - | - | - | - | - | - | - | - | - | - | - | - | - | H | L | C | S | P | V | T | V | D | - | - | - | - | - | - | - | - | - | - | - | - | - | - | - | - | - | - | - | - | E | G | E | S | T | H | G | G | W | - | - | - | - | - | - | - | - | V | T | D | L | C | F | S | P | E | S | N | - | - | - | - | - | - | - | - | - | - | - | - | - | - | - | - | - | - | - | - | - | - | - | - | - | - | - | - | - | - | - | - | - | - | - | - | - | - | - | - | - | - | - | - | - | M | L | V | S | - | - | C | G | G | Y | L | K | - | - | - | - | - | - | - | - | W | W | N | V | T | T | - | - | - | - | - | - | - | - | - | - | - | - | - | - | - | - | - | - | - | - | - | - | - | - | - | - | - | - | - | - | - | - | - | - | - | - | - | - | - | - | - | - | - | - | - | - | - | - | - | - | - | - | - | - | - | - | - | - | - | - | - | - | - | - | - | - | - | - | - | - | - | - | - | - | - | - | - | - | - | - | - | - | - | - | - | - | - | - | - | - | - | - | - | - | - | - | - | - | - | - | - | - | - | - | - | - | - | - | - | - | - | - | - | - | - | - | - | - | - | - | - | - | - | - | - | - | - | - | - | - | - | - | - | - | - | - | - | - | - | - | - | - | - | - | - | - | - | - | - | - | - | - | - | - | - | - | - | - | - | - | - | - | - | - | - | - | - | - | - | - | - | - | - | - | - | - | - | - | - | - | - | - | - | - | - | - | - | - | - | - | - | - | - | - | - | - | - | - | - | - | - | - | - | - | - | - | - | - | - | - | - | - | - | - | - | - | - | - | - | - | - | - | - | - | - | - | - | - | - | - | - | - | - | - | - | - | - | - | - | - | - | - | - | - | - | - | - | - | - | - | - | - | - | - | - | - | - | - | - | - | - | - | - | - | - | - | - | G | E | S | L | Q | T | F | Y | T | - | - | - | - | - | N | G | - | - | - | T | N | L | K | S | I | - | - | - | Y | V | S | P | - | - | - | - | N | F | K | V | - | - | - | - | - | - | - | - | - | - | - | - | - | - | - | - | - | - | - | - | - | - | - | - | - | - | Y | V | T | I | D | - | N | L | G | I | L | - | - | Y | V | L | Q | M | - | - | - | - | - | - | - | - | - | - | - | - | - | - | - | - | - | - | - | - | - | - | - | - | - | - | - | - | - | - | - | - | - | - | L | E | - | - | - | - | - | - | - | - | - | - | - | - | - | - | - | - | - | - | - | - | - | - | - | - | - | - | - | - | - | - | - | - | - | - | - | - | - | - | - | - | - | - | - | - | - | - | - | - | - | - | - | - | M | - | - | - | - | - | - | - | - | - | - | - | - | - | - | - | - | - | - | - | - | - | - | - | - | - | - | - | - | - | - | - | - | - | - | - | - | - | - | - | - | - | - | - | - | - | - | - | - | - | - | - | - | - | - | S | Y | T | T | K | - | - | - | - | - | - | - | - | - | - | - | - | - | - | - | - | - | - | - | - | - | - | - | - | - | - | - | - | - | - | - | - | - | - | - | - | - | - | - | - | - | - | - | - | - | - | - | - | - | - | - | - | - | - | - | - | - | - | - | - | - | - | - | - | - | - | - | - | - | - | - | - | - | - | - | - | - | - | - | - | - | - | - | - | - | - | - | - | - | - | - | - | - | - | - | - | - | - | - | - | - | - | - | - | - | - | - | - | - | - | - | - | - | - | - | - | - | - | - | - | - | - | - | - | - | - | - | - | - | - | - | - | - | - | - | - | - | - | - | - | - | - | - | - | - | - | - | - | - | - | - | - | - | - | - | - | - | - | - | - | - | - | - | - | - | - | - | - | - | - | - | - | - | - | - | - | - | - | - | - | - | - | - | - | - | - | - | - | - | - | K | R | A | - | - | - | - | - | - | - | - | - | - | G | T | R | R | D | Q | - | A | R | Q | L | V | T | D | L | Q | T | R | G | - | R | H | A | F | P | A | F | V | S | S | L | R | E | T | G | Q | N | D | L | A | T | L | L | S | E | G | D | K | F | P | - | - | - | - | Q | L | R | P | - | - | - | - | - | - | - | - | - | - | - | - | - | - | - | - | - | - | - | - | - | - | - | - | - | - | - | - | - | - | - | - | V | D | - | - | - | - | - | - | - | - | - | - | - | - | - | - | - | L | K | P | V | - | - | - | - | - | - | - | - | - | - | - | - | - | - | - | - | - | - | - | - | - | - | - | - | - | - | - | - | - | - | - | - | - | - | - | - | - | - | - | E | I | G | L | R | G | G | K | G | D | T | Y | G | - | - | - | - | - | - | - | - | - | - | - | K | - | - | - | - | - | - | - | - | - | - | - | - | - | - | - | - | - | - | - | - | - | - | - | - | - | - | - | - | - | - | - | - | - | - | - | - | - | - | - | - | - | - | - | - | - | - | - | - | - | - | - | - | - | - | - | - | - | - | - | - | - | - | - | - | - | - | - | - | - | - | - | - | - | - | - | - | - | - | - | - | - | - | - | - | - | - | - | - | - | - | - | - | - | - | - | - | - | - | - | - | - | - | - | - | - | - | - | - | - | - | - | - | - | - | - | - | - | - | - | - | - | - | - | - | - | - | - | - | - | - | - | - | - | - | - | - | - | - | - | - | - | - | - | - | - | - | - | - | - | - | - | - | - | - | - | - | A | H | E | Y | L | P | - | - | - | - | - | - | - | - | - | - | - | - | - | - | - | - | - | - | - | - | - | - | - | - | - | - | T | P | I | Q | A | - | - | - | - | - | - | V | T | E | R | G | - | Q | V | S | T | - | - | - | - | - | - | - | - | - | - | - | - | - | - | - | C | P | S | A | A | A | P | G | - | - | - | - | - | - | - | - | - | - | - | - | - | - | - | - | - | - | - | - | S | - | - | - | - | - | - | - | - | - | - | A | - | A | E | K | S | R | R | N | S | D | - | - | - | - | - | M | V | Y | V | - | - | L | K | A | N | - | P | C | G | Y | C | L | I | I | N | N | V | R | F | T | E | E | S | - | - | D | L | S | S | R | E | G | S | N | V | D | C | E | K | L | E | K | R | F | - | K | S | L | H | F | D | V | L | V | Q | R | D | L | K | A | Q | E | I | A | R | E | L | Q | H | L | A | R | R | D | H | - | S | A | L | D | C | C | L | V | V | I | L | S | H | G | C | Q | T | S | H | I | Q | F | P | G | G | - | I | Y | G | T | D | G | K | L | I | P | V | E | K | I | V | N | F | F | N | G | S | H | C | P | S | L | R | G | K | P | K | L | F | F | I | Q | A | C | - | - | G | G | E | Q | K | D | R | G | F | E | V | D | S | D | - | - | - | S | P | V | N | - | Q | P | R | G | G | S | - | - | - | - | - | - | - | V | E | S | D | A | T | P | F | Q | I | Q | P | G | N | L | - | - | - | - | - | - | - | - | D | E | P | D | A | V | - | - | - | - | - | - | - | A | S | L | - | - | - | P | T | P | S | D | I | F | V | S | Y | S | T | F | P | - | - | G | - | - | - | - | - | - | - | - | F | V | S | W | R | D | T | Q | T | G | S | W | Y | V | E | T | L | D | S | V | L | E | Q | Y | A | P | - | T | - | - | - | - | - | - | - | - | - | - | - | - | - | - | - | - | - | - | - | - | - | - | - | - | - | - | - | - | - | - | - | E | D | L | L | C | M | L | - | - | - | - | - | V | R | - | - | - | - | - | - | - | - | - | - | - | - | V | A | H | A | V | - | - | - | - | - | S | A | K | - | - | - | - | - | - | - | G | K | Y | - | K | Q | M | - | - | P | G | - | - | - | - | - | - | - | - | - | - | - | - | - | - | - | T | F | N | F | - | - | - | - | - | - | - | - | - | L | R | K | Q | F | F | F | L | S | D | - | - | - | - | - | - | - | - | - | - | - | - | - | - | - | - | - | - | - | - | - | - | - | - | - | - | - | - | - | - | - | - | - | - | - | - | - | M | - | - | - | - | - | - | - | - | - | - | - | - | - | - | - | - | - | - | - | - | - | - | - | - | - | - | - | - | - | - | - | - | - | - | - | - | - | - | - | - | - | - | - | - | - | - | - | - | - | - | - | - | G | D | I | E | K | G | K | K | I | F | V | Q | K | C | A | Q | C | H | T | V | E | K | G | G | K | H | K | T | G | P | N | L | N | G | L | I | G | R | K | T | G | Q | A | E | N | F | S | Y | T | E | A | N | K | N | K | G | I | T | W | N | E | E | T | L | M | E | Y | L | E | N | P | K | K | Y | I | P | G | T | K | M | I | F | A | G | I | K | K | K | T | E | R | A | D | L | I | A | Y | L | K | D | A | T | S | K | - | - | - | - | - | - |
| Chinchilla\_lanigera | M | - | - | - | - | - | - | - | - | - | - | - | - | - | - | - | - | - | - | - | - | - | - | - | - | - | - | - | - | - | - | - | - | - | - | - | - | - | - | - | - | - | - | - | - | - | - | - | - | - | - | - | - | - | - | - | - | - | - | - | - | - | - | - | - | - | - | - | - | - | - | - | - | - | - | - | - | - | - | - | - | - | - | - | - | - | - | - | - | - | - | - | - | - | - | - | - | - | D | A | K | A | - | R | N | - | - | - | - | - | - | - | - | - | - | - | - | - | - | - | - | C | L | L | - | - | - | - | - | - | - | - | - | - | - | - | - | - | - | - | - | - | - | - | - | - | - | - | - | - | - | - | - | - | - | - | - | - | - | - | - | - | - | - | - | - | - | - | - | - | - | - | - | - | - | - | - | - | - | - | - | - | - | - | - | - | - | - | - | - | - | - | - | - | - | - | - | - | - | - | - | - | - | - | - | - | - | - | - | - | - | - | - | - | - | - | - | - | - | - | - | - | - | - | - | - | - | - | - | - | - | - | - | - | - | - | - | - | - | - | - | - | - | - | - | Q | H | R | E | A | L | E | K | D | I | K | T | S | Y | - | I | M | D | H | M | I | S | D | G | V | L | T | I | S | E | E | E | R | V | K | N | Q | - | - | - | - | - | - | - | - | - | - | - | - | - | - | - | - | - | P | S | N | H | Q | R | A | A | T | L | I | K | M | I | L | K | K | - | - | - | - | D | N | Y | S | Y | I | S | - | - | - | - | - | F | Y | N | A | L | - | L | H | E | G | - | - | - | - | - | - | Y | K | D | L | A | A | L | L | H | S | G | M | P | V | L | S | S | S | S | G | - | K | - | - | - | - | - | - | - | - | - | - | - | - | - | - | - | - | - | - | - | - | - | - | D | S | A | D | G | I | T | - | - | - | - | - | - | - | - | - | - | - | S | Y | V | R | T | V | L | C | E | G | G | - | - | - | - | - | - | - | V | P | Q | R | P | V | V | F | V | T | R | K | K | L | V | N | A | I | - | - | - | - | - | - | - | - | - | - | Q | Q | K | L | L | - | - | - | - | - | - | - | - | - | - | - | - | - | - | - | - | - | - | - | - | - | - | - | - | - | - | - | - | - | - | - | - | K | L | N | G | E | P | - | - | - | - | - | - | - | - | - | - | - | - | - | - | - | - | - | - | - | - | - | - | - | - | - | - | - | - | - | - | - | - | - | - | - | - | - | - | G | W | V | S | I | Y | G | M | A | G | C | G | K | S | V | L | A | A | E | A | V | - | R | D | H | S | L | L | E | G | C | F | P | G | G | V | H | W | V | S | V | G | - | - | - | - | - | - | - | K | - | - | - | - | - | - | Q | D | K | S | - | - | - | - | - | - | - | - | - | - | - | - | G | L | L | M | K | L | Q | N | L | C | T | R | L | D | - | - | - | - | Q | - | - | - | - | - | - | - | - | - | - | - | - | - | - | - | D | E | S | F | S | Q | R | L | P | L | N | I | - | - | - | - | - | - | - | - | - | - | - | - | - | - | - | - | E | E | A | K | D | R | L | R | I | L | M | L | R | - | - | - | - | - | - | - | - | - | - | - | - | K | H | P | R | S | L | L | - | - | I | L | D | D | I | W | D | P | W | V | L | K | A | F | D | - | N | Q | C | Q | I | L | L | T | T | R | D | K | S | V | T | D | S | V | - | M | G | P | - | - | K | Y | V | V | P | V | E | S | - | - | - | G | L | G | K | E | K | G | L | E | I | L | S | L | F | V | N | R | - | K | K | A | D | L | P | E | - | - | - | - | - | - | - | - | - | - | - | - | - | - | E | A | H | S | I | I | K | E | C | K | - | - | - | - | - | - | - | - | - | - | - | - | - | - | - | - | - | - | - | G | - | S | P | L | V | V | S | L | I | G | A | L | L | R | D | F | P | N | - | - | - | - | - | - | - | - | - | - | R | W | E | Y | Y | L | - | R | Q | L | Q | N | K | Q | F | K | R | I | - | - | - | - | - | - | - | - | - | - | - | - | - | - | - | - | - | - | - | - | - | - | R | K | S | S | S | Y | D | - | - | - | - | Y | E | A | L | D | E | A | M | S | I | S | V | E | - | M | L | R | - | - | - | E | D | I | K | D | Y | - | Y | T | D | L | S | I | L | Q | K | D | V | K | V | P | T | Q | V | L | C | I | L | W | D | - | - | - | - | M | E | T | E | E | V | E | D | I | L | Q | E | F | - | - | - | - | - | - | - | - | - | - | - | V | N | K | S | L | L | F | C | D | R | N | G | K | S | - | - | - | - | - | - | - | - | - | F | C | Y | Y | L | H | D | L | Q | V | D | F | L | T | E | K | - | - | N | R | S | Q | - | L | - | Q | D | L | H | K | K | M | I | T | Q | F | Q | R | - | Y | H | - | - | - | - | - | - | Q | P | H | T | L | S | P | D | - | Q | E | - | - | - | - | - | - | - | D | C | M | Y | W | Y | N | F | L | A | Y | H | M | A | S | A | N | - | - | - | - | M | H | K | E | L | C | T | L | M | F | S | L | D | W | I | K | A | K | T | E | L | V | G | P | A | - | - | - | - | - | - | - | - | H | L | I | H | E | F | V | E | Y | R | H | I | L | D | E | K | - | - | - | - | - | - | - | D | C | A | V | C | E | N | F | Q | - | - | - | E | F | L | S | L | N | G | H | L | L | G | R | Q | P | F | - | - | - | - | - | - | - | - | - | - | - | - | - | - | - | - | - | - | - | - | - | - | - | - | - | - | - | - | - | - | - | - | - | - | - | - | - | - | - | - | - | - | - | - | - | - | - | - | - | - | - | - | - | - | - | - | - | - | - | - | - | - | - | - | - | - | - | - | - | - | - | P | N | I | V | Q | L | G | L | C | E | P | E | T | S | E | V | Y | Q | Q | A | K | L | Q | A | K | Q | E | V | D | N | G | - | - | - | - | - | - | - | - | - | - | - | - | - | - | - | - | - | - | - | - | - | - | - | - | - | - | - | - | - | M | L | Y | L | - | - | - | - | - | - | - | - | - | - | E | W | I | N | K | - | - | - | - | - | - | - | K | T | I | K | N | L | S | R | - | - | - | - | - | - | - | - | - | - | L | V | V | R | P | H | T | D | - | - | - | - | - | - | - | - | - | - | - | - | - | - | - | A | V | Y | H | A | C | - | - | - | F | S | - | E | D | G | Q | R | I | A | - | - | - | - | - | - | - | - | - | S | C | G | A | D | K | T | L | Q | V | F | K | - | - | - | - | - | A | E | T | G | E | K | L | L | E | I | - | N | A | H | E | D | E | - | - | - | V | L | C | C | A | F | S | - | T | - | - | - | - | - | - | - | - | - | - | - | - | - | D | D | R | F | I | A | T | C | S | V | D | K | K | V | K | I | - | - | - | W | N | S | V | - | - | - | - | - | - | - | - | - | - | - | - | - | - | - | - | - | - | - | - | - | - | - | - | - | - | - | - | - | - | - | - | - | - | - | - | - | - | - | - | - | - | - | - | - | - | - | - | - | - | - | - | - | - | - | - | - | - | - | - | - | - | - | - | - | - | - | - | - | - | - | - | - | - | - | - | - | - | - | - | - | - | - | - | - | - | - | - | - | - | - | - | - | - | - | - | - | - | - | - | - | - | - | - | - | - | - | - | - | - | - | - | - | - | - | - | - | - | - | - | - | - | - | - | - | - | - | - | - | - | - | - | - | - | - | - | - | - | - | - | T | G | E | L | V | H | T | Y | - | D | E | H | S | - | - | - | - | - | E | Q | V | N | - | - | - | - | - | - | - | - | - | - | - | - | - | C | C | C | - | - | - | - | - | - | - | - | - | - | - | - | - | F | T | N | N | S | - | - | - | - | - | - | - | - | - | - | - | - | - | - | - | - | Q | N | - | L | L | L | A | T | A | S | N | - | - | - | - | D | S | F | L | - | - | - | - | - | - | - | - | - | - | - | - | K | L | W | D | L | - | - | - | N | Q | K | Q | C | R | N | T | L | F | - | - | - | - | - | - | - | - | - | - | - | - | - | - | - | - | - | G | H | T | N | S | - | - | - | - | - | - | - | - | - | - | - | - | - | - | - | - | V | T | H | C | K | F | S | P | D | D | E | - | - | - | - | - | - | - | - | - | - | - | - | - | - | - | - | - | - | - | - | - | - | - | - | - | - | - | - | L | L | A | S | C | S | A | D | G | T | L | K | L | - | - | - | - | - | - | - | - | - | - | - | - | - | - | - | - | - | - | - | W | D | - | V | R | S | A | N | E | R | K | S | - | - | - | - | - | - | V | N | - | - | - | - | - | - | - | - | - | - | - | - | - | - | - | - | - | - | - | - | - | - | V | K | Q | F | - | - | - | - | - | - | - | - | F | L | N | P | E | D | C | Q | E | - | - | - | - | - | - | - | - | - | - | - | - | - | - | - | - | - | - | - | - | - | - | - | - | - | - | - | E | M | E | - | V | I | V | K | C | C | - | - | - | - | - | - | - | - | - | - | - | - | - | - | - | - | S | W | S | A | D | G | A | R | V | M | A | A | A | K | N | K | I | - | - | - | - | - | - | - | - | - | - | - | F | L | L | D | V | H | T | - | - | - | S | G | - | - | - | - | - | - | - | - | - | - | - | - | - | - | - | - | - | - | - | - | - | - | - | - | - | - | - | - | - | - | - | - | - | - | - | - | - | - | - | - | - | - | - | - | - | - | - | - | - | - | - | - | - | - | - | - | - | - | - | - | - | - | - | - | - | - | - | - | - | - | - | - | - | - | - | - | - | - | - | - | - | - | - | - | - | - | - | - | - | - | - | - | - | - | - | - | - | - | - | - | - | - | - | - | - | - | - | - | - | - | - | - | - | - | - | - | - | - | - | - | - | - | - | - | - | - | L | V | A | - | - | - | - | - | - | - | - | - | - | - | - | - | - | - | - | - | - | - | - | - | - | - | - | - | - | - | - | - | - | - | - | - | - | - | - | - | - | - | - | - | - | - | - | - | - | - | - | - | - | - | - | - | - | - | - | - | - | - | - | - | - | - | - | - | - | - | - | - | - | - | - | - | - | - | - | - | - | - | - | - | - | - | - | - | - | - | - | - | - | - | - | - | - | - | - | - | - | - | - | - | - | - | - | - | - | - | - | - | - | - | - | - | - | - | - | - | - | - | - | - | - | - | - | - | - | - | - | - | - | - | - | - | - | - | - | - | - | - | - | - | - | - | - | - | - | - | - | - | - | - | - | - | - | - | - | - | - | - | - | - | - | - | - | - | - | - | - | - | - | - | - | - | - | - | - | - | - | - | - | - | - | - | E | I | - | - | - | H | T | G | H | H | S | T | I | Q | Y | C | - | - | - | - | - | - | - | - | - | - | - | - | - | - | - | - | - | - | - | - | - | - | - | - | - | D | F | S | P | Q | N | H | L | A | V | V | A | L | S | Q | - | - | - | - | - | - | F | C | V | E | - | - | - | - | - | - | - | - | - | - | - | - | - | - | - | - | - | L | W | N | T | D | S | C | - | - | - | - | - | - | - | - | - | - | - | - | - | - | - | - | - | - | - | - | - | - | - | - | - | - | - | - | - | V | K | V | A | D | C | R | G | H | F | S | - | - | - | - | - | - | - | - | - | - | - | - | - | - | - | - | - | - | - | - | - | - | - | - | - | - | - | - | - | - | - | - | - | - | - | - | - | - | - | - | - | - | - | - | W | V | H | G | V | M | F | S | - | - | - | - | - | - | - | - | - | - | - | P | D | G | S | - | - | - | - | - | - | - | - | - | - | S | F | L | T | S | S | D | D | Q | - | - | - | - | - | - | - | - | - | - | - | - | - | - | T | I | R | V | - | W | - | - | - | - | - | - | - | - | - | - | - | - | - | - | - | - | - | - | - | - | - | - | - | - | - | - | - | - | - | - | - | - | - | - | - | - | - | - | E | T | K | K | M | C | K | N | S | A | V | V | L | K | Q | E | - | - | - | - | - | - | - | - | - | - | - | - | - | - | - | - | I | D | V | V | F | - | - | - | - | - | - | - | - | - | - | - | - | - | - | - | - | - | - | - | - | - | - | - | - | - | - | - | - | - | - | - | - | - | - | - | - | - | - | - | - | - | - | - | - | - | - | - | - | - | - | - | - | - | - | Q | D | S | E | V | M | - | - | - | - | - | - | - | - | - | - | - | - | - | - | - | - | - | - | - | - | - | - | - | - | - | - | - | - | - | - | - | - | - | - | - | - | V | L | A | A | D | N | V | R | R | L | Q | L | - | I | N | G | K | T | - | - | - | - | - | - | - | - | - | - | G | Q | T | D | Y | L | T | E | A | - | - | - | - | - | - | - | - | - | - | - | - | - | - | - | - | - | - | - | - | - | - | Q | - | - | - | - | - | - | - | - | - | - | - | V | S | C | C | C | L | N | P | H | L | Q | - | - | - | - | - | - | - | - | - | H | I | A | F | G | G | - | - | - | - | - | - | - | - | - | - | - | - | - | - | - | - | - | - | - | - | - | - | - | - | - | - | - | - | - | - | - | - | - | - | - | - | E | - | - | - | - | - | - | - | - | - | D | G | A | - | - | - | - | - | - | - | - | - | - | V | - | - | - | - | - | - | - | - | - | - | - | - | - | - | - | - | - | - | - | - | - | - | - | - | - | - | - | - | - | - | - | - | - | - | - | - | - | - | - | E | I | L | E | L | L | N | N | K | V | F | Q | S | R | I | G | - | - | - | - | - | H | - | - | - | - | - | - | - | - | - | - | - | - | K | K | A | V | R | H | I | Q | F | S | - | - | - | - | - | - | - | - | - | - | - | - | - | - | - | - | - | - | - | - | - | - | - | - | - | - | - | - | - | - | - | - | - | - | - | - | - | A | D | G | K | T | L | I | S | S | S | - | D | D | S | - | - | - | - | - | - | - | - | A | I | Q | - | V | - | - | - | - | - | - | - | - | - | - | - | - | - | - | - | - | - | - | - | - | - | - | - | - | - | - | - | - | - | - | - | - | - | - | - | - | - | - | - | W | N | - | - | - | - | - | - | - | - | - | - | W | Q | S | E | E | C | V | L | - | - | - | - | - | - | - | - | - | - | - | - | - | - | - | - | - | - | - | - | - | - | - | - | - | - | - | - | - | - | - | - | - | - | - | - | - | - | - | - | - | - | - | - | - | - | - | - | - | - | L | R | A | H | Q | - | - | - | - | - | - | - | - | - | - | - | - | - | - | - | - | - | - | - | - | - | - | - | - | - | - | E | T | V | T | D | F | R | L | L | - | - | - | - | - | - | - | - | - | - | - | - | - | - | - | - | - | - | - | - | - | - | - | E | N | - | S | R | L | L | - | S | W | - | - | - | S | F | D | G | T | V | - | - | - | - | - | - | - | - | - | - | - | - | - | - | - | - | - | - | - | - | - | - | - | - | - | - | - | - | - | - | - | - | - | - | - | K | V | W | N | I | I | T | G | K | I | - | - | - | - | - | E | K | D | F | V | C | - | - | - | - | - | - | - | H | Q | - | - | - | - | - | - | - | - | - | - | - | - | - | - | - | - | - | - | - | G | T | V | L | - | - | - | - | - | - | - | - | - | - | - | - | - | - | - | S | C | A | - | - | - | - | - | - | - | - | - | - | - | - | - | - | - | I | S | S | D | A | T | - | K | F | S | S | T | - | S | A | D | K | T | - | - | - | - | - | - | A | K | I | W | S | F | E | L | P | - | - | - | - | - | - | - | - | - | - | - | - | - | - | - | - | - | - | - | - | - | - | - | - | S | P | L | H | E | - | - | - | - | - | - | - | - | L | - | - | - | - | K | G | H | K | G | C | V | - | - | - | - | - | - | - | - | - | R | C | S | - | A | F | S | - | - | - | - | - | - | - | - | - | E | D | G | S | - | L | L | A | T | - | - | - | - | - | - | - | - | G | D | D | N | G | E | V | R | I | W | S | V | S | D | G | E | L | L | - | - | - | - | - | - | - | - | - | - | - | - | - | - | - | - | - | - | - | - | - | - | - | - | - | - | - | - | - | - | - | - | - | - | - | - | - | - | - | - | - | - | - | - | - | - | - | - | - | - | - | - | - | - | - | - | H | L | C | A | P | F | S | V | E | - | - | - | - | - | - | - | - | - | - | - | - | - | - | - | - | - | - | - | - | E | G | A | A | T | H | A | G | W | - | - | - | - | - | - | - | - | V | T | D | L | C | F | S | P | D | S | K | - | - | - | - | - | - | - | - | - | - | - | - | - | - | - | - | - | - | - | - | - | - | - | - | - | - | - | - | - | - | - | - | - | - | - | - | - | - | - | - | - | - | - | - | - | M | L | V | S | - | - | A | G | G | Y | L | K | - | - | - | - | - | - | - | - | W | W | N | V | I | T | - | - | - | - | - | - | - | - | - | - | - | - | - | - | - | - | - | - | - | - | - | - | - | - | - | - | - | - | - | - | - | - | - | - | - | - | - | - | - | - | - | - | - | - | - | - | - | - | - | - | - | - | - | - | - | - | - | - | - | - | - | - | - | - | - | - | - | - | - | - | - | - | - | - | - | - | - | - | - | - | - | - | - | - | - | - | - | - | - | - | - | - | - | - | - | - | - | - | - | - | - | - | - | - | - | - | - | - | - | - | - | - | - | - | - | - | - | - | - | - | - | - | - | - | - | - | - | - | - | - | - | - | - | - | - | - | - | - | - | - | - | - | - | - | - | - | - | - | - | - | - | - | - | - | - | - | - | - | - | - | - | - | - | - | - | - | - | - | - | - | - | - | - | - | - | - | - | - | - | - | - | - | - | - | - | - | - | - | - | - | - | - | - | - | - | - | - | - | - | - | - | - | - | - | - | - | - | - | - | - | - | - | - | - | - | - | - | - | - | - | - | - | - | - | - | - | - | - | - | - | - | - | - | - | - | - | - | - | - | - | - | - | - | - | - | - | - | - | - | - | - | - | - | - | - | - | - | - | - | - | - | - | - | - | - | - | - | G | E | S | A | Q | T | F | Y | T | - | - | - | - | - | N | G | - | - | - | T | N | L | K | K | I | - | - | - | H | V | S | P | - | - | - | - | D | F | K | T | - | - | - | - | - | - | - | - | - | - | - | - | - | - | - | - | - | - | - | - | - | - | - | - | - | - | Y | V | T | V | D | - | N | L | G | I | L | - | - | Y | I | L | Q | I | - | - | - | - | - | - | - | - | - | - | - | - | - | - | - | - | - | - | - | - | - | - | - | - | - | - | - | - | - | - | - | - | - | - | L | E | - | - | - | - | - | - | - | - | - | - | - | - | - | - | - | - | - | - | - | - | - | - | - | - | - | - | - | - | - | - | - | - | - | - | - | - | - | - | - | - | - | - | - | - | - | - | - | - | - | - | - | - | M | - | - | - | - | - | - | - | - | - | - | - | - | - | - | - | - | - | - | - | - | - | - | - | - | - | - | - | - | - | - | - | - | - | - | - | - | - | - | - | - | - | - | - | - | - | - | - | - | - | - | - | - | - | - | D | Q | E | D | R | E | - | - | - | - | - | - | - | - | - | - | - | - | - | - | - | - | - | - | - | - | L | L | R | R | - | - | - | - | - | - | - | - | - | - | - | - | - | - | - | - | - | - | - | - | - | - | - | - | - | - | - | - | - | - | C | R | - | - | - | - | - | - | - | - | L | R | L | - | V | G | D | L | Q | V | A | P | L | W | - | - | - | - | - | - | - | - | - | - | - | - | - | - | - | - | - | - | - | - | - | - | - | - | - | - | - | - | - | - | - | - | - | - | - | - | - | - | - | - | - | - | - | - | - | D | A | L | L | N | R | E | L | - | - | - | - | - | - | - | - | - | - | - | - | - | - | - | - | - | - | - | - | - | - | - | - | F | T | P | D | M | I | E | D | - | - | - | - | - | - | - | - | - | - | - | - | - | - | - | - | - | - | - | - | - | - | - | - | - | I | Q | R | A | G | - | - | - | - | - | - | - | - | S | G | S | R | R | D | Q | - | A | R | Q | L | I | T | D | L | E | T | R | G | - | R | Q | A | L | P | L | F | I | S | C | L | K | D | T | G | Q | D | T | L | A | S | L | L | C | A | S | R | Q | A | I | - | - | - | - | R | R | D | P | - | E | A | I | - | - | - | - | - | - | - | - | - | - | - | - | - | - | - | - | - | - | - | - | - | - | - | - | - | - | R | P | L | D | - | - | - | - | - | - | - | - | - | - | - | - | - | - | - | P | E | P | A | - | - | - | - | - | - | - | - | - | - | - | - | - | - | - | - | - | - | - | - | - | - | - | - | - | - | - | - | - | - | - | - | - | - | - | - | - | - | - | - | - | - | - | - | - | - | - | - | - | - | - | - | - | - | - | - | - | - | - | - | - | - | - | - | - | - | - | - | - | - | - | - | - | - | - | - | - | - | - | - | - | - | - | - | - | - | - | - | - | - | - | - | - | - | - | - | - | - | - | - | - | - | - | - | - | - | - | - | - | - | - | - | - | - | - | - | - | - | - | - | - | - | - | - | - | - | - | - | - | - | - | - | - | - | - | - | - | - | - | - | - | - | - | - | - | - | - | - | - | - | - | - | - | - | - | - | - | - | - | - | - | - | - | - | - | - | - | - | - | - | - | - | - | - | - | - | - | - | - | - | - | - | - | - | - | - | - | - | - | - | - | - | - | - | - | - | - | - | - | - | - | - | - | - | - | - | - | - | - | - | - | - | - | - | - | - | - | - | - | - | - | - | - | - | - | - | - | - | - | - | - | - | - | - | - | - | - | - | - | - | - | - | - | - | - | - | - | - | - | - | - | - | - | - | - | - | - | - | - | - | - | - | - | - | V | M | E | P | - | M | V | - | - | - | - | - | - | - | - | - | - | - | - | - | - | - | - | - | - | P | K | P | S | - | - | - | - | - | - | - | - | - | - | - | - | - | - | - | - | - | - | - | - | - | - | - | - | - | - | - | - | - | - | - | - | - | - | S | - | Q | E | T | M | K | G | N | A | D | - | - | - | - | - | L | A | Y | V | - | - | L | S | A | E | - | P | C | G | H | C | L | I | I | N | N | V | R | F | C | P | T | S | - | - | G | L | S | T | R | T | G | S | N | I | D | C | E | K | M | Q | R | R | F | - | C | R | L | H | F | Q | V | E | V | R | Y | D | L | T | A | K | E | M | V | Q | A | L | A | Q | L | A | Q | R | D | H | - | H | A | L | D | C | C | V | V | I | I | L | S | H | G | C | Q | A | S | H | R | Q | F | P | G | A | - | V | Y | G | T | D | G | C | S | V | S | V | E | K | I | V | N | I | F | N | G | T | N | C | P | S | L | G | G | K | P | K | L | F | F | I | Q | A | C | - | - | G | G | E | Q | K | D | H | G | F | D | V | A | F | V | - | - | - | S | P | D | T | - | E | A | P | G | S | S | - | - | - | - | - | - | - | D | E | A | D | A | V | P | F | Q | G | G | P | E | P | T | - | - | - | - | - | - | - | - | D | Q | P | D | A | V | - | - | - | - | - | - | - | S | S | L | - | - | - | P | T | P | S | D | I | L | V | S | Y | S | T | F | P | - | - | G | - | - | - | - | - | - | - | - | F | V | S | W | R | H | T | T | R | G | S | W | Y | I | E | T | L | D | G | I | L | E | Q | W | A | P | - | S | - | - | - | - | - | - | - | - | - | - | - | - | - | - | - | - | - | - | - | - | - | - | - | - | - | - | - | - | - | - | - | E | D | L | Q | D | L | L | - | - | - | - | - | L | R | - | - | - | - | - | - | - | - | - | - | - | - | V | A | N | A | V | - | - | - | - | - | S | E | R | - | - | - | - | - | - | - | G | I | Y | - | K | Q | I | - | - | P | G | - | - | - | - | - | - | - | - | - | - | - | - | - | - | - | C | F | N | F | - | - | - | - | - | - | - | - | - | L | R | K | K | L | F | F | K | T | K | - | - | - | - | - | - | - | - | - | - | - | - | - | - | - | - | - | - | - | - | - | - | - | - | - | - | - | - | - | - | - | - | - | - | - | - | - | M | - | - | - | - | - | - | - | - | - | - | - | - | - | - | - | - | - | - | - | - | - | - | - | - | - | - | - | - | - | - | - | - | - | - | - | - | - | - | - | - | - | - | - | - | - | - | - | - | - | - | - | - | G | D | V | E | K | S | K | K | I | F | I | Q | K | C | A | Q | C | H | T | V | E | K | G | G | R | H | K | T | R | P | N | L | H | G | L | F | G | Q | E | R | G | Q | A | V | G | F | S | C | T | D | A | N | K | N | K | G | I | T | W | G | E | D | T | L | R | E | Y | L | E | N | T | K | K | Y | I | P | G | T | K | M | I | F | A | G | I | T | K | K | G | E | R | A | D | L | I | E | Y | L | K | K | A | T | N | E | - | - | - | - | - | - |
| Colobus\_angolensis\_palliatus | M | - | - | - | - | - | - | - | - | - | - | - | - | - | - | - | - | - | - | - | - | - | - | - | - | - | - | - | - | - | - | - | - | - | - | - | - | - | - | - | - | - | - | - | - | - | - | - | - | - | - | - | - | - | - | - | - | - | - | - | - | - | - | - | - | - | - | - | - | - | - | - | - | - | - | - | - | - | - | - | - | - | - | - | - | - | - | - | - | - | - | - | - | - | - | - | - | - | D | A | K | A | - | R | N | - | - | - | - | - | - | - | - | - | - | - | - | - | - | - | - | C | L | L | - | - | - | - | - | - | - | - | - | - | - | - | - | - | - | - | - | - | - | - | - | - | - | - | - | - | - | - | - | - | - | - | - | - | - | - | - | - | - | - | - | - | - | - | - | - | - | - | - | - | - | - | - | - | - | - | - | - | - | - | - | - | - | - | - | - | - | - | - | - | - | - | - | - | - | - | - | - | - | - | - | - | - | - | - | - | - | - | - | - | - | - | - | - | - | - | - | - | - | - | - | - | - | - | - | - | - | - | - | - | - | - | - | - | - | - | - | - | - | - | - | Q | H | R | E | A | L | E | K | D | I | K | T | S | Y | - | I | M | D | H | M | I | S | D | G | I | L | T | I | S | E | E | E | K | V | R | I | E | - | - | - | - | - | - | - | - | - | - | - | - | - | - | - | - | - | P | T | Q | Q | Q | R | A | A | M | L | I | K | M | I | L | K | K | - | - | - | - | D | N | D | S | Y | I | S | - | - | - | - | - | F | Y | N | A | L | - | L | H | E | G | - | - | - | - | - | - | Y | K | D | L | A | A | L | L | H | D | G | I | P | V | V | S | S | S | S | G | - | K | - | - | - | - | - | - | - | - | - | - | - | - | - | - | - | - | - | - | - | - | - | - | D | S | V | S | G | I | T | - | - | - | - | - | - | - | - | - | - | - | S | Y | V | R | T | V | L | C | E | G | G | - | - | - | - | - | - | - | V | P | Q | R | P | V | V | F | V | T | R | K | K | L | V | N | T | I | - | - | - | - | - | - | - | - | - | - | Q | Q | K | L | S | - | - | - | - | - | - | - | - | - | - | - | - | - | - | - | - | - | - | - | - | - | - | - | - | - | - | - | - | - | - | - | - | K | L | K | G | E | P | - | - | - | - | - | - | - | - | - | - | - | - | - | - | - | - | - | - | - | - | - | - | - | - | - | - | - | - | - | - | - | - | - | - | - | - | - | - | G | W | V | T | I | H | G | M | A | G | C | G | K | S | V | L | A | A | E | A | V | - | R | D | H | S | L | L | E | D | C | F | P | G | G | V | H | W | V | S | V | G | - | - | - | - | - | - | - | K | - | - | - | - | - | - | Q | D | K | S | - | - | - | - | - | - | - | - | - | - | - | - | G | L | L | M | K | L | Q | N | L | C | T | R | L | D | - | - | - | - | Q | - | - | - | - | - | - | - | - | - | - | - | - | - | - | - | D | E | S | F | S | Q | R | L | P | L | N | I | - | - | - | - | - | - | - | - | - | - | - | - | - | - | - | - | E | E | A | K | D | R | L | R | I | L | M | L | R | - | - | - | - | - | - | - | - | - | - | - | - | K | H | P | R | S | L | L | - | - | I | L | D | D | V | W | D | S | W | V | L | K | A | F | D | - | N | Q | C | Q | I | L | L | T | T | R | D | K | S | V | T | D | S | V | - | M | G | P | - | - | K | Y | V | V | P | V | E | S | - | - | - | S | L | G | K | E | K | G | L | E | I | L | S | L | F | V | N | M | - | K | K | A | D | L | P | E | - | - | - | - | - | - | - | - | - | - | - | - | - | - | Q | A | H | S | I | I | K | E | C | K | - | - | - | - | - | - | - | - | - | - | - | - | - | - | - | - | - | - | - | G | - | S | P | L | V | V | S | L | I | G | A | L | L | R | D | F | P | N | - | - | - | - | - | - | - | - | - | - | R | W | E | Y | Y | L | - | K | Q | L | Q | N | K | Q | F | K | R | I | - | - | - | - | - | - | - | - | - | - | - | - | - | - | - | - | - | - | - | - | - | - | R | K | S | S | S | Y | D | - | - | - | - | Y | E | A | L | D | E | A | M | S | I | S | V | E | - | M | L | R | - | - | - | E | D | I | K | Y | Y | - | Y | T | D | L | S | I | L | Q | K | D | V | K | V | P | T | K | V | L | C | I | L | W | D | - | - | - | - | M | E | T | E | E | V | E | D | I | L | Q | E | F | - | - | - | - | - | - | - | - | - | - | - | V | N | K | S | L | L | F | C | D | R | N | G | K | S | - | - | - | - | - | - | - | - | - | F | R | Y | Y | L | H | D | L | Q | V | D | F | L | T | E | K | - | - | N | C | S | Q | - | L | - | Q | D | L | H | K | K | I | I | T | Q | F | Q | R | - | Y | Y | - | - | - | - | - | - | Q | P | H | T | L | S | P | D | - | Q | E | - | - | - | - | - | - | - | D | C | M | Y | W | Y | N | F | L | A | Y | H | M | A | S | A | K | - | - | - | - | M | H | K | E | L | C | A | L | M | F | S | L | D | W | I | K | A | K | T | E | L | V | G | P | A | - | - | - | - | - | - | - | - | H | L | I | H | E | F | V | E | Y | R | H | I | L | D | E | K | - | - | - | - | - | - | - | D | C | A | V | C | E | N | F | Q | - | - | - | E | F | L | S | L | N | G | H | L | L | G | R | Q | P | F | - | - | - | - | - | - | - | - | - | - | - | - | - | - | - | - | - | - | - | - | - | - | - | - | - | - | - | - | - | - | - | - | - | - | - | - | - | - | - | - | - | - | - | - | - | - | - | - | - | - | - | - | - | - | - | - | - | - | - | - | - | - | - | - | - | - | - | - | - | - | - | P | N | I | V | Q | L | G | L | C | E | P | E | T | S | E | V | Y | Q | Q | A | K | L | Q | A | K | Q | E | V | D | N | G | - | - | - | - | - | - | - | - | - | - | - | - | - | - | - | - | - | - | - | - | - | - | - | - | - | - | - | - | - | M | L | Y | L | - | - | - | - | - | - | - | - | - | - | E | W | I | N | K | - | - | - | - | - | - | - | K | N | I | T | N | L | S | R | - | - | - | - | - | - | - | - | - | - | L | V | V | R | P | H | T | D | - | - | - | - | - | - | - | - | - | - | - | - | - | - | - | A | V | Y | H | A | C | - | - | - | F | S | - | E | D | G | Q | R | I | A | - | - | - | - | - | - | - | - | - | S | C | G | A | D | K | T | L | Q | V | F | K | - | - | - | - | - | A | E | T | G | E | K | L | L | E | I | - | K | A | H | E | D | E | - | - | - | V | L | C | C | A | F | S | - | T | - | - | - | - | - | - | - | - | - | - | - | - | - | D | D | R | F | I | A | T | C | S | V | D | K | K | V | K | I | - | - | - | W | N | A | V | - | - | - | - | - | - | - | - | - | - | - | - | - | - | - | - | - | - | - | - | - | - | - | - | - | - | - | - | - | - | - | - | - | - | - | - | - | - | - | - | - | - | - | - | - | - | - | - | - | - | - | - | - | - | - | - | - | - | - | - | - | - | - | - | - | - | - | - | - | - | - | - | - | - | - | - | - | - | - | - | - | - | - | - | - | - | - | - | - | - | - | - | - | - | - | - | - | - | - | - | - | - | - | - | - | - | - | - | - | - | - | - | - | - | - | - | - | - | - | - | - | - | - | - | - | - | - | - | - | - | - | - | - | - | - | - | - | - | - | - | T | G | E | L | V | H | T | Y | - | D | E | H | S | - | - | - | - | - | E | Q | V | N | - | - | - | - | - | - | - | - | - | - | - | - | - | C | C | H | - | - | - | - | - | - | - | - | - | - | - | - | - | F | T | N | S | S | - | - | - | - | - | - | - | - | - | - | - | - | - | - | - | - | H | H | - | L | L | L | A | T | G | S | S | - | - | - | - | D | C | F | L | - | - | - | - | - | - | - | - | - | - | - | - | K | L | W | D | L | - | - | - | N | Q | K | E | C | R | N | T | M | F | - | - | - | - | - | - | - | - | - | - | - | - | - | - | - | - | - | G | H | T | N | S | - | - | - | - | - | - | - | - | - | - | - | - | - | - | - | - | V | N | H | C | R | F | S | P | D | D | K | - | - | - | - | - | - | - | - | - | - | - | - | - | - | - | - | - | - | - | - | - | - | - | - | - | - | - | - | L | L | A | S | C | S | A | D | G | T | L | K | L | - | - | - | - | - | - | - | - | - | - | - | - | - | - | - | - | - | - | - | W | D | - | V | T | S | A | N | E | R | K | S | - | - | - | - | - | - | F | N | - | - | - | - | - | - | - | - | - | - | - | - | - | - | - | - | - | - | - | - | - | - | V | K | Q | F | - | - | - | - | - | - | - | - | F | L | N | S | E | D | P | Q | E | - | - | - | - | - | - | - | - | - | - | - | - | - | - | - | - | - | - | - | - | - | - | - | - | - | - | - | D | M | E | - | V | I | V | K | C | C | - | - | - | - | - | - | - | - | - | - | - | - | - | - | - | - | S | W | F | A | D | G | A | R | I | M | V | A | A | K | N | K | I | - | - | - | - | - | - | - | - | - | - | - | F | L | F | D | I | H | T | - | - | - | S | G | - | - | - | - | - | - | - | - | - | - | - | - | - | - | - | - | - | - | - | - | - | - | - | - | - | - | - | - | - | - | - | - | - | - | - | - | - | - | - | - | - | - | - | - | - | - | - | - | - | - | - | - | - | - | - | - | - | - | - | - | - | - | - | - | - | - | - | - | - | - | - | - | - | - | - | - | - | - | - | - | - | - | - | - | - | - | - | - | - | - | - | - | - | - | - | - | - | - | - | - | - | - | - | - | - | - | - | - | - | - | - | - | - | - | - | - | - | - | - | - | - | - | - | - | - | - | L | L | G | - | - | - | - | - | - | - | - | - | - | - | - | - | - | - | - | - | - | - | - | - | - | - | - | - | - | - | - | - | - | - | - | - | - | - | - | - | - | - | - | - | - | - | - | - | - | - | - | - | - | - | - | - | - | - | - | - | - | - | - | - | - | - | - | - | - | - | - | - | - | - | - | - | - | - | - | - | - | - | - | - | - | - | - | - | - | - | - | - | - | - | - | - | - | - | - | - | - | - | - | - | - | - | - | - | - | - | - | - | - | - | - | - | - | - | - | - | - | - | - | - | - | - | - | - | - | - | - | - | - | - | - | - | - | - | - | - | - | - | - | - | - | - | - | - | - | - | - | - | - | - | - | - | - | - | - | - | - | - | - | - | - | - | - | - | - | - | - | - | - | - | - | - | - | - | - | - | - | - | - | - | - | - | E | I | - | - | - | H | M | G | H | H | S | T | I | Q | Y | C | - | - | - | - | - | - | - | - | - | - | - | - | - | - | - | - | - | - | - | - | - | - | - | - | - | D | F | S | P | Q | N | H | L | A | V | V | A | L | S | Q | - | - | - | - | - | - | Y | C | V | E | - | - | - | - | - | - | - | - | - | - | - | - | - | - | - | - | - | L | W | N | I | D | S | C | - | - | - | - | - | - | - | - | - | - | - | - | - | - | - | - | - | - | - | - | - | - | - | - | - | - | - | - | - | S | K | V | A | D | C | R | G | H | L | S | - | - | - | - | - | - | - | - | - | - | - | - | - | - | - | - | - | - | - | - | - | - | - | - | - | - | - | - | - | - | - | - | - | - | - | - | - | - | - | - | - | - | - | - | W | V | H | G | V | K | F | S | - | - | - | - | - | - | - | - | - | - | - | P | D | G | S | - | - | - | - | - | - | - | - | - | - | S | F | L | T | S | S | D | D | Q | - | - | - | - | - | - | - | - | - | - | - | - | - | - | T | I | R | L | - | W | - | - | - | - | - | - | - | - | - | - | - | - | - | - | - | - | - | - | - | - | - | - | - | - | - | - | - | - | - | - | - | - | - | - | - | - | - | - | E | T | K | K | V | C | K | N | S | A | I | M | L | K | Q | E | - | - | - | - | - | - | - | - | - | - | - | - | - | - | - | - | V | D | V | V | F | - | - | - | - | - | - | - | - | - | - | - | - | - | - | - | - | - | - | - | - | - | - | - | - | - | - | - | - | - | - | - | - | - | - | - | - | - | - | - | - | - | - | - | - | - | - | - | - | - | - | - | - | - | - | Q | E | N | E | V | M | - | - | - | - | - | - | - | - | - | - | - | - | - | - | - | - | - | - | - | - | - | - | - | - | - | - | - | - | - | - | - | - | - | - | - | - | V | L | A | V | D | H | I | R | R | L | Q | L | - | I | N | G | K | T | - | - | - | - | - | - | - | - | - | - | G | Q | I | D | Y | L | T | E | A | - | - | - | - | - | - | - | - | - | - | - | - | - | - | - | - | - | - | - | - | - | - | Q | - | - | - | - | - | - | - | - | - | - | - | V | S | C | C | C | L | S | P | H | L | Q | - | - | - | - | - | - | - | - | - | Y | I | A | F | G | D | - | - | - | - | - | - | - | - | - | - | - | - | - | - | - | - | - | - | - | - | - | - | - | - | - | - | - | - | - | - | - | - | - | - | - | - | E | - | - | - | - | - | - | - | - | - | N | G | T | - | - | - | - | - | - | - | - | - | - | I | - | - | - | - | - | - | - | - | - | - | - | - | - | - | - | - | - | - | - | - | - | - | - | - | - | - | - | - | - | - | - | - | - | - | - | - | - | - | - | E | I | L | E | L | V | N | N | R | I | F | Q | S | R | I | R | - | - | - | - | - | H | - | - | - | - | - | - | - | - | - | - | - | - | K | K | T | V | W | H | I | Q | F | T | - | - | - | - | - | - | - | - | - | - | - | - | - | - | - | - | - | - | - | - | - | - | - | - | - | - | - | - | - | - | - | - | - | - | - | - | - | A | D | E | K | T | L | I | S | S | S | - | D | D | S | - | - | - | - | - | - | - | - | A | I | Q | - | V | - | - | - | - | - | - | - | - | - | - | - | - | - | - | - | - | - | - | - | - | - | - | - | - | - | - | - | - | - | - | - | - | - | - | - | - | - | - | - | W | N | - | - | - | - | - | - | - | - | - | - | W | Q | L | E | E | Y | V | F | - | - | - | - | - | - | - | - | - | - | - | - | - | - | - | - | - | - | - | - | - | - | - | - | - | - | - | - | - | - | - | - | - | - | - | - | - | - | - | - | - | - | - | - | - | - | - | - | - | - | L | Q | A | H | Q | - | - | - | - | - | - | - | - | - | - | - | - | - | - | - | - | - | - | - | - | - | - | - | - | - | - | E | T | V | K | D | F | R | L | L | - | - | - | - | - | - | - | - | - | - | - | - | - | - | - | - | - | - | - | - | - | - | - | K | N | - | S | R | L | L | - | S | W | - | - | - | S | F | D | G | T | V | - | - | - | - | - | - | - | - | - | - | - | - | - | - | - | - | - | - | - | - | - | - | - | - | - | - | - | - | - | - | - | - | - | - | - | K | V | W | N | I | I | T | G | K | I | - | - | - | - | - | E | K | D | F | V | C | - | - | - | - | - | - | - | H | Q | - | - | - | - | - | - | - | - | - | - | - | - | - | - | - | - | - | - | - | G | T | V | L | - | - | - | - | - | - | - | - | - | - | - | - | - | - | - | S | C | D | - | - | - | - | - | - | - | - | - | - | - | - | - | - | - | I | S | H | D | A | T | - | K | F | S | S | T | - | S | A | D | K | T | - | - | - | - | - | - | A | K | I | W | S | F | D | L | L | - | - | - | - | - | - | - | - | - | - | - | - | - | - | - | - | - | - | - | - | - | - | - | - | L | P | L | H | E | - | - | - | - | - | - | - | - | L | - | - | - | - | R | G | H | S | G | C | V | - | - | - | - | - | - | - | - | - | R | C | S | - | A | F | S | - | - | - | - | - | - | - | - | - | V | D | G | T | - | L | L | A | T | - | - | - | - | - | - | - | - | G | D | D | N | G | E | I | R | I | W | N | V | S | N | G | E | L | L | - | - | - | - | - | - | - | - | - | - | - | - | - | - | - | - | - | - | - | - | - | - | - | - | - | - | - | - | - | - | - | - | - | - | - | - | - | - | - | - | - | - | - | - | - | - | - | - | - | - | - | - | - | - | - | - | H | L | C | A | P | L | S | - | E | - | - | - | - | - | - | - | - | - | - | - | - | - | - | - | - | - | - | - | - | E | G | A | A | T | H | G | G | W | - | - | - | - | - | - | - | - | V | T | D | L | C | F | S | P | D | G | K | - | - | - | - | - | - | - | - | - | - | - | - | - | - | - | - | - | - | - | - | - | - | - | - | - | - | - | - | - | - | - | - | - | - | - | - | - | - | - | - | - | - | - | - | - | M | L | I | S | - | - | A | G | G | Y | I | K | - | - | - | - | - | - | - | - | W | W | N | V | V | T | - | - | - | - | - | - | - | - | - | - | - | - | - | - | - | - | - | - | - | - | - | - | - | - | - | - | - | - | - | - | - | - | - | - | - | - | - | - | - | - | - | - | - | - | - | - | - | - | - | - | - | - | - | - | - | - | - | - | - | - | - | - | - | - | - | - | - | - | - | - | - | - | - | - | - | - | - | - | - | - | - | - | - | - | - | - | - | - | - | - | - | - | - | - | - | - | - | - | - | - | - | - | - | - | - | - | - | - | - | - | - | - | - | - | - | - | - | - | - | - | - | - | - | - | - | - | - | - | - | - | - | - | - | - | - | - | - | - | - | - | - | - | - | - | - | - | - | - | - | - | - | - | - | - | - | - | - | - | - | - | - | - | - | - | - | - | - | - | - | - | - | - | - | - | - | - | - | - | - | - | - | - | - | - | - | - | - | - | - | - | - | - | - | - | - | - | - | - | - | - | - | - | - | - | - | - | - | - | - | - | - | - | - | - | - | - | - | - | - | - | - | - | - | - | - | - | - | - | - | - | - | - | - | - | - | - | - | - | - | - | - | - | - | - | - | - | - | - | - | - | - | - | - | - | - | - | - | - | - | - | - | - | - | - | - | - | - | G | E | S | S | Q | T | F | Y | T | - | - | - | - | - | N | G | - | - | - | T | N | L | K | K | I | - | - | - | H | V | S | P | - | - | - | - | D | F | K | T | - | - | - | - | - | - | - | - | - | - | - | - | - | - | - | - | - | - | - | - | - | - | - | - | - | - | Y | V | T | V | D | - | N | L | G | I | L | - | - | Y | I | L | Q | T | - | - | - | - | - | - | - | - | - | - | - | - | - | - | - | - | - | - | - | - | - | - | - | - | - | - | - | - | - | - | - | - | - | - | L | E | - | - | - | - | - | - | - | - | - | - | - | - | - | - | - | - | - | - | - | - | - | - | - | - | - | - | - | - | - | - | - | - | - | - | - | - | - | - | - | - | - | - | - | - | - | - | - | - | - | - | - | - | M | - | - | - | - | - | - | - | - | - | - | - | - | - | - | - | - | - | - | - | - | - | - | - | - | - | - | - | - | - | - | - | - | - | - | - | - | - | - | - | - | - | - | - | - | - | - | - | - | - | - | - | - | - | - | D | E | A | D | R | R | - | - | - | - | - | - | - | - | - | - | - | - | - | - | - | - | - | - | - | - | L | L | R | R | - | - | - | - | - | - | - | - | - | - | - | - | - | - | - | - | - | - | - | - | - | - | - | - | - | - | - | - | - | - | C | R | - | - | - | - | - | - | - | - | L | R | L | - | V | E | E | L | Q | V | D | Q | L | W | - | - | - | - | - | - | - | - | - | - | - | - | - | - | - | - | - | - | - | - | - | - | - | - | - | - | - | - | - | - | - | - | - | - | - | - | - | - | - | - | - | - | - | - | - | D | A | L | L | S | R | E | L | - | - | - | - | - | - | - | - | - | - | - | - | - | - | - | - | - | - | - | - | - | - | - | - | F | K | P | H | M | I | E | D | - | - | - | - | - | - | - | - | - | - | - | - | - | - | - | - | - | - | - | - | - | - | - | - | - | I | Q | R | A | G | - | - | - | - | - | - | - | - | S | G | S | R | R | D | Q | - | A | R | Q | L | I | T | D | L | E | T | R | G | - | S | Q | A | L | P | L | F | I | S | C | L | E | D | T | G | Q | D | M | L | A | S | F | L | R | T | N | R | Q | A | G | - | - | - | - | K | L | - | - | - | - | - | - | - | - | - | - | - | - | - | - | - | - | - | - | - | - | - | - | - | - | - | - | - | - | - | - | - | - | - | - | - | - | - | - | - | - | - | - | - | - | - | - | - | - | - | - | - | - | - | - | - | - | - | - | - | - | - | - | - | - | - | - | - | - | - | - | - | - | - | - | - | - | - | - | - | - | - | - | - | - | - | - | - | - | - | - | - | - | - | - | - | - | - | - | - | - | - | - | - | - | - | - | - | - | - | - | - | - | - | - | S | K | P | - | T | - | L | - | - | - | - | - | - | - | - | - | - | - | - | - | - | - | - | - | - | - | - | - | - | - | - | - | - | - | - | - | - | - | - | - | - | - | - | - | - | - | - | - | - | - | - | - | - | - | - | - | - | - | E | N | L | T | P | V | - | - | - | - | - | - | - | - | - | - | - | - | - | - | - | - | - | - | - | - | - | - | - | - | - | - | V | L | - | - | - | - | - | - | - | - | - | - | - | - | - | - | - | - | - | - | - | - | - | - | - | - | - | - | - | - | - | - | - | - | - | - | - | - | - | - | - | - | - | - | - | - | - | - | - | - | - | - | - | - | - | - | - | - | - | - | - | - | - | - | - | - | - | - | - | - | - | - | - | - | - | R | P | - | - | - | - | - | - | - | - | - | - | - | - | - | - | - | - | - | - | - | - | - | - | - | - | - | - | E | I | R | K | P | - | - | - | - | - | - | E | V | L | R | P | - | E | T | - | - | - | - | - | - | - | - | - | - | - | - | - | - | - | - | - | - | P | R | P | V | D | I | G | S | G | G | F | - | - | - | - | - | - | - | - | - | - | - | - | - | G | D | V | G | - | - | - | - | - | - | - | - | - | - | A | - | P | E | S | L | R | G | N | A | D | - | - | - | - | - | L | A | Y | V | - | - | L | S | M | E | - | P | C | G | H | C | L | I | I | N | N | V | N | F | C | R | E | S | - | - | G | L | S | T | R | T | G | S | S | I | D | C | E | K | L | Q | R | R | F | - | S | S | L | H | F | M | V | E | V | K | H | D | L | T | A | K | E | M | V | L | A | L | L | E | L | A | R | R | D | H | - | S | A | L | D | C | C | V | V | V | I | L | S | H | G | C | Q | A | S | H | L | Q | F | P | G | A | - | V | Y | G | T | D | G | C | P | V | S | V | E | K | I | V | N | I | F | N | G | T | S | C | P | S | L | G | G | K | P | K | L | F | F | I | Q | A | C | - | - | G | G | D | Q | K | D | R | G | F | E | V | A | S | T | - | - | - | S | P | E | D | - | E | S | S | G | S | N | - | - | - | - | - | - | - | P | E | P | D | A | T | P | F | Q | E | D | L | R | T | F | - | - | - | - | - | - | - | - | D | Q | P | D | A | V | - | - | - | - | - | - | - | S | S | L | - | - | - | P | T | P | S | D | I | F | V | S | Y | S | T | F | P | - | - | G | - | - | - | - | - | - | - | - | F | V | S | L | R | D | P | K | S | G | S | W | Y | V | E | T | L | D | D | I | F | E | Q | W | A | R | - | S | - | - | - | - | - | - | - | - | - | - | - | - | - | - | - | - | - | - | - | - | - | - | - | - | - | - | - | - | - | - | - | E | D | L | Q | S | L | L | - | - | - | - | - | L | R | - | - | - | - | - | - | - | - | - | - | - | - | V | A | N | A | V | - | - | - | - | - | S | V | K | - | - | - | - | - | - | - | G | I | Y | - | K | Q | M | - | - | P | G | - | - | - | - | - | - | - | - | - | - | - | - | - | - | - | C | F | N | F | - | - | - | - | - | - | - | - | - | L | R | K | K | L | F | F | K | T | S | - | - | - | - | - | - | - | - | - | - | - | - | - | - | - | - | - | - | - | - | - | - | - | - | - | - | - | - | - | - | - | - | - | - | - | - | - | M | - | - | - | - | - | - | - | - | - | - | - | - | - | - | - | - | - | - | - | - | - | - | - | - | - | - | - | - | - | - | - | - | - | - | - | - | - | - | - | - | - | - | - | - | - | - | - | - | - | - | - | - | G | D | V | E | K | G | K | K | I | F | I | M | K | C | S | Q | C | H | T | V | E | K | G | G | K | H | K | T | G | P | N | L | H | G | L | F | G | R | K | T | G | Q | A | P | G | Y | S | Y | T | A | A | N | K | N | K | G | I | T | W | G | E | D | T | L | M | E | Y | L | E | N | P | K | K | Y | I | P | G | T | K | M | I | F | V | G | I | K | K | K | E | E | R | A | D | L | I | A | Y | L | K | K | A | T | N | E | - | - | - | - | - | - |
| Columba\_livia | M | - | - | - | - | - | - | - | - | - | - | - | - | - | - | - | - | - | - | - | - | - | - | - | - | - | - | - | - | - | - | - | - | - | - | - | - | - | - | - | - | - | - | - | - | - | - | - | - | - | - | - | - | - | - | - | - | - | - | - | - | - | - | - | - | - | - | - | - | - | - | - | - | - | - | - | - | - | - | - | - | - | - | - | - | - | - | - | - | - | - | - | - | - | - | - | - | - | D | V | K | S | - | R | N | - | - | - | - | - | - | - | - | - | - | - | - | - | - | - | - | Y | L | L | - | - | - | - | - | - | - | - | - | - | - | - | - | - | - | - | - | - | - | - | - | - | - | - | - | - | - | - | - | - | - | - | - | - | - | - | - | - | - | - | - | - | - | - | - | - | - | - | - | - | - | - | - | - | - | - | - | - | - | - | - | - | - | - | - | - | - | - | - | - | - | - | - | - | - | - | - | - | - | - | - | - | - | - | - | - | - | - | - | - | - | - | - | - | - | - | - | - | - | - | - | - | - | - | - | - | - | - | - | - | - | - | - | - | - | - | - | - | - | - | - | K | N | R | Q | A | L | E | K | D | I | K | T | S | Y | - | I | M | D | C | M | I | S | D | E | V | L | T | L | E | E | E | E | R | V | K | Q | Q | - | - | - | - | - | - | - | - | - | - | - | - | - | - | - | - | - | N | T | Q | K | E | Q | A | A | M | L | I | N | I | I | L | T | K | - | - | - | - | D | N | N | S | Y | R | S | - | - | - | - | - | F | Y | N | A | L | - | L | H | E | G | - | - | - | - | - | - | Y | R | D | L | A | E | L | L | Q | D | G | I | P | A | I | S | S | D | N | G | - | K | - | - | - | - | - | - | - | - | - | - | - | - | - | - | - | - | - | - | - | - | - | - | T | S | M | D | G | M | T | - | - | - | - | - | - | - | - | - | - | - | S | D | V | R | M | I | L | C | E | G | G | - | - | - | - | - | - | - | V | P | Q | R | P | V | V | F | V | T | R | P | K | L | V | D | A | I | - | - | - | - | - | - | - | - | - | - | K | Q | K | L | Y | - | - | - | - | - | - | - | - | - | - | - | - | - | - | - | - | - | - | - | - | - | - | - | - | - | - | - | - | - | - | - | - | C | L | G | N | D | P | - | - | - | - | - | - | - | - | - | - | - | - | - | - | - | - | - | - | - | - | - | - | - | - | - | - | - | - | - | - | - | - | - | - | - | - | - | - | G | W | V | T | V | Y | G | M | A | G | C | G | K | T | V | L | T | A | E | A | L | - | R | D | H | Q | L | L | K | D | Y | F | P | G | G | V | H | W | I | S | V | G | - | - | - | - | - | - | - | K | - | - | - | - | - | - | Q | D | K | A | - | - | - | - | - | - | - | - | - | - | - | - | G | L | L | I | K | L | Q | N | L | C | S | R | L | E | - | - | - | - | H | - | - | - | - | - | - | - | - | - | - | - | - | - | - | - | D | S | A | L | S | Q | R | P | P | L | N | I | - | - | - | - | - | - | - | - | - | - | - | - | - | - | - | - | E | E | A | K | D | R | L | R | L | L | M | L | R | - | - | - | - | - | - | - | - | - | - | - | - | K | Y | A | R | S | L | L | - | - | I | L | D | D | I | W | D | S | W | V | L | K | A | F | D | - | N | Q | C | Q | V | L | I | T | S | R | D | R | S | V | T | D | A | V | - | S | G | K | - | - | K | Y | E | V | H | V | E | S | - | - | - | G | L | A | H | E | K | G | L | E | I | L | S | L | F | V | N | M | - | K | I | S | E | L | P | E | - | - | - | - | - | - | - | - | - | - | - | - | - | - | Q | A | N | C | L | V | R | E | C | K | - | - | - | - | - | - | - | - | - | - | - | - | - | - | - | - | - | - | - | G | - | S | P | L | V | I | S | L | I | G | A | L | L | R | D | F | P | S | - | - | - | - | - | - | - | - | - | - | R | W | D | Y | Y | L | - | K | Q | L | Q | N | K | Q | F | K | R | I | - | - | - | - | - | - | - | - | - | - | - | - | - | - | - | - | - | - | - | - | - | - | R | K | S | S | S | Y | D | - | - | - | - | Y | E | A | L | D | E | A | M | S | I | S | V | E | - | Q | L | N | - | - | - | D | N | Y | K | D | Y | - | Y | K | D | L | S | I | L | P | K | D | V | R | V | P | T | K | V | L | C | I | L | W | D | - | - | - | - | M | E | T | E | E | V | E | D | I | L | Q | E | F | - | - | - | - | - | - | - | - | - | - | - | V | N | K | S | L | L | F | C | D | H | N | G | K | S | - | - | - | - | - | - | - | - | - | F | H | Y | Y | L | H | D | L | Q | L | D | F | L | T | E | K | - | - | N | R | N | Q | - | L | - | Q | E | L | H | K | N | I | V | N | Q | Y | K | K | - | Y | Y | - | - | - | - | - | - | K | L | D | T | P | V | P | S | - | Q | E | - | - | - | - | - | - | - | D | C | M | Y | W | Y | N | F | L | A | Y | H | M | A | G | A | N | - | - | - | - | M | Q | K | E | L | C | D | L | M | F | S | L | D | W | I | K | A | K | T | E | L | V | G | P | A | - | - | - | - | - | - | - | - | H | L | I | H | E | Y | V | E | Y | S | S | V | L | D | Q | K | - | - | - | - | - | - | - | D | S | T | V | R | E | N | F | Q | - | - | - | E | F | L | S | L | N | G | H | L | L | G | R | K | P | F | - | - | - | - | - | - | - | - | - | - | - | - | - | - | - | - | - | - | - | - | - | - | - | - | - | - | - | - | - | - | - | - | - | - | - | - | - | - | - | - | - | - | - | - | - | - | - | - | - | - | - | - | - | - | - | - | - | - | - | - | - | - | - | - | - | - | - | - | - | - | - | P | D | I | I | Q | L | G | L | C | Q | P | E | T | S | E | V | Y | Q | Q | A | K | L | R | A | Q | R | Q | - | - | T | G | - | - | - | - | - | - | - | - | - | - | - | - | - | - | - | - | - | - | - | - | - | - | - | - | - | - | - | - | - | A | F | Y | L | - | - | - | - | - | - | - | - | - | - | E | W | I | N | K | - | - | - | - | - | - | - | K | S | L | K | N | L | Y | R | - | - | - | - | - | - | - | - | - | - | L | V | V | R | P | H | R | D | - | - | - | - | - | - | - | - | - | - | - | - | - | - | - | A | V | Y | H | A | C | - | - | - | F | S | - | K | D | R | Q | R | I | A | - | - | - | - | - | - | - | - | - | S | C | G | A | D | K | T | L | Q | V | F | K | - | - | - | - | - | A | G | S | G | E | R | L | L | E | I | - | N | A | H | D | D | E | - | - | - | I | L | C | C | A | F | S | - | A | - | - | - | - | - | - | - | - | - | - | - | - | - | D | G | E | F | V | A | T | C | S | A | D | K | K | V | K | I | - | - | - | W | N | S | R | - | - | - | - | - | - | - | - | - | - | - | - | - | - | - | - | - | - | - | - | - | - | - | - | - | - | - | - | - | - | - | - | - | - | - | - | - | - | - | - | - | - | - | - | - | - | - | - | - | - | - | - | - | - | - | - | - | - | - | - | - | - | - | - | - | - | - | - | - | - | - | - | - | - | - | - | - | - | - | - | - | - | - | - | - | - | - | - | - | - | - | - | - | - | - | - | - | - | - | - | - | - | - | - | - | - | - | - | - | - | - | - | - | - | - | - | - | - | - | - | - | - | - | - | - | - | - | - | - | - | - | - | - | - | - | - | - | - | - | - | T | G | Q | C | R | C | V | Y | - | E | E | H | T | - | - | - | - | - | E | Q | V | N | - | - | - | - | - | - | - | - | - | - | - | - | - | C | C | Q | - | - | - | - | - | - | - | - | - | - | - | - | - | F | N | N | R | S | - | - | - | - | - | - | - | - | - | - | - | - | - | - | - | - | G | Q | - | Y | V | L | A | T | C | S | N | - | - | - | - | D | T | F | I | - | - | - | - | - | - | - | - | - | - | - | - | K | L | W | D | L | - | - | - | N | K | K | Y | C | R | N | T | M | M | - | - | - | - | - | - | - | - | - | - | - | - | - | - | - | - | - | G | H | V | N | S | - | - | - | - | - | - | - | - | - | - | - | - | - | - | - | - | V | N | H | C | R | F | S | P | N | D | E | - | - | - | - | - | - | - | - | - | - | - | - | - | - | - | - | - | - | - | - | - | - | - | - | - | - | - | - | Y | V | A | S | C | S | T | D | G | T | V | K | L | - | - | - | - | - | - | - | - | - | - | - | - | - | - | - | - | - | - | - | W | E | - | T | R | S | A | N | E | L | K | S | - | - | - | - | - | - | I | E | - | - | - | - | - | - | - | - | - | - | - | - | - | - | - | - | - | - | - | - | - | - | I | K | D | F | - | - | - | - | - | - | - | - | F | K | N | A | D | E | Q | P | D | - | - | - | - | - | - | - | - | - | - | - | - | - | - | - | - | - | - | - | - | - | - | - | - | - | - | - | D | V | E | - | I | L | V | K | C | C | - | - | - | - | - | - | - | - | - | - | - | - | - | - | - | - | S | W | S | G | N | G | D | M | I | L | V | V | A | K | N | K | L | - | - | - | - | - | - | - | - | - | - | - | L | L | F | D | V | K | T | - | - | - | C | V | - | - | - | - | - | - | - | - | - | - | - | - | - | - | - | - | - | - | - | - | - | - | - | - | - | - | - | - | - | - | - | - | - | - | - | - | - | - | - | - | - | - | - | - | - | - | - | - | - | - | - | - | - | - | - | - | - | - | - | - | - | - | - | - | - | - | - | - | - | - | - | - | - | - | - | - | - | - | - | - | - | - | - | - | - | - | - | - | - | - | - | - | - | - | - | - | - | - | - | - | - | - | - | - | - | - | - | - | - | - | - | - | - | - | - | - | - | - | - | - | - | - | - | - | - | - | L | L | T | - | - | - | - | - | - | - | - | - | - | - | - | - | - | - | - | - | - | - | - | - | - | - | - | - | - | - | - | - | - | - | - | - | - | - | - | - | - | - | - | - | - | - | - | - | - | - | - | - | - | - | - | - | - | - | - | - | - | - | - | - | - | - | - | - | - | - | - | - | - | - | - | - | - | - | - | - | - | - | - | - | - | - | - | - | - | - | - | - | - | - | - | - | - | - | - | - | - | - | - | - | - | - | - | - | - | - | - | - | - | - | - | - | - | - | - | - | - | - | - | - | - | - | - | - | - | - | - | - | - | - | - | - | - | - | - | - | - | - | - | - | - | - | - | - | - | - | - | - | - | - | - | - | - | - | - | - | - | - | - | - | - | - | - | - | - | - | - | - | - | - | - | - | - | - | - | - | - | - | - | - | - | - | Q | V | - | - | - | I | E | S | H | H | S | T | I | Q | Y | C | - | - | - | - | - | - | - | - | - | - | - | - | - | - | - | - | - | - | - | - | - | - | - | - | - | D | F | C | P | G | D | E | L | V | A | V | A | L | S | H | - | - | - | - | - | - | C | S | V | E | - | - | - | - | - | - | - | - | - | - | - | - | - | - | - | - | - | L | W | N | I | K | S | L | - | - | - | - | - | - | - | - | - | - | - | - | - | - | - | - | - | - | - | - | - | - | - | - | - | - | - | - | - | S | K | V | A | Y | C | R | G | H | M | S | - | - | - | - | - | - | - | - | - | - | - | - | - | - | - | - | - | - | - | - | - | - | - | - | - | - | - | - | - | - | - | - | - | - | - | - | - | - | - | - | - | - | - | - | W | V | H | C | V | T | F | S | - | - | - | - | - | - | - | - | - | - | - | P | D | G | S | - | - | - | - | - | - | - | - | - | - | L | F | L | T | S | S | D | D | Q | - | - | - | - | - | - | - | - | - | - | - | - | - | - | T | V | R | I | - | W | - | - | - | - | - | - | - | - | - | - | - | - | - | - | - | - | - | - | - | - | - | - | - | - | - | - | - | - | - | - | - | - | - | - | - | - | - | - | E | T | K | K | V | C | K | S | S | D | A | V | L | K | S | E | - | - | - | - | - | - | - | - | - | - | - | - | - | - | - | - | L | D | V | V | F | - | - | - | - | - | - | - | - | - | - | - | - | - | - | - | - | - | - | - | - | - | - | - | - | - | - | - | - | - | - | - | - | - | - | - | - | - | - | - | - | - | - | - | - | - | - | - | - | - | - | - | - | - | - | Y | N | G | E | V | M | - | - | - | - | - | - | - | - | - | - | - | - | - | - | - | - | - | - | - | - | - | - | - | - | - | - | - | - | - | - | - | - | - | - | - | - | I | L | A | I | Y | N | Q | K | H | L | Q | L | - | I | N | G | N | T | - | - | - | - | - | - | - | - | - | - | D | S | V | I | M | Q | T | A | A | - | - | - | - | - | - | - | - | - | - | - | - | - | - | - | - | - | - | - | - | - | - | Q | E | S | - | - | - | - | - | - | - | - | P | I | S | C | C | C | L | S | E | D | L | Q | - | - | - | - | - | - | - | - | - | F | A | A | F | G | Q | - | - | - | - | - | - | - | - | - | - | - | - | - | - | - | - | - | - | - | - | - | - | - | - | - | - | - | - | - | - | - | - | - | - | - | - | E | - | - | - | - | - | - | - | - | - | S | G | T | - | - | - | - | - | - | - | - | - | - | I | - | - | - | - | - | - | - | - | - | - | - | - | - | - | - | - | - | - | - | - | - | - | - | - | - | - | - | - | - | - | - | - | - | - | - | - | - | - | - | K | V | L | Q | L | L | D | G | K | V | L | K | S | R | E | A | - | - | - | - | - | Y | - | - | - | - | - | - | - | - | - | - | - | - | R | T | S | V | Q | H | C | R | F | T | - | - | - | - | - | - | - | - | - | - | - | - | - | - | - | - | - | - | - | - | - | - | - | - | - | - | - | - | - | - | - | - | - | - | - | - | - | S | D | C | Q | T | L | I | S | S | A | - | H | D | S | - | - | - | - | - | - | - | - | I | I | Q | - | V | - | - | - | - | - | - | - | - | - | - | - | - | - | - | - | - | - | - | - | - | - | - | - | - | - | - | - | - | - | - | - | - | - | - | - | - | - | - | - | W | N | - | - | - | - | - | - | - | - | - | - | W | Q | S | S | E | C | V | F | - | - | - | - | - | - | - | - | - | - | - | - | - | - | - | - | - | - | - | - | - | - | - | - | - | - | - | - | - | - | - | - | - | - | - | - | - | - | - | - | - | - | - | - | - | - | - | - | - | - | L | R | G | H | K | - | - | - | - | - | - | - | - | - | - | - | - | - | - | - | - | - | - | - | - | - | - | - | - | - | - | E | A | V | K | D | F | R | L | L | - | - | - | - | - | - | - | - | - | - | - | - | - | - | - | - | - | - | - | - | - | - | - | E | N | - | S | E | L | L | - | S | W | - | - | - | S | F | D | G | T | V | - | - | - | - | - | - | - | - | - | - | - | - | - | - | - | - | - | - | - | - | - | - | - | - | - | - | - | - | - | - | - | - | - | - | - | K | V | W | N | L | T | T | G | N | T | - | - | - | - | - | E | K | D | F | A | C | - | - | - | - | - | - | - | H | G | - | - | - | - | - | - | - | - | - | - | - | - | - | - | - | - | - | - | - | D | A | V | L | - | - | - | - | - | - | - | - | - | - | - | - | - | - | - | S | C | A | - | - | - | - | - | - | - | - | - | - | - | - | - | - | - | V | S | P | D | G | S | - | K | F | S | S | T | - | S | A | D | K | T | - | - | - | - | - | - | A | K | I | W | S | F | E | S | S | - | - | - | - | - | - | - | - | - | - | - | - | - | - | - | - | - | - | - | - | - | - | - | - | S | V | L | H | K | - | - | - | - | - | - | - | - | L | - | - | - | - | K | G | H | E | A | C | V | - | - | - | - | - | - | - | - | - | R | C | C | - | T | F | S | - | - | - | - | - | - | - | - | - | P | N | N | K | - | L | L | A | T | - | - | - | - | - | - | - | - | G | D | D | K | G | E | I | R | I | W | D | I | L | T | G | A | L | L | - | - | - | - | - | - | - | - | - | - | - | - | - | - | - | - | - | - | - | - | - | - | - | - | - | - | - | - | - | - | - | - | - | - | - | - | - | - | - | - | - | - | - | - | - | - | - | - | - | - | - | - | - | - | - | - | H | F | C | S | P | V | T | A | D | - | - | - | - | - | - | - | - | - | - | - | - | - | - | - | - | - | - | - | - | E | - | E | P | T | H | G | G | W | - | - | - | - | - | - | - | - | V | T | D | L | S | F | S | P | D | S | K | - | - | - | - | - | - | - | - | - | - | - | - | - | - | - | - | - | - | - | - | - | - | - | - | - | - | - | - | - | - | - | - | - | - | - | - | - | - | - | - | - | - | - | - | - | M | L | V | S | - | - | S | G | G | F | L | K | - | - | - | - | - | - | - | - | W | W | N | V | T | T | - | - | - | - | - | - | - | - | - | - | - | - | - | - | - | - | - | - | - | - | - | - | - | - | - | - | - | - | - | - | - | - | - | - | - | - | - | - | - | - | - | - | - | - | - | - | - | - | - | - | - | - | - | - | - | - | - | - | - | - | - | - | - | - | - | - | - | - | - | - | - | - | - | - | - | - | - | - | - | - | - | - | - | - | - | - | - | - | - | - | - | - | - | - | - | - | - | - | - | - | - | - | - | - | - | - | - | - | - | - | - | - | - | - | - | - | - | - | - | - | - | - | - | - | - | - | - | - | - | - | - | - | - | - | - | - | - | - | - | - | - | - | - | - | - | - | - | - | - | - | - | - | - | - | - | - | - | - | - | - | - | - | - | - | - | - | - | - | - | - | - | - | - | - | - | - | - | - | - | - | - | - | - | - | - | - | - | - | - | - | - | - | - | - | - | - | - | - | - | - | - | - | - | - | - | - | - | - | - | - | - | - | - | - | - | - | - | - | - | - | - | - | - | - | - | - | - | - | - | - | - | - | - | - | - | - | - | - | - | - | - | - | - | - | - | - | - | - | - | - | - | - | - | - | - | - | - | - | - | - | - | - | - | - | - | - | - | G | E | S | L | Q | T | F | Y | T | - | - | - | - | - | N | G | - | - | - | T | N | L | K | S | I | - | - | - | H | V | S | P | - | - | - | - | N | F | R | M | - | - | - | - | - | - | - | - | - | - | - | - | - | - | - | - | - | - | - | - | - | - | - | - | - | - | Y | V | T | V | D | - | N | L | G | I | L | - | - | Y | V | L | Q | K | - | - | - | - | - | - | - | - | - | - | - | - | - | - | - | - | - | - | - | - | - | - | - | - | - | - | - | - | - | - | - | - | - | - | L | - | - | - | - | - | - | - | - | - | - | - | - | - | - | - | - | - | - | - | - | - | - | - | - | - | - | - | - | - | - | - | - | - | - | - | - | - | - | - | - | - | - | - | - | - | - | - | - | - | - | - | - | - | M | - | - | - | - | - | - | - | - | - | - | - | - | - | - | - | - | - | - | - | - | - | - | - | - | - | - | - | - | - | - | - | - | - | - | - | - | - | - | - | - | - | - | - | - | - | - | - | - | - | - | - | - | - | - | - | - | - | - | - | - | - | - | - | - | - | - | - | - | - | - | - | - | - | - | - | - | - | - | - | - | - | - | - | - | - | - | - | - | - | - | - | - | - | - | - | - | - | - | - | - | - | - | - | - | - | - | - | - | - | - | - | - | - | - | - | - | - | - | - | - | - | - | - | - | - | - | - | - | - | - | - | - | - | - | - | - | - | - | - | - | - | - | - | - | - | - | - | - | - | - | - | - | - | - | - | - | - | - | - | - | - | - | - | - | - | - | - | - | - | - | - | - | - | - | - | - | - | - | - | - | - | - | - | - | - | - | - | - | - | - | - | - | - | - | - | - | - | - | - | - | - | - | - | - | - | - | - | - | - | - | - | - | - | - | - | - | - | - | - | - | - | - | - | - | - | - | - | - | - | - | - | - | - | - | - | - | - | - | - | - | - | - | - | - | - | - | - | - | - | - | - | - | - | - | - | - | - | - | - | - | - | - | - | - | - | - | - | - | - | - | - | - | - | - | - | - | - | - | - | - | - | - | - | - | - | - | - | - | - | - | - | - | - | - | - | - | - | - | - | - | - | - | - | - | L | M | Q | E | C | G | H | L | - | - | - | - | P | A | P | P | - | Q | - | - | - | - | - | - | - | - | - | - | - | - | - | - | - | - | - | - | - | - | - | - | - | - | - | - | - | - | - | L | G | D | - | - | - | - | - | - | - | - | - | - | - | - | - | - | - | L | R | P | V | - | - | - | - | - | - | - | - | - | - | - | - | - | - | - | - | - | - | - | - | - | - | - | - | - | - | - | - | - | - | - | - | - | - | - | - | - | - | - | E | L | D | L | R | G | E | K | H | S | K | N | V | - | - | - | - | - | - | - | - | - | - | - | S | - | - | - | - | - | - | - | - | - | - | - | - | - | - | - | - | - | - | - | - | - | - | - | - | - | - | - | - | - | - | - | - | - | - | - | - | - | - | - | - | - | - | - | - | - | - | - | - | - | - | - | - | - | - | - | - | - | - | - | - | - | - | - | - | - | - | - | - | - | - | - | - | - | - | - | - | - | - | - | - | - | - | - | - | - | - | - | - | - | - | - | - | - | - | - | - | - | - | - | - | - | - | - | - | - | - | - | - | - | - | - | - | - | - | - | - | - | - | - | - | - | - | - | - | - | - | - | - | - | - | - | - | - | - | - | - | - | - | - | - | - | - | - | - | - | - | - | - | - | - | - | - | - | - | - | - | F | P | G | R | L | S | - | - | - | - | - | - | - | - | - | - | - | - | - | - | - | - | - | - | - | - | - | - | - | - | - | - | V | P | I | Q | A | - | - | - | - | - | - | E | S | E | R | P | - | Q | M | - | - | - | - | - | - | - | - | - | - | - | - | - | - | - | - | - | - | P | P | M | P | A | Q | G | - | - | - | - | - | - | - | - | - | - | - | - | - | - | - | - | - | - | - | - | S | - | - | - | - | - | - | - | - | - | - | A | - | V | D | - | - | K | R | N | R | D | - | - | - | - | - | L | V | Y | E | - | - | L | K | S | D | - | P | C | G | R | C | L | I | L | N | N | V | N | F | S | R | D | S | - | - | D | L | S | T | R | D | G | S | D | V | D | C | E | K | L | E | K | R | F | - | K | A | L | S | F | D | V | L | T | R | R | D | L | P | A | Q | K | M | V | S | E | L | Q | E | L | A | R | Q | D | H | - | S | A | L | D | C | C | V | V | V | I | L | S | H | G | C | Q | T | S | H | I | Q | F | P | G | G | - | V | Y | G | T | D | G | K | A | I | P | V | E | K | I | V | N | Y | F | N | G | S | H | C | P | S | L | R | G | K | P | K | L | F | F | I | Q | A | C | - | - | G | G | E | Q | R | D | R | G | F | V | V | D | G | D | - | - | - | S | P | G | D | - | E | A | P | G | G | S | - | - | - | - | - | - | - | L | E | S | D | T | T | P | F | Q | T | P | A | G | N | V | - | - | - | - | - | - | - | - | D | Q | P | D | A | V | - | - | - | - | - | - | - | A | S | L | - | - | - | P | T | P | S | D | I | L | V | S | Y | S | T | F | P | - | - | G | - | - | - | - | - | - | - | - | F | V | S | W | R | D | A | S | S | G | S | W | Y | V | E | T | L | D | S | V | L | E | Q | Y | A | R | - | S | - | - | - | - | - | - | - | - | - | - | - | - | - | - | - | - | - | - | - | - | - | - | - | - | - | - | - | - | - | - | - | E | D | L | L | N | M | L | - | - | - | - | - | L | R | - | - | - | - | - | - | - | - | - | - | - | - | V | A | N | A | V | - | - | - | - | - | S | A | K | - | - | - | - | - | - | - | G | R | Y | - | K | Q | I | - | - | P | G | - | - | - | - | - | - | - | - | - | - | - | - | - | - | - | C | F | N | F | - | - | - | - | - | - | - | - | - | L | R | K | K | F | F | F | R | C | K | - | - | - | - | - | - | - | - | - | - | - | - | - | - | - | - | - | - | - | - | - | - | - | - | - | - | - | - | - | - | - | - | - | - | - | - | - | M | - | - | - | - | - | - | - | - | - | - | - | - | - | - | - | - | - | - | - | - | - | - | - | - | - | - | - | - | - | - | - | - | - | - | - | - | - | - | - | - | - | - | - | - | - | - | - | - | - | - | - | - | G | D | I | E | K | G | K | K | I | F | V | Q | K | C | S | Q | C | H | T | V | E | K | G | G | K | H | K | T | G | P | N | L | H | G | L | F | G | R | K | T | G | Q | A | E | G | F | S | Y | T | D | A | N | K | N | K | G | I | T | W | G | E | D | T | L | M | E | Y | L | E | N | P | K | K | Y | I | P | G | T | K | M | I | F | A | G | I | K | K | K | A | E | R | A | D | L | I | A | Y | L | K | Q | A | T | A | K | - | - | - | - | - | - |
| Condylura\_cristata | M | - | - | - | - | - | - | - | - | - | - | - | - | - | - | - | - | - | - | - | - | - | - | - | - | - | - | - | - | - | - | - | - | - | - | - | - | - | - | - | - | - | - | - | - | - | - | - | - | - | - | - | - | - | - | - | - | - | - | - | - | - | - | - | - | - | - | - | - | - | - | - | - | - | - | - | - | - | - | - | - | - | - | - | - | - | - | - | - | - | - | - | - | - | - | - | - | - | D | A | K | A | - | R | N | - | - | - | - | - | - | - | - | - | - | - | - | - | - | - | - | C | L | L | - | - | - | - | - | - | - | - | - | - | - | - | - | - | - | - | - | - | - | - | - | - | - | - | - | - | - | - | - | - | - | - | - | - | - | - | - | - | - | - | - | - | - | - | - | - | - | - | - | - | - | - | - | - | - | - | - | - | - | - | - | - | - | - | - | - | - | - | - | - | - | - | - | - | - | - | - | - | - | - | - | - | - | - | - | - | - | - | - | - | - | - | - | - | - | - | - | - | - | - | - | - | - | - | - | - | - | - | - | - | - | - | - | - | - | - | - | - | - | - | - | Q | H | R | E | A | L | E | Q | D | I | K | T | S | Y | - | I | M | D | H | M | I | S | D | G | V | L | T | V | S | E | E | E | K | V | K | N | E | - | - | - | - | - | - | - | - | - | - | - | - | - | - | - | - | - | P | T | Q | Q | Q | R | A | A | M | L | I | K | T | I | L | K | K | - | - | - | - | D | N | H | S | Y | I | S | - | - | - | - | - | F | Y | N | A | L | - | L | H | E | G | - | - | - | - | - | - | Y | K | D | L | A | A | L | L | H | S | G | V | P | A | V | S | S | S | N | G | - | K | - | - | - | - | - | - | - | - | - | - | - | - | - | - | - | - | - | - | - | - | - | - | D | S | A | G | G | I | T | - | - | - | - | - | - | - | - | - | - | - | S | Y | V | R | T | V | L | C | E | G | G | - | - | - | - | - | - | - | V | P | Q | R | P | V | V | F | V | T | R | K | K | L | V | N | A | I | - | - | - | - | - | - | - | - | - | - | Q | Q | K | L | Y | - | - | - | - | - | - | - | - | - | - | - | - | - | - | - | - | - | - | - | - | - | - | - | - | - | - | - | - | - | - | - | - | K | L | N | G | E | P | - | - | - | - | - | - | - | - | - | - | - | - | - | - | - | - | - | - | - | - | - | - | - | - | - | - | - | - | - | - | - | - | - | - | - | - | - | - | G | W | V | T | I | Y | G | M | A | G | C | G | K | S | V | L | A | A | E | A | V | - | R | D | H | S | L | L | Q | S | C | F | P | G | G | V | N | W | V | S | V | G | - | - | - | - | - | - | - | K | - | - | - | - | - | - | Q | D | K | S | - | - | - | - | - | - | - | - | - | - | - | - | G | L | L | M | K | L | Q | N | L | C | T | R | L | D | - | - | - | - | Q | - | - | - | - | - | - | - | - | - | - | - | - | - | - | - | D | E | S | F | S | Q | R | L | P | L | N | I | - | - | - | - | - | - | - | - | - | - | - | - | - | - | - | - | E | E | A | K | D | R | L | R | I | L | M | L | R | - | - | - | - | - | - | - | - | - | - | - | - | K | H | P | R | S | L | L | - | - | I | L | D | D | V | W | D | P | W | V | L | K | A | F | D | - | N | Q | C | Q | I | L | L | T | T | R | D | K | S | V | T | D | S | V | - | M | G | P | - | - | K | Y | V | V | P | V | E | S | - | - | - | S | L | G | K | E | K | G | L | E | I | L | S | L | F | V | N | M | - | K | K | A | D | L | P | E | - | - | - | - | - | - | - | - | - | - | - | - | - | - | Q | A | H | S | I | I | K | E | C | K | - | - | - | - | - | - | - | - | - | - | - | - | - | - | - | - | - | - | - | G | - | S | P | L | V | V | S | L | I | G | A | L | L | R | D | F | P | N | - | - | - | - | - | - | - | - | - | - | R | W | E | Y | Y | L | - | R | Q | L | Q | N | K | Q | F | K | R | I | - | - | - | - | - | - | - | - | - | - | - | - | - | - | - | - | - | - | - | - | - | - | R | K | S | S | F | Y | D | - | - | - | - | Y | E | A | L | D | E | A | M | S | I | S | V | E | - | M | L | R | - | - | - | E | D | I | K | G | Y | - | Y | T | D | L | S | I | L | Q | K | D | V | K | V | P | T | K | V | L | C | I | L | W | D | - | - | - | - | M | E | T | E | E | V | E | D | I | L | Q | E | F | - | - | - | - | - | - | - | - | - | - | - | V | N | K | S | L | L | F | C | D | R | N | G | K | S | - | - | - | - | - | - | - | - | - | F | H | Y | Y | L | H | D | L | Q | V | D | F | L | T | E | K | - | - | N | R | N | Q | - | L | - | Q | D | L | H | K | K | M | I | T | Q | F | Q | R | - | H | H | - | - | - | - | - | - | Q | P | H | T | L | S | P | D | - | Q | E | - | - | - | - | - | - | - | D | C | M | Y | W | Y | N | F | L | A | Y | H | M | A | N | A | N | - | - | - | - | M | H | K | E | L | C | A | L | M | F | S | L | D | W | I | K | A | K | T | E | L | V | G | P | A | - | - | - | - | - | - | - | - | H | L | I | H | E | F | V | E | Y | S | H | I | L | D | E | K | - | - | - | - | - | - | - | C | S | A | V | C | E | N | F | Q | - | - | - | E | F | L | S | L | N | G | H | L | L | G | R | Q | P | F | - | - | - | - | - | - | - | - | - | - | - | - | - | - | - | - | - | - | - | - | - | - | - | - | - | - | - | - | - | - | - | - | - | - | - | - | - | - | - | - | - | - | - | - | - | - | - | - | - | - | - | - | - | - | - | - | - | - | - | - | - | - | - | - | - | - | - | - | - | - | - | P | N | I | V | Q | L | G | L | C | E | P | E | T | S | E | V | Y | R | Q | A | K | L | Q | A | K | Q | E | V | D | N | G | - | - | - | - | - | - | - | - | - | - | - | - | - | - | - | - | - | - | - | - | - | - | - | - | - | - | - | - | - | M | L | Y | L | - | - | - | - | - | - | - | - | - | - | E | W | I | N | K | - | - | - | - | - | - | - | K | N | I | K | N | L | S | R | - | - | - | - | - | - | - | - | - | - | L | V | V | R | P | H | T | D | - | - | - | - | - | - | - | - | - | - | - | - | - | - | - | A | V | Y | H | A | C | - | - | - | F | S | - | E | D | G | Q | R | I | A | - | - | - | - | - | - | - | - | - | S | C | G | A | D | K | T | L | Q | V | F | K | - | - | - | - | - | A | E | T | G | E | K | L | L | E | I | - | K | A | H | E | D | E | - | - | - | V | L | C | C | A | F | S | - | A | - | - | - | - | - | - | - | - | - | - | - | - | - | D | N | R | F | I | A | T | C | S | V | D | R | K | V | K | I | - | - | - | W | N | S | L | - | - | - | - | - | - | - | - | - | - | - | - | - | - | - | - | - | - | - | - | - | - | - | - | - | - | - | - | - | - | - | - | - | - | - | - | - | - | - | - | - | - | - | - | - | - | - | - | - | - | - | - | - | - | - | - | - | - | - | - | - | - | - | - | - | - | - | - | - | - | - | - | - | - | - | - | - | - | - | - | - | - | - | - | - | - | - | - | - | - | - | - | - | - | - | - | - | - | - | - | - | - | - | - | - | - | - | - | - | - | - | - | - | - | - | - | - | - | - | - | - | - | - | - | - | - | - | - | - | - | - | - | - | - | - | - | - | - | - | - | T | G | E | L | V | H | T | Y | - | E | E | H | S | - | - | - | - | - | E | Q | V | N | - | - | - | - | - | - | - | - | - | - | - | - | - | C | C | H | - | - | - | - | - | - | - | - | - | - | - | - | - | F | T | N | S | S | - | - | - | - | - | - | - | - | - | - | - | - | - | - | - | - | H | H | - | I | L | L | A | T | G | S | S | - | - | - | - | D | S | C | L | - | - | - | - | - | - | - | - | - | - | - | - | K | L | W | D | L | - | - | - | N | Q | E | S | C | R | N | T | L | F | - | - | - | - | - | - | - | - | - | - | - | - | - | - | - | - | - | D | H | R | N | A | - | - | - | - | - | - | - | - | - | - | - | - | - | - | - | - | V | T | H | C | R | F | S | P | D | D | K | - | - | - | - | - | - | - | - | - | - | - | - | - | - | - | - | - | - | - | - | - | - | - | - | - | - | - | - | F | L | A | S | C | S | A | D | G | T | L | K | L | - | - | - | - | - | - | - | - | - | - | - | - | - | - | - | - | - | - | - | W | D | - | V | K | S | A | N | L | R | K | S | - | - | - | - | - | - | I | D | - | - | - | - | - | - | - | - | - | - | - | - | - | - | - | - | - | - | - | - | - | - | V | K | Q | F | - | - | - | - | - | - | - | - | F | L | N | S | E | E | P | Q | E | - | - | - | - | - | - | - | - | - | - | - | - | - | - | - | - | - | - | - | - | - | - | - | - | - | - | - | D | M | E | - | V | I | V | K | C | C | - | - | - | - | - | - | - | - | - | - | - | - | - | - | - | - | S | W | S | A | D | G | A | R | I | M | V | A | A | K | N | K | I | - | - | - | - | - | - | - | - | - | - | - | F | L | F | D | I | H | T | - | - | - | R | N | - | - | - | - | - | - | - | - | - | - | - | - | - | - | - | - | - | - | - | - | - | - | - | - | - | - | - | - | - | - | - | - | - | - | - | - | - | - | - | - | - | - | - | - | - | - | - | - | - | - | - | - | - | - | - | - | - | - | - | - | - | - | - | - | - | - | - | - | - | - | - | - | - | - | - | - | - | - | - | - | - | - | - | - | - | - | - | - | - | - | - | - | - | - | - | - | - | - | - | - | - | - | - | - | - | - | - | - | - | - | - | - | - | - | - | - | - | - | - | - | - | - | - | - | - | - | L | L | A | - | - | - | - | - | - | - | - | - | - | - | - | - | - | - | - | - | - | - | - | - | - | - | - | - | - | - | - | - | - | - | - | - | - | - | - | - | - | - | - | - | - | - | - | - | - | - | - | - | - | - | - | - | - | - | - | - | - | - | - | - | - | - | - | - | - | - | - | - | - | - | - | - | - | - | - | - | - | - | - | - | - | - | - | - | - | - | - | - | - | - | - | - | - | - | - | - | - | - | - | - | - | - | - | - | - | - | - | - | - | - | - | - | - | - | - | - | - | - | - | - | - | - | - | - | - | - | - | - | - | - | - | - | - | - | - | - | - | - | - | - | - | - | - | - | - | - | - | - | - | - | - | - | - | - | - | - | - | - | - | - | - | - | - | - | - | - | - | - | - | - | - | - | - | - | - | - | - | - | - | - | - | - | E | I | - | - | - | H | T | G | H | Q | S | T | I | Q | Y | C | - | - | - | - | - | - | - | - | - | - | - | - | - | - | - | - | - | - | - | - | - | - | - | - | - | D | F | S | P | H | N | H | L | V | I | V | A | L | S | Q | - | - | - | - | - | - | Y | C | V | E | - | - | - | - | - | - | - | - | - | - | - | - | - | - | - | - | - | L | W | N | M | D | S | C | - | - | - | - | - | - | - | - | - | - | - | - | - | - | - | - | - | - | - | - | - | - | - | - | - | - | - | - | - | L | K | V | A | D | C | R | G | H | L | S | - | - | - | - | - | - | - | - | - | - | - | - | - | - | - | - | - | - | - | - | - | - | - | - | - | - | - | - | - | - | - | - | - | - | - | - | - | - | - | - | - | - | - | - | W | V | H | C | V | M | F | S | - | - | - | - | - | - | - | - | - | - | - | P | D | G | S | - | - | - | - | - | - | - | - | - | - | S | F | L | T | S | S | D | D | Q | - | - | - | - | - | - | - | - | - | - | - | - | - | - | T | I | R | L | - | W | - | - | - | - | - | - | - | - | - | - | - | - | - | - | - | - | - | - | - | - | - | - | - | - | - | - | - | - | - | - | - | - | - | - | - | - | - | - | E | T | R | K | V | C | Q | N | S | A | T | V | L | K | Q | E | - | - | - | - | - | - | - | - | - | - | - | - | - | - | - | - | I | D | V | M | F | - | - | - | - | - | - | - | - | - | - | - | - | - | - | - | - | - | - | - | - | - | - | - | - | - | - | - | - | - | - | - | - | - | - | - | - | - | - | - | - | - | - | - | - | - | - | - | - | - | - | - | - | - | - | Q | E | N | E | V | M | - | - | - | - | - | - | - | - | - | - | - | - | - | - | - | - | - | - | - | - | - | - | - | - | - | - | - | - | - | - | - | - | - | - | - | - | V | L | A | A | D | N | L | R | H | L | Q | L | - | I | N | G | K | T | - | - | - | - | - | - | - | - | - | - | G | Q | T | D | H | L | T | E | A | - | - | - | - | - | - | - | - | - | - | - | - | - | - | - | - | - | - | - | - | - | - | Q | - | - | - | - | - | - | - | - | - | - | - | V | S | C | C | C | L | S | P | H | L | Q | - | - | - | - | - | - | - | - | - | Y | V | A | L | G | D | - | - | - | - | - | - | - | - | - | - | - | - | - | - | - | - | - | - | - | - | - | - | - | - | - | - | - | - | - | - | - | - | - | - | - | - | E | - | - | - | - | - | - | - | - | - | E | G | A | - | - | - | - | - | - | - | - | - | - | I | - | - | - | - | - | - | - | - | - | - | - | - | - | - | - | - | - | - | - | - | - | - | - | - | - | - | - | - | - | - | - | - | - | - | - | - | - | - | - | K | I | L | E | P | L | N | N | R | T | F | Q | S | R | I | G | - | - | - | - | - | H | - | - | - | - | - | - | - | - | - | - | - | - | K | S | T | V | R | H | M | Q | F | T | - | - | - | - | - | - | - | - | - | - | - | - | - | - | - | - | - | - | - | - | - | - | - | - | - | - | - | - | - | - | - | - | - | - | - | - | - | A | D | G | K | T | L | I | S | S | A | - | D | D | P | - | - | - | - | - | - | - | - | V | I | Q | - | V | - | - | - | - | - | - | - | - | - | - | - | - | - | - | - | - | - | - | - | - | - | - | - | - | - | - | - | - | - | - | - | - | - | - | - | - | - | - | - | W | N | - | - | - | - | - | - | - | - | - | - | W | Q | S | E | D | Y | V | F | - | - | - | - | - | - | - | - | - | - | - | - | - | - | - | - | - | - | - | - | - | - | - | - | - | - | - | - | - | - | - | - | - | - | - | - | - | - | - | - | - | - | - | - | - | - | - | - | - | - | L | Q | A | H | Q | - | - | - | - | - | - | - | - | - | - | - | - | - | - | - | - | - | - | - | - | - | - | - | - | - | - | E | A | V | K | D | F | R | L | L | - | - | - | - | - | - | - | - | - | - | - | - | - | - | - | - | - | - | - | - | - | - | - | K | N | - | S | S | L | L | - | S | W | - | - | - | S | F | D | G | T | V | - | - | - | - | - | - | - | - | - | - | - | - | - | - | - | - | - | - | - | - | - | - | - | - | - | - | - | - | - | - | - | - | - | - | - | K | V | W | N | I | I | N | G | R | I | - | - | - | - | - | E | K | D | F | V | C | - | - | - | - | - | - | - | H | Q | - | - | - | - | - | - | - | - | - | - | - | - | - | - | - | - | - | - | - | D | T | V | L | - | - | - | - | - | - | - | - | - | - | - | - | - | - | - | S | C | D | - | - | - | - | - | - | - | - | - | - | - | - | - | - | - | I | S | P | D | A | T | - | K | F | S | S | T | - | S | A | D | K | T | - | - | - | - | - | - | A | K | I | W | N | F | E | D | F | - | - | - | - | - | - | - | - | - | - | - | - | - | - | - | - | - | - | - | - | - | - | - | - | S | L | L | H | E | - | - | - | - | - | - | - | - | L | - | - | - | - | K | D | H | K | G | C | V | - | - | - | - | - | - | - | - | - | R | C | S | - | V | F | S | - | - | - | - | - | - | - | - | - | V | D | S | T | - | L | L | A | T | - | - | - | - | - | - | - | - | G | D | D | N | G | E | V | R | I | W | N | V | S | N | G | K | L | L | - | - | - | - | - | - | - | - | - | - | - | - | - | - | - | - | - | - | - | - | - | - | - | - | - | - | - | - | - | - | - | - | - | - | - | - | - | - | - | - | - | - | - | - | - | - | - | - | - | - | - | - | - | - | - | - | H | L | C | A | P | I | S | I | E | - | - | - | - | - | - | - | - | - | - | - | - | - | - | - | - | - | - | - | - | G | G | A | A | T | H | G | G | W | - | - | - | - | - | - | - | - | V | T | D | L | R | F | S | P | D | N | K | - | - | - | - | - | - | - | - | - | - | - | - | - | - | - | - | - | - | - | - | - | - | - | - | - | - | - | - | - | - | - | - | - | - | - | - | - | - | - | - | - | - | - | - | - | M | L | V | S | - | - | A | G | G | Y | L | K | - | - | - | - | - | - | - | - | W | W | N | V | V | T | - | - | - | - | - | - | - | - | - | - | - | - | - | - | - | - | - | - | - | - | - | - | - | - | - | - | - | - | - | - | - | - | - | - | - | - | - | - | - | - | - | - | - | - | - | - | - | - | - | - | - | - | - | - | - | - | - | - | - | - | - | - | - | - | - | - | - | - | - | - | - | - | - | - | - | - | - | - | - | - | - | - | - | - | - | - | - | - | - | - | - | - | - | - | - | - | - | - | - | - | - | - | - | - | - | - | - | - | - | - | - | - | - | - | - | - | - | - | - | - | - | - | - | - | - | - | - | - | - | - | - | - | - | - | - | - | - | - | - | - | - | - | - | - | - | - | - | - | - | - | - | - | - | - | - | - | - | - | - | - | - | - | - | - | - | - | - | - | - | - | - | - | - | - | - | - | - | - | - | - | - | - | - | - | - | - | - | - | - | - | - | - | - | - | - | - | - | - | - | - | - | - | - | - | - | - | - | - | - | - | - | - | - | - | - | - | - | - | - | - | - | - | - | - | - | - | - | - | - | - | - | - | - | - | - | - | - | - | - | - | - | - | - | - | - | - | - | - | - | - | - | - | - | - | - | - | - | - | - | - | - | - | - | - | - | - | - | G | E | S | S | Q | T | F | Y | T | - | - | - | - | - | N | G | - | - | - | T | N | L | K | K | I | - | - | - | H | V | S | P | - | - | - | - | D | F | K | T | - | - | - | - | - | - | - | - | - | - | - | - | - | - | - | - | - | - | - | - | - | - | - | - | - | - | C | V | T | V | D | - | N | L | G | I | L | - | - | Y | I | L | Q | I | - | - | - | - | - | - | - | - | - | - | - | - | - | - | - | - | - | - | - | - | - | - | - | - | - | - | - | - | - | - | - | - | - | - | L | E | - | - | - | - | - | - | - | - | - | - | - | - | - | - | - | - | - | - | - | - | - | - | - | - | - | - | - | - | - | - | - | - | - | - | - | - | - | - | - | - | - | - | - | - | - | - | - | - | - | - | - | - | M | - | - | - | - | - | - | - | - | - | - | - | - | - | - | - | - | - | - | - | - | - | - | - | - | - | - | - | - | - | - | - | - | - | - | - | - | - | - | - | - | - | - | - | - | - | - | - | - | - | - | - | - | - | - | D | E | A | H | R | Q | - | - | - | - | - | - | - | - | - | - | - | - | - | - | - | - | - | - | - | - | V | L | R | R | - | - | - | - | - | - | - | - | - | - | - | - | - | - | - | - | - | - | - | - | - | - | - | - | - | - | - | - | - | - | C | R | - | - | - | - | - | - | - | - | L | Q | L | - | V | R | E | L | Q | V | D | L | L | W | - | - | - | - | - | - | - | - | - | - | - | - | - | - | - | - | - | - | - | - | - | - | - | - | - | - | - | - | - | - | - | - | - | - | - | - | - | - | - | - | - | - | - | - | - | D | H | L | L | D | R | D | L | - | - | - | - | - | - | - | - | - | - | - | - | - | - | - | - | - | - | - | - | - | - | - | - | F | T | R | D | M | I | E | D | - | - | - | - | - | - | - | - | - | - | - | - | - | - | - | - | - | - | - | - | - | - | - | - | - | I | Q | R | A | G | - | - | - | - | - | - | - | - | S | G | S | R | R | D | Q | - | A | R | Q | L | I | T | D | L | E | T | R | G | - | S | R | A | L | P | L | F | I | S | C | L | E | D | A | G | Q | D | A | L | A | S | L | M | R | T | S | C | Q | T | A | - | - | - | - | K | Q | D | L | - | E | V | I | - | - | - | - | - | - | - | - | - | - | - | - | - | - | - | - | - | - | - | - | - | - | - | - | - | - | R | P | L | D | - | - | - | - | - | - | - | - | - | - | - | - | - | - | - | I | E | P | A | - | - | - | - | - | - | - | - | - | - | - | - | - | - | - | - | - | - | - | - | - | - | - | - | - | - | - | - | - | - | - | - | - | - | - | - | - | - | - | V | - | - | - | - | - | - | - | - | - | - | - | - | - | - | - | - | - | - | - | - | - | - | - | - | - | - | - | - | - | - | - | - | - | - | - | - | - | - | - | - | - | - | - | - | - | - | - | - | - | - | - | - | - | - | - | - | - | - | - | - | - | - | - | - | - | - | - | - | - | - | - | - | - | - | - | - | - | - | - | - | - | - | - | - | - | - | - | - | - | - | - | - | - | - | - | - | - | - | - | - | - | - | - | - | - | - | - | - | - | - | - | - | - | - | - | - | - | - | - | - | - | - | - | - | - | - | - | - | - | - | - | - | - | - | - | - | - | - | - | - | - | - | - | - | - | - | - | - | - | - | - | - | - | - | - | - | - | - | - | - | - | - | - | - | - | - | - | - | - | - | - | - | - | - | - | - | - | - | - | - | - | - | - | - | - | - | - | - | - | - | - | - | - | - | - | - | - | - | - | - | - | - | - | - | - | - | - | - | - | - | - | - | - | - | L | S | P | - | - | - | - | - | - | R | D | L | R | P | - | V | V | - | - | - | - | - | - | - | - | - | - | - | - | - | - | - | - | - | - | S | R | P | I | D | I | S | P | P | G | F | - | - | - | - | - | - | - | - | - | - | - | - | - | S | N | V | R | - | - | - | - | - | - | - | - | - | - | T | - | L | E | T | W | K | G | N | A | D | - | - | - | - | - | L | A | Y | I | - | - | L | K | A | D | - | P | C | G | H | C | L | I | I | N | N | V | N | F | H | P | E | S | - | - | K | L | S | T | R | K | G | S | N | I | D | C | E | K | L | Q | R | R | F | - | G | L | L | H | F | M | V | E | V | K | S | D | L | T | A | K | Q | I | D | R | A | L | M | Q | L | A | K | Q | D | H | - | S | A | L | D | C | C | V | V | V | I | L | S | H | G | C | Q | A | S | H | R | Q | F | P | G | A | - | V | Y | G | A | D | G | C | P | V | S | V | E | R | I | V | N | I | L | N | G | A | G | C | P | S | L | G | G | K | P | K | L | F | F | I | Q | A | C | - | - | G | G | E | Q | K | D | H | G | F | S | V | A | S | V | - | - | - | P | H | K | D | - | R | A | P | E | D | D | - | - | - | - | - | - | - | S | E | S | D | A | A | P | F | Q | D | G | L | G | P | F | - | - | - | - | - | - | - | - | D | Q | T | D | A | V | - | - | - | - | - | - | - | A | S | L | - | - | - | P | T | P | S | D | I | F | V | S | Y | S | T | F | P | - | - | G | - | - | - | - | - | - | - | - | F | V | S | W | R | D | T | N | S | G | S | W | Y | V | E | T | L | D | R | V | L | E | Q | W | A | H | - | S | - | - | - | - | - | - | - | - | - | - | - | - | - | - | - | - | - | - | - | - | - | - | - | - | - | - | - | - | - | - | - | E | D | L | Q | T | L | L | - | - | - | - | - | L | R | - | - | - | - | - | - | - | - | - | - | - | - | V | A | D | A | V | - | - | - | - | - | S | E | K | - | - | - | - | - | - | - | G | V | F | - | K | Q | M | - | - | P | G | - | - | - | - | - | - | - | - | - | - | - | - | - | - | - | C | F | N | F | - | - | - | - | - | - | - | - | - | L | R | K | K | L | F | F | K | T | - | - | - | - | - | - | - | - | - | - | - | - | - | - | - | - | - | - | - | - | - | - | - | - | - | - | - | - | - | - | - | - | - | - | - | - | - | - | M | - | - | - | - | - | - | - | - | - | - | - | - | - | - | - | - | - | - | - | - | - | - | - | - | - | - | - | - | - | - | - | - | - | - | - | - | - | - | - | - | - | - | - | - | - | - | - | - | - | - | - | - | S | D | I | E | K | G | K | K | I | F | V | Q | K | C | A | Q | C | H | T | V | E | K | G | G | K | H | K | T | G | P | N | L | H | G | L | F | G | R | K | T | G | Q | A | P | G | F | S | Y | T | D | A | N | K | N | K | G | I | T | W | K | E | D | T | L | M | E | Y | L | E | N | P | K | K | Y | I | P | G | T | K | M | I | F | A | G | I | K | K | K | S | E | R | A | E | L | I | A | Y | L | K | K | A | T | N | E | - | - | - | - | - | - |
| Corvus\_cornix\_cornix | M | - | - | - | - | - | - | - | - | - | - | - | - | - | - | - | - | - | - | - | - | - | - | - | - | - | - | - | - | - | - | - | - | - | - | - | - | - | - | - | - | - | - | - | - | - | - | - | - | - | - | - | - | - | - | - | - | - | - | - | - | - | - | - | - | - | - | - | - | - | - | - | - | - | - | - | - | - | - | - | - | - | - | - | - | - | - | - | - | - | - | - | - | - | - | - | - | - | D | V | R | S | - | R | N | - | - | - | - | - | - | - | - | - | - | - | - | - | - | - | - | Y | L | L | - | - | - | - | - | - | - | - | - | - | - | - | - | - | - | - | - | - | - | - | - | - | - | - | - | - | - | - | - | - | - | - | - | - | - | - | - | - | - | - | - | - | - | - | - | - | - | - | - | - | - | - | - | - | - | - | - | - | - | - | - | - | - | - | - | - | - | - | - | - | - | - | - | - | - | - | - | - | - | - | - | - | - | - | - | - | - | - | - | - | - | - | - | - | - | - | - | - | - | - | - | - | - | - | - | - | - | - | - | - | - | - | - | - | - | - | - | - | - | - | - | M | N | R | Q | A | L | E | N | D | I | K | T | S | Y | - | I | M | D | R | M | I | S | D | E | V | L | T | L | Q | E | E | E | R | V | R | Q | Q | - | - | - | - | - | - | - | - | - | - | - | - | - | - | - | - | - | N | T | R | K | E | R | A | A | M | L | I | N | I | I | L | T | K | - | - | - | - | D | N | N | S | Y | R | S | - | - | - | - | - | F | Y | N | A | L | - | L | H | E | G | - | - | - | - | - | - | Y | R | D | L | A | A | L | L | Q | D | G | I | P | A | I | S | S | G | N | R | - | K | - | - | - | - | - | - | - | - | - | - | - | - | - | - | - | - | - | - | - | - | - | - | S | S | M | D | G | M | T | - | - | - | - | - | - | - | - | - | - | - | S | H | V | K | T | I | L | C | E | G | G | - | - | - | - | - | - | - | V | P | Q | R | P | V | V | F | V | T | R | P | K | L | V | D | A | I | - | - | - | - | - | - | - | - | - | - | K | K | K | L | Y | - | - | - | - | - | - | - | - | - | - | - | - | - | - | - | - | - | - | - | - | - | - | - | - | - | - | - | - | - | - | - | - | C | L | G | S | E | P | - | - | - | - | - | - | - | - | - | - | - | - | - | - | - | - | - | - | - | - | - | - | - | - | - | - | - | - | - | - | - | - | - | - | - | - | - | - | G | W | V | T | V | Y | G | M | A | G | C | G | K | T | V | L | T | A | E | A | L | - | R | D | P | Q | L | L | E | D | Y | F | P | G | G | V | H | W | I | S | V | G | - | - | - | - | - | - | - | K | - | - | - | - | - | - | Q | D | K | A | - | - | - | - | - | - | - | - | - | - | - | - | G | L | L | I | K | L | Q | N | L | C | S | R | L | E | - | - | - | - | H | - | - | - | - | - | - | - | - | - | - | - | - | - | - | - | D | S | T | L | P | Q | R | - | P | L | N | I | - | - | - | - | - | - | - | - | - | - | - | - | - | - | - | - | E | E | A | K | D | R | L | R | L | L | M | L | R | - | - | - | - | - | - | - | - | - | - | - | - | K | Y | P | R | S | L | L | - | - | V | L | D | D | I | W | D | S | W | V | L | K | A | F | D | - | N | Q | C | Q | V | L | I | T | S | R | D | R | S | V | T | D | A | V | - | A | G | N | - | - | K | Y | E | V | H | V | E | S | - | - | - | G | L | A | H | E | K | G | L | E | I | L | S | L | F | V | N | M | - | K | I | S | E | L | P | E | - | - | - | - | - | - | - | - | - | - | - | - | - | - | Q | A | N | S | L | V | R | E | C | K | - | - | - | - | - | - | - | - | - | - | - | - | - | - | - | - | - | - | - | G | - | S | P | L | V | I | S | L | I | G | A | L | L | R | D | F | P | N | - | - | - | - | - | - | - | - | - | - | R | W | E | Y | Y | L | - | R | Q | L | Q | N | K | Q | F | K | R | I | - | - | - | - | - | - | - | - | - | - | - | - | - | - | - | - | - | - | - | - | - | - | R | K | S | S | S | Y | D | - | - | - | - | Y | E | A | L | D | E | A | M | S | I | S | V | E | - | Q | L | D | - | - | - | D | N | Y | K | D | Y | - | Y | K | D | L | S | I | L | P | K | D | V | K | V | P | T | K | V | L | C | I | L | W | D | - | - | - | - | M | E | T | E | E | V | E | D | I | L | Q | E | F | - | - | - | - | - | - | - | - | - | - | - | V | N | K | S | L | L | F | C | D | R | N | G | K | S | - | - | - | - | - | - | - | - | - | F | H | Y | Y | L | H | D | L | Q | L | D | F | L | T | E | K | - | - | N | R | N | Q | - | L | - | Q | E | L | H | K | N | I | V | N | Q | Y | K | K | - | Y | Y | - | - | - | - | - | - | K | L | N | T | P | V | L | S | - | Q | E | - | - | - | - | - | - | - | D | C | M | Y | W | Y | N | F | L | A | Y | H | M | A | G | A | N | - | - | - | - | M | Q | Q | E | L | R | D | L | M | F | S | L | D | W | I | K | A | K | T | E | L | V | G | P | A | - | - | - | - | - | - | - | - | H | L | I | H | E | Y | V | E | Y | S | S | I | L | D | Q | K | - | - | - | - | - | - | - | D | S | T | V | R | E | N | F | Q | - | - | - | E | F | L | S | L | N | G | H | L | L | G | R | Q | P | F | - | - | - | - | - | - | - | - | - | - | - | - | - | - | - | - | - | - | - | - | - | - | - | - | - | - | - | - | - | - | - | - | - | - | - | - | - | - | - | - | - | - | - | - | - | - | - | - | - | - | - | - | - | - | - | - | - | - | - | - | - | - | - | - | - | - | - | - | - | - | - | P | D | I | I | Q | L | G | L | C | Q | P | E | T | S | E | V | Y | Q | Q | A | K | L | H | A | Q | R | E | - | - | T | G | - | - | - | - | - | - | - | - | - | - | - | - | - | - | - | - | - | - | - | - | - | - | - | - | - | - | - | - | - | A | F | Y | L | - | - | - | - | - | - | - | - | - | - | E | W | I | N | K | - | - | - | - | - | - | - | K | S | L | K | N | L | Y | R | - | - | - | - | - | - | - | - | - | - | L | V | V | R | P | H | R | D | - | - | - | - | - | - | - | - | - | - | - | - | - | - | - | A | V | Y | H | A | C | - | - | - | F | S | - | K | D | R | Q | R | I | A | - | - | - | - | - | - | - | - | - | S | C | G | A | D | K | T | L | Q | V | F | K | - | - | - | - | - | A | E | S | G | E | R | L | L | E | I | - | N | A | H | D | D | E | - | - | - | I | L | C | C | A | F | S | - | A | - | - | - | - | - | - | - | - | - | - | - | - | - | D | G | E | F | V | A | T | C | S | S | D | K | K | V | K | V | - | - | - | W | N | S | R | - | - | - | - | - | - | - | - | - | - | - | - | - | - | - | - | - | - | - | - | - | - | - | - | - | - | - | - | - | - | - | - | - | - | - | - | - | - | - | - | - | - | - | - | - | - | - | - | - | - | - | - | - | - | - | - | - | - | - | - | - | - | - | - | - | - | - | - | - | - | - | - | - | - | - | - | - | - | - | - | - | - | - | - | - | - | - | - | - | - | - | - | - | - | - | - | - | - | - | - | - | - | - | - | - | - | - | - | - | - | - | - | - | - | - | - | - | - | - | - | - | - | - | - | - | - | - | - | - | - | - | - | - | - | - | - | - | - | - | - | T | G | Q | C | R | C | V | Y | - | E | E | H | S | - | - | - | - | - | E | Q | V | N | - | - | - | - | - | - | - | - | - | - | - | - | - | C | C | Q | - | - | - | - | - | - | - | - | - | - | - | - | - | F | N | N | R | S | - | - | - | - | - | - | - | - | - | - | - | - | - | - | - | - | G | Q | - | Y | L | L | A | T | C | S | N | - | - | - | - | D | T | F | I | - | - | - | - | - | - | - | - | - | - | - | - | K | V | W | D | L | - | - | - | N | K | K | Y | C | R | N | T | M | I | - | - | - | - | - | - | - | - | - | - | - | - | - | - | - | - | - | G | H | E | N | S | - | - | - | - | - | - | - | - | - | - | - | - | - | - | - | - | V | N | H | C | R | F | S | P | N | D | E | - | - | - | - | - | - | - | - | - | - | - | - | - | - | - | - | - | - | - | - | - | - | - | - | - | - | - | - | Y | V | A | S | C | S | T | D | G | T | V | K | L | - | - | - | - | - | - | - | - | - | - | - | - | - | - | - | - | - | - | - | W | E | - | A | R | S | G | N | E | L | K | S | - | - | - | - | - | - | I | E | - | - | - | - | - | - | - | - | - | - | - | - | - | - | - | - | - | - | - | - | - | - | I | K | D | F | - | - | - | - | - | - | - | - | F | K | N | A | D | E | Q | P | D | - | - | - | - | - | - | - | - | - | - | - | - | - | - | - | - | - | - | - | - | - | - | - | - | - | - | - | D | V | E | - | V | L | V | K | C | C | - | - | - | - | - | - | - | - | - | - | - | - | - | - | - | - | S | W | S | R | N | G | D | M | I | L | V | G | A | K | N | K | L | - | - | - | - | - | - | - | - | - | - | - | L | L | F | D | I | K | T | - | - | - | C | G | - | - | - | - | - | - | - | - | - | - | - | - | - | - | - | - | - | - | - | - | - | - | - | - | - | - | - | - | - | - | - | - | - | - | - | - | - | - | - | - | - | - | - | - | - | - | - | - | - | - | - | - | - | - | - | - | - | - | - | - | - | - | - | - | - | - | - | - | - | - | - | - | - | - | - | - | - | - | - | - | - | - | - | - | - | - | - | - | - | - | - | - | - | - | - | - | - | - | - | - | - | - | - | - | - | - | - | - | - | - | - | - | - | - | - | - | - | - | - | - | - | - | - | - | - | - | L | L | T | - | - | - | - | - | - | - | - | - | - | - | - | - | - | - | - | - | - | - | - | - | - | - | - | - | - | - | - | - | - | - | - | - | - | - | - | - | - | - | - | - | - | - | - | - | - | - | - | - | - | - | - | - | - | - | - | - | - | - | - | - | - | - | - | - | - | - | - | - | - | - | - | - | - | - | - | - | - | - | - | - | - | - | - | - | - | - | - | - | - | - | - | - | - | - | - | - | - | - | - | - | - | - | - | - | - | - | - | - | - | - | - | - | - | - | - | - | - | - | - | - | - | - | - | - | - | - | - | - | - | - | - | - | - | - | - | - | - | - | - | - | - | - | - | - | - | - | - | - | - | - | - | - | - | - | - | - | - | - | - | - | - | - | - | - | - | - | - | - | - | - | - | - | - | - | - | - | - | - | - | - | - | - | Q | V | - | - | - | I | V | S | H | H | S | T | I | Q | Y | C | - | - | - | - | - | - | - | - | - | - | - | - | - | - | - | - | - | - | - | - | - | - | - | - | - | D | F | C | P | G | D | E | L | V | A | V | A | L | S | H | - | - | - | - | - | - | C | S | V | E | - | - | - | - | - | - | - | - | - | - | - | - | - | - | - | - | - | L | W | N | I | K | S | L | - | - | - | - | - | - | - | - | - | - | - | - | - | - | - | - | - | - | - | - | - | - | - | - | - | - | - | - | - | S | K | V | A | Y | C | R | G | H | M | S | - | - | - | - | - | - | - | - | - | - | - | - | - | - | - | - | - | - | - | - | - | - | - | - | - | - | - | - | - | - | - | - | - | - | - | - | - | - | - | - | - | - | - | - | W | V | H | C | V | T | F | S | - | - | - | - | - | - | - | - | - | - | - | P | D | G | S | - | - | - | - | - | - | - | - | - | - | L | F | L | T | S | S | D | D | Q | - | - | - | - | - | - | - | - | - | - | - | - | - | - | T | I | R | I | - | W | - | - | - | - | - | - | - | - | - | - | - | - | - | - | - | - | - | - | - | - | - | - | - | - | - | - | - | - | - | - | - | - | - | - | - | - | - | - | E | T | K | K | V | C | K | S | S | D | A | V | L | K | S | E | - | - | - | - | - | - | - | - | - | - | - | - | - | - | - | - | L | D | V | V | F | - | - | - | - | - | - | - | - | - | - | - | - | - | - | - | - | - | - | - | - | - | - | - | - | - | - | - | - | - | - | - | - | - | - | - | - | - | - | - | - | - | - | - | - | - | - | - | - | - | - | - | - | - | - | H | N | G | E | V | M | - | - | - | - | - | - | - | - | - | - | - | - | - | - | - | - | - | - | - | - | - | - | - | - | - | - | - | - | - | - | - | - | - | - | - | - | I | L | A | I | Y | N | Q | K | H | L | Q | L | - | I | N | G | N | T | - | - | - | - | - | - | - | - | - | - | D | N | V | I | M | Q | T | A | A | - | - | - | - | - | - | - | - | - | - | - | - | - | - | - | - | - | - | - | - | - | - | Q | E | S | - | - | - | - | - | - | - | - | P | I | S | C | C | C | L | S | E | D | L | K | - | - | - | - | - | - | - | - | - | F | A | A | F | G | Q | - | - | - | - | - | - | - | - | - | - | - | - | - | - | - | - | - | - | - | - | - | - | - | - | - | - | - | - | - | - | - | - | - | - | - | - | E | - | - | - | - | - | - | - | - | - | S | G | T | - | - | - | - | - | - | - | - | - | - | I | - | - | - | - | - | - | - | - | - | - | - | - | - | - | - | - | - | - | - | - | - | - | - | - | - | - | - | - | - | - | - | - | - | - | - | - | - | - | - | K | V | L | Q | L | S | D | G | K | V | L | K | S | W | E | A | - | - | - | - | - | C | - | - | - | - | - | - | - | - | - | - | - | - | R | T | S | V | Q | H | C | R | F | T | - | - | - | - | - | - | - | - | - | - | - | - | - | - | - | - | - | - | - | - | - | - | - | - | - | - | - | - | - | - | - | - | - | - | - | - | - | P | D | C | Q | T | L | I | S | S | A | - | H | D | S | - | - | - | - | - | - | - | - | V | I | Q | - | V | - | - | - | - | - | - | - | - | - | - | - | - | - | - | - | - | - | - | - | - | - | - | - | - | - | - | - | - | - | - | - | - | - | - | - | - | - | - | - | W | N | - | - | - | - | - | - | - | - | - | - | W | Q | L | S | E | C | A | L | - | - | - | - | - | - | - | - | - | - | - | - | - | - | - | - | - | - | - | - | - | - | - | - | - | - | - | - | - | - | - | - | - | - | - | - | - | - | - | - | - | - | - | - | - | - | - | - | - | - | L | R | G | H | K | - | - | - | - | - | - | - | - | - | - | - | - | - | - | - | - | - | - | - | - | - | - | - | - | - | - | E | A | I | K | D | F | R | L | L | - | - | - | - | - | - | - | - | - | - | - | - | - | - | - | - | - | - | - | - | - | - | - | E | N | - | S | K | L | L | - | S | W | - | - | - | S | F | D | G | T | V | - | - | - | - | - | - | - | - | - | - | - | - | - | - | - | - | - | - | - | - | - | - | - | - | - | - | - | - | - | - | - | - | - | - | - | K | V | W | N | I | T | T | G | N | P | - | - | - | - | - | E | K | D | F | A | C | - | - | - | - | - | - | - | H | G | - | - | - | - | - | - | - | - | - | - | - | - | - | - | - | - | - | - | - | G | A | V | L | - | - | - | - | - | - | - | - | - | - | - | - | - | - | - | S | C | A | - | - | - | - | - | - | - | - | - | - | - | - | - | - | - | V | S | P | D | G | S | - | K | F | S | S | T | - | S | A | D | K | T | - | - | - | - | - | - | A | K | I | W | S | F | E | S | S | - | - | - | - | - | - | - | - | - | - | - | - | - | - | - | - | - | - | - | - | - | - | - | - | S | V | L | H | E | - | - | - | - | - | - | - | - | L | - | - | - | - | K | D | H | E | A | C | V | - | - | - | - | - | - | - | - | - | R | C | C | - | T | F | S | - | - | - | - | - | - | - | - | - | P | N | N | K | - | L | L | A | T | - | - | - | - | - | - | - | - | G | D | D | K | G | E | I | R | I | W | D | I | L | T | G | A | L | L | - | - | - | - | - | - | - | - | - | - | - | - | - | - | - | - | - | - | - | - | - | - | - | - | - | - | - | - | - | - | - | - | - | - | - | - | - | - | - | - | - | - | - | - | - | - | - | - | - | - | - | - | - | - | - | - | H | F | C | S | P | V | T | V | D | - | - | - | - | - | - | - | - | - | - | - | - | - | - | - | - | - | - | - | - | E | G | E | P | T | H | G | G | W | - | - | - | - | - | - | - | - | V | T | D | L | S | F | S | P | D | S | K | - | - | - | - | - | - | - | - | - | - | - | - | - | - | - | - | - | - | - | - | - | - | - | - | - | - | - | - | - | - | - | - | - | - | - | - | - | - | - | - | - | - | - | - | - | M | L | V | S | - | - | S | G | G | Y | L | K | - | - | - | - | - | - | - | - | W | W | N | V | T | T | - | - | - | - | - | - | - | - | - | - | - | - | - | - | - | - | - | - | - | - | - | - | - | - | - | - | - | - | - | - | - | - | - | - | - | - | - | - | - | - | - | - | - | - | - | - | - | - | - | - | - | - | - | - | - | - | - | - | - | - | - | - | - | - | - | - | - | - | - | - | - | - | - | - | - | - | - | - | - | - | - | - | - | - | - | - | - | - | - | - | - | - | - | - | - | - | - | - | - | - | - | - | - | - | - | - | - | - | - | - | - | - | - | - | - | - | - | - | - | - | - | - | - | - | - | - | - | - | - | - | - | - | - | - | - | - | - | - | - | - | - | - | - | - | - | - | - | - | - | - | - | - | - | - | - | - | - | - | - | - | - | - | - | - | - | - | - | - | - | - | - | - | - | - | - | - | - | - | - | - | - | - | - | - | - | - | - | - | - | - | - | - | - | - | - | - | - | - | - | - | - | - | - | - | - | - | - | - | - | - | - | - | - | - | - | - | - | - | - | - | - | - | - | - | - | - | - | - | - | - | - | - | - | - | - | - | - | - | - | - | - | - | - | - | - | - | - | - | - | - | - | - | - | - | - | - | - | - | - | - | - | - | - | - | - | - | - | G | E | S | L | Q | T | F | Y | T | - | - | - | - | - | N | G | - | - | - | T | N | L | K | S | I | - | - | - | H | V | S | P | - | - | - | - | D | F | K | V | - | - | - | - | - | - | - | - | - | - | - | - | - | - | - | - | - | - | - | - | - | - | - | - | - | - | Y | V | T | V | D | - | N | L | G | I | L | - | - | Y | V | L | Q | K | - | - | - | - | - | - | - | - | - | - | - | - | - | - | - | - | - | - | - | - | - | - | - | - | - | - | - | - | - | - | - | - | - | - | F | - | - | - | - | - | - | - | - | - | - | - | - | - | - | - | - | - | - | - | - | - | - | - | - | - | - | - | - | - | - | - | - | - | - | - | - | - | - | - | - | - | - | - | - | - | - | - | - | - | - | - | - | - | M | - | - | - | - | - | - | - | - | - | - | - | - | - | - | - | - | - | - | - | - | - | - | - | - | - | - | - | - | - | - | - | - | - | - | - | - | - | - | - | - | - | - | - | - | - | - | - | - | - | - | - | - | - | - | - | - | - | - | - | - | - | - | - | - | - | - | - | - | - | - | - | - | - | - | - | - | - | - | - | - | - | - | - | - | - | - | - | - | - | - | - | - | - | - | - | - | - | - | - | - | - | - | - | - | - | - | - | - | - | - | - | - | - | - | - | - | - | - | - | - | - | - | - | - | - | - | - | - | - | - | - | - | - | - | - | - | - | - | - | - | - | - | - | - | - | - | - | - | - | - | - | - | - | - | - | - | - | - | - | - | - | - | - | - | - | - | - | - | - | - | - | - | - | - | - | - | - | - | - | - | - | - | - | - | - | - | - | - | - | - | - | - | - | - | - | - | - | - | - | - | - | - | - | - | - | - | - | - | - | - | - | - | - | - | - | - | - | - | - | - | - | - | - | - | - | - | - | - | - | - | - | - | - | - | - | - | - | - | - | - | - | - | - | - | - | - | - | - | - | - | - | - | - | - | - | - | - | - | - | - | - | - | - | - | - | - | - | - | - | - | - | - | - | - | - | - | - | - | - | - | - | - | - | - | - | - | - | - | - | - | - | - | - | - | - | - | - | - | - | - | - | - | - | - | L | I | Q | E | C | E | S | R | - | - | - | - | P | V | L | P | - | Q | - | - | - | - | - | - | - | - | - | - | - | - | - | - | - | - | - | - | - | - | - | - | - | - | - | - | - | - | - | L | P | D | - | - | - | - | - | - | - | - | - | - | - | - | - | - | - | L | R | P | V | - | - | - | - | - | - | - | - | - | - | - | - | - | - | - | - | - | - | - | - | - | - | - | - | - | - | - | - | - | - | - | - | - | - | - | - | - | - | - | E | L | E | L | R | E | E | K | Q | R | K | N | V | - | - | - | - | - | - | - | - | - | - | - | T | - | - | - | - | - | - | - | - | - | - | - | - | - | - | - | - | - | - | - | - | - | - | - | - | - | - | - | - | - | - | - | - | - | - | - | - | - | - | - | - | - | - | - | - | - | - | - | - | - | - | - | - | - | - | - | - | - | - | - | - | - | - | - | - | - | - | - | - | - | - | - | - | - | - | - | - | - | - | - | - | - | - | - | - | - | - | - | - | - | - | - | - | - | - | - | - | - | - | - | - | - | - | - | - | - | - | - | - | - | - | - | - | - | - | - | - | - | - | - | - | - | - | - | - | - | - | - | - | - | - | - | - | - | - | - | - | - | - | - | - | - | - | - | - | - | - | - | - | - | - | - | - | - | - | - | - | P | H | E | H | L | S | - | - | - | - | - | - | - | - | - | - | - | - | - | - | - | - | - | - | - | - | - | - | - | - | - | - | I | S | V | Q | A | - | - | - | - | - | - | E | S | E | R | P | - | R | I | - | - | - | - | - | - | - | - | - | - | - | - | - | - | - | - | - | - | H | P | V | P | A | R | G | - | - | - | - | - | - | - | - | - | - | - | - | - | - | - | - | - | - | - | - | S | - | - | - | - | - | - | - | - | - | - | A | - | V | D | - | - | K | R | S | R | D | - | - | - | - | - | L | V | Y | E | - | - | M | K | T | D | - | P | C | G | H | C | L | I | L | N | N | V | N | F | S | R | D | S | - | - | G | L | S | T | R | E | G | S | D | I | D | C | E | K | L | E | R | R | F | - | K | A | L | H | F | T | V | L | T | R | R | D | L | K | A | Q | E | I | V | L | E | L | Q | K | L | S | R | Q | D | H | - | S | A | L | D | C | C | I | V | V | I | L | S | H | G | C | Q | T | S | H | I | Q | F | P | G | G | - | I | Y | G | T | D | G | K | P | I | P | I | E | K | I | V | N | Y | F | N | G | S | N | S | P | S | L | R | G | K | P | K | L | F | F | I | Q | A | C | - | - | G | G | E | Q | R | D | Q | G | F | V | V | D | C | D | - | - | - | S | P | E | E | - | E | A | P | G | G | S | - | - | - | - | - | - | - | L | E | S | D | A | T | P | F | R | V | P | S | D | N | V | - | - | - | - | - | - | - | - | D | E | P | D | A | I | - | - | - | - | - | - | - | A | C | L | - | - | - | P | T | P | S | D | I | L | V | S | Y | S | T | F | P | - | - | G | - | - | - | - | - | - | - | - | F | V | S | W | R | E | K | S | S | G | S | W | Y | V | E | M | L | D | S | V | L | E | Q | Y | A | H | - | S | - | - | - | - | - | - | - | - | - | - | - | - | - | - | - | - | - | - | - | - | - | - | - | - | - | - | - | - | - | - | - | E | D | L | L | G | M | L | - | - | - | - | - | V | R | - | - | - | - | - | - | - | - | - | - | - | - | V | A | G | A | V | - | - | - | - | - | S | A | K | - | - | - | - | - | - | - | G | R | Y | - | K | Q | M | - | - | P | G | - | - | - | - | - | - | - | - | - | - | - | - | - | - | - | C | F | N | F | - | - | - | - | - | - | - | - | - | L | R | K | K | F | F | F | A | C | D | - | - | - | - | - | - | - | - | - | - | - | - | - | - | - | - | - | - | - | - | - | - | - | - | - | - | - | - | - | - | - | - | - | - | - | - | - | M | - | - | - | - | - | - | - | - | - | - | - | - | - | - | - | - | - | - | - | - | - | - | - | - | - | - | - | - | - | - | - | - | - | - | - | - | - | - | - | - | - | - | - | - | - | - | - | - | - | - | - | - | G | D | V | E | K | G | K | K | I | F | V | Q | K | C | S | Q | C | H | T | V | E | K | G | G | K | H | K | T | G | P | N | L | N | G | L | F | G | R | K | T | G | Q | A | E | G | F | S | Y | T | D | A | N | K | N | K | G | I | T | W | S | E | D | T | L | M | E | Y | L | E | N | P | K | K | Y | I | P | G | T | K | M | I | F | A | G | I | K | K | K | G | E | R | A | D | L | I | A | Y | L | K | V | A | T | A | K | - | - | - | - | - | - |
| Coturnix\_japonica | M | - | - | - | - | - | - | - | - | - | - | - | - | - | - | - | - | - | - | - | - | - | - | - | - | - | - | - | - | - | - | - | - | - | - | - | - | - | - | - | - | - | - | - | - | - | - | - | - | - | - | - | - | - | - | - | - | - | - | - | - | - | - | - | - | - | - | - | - | - | - | - | - | - | - | - | - | - | - | - | - | - | - | - | - | - | - | - | - | - | - | - | - | - | - | - | - | - | D | A | K | S | - | R | M | - | - | - | - | - | - | - | - | - | - | - | - | - | - | - | - | C | L | I | - | - | - | - | - | - | - | - | - | - | - | - | - | - | - | - | - | - | - | - | - | - | - | - | - | - | - | - | - | - | - | - | - | - | - | - | - | - | - | - | - | - | - | - | - | - | - | - | - | - | - | - | - | - | - | - | - | - | - | - | - | - | - | - | - | - | - | - | - | - | - | - | - | - | - | - | - | - | - | - | - | - | - | - | - | - | - | - | - | - | - | - | - | - | - | - | - | - | - | - | - | - | - | - | - | - | - | - | - | - | - | - | - | - | - | - | - | - | - | - | - | M | N | R | Q | A | L | E | K | D | I | K | T | S | Y | - | I | M | D | C | M | V | A | D | Q | V | L | T | L | Q | E | E | E | K | V | K | Q | Q | - | - | - | - | - | - | - | - | - | - | - | - | - | - | - | - | - | N | T | Q | K | E | R | A | A | M | L | I | N | T | L | L | T | K | - | - | - | - | D | N | N | A | Y | R | S | - | - | - | - | - | F | Y | N | A | L | - | L | H | E | G | - | - | - | - | - | - | Y | R | D | L | A | A | L | L | Q | D | G | I | P | V | F | S | S | R | N | G | - | K | - | - | - | - | - | - | - | - | - | - | - | - | - | - | - | - | - | - | - | - | - | - | N | A | M | D | E | M | T | - | - | - | - | - | - | - | - | - | - | - | S | Y | V | K | T | T | L | C | E | G | G | - | - | - | - | - | - | - | V | P | Q | R | P | V | V | F | V | T | R | P | K | L | V | E | A | I | - | - | - | - | - | - | - | - | - | - | K | Q | K | L | C | - | - | - | - | - | - | - | - | - | - | - | - | - | - | - | - | - | - | - | - | - | - | - | - | - | - | - | - | - | - | - | - | S | L | G | N | D | P | - | - | - | - | - | - | - | - | - | - | - | - | - | - | - | - | - | - | - | - | - | - | - | - | - | - | - | - | - | - | - | - | - | - | - | - | - | - | G | W | V | T | V | Y | G | M | A | G | C | G | K | T | V | L | T | A | E | A | L | - | R | D | H | Q | L | L | E | G | Y | F | P | G | G | V | H | W | I | S | V | G | - | - | - | - | - | - | - | K | - | - | - | - | - | - | Q | D | K | A | - | - | - | - | - | - | - | - | - | - | - | - | G | L | L | I | K | L | Q | N | L | C | S | R | L | E | - | - | - | - | Q | - | - | - | - | - | - | - | - | - | - | - | - | - | - | - | D | S | T | L | S | Q | R | P | P | L | N | I | - | - | - | - | - | - | - | - | - | - | - | - | - | - | - | - | E | E | A | K | D | R | L | R | L | L | M | L | R | - | - | - | - | - | - | - | - | - | - | - | - | K | Y | P | R | S | L | L | - | - | V | L | D | D | I | W | D | S | W | V | L | K | A | F | D | - | N | Q | C | Q | V | L | I | T | S | R | D | R | S | V | T | D | A | V | - | S | G | N | - | - | K | Y | E | I | H | V | E | S | - | - | - | G | L | T | H | E | K | G | L | E | V | L | A | L | F | V | N | M | - | K | I | S | E | L | P | E | - | - | - | - | - | - | - | - | - | - | - | - | - | - | Q | A | N | C | I | I | R | E | C | K | - | - | - | - | - | - | - | - | - | - | - | - | - | - | - | - | - | - | - | G | - | S | P | L | V | I | S | L | I | G | A | L | L | R | D | F | P | S | - | - | - | - | - | - | - | - | - | - | R | W | E | Y | Y | L | - | K | Q | L | Q | N | K | Q | F | K | R | I | - | - | - | - | - | - | - | - | - | - | - | - | - | - | - | - | - | - | - | - | - | - | R | K | S | S | S | Y | D | - | - | - | - | Y | E | A | L | D | E | A | M | S | I | S | V | E | - | Q | L | S | - | - | - | N | D | L | K | D | Y | - | Y | K | D | L | S | I | L | P | K | D | V | K | V | P | T | K | V | L | C | I | L | W | D | - | - | - | - | M | E | T | E | E | V | E | D | I | L | Q | E | F | - | - | - | - | - | - | - | - | - | - | - | V | N | K | S | L | L | F | C | D | R | N | G | N | S | - | - | - | - | - | - | - | - | - | F | H | Y | Y | L | H | D | L | Q | L | D | F | L | T | E | Q | - | - | N | R | N | Q | - | L | - | Q | E | L | H | K | N | I | V | N | Q | Y | K | K | - | Y | Y | - | - | - | - | - | - | K | L | H | V | P | V | L | S | - | Q | E | - | - | - | - | - | - | - | D | C | M | Y | W | Y | N | F | L | A | Y | H | M | A | G | A | N | - | - | - | - | M | Q | K | E | L | R | D | L | M | F | S | L | D | W | I | K | A | K | T | E | L | V | G | P | A | - | - | - | - | - | - | - | - | H | L | I | H | E | Y | V | E | Y | S | S | V | L | D | Q | K | - | - | - | - | - | - | - | D | T | I | I | R | E | N | F | Q | - | - | - | E | F | L | S | L | N | G | H | L | L | G | R | Q | P | F | - | - | - | - | - | - | - | - | - | - | - | - | - | - | - | - | - | - | - | - | - | - | - | - | - | - | - | - | - | - | - | - | - | - | - | - | - | - | - | - | - | - | - | - | - | - | - | - | - | - | - | - | - | - | - | - | - | - | - | - | - | - | - | - | - | - | - | - | - | - | - | P | D | I | T | Q | L | G | L | C | Q | P | E | A | S | E | V | Y | Q | Q | A | K | L | R | A | Q | R | E | - | - | T | G | - | - | - | - | - | - | - | - | - | - | - | - | - | - | - | - | - | - | - | - | - | - | - | - | - | - | - | - | - | T | F | Y | L | - | - | - | - | - | - | - | - | - | - | E | W | I | N | K | - | - | - | - | - | - | - | K | S | L | K | N | L | S | R | - | - | - | - | - | - | - | - | - | - | L | V | V | R | P | H | R | D | - | - | - | - | - | - | - | - | - | - | - | - | - | - | - | A | V | Y | H | A | C | - | - | - | F | S | - | R | D | R | Q | R | I | A | - | - | - | - | - | - | - | - | - | S | C | G | A | D | K | T | L | Q | V | F | K | - | - | - | - | - | A | E | S | G | E | R | L | L | E | I | - | S | A | H | D | D | E | - | - | - | I | L | C | C | T | F | S | - | A | - | - | - | - | - | - | - | - | - | - | - | - | - | D | D | K | Y | V | A | T | C | S | A | D | K | K | V | K | I | - | - | - | W | N | S | R | - | - | - | - | - | - | - | - | - | - | - | - | - | - | - | - | - | - | - | - | - | - | - | - | - | - | - | - | - | - | - | - | - | - | - | - | - | - | - | - | - | - | - | - | - | - | - | - | - | - | - | - | - | - | - | - | - | - | - | - | - | - | - | - | - | - | - | - | - | - | - | - | - | - | - | - | - | - | - | - | - | - | - | - | - | - | - | - | - | - | - | - | - | - | - | - | - | - | - | - | - | - | - | - | - | - | - | - | - | - | - | - | - | - | - | - | - | - | - | - | - | - | - | - | - | - | - | - | - | - | - | - | - | - | - | - | - | - | - | - | T | G | E | C | L | Y | V | Y | - | E | E | H | T | - | - | - | - | - | E | Q | V | N | - | - | - | - | - | - | - | - | - | - | - | - | - | C | C | Q | - | - | - | - | - | - | - | - | - | - | - | - | - | F | N | N | R | R | - | - | - | - | - | - | - | - | - | - | - | - | - | - | - | - | G | Q | - | Y | L | L | A | T | C | S | D | - | - | - | - | D | T | Y | I | - | - | - | - | - | - | - | - | - | - | - | - | K | L | W | D | L | - | - | - | K | E | K | Y | C | R | N | T | M | F | - | - | - | - | - | - | - | - | - | - | - | - | - | - | - | - | - | G | H | I | N | A | - | - | - | - | - | - | - | - | - | - | - | - | - | - | - | - | V | S | H | C | R | F | S | P | N | D | D | - | - | - | - | - | - | - | - | - | - | - | - | - | - | - | - | - | - | - | - | - | - | - | - | - | - | - | - | Y | I | A | S | C | S | T | D | G | T | V | K | L | - | - | - | - | - | - | - | - | - | - | - | - | - | - | - | - | - | - | - | W | D | - | V | R | S | A | N | E | L | K | T | - | - | - | - | - | - | I | E | - | - | - | - | - | - | - | - | - | - | - | - | - | - | - | - | - | - | - | - | - | - | I | K | D | F | - | - | - | - | - | - | - | - | F | R | N | A | D | E | H | Q | D | - | - | - | - | - | - | - | - | - | - | - | - | - | - | - | - | - | - | - | - | - | - | - | - | - | - | - | E | V | E | - | V | L | V | K | C | C | - | - | - | - | - | - | - | - | - | - | - | - | - | - | - | - | S | W | S | R | S | S | D | T | I | L | V | V | A | K | N | K | L | - | - | - | - | - | - | - | - | - | - | - | L | L | F | D | V | T | T | - | - | - | C | N | - | - | - | - | - | - | - | - | - | - | - | - | - | - | - | - | - | - | - | - | - | - | - | - | - | - | - | - | - | - | - | - | - | - | - | - | - | - | - | - | - | - | - | - | - | - | - | - | - | - | - | - | - | - | - | - | - | - | - | - | - | - | - | - | - | - | - | - | - | - | - | - | - | - | - | - | - | - | - | - | - | - | - | - | - | - | - | - | - | - | - | - | - | - | - | - | - | - | - | - | - | - | - | - | - | - | - | - | - | - | - | - | - | - | - | - | - | - | - | - | - | - | - | - | - | - | L | L | T | - | - | - | - | - | - | - | - | - | - | - | - | - | - | - | - | - | - | - | - | - | - | - | - | - | - | - | - | - | - | - | - | - | - | - | - | - | - | - | - | - | - | - | - | - | - | - | - | - | - | - | - | - | - | - | - | - | - | - | - | - | - | - | - | - | - | - | - | - | - | - | - | - | - | - | - | - | - | - | - | - | - | - | - | - | - | - | - | - | - | - | - | - | - | - | - | - | - | - | - | - | - | - | - | - | - | - | - | - | - | - | - | - | - | - | - | - | - | - | - | - | - | - | - | - | - | - | - | - | - | - | - | - | - | - | - | - | - | - | - | - | - | - | - | - | - | - | - | - | - | - | - | - | - | - | - | - | - | - | - | - | - | - | - | - | - | - | - | - | - | - | - | - | - | - | - | - | - | - | - | - | - | - | Q | V | - | - | - | I | I | S | H | H | S | T | I | Q | Y | C | - | - | - | - | - | - | - | - | - | - | - | - | - | - | - | - | - | - | - | - | - | - | - | - | - | D | F | C | A | G | D | E | L | V | A | V | A | L | S | H | - | - | - | - | - | - | C | S | I | E | - | - | - | - | - | - | - | - | - | - | - | - | - | - | - | - | - | L | W | N | M | K | S | L | - | - | - | - | - | - | - | - | - | - | - | - | - | - | - | - | - | - | - | - | - | - | - | - | - | - | - | - | - | S | K | V | A | D | C | R | G | H | M | S | - | - | - | - | - | - | - | - | - | - | - | - | - | - | - | - | - | - | - | - | - | - | - | - | - | - | - | - | - | - | - | - | - | - | - | - | - | - | - | - | - | - | - | - | W | V | H | C | V | A | F | S | - | - | - | - | - | - | - | - | - | - | - | S | D | G | S | - | - | - | - | - | - | - | - | - | - | L | F | L | T | S | S | E | D | Q | - | - | - | - | - | - | - | - | - | - | - | - | - | - | T | I | R | I | - | W | - | - | - | - | - | - | - | - | - | - | - | - | - | - | - | - | - | - | - | - | - | - | - | - | - | - | - | - | - | - | - | - | - | - | - | - | - | - | E | T | N | K | V | C | K | S | S | D | A | V | L | K | S | E | - | - | - | - | - | - | - | - | - | - | - | - | - | - | - | - | L | D | V | V | F | - | - | - | - | - | - | - | - | - | - | - | - | - | - | - | - | - | - | - | - | - | - | - | - | - | - | - | - | - | - | - | - | - | - | - | - | - | - | - | - | - | - | - | - | - | - | - | - | - | - | - | - | - | - | H | N | G | E | V | M | - | - | - | - | - | - | - | - | - | - | - | - | - | - | - | - | - | - | - | - | - | - | - | - | - | - | - | - | - | - | - | - | - | - | - | - | I | L | A | I | Y | N | Q | K | H | I | Q | L | - | I | N | G | N | T | - | - | - | - | - | - | - | - | - | - | D | S | I | I | M | Q | T | E | P | - | - | - | - | - | - | - | - | - | - | - | - | - | - | - | - | - | - | - | - | - | - | Q | E | S | - | - | - | - | - | - | - | - | L | I | C | C | C | C | L | S | E | D | L | K | - | - | - | - | - | - | - | - | - | F | A | A | F | G | Q | - | - | - | - | - | - | - | - | - | - | - | - | - | - | - | - | - | - | - | - | - | - | - | - | - | - | - | - | - | - | - | - | - | - | - | - | E | - | - | - | - | - | - | - | - | - | N | G | A | - | - | - | - | - | - | - | - | - | - | I | - | - | - | - | - | - | - | - | - | - | - | - | - | - | - | - | - | - | - | - | - | - | - | - | - | - | - | - | - | - | - | - | - | - | - | - | - | - | - | K | V | L | Q | L | L | D | G | R | F | L | A | S | R | E | A | - | - | - | - | - | Y | - | - | - | - | - | - | - | - | - | - | - | - | R | T | S | V | Q | H | C | R | F | T | - | - | - | - | - | - | - | - | - | - | - | - | - | - | - | - | - | - | - | - | - | - | - | - | - | - | - | - | - | - | - | - | - | - | - | - | - | T | D | C | Q | T | L | I | S | S | A | - | H | D | A | - | - | - | - | - | - | - | - | V | I | Q | - | V | - | - | - | - | - | - | - | - | - | - | - | - | - | - | - | - | - | - | - | - | - | - | - | - | - | - | - | - | - | - | - | - | - | - | - | - | - | - | - | W | N | - | - | - | - | - | - | - | - | - | - | W | Q | L | N | E | C | V | F | - | - | - | - | - | - | - | - | - | - | - | - | - | - | - | - | - | - | - | - | - | - | - | - | - | - | - | - | - | - | - | - | - | - | - | - | - | - | - | - | - | - | - | - | - | - | - | - | - | - | L | R | G | H | E | - | - | - | - | - | - | - | - | - | - | - | - | - | - | - | - | - | - | - | - | - | - | - | - | - | - | E | A | I | K | D | F | A | L | L | - | - | - | - | - | - | - | - | - | - | - | - | - | - | - | - | - | - | - | - | - | - | - | E | N | - | S | K | L | L | - | S | W | - | - | - | S | F | D | G | T | V | - | - | - | - | - | - | - | - | - | - | - | - | - | - | - | - | - | - | - | - | - | - | - | - | - | - | - | - | - | - | - | - | - | - | - | K | V | W | N | I | T | T | G | K | T | - | - | - | - | - | E | K | D | F | A | C | - | - | - | - | - | - | - | H | R | - | - | - | - | - | - | - | - | - | - | - | - | - | - | - | - | - | - | - | G | A | V | L | - | - | - | - | - | - | - | - | - | - | - | - | - | - | - | S | C | A | - | - | - | - | - | - | - | - | - | - | - | - | - | - | - | V | S | P | D | G | S | - | R | F | S | S | T | - | S | A | D | K | T | - | - | - | - | - | - | A | K | I | W | S | F | E | S | S | - | - | - | - | - | - | - | - | - | - | - | - | - | - | - | - | - | - | - | - | - | - | - | - | S | V | L | H | E | - | - | - | - | - | - | - | - | L | - | - | - | - | K | G | H | E | A | C | V | - | - | - | - | - | - | - | - | - | R | C | S | - | A | F | S | - | - | - | - | - | - | - | - | - | P | N | N | K | - | L | L | A | T | - | - | - | - | - | - | - | - | G | D | D | K | G | D | I | R | I | W | D | I | L | T | G | E | L | L | - | - | - | - | - | - | - | - | - | - | - | - | - | - | - | - | - | - | - | - | - | - | - | - | - | - | - | - | - | - | - | - | - | - | - | - | - | - | - | - | - | - | - | - | - | - | - | - | - | - | - | - | - | - | - | - | H | F | C | S | S | V | T | V | D | - | - | - | - | - | - | - | - | - | - | - | - | - | - | - | - | - | - | - | - | E | G | E | P | T | H | G | G | W | - | - | - | - | - | - | - | - | V | T | A | L | S | F | S | P | D | S | K | - | - | - | - | - | - | - | - | - | - | - | - | - | - | - | - | - | - | - | - | - | - | - | - | - | - | - | - | - | - | - | - | - | - | - | - | - | - | - | - | - | - | - | - | - | M | L | V | S | - | - | S | G | G | Y | L | K | - | - | - | - | - | - | - | - | W | W | D | V | T | T | - | - | - | - | - | - | - | - | - | - | - | - | - | - | - | - | - | - | - | - | - | - | - | - | - | - | - | - | - | - | - | - | - | - | - | - | - | - | - | - | - | - | - | - | - | - | - | - | - | - | - | - | - | - | - | - | - | - | - | - | - | - | - | - | - | - | - | - | - | - | - | - | - | - | - | - | - | - | - | - | - | - | - | - | - | - | - | - | - | - | - | - | - | - | - | - | - | - | - | - | - | - | - | - | - | - | - | - | - | - | - | - | - | - | - | - | - | - | - | - | - | - | - | - | - | - | - | - | - | - | - | - | - | - | - | - | - | - | - | - | - | - | - | - | - | - | - | - | - | - | - | - | - | - | - | - | - | - | - | - | - | - | - | - | - | - | - | - | - | - | - | - | - | - | - | - | - | - | - | - | - | - | - | - | - | - | - | - | - | - | - | - | - | - | - | - | - | - | - | - | - | - | - | - | - | - | - | - | - | - | - | - | - | - | - | - | - | - | - | - | - | - | - | - | - | - | - | - | - | - | - | - | - | - | - | - | - | - | - | - | - | - | - | - | - | - | - | - | - | - | - | - | - | - | - | - | - | - | - | - | - | - | - | - | - | - | - | G | E | S | L | Q | T | F | Y | T | - | - | - | - | - | N | G | - | - | - | T | N | L | K | S | I | - | - | - | H | V | S | P | - | - | - | - | S | F | E | V | - | - | - | - | - | - | - | - | - | - | - | - | - | - | - | - | - | - | - | - | - | - | - | - | - | - | Y | V | T | V | D | - | N | L | G | I | L | - | - | Y | V | L | R | K | - | - | - | - | - | - | - | - | - | - | - | - | - | - | - | - | - | - | - | - | - | - | - | - | - | - | - | - | - | - | - | - | - | - | L | - | - | - | - | - | - | - | - | - | - | - | - | - | - | - | - | - | - | - | - | - | - | - | - | - | - | - | - | - | - | - | - | - | - | - | - | - | - | - | - | - | - | - | - | - | - | - | - | - | - | - | - | - | M | - | - | - | - | - | - | - | - | - | - | - | - | - | - | - | - | - | - | - | - | - | - | - | - | - | - | - | - | - | - | - | - | - | - | - | - | - | - | - | - | - | - | - | - | - | - | - | - | - | - | - | - | - | - | E | E | A | R | R | S | - | - | - | - | - | - | - | - | - | - | - | - | - | - | - | - | - | - | - | - | A | L | R | R | - | - | - | - | - | - | - | - | - | - | - | - | - | - | - | - | - | - | - | - | - | - | - | - | - | - | - | - | - | - | G | R | - | - | - | - | - | - | - | - | L | R | L | - | V | E | G | L | R | P | D | V | L | W | - | - | - | - | - | - | - | - | - | - | - | - | - | - | - | - | - | - | - | - | - | - | - | - | - | - | - | - | - | - | - | - | - | - | - | - | - | - | - | - | - | - | - | - | - | G | P | L | R | E | R | G | V | - | - | - | - | - | - | - | - | - | - | - | - | - | - | - | - | - | - | - | - | - | - | - | - | F | T | S | A | M | V | E | D | - | - | - | - | - | - | - | - | - | - | - | - | - | - | - | - | - | - | - | - | - | - | - | - | - | M | Q | S | A | - | - | - | - | - | - | - | - | - | - | G | T | R | R | E | Q | - | A | R | Q | L | V | I | D | L | E | S | R | G | - | K | Q | A | F | P | I | F | L | S | I | L | R | D | T | G | H | G | D | L | A | D | M | L | D | E | G | C | G | S | P | - | - | - | - | M | S | P | P | - | - | - | - | - | - | - | - | - | - | - | - | - | - | - | - | - | - | - | - | - | - | - | - | - | - | - | - | - | - | - | - | V | D | - | - | - | - | - | - | - | - | - | - | - | - | - | - | - | L | R | P | V | - | - | - | - | - | - | - | - | - | - | - | - | - | - | - | - | - | - | - | - | - | - | - | - | - | - | - | - | - | - | - | - | - | - | - | - | - | - | - | Q | L | E | L | P | G | E | R | R | D | K | S | V | - | - | - | - | - | - | - | - | - | - | - | R | - | - | - | - | - | - | - | - | - | - | - | - | - | - | - | - | - | - | - | - | - | - | - | - | - | - | - | - | - | - | - | - | - | - | - | - | - | - | - | - | - | - | - | - | - | - | - | - | - | - | - | - | - | - | - | - | - | - | - | - | - | - | - | - | - | - | - | - | - | - | - | - | - | - | - | - | - | - | - | - | - | - | - | - | - | - | - | - | - | - | - | - | - | - | - | - | - | - | - | - | - | - | - | - | - | - | - | - | - | - | - | - | - | - | - | - | - | - | - | - | - | - | - | - | - | - | - | - | - | - | - | - | - | - | - | - | - | - | - | - | - | - | - | - | - | - | - | - | - | - | - | - | - | - | - | - | T | T | E | Y | L | S | - | - | - | - | - | - | - | - | - | - | - | - | - | - | - | - | - | - | - | - | - | - | - | - | - | - | I | P | V | Q | A | - | - | - | - | - | - | E | S | E | R | F | - | R | M | - | - | - | - | - | - | - | - | - | - | - | - | - | - | - | - | - | - | P | P | A | P | A | Q | G | - | - | - | - | - | - | - | - | - | - | - | - | - | - | - | - | - | - | - | - | S | - | - | - | - | - | - | - | - | - | - | A | - | V | D | - | - | K | S | D | H | R | - | - | - | - | - | L | V | Y | Q | - | - | L | R | A | D | - | P | C | G | H | C | L | I | F | N | N | V | S | F | S | P | D | S | - | - | N | L | S | P | R | A | G | S | D | I | D | C | E | K | L | E | K | R | F | - | R | S | L | C | F | N | V | Q | T | L | R | N | L | K | A | Q | E | I | D | L | E | L | R | K | L | A | R | L | D | H | - | S | A | L | D | C | C | L | V | V | I | L | S | H | G | C | Q | T | S | H | I | Q | F | P | G | G | - | I | Y | G | T | D | G | K | T | I | P | I | E | K | I | V | N | Y | F | N | G | S | Q | C | P | S | L | R | G | K | P | K | L | F | F | I | Q | A | C | - | - | G | G | E | Q | K | D | Q | G | F | E | V | D | F | E | - | - | - | S | P | R | D | - | E | T | C | Q | G | S | - | - | - | - | - | - | - | I | E | S | D | A | I | P | F | Q | A | P | S | G | S | V | - | - | - | - | - | - | - | - | D | E | L | D | A | V | - | - | - | - | - | - | - | A | S | L | - | - | - | P | T | P | G | D | I | L | V | S | Y | S | T | F | P | - | - | G | - | - | - | - | - | - | - | - | F | V | S | W | R | D | R | V | S | G | S | W | Y | V | E | T | L | D | S | V | L | E | C | Y | A | S | - | S | - | - | - | - | - | - | - | - | - | - | - | - | - | - | - | - | - | - | - | - | - | - | - | - | - | - | - | - | - | - | - | E | D | L | L | T | M | L | - | - | - | - | - | V | R | - | - | - | - | - | - | - | - | - | - | - | - | V | S | D | I | V | - | - | - | - | - | S | T | K | - | - | - | - | - | - | - | G | K | Y | - | K | Q | I | - | - | P | G | - | - | - | - | - | - | - | - | - | - | - | - | - | - | - | C | F | N | F | - | - | - | - | - | - | - | - | - | L | R | K | K | F | F | F | L | C | K | - | - | - | - | - | - | - | - | - | - | - | - | - | - | - | - | - | - | - | - | - | - | - | - | - | - | - | - | - | - | - | - | - | - | - | - | - | M | - | - | - | - | - | - | - | - | - | - | - | - | - | - | - | - | - | - | - | - | - | - | - | - | - | - | - | - | - | - | - | - | - | - | - | - | - | - | - | - | - | - | - | - | - | - | - | - | - | - | - | - | G | D | I | E | K | G | K | K | I | F | V | Q | K | C | S | Q | C | H | T | V | E | K | G | G | K | H | K | T | G | P | N | L | H | G | L | F | G | R | K | T | G | Q | A | E | G | F | S | Y | T | D | A | N | K | N | K | G | I | T | W | G | E | D | T | L | M | E | Y | L | E | N | P | K | K | Y | I | P | G | T | K | M | I | F | A | G | I | K | K | K | S | E | R | V | D | L | I | A | Y | L | K | D | A | T | K | K | - | - | - | - | - | - |
| Crocodylus\_porosus | M | - | - | - | - | - | - | - | - | - | - | - | - | - | - | - | - | - | - | - | - | - | - | - | - | - | - | - | - | - | - | - | - | - | - | - | - | - | - | - | - | - | - | - | - | - | - | - | - | - | - | - | - | - | - | - | - | - | - | - | - | - | - | - | - | - | - | - | - | - | - | - | - | - | - | - | - | - | - | - | - | - | - | - | - | - | - | - | - | - | - | - | - | - | - | - | - | - | N | V | K | S | - | R | N | - | - | - | - | - | - | - | - | - | - | - | - | - | - | - | - | H | L | V | - | - | - | - | - | - | - | - | - | - | - | - | - | - | - | - | - | - | - | - | - | - | - | - | - | - | - | - | - | - | - | - | - | - | - | - | - | - | - | - | - | - | - | - | - | - | - | - | - | - | - | - | - | - | - | - | - | - | - | - | - | - | - | - | - | - | - | - | - | - | - | - | - | - | - | - | - | - | - | - | - | - | - | - | - | - | - | - | - | - | - | - | - | - | - | - | - | - | - | - | - | - | - | - | - | - | - | - | - | - | - | - | - | - | - | - | - | - | - | - | - | L | N | R | V | A | L | E | R | D | I | Q | T | S | Y | - | I | M | D | H | M | I | S | D | Q | V | L | T | L | Q | E | E | E | K | V | K | T | Q | - | - | - | - | - | - | - | - | - | - | - | - | - | - | - | - | - | A | T | R | R | D | R | A | A | M | L | I | S | I | I | L | K | K | - | - | - | - | D | N | N | S | Y | I | S | - | - | - | - | - | F | Y | N | A | L | - | L | H | E | G | - | - | - | - | - | - | Y | K | D | L | A | A | L | L | H | D | G | I | P | S | I | T | S | Y | D | K | - | K | - | - | - | - | - | - | - | - | - | - | - | - | - | - | - | - | - | - | - | - | - | - | N | S | T | D | G | V | A | - | - | - | - | - | - | - | - | - | - | - | F | C | V | R | T | I | L | C | E | G | G | - | - | - | - | - | - | - | V | P | Q | R | P | V | V | F | V | T | R | P | E | L | V | N | E | I | - | - | - | - | - | - | - | - | - | - | R | Q | K | L | Y | - | - | - | - | - | - | - | - | - | - | - | - | - | - | - | - | - | - | - | - | - | - | - | - | - | - | - | - | - | - | - | - | N | L | R | D | D | P | - | - | - | - | - | - | - | - | - | - | - | - | - | - | - | - | - | - | - | - | - | - | - | - | - | - | - | - | - | - | - | - | - | - | - | - | - | - | G | W | V | T | V | Y | G | M | A | G | S | G | K | S | V | L | T | A | E | A | L | - | R | D | E | R | L | L | E | E | C | F | P | G | G | V | H | W | I | S | V | G | - | - | - | - | - | - | - | K | - | - | - | - | - | - | Q | D | D | A | - | - | - | - | - | - | - | - | - | - | - | - | G | L | L | M | K | L | H | N | L | C | S | R | L | D | - | - | - | - | L | - | - | - | - | - | - | - | - | - | - | - | - | - | - | - | D | F | T | Y | A | Q | R | P | P | L | N | I | - | - | - | - | - | - | - | - | - | - | - | - | - | - | - | - | G | E | A | K | D | R | L | R | L | L | M | L | H | - | - | - | - | - | - | - | - | - | - | - | - | K | Y | P | R | S | L | L | - | - | V | L | D | D | I | W | D | S | Q | T | L | E | A | F | D | - | N | H | C | K | V | L | I | T | S | R | D | R | S | V | T | D | H | V | - | T | G | R | - | - | K | Y | E | V | H | M | E | S | - | - | - | G | L | S | N | E | K | G | L | E | I | L | S | L | F | V | D | M | - | K | I | N | D | M | P | A | - | - | - | - | - | - | - | - | - | - | - | - | - | - | Q | A | D | K | I | V | R | E | C | K | - | - | - | - | - | - | - | - | - | - | - | - | - | - | - | - | - | - | - | G | - | S | P | L | V | L | S | L | I | G | A | L | L | R | D | F | P | N | - | - | - | - | - | - | - | - | - | - | R | W | E | Y | Y | V | - | R | Q | L | Q | N | K | Q | F | K | R | I | - | - | - | - | - | - | - | - | - | - | - | - | - | - | - | - | - | - | - | - | - | - | K | K | S | S | S | Y | D | - | - | - | - | Y | E | A | L | D | E | A | M | S | I | S | V | E | - | M | L | N | - | - | - | E | N | F | K | D | F | - | Y | K | D | L | S | I | L | P | K | D | V | K | V | P | T | K | V | F | C | I | L | W | D | - | - | - | - | M | E | P | E | E | V | E | D | M | Q | Q | E | F | - | - | - | - | - | - | - | - | - | - | - | V | N | K | S | L | L | F | C | D | R | N | G | K | S | - | - | - | - | - | - | - | - | - | F | H | Y | Y | L | H | D | L | Q | L | D | F | L | I | E | K | - | - | N | R | E | Q | - | L | - | Q | E | L | H | E | K | I | V | K | Q | Y | K | K | - | H | Y | - | - | - | - | - | - | K | L | C | I | P | T | P | G | - | E | E | - | - | - | - | - | - | - | D | C | M | Y | W | Y | N | F | L | A | F | H | M | A | S | A | K | - | - | - | - | M | Q | E | E | L | C | D | L | M | F | S | L | D | W | I | K | A | K | T | E | L | V | G | P | A | - | - | - | - | - | - | - | - | H | L | I | H | E | Y | V | E | Y | S | H | I | L | D | Q | K | - | - | - | - | - | - | - | N | S | L | A | R | G | N | F | Q | - | - | - | K | F | L | S | L | N | G | H | L | L | G | R | Q | P | F | - | - | - | - | - | - | - | - | - | - | - | - | - | - | - | - | - | - | - | - | - | - | - | - | - | - | - | - | - | - | - | - | - | - | - | - | - | - | - | - | - | - | - | - | - | - | - | - | - | - | - | - | - | - | - | - | - | - | - | - | - | - | - | - | - | - | - | - | - | - | - | P | D | I | L | Q | L | G | L | C | Q | P | E | T | S | E | V | Y | Q | Q | A | K | L | K | A | Q | R | E | V | D | K | G | - | - | - | - | - | - | - | - | - | - | - | - | - | - | - | - | - | - | - | - | - | - | - | - | - | - | - | - | - | A | F | Y | V | - | - | - | - | - | - | - | - | - | - | E | W | I | N | K | - | - | - | - | - | - | - | - | G | M | K | N | L | S | R | - | - | - | - | - | - | - | - | - | - | L | V | V | R | P | H | T | D | - | - | - | - | - | - | - | - | - | - | - | - | - | - | - | A | V | Y | H | A | C | - | - | - | F | S | - | Q | D | G | Q | K | I | A | - | - | - | - | - | - | - | - | - | S | C | G | A | D | R | T | V | Q | V | F | K | - | - | - | - | - | A | E | T | G | E | K | L | Q | S | I | - | E | A | H | D | D | E | - | - | - | V | L | Y | C | A | L | S | - | P | - | - | - | - | - | - | - | - | - | - | - | - | - | D | G | R | F | L | A | T | C | S | A | D | N | K | A | K | I | - | - | - | W | C | L | R | - | - | - | - | - | - | - | - | - | - | - | - | - | - | - | - | - | - | - | - | - | - | - | - | - | - | - | - | - | - | - | - | - | - | - | - | - | - | - | - | - | - | - | - | - | - | - | - | - | - | - | - | - | - | - | - | - | - | - | - | - | - | - | - | - | - | - | - | - | - | - | - | - | - | - | - | - | - | - | - | - | - | - | - | - | - | - | - | - | - | - | - | - | - | - | - | - | - | - | - | - | - | - | - | - | - | - | - | - | - | - | - | - | - | - | - | - | - | - | - | - | - | - | - | - | - | - | - | - | - | - | - | - | - | - | - | - | - | - | - | T | G | Q | S | I | Y | D | C | - | E | E | H | T | - | - | - | - | - | E | Q | V | N | - | - | - | - | - | - | - | - | - | - | - | - | - | C | C | H | - | - | - | - | - | - | - | - | - | - | - | - | - | F | N | S | G | N | - | - | - | - | - | - | - | - | - | - | - | - | - | - | - | - | N | K | - | L | L | L | T | T | C | S | N | - | - | - | - | D | S | L | L | - | - | - | - | - | - | - | - | - | - | - | - | K | L | W | D | V | - | - | - | R | K | K | H | C | Q | N | T | M | F | - | - | - | - | - | - | - | - | - | - | - | - | - | - | - | - | - | G | H | T | S | S | - | - | - | - | - | - | - | - | - | - | - | - | - | - | - | - | V | N | H | C | R | F | S | P | D | D | K | - | - | - | - | - | - | - | - | - | - | - | - | - | - | - | - | - | - | - | - | - | - | - | - | - | - | - | - | Y | V | A | S | C | S | A | D | G | T | V | K | L | - | - | - | - | - | - | - | - | - | - | - | - | - | - | - | - | - | - | - | W | V | - | T | S | S | G | N | E | L | K | C | - | - | - | - | - | - | I | K | - | - | - | - | - | - | - | - | - | - | - | - | - | - | - | - | - | - | - | - | - | - | I | K | D | F | - | - | - | - | - | - | - | - | F | R | N | A | D | E | L | - | - | - | - | - | - | - | - | - | - | - | - | - | - | - | - | - | - | - | - | - | - | - | - | - | - | - | - | - | E | V | E | - | V | L | P | K | C | C | - | - | - | - | - | - | - | - | - | - | - | - | - | - | - | - | S | W | S | T | D | S | A | T | V | M | V | A | A | K | N | K | L | - | - | - | - | - | - | - | - | - | - | - | L | L | L | D | V | E | T | - | - | - | S | D | - | - | - | - | - | - | - | - | - | - | - | - | - | - | - | - | - | - | - | - | - | - | - | - | - | - | - | - | - | - | - | - | - | - | - | - | - | - | - | - | - | - | - | - | - | - | - | - | - | - | - | - | - | - | - | - | - | - | - | - | - | - | - | - | - | - | - | - | - | - | - | - | - | - | - | - | - | - | - | - | - | - | - | - | - | - | - | - | - | - | - | - | - | - | - | - | - | - | - | - | - | - | - | - | - | - | - | - | - | - | - | - | - | - | - | - | - | - | - | - | - | - | - | - | - | - | L | L | E | - | - | - | - | - | - | - | - | - | - | - | - | - | - | - | - | - | - | - | - | - | - | - | - | - | - | - | - | - | - | - | - | - | - | - | - | - | - | - | - | - | - | - | - | - | - | - | - | - | - | - | - | - | - | - | - | - | - | - | - | - | - | - | - | - | - | - | - | - | - | - | - | - | - | - | - | - | - | - | - | - | - | - | - | - | - | - | - | - | - | - | - | - | - | - | - | - | - | - | - | - | - | - | - | - | - | - | - | - | - | - | - | - | - | - | - | - | - | - | - | - | - | - | - | - | - | - | - | - | - | - | - | - | - | - | - | - | - | - | - | - | - | - | - | - | - | - | - | - | - | - | - | - | - | - | - | - | - | - | - | - | - | - | - | - | - | - | - | - | - | - | - | - | - | - | - | - | - | - | - | - | - | - | E | V | - | - | - | L | - | S | Y | Q | N | S | I | Q | Y | Y | - | - | - | - | - | - | - | - | - | - | - | - | - | - | - | - | - | - | - | - | - | - | - | - | - | D | F | C | P | S | N | Q | L | V | A | V | V | L | S | H | - | - | - | - | - | - | Y | F | V | E | - | - | - | - | - | - | - | - | - | - | - | - | - | - | - | - | - | L | W | N | T | K | T | K | - | - | - | - | - | - | - | - | - | - | - | - | - | - | - | - | - | - | - | - | - | - | - | - | - | - | - | - | - | T | K | V | A | D | C | R | E | H | M | S | - | - | - | - | - | - | - | - | - | - | - | - | - | - | - | - | - | - | - | - | - | - | - | - | - | - | - | - | - | - | - | - | - | - | - | - | - | - | - | - | - | - | - | - | W | V | H | C | V | T | F | S | - | - | - | - | - | - | - | - | - | - | - | P | D | G | S | - | - | - | - | - | - | - | - | - | - | L | F | L | T | S | S | D | D | Q | - | - | - | - | - | - | - | - | - | - | - | - | - | - | T | I | R | I | - | W | - | - | - | - | - | - | - | - | - | - | - | - | - | - | - | - | - | - | - | - | - | - | - | - | - | - | - | - | - | - | - | - | - | - | - | - | - | - | E | T | S | K | V | C | K | P | P | E | I | A | L | K | S | E | - | - | - | - | - | - | - | - | - | - | - | - | - | - | - | - | L | D | V | V | F | - | - | - | - | - | - | - | - | - | - | - | - | - | - | - | - | - | - | - | - | - | - | - | - | - | - | - | - | - | - | - | - | - | - | - | - | - | - | - | - | - | - | - | - | - | - | - | - | - | - | - | - | - | - | Q | D | D | E | V | M | - | - | - | - | - | - | - | - | - | - | - | - | - | - | - | - | - | - | - | - | - | - | - | - | - | - | - | - | - | - | - | - | - | - | - | - | I | L | G | I | L | N | R | T | H | L | Q | L | - | I | N | G | N | T | - | - | - | - | - | - | - | - | - | - | G | H | T | V | S | L | T | D | V | - | - | - | - | - | - | - | - | - | - | - | - | - | - | - | - | - | - | - | - | - | - | E | E | S | - | - | - | - | - | - | - | - | - | I | S | C | C | C | L | S | E | D | L | Q | - | - | - | - | - | - | - | - | - | F | A | A | F | G | Q | - | - | - | - | - | - | - | - | - | - | - | - | - | - | - | - | - | - | - | - | - | - | - | - | - | - | - | - | - | - | - | - | - | - | - | - | E | - | - | - | - | - | - | - | - | - | N | G | T | - | - | - | - | - | - | - | - | - | - | V | - | - | - | - | - | - | - | - | - | - | - | - | - | - | - | - | - | - | - | - | - | - | - | - | - | - | - | - | - | - | - | - | - | - | - | - | - | - | - | K | V | L | E | V | S | G | G | R | V | L | K | S | R | R | G | - | - | - | - | - | H | - | - | - | - | - | - | - | - | - | - | - | - | K | T | S | V | K | H | C | L | F | T | - | - | - | - | - | - | - | - | - | - | - | - | - | - | - | - | - | - | - | - | - | - | - | - | - | - | - | - | - | - | - | - | - | - | - | - | - | S | D | G | Q | T | L | I | S | S | D | - | D | D | P | - | - | - | - | - | - | - | - | V | I | Q | - | V | - | - | - | - | - | - | - | - | - | - | - | - | - | - | - | - | - | - | - | - | - | - | - | - | - | - | - | - | - | - | - | - | - | - | - | - | - | - | - | W | N | - | - | - | - | - | - | - | - | - | - | W | E | L | E | E | C | L | F | - | - | - | - | - | - | - | - | - | - | - | - | - | - | - | - | - | - | - | - | - | - | - | - | - | - | - | - | - | - | - | - | - | - | - | - | - | - | - | - | - | - | - | - | - | - | - | - | - | - | L | E | G | H | N | - | - | - | - | - | - | - | - | - | - | - | - | - | - | - | - | - | - | - | - | - | - | - | - | - | - | E | T | I | K | N | F | R | L | L | - | - | - | - | - | - | - | - | - | - | - | - | - | - | - | - | - | - | - | - | - | - | - | E | N | - | S | R | L | L | - | S | W | - | - | - | S | S | D | G | T | V | - | - | - | - | - | - | - | - | - | - | - | - | - | - | - | - | - | - | - | - | - | - | - | - | - | - | - | - | - | - | - | - | - | - | - | K | V | W | N | I | I | S | G | K | I | - | - | - | - | - | E | V | N | F | A | C | - | - | - | - | - | - | - | H | K | - | - | - | - | - | - | - | - | - | - | - | - | - | - | - | - | - | - | - | D | T | V | L | - | - | - | - | - | - | - | - | - | - | - | - | - | - | - | S | C | A | - | - | - | - | - | - | - | - | - | - | - | - | - | - | - | I | S | P | D | A | T | - | K | F | S | S | T | - | S | V | D | K | T | - | - | - | - | - | - | A | K | I | W | S | F | K | S | S | - | - | - | - | - | - | - | - | - | - | - | - | - | - | - | - | - | - | - | - | - | - | - | - | S | I | L | H | E | - | - | - | - | - | - | - | - | L | - | - | - | - | K | G | H | E | A | C | V | - | - | - | - | - | - | - | - | - | R | C | C | - | T | F | S | - | - | - | - | - | - | - | - | - | F | D | N | K | - | L | L | A | T | - | - | - | - | - | - | - | - | G | D | D | K | R | E | I | R | I | W | D | V | S | S | G | E | L | L | - | - | - | - | - | - | - | - | - | - | - | - | - | - | - | - | - | - | - | - | - | - | - | - | - | - | - | - | - | - | - | - | - | - | - | - | - | - | - | - | - | - | - | - | - | - | - | - | - | - | - | - | - | - | - | - | H | L | C | S | P | V | T | I | D | - | - | - | - | - | - | - | - | - | - | - | - | - | - | - | - | - | - | - | - | E | G | G | P | T | H | D | Y | W | - | - | - | - | - | - | - | - | V | T | D | L | C | F | S | P | D | S | K | - | - | - | - | - | - | - | - | - | - | - | - | - | - | - | - | - | - | - | - | - | - | - | - | - | - | - | - | - | - | - | - | - | - | - | - | - | - | - | - | - | - | - | - | - | M | L | V | S | - | - | S | G | G | Y | L | K | - | - | - | - | - | - | - | - | W | W | N | A | V | T | - | - | - | - | - | - | - | - | - | - | - | - | - | - | - | - | - | - | - | - | - | - | - | - | - | - | - | - | - | - | - | - | - | - | - | - | - | - | - | - | - | - | - | - | - | - | - | - | - | - | - | - | - | - | - | - | - | - | - | - | - | - | - | - | - | - | - | - | - | - | - | - | - | - | - | - | - | - | - | - | - | - | - | - | - | - | - | - | - | - | - | - | - | - | - | - | - | - | - | - | - | - | - | - | - | - | - | - | - | - | - | - | - | - | - | - | - | - | - | - | - | - | - | - | - | - | - | - | - | - | - | - | - | - | - | - | - | - | - | - | - | - | - | - | - | - | - | - | - | - | - | - | - | - | - | - | - | - | - | - | - | - | - | - | - | - | - | - | - | - | - | - | - | - | - | - | - | - | - | - | - | - | - | - | - | - | - | - | - | - | - | - | - | - | - | - | - | - | - | - | - | - | - | - | - | - | - | - | - | - | - | - | - | - | - | - | - | - | - | - | - | - | - | - | - | - | - | - | - | - | - | - | - | - | - | - | - | - | - | - | - | - | - | - | - | - | - | - | - | - | - | - | - | - | - | - | - | - | - | - | - | - | - | - | - | - | - | G | E | S | L | Q | T | F | Y | T | - | - | - | - | - | N | G | - | - | - | T | N | L | Q | S | I | - | - | - | Y | V | S | S | - | - | - | - | N | F | K | I | - | - | - | - | - | - | - | - | - | - | - | - | - | - | - | - | - | - | - | - | - | - | - | - | - | - | F | V | T | V | D | - | N | L | G | I | L | - | - | Y | V | L | Q | M | - | - | - | - | - | - | - | - | - | - | - | - | - | - | - | - | - | - | - | - | - | - | - | - | - | - | - | - | - | - | - | - | - | - | M | E | - | - | - | - | - | - | - | - | - | - | - | - | - | - | - | - | - | - | - | - | - | - | - | - | - | - | - | - | - | - | - | - | - | - | - | - | - | - | - | - | - | - | - | - | - | - | - | - | - | - | - | - | M | - | - | - | - | - | - | - | - | - | - | - | - | - | - | - | - | - | - | - | - | - | - | - | - | - | - | - | - | - | - | - | - | - | - | - | - | - | - | - | - | - | - | - | - | - | - | - | - | - | - | - | - | - | - | E | A | E | Q | R | R | - | - | - | - | - | - | - | - | - | - | - | - | - | - | - | - | - | - | - | - | L | L | R | L | - | - | - | - | - | - | - | - | - | - | - | - | - | - | - | - | - | - | - | - | - | - | - | - | - | - | - | - | - | - | G | R | - | - | - | - | - | - | - | - | V | R | L | - | V | R | G | L | Q | L | E | A | L | W | - | - | - | - | - | - | - | - | - | - | - | - | - | - | - | - | - | - | - | - | - | - | - | - | - | - | - | - | - | - | - | - | - | - | - | - | - | - | - | - | - | - | - | - | - | E | P | L | A | R | E | G | V | - | - | - | - | - | - | - | - | - | - | - | - | - | - | - | - | - | - | - | - | - | - | - | - | F | S | Q | D | M | L | D | D | - | - | - | - | - | - | - | - | - | - | - | - | - | - | - | - | - | - | - | - | - | - | - | - | - | I | Q | R | A | - | - | - | - | - | - | - | - | - | - | G | T | P | R | D | Q | - | A | R | Q | L | L | I | D | L | E | T | R | G | - | K | Q | A | F | P | T | F | V | S | I | L | H | E | T | G | Q | K | E | L | A | D | F | L | S | E | G | C | T | S | P | - | - | - | - | Q | P | S | S | - | - | - | - | - | - | - | - | - | - | - | - | - | - | - | - | - | - | - | - | - | - | - | - | - | - | - | - | - | - | - | - | I | D | - | - | - | - | - | - | - | - | - | - | - | - | - | - | - | I | K | P | I | - | - | - | - | - | - | - | - | - | - | - | - | - | - | - | - | - | - | - | - | - | - | - | - | - | - | - | - | - | - | - | - | - | - | - | - | - | - | - | E | I | G | L | R | G | G | K | G | D | A | S | G | - | - | - | - | - | - | - | - | - | - | - | R | - | - | - | - | - | - | - | - | - | - | - | - | - | - | - | - | - | - | - | - | - | - | - | - | - | - | - | - | - | - | - | - | - | - | - | - | - | - | - | - | - | - | - | - | - | - | - | - | - | - | - | - | - | - | - | - | - | - | - | - | - | - | - | - | - | - | - | - | - | - | - | - | - | - | - | - | - | - | - | - | - | - | - | - | - | - | - | - | - | - | - | - | - | - | - | - | - | - | - | - | - | - | - | - | - | - | - | - | - | - | - | - | - | - | - | - | - | - | - | - | - | - | - | - | - | - | - | - | - | - | - | - | - | - | - | - | - | - | - | - | - | - | - | - | - | - | - | - | - | - | - | - | - | - | - | - | - | - | - | - | - | P | - | - | - | - | - | - | - | - | - | - | - | - | - | - | - | - | - | - | - | - | - | - | - | - | - | - | L | P | V | Q | A | - | - | - | - | - | - | R | T | E | K | C | - | Q | E | L | D | - | - | - | - | - | - | - | - | - | - | - | - | - | - | - | H | P | G | T | V | S | H | S | - | - | - | - | - | - | - | - | - | - | - | - | - | - | - | - | - | - | - | - | P | - | - | - | - | - | - | - | - | - | - | A | - | A | D | R | F | R | R | N | S | D | - | - | - | - | - | M | V | Y | V | - | - | L | K | S | R | - | P | C | G | Y | C | F | I | I | N | N | V | H | F | S | R | E | S | - | - | G | L | S | L | R | Q | G | S | N | L | D | Y | E | K | L | K | R | R | F | - | S | L | F | H | F | E | V | L | T | R | Q | D | L | T | A | K | K | M | R | W | E | L | S | E | L | A | K | K | D | H | - | S | A | M | D | C | C | L | V | V | I | L | S | H | G | R | Q | R | S | H | I | Q | F | P | G | E | - | V | C | G | T | D | G | K | S | I | P | I | E | E | I | V | N | Y | F | N | G | L | N | C | W | S | L | R | G | K | P | K | L | F | I | I | Q | A | C | - | - | G | G | E | E | R | D | Y | G | F | Q | L | D | S | D | - | - | - | S | S | G | N | - | Q | G | Y | G | A | S | - | - | - | - | - | - | - | L | E | S | D | A | S | P | L | K | P | S | S | G | N | L | - | - | - | - | - | - | - | - | D | E | P | D | A | V | - | - | - | - | - | - | - | A | S | L | - | - | - | P | T | A | S | D | I | L | V | S | Y | S | T | F | P | - | - | G | - | - | - | - | - | - | - | - | F | V | S | W | R | D | R | N | S | G | S | W | F | V | E | T | L | D | R | I | L | E | Q | Y | A | D | - | S | - | - | - | - | - | - | - | - | - | - | - | - | - | - | - | - | - | - | - | - | - | - | - | - | - | - | - | - | - | - | - | E | D | L | Q | K | L | L | - | - | - | - | - | L | R | - | - | - | - | - | - | - | - | - | - | - | - | V | A | N | E | V | - | - | - | - | - | S | A | K | - | - | - | - | - | - | - | E | K | Y | - | K | Q | M | - | - | P | G | - | - | - | - | - | - | - | - | - | - | - | - | - | - | - | C | Y | H | Y | - | - | - | - | - | - | - | - | - | L | R | K | A | F | F | F | M | T | K | - | - | - | - | - | - | - | - | - | - | - | - | - | - | - | - | - | - | - | - | - | - | - | - | - | - | - | - | - | - | - | - | - | - | - | - | - | M | - | - | - | - | - | - | - | - | - | - | - | - | - | - | - | - | - | - | - | - | - | - | - | - | - | - | - | - | - | - | - | - | - | - | - | - | - | - | - | - | - | - | - | - | - | - | - | - | - | - | - | - | G | D | V | E | K | G | K | K | I | F | V | Q | K | C | A | Q | C | H | T | V | E | K | G | G | K | H | K | T | G | P | N | L | H | G | L | F | G | R | K | T | G | Q | A | P | G | F | S | Y | T | E | A | N | K | N | K | G | I | T | W | D | E | D | T | L | M | E | Y | L | E | N | P | K | K | Y | I | P | G | T | K | M | I | F | A | G | I | K | K | K | P | E | R | A | D | L | I | A | Y | L | K | D | A | T | A | N | - | - | - | - | - | - |
| and 88 other sequences |  |
| conservation |  |  |  |  |  |  |  |  |  |  |  |  |  |  |  |  |  |  |  |  |  |  |  |  |  |  |  |  |  |  |  |  |  |  |  |  |  |  |  |  |  |  |  |  |  |  |  |  |  |  |  |  |  |  |  |  |  |  |  |  |  |  |  |  |  |  |  |  |  |  |  |  |  |  |  |  |  |  |  |  |  |  |  |  |  |  |  |  |  |  |  |  |  |  |  |  |  |  |  |  |  |  |  |  |  |  |  |  |  |  |  |  |  |  |  |  |  |  |  |  |  |  |  |  |  |  |  |  |  |  |  |  |  |  |  |  |  |  |  |  |  |  |  |  |  |  |  |  |  |  |  |  |  |  |  |  |  |  |  |  |  |  |  |  |  |  |  |  |  |  |  |  |  |  |  |  |  |  |  |  |  |  |  |  |  |  |  |  |  |  |  |  |  |  |  |  |  |  |  |  |  |  |  |  |  |  |  |  |  |  |  |  |  |  |  |  |  |  |  |  |  |  |  |  |  |  |  |  |  |  |  |  |  |  |  |  |  |  |  |  |  |  |  |  |  |  |  |  |  |  |  |  |  |  |  |  |  |  |  |  |  |  |  |  |  |  |  |  |  |  |  |  |  |  |  |  |  |  |  |  |  |  |  |  |  |  |  |  |  |  |  |  |  |  |  |  |  |  |  |  |  |  |  |  |  |  |  |  |  |  |  |  |  |  |  |  |  |  |  |  |  |  |  |  |  |  |  |  |  |  |  |  |  |  |  |  |  |  |  |  |  |  |  |  |  |  |  |  |  |  |  |  |  |  |  |  |  |  |  |  |  |  |  |  |  |  |  |  |  |  |  |  |  |  |  |  |  |  |  |  |  |  |  |  |  |  |  |  |  |  |  |  |  |  |  |  |  |  |  |  |  |  |  |  |  |  |  |  |  |  |  |  |  |  |  |  |  |  |  |  |  |  |  |  |  |  |  |  |  |  |  |  |  |  |  |  |  |  |  |  |  |  |  |  |  |  |  |  |  |  |  |  |  |  |  |  |  |  |  |  |  |  |  |  |  |  |  |  |  |  |  |  |  |  |  |  |  |  |  |  |  |  |  |  |  |  |  |  |  |  |  |  |  |  |  |  |  |  |  |  |  |  |  |  |  |  |  |  |  |  |  |  |  |  |  |  |  |  |  |  |  |  |  |  |  |  |  |  |  |  |  |  |  |  |  |  |  |  |  |  |  |  |  |  |  |  |  |  |  |  |  |  |  |  |  |  |  |  |  |  |  |  |  |  |  |  |  |  |  |  |  |  |  |  |  |  |  |  |  |  |  |  |  |  |  |  |  |  |  |  |  |  |  |  |  |  |  |  |  |  |  |  |  |  |  |  |  |  |  |  |  |  |  |  |  |  |  |  |  |  |  |  |  |  |  |  |  |  |  |  |  |  |  |  |  |  |  |  |  |  |  |  |  |  |  |  |  |  |  |  |  |  |  |  |  |  |  |  |  |  |  |  |  |  |  |  |  |  |  |  |  |  |  |  |  |  |  |  |  |  |  |  |  |  |  |  |  |  |  |  |  |  |  |  |  |  |  |  |  |  |  |  |  |  |  |  |  |  |  |  |  |  |  |  |  |  |  |  |  |  |  |  |  |  |  |  |  |  |  |  |  |  |  |  |  |  |  |  |  |  |  |  |  |  |  |  |  |  |  |  |  |  |  |  |  |  |  |  |  |  |  |  |  |  |  |  |  |  |  |  |  |  |  |  |  |  |  |  |  |  |  |  |  |  |  |  |  |  |  |  |  |  |  |  |  |  |  |  |  |  |  |  |  |  |  |  |  |  |  |  |  |  |  |  |  |  |  |  |  |  |  |  |  |  |  |  |  |  |  |  |  |  |  |  |  |  |  |  |  |  |  |  |  |  |  |  |  |  |  |  |  |  |  |  |  |  |  |  |  |  |  |  |  |  |  |  |  |  |  |  |  |  |  |  |  |  |  |  |  |  |  |  |  |  |  |  |  |  |  |  |  |  |  |  |  |  |  |  |  |  |  |  |  |  |  |  |  |  |  |  |  |  |  |  |  |  |  |  |  |  |  |  |  |  |  |  |  |  |  |  |  |  |  |  |  |  |  |  |  |  |  |  |  |  |  |  |  |  |  |  |  |  |  |  |  |  |  |  |  |  |  |  |  |  |  |  |  |  |  |  |  |  |  |  |  |  |  |  |  |  |  |  |  |  |  |  |  |  |  |  |  |  |  |  |  |  |  |  |  |  |  |  |  |  |  |  |  |  |  |  |  |  |  |  |  |  |  |  |  |  |  |  |  |  |  |  |  |  |  |  |  |  |  |  |  |  |  |  |  |  |  |  |  |  |  |  |  |  |  |  |  |  |  |  |  |  |  |  |  |  |  |  |  |  |  |  |  |  |  |  |  |  |  |  |  |  |  |  |  |  |  |  |  |  |  |  |  |  |  |  |  |  |  |  |  |  |  |  |  |  |  |  |  |  |  |  |  |  |  |  |  |  |  |  |  |  |  |  |  |  |  |  |  |  |  |  |  |  |  |  |  |  |  |  |  |  |  |  |  |  |  |  |  |  |  |  |  |  |  |  |  |  |  |  |  |  |  |  |  |  |  |  |  |  |  |  |  |  |  |  |  |  |  |  |  |  |  |  |  |  |  |  |  |  |  |  |  |  |  |  |  |  |  |  |  |  |  |  |  |  |  |  |  |  |  |  |  |  |  |  |  |  |  |  |  |  |  |  |  |  |  |  |  |  |  |  |  |  |  |  |  |  |  |  |  |  |  |  |  |  |  |  |  |  |  |  |  |  |  |  |  |  |  |  |  |  |  |  |  |  |  |  |  |  |  |  |  |  |  |  |  |  |  |  |  |  |  |  |  |  |  |  |  |  |  |  |  |  |  |  |  |  |  |  |  |  |  |  |  |  |  |  |  |  |  |  |  |  |  |  |  |  |  |  |  |  |  |  |  |  |  |  |  |  |  |  |  |  |  |  |  |  |  |  |  |  |  |  |  |  |  |  |  |  |  |  |  |  |  |  |  |  |  |  |  |  |  |  |  |  |  |  |  |  |  |  |  |  |  |  |  |  |  |  |  |  |  |  |  |  |  |  |  |  |  |  |  |  |  |  |  |  |  |  |  |  |  |  |  |  |  |  |  |  |  |  |  |  |  |  |  |  |  |  |  |  |  |  |  |  |  |  |  |  |  |  |  |  |  |  |  |  |  |  |  |  |  |  |  |  |  |  |  |  |  |  |  |  |  |  |  |  |  |  |  |  |  |  |  |  |  |  |  |  |  |  |  |  |  |  |  |  |  |  |  |  |  |  |  |  |  |  |  |  |  |  |  |  |  |  |  |  |  |  |  |  |  |  |  |  |  |  |  |  |  |  |  |  |  |  |  |  |  |  |  |  |  |  |  |  |  |  |  |  |  |  |  |  |  |  |  |  |  |  |  |  |  |  |  |  |  |  |  |  |  |  |  |  |  |  |  |  |  |  |  |  |  |  |  |  |  |  |  |  |  |  |  |  |  |  |  |  |  |  |  |  |  |  |  |  |  |  |  |  |  |  |  |  |  |  |  |  |  |  |  |  |  |  |  |  |  |  |  |  |  |  |  |  |  |  |  |  |  |  |  |  |  |  |  |  |  |  |  |  |  |  |  |  |  |  |  |  |  |  |  |  |  |  |  |  |  |  |  |  |  |  |  |  |  |  |  |  |  |  |  |  |  |  |  |  |  |  |  |  |  |  |  |  |  |  |  |  |  |  |  |  |  |  |  |  |  |  |  |  |  |  |  |  |  |  |  |  |  |  |  |  |  |  |  |  |  |  |  |  |  |  |  |  |  |  |  |  |  |  |  |  |  |  |  |  |  |  |  |  |  |  |  |  |  |  |  |  |  |  |  |  |  |  |  |  |  |  |  |  |  |  |  |  |  |  |  |  |  |  |  |  |  |  |  |  |  |  |  |  |  |  |  |  |  |  |  |  |  |  |  |  |  |  |  |  |  |  |  |  |  |  |  |  |  |  |  |  |  |  |  |  |  |  |  |  |  |  |  |  |  |  |  |  |  |  |  |  |  |  |  |  |  |  |  |  |  |  |  |  |  |  |  |  |  |  |  |  |  |  |  |  |  |  |  |  |  |  |  |  |  |  |  |  |  |  |  |  |  |  |  |  |  |  |  |  |  |  |  |  |  |  |  |  |  |  |  |  |  |  |  |  |  |  |  |  |  |  |  |  |  |  |  |  |  |  |  |  |  |  |  |  |  |  |  |  |  |  |  |  |  |  |  |  |  |  |  |  |  |  |  |  |  |  |  |  |  |  |  |  |  |  |  |  |  |  |  |  |  |  |  |  |  |  |  |  |  |  |  |  |  |  |  |  |  |  |  |  |  |  |  |  |  |  |  |  |  |  |  |  |  |  |  |  |  |  |  |  |  |  |  |  |  |  |  |  |  |  |  |  |  |  |  |  |  |  |  |  |  |  |  |  |  |  |  |  |  |  |  |  |  |  |  |  |  |  |  |  |  |  |  |  |  |  |  |  |  |  |  |  |  |  |  |  |  |  |  |  |  |  |  |  |  |  |  |  |  |  |  |  |  |  |  |  |  |  |  |  |  |  |  |  |  |  |  |  |  |  |  |  |  |  |  |  |  |  |  |  |  |  |  |  |  |  |  |  |  |  |  |  |  |  |  |  |  |  |  |  |  |  |  |  |  |  |  |  |  |  |  |  |  |  |  |  |  |  |  |  |  |  |  |  |  |  |  |  |  |  |  |  |  |  |  |  |  |  |  |  |  |  |  |  |  |  |  |  |  |  |  |  |  |  |  |  |  |  |  |  |  |  |  |  |  |  |  |  |  |  |  |  |  |  |  |  |  |  |  |  |  |  |  |  |  |  |  |  |  |  |  |  |  |  |  |  |  |  |  |  |  |  |  |  |  |  |  |  |  |  |  |  |  |  |  |  |  |  |  |  |  |  |  |  |  |  |  |  |  |  |  |  |  |  |  |  |  |  |  |  |  |  |  |  |  |  |  |  |  |  |  |  |  |  |  |  |  |  |  |  |  |  |  |  |  |  |  |  |  |  |  |  |  |  |  |  |  |  |  |  |  |  |  |  |  |  |  |  |  |  |  |  |  |  |  |  |  |  |  |  |  |  |  |  |  |  |  |  |  |  |  |  |  |  |  |  |  |  |  |  |  |  |  |  |  |  |  |  |  |  |  |  |  |  |  |  |  |  |  |  |  |  |  |  |  |  |  |  |  |  |  |  |  |  |  |  |  |  |  |  |  |  |  |  |  |  |  |  |  |  |  |  |  |  |  |  |  |  |  |  |  |  |  |  |  |  |  |  |  |  |  |  |  |  |  |  |  |  |  |  |  |  |  |  |  |  |  |  |  |  |  |  |  |  |  |  |  |  |  |  |  |  |  |  |  |  |  |  |  |  |  |  |  |  |  |  |  |  |  |  |  |  |  |  |  |  |  |  |  |  |  |  |  |  |  |  |  |  |  |  |  |  |  |  |  |  |  |  |  |  |  |  |  |  |  |  |  |  |  |  |  |  |  |  |  |  |  |  |  |  |  |  |  |  |  |  |  |  |  |  |  |  |  |  |  |  |  |  |  |  |  |  |  |  |  |  |  |  |  |  |  |  |  |  |  |  |  |  |  |  |  |  |  |  |  |  |  |  |  |  |  |  |  |  |  |  |  |  |  |  |  |  |  |  |  |  |  |  |  |  |  |  |  |  |  |  |  |  |  |  |  |  |  |  |  |  |  |  |  |  |  |  |  |  |  |  |  |  |  |  |  |  |  |  |  |  |  |  |  |  |  |  |  |  |  |  |  |  |  |  |  |  |  |  |  |  |  |  |  |  |  |  |  |  |  |  |  |  |  |  |  |  |  |  |  |  |  |  |  |  |  |  |  |  |  |  |  |  |  |  |  |  |  |  |  |  |  |  |  |  |  |  |  |  |  |  |  |  |  |  |  |  |  |  |  |  |  |  |  |  |  |  |  |  |  |  |  |  |  |  |  |  |  |  |  |  |  |  |  |  |  |  |  |  |  |  |  |  |  |  |  |  |  |  |  |  |  |  |  |  |  |  |  |  |  |  |  |  |  |  |  |  |  |  |  |  |  |  |  |  |  |  |  |  |  |  |  |  |  |  |  |  |  |  |  |  |  |  |  |  |  |  |  |  |  |  |  |  |  |  |  |  |  |  |  |  |  |  |  |  |  |  |  |  |  |  |  |  |  |  |  |  |  |  |  |  |  |  |  |  |  |  |  |  |  |  |  |  |  |  |  |  |  |  |  |  |  |  |  |  |  |  |  |  |  |  |  |  |  |  |  |  |  |  |  |  |  |  |  |  |  |  |  |  |  |  |  |  |  |  |  |  |  |  |  |  |  |  |  |  |  |  |  |  |  |  |  |  |  |  |  |  |  |  |  |  |  |  |  |  |  |  |  |  |  |  |  |  |  |  |  |  |  |  |  |  |  |  |  |  |  |  |  |  |  |  |  |  |  |  |  |  |  |  |  |  |  |  |  |  |  |  |  |  |  |  |  |  |  |  |  |  |  |  |  |  |  |  |  |  |  |  |  |  |  |  |  |  |  |  |  |  |  |  |  |  |  |  |  |  |  |  |  |  |  |  |  |  |  |  |  |  |  |  |  |  |  |  |  |  |  |  |  |  |  |  |  |  |  |  |  |  |  |  |  |  |  |  |  |  |  |  |  |  |  |  |  |  |  |  |  |  |  |  |  |  |  |  |  |  |  |  |  |  |  |  |  |  |  |  |  |  |  |  |  |  |  |  |  |  |  |  |  |  |  |  |  |  |  |  |  |  |  |  |  |  |  |  |  |  |  |  |  |  |  |  |  |  |  |  |  |  |  |  |  |  |  |  |  |  |  |  |  |  |  |  |  |  |  |  |  |  |  |  |  |  |  |  |  |  |  |  |  |  |  |  |  |  |  |  |  |  |  |  |  |  |  |  |  |  |  |  |  |  |  |  |  |  |  |  |  |  |  |  |  |  |  |  |  |  |  |  |  |  |  |  |  |  |  |  |  |  |  |  |  |  |  |  |  |  |  |  |  |  |  |  |  |  |  |  |  |  |  |  |  |  |  |  |  |  |  |  |  |  |  |  |  |  |  |  |  |  |  |  |  |  |  |  |  |  |  |  |  |  |  |  |  |  |  |  |  |  |  |  |  |  |  |  |  |  |  |  |  |  |  |  |  |  |  |  |  |  |  |  |  |  |  |  |  |  |  |  |  |  |  |  |  |  |  |  |  |  |  |  |  |  |  |  |  |  |  |  |  |  |  |  |  |  |  |  |  |  |  |  |  |  |  |  |  |  |  |  |  |  |  |  |  |  |  |  |  |  |  |  |  |  |  |  |  |  |  |  |  |  |  |  |  |  |  |  |  |  |  |  |  |  |  |  |  |  |  |  |  |  |  |  |  |  |  |  |  |  |  |  |  |  |  |  |  |  |  |  |  |  |  |  |  |  |  |  |  |  |  |  |  |  |  |  |  |  |  |  |  |  |  |  |  |  |  |  |  |  |  |  |  |  |  |  |  |  |  |  |  |  |  |  |  |  |  |  |  |  |  |  |  |  |  |  |  |  |  |  |  |  |  |  |  |  |  |  |  |  |  |  |  |  |  |  |  |  |  |  |  |  |  |  |  |  |  |  |  |  |  |  |  |  |  |  |  |  |  |  |  |  |  |  |  |  |  |  |  |  |  |  |  |  |  |  |  |  |  |  |  |  |  |  |  |  |  |  |  |  |  |  |  |  |  |  |  |  |  |  |  |  |  |  |  |  |  |  |  |  |  |  |  |  |  |  |  |  |  |  |  |  |  |  |  |  |  |  |  |  |  |  |  |  |  |  |  |  |  |  |  |  |  |  |  |  |  |  |  |  |  |  |  |  |  |  |  |  |  |  |  |  |  |  |  |  |  |  |  |  |  |  |  |  |  |  |  |  |  |  |  |  |  |  |  |  |  |  |  |  |  |  |  |  |  |  |  |  |  |  |  |  |  |  |  |  |  |  |  |  |  |  |  |  |  |  |  |  |  |  |  |  |  |  |  |  |  |  |  |  |  |  |  |  |  |  |  |  |  |  |  |  |  |  |  |  |  |  |  |  |  |  |  |  |  |  |  |  |  |  |  |  |  |  |  |  |  |  |  |  |  |  |  |  |  |  |  |  |  |  |  |  |  |  |  |  |  |  |  |  |  |  |  |  |  |  |  |  |  |  |  |  |  |  |  |  |  |  |  |  |  |  |  |  |  |  |  |  |  |  |  |  |  |  |  |  |  |  |  |  |  |  |  |  |  |  |  |  |  |  |  |  |  |  |  |  |  |  |  |  |  |  |  |  |  |  |  |  |  |  |  |  |  |  |  |  |  |  |  |  |  |  |  |  |  |  |  |  |  |  |  |  |  |  |  |  |  |  |  |  |  |  |  |  |  |  |  |  |  |  |  |  |  |  |  |  |  |  |  |  |  |  |  |  |  |  |  |  |  |  |  |  |  |  |  |  |  |  |  |  |  |  |  |  |  |  |  |  |  |  |  |  |  |  |  |  |  |  |  |  |  |  |  |  |  |  |  |  |  |  |  |  |  |  |  |  |  |  |  |  |  |  |  |  |  |  |  |  |  |  |  |  |  |  |  |  |  |  |  |  |  |  |  |  |  |  |  |  |  |  |  |  |  |  |  |  |  |  |  |  |  |  |  |  |  |  |  |  |  |  |  |  |  |  |  |  |  |  |  |  |  |  |  |  |  |  |  |  |  |  |  |  |  |  |  |  |  |  |  |  |  |  |  |  |  |  |  |  |  |  |  |  |  |  |  |  |  |  |  |  |  |  |  |  |  |  |  |  |  |  |  |  |  |  |  |  |  |  |  |  |  |  |  |  |  |  |  |  |  |  |  |  |  |  |  |  |  |  |  |  |  |  |  |  |  |  |  |  |  |  |  |  |  |  |  |  |  |  |  |  |  |  |  |  |  |  |  |  |  |  |  |  |  |  |  |  |  |  |  |  |  |  |  |  |  |  |  |  |  |  |  |  |  |  |  |  |  |  |  |  |  |  |  |  |  |  |  |  |  |  |  |  |  |  |  |  |  |  |  |  |  |  |  |  |  |  |  |  |  |  |  |  |  |  |  |  |  |  |  |  |  |  |  |  |  |  |  |  |  |  |  |  |  |  |  |  |  |  |  |  |  |  |  |  |  |  |  |  |  |  |  |  |  |  |  |  |  |  |  |  |  |  |  |  |  |  |  |  |  |  |  |  |  |  |  |  |  |  |  |  |  |  |  |  |  |  |  |  |  |  |  |  |  |  |  |  |  |  |  |  |  |  |  |  |  |  |  |  |  |  |  |  |  |  |  |  |  |  |  |  |  |  |  |  |  |  |  |  |  |  |  |  |  |  |  |  |  |  |  |  |  |  |  |  |  |  |  |  |  |  |  |  |  |  |  |  |  |  |  |  |  |  |  |  |  |  |  |  |  |  |  |  |  |  |  |  |  |  |  |  |  |  |  |  |  |  |  |  |  |  |  |  |  |  |  |  |  |  |  |  |  |  |  |  |  |  |  |  |  |  |  |  |  |  |  |  |  |  |  |  |  |  |  |  |  |  |  |  |  |  |  |  |  |  |  |  |  |  |  |  |  |  |  |  |  |  |  |  |  |  |  |  |  |  |  |  |  |  |  |  |  |  |  |  |  |  |  |  |  |  |  |  |  |  |  |  |  |  |  |  |  |  |  |  |  |  |  |  |  |  |  |  |  |  |  |  |  |  |  |  |  |  |  |  |  |  |  |  |  |  |  |  |  |  |  |  |  |  |  |  |  |  |  |  |  |  |  |  |  |  |  |  |  |  |  |  |  |  |  |  |  |  |  |  |  |  |  |  |  |  |  |  |  |  |  |  |  |  |  |  |  |  |  |  |  |  |  |  |  |  |  |  |  |  |  |  |  |  |  |  |  |  |  |  |  |  |  |  |  |  |  |  |  |  |  |  |  |  |  |  |  |  |  |  |  |  |  |  |  |  |  |  |  |  |  |  |  |  |  |  |  |  |  |  |  |  |  |  |  |  |  |  |  |  |  |  |  |  |  |  |  |  |  |  |  |  |  |  |  |  |  |  |  |  |  |  |  |  |  |  |  |  |  |  |  |  |  |  |  |  |  |  |  |  |  |  |  |  |  |  |  |  |  |  |  |  |  |  |  |  |  |  |  |  |  |  |  |  |  |  |  |  |  |  |  |  |  |  |  |  |  |  |  |  |  |  |  |  |  |  |  |  |  |  |  |  |  |  |  |  |  |  |  |  |  |  |  |  |  |  |  |  |  |  |  |  |  |  |  |  |  |  |  |  |  |  |  |  |  |  |  |  |  |  |  |  |  |  |  |  |  |  |  |  |  |  |  |  |  |  |  |  |  |  |  |  |  |  |  |  |  |  |  |  |  |  |  |  |  |  |  |  |  |  |  |  |  |  |  |  |  |  |  |  |  |  |  |  |  |  |  |  |  |  |  |  |  |  |  |  |  |  |  |  |  |  |  |  |  |  |  |  |  |  |  |  |  |  |  |  |  |  |  |  |  |  |  |  |  |  |  |  |  |  |  |  |  |  |  |  |  |  |  |  |  |  |  |  |  |  |  |  |  |  |  |  |  |  |  |  |  |  |  |  |  |  |  |  |  |  |  |  |  |  |  |  |  |  |  |  |  |  |  |  |  |  |  |  |  |  |  |  |  |  |  |  |  |  |  |  |  |  |  |  |  |  |  |  |  |  |  |  |  |  |  |  |  |  |  |  |  |  |  |  |  |  |  |  |  |  |  |  |  |  |  |  |  |  |  |  |  |  |  |  |  |  |  |  |  |  |  |  |  |  |  |  |  |  |  |  |  |  |  |  |  |  |  |  |  |  |  |  |  |  |  |  |  |  |  |  |  |  |  |  |  |  |  |  |  |  |  |  |  |  |  |  |  |  |  |  |  |  |  |  |  |  |  |  |  |  |  |  |  |  |  |  |  |  |  |  |  |  |  |  |  |  |  |  |  |  |  |  |  |  |  |  |  |  |  |  |  |  |  |  |  |  |  |  |  |  |  |  |  |  |  |  |  |  |  |  |  |  |  |  |  |  |  |  |  |  |  |  |  |  |  |  |  |  |  |  |  |  |  |  |  |  |  |  |  |  |  |  |  |  |  |  |  |  |  |  |  |  |  |  |  |  |  |  |  |  |  |  |  |  |  |  |  |  |  |  |  |  |  |  |  |  |  |  |  |  |  |  |  |  |  |  |  |  |  |  |  |  |  |  |  |  |  |  |  |  |  |  |  |  |  |  |  |  |  |  |  |  |  |  |  |  |  |  |  |  |  |  |  |  |  |  |  |  |  |  |  |  |  |  |  |  |  |  |  |  |  |  |  |  |  |  |  |  |  |  |  |  |  |  |  |  |  |  |  |  |  |  |  |  |  |  |  |  |  |  |  |  |  |  |  |  |  |  |  |  |  |  |  |  |  |  |  |  |  |  |  |  |  |  |  |  |  |  |  |  |  |  |  |  |  |  |  |  |  |  |  |  |  |  |  |  |  |  |  |  |  |  |  |  |  |  |  |  |  |  |  |  |  |  |  |  |  |  |  |  |  |  |  |  |  |  |  |  |  |  |  |  |  |  |  |  |  |  |  |  |  |  |  |  |  |  |  |  |  |  |  |  |  |  |  |  |  |  |  |  |  |  |  |  |  |  |  |  |  |  |  |  |  |  |  |  |  |  |  |  |  |  |  |  |  |  |  |  |  |  |  |  |  |  |  |  |  |  |  |  |  |  |  |  |  |  |  |  |  |  |  |  |  |  |  |  |  |  |  |  |  |  |  |  |  |  |  |  |  |  |  |  |  |  |  |  |  |  |  |  |  |  |  |  |  |  |  |  |  |  |  |  |  |  |  |  |  |  |  |  |  |  |  |  |  |  |  |  |  |  |  |  |  |  |  |  |  |  |  |  |  |  |  |  |  |  |  |  |  |  |  |  |  |  |  |  |  |  |  |  |  |  |  |  |  |  |  |  |  |  |  |  |  |  |  |  |  |  |  |  |  |  |  |  |  |  |  |  |  |  |  |  |  |  |  |  |  |  |  |  |  |  |  |  |  |  |  |  |  |  |  |  |  |  |  |  |  |  |  |  |  |  |  |  |
| BIS d=0 cl.8 sym=1 env=1 | - | - | - | - | - | - | - | - | - | - | - | - | - | - | - | - | - | - | - | - | - | - | - | - | - | - | - | - | - | - | - | - | - | - | - | - | - | - | - | - | - | - | - | - | - | - | - | - | - | - | - | - | - | - | - | - | - | - | - | - | - | - | - | - | - | - | - | - | - | - | - | - | - | - | - | - | - | - | - | - | - | - | - | - | - | - | - | - | - | - | - | - | - | - | - | - | - | - | - | - | - | - | - | - | - | - | - | - | - | - | - | - | - | - | - | - | - | - | - | - | - | - | - | - | - | - | - | - | - | - | - | - | - | - | - | - | - | - | - | - | - | - | - | - | - | - | - | - | - | - | - | - | - | - | - | - | - | - | - | - | - | - | - | - | - | - | - | - | - | - | - | - | - | - | - | - | - | - | - | - | - | - | - | - | - | - | - | - | - | - | - | - | - | - | - | - | - | - | - | - | - | - | - | - | - | - | - | - | - | - | - | - | - | - | - | - | - | - | - | - | - | - | - | - | - | - | - | - | - | - | - | - | - | - | - | - | - | - | - | - | - | - | - | - | - | - | - | - | - | - | - | - | - | - | - | - | - | - | - | - | - | - | - | - | - | - | - | - | - | - | - | - | - | - | - | - | - | - | - | - | - | - | - | - | - | - | - | - | - | - | - | - | - | - | - | - | - | - | - | - | - | - | - | - | - | - | - | - | - | - | - | - | - | - | - | - | - | - | - | - | - | - | - | - | - | - | - | - | - | - | - | - | - | - | - | - | - | - | - | - | - | - | - | - | - | - | - | - | - | - | - | - | - | - | - | - | - | - | - | - | - | - | - | - | - | - | - | - | - | - | - | - | - | - | - | - | - | - | - | - | - | - | - | - | - | - | - | - | - | - | - | - | - | - | - | - | - | - | - | - | - | - | - | - | - | - | - | - | - | - | - | - | - | - | - | - | - | - | - | - | - | - | - | - | - | - | - | - | - | - | - | - | - | - | - | - | - | - | - | - | - | - | - | - | - | - | - | - | - | - | - | - | - | - | - | - | - | - | - | - | - | - | - | - | - | - | - | - | - | - | - | - | - | - | - | - | - | - | - | - | - | - | - | - | - | - | - | - | - | - | - | - | - | - | - | - | - | - | - | - | - | - | - | - | - | - | - | - | - | - | - | - | - | - | - | - | - | - | - | - | - | - | - | - | - | - | - | - | - | - | - | - | - | - | - | - | - | - | - | - | - | - | - | - | - | - | - | - | - | - | - | - | - | - | - | - | - | - | - | - | - | - | - | - | - | - | - | - | - | - | - | - | - | - | - | - | - | - | - | - | - | - | - | - | - | - | - | - | - | - | - | - | - | - | - | - | - | - | - | - | - | - | - | - | - | - | - | - | - | - | - | - | - | - | - | - | - | - | - | - | - | - | - | - | - | - | - | - | - | - | - | - | - | - | - | - | - | - | - | - | - | - | - | - | - | - | - | - | - | - | - | - | - | - | - | - | - | - | - | - | - | - | - | - | - | - | - | - | - | - | - | - | - | - | - | - | - | - | - | - | - | - | - | - | - | - | - | - | - | - | - | - | - | - | - | - | - | - | - | - | - | - | - | - | - | - | - | - | - | - | - | - | - | - | - | - | - | - | - | - | - | - | - | - | - | - | - | - | - | - | - | - | - | - | - | - | - | - | - | - | - | - | - | - | - | - | - | - | - | - | - | - | - | - | - | - | - | - | - | - | - | - | - | - | - | - | - | - | - | - | - | - | - | - | - | - | - | - | - | - | - | - | - | - | - | - | - | - | - | - | - | - | - | - | - | - | - | - | - | - | - | - | - | - | - | - | - | - | - | - | - | - | - | - | - | - | - | - | - | - | - | - | - | - | - | - | - | - | - | - | - | - | - | - | - | - | - | - | - | - | - | - | - | - | - | - | - | - | - | - | - | - | - | - | - | - | - | - | - | - | - | - | - | - | - | - | - | - | - | - | - | - | - | - | - | - | - | - | - | - | - | - | - | - | - | - | - | - | - | - | - | - | - | - | - | - | - | - | - | - | - | - | - | - | - | - | - | - | - | - | - | - | - | - | - | - | - | - | - | - | - | - | - | - | - | - | - | - | - | - | - | - | - | - | - | - | - | - | - | - | - | - | - | - | - | - | - | - | - | - | - | - | - | - | - | - | - | - | - | - | - | - | - | - | - | - | - | - | - | - | - | - | - | - | - | - | - | - | - | - | - | - | - | - | - | - | - | - | - | - | - | - | - | - | - | - | - | - | - | - | - | - | - | - | - | - | - | - | - | - | - | - | - | - | - | - | - | - | - | - | - | - | - | - | - | - | - | - | - | - | - | - | - | - | - | - | - | - | - | - | - | - | - | - | - | - | - | - | - | - | - | - | - | - | - | - | - | - | - | - | - | - | - | - | - | - | - | - | - | - | - | - | - | - | - | - | - | - | - | - | - | - | - | - | - | - | - | - | - | - | - | - | - | - | - | - | - | - | - | - | - | - | - | - | - | - | - | - | - | - | - | - | - | - | - | - | - | - | - | - | - | - | - | - | - | - | - | - | - | - | - | - | - | - | - | - | - | - | - | - | - | - | - | - | - | - | - | - | - | - | - | - | - | - | - | - | - | - | - | - | - | - | - | - | - | - | - | - | - | - | - | - | - | - | - | - | - | - | - | - | - | - | - | - | - | - | - | - | - | - | - | - | - | - | - | - | - | - | - | - | - | - | - | - | - | - | - | - | - | - | - | - | - | - | - | - | - | - | - | - | - | - | - | - | - | - | - | - | - | - | - | - | - | - | - | - | - | - | - | - | - | - | - | - | - | - | - | - | - | - | - | - | - | - | - | - | - | - | - | - | - | - | - | - | - | - | - | - | - | - | - | - | - | - | - | - | - | - | - | - | - | - | - | - | - | - | - | - | - | - | - | - | - | - | - | - | - | - | - | - | - | - | - | - | - | - | - | - | - | - | - | - | - | - | - | - | - | - | - | - | - | - | - | - | - | - | - | - | - | - | - | - | - | - | - | - | - | - | - | - | - | - | - | - | - | - | - | - | - | - | - | - | - | - | - | - | - | - | - | - | - | - | - | - | - | - | - | - | - | - | - | - | - | - | - | - | - | - | - | - | - | - | - | - | - | - | - | - | - | - | - | - | - | - | - | - | - | - | - | - | - | - | - | - | - | - | - | - | - | - | - | - | - | - | - | - | - | - | - | - | - | - | - | - | - | - | - | - | - | - | - | - | - | - | - | - | - | - | - | - | - | - | - | - | - | - | - | - | - | - | - | - | - | - | - | - | - | - | - | - | - | - | - | - | - | - | - | - | - | - | - | - | - | - | - | - | - | - | - | - | - | - | - | - | - | - | - | - | - | - | - | - | - | - | - | - | - | - | - | - | - | - | - | - | - | - | - | - | - | - | - | - | - | - | - | - | - | - | - | - | - | - | - | - | - | - | - | - | - | - | - | - | - | - | - | - | - | - | - | - | - | - | - | - | - | - | - | - | - | - | - | - | - | - | - | - | - | - | - | - | - | - | - | - | - | - | - | - | - | - | - | - | - | - | - | - | - | - | - | - | - | - | - | - | - | - | - | - | - | - | - | - | - | - | - | - | - | - | - | - | - | - | - | - | - | - | - | - | - | - | - | - | - | - | - | - | - | - | - | - | - | - | - | - | - | - | - | - | - | - | - | - | - | - | - | - | - | - | - | - | - | - | - | - | - | - | - | - | - | - | - | - | - | - | - | - | - | - | - | - | - | - | - | - | - | - | - | - | - | - | - | - | - | - | - | - | - | - | - | - | - | - | - | - | - | - | - | - | - | - | - | - | - | - | - | - | - | - | - | - | - | - | - | - | - | - | - | - | - | - | - | - | - | - | - | - | - | - | - | - | - | - | - | - | - | - | - | - | - | - | - | - | - | - | - | - | - | - | - | - | - | - | - | - | - | - | - | - | - | - | - | - | - | - | - | - | - | - | - | - | - | - | - | - | - | - | - | - | - | - | - | - | - | - | - | - | - | - | - | - | - | - | - | - | - | - | - | - | - | - | - | - | - | - | - | - | - | - | - | - | - | - | - | - | - | - | - | - | - | - | - | - | - | - | - | - | - | - | - | - | - | - | - | - | - | - | - | - | - | - | - | - | - | - | - | - | - | - | - | - | - | - | - | - | - | - | - | - | - | - | - | - | - | - | - | - | - | - | - | - | - | - | - | - | - | - | - | - | - | - | - | - | - | - | - | - | - | - | - | - | - | - | - | - | - | - | - | - | - | - | - | - | - | - | - | - | - | - | - | - | - | - | - | - | - | - | - | - | - | - | - | - | - | - | - | - | - | - | - | - | - | - | - | - | - | - | - | - | - | - | - | - | - | - | - | - | - | - | - | - | - | - | - | - | - | - | - | - | - | - | - | - | - | - | - | - | - | - | - | - | - | - | - | - | - | - | - | - | - | - | - | - | - | - | - | - | - | - | - | - | - | - | - | - | - | - | - | - | - | - | - | - | - | - | - | - | - | - | - | - | - | - | - | - | - | - | - | - | - | - | - | - | - | - | - | - | - | - | - | - | - | - | - | - | - | - | - | - | - | - | - | - | - | - | - | - | - | - | - | - | - | - | - | - | - | - | - | - | - | - | - | - | - | - | - | - | - | - | - | - | - | - | - | - | - | - | - | - | - | - | - | - | - | - | - | - | - | - | - | - | - | - | - | - | - | - | - | - | - | - | - | - | - | - | - | - | - | - | - | - | - | - | - | - | - | - | - | - | - | - | - | - | - | - | - | - | - | - | - | - | - | - | - | - | - | - | - | - | - | - | - | - | - | - | - | - | - | - | - | - | - | - | - | - | - | - | - | - | - | - | - | - | - | - | - | - | - | - | - | - | - | - | - | - | - | - | - | - | - | - | - | - | - | - | - | - | - | - | - | - | - | - | - | - | - | - | - | - | - | - | - | - | - | - | - | - | - | - | - | - | - | - | - | - | - | - | - | - | - | - | - | - | - | - | - | - | - | - | - | - | - | - | - | - | - | - | - | - | - | - | - | - | - | - | - | - | - | - | - | - | - | - | - | - | - | - | - | - | - | - | - | - | - | - | - | - | - | - | - | - | - | - | - | - | - | - | - | - | - | - | - | - | - | - | - | - | - | - | - | - | - | - | - | - | - | - | - | - | - | - | - | - | - | - | - | - | - | - | - | - | - | - | - | - | - | - | - | - | - | - | - | - | - | - | - | - | - | - | - | - | - | - | - | - | - | - | - | - | - | - | - | - | - | - | - | - | - | - | - | - | - | - | - | - | - | - | - | - | - | - | - | - | - | - | - | - | - | - | - | - | - | - | - | - | - | - | - | - | - | - | - | - | - | - | - | - | - | - | - | - | - | - | - | - | - | - | - | - | - | - | - | - | - | - | - | - | - | - | - | - | - | - | - | - | - | - | - | - | - | - | - | - | - | - | - | - | - | - | - | - | - | - | - | - | - | - | - | - | - | - | - | - | - | - | - | - | - | - | - | - | - | - | - | - | - | - | - | - | - | - | - | - | - | - | - | - | - | - | - | - | - | - | - | - | - | - | - | - | - | - | - | - | - | - | - | - | - | - | - | - | - | - | - | - | - | - | - | - | - | - | - | - | - | - | - | - | - | - | - | - | - | - | - | - | - | - | - | - | - | - | - | - | - | - | - | - | - | - | - | - | - | - | - | - | - | - | - | - | - | - | - | - | - | - | - | - | - | - | - | - | - | - | - | - | - | - | - | - | - | - | - | - | - | - | - | - | - | - | - | - | - | - | - | - | - | - | - | - | - | - | - | - | - | - | - | - | - | - | - | - | - | - | - | - | - | - | - | - | - | - | - | - | - | - | - | - | - | - | - | - | - | - | - | - | - | - | - | - | - | - | - | - | - | - | - | - | - | - | - | - | - | - | - | - | - | - | - | - | - | - | - | - | - | - | - | - | - | - | - | - | - | - | - | - | - | - | - | - | - | - | - | - | - | - | - | - | - | - | - | - | - | - | - | - | - | - | - | - | - | - | - | - | - | - | - | - | - | - | - | - | - | - | - | - | - | - | - | - | - | - | - | - | - | - | - | - | - | - | - | - | - | - | - | - | - | - | - | - | - | - | - | - | - | - | - | - | - | - | - | - | - | - | - | - | - | - | - | - | - | - | - | - | - | - | - | - | - | - | - | - | - | - | - | - | - | - | - | - | - | - | - | - | - | - | - | - | - | - | - | - | - | - | - | - | - | - | - | - | - | - | - | - | - | - | - | - | - | - | - | - | - | - | - | - | - | - | - | - | - | - | - | - | - | - | - | - | - | - | - | - | - | - | - | - | - | - | - | - | - | - | - | - | - | - | - | - | - | - | - | - | - | - | - | - | - | - | - | - | - | - | - | - | - | - | - | - | - | - | - | - | - | - | - | - | - | - | - | - | - | - | - | - | - | - | - | - | - | - | - | - | - | - | - | - | - | - | - | - | - | - | - | - | - | - | - | - | - | - | - | - | - | - | - | - | - | - | - | - | - | - | - | - | - | - | - | - | - | - | - | - | - | - | - | - | - | - | - | - | - | - | - | - | - | - | - | - | - | - | - | - | - | - | - | - | - | - | - | - | - | - | - | - | - | - | - | - | - | - | - | - | - | - | - | - | - | - | - | - | - | - | - | - | - | - | - | - | - | - | - | - | - | - | - | - | - | - | - | - | - | - | - | - | - | - | - | - | - | - | - | - | - | - | - | - | - | - | - | - | - | - | - | - | - | - | - | - | - | - | - | - | - | - | - | - | - | - | - | - | - | - | - | - | - | - | - | - | - | - | - | - | - | - | - | - | - | - | - | - | - | - | - | - | - | - | - | - | - | - | - | - | - | - | - | - | - | - | - | - | - | - | - | - | - | - | - | - | - | - | - | - | - | - | - | - | - | - | - | - | - | - | - | - | - | - | - | - | - | - | - | - | - | - | - | - | - | - | - | - | - | - | - | - | - | - | - | - | - | - | - | - | - | - | - | - | - | - | - | - | - | - | - | - | - | - | - | - | - | - | - | - | - | - | - | - | - | - | - | - | - | - | - | - | - | - | - | - | - | - | - | - | - | - | - | - | - | - | - | - | - | - | - | - | - | - | - | - | - | - | - | - | - | - | - | - | - | - | - | - | - | - | - | - | - | - | - | - | - | - | - | - | - | - | - | - | - | - | - | - | - | - | - | - | - | - | - | - | - | - | - | - | - | - | - | - | - | - | - | - | - | - | - | - | - | - | - | - | - | - | - | - | - | - | - | - | - | - | - | - | - | - | - | - | - | - | - | - | - | - | - | - | - | - | - | - | - | - | - | - | - | - | - | - | - | - | - | - | - | - | - | - | - | - | - | - | - | - | - | - | - | - | - | - | - | - | - | - | - | - | - | - | - | - | - | - | - | - | - | - | - | - | - | - | - | - | - | - | - | - | - | - | - | - | - | - | - | - | - | - | - | - | - | - | - | - | - | - | - | - | - | - | - | - | - | - | - | - | - | - | - | - | - | - | - | - | - | - | - | - | - | - | - | - | - | - | - | - | - | - | - | - | - | - | - | - | - | - | - | - | - | - | - | - | - | - | - | - | - | - | - | - | - | - | - | - | - | - | - | - | - | - | - | - | - | - | - | - | - | - | - | - | - | - | - | - | - | - | - | - | - | - | - | - | - | - | - | - | - | - | - | - | - | - | - | - | - | - | - | - | - | - | - | - | - | - | - | - | - | - | - | - | - | - | - | - | - | - | - | - | - | - | - | - | - | - | - | - | - | - | - | - | - | - | - | - | - | - | - | - | - | - | - | - | - | - | - | - | - | - | - | - | - | - | - | - | - | - | - | - | - | - | - | - | - | - | - | - | - | - | - | - | - | - | - | - | - | - | - | - | - | - | - | - | - | - | - | - | - | - | - | - | - | - | - | - | - | - | - | - | - | - | - | - | - | - | - | - | - | - | - | - | - | - | - | - | - | - | - | - | - | - | - | - | - | - | - | - | - | - | - | - | - | - | - | - | - | - | - | - | - | - | - | - | - | - | - | - | - | - | - | - | - | - | - | - | - | - | - | - | - | - | - | - | - | - | - | - | - | - | - | - | - | - | - | - | - | - | - | - | - | - | - | - | - | - | - | - | - | - | - | - | - | - | - | - | - | - | - | - | - | - | - | - | - | - | - | - | - | - | - | - | - | - | - | - | - | - | - | - | - | - | - | - | - | - | - | - | - | - | - | - | - | - | - | - | - | - | - | - | - | - | - | - | - | - | - | - | - | - | - | - | - | - | - | - | - | - | - | - | - | - | - | - | - | - | - | - | - | - | - | - | - | - | - | - | - | - | - | - | - | - | - | - | - | - | - | - | - | - | - | - | - | - | - | - | - | - | - | - | - | - | - | - | - | - | - | - | - | - | - | - | - | - | - | - | - | - | - | - | - | - | - | - | - | - | - | - | - | - | - | - | - | - | - | - | - | - | - | - | - | - | - | - | - | - | - | - | - | - | - | - | - | - | - | - | - | - | - | - | - | - | - | - | - | - | - | - | - | - | - | - | - | - | - | - | - | - | - | - | - | - | - | - | - | - | - | - | - | - | - | - | - | - | - | - | - | - | - | - | - | - | - | - | - | - | - | - | - | - | - | - | - | - | - | - | - | - | - | - | - | - | - | - | - | - | - | - | - | - | - | - | - | - | - | - | - | - | - | - | - | - | - | - | - | - | - | - | - | - | - | - | - | - | - | - | - | - | - | - | - | - | - | - | - | - | - | - | - | - | - | - | - | - | - | - | - | - | - | - | - | - | - | - | - | - | - | - | - | - | - | - | - | - | - | - | - | - | - | - | - | - | - | - | - | - | - | - | - | - | - | - | - | - | - | - | - | - | - | - | - | - | - | - | - | - | - | - | - | - | - | - | - | - | - | - | - | - | - | - | - | - | - | - | - | - | - | - | - | - | - | - | - | - | - | - | - | - | - | - | - | - | - | - | - | - | - | - | - | - | - | - | - | - | - | - | - | - | - | - | - | - | - | - | - | - | - | - | - | - | - | - | - | - | - | - | - | - | - | - | - | - | - | - | - | - | - | - | - | - | - | - | - | - | - | - | - | - | - | - | - | - | - | - | - | - | - | - | - | - | - | - | - | - | - | - | - | - | - | - | - | - | - | - | - | - | - | - | - | - | - | - | - | - | - | - | - | - | - | - | - | - | - | - | - | - | - | - | - | - | - | - | - | - | - | - | - | - | - | - | - | - | - | - | - | - | - | - | - | - | - | - | - | - | - | - | - | - | - | - | - | - | - | - | - | - | - | - | - | - | - | - | - | - | - | - | - | - | - | - | - | - | - | - | - | - | - | - | - | - | - | - | - | - | - | - | - | - | - | - | - | - | - | - | - | - | - | - | - | - | - | - | - | - | - | - | - | - | - | - | - | - | - | - | - | - | - | - | - | - | - | - | - | - | - | - | - | - | - | - | - | - | - | - | - | - | - | - | - | - | - | - | - | - | - | - | - | - | - | - | - | - | - | - | - | - | - | - | - | - | - | - | - | - | - | - | - | - | - | - | - | - | - | - | - | - | - | - | - | - | - | - | - | - | - | - | - | - | - | - | - | - | - | - | - | - | - | - | - | - | - | - | - | - | - | - | - | - | - | - | - | - | - | - | - | - | - | - | - | - | - | - | - | - | - | - | - | - | - | - | - | - | - | - | - | - | - | - | - | - | - | - | - | - | - | - | - | - | - | - | - | - | - | - | - | - | - | - | - | - | - | - | - | - | - | - | - | - | - | - | - | - | - | - | - | - | - | - | - | - | - | - | - | - | - | - | - | - | - | - | - | - | - | - | - | - | - | - | - | - | - | - | - | - | - | - | - | - | - | - | - | - | - | H | x | x | x | H | x | x | x | x | x | x | x | H | H | x | x | H | H | - | - | - | - | - | - | - | - | - | - | - | - | - | - | - | - | - | - | - | - | - | - | - | - | - | - | - | - | - | - | - | - | - | - | - | - | - | - | - | - | - | - | - | - | - | - | - | - | - | - | - | - | - | - | - | - | - | - | - | - | - | - | - | - | - | - | - | - | - | - | - | - | - | - | - | - | - | - | - | - | - | - | - | - | - | - | - | - | - | - | - | - | - | - | - | - | - | - | - | - | - | - | - | - | - | - | - | - | - | - | - | - | - | - | - | - | - | - | - | - | - | - | - | - | - | - | - | - | - | - | - | - | - | - | - | - | - | - | - | - | - | - | - | - | - | - | - | - | - | - | - | - | - | - | - | - | - | - | - | - | - | - | - | - | - | - | - | - | - | - | - | - | - | - | - | - | - | - | - | - | - | - | - | - | - | - | - | - | - | - | - | - | - | - | - | - | - | - | - | - | - | - | - | - | - | - | - | - | - | - | - | - | - | - | - | - | - | - | - | - | - | - | - | - | - | - | - | - | - | - | - | - | - | - | - | - | - | - | - | - | - | - | - | - | - | - | - | - | - | - | - | - | - | - | - | - | - | - | - | - | - | - | - | - | - | - | - | - | - | - | - | - | - | - | - | - | - | - | - | - | - | - | - | - | - | - | - | - | - | - | - | - | - | - | - | - | - | - | - | - | - | - | - | - | - | - | - | - | - | - | - | - | - | - | - | - | - | - | - | - | - | - | - | - | - | - | - | - | - | - | - | - | - | - | - | - | - | - | - | - | - | - | - | - | - | - | - | - | - | - | - | - | - | - | - | - | - | - | - | - | - | - | - | - | - | - | - | - | - | - | - | - | - | - | - | - | - | - | - | - | - | - | - | - | - | - | - | - | - | - | - | - | - | - | - | - | - | - | - | - | - | - | - | - | - | - | - | - | - | - | - | - | - | - | - | - | - | - | - | - | - | - | - | - | - | - | - | - | - | - | - | - | - | - | - | - | - | - | - | - | - | - | - | - | - | - | - | - | - | - | - | - | - | - | - | - | - | - | - | - | - | - | - | - | - | - | - | - | - | - | - | - | - | - | - | - | - | - | - | - | - | - | - | - | - | - | - | - | - | - | - | - | - | - | - | - | - | - | - | - | - | - | - | - | - | - | - | - | - | - | - | - | - | - | - | - | - | - | - | - | - | - | - | - | - | - | - | - | - | - | - | - | - | - | - | - | - | - | - | - | - | - | - | - | - | - | - | - | - | - | - | - | - | - | - | - | - | - | - | - | - | - | - | - | - | - | - | - | - | - | - | - | - | - | - | - | - | - | - | - | - | - | - | - | - | - | - | - | - | - | - | - | - | - | - | - | - | - | - | - | - | - | - | - | - | - | - | - | - | - | - | - | - | - | - | - | - | - | - | - | - | - | - | - | - | - | - | - | - | - | - | - | - | - | - | - | - | - | - | - | - | - | - | - | - | - | - | - | - | - | - | - | - | - | - | - | - | - | - | - | - | - | - | - | - | - | - | - | - | - | - | - | - | - | - | - | - | - | - | - | - | - | - | - | - | - | - | - | - | - | - | - | - | - | - | - | - | - | - | - | - | - | - | - | - | - | - | - | - | - | - | - | - | - | - | - | - | - | - | - | - | - | - | - | - | - | - | - | - | - | - | - | - | - | - | - | - | - | - | - | - | - | - | - | - | - | - | - | - | - | - | - | - | - | - | - | - | - | - | - | - | - | - | - | - | - | - | - | - | - | - | - | - | - | - | - | - | - | - | - | - | - | - | - | - | - | - | - | - | - | - | - | - | - | - | - | - | - | - | - | - | - | - | - | - | - | - | - | - | - | - | - | - | - | - | - | - | - | - | - | - | - | - | - | - | - | - | - | - | - | - | - | - | - | - | - | - | - | - | - | - | - | - | - | - | - | - | - | - | - | - | - | - | - | - | - | - | - | - | - | - | - | - | - | - | - | - | - | - | - | - | - | - | - | - | - | - | - | - | - | - | - | - | - | - | - | - | - | - | - | - | - | - | - | - | - | - | - | - | - | - | - | - | - | - | - | - | - | - | - | - | - | - | - | - | - | - | - | - | - | - | - | - | - | - | - | - | - | - | - | - | - | - | - | - | - | - | - | - | - | - | - | - | - | - | - | - | - | - | - | - | - | - | - | - | - | - | - | - | - | - | - | - | - | - | - | - | - | - | - | - | - | - | - | - | - | - | - | - | - | - | - | - | - | - | - | - | - | - | - | - | - | - | - | - | - | - | - | - | - | - | - | - | - | - | - | - | - | - | - | - | - | - | - | - | - | - | - | - | - | - | - | - | - | - | - | - | - | - |
| BIS d=0 cl.9 sym=1 env=1 | - | - | - | - | - | - | - | - | - | - | - | - | - | - | - | - | - | - | - | - | - | - | - | - | - | - | - | - | - | - | - | - | - | - | - | - | - | - | - | - | - | - | - | - | - | - | - | - | - | - | - | - | - | - | - | - | - | - | - | - | - | - | - | - | - | - | - | - | - | - | - | - | - | - | - | - | - | - | - | - | - | - | - | - | - | - | - | - | - | - | - | - | - | - | - | - | - | - | - | - | - | - | - | - | - | - | - | - | - | - | - | - | - | - | - | - | - | - | - | - | - | - | - | - | - | - | - | - | - | - | - | - | - | - | - | - | - | - | - | - | - | - | - | - | - | - | - | - | - | - | - | - | - | - | - | - | - | - | - | - | - | - | - | - | - | - | - | - | - | - | - | - | - | - | - | - | - | - | - | - | - | - | - | - | - | - | - | - | - | - | - | - | - | - | - | - | - | - | - | - | - | - | - | - | - | - | - | - | - | - | - | - | - | - | - | - | - | - | - | - | - | - | - | - | - | - | - | - | - | - | - | - | - | - | - | - | - | - | - | - | - | - | - | - | - | - | - | - | - | - | - | - | - | - | - | - | - | - | - | - | - | - | - | - | - | - | - | - | - | - | - | - | - | - | - | - | - | - | - | - | - | - | - | - | - | - | - | - | - | - | - | - | - | - | - | - | - | - | - | - | - | - | - | - | - | - | - | - | - | - | - | - | - | - | - | - | - | - | - | - | - | - | - | - | - | - | - | - | - | - | - | - | - | - | - | - | - | - | - | - | - | - | - | - | - | - | - | - | - | - | - | - | - | - | - | - | - | - | - | - | - | - | - | - | - | - | - | - | - | - | - | - | - | - | - | - | - | - | - | - | - | - | - | - | - | - | - | - | - | - | - | - | - | - | - | - | - | - | - | - | - | - | - | - | - | - | - | - | - | - | - | - | - | - | - | - | - | - | H | E | x | x | x | x | x | x | x | x | x | x | x | H | E | - | - | - | - | - | - | - | - | - | - | - | - | - | - | - | - | - | - | - | - | - | - | - | - | - | - | - | - | - | - | - | - | - | - | - | - | - | - | - | - | - | - | - | - | - | - | - | - | - | - | - | - | - | - | - | - | - | - | - | - | - | - | - | - | - | - | - | - | - | - | - | - | - | - | - | - | - | - | - | - | - | - | - | - | - | - | - | - | - | - | - | - | - | - | - | - | - | - | - | - | - | - | - | - | - | - | - | - | - | - | - | - | - | - | - | - | - | - | - | - | - | - | - | - | - | - | - | - | - | - | - | - | - | - | - | - | - | - | - | - | - | - | - | - | - | - | - | - | - | - | - | - | - | - | - | - | - | - | - | - | - | - | - | - | - | - | - | - | - | - | - | - | - | - | - | - | - | - | - | - | - | - | - | - | - | - | - | - | - | - | - | - | - | - | - | - | - | - | - | - | - | - | - | - | - | - | - | - | - | - | - | - | - | - | - | - | - | - | - | - | - | - | - | - | - | - | - | - | - | - | - | - | - | - | - | - | - | - | - | - | - | - | - | - | - | - | - | - | - | - | - | - | - | - | - | - | - | - | - | - | - | - | - | - | - | - | - | - | - | - | - | - | - | - | - | - | - | - | - | - | - | - | - | - | - | - | - | - | - | - | - | - | - | - | - | - | - | - | - | - | - | - | - | - | - | - | - | - | - | - | - | - | - | - | - | - | - | - | - | - | - | - | - | - | - | - | - | - | - | - | - | - | - | - | - | - | - | - | - | - | - | - | - | - | - | - | - | - | - | - | - | - | - | - | - | - | - | - | - | - | - | - | - | - | - | - | - | - | - | - | - | - | - | - | - | - | - | - | - | - | - | - | - | - | - | - | - | - | - | - | - | - | - | - | - | - | - | - | - | - | - | - | - | - | - | - | - | - | - | - | - | - | - | - | - | - | - | - | - | - | - | - | - | - | - | - | - | - | - | - | - | - | - | - | - | - | - | - | - | - | - | - | - | - | - | - | - | - | - | - | - | - | - | - | - | - | - | - | - | - | - | - | - | - | - | - | - | - | - | - | - | - | - | - | - | - | - | - | - | - | - | - | - | - | - | - | - | - | - | - | - | - | - | - | - | - | - | - | - | - | - | - | - | - | - | - | - | - | - | - | - | - | - | - | - | - | - | - | - | - | - | - | - | - | - | - | - | - | - | - | - | - | - | - | - | - | - | - | - | - | - | - | - | - | - | - | - | - | - | - | - | - | - | - | - | - | - | - | - | - | - | - | - | - | - | - | - | - | - | - | - | - | - | - | - | - | - | - | - | - | - | - | - | - | - | - | - | - | - | - | - | - | - | - | - | - | - | - | - | - | - | - | - | - | - | - | - | - | - | - | - | - | - | - | - | - | - | - | - | - | - | - | - | - | - | - | - | - | - | - | - | - | - | - | - | - | - | - | - | - | - | - | - | - | - | - | - | - | - | - | - | - | - | - | - | - | - | - | - | - | - | - | - | - | - | - | - | - | - | - | - | - | - | - | - | - | - | - | - | - | - | - | - | - | - | - | - | - | - | - | - | - | - | - | - | - | - | - | - | - | - | - | - | - | - | - | - | - | - | - | - | - | - | - | - | - | - | - | - | - | - | - | - | - | - | - | - | - | - | - | - | - | - | - | - | - | - | - | - | - | - | - | - | - | - | - | - | - | - | - | - | - | - | - | - | - | - | - | - | - | - | - | - | - | - | - | - | - | - | - | - | - | - | - | - | - | - | - | - | - | - | - | - | - | - | - | - | - | - | - | - | - | - | - | - | - | - | - | - | - | - | - | - | - | - | - | - | - | - | - | - | - | - | - | - | - | - | - | - | - | - | - | - | - | - | - | - | - | - | - | - | - | - | - | - | - | - | - | - | - | - | - | - | - | - | - | - | - | - | - | - | - | - | - | - | - | - | - | - | - | - | - | - | - | - | - | - | - | - | - | - | - | - | - | - | - | - | - | - | - | - | - | - | - | - | - | - | - | - | - | - | - | - | - | - | - | - | - | - | - | - | - | - | - | - | - | - | - | - | - | - | - | - | - | - | - | - | - | - | - | - | - | - | - | - | - | - | - | - | - | - | - | - | - | - | - | - | - | - | - | - | - | - | - | - | - | - | - | - | - | - | - | - | - | - | - | - | - | - | - | - | - | - | - | - | - | - | - | - | - | - | - | - | - | - | - | - | - | - | - | - | - | - | - | - | - | - | - | - | - | - | - | - | - | - | - | - | - | - | - | - | - | - | - | - | - | - | - | - | - | - | - | - | - | - | - | - | - | - | - | - | - | - | - | - | - | - | - | - | - | - | - | - | - | - | - | - | - | - | - | - | - | - | - | - | - | - | - | - | - | - | - | - | - | - | - | - | - | - | - | - | - | - | - | - | - | - | - | - | - | - | - | - | - | - | - | - | - | - | - | - | - | - | - | - | - | - | - | - | - | - | - | - | - | - | - | - | - | - | - | - | - | - | - | - | - | - | - | - | - | - | - | - | - | - | - | - | - | - | - | - | - | - | - | - | - | - | - | - | - | - | - | - | - | - | - | - | - | - | - | - | - | - | - | - | - | - | - | - | - | - | - | - | - | - | - | - | - | - | - | - | - | - | - | - | - | - | - | - | - | - | - | - | - | - | - | - | - | - | - | - | - | - | - | - | - | - | - | - | - | - | - | - | - | - | - | - | - | - | - | - | - | - | - | - | - | - | - | - | - | - | - | - | - | - | - | - | - | - | - | - | - | - | - | - | - | - | - | - | - | - | - | - | - | - | - | - | - | - | - | - | - | - | - | - | - | - | - | - | - | - | - | - | - | - | - | - | - | - | - | - | - | - | - | - | - | - | - | - | - | - | - | - | - | - | - | - | - | - | - | - | - | - | - | - | - | - | - | - | - | - | - | - | - | - | - | - | - | - | - | - | - | - | - | - | - | - | - | - | - | - | - | - | - | - | - | - | - | - | - | - | - | - | - | - | - | - | - | - | - | - | - | - | - | - | - | - | - | - | - | - | - | - | - | - | - | - | - | - | - | - | - | - | - | - | - | - | - | - | - | - | - | - | - | - | - | - | - | - | - | - | - | - | - | - | - | - | - | - | - | - | - | - | - | - | - | - | - | - | - | - | - | - | - | - | - | - | - | - | - | - | - | - | - | - | - | - | - | - | - | - | - | - | - | - | - | - | - | - | - | - | - | - | - | - | - | - | - | - | - | - | - | - | - | - | - | - | - | - | - | - | - | - | - | - | - | - | - | - | - | - | - | - | - | - | - | - | - | - | - | - | - | - | - | - | - | - | - | - | - | - | - | - | - | - | - | - | - | - | - | - | - | - | - | - | - | - | - | - | - | - | - | - | - | - | - | - | - | - | - | - | - | - | - | - | - | - | - | - | - | - | - | - | - | - | - | - | - | - | - | - | - | - | - | - | - | - | - | - | - | - | - | - | - | - | - | - | - | - | - | - | - | - | - | - | - | - | - | - | - | - | - | - | - | - | - | - | - | - | - | - | - | - | - | - | - | - | - | - | - | - | - | - | - | - | - | - | - | - | - | - | - | - | - | - | - | - | - | - | - | - | - | - | - | - | - | - | - | - | - | - | - | - | - | - | - | - | - | - | - | - | - | - | - | - | - | - | - | - | - | - | - | - | - | - | - | - | - | - | - | - | - | - | - | - | - | - | - | - | - | - | - | - | - | - | - | - | - | - | - | - | - | - | - | - | - | - | - | - | - | - | - | - | - | - | - | - | - | - | - | - | - | - | - | - | - | - | - | - | - | - | - | - | - | - | - | - | - | - | - | - | - | - | - | - | - | - | - | - | - | - | - | - | - | - | - | - | - | - | - | - | - | - | - | - | - | - | - | - | - | - | - | - | - | - | - | - | - | - | - | - | - | - | - | - | - | - | - | - | - | - | - | - | - | - | - | - | - | - | - | - | - | - | - | - | - | - | - | - | - | - | - | - | - | - | - | - | - | - | - | - | - | - | - | - | - | - | - | - | - | - | - | - | - | - | - | - | - | - | - | - | - | - | - | - | - | - | - | - | - | - | - | - | - | - | - | - | - | - | - | - | - | - | - | - | - | - | - | - | - | - | - | - | - | - | - | - | - | - | - | - | - | - | - | - | - | - | - | - | - | - | - | - | - | - | - | - | - | - | - | - | - | - | - | - | - | - | - | - | - | - | - | - | - | - | - | - | - | - | - | - | - | - | - | - | - | - | - | - | - | - | - | - | - | - | - | - | - | - | - | - | - | - | - | - | - | - | - | - | - | - | - | - | - | - | - | - | - | - | - | - | - | - | - | - | - | - | - | - | - | - | - | - | - | - | - | - | - | - | - | - | - | - | - | - | - | - | - | - | - | - | - | - | - | - | - | - | - | - | - | - | - | - | - | - | - | - | - | - | - | - | - | - | - | - | - | - | - | - | - | - | - | - | - | - | - | - | - | - | - | - | - | - | - | - | - | - | - | - | - | - | - | - | - | - | - | - | - | - | - | - | - | - | - | - | - | - | - | - | - | - | - | - | - | - | - | - | - | - | - | - | - | - | - | - | - | - | - | - | - | - | - | - | - | - | - | - | - | - | - | - | - | - | - | - | - | - | - | - | - | - | - | - | - | - | - | - | - | - | - | - | - | - | - | - | - | - | - | - | - | - | - | - | - | - | - | - | - | - | - | - | - | - | - | - | - | - | - | - | - | - | - | - | - | - | - | - | - | - | - | - | - | - | - | - | - | - | - | - | - | - | - | - | - | - | - | - | - | - | - | - | - | - | - | - | - | - | - | - | - | - | - | - | - | - | - | - | - | - | - | - | - | - | - | - | - | - | - | - | - | - | - | - | - | - | - | - | - | - | - | - | - | - | - | - | - | - | - | - | - | - | - | - | - | - | - | - | - | - | - | - | - | - | - | - | - | - | - | - | - | - | - | - | - | - | - | - | - | - | - | - | - | - | - | - | - | - | - | - | - | - | - | - | - | - | - | - | - | - | - | - | - | - | - | - | - | - | - | - | - | - | - | - | - | - | - | - | - | - | - | - | - | - | - | - | - | - | - | - | - | - | - | - | - | - | - | - | - | - | - | - | - | - | - | - | - | - | - | - | - | - | - | - | - | - | - | - | - | - | - | - | - | - | - | - | - | - | - | - | - | - | - | - | - | - | - | - | - | - | - | - | - | - | - | - | - | - | - | - | - | - | - | - | - | - | - | - | - | - | - | - | - | - | - | - | - | - | - | - | - | - | - | - | - | - | - | - | - | - | - | - | - | - | - | - | - | - | - | - | - | - | - | - | - | - | - | - | - | - | - | - | - | - | - | - | - | - | - | - | - | - | - | - | - | - | - | - | - | - | - | - | - | - | - | - | - | - | - | - | - | - | - | - | - | - | - | - | - | - | - | - | - | - | - | - | - | - | - | - | - | - | - | - | - | - | - | - | - | - | - | - | - | - | - | - | - | - | - | - | - | - | - | - | - | - | - | - | - | - | - | - | - | - | - | - | - | - | - | - | - | - | - | - | - | - | - | - | - | - | - | - | - | - | - | - | - | - | - | - | - | - | - | - | - | - | - | - | - | - | - | - | - | - | - | - | - | - | - | - | - | - | - | - | - | - | - | - | - | - | - | - | - | - | - | - | - | - | - | - | - | - | - | - | - | - | - | - | - | - | - | - | - | - | - | - | - | - | - | - | - | - | - | - | - | - | - | - | - | - | - | - | - | - | - | - | - | - | - | - | - | - | - | - | - | - | - | - | - | - | - | - | - | - | - | - | - | - | - | - | - | - | - | - | - | - | - | - | - | - | - | - | - | - | - | - | - | - | - | - | - | - | - | - | - | - | - | - | - | - | - | - | - | - | - | - | - | - | - | - | - | - | - | - | - | - | - | - | - | - | - | - | - | - | - | - | - | - | - | - | - | - | - | - | - | - | - | - | - | - | - | - | - | - | - | - | - | - | - | - | - | - | - | - | - | - | - | - | - | - | - | - | - | - | - | - | - | - | - | - | - | - | - | - | - | - | - | - | - | - | - | - | - | - | - | - | - | - | - | - | - | - | - | - | - | - | - | - | - | - | - | - | - | - | - | - | - | - | - | - | - | - | - | - | - | - | - | - | - | - | - | - | - | - | - | - | - | - | - | - | - | - | - | - | - | - | - | - | - | - | - | - | - | - | - | - | - | - | - | - | - | - | - | - | - | - | - | - | - | - | - | - | - | - | - | - | - | - | - | - | - | - | - | - | - | - | - | - | - | - | - | - | - | - | - | - | - | - | - | - | - | - | - | - | - | - | - | - | - | - | - | - | - | - | - | - | - | - | - | - | - | - | - | - | - | - | - | - | - | - | - | - | - | - | - | - | - | - | - | - | - | - | - | - | - | - | - | - | - | - | - | - | - | - | - | - | - | - | - | - | - | - | - | - | - | - | - | - | - | - | - | - | - | - | - | - | - | - | - | - | - | - | - | - | - | - | - | - | - | - | - | - | - | - | - | - | - | - | - | - | - | - | - | - | - | - | - | - | - | - | - | - | - | - | - | - | - | - | - | - | - | - | - | - | - | - | - | - | - | - | - | - | - | - | - | - | - | - | - | - | - | - | - | - | - | - | - | - | - | - | - | - | - | - | - | - | - | - | - | - | - | - | - | - | - | - | - | - | - | - | - | - | - | - | - | - | - | - | - | - | - | - | - | - | - | - | - | - | - | - | - | - | - | - | - | - | - | - | - | - | - | - | - | - | - | - | - | - | - | - | - | - | - | - | - | - | - | - | - | - | - | - | - | - | - | - | - | - | - | - | - | - | - | - | - | - | - | - | - | - | - | - | - | - | - | - | - | - | - | - | - | - | - | - | - | - | - | - | - | - | - | - | - | - | - | - | - | - | - | - | - | - | - | - | - | - | - | - | - | - | - | - | - | - | - | - | - | - | - | - | - | - | - | - | - | - | - | - | - | - | - | - | - | - | - | - | - | - | - | - | - | - | - | - | - | - | - | - | - | - | - | - | - | - | - | - | - | - | - | - | - | - | - | - | - | - | - | - | - | - | - | - | - | - | - | - | - | - | - | - | - | - | - | - | - | - | - | - | - | - | - | - | - | - | - | - | - | - | - | - | - | - | - | - | - | - | - | - | - | - | - | - | - | - | - | - | - | - | - | - | - | - | - | - | - | - | - | - | - | - | - | - | - | - | - | - | - | - | - | - | - | - | - | - | - | - | - | - | - | - | - | - | - | - | - | - | - | - | - | - | - | - | - | - | - | - | - | - | - | - | - | - | - | - | - | - | - | - | - | - | - | - | - | - | - | - | - | - | - | - | - | - | - | - | - | - | - | - | - | - | - | - | - | - | - | - | - | - | - | - | - | - | - | - | - | - | - | - | - | - | - | - | - | - | - | - | - | - | - | - | - | - | - | - | - | - | - | - | - | - | - | - | - | - | - | - | - | - | - | - | - | - | - | - | - | - | - | - | - | - | - | - | - | - | - | - | - | - | - | - | - | - | - | - | - | - | - | - | - | - | - | - | - | - | - | - | - | - | - | - | - | - | - | - | - | - | - | - | - | - | - | - | - | - | - | - | - | - | - | - | - | - | - | - | - | - | - | - | - | - | - | - | - | - | - | - | - | - | - | - | - | - | - | - | - | - | - | - | - | - | - | - | - | - | - | - | - | - | - | - | - | - | - | - | - | - | - | - | - | - | - | - | - | - | - | - | - | - | - | - | - | - | - | - | - | - | - | - | - | - | - | - | - | - | - | - | - | - | - | - | - | - | - | - | - | - | - | - | - | - | - | - | - | - | - | - | - | - | - | - | - | - | - | - | - | - | - | - | - | - | - | - | - | - | - | - | - | - | - | - | - | - | - | - | - | - | - | - | - | - | - | - | - | - | - | - | - | - | - | - | - | - | - | - | - | - | - | - | - | - | - | - | - | - | - | - | - | - | - | - | - | - | - | - | - | - | - | - | - | - | - | - | - | - | - | - | - | - | - | - | - | - | - | - | - | - | - | - | - | - | - | - | - | - | - | - | - | - | - | - | - | - | - | - | - | - | - | - | - | - | - | - | - | - | - | - | - | - | - | - | - | - | - | - | - | - | - | - | - | - | - | - | - | - | - | - | - | - | - | - | - | - | - | - | - | - | - | - | - | - | - | - | - | - | - | - | - | - | - | - | - | - | - | - | - | - | - | - | - | - | - | - | - | - | - | - | - | - | - | - | - | - | - | - | - | - | - | - | - | - | - | - | - | - | - | - | - | - | - | - | - | - | - | - | - | - | - | - | - | - | - | - | - | - | - | - | - | - | - | - | - | - | - | - | - | - | - | - | - | - | - | - | - | - | - | - | - | - | - | - | - | - | - | - | - | - | - | - | - | - | - | - | - | - | - | - | - | - | - | - | - | - | - | - | - | - | - | - | - | - | - | - | - | - | - | - | - | - | - | - | - | - | - | - | - | - | - | - | - | - | - | - | - | - | - | - | - | - | - | - | - | - | - | - | - | - | - | - | - | - | - | - | - | - | - | - | - | - | - | - | - | - | - | - | - | - | - | - | - | - | - | - | - | - | - | - | - | - | - | - | - | - | - | - | - | - | - | - | - | - | - | - | - | - | - | - | - | - | - | - | - | - | - | - | - | - | - | - | - | - | - | - | - | - | - | - | - | - | - | - | - | - | - | - | - | - | - | - | - | - | - | - | - | - | - | - | - | - | - | - | - | - | - | - | - | - | - | - | - | - | - | - | - | - | - | - | - | - | - | - | - | - | - | - | - | - | - | - | - | - | - | - | - | - | - | - | - | - | - | - | - | - | - | - | - | - | - | - | - | - | - | - | - | - | - | - | - | - | - | - | - | - | - | - | - | - | - | - | - | - | - | - | - | - | - | - | - | - | - | - | - | - | - | - | - | - | - | - | - | - | - | - | - | - | - | - | - | - | - | - | - | - | - | - | - | - | - | - | - | - | - | - | - | - | - | - | - | - | - | - | - | - | - | - | - | - | - | - | - | - | - | - | - | - | - | - | - | - | - | - | - | - | - | - | - | - | - | - | - | - | - | - | - | - | - | - | - | - | - | - | - | - | - | - | - | - | - | - | - | - | - | - | - | - | - | - | - | - | - | - | - | - | - | - | - | - | - | - | - | - | - | - | - | - | - | - | - | - | - | - | - | - | - | - | - | - | - | - | - | - | - | - | - | - | - | - | - | - | - | - | - | - | - | - | - | - | - | - | - | - | - | - | - | - | - | - | - | - | - | - | - | - | - | - | - | - | - | - | - | - | - | - | - | - | - | - | - | - | - | - | - | - | - | - | - | - | - | - | - | - | - | - | - | - | - | - | - | - | - | - | - | - | - | - | - | - | - | - | - | - | - | - | - | - | - | - | - | - | - | - | - | - | - | - | - | - | - | - | - | - | - | - | - | - | - | - | - | - | - | - | - | - | - | - | - | - | - | - | - | - | - | - | - | - | - | - | - | - | - | - | - | - | - | - | - | - | - | - | - | - | - | - | - | - | - | - | - | - | - | - | - | - | - | - | - | - | - | - | - | - | - | - | - | - | - | - | - | - | - | - | - | - | - | - | - | - | - | - | - | - | - | - | - | - | - | - | - | - | - | - | - | - | - | - | - | - | - | - | - | - | - | - | - | - | - | - | - | - | - | - | - | - | - | - | - | - | - | - | - | - | - | - | - | - | - | - | - | - | - | - | - | - | - | - | - | - | - | - | - | - | - | - | - | - | - | - | - | - | - | - | - | - | - | - | - | - | - | - | - | - | - | - | - | - | - | - | - | - | - | - | - | - | - | - | - | - | - | - | - | - | - | - | - | - | - | - | - | - | - | - | - | - | - | - | - | - | - | - | - | - | - | - | - | - | - | - | - | - | - | - | - | - | - | - | - | - | - | - | - | - | - | - | - | - | - | - | - | - | - | - | - | - | - | - | - | - | - | - | - | - | - | - | - | - | - | - | - | - | - | - | - | - | - | - | - | - | - | - | - | - | - | - | - | - | - | - | - | - | - | - | - | - | - | - | - | - | - | - | - | - | - | - | - | - | - | - | - | - | - | - | - | - | - | - | - | - | - | - | - | - | - | - | - | - | - | - | - | - | - | - | - | - | - | - | - | - | - | - | - | - | - | - | - | - | - | - | - | - | - | - | - | - | - | - | - | - | - | - | - | - | - | - | - | - | - | - | - | - | - | - | - | - | - | - | - | - | - | - | - | - | - | - | - | - | - | - | - | - | - | - | - | - | - | - | - | - | - | - | - | - | - | - | - | - | - | - | - | - | - | - | - | - | - | - | - | - | - | - | - | - | - | - | - | - | - | - | - | - | - | - | - | - | - | - | - | - | - | - | - | - | - | - | - | - | - | - | - | - | - | - | - | - | - | - | - | - | - | - | - | - | - | - | - | - | - | - | - | - | - | - | - | - | - | - | - | - | - | - | - | - | - | - | - | - | - | - | - | - | - | - | - | - | - | - | - | - | - | - | - | - | - | - | - | - | - | - | - | - | - | - | - | - | - | - | - | - | - | - | - | - | - | - | - | - | - | - | - | - | - | - | - | - | - | - | - | - | - | - | - | - | - | - | - | - | - | - | - | - | - | - | - | - | - | - | - | - | - | - | - | - | - | - | - | - | - | - | - | - | - | - | - | - | - | - | - | - | - | - | - | - | - | - | - | - | - | - | - | - | - | - | - | - | - | - | - | - | - | - | - | - | - | - | - | - | - | - | - | - | - | - | - | - | - | - | - | - | - | - | - | - | - | - | - | - | - | - | - | - | - | - | - | - | - | - | - | - | - | - | - | - | - | - | - | - | - | - | - | - | - | - | - | - | - | - | - | - | - | - | - | - | - | - | - | - | - | - | - | - | - | - | - | - | - | - | - | - | - | - | - | - | - | - | - | - | - | - | - | - | - | - | - | - | - | - | - | - | - | - | - | - | - | - | - | - | - | - | - | - | - | - | - | - | - | - | - | - | - | - | - | - | - | - | - | - | - | - | - | - | - | - | - | - | - | - | - | - | - | - | - | - | - | - | - | - | - | - | - | - | - | - | - | - | - | - | - | - | - | - | - | - | - | - | - | - | - | - | - | - | - | - | - | - | - | - | - | - | - | - | - | - | - | - | - | - | - | - | - | - | - | - | - | - | - | - | - | - | - | - | - | - | - | - | - | - | - | - | - | - | - | - | - | - | - | - | - | - | - | - | - | - | - | - | - | - | - | - | - | - | - | - | - | - | - | - | - | - |
| BIS d=0 cl.5 sym=1 env=1 | - | - | - | - | - | - | - | - | - | - | - | - | - | - | - | - | - | - | - | - | - | - | - | - | - | - | - | - | - | - | - | - | - | - | - | - | - | - | - | - | - | - | - | - | - | - | - | - | - | - | - | - | - | - | - | - | - | - | - | - | - | - | - | - | - | - | - | - | - | - | - | - | - | - | - | - | - | - | - | - | - | - | - | - | - | - | - | - | - | - | - | - | - | - | - | - | - | - | - | - | - | - | - | - | - | - | - | - | - | - | - | - | - | - | - | - | - | - | - | - | - | - | - | - | - | - | - | - | - | - | - | - | - | - | - | - | - | - | - | - | - | - | - | - | - | - | - | - | - | - | - | - | - | - | - | - | - | - | - | - | - | - | - | - | - | - | - | - | - | - | - | - | - | - | - | - | - | - | - | - | - | - | - | - | - | - | - | - | - | - | - | - | - | - | - | - | - | - | - | - | - | - | - | - | - | - | - | - | - | - | - | - | - | - | - | - | - | - | - | - | - | - | - | - | - | - | - | - | - | - | - | - | - | - | - | - | - | - | - | - | - | - | - | - | - | - | - | - | - | - | - | - | - | - | - | - | - | - | - | - | - | - | - | - | - | - | - | - | - | - | - | - | - | - | - | - | - | - | - | - | - | - | - | - | - | - | - | - | - | - | - | - | - | - | - | - | - | - | - | - | - | - | - | - | - | - | - | - | - | - | - | - | - | - | - | - | - | - | - | - | - | - | - | - | - | - | - | - | - | - | - | - | - | - | - | - | - | - | - | - | - | - | - | - | - | - | - | - | - | - | - | - | - | - | - | - | - | - | - | - | - | - | - | - | - | - | - | - | - | - | - | - | - | - | - | - | - | - | - | - | - | - | - | - | - | - | - | - | - | - | - | - | - | - | - | - | - | - | - | - | - | - | - | - | - | - | - | - | - | - | - | - | - | - | - | - | - | - | - | - | - | - | - | - | - | - | - | - | - | - | - | - | - | - | - | - | - | - | - | - | - | - | - | - | - | - | - | - | - | - | - | - | - | - | - | - | - | - | - | - | - | - | - | - | - | - | - | - | - | - | - | - | - | - | - | - | - | - | - | - | - | - | - | - | - | - | - | - | - | - | - | - | - | - | - | - | - | - | - | - | - | - | - | - | - | - | - | - | - | - | - | - | - | - | - | - | - | - | - | - | - | - | - | - | - | - | - | - | - | - | - | - | - | - | - | - | - | - | - | - | - | - | - | - | - | - | - | - | - | - | - | - | - | - | - | - | - | - | - | - | - | - | - | - | - | - | - | - | - | - | - | - | - | - | - | - | - | - | - | - | - | - | - | - | - | - | - | - | - | - | - | - | - | - | - | - | - | - | - | - | - | - | - | - | - | - | - | - | - | - | - | - | - | - | - | - | - | - | - | - | - | - | - | - | - | - | - | - | - | - | - | - | - | - | - | - | - | - | - | - | - | - | - | - | - | - | - | - | - | - | - | - | - | - | - | - | - | - | - | - | - | - | - | - | - | - | - | - | - | - | - | - | - | - | - | - | - | - | - | - | - | - | - | - | - | - | - | - | - | - | - | - | - | - | - | - | - | - | - | - | - | - | - | - | - | - | - | - | - | - | - | - | - | - | - | - | - | - | - | - | - | - | - | - | - | - | - | - | - | - | - | - | - | - | - | - | - | - | - | - | - | - | - | - | - | - | - | - | - | - | - | - | - | - | - | - | - | - | - | - | - | - | - | - | - | - | - | - | - | - | - | - | - | - | - | - | - | - | - | - | - | - | - | - | - | - | - | - | - | - | - | - | - | - | - | - | - | - | - | - | - | - | - | - | - | - | - | - | - | - | - | - | - | - | - | - | - | - | - | - | - | - | - | - | - | - | - | - | - | - | - | - | - | - | - | - | - | - | - | - | - | - | - | - | - | - | - | - | - | - | - | - | - | - | - | - | - | - | - | - | - | - | - | - | - | - | - | - | - | - | - | - | - | - | - | - | - | - | - | - | - | - | - | - | - | - | - | - | - | - | - | - | - | - | - | - | - | - | - | - | - | - | - | - | - | - | - | - | - | - | - | - | - | - | - | - | - | - | - | - | - | - | - | - | - | - | - | - | - | - | - | - | - | - | - | - | - | - | - | - | - | - | - | - | - | - | - | - | - | - | - | - | - | - | - | - | - | - | - | - | - | - | - | - | - | - | - | - | - | - | - | - | - | - | - | - | - | - | - | - | - | - | - | - | - | - | - | - | - | - | - | - | - | - | - | - | - | - | - | - | - | - | - | - | - | - | - | - | - | - | - | - | - | - | - | - | - | - | - | - | - | - | - | - | - | - | - | - | - | - | - | - | - | - | - | - | - | - | - | - | - | - | - | - | - | - | - | - | - | - | - | - | - | - | - | - | - | - | - | - | - | - | - | - | - | - | - | - | - | - | - | - | - | - | - | - | - | - | - | - | - | - | - | - | - | - | - | - | - | - | - | - | - | - | - | - | - | - | - | - | - | - | - | - | - | - | - | - | - | - | - | - | - | - | - | - | - | - | - | - | - | - | - | - | - | - | - | - | - | - | - | - | - | - | - | - | - | - | - | - | - | - | - | - | - | - | - | - | - | - | - | - | - | - | - | - | - | - | - | - | - | - | - | - | - | - | - | - | - | - | - | - | - | - | - | - | - | - | - | - | - | - | - | - | - | - | - | - | - | - | - | - | - | - | - | - | - | - | - | - | - | - | - | - | - | - | - | - | - | - | - | - | - | - | - | - | - | - | - | - | - | - | - | - | - | - | - | - | - | - | - | - | - | - | - | - | - | - | - | - | - | - | - | - | - | - | - | - | - | - | - | - | - | - | - | - | - | - | - | - | - | - | - | - | - | - | - | - | - | - | - | - | - | - | - | - | - | - | - | - | - | - | - | - | - | - | - | - | - | - | - | - | - | - | - | - | - | - | - | - | - | - | - | - | - | - | - | - | - | - | - | - | - | - | - | - | - | - | - | - | - | - | - | - | - | - | - | - | - | - | - | - | - | - | - | - | - | - | - | - | - | - | - | - | - | - | - | - | - | - | - | - | - | - | - | - | - | - | - | - | - | - | - | - | - | - | - | - | - | - | - | - | - | - | - | - | - | - | - | - | - | - | - | - | - | - | - | - | - | - | - | - | - | - | - | - | - | - | - | - | - | - | - | - | - | - | - | - | - | - | - | - | - | - | - | - | - | - | - | - | - | - | - | - | - | - | - | - | - | - | - | - | - | - | - | - | - | - | - | - | - | - | - | - | - | - | - | - | - | - | - | - | - | - | - | - | - | - | - | - | - | - | - | - | - | - | - | - | - | - | - | - | - | - | - | - | - | - | - | - | - | - | - | - | - | - | - | - | - | - | - | - | - | - | - | - | - | - | - | - | - | - | - | - | - | - | - | - | - | - | - | - | - | - | - | - | - | - | - | - | - | - | - | - | - | - | - | - | - | - | - | - | - | - | - | - | - | - | - | - | - | - | - | - | - | - | - | - | - | - | - | - | - | - | - | - | - | - | - | - | - | - | - | - | - | - | - | - | - | - | - | - | - | - | - | - | - | - | - | - | - | - | - | - | - | - | - | - | - | - | - | - | - | - | - | - | - | - | - | - | - | - | - | - | - | - | - | - | - | - | - | - | - | - | - | - | - | - | - | - | - | - | - | - | - | - | - | - | - | - | - | - | - | - | - | - | - | - | - | - | - | - | - | - | - | - | - | - | - | - | - | - | - | - | - | - | - | - | - | - | - | - | - | - | - | - | - | - | - | - | - | - | - | - | - | - | - | - | - | - | - | - | - | - | - | - | - | - | - | - | - | - | - | - | - | - | - | - | - | - | - | - | - | - | - | - | - | - | - | - | - | - | - | - | - | - | - | - | - | - | - | - | - | - | - | - | - | - | - | - | - | - | - | - | - | - | - | - | - | - | - | - | - | - | - | - | - | - | - | - | - | - | - | - | - | - | - | - | - | - | - | - | - | - | - | - | - | - | - | - | - | - | - | - | - | - | - | - | - | - | - | - | - | - | - | - | - | - | - | - | - | - | - | - | - | - | - | - | - | - | - | - | - | - | - | - | - | - | - | - | - | - | - | - | - | - | - | - | - | - | - | - | - | - | - | - | - | - | - | - | - | - | - | - | - | - | - | - | - | - | - | - | - | - | - | - | - | - | - | - | - | - | - | - | - | - | - | - | - | - | - | - | - | - | - | - | - | - | - | - | - | - | - | - | - | - | - | - | - | - | - | - | - | - | - | - | - | - | - | - | - | - | - | - | - | - | - | - | - | - | - | - | - | - | - | - | - | - | - | - | - | - | - | - | - | - | - | - | - | - | - | - | - | - | - | - | - | - | - | - | - | - | - | - | - | - | - | - | - | - | - | - | - | - | - | - | - | - | - | - | - | - | - | - | - | - | - | - | - | - | - | - | - | - | - | - | - | - | - | - | - | - | - | - | - | - | - | - | - | - | - | - | - | - | - | - | - | - | - | - | - | - | - | - | - | - | - | - | - | - | - | - | - | - | - | - | - | - | - | - | - | - | - | - | - | - | - | - | - | - | - | - | - | - | - | - | - | - | - | - | - | - | - | - | - | - | - | - | - | - | - | - | - | - | - | - | - | - | - | - | - | - | - | - | - | - | - | - | - | - | - | - | - | - | - | - | - | - | - | - | - | - | - | - | - | - | - | - | - | - | - | - | - | - | - | - | - | - | - | - | - | - | - | - | - | - | - | - | - | - | - | - | - | - | - | - | - | - | - | - | - | - | - | - | - | - | - | - | - | - | - | - | - | - | - | - | - | - | - | - | - | - | - | - | - | - | - | - | - | - | - | - | - | - | - | - | - | - | - | - | - | - | - | - | - | - | - | - | - | - | - | - | - | - | - | - | - | - | - | - | - | - | - | - | - | - | - | - | - | - | - | - | - | - | - | - | - | - | - | - | - | - | - | - | - | - | - | - | - | - | - | - | - | - | - | - | - | - | - | - | - | - | - | - | - | - | - | - | - | - | - | - | - | - | - | - | - | - | - | - | - | - | - | - | - | - | - | - | - | - | - | - | - | - | - | - | - | - | - | - | - | - | - | - | - | - | - | - | - | - | - | - | - | - | - | - | - | - | - | - | - | - | - | - | - | - | - | - | - | - | - | - | - | - | - | - | - | - | - | - | - | - | - | - | - | - | - | - | - | - | - | - | - | - | - | - | - | - | - | - | - | - | - | - | - | - | - | - | - | - | - | - | - | - | - | - | - | - | - | - | - | - | - | - | - | - | - | - | - | - | - | - | - | - | - | - | - | - | - | - | - | - | - | - | - | - | - | - | - | - | - | - | - | - | - | - | - | - | - | - | - | - | - | - | - | - | - | - | - | - | - | - | - | - | - | - | - | - | - | - | - | - | - | - | - | - | - | - | - | - | - | - | - | - | - | - | - | - | - | - | - | - | - | - | - | - | - | - | - | - | - | - | - | - | - | - | - | - | - | - | - | - | - | - | - | - | - | - | - | - | - | - | - | - | - | - | - | - | - | - | - | - | - | - | - | - | - | - | - | - | - | - | - | - | - | - | - | - | - | - | - | - | - | - | - | - | - | - | - | - | - | - | - | - | - | - | - | - | - | - | - | - | - | - | - | - | - | - | - | - | - | - | - | - | - | - | - | - | - | - | - | - | - | - | - | - | - | - | - | - | - | - | - | - | - | - | - | - | - | - | - | - | - | - | - | - | - | - | - | - | - | - | - | - | - | - | - | - | - | - | - | - | - | - | - | - | - | - | - | - | - | - | - | - | - | - | - | - | - | - | - | - | - | - | - | - | - | - | - | - | - | - | - | - | - | - | - | - | - | - | - | - | - | - | - | - | - | - | - | - | - | - | - | - | - | - | - | - | - | - | - | - | - | - | - | - | - | - | - | - | - | - | - | - | - | - | - | - | - | - | - | - | - | - | - | - | - | - | - | - | - | - | - | - | - | - | - | - | - | - | - | - | - | - | - | - | - | - | - | - | - | - | - | - | - | - | - | - | - | - | - | - | - | - | - | - | - | - | - | - | - | - | - | - | - | - | - | - | - | - | - | - | - | - | - | - | - | - | - | - | - | - | - | - | - | - | - | - | - | - | - | - | - | - | - | - | - | - | - | - | - | - | - | - | - | - | - | - | - | - | - | - | - | - | - | - | - | - | - | - | - | - | - | - | - | - | - | - | - | - | - | - | - | - | - | - | - | - | - | - | - | - | - | - | - | - | - | - | - | - | - | - | - | - | - | - | - | - | - | - | - | - | - | - | - | - | - | - | - | - | - | - | - | - | - | - | - | - | - | - | - | - | - | - | - | - | - | - | - | - | - | - | - | - | - | - | - | - | - | - | - | - | - | - | - | - | - | - | - | - | - | - | - | - | - | - | - | - | - | - | - | - | - | - | - | - | - | - | - | - | - | - | - | - | - | - | - | - | - | - | - | - | - | - | - | - | - | - | - | - | - | - | - | - | - | - | - | - | - | - | - | - | - | - | - | - | - | - | - | - | - | - | - | - | - | - | - | - | - | - | - | - | - | - | - | - | - | - | - | - | - | - | - | - | - | - | - | - | - | - | - | - | - | - | - | - | - | - | - | - | - | - | - | - | - | - | - | - | - | - | - | - | - | - | - | - | - | - | - | - | - | - | - | - | - | - | - | - | - | - | - | - | - | - | - | - | - | - | - | - | - | - | - | - | - | - | - | - | - | - | - | - | - | - | - | - | - | - | - | - | - | - | - | - | - | - | - | - | - | - | - | - | - | - | - | - | - | - | - | - | - | - | - | - | - | - | - | - | - | - | - | - | - | - | - | - | - | - | - | - | - | - | - | - | - | - | - | - | - | - | - | - | - | - | - | - | - | - | - | - | - | - | - | - | - | - | - | - | - | - | - | - | - | - | - | - | - | - | - | - | - | - | - | - | - | - | - | - | - | - | - | - | - | - | - | - | - | - | - | - | - | - | - | - | - | - | - | - | - | - | - | - | - | - | - | - | - | - | - | - | - | - | - | - | - | - | - | - | - | - | - | - | - | - | - | - | - | - | - | - | - | - | - | - | - | - | - | - | - | - | - | - | - | - | - | - | - | - | - | - | - | - | - | - | - | - | - | - | - | - | - | - | - | - | - | - | - | - | - | - | - | - | - | - | - | - | - | - | - | - | - | - | - | - | - | - | - | - | - | - | - | - | - | - | - | - | - | - | - | - | - | - | - | - | - | - | - | - | - | - | - | - | - | - | - | - | - | - | - | - | - | - | - | - | - | - | - | - | - | - | - | - | - | - | - | - | - | - | - | - | - | - | - | - | - | - | - | - | - | - | - | - | - | - | - | - | - | - | - | - | - | - | - | - | - | - | - | - | - | - | - | - | - | - | - | - | - | - | - | - | - | - | - | - | - | - | - | - | - | - | - | - | - | - | - | - | - | - | - | - | - | - | - | - | - | - | - | - | - | - | - | - | - | - | - | - | - | - | - | - | - | - | - | - | - | - | - | - | - | - | - | - | - | - | - | - | - | - | - | - | - | - | - | - | - | - | - | - | - | - | - | - | - | - | - | - | - | - | - | - | - | - | - | - | - | - | - | - | - | - | - | - | - | - | - | - | - | - | - | - | - | - | - | - | - | - | - | - | - | - | - | - | - | - | - | - | - | - | - | - | - | - | - | - | - | - | - | - | - | - | - | - | - | - | - | - | - | - | - | - | - | - | - | - | - | - | - | - | - | - | - | - | - | - | - | - | - | - | - | - | - | - | - | - | - | - | - | - | - | - | - | - | - | - | - | - | - | - | - | - | - | - | - | - | - | - | - | - | - | - | - | - | - | - | - | - | - | - | - | - | - | - | - | - | - | - | - | - | - | - | - | - | - | - | - | - | - | - | - | - | - | - | - | - | - | - | - | - | - | - | - | - | - | - | - | - | - | - | - | - | - | - | - | - | - | - | - | - | - | - | - | - | - | - | - | - | - | - | - | - | - | - | - | - | - | - | - | - | - | - | - | - | - | - | - | - | - | - | - | - | - | - | - | - | - | - | - | - | - | - | - | - | - | - | - | - | - | - | - | - | - | - | - | - | - | - | - | - | - | - | - | - | - | - | - | - | - | - | - | - | - | - | - | - | - | - | - | - | - | - | - | - | - | - | - | - | - | - | - | - | - | - | - | - | - | - | - | - | - | - | - | - | - | - | - | - | - | - | - | - | - | - | - | - | - | - | - | - | - | - | - | - | - | - | - | - | - | - | - | - | - | - | - | - | - | - | - | - | - | - | - | - | - | - | - | - | - | - | - | - | - | - | - | - | - | - | - | - | - | - | - | - | - | - | - | - | - | - | - | - | - | - | - | - | - | - | - | - | - | - | - | - | - | - | - | - | - | - | - | - | - | - | - | - | - | - | - | - | - | - | - | - | - | - | - | - | - | - | - | - | - | - | - | - | - | - | - | - | - | - | - | - | - | - | - | - | - | - | - | - | - | - | - | - | - | - | - | - | - | - | - | - | - | - | - | - | - | - | - | - | - | - | - | - | - | - | - | - | - | - | - | - | - | - | - | - | - | - | - | - | - | - | - | - | - | - | - | - | - | - | - | - | - | - | - | - | - | - | - | - | - | - | - | - | - | - | - | - | - | - | - | - | - | - | - | - | - | - | - | - | - | - | - | - | - | - | - | - | - | - | - | - | - | - | - | - | - | - | - | - | - | - | - | - | - | - | - | - | - | - | - | - | - | - | - | - | - | - | - | - | - | - | - | - | - | - | - | - | - | - | - | - | - | - | - | - | - | - | - | - | - | - | - | - | - | - | - | - | - | - | - | - | - | - | - | - | - | - | - | - | - | - | - | - | - | - | - | - | - | - | - | - | - | - | - | - | - | - | - | - | - | - | - | - | - | - | - | - | - | - | - | - | - | - | - | - | - | - | - | - | - | - | - | - | - | - | - | - | - | - | - | - | - | - | - | - | - | - | - | - | - | - | - | - | - | - | - | - | - | - | - | - | - | - | - | - | - | - | - | - | - | - | - | - | - | - | - | - | - | - | - | - | - | - | - | - | - | - | - | - | - | - | - | - | - | - | - | - | - | - | - | - | - | - | - | - | - | - | - | - | - | - | - | - | - | - | - | - | - | - | - | - | - | - | - | - | - | - | - | - | - | - | - | - | - | - | - | - | - | - | - | - | - | - | - | - | - | - | - | - | - | - | - | - | - | - | - | - | - | - | - | - | - | - | - | - | - | - | - | - | - | - | - | - | - | - | - | - | - | - | - | - | - | - | - | - | - | - | - | - | - | - | - | - | - | - | - | - | - | - | - | - | - | - | - | - | - | - | - | - | - | - | - | - | - | - | - | - | - | - | - | - | - | - | - | - | - | - | - | - | - | - | - | - | - | - | - | - | - | - | - | - | - | - | - | - | - | - | - | - | - | - | - | - | - | - | - | - | - | - | - | - | - | - | - | - | - | - | - | - | - | - | - | - | - | - | - | - | - | - | - | - | - | - | - | - | - | - | - | - | - | - | - | - | - | - | - | - | - | - | - | - | - | - | - | - | - | - | - | - | - | - | - | - | - | - | - | - | - | - | - | - | - | - | - | - | - | - | - | - | - | - | - | - | - | - | - | - | - | - | - | - | - | - | - | - | - | - | - | - | - | - | - | - | - | - | - | - | - | - | - | - | - | - | - | - | - | - | - | - | - | - | - | - | - | - | - | - | - | - | - | - | - | - | - | - | - | - | - | - | - | - | - | - | - | - | - | - | - | - | - | - | - | - | - | - | - | - | - | - | - | - | - | - | - | - | - | - | - | - | - | - | - | - | - | - | - | - | - | - | - | - | - | - | - | - | - | - | - | - | - | - | - | - | - | - | - | - | - | - | - | - | - | - | - | - | - | - | - | - | - | - | - | - | - | - | - | - | - | - | - | - | - | - | - | - | - | - | - | - | - | - | - | - | - | - | - | - | - | - | - | - | - | - | - | - | - | - | - | - | - | - | - | - | - | - | - | - | - | - | - | - | - | - | - | - | - | - | - | - | - | - | - | - | - | - | - | - | - | - | - | - | - | - | - | - | - | - | - | - | - | - | - | - | - | - | - | - | - | - | - | - | - | - | - | - | - | - | - | - | - | - | - | - | - | - | - | - | - | - | - | - | - | - | - | - | - | - | - | - | - | - | - | - | - | - | - | - | - | - | - | - | - | - | - | - | - | - | - | - | - | - | - | - | - | - | - | - | - | - | - | - | - | - | - | - | - | - | - | - | - | - | - | - | - | - | - | - | - | - | - | - | - | - | - | - | - | - | - | - | - | - | - | - | - | - | - | - | - | - | - | - | - | - | - | - | - | - | - | - | - | - | - | - | - | - | - | - | - | - | - | - | - | - | - | - | - | - | - | - | - | - | - | - | - | - | - | - | - | - | - | - | - | - | - | - | - | - | - | - | - | - | - | - | - | - | - | - | - | - | - | - | - | - | - | - | - | - | - | - | - | - | - | - | - | - | - | - | - | - | - | - | - | - | - | - | - | - | - | - | - | - | - | - | - | - | - | - | - | - | - | - | - | - | - | - | - | - | - | - | - | - | - | - | - | - | - | - | - | - | - | - | - | - | - | - | - | - | - | - | - | - | - | - | - | - | - | - | - | - | - | - | - | - | - | - | - | - | - | - | - | - | - | - | - | - | - | - | - | - | - | - | - | - | - | - | - | - | - | - | - | - | - | - | - | - | - | - | - | - | - | - | - | - | - | - | - | - | - | - | - | - | - | - | - | - | - | - | - | - | - | - | - | - | - | - | - | - | - | - | - | - | - | - | - | - | - | - | - | - | - | - | - | - | - | - | - | - | - | - | - | - | - | - | - | - | - | - | - | - | - | - | - | - | - | - | - | - | - | - | - | - | - | - | - | - | - | - | - | - | - | - | - | - | - | - | - | - | - | - | - | - | - | - | - | - | - | - | - | - | - | - | - | - | - | - | - | - | - | - | - | - | - | - | - | - | - | - | - | - | - | - | - | - | - | - | - | - | - | - | - | - | - | - | - | - | - | - | - | - | - | - | - | - | - | - | - | - | - | - | - | - | - | - | - | - | - | - | - | - | - | - | - | - | H | x | x | H | H | - | - | - | - | - | - | - | - | - | - | - | - | - | - | - | - | - | - | - | - | - | - | - | - | - | - | - | - | - | - | - | - | - | - | - | - | - | - | - | - | - | - | - | - | - | - | - | - | - | - | - | - | - | - | - | - | - | - | - | - | - | - | - | - | - | - | - | - | - | - | - | - | - | - | - | - | - | - | - | - | - | - | - | - | - | - | - | - | - | - | - | - | - | - | - | - | - | - | - | - | - | - | - | - | - | - | - | - | - | - | - | - | - | - | - | - | - | - | - | - | - | - | - | - | - | - | - | - | - | - | - | - | - | - | - | - | - | - | - | - | - | - | - | - | - | - | - | - | - | - | - | - | - | - | - | - | - | - | - | - | - | - | - | - | - | - | - | - | - | - | - | - | - | - | - | - | - | - | - | - | - | - | - | - | - | - | - | - | - | - | - | - | - | - | - | - | - | - | - | - | - | - | - | - | - | - | - | - | - | - | - | - | - | - | - | - | - | - | - | - | - | - | - | - | - | - | - | - | - | - | - | - | - | - | - | - | - | - | - | - | - | - | - | - | - | - | - | - | - | - | - | - | - | - | - | - | - | - | - | - | - | - | - | - | - | - | - | - | - | - | - | - | - | - | - | - | - | - | - | - | - | - | - | - | - | - | - | - | - | - | - | - | - | - | - | - | - | - | - | - | - | - | - | - | - | - | - | - | - | - | - | - | - | - | - | - | - | - | - | - | - | - | - | - | - | - | - | - | - | - | - | - | - | - | - | - | - | - | - | - | - | - | - | - | - | - | - | - | - | - | - | - | - | - | - | - | - | - | - | - | - | - | - | - | - | - | - | - | - | - | - | - | - | - | - | - | - | - | - | - | - | - | - | - | - | - | - | - | - | - | - | - | - | - | - | - | - | - | - | - | - | - | - | - | - | - | - | - | - | - | - | - | - | - | - | - | - | - | - | - | - | - | - | - | - | - | - | - | - | - | - | - | - | - | - | - | - | - | - | - | - | - | - | - | - | - | - | - | - | - | - | - | - | - | - | - | - | - | - | - | - | - | - | - | - | - | - | - | - | - | - | - | - | - | - | - | - | - | - | - | - | - | - | - | - | - | - | - | - | - | - | - | - | - | - | - | - | - | - |
| BIS d=0 cl.4 sym=1 env=1 | - | - | - | - | - | - | - | - | - | - | - | - | - | - | - | - | - | - | - | - | - | - | - | - | - | - | - | - | - | - | - | - | - | - | - | - | - | - | - | - | - | - | - | - | - | - | - | - | - | - | - | - | - | - | - | - | - | - | - | - | - | - | - | - | - | - | - | - | - | - | - | - | - | - | - | - | - | - | - | - | - | - | - | - | - | - | - | - | - | - | - | - | - | - | - | - | - | - | - | - | - | - | - | - | - | - | - | - | - | - | - | - | - | - | - | - | - | - | - | - | - | - | - | - | - | - | - | - | - | - | - | - | - | - | - | - | - | - | - | - | - | - | - | - | - | - | - | - | - | - | - | - | - | - | - | - | - | - | - | - | - | - | - | - | - | - | - | - | - | - | - | - | - | - | - | - | - | - | - | - | - | - | - | - | - | - | - | - | - | - | - | - | - | - | - | - | - | - | - | - | - | - | - | - | - | - | - | - | - | - | - | - | - | - | - | - | - | - | - | - | - | - | - | - | - | - | - | - | - | - | - | - | - | - | - | - | - | - | - | - | - | - | - | - | - | - | - | - | - | - | - | - | - | - | - | - | - | - | - | - | - | - | - | - | - | - | - | - | - | - | - | - | - | - | - | - | - | - | - | - | - | - | - | - | - | - | - | - | - | - | - | - | - | - | - | - | - | - | - | - | - | - | - | - | - | - | - | - | - | - | - | - | - | - | - | - | - | - | - | - | - | - | - | - | - | - | - | - | - | - | - | - | - | - | - | - | - | - | - | - | - | - | - | - | - | - | - | - | - | - | - | - | - | - | - | - | - | - | - | - | - | - | - | - | - | - | - | - | - | - | - | - | - | - | - | - | - | - | - | - | - | - | - | - | - | - | - | - | - | - | - | - | - | - | - | - | - | - | - | - | - | - | - | - | - | - | - | - | - | - | - | - | - | - | - | - | - | - | - | - | - | - | - | - | - | - | - | - | - | - | - | - | - | - | - | - | - | - | - | - | - | - | - | - | - | - | - | - | - | - | - | - | - | - | - | - | - | - | - | - | - | - | - | - | - | - | - | - | - | - | - | - | - | - | - | - | - | - | - | - | - | - | - | - | - | - | - | - | - | - | - | - | - | - | - | - | - | - | - | - | - | - | - | - | - | - | - | - | - | - | - | - | - | - | - | - | - | - | - | - | - | - | - | - | - | - | - | - | - | - | - | - | - | - | - | - | - | - | - | - | - | - | - | - | - | - | - | - | - | - | - | - | - | - | - | - | - | - | - | - | - | - | - | - | - | - | - | - | - | - | - | - | - | - | - | - | - | - | - | - | - | - | - | - | - | - | - | - | - | - | - | - | - | - | - | - | - | - | - | - | - | - | - | - | - | - | - | - | - | - | - | - | - | - | - | - | - | - | - | - | - | - | - | - | - | - | - | - | - | - | - | - | - | - | - | - | - | - | - | - | - | - | - | - | - | - | - | - | - | - | - | - | - | - | - | - | - | - | - | - | - | - | - | - | - | - | - | - | - | - | - | - | - | - | - | - | - | - | - | - | - | - | - | - | - | - | - | - | - | - | - | - | - | - | - | - | - | - | - | - | - | - | - | - | - | - | - | - | - | - | - | - | - | - | - | - | - | - | - | - | - | - | - | - | - | - | - | - | - | - | - | - | - | - | - | - | - | - | - | - | - | - | - | - | - | - | - | - | - | - | - | - | - | - | - | - | - | - | - | - | - | - | - | - | - | - | - | - | - | - | - | - | - | - | - | - | - | - | - | - | - | - | - | - | - | - | - | - | - | - | - | - | - | - | - | - | - | - | - | - | - | - | - | - | - | - | - | - | - | - | - | - | - | - | - | - | - | - | - | - | - | - | - | - | - | - | - | - | - | - | - | - | - | - | - | - | - | - | - | - | - | - | - | - | - | - | - | - | - | - | - | - | - | - | - | - | - | - | - | - | - | - | - | - | - | - | - | - | - | - | - | - | - | - | - | - | - | - | - | - | - | - | - | - | - | - | - | - | - | - | - | - | - | - | - | - | - | - | - | - | - | - | - | - | - | - | - | - | - | - | - | - | - | - | - | - | - | - | - | - | - | - | - | - | - | - | - | - | - | - | - | - | - | - | - | - | - | - | - | - | - | - | - | - | - | - | - | - | - | - | - | - | - | - | - | - | - | - | - | - | - | - | - | - | - | - | - | - | - | - | - | - | - | - | - | - | - | - | - | - | - | - | - | - | - | - | - | - | - | - | - | - | - | - | - | - | - | - | - | - | - | - | - | - | - | - | - | - | - | - | - | - | - | - | - | - | - | - | - | - | - | - | - | - | - | - | - | - | - | - | - | - | - | - | - | - | - | - | - | - | - | - | - | - | - | - | - | - | - | - | - | - | - | - | - | - | - | - | - | - | - | - | - | - | - | - | - | - | - | - | - | - | - | - | - | - | - | - | - | - | - | - | - | - | - | - | - | - | - | - | - | - | - | - | - | - | - | - | - | - | - | - | - | - | - | - | - | - | - | - | - | - | - | - | - | - | - | - | - | - | - | - | - | - | - | - | - | - | - | - | - | - | - | - | - | - | - | - | - | - | - | - | - | - | - | - | - | - | - | - | - | - | - | - | - | - | - | - | - | - | - | - | - | - | - | - | - | - | - | - | - | - | - | - | - | - | - | - | - | - | - | - | - | - | - | - | - | - | - | - | - | - | - | - | - | - | - | - | - | - | - | - | - | - | - | - | - | - | - | - | - | - | - | - | - | - | - | - | - | - | - | - | - | - | - | - | - | - | - | - | - | - | - | - | - | - | - | - | - | - | - | - | - | - | - | - | - | - | - | - | - | - | - | - | - | - | - | - | - | - | - | - | - | - | - | - | - | - | - | - | - | - | - | - | - | - | - | - | - | - | - | - | - | - | - | - | - | - | - | - | - | - | - | - | - | - | - | - | - | - | - | - | - | - | - | - | - | - | - | - | - | - | - | - | - | - | - | - | - | - | - | - | - | - | - | - | - | - | - | - | - | - | - | - | - | - | - | - | - | - | - | - | - | - | - | - | - | - | - | - | - | - | - | - | - | - | - | - | - | - | - | - | - | - | - | - | - | - | - | - | - | - | - | - | - | - | - | - | - | - | - | - | - | - | - | - | - | - | - | - | - | - | - | - | - | - | - | - | - | - | - | - | - | - | - | - | - | - | - | - | - | - | - | - | - | - | - | - | - | - | - | - | - | - | - | - | - | - | - | - | - | - | - | - | - | - | - | - | - | - | - | - | - | - | - | - | - | - | - | - | - | - | - | - | - | - | - | - | - | - | - | - | - | - | - | - | - | - | - | - | - | - | - | - | - | - | - | - | - | - | - | - | - | - | - | - | - | - | - | - | - | - | - | - | - | - | - | - | - | - | - | - | - | - | - | - | - | - | - | - | - | - | - | - | - | - | - | - | - | - | - | - | - | - | - | - | - | - | - | - | - | - | - | - | - | - | - | - | - | - | - | - | - | - | - | - | - | - | - | - | - | - | - | - | - | - | - | - | - | - | - | - | - | - | - | - | - | - | - | - | - | - | - | - | - | - | - | - | - | - | - | - | - | - | - | - | - | - | - | - | - | - | - | - | - | - | - | - | - | - | - | - | - | - | - | - | - | - | - | - | - | - | - | - | - | - | - | - | - | - | - | - | - | - | - | - | - | - | - | - | - | - | - | - | - | - | - | - | - | - | - | - | - | - | - | - | - | - | - | - | - | - | - | - | - | - | - | - | - | - | - | - | - | - | - | - | - | - | - | - | - | - | - | - | - | - | - | - | - | - | - | - | - | - | - | - | - | - | - | - | - | - | - | - | - | - | - | - | - | - | - | - | - | - | - | - | - | - | - | - | - | - | - | - | - | - | - | - | - | - | - | - | - | - | - | - | - | - | - | - | - | - | - | - | - | - | - | - | - | - | - | - | - | - | - | - | - | - | - | - | - | - | - | - | - | - | - | - | - | - | - | - | - | - | - | - | - | - | - | - | - | - | - | - | - | - | - | - | - | - | - | - | - | - | - | - | - | - | - | - | - | - | - | - | - | - | - | - | - | - | - | - | - | - | - | - | - | - | - | - | - | - | - | - | - | - | - | - | - | - | - | - | - | - | - | - | - | - | - | - | - | - | - | - | - | - | - | - | - | - | - | - | - | - | - | - | - | - | - | - | - | - | - | - | - | - | - | - | - | - | - | - | - | - | - | - | - | - | - | - | - | - | - | - | - | - | - | - | - | - | - | - | - | - | - | - | - | - | - | - | - | - | - | - | - | - | - | - | - | - | - | - | - | - | - | - | - | - | - | - | - | - | - | - | - | - | - | - | - | - | - | - | - | - | - | - | - | - | - | - | - | - | - | - | - | - | - | - | - | - | - | - | - | - | - | - | - | - | - | - | - | - | - | - | - | - | - | - | - | - | - | - | - | - | - | - | - | - | - | - | - | - | - | - | - | - | - | - | - | - | - | - | - | - | - | - | - | - | - | - | - | - | - | - | - | - | - | - | - | - | - | - | - | - | - | - | - | - | - | - | - | - | - | - | - | - | - | - | - | - | - | - | - | - | - | - | - | - | - | - | - | - | - | - | - | - | - | - | - | - | - | - | - | - | - | - | - | - | - | - | - | - | - | - | - | - | - | - | - | - | - | - | - | - | - | - | - | - | - | - | - | - | - | - | - | - | - | - | - | - | - | - | - | - | - | - | - | - | - | - | - | - | - | - | - | - | - | - | - | - | - | - | - | - | - | - | - | - | - | - | - | - | - | - | - | - | - | - | - | - | - | - | - | - | - | - | - | - | - | - | - | - | - | - | - | - | - | - | - | - | - | - | - | - | - | - | - | - | - | - | - | - | - | - | - | - | - | - | - | - | - | - | - | - | - | - | - | - | - | - | - | - | - | - | - | - | - | - | - | - | - | - | - | - | - | - | - | - | - | - | - | - | - | - | - | - | - | - | - | - | - | - | - | - | - | - | - | - | - | - | - | - | - | - | - | - | - | - | - | - | - | - | - | - | - | - | - | - | - | - | - | - | - | - | - | - | - | - | - | - | - | - | - | - | - | - | - | - | - | - | - | - | - | - | - | - | - | - | - | - | - | - | - | - | - | - | - | - | - | - | - | - | - | - | - | - | - | - | - | - | - | - | - | - | - | - | - | - | - | - | - | - | - | - | - | - | - | - | - | - | - | - | - | - | - | - | - | - | - | - | - | - | - | - | - | - | - | - | - | - | - | - | - | - | - | - | - | - | - | - | - | - | - | - | - | - | - | - | - | - | - | - | - | - | - | - | - | - | - | - | - | - | - | - | - | - | - | - | - | - | - | - | - | - | - | - | - | - | - | - | - | - | - | - | - | - | - | - | - | - | - | - | - | - | - | - | - | - | - | - | - | - | - | - | - | - | - | - | - | - | - | - | - | - | - | - | - | - | - | - | - | - | - | - | - | - | - | - | - | - | - | - | - | - | - | - | - | - | - | - | - | - | - | - | - | - | - | - | - | - | - | - | - | - | - | - | - | - | - | - | - | - | - | - | - | - | - | - | - | - | - | - | - | - | - | - | - | - | - | - | - | - | - | - | - | - | - | - | - | - | - | - | - | - | - | - | - | - | - | - | - | - | - | - | - | - | - | - | - | - | - | - | - | - | - | - | - | - | - | - | - | - | - | - | - | - | - | - | - | - | - | - | - | - | - | - | - | - | - | - | - | - | - | - | - | - | - | - | - | - | - | - | - | - | - | - | - | - | - | - | - | - | - | - | - | - | - | - | - | - | - | - | - | - | - | - | - | - | - | - | - | - | - | - | - | - | - | - | - | - | - | - | - | - | - | - | - | - | - | - | - | - | - | - | - | - | - | - | - | - | - | - | - | - | - | - | - | - | - | - | - | - | - | - | - | - | - | - | - | - | - | - | - | - | - | - | - | - | - | - | - | - | - | - | - | - | - | - | - | - | - | - | - | - | - | - | - | - | - | - | - | - | - | - | - | - | - | - | - | - | - | - | - | - | - | - | - | - | - | - | - | - | - | - | - | - | - | - | - | - | - | - | - | - | - | - | - | - | - | - | - | - | - | - | - | - | - | - | - | - | - | - | - | - | - | - | - | - | - | - | - | - | - | - | - | - | - | - | - | - | - | - | - | - | - | - | - | - | - | - | - | - | - | - | - | - | - | - | - | - | - | - | - | - | - | - | - | - | - | - | - | - | - | - | - | - | - | - | - | - | - | - | - | - | - | - | - | - | - | - | - | - | - | - | - | - | - | - | - | - | - | - | - | - | - | - | - | - | - | - | - | - | - | - | - | - | - | - | - | - | - | - | - | - | - | - | - | - | - | - | - | - | - | - | - | - | - | - | - | - | - | - | - | - | - | - | - | - | - | - | - | - | - | - | - | - | - | - | - | - | - | - | - | - | - | - | - | - | - | - | - | - | - | - | - | - | - | - | - | - | - | - | - | - | - | - | - | - | - | - | - | - | - | - | - | - | - | - | - | - | - | - | - | - | - | - | - | - | - | - | - | - | - | - | - | - | - | - | - | - | - | - | - | - | - | - | - | - | - | - | - | - | - | - | - | - | - | - | - | - | - | - | - | - | - | - | - | - | - | - | - | - | - | - | - | - | - | - | - | - | - | - | - | - | - | - | - | - | - | - | - | - | - | - | - | - | - | - | - | - | - | - | - | - | - | - | - | - | - | - | - | - | - | - | - | - | - | - | - | - | - | - | - | - | - | - | - | - | - | - | - | - | - | - | - | - | - | - | - | - | - | - | - | - | - | - | - | - | - | - | - | - | - | - | - | - | - | - | - | - | - | - | - | - | - | - | - | - | - | - | - | - | - | - | - | - | - | - | - | - | - | - | - | - | - | - | - | - | - | - | - | - | - | - | - | - | - | - | - | - | - | - | - | - | - | - | - | - | - | - | - | - | - | - | - | - | - | - | - | - | - | - | - | - | - | - | - | - | - | - | - | - | - | - | - | - | - | - | - | - | - | - | - | - | - | - | - | - | - | - | - | - | - | - | - | - | - | - | - | - | - | - | - | - | - | - | - | - | - | - | - | - | - | - | - | - | - | - | - | - | - | - | - | - | - | - | - | - | - | - | - | - | - | - | - | - | - | - | - | - | - | - | - | - | - | - | - | - | - | - | - | - | - | - | - | - | - | - | - | - | - | - | - | - | - | - | - | - | - | - | - | - | - | - | - | - | - | - | - | - | - | - | - | - | - | - | - | - | - | - | - | - | - | - | - | - | - | - | - | - | - | - | - | - | - | - | - | - | - | - | - | - | - | - | - | - | - | - | - | - | - | - | - | - | - | - | - | - | - | - | - | - | - | - | - | - | - | - | - | - | - | - | - | - | - | - | - | - | - | - | - | - | - | - | - | - | - | - | - | - | - | - | - | - | - | - | - | - | - | - | - | - | - | - | - | - | - | - | - | - | - | - | - | - | - | - | - | - | - | - | - | - | - | - | - | - | - | - | - | - | - | - | - | - | - | - | - | - | - | - | - | - | - | - | - | - | - | - | - | - | - | - | - | - | - | - | - | - | - | - | - | - | - | - | - | - | - | - | - | - | - | - | - | - | - | - | - | - | - | - | - | - | - | - | - | - | - | - | - | - | - | - | - | - | - | - | - | - | - | - | - | - | - | - | - | - | - | - | - | - | - | - | - | - | - | - | - | - | - | - | - | - | - | - | - | - | - | - | - | - | - | - | - | - | - | - | - | - | - | - | - | - | - | - | - | - | - | - | - | - | - | - | - | - | - | - | - | - | - | - | - | - | - | - | - | - | - | - | - | - | - | - | - | - | - | - | - | - | - | - | - | - | - | - | - | - | - | - | - | - | - | - | - | - | - | - | - | - | - | - | - | - | - | - | - | - | - | - | - | - | - | - | - | - | - | - | - | - | - | - | - | - | - | - | - | - | - | - | - | - | - | - | - | - | - | - | - | - | - | - | - | - | - | - | - | - | - | - | - | - | - | - | - | - | - | - | - | - | - | - | - | - | - | - | - | - | - | - | - | - | - | - | - | - | - | - | - | - | - | - | - | - | - | - | - | - | - | - | - | - | - | - | - | - | - | - | - | - | - | - | - | - | - | - | - | - | - | - | - | - | - | - | - | - | - | - | - | - | - | - | - | - | - | - | - | - | - | - | - | - | - | - | - | - | - | - | - | - | - | - | - | - | - | - | - | - | - | - | - | - | - | - | - | - | - | - | - | - | - | - | - | - | - | - | - | - | - | - | - | - | - | - | - | - | - | - | - | - | - | - | - | - | - | - | - | - | - | - | - | - | - | - | - | - | - | - | - | - | - | - | - | - | - | - | - | - | - | - | - | - | - | - | - | - | - | - | - | - | - | - | - | - | - | - | - | - | - | - | - | - | - | - | - | - | - | - | - | - | - | - | - | - | - | - | - | - | - | - | - | - | - | - | - | - | - | - | - | - | - | - | - | - | - | - | - | - | - | - | - | - | - | - | - | - | - | - | - | - | - | - | - | - | - | - | - | - | - | - | - | - | - | - | - | - | - | - | - | - | - | - | - | - | - | - | - | - | - | - | - | - | - | - | - | - | - | - | - | - | - | - | - | - | - | - | - | - | - | - | - | - | - | - | - | - | - | - | - | - | - | - | - | - | - | - | - | - | - | - | - | - | - | - | - | - | - | - | - | - | - | - | - | - | - | - | - | - | - | - | - | - | - | - | - | - | - | - | - | - | - | - | - | - | - | - | - | - | - | - | - | - | - | - | - | - | - | - | - | - | - | - | - | - | - | - | - | - | - | - | - | - | - | - | - | - | - | - | - | - | - | - | - | - | - | - | - | - | - | - | - | - | - | - | - | - | - | - | - | - | - | - | - | - | - | - | - | - | - | - | - | - | - | - | - | - | - | - | - | - | - | - | - | - | - | - | - | - | - | - | - | - | - | - | - | - | - | - | - | - | - | - | - | - | - | - | - | - | - | - | - | - | - | - | - | - | - | - | - | - | - | - | - | - | - | - | - | - | - | - | - | - | - | - | - | - | - | - | - | - | - | - | - | - | - | - | - | - | - | - | - | - | - | - | - | - | - | - | - | - | - | - | - | - | - | - | - | - | - | - | - | - | - | - | - | - | - | - | - | - | - | - | - | - | - | - | - | - | - | - | - | - | - | - | - | - | - | - | - | - | - | - | - | - | - | - | - | - | - | - | - | - | - | - | - | - | - | - | - | - | - | - | - | - | - | - | - | - | - | - | - | - | - | - | - | - | - | - | - | - | - | - | - | - | - | - | - | - | - | - | - | - | - | - | - | - | - | - | - | - | - | - | - | - | - | - | - | - | - | - | - | - | - | - | - | - | - | - | - | - | - | - | - | - | - | - | - | - | - | - | - | - | - | - | - | - | - | - | - | - | - | - | - | - | - | - | - | - | - | - | - | - | - | - | - | - | - | - | - | - | - | - | - | - | - | - | - | - | - | - | - | - | - | - | - | - | - | - | - | - | - | - | - | - | - | - | - | - | - | - | - | - | - | - | - | - | - | - | - | - | - | - | - | - | - | - | - | - | - | - | - | - | - | - | - | - | - | - | - | - | - | - | - | - | - | - | - | - | - | - | - | - | - | - | - | - | - | - | - | - | - | - | - | - | - | - | - | - | - | - | - | - | - | - | - | - | - | - | - | - | - | - | - | - | - | - | - | - | - | - | - | - | - | - | - | - | - | - | - | - | - | - | - | - | - | - | - | - | - | - | - | - | - | - | - | - | - | - | - | - | - | - | - | - | - | - | - | - | - | - | - | - | - | - | - | - | - | - | - | - | - | - | - | - | - | - | - | - | - | - | - | - | - | - | - | - | - | - | - | - | - | - | - | - | - | - | - | - | - | - | - | - | - | - | - | - | - | - | - | - | - | - | - | - | - | - | - | - | - | - | - | - | - | - | - | - | - | - | - | - | - | - | - | - | - | - | - | - | - | - | - | - | - | - | - | - | - | - | - | - | - | - | - | - | - | - | - | - | - | - | - | - | - | - | - | - | - | - | - | - | - | - | - | - | - | - | - | - | - | - | - | - | - | - | - | - | - | - | - | - | - | - | - | - | - | - | - | - | - | - | - | - | - | - | - | - | - | - | - | - | - | - | - | - | - | - | - | - | - | - | - | - | - | - | - | - | - | - | - | - | - | - | - | - | - | - | - | - | - | - | - | - | - | - | - | - | - | - | - | - | - | - | - | - | - | - | - | - | - | - | - | - | - | - | - | - | - | - | - | - | - | - | - | - | - | - | - | - | - | - | - | - | - | - | - | - | - | - | - | - | - | - | - | - | - | - | - | - | - | - | - | - | - | - | - | - | - | - | - | - | - | - | - | - | - | - | - | - | - | - | - | - | - | - | - | - | - | - | - | - | - | - | - | - | - | - | - | - | - | - | - | - | - | - | - | - | - | - | - | - | - | - | - | - | - | - | - | - | - | - | - | - | - | - | - | - | - | - | - | - | - | - | - | - | - | - | - | - | - | - | - | - | - | - | - | - | - | - | - | - | - | - | - | - | - | - | - | - | - | - | - | - | - | - | - | - | - | - | - | - | - | - | - | - | - | - | - | - | - | - | - | - | - | - | - | - | - | - | - | - | - | - | - | - | - | - | - | - | - | - | - | - | - | - | - | - | - | - | - | - | - | - | - | - | - | - | - | - | - | - | - | - | - | - | - | - | - | - | - | - | - | - | - | - | - | - | - | - | - | - | - | - | - | - | - | - | - | - | - | - | - | - | - | - | - | - | - | - | - | - | - | - | - | - | - | - | - | - | - | - | - | - | - | - | - | - | - | - | - | - | - | - | - | - | - | - | - | - | - | - | - | - | - | - | - | - | - | - | - | - | - | - | - | - | - | - | - | - | - | - | - | - | - | - | - | - | - | - | - | - | - | - | - | - | - | - | - | - | - | - | - | - | - | - | - | - | - | - | - | - | - | - | - | - | - | - | - | - | - | - | - | - | - | - | - | - | - | - | - | - | - | - | - | - | - | - | - | - | - | - | - | - | - | - | - | - | - | - | - | - | - | - | - | - | - | - | - | - | - | - | - | - | - | - | - | - | - | - | - | - | - | - | - | - | - | - | - | - | - | - | - | - | - | - | - | - | - | - | - | - | - | - | - | - | - | - | - | - | - | - | - | - | - | H | x | x | x | x | x | x | x | x | x | x | x | x | x | x | x | x | x | x | x | x | H | E | - | - | - | - | - | - | - | - | - | - | - | - | - | - | - | - | - | - | - | - | - | - | - | - | - | - | - | - | - | - | - | - | - | - | - | - | - | - | - | - | - | - | - | - | - | - | - | - | - | - | - | - | - | - | - | - | - | - | - | - | - | - | - | - | - | - | - | - | - | - | - | - | - | - | - | - | - | - | - | - | - | - | - | - | - | - | - | - | - | - | - | - | - | - | - | - | - | - | - | - | - | - | - | - | - | - | - | - | - | - | - | - | - | - | - | - | - | - | - | - | - | - | - | - | - | - | - | - | - | - | - | - | - | - | - | - | - | - | - | - | - | - | - | - | - | - | - | - | - | - | - | - | - | - | - | - | - | - | - | - | - | - | - | - | - | - | - | - | - | - | - | - | - | - | - | - | - | - | - | - | - | - | - | - | - | - | - | - | - | - | - | - | - | - | - | - | - | - | - | - | - | - | - | - | - | - | - | - | - | - | - | - | - | - | - | - | - | - | - | - | - | - | - | - | - | - | - | - | - | - | - | - | - | - | - | - | - | - | - | - | - | - | - | - | - | - | - | - | - | - | - | - | - | - | - | - | - | - | - | - | - | - | - | - | - | - | - | - | - | - | - | - | - | - | - | - | - | - | - | - | - | - | - | - | - | - | - | - | - | - | - | - | - | - | - | - | - | - | - | - | - | - | - | - | - | - | - | - | - | - | - | - | - | - | - | - | - | - | - | - | - | - | - | - | - | - | - | - | - | - | - | - | - | - | - | - | - | - | - | - | - | - | - | - | - | - | - | - | - | - | - | - | - | - | - | - | - | - | - | - | - | - | - | - | - | - | - | - | - | - | - | - | - | - | - | - | - | - | - | - | - | - | - | - | - | - | - | - | - | - | - | - | - | - | - | - | - | - | - | - | - | - | - | - | - | - | - | - | - | - | - | - | - | - | - | - | - | - | - | - | - | - | - | - | - | - | - | - | - | - | - | - | - | - | - | - | - | - | - | - | - | - | - | - | - | - | - | - | - | - | - | - | - | - |
| BIS d=0 cl.23 sym=0.73 env=0.5 | - | - | - | - | - | - | - | - | - | - | - | - | - | - | - | - | - | - | - | - | - | - | - | - | - | - | - | - | - | - | - | - | - | - | - | - | - | - | - | - | - | - | - | - | - | - | - | - | - | - | - | - | - | - | - | - | - | - | - | - | - | - | - | - | - | - | - | - | - | - | - | - | - | - | - | - | - | - | - | - | - | - | - | - | - | - | - | - | - | - | - | - | - | - | - | - | - | - | - | - | - | - | - | - | - | - | - | - | - | - | - | - | - | - | - | - | - | - | - | - | - | - | - | - | - | - | - | - | - | - | - | - | - | - | - | - | - | - | - | - | - | - | - | - | - | - | - | - | - | - | - | - | - | - | - | - | - | - | - | - | - | - | - | - | - | - | - | - | - | - | - | - | - | - | - | - | - | - | - | - | - | - | - | - | - | - | - | - | - | - | - | - | - | - | - | - | - | - | - | - | - | - | - | - | - | - | - | - | - | - | - | - | - | - | - | - | - | - | - | - | - | - | - | - | - | - | - | - | - | - | - | - | - | - | - | - | - | - | - | - | - | - | - | - | - | - | - | - | - | - | - | - | - | - | - | - | - | - | - | - | - | - | - | - | - | - | - | - | - | - | - | - | - | - | - | - | - | - | - | - | - | - | - | - | - | - | - | - | - | - | - | - | - | - | - | - | - | - | - | - | - | - | - | - | - | - | - | - | - | - | - | - | - | - | - | - | - | - | - | - | - | - | - | - | - | - | - | - | - | - | - | - | - | - | - | - | - | - | - | - | - | - | - | - | - | - | - | - | - | - | - | - | - | - | - | - | - | - | - | - | - | - | - | - | - | - | - | - | - | - | - | - | - | - | - | - | - | - | - | - | - | - | - | - | - | - | - | - | - | - | - | - | - | - | - | - | - | - | - | - | - | - | - | - | - | - | - | - | - | - | - | - | - | - | - | - | - | - | - | - | - | - | - | - | - | - | - | - | - | - | - | - | - | - | - | - | - | - | - | - | - | - | - | - | - | - | - | - | - | - | - | - | - | - | - | - | - | - | - | - | - | - | - | - | - | - | - | - | - | - | - | - | - | - | - | - | - | - | - | - | - | - | - | - | - | - | - | - | - | - | - | - | - | - | - | - | - | - | - | - | - | - | - | - | - | - | - | - | - | - | - | - | - | - | - | - | - | - | - | - | - | - | - | - | - | - | - | - | - | - | - | - | - | - | - | - | - | - | - | - | - | - | - | - | - | - | - | - | - | - | - | - | - | - | - | - | - | - | - | - | - | - | - | - | - | - | - | - | - | - | - | - | - | - | - | - | - | - | - | - | - | - | - | - | - | - | - | - | - | - | - | - | - | - | - | - | - | - | - | - | - | - | - | - | - | - | - | - | - | - | - | - | - | - | - | - | - | - | - | - | - | - | - | - | - | - | - | - | - | - | - | - | - | - | - | - | - | - | - | - | - | - | - | - | - | - | - | - | - | - | - | - | - | - | - | - | - | - | - | - | - | - | - | - | - | - | - | - | - | - | - | - | - | - | - | - | - | - | - | - | - | - | - | - | - | - | - | - | - | - | - | - | - | - | - | - | - | - | - | - | - | - | - | - | - | - | - | - | - | - | - | - | - | - | - | - | - | - | - | - | - | - | - | - | - | - | - | - | - | - | - | - | - | - | - | - | - | - | - | - | - | - | - | - | - | - | - | - | - | - | - | - | - | - | - | - | - | - | - | - | - | - | - | - | - | - | - | - | - | - | - | - | - | - | - | - | - | - | - | - | - | - | - | - | - | - | - | - | - | - | - | - | - | - | - | - | - | - | - | - | - | - | - | - | - | - | - | - | - | - | - | - | - | - | - | - | - | - | - | - | - | - | - | - | - | - | - | - | - | - | - | - | - | - | - | - | - | - | - | - | - | - | - | - | - | - | - | - | - | - | - | - | - | - | - | - | - | - | - | - | - | - | - | - | - | - | - | - | - | - | - | - | - | - | - | - | - | - | - | - | - | - | - | - | - | - | - | - | - | - | - | - | - | - | - | - | - | - | - | - | - | - | - | - | - | - | - | - | - | - | - | - | - | - | - | - | - | - | - | - | - | - | - | - | - | - | - | - | - | - | - | - | - | - | - | - | - | - | - | - | - | - | - | - | - | - | - | - | - | - | - | - | - | - | - | - | - | - | - | - | - | - | - | - | - | - | - | - | - | - | - | - | - | - | - | - | - | - | - | - | - | - | - | - | - | - | - | - | - | - | - | - | - | - | - | - | - | - | - | - | - | - | - | - | - | - | - | - | - | - | - | - | - | - | - | - | - | - | - | - | - | - | - | - | - | - | - | - | - | - | - | - | - | - | - | - | - | - | - | - | - | - | - | - | - | - | - | - | - | - | - | - | - | - | - | - | - | - | - | - | - | - | - | - | - | - | - | - | - | - | - | - | - | - | - | - | - | - | - | - | - | - | - | - | - | - | - | - | - | - | - | - | - | - | - | - | - | - | - | - | - | - | - | - | - | - | - | - | - | - | - | - | - | - | - | - | - | - | - | - | - | - | - | - | - | - | - | - | - | - | - | - | - | - | - | - | - | - | - | - | - | - | - | - | - | - | - | - | - | - | - | - | - | - | - | - | - | - | - | - | - | - | - | - | - | - | - | - | - | - | - | - | - | - | - | - | - | - | - | - | - | - | - | - | - | - | - | - | - | - | - | - | - | - | - | - | - | - | - | - | - | - | - | - | - | - | - | - | - | - | - | - | - | - | - | - | - | - | - | - | - | - | - | - | - | - | - | - | - | - | - | - | - | - | - | - | - | - | - | - | - | - | - | - | - | - | - | - | - | - | - | - | - | - | - | - | - | - | - | - | - | - | - | - | - | - | - | - | - | - | - | - | - | - | - | - | - | - | - | - | - | - | - | - | - | - | - | - | - | - | - | - | - | - | - | - | - | - | - | - | - | - | - | - | - | - | - | - | - | - | - | - | - | - | - | - | - | - | - | - | - | - | - | - | - | - | - | - | - | - | - | - | - | - | - | - | - | - | - | - | - | - | - | - | - | - | - | - | - | - | - | - | - | - | - | - | - | - | - | - | - | - | - | - | - | - | - | - | - | - | - | - | - | - | - | - | - | - | - | - | - | - | - | - | - | - | - | - | - | - | - | - | - | - | - | - | - | - | - | - | - | - | - | - | - | - | - | - | - | - | - | - | - | - | - | - | - | - | - | - | - | - | - | - | - | - | - | - | - | - | - | - | - | - | - | - | - | - | - | - | - | - | - | - | - | - | - | - | - | - | - | - | - | - | - | - | - | - | - | - | - | - | - | - | - | - | - | - | - | - | - | - | - | - | - | - | - | - | - | - | - | - | - | - | - | - | - | - | - | - | - | - | - | - | - | - | - | - | - | - | - | - | - | - | - | - | - | - | - | - | - | - | - | - | - | - | - | - | - | - | - | - | - | - | - | - | - | - | - | - | - | - | - | - | - | - | - | - | - | - | - | - | - | - | - | - | - | - | - | - | - | - | - | - | - | - | - | - | - | - | - | - | - | - | - | - | - | - | - | - | - | - | - | - | - | - | - | - | - | - | - | - | - | - | - | - | - | - | - | - | - | - | - | - | - | - | - | - | - | - | - | - | - | - | - | - | - | - | - | - | - | - | - | - | - | - | - | - | - | - | - | - | - | - | - | - | - | - | - | - | - | - | - | - | - | - | - | - | - | - | - | - | - | - | - | - | - | - | - | - | - | - | - | - | - | - | - | - | - | - | - | - | - | - | - | - | - | - | - | - | - | - | - | - | - | - | - | - | - | - | - | - | - | - | - | - | - | - | - | - | - | - | - | - | - | - | - | - | - | - | - | - | - | - | - | - | - | - | - | - | - | - | - | - | - | - | - | - | - | - | - | - | - | - | - | - | - | - | - | - | - | - | - | - | - | - | - | - | - | - | - | - | - | - | - | - | - | - | - | - | - | - | - | - | - | - | - | - | - | - | - | - | - | - | - | - | - | - | - | - | - | - | - | - | - | - | - | - | - | - | - | - | - | - | - | - | H | E | x | x | H | - | - | - | - | - | - | - | - | - | - | - | - | - | - | - | - | - | - | - | - | - | - | - | - | - | - | - | - | - | - | - | - | - | - | - | - | - | - | - | - | - | - | - | - | - | - | - | - | - | - | - | - | - | - | - | - | - | - | - | - | - | - | - | - | - | - | - | - | - | - | - | - | - | - | - | - | - | - | - | - | - | - | - | - | - | - | - | - | - | - | - | - | - | - | - | - | - | - | - | - | - | - | - | - | - | - | - | - | - | - | - | - | - | - | - | - | - | - | - | - | - | - | - | - | - | - | - | - | - | - | - | - | - | - | - | - | - | - | - | - | - | - | - | - | - | - | - | - | - | - | - | - | - | - | - | - | - | - | - | - | - | - | - | - | - | - | - | - | - | - | - | - | - | - | - | - | - | - | - | - | - | - | - | - | - | - | - | - | - | - | - | - | - | - | - | - | - | - | - | - | - | - | - | - | - | - | - | - | - | - | - | - | - | - | - | - | - | - | - | - | - | - | - | - | - | - | - | - | - | - | - | - | - | - | - | - | - | - | - | - | - | - | - | - | - | - | - | - | - | - | - | - | - | - | - | - | - | - | - | - | - | - | - | - | - | - | - | - | - | - | - | - | - | - | - | - | - | - | - | - | - | - | - | - | - | - | - | - | - | - | - | - | - | - | - | - | - | - | - | - | - | - | - | - | - | - | - | - | - | - | - | - | - | - | - | - | - | - | - | - | - | - | - | - | - | - | - | - | - | - | - | - | - | - | - | - | - | - | - | - | - | - | - | - | - | - | - | - | - | - | - | - | - | - | - | - | - | - | - | - | - | - | - | - | - | - | - | - | - | - | - | - | - | - | - | - | - | - | - | - | - | - | - | - | - | - | - | - | - | - | - | - | - | - | - | - | - | - | - | - | - | - | - | - | - | - | - | - | - | - | - | - | - | - | - | - | - | - | - | - | - | - | - | - | - | - | - | - | - | - | - | - | - | - | - | - | - | - | - | - | - | - | - | - | - | - | - | - | - | - | - | - | - | - | - | - | - | - | - | - | - | - | - | - | - | - | - | - | - | - | - | - | - | - | - | - | - | - | - | - | - | - | - | - | - | - | - | - | - | - | - | - | - | - | - | - | - | - | - | - | - | - | - | - | - | - | - | - | - | - | - | - | - | - | - | - | - | - | - | - | - | - | - | - | - | - | - | - | - | - | - | - | - | - | - | - | - | - | - | - | - | - | - | - | - | - | - | - | - | - | - | - | - | - | - | - | - | - | - | - | - | - | - | - | - | - | - | - | - | - | - | - | - | - | - | - | - | - | - | - | - | - | - | - | - | - | - | - | - | - | - | - | - | - | - | - | - | - | - | - | - | - | - | - | - | - | - | - | - | - | - | - | - | - | - | - | - | - | - | - | - | - | - | - | - | - | - | - | - | - | - | - | - | - | - | - | - | - | - | - | - | - | - | - | - | - | - | - | - | - | - | - | - | - | - | - | - | - | - | - | - | - | - | - | - | - | - | - | - | - | - | - | - | - | - | - | - | - | - | - | - | - | - | - | - | - | - | - | - | - | - | - | - | - | - | - | - | - | - | - | - | - | - | - | - | - | - | - | - | - | - | - | - | - | - | - | - | - | - | - | - | - | - | - | - | - | - | - | - | - | - | - | - | - | - | - | - | - | - | - | - | - | - | - | - | - | - | - | - | - | - | - | - | - | - | - | - | - | - | - | - | - | - | - | - | - | - | - | - | - | - | - | - | - | - | - | - | - | - | - | - | - | - | - | - | - | - | - | - | - | - | - | - | - | - | - | - | - | - | - | - | - | - | - | - | - | - | - | - | - | - | - | - | - | - | - | - | - | - | - | - | - | - | - | - | - | - | - | - | - | - | - | - | - | - | - | - | - | - | - | - | - | - | - | - | - | - | - | - | - | - | - | - | - | - | - | - | - | - | - | - | - | - | - | - | - | - | - | - | - | - | - | - | - | - | - | - | - | - | - | - | - | - | - | - | - | - | - | - | - | - | - | - | - | - | - | - | - | - | - | - | - | - | - | - | - | - | - | - | - | - | - | - | - | - | - | - | - | - | - | - | - | - | - | - | - | - | - | - | - | - | - | - | - | - | - | - | - | - | - | - | - | - | - | - | - | - | - | - | - | - | - | - | - | - | - | - | - | - | - | - | - | - | - | - | - | - | - | - | - | - | - | - | - | - | - | - | - | - | - | - | - | - | - | - | - | - | - | - | - | - | - | - | - | - | - | - | - | - | - | - | - | - | - | - | - | - | - | - | - | - | - | - | - | - | - | - | - | - | - | - | - | - | - | - | - | - | - | - | - | - | - | - | - | - | - | - | - | - | - | - | - | - | - | - | - | - | - | - | - | - | - | - | - | - | - | - | - | - | - | - | - | - | - | - | - | - | - | - | - | - | - | - | - | - | - | - | - | - | - | - | - | - | - | - | - | - | - | - | - | - | - | - | - | - | - | - | - | - | - | - | - | - | - | - | - | - | - | - | - | - | - | - | - | - | - | - | - | - | - | - | - | - | - | - | - | - | - | - | - | - | - | - | - | - | - | - | - | - | - | - | - | - | - | - | - | - | - | - | - | - | - | - | - | - | - | - | - | - | - | - | - | - | - | - | - | - | - | - | - | - | - | - | - | - | - | - | - | - | - | - | - | - | - | - | - | - | - | - | - | - | - | - | - | - | - | - | - | - | - | - | - | - | - | - | - | - | - | - | - | - | - | - | - | - | - | - | - | - | - | - | - | - | - | - | - | - | - | - | - | - | - | - | - | - | - | - | - | - | - | - | - | - | - | - | - | - | - | - | - | - | - | - | - | - | - | - | - | - | - | - | - | - | - | - | - | - | - | - | - | - | - | - | - | - | - | - | - | - | - | - | - | - | - | - | - | - | - | - | - | - | - | - | - | - | - | - | - | - | - | - | - | - | - | - | - | - | - | - | - | - | - | - | - | - | - | - | - | - | - | - | - | - | - | - | - | - | - | - | - | - | - | - | - | - | - | - | - | - | - | - | - | - | - | - | - | - | - | - | - | - | - | - | - | - | - | - | - | - | - | - | - | - | - | - | - | - | - | - | - | - | - | - | - | - | - | - | - | - | - | - | - | - | - | - | - | - | - | - | - | - | - | - | - | - | - | - | - | - | - | - | - | - | - | - | - | - | - | - | - | - | - | - | - | - | - | - | - | - | - | - | - | - | - | - | - | - | - | - | - | - | - | - | - | - | - | - | - | - | - | - | - | - | - | - | - | - | - | - | - | - | - | - | - | - | - | - | - | - | - | - | - | - | - | - | - | - | - | - | - | - | - | - | - | - | - | - | - | - | - | - | - | - | - | - | - | - | - | - | - | - | - | - | - | - | - | - | - | - | - | - | - | - | - | - | - | - | - | - | - | - | - | - | - | - | - | - | - | - | - | - | - | - | - | - | - | - | - | - | - | - | - | - | - | - | - | - | - | - | - | - | - | - | - | - | - | - | - | - | - | - | - | - | - | - | - | - | - | - | - | - | - | - | - | - | - | - | - | - | - | - | - | - | - | - | - | - | - | - | - | - | - | - | - | - | - | - | - | - | - | - | - | - | - | - | - | - | - | - | - | - | - | - | - | - | - | - | - | - | - | - | - | - | - | - | - | - | - | - | - | - | - | - | - | - | - | - | - | - | - | - | - | - | - | - | - | - | - | - | - | - | - | - | - | - | - | - | - | - | - | - | - | - | - | - | - | - | - | - | - | - | - | - | - | - | - | - | - | - | - | - | - | - | - | - | - | - | - | - | - | - | - | - | - | - | - | - | - | - | - | - | - | - | - | - | - | - | - | - | - | - | - | - | - | - | - | - | - | - | - | - | - | - | - | - | - | - | - | - | - | - | - | - | - | - | - | - | - | - | - | - | - | - | - | - | - | - | - | - | - | - | - | - | - | - | - | - | - | - | - | - | - | - | - | - | - | - | - | - | - | - | - | - | - | - | - | - | - | - | - | - | - | - | - | - | - | - | - | - | - | - | - | - | - | - | - | - | - | - | - | - | - | - | - | - | - | - | - | - | - | - | - | - | - | - | - | - | - | - | - | - | - | - | - | - | - | - | - | - | - | - | - | - | - | - | - | - | - | - | - | - | - | - | - | - | - | - | - | - | - | - | - | - | - | - | - | - | - | - | - | - | - | - | - | - | - | - | - | - | - | - | - | - | - | - | - | - | - | - | - | - | - | - | - | - | - | - | - | - | - | - | - | - | - | - | - | - | - | - | - | - | - | - | - | - | - | - | - | - | - | - | - | - | - | - | - | - | - | - | - | - | - | - | - | - | - | - | - | - | - | - | - | - | - | - | - | - | - | - | - | - | - | - | - | - | - | - | - | - | - | - | - | - | - | - | - | - | - | - | - | - | - | - | - | - | - | - | - | - | - | - | - | - | - | - | - | - | - | - | - | - | - | - | - | - | - | - | - | - | - | - | - | - | - | - | - | - | - | - | - | - | - | - | - | - | - | - | - | - | - | - | - | - | - | - | - | - | - | - | - | - | - | - | - | - | - | - | - | - | - | - | - | - | - | - | - | - | - | - | - | - | - | - | - | - | - | - | - | - | - | - | - | - | - | - | - | - | - | - | - | - | - | - | - | - | - | - | - | - | - | - | - | - | - | - | - | - | - | - | - | - | - | - | - | - | - | - | - | - | - | - | - | - | - | - | - | - | - | - | - | - | - | - | - | - | - | - | - | - | - | - | - | - | - | - | - | - | - | - | - | - | - | - | - | - | - | - | - | - | - | - | - | - | - | - | - | - | - | - | - | - | - | - | - | - | - | - | - | - | - | - | - | - | - | - | - | - | - | - | - | - | - | - | - | - | - | - | - | - | - | - | - | - | - | - | - | - | - | - | - | - | - | - | - | - | - | - | - | - | - | - | - | - | - | - | - | - | - | - | - | - | - | - | - | - | - | - | - | - | - | - | - | - | - | - | - | - | - | - | - | - | - | - | - | - | - | - | - | - | - | - | - | - | - | - | - | - | - | - | - | - | - | - | - | - | - | - | - | - | - | - | - | - | - | - | - | - | - | - | - | - | - | - | - | - | - | - | - | - | - | - | - | - | - | - | - | - | - | - | - | - | - | - | - | - | - | - | - | - | - | - | - | - | - | - | - | - | - | - | - | - | - | - | - | - | - | - | - | - | - | - | - | - | - | - | - | - | - | - | - | - | - | - | - | - | - | - | - | - | - | - | - | - | - | - | - | - | - | - | - | - | - | - | - | - | - | - | - | - | - | - | - | - | - | - | - | - | - | - | - | - | - | - | - | - | - | - | - | - | - | - | - | - | - | - | - | - | - | - | - | - | - | - | - | - | - | - | - | - | - | - | - | - | - | - | - | - | - | - | - | - | - | - | - | - | - | - | - | - | - | - | - | - | - | - | - | - | - | - | - | - | - | - | - | - | - | - | - | - | - | - | - | - | - | - | - | - | - | - | - | - | - | - | - | - | - | - | - | - | - | - | - | - | - | - | - | - | - | - | - | - | - | - | - | - | - | - | - | - | - | - | - | - | - | - | - | - | - | - | - | - | - | - | - | - | - | - | - | - | - | - | - | - | - | - | - | - | - | - | - | - | - | - | - | - | - | - | - | - | - | - | - | - | - | - | - | - | - | - | - | - | - | - | - | - | - | - | - | - | - | - | - | - | - | - | - | - | - | - | - | - | - | - | - | - | - | - | - | - | - | - | - | - | - | - | - | - | - | - | - | - | - | - | - | - | - | - | - | - | - | - | - | - | - | - | - | - | - | - | - | - | - | - | - | - | - | - | - | - | - | - | - | - | - | - | - | - | - | - | - | - | - | - | - | - | - | - | - | - | - | - | - | - | - | - | - | - | - | - | - | - | - | - | - | - | - | - | - | - | - | - | - | - | - | - | - | - | - | - | - | - | - | - | - | - | - | - | - | - | - | - | - | - | - | - | - | - | - | - | - | - | - | - | - | - | - | - | - | - | - | - | - | - | - | - | - | - | - | - | - | - | - | - | - | - | - | - | - | - | - | - | - | - | - | - | - | - | - | - | - | - | - | - | - | - | - | - | - | - | - | - | - | - | - | - | - | - | - | - | - | - | - | - | - | - | - | - | - | - | - | - | - | - | - | - | - | - | - | - | - | - | - | - | - | - | - | - | - | - | - | - | - | - | - | - | - | - | - | - | - | - | - | - | - | - | - | - | - | - | - | - | - | - | - | - | - | - | - | - | - | - | - | - | - | - | - | - | - | - | - | - | - | - | - | - | - | - | - | - | - | - | - | - | - | - | - | - | - | - | - | - | - | - | - | - | - | - | - | - | - | - | - | - | - | - | - | - | - | - | - | - | - | - | - | - | - | - | - | - | - | - | - | - | - | - | - | - | - | - | - | - | - | - | - | - | - | - | - | - | - | - | - | - | - | - | - | - | - | - | - | - | - | - | - | - | - | - | - | - | - | - | - | - | - | - | - | - | - | - | - | - | - | - | - | - | - | - | - | - | - | - | - | - | - | - | - | - | - | - | - | - | - | - | - | - | - | - | - | - | - | - | - | - | - | - | - | - | - | - | - | - | - | - | - | - | - | - | - | - | - | - | - | - | - | - | - | - | - | - | - | - | - | - | - | - | - | - | - | - | - | - | - | - | - | - | - | - | - | - | - | - | - | - | - | - | - | - | - | - | - | - | - | - | - | - | - | - | - | - | - | - | - | - | - | - | - | - | - | - | - | - | - | - | - | - | - | - | - | - | - | - | - | - | - | - | - | - | - | - | - | - | - | - | - | - | - | - | - | - | - | - | - | - | - | - | - | - | - | - | - | - | - | - | - | - | - | - | - | - | - | - | - | - | - | - | - | - | - | - | - | - | - | - | - | - | - | - | - | - | - | - | - | - | - | - | - | - | - | - | - | - | - | - | - | - | - | - | - | - | - | - | - | - | - | - | - | - | - | - | - | - | - | - | - | - | - | - | - | - | - | - | - | - | - | - | - | - | - | - | - | - | - | - | - | - | - | - | - | - | - | - | - | - | - | - | - | - | - | - | - | - | - | - | - | - | - | - | - | - | - | - | - | - | - | - | - | - | - | - | - | - | - | - | - | - | - | - | - | - | - | - | - | - | - | - | - | - | - | - | - | - | - | - | - | - | - | - | - | - | - | - | - | - | - | - | - | - | - | - | - | - | - | - | - | - | - | - | - | - | - | - | - | - | - | - | - | - | - | - | - | - | - | - | - | - | - | - | - | - | - | - | - | - | - | - | - | - | - | - | - | - | - | - | - | - | - | - | - | - | - | - | - | - | - | - | - | - | - | - | - | - | - | - | - | - | - | - | - | - | - | - | - | - | - | - | - | - | - | - | - | - | - | - | - | - | - | - | - | - | - | - | - | - | - | - | - | - | - | - | - | - | - | - | - | - | - | - | - | - | - | - | - | - | - | - | - | - | - | - | - | - | - | - | - | - | - | - | - | - | - | - | - | - | - | - | - | - | - | - | - | - | - | - | - | - | - | - | - | - | - | - | - | - | - | - | - | - | - | - | - | - | - | - | - | - | - | - | - | - | - | - | - | - | - | - | - | - | - | - | - | - | - | - | - | - | - | - | - | - | - | - | - | - | - | - | - | - | - | - | - | - | - | - | - | - | - | - | - | - | - | - | - | - | - | - | - | - | - | - | - | - | - | - | - | - | - | - | - | - | - | - | - | - | - | - | - | - | - | - | - | - | - | - | - | - | - | - | - | - | - | - | - | - | - | - | - | - | - | - | - | - | - | - | - | - | - | - | - | - | - | - | - | - | - | - | - | - | - | - | - | - | - | - | - | - | - | - | - | - | - | - | - | - | - | - | - | - | - | - | - | - | - | - | - | - | - | - | - | - | - | - | - | - | - | - | - | - | - | - | - | - | - | - | - | - | - | - | - | - | - | - | - | - | - | - | - | - | - | - | - | - | - | - | - | - | - | - | - | - | - | - | - | - | - | - | - | - | - | - | - | - | - | - | - | - | - | - | - | - | - | - | - | - | - | - | - | - | - | - | - | - | - | - | - | - | - | - | - | - | - | - | - | - | - | - | - | - | - | - | - | - | - | - | - | - | - | - | - | - | - | - | - | - | - | - | - | - | - | - | - | - | - | - | - | - | - | - | - | - | - | - | - | - | - | - | - | - | - | - | - | - | - | - | - | - | - | - |
| BIS d=0 cl.10 sym=1 env=1 | - | - | - | - | - | - | - | - | - | - | - | - | - | - | - | - | - | - | - | - | - | - | - | - | - | - | - | - | - | - | - | - | - | - | - | - | - | - | - | - | - | - | - | - | - | - | - | - | - | - | - | - | - | - | - | - | - | - | - | - | - | - | - | - | - | - | - | - | - | - | - | - | - | - | - | - | - | - | - | - | - | - | - | - | - | - | - | - | - | - | - | - | - | - | - | - | - | - | - | - | - | - | - | - | - | - | - | - | - | - | - | - | - | - | - | - | - | - | - | - | - | - | - | - | - | - | - | - | - | - | - | - | - | - | - | - | - | - | - | - | - | - | - | - | - | - | - | - | - | - | - | - | - | - | - | - | - | - | - | - | - | - | - | - | - | - | - | - | - | - | - | - | - | - | - | - | - | - | - | - | - | - | - | - | - | - | - | - | - | - | - | - | - | - | - | - | - | - | - | - | - | - | - | - | - | - | - | - | - | - | - | - | - | - | - | - | - | - | - | - | - | - | - | - | - | - | - | - | - | - | - | - | - | - | - | - | - | - | - | - | - | - | - | - | - | - | - | - | - | - | - | - | - | - | - | - | - | - | - | - | - | - | - | - | - | - | - | - | - | - | - | - | - | - | - | - | - | - | - | - | - | - | - | - | - | - | - | - | - | - | - | - | - | - | - | - | - | - | - | - | - | - | - | - | - | - | - | - | - | - | - | - | - | - | - | - | - | - | - | - | - | - | - | - | - | - | - | - | - | - | - | - | - | - | - | - | - | - | - | - | - | - | - | - | - | - | - | - | - | - | - | - | - | - | - | - | - | - | - | - | - | - | - | - | - | - | - | - | - | - | - | - | - | - | - | - | - | - | - | - | - | - | - | - | - | - | - | - | - | - | - | - | - | - | - | - | - | - | - | - | - | - | - | - | - | - | - | - | - | - | - | - | - | - | - | - | - | - | - | - | - | - | - | - | - | - | - | - | - | - | - | - | - | - | - | - | - | - | - | - | - | - | - | - | - | - | - | - | - | - | - | - | - | - | - | - | - | - | - | - | - | - | - | - | - | - | - | - | - | - | - | - | - | - | - | - | - | - | - | - | - | - | - | - | - | - | - | - | - | - | - | - | - | - | - | - | - | - | - | - | - | - | - | - | - | - | - | - | - | - | - | - | - | - | - | - | - | - | - | - | - | - | - | - | - | - | - | - | - | - | - | - | - | - | - | - | - | - | - | - | - | - | - | - | - | - | - | - | - | - | - | - | - | - | - | - | - | - | - | - | - | - | - | - | - | - | - | - | - | - | - | - | - | - | - | - | - | - | - | - | - | - | - | - | - | - | - | - | - | - | - | - | - | - | - | - | - | - | - | - | - | - | - | - | - | - | - | - | - | - | - | - | - | - | - | - | - | - | - | - | - | - | - | - | - | - | - | - | - | - | - | - | - | - | - | - | - | - | - | - | - | - | - | - | - | - | - | - | - | - | - | - | - | - | - | - | - | - | - | - | - | - | - | - | - | - | - | - | - | - | - | - | - | - | - | - | - | - | - | - | - | - | - | - | - | - | - | - | - | - | - | - | - | - | - | - | - | - | - | - | - | - | - | - | - | - | - | - | - | - | - | - | - | - | - | - | - | - | - | - | - | - | - | - | - | - | - | - | - | - | - | - | - | - | - | - | - | - | - | - | - | - | - | - | - | - | - | - | - | - | - | - | - | - | - | - | - | - | - | - | - | - | - | - | - | - | - | - | - | - | - | - | - | - | - | - | - | - | - | - | - | - | - | - | - | - | - | - | - | - | - | - | - | - | - | - | - | - | - | - | - | - | - | - | - | - | - | - | - | - | - | - | - | - | - | - | - | - | - | - | - | - | - | - | - | - | - | - | - | - | - | - | - | - | - | - | - | - | - | - | - | - | - | - | - | - | - | - | - | - | - | - | - | - | - | - | - | - | - | - | - | - | - | - | - | - | - | - | - | - | - | - | - | - | - | - | - | - | - | - | - | - | - | - | - | - | - | - | - | - | - | - | - | - | - | - | - | - | - | - | - | - | - | - | - | - | - | - | - | - | - | - | - | - | - | - | - | - | - | - | - | - | - | - | - | - | - | - | - | - | - | - | - | - | - | - | - | - | - | - | - | - | - | - | - | - | - | - | - | - | - | - | - | - | - | - | - | - | - | - | - | - | - | - | - | - | - | - | - | - | - | - | - | - | - | - | - | - | - | - | - | - | - | - | - | - | - | - | - | - | - | - | - | - | - | - | - | - | - | - | - | - | - | - | - | - | - | - | - | - | - | - | - | - | - | - | - | - | - | - | - | - | - | - | - | - | - | - | - | - | - | - | - | - | - | - | - | - | - | - | - | - | - | - | - | - | - | - | - | - | - | - | - | - | - | - | - | - | - | - | - | - | - | - | - | - | - | - | - | - | - | - | - | - | - | - | - | - | - | - | - | - | - | - | - | - | - | - | - | - | - | - | - | - | - | - | - | - | - | - | - | - | - | - | - | - | - | - | - | - | - | - | - | - | - | - | - | - | - | - | - | - | - | - | - | - | - | - | - | - | - | - | - | - | - | - | - | - | - | - | - | - | - | - | - | - | - | - | - | - | - | - | - | - | - | - | - | - | - | - | - | - | - | - | - | - | - | - | - | - | - | - | - | - | - | - | - | - | - | - | - | - | - | - | - | - | - | - | - | - | - | - | - | - | - | - | - | - | - | - | - | - | - | - | - | - | - | - | - | - | - | - | - | - | - | - | - | - | - | - | - | - | - | - | - | - | - | - | - | - | - | - | - | - | - | - | - | - | - | - | - | - | - | - | - | - | - | - | - | - | - | - | - | - | - | - | - | - | - | - | - | - | - | - | - | - | - | - | - | - | - | - | - | - | - | - | - | - | - | - | - | - | - | - | - | - | - | - | - | - | - | - | - | - | - | - | - | - | - | - | - | - | - | - | - | - | - | - | - | - | - | - | - | - | - | - | - | - | - | - | - | - | - | - | - | - | - | - | - | - | - | - | - | - | - | - | - | - | - | - | - | - | - | - | - | - | - | - | - | - | - | - | - | - | - | - | - | - | - | - | - | - | - | - | - | - | - | - | - | - | - | - | - | - | - | - | - | - | - | - | - | - | - | - | - | - | - | - | - | - | - | - | - | - | - | - | - | - | - | - | - | - | - | - | - | - | - | - | - | - | - | - | - | - | - | - | - | - | - | - | - | - | - | - | - | - | - | - | - | - | - | - | - | - | - | - | - | - | - | - | - | - | - | - | - | - | - | - | - | - | - | - | - | - | - | - | - | - | - | - | - | - | - | - | - | - | - | - | - | - | - | - | - | - | - | - | - | - | - | - | - | - | - | - | - | - | - | - | - | - | - | - | - | - | - | - | - | - | - | - | - | - | - | - | - | - | - | - | - | - | - | - | - | - | - | - | - | - | - | - | - | - | - | - | - | - | - | - | - | - | - | - | - | - | - | - | - | - | - | - | - | - | - | - | - | - | - | - | - | - | - | - | - | - | - | - | - | - | - | - | - | - | - | - | - | - | - | - | - | - | - | - | - | - | - | - | - | - | - | - | - | - | - | - | - | - | - | - | - | - | - | - | - | - | - | - | - | - | - | - | - | - | - | - | - | - | - | - | - | - | - | - | - | - | - | - | - | - | - | - | - | - | - | - | - | - | - | - | - | - | - | - | - | - | - | - | - | - | - | - | - | - | - | - | - | - | - | - | - | - | - | - | - | - | - | - | - | - | - | - | - | - | - | - | - | - | - | - | - | - | - | - | - | - | - | - | - | - | - | - | - | - | - | - | - | - | - | - | - | - | - | - | - | - | - | - | - | - | - | - | - | - | - | - | - | - | - | - | - | - | - | - | - | - | - | - | - | - | - | - | - | - | - | - | - | - | - | - | - | - | - | - | - | - | - | - | - | - | - | - | - | - | - | - | - | - | - | - | - | - | - | - | - | - | - | - | - | - | - | - | - | - | - | - | - | - | - | - | - | - | - | - | - | - | - | - | - | - | - | - | - | - | - | - | - | - | - | - | - | - | - | - | - | - | - | - | - | - | - | - | - | - | - | - | - | - | - | - | - | - | - | - | - | - | - | - | - | - | - | - | - | - | - | - | - | - | - | - | - | - | - | - | - | - | - | - | - | - | - | - | - | - | - | - | - | - | - | - | - | - | - | - | - | - | - | - | - | - | - | - | - | - | - | - | - | - | - | - | - | - | - | - | - | - | - | - | - | - | - | - | - | - | - | - | - | - | - | - | - | - | - | - | - | - | - | - | - | - | - | - | - | - | - | - | - | - | - | - | - | - | - | - | - | - | - | - | - | - | - | - | - | - | - | - | - | - | - | - | - | - | - | - | - | - | - | - | - | - | - | - | - | - | - | - | - | - | - | - | - | - | - | - | - | - | - | - | - | - | - | - | - | - | - | - | - | - | - | - | - | - | - | - | - | - | - | - | - | - | - | - | - | - | - | - | - | - | - | - | - | - | - | - | - | - | - | - | - | - | - | - | - | - | - | - | - | - | - | - | - | - | - | - | - | - | - | - | - | - | - | - | - | - | - | - | - | - | - | - | - | - | - | - | - | - | - | - | - | - | - | - | - | - | - | - | - | - | - | - | - | - | - | - | - | - | - | - | - | - | - | - | - | - | - | - | - | - | - | - | - | - | - | - | - | - | - | - | - | - | - | - | - | - | - | - | - | - | - | - | - | - | - | - | - | - | - | - | - | - | - | - | - | - | - | - | - | - | - | - | - | - | - | - | - | - | - | - | - | - | - | - | - | - | - | - | - | - | - | - | - | - | - | - | - | - | - | - | - | - | - | - | - | - | - | - | - | - | - | - | - | - | - | - | - | - | - | - | - | - | - | - | - | - | - | - | - | - | - | - | - | - | - | - | - | - | - | - | - | - | - | - | - | - | - | - | - | - | - | - | - | - | - | - | - | - | - | - | - | - | - | - | - | - | - | - | - | - | - | - | - | - | - | - | - | - | - | - | - | - | - | - | - | - | - | - | - | - | - | - | - | - | - | - | - | - | - | - | - | - | - | - | - | - | - | - | - | - | - | - | - | - | - | - | - | - | - | - | - | - | - | - | - | - | - | - | - | - | - | - | - | - | - | - | - | - | - | - | - | - | - | - | - | - | - | - | - | - | - | - | - | - | - | - | - | - | - | - | - | - | - | - | - | - | - | - | - | - | - | - | - | - | - | - | - | - | - | - | - | - | - | - | - | - | - | - | - | - | - | - | - | - | - | - | - | - | - | - | - | - | - | - | - | - | - | - | - | - | - | - | - | - | - | - | - | - | - | - | - | - | - | - | - | - | - | - | - | - | - | - | - | - | - | - | - | - | - | - | - | - | - | - | - | - | - | - | - | - | - | - | - | - | - | - | - | - | - | - | - | - | - | - | - | - | - | - | - | - | - | - | - | - | - | - | - | - | - | - | - | - | - | - | - | - | - | - | - | - | - | - | - | - | - | - | - | - | - | - | - | - | - | - | - | - | - | - | - | - | - | - | - | - | - | - | - | - | - | - | - | - | - | - | - | - | - | - | - | - | - | - | - | - | - | - | - | - | - | - | - | - | - | - | - | - | - | - | - | - | - | - | - | - | - | - | - | - | - | - | - | - | - | - | - | - | - | - | - | - | - | - | - | - | - | - | - | - | - | - | - | - | - | - | - | - | - | - | - | - | - | - | - | - | - | - | - | - | - | - | - | - | - | - | - | - | - | - | - | - | - | - | - | - | - | - | - | - | - | - | - | - | - | - | - | - | - | - | - | - | - | - | - | - | - | - | - | - | - | - | - | - | - | - | - | - | - | - | - | - | - | - | - | - | - | - | - | - | - | - | - | - | - | - | - | - | - | - | - | - | - | - | - | - | - | - | - | - | - | - | - | - | - | - | - | - | - | - | - | - | - | - | - | - | - | - | - | - | - | - | - | - | - | - | - | - | - | - | - | - | - | - | - | - | - | - | - | - | - | - | - | - | - | - | - | - | - | - | - | - | - | - | - | - | - | - | - | - | - | - | - | - | - | - | - | - | - | - | - | - | - | - | - | - | - | - | - | - | - | - | - | - | - | - | - | - | - | - | - | - | - | - | - | - | - | - | - | - | - | - | - | - | - | - | - | - | - | - | - | - | - | - | - | - | - | - | - | - | - | - | - | - | - | - | - | - | - | - | - | - | - | - | - | - | - | - | - | - | - | - | - | - | - | - | - | - | - | - | - | - | - | - | - | - | - | - | - | - | - | - | - | - | - | - | - | - | - | - | - | - | - | - | - | - | - | - | - | - | - | - | - | - | - | - | - | - | - | - | - | - | - | - | - | - | - | - | - | - | - | - | - | - | - | - | - | - | - | - | - | - | - | - | - | - | - | - | - | - | - | - | - | - | - | - | - | - | - | - | - | - | - | - | - | - | - | - | - | - | - | - | - | - | - | - | - | - | - | - | - | - | - | - | - | - | - | - | - | - | - | - | - | - | - | - | - | - | - | - | - | - | - | - | - | - | - | - | - | - | - | - | - | - | - | - | - | - | - | - | - | - | - | - | - | - | - | - | - | - | - | - | - | - | - | - | - | - | - | - | - | - | - | - | - | - | - | - | - | - | - | - | - | - | - | - | - | - | - | - | - | - | - | - | - | - | - | - | - | - | - | - | - | - | - | - | - | - | - | - | - | - | - | - | - | - | - | - | - | - | - | - | - | - | - | - | - | - | - | - | - | - | - | - | - | - | - | - | - | - | - | - | - | - | - | - | - | - | - | - | - | - | - | - | - | - | - | - | - | - | - | - | - | - | - | - | - | - | - | - | - | - | - | - | - | - | - | - | - | - | - | - | - | - | - | - | - | - | - | - | - | - | - | - | - | - | - | - | - | - | - | - | - | - | - | - | - | - | - | - | - | - | - | - | - | - | - | - | - | - | - | - | - | - | - | - | - | - | - | - | - | - | - | - | - | - | - | - | - | - | - | - | - | - | - | - | - | - | - | - | - | - | - | - | - | - | - | - | - | - | - | - | - | - | - | - | - | - | - | - | - | - | - | - | - | - | - | - | - | - | - | - | - | - | - | - | - | - | - | - | - | - | - | - | - | - | - | - | - | - | - | - | - | - | - | - | - | - | - | - | - | - | - | - | - | - | - | - | - | - | - | - | - | - | - | - | - | - | - | - | - | - | - | - | - | - | - | - | - | - | - | - | - | - | - | - | - | - | - | - | - | - | - | - | - | - | - | - | - | - | - | - | - | - | - | - | - | - | - | - | - | - | - | - | - | - | - | - | - | - | - | - | - | - | - | - | - | - | - | - | - | - | - | - | - | - | - | - | - | - | - | - | - | - | - | - | - | - | - | - | - | - | - | - | - | - | - | - | - | - | - | - | - | - | - | - | - | - | - | - | - | - | - | - | - | - | - | - | - | - | - | - | - | - | - | - | - | - | - | - | - | - | - | - | - | - | - | - | - | - | - | - | - | - | - | - | - | - | - | - | - | - | - | - | - | - | - | - | - | - | - | - | - | - | - | - | - | - | - | - | - | - | - | - | - | - | - | - | - | - | - | - | - | - | - | - | - | - | - | - | - | - | - | - | - | - | - | - | - | - | - | - | - | - | - | - | - | - | - | - | - | - | - | - | - | - | - | - | - | - | - | - | - | - | - | - | - | - | - | - | - | - | - | - | - | - | - | - | - | - | - | - | - | - | - | - | - | - | - | - | - | - | - | - | - | - | - | - | - | - | - | - | - | - | - | - | - | - | - | - | - | - | - | - | - | - | - | - | - | - | - | - | - | - | - | - | - | - | - | - | - | - | - | - | - | - | - | - | - | - | - | - | - | - | - | - | - | - | - | - | - | - | - | - | - | - | - | - | - | - | - | - | - | - | - | - | - | - | - | - | - | - | - | - | - | - | - | - | - | - | - | - | - | - | - | - | - | - | - | - | - | - | - | - | - | - | - | - | - | - | - | - | - | - | - | - | - | - | - | - | - | - | - | - | - | - | - | - | - | - | - | - | - | - | - | - | - | - | - | - | - | - | - | - | - | - | - | - | - | - | - | - | - | - | - | - | - | - | - | - | - | - | - | - | - | - | - | - | - | - | - | - | - | - | - | - | - | - | - | - | - | - | - | - | - | - | - | - | - | - | - | - | - | - | - | - | - | - | - | - | - | - | - | - | - | - | - | - | - | - | - | - | - | - | - | - | - | - | - | - | - | - | - | - | - | - | - | - | - | - | - | - | - | - | - | - | - | - | - | - | - | - | - | - | - | - | - | - | - | - | - | - | - | - | - | - | - | - | - | - | - | - | - | - | - | - | - | - | - | - | - | - | - | - | - | - | - | - | - | - | - | - | - | - | - | - | - | - | - | - | - | - | - | - | - | - | - | - | - | - | - | - | - | - | - | - | - | - | - | - | - | - | - | - | - | - | - | - | - | - | - | - | - | - | - | - | - | - | - | - | - | - | - | - | - | - | - | - | - | - | - | - | - | - | - | - | - | - | - | - | - | - | - | - | - | - | - | - | - | - | - | - | - | - | - | - | - | - | - | - | - | - | - | - | - | - | - | - | - | - | - | - | - | - | - | - | - | - | - | - | - | - | - | - | - | - | - | - | - | - | - | - | - | - | - | - | - | - | - | - | - | - | - | - | - | - | - | - | - | - | - | - | - | - | - | - | - | - | - | - | - | - | - | - | - | - | - | - | - | - | - | - | - | - | - | - | - | - | - | - | - | - | - | - | - | - | - | - | - | - | - | - | - | - | - | - | - | - | - | - | - | - | - | - | - | - | - | - | - | - | - | - | - | - | - | - | - | - | - | - | - | - | - | - | - | - | - | - | - | - | - | - | - | - | - | - | - | - | - | - | - | - | - | - | - | - | - | - | - | - | - | - | - | - | - | - | - | - | - | - | - | - | - | - | - | - | - | - | - | - | - | - | - | - | - | - | - | - | - | - | - | - | - | - | - | - | - | - | - | - | - | - | - | - | - | - | - | - | - | - | - | - | - | - | - | - | - | - | - | H | x | H | x | x | x | x | x | x | H | - | - | - | - | - | - | - | - | - | - | - | - | - | - | - | - | - | - | - | - | - | - | - | - | - | - | - | - | - | - | - | - | - | - | - | - | - | - | - | - | - | - | - | - | - | - | - | - | - | - | - | - | - | - | - | - | - | - | - | - | - | - | - | - | - | - | - | - | - | - | - | - | - | - | - | - | - | - | - | - | - | - | - | - | - | - | - | - | - | - | - | - | - | - | - | - | - | - | - | - | - | - | - | - | - | - | - | - | - | - | - | - | - | - | - | - | - | - | - | - | - | - | - | - | - | - | - | - | - | - | - | - | - | - | - | - | - | - | - | - | - | - | - | - | - | - | - | - | - | - | - | - | - | - | - | - | - | - | - | - | - | - | - | - | - | - | - | - | - | - | - | - | - | - | - | - | - | - | - | - | - | - | - | - | - | - | - | - | - | - | - | - | - | - | - | - | - | - | - | - | - | - | - | - | - | - | - | - | - | - | - | - | - | - | - | - | - | - | - | - | - | - | - | - | - | - | - | - | - | - | - | - | - | - | - | - | - | - | - | - | - | - | - | - | - | - | - | - | - | - | - | - | - | - | - | - | - | - | - | - | - | - | - | - | - | - | - | - | - | - | - | - | - | - | - | - | - | - | - | - | - | - | - | - | - | - | - | - | - | - | - | - | - | - | - | - | - | - | - | - | - | - | - | - | - | - | - | - | - | - | - | - | - | - | - | - | - | - | - | - | - | - | - | - | - | - | - | - | - | - | - | - | - | - | - | - | - | - | - | - | - | - | - | - | - | - | - | - | - | - | - | - | - | - | - | - | - | - | - | - | - | - | - | - | - | - | - | - | - | - | - | - | - | - | - | - | - | - | - | - | - | - | - | - | - | - | - | - | - | - | - | - | - | - | - | - | - | - | - | - | - | - | - | - | - | - | - | - | - | - | - | - | - | - | - | - | - | - | - | - | - | - | - | - | - | - | - | - | - | - | - | - | - | - | - | - | - | - | - | - | - | - | - | - | - | - | - | - | - | - | - | - | - | - | - | - | - | - | - | - | - | - | - | - | - | - | - | - | - | - | - | - | - | - | - | - | - | - | - | - | - | - | - | - | - | - | - | - | - | - | - | - | - | - | - | - | - | - | - | - | - | - | - | - | - | - | - | - | - | - | - | - | - | - | - | - | - | - | - | - | - | - | - | - | - | - | - | - | - | - | - | - | - | - | - | - | - | - | - | - | - | - | - | - | - | - | - | - | - | - | - | - | - | - | - | - | - | - | - | - | - | - | - | - | - | - | - | - | - | - | - | - | - | - | - | - | - | - | - | - | - | - | - | - | - | - | - | - | - | - | - | - | - | - | - | - | - | - | - | - | - | - | - | - | - | - | - | - | - | - | - | - | - | - | - | - | - | - | - | - | - | - | - | - | - | - | - | - | - | - | - | - | - | - | - | - | - | - | - | - | - | - | - | - | - | - | - | - | - | - | - | - | - | - | - | - | - | - | - | - | - | - | - | - | - | - | - | - | - | - | - | - | - | - | - | - | - | - | - | - | - | - | - | - | - | - | - | - | - | - | - | - | - | - | - | - | - | - | - | - | - | - | - | - | - | - | - | - | - | - | - | - | - | - | - | - | - | - | - | - | - | - | - | - | - | - | - | - | - | - | - | - | - | - | - | - | - | - | - | - | - | - | - | - | - | - | - | - | - | - | - | - | - | - | - | - | - | - | - | - | - | - | - | - | - | - | - | - | - | - | - | - | - | - | - | - | - | - | - | - | - | - | - | - | - | - | - | - | - | - | - | - | - | - | - | - | - | - | - | - | - | - | - | - | - | - | - | - | - | - | - | - | - | - | - | - | - | - | - | - | - | - | - | - | - | - | - | - | - | - | - | - | - | - | - | - | - | - | - | - | - | - | - | - | - | - | - | - | - | - | - | - | - | - | - | - | - | - | - | - | - | - | - | - | - | - | - | - | - | - | - | - | - | - | - | - | - | - | - | - | - | - | - | - | - | - | - | - | - | - | - | - | - | - | - | - | - | - | - | - | - | - | - | - | - | - | - | - | - | - | - | - | - | - | - | - | - | - | - | - | - | - | - | - | - | - | - | - | - | - | - | - | - | - | - | - | - | - | - | - | - | - | - | - | - | - | - | - | - | - | - | - | - | - | - | - | - | - | - | - | - | - | - | - | - | - | - | - | - | - | - | - | - | - | - | - | - | - | - | - | - | - | - | - | - | - | - | - | - | - | - | - | - | - | - | - | - | - | - | - | - | - | - | - | - | - | - | - | - | - | - | - | - | - | - | - | - | - | - | - | - | - | - | - | - | - | - | - | - | - | - | - | - | - | - | - | - | - | - | - | - | - | - | - | - | - | - | - | - | - | - | - | - | - | - | - | - | - | - | - | - | - | - | - | - | - | - | - | - | - | - | - | - | - | - | - | - | - | - | - | - | - | - | - | - | - | - | - | - | - | - | - | - | - | - | - | - | - | - | - | - | - | - | - | - | - | - | - | - | - | - | - | - | - | - | - | - | - | - | - | - | - | - | - | - | - | - | - | - | - | - | - | - | - | - | - | - | - | - | - | - | - | - | - | - | - | - | - | - | - | - | - | - | - | - | - | - | - | - | - | - | - | - | - | - | - | - | - | - | - | - | - | - | - | - | - | - | - | - | - | - | - | - | - | - | - | - | - | - | - | - | - | - | - | - | - | - | - | - | - | - | - | - | - | - | - | - | - | - | - | - | - | - | - | - | - | - | - | - | - | - | - | - | - | - | - | - | - | - | - | - | - | - | - | - | - | - | - | - | - | - | - | - | - | - | - | - | - | - | - | - | - | - | - | - | - | - | - | - | - | - | - | - | - | - | - | - | - | - | - | - | - | - | - | - | - | - | - | - | - | - | - | - | - | - | - | - | - | - | - | - | - | - | - | - | - | - | - | - | - | - | - | - | - | - | - | - | - | - | - | - | - | - | - | - | - | - | - | - | - | - | - | - | - | - | - | - | - | - | - | - | - | - | - | - | - | - | - | - | - | - | - | - | - | - | - | - | - | - | - | - | - | - | - | - | - | - | - | - | - | - | - | - | - | - | - | - | - | - | - | - | - | - | - | - | - | - | - | - | - | - | - | - | - | - | - | - | - | - | - | - | - | - | - | - | - | - | - | - | - | - | - | - | - | - | - | - | - | - | - | - | - | - | - | - |
| BIS d=0 cl.15 sym=1 env=1 | - | - | - | - | - | - | - | - | - | - | - | - | - | - | - | - | - | - | - | - | - | - | - | - | - | - | - | - | - | - | - | - | - | - | - | - | - | - | - | - | - | - | - | - | - | - | - | - | - | - | - | - | - | - | - | - | - | - | - | - | - | - | - | - | - | - | - | - | - | - | - | - | - | - | - | - | - | - | - | - | - | - | - | - | - | - | - | - | - | - | - | - | - | - | - | - | - | - | - | - | - | - | - | - | - | - | - | - | - | - | - | - | - | - | - | - | - | - | - | - | - | - | - | - | - | - | - | - | - | - | - | - | - | - | - | - | - | - | - | - | - | - | - | - | - | - | - | - | - | - | - | - | - | - | - | - | - | - | - | - | - | - | - | - | - | - | - | - | - | - | - | - | - | - | - | - | - | - | - | - | - | - | - | - | - | - | - | - | - | - | - | - | - | - | - | - | - | - | - | - | - | - | - | - | - | - | - | - | - | - | - | - | - | - | - | - | - | - | - | - | - | - | - | - | - | - | - | - | - | - | - | - | - | - | - | - | - | - | - | - | - | - | - | - | - | - | - | - | - | - | - | - | - | - | - | - | - | - | - | - | - | - | - | - | - | - | - | - | - | - | - | - | - | - | - | - | - | - | - | - | - | - | - | - | - | - | - | - | - | - | - | - | - | - | - | - | - | - | - | - | - | - | - | - | - | - | - | - | - | - | - | - | - | - | - | - | - | - | - | - | - | - | - | - | - | - | - | - | - | - | - | - | - | - | - | - | - | - | - | - | - | - | - | - | - | - | - | - | - | - | - | - | - | - | - | - | - | - | - | - | - | - | - | - | - | - | - | - | - | - | - | - | - | - | - | - | - | - | - | - | - | - | - | - | - | - | - | - | - | - | - | - | - | - | - | - | - | - | - | - | - | - | - | - | - | - | - | - | - | - | - | - | - | - | - | - | - | - | - | - | - | - | - | - | - | - | - | - | - | - | - | - | - | - | - | - | - | - | - | - | - | - | - | - | - | - | - | - | - | - | - | - | - | - | - | - | - | - | - | - | - | - | - | - | - | - | - | - | - | - | - | - | - | - | - | - | - | - | - | - | - | - | - | - | - | - | - | - | - | - | - | - | - | - | - | - | - | - | - | - | - | - | - | - | - | - | - | - | - | - | - | - | - | - | - | - | - | - | - | - | - | - | - | - | - | - | - | - | - | - | - | - | - | - | - | - | - | - | - | - | - | - | - | - | - | - | - | - | - | - | - | - | - | - | - | - | - | - | - | - | - | - | - | - | - | - | - | - | - | - | - | - | - | - | - | - | - | - | - | - | - | - | - | - | - | - | - | - | - | - | - | - | - | - | - | - | - | - | - | - | - | - | - | - | - | - | - | - | - | - | - | - | - | - | - | - | - | - | - | - | - | - | - | - | - | - | - | - | - | - | - | - | - | - | - | - | - | - | - | - | - | - | - | - | - | - | - | - | - | - | - | - | - | - | - | - | - | - | - | - | - | - | - | - | - | - | - | - | - | - | - | - | - | - | - | - | - | - | - | - | - | - | - | - | - | - | - | - | - | - | - | - | - | - | - | - | - | - | - | - | - | - | - | - | - | - | - | - | - | - | - | - | - | - | - | - | - | - | - | - | - | - | - | - | - | - | - | - | - | - | - | - | - | - | - | - | - | - | - | - | - | - | - | - | - | - | - | - | - | - | - | - | - | - | - | - | - | - | - | - | - | - | - | - | - | - | - | - | - | - | - | - | - | - | - | - | - | - | - | - | - | - | - | - | - | - | - | - | - | - | - | - | - | - | - | - | - | - | - | - | - | - | - | - | - | - | - | - | - | - | - | - | - | - | - | - | - | - | - | - | - | - | - | - | - | - | - | - | - | - | - | - | - | - | - | - | - | - | - | - | - | - | - | - | - | - | - | - | - | - | - | - | - | - | - | - | - | - | - | - | - | - | - | - | - | - | - | - | - | - | - | - | - | - | - | - | - | - | - | - | - | - | - | - | - | - | - | - | - | - | - | - | - | - | - | - | - | - | - | - | - | - | - | - | - | - | - | - | - | - | - | - | - | - | - | - | - | - | - | - | - | - | - | - | - | - | - | - | - | - | - | - | - | - | - | - | - | - | - | - | - | - | - | - | - | - | - | - | - | - | - | - | - | - | - | - | - | - | - | - | - | - | - | - | - | - | - | - | - | - | - | - | - | - | - | - | - | - | - | - | - | - | - | - | - | - | - | - | - | - | - | - | - | - | - | - | - | - | - | - | - | - | - | - | - | - | - | - | - | - | - | - | - | - | - | - | - | - | - | - | - | - | - | - | - | - | - | - | - | - | - | - | - | - | - | - | - | - | - | - | - | - | - | - | - | - | - | - | - | - | - | - | - | - | - | - | - | - | - | - | - | - | - | - | - | - | - | - | - | - | - | - | - | - | - | - | - | - | - | - | - | - | - | - | - | - | - | - | - | - | - | - | - | - | - | - | - | - | - | - | - | - | - | - | - | - | - | - | - | - | - | - | - | - | - | - | - | - | - | - | - | - | - | - | - | - | - | - | - | - | - | - | - | - | - | - | - | - | - | - | - | - | - | - | - | - | - | - | - | - | - | - | - | - | - | - | - | - | - | - | - | - | - | - | - | - | - | - | - | - | - | - | - | - | - | - | - | - | - | - | - | - | - | - | - | - | - | - | - | - | - | - | - | - | - | - | - | - | - | - | - | - | - | - | - | - | - | - | - | - | - | - | - | - | - | - | - | - | - | - | - | - | - | - | - | - | - | - | - | - | - | - | - | - | - | - | - | - | - | - | - | - | - | - | - | - | - | - | - | - | - | - | - | - | - | - | - | - | - | - | - | - | - | - | - | - | - | - | - | - | - | - | - | - | - | - | - | - | - | - | - | - | - | - | - | - | - | - | - | - | - | - | - | - | - | - | - | - | - | - | - | - | - | - | - | - | - | - | - | - | - | - | - | - | - | - | - | - | - | - | - | - | - | - | - | - | - | - | - | - | - | - | - | - | - | - | - | - | - | - | - | - | - | - | - | - | - | - | - | - | - | - | - | - | - | - | - | - | - | - | - | - | - | - | - | - | - | - | - | - | - | - | - | - | - | - | - | - | - | - | - | - | - | - | - | - | - | - | - | - | - | - | - | - | - | - | - | - | - | - | - | - | - | - | - | - | - | - | - | - | - | - | - | - | - | - | - | - | - | - | - | - | - | - | - | - | - | - | - | - | - | - | - | - | - | - | - | - | - | - | - | - | - | - | - | - | - | - | - | - | - | - | - | - | - | - | - | - | - | - | - | - | - | - | - | - | - | - | - | - | - | - | - | - | - | - | - | - | - | - | - | - | - | - | - | - | - | - | - | - | - | - | - | - | - | - | - | - | - | - | - | - | - | - | - | - | - | - | - | - | - | - | - | - | - | - | - | - | - | - | - | - | - | - | - | - | - | - | - | - | - | - | - | - | - | - | - | - | - | - | - | - | - | - | - | - | - | - | - | - | - | - | - | - | - | - | - | - | - | - | - | - | - | - | - | - | - | - | - | - | - | - | - | - | - | - | - | - | - | - | - | - | - | - | - | - | - | - | - | - | - | - | - | - | - | - | - | - | - | - | - | - | - | - | - | - | - | - | - | - | - | - | - | - | - | - | - | - | - | - | - | - | - | - | - | - | - | - | - | - | - | - | - | - | - | - | - | - | - | - | - | - | - | - | - | - | - | - | - | - | - | - | - | - | - | - | - | - | - | - | - | - | - | - | - | - | - | - | - | - | - | - | - | - | - | - | - | - | - | - | - | - | - | - | - | - | - | - | - | - | - | - | - | - | - | - | - | - | - | - | - | - | - | - | - | - | - | - | - | - | - | - | - | - | - | - | - | - | - | - | - | - | - | - | - | - | - | - | - | - | - | - | - | - | - | - | - | - | - | - | - | - | - | - | - | - | - | - | - | - | - | - | - | - | - | - | - | - | - | - | - | - | - | - | - | - | - | - | - | - | - | - | - | - | - | - | - | - | - | - | - | - | - | - | - | - | - | - | - | - | - | - | - | - | - | - | - | - | - | - | - | - | - | - | - | - | - | - | - | - | - | - | - | - | - | - | - | - | - | - | - | - | - | - | - | - | - | - | - | - | - | - | - | - | - | - | - | - | - | - | - | - | - | - | - | - | - | - | - | - | - | - | - | - | - | - | - | - | - | - | - | - | - | - | - | - | - | - | - | - | - | - | - | - | - | - | - | - | - | - | - | - | - | - | - | - | - | - | - | - | - | - | - | - | - | - | - | - | - | - | - | - | - | - | - | - | - | - | - | - | - | - | - | - | - | - | - | - | - | - | - | - | - | - | - | - | - | - | - | - | - | - | - | - | - | - | - | - | - | - | - | - | - | - | - | - | - | - | - | - | - | - | - | - | - | - | - | - | - | - | - | - | - | - | - | - | - | - | - | - | - | - | - | - | - | - | - | - | - | - | - | - | - | - | - | - | - | - | - | - | - | - | - | - | - | - | - | - | - | - | - | - | - | - | - | - | - | - | - | - | - | - | - | - | - | - | - | - | - | - | - | - | - | - | - | - | - | - | - | - | - | - | - | - | - | - | - | - | - | - | - | - | - | - | - | - | - | - | - | - | - | - | - | - | - | - | - | - | - | - | - | - | - | - | - | - | - | - | - | - | - | - | - | - | - | - | - | - | - | - | - | - | - | - | - | - | - | - | - | - | - | - | - | - | - | - | - | - | - | - | - | - | - | - | - | - | - | - | - | - | - | - | - | - | - | - | - | - | - | - | - | - | - | - | - | - | - | - | - | - | - | - | - | - | - | - | - | - | - | - | - | - | - | - | - | - | - | - | - | - | - | - | - | - | - | - | - | - | - | - | - | - | - | - | - | - | - | - | - | - | - | - | - | - | - | - | - | - | - | - | - | - | - | - | - | - | - | - | - | - | - | - | - | - | - | - | - | - | - | - | - | - | - | - | - | - | - | - | - | - | - | - | - | - | - | - | - | - | - | - | - | - | - | - | - | - | - | - | - | - | - | - | - | - | - | - | - | - | - | - | - | - | - | - | - | - | - | - | - | - | - | - | - | - | - | - | - | - | - | - | - | - | - | - | - | - | - | - | - | - | - | - | - | - | - | - | - | - | - | - | - | - | - | - | - | - | - | - | - | - | - | - | - | - | - | - | - | - | - | - | - | - | - | - | - | - | - | - | - | - | - | - | - | - | - | - | - | - | - | - | - | - | - | - | - | - | - | - | - | - | - | - | - | - | - | - | - | - | - | - | - | - | - | - | - | - | - | - | - | - | - | - | - | - | - | - | - | - | - | - | - | - | - | - | - | - | - | - | - | - | - | - | - | - | - | - | - | - | - | - | - | - | - | - | - | - | - | - | - | - | - | - | - | - | - | - | - | - | - | - | - | - | - | - | - | - | - | - | - | - | - | - | - | - | - | - | - | - | - | - | - | - | - | - | - | - | - | - | - | - | - | - | - | - | - | - | - | - | - | - | - | - | H | x | x | x | x | H | x | x | x | x | x | x | x | x | x | x | x | x | x | x | x | x | x | x | x | x | x | x | x | x | x | x | x | x | x | x | x | x | x | x | x | x | x | x | x | x | x | x | x | x | x | x | x | x | x | x | x | x | x | x | x | x | x | x | x | x | x | x | x | x | x | x | x | x | x | x | x | x | x | x | x | x | x | x | x | x | x | x | x | x | x | x | x | x | x | x | x | x | x | x | x | x | x | x | x | x | x | x | x | H | - | - | - | - | - | - | - | - | - | - | - | - | - | - | - | - | - | - | - | - | - | - | - | - | - | - | - | - | - | - | - | - | - | - | - | - | - | - | - | - | - | - | - | - | - | - | - | - | - | - | - | - | - | - | - | - | - | - | - | - | - | - | - | - | - | - | - | - | - | - | - | - | - | - | - | - | - | - | - | - | - | - | - | - | - | - | - | - | - | - | - | - | - | - | - | - | - | - | - | - | - | - | - | - | - | - | - | - | - | - | - | - | - | - | - | - | - | - | - | - | - | - | - | - | - | - | - | - | - | - | - | - | - | - | - | - | - | - | - | - | - | - | - | - | - | - | - | - | - | - | - | - | - | - | - | - | - | - | - | - | - | - | - | - | - | - | - | - | - | - | - | - | - | - | - | - | - | - | - | - | - | - | - | - | - | - | - | - | - | - | - | - | - | - | - | - | - | - | - | - | - | - | - | - | - | - | - | - | - | - | - | - | - | - | - | - | - | - | - | - | - | - | - | - | - | - | - | - | - | - | - | - | - | - | - | - | - | - | - | - | - | - | - | - | - | - | - | - | - | - | - | - | - | - | - | - | - | - | - | - | - | - | - | - | - | - | - | - | - | - | - | - | - | - | - | - | - | - | - | - | - | - | - | - | - | - | - | - | - | - | - | - | - | - | - | - | - | - | - | - | - | - | - | - | - | - | - | - | - | - | - | - | - | - | - | - | - | - | - | - | - | - | - | - | - | - | - | - | - | - | - | - | - | - | - | - | - | - | - | - | - | - | - | - | - | - | - | - | - | - | - | - | - | - | - | - | - | - | - | - | - | - | - | - | - | - | - | - | - | - | - | - | - | - | - | - | - | - | - | - | - | - | - | - | - | - | - | - | - | - | - | - | - | - | - | - | - | - | - | - | - | - | - | - | - | - | - | - | - | - | - | - | - | - | - | - | - | - | - | - | - | - | - | - | - | - | - | - | - | - | - | - | - | - | - | - | - | - | - | - | - | - | - | - | - | - | - | - | - | - | - | - | - | - | - | - | - | - | - | - | - | - | - | - | - | - | - | - | - | - | - | - | - | - | - | - | - | - | - | - | - | - | - | - | - | - | - | - | - | - | - | - | - | - | - | - | - | - | - | - | - | - | - | - | - | - | - | - | - | - | - | - | - | - | - | - | - | - | - | - | - | - | - | - | - | - | - | - | - | - | - | - | - | - | - | - | - | - | - | - | - | - | - | - | - | - | - | - | - | - | - | - | - | - | - | - | - | - | - | - | - | - | - | - | - | - | - | - | - | - | - | - | - | - | - | - | - | - | - | - | - | - | - | - | - | - | - | - | - | - | - | - | - | - | - | - | - | - | - | - | - | - | - | - | - | - | - | - | - | - | - | - | - | - | - | - | - | - | - | - | - | - | - | - | - | - | - | - | - | - | - | - | - | - | - | - | - | - | - | - | - | - | - | - | - | - | - | - | - | - | - | - | - | - | - | - | - | - | - | - | - | - | - | - | - | - | - | - | - | - | - | - | - | - | - | - | - | - | - | - | - | - | - | - | - | - | - | - | - | - | - | - | - | - | - | - | - | - | - | - | - | - | - | - | - | - | - | - | - | - | - | - | - | - | - | - | - | - | - | - | - | - | - | - | - | - | - | - | - | - | - | - | - | - | - | - | - | - | - | - | - | - | - | - | - | - | - | - | - | - | - | - | - | - | - | - | - | - | - | - | - | - | - | - | - | - | - | - | - | - | - | - | - | - | - | - | - | - | - | - | - | - | - | - | - | - | - | - | - | - | - | - | - | - | - | - | - | - | - | - | - | - | - | - | - | - | - | - | - | - | - | - | - | - | - | - | - | - | - | - | - | - | - | - | - | - | - | - | - | - | - | - | - | - | - | - | - | - | - | - | - | - | - | - | - | - | - | - | - | - | - | - | - | - | - | - | - | - | - | - | - | - | - | - | - | - | - | - | - | - | - | - | - | - | - | - | - | - | - | - | - | - | - | - | - | - | - | - | - | - | - | - | - | - | - | - | - | - | - | - | - | - | - | - | - | - | - | - | - | - | - | - | - | - | - | - | - | - | - | - | - | - | - | - | - | - | - | - | - | - | - | - | - | - | - | - | - | - | - | - | - | - | - | - | - | - | - | - | - | - | - | - | - | - | - | - | - | - | - | - | - | - | - | - | - | - | - | - | - | - | - | - | - | - | - | - | - | - | - | - | - | - | - | - | - | - | - | - | - | - | - | - | - | - | - | - | - | - | - | - | - | - | - | - | - | - | - | - | - | - | - | - | - | - | - | - | - | - | - | - | - | - | - | - | - | - | - | - | - | - | - | - | - | - | - | - | - | - | - | - | - | - | - | - | - | - | - | - | - | - | - | - | - | - | - | - | - | - | - | - | - | - | - | - | - | - | - | - | - | - | - | - | - | - | - | - | - | - | - | - | - | - | - | - | - | - | - | - | - | - | - | - | - | - | - | - | - | - | - | - | - | - | - | - | - | - | - | - | - | - | - | - | - | - | - | - | - | - | - | - | - | - | - | - | - | - | - | - | - | - | - | - | - | - | - | - | - | - | - | - | - | - | - | - | - | - | - | - | - | - | - | - | - | - | - | - | - | - | - | - | - | - | - | - | - | - | - | - | - | - | - | - | - | - | - | - | - | - | - | - | - | - | - | - | - | - | - | - | - | - | - | - | - | - | - | - | - | - | - | - | - | - | - | - | - | - | - | - | - | - | - | - | - | - | - | - | - | - | - | - | - | - | - | - | - | - | - | - | - | - | - | - | - | - | - | - | - | - | - | - | - | - | - | - | - | - | - | - | - | - | - | - | - | - | - | - | - | - | - | - | - | - | - | - | - | - | - | - | - | - | - | - | - | - | - | - | - | - | - | - | - | - | - | - | - | - | - | - | - | - | - | - | - | - | - | - | - | - | - | - | - | - | - | - | - | - | - | - | - | - | - | - | - | - | - | - | - | - | - | - | - | - | - | - | - | - | - | - | - | - | - | - | - | - | - | - | - | - | - | - | - | - | - | - | - | - | - | - | - | - | - | - | - | - | - | - | - | - | - | - | - | - | - | - | - | - | - | - | - | - | - | - | - | - | - | - | - | - | - | - | - | - | - | - | - | - | - | - | - | - | - | - | - | - | - | - | - | - | - | - | - | - | - | - | - | - | - | - | - | - | - | - | - | - | - | - | - | - | - | - | - | - | - | - | - | - | - | - | - | - | - | - | - | - | - | - | - | - | - | - | - | - | - | - | - | - | - | - | - | - | - | - | - | - | - | - | - | - | - | - | - | - | - | - | - | - | - | - | - | - | - | - | - | - | - | - | - | - | - | - | - | - | - | - | - | - | - | - | - | - | - | - | - | - | - | - | - | - | - | - | - | - | - | - | - | - | - | - | - | - | - | - | - | - | - | - | - | - | - | - | - | - | - | - | - | - | - | - | - | - | - | - | - | - | - | - | - | - | - | - | - | - | - | - | - | - | - | - | - | - | - | - | - | - | - | - | - | - | - | - | - | - | - | - | - | - | - | - | - | - | - | - | - | - | - | - | - | - | - | - | - | - | - | - | - | - | - | - | - | - | - | - | - | - | - | - | - | - | - | - | - | - | - | - | - | - | - | - | - | - | - | - | - | - | - | - | - | - | - | - | - | - | - | - | - | - | - | - | - | - | - | - | - | - | - | - | - | - | - | - | - | - | - | - | - | - | - | - | - | - | - | - | - | - | - | - | - | - | - | - | - | - | - | - | - | - | - | - | - | - | - | - | - | - | - | - | - | - | - | - | - | - | - | - | - | - | - | - | - | - | - | - | - | - | - | - | - | - | - | - | - | - | - | - | - | - | - | - | - | - | - | - | - | - | - | - | - | - | - | - | - | - | - | - | - | - | - | - | - | - | - | - | - | - | - | - | - | - | - | - | - | - | - | - | - | - | - | - | - | - | - | - | - | - | - | - | - | - | - | - | - | - | - | - | - | - | - | - | - | - | - | - | - | - | - | - | - | - | - | - | - | - | - | - | - | - | - | - | - | - | - | - | - | - | - | - | - | - | - | - | - | - | - | - | - | - | - | - | - | - | - | - | - | - | - | - | - | - | - | - | - | - | - | - | - | - | - | - | - | - | - | - | - | - | - | - | - | - | - | - | - | - | - | - | - | - | - | - | - | - | - | - | - | - | - | - | - | - | - | - | - | - | - | - | - | - | - | - | - | - | - | - | - | - | - | - | - | - | - | - | - | - | - | - | - | - | - | - | - | - | - | - | - | - | - | - | - | - | - | - | - | - | - | - | - | - | - | - | - | - | - | - | - | - | - | - | - | - | - | - | - | - | - | - | - | - | - | - | - | - | - | - | - | - | - | - | - | - | - | - | - | - | - | - | - | - | - | - | - | - | - | - | - | - | - | - | - | - | - | - | - | - | - | - | - | - | - | - | - | - | - | - | - | - | - | - | - | - | - | - | - | - | - | - | - | - | - | - | - | - | - | - | - | - | - | - | - | - | - | - | - | - | - | - | - | - | - | - | - | - | - | - | - | - | - | - | - | - | - | - | - | - | - | - | - | - | - | - | - | - | - | - | - | - | - | - | - | - | - | - | - | - | - | - | - | - | - | - | - | - | - | - | - | - | - | - | - | - | - | - | - | - | - | - | - | - | - | - | - | - | - | - | - | - | - | - | - | - | - | - | - | - | - | - | - | - | - | - | - | - | - | - | - | - | - | - | - | - | - | - | - | - | - | - | - | - | - | - | - | - | - | - | - | - | - | - | - | - | - | - | - | - | - | - | - | - | - | - | - | - | - | - | - | - | - | - | - | - | - | - | - | - | - | - | - | - | - | - | - | - | - | - | - | - | - | - | - | - | - | - | - | - | - | - | - | - | - | - | - | - | - | - | - | - | - | - | - | - | - | - | - | - | - | - | - | - | - | - | - | - | - | - | - | - | - | - | - | - | - | - | - | - | - | - | - | - | - | - | - | - | - | - | - | - | - | - | - | - | - | - | - | - | - | - | - | - | - | - | - | - | - | - | - | - | - | - | - | - | - | - | - | - | - | - | - | - | - | - | - | - | - | - | - | - | - | - | - | - | - | - | - | - | - | - | - | - | - | - | - | - | - | - | - | - | - | - | - | - | - | - | - | - | - | - | - | - | - | - | - | - | - | - | - | - | - | - | - | - | - | - | - | - | - | - | - | - | - | - | - | - | - | - | - | - | - | - | - | - | - | - | - | - | - | - | - | - | - | - | - | - | - | - | - | - | - | - | - | - | - | - | - | - | - | - | - | - | - | - | - | - | - | - | - | - | - | - | - | - | - | - | - | - | - | - | - | - | - | - | - | - | - | - | - | - | - | - | - | - | - | - | - | - | - | - | - | - | - | - | - | - | - | - | - | - | - | - | - | - | - | - | - | - | - | - | - | - | - | - | - | - | - | - | - | - | - | - | - | - | - | - | - | - | - | - | - | - | - | - | - | - | - | - | - | - | - | - | - | - | - | - | - | - | - | - | - | - | - | - | - | - | - | - | - | - | - | - | - | - | - | - | - | - | - | - | - | - | - | - | - | - | - | - | - | - | - | - | - | - | - | - | - | - | - | - | - | - | - | - | - | - | - | - | - | - | - | - | - | - | - | - | - | - | - | - | - | - | - | - | - | - | - | - | - | - | - | - | - | - | - | - | - | - | - | - | - | - | - | - | - | - | - | - | - | - | - | - | - | - | - | - | - | - | - | - | - | - | - | - | - | - | - | - | - | - | - | - | - | - | - | - | - | - | - | - | - | - | - | - | - | - | - | - | - | - | - | - | - | - | - | - | - | - | - | - | - | - | - | - | - | - | - | - | - | - | - | - | - | - | - | - | - | - | - | - | - | - | - | - | - | - | - | - | - | - | - | - | - | - | - | - | - | - | - | - | - | - | - | - | - | - | - | - | - | - | - | - | - | - | - | - | - | - | - | - | - | - | - | - | - | - | - | - | - | - | - | - | - | - | - | - | - | - | - | - | - | - | - | - | - | - | - | - | - | - | - | - | - | - | - | - | - | - | - | - | - | - | - | - | - | - | - | - | - | - | - | - | - | - | - | - | - | - | - | - | - | - | - | - | - | - | - | - | - | - | - | - | - | - | - | - | - | - | - | - | - | - | - | - | - | - | - | - | - | - | - | - | - | - | - | - | - | - | - | - | - | - | - | - | - | - | - | - | - | - | - | - | - | - | - | - | - | - | - | - | - | - | - | - | - | - | - | - | - | - | - | - | - | - | - | - | - | - | - | - | - | - | - | - | - | - | - | - | - | - | - | - | - | - | - | - | - | - | - | - | - | - | - | - | - | - | - | - | - | - | - | - | - | - | - | - | - | - | - | - | - | - | - | - | - | - | - | - | - | - | - | - | - | - | - | - | - | - | - | - | - | - | - | - | - | - | - | - | - | - | - | - | - | - | - | - | - | - | - | - | - | - | - | - | - | - | - | - | - | - | - | - |
| BIS d=0 cl.13 sym=1 env=1 | - | - | - | - | - | - | - | - | - | - | - | - | - | - | - | - | - | - | - | - | - | - | - | - | - | - | - | - | - | - | - | - | - | - | - | - | - | - | - | - | - | - | - | - | - | - | - | - | - | - | - | - | - | - | - | - | - | - | - | - | - | - | - | - | - | - | - | - | - | - | - | - | - | - | - | - | - | - | - | - | - | - | - | - | - | - | - | - | - | - | - | - | - | - | - | - | - | - | - | - | - | - | - | H | x | x | x | x | x | x | x | x | x | x | x | x | x | x | x | x | x | x | H | x | x | x | x | x | x | x | x | x | x | x | x | x | x | x | x | x | x | x | x | x | x | x | x | x | x | x | x | x | x | x | x | x | x | x | x | x | x | x | x | x | x | x | x | x | x | x | x | x | x | x | x | x | x | x | x | x | x | x | x | x | x | x | x | x | x | x | x | x | x | x | x | x | x | x | x | x | x | x | x | x | x | x | x | x | x | x | x | x | x | x | x | x | x | x | x | x | x | x | x | x | x | x | x | x | x | x | x | x | x | x | x | x | x | x | x | x | x | x | x | x | x | x | x | x | x | x | H | x | x | x | H | x | x | x | x | x | H | x | x | x | H | H | x | x | x | x | H | x | x | x | H | x | x | x | H | - | - | - | - | - | - | - | - | - | - | - | - | - | - | - | - | - | - | - | - | - | - | - | - | - | - | - | - | - | - | - | - | - | - | - | - | - | - | - | - | - | - | - | - | - | - | - | - | - | - | - | - | - | - | - | - | - | - | - | - | - | - | - | - | - | - | - | - | - | - | - | - | - | - | - | - | - | - | - | - | - | - | - | - | - | - | - | - | - | - | - | - | - | - | - | - | - | - | - | - | - | - | - | - | - | - | - | - | - | - | - | - | - | - | - | - | - | - | - | - | - | - | - | - | - | - | - | - | - | - | - | - | - | - | - | - | - | - | - | - | - | - | - | - | - | - | - | - | - | - | - | - | - | - | - | - | - | - | - | - | - | - | - | - | - | - | - | - | - | - | - | - | - | - | - | - | - | - | - | - | - | - | - | - | - | - | - | - | - | - | - | - | - | - | - | - | - | - | - | - | - | - | - | - | - | - | - | - | - | - | - | - | - | - | - | - | - | - | - | - | - | - | - | - | - | - | - | - | - | - | - | - | - | - | - | - | - | - | - | - | - | - | - | - | - | - | - | - | - | - | - | - | - | - | - | - | - | - | - | - | - | - | - | - | - | - | - | - | - | - | - | - | - | - | - | - | - | - | - | - | - | - | - | - | - | - | - | - | - | - | - | - | - | - | - | - | - | - | - | - | - | - | - | - | - | - | - | - | - | - | - | - | - | - | - | - | - | - | - | - | - | - | - | - | - | - | - | - | - | - | - | - | - | - | - | - | - | - | - | - | - | - | - | - | - | - | - | - | - | - | - | - | - | - | - | - | - | - | - | - | - | - | - | - | - | - | - | - | - | - | - | - | - | - | - | - | - | - | - | - | - | - | - | - | - | - | - | - | - | - | - | - | - | - | - | - | - | - | - | - | - | - | - | - | - | - | - | - | - | - | - | - | - | - | - | - | - | - | - | - | - | - | - | - | - | - | - | - | - | - | - | - | - | - | - | - | - | - | - | - | - | - | - | - | - | - | - | - | - | - | - | - | - | - | - | - | - | - | - | - | - | - | - | - | - | - | - | - | - | - | - | - | - | - | - | - | - | - | - | - | - | - | - | - | - | - | - | - | - | - | - | - | - | - | - | - | - | - | - | - | - | - | - | - | - | - | - | - | - | - | - | - | - | - | - | - | - | - | - | - | - | - | - | - | - | - | - | - | - | - | - | - | - | - | - | - | - | - | - | - | - | - | - | - | - | - | - | - | - | - | - | - | - | - | - | - | - | - | - | - | - | - | - | - | - | - | - | - | - | - | - | - | - | - | - | - | - | - | - | - | - | - | - | - | - | - | - | - | - | - | - | - | - | - | - | - | - | - | - | - | - | - | - | - | - | - | - | - | - | - | - | - | - | - | - | - | - | - | - | - | - | - | - | - | - | - | - | - | - | - | - | - | - | - | - | - | - | - | - | - | - | - | - | - | - | - | - | - | - | - | - | - | - | - | - | - | - | - | - | - | - | - | - | - | - | - | - | - | - | - | - | - | - | - | - | - | - | - | - | - | - | - | - | - | - | - | - | - | - | - | - | - | - | - | - | - | - | - | - | - | - | - | - | - | - | - | - | - | - | - | - | - | - | - | - | - | - | - | - | - | - | - | - | - | - | - | - | - | - | - | - | - | - | - | - | - | - | - | - | - | - | - | - | - | - | - | - | - | - | - | - | - | - | - | - | - | - | - | - | - | - | - | - | - | - | - | - | - | - | - | - | - | - | - | - | - | - | - | - | - | - | - | - | - | - | - | - | - | - | - | - | - | - | - | - | - | - | - | - | - | - | - | - | - | - | - | - | - | - | - | - | - | - | - | - | - | - | - | - | - | - | - | - | - | - | - | - | - | - | - | - | - | - | - | - | - | - | - | - | - | - | - | - | - | - | - | - | - | - | - | - | - | - | - | - | - | - | - | - | - | - | - | - | - | - | - | - | - | - | - | - | - | - | - | - | - | - | - | - | - | - | - | - | - | - | - | - | - | - | - | - | - | - | - | - | - | - | - | - | - | - | - | - | - | - | - | - | - | - | - | - | - | - | - | - | - | - | - | - | - | - | - | - | - | - | - | - | - | - | - | - | - | - | - | - | - | - | - | - | - | - | - | - | - | - | - | - | - | - | - | - | - | - | - | - | - | - | - | - | - | - | - | - | - | - | - | - | - | - | - | - | - | - | - | - | - | - | - | - | - | - | - | - | - | - | - | - | - | - | - | - | - | - | - | - | - | - | - | - | - | - | - | - | - | - | - | - | - | - | - | - | - | - | - | - | - | - | - | - | - | - | - | - | - | - | - | - | - | - | - | - | - | - | - | - | - | - | - | - | - | - | - | - | - | - | - | - | - | - | - | - | - | - | - | - | - | - | - | - | - | - | - | - | - | - | - | - | - | - | - | - | - | - | - | - | - | - | - | - | - | - | - | - | - | - | - | - | - | - | - | - | - | - | - | - | - | - | - | - | - | - | - | - | - | - | - | - | - | - | - | - | - | - | - | - | - | - | - | - | - | - | - | - | - | - | - | - | - | - | - | - | - | - | - | - | - | - | - | - | - | - | - | - | - | - | - | - | - | - | - | - | - | - | - | - | - | - | - | - | - | - | - | - | - | - | - | - | - | - | - | - | - | - | - | - | - | - | - | - | - | - | - | - | - | - | - | - | - | - | - | - | - | - | - | - | - | - | - | - | - | - | - | - | - | - | - | - | - | - | - | - | - | - | - | - | - | - | - | - | - | - | - | - | - | - | - | - | - | - | - | - | - | - | - | - | - | - | - | - | - | - | - | - | - | - | - | - | - | - | - | - | - | - | - | - | - | - | - | - | - | - | - | - | - | - | - | - | - | - | - | - | - | - | - | - | - | - | - | - | - | - | - | - | - | - | - | - | - | - | - | - | - | - | - | - | - | - | - | - | - | - | - | - | - | - | - | - | - | - | - | - | - | - | - | - | - | - | - | - | - | - | - | - | - | - | - | - | - | - | - | - | - | - | - | - | - | - | - | - | - | - | - | - | - | - | - | - | - | - | - | - | - | - | - | - | - | - | - | - | - | - | - | - | - | - | - | - | - | - | - | - | - | - | - | - | - | - | - | - | - | - | - | - | - | - | - | - | - | - | - | - | - | - | - | - | - | - | - | - | - | - | - | - | - | - | - | - | - | - | - | - | - | - | - | - | - | - | - | - | - | - | - | - | - | - | - | - | - | - | - | - | - | - | - | - | - | - | - | - | - | - | - | - | - | - | - | - | - | - | - | - | - | - | - | - | - | - | - | - | - | - | - | - | - | - | - | - | - | - | - | - | - | - | - | - | - | - | - | - | - | - | - | - | - | - | - | - | - | - | - | - | - | - | - | - | - | - | - | - | - | - | - | - | - | - | - | - | - | - | - | - | - | - | - | - | - | - | - | - | - | - | - | - | - | - | - | - | - | - | - | - | - | - | - | - | - | - | - | - | - | - | - | - | - | - | - | - | - | - | - | - | - | - | - | - | - | - | - | - | - | - | - | - | - | - | - | - | - | - | - | - | - | - | - | - | - | - | - | - | - | - | - | - | - | - | - | - | - | - | - | - | - | - | - | - | - | - | - | - | - | - | - | - | - | - | - | - | - | - | - | - | - | - | - | - | - | - | - | - | - | - | - | - | - | - | - | - | - | - | - | - | - | - | - | - | - | - | - | - | - | - | - | - | - | - | - | - | - | - | - | - | - | - | - | - | - | - | - | - | - | - | - | - | - | - | - | - | - | - | - | - | - | - | - | - | - | - | - | - | - | - | - | - | - | - | - | - | - | - | - | - | - | - | - | - | - | - | - | - | - | - | - | - | - | - | - | - | - | - | - | - | - | - | - | - | - | - | - | - | - | - | - | - | - | - | - | - | - | - | - | - | - | - | - | - | - | - | - | - | - | - | - | - | - | - | - | - | - | - | - | - | - | - | - | - | - | - | - | - | - | - | - | - | - | - | - | - | - | - | - | - | - | - | - | - | - | - | - | - | - | - | - | - | - | - | - | - | - | - | - | - | - | - | - | - | - | - | - | - | - | - | - | - | - | - | - | - | - | - | - | - | - | - | - | - | - | - | - | - | - | - | - | - | - | - | - | - | - | - | - | - | - | - | - | - | - | - | - | - | - | - | - | - | - | - | - | - | - | - | - | - | - | - | - | - | - | - | - | - | - | - | - | - | - | - | - | - | - | - | - | - | - | - | - | - | - | - | - | - | - | - | - | - | - | - | - | - | - | - | - | - | - | - | - | - | - | - | - | - | - | - | - | - | - | - | - | - | - | - | - | - | - | - | - | - | - | - | - | - | - | - | - | - | - | - | - | - | - | - | - | - | - | - | - | - | - | - | - | - | - | - | - | - | - | - | - | - | - | - | - | - | - | - | - | - | - | - | - | - | - | - | - | - | - | - | - | - | - | - | - | - | - | - | - | - | - | - | - | - | - | - | - | - | - | - | - | - | - | - | - | - | - | - | - | - | - | - | - | - | - | - | - | - | - | - | - | - | - | - | - | - | - | - | - | - | - | - | - | - | - | - | - | - | - | - | - | - | - | - | - | - | - | - | - | - | - | - | - | - | - | - | - | - | - | - | - | - | - | - | - | - | - | - | - | - | - | - | - | - | - | - | - | - | - | - | - | - | - | - | - | - | - | - | - | - | - | - | - | - | - | - | - | - | - | - | - | - | - | - | - | - | - | - | - | - | - | - | - | - | - | - | - | - | - | - | - | - | - | - | - | - | - | - | - | - | - | - | - | - | - | - | - | - | - | - | - | - | - | - | - | - | - | - | - | - | - | - | - | - | - | - | - | - | - | - | - | - | - | - | - | - | - | - | - | - | - | - | - | - | - | - | - | - | - | - | - | - | - | - | - | - | - | - | - | - | - | - | - | - | - | - | - | - | - | - | - | - | - | - | - | - | - | - | - | - | - | - | - | - | - | - | - | - | - | - | - | - | - | - | - | - | - | - | - | - | - | - | - | - | - | - | - | - | - | - | - | - | - | - | - | - | - | - | - | - | - | - | - | - | - | - | - | - | - | - | - | - | - | - | - | - | - | - | - | - | - | - | - | - | - | - | - | - | - | - | - | - | - | - | - | - | - | - | - | - | - | - | - | - | - | - | - | - | - | - | - | - | - | - | - | - | - | - | - | - | - | - | - | - | - | - | - | - | - | - | - | - | - | - | - | - | - | - | - | - | - | - | - | - | - | - | - | - | - | - | - | - | - | - | - | - | - | - | - | - | - | - | - | - | - | - | - | - | - | - | - | - | - | - | - | - | - | - | - | - | - | - | - | - | - | - | - | - | - | - | - | - | - | - | - | - | - | - | - | - | - | - | - | - | - | - | - | - | - | - | - | - | - | - | - | - | - | - | - | - | - | - | - | - | - | - | - | - | - | - | - | - | - | - | - | - | - | - | - | - | - | - | - | - | - | - | - | - | - | - | - | - | - | - | - | - | - | - | - | - | - | - | - | - | - | - | - | - | - | - | - | - | - | - | - | - | - | - | - | - | - | - | - | - | - | - | - | - | - | - | - | - | - | - | - | - | - | - | - | - | - | - | - | - | - | - | - | - | - | - | - | - | - | - | - | - | - | - | - | - | - | - | - | - | - | - | - | - | - | - | - | - | - | - | - | - | - | - | - | - | - | - | - | - | - | - | - | - | - | - | - | - | - | - | - | - | - | - | - | - | - | - | - | - | - | - | - | - | - | - | - | - | - | - | - | - | - | - | - | - | - | - | - | - | - | - | - | - | - | - | - | - | - | - | - | - | - | - | - | - | - | - | - | - | - | - | - | - | - | - | - | - | - | - | - | - | - | - | - | - | - | - | - | - | - | - | - | - | - | - | - | - | - | - | - | - | - | - | - | - | - | - | - | - | - | - | - | - | - | - | - | - | - | - | - | - | - | - | - | - | - | - | - | - | - | - | - | - | - | - | - | - | - | - | - | - | - | - | - | - | - | - | - | - | - | - | - | - | - | - | - | - | - | - | - | - | - | - | - | - | - | - | - | - | - | - | - | - | - | - | - | - | - | - | - | - | - | - | - | - | - | - | - | - | - | - | - | - | - | - | - | - | - | - | - | - | - | - | - | - | - | - | - | - | - | - | - | - | - | - | - | - | - | - | - | - | - | - | - | - | - | - | - | - | - | - | - | - | - | - | - | - | - | - | - | - | - | - | - | - | - | - | - | - | - | - | - | - | - | - | - | - | - | - | - | - | - | - | - | - | - | - | - | - | - | - | - | - | - | - | - | - | - | - | - | - | - | - | - | - | - | - | - | - | - | - | - | - | - | - | - | - | - | - | - | - | - | - | - | - | - | - | - | - | - | - | - | - | - | - | - | - | - | - | - | - | - | - | - | - | - | - | - | - | - | - | - | - | - | - | - | - | - | - | - | - | - | - | - | - | - | - | - | - | - | - | - | - | - | - | - | - | - | - | - | - | - | - | - | - | - | - | - | - | - | - | - | - | - | - | - | - | - | - | - | - | - | - | - | - | - | - | - | - | - | - | - | - | - | - | - | - | - | - | - | - | - | - | - | - | - | - | - | - | - | - | - | - | - | - | - | - | - | - | - | - | - | - | - | - | - | - | - | - | - | - | - | - | - | - | - | - | - | - | - | - | - | - | - | - | - | - | - | - | - | - | - | - | - | - | - | - | - | - | - | - | - | - | - | - | - | - | - | - | - | - | - | - | - | - | - | - | - | - | - | - | - | - | - | - | - | - | - | - | - | - | - | - | - | - | - | - | - | - | - | - | - | - | - | - | - | - | - | - | - | - | - | - | - | - | - | - | - | - | - | - | - | - | - | - | - | - | - | - | - | - | - | - | - | - | - | - | - | - | - | - | - | - | - | - | - | - | - | - | - | - | - | - | - | - | - | - | - | - | - | - | - | - | - | - | - | - | - | - | - | - | - | - | - | - | - | - | - | - | - | - | - | - | - | - | - | - | - | - | - | - | - | - | - | - | - | - | - | - | - | - | - | - | - | - | - | - | - | - | - | - | - | - | - | - | - | - | - | - | - | - | - | - | - | - | - | - | - | - | - | - | - | - | - | - | - | - | - | - | - | - | - | - | - | - | - | - | - | - | - | - | - | - | - | - | - | - | - | - | - | - | - | - | - | - | - | - | - | - | - | - | - | - | - | - | - | - | - | - | - | - | - | - | - | - | - | - | - | - | - | - | - | - | - | - | - | - | - | - | - | - | - | - | - | - | - | - | - | - | - | - | - | - | - | - | - | - | - | - | - | - | - | - | - | - | - | - | - | - | - | - | - | - | - | - | - | - | - | - | - | - | - | - | - | - | - | - | - | - | - | - | - | - | - | - | - | - | - | - | - | - | - | - | - | - | - | - | - | - | - | - | - | - | - | - | - | - | - | - | - | - | - | - | - | - | - | - | - | - | - | - | - | - | - | - | - | - | - | - | - | - | - | - | - | - | - | - | - | - | - | - | - | - | - | - | - | - | - | - | - | - | - | - | - | - | - | - | - | - | - | - | - | - | - | - | - | - | - | - | - | - | - | - | - | - | - | - | - | - | - | - | - | - | - | - | - | - | - | - | - | - | - | - | - | - | - | - | - | - | - | - | - | - | - | - | - | - | - | - | - | - | - | - | - | - | - | - | - | - | - | - | - | - | - | - | - | - | - | - | - | - | - | - | - | - | - | - | - | - | - | - | - | - | - | - | - | - | - | - | - | - | - | - | - | - | - | - | - | - | - | - | - | - | - | - | - | - | - | - | - | - | - | - | - | - | - | - | - | - | - | - | - | - | - | - | - | - | - | - | - | - | - | - | - | - | - | - | - | - | - | - | - | - | - | - | - | - | - | - | - | - | - | - | - | - | - | - | - | - | - | - | - | - | - | - | - | - | - | - | - | - | - | - | - | - | - | - | - | - | - | - | - | - | - | - | - | - | - | - | - | - | - | - | - | - | - | - | - | - | - | - | - | - | - | - | - | - | - | - | - | - | - | - | - | - | - | - | - | - | - | - | - | - | - | - | - | - | - | - | - | - | - | - | - | - | - | - | - | - | - | - | - | - | - | - | - | - | - | - | - | - | - | - | - | - | - | - | - | - | - | - | - | - | - | - | - | - | - | - | - | - | - | - | - | - | - | - | - | - | - | - | - | - | - | - | - | - | - | - | - | - | - | - | - | - | - | - | - | - | - | - | - | - | - | - | - | - | - | - | - | - | - | - | - | - | - | - | - | - | - | - | - | - | - | - | - | - | - | - | - | - | - | - | - | - | - | - | - | - | - | - | - | - | - | - | - | - | - | - | - | - | - | - | - | - | - | - | - | - | - | - | - | - | - | - | - | - | - | - | - | - | - | - | - | - | - | - | - | - | - | - | - | - | - | - | - | - | - | - | - | - | - | - | - | - | - | - | - | - | - | - | - | - | - | - | - | - | - | - | - | - | - | - | - | - | - | - | - | - | - | - | - | - | - | - | - | - | - | - | - | - | - | - | - | - | - | - | - | - | - | - | - | - | - | - | - | - | - | - | - | - | - | - | - | - | - | - | - | - | - | - | - | - | - | - | - | - | - | - | - | - | - | - | - | - | - | - | - | - | - | - | - | - | - | - | - | - | - | - | - | - | - | - | - | - | - | - | - | - | - | - | - | - | - | - | - | - | - | - | - | - | - | - | - | - | - | - | - | - | - | - | - | - | - | - | - | - | - | - | - | - | - | - | - | - | - | - | - | - | - | - | - | - | - | - | - | - | - | - | - | - | - | - | - | - | - | - | - | - | - | - | - | - | - | - | - | - | - | - | - | - | - | - | - | - | - | - | - | - | - | - | - | - | - | - | - | - | - | - | - | - | - | - | - | - | - | - | - | - | - | - | - | - | - | - | - | - | - | - | - | - | - | - | - | - | - | - | - | - | - | - | - | - | - | - | - | - | - | - | - | - | - | - | - | - | - | - | - | - | - | - | - | - | - | - | - | - | - | - | - | - | - | - | - | - | - | - | - | - | - | - | - | - | - | - | - | - | - | - | - | - | - | - | - | - | - | - | - | - | - | - | - | - | - | - | - | - | - | - | - | - | - | - | - | - | - | - | - | - | - | - | - | - | - | - | - | - | - | - | - | - | - | - | - | - | - | - | - | - | - | - | - | - | - | - | - | - | - | - | - | - | - | - | - | - | - | - | - | - | - | - | - | - | - | - | - | - | - | - | - | - | - | - | - | - | - | - | - | - | - | - | - | - | - | - | - | - | - | - | - | - | - | - | - | - | - | - | - | - | - | - | - | - | - | - | - | - | - | - | - | - | - | - | - | - | - | - | - | - | - | - | - | - | - | - | - | - | - | - | - | - | - | - | - | - | - | - | - | - | - | - | - | - | - | - | - | - | - | - | - | - | - | - | - | - | - | - | - | - | - | - | - | - | - | - | - | - | - | - | - | - | - | - | - | - | - | - | - | - | - | - | - | - | - | - | - | - | - | - | - | - | - | - | - | - | - | - | - | - | - | - | - | - | - | - | - | - | - | - | - | - | - | - | - | - | - | - | - | - | - | - | - | - | - | - | - | - | - | - | - | - | - | - | - | - | - | - | - | - | - | - | - | - | - | - | - | - | - | - | - | - | - | - | - | - | - | - | - | - | - | - | - | - | - | - | - | - | - | - | - | - | - | - | - | - | - | - | - | - | - | - | - | - | - | - | - | - | - | - | - | - | - | - | - | - | - | - | - | - | - | - | - | - | - | - | - | - | - | - | - | - | - | - | - | - | - | - | - | - | - | - | - | - | - | - | - | - | - | - | - | - | - | - | - | - | - | - | - | - | - | - | - | - | - | - | - | - | - | - | - | - | - | - | - | - | - | - | - | - | - | - | - | - | - | - | - | - | - | - | - | - | - | - | - | - | - | - | - | - | - | - | - | - | - | - | - | - | - | - | - | - | - | - | - | - | - | - | - | - | - | - | - | - | - | - | - | - | - | - | - | - | - | - | - | - | - | - | - | - | - | - | - | - | - | - | - | - | - | - | - | - | - | - | - | - | - | - | - | - | - | - | - | - | - | - | - | - | - | - | - | - | - | - | - | - | - | - | - | - | - | - | - | - | - | - | - | - | - | - | - | - | - | - | - | - | - | - | - | - | - | - | - | - | - | - | - | - | - | - | - | - | - | - | - | - | - | - | - | - | - | - | - | - | - | - | - | - | - | - | - | - | - | - | - | - | - | - | - | - | - | - | - | - | - | - | - | - | - | - | - | - | - | - | - | - | - | - | - | - | - | - | - | - | - | - | - | - | - | - | - | - | - | - | - | - | - | - | - | - | - | - | - | - | - | - | - | - | - | - | - | - | - | - | - | - | - | - | - | - | - | - | - | - | - | - | - | - | - | - | - | - | - | - | - | - | - | - | - | - | - | - | - | - | - | - | - | - | - | - | - | - | - | - | - | - | - | - | - | - | - | - | - | - | - | - | - | - | - | - | - | - | - | - | - | - | - | - | - | - | - | - | - | - | - | - | - | - | - | - | - | - | - | - | - | - | - | - | - | - | - | - | - | - | - | - | - | - | - | - | - | - | - | - | - | - | - | - | - | - | - | - | - | - | - | - | - | - | - | - | - | - | - | - | - | - | - | - | - | - | - | - | - | - | - | - | - | - | - | - | - | - | - | - | - | - | - | - | - | - | - | - | - | - | - | - | - | - | - | - | - | - | - | - | - | - | - | - | - | - | - | - | - | - | - | - | - | - | - | - | - | - | - | - | - | - | - | - | - | - | - | - | - | - | - | - | - | - | - | - | - | - | - | - | - | - | - | - | - | - | - | - | - | - | - | - | - | - | - | - | - | - | - | - | - | - | - | - | - | - | - | - | - | - | - | - | - | - | - | - | - | - | - | - | - | - | - | - | - | - | - | - | - | - | - | - | - | - | - | - | - | - | - | - | - | - | - | - | - | - | - | - | - | - | - | - | - | - | - | - | - | - | - | - | - | - | - | - | - | - | - | - | - | - | - | - | - | - | - | - | - | - | - | - | - | - | - | - | - | - | - | - | - | - | - | - | - | - | - | - | - | - | - | - | - | - | - | - | - | - | - | - | - | - | - | - | - | - | - | - | - | - | - | - | - | - | - | - | - | - | - | - | - | - | - | - | - | - | - | - | - | - | - | - | - | - | - | - | - | - | - | - | - | - | - | - | - | - | - | - | - | - | - | - | - | - | - | - | - | - | - | - | - | - | - | - | - | - | - | - | - | - | - | - | - | - | - | - | - | - | - | - | - | - | - | - | - | - | - | - | - | - | - | - | - | - | - | - | - | - | - | - | - | - | - | - | - | - | - | - | - | - | - | - | - | - | - | - | - | - | - | - | - | - | - | - | - | - | - | - | - | - | - | - | - | - | - | - | - | - | - | - | - | - | - | - | - | - | - | - | - | - | - | - | - | - | - | - | - | - | - | - | - | - | - | - | - | - | - | - | - | - | - | - | - | - | - | - | - | - | - | - | - | - | - | - | - | - | - | - | - | - | - | - | - | - | - | - | - | - | - | - | - | - | - | - | - | - | - | - | - | - | - | - | - | - | - | - | - | - | - | - | - | - | - | - | - | - | - | - |
| BIS d=0 cl.20 sym=1 env=1 | - | - | - | - | - | - | - | - | - | - | - | - | - | - | - | - | - | - | - | - | - | - | - | - | - | - | - | - | - | - | - | - | - | - | - | - | - | - | - | - | - | - | - | - | - | - | - | - | - | - | - | - | - | - | - | - | - | - | - | - | - | - | - | - | - | - | - | - | - | - | - | - | - | - | - | - | - | - | - | - | - | - | - | - | - | - | - | - | - | - | - | - | - | - | - | - | - | - | - | - | - | - | - | - | - | - | - | - | - | - | - | - | - | - | - | - | - | - | - | - | - | - | - | - | - | - | - | - | - | - | - | - | - | - | - | - | - | - | - | - | - | - | - | - | - | - | - | - | - | - | - | - | - | - | - | - | - | - | - | - | - | - | - | - | - | - | - | - | - | - | - | - | - | - | - | - | - | - | - | - | - | - | - | - | - | - | - | - | - | - | - | - | - | - | - | - | - | - | - | - | - | - | - | - | - | - | - | - | - | - | - | - | - | - | - | - | - | - | - | - | - | - | - | - | - | - | - | - | - | - | - | - | - | - | - | - | - | - | - | - | - | - | - | - | - | - | - | - | - | - | - | - | - | - | - | - | - | - | - | - | - | - | - | - | - | - | - | - | - | - | - | - | - | - | - | - | - | - | - | - | - | - | - | - | - | - | - | - | - | - | - | - | - | - | - | - | - | - | - | - | - | - | - | - | - | - | - | - | - | - | - | - | - | - | - | - | - | - | - | - | - | - | - | - | - | - | - | - | - | - | - | - | - | - | - | - | - | - | - | - | - | - | - | - | - | - | - | - | - | - | - | - | - | - | - | - | - | - | - | - | - | - | - | - | - | - | - | - | - | - | - | - | - | - | - | - | - | - | - | - | - | - | - | - | - | - | - | - | - | - | - | - | - | - | - | - | - | - | - | - | - | - | - | - | - | - | - | - | - | - | - | - | - | - | - | - | - | - | - | - | - | - | - | - | - | - | - | - | - | - | - | - | - | - | - | - | - | - | - | - | - | - | - | - | - | - | - | - | - | - | - | - | - | - | - | - | - | - | - | - | - | - | - | - | - | - | - | - | - | - | - | - | - | - | - | - | - | - | - | - | - | - | - | - | - | - | - | - | - | - | - | - | - | - | - | - | - | - | - | - | - | - | - | - | - | - | - | - | - | - | - | - | - | - | - | - | - | - | - | - | - | - | - | - | - | - | - | - | - | - | - | - | - | - | - | - | - | - | - | - | - | - | - | - | - | - | - | - | - | - | - | - | - | - | - | - | - | - | - | - | - | - | - | - | - | - | - | - | - | - | - | - | - | - | - | - | - | - | - | - | - | - | - | - | - | - | - | - | - | - | - | - | - | - | - | - | - | - | - | - | - | - | - | - | - | - | - | - | - | - | - | - | - | - | - | - | - | - | - | - | - | - | - | - | - | - | - | - | - | - | - | - | - | - | - | - | - | - | - | - | - | - | - | - | - | - | - | - | - | - | - | - | - | - | - | - | - | - | - | - | - | - | - | - | - | - | - | - | - | - | - | - | - | - | - | - | - | - | - | - | - | - | - | - | - | - | - | - | - | - | - | - | - | - | - | - | - | - | - | - | - | - | - | - | - | - | - | - | - | - | - | - | - | - | - | - | - | - | - | - | - | - | - | - | - | - | - | - | - | - | - | - | - | - | - | - | - | - | - | - | - | - | - | - | - | - | - | - | - | - | - | - | - | - | - | - | - | - | - | - | - | - | - | - | - | - | - | - | - | - | - | - | - | - | - | - | - | - | - | - | - | - | - | - | - | - | - | - | - | - | - | - | - | - | - | - | - | - | - | - | - | - | - | - | - | - | - | - | - | - | - | - | - | - | - | - | - | - | - | - | - | - | - | - | - | - | - | - | - | - | - | - | - | - | - | - | - | - | - | - | - | - | - | - | - | - | - | - | - | - | - | - | - | - | - | - | - | - | - | - | - | - | - | - | - | - | - | - | - | - | - | - | - | - | - | - | - | - | - | - | - | - | - | - | - | - | - | - | - | - | - | - | - | - | - | - | - | - | - | - | - | - | - | - | - | - | - | - | - | - | - | - | - | - | - | - | - | - | - | - | - | - | - | - | - | - | - | - | - | - | - | - | - | - | - | - | - | - | - | - | - | - | - | - | - | - | - | - | - | - | - | - | - | - | - | - | - | - | - | - | - | - | - | - | - | - | - | - | - | - | - | - | - | - | - | - | - | - | - | - | - | - | - | - | - | - | - | - | - | - | - | - | - | - | - | - | - | - | - | - | - | - | - | - | - | - | - | - | - | - | - | - | - | - | - | - | - | - | - | - | - | - | - | - | - | - | - | - | - | - | - | - | - | - | - | - | H | H | x | x | x | x | x | x | x | x | x | x | x | x | x | x | x | x | x | x | x | x | x | x | x | x | x | x | x | x | x | x | x | x | x | x | x | H | x | x | x | x | H | x | x | H | H | - | - | - | - | - | - | - | - | - | - | - | - | - | - | - | - | - | - | - | - | - | - | - | - | - | - | - | - | - | - | - | - | - | - | - | - | - | - | - | - | - | - | - | - | - | - | - | - | - | - | - | - | - | - | - | - | - | - | - | - | - | - | - | - | - | - | - | - | - | - | - | - | - | - | - | - | - | - | - | - | - | - | - | - | - | - | - | - | - | - | - | - | - | - | - | - | - | - | - | - | - | - | - | - | - | - | - | - | - | - | - | - | - | - | - | - | - | - | - | - | - | - | - | - | - | - | - | - | - | - | - | - | - | - | - | - | - | - | - | - | - | - | - | - | - | - | - | - | - | - | - | - | - | - | - | - | - | - | - | - | - | - | - | - | - | - | - | - | - | - | - | - | - | - | - | - | - | - | - | - | - | - | - | - | - | - | - | - | - | - | - | - | - | - | - | - | - | - | - | - | - | - | - | - | - | - | - | - | - | - | - | - | - | - | - | - | - | - | - | - | - | - | - | - | - | - | - | - | - | - | - | - | - | - | - | - | - | - | - | - | - | - | - | - | - | - | - | - | - | - | - | - | - | - | - | - | - | - | - | - | - | - | - | - | - | - | - | - | - | - | - | - | - | - | - | - | - | - | - | - | - | - | - | - | - | - | - | - | - | - | - | - | - | - | - | - | - | - | - | - | - | - | - | - | - | - | - | - | - | - | - | - | - | - | - | - | - | - | - | - | - | - | - | - | - | - | - | - | - | - | - | - | - | - | - | - | - | - | - | - | - | - | - | - | - | - | - | - | - | - | - | - | - | - | - | - | - | - | - | - | - | - | - | - | - | - | - | - | - | - | - | - | - | - | - | - | - | - | - | - | - | - | - | - | - | - | - | - | - | - | - | - | - | - | - | - | - | - | - | - | - | - | - | - | - | - | - | - | - | - | - | - | - | - | - | - | - | - | - | - | - | - | - | - | - | - | - | - | - | - | - | - | - | - | - | - | - | - | - | - | - | - | - | - | - | - | - | - | - | - | - | - | - | - | - | - | - | - | - | - | - | - | - | - | - | - | - | - | - | - | - | - | - | - | - | - | - | - | - | - | - | - | - | - | - | - | - | - | - | - | - | - | - | - | - | - | - | - | - | - | - | - | - | - | - | - | - | - | - | - | - | - | - | - | - | - | - | - | - | - | - | - | - | - | - | - | - | - | - | - | - | - | - | - | - | - | - | - | - | - | - | - | - | - | - | - | - | - | - | - | - | - | - | - | - | - | - | - | - | - | - | - | - | - | - | - | - | - | - | - | - | - | - | - | - | - | - | - | - | - | - | - | - | - | - | - | - | - | - | - | - | - | - | - | - | - | - | - | - | - | - | - | - | - | - | - | - | - | - | - | - | - | - | - | - | - | - | - | - | - | - | - | - | - | - | - | - | - | - | - | - | - | - | - | - | - | - | - | - | - | - | - | - | - | - | - | - | - | - | - | - | - | - | - | - | - | - | - | - | - | - | - | - | - | - | - | - | - | - | - | - | - | - | - | - | - | - | - | - | - | - | - | - | - | - | - | - | - | - | - | - | - | - | - | - | - | - | - | - | - | - | - | - | - | - | - | - | - | - | - | - | - | - | - | - | - | - | - | - | - | - | - | - | - | - | - | - | - | - | - | - | - | - | - | - | - | - | - | - | - | - | - | - | - | - | - | - | - | - | - | - | - | - | - | - | - | - | - | - | - | - | - | - | - | - | - | - | - | - | - | - | - | - | - | - | - | - | - | - | - | - | - | - | - | - | - | - | - | - | - | - | - | - | - | - | - | - | - | - | - | - | - | - | - | - | - | - | - | - | - | - | - | - | - | - | - | - | - | - | - | - | - | - | - | - | - | - | - | - | - | - | - | - | - | - | - | - | - | - | - | - | - | - | - | - | - | - | - | - | - | - | - | - | - | - | - | - | - | - | - | - | - | - | - | - | - | - | - | - | - | - | - | - | - | - | - | - | - | - | - | - | - | - | - | - | - | - | - | - | - | - | - | - | - | - | - | - | - | - | - | - | - | - | - | - | - | - | - | - | - | - | - | - | - | - | - | - | - | - | - | - | - | - | - | - | - | - | - | - | - | - | - | - | - | - | - | - | - | - | - | - | - | - | - | - | - | - | - | - | - | - | - | - | - | - | - | - | - | - | - | - | - | - | - | - | - | - | - | - | - | - | - | - | - | - | - | - | - | - | - | - | - | - | - | - | - | - | - | - | - | - | - | - | - | - | - | - | - | - | - | - | - | - | - | - | - | - | - | - | - | - | - | - | - | - | - | - | - | - | - | - | - | - | - | - | - | - | - | - | - | - | - | - | - | - | - | - | - | - | - | - | - | - | - | - | - | - | - | - | - | - | - | - | - | - | - | - | - | - | - | - | - | - | - | - | - | - | - | - | - | - | - | - | - | - | - | - | - | - | - | - | - | - | - | - | - | - | - | - | - | - | - | - | - | - | - | - | - | - | - | - | - | - | - | - | - | - | - | - | - | - | - | - | - | - | - | - | - | - | - | - | - | - | - | - | - | - | - | - | - | - | - | - | - | - | - | - | - | - | - | - | - | - | - | - | - | - | - | - | - | - | - | - | - | - | - | - | - | - | - | - | - | - | - | - | - | - | - | - | - | - | - | - | - | - | - | - | - | - | - | - | - | - | - | - | - | - | - | - | - | - | - | - | - | - | - | - | - | - | - | - | - | - | - | - | - | - | - | - | - | - | - | - | - | - | - | - | - | - | - | - | - | - | - | - | - | - | - | - | - | - | - | - | - | - | - | - | - | - | - | - | - | - | - | - | - | - | - | - | - | - | - | - | - | - | - | - | - | - | - | - | - | - | - | - | - | - | - | - | - | - | - | - | - | - | - | - | - | - | - | - | - | - | - | - | - | - | - | - | - | - | - | - | - | - | - | - | - | - | - | - | - | - | - | - | - | - | - | - | - | - | - | - | - | - | - | - | - | - | - | - | - | - | - | - | - | - | - | - | - | - | - | - | - | - | - | - | - | - | - | - | - | - | - | - | - | - | - | - | - | - | - | - | - | - | - | - | - | - | - | - | - | - | - | - | - | - | - | - | - | - | - | - | - | - | - | - | - | - | - | - | - | - | - | - | - | - | - | - | - | - | - | - | - | - | - | - | - | - | - | - | - | - | - | - | - | - | - | - | - | - | - | - | - | - | - | - | - | - | - | - | - | - | - | - | - | - | - | - | - | - | - | - | - | - | - | - | - | - | - | - | - | - | - | - | - | - | - | - | - | - | - | - | - | - | - | - | - | - | - | - | - | - | - | - | - | - | - | - | - | - | - | - | - | - | - | - | - | - | - | - | - | - | - | - | - | - | - | - | - | - | - | - | - | - | - | - | - | - | - | - | - | - | - | - | - | - | - | - | - | - | - | - | - | - | - | - | - | - | - | - | - | - | - | - | - | - | - | - | - | - | - | - | - | - | - | - | - | - | - | - | - | - | - | - | - | - | - | - | - | - | - | - | - | - | - | - | - | - | - | - | - | - | - | - | - | - | - | - | - | - | - | - | - | - | - | - | - | - | - | - | - | - | - | - | - | - | - | - | - | - | - | - | - | - | - | - | - | - | - | - | - | - | - | - | - | - | - | - | - | - | - | - | - | - | - | - | - | - | - | - | - | - | - | - | - | - | - | - | - | - | - | - | - | - | - | - | - | - | - | - | - | - | - | - | - | - | - | - | - | - | - | - | - | - | - | - | - | - | - | - | - | - | - | - | - | - | - | - | - | - | - | - | - | - | - | - | - | - | - | - | - | - | - | - | - | - | - | - | - | - | - | - | - | - | - | - | - | - | - | - | - | - | - | - | - | - | - | - | - | - | - | - | - | - | - | - | - | - | - | - | - | - | - | - | - | - | - | - | - | - | - | - | - | - | - | - | - | - | - | - | - | - | - | - | - | - | - | - | - | - | - | - | - | - | - | - | - | - | - | - | - | - | - | - | - | - | - | - | - | - | - | - | - | - | - | - | - | - | - | - | - | - | - | - | - | - | - | - | - | - | - | - | - | - | - | - | - | - | - | - | - | - | - | - | - | - | - | - | - | - | - | - | - | - | - | - | - | - | - | - | - | - | - | - | - | - | - | - | - | - | - | - | - | - | - | - | - | - | - | - | - | - | - | - | - | - | - | - | - | - | - | - | - | - | - | - | - | - | - | - | - | - | - | - | - | - | - | - | - | - | - | - | - | - | - | - | - | - | - | - | - | - | - | - | - | - | - | - | - | - | - | - | - | - | - | - | - | - | - | - | - | - | - | - | - | - | - | - | - | - | - | - | - | - | - | - | - | - | - | - | - | - | - | - | - | - | - | - | - | - | - | - | - | - | - | - | - | - | - | - | - | - | - | - | - | - | - | - | - | - | - | - | - | - | - | - | - | - | - | - | - | - | - | - | - | - | - | - | - | - | - | - | - | - | - | - | - | - | - | - | - | - | - | - | - | - | - | - | - | - | - | - | - | - | - | - | - | - | - | - | - | - | - | - | - | - | - | - | - | - | - | - | - | - | - | - | - | - | - | - | - | - | - | - | - | - | - | - | - | - | - | - | - | - | - | - | - | - | - | - | - | - | - | - | - | - | - | - | - | - | - | - | - | - | - | - | - | - | - | - | - | - | - | - | - | - | - | - | - | - | - | - | - | - | - | - | - | - | - | - | - | - | - | - | - | - | - | - | - | - | - | - | - | - | - | - | - | - | - | - | - | - | - | - | - | - | - | - | - | - | - | - | - | - | - | - | - | - | - | - | - | - | - | - | - | - | - | - | - | - | - | - | - | - | - | - | - | - | - | - | - | - | - | - | - | - | - | - | - | - | - | - | - | - | - | - | - | - | - | - | - | - | - | - | - | - | - | - | - | - | - | - | - | - | - | - | - | - | - | - | - | - | - | - | - | - | - | - | - | - | - | - | - | - | - | - | - | - | - | - | - | - | - | - | - | - | - | - | - | - | - | - | - | - | - | - | - | - | - | - | - | - | - | - | - | - | - | - | - | - | - | - | - | - | - | - | - | - | - | - | - | - | - | - | - | - | - | - | - | - | - | - | - | - | - | - | - | - | - | - | - | - | - | - | - | - | - | - | - | - | - | - | - | - | - | - | - | - | - | - | - | - | - | - | - | - | - | - | - | - | - | - | - | - | - | - | - | - | - | - | - | - | - | - | - | - | - | - | - | - | - | - | - | - | - | - | - | - | - | - | - | - | - | - | - | - | - | - | - | - | - | - | - | - | - | - | - | - | - | - | - | - | - | - | - | - | - | - | - | - | - | - | - | - | - | - | - | - | - | - | - | - | - | - | - | - | - | - | - | - | - | - | - | - | - | - | - | - | - | - | - | - | - | - | - | - | - | - | - | - | - | - | - | - | - | - | - | - | - | - | - | - | - | - | - | - | - | - | - | - | - | - | - | - | - | - | - | - | - | - | - | - | - | - | - | - | - | - | - | - | - | - | - | - | - | - | - | - | - | - | - | - | - | - | - | - | - | - | - | - | - | - | - | - | - | - | - | - | - | - | - | - | - | - | - | - | - | - | - | - | - | - | - | - | - | - | - | - | - | - | - | - | - | - | - | - | - | - | - | - | - | - | - | - | - | - | - | - | - | - | - | - | - | - | - | - | - | - | - | - | - | - | - | - | - | - | - | - | - | - | - | - | - | - | - | - | - | - | - | - | - | - | - | - | - | - | - | - | - | - | - | - | - | - | - | - | - | - | - | - | - | - | - | - | - | - | - | - | - | - | - | - | - | - | - | - | - | - | - | - | - | - | - | - | - | - | - | - | - | - | - | - | - | - | - | - | - | - | - | - | - | - | - | - | - | - | - | - | - | - | - | - | - | - | - | - | - | - | - | - | - | - | - | - | - | - | - | - | - | - | - | - | - | - | - | - | - | - | - | - | - | - | - | - | - | - | - | - | - | - | - | - | - | - | - | - | - | - | - | - | - | - | - | - | - | - | - | - | - | - | - | - | - | - | - | - | - | - | - | - | - | - | - | - | - | - | - | - | - | - | - | - | - | - | - | - | - | - | - | - | - | - | - | - | - | - | - | - | - | - | - | - | - | - | - | - | - | - | - | - | - | - | - | - | - | - | - | - | - | - | - | - | - | - | - | - | - | - | - | - | - | - | - | - | - | - | - | - | - | - | - | - | - | - | - | - | - | - | - | - | - | - | - | - | - | - | - | - | - | - | - | - | - | - | - | - | - | - | - | - | - | - | - | - | - | - | - | - | - | - | - | - | - | - | - | - | - | - | - | - | - | - | - | - | - | - | - | - | - | - | - | - | - | - | - | - | - | - | - | - | - | - | - | - | - | - | - | - | - | - | - | - | - | - | - | - | - | - | - | - | - | - | - | - | - | - | - | - | - | - | - | - | - | - | - | - | - | - | - | - | - | - | - | - | - | - | - | - | - | - | - | - | - | - | - | - | - | - | - | - | - | - | - | - | - | - | - | - | - | - | - | - | - | - | - | - | - | - | - | - | - | - | - | - | - | - | - | - | - | - | - | - | - | - | - | - | - | - | - | - | - | - | - | - | - | - | - | - | - | - | - | - | - | - | - | - | - | - | - | - | - | - | - | - | - | - | - | - | - | - | - | - | - | - | - | - | - | - | - | - | - | - | - | - | - | - | - | - | - | - | - | - | - | - | - | - | - | - | - | - | - | - | - | - | - | - | - | - | - | - | - | - | - | - | - | - | - | - | - | - | - | - | - | - | - | - | - | - | - | - | - | - | - | - | - | - | - | - | - | - | - | - | - | - | - | - | - | - | - | - | - | - | - | - | - | - | - | - | - | - | - | - | - | - | - | - | - | - | - | - | - | - | - | - | - | - | - | - | - | - | - | - | - | - | - | - | - | - | - | - | - | - | - | - | - | - | - | - | - | - | - | - | - | - | - | - | - | - | - | - | - | - | - | - | - | - | - | - | - | - | - | - | - | - | - | - | - | - | - | - | - | - | - | - | - | - | - | - | - | - | - | - | - | - | - | - | - | - | - | - | - | - | - | - | - | - | - | - | - | - | - | - | - | - | - | - | - | - | - | - | - | - | - | - | - | - | - | - | - | - | - | - | - | - | - | - | - | - | - | - | - | - | - | - | - | - | - | - | - | - | - | - | - | - | - | - | - | - | - | - | - | - | - | - | - | - | - | - | - | - | - | - | - | - | - | - | - | - | - | - | - | - | - | - | - | - | - | - | - | - | - | - | - | - | - | - | - | - | - | - | - | - | - | - | - | - | - | - | - | - | - | - | - | - | - | - | - | - | - | - | - | - | - | - | - | - | - | - | - | - | - | - | - | - | - | - | - | - | - | - | - | - | - | - | - | - | - | - | - | - | - | - | - | - | - | - | - | - | - | - | - | - | - | - | - | - | - | - | - | - | - | - | - | - | - | - | - | - | - | - | - | - | - | - | - | - | - | - | - | - | - | - | - | - | - | - | - | - | - | - | - | - | - | - | - | - | - | - | - | - | - | - | - | - | - | - | - | - | - | - | - | - | - | - | - | - | - | - | - | - | - | - | - | - | - | - | - | - | - | - | - | - | - | - | - | - | - | - | - | - | - | - | - | - | - | - | - | - | - | - | - | - | - | - | - | - | - | - | - | - | - | - | - | - | - | - | - | - | - | - | - | - | - | - | - | - | - | - | - | - | - | - | - | - | - | - | - | - | - | - | - | - | - | - | - | - | - | - | - | - | - | - | - | - | - | - | - | - | - | - | - | - | - | - | - | - | - | - | - | - | - | - | - | - | - | - | - | - | - | - | - | - | - | - | - | - | - | - | - | - | - | - | - | - | - | - | - | - | - | - | - | - | - | - | - | - | - | - | - | - | - | - | - | - | - | - | - | - | - | - | - | - | - | - | - | - | - | - | - | - | - | - | - | - | - | - | - | - | - | - | - | - | - | - | - | - | - | - | - | - | - | - | - | - | - | - | - | - | - | - | - | - | - | - | - | - | - | - | - | - | - | - | - | - | - | - | - | - | - | - | - | - | - | - | - | - | - | - | - | - | - | - | - | - | - | - | - | - | - | - | - | - | - | - | - | - | - | - | - | - | - | - | - | - | - | - | - | - | - | - | - | - | - | - | - | - | - | - | - | - | - | - | - | - | - | - | - | - | - | - | - | - | - | - | - | - | - | - | - | - | - | - | - | - | - | - | - | - | - | - | - | - | - | - | - | - | - | - | - | - | - | - | - | - | - | - | - | - | - | - | - | - | - | - | - | - | - | - | - | - | - | - | - | - | - | - | - | - | - | - | - | - | - | - | - | - | - | - | - | - | - | - | - | - | - | - | - | - | - | - | - | - | - | - | - | - | - | - | - | - | - | - | - | - | - | - | - | - | - | - | - | - | - | - | - | - | - | - | - | - | - | - | - | - | - | - | - | - | - | - | - | - | - | - | - | - | - | - | - | - | - | - | - | - | - | - | - | - | - | - | - | - | - | - | - | - | - | - | - | - | - | - | - | - | - | - | - | - | - | - | - | - | - | - | - | - | - | - | - | - | - | - | - | - | - | - | - | - | - | - | - | - | - | - | - | - | - | - | - | - | - | - | - | - | - | - | - | - | - | - | - | - | - | - | - | - | - | - | - | - | - | - | - | - | - | - | - | - | - | - | - | - | - | - | - | - | - | - | - | - | - | - | - | - | - | - | - | - | - | - | - | - | - | - | - | - | - | - | - | - | - | - | - | - | - | - | - | - | - | - | - | - | - | - | - | - | - | - | - | - | - | - | - | - | - | - | - | - | - | - | - | - | - | - | - | - | - | - | - | - | - | - | - | - | - | - | - | - | - | - | - | - | - | - | - | - | - | - | - | - | - | - | - | - | - | - | - | - | - | - | - | - | - | - | - | - | - | - | - | - | - | - | - | - | - | - | - | - | - | - | - | - | - | - | - | - | - | - | - | - | - | - | - | - | - | - | - | - | - | - | - | - | - | - | - | - | - | - | - | - | - | - | - | - | - | - | - | - | - | - | - | - | - | - | - | - | - | - | - | - | - | - | - | - | - | - | - | - | - | - | - | - | - | - | - | - | - | - | - | - | - | - | - | - | - | - | - | - | - | - | - | - | - | - | - | - | - | - | - | - | - | - | - | - | - | - | - | - | - | - | - | - | - | - | - | - | - | - | - | - | - | - | - | - | - | - | - | - | - | - | - | - | - | - | - | - | - | - | - | - | - | - | - | - | - | - | - | - | - | - | - | - | - | - | - | - | - | - | - | - | - | - | - | - | - | - | - | - | - | - | - | - | - | - | - | - | - | - | - | - | - | - | - | - | - | - | - | - | - | - | - | - | - | - | - | - | - | - | - | - | - | - | - | - | - | - | - | - | - | - | - | - | - | - | - | - | - | - | - | - | - | - | - | - | - | - | - | - | - | - | - | - | - | - | - | - | - | - | - | - | - | - | - | - | - | - | - | - | - | - | - | - | - | - | - | - | - | - | - | - | - | - | - | - | - | - | - | - | - | - | - | - | - | - | - | - | - | - | - | - | - | - | - | - | - | - | - | - | - | - | - | - | - | - | - | - | - | - | - | - | - | - | - | - | - | - | - | - | - | - | - | - | - | - | - | - | - | - | - | - | - | - | - | - | - | - | - | - | - | - | - | - | - | - | - | - | - | - | - | - | - | - | - | - | - | - | - | - | - | - | - | - | - | - | - | - | - | - | - | - | - | - | - | - | - | - | - | - | - | - | - | - | - | - | - | - | - | - | - | - | - | - | - | - | - | - | - | - | - | - | - | - | - | - |
| BIS d=0 cl.1 sym=1 env=1 | - | - | - | - | - | - | - | - | - | - | - | - | - | - | - | - | - | - | - | - | - | - | - | - | - | - | - | - | - | - | - | - | - | - | - | - | - | - | - | - | - | - | - | - | - | - | - | - | - | - | - | - | - | - | - | - | - | - | - | - | - | - | - | - | - | - | - | - | - | - | - | - | - | - | - | - | - | - | - | - | - | - | - | - | - | - | - | - | - | - | - | - | - | - | - | - | - | - | - | - | - | - | - | - | - | - | - | - | - | - | - | - | - | - | - | - | - | - | - | - | - | - | - | - | - | - | - | - | - | - | - | - | - | - | - | - | - | - | - | - | - | - | - | - | - | - | - | - | - | - | - | - | - | - | - | - | - | - | - | - | - | - | - | - | - | - | - | - | - | - | - | - | - | - | - | - | - | - | - | - | - | - | - | - | - | - | - | - | - | - | - | - | - | - | - | - | - | - | - | - | - | - | - | - | - | - | - | - | - | - | - | - | - | - | - | - | - | - | - | - | - | - | - | - | - | - | - | - | - | - | - | - | - | - | - | - | - | - | - | - | - | - | - | - | - | - | - | - | - | - | - | - | - | - | - | - | - | - | - | - | - | - | - | - | - | - | - | - | - | - | - | - | - | - | - | - | - | - | - | - | - | - | - | - | - | - | - | - | - | - | - | - | - | - | - | - | - | - | - | - | - | - | - | - | - | - | - | - | - | - | - | - | - | - | - | - | - | - | - | - | - | - | - | - | - | - | - | - | - | - | - | - | - | - | - | - | - | - | - | - | - | - | - | - | - | - | - | - | - | - | - | - | - | - | - | - | - | - | - | - | - | - | - | - | - | - | - | - | - | - | - | - | - | - | - | - | - | - | - | - | - | - | - | - | - | - | - | - | - | - | - | - | - | - | - | - | - | - | - | - | - | - | - | - | - | - | - | - | - | - | - | - | - | - | - | - | - | - | - | - | - | - | - | - | - | - | - | - | - | - | - | - | - | - | - | - | - | - | - | - | - | - | - | - | - | - | - | - | - | - | - | - | - | - | - | - | - | - | - | - | - | - | - | - | - | - | - | - | - | - | - | - | - | - | - | - | - | - | - | - | - | - | - | - | - | - | - | - | - | - | - | - | - | - | - | - | - | - | - | - | - | - | - | - | - | - | - | - | - | - | - | - | - | - | - | - | - | - | - | - | - | - | - | - | - | - | - | - | - | - | - | - | - | - | - | - | - | - | - | - | - | - | - | - | - | - | - | - | - | - | - | - | - | - | - | - | - | - | - | - | - | - | - | - | - | - | - | - | - | - | - | - | - | - | - | - | - | - | - | - | - | - | - | - | - | - | - | - | - | - | - | - | - | - | - | - | - | - | - | - | - | - | - | - | - | - | - | - | - | - | - | - | - | - | - | - | - | - | - | - | - | - | - | - | - | - | - | - | - | - | - | - | - | - | - | - | - | - | - | - | - | - | - | - | - | - | - | - | - | - | - | - | - | - | - | - | - | - | - | - | - | - | - | - | - | - | - | - | - | - | - | - | - | - | - | - | - | - | - | - | - | - | - | - | - | - | - | - | - | - | - | - | - | - | - | - | - | - | - | - | - | - | - | - | - | - | - | - | - | - | - | - | - | - | - | - | - | - | - | - | - | - | - | - | - | - | - | - | - | - | - | - | - | - | - | - | - | - | - | - | - | - | - | - | - | - | - | - | - | - | - | - | - | - | - | - | - | - | - | - | - | - | - | - | - | - | - | - | - | - | - | - | - | - | - | - | - | - | - | - | - | - | - | - | - | - | - | - | - | - | - | - | - | - | - | - | - | - | - | - | - | - | - | - | - | - | - | - | - | - | - | - | - | - | - | - | - | - | - | - | - | - | - | - | - | - | - | - | - | - | - | - | - | - | - | - | - | - | - | - | - | - | - | - | - | - | - | - | - | - | - | - | - | - | - | - | - | - | - | - | - | - | - | - | - | - | - | - | - | - | - | - | - | - | - | - | - | - | - | - | - | - | - | - | - | - | - | - | - | - | - | - | - | - | - | - | - | - | - | - | - | - | - | - | - | - | - | - | - | - | - | - | - | - | - | - | - | - | - | - | - | - | - | - | - | - | - | - | - | - | - | - | - | - | - | - | - | - | - | - | - | - | - | - | - | - | - | - | - | - | - | - | - | - | - | - | - | - | - | - | - | - | - | - | - | - | - | - | - | - | - | - | - | - | - | - | - | - | - | - | - | - | - | - | - | - | - | - | - | - | - | - | - | - | - | - | - | - | - | - | - | - | - | - | - | - | - | - | - | - | - | - | - | - | - | - | - | - | - | - | - | - | - | - | - | - | - | - | - | - | - | - | - | - | - | - | - | - | - | - | - | - | - | - | - | - | - | - | - | - | - | - | - | - | - | - | - | - | - | - | - | - | - | - | - | - | - | - | - | - | - | - | - | - | - | - | - | - | - | - | - | - | - | - | - | - | - | - | - | - | - | - | - | - | - | - | - | - | - | - | - | - | - | - | - | - | - | - | - | - | - | - | - | - | - | - | - | - | - | - | - | - | - | - | - | - | - | - | - | - | - | - | - | - | - | - | - | - | - | - | - | - | - | - | - | - | - | - | - | - | - | - | - | - | - | - | - | - | - | - | - | - | - | - | - | - | - | - | - | - | - | - | - | - | - | - | - | - | - | - | - | - | - | - | - | - | - | - | - | - | - | - | - | - | - | - | - | - | - | - | - | - | - | - | - | - | - | - | - | - | - | - | - | - | - | - | - | - | - | - | - | - | - | - | - | - | - | - | - | - | - | - | - | - | - | - | - | - | - | - | - | - | - | - | - | - | - | - | - | - | - | - | - | - | - | - | - | - | - | - | - | - | - | - | - | - | - | - | - | - | - | - | - | - | - | - | - | - | - | - | - | - | - | - | - | - | - | - | - | - | - | - | - | - | - | - | - | - | - | - | - | - | - | - | - | - | - | - | - | - | - | - | - | - | - | - | - | - | - | - | - | - | - | - | - | - | - | - | - | - | - | - | - | - | - | - | - | - | - | - | - | - | - | - | - | - | - | - | - | - | - | - | - | - | - | - | - | - | - | - | - | - | - | - | - | - | - | - | - | - | - | - | - | - | - | - | - | - | - | - | - | - | - | - | - | - | - | - | - | - | - | - | - | - | - | - | - | - | - | - | - | - | - | - | - | - | - | - | - | - | - | - | - | - | - | - | - | - | - | - | - | - | - | - | - | - | - | - | - | - | - | - | - | - | - | - | - | - | - | - | - | - | - | - | - | - | - | - | - | - | - | - | - | - | - | - | - | - | - | - | - | - | - | - | - | - | - | - | - | - | - | - | - | - | - | - | - | - | - | - | - | - | - | - | - | - | - | - | - | - | - | - | - | - | - | - | - | - | - | - | - | - | - | - | - | - | - | - | - | - | - | - | - | - | - | - | - | - | - | - | - | - | - | - | - | - | - | - | - | - | - | - | - | - | - | - | - | - | - | - | - | - | - | - | - | - | - | - | - | - | - | - | - | - | - | - | - | - | - | - | - | - | - | - | - | - | - | - | - | - | - | - | - | - | - | - | - | - | - | - | - | - | - | - | - | - | - | - | - | - | - | - | - | - | - | - | - | - | - | - | - | - | - | - | - | - | - | - | - | - | - | - | - | - | - | - | - | - | - | - | - | - | - | - | - | - | - | - | - | - | - | - | - | - | - | - | - | - | - | - | - | - | - | - | - | - | - | - | - | - | - | - | - | - | - | - | - | - | - | - | - | - | - | - | - | - | - | - | - | - | - | - | - | - | - | - | - | - | - | - | - | - | - | - | - | - | - | - | - | - | - | - | - | - | - | - | - | - | - | - | - | - | - | - | - | - | - | - | - | - | - | - | - | - | - | - | - | - | - | - | - | - | - | - | - | - | - | - | - | - | - | - | - | - | - | - | - | - | - | - | - | - | - | - | - | - | - | - | - | - | - | - | - | - | - | - | - | - | - | - | - | - | - | - | - | - | - | - | - | - | - | - | - | - | - | - | - | - | - | - | - | - | - | - | - | - | - | - | - | - | - | - | - | - | - | - | - | - | - | - | - | - | - | - | - | - | - | - | - | - | - | - | - | - | - | - | - | - | - | - | - | - | - | - | - | - | - | - | - | - | - | - | - | - | - | - | - | - | - | - | - | - | - | - | - | - | - | - | - | - | - | - | - | - | - | - | - | - | - | - | - | - | - | - | - | - | - | - | - | - | - | - | - | - | - | - | - | - | - | - | - | - | - | - | - | - | - | - | - | - | - | - | - | - | - | - | - | - | - | - | - | - | - | - | - | - | - | - | - | - | - | - | - | - | - | - | - | - | - | - | - | - | - | - | - | - | - | - | - | - | - | - | - | - | - | - | - | - | - | - | - | - | - | - | - | - | - | - | - | - | - | - | - | - | - | - | - | - | - | - | - | - | - | - | - | - | - | - | - | - | - | - | - | - | - | - | - | - | - | - | - | - | - | - | - | - | - | - | - | - | - | - | - | - | - | - | - | - | - | - | - | - | - | - | - | - | - | - | - | - | - | - | - | - | - | - | - | - | - | - | - | - | - | - | - | - | - | - | - | - | - | - | - | - | - | - | - | - | - | - | - | - | - | - | - | - | - | - | - | - | - | - | - | - | - | - | - | - | - | - | - | - | - | - | - | - | - | - | - | - | - | - | - | - | - | - | - | - | - | - | - | - | - | - | - | - | - | - | - | - | - | - | - | - | - | - | - | - | - | - | - | - | - | - | - | - | - | - | - | - | - | - | - | - | - | - | - | - | - | - | - | - | - | - | - | - | - | - | - | - | - | - | - | - | - | - | - | - | - | - | - | - | - | - | - | - | - | - | - | - | - | - | - | - | - | - | - | - | - | - | - | - | - | - | - | - | - | - | - | - | - | - | - | - | - | - | - | - | - | - | - | - | - | - | - | - | - | - | - | - | - | - | - | - | - | - | - | - | - | - | - | - | - | - | - | - | - | - | - | - | - | - | - | - | - | - | - | - | - | - | - | - | - | - | - | - | - | - | - | - | - | - | - | - | - | - | - | - | - | - | - | - | - | - | - | - | - | - | - | - | - | - | - | - | - | - | - | - | - | - | - | - | - | - | - | - | - | - | - | - | - | - | - | - | - | - | - | - | - | - | - | - | - | - | - | - | - | - | - | - | - | - | - | - | - | - | - | - | - | - | - | - | - | - | - | - | - | - | - | - | - | - | - | - | - | - | - | - | - | - | - | - | - | - | - | - | - | - | - | - | - | - | - | - | - | - | - | - | - | - | - | - | - | - | - | - | - | - | - | - | - | - | - | - | - | - | - | - | - | - | - | - | - | - | - | - | - | - | - | - | - | - | - | - | - | - | - | - | - | - | - | - | - | - | - | - | - | - | - | - | - | - | - | - | - | - | - | - | - | - | - | - | - | - | - | - | - | - | - | - | - | - | - | - | - | - | - | - | - | - | - | - | - | - | - | - | - | - | - | - | - | - | - | - | - | - | - | - | - | - | - | - | - | - | - | - | - | - | - | - | - | - | - | - | - | - | - | - | - | - | - | - | - | - | - | - | - | - | - | - | - | - | - | - | - | - | - | - | - | - | - | - | - | - | - | - | - | - | - | - | - | - | - | - | - | - | - | - | - | - | - | - | - | - | - | - | - | - | - | - | - | - | - | - | - | - | - | - | - | - | - | - | - | - | - | - | - | - | - | - | - | - | - | - | - | - | - | - | - | - | - | - | - | - | - | - | - | - | - | - | - | - | - | - | - | - | - | - | - | - | - | - | - | - | - | - | - | - | - | - | - | - | - | - | - | - | - | - | - | - | - | - | - | - | - | - | - | - | - | - | - | - | - | - | - | - | - | - | - | - | - | - | - | - | - | - | - | - | - | - | - | - | - | - | - | - | - | - | - | - | - | - | - | - | - | - | - | - | - | - | - | - | - | - | - | - | - | - | - | - | - | - | - | - | - | - | - | - | - | - | - | - | - | - | - | - | - | - | - | - | - | - | - | - | - | - | - | - | - | - | - | - | - | - | - | - | - | - | - | - | - | - | - | - | - | - | - | - | - | - | - | - | - | - | - | - | - | - | - | - | - | - | - | - | - | - | - | - | - | - | - | - | - | - | - | - | - | - | - | - | - | - | - | - | - | - | - | - | - | - | - | - | - | - | - | - | - | - | - | - | - | - | - | - | - | - | - | - | - | - | - | - | - | - | - | - | - | - | - | - | - | - | - | - | - | - | - | - | - | - | - | - | - | - | - | - | - | - | - | - | - | - | - | - | - | - | - | - | - | - | - | - | - | - | - | - | - | - | - | - | - | - | - | - | - | - | - | - | - | - | - | - | - | - | - | - | - | - | - | - | - | - | - | - | - | - | - | - | - | - | - | - | - | - | - | - | - | - | - | - | - | - | - | - | - | - | - | - | - | - | - | - | - | - | - | - | - | - | - | - | - | - | - | - | - | - | - | - | - | - | - | - | - | - | - | - | - | - | - | - | - | - | - | - | - | - | - | - | - | - | - | - | - | - | - | - | - | - | - | - | - | - | - | - | - | - | - | - | - | - | - | - | - | - | - | - | - | - | - | - | - | - | - | - | - | - | - | - | - | - | - | - | - | - | - | - | - | - | - | - | - | - | - | - | - | - | - | - | - | - | - | - | - | - | - | - | - | - | - | - | - | - | - | - | - | - | - | - | - | - | - | - | - | - | - | - | - | - | - | - | - | - | - | - | - | - | - | - | - | - | - | - | - | - | - | - | - | - | - | - | - | - | - | - | - | - | - | - | - | - | - | - | - | - | - | - | - | - | - | - | - | - | - | - | - | - | - | - | - | - | - | - | - | - | - | - | - | - | - | - | - | - | - | - | - | - | - | - | - | - | - | - | - | - | - | - | - | - | - | - | - | - | - | - | - | - | - | - | - | - | - | - | - | - | - | - | - | - | - | - | - | - | - | - | - | - | - | - | - | - | - | - | - | - | - | - | - | - | - | - | - | - | - | - | - | - | - | - | - | - | - | - | - | - | - | - | - | - | - | - | - | - | - | - | - | - | - | - | - | - | - | - | - | - | - | - | - | - | - | - | - | - | - | - | - | - | - | - | - | - | - | - | - | - | - | - | - | - | - | - | - | - | - | - | - | - | - | - | - | - | - | - | - | - | - | - | - | - | - | - | - | - | - | - | - | - | - | - | - | - | - | - | - | - | - | - | - | - | - | - | - | - | - | - | - | - | - | - | - | - | - | - | - | - | - | - | - | - | - | - | - | - | - | - | - | - | - | - | - | - | - | - | - | - | - | - | - | - | - | - | - | - | - | - | - | - | - | - | - | - | - | - | - | - | - | - | - | - | - | - | - | - | - | - | - | - | - | - | - | - | - | - | - | - | - | - | - | - | - | - | - | - | - | - | - | - | - | - | - | - | - | - | - | - | - | - | - | - | - | - | - | - | - | - | - | - | - | - | - | - | - | - | - | - | - | - | - | - | - | - | - | - | - | - | - | - | - | - | - | - | - | - | - | - | - | - | - | - | - | - | - | - | - | - | - | - | - | - | - | - | - | - | - | - | - | - | - | - | - | - | - | - | - | - | - | - | - | - | - | - | - | - | - | - | - | - | - | - | - | - | - | - | - | - | - | - | - | - | - | - | - | - | - | - | - | - | - | - | - | - | - | - | - | - | - | - | - | - | - | - | - | - | - | - | - | - | - | - | - | - | - | - | - | - | - | - | - | - | - | - | - | - | - | - | - | - | - | - | - | - | - | - | - | - | - | - | - | - | - | - | - | - | - | - | - | - | - | - | - | - | - | - | - | - | - | - | - | - | - | - | - | - | - | - | - | - | - | - | - | - | - | - | - | - | - | - | - | - | - | - | - | - | - | - | - | - | - | - | - | - | - | - | - | - | - | - | - | - | - | - | - | - | - | - | - | - | - | - | - | - | - | - | - | - | - | - | - | - | - | - | - | - | - | - | - | - | - | - | - | - | - | - | - | - | - | - | - | - | - | - | - | - | - | - | - | - | - | - | - | - | - | - | - | - | - | - | - | - | - | - | - | - | - | - | - | - | - | - | - | - | - | - | - | - | - | - | - | - | - | - | - | - | - | - | - | - | - | - | - | - | - | - | - | - | - | - | - | - | - | - | - | - | - | - | - | - | - | - | - | - | - | - | - | - | - | - | - | - | - | - | - | - | - | - | - | - | - | - | - | - | - | - | - | - | - | - | - | - | - | - | - | - | - | - | - | - | - | - | - | - | - | - | - | - | - | - | - | - | - | - | - | - | - | - | - | - | - | - | - | - | - | - | - | - | - | - | - | - | - | - | - | - | - | - | - | - | - | - | - | - | - | - | - | - | - | - | - | - | - | - | - | - | - | - | - | - | - | - | - | - | - | - | - | - | - | - | - | - | - | - | - | - | - | - | - | - | - | - | - | - | - | - | - | - | - | - | - | - | - | - | - | - | - | - | - | - | - | - | - | - | - | - | - | - | - | - | - | - | - | - | - | - | - | - | - | - | - | - | - | - | - | - | - | - | - | - | - | - | - | - | - | - | - | - | - | - | - | - | - | - | - | - | - | - | - | - | - | - | - | - | - | - | - | - | - | - | - | - | - | - | - | - | - | - | - | - | - | - | - | - | - | - | - | - | - | - | - | - | - | - | - | - | - | - | - | - | - | - | - | - | - | - | - | - | - | - | - | - | - | - | - | - | - | - | - | - | - | - | - | - | - | - | - | - | - | - | - | - | - | - | - | - | - | - | - | - | - | - | - | - | - | - | - | - | - | - | - | - | - | - | - | - | - | - | - | - | - | - | - | - | - | - | - | - | - | - | - | - | - | - | - | - | - | - | - | - | - | - | - | - | - | - | - | - | - | - | - | - | - | - | - | - | - | - | - | - | - | - | - | - | - | - | - | - | - | - | - | - | - | - | - | - | - | - | - | - | - | - | - | - | - | - | - | - | - | - | - | - | - | - | - | - | - | - | - | - | - | - | - | - | - | - | - | - | - | - | - | - | - | - | - | - | - | - | - | - | - | - | - | - | - | - | - | - | - | - | - | - | - | - | - | - | - | - | - | - | - | - | - | - | - | - | - | - | - | - | - | - | - | - | - | - | - | - | - | - | - | - | - | - | - | - | - | - | - | - | - | - | - | - | - | - | - | - | - | - | - | - | - | - | - | - | - | - | - | - | - | - | - | - | - | - | - | - | - | - | - | - | - | - | - | - | - | - | - | - | - | - | - | - | - | - | - | - | - | - | - | - | - | - | - | - | - | - | - | - | - | - | - | - | - | - | - | - | - | - | - | - | - | - | - | - | - | - | - | - | - | - | - | - | - | - | - | - | - | - | - | - | - | - | - | - | - | - | - | - | - | - | - | - | - | - | - | - | - | - | - | - | - | - | - | - | - | - | - | - | - | - | - | - | - | - | - | - | - | - | - | - | - | - | - | - | - | - | - | - | - | - | - | - | - | - | - | - | - | - | - | - | - | - | - | - | - | - | - | - | - | - | - | - | - | - | - | - | - | - | - | - | - | - | - | - | - | - | - | - | - | - | - | - | - | - | - | - | - | - | - | - | - | - | - | - | - | - | - | - | - | - | - | - | - | - | - | - | - | - | - | - | - | - | - | - | - | - | - | - | - | - | - | - | - | - | - | - | - | - | - | - | - | - | - | - | - | - | - | - | - | - | - | - | - | - | - | - | - | - | - | - | - | - | - | - | - | - | - | - | - | - | - | - | - | - | - | - | - | - | - | - | - | - | - | - | - | - | - | - | - | - | - | - | - | - | - | - | - | - | - | - | - | - | - | - | - | - | - | - | - | - | - | - | - | - | - | - | - | - | - | - | - | - | - | - | - | - | - | - | - | - | - | - | - | - | - | - | - | - | - | - | - | - | - | - | - | - | - | - | - | - | - | - | - | - | - | - | - | - | - | - | - | - | - | - | - | - | - | - | - | - | - | - | - | - | - | - | - | - | - | - | - | - | - | - | - | - | - | - | - | - | - | - | - | - | - | - | - | - | - | - | - | - | - | - | - | - | - | - | - | - | - | - | - | - | - | - | - | - | - | - | - | - | - | - | - | - | - | - | - | - | - | - | - | - | - | - | - | - | - | - | - | - | - | - | - | - | - | - | - | - | - | - | - | - | - | - | - | - | - | - | - | - | - | - | - | - | - | - | - | - | - | - | - | - | - | - | - | - | - | - | - | - | - | - | - | - | - | - | - | - | - | - | - | - | - | - | - | - | - | - | - | - | - | - | - | - | - | - | - | - | - | - | - | - | - | - | - | - | - | - | - | - | - | - | - | - | - | - | - | - | - | - | - | - | - | - | - | - | - | - | - | - | - | - | - | - | - | - | - | - | - | - | - | - | - | - | - | - | - | - | - | - | - | - | - | - | - | - | - | - | - | - | - | - | - | - | - | - | - | - | - | - | - | - | - | - | - | - | - | - | - | - | - | - | - | - | - | - | - | - | - | - | - | - | - | - | - | - | - | - | - | - | - | - | - | - | - | - | - | - | - | - | - | - | - | - | - | - | - | - | - | - | - | - | - | - | - | - | - | - | - | - | - | - | - | - | - | - | - | - | - | - | - | - | - | - | - | - | - | - | - | - | - | - | - | - | - | - | - | - | - | - | - | - | - | - | - | - | - | - | - | - | - | - | - | - | - | - | - | - | - | - | - | - | - | - | - | - | - | - | - | - | - | - | - | - | - | - | - | - | - | - | - | - | - | - | - | - | - | - | - | - | - | - | - | - | - | - | - | - | - | - | - | - | - | - | - | - | - | - | - | - | - | - | - | - | - | - | - | - | - | - | - | - | - | - | - | - | - | - | - | - | - | - | - | - | - | - | - | - | - | - | - | - | - | - | - | - | - | - | - | - | - | - | - | - | - | - | - | - | - | - | - | - | - | - | - | - | - | - | - | - | - | - | - | - | - | - | - | - | - | - | - | - | - | - | - | - | - | - | - | - | - | - | - | - | - | - | - | - | - | - | - | - | - | - | - | - | - | - | - | - | - | - | - | - | - | - | - | - | - | - | - | - | - | - | - | - | - | - | - | - | - | - | - | - | - | - | - | - | - | - | - | - | - | - | - | - | - | - | - | - | - | - | - | - | - | - | - | - | - | - | - | - | - | - | - | - | - | - | - | - | - | - | - | - | - | - | - | - | - | - | - | - | - | - | - | - | - | - | - | - | - | - | - | - | - | - | - | - | - | - | - | - | - | - | - | - | - | - | - | - | - | - | - | - | - | - | - | - | - | - | - | - | - | - | - | - | - | - | - | - | - | - | - | - | - | - | - | - | - | - | - | - | - | - | - | - | - | - | - | - | - | - | - | - | - | - | - | - | - | - | - | - | - | - | - | - | - | - | - | - | - | - | - | - | - | - | - | - | - | - | - | - | - | - | - | - | - | - | - | - | - | - | - | - | - | - | - | - | - | - | - | - | - | - | - | - | - | - | - | - | - | - | - | - | - | - | - | - | - | - | - | - | - | - | - | - | - | - | - | - | - | - | - | - | - | - | - | - | - | - | - | - | - | - | - | - | - | - | - | - | - | - | - | - | - | - | - | - | - | - | - | - | - | - | - | - | - | - | - | - | - | - | - | - | - | - | - | - | - | - | - | - | - | - | - | - | - | - | - | - | - | - | - | - | - | - | - | - | - | - | - | - | - | - | - | - | - | - | - | - | - | - | - | - | - | - | - | - | - | - | - | - | - | - | - | - | - | - | - | - | - | - | - | - | - | - | - | - | - | - | - | - | - | - | - | - | - | - | - | - | - | - | - | - | - | - | - | - | - | - | - | - | - | - | - | - | - | - | - | - | - | - | - | - | - | - | - | - | - | - | - | - | - | - | - | - | - | - | - | - | - | - | - | - | - | - | - | - | - | - | - | - | - | - | - | - | - | - | - | - | - | - | - | - | - | - | - | - | - | - | - | - | - | - | - | - | - | - | - | - | - | - | - | - | - | - | - | - | - | - | - | - | - | - | - | - | - | - | - | - | - | - | - | - | - | - | - | - | - | - | - | - | - | - | - | - | - | - | - | - | - | - | - | - | - | - | - | - | - | - | - | - | - | - | - | - | - | - | - | - | - | - | - | - | - | - | - | - | - | - | - | - | - | - | - | - | - | - | - | - | - | - | - | - | - | - | - | - | - | - | - | - | - | - | - | - | - | - | - | - | - | - | - | - | - | - | H | x | x | H | - | - | - | - | - | - | - | - |
| BIS d=0 cl.17 sym=1 env=1 | - | - | - | - | - | - | - | - | - | - | - | - | - | - | - | - | - | - | - | - | - | - | - | - | - | - | - | - | - | - | - | - | - | - | - | - | - | - | - | - | - | - | - | - | - | - | - | - | - | - | - | - | - | - | - | - | - | - | - | - | - | - | - | - | - | - | - | - | - | - | - | - | - | - | - | - | - | - | - | - | - | - | - | - | - | - | - | - | - | - | - | - | - | - | - | - | - | - | - | - | - | - | - | - | - | - | - | - | - | - | - | - | - | - | - | - | - | - | - | - | - | - | - | - | - | - | - | - | - | - | - | - | - | - | - | - | - | - | - | - | - | - | - | - | - | - | - | - | - | - | - | - | - | - | - | - | - | - | - | - | - | - | - | - | - | - | - | - | - | - | - | - | - | - | - | - | - | - | - | - | - | - | - | - | - | - | - | - | - | - | - | - | - | - | - | - | - | - | - | - | - | - | - | - | - | - | - | - | - | - | - | - | - | - | - | - | - | - | - | - | - | - | - | - | - | - | - | - | - | - | - | - | - | - | - | - | - | - | - | - | - | - | - | - | - | - | - | - | - | - | - | - | - | - | - | - | - | - | - | - | - | - | - | - | - | - | - | - | - | - | - | - | - | - | - | - | - | - | - | - | - | - | - | - | - | - | - | - | - | - | - | - | - | - | - | - | - | - | - | - | - | - | - | - | - | - | - | - | - | - | - | - | - | - | - | - | - | - | - | - | - | - | - | - | - | - | - | - | - | - | - | - | - | - | - | - | - | - | - | - | - | - | - | - | - | - | - | - | - | - | - | - | - | - | - | - | - | - | - | - | - | - | - | - | - | - | - | - | - | - | - | - | - | - | - | - | - | - | - | - | - | - | - | - | - | - | - | - | - | - | - | - | - | - | - | - | - | - | - | - | - | - | - | - | - | - | - | - | - | - | - | - | - | - | - | - | - | - | - | - | - | - | - | - | - | - | - | - | - | - | - | - | - | - | - | - | - | - | - | - | - | - | - | - | - | - | - | - | - | - | - | - | - | - | - | - | - | - | - | - | - | - | - | - | - | - | - | - | - | - | - | - | - | - | - | - | - | - | - | - | - | - | - | - | - | - | - | - | - | - | - | - | - | - | - | - | - | - | - | - | - | - | - | - | - | - | - | - | - | - | - | - | - | - | - | - | - | - | - | - | - | - | - | - | - | - | - | - | - | - | - | - | - | - | - | - | - | - | - | - | - | - | - | - | - | - | - | - | - | - | - | - | - | - | - | - | - | - | - | - | - | - | - | - | - | - | - | - | - | - | - | - | - | - | - | - | - | - | - | - | - | - | - | - | - | - | - | - | - | - | - | - | - | - | - | - | - | - | - | - | - | - | - | - | - | - | - | - | - | - | - | - | - | - | - | - | - | - | - | - | - | - | - | - | - | - | - | - | - | - | - | - | - | - | - | - | - | - | - | - | - | - | - | - | - | - | - | - | - | - | - | - | - | - | - | - | - | - | - | - | - | - | - | - | - | - | - | - | - | - | - | - | - | - | - | - | - | - | - | - | - | - | - | - | - | - | - | - | - | - | - | - | - | - | - | - | - | - | - | - | - | - | - | - | - | - | - | - | - | - | - | - | - | - | - | - | - | - | - | - | - | - | - | - | - | - | - | - | - | - | - | - | - | - | - | - | - | - | - | - | - | - | - | - | - | - | - | - | - | - | - | - | - | - | - | - | - | - | - | - | - | - | - | - | - | - | - | - | - | - | - | - | - | - | - | - | - | - | - | - | - | - | - | - | - | - | - | - | - | - | - | - | - | - | - | - | - | - | - | - | - | - | - | - | - | - | - | - | - | - | - | - | - | - | - | - | - | - | - | - | - | - | - | - | - | - | - | - | - | - | - | - | - | - | - | - | - | - | - | - | - | - | - | - | - | - | - | - | - | - | - | - | - | - | - | - | - | - | - | - | - | - | - | - | - | - | - | - | - | - | - | - | - | - | - | - | - | - | - | - | - | - | - | - | - | - | - | - | - | - | - | - | - | - | - | - | - | - | - | - | - | - | - | - | - | - | - | - | - | - | - | - | - | - | - | - | - | - | - | - | - | - | - | - | - | - | - | - | - | - | - | - | - | - | - | - | - | - | - | - | - | - | - | - | - | - | - | - | - | - | - | - | - | - | - | - | - | - | - | - | - | - | - | - | - | - | - | - | - | - | - | - | - | - | - | - | - | - | - | - | - | - | - | - | - | - | - | - | - | - | - | - | - | - | - | - | - | - | - | - | - | - | - | - | - | - | - | - | - | - | - | - | - | - | - | - | - | - | - | - | - | - | - | - | - | - | - | - | - | - | - | - | - | - | - | - | - | - | - | - | - | - | - | - | - | - | - | - | - | - | - | - | - | - | - | - | - | - | - | - | - | - | - | - | - | - | - | - | - | - | - | - | - | - | - | - | - | - | - | - | - | - | - | - | - | - | - | - | - | - | - | - | - | - | - | - | - | - | - | - | - | - | - | - | - | - | - | - | - | - | - | - | - | - | - | - | - | - | - | - | - | - | - | - | - | - | - | - | - | - | - | - | - | - | - | - | - | - | - | - | - | - | - | - | - | - | - | - | - | - | - | - | - | - | - | - | - | - | - | - | - | - | - | - | - | - | - | - | - | - | - | - | - | - | - | - | - | - | - | - | - | - | - | - | - | - | - | - | - | - | - | - | - | - | - | - | - | - | - | - | - | - | - | - | - | - | - | - | - | - | - | - | - | - | - | - | - | - | - | - | - | - | - | - | - | - | - | - | - | - | - | - | - | - | - | - | - | - | - | - | - | - | - | - | - | - | - | - | - | - | - | - | - | - | - | - | - | - | - | - | - | - | - | - | - | - | - | - | - | - | - | - | - | - | - | - | - | - | - | - | - | - | - | - | - | - | - | - | - | - | - | - | - | - | - | - | - | - | - | - | - | - | - | - | - | - | - | - | - | - | - | - | - | - | - | - | - | - | - | - | - | - | - | - | - | - | - | - | - | - | - | - | - | - | - | - | - | - | - | - | - | - | - | - | - | - | - | - | - | - | - | - | - | - | - | - | - | - | - | - | - | - | - | - | - | - | - | - | - | - | - | - | - | - | - | - | - | - | - | - | - | - | - | - | - | - | - | - | - | - | - | - | - | - | - | - | - | - | - | - | - | - | - | - | - | - | - | - | - | - | - | - | - | - | - | - | - | - | - | - | - | - | - | - | - | - | - | - | - | - | - | - | - | - | - | - | - | - | - | - | - | - | - | - | - | - | - | - | - | - | - | - | - | - | - | - | - | - | - | - | - | - | - | - | - | - | - | - | - | - | - | - | - | - | - | - | - | - | - | - | - | - | - | - | - | - | - | - | - | - | - | - | - | - | - | - | - | - | - | - | - | - | - | - | - | - | - | - | - | - | - | - | - | - | - | - | - | - | - | - | - | - | - | - | - | - | - | - | - | - | - | - | - | - | - | - | - | - | - | - | - | - | - | - | - | - | - | - | - | - | - | - | - | - | - | - | - | - | - | - | - | - | - | - | - | - | - | - | - | - | - | - | - | - | - | - | - | - | - | - | - | - | - | - | - | - | - | - | - | - | - | - | - | - | - | - | - | - | - | - | - | - | - | - | - | - | - | - | - | - | - | - | - | - | - | - | - | - | - | - | - | - | - | - | - | - | - | - | - | - | - | - | - | H | x | x | x | x | x | x | x | x | x | x | x | x | x | x | x | x | x | x | x | x | x | x | x | x | x | x | x | x | x | x | x | x | x | x | x | x | x | x | x | x | x | x | x | x | x | x | x | x | x | x | x | x | x | x | x | x | x | x | x | x | x | x | x | x | x | H | - | - | - | - | - | - | - | - | - | - | - | - | - | - | - | - | - | - | - | - | - | - | - | - | - | - | - | - | - | - | - | - | - | - | - | - | - | - | - | - | - | - | - | - | - | - | - | - | - | - | - | - | - | - | - | - | - | - | - | - | - | - | - | - | - | - | - | - | - | - | - | - | - | - | - | - | - | - | - | - | - | - | - | - | - | - | - | - | - | - | - | - | - | - | - | - | - | - | - | - | - | - | - | - | - | - | - | - | - | - | - | - | - | - | - | - | - | - | - | - | - | - | - | - | - | - | - | - | - | - | - | - | - | - | - | - | - | - | - | - | - | - | - | - | - | - | - | - | - | - | - | - | - | - | - | - | - | - | - | - | - | - | - | - | - | - | - | - | - | - | - | - | - | - | - | - | - | - | - | - | - | - | - | - | - | - | - | - | - | - | - | - | - | - | - | - | - | - | - | - | - | - | - | - | - | - | - | - | - | - | - | - | - | - | - | - | - | - | - | - | - | - | - | - | - | - | - | - | - | - | - | - | - | - | - | - | - | - | - | - | - | - | - | - | - | - | - | - | - | - | - | - | - | - | - | - | - | - | - | - | - | - | - | - | - | - | - | - | - | - | - | - | - | - | - | - | - | - | - | - | - | - | - | - | - | - | - | - | - | - | - | - | - | - | - | - | - | - | - | - | - | - | - | - | - | - | - | - | - | - | - | - | - | - | - | - | - | - | - | - | - | - | - | - | - | - | - | - | - | - | - | - | - | - | - | - | - | - | - | - | - | - | - | - | - | - | - | - | - | - | - | - | - | - | - | - | - | - | - | - | - | - | - | - | - | - | - | - | - | - | - | - | - | - | - | - | - | - | - | - | - | - | - | - | - | - | - | - | - | - | - | - | - | - | - | - | - | - | - | - | - | - | - | - | - | - | - | - | - | - | - | - | - | - | - | - | - | - | - | - | - | - | - | - | - | - | - | - | - | - | - | - | - | - | - | - | - | - | - | - | - | - | - | - | - | - | - | - | - | - | - | - | - | - | - | - | - | - | - | - | - | - | - | - | - | - | - | - | - | - | - | - | - | - | - | - | - | - | - | - | - | - | - | - | - | - | - | - | - | - | - | - | - | - | - | - | - | - | - | - | - | - | - | - | - | - | - | - | - | - | - | - | - | - | - | - | - | - | - | - | - | - | - | - | - | - | - | - | - | - | - | - | - | - | - | - | - | - | - | - | - | - | - | - | - | - | - | - | - | - | - | - | - | - | - | - | - | - | - | - | - | - | - | - | - | - | - | - | - | - | - | - | - | - | - | - | - | - | - | - | - | - | - | - | - | - | - | - | - | - | - | - | - | - | - | - | - | - | - | - | - | - | - | - | - | - | - | - | - | - | - | - | - | - | - | - | - | - | - | - | - | - | - | - | - | - | - | - | - | - | - | - | - | - | - | - | - | - | - | - | - | - | - | - | - | - | - | - | - | - | - | - | - | - | - | - | - | - | - | - | - | - | - | - | - | - | - | - | - | - | - | - | - | - | - | - | - | - | - | - | - | - | - | - | - | - | - | - | - | - | - | - | - | - | - | - | - | - | - | - | - | - | - | - | - | - | - | - | - | - | - | - | - | - | - | - | - | - | - | - | - | - | - | - | - | - | - | - | - | - | - | - | - | - | - | - | - | - | - | - | - | - | - | - | - | - | - | - | - | - | - | - | - | - | - | - | - | - | - | - | - | - | - | - | - | - | - | - | - | - | - | - | - | - | - | - | - | - | - | - | - | - | - | - | - | - | - | - | - | - | - | - | - | - | - | - | - | - | - | - | - | - | - | - | - | - | - | - | - | - | - | - | - | - | - | - | - | - | - | - | - | - | - | - | - | - | - | - | - | - | - | - | - | - | - | - | - | - | - | - | - | - | - | - | - | - | - | - | - | - | - | - | - | - | - | - | - | - | - | - | - | - | - | - | - | - | - | - | - | - | - | - | - | - | - | - | - | - | - | - | - | - | - | - | - | - | - | - | - | - | - | - | - | - | - | - | - | - | - | - | - | - | - | - | - | - | - | - | - | - | - | - | - | - | - | - | - | - | - | - | - | - | - | - | - | - | - | - | - | - | - | - | - | - | - | - | - | - | - | - | - | - | - | - | - | - | - | - | - | - | - | - | - | - | - | - | - | - | - | - | - | - | - | - | - | - | - | - | - | - | - | - | - | - | - | - | - | - | - | - | - | - | - | - | - | - | - | - | - | - | - | - | - | - | - | - | - | - | - | - | - | - | - | - | - | - | - | - | - | - | - | - | - | - | - | - | - | - | - | - | - | - | - | - | - | - | - | - | - | - | - | - | - | - | - | - | - | - | - | - | - | - | - | - | - | - | - | - | - | - | - | - | - | - | - | - | - | - | - | - | - | - | - | - | - | - | - | - | - | - | - | - | - | - | - | - | - | - | - | - | - | - | - | - | - | - | - | - | - | - | - | - | - | - | - | - | - | - | - | - | - | - | - | - | - | - | - | - | - | - | - | - | - | - | - | - | - | - | - | - | - | - | - | - | - | - | - | - | - | - | - | - | - | - | - | - | - | - | - | - | - | - | - | - | - | - | - | - | - | - | - | - | - | - | - | - | - | - | - | - | - | - | - | - | - | - | - | - | - | - | - | - | - | - | - | - | - | - | - | - | - | - | - | - | - | - | - | - | - | - | - | - | - | - | - | - | - | - | - | - | - | - | - | - | - | - | - | - | - | - | - | - | - | - | - | - | - | - | - | - | - | - | - | - | - | - | - | - | - | - | - | - | - | - | - | - | - | - | - | - | - | - | - | - | - | - | - | - | - | - | - | - | - | - | - | - | - | - | - | - | - | - | - | - | - | - | - | - | - | - | - | - | - | - | - | - | - | - | - | - | - | - | - | - | - | - | - | - | - | - | - | - | - | - | - | - | - | - | - | - | - | - | - | - | - | - | - | - | - | - | - | - | - | - | - | - | - | - | - | - | - | - | - | - | - | - | - | - | - | - | - | - | - | - | - | - | - | - | - | - | - | - | - | - | - | - | - | - | - | - | - | - | - | - | - | - | - | - | - | - | - | - | - | - | - | - | - | - | - | - | - | - | - | - | - | - | - | - | - | - | - | - | - | - | - | - | - | - | - | - | - | - | - | - | - | - | - | - | - | - | - | - | - | - | - | - | - | - | - | - | - | - | - | - | - | - | - | - | - | - | - | - | - | - | - | - | - | - | - | - | - | - | - | - | - | - | - | - | - | - | - | - | - | - | - | - | - | - | - | - | - | - | - | - | - | - | - | - | - | - | - | - | - | - | - | - | - | - | - | - | - | - | - | - | - | - | - | - | - | - | - | - | - | - | - | - | - | - | - | - | - | - | - | - | - | - | - | - | - | - | - | - | - | - | - | - | - | - | - | - | - | - | - | - | - | - | - | - | - | - | - | - | - | - | - | - | - | - | - | - | - | - | - | - | - | - | - | - | - | - | - | - | - | - | - | - | - | - | - | - | - | - | - | - | - | - | - | - | - | - | - | - | - | - | - | - | - | - | - | - | - | - | - | - | - | - | - | - | - | - | - | - | - | - | - | - | - | - | - | - | - | - | - | - | - | - | - | - | - | - | - | - | - | - | - | - | - | - | - | - | - | - | - | - | - | - | - | - | - | - | - | - | - | - | - | - | - | - | - | - | - | - | - | - | - | - | - | - | - | - | - | - | - | - | - | - | - | - | - | - | - | - | - | - | - | - | - | - | - | - | - | - | - | - | - | - | - | - | - | - | - | - | - | - | - | - | - | - | - | - | - | - | - | - | - | - | - | - | - | - | - | - | - | - | - | - | - | - | - | - | - | - | - | - | - | - | - | - | - | - | - | - | - | - | - | - | - | - | - | - | - | - | - | - | - | - | - | - | - | - | - | - | - | - | - | - | - | - | - | - | - | - | - | - | - | - | - | - | - | - | - | - | - | - | - | - | - | - | - | - | - | - | - | - | - | - | - | - | - | - | - | - | - | - | - | - | - | - | - | - | - | - | - | - | - | - | - | - | - | - | - | - | - | - | - | - | - | - | - | - | - | - | - | - | - | - | - | - | - | - | - | - | - | - | - | - | - | - | - | - | - | - | - | - | - | - | - | - | - | - | - | - | - | - | - | - | - | - | - | - | - | - | - | - | - | - | - | - | - | - | - | - | - | - | - | - | - | - | - | - | - | - | - | - | - | - | - | - | - | - | - | - | - | - | - | - | - | - | - | - | - | - | - | - | - | - | - | - | - | - | - | - | - | - | - | - | - | - | - | - | - | - | - | - | - | - | - | - | - | - | - | - | - | - | - | - | - | - | - | - | - | - | - | - | - | - | - | - | - | - | - | - | - | - | - | - | - | - | - | - | - | - | - | - | - | - | - | - | - | - | - | - | - | - | - | - | - | - | - | - | - | - | - | - | - | - | - | - | - | - | - | - | - | - | - | - | - | - | - | - | - | - | - | - | - | - | - | - | - | - | - | - | - | - | - | - | - | - | - | - | - | - | - | - | - | - | - | - | - | - | - | - | - | - | - | - | - | - | - | - | - | - | - | - | - | - | - | - | - | - | - | - | - | - | - | - | - | - | - | - | - | - | - | - | - | - | - | - | - | - | - | - | - | - | - | - | - | - | - | - | - | - | - | - | - | - | - | - | - | - | - | - | - | - | - | - | - | - | - | - | - | - | - | - | - | - | - | - | - | - | - | - | - | - | - | - | - | - | - | - | - | - | - | - | - | - | - | - | - | - | - | - | - | - | - | - | - | - | - | - | - | - | - | - | - | - | - | - | - | - | - | - | - | - | - | - | - | - | - | - | - | - | - | - | - | - | - | - | - | - | - | - | - | - | - | - | - | - | - | - | - | - | - | - | - | - | - | - | - | - | - | - | - | - | - | - | - | - | - | - | - | - | - | - | - | - | - | - | - | - | - | - | - | - | - | - | - | - | - | - | - | - | - | - | - | - | - | - | - | - | - | - | - | - | - | - | - | - | - | - | - | - | - | - | - | - | - | - | - | - | - | - | - | - | - | - | - | - | - | - | - | - | - | - | - | - | - | - | - | - | - | - | - | - | - | - | - | - | - | - | - | - | - | - | - | - | - | - | - | - | - | - | - | - | - | - | - | - | - | - | - | - | - | - | - | - | - | - | - | - | - | - | - | - | - | - | - | - | - | - | - | - | - | - | - | - | - | - | - | - | - | - | - | - | - | - | - | - | - | - | - | - | - | - | - | - | - | - | - | - | - | - | - | - | - | - | - | - | - | - | - | - | - | - | - | - | - | - | - | - | - | - | - | - | - | - | - | - | - | - | - | - | - | - | - | - | - | - | - | - | - | - | - | - | - | - | - | - | - | - | - | - | - | - | - | - | - | - | - | - | - | - | - | - | - | - | - | - | - | - | - | - | - | - | - | - | - | - | - | - | - | - | - | - | - | - | - | - | - | - | - | - | - | - | - | - | - | - | - | - | - | - | - | - | - | - | - | - | - | - | - | - | - | - | - | - | - | - | - | - | - | - | - | - | - | - | - | - | - | - | - | - | - | - | - | - | - | - | - | - | - | - | - | - | - | - | - | - | - | - | - | - | - | - | - | - | - | - | - | - | - | - | - | - | - | - | - | - | - | - | - | - | - | - | - | - | - | - | - | - | - | - | - | - | - | - | - | - | - | - | - | - | - | - | - | - | - | - | - | - | - | - | - | - | - | - | - | - | - | - | - | - | - | - | - | - | - | - | - | - | - | - | - | - | - | - | - | - | - | - | - | - | - | - | - | - | - | - | - | - | - | - | - | - | - | - | - | - | - | - | - | - | - | - | - | - | - | - | - | - | - | - | - | - | - | - | - | - | - | - | - | - | - | - | - | - | - | - | - | - | - | - | - | - | - | - | - | - | - | - | - | - | - | - | - | - | - | - | - | - | - | - | - | - | - | - | - | - | - | - | - | - | - | - | - | - | - | - | - | - | - | - | - | - | - | - | - | - | - | - | - | - | - | - | - | - | - | - | - | - | - | - | - | - | - | - | - | - | - | - | - | - | - | - | - | - | - | - | - | - | - | - | - | - | - | - | - | - | - | - | - | - | - | - | - | - | - | - | - | - | - | - | - | - | - | - | - | - | - | - | - | - | - | - | - | - | - | - | - | - | - | - | - | - | - | - | - | - | - | - | - | - | - | - | - | - | - | - | - | - | - | - | - | - | - | - | - | - | - | - | - | - | - | - | - | - | - | - | - | - | - | - | - | - | - | - | - | - | - | - | - | - | - | - | - | - | - | - | - | - | - | - | - | - | - | - | - | - | - | - | - | - | - | - | - | - | - | - | - | - | - | - | - | - | - | - | - | - | - | - | - | - | - | - | - | - | - | - | - | - | - | - | - | - | - | - | - | - | - | - | - | - | - | - | - | - | - | - | - | - | - | - | - | - | - | - | - | - | - | - | - | - | - | - | - | - | - | - | - | - | - | - | - | - | - | - | - | - | - | - | - | - | - | - | - | - | - | - | - | - | - | - | - | - | - | - | - | - | - | - | - | - | - | - | - | - | - | - | - | - | - | - | - | - | - | - | - | - | - | - | - | - | - | - | - | - | - | - | - | - | - | - | - | - | - | - | - | - | - | - | - | - | - | - | - | - | - | - | - | - | - | - | - | - | - | - | - | - | - | - | - | - | - | - | - | - | - | - | - | - | - | - | - | - | - | - | - | - | - | - | - | - | - | - | - | - | - | - | - | - | - | - | - | - | - | - | - | - | - | - | - | - | - | - | - | - | - | - | - | - | - | - | - | - | - | - | - | - | - | - | - | - | - | - | - | - | - | - | - | - | - | - | - | - | - | - | - | - | - | - | - | - | - | - | - | - | - | - | - | - | - | - | - | - | - | - | - | - | - | - | - | - | - | - | - | - | - | - | - | - | - | - | - | - | - | - | - | - | - | - | - | - | - | - | - | - | - | - | - | - | - | - | - | - | - | - | - | - | - | - | - | - | - | - | - | - | - | - | - | - | - | - | - | - | - | - | - | - | - | - | - | - | - | - | - | - | - | - | - | - | - | - | - | - | - | - | - | - | - | - | - | - | - | - | - | - | - | - | - | - | - | - | - | - | - | - | - | - | - | - | - | - | - | - | - | - | - | - | - | - | - | - | - | - | - | - | - | - | - | - | - | - | - | - | - | - | - | - | - | - | - | - | - | - | - | - | - | - | - | - | - | - | - | - | - | - | - | - | - | - | - | - | - | - | - | - | - | - | - | - | - | - | - | - | - | - | - | - | - | - | - | - | - | - | - | - | - | - | - | - | - | - | - | - | - | - | - | - | - | - | - | - | - | - | - | - | - | - | - | - | - | - | - | - | - | - | - | - | - | - | - | - | - | - | - | - | - | - | - | - | - | - | - | - | - | - | - | - | - | - | - | - | - | - | - | - | - | - | - | - | - | - | - | - | - | - | - | - | - | - | - | - | - | - | - | - | - | - | - | - | - | - | - | - | - | - | - | - | - | - | - | - | - | - | - | - | - | - | - | - | - | - | - | - | - | - | - | - | - | - | - | - | - | - | - | - | - | - | - | - | - | - | - | - | - | - | - | - | - | - | - | - | - | - | - | - | - | - | - | - | - | - | - | - | - | - | - | - | - | - | - | - | - | - | - | - | - | - | - | - | - | - | - | - | - | - | - | - | - | - | - | - | - | - | - | - | - | - | - | - | - | - | - | - | - | - | - | - | - | - | - | - | - | - | - | - | - | - | - | - | - | - | - | - | - | - | - | - | - | - | - | - | - | - | - | - | - | - | - | - | - | - | - | - | - | - | - | - | - | - | - | - | - | - | - | - | - | - | - | - | - | - | - | - | - | - | - | - | - | - | - | - | - | - | - | - | - | - | - | - | - | - | - | - | - | - | - | - | - | - | - | - | - | - | - | - | - | - | - | - | - | - | - | - | - | - | - | - | - | - | - | - | - | - | - | - | - | - | - | - | - | - | - | - | - | - | - | - | - | - | - | - | - | - | - | - | - | - | - | - | - | - | - | - | - | - | - | - | - | - | - | - | - | - | - | - | - | - | - | - | - | - | - | - | - | - | - | - | - | - | - | - | - | - | - | - | - | - | - | - | - | - | - | - | - | - | - | - | - | - | - | - | - | - | - | - | - | - | - | - | - | - | - | - | - | - | - | - | - | - | - | - | - | - | - | - | - | - | - | - | - | - | - | - | - | - | - | - | - | - | - | - | - | - | - | - | - | - | - | - | - | - | - | - | - | - | - | - | - | - | - | - | - | - | - | - | - | - | - | - | - | - | - | - | - | - | - | - | - | - | - | - | - | - | - | - | - | - | - | - | - | - | - | - | - | - | - | - | - | - | - | - | - | - | - | - | - | - | - | - | - | - | - | - | - | - | - | - |
| BIS d=0 cl.12 sym=1 env=1 | - | - | - | - | - | - | - | - | - | - | - | - | - | - | - | - | - | - | - | - | - | - | - | - | - | - | - | - | - | - | - | - | - | - | - | - | - | - | - | - | - | - | - | - | - | - | - | - | - | - | - | - | - | - | - | - | - | - | - | - | - | - | - | - | - | - | - | - | - | - | - | - | - | - | - | - | - | - | - | - | - | - | - | - | - | - | - | - | - | - | - | - | - | - | - | - | - | - | - | - | - | - | - | - | - | - | - | - | - | - | - | - | - | - | - | - | - | - | - | - | - | - | - | - | - | - | - | - | - | - | - | - | - | - | - | - | - | - | - | - | - | - | - | - | - | - | - | - | - | - | - | - | - | - | - | - | - | - | - | - | - | - | - | - | - | - | - | - | - | - | - | - | - | - | - | - | - | - | - | - | - | - | - | - | - | - | - | - | - | - | - | - | - | - | - | - | - | - | - | - | - | - | - | - | - | - | - | - | - | - | - | - | - | - | - | - | - | - | - | - | - | - | - | - | - | - | - | - | - | - | - | - | - | - | - | - | - | - | - | - | - | - | - | - | - | - | - | - | - | - | - | - | - | - | - | - | - | - | - | - | - | - | - | - | - | - | - | - | - | - | - | - | - | - | - | - | - | - | - | - | - | - | - | - | - | - | - | - | - | - | - | - | - | - | - | - | - | - | - | - | - | - | - | - | - | - | - | - | - | - | - | - | - | - | - | - | - | - | - | - | - | - | - | - | - | - | - | - | - | - | - | - | - | - | - | - | - | - | - | - | - | - | - | - | - | - | - | - | - | - | - | - | - | - | - | - | - | - | - | - | - | - | - | - | - | - | - | - | - | - | - | - | - | - | - | - | - | - | - | - | - | - | - | - | - | - | - | - | - | - | - | - | - | - | - | - | - | - | - | - | - | - | - | - | - | - | - | - | - | - | - | - | - | - | - | - | - | - | - | - | - | - | - | - | - | - | - | - | - | - | - | - | - | - | - | - | - | - | - | - | - | - | - | - | - | - | - | - | - | - | - | - | - | - | - | - | - | - | - | - | - | - | - | - | - | - | - | - | - | - | - | - | - | - | - | - | - | - | - | - | - | - | - | - | - | - | - | - | - | - | - | - | - | - | - | - | - | - | - | - | - | - | - | - | - | - | - | - | - | - | - | - | - | - | - | - | - | - | - | - | - | - | - | - | - | - | - | - | - | - | - | - | - | - | - | - | - | - | - | - | - | - | - | - | - | - | - | - | - | - | - | - | - | - | - | - | - | - | - | - | - | - | - | - | - | - | - | - | - | - | - | - | - | - | - | - | - | - | - | - | - | - | - | - | - | - | - | - | - | - | - | - | - | - | - | - | - | - | - | - | - | - | - | - | - | - | - | - | - | - | - | - | - | - | - | - | - | - | - | - | - | - | - | - | - | - | - | - | - | - | - | - | - | - | - | - | - | - | - | - | - | - | - | - | - | - | - | - | - | - | - | - | - | - | - | - | - | - | - | - | - | - | - | - | - | - | - | - | - | - | - | - | - | - | - | - | - | - | - | - | - | - | - | - | - | - | - | - | - | - | - | - | - | - | - | - | - | - | - | - | - | - | - | - | - | - | - | - | - | - | - | - | - | - | - | - | - | - | - | - | - | - | - | - | - | - | - | - | - | - | - | - | - | - | - | - | - | - | - | - | - | - | - | - | - | - | - | - | - | - | - | - | - | - | - | - | - | - | - | - | - | - | - | - | - | - | - | - | - | - | - | - | - | - | - | - | - | - | - | - | - | - | - | - | - | - | - | - | - | - | - | - | - | - | - | - | - | - | - | - | - | - | - | - | - | - | - | - | - | - | - | - | - | - | - | - | - | - | - | - | - | - | - | - | - | - | - | - | - | - | - | - | - | - | - | - | - | - | - | - | - | - | - | - | - | - | - | - | - | - | - | - | - | - | - | - | - | - | - | - | - | - | - | - | - | - | - | - | - | - | - | - | - | - | - | - | - | - | - | - | - | - | - | - | - | - | - | - | - | - | - | - | - | - | - | - | - | - | - | - | - | - | - | - | - | - | - | - | - | - | - | - | - | - | - | - | - | - | - | - | - | - | - | - | - | - | - | - | - | - | - | - | - | - | - | - | - | - | - | - | - | - | - | - | - | - | - | - | - | - | - | - | - | - | - | - | - | - | - | - | - | - | - | - | - | - | - | - | - | - | - | - | - | - | - | - | - | - | - | - | - | - | - | - | - | - | - | - | - | - | - | - | - | - | - | - | - | - | - | - | - | - | - | - | - | - | - | - | - | - | - | - | - | - | - | - | - | - | - | - | - | - | - | - | - | - | - | - | - | - | - | - | - | - | - | - | - | - | - | - | - | - | - | - | - | - | - | - | - | - | - | - | - | - | - | - | - | - | - | - | - | - | - | - | - | - | - | - | - | - | - | - | - | - | - | - | - | - | - | - | - | - | - | - | - | - | - | - | - | - | - | - | - | - | - | - | - | - | - | - | - | - | - | - | - | - | - | - | - | - | - | - | - | - | - | - | - | - | - | - | - | - | - | - | - | - | - | - | - | - | - | - | - | - | - | - | - | - | - | - | - | - | - | - | - | - | - | - | - | - | - | - | - | - | - | - | - | - | - | - | - | - | - | - | - | - | - | - | - | - | - | - | - | - | - | - | - | - | - | - | - | - | - | - | - | - | - | - | - | - | - | - | - | - | - | - | - | - | - | - | - | - | - | - | - | - | - | - | - | - | - | - | - | - | - | - | - | - | - | - | - | - | - | - | - | - | - | - | - | - | - | - | - | - | - | - | - | - | - | - | - | - | - | - | - | - | - | - | - | - | - | - | - | - | - | - | - | - | - | - | - | - | - | - | - | - | - | - | - | - | - | - | - | - | - | - | - | - | - | - | - | - | - | - | - | - | - | - | - | - | - | - | - | - | - | - | - | - | - | - | - | - | - | - | - | - | - | - | - | - | - | - | - | - | - | - | - | - | - | - | - | - | - | - | - | - | - | - | - | - | - | - | - | - | - | - | - | - | - | - | - | - | - | - | - | - | - | - | - | - | - | - | - | - | - | - | - | - | - | - | - | - | - | - | - | - | - | - | - | - | - | - | - | - | - | - | - | - | - | - | - | - | - | - | - | - | - | - | - | - | - | - | - | - | - | - | - | - | - | - | - | - | - | - | - | - | - | - | - | - | - | - | - | - | - | - | - | - | - | - | - | - | - | - | - | - | - | - | - | - | - | - | - | - | - | - | - | - | - | - | - | - | - | - | - | - | - | - | - | - | - | - | - | - | - | - | - | - | - | - | - | - | - | - | - | - | - | - | - | - | - | - | - | - | - | - | - | - | - | - | - | - | - | - | - | - | - | - | - | - | - | - | - | - | - | - | - | - | - | - | - | - | - | - | - | - | - | - | - | - | - | - | - | - | - | - | - | - | - | - | - | - | - | - | - | - | - | - | - | - | - | - | - | - | - | - | - | - | - | - | - | - | - | - | - | - | - | - | - | - | - | - | - | - | - | - | - | - | - | - | - | - | - | - | - | - | - | - | - | - | - | - | - | - | - | - | - | - | - | - | - | - | - | - | - | - | - | - | - | - | - | - | - | - | - | - | - | - | - | - | - | - | - | - | - | - | - | - | - | - | - | - | - | - | - | - | - | - | - | - | - | - | - | - | - | - | - | - | - | - | - | - | - | - | - | - | - | - | - | - | - | - | - | - | - | - | - | - | - | - | - | - | - | - | - | - | - | - | - | - | - | - | - | - | - | - | - | - | - | - | - | - | - | - | - | - | - | - | - | - | - | - | - | - | - | - | - | - | - | - | - | - | - | - | - | - | - | - | - | - | - | - | - | - | - | - | - | - | - | - | - | - | - | - | - | - | - | - | - | - | - | - | - | - | - | - | - | - | - | - | - | - | - | - | - | - | - | - | - | - | - | - | - | - | - | - | - | - | - | - | - | - | - | - | - | - | - | - | - | - | - | - | - | - | - | - | - | - | - | - | - | - | - | - | - | - | - | - | - | - | - | - | - | - | - | - | - | - | - | - | - | - | - | - | - | - | - | - | - | - | - | - | - | - | - | - | - | - | - | - | - | - | - | - | - | - | - | - | - | - | - | - | - | - | - | - | - | - | - | - | - | - | - | - | - | - | - | - | - | - | - | - | - | - | - | - | - | - | - | - | - | - | - | - | - | - | - | - | - | - | - | - | - | - | - | - | - | - | - | - | - | - | - | - | - | - | - | - | - | - | - | - | - | - | - | - | - | - | - | - | - | - | - | - | - | - | - | - | - | - | - | - | - | - | - | - | - | - | - | - | - | - | - | - | - | - | - | - | - | - | - | - | - | - | - | - | - | - | - | - | - | - | - | - | - | - | - | - | - | - | - | - | - | - | - | - | - | - | - | - | - | - | - | - | - | - | - | - | - | - | - | - | - | - | - | - | - | - | - | - | - | - | - | - | - | - | - | - | - | - | - | - | - | - | - | - | - | - | - | - | - | - | - | - | - | - | - | - | - | - | - | - | - | - | - | - | - | - | - | - | - | - | - | - | - | - | - | - | - | - | - | - | - | - | - | - | - | - | - | - | - | - | - | - | - | - | - | - | - | - | - | - | - | - | - | - | - | - | - | - | - | - | - | - | - | - | - | - | - | - | - | - | - | - | - | - | - | - | - | - | - | - | - | - | - | - | - | - | - | - | - | - | - | - | - | - | - | - | - | - | - | - | - | - | - | - | - | - | - | - | - | - | - | - | - | - | - | - | - | - | - | - | - | - | - | - | - | - | - | - | - | - | - | - | - | - | - | - | - | - | - | - | - | - | - | - | - | - | - | - | - | - | - | - | - | - | - | - | - | - | - | - | - | - | - | - | - | - | - | - | - | - | - | - | - | - | - | - | - | - | - | - | - | - | - | - | - | - | - | - | - | - | - | - | - | - | - | - | - | - | - | - | - | - | - | - | - | - | - | - | - | - | - | - | - | - | - | - | - | - | - | - | - | - | - | - | - | - | - | - | - | - | - | - | - | - | - | - | - | - | - | - | - | - | - | - | - | - | - | - | - | - | - | - | - | - | - | - | - | - | - | - | - | - | - | - | - | - | - | - | - | - | - | - | - | - | - | - | - | - | - | - | - | - | - | - | - | - | - | - | - | - | - | - | - | - | - | - | - | - | - | - | - | - | - | - | - | - | - | - | - | - | - | - | - | - | - | - | - | - | - | - | - | - | - | - | - | - | - | - | - | - | - | - | - | - | - | - | - | - | - | - | - | - | - | - | - | - | - | - | - | - | - | - | - | - | - | - | - | - | - | - | - | - | - | - | - | - | - | - | - | - | - | - | - | - | - | - | - | - | - | - | - | - | - | - | - | - | - | - | - | - | - | - | - | - | - | - | - | - | - | - | - | - | - | - | - | - | - | - | - | - | - | - | - | - | - | - | - | - | - | - | - | - | - | - | - | - | - | - | - | - | - | - | - | - | - | - | - | - | - | - | - | - | - | - | - | - | - | - | - | - | - | - | - | - | - | - | - | - | - | - | - | - | - | - | - | - | - | - | - | - | - | - | - | - | - | - | - | - | - | - | - | - | - | - | - | - | - | - | - | - | - | - | - | - | - | - | - | - | - | - | - | - | - | - | - | - | - | - | - | - | - | - | - | - | - | - | - | - | - | - | - | - | - | - | - | - | - | - | - | - | - | - | - | - | - | - | - | - | - | - | - | - | - | - | - | - | - | - | - | - | - | - | - | - | - | - | - | - | - | - | - | - | - | - | - | - | - | - | - | - | - | - | - | - | - | - | - | - | - | - | - | - | - | - | - | - | - | - | - | - | - | - | - | - | - | - | - | - | - | - | - | - | - | - | - | - | - | - | - | - | - | - | - | - | - | - | - | - | - | - | - | - | - | - | - | - | - | - | - | - | - | - | - | - | - | - | - | - | - | - | - | - | - | - | - | - | - | - | - | - | - | - | - | - | - | - | - | - | - | - | - | - | - | - | - | - | - | - | - | - | - | - | - | - | - | - | - | - | - | - | - | - | - | - | - | - | - | - | - | - | - | - | - | - | - | - | - | - | - | - | - | - | - | - | - | - | - | - | - | - | - | - | - | - | - | - | - | - | - | - | - | - | - | - | - | - | - | - | - | - | - | - | - | - | - | - | - | - | - | - | - | - | - | - | - | - | - | - | - | - | - | - | - | - | - | - | - | - | - | - | - | - | - | - | - | - | - | - | - | - | - | - | - | - | - | - | - | - | - | - | - | - | - | - | - | - | - | - | - | - | - | - | - | - | - | - | - | - | - | - | - | - | - | - | - | - | - | - | - | - | - | - | - | - | - | - | - | - | - | - | - | - | - | - | - | - | - | - | - | - | - | - | - | - | - | - | - | - | - | - | - | - | - | - | - | - | - | - | - | - | - | - | - | - | - | - | - | - | - | - | - | - | - | - | - | - | - | - | - | - | - | - | - | - | - | - | - | - | - | - | - | - | - | - | - | - | - | - | - | - | - | - | - | - | - | - | - | - | - | - | - | - | - | - | - | - | - | - | - | - | - | - | - | - | - | - | - | - | - | - | - | - | - | - | - | - | - | - | - | - | - | - | - | - | - | - | - | - | - | - | - | - | - | - | - | - | - | - | - | - | - | - | - | - | - | - | - | - | - | - | - | - | - | - | - | - | - | - | - | - | - | - | - | E | H | x | x | x | x | x | x | x | x | x | x | x | x | x | x | x | x | x | x | x | x | x | x | x | x | x | x | x | x | x | x | x | x | x | x | x | x | x | x | x | x | x | x | x | x | x | x | x | x | x | x | x | x | x | x | x | x | x | x | x | x | x | x | x | x | x | x | x | x | x | x | x | x | x | x | x | x | x | x | x | x | x | x | x | x | x | x | x | x | x | x | x | x | x | x | x | x | x | x | x | x | x | x | x | x | x | x | x | x | x | x | x | x | x | x | x | x | x | x | x | x | x | x | x | x | x | x | x | x | x | x | x | x | x | x | x | x | x | x | x | x | x | x | x | x | x | x | x | x | x | x | x | x | x | x | x | x | x | x | x | x | x | x | x | x | x | x | x | x | x | x | x | x | x | x | x | x | x | x | x | x | x | x | x | x | x | x | x | x | x | x | x | x | x | x | x | x | x | x | x | x | x | x | x | x | x | x | x | x | x | x | x | x | x | x | x | x | x | x | x | x | x | x | x | x | x | x | x | x | x | x | x | x | x | x | x | x | x | x | x | x | x | x | x | x | x | x | x | x | x | x | x | x | x | x | x | x | x | x | x | x | x | x | x | x | x | x | x | x | x | x | x | x | x | x | x | x | x | x | x | x | x | x | x | x | x | x | x | x | x | x | x | x | x | x | x | x | x | x | x | x | x | x | x | x | x | x | x | x | x | x | x | x | x | x | x | x | x | x | x | x | x | x | x | x | x | x | x | x | x | x | x | x | x | x | x | x | x | x | x | x | x | x | x | x | x | x | x | x | x | x | x | x | x | x | x | x | x | x | x | x | x | x | x | x | x | x | x | x | x | x | x | x | x | x | x | x | x | x | x | x | x | x | x | x | x | x | x | x | x | x | x | x | x | x | x | x | x | x | x | x | x | x | x | x | x | x | x | x | x | x | x | x | x | x | x | x | x | x | x | x | x | x | x | x | x | x | x | x | x | x | x | x | x | x | x | x | x | x | x | x | x | x | x | x | x | x | x | x | x | x | x | x | x | x | x | x | x | x | x | x | x | x | x | x | x | x | x | x | x | x | x | x | x | x | x | x | x | x | x | x | x | x | x | x | x | x | x | x | x | x | x | x | x | x | x | x | x | x | x | x | x | x | x | x | x | x | x | x | x | x | x | x | x | x | x | x | x | x | x | x | x | x | x | x | x | x | x | x | x | x | x | x | x | x | x | x | x | x | x | x | x | x | x | x | x | x | x | x | x | x | x | x | x | x | x | x | x | x | x | x | x | x | x | x | x | x | x | x | x | x | x | x | x | x | x | x | x | x | x | x | x | x | x | x | x | x | x | x | x | x | x | x | x | x | x | x | x | x | x | x | x | x | x | x | x | x | x | x | x | x | x | x | x | x | x | x | x | x | x | x | x | x | x | x | x | x | x | x | x | x | x | x | x | x | x | x | x | x | x | x | x | x | x | x | x | x | x | x | x | x | x | x | x | x | x | x | x | x | x | x | x | x | x | x | x | x | x | x | x | x | x | x | x | x | x | x | x | x | x | x | x | x | x | x | x | x | x | x | x | x | x | x | x | x | x | x | x | x | x | x | x | x | x | x | x | x | x | x | x | x | x | x | x | x | x | x | x | x | x | x | x | x | x | x | x | x | x | x | x | x | x | x | x | x | x | x | x | x | x | x | x | x | x | x | x | x | x | x | x | x | x | x | x | x | x | x | x | x | x | x | x | x | x | x | x | x | x | x | x | x | x | x | x | x | x | x | x | x | x | x | x | x | x | x | x | x | x | x | x | x | x | x | x | x | x | x | x | x | x | x | x | x | x | x | x | x | x | x | x | x | x | x | x | x | x | x | x | x | x | x | x | x | x | x | x | x | x | x | x | x | x | x | x | x | x | x | x | x | x | x | x | x | x | x | x | x | x | x | x | x | x | x | x | x | x | x | x | x | x | x | x | x | x | x | x | x | x | x | x | x | x | x | x | x | x | x | x | x | x | x | x | x | x | x | x | x | x | x | x | x | x | x | x | x | x | x | x | x | x | x | x | x | x | x | x | x | x | x | x | x | x | x | x | x | x | x | x | x | x | x | x | x | x | x | x | x | x | x | x | x | x | x | x | x | x | x | x | x | x | x | x | x | x | x | x | x | x | x | x | x | x | x | x | x | x | x | x | x | x | x | x | x | x | x | x | x | x | x | x | x | x | x | x | x | x | x | x | x | x | x | x | x | x | x | x | x | x | x | x | x | x | x | x | x | x | x | x | x | x | x | x | x | x | x | x | x | x | x | x | x | x | x | x | x | x | x | x | x | x | x | x | x | x | x | x | x | x | x | x | x | x | x | x | x | x | x | x | x | x | x | x | x | x | x | x | x | x | x | x | x | x | x | x | x | x | x | x | x | x | x | x | x | x | x | x | x | x | x | x | x | x | x | x | x | x | x | x | x | x | x | x | x | x | x | x | x | x | x | x | x | x | x | x | x | x | x | x | x | x | x | x | x | x | x | x | x | x | x | x | x | x | x | x | x | x | x | x | x | x | x | x | x | x | x | x | x | x | x | x | x | x | x | x | x | x | x | x | x | x | x | x | x | x | x | x | x | x | x | x | x | x | x | x | x | x | x | x | x | x | x | x | x | x | x | x | x | x | x | x | x | x | x | x | x | x | x | x | x | x | x | x | x | x | x | x | x | x | x | x | x | x | x | x | x | x | x | x | x | x | x | x | x | x | x | x | x | x | x | x | x | x | x | x | x | x | x | x | x | x | x | x | x | x | x | x | x | x | x | x | x | x | x | x | x | x | x | x | x | x | x | x | x | x | x | x | x | x | x | x | x | x | x | x | x | x | x | x | x | x | x | x | x | x | x | x | x | x | x | x | x | x | x | x | x | x | x | x | x | x | x | x | x | x | x | x | x | x | x | x | x | x | x | x | x | x | x | x | x | x | x | x | x | x | x | x | x | x | x | x | x | x | x | x | x | x | x | x | x | x | x | x | x | x | x | x | x | x | x | x | x | x | x | x | x | x | x | x | x | x | x | x | x | x | x | x | x | x | x | x | x | x | x | x | x | x | x | x | x | x | x | x | x | x | x | x | x | x | x | x | x | x | x | x | x | x | x | x | x | x | x | x | x | x | x | x | x | x | x | x | x | x | x | x | x | x | x | x | x | x | x | x | x | x | x | x | x | x | x | x | x | x | x | x | x | x | x | x | x | x | x | x | x | x | x | x | x | x | x | x | x | x | x | x | x | x | x | x | x | x | x | x | x | x | x | x | x | x | x | x | x | x | x | x | x | x | x | x | x | x | x | x | x | x | x | x | x | x | x | x | x | x | x | x | x | x | x | x | x | x | x | x | x | x | x | x | x | x | x | x | x | x | x | x | x | x | x | x | x | x | x | x | x | x | x | x | x | x | x | x | x | x | x | x | x | x | x | x | x | x | x | x | x | x | x | x | x | x | x | x | x | x | x | x | x | x | x | x | x | x | x | x | x | x | x | x | x | x | x | x | x | x | x | x | x | x | x | x | x | x | x | x | x | x | x | x | x | x | x | x | x | x | x | x | x | x | x | x | x | x | x | x | x | x | x | x | x | x | x | x | x | x | x | x | x | x | x | x | x | x | x | x | x | x | x | x | x | x | x | x | x | x | x | x | x | x | x | x | x | x | x | x | x | x | x | x | x | x | x | x | x | x | x | x | x | x | x | x | x | x | x | x | x | x | x | x | x | x | x | x | x | x | x | x | x | x | x | x | x | x | x | x | x | x | x | x | x | x | x | x | x | x | x | x | x | x | x | x | x | x | x | x | x | x | x | x | x | x | x | x | x | x | x | x | x | x | x | x | x | x | x | x | x | x | x | x | x | x | x | x | x | x | x | x | x | x | x | x | x | x | x | x | x | x | x | x | x | x | x | x | x | x | x | x | x | x | x | x | x | x | x | x | x | x | x | x | x | x | x | x | x | x | x | x | x | x | x | x | x | x | x | x | x | x | x | x | x | x | x | x | x | x | x | x | x | x | x | x | x | x | x | x | x | x | x | x | x | x | x | x | x | x | x | x | x | x | x | x | x | x | x | x | x | x | x | x | x | x | x | x | x | x | x | x | x | x | x | x | x | x | x | x | x | x | x | x | x | x | x | x | x | x | x | x | x | x | x | x | x | x | x | x | x | x | x | x | x | x | x | x | x | x | x | x | x | x | x | x | x | x | x | x | x | x | x | x | x | x | x | x | x | x | x | x | x | x | x | x | x | x | x | x | x | x | x | x | x | x | x | x | x | x | x | x | x | x | x | x | x | x | x | x | x | x | x | x | x | x | x | x | x | x | x | x | x | x | x | x | x | x | x | x | x | x | x | x | x | x | x | x | x | x | x | x | x | x | x | x | x | x | x | x | x | x | x | x | x | x | x | x | x | x | x | x | x | x | x | x | x | x | x | x | x | x | x | x | x | x | x | x | x | x | x | x | x | x | x | x | x | x | x | x | x | x | x | x | x | x | x | x | x | x | x | x | x | x | x | x | x | x | x | x | x | x | x | x | x | x | x | x | x | x | x | x | x | x | x | x | x | x | x | x | x | x | x | x | x | x | x | x | x | x | x | x | x | x | x | x | x | x | x | x | x | x | x | x | x | x | x | x | x | x | x | x | x | x | x | x | x | x | x | x | x | x | x | x | x | x | x | x | x | x | x | x | x | x | x | x | x | x | x | x | x | x | x | x | x | x | x | x | x | x | x | x | x | x | x | x | x | x | x | x | x | x | x | x | x | x | x | x | x | x | x | x | x | x | x | x | x | x | x | x | x | x | x | x | x | x | x | x | x | x | x | x | x | x | x | x | x | x | x | x | x | x | x | x | x | x | x | x | x | x | x | x | x | x | x | x | x | x | x | x | x | x | x | x | x | x | x | x | x | x | x | x | x | x | x | x | x | x | x | x | x | x | x | x | x | x | x | x | x | x | x | x | x | x | x | x | x | x | x | x | x | x | x | x | x | x | x | x | x | x | x | x | x | x | x | x | x | x | x | x | x | x | x | x | x | x | x | x | x | x | x | x | x | x | x | x | x | x | x | x | x | x | x | x | x | x | x | x | x | x | x | x | x | x | x | x | x | x | x | x | x | x | x | x | x | x | x | x | x | x | x | x | x | x | x | x | x | x | x | x | x | x | x | x | x | x | x | x | x | x | x | x | x | x | x | x | x | x | x | x | x | x | x | x | x | x | x | x | x | x | x | x | x | x | x | x | x | x | x | x | x | x | x | x | x | x | x | x | x | x | x | x | x | x | x | x | x | x | x | x | x | x | x | x | x | x | x | x | x | x | x | x | x | x | x | x | x | x | x | x | x | x | x | x | x | x | x | x | x | x | x | x | x | x | x | x | x | x | x | x | x | x | x | x | x | x | x | x | x | x | x | x | x | x | x | x | x | x | x | x | E | E | H | H | E | E | - | - | - | - | - | - | - | - | - | - | - | - | - | - | - | - | - | - | - | - | - | - | - | - | - | - | - | - | - | - | - | - | - | - | - | - | - | - | - | - | - | - | - | - | - | - | - | - | - | - | - | - | - | - |
| BIS d=0 cl.11 sym=1 env=1 | - | - | - | - | - | - | - | - | - | - | - | - | - | - | - | - | - | - | - | - | - | - | - | - | - | - | - | - | - | - | - | - | - | - | - | - | - | - | - | - | - | - | - | - | - | - | - | - | - | - | - | - | - | - | - | - | - | - | - | - | - | - | - | - | - | - | - | - | - | - | - | - | - | - | - | - | - | - | - | - | - | - | - | - | - | - | - | - | - | - | - | - | - | - | - | - | - | - | - | - | - | - | - | - | - | - | - | - | - | - | - | - | - | - | - | - | - | - | - | - | - | - | - | - | - | - | - | - | - | - | - | - | - | - | - | - | - | - | - | - | - | - | - | - | - | - | - | - | - | - | - | - | - | - | - | - | - | - | - | - | - | - | - | - | - | - | - | - | - | - | - | - | - | - | - | - | - | - | - | - | - | - | - | - | - | - | - | - | - | - | - | - | - | - | - | - | - | - | - | - | - | - | - | - | - | - | - | - | - | - | - | - | - | - | - | - | - | - | - | - | - | - | - | - | - | - | - | - | - | - | - | - | - | - | - | - | - | - | - | - | - | - | - | - | - | - | - | - | - | - | - | - | - | - | - | - | - | - | - | - | - | - | - | - | - | - | - | - | - | - | - | - | - | - | - | - | - | - | - | - | - | - | - | - | - | - | - | - | - | - | - | - | - | - | - | - | - | - | - | - | - | - | - | - | - | - | - | - | - | - | - | - | - | - | - | - | - | - | - | - | - | - | - | - | - | - | - | - | - | - | - | - | - | - | - | - | - | - | - | - | - | - | - | - | - | - | - | - | - | - | - | - | - | - | - | - | - | - | - | - | - | - | - | - | - | - | - | - | - | - | - | - | - | - | - | - | - | - | - | - | - | - | - | - | - | - | - | - | - | - | - | - | - | - | - | - | - | - | - | - | - | - | - | - | - | - | - | - | - | - | - | - | - | - | - | - | - | - | - | - | - | - | - | - | - | - | - | - | - | - | - | - | - | - | - | - | - | - | - | - | - | - | - | - | - | - | - | - | - | - | - | - | - | - | - | - | - | - | - | - | - | - | - | - | - | - | - | - | - | - | - | - | - | - | - | - | - | - | - | - | - | - | - | - | - | - | - | - | - | - | - | - | - | - | - | - | - | - | - | - | - | - | - | - | - | - | - | - | - | - | - | - | - | - | - | - | - | - | - | - | - | - | - | - | - | - | - | - | - | - | - | - | - | - | - | - | - | - | - | - | - | - | - | - | - | - | - | - | - | - | - | - | - | - | - | - | - | - | - | - | - | - | - | - | - | - | - | - | - | - | - | - | - | - | - | - | - | - | - | - | - | - | - | - | - | - | - | - | - | - | - | - | - | - | - | - | - | - | - | - | - | - | - | - | - | - | - | - | - | - | - | - | - | - | - | - | - | - | - | - | - | - | - | - | - | - | - | - | - | - | - | - | - | - | - | - | - | - | - | - | - | - | - | - | - | - | - | - | - | - | - | - | - | - | - | - | - | - | - | - | - | - | - | - | - | - | - | - | - | - | - | - | - | - | - | - | - | - | - | - | - | - | - | - | - | - | - | - | - | - | - | - | - | - | - | - | - | - | - | - | - | - | - | - | - | - | - | - | - | - | - | - | - | - | - | - | - | - | - | - | - | - | - | - | - | - | - | - | - | - | - | - | - | - | - | - | - | - | - | - | - | - | - | - | - | - | - | - | - | - | - | - | - | - | - | - | - | - | - | - | - | - | - | - | - | - | - | - | - | - | - | - | - | - | - | - | - | - | - | - | - | - | - | - | - | - | - | - | - | - | - | - | - | - | - | - | - | - | - | - | - | - | - | - | - | - | - | - | - | - | - | - | - | - | - | - | - | - | - | - | - | - | - | - | - | - | - | - | - | - | - | - | - | - | - | - | - | - | - | - | - | - | - | - | - | - | - | - | - | - | - | - | - | - | - | - | - | - | - | - | - | - | - | - | - | - | - | - | - | - | - | - | - | - | - | - | - | - | - | - | - | - | - | - | - | - | - | - | - | - | - | - | - | - | - | - | - | - | - | - | - | - | - | - | - | - | - | - | - | - | - | - | - | - | - | - | - | - | - | - | - | - | - | - | - | - | - | - | - | - | - | - | - | - | - | - | - | - | - | - | - | - | - | - | - | - | - | - | - | - | - | - | - | - | - | - | - | - | - | - | - | - | - | - | - | - | - | - | - | - | - | - | - | - | - | - | - | - | - | - | - | - | - | - | - | - | - | - | - | - | - | - | - | - | - | - | - | - | - | - | - | - | - | - | - | - | - | - | - | - | - | - | - | - | - | - | - | - | - | - | - | - | - | - | - | - | - | - | - | - | - | - | - | - | - | - | - | - | - | - | - | - | - | - | - | - | - | - | - | - | - | - | - | - | - | - | - | - | - | - | - | - | - | - | - | - | - | - | - | - | - | - | - | - | - | - | - | - | - | - | - | - | - | - | - | - | - | - | - | - | - | - | - | - | - | - | - | - | - | - | - | - | - | - | - | - | - | - | - | - | - | - | - | - | - | - | - | - | - | - | - | - | - | - | - | - | - | - | - | - | - | - | - | - | - | - | - | - | - | - | - | - | - | - | - | - | - | - | - | - | - | - | - | - | - | - | - | - | - | - | - | - | - | - | - | - | - | - | - | - | - | - | - | - | - | - | - | - | - | - | - | - | - | - | - | - | - | - | - | - | - | - | - | - | - | - | - | - | - | - | - | - | - | - | - | - | - | - | - | - | - | - | - | - | - | - | - | - | - | - | - | - | - | - | - | - | - | - | - | - | - | - | - | - | - | - | - | - | - | - | - | - | - | - | - | - | - | - | - | - | - | - | - | - | - | - | - | - | - | - | - | - | - | - | - | - | - | - | - | - | - | - | - | - | - | - | - | - | - | - | - | - | - | - | - | - | - | - | - | - | - | - | - | - | - | - | - | - | - | - | - | - | - | - | - | - | - | - | - | - | - | - | - | - | - | - | - | - | - | - | - | - | - | - | - | - | - | - | - | - | - | - | - | - | - | - | - | - | - | - | - | - | - | - | - | - | - | - | - | - | - | - | - | - | - | - | - | - | - | - | - | - | - | - | - | - | - | - | - | - | - | - | - | - | - | - | - | - | - | - | - | - | - | - | - | - | - | - | - | - | - | - | - | - | - | - | - | - | - | - | - | - | - | - | - | - | - | - | - | - | - | - | - | - | - | - | - | - | - | - | - | - | - | - | - | - | - | - | - | - | - | - | - | - | - | - | - | - | - | - | - | - | - | - | - | - | - | - | - | - | - | - | - | - | - | - | - | - | - | - | - | - | - | - | - | - | - | - | - | - | - | - | - | - | - | - | - | - | - | - | - | - | - | - | - | - | - | - | - | - | - | - | - | - | - | - | - | - | - | - | - | - | - | - | - | - | - | - | - | - | - | - | - | - | - | - | - | - | - | - | - | - | - | - | - | - | - | - | - | - | - | - | - | - | - | - | - | - | - | - | - | - | - | - | - | - | - | - | - | - | - | - | - | - | - | - | - | - | - | - | - | - | - | - | - | - | - | - | - | - | - | - | - | - | - | - | - | - | - | - | - | - | - | - | - | - | - | - | - | - | - | - | - | - | - | - | - | - | - | - | - | - | - | - | - | - | - | - | - | - | - | - | - | - | - | - | - | - | - | - | - | - | - | - | - | - | - | - | - | - | - | - | - | - | - | - | - | - | - | - | - | - | - | - | - | - | - | - | - | - | - | - | - | - | - | - | - | - | - | - | - | - | - | - | - | - | - | - | - | - | - | - | - | - | - | - | - | - | - | - | - | - | - | - | - | - | - | - | - | - | - | - | - | - | - | - | - | - | - | - | - | - | - | - | - | - | - | - | - | - | - | - | - | - | - | - | - | - | - | - | - | - | - | - | - | - | - | - | - | - | - | - | - | - | - | - | - | - | - | - | - | - | - | - | - | - | - | - | - | - | - | - | - | - | - | - | - | - | - | - | - | - | - | - | - | - | - | - | - | - | - | - | - | - | - | - | - | - | - | - | - | - | - | - | - | - | - | - | - | - | - | - | - | - | - | - | - | - | - | - | - | - | - | - | - | - | - | - | - | - | - | - | - | - | - | - | - | - | - | - | - | - | - | - | - | - | - | - | - | - | - | - | - | - | - | - | - | - | - | - | - | - | - | - | - | - | - | - | - | - | - | - | - | - | - | - | - | - | - | - | - | - | - | - | - | - | - | - | - | - | - | - | - | - | - | - | - | - | - | - | - | - | - | - | - | - | - | - | - | - | - | - | - | - | - | - | - | - | - | - | - | - | - | - | - | - | - | - | - | - | - | - | - | - | - | - | - | - | - | - | - | - | - | - | - | - | - | - | - | - | - | - | - | - | - | - | - | - | - | - | - | - | - | - | - | - | - | - | - | - | - | - | - | - | - | - | - | - | - | - | - | - | - | - | - | - | - | - | - | - | - | - | - | - | - | - | - | - | - | - | - | - | - | - | - | - | - | - | - | - | - | - | - | - | - | - | - | - | - | - | - | - | - | - | - | - | - | - | - | - | - | - | - | - | - | - | - | - | - | - | - | - | - | - | - | - | - | - | - | - | - | - | - | - | - | - | - | - | - | - | - | - | - | - | - | - | - | - | - | - | - | - | - | - | - | - | - | - | - | - | - | - | - | - | - | - | - | - | - | - | - | - | - | - | - | - | - | - | - | - | - | - | - | - | - | - | - | - | - | - | - | - | - | - | - | - | - | - | - | - | - | - | - | - | - | - | - | - | - | - | - | - | - | - | - | - | - | - | - | - | - | - | - | - | - | - | - | - | - | - | - | - | - | - | - | - | - | - | - | - | - | - | - | - | - | - | - | - | - | - | - | - | - | - | - | - | - | - | - | - | - | - | - | - | - | - | - | - | - | - | - | - | - | - | - | - | - | - | - | - | - | - | - | - | - | - | - | - | - | - | - | - | - | - | - | - | - | - | - | - | - | - | - | - | - | - | - | - | - | - | - | - | - | - | - | - | - | - | - | - | - | - | - | - | - | - | - | - | - | - | - | - | - | - | - | - | - | - | - | - | - | - | - | - | - | - | - | - | - | - | - | - | - | - | - | - | - | - | - | - | - | - | - | - | - | - | - | - | - | - | - | - | - | - | - | - | - | - | - | - | - | - | - | - | - | - | - | - | - | - | - | - | - | - | - | - | - | - | - | - | - | - | - | - | - | - | - | - | - | - | - | - | - | - | - | - | - | - | - | - | - | - | - | - | - | - | - | - | - | - | - | - | - | - | - | - | - | - | - | - | - | - | - | - | - | - | - | - | - | - | - | - | - | - | - | - | - | - | - | - | - | - | - | - | - | - | - | - | - | - | - | - | - | - | - | - | - | - | - | - | - | - | - | - | - | - | - | - | - | - | - | - | - | - | - | - | - | - | - | - | - | - | - | - | - | - | - | - | - | - | - | - | - | - | - | - | - | - | - | - | - | - | - | - | - | - | - | - | - | - | - | - | - | - | - | - | - | - | - | - | - | - | - | - | - | - | - | - | - | - | - | - | - | - | - | - | - | - | - | - | - | - | - | - | - | - | - | - | - | - | - | - | - | - | - | - | - | - | - | - | - | - | - | - | - | - | - | - | - | - | - | - | - | - | - | - | - | - | - | - | - | - | - | - | - | - | - | - | - | - | - | - | - | - | - | - | - | - | - | - | - | - | - | - | - | - | - | - | - | - | - | - | - | - | - | - | - | - | - | - | - | - | - | - | - | - | - | - | - | - | - | - | - | - | - | - | - | - | - | - | - | - | - | - | - | - | - | - | - | - | - | - | - | - | - | - | - | - | - | - | - | - | - | - | - | - | - | - | - | - | - | - | - | - | - | - | - | - | - | - | - | - | - | - | - | - | - | - | - | - | - | - | - | - | - | - | - | - | - | - | - | - | - | - | - | - | - | - | - | - | - | - | - | - | - | - | - | - | - | - | - | - | - | - | - | - | - | - | - | - | - | - | - | - | - | - | - | - | - | - | - | - | - | - | - | - | - | - | - | - | - | - | - | - | - | - | - | - | - | - | - | - | - | - | - | - | - | - | - | - | - | - | - | - | - | - | - | - | - | - | - | - | - | - | - | - | - | - | - | - | - | - | - | - | - | - | - | - | - | - | - | - | - | - | - | - | - | - | - | - | - | - | - | - | - | - | - | - | - | - | - | - | - | - | - | - | - | - | - | - | - | - | - | - | - | - | - | - | - | - | - | - | - | - | - | - | - | - | - | - | - | - | - | - | - | - | - | - | - | - | - | - | - | - | - | - | - | - | - | - | - | - | - | - | - | - | - | - | - | - | - | - | - | - | - | - | - | - | - | - | - | - | - | - | - | - | - | - | - | - | - | - | - | - | - | - | - | - | - | - | - | - | - | - | - | - | - | - | - | - | - | - | - | - | - | - | - | - | - | - | - | - | - | - | - | - | - | - | - | - | - | - | - | - | - | - | - | - | - | - | - | - | - | - | - | - | - | - | - | - | - | - | - | - | - | - | - | - | - | - | - | - | - | - | - | - | - | - | - | - | - | - | - | - | - | - | - | - | - | - | - | - | - | - | - | - | - | - | - | - | - | - | - | - | - | - | - | - | - | - | - | - | - | - | - | - | - | - | - | - | - | - | - | - | - | - | - | - | - | - | - | - | - | - | - | - | - | - | - | - | - | - | - | - | - | - | - | - | - | - | - | - | - | - | - | - | - | - | - | - | - | - | - | - | - | - | - | - | - | - | - | - | - | - | - | - | - | - | - | - | - | - | - | - | - | - | - | - | - | - | - | - | - | - | - | - | - | - | - | - | - | - | - | - | - | - | - | - | - | - | - | - | - | - | - | - | - | - | - | - | - | - | - | - | - | - | - | - | - | - | - | - | - | - | - | - | - | - | - | - | - | - | - | - | - | - | - | - | - | - | - | - | - | - | - | - | - | - | - | - | - | - | - | - | - | - | - | - | - | - | - | - | - | - | - | - | - | - | - | - | - | - | - | - | - | - | - | - | - | - | - | - | - | - | - | - | - | - | - | - | - | - | - | - | - | - | - | - | - | - | - | - | - | - | - | - | - | - | - | - | - | - | - | - | - | - | - | - | - | - | - | - | - | - | - | - | - | - | - | - | - | - | - | - | - | - | - | - | - | - | - | - | - | - | - | - | - | - | - | - | - | - | - | - | - | - | - | - | - | - | - | - | - | - | - | - | - | - | - | - | - | - | - | - | - | - | - | - | - | - | - | H | x | x | x | x | x | x | x | x | x | x | x | x | x | x | x | x | x | x | x | x | x | x | x | x | x | x | x | x | x | x | x | x | x | x | x | x | x | x | x | x | x | x | x | x | x | x | x | x | x | x | x | x | x | x | x | x | x | x | x | x | x | x | x | x | x | x | x | x | x | x | x | x | x | x | x | x | x | x | x | x | x | x | x | x | x | x | x | x | x | x | x | x | x | x | x | x | H | H | x | x | x | x | x | x | x | x | x | x | x | x | x | x | x | x | x | x | x | x | x | x | x | x | x | x | x | x | x | x | x | x | x | x | x | x | x | x | x | x | x | x | x | x | x | x | x | x | x | x | x | x | x | x | x | x | x | x | x | x | x | x | x | x | x | x | x | x | x | x | x | x | x | x | x | x | x | x | x | x | x | x | x | x | x | x | x | x | x | x | x | x | x | x | x | x | x | x | x | x | x | x | x | H | x | x | x | x | x | x | x | x | x | x | x | x | x | x | x | x | x | x | x | x | x | x | x | x | x | x | x | x | x | x | x | x | x | x | x | x | x | x | x | x | x | x | x | x | x | x | x | x | x | x | x | x | x | x | x | x | x | x | x | x | x | x | x | x | x | x | x | x | x | x | x | x | x | x | x | x | x | x | x | x | x | x | x | x | x | x | x | x | x | x | x | x | x | x | x | x | x | x | x | x | x | x | x | x | x | x | x | x | x | x | x | x | x | x | x | x | x | x | x | x | x | x | x | x | x | x | x | x | x | x | x | x | x | x | x | x | x | x | x | x | x | x | x | x | x | x | x | x | x | x | x | x | x | x | x | x | x | x | x | x | x | x | x | x | x | x | x | x | x | x | x | x | x | x | x | x | x | x | x | x | x | x | x | x | x | x | x | x | x | x | x | x | x | x | x | x | x | x | x | x | x | x | x | x | x | x | x | x | x | x | x | x | x | x | x | x | x | x | x | x | x | x | x | x | x | x | x | x | x | x | x | x | x | x | x | x | x | x | x | x | x | x | x | x | x | x | x | x | x | x | x | x | x | x | x | x | x | x | x | x | x | x | x | x | x | x | x | x | x | x | x | x | x | x | x | x | x | x | x | x | x | x | x | x | x | x | x | x | x | x | x | x | x | x | x | x | x | x | x | x | x | x | x | x | x | x | x | x | x | x | x | x | x | x | x | x | x | x | x | x | x | x | x | x | x | x | x | x | x | x | x | x | x | x | x | x | x | x | x | x | x | x | x | x | x | x | x | x | x | x | x | x | x | x | x | x | x | x | x | x | x | x | x | x | x | x | x | x | x | x | x | x | x | x | x | x | x | x | x | x | x | x | x | x | x | x | x | x | x | x | x | x | x | x | x | x | x | x | x | x | x | x | x | x | x | x | x | x | x | x | x | x | x | x | x | x | x | x | x | x | x | x | x | x | x | x | x | x | x | x | x | x | x | x | x | x | x | x | x | x | x | x | x | x | x | x | x | x | x | x | x | x | x | x | x | x | x | x | x | x | x | x | x | x | x | x | x | x | x | x | x | x | x | x | x | x | x | x | x | x | x | x | x | x | x | x | x | x | x | x | x | x | x | x | x | x | x | x | x | x | x | x | x | x | x | x | x | x | x | x | x | x | x | x | x | x | x | x | x | x | x | x | x | x | x | x | H | - | - | - | - | - | - | - | - | - | - | - | - | - | - | - | - | - | - | - | - | - | - | - | - | - | - | - | - | - | - | - | - | - | - | - | - | - | - | - | - | - | - | - | - | - | - | - | - | - | - | - | - | - | - | - | - | - | - | - | - | - | - | - | - | - | - | - | - | - | - | - | - | - | - | - | - | - | - | - | - | - | - | - | - | - | - | - | - | - | - | - | - | - | - | - | - | - | - | - | - | - | - | - | - | - | - | - | - | - | - | - | - | - | - | - | - | - | - | - | - | - | - | - | - | - | - | - | - | - | - | - | - | - | - | - | - | - | - | - | - | - | - | - | - | - | - | - | - | - | - | - | - | - | - | - | - | - | - | - | - | - | - | - | - | - | - | - | - | - | - | - | - | - | - | - | - | - | - | - | - | - | - | - | - | - | - | - | - | - | - | - | - | - | - | - | - | - | - | - | - | - | - | - | - | - | - | - | - | - | - | - | - | - | - | - | - | - | - | - | - | - | - | - | - | - | - | - | - | - | - | - | - | - | - | - | - | - | - | - | - | - | - | - | - | - | - | - | - | - | - | - | - | - | - | - | - | - | - | - | - | - | - | - | - | - | - | - | - | - | - | - | - | - | - | - | - | - | - | - | - | - | - | - | - | - | - | - | - | - | - | - | - | - | - | - | - | - | - | - | - | - | - | - | - | - | - | - | - | - | - | - | - | - | - | - | - | - | - | - | - | - | - | - | - | - | - | - | - | - | - | - | - | - | - | - | - | - | - | - | - | - | - | - | - | - | - | - | - | - | - | - | - | - | - | - | - | - | - | - | - | - | - | - | - | - | - | - | - | - | - | - | - | - | - | - | - | - | - | - | - | - | - | - | - | - | - | - | - | - | - | - | - | - | - | - | - | - | - | - | - | - | - | - | - | - | - | - | - | - | - | - | - | - | - | - | - | - | - | - | - | - | - | - | - | - | - | - | - | - | - | - | - | - | - | - | - | - | - | - | - | - | - | - | - | - | - | - | - | - | - | - | - | - | - | - | - | - | - | - | - | - | - | - | - | - | - | - | - | - | - | - | - | - | - | - | - | - | - | - | - | - | - | - | - | - | - | - | - | - | - | - | - | - | - | - | - | - | - | - | - | - | - | - | - | - | - | - | - | - | - | - | - | - | - | - | - | - | - | - | - | - | - | - | - | - | - | - | - | - | - | - | - | - | - | - | - | - | - | - | - | - | - | - | - | - | - | - | - | - | - | - | - | - | - | - | - | - | - | - | - | - | - | - | - | - | - | - | - | - | - | - | - | - | - | - | - | - | - | - | - | - | - | - | - | - | - | - | - | - | - | - | - | - | - | - | - | - | - | - | - | - | - | - | - | - | - | - | - | - | - | - | - | - | - | - | - | - | - | - | - | - | - | - | - | - | - | - | - | - | - | - | - | - | - | - | - | - | - | - | - | - | - | - | - | - | - | - | - | - | - | - | - | - | - | - | - | - | - | - | - | - | - | - | - | - | - | - | - | - | - | - | - | - | - | - | - | - | - | - | - | - | - | - | - | - | - | - | - | - | - | - | - | - | - | - | - | - | - | - | - | - | - | - | - | - | - | - | - | - | - | - | - | - | - | - | - | - | - | - | - | - | - | - | - | - | - | - | - | - | - | - | - | - | - | - | - | - | - | - | - | - | - | - | - | - | - | - | - | - | - | - | - | - | - | - | - | - | - | - | - | - | - | - | - | - | - | - | - | - | - | - | - | - | - | - | - | - | - | - | - | - | - | - | - | - | - | - | - | - | - | - | - | - | - | - | - | - | - | - | - | - | - | - | - | - | - | - | - | - | - | - | - | - | - | - | - | - | - | - | - | - | - | - | - | - | - | - | - | - | - | - | - | - | - | - | - | - | - | - | - | - | - | - | - | - | - | - | - | - | - | - | - | - | - | - | - | - | - | - | - | - | - | - | - | - | - | - | - | - | - | - | - | - | - | - | - | - | - | - | - | - | - | - | - | - | - | - | - | - | - | - | - | - | - | - | - | - | - | - | - | - | - | - | - | - | - | - | - | - | - | - | - | - | - | - | - | - | - | - | - | - | - | - | - | - | - | - | - | - | - | - | - | - | - | - | - | - | - | - | - | - | - | - | - | - | - | - | - | - | - | - | - | - | - | - | - | - | - | - | - | - | - | - | - | - | - | - | - | - | - | - | - | - | - | - | - | - | - | - | - | - | - | - | - | - | - | - | - | - | - | - | - | - | - | - | - | - | - | - | - | - | - | - | - | - | - | - | - | - | - | - | - | - | - | - | - | - | - | - | - | - | - | - | - | - | - | - | - | - | - | - | - | - | - | - | - | - | - | - | - | - | - | - | - | - | - | - | - | - | - | - | - | - | - | - | - | - | - | - | - | - | - | - | - | - | - | - | - | - | - | - | - | - | - | - | - | - | - | - | - | - | - | - | - | - | - | - | - | - | - | - | - | - | - | - | - | - | - | - | - | - | - | - | - | - | - | - | - | - | - | - | - | - | - | - | - | - | - | - | - | - | - | - | - | - | - | - | - | - | - | - | - | - | - | - | - | - | - | - | - | - | - | - | - | - | - | - | - | - | - | - | - | - | - | - | - | - | - | - | - | - | - | - | - | - | - | - | - | - | - | - | - | - | - | - | - | - | - | - | - | - | - | - | - | - | - | - | - | - | - | - | - | - | - | - | - | - | - | - | - | - | - | - | - | - | - | - | - | - | - | - | - | - | - | - | - | - | - | - | - | - | - | - | - | - | - | - | - | - | - | - | - | - | - | - | - | - | - | - | - | - | - | - | - | - | - | - | - | - | - | - | - | - | - | - | - | - | - | - | - | - | - | - | - | - | - | - | - | - | - | - | - | - | - | - | - | - | - | - | - | - | - | - | - | - | - | - | - | - | - | - | - | - | - | - | - | - | - | - | - | - | - | - | - | - | - | - | - | - | - | - | - | - | - | - | - | - | - | - | - | - | - | - | - | - | - | - | - | - | - | - | - | - | - | - | - | - | - | - | - | - | - | - | - | - | - | - | - | - | - | - | - | - | - | - | - | - | - | - | - | - | - | - | - | - | - | - | - | - | - | - | - | - | - | - | - | - | - | - | - | - | - | - | - | - | - | - | - | - | - | - | - | - | - | - | - | - | - | - | - | - | - | - | - | - | - | - | - | - | - | - | - | - | - | - | - | - | - | - | - | - | - | - | - | - | - | - | - | - | - | - | - | - | - | - | - | - | - | - | - | - | - | - | - | - | - | - | - | - | - | - | - | - | - | - | - | - | - | - | - | - | - | - | - | - | - | - | - | - | - | - | - | - | - | - | - | - | - | - | - | - | - | - | - | - | - | - |
| BIS d=0 cl.19 sym=1 env=1 | - | - | - | - | - | - | - | - | - | - | - | - | - | - | - | - | - | - | - | - | - | - | - | - | - | - | - | - | - | - | - | - | - | - | - | - | - | - | - | - | - | - | - | - | - | - | - | - | - | - | - | - | - | - | - | - | - | - | - | - | - | - | - | - | - | - | - | - | - | - | - | - | - | - | - | - | - | - | - | - | - | - | - | - | - | - | - | - | - | - | - | - | - | - | - | - | - | - | - | - | - | - | - | - | - | - | - | - | - | - | - | - | - | - | - | - | - | - | - | - | - | - | - | - | - | - | - | - | - | - | - | - | - | - | - | - | - | - | - | - | - | - | - | - | - | - | - | - | - | - | - | - | - | - | - | - | - | - | - | - | - | - | - | - | - | - | - | - | - | - | - | - | - | - | - | - | - | - | - | - | - | - | - | - | - | - | - | - | - | - | - | - | - | - | - | - | - | - | - | - | - | - | - | - | - | - | - | - | - | - | - | - | - | - | - | - | - | - | - | - | - | - | - | - | - | - | - | - | - | - | - | - | - | - | - | - | - | - | - | - | - | - | - | - | - | - | - | - | - | - | - | - | - | - | - | - | - | - | - | - | - | - | - | - | - | - | - | - | - | - | - | - | - | - | - | - | - | - | - | - | - | - | - | - | - | - | - | - | - | - | - | - | - | - | - | - | - | - | - | - | - | - | - | - | - | - | - | - | - | - | - | - | - | - | - | - | - | - | - | - | - | - | - | - | - | - | - | - | - | - | - | - | - | - | - | - | - | - | - | - | - | - | - | - | - | - | - | - | - | - | - | - | - | - | - | - | - | - | - | - | - | - | - | - | - | - | - | - | - | - | - | - | - | - | - | - | - | - | - | - | - | - | - | - | - | - | - | - | - | - | - | - | - | - | - | - | - | - | - | - | - | - | - | - | - | - | - | - | - | - | - | - | - | - | - | - | - | - | - | - | - | - | - | - | - | - | - | - | - | - | - | - | - | - | - | - | - | - | - | - | - | - | - | - | - | - | - | - | - | - | - | - | - | - | - | - | - | - | - | - | - | - | - | - | - | - | - | - | - | - | - | - | - | - | - | - | - | - | - | - | - | - | - | - | - | - | - | - | - | - | - | - | - | - | - | - | - | - | - | - | - | - | - | - | - | - | - | - | - | - | - | - | - | - | - | - | - | - | - | - | - | - | - | - | - | - | - | - | - | - | - | - | - | - | - | - | - | - | - | - | - | - | - | - | - | - | - | - | - | - | - | - | - | - | - | - | - | - | - | - | - | - | - | - | - | - | - | - | - | - | - | - | - | - | - | - | - | - | - | - | - | - | - | - | - | - | - | - | - | - | - | - | - | - | - | - | - | - | - | - | - | - | - | - | - | - | - | - | - | - | - | - | - | - | - | - | - | - | - | - | - | - | - | - | - | - | - | - | - | - | - | - | - | - | - | - | - | - | - | - | - | - | - | - | - | - | - | - | - | - | - | - | - | - | - | - | - | - | - | - | - | - | - | - | - | - | - | - | - | - | - | - | - | - | - | - | - | - | - | - | - | - | - | - | - | - | - | - | - | - | - | - | - | - | - | - | - | - | - | - | - | - | - | - | - | - | - | - | - | - | - | - | - | - | - | - | - | - | - | - | - | - | - | - | - | - | - | - | - | - | - | - | - | - | - | - | - | - | - | - | - | - | - | - | - | - | - | - | - | - | - | - | - | - | - | - | - | - | - | - | - | - | - | - | - | - | - | - | - | - | - | - | - | - | - | - | - | - | - | - | - | - | - | - | - | - | - | - | - | - | - | - | - | - | - | - | - | - | - | - | - | - | - | - | - | - | - | - | - | - | - | - | - | - | - | - | - | - | - | - | - | - | - | - | - | - | - | - | - | - | - | - | - | - | - | - | - | - | - | - | - | - | - | - | - | - | - | - | - | - | - | - | - | - | - | - | - | - | - | - | - | - | - | - | - | - | - | - | - | - | - | - | - | - | - | - | - | - | - | - | - | - | - | - | - | - | - | - | - | - | - | - | - | - | - | - | - | - | - | - | - | - | - | - | - | - | - | - | - | - | - | - | - | - | - | - | - | - | - | - | - | - | - | - | - | - | - | - | - | - | - | - | - | - | - | - | - | - | - | - | - | - | - | - | - | - | - | - | - | - | - | - | - | - | - | - | - | - | - | - | - | - | - | - | - | - | - | - | - | - | - | - | - | - | - | - | - | - | - | - | - | - | - | - | - | - | - | - | - | - | - | - | - | - | - | - | - | - | - | - | - | - | - | - | - | - | - | - | - | - | - | - | - | - | - | - | - | - | - | - | - | - | - | - | - | - | - | - | - | - | - | - | - | - | - | - | - | - | - | - | - | - | - | - | - | - | - | - | - | - | - | - | - | - | - | - | - | - | - | - | - | - | - | - | - | - | - | - | - | - | - | - | - | - | - | - | - | - | - | - | - | - | - | - | - | - | - | - | - | - | - | - | - | - | H | H | x | x | H | H | x | x | H | x | H | H | x | x | x | x | x | x | x | x | x | x | x | x | x | x | x | x | x | x | x | x | H | x | x | x | x | x | H | x | x | x | H | - | - | - | - | - | - | - | - | - | - | - | - | - | - | - | - | - | - | - | - | - | - | - | - | - | - | - | - | - | - | - | - | - | - | - | - | - | - | - | - | - | - | - | - | - | - | - | - | - | - | - | - | - | - | - | - | - | - | - | - | - | - | - | - | - | - | - | - | - | - | - | - | - | - | - | - | - | - | - | - | - | - | - | - | - | - | - | - | - | - | - | - | - | - | - | - | - | - | - | - | - | - | - | - | - | - | - | - | - | - | - | - | - | - | - | - | - | - | - | - | - | - | - | - | - | - | - | - | - | - | - | - | - | - | - | - | - | - | - | - | - | - | - | - | - | - | - | - | - | - | - | - | - | - | - | - | - | - | - | - | - | - | - | - | - | - | - | - | - | - | - | - | - | - | - | - | - | - | - | - | - | - | - | - | - | - | - | - | - | - | - | - | - | - | - | - | - | - | - | - | - | - | - | - | - | - | - | - | - | - | - | - | - | - | - | - | - | - | - | - | - | - | - | - | - | - | - | - | - | - | - | - | - | - | - | - | - | - | - | - | - | - | - | - | - | - | - | - | - | - | - | - | - | - | - | - | - | - | - | - | - | - | - | - | - | - | - | - | - | - | - | - | - | - | - | - | - | - | - | - | - | - | - | - | - | - | - | - | - | - | - | - | - | - | - | - | - | - | - | - | - | - | - | - | - | - | - | - | - | - | - | - | - | - | - | - | - | - | - | - | - | - | - | - | - | - | - | - | - | - | - | - | - | - | - | - | - | - | - | - | - | - | - | - | - | - | - | - | - | - | - | - | - | - | - | - | - | - | - | - | - | - | - | - | - | - | - | - | - | - | - | - | - | - | - | - | - | - | - | - | - | - | - | - | - | - | - | - | - | - | - | - | - | - | - | - | - | - | - | - | - | - | - | - | - | - | - | - | - | - | - | - | - | - | - | - | - | - | - | - | - | - | - | - | - | - | - | - | - | - | - | - | - | - | - | - | - | - | - | - | - | - | - | - | - | - | - | - | - | - | - | - | - | - | - | - | - | - | - | - | - | - | - | - | - | - | - | - | - | - | - | - | - | - | - | - | - | - | - | - | - | - | - | - | - | - | - | - | - | - | - | - | - | - | - | - | - | - | - | - | - | - | - | - | - | - | - | - | - | - | - | - | - | - | - | - | - | - | - | - | - | - | - | - | - | - | - | - | - | - | - | - | - | - | - | - | - | - | - | - | - | - | - | - | - | - | - | - | - | - | - | - | - | - | - | - | - | - | - | - | - | - | - | - | - | - | - | - | - | - | - | - | - | - | - | - | - | - | - | - | - | - | - | - | - | - | - | - | - | - | - | - | - | - | - | - | - | - | - | - | - | - | - | - | - | - | - | - | - | - | - | - | - | - | - | - | - | - | - | - | - | - | - | - | - | - | - | - | - | - | - | - | - | - | - | - | - | - | - | - | - | - | - | - | - | - | - | - | - | - | - | - | - | - | - | - | - | - | - | - | - | - | - | - | - | - | - | - | - | - | - | - | - | - | - | - | - | - | - | - | - | - | - | - | - | - | - | - | - | - | - | - | - | - | - | - | - | - | - | - | - | - | - | - | - | - | - | - | - | - | - | - | - | - | - | - | - | - | - | - | - | - | - | - | - | - | - | - | - | - | - | - | - | - | - | - | - | - | - | - | - | - | - | - | - | - | - | - | - | - | - | - | - | - | - | - | - | - | - | - | - | - | - | - | - | - | - | - | - | - | - | - | - | - | - | - | - | - | - | - | - | - | - | - | - | - | - | - | - | - | - | - | - | - | - | - | - | - | - | - | - | - | - | - | - | - | - | - | - | - | - | - | - | - | - | - | - | - | - | - | - | - | - | - | - | - | - | - | - | - | - | - | - | - | - | - | - | - | - | - | - | - | - | - | - | - | - | - | - | - | - | - | - | - | - | - | - | - | - | - | - | - | - | - | - | - | - | - | - | - | - | - | - | - | - | - | - | - | - | - | - | - | - | - | - | - | - | - | - | - | - | - | - | - | - | - | - | - | - | - | - | - | - | - | - | - | - | - | - | - | - | - | - | - | - | - | - | - | - | - | - | - | - | - | - | - | - | - | - | - | - | - | - | - | - | - | - | - | - | - | - | - | - | - | - | - | - | - | - | - | - | - | - | - | - | - | - | - | - | - | - | - | - | - | - | - | - | - | - | - | - | - | - | - | - | - | - | - | - | - | - | - | - | - | - | - | - | - | - | - | - | - | - | - | - | - | - | - | - | - | - | - | - | - | - | - | - | - | - | - | - | - | - | - | - | - | - | - | - | - | - | - | - | - | - | - | - | - | - | - | - | - | - | - | - | - | - | - | - | - | - | - | - | - | - | - | - | - | - | - | - | - | - | - | - | - | - | - | - | - | - | - | - | - | - | - | - | - | - | - | - | - | - | - | - | - | - | - | - | - | - | - | - | - | - | - | - | - | - | - | - | - | - | - | - | - | - | - | - | - | - | - | - | - | - | - | - | - | - | - | - | - | - | - | - | - | - | - | - | - | - | - | - | - | - | - | - | - | - | - | - | - | - | - | - | - | - | - | - | - | - | - | - | - | - | - | - | - | - | - | - | - | - | - | - | - | - | - | - | - | - | - | - | - | - | - | - | - | - | - | - | - | - | - | - | - | - | - | - | - | - | - | - | - | - | - | - | - | - | - | - | - | - | - | - | - | - | - | - | - | - | - | - | - | - | - | - | - | - | - | - | - | - | - | - | - | - | - | - | - | - | - | - | - | - | - | - | - | - | - | - | - | - | - | - | - | - | - | - | - | - | - | - | - | - | - | - | - | - | - | - | - | - | - | - | - | - | - | - | - | - | - | - | - | - | - | - | - | - | - | - | - | - | - | - | - | - | - | - | - | - | - | - | - | - | - | - | - | - | - | - | - | - | - | - | - | - | - | - | - | - | - | - | - | - | - | - | - | - | - | - | - | - | - | - | - | - | - | - | - | - | - | - | - | - | - | - | - | - | - | - | - | - | - | - | - | - | - | - | - | - | - | - | - | - | - | - | - | - | - | - | - | - | - | - | - | - | - | - | - | - | - | - | - | - | - | - | - | - | - | - | - | - | - | - | - | - | - | - | - | - | - | - | - | - | - | - | - | - | - | - | - | - | - | - | - | - | - | - | - | - | - | - | - | - | - | - | - | - | - | - | - | - | - | - | - | - | - | - | - | - | - | - | - | - | - | - | - | - | - | - | - | - | - | - | - | - | - | - | - | - | - | - | - | - | - | - | - | - | - | - | - | - | - | - | - | - | - | - | - | - | - | - | - | - | - | - | - | - | - | - | - | - | - | - | - | - | - | - | - | - | - | - | - | - | - | - | - | - | - | - | - | - | - | - | - | - | - | - | - | - | - | - | - | - | - | - | - | - | - | - | - | - | - | - | - | - | - | - | - | - | - | - | - | - | - | - | - | - | - | - | - | - | - | - | - | - | - | - | - | - | - | - | - | - | - | - | - | - | - | - | - | - | - | - | - | - | - | - | - | - | - | - | - | - | - | - | - | - | - | - | - | - | - | - | - | - | - | - | - | - | - | - | - | - | - | - | - | - | - | - | - | - | - | - | - | - | - | - | - | - | - | - | - | - | - | - | - | - | - | - | - | - | - | - | - | - | - | - | - | - | - | - | - | - | - | - | - | - | - | - | - | - | - | - | - | - | - | - | - | - | - | - | - | - | - | - | - | - | - | - | - | - | - | - | - | - | - | - | - | - | - | - | - | - | - | - | - | - | - | - | - | - | - | - | - | - | - | - | - | - | - | - | - | - | - | - | - | - | - | - | - | - | - | - | - | - | - | - | - | - | - | - | - | - | - | - | - | - | - | - | - | - | - | - | - | - | - | - | - | - | - | - | - | - | - | - | - | - | - | - | - | - | - | - | - | - | - | - | - | - | - | - | - | - | - | - | - | - | - | - | - | - | - | - | - | - | - | - | - | - | - | - | - | - | - | - | - | - | - | - | - | - | - | - | - | - | - | - | - | - | - | - | - | - | - | - | - | - | - | - | - | - | - | - | - | - | - | - | - | - | - | - | - | - | - | - | - | - | - | - | - | - | - | - | - | - | - | - | - | - | - | - | - | - | - | - | - | - | - | - | - | - | - | - | - | - | - | - | - | - | - | - | - | - | - | - | - | - | - | - | - | - | - | - | - | - | - | - | - | - | - | - | - | - | - | - | - | - | - | - | - | - | - | - | - | - | - | - | - | - | - | - | - | - | - | - | - | - | - | - | - | - | - | - | - | - | - | - | - | - | - | - | - | - | - | - | - | - | - | - | - | - | - | - | - | - | - | - | - | - | - | - | - | - | - | - | - | - | - | - | - | - | - | - | - | - | - | - | - | - | - | - | - | - | - | - | - | - | - | - | - | - | - | - | - | - | - | - | - | - | - | - | - | - | - | - | - | - | - | - | - | - | - | - | - | - | - | - | - | - | - | - | - | - | - | - | - | - | - | - | - | - | - | - | - | - | - | - | - | - | - | - | - | - | - | - | - | - | - | - | - | - | - | - | - | - | - | - | - | - | - | - | - | - | - | - | - | - | - | - | - | - | - | - | - | - | - | - | - | - | - | - | - | - | - | - | - | - | - | - | - | - | - | - | - | - | - | - | - | - | - | - | - | - | - | - | - | - | - | - | - | - | - | - | - | - | - | - | - | - | - | - | - | - | - | - | - | - | - | - | - | - | - | - | - | - | - | - | - | - | - | - | - | - | - | - | - | - | - | - | - | - | - | - | - | - | - | - | - | - | - | - | - | - | - | - | - | - | - | - | - | - | - | - | - | - | - | - | - | - | - | - | - | - | - | - | - | - | - | - | - | - | - | - | - | - | - | - | - | - | - | - | - | - | - | - | - | - | - | - | - | - | - | - | - | - | - | - | - | - | - | - | - | - | - | - | - | - | - | - | - | - | - | - | - | - | - | - | - | - | - | - | - | - | - | - | - | - | - | - | - | - | - | - | - | - | - | - | - | - | - | - | - | - | - | - | - | - | - | - | - | - | - | - | - | - | - | - | - | - | - | - | - | - | - | - | - | - | - | - | - | - | - | - | - | - | - | - | - | - | - | - | - | - | - | - | - | - | - | - | - | - | - | - | - | - | - | - | - | - | - | - | - | - | - | - | - | - | - | - | - | - | - | - | - | - | - | - | - | - | - | - | - | - | - | - | - | - | - | - | - | - | - | - | - | - | - | - | - | - | - | - | - | - | - | - | - | - | - | - | - | - | - | - | - | - | - | - | - | - | - | - | - | - | - | - | - | - | - | - | - | - | - | - | - | - | - | - | - | - | - | - | - | - | - | - | - | - | - | - | - | - | - | - | - | - | - | - | - | - | - | - | - | - | - | - | - | - | - | - | - | - | - | - | - | - | - | - | - | - | - | - | - | - | - | - | - | - | - | - | - | - | - | - | - | - | - | - | - | - | - | - | - | - | - | - | - | - | - | - | - | - | - | - | - | - | - | - | - | - | - | - | - | - | - | - | - | - | - | - | - | - | - | - | - | - | - | - | - | - | - | - | - | - | - | - | - | - | - | - | - | - | - | - | - | - | - | - | - | - | - | - | - | - | - | - | - | - | - | - | - | - | - | - | - | - | - | - | - | - | - | - | - | - | - | - | - | - | - | - | - | - | - | - | - | - | - | - | - | - | - | - | - | - | - | - | - | - | - | - | - | - | - | - | - | - | - | - | - | - | - | - | - | - | - | - | - | - | - | - | - | - | - | - | - | - | - | - | - | - | - | - | - | - | - | - | - | - | - | - | - | - | - | - | - | - | - | - | - | - | - | - | - | - | - | - | - | - | - | - | - | - | - | - | - | - | - | - | - | - | - | - | - | - | - | - | - | - | - | - | - | - | - | - | - | - | - | - | - | - | - | - | - | - | - | - | - | - | - | - | - | - | - | - | - | - | - | - | - | - | - | - | - | - | - | - | - | - | - | - | - | - | - | - | - | - | - | - | - | - | - | - | - | - | - | - | - | - | - | - | - | - | - | - | - | - | - | - | - | - | - | - | - | - | - | - | - | - | - | - | - | - | - | - | - | - | - | - | - | - | - | - | - | - | - | - | - | - | - | - | - | - | - | - | - | - | - | - | - | - | - | - | - | - | - | - | - | - | - | - | - | - | - | - | - | - | - | - | - | - | - | - | - | - | - | - | - | - | - | - | - | - | - | - | - | - | - | - | - | - | - | - | - | - | - | - | - | - | - | - | - | - | - | - | - | - | - | - | - | - | - | - | - | - | - | - | - | - | - | - | - | - | - | - | - | - | - | - | - | - | - | - | - | - | - | - | - | - | - | - | - | - | - | - | - | - | - | - | - | - | - | - | - | - | - | - | - | - | - | - | - | - | - | - | - | - | - | - | - | - | - | - | - | - | - | - | - | - | - | - | - | - | - | - | - | - | - | - | - | - | - | - | - | - | - | - | - | - | - | - | - | - | - | - | - | - | - | - | - | - | - | - | - | - | - | - | - | - | - | - | - | - | - | - | - | - | - | - | - | - | - | - | - | - | - | - | - | - | - | - | - | - | - | - | - | - | - | - | - | - | - | - | - | - | - | - | - | - | - | - | - | - | - | - | - | - | - | - | - | - | - | - | - | - | - | - | - | - | - | - | - | - | - | - | - | - | - | - | - | - | - | - | - | - | - | - | - | - | - | - | - | - | - | - | - | - | - | - | - | - | - | - | - | - | - | - | - | - | - | - | - | - | - | - | - | - | - | - | - | - | - | - | - | - | - | - | - | - | - | - | - | - | - | - | - | - | - | - | - | - | - | - | - | - | - | - | - | - | - | - | - | - | - | - | - | - | - | - | - | - | - | - | - | - | - | - | - | - | - | - | - | - | - | - | - | - | - | - | - | - | - | - | - | - | - | - | - | - | - | - | - | - | - | - | - | - | - | - | - | - | - | - | - | - | - | - | - | - | - | - | - | - | - | - | - | - | - | - | - | - | - | - | - | - | - | - | - | - | - | - | - | - | - | - | - | - | - | - | - | - | - | - | - | - | - | - | - | - | - | - | - | - | - | - | - | - | - | - | - | - | - | - | - | - | - | - | - | - | - | - | - | - | - | - | - | - | - | - | - | - | - | - | - | - | - | - | - | - | - | - | - | - | - | - | - | - | - | - | - | - | - | - | - | - | - | - | - | - | - | - | - | - | - | - | - | - | - | - | - | - | - | - | - | - | - | - | - | - | - | - | - | - | - | - | - | - | - | - | - | - | - | - | - | - | - | - | - | - | - | - | - | - | - | - | - | - | - | - | - | - | - | - | - | - | - | - | - | - | - | - | - | - | - | - | - | - | - | - | - | - | - | - | - | - | - | - | - | - | - | - | - | - | - | - | - | - | - | - | - | - | - | - | - | - | - | - | - | - | - | - | - | - | - | - | - | - | - | - | - | - | - | - | - | - | - | - | - | - | - | - | - | - | - | - | - | - | - | - | - | - | - | - | - | - | - | - | - | - | - | - | - | - | - | - | - | - | - | - | - | - | - | - | - | - | - | - | - | - | - | - | - | - | - | - | - | - | - | - | - | - | - | - | - | - | - | - | - | - | - | - | - | - | - | - | - | - | - | - | - | - | - | - | - | - | - | - | - | - | - | - | - | - | - | - | - | - | - | - | - | - | - | - | - | - | - | - | - | - | - | - | - | - | - | - | - | - | - | - | - | - | - | - | - | - | - | - | - | - | - | - | - | - | - | - | - | - | - | - | - | - | - | - | - | - | - | - | - | - | - | - | - | - | - | - | - | - | - | - | - | - | - | - | - | - | - | - | - | - | - | - | - | - | - | - | - | - | - | - | - | - | - | - | - | - | - | - | - | - | - | - | - | - | - | - | - | - | - | - | - | - | - | - | - | - | - | - | - | - | - | - | - | - | - | - | - | - | - | - | - | - | - | - | - | - | - | - | - | - | - | - | - | - | - | - | - | - | - | - | - | - | - | - | - | - | - | - | - | - | - | - | - | - | - | - | - | - | - | - | - | - | - | - | - | - | - | - | - | - | - | - | - | - | - | - | - | - | - | - | - | - | - | - | - | - | - | - | - | - | - | - | - | - | - | - | - | - | - | - | - | - | - | - | - | - | - | - | - | - | - | - | - | - | - | - | - | - | - | - | - | - | - | - | - | - | - | - | - | - | - | - | - | - | - | - | - | - | - | - | - | - | - | - | - | - | - | - | - | - | - | - | - | - | - | - | - | - | - | - | - | - | - | - | - | - | - | - | - | - | - | - | - | - | - | - | - | - | - | - | - | - | - | - | - | - | - | - | - | - | - | - | - | - | - | - | - | - | - | - | - | - | - | - | - | - | - | - | - | - | - | - | - | - | - | - | - | - | - | - | - | - | - | - | - | - | - | - | - | - | - | - | - | - | - | - | - | - | - | - | - | - | - | - | - | - | - | - | - | - | - | - | - | - | - | - | - | - | - | - | - | - | - | - | - | - | - | - | - | - | - | - | - | - | - | - | - | - | - | - | - | - | - | - | - | - | - | - | - | - | - | - | - | - | - | - | - | - | - | - | - | - | - | - | - | - | - | - | - | - | - | - | - | - | - | - | - | - | - | - | - | - | - | - | - | - | - | - | - | - | - | - | - | - | - | - | - | - | - | - | - | - | - | - | - | - | - | - | - | - | - | - | - | - | - | - | - | - | - | - | - | - | - | - | - | - | - | - | - | - | - | - | - | - | - | - | - | - | - | - | - | - | - | - | - | - | - | - | - | - | - | - | - | - | - | - | - | - | - | - | - | - | - | - | - | - | - | - | - | - | - | - | - | - | - | - | - | - | - | - | - | - | - | - | - | - | - | - | - | - | - | - | - | - | - | - | - | - | - | - | - | - | - | - | - | - | - | - | - | - | - | - | - | - | - | - | - | - | - | - | - | - | - | - | - | - | - | - | - | - | - | - | - | - | - | - | - | - | - | - | - | - | - | - | - | - | - | - | - | - | - | - | - | - | - | - | - | - | - | - | - | - | - | - | - | - | - | - | - | - | - | - | - | - | - | - | - | - | - | - | - | - | - | - | - | - | - | - | - | - | - | - | - | - | - | - | - | - | - | - | - | - | - | - | - | - | - | - | - | - | - | - | - | - | - | - | - | - | - | - | - | - | - | - | - | - | - | - | - | - | - | - | - | - | - | - | - | - | - | - | - | - | - | - | - | - | - | - | - | - | - | - | - | - | - | - | - | - | - | - | - | - | - | - | - | - | - | - | - | - | - | - | - | - | - | - | - | - | - | - | - | - | - | - | - | - | - | - | - | - | - | - | - | - | - | - | - | - | - | - | - | - | - | - | - | - | - | - | - | - | - | - | - | - | - | - | - | - | - | - | - | - | - | - | - | - | - | - | - | - | - | - | - | - | - | - | - | - | - | - | - | - | - | - | - | - | - | - | - | - | - | - | - | - | - | - | - | - | - | - | - | - | - | - | - | - | - | - | - | - | - | - | - | - | - | - | - | - | - | - | - | - | - | - | - | - | - | - | - | - | - | - | - | - | - | - | - | - | - | - | - | - | - | - | - | - | - | - | - | - | - | - | - | - | - | - | - | - | - | - | - | - | - | - | - | - | - | - | - | - | - | - | - | - | - | - | - | - | - | - | - | - | - | - | - | - | - | - | - | - | - | - |
| BIS d=0 cl.18 sym=1 env=1 | - | - | - | - | - | - | - | - | - | - | - | - | - | - | - | - | - | - | - | - | - | - | - | - | - | - | - | - | - | - | - | - | - | - | - | - | - | - | - | - | - | - | - | - | - | - | - | - | - | - | - | - | - | - | - | - | - | - | - | - | - | - | - | - | - | - | - | - | - | - | - | - | - | - | - | - | - | - | - | - | - | - | - | - | - | - | - | - | - | - | - | - | - | - | - | - | - | - | - | - | - | - | - | - | - | - | - | - | - | - | - | - | - | - | - | - | - | - | - | - | - | - | - | - | - | - | - | - | - | - | - | - | - | - | - | - | - | - | - | - | - | - | - | - | - | - | - | - | - | - | - | - | - | - | - | - | - | - | - | - | - | - | - | - | - | - | - | - | - | - | - | - | - | - | - | - | - | - | - | - | - | - | - | - | - | - | - | - | - | - | - | - | - | - | - | - | - | - | - | - | - | - | - | - | - | - | - | - | - | - | - | - | - | - | - | - | - | - | - | - | - | - | - | - | - | - | - | - | - | - | - | - | - | - | - | - | - | - | - | - | - | - | - | - | - | - | - | - | - | - | - | - | - | - | - | - | - | - | - | - | - | - | - | - | - | - | - | - | - | - | - | - | - | - | - | - | - | - | - | - | - | - | - | - | - | - | - | - | - | - | - | - | - | - | - | - | - | - | - | - | - | - | - | - | - | - | - | - | - | - | - | - | - | - | - | - | - | - | - | - | - | - | - | - | - | - | - | - | - | - | - | - | - | - | - | - | - | - | - | - | - | - | - | - | - | - | - | - | - | - | - | - | - | - | - | - | - | - | - | - | - | - | - | - | - | - | - | - | - | - | - | - | - | - | - | - | - | - | - | - | - | - | - | - | - | - | - | - | - | - | - | - | - | - | - | - | - | - | - | - | - | - | - | - | - | - | - | - | - | - | - | - | - | - | - | - | - | - | - | - | - | - | - | - | - | - | - | - | - | - | - | - | - | - | - | - | - | - | - | - | - | - | - | - | - | - | - | - | - | - | - | - | - | - | - | - | - | - | - | - | - | - | - | - | - | - | - | - | - | - | - | - | - | - | - | - | - | - | - | - | - | - | - | - | - | - | - | - | - | - | - | - | - | - | - | - | - | - | - | - | - | - | - | - | - | - | - | - | - | - | - | - | - | - | - | - | - | - | - | - | - | - | - | - | - | - | - | - | - | - | - | - | - | - | - | - | - | - | - | - | - | - | - | - | - | - | - | - | - | - | - | - | - | - | - | - | - | - | - | - | - | - | - | - | - | - | - | - | - | - | - | - | - | - | - | - | - | - | - | - | - | - | - | - | - | - | - | - | - | - | - | - | - | - | - | - | - | - | - | - | - | - | - | - | - | - | - | - | - | - | - | - | - | - | - | - | - | - | - | - | - | - | - | - | - | - | - | - | - | - | - | - | - | - | - | - | - | - | - | - | - | - | - | - | - | - | - | - | - | - | - | - | - | - | - | - | - | - | - | - | - | - | - | - | - | - | - | - | - | - | - | - | - | - | - | - | - | - | - | - | - | - | - | - | - | - | - | - | - | - | - | - | - | - | - | - | - | - | - | - | - | - | - | - | - | - | - | - | - | - | - | - | - | - | - | - | - | - | - | - | - | - | - | - | - | - | - | - | - | - | - | - | - | - | - | - | - | - | - | - | - | - | - | - | - | - | - | - | - | - | - | - | - | - | - | - | - | - | - | - | - | - | - | - | - | - | - | - | - | - | - | - | - | - | - | - | - | - | - | - | - | - | - | - | - | - | - | - | - | - | - | - | - | - | - | - | - | - | - | - | - | - | - | - | - | - | - | - | - | - | - | - | - | - | - | - | - | - | - | - | - | - | - | - | - | - | - | - | - | - | - | - | - | - | - | - | - | - | - | - | - | - | - | - | - | - | - | - | - | - | - | - | - | - | - | - | - | - | - | - | - | - | - | - | - | - | - | - | - | - | - | - | - | - | - | - | - | - | - | - | - | - | - | - | - | - | - | - | - | - | - | - | - | - | - | - | - | - | - | - | - | - | - | - | - | - | - | - | - | - | - | - | - | - | - | - | - | - | - | - | - | - | - | - | - | - | - | - | - | - | - | - | - | - | - | - | - | - | - | - | - | - | - | - | - | - | - | - | - | - | - | - | - | - | - | - | - | - | - | - | - | - | - | - | - | - | - | - | - | - | - | - | - | - | - | - | - | - | - | - | - | - | - | - | - | - | - | - | - | - | - | - | - | - | - | - | - | - | - | - | - | - | - | - | - | - | - | - | - | - | - | - | - | - | - | - | - | - | - | - | - | - | - | - | - | - | - | - | - | - | - | - | - | - | - | - | - | - | - | - | - | - | - | - | - | - | - | - | - | - | - | - | - | - | - | - | - | - | - | - | - | - | - | - | - | - | - | - | - | - | - | - | - | - | - | - | - | - | - | - | - | - | - | - | - | - | - | - | - | - | - | - | - | - | - | - | - | - | - | - | - | - | - | - | - | - | - | - | - | - | - | - | - | - | - | - | - | - | - | - | - | - | - | - | - | - | - | - | - | - | - | - | - | - | - | - | - | - | - | - | - | - | - | - | - | - | - | - | - | - | - | - | - | - | - | - | - | - | - | - | - | - | - | - | - | - | - | - | - | - | - | - | - | - | - | - | - | - | - | - | - | - | - | - | - | - | - | - | - | - | - | - | - | - | - | - | - | - | - | - | - | - | - | - | - | - | - | - | - | - | - | - | - | - | - | - | - | - | - | - | - | - | - | - | - | - | - | - | - | - | - | - | - | - | - | - | - | - | - | - | - | - | - | - | - | - | - | - | - | - | - | - | - | - | - | - | - | - | - | - | H | x | x | x | x | x | x | x | H | x | x | x | x | x | x | x | x | x | x | x | x | x | x | x | x | x | x | x | x | x | x | x | x | x | x | x | x | x | x | x | x | x | x | x | x | x | x | x | H | x | x | x | x | x | x | x | x | x | x | x | H | - | - | - | - | - | - | - | - | - | - | - | - | - | - | - | - | - | - | - | - | - | - | - | - | - | - | - | - | - | - | - | - | - | - | - | - | - | - | - | - | - | - | - | - | - | - | - | - | - | - | - | - | - | - | - | - | - | - | - | - | - | - | - | - | - | - | - | - | - | - | - | - | - | - | - | - | - | - | - | - | - | - | - | - | - | - | - | - | - | - | - | - | - | - | - | - | - | - | - | - | - | - | - | - | - | - | - | - | - | - | - | - | - | - | - | - | - | - | - | - | - | - | - | - | - | - | - | - | - | - | - | - | - | - | - | - | - | - | - | - | - | - | - | - | - | - | - | - | - | - | - | - | - | - | - | - | - | - | - | - | - | - | - | - | - | - | - | - | - | - | - | - | - | - | - | - | - | - | - | - | - | - | - | - | - | - | - | - | - | - | - | - | - | - | - | - | - | - | - | - | - | - | - | - | - | - | - | - | - | - | - | - | - | - | - | - | - | - | - | - | - | - | - | - | - | - | - | - | - | - | - | - | - | - | - | - | - | - | - | - | - | - | - | - | - | - | - | - | - | - | - | - | - | - | - | - | - | - | - | - | - | - | - | - | - | - | - | - | - | - | - | - | - | - | - | - | - | - | - | - | - | - | - | - | - | - | - | - | - | - | - | - | - | - | - | - | - | - | - | - | - | - | - | - | - | - | - | - | - | - | - | - | - | - | - | - | - | - | - | - | - | - | - | - | - | - | - | - | - | - | - | - | - | - | - | - | - | - | - | - | - | - | - | - | - | - | - | - | - | - | - | - | - | - | - | - | - | - | - | - | - | - | - | - | - | - | - | - | - | - | - | - | - | - | - | - | - | - | - | - | - | - | - | - | - | - | - | - | - | - | - | - | - | - | - | - | - | - | - | - | - | - | - | - | - | - | - | - | - | - | - | - | - | - | - | - | - | - | - | - | - | - | - | - | - | - | - | - | - | - | - | - | - | - | - | - | - | - | - | - | - | - | - | - | - | - | - | - | - | - | - | - | - | - | - | - | - | - | - | - | - | - | - | - | - | - | - | - | - | - | - | - | - | - | - | - | - | - | - | - | - | - | - | - | - | - | - | - | - | - | - | - | - | - | - | - | - | - | - | - | - | - | - | - | - | - | - | - | - | - | - | - | - | - | - | - | - | - | - | - | - | - | - | - | - | - | - | - | - | - | - | - | - | - | - | - | - | - | - | - | - | - | - | - | - | - | - | - | - | - | - | - | - | - | - | - | - | - | - | - | - | - | - | - | - | - | - | - | - | - | - | - | - | - | - | - | - | - | - | - | - | - | - | - | - | - | - | - | - | - | - | - | - | - | - | - | - | - | - | - | - | - | - | - | - | - | - | - | - | - | - | - | - | - | - | - | - | - | - | - | - | - | - | - | - | - | - | - | - | - | - | - | - | - | - | - | - | - | - | - | - | - | - | - | - | - | - | - | - | - | - | - | - | - | - | - | - | - | - | - | - | - | - | - | - | - | - | - | - | - | - | - | - | - | - | - | - | - | - | - | - | - | - | - | - | - | - | - | - | - | - | - | - | - | - | - | - | - | - | - | - | - | - | - | - | - | - | - | - | - | - | - | - | - | - | - | - | - | - | - | - | - | - | - | - | - | - | - | - | - | - | - | - | - | - | - | - | - | - | - | - | - | - | - | - | - | - | - | - | - | - | - | - | - | - | - | - | - | - | - | - | - | - | - | - | - | - | - | - | - | - | - | - | - | - | - | - | - | - | - | - | - | - | - | - | - | - | - | - | - | - | - | - | - | - | - | - | - | - | - | - | - | - | - | - | - | - | - | - | - | - | - | - | - | - | - | - | - | - | - | - | - | - | - | - | - | - | - | - | - | - | - | - | - | - | - | - | - | - | - | - | - | - | - | - | - | - | - | - | - | - | - | - | - | - | - | - | - | - | - | - | - | - | - | - | - | - | - | - | - | - | - | - | - | - | - | - | - | - | - | - | - | - | - | - | - | - | - | - | - | - | - | - | - | - | - | - | - | - | - | - | - | - | - | - | - | - | - | - | - | - | - | - | - | - | - | - | - | - | - | - | - | - | - | - | - | - | - | - | - | - | - | - | - | - | - | - | - | - | - | - | - | - | - | - | - | - | - | - | - | - | - | - | - | - | - | - | - | - | - | - | - | - | - | - | - | - | - | - | - | - | - | - | - | - | - | - | - | - | - | - | - | - | - | - | - | - | - | - | - | - | - | - | - | - | - | - | - | - | - | - | - | - | - | - | - | - | - | - | - | - | - | - | - | - | - | - | - | - | - | - | - | - | - | - | - | - | - | - | - | - | - | - | - | - | - | - | - | - | - | - | - | - | - | - | - | - | - | - | - | - | - | - | - | - | - | - | - | - | - | - | - | - | - | - | - | - | - | - | - | - | - | - | - | - | - | - | - | - | - | - | - | - | - | - | - | - | - | - | - | - | - | - | - | - | - | - | - | - | - | - | - | - | - | - | - | - | - | - | - | - | - | - | - | - | - | - | - | - | - | - | - | - | - | - | - | - | - | - | - | - | - | - | - | - | - | - | - | - | - | - | - | - | - | - | - | - | - | - | - | - | - | - | - | - | - | - | - | - | - | - | - | - | - | - | - | - | - | - | - | - | - | - | - | - | - | - | - | - | - | - | - | - | - | - | - | - | - | - | - | - | - | - | - | - | - | - | - | - | - | - | - | - | - | - | - | - | - | - | - | - | - | - | - | - | - | - | - | - | - | - | - | - | - | - | - | - | - | - | - | - | - | - | - | - | - | - | - | - | - | - | - | - | - | - | - | - | - | - | - | - | - | - | - | - | - | - | - | - | - | - | - | - | - | - | - | - | - | - | - | - | - | - | - | - | - | - | - | - | - | - | - | - | - | - | - | - | - | - | - | - | - | - | - | - | - | - | - | - | - | - | - | - | - | - | - | - | - | - | - | - | - | - | - | - | - | - | - | - | - | - | - | - | - | - | - | - | - | - | - | - | - | - | - | - | - | - | - | - | - | - | - | - | - | - | - | - | - | - | - | - | - | - | - | - | - | - | - | - | - | - | - | - | - | - | - | - | - | - | - | - | - | - | - | - | - | - | - | - | - | - | - | - | - | - | - | - | - | - | - | - | - | - | - | - | - | - | - | - | - | - | - | - | - | - | - | - | - | - | - | - | - | - | - | - | - | - | - | - | - | - | - | - | - | - | - | - | - | - | - | - | - | - | - | - | - | - | - | - | - | - | - | - | - | - | - | - | - | - | - | - | - | - | - | - | - | - | - | - | - | - | - | - | - | - | - | - | - | - | - | - | - | - | - | - | - | - | - | - | - | - | - | - | - | - | - | - | - | - | - | - | - | - | - | - | - | - | - | - | - | - | - | - | - | - | - | - | - | - | - | - | - | - | - | - | - | - | - | - | - | - | - | - | - | - | - | - | - | - | - | - | - | - | - | - | - | - | - | - | - | - | - | - | - | - | - | - | - | - | - | - | - | - | - | - | - | - | - | - | - | - | - | - | - | - | - | - | - | - | - | - | - | - | - | - | - | - | - | - | - | - | - | - | - | - | - | - | - | - | - | - | - | - | - | - | - | - | - | - | - | - | - | - | - | - | - | - | - | - | - | - | - | - | - | - | - | - | - | - | - | - | - | - | - | - | - | - | - | - | - | - | - | - | - | - | - | - | - | - | - | - | - | - | - | - | - | - | - | - | - | - | - | - | - | - | - | - | - | - | - | - | - | - | - | - | - | - | - | - | - | - | - | - | - | - | - | - | - | - | - | - | - | - | - | - | - | - | - | - | - | - | - | - | - | - | - | - | - | - | - | - | - | - | - | - | - | - | - | - | - | - | - | - | - | - | - | - | - | - | - | - | - | - | - | - | - | - | - | - | - | - | - | - | - | - | - | - | - | - | - | - | - | - | - | - | - | - | - | - | - | - | - | - | - | - | - | - | - | - | - | - | - | - | - | - | - | - | - | - | - | - | - | - | - | - | - | - | - | - | - | - | - | - | - | - | - | - | - | - | - | - | - | - | - | - | - | - | - | - | - | - | - | - | - | - | - | - | - | - | - | - | - | - | - | - | - | - | - | - | - | - | - | - | - | - | - | - | - | - | - | - | - | - | - | - | - | - | - | - | - | - | - | - | - | - | - | - | - | - | - | - | - | - | - | - | - | - | - | - | - | - | - | - | - | - | - | - | - | - | - | - | - | - | - | - | - | - | - | - | - | - | - | - | - | - | - | - | - | - | - | - | - | - | - | - | - | - | - | - | - | - | - | - | - | - | - | - | - | - | - | - | - | - | - | - | - | - | - | - | - | - | - | - | - | - | - | - | - | - | - | - | - | - | - | - | - | - | - | - | - | - | - | - | - | - | - | - | - | - | - | - | - | - | - | - | - | - | - | - | - | - | - | - | - | - | - | - | - | - | - | - | - | - | - | - | - | - | - | - | - | - | - | - | - | - | - | - | - | - | - | - | - | - | - | - | - | - | - | - | - | - | - | - | - | - | - | - | - | - | - | - | - | - | - | - | - | - | - | - | - | - | - | - | - | - | - | - | - | - | - | - | - | - | - | - | - | - | - | - | - | - | - | - | - | - | - | - | - | - | - | - | - | - | - | - | - | - | - | - | - | - | - | - | - | - | - | - | - | - | - | - | - | - | - | - | - | - | - | - | - | - | - | - | - | - | - | - | - | - | - | - | - | - | - | - | - | - | - | - | - | - | - | - | - | - | - | - | - | - | - | - | - | - | - | - | - | - | - | - | - | - | - | - | - | - | - | - | - | - | - | - | - | - | - | - | - | - | - | - | - | - | - | - | - | - | - | - | - | - | - | - | - | - | - | - | - | - | - | - | - | - | - | - | - | - | - | - | - | - | - | - | - | - | - | - | - | - | - | - | - | - | - | - | - | - | - | - | - | - | - | - | - | - | - | - | - | - | - | - | - | - | - | - | - | - | - | - | - | - | - | - | - | - | - | - | - | - | - | - | - | - | - | - | - | - | - | - | - | - | - | - | - | - | - | - | - | - | - | - | - | - | - | - | - | - | - | - | - | - | - | - | - | - | - | - | - | - | - | - | - | - | - | - | - | - | - | - | - | - | - | - | - | - | - | - | - | - | - | - | - | - | - | - | - | - | - | - | - | - | - | - | - | - | - | - | - | - | - | - | - | - | - | - | - | - | - | - | - | - | - | - | - | - | - | - | - | - | - | - | - | - | - | - | - | - | - | - | - | - | - | - | - | - | - | - | - | - | - | - | - | - | - | - | - | - | - | - | - | - | - | - | - | - | - | - | - | - | - | - | - | - | - | - | - | - | - | - | - | - | - | - | - | - | - | - | - | - | - | - | - | - | - | - | - | - | - | - | - | - | - | - | - | - | - | - | - | - | - | - | - | - | - | - | - | - | - | - | - | - | - | - | - | - | - | - | - | - | - | - | - | - | - | - | - | - | - | - | - | - | - | - | - | - | - | - | - | - | - | - | - | - | - | - | - | - | - | - | - | - | - | - | - | - | - | - | - | - | - | - | - | - | - | - | - | - | - | - | - | - | - | - | - | - | - | - | - | - | - | - | - | - | - | - | - | - | - | - | - | - | - | - | - | - | - | - | - | - | - | - | - | - | - | - | - | - | - | - | - | - | - | - | - | - | - | - | - | - | - | - | - | - | - | - | - | - | - | - | - | - | - | - | - | - | - | - | - | - | - | - | - | - | - | - | - | - | - | - | - | - | - | - | - | - | - | - | - | - | - | - | - | - | - | - | - | - | - | - | - | - | - | - | - | - | - | - | - | - | - | - | - | - | - | - | - | - | - | - | - | - | - | - | - | - | - | - | - | - | - | - | - | - | - | - | - | - | - | - | - | - | - | - | - | - | - | - | - | - | - | - | - | - | - | - | - | - | - | - | - | - | - | - | - | - | - | - | - | - | - | - | - | - | - | - | - | - | - | - | - | - | - | - | - | - | - | - | - | - | - | - | - | - | - | - | - | - | - | - | - | - | - | - | - | - | - | - | - | - | - | - | - | - | - | - | - | - | - | - | - | - | - | - | - | - | - | - | - | - | - | - | - | - | - | - | - | - | - | - | - | - | - | - | - | - | - | - | - | - | - | - | - | - | - | - | - | - | - | - | - | - | - | - | - | - | - | - | - | - | - | - | - | - | - | - | - | - | - | - | - | - | - | - | - | - | - | - | - | - | - | - | - | - | - | - | - | - | - | - | - | - | - | - | - | - | - | - | - | - | - | - | - | - | - | - | - | - | - | - | - | - | - | - | - | - | - | - | - | - | - | - | - | - | - | - | - | - | - | - | - | - | - | - | - | - | - | - | - | - | - | - | - | - | - | - | - | - | - | - | - | - | - | - | - | - | - | - | - | - | - | - | - | - | - | - | - | - | - | - | - | - | - | - | - | - | - | - | - | - | - | - | - | - | - | - | - | - | - | - | - | - | - | - | - | - | - | - | - | - | - | - | - | - | - | - | - | - | - | - | - | - | - | - | - | - | - | - | - | - | - | - | - | - | - | - | - | - | - | - | - | - | - | - | - | - | - | - | - | - | - | - | - | - | - | - | - | - | - | - | - | - | - | - | - | - | - | - | - | - | - | - | - | - | - | - | - | - | - | - | - | - | - | - | - | - | - | - | - | - | - | - | - | - | - | - | - | - | - | - | - | - | - | - | - | - | - | - | - | - | - | - | - | - | - | - | - | - | - | - | - | - | - | - | - | - | - | - | - | - | - | - | - | - | - | - | - | - | - | - | - | - | - | - | - | - | - | - | - | - | - | - | - | - | - | - | - | - | - | - | - | - | - | - | - | - | - | - | - | - | - | - | - | - | - | - | - | - | - | - | - | - | - | - | - | - | - | - | - | - | - | - | - | - | - | - | - | - | - | - | - | - | - | - | - | - | - | - | - | - | - | - | - | - | - | - | - | - | - | - | - | - | - | - | - | - | - | - | - | - | - | - | - | - | - | - | - | - | - | - | - | - | - | - | - | - | - | - | - | - | - | - | - | - | - | - | - | - | - | - | - | - | - | - | - | - | - | - | - | - | - | - | - | - | - | - | - | - | - | - | - | - | - | - | - | - | - | - | - | - | - | - | - | - | - | - | - | - | - | - | - | - | - | - | - | - | - | - | - | - | - | - | - | - | - | - | - | - | - | - | - | - | - | - | - | - | - | - | - | - | - | - | - | - | - | - | - | - | - | - | - | - | - | - | - | - | - | - | - | - | - | - | - | - | - | - | - | - | - | - | - | - | - | - | - | - | - | - | - | - | - | - | - | - | - | - | - | - | - | - | - | - | - | - | - | - | - | - | - | - | - | - | - | - | - | - | - | - | - | - | - | - | - | - | - | - | - | - | - | - | - | - | - | - | - | - | - | - | - | - | - | - | - | - | - | - | - | - | - | - | - | - | - | - | - | - | - | - | - | - | - | - | - | - | - | - | - | - | - | - | - | - | - | - | - | - | - | - | - | - | - | - | - | - | - | - | - | - | - | - | - | - | - | - | - | - | - | - | - | - | - | - | - | - | - | - | - | - | - | - | - | - | - | - | - | - | - | - | - | - | - | - | - | - | - | - | - | - | - | - | - | - | - | - | - | - | - | - | - | - | - | - | - | - | - | - | - | - | - | - | - | - | - | - | - | - | - | - | - | - | - | - | - | - | - | - | - | - | - | - | - | - | - | - | - | - | - | - | - | - | - | - | - | - | - | - | - | - | - | - | - | - | - | - | - | - | - | - | - | - | - | - | - | - | - | - | - | - | - | - | - | - | - | - | - | - | - | - | - | - | - | - | - | - | - | - | - | - | - | - | - | - | - | - | - | - | - | - | - | - | - | - | - | - | - | - | - | - | - | - | - | - | - | - | - | - | - | - | - | - | - | - | - | - | - | - | - | - | - | - | - | - | - | - | - | - | - | - | - | - | - | - | - | - | - | - | - | - | - | - | - | - | - | - | - | - | - | - | - | - | - | - | - | - | - | - | - | - | - | - | - | - | - | - | - | - | - | - | - | - | - | - | - | - | - | - | - | - | - | - | - | - | - | - | - | - | - | - | - | - | - | - | - | - | - | - | - | - | - | - | - | - | - | - | - | - | - | - | - | - | - | - | - | - | - | - | - | - | - | - | - | - | - | - | - | - | - | - | - | - | - | - | - | - | - | - | - | - | - | - | - | - | - | - | - | - | - | - | - | - | - | - | - | - | - | - | - | - | - | - | - | - | - | - | - | - | - | - | - | - | - | - | - | - | - | - | - | - | - | - | - | - | - | - | - | - | - | - | - | - | - | - | - | - | - | - | - | - | - | - | - | - | - | - | - | - | - | - | - | - | - | - | - | - | - | - | - | - | - | - | - | - | - | - | - | - | - | - | - | - | - | - | - | - | - | - | - | - | - | - | - | - | - | - | - | - | - | - | - | - | - | - | - | - | - | - | - | - | - | - | - | - | - | - | - | - | - | - | - | - | - | - | - | - | - | - | - | - | - | - | - | - | - | - | - | - | - | - | - | - | - | - | - | - | - | - | - | - | - | - | - | - | - | - | - | - | - | - | - | - | - | - | - | - | - | - | - | - | - | - | - | - | - | - | - | - | - | - | - | - | - | - | - | - | - | - | - | - | - | - | - | - | - | - | - | - | - | - | - | - | - | - | - | - | - | - | - | - | - | - | - | - | - | - | - | - | - | - | - | - | - | - | - | - | - | - | - | - | - | - | - | - | - | - | - | - | - | - | - | - | - | - | - | - | - | - | - | - | - | - | - | - | - | - | - | - | - | - | - | - | - | - | - | - | - | - | - | - | - | - | - | - | - | - | - | - | - | - | - | - | - | - | - | - | - | - | - | - | - | - | - | - | - | - | - | - | - | - | - | - | - | - | - | - | - | - | - | - | - | - | - | - | - | - | - | - | - | - | - | - | - | - | - | - | - | - | - | - | - | - | - | - | - | - | - | - | - | - | - | - | - | - | - | - | - | - | - | - | - | - | - | - | - | - | - | - | - | - | - | - | - | - | - | - | - | - | - | - | - | - | - | - | - | - | - | - | - | - | - | - | - | - | - | - | - | - | - | - | - | - | - | - | - | - | - | - | - | - | - | - | - | - | - | - | - | - | - | - | - | - | - | - | - | - | - | - | - | - | - | - | - | - | - | - | - | - | - | - | - | - | - | - | - | - | - | - | - | - | - | - | - | - | - | - | - | - | - | - | - | - | - | - | - | - | - | - | - | - | - | - |
| BIS d=0 cl.14 sym=1 env=1 | - | - | - | - | - | - | - | - | - | - | - | - | - | - | - | - | - | - | - | - | - | - | - | - | - | - | - | - | - | - | - | - | - | - | - | - | - | - | - | - | - | - | - | - | - | - | - | - | - | - | - | - | - | - | - | - | - | - | - | - | - | - | - | - | - | - | - | - | - | - | - | - | - | - | - | - | - | - | - | - | - | - | - | - | - | - | - | - | - | - | - | - | - | - | - | - | - | - | - | - | - | - | - | - | - | - | - | - | - | - | - | - | - | - | - | - | - | - | - | - | - | - | - | - | - | - | - | - | - | - | - | - | - | - | - | - | - | - | - | - | - | - | - | - | - | - | - | - | - | - | - | - | - | - | - | - | - | - | - | - | - | - | - | - | - | - | - | - | - | - | - | - | - | - | - | - | - | - | - | - | - | - | - | - | - | - | - | - | - | - | - | - | - | - | - | - | - | - | - | - | - | - | - | - | - | - | - | - | - | - | - | - | - | - | - | - | - | - | - | - | - | - | - | - | - | - | - | - | - | - | - | - | - | - | - | - | - | - | - | - | - | - | - | - | - | - | - | - | - | - | - | - | - | - | - | - | - | - | - | - | - | - | - | - | - | - | - | - | - | - | - | - | - | - | - | - | - | - | - | - | - | - | - | - | - | - | - | - | - | - | - | - | - | - | - | - | - | - | - | - | - | - | - | - | - | - | - | - | - | - | - | - | - | - | - | - | - | - | - | - | - | - | - | - | - | - | - | - | - | - | - | - | - | - | - | - | - | - | - | - | - | - | - | - | - | - | - | - | - | - | - | - | - | - | - | - | - | - | - | - | - | - | - | - | - | - | - | - | - | - | - | - | - | - | - | - | - | - | - | - | - | - | - | - | - | - | - | - | - | - | - | - | - | - | - | - | - | - | - | - | - | - | - | - | - | - | - | - | - | - | - | - | - | - | - | - | - | - | - | - | - | - | - | - | - | - | - | - | - | - | - | - | - | - | - | - | - | - | - | - | - | - | - | - | - | - | - | - | - | - | - | - | - | - | - | - | - | - | - | - | - | - | - | - | - | - | - | - | - | - | - | - | - | - | - | - | - | - | - | - | - | - | - | - | - | - | - | - | - | - | - | - | - | - | - | - | - | - | - | - | - | - | - | - | - | - | - | - | - | - | - | - | - | - | - | - | - | - | - | - | - | - | - | - | - | - | - | - | - | - | - | - | - | - | - | - | - | - | - | - | - | - | - | - | - | - | - | - | - | - | - | - | - | - | - | - | - | - | - | - | - | - | - | - | - | - | - | - | - | - | - | - | - | - | - | - | - | - | - | - | - | - | - | - | - | - | - | - | - | - | - | - | - | - | - | - | - | - | - | - | - | - | - | - | - | - | - | - | - | - | - | - | - | - | - | - | - | - | - | - | - | - | - | - | - | - | - | - | - | - | - | - | - | - | - | - | - | - | - | - | - | - | - | - | - | - | - | - | - | - | - | - | - | - | - | - | - | - | - | - | - | - | - | - | - | - | - | - | - | - | - | - | - | - | - | - | - | - | - | - | - | - | - | - | - | - | - | - | - | - | - | - | - | - | - | - | - | - | - | - | - | - | - | - | - | - | - | - | - | - | - | - | - | - | - | - | - | - | - | - | - | - | - | - | - | - | - | - | - | - | - | - | - | - | - | - | - | - | - | - | - | - | - | - | - | - | - | - | - | - | - | - | - | - | - | - | - | - | - | - | - | - | - | - | - | - | - | - | - | - | - | - | - | - | - | - | - | - | - | - | - | - | - | - | - | - | - | - | - | - | - | - | - | - | - | - | - | - | - | - | - | - | - | - | - | - | - | - | - | - | - | - | - | - | - | - | - | - | - | - | - | - | - | - | - | - | - | - | - | - | - | - | - | - | - | - | - | - | - | - | - | - | - | - | - | - | - | - | - | - | - | - | - | - | - | - | - | - | - | - | - | - | - | - | - | - | - | - | - | - | - | - | - | - | - | - | - | - | - | - | - | - | - | - | - | - | - | - | - | - | - | - | - | - | - | - | - | - | - | - | - | - | - | - | - | - | - | - | - | - | - | - | - | - | - | - | - | - | - | - | - | - | - | - | - | - | - | - | - | - | - | - | - | - | - | - | - | - | - | - | - | - | - | - | - | - | - | - | - | - | - | - | - | - | - | - | - | - | - | - | - | - | - | - | - | - | - | - | - | - | - | - | - | - | - | - | - | - | - | - | - | - | - | - | - | - | - | - | - | - | - | - | - | - | - | - | - | - | - | - | - | - | - | - | - | - | - | - | - | - | - | - | - | - | - | - | - | - | - | - | - | - | - | - | - | - | - | - | - | - | - | - | - | - | - | - | - | - | - | - | - | - | - | - | - | - | - | - | - | - | - | - | - | - | - | - | - | - | - | - | - | - | - | - | - | - | - | - | - | - | - | - | - | - | - | - | - | - | - | - | - | - | - | - | - | - | - | - | - | - | - | - | - | - | - | - | - | - | - | - | - | - | - | - | - | - | - | - | - | - | - | - | - | - | - | - | - | - | - | - | - | - | - | - | - | - | - | - | - | - | - | - | - | - | - | - | - | - | - | - | - | - | - | - | - | - | - | - | - | - | - | - | - | - | - | - | - | - | - | - | - | - | - | - | - | - | - | - | - | - | - | - | - | - | - | - | - | - | - | - | - | - | - | - | - | - | - | - | - | - | - | - | - | - | - | - | - | - | - | - | - | - | - | - | - | - | - | - | - | - | - | - | - | - | - | - | - | - | - | - | - | - | - | - | - | - | - | - | - | - | - | - | - | - | - | - | - | - | - | - | - | - | - | - | - | - | - | - | - | - | - | - | - | - | - | - | - | - | - | - | - | - | - | - | - | - | - | - | - | - | - | - | - | - | - | - | - | - | - | - | - | - | - | - | - | - | - | - | - | - | - | - | - | - | - | - | - | - | - | - | - | - | - | - | - | - | - | - | - | - | - | - | - | - | - | - | - | - | - | - | - | - | - | - | - | - | - | - | - | - | - | - | - | - | - | - | - | - | - | - | - | - | - | - | - | - | - | - | - | - | - | - | - | - | - | - | - | - | - | - | - | - | - | - | - | - | - | - | - | - | - | - | - | - | - | - | - | - | - | - | - | - | - | - | - | - | - | - | - | - | - | - | - | - | - | - | - | - | - | - | - | - | - | - | - | - | - | - | - | - | - | - | - | - | - | - | - | - | - | - | - | - | - | - | - | - | - | - | - | - | - | - | - | - | - | - | - | - | - | - | - | - | - | - | - | - | - | - | - | - | - | - | - | - | - | - | - | - | - | - | - | - | - | - | - | - | - | - | - | - | - | - | - | - | - | - | - | - | - | - | - | - | - | - | - | - | - | - | - | - | - | - | - | - | - | - | - | - | - | - | - | - | - | - | - | - | - | - | - | - | - | - | - | - | - | - | - | - | - | - | - | - | - | - | - | - | - | - | - | - | - | - | - | - | - | - | - | - | - | - | - | - | - | - | - | - | - | - | - | - | - | - | - | - | - | - | - | - | - | - | - | - | - | - | - | - | - | - | - | - | - | - | - | - | - | - | - | - | - | - | - | - | - | - | - | - | - | - | - | - | - | - | - | - | - | - | - | - | - | - | - | - | - | - | - | - | - | - | - | - | - | - | - | - | - | - | - | - | - | - | - | - | - | - | - | - | - | - | - | - | - | - | - | - | - | - | - | - | - | - | - | - | - | - | - | - | - | - | - | - | - | - | - | - | - | - | - | - | - | - | - | - | - | - | - | - | - | - | - | - | - | - | - | - | - | - | - | - | - | - | - | - | - | - | - | - | - | - | - | - | - | - | - | - | - | - | - | - | - | - | - | - | - | - | - | - | - | - | - | - | - | - | - | - | - | - | - | - | - | - | - | - | - | - | - | - | - | - | - | - | - | - | - | - | - | - | - | - | - | - | - | - | - | - | - | - | - | - | - | - | - | - | - | - | - | - | - | - | - | - | - | - | - | - | - | - | - | - | - | - | - | - | - | - | - | - | - | - | - | - | - | - | - | - | - | - | - | - | - | - | - | - | - | - | - | - | - | - | - | - | - | - | - | - | - | - | - | - | - | - | - | - | - | - | - | - | - | - | - | - | - | - | - | - | - | - | - | - | - | - | - | - | - | - | - | - | - | - | - | - | - | - | - | - | - | - | - | - | - | - | - | - | - | - | - | - | - | - | - | - | - | - | - | - | - | - | - | - | - | - | - | - | - | - | - | - | - | - | - | - | - | - | - | - | - | - | - | - | - | - | - | - | - | - | - | - | - | - | - | - | - | - | - | - | - | - | - | - | - | - | - | - | - | - | - | - | - | - | - | - | - | - | - | - | - | - | - | - | - | - | - | - | - | - | - | - | - | - | - | - | - | - | - | - | - | - | - | - | - | - | - | - | - | - | - | - | - | - | - | - | - | - | - | - | - | - | - | - | - | - | - | - | - | - | - | - | - | - | - | - | - | - | - | - | - | - | - | - | - | - | - | - | - | - | - | - | - | - | - | - | - | - | - | - | - | - | - | - | - | - | - | - | - | - | - | - | - | - | - | - | - | - | - | - | - | - | - | - | - | - | - | - | - | - | - | - | - | - | - | - | - | - | - | - | - | - | - | - | - | - | - | - | - | - | - | - | - | - | - | - | - | - | - | - | - | - | - | - | - | - | - | - | - | - | - | - | - | - | - | - | - | - | - | - | - | - | - | - | - | - | - | - | - | - | - | - | - | - | - | - | - | - | - | - | - | - | - | - | - | - | - | - | - | - | - | - | - | - | - | - | - | - | - | - | - | - | - | - | - | - | - | - | - | - | - | - | - | - | - | - | - | - | - | - | - | - | - | - | - | - | - | - | - | - | - | - | - | - | - | - | - | - | - | - | - | - | - | - | - | - | - | - | - | - | - | - | - | - | - | - | - | - | - | - | - | - | - | - | - | - | - | - | - | - | - | - | - | - | - | - | - | - | - | - | - | - | - | - | - | - | - | - | - | - | - | - | - | - | - | - | - | - | - | - | - | - | - | - | - | - | - | - | - | - | - | - | - | - | - | - | - | - | - | - | - | - | - | - | - | - | - | - | - | - | - | - | - | - | - | - | - | - | - | - | - | - | - | - | - | - | - | - | - | - | - | - | - | - | - | - | - | - | - | - | - | - | - | - | - | - | - | - | - | - | - | - | - | - | - | - | - | - | - | - | - | - | - | - | - | - | - | - | - | - | - | - | - | - | - | - | - | - | - | - | - | - | - | - | - | - | - | - | - | - | - | - | - | - | - | - | - | - | - | - | - | - | - | - | - | - | - | - | - | - | - | - | - | - | - | - | - | - | - | - | - | - | - | - | - | - | - | - | - | - | - | - | - | - | - | - | - | - | - | - | - | - | - | - | - | - | - | - | - | - | - | - | - | - | - | - | - | - | - | H | H | x | x | x | x | x | x | x | x | x | x | x | x | x | x | x | x | x | x | x | x | x | x | x | x | x | x | x | x | x | x | x | x | x | x | x | x | x | x | x | x | x | x | x | x | x | x | x | x | x | x | x | x | x | x | x | x | x | x | x | x | x | x | x | x | x | x | x | x | x | x | x | x | x | x | x | x | x | x | x | x | x | x | x | x | x | x | x | x | x | x | H | x | x | H | x | H | x | x | x | x | x | x | x | x | x | x | x | x | x | x | x | x | x | x | x | x | x | x | x | x | x | x | x | H | H | H | x | H | - | - | - | - | - | - | - | - | - | - | - | - | - | - | - | - | - | - | - | - | - | - | - | - | - | - | - | - | - | - | - | - | - | - | - | - | - | - | - | - | - | - | - | - | - | - | - | - | - | - | - | - | - | - | - | - | - | - | - | - | - | - | - | - | - | - | - | - | - | - | - | - | - | - | - | - | - | - | - | - | - | - | - | - | - | - | - | - | - | - | - | - | - | - | - | - | - | - | - | - | - | - | - | - | - | - | - | - | - | - | - | - | - | - | - | - | - | - | - | - | - | - | - | - | - | - | - | - | - | - | - | - | - | - | - | - | - | - | - | - | - | - | - | - | - | - | - | - | - | - | - | - | - | - | - | - | - | - | - | - | - | - | - | - | - | - | - | - | - | - | - | - | - | - | - | - | - | - | - | - | - | - | - | - | - | - | - | - | - | - | - | - | - | - | - | - | - | - | - | - | - | - | - | - | - | - | - | - | - | - | - | - | - | - | - | - | - | - | - | - | - | - | - | - | - | - | - | - | - | - | - | - | - | - | - | - | - | - | - | - | - | - | - | - | - | - | - | - | - | - | - | - | - | - | - | - | - | - | - | - | - | - | - | - | - | - | - | - | - | - | - | - | - | - | - | - | - | - | - | - | - | - | - | - | - | - | - | - | - | - | - | - | - | - | - | - | - | - | - | - | - | - | - | - | - | - | - | - | - | - | - | - | - | - | - | - | - | - | - | - | - | - | - | - | - | - | - | - | - | - | - | - | - | - | - | - | - | - | - | - | - | - | - | - | - | - | - | - | - | - | - | - | - | - | - | - | - | - | - | - | - | - | - | - | - | - | - | - | - | - | - | - | - | - | - | - | - | - | - | - | - | - | - | - | - | - | - | - | - | - | - | - | - | - | - | - | - | - | - | - | - | - | - | - | - | - | - | - | - | - | - | - | - | - | - | - | - | - | - | - | - | - | - | - | - | - | - | - | - | - | - | - | - | - | - | - | - | - | - | - | - | - | - | - | - | - | - | - | - | - | - | - | - | - | - | - | - | - | - | - | - | - | - | - | - | - | - | - | - | - | - | - | - | - | - | - | - | - | - | - | - | - | - | - | - | - | - | - | - | - | - | - | - | - | - | - | - | - | - | - | - | - | - | - | - | - | - | - | - | - | - | - | - | - | - | - | - | - | - | - | - | - | - | - | - | - | - | - | - | - | - | - | - | - | - | - | - | - | - | - | - | - | - | - | - | - | - | - | - | - | - | - | - | - | - | - | - | - | - | - | - | - | - | - | - | - | - | - | - | - | - | - | - | - | - | - | - | - | - | - | - | - | - | - | - | - | - | - | - | - | - | - | - | - | - | - | - | - | - | - | - | - | - | - | - | - | - | - | - | - | - | - | - | - | - | - | - | - | - | - | - | - | - | - | - | - | - | - | - | - | - | - | - | - | - | - | - | - | - | - | - | - | - | - | - | - | - | - | - | - | - | - | - | - | - | - | - | - | - | - | - | - | - | - | - | - | - | - | - | - | - | - | - | - | - | - | - | - | - | - | - | - | - | - | - | - | - | - | - | - | - | - | - | - | - | - | - | - | - | - | - | - | - | - | - | - | - | - | - | - | - | - | - | - | - | - | - | - | - | - | - | - | - | - | - | - | - | - | - | - | - | - | - | - | - | - | - | - | - | - | - | - | - | - | - | - | - | - | - | - | - | - | - | - | - | - | - | - | - | - | - | - | - | - | - | - | - | - | - | - | - | - | - | - | - | - | - | - | - | - | - | - | - | - | - | - | - | - | - | - | - | - | - | - | - | - | - | - | - | - | - | - | - | - | - | - | - | - | - | - | - | - | - | - | - | - | - | - | - | - | - | - | - | - | - | - | - | - | - | - | - | - | - | - | - | - | - | - | - | - | - | - | - | - | - | - | - | - | - | - | - | - | - | - | - | - | - | - | - | - | - | - | - | - | - | - | - | - | - | - | - | - | - | - | - | - | - | - | - | - | - | - | - | - | - | - | - | - | - | - | - | - | - | - | - | - | - | - | - | - | - | - | - | - | - | - | - | - | - | - | - | - | - | - | - | - | - | - | - | - | - | - | - | - | - | - | - | - | - | - | - | - | - | - | - | - | - | - | - | - | - | - | - | - | - | - | - | - | - | - | - | - | - | - | - | - | - | - | - | - | - | - | - | - | - | - | - | - | - | - | - | - | - | - | - | - | - | - | - | - | - | - | - | - | - | - | - | - | - | - | - | - | - | - | - | - | - | - | - | - | - | - | - | - | - | - | - | - | - | - | - | - | - | - | - | - | - | - | - | - | - | - | - | - | - | - | - | - | - | - | - | - | - | - | - | - | - | - | - | - | - | - | - | - | - | - | - | - | - | - | - | - | - | - | - | - | - | - | - | - | - | - | - | - | - | - | - | - | - | - | - | - | - | - | - | - | - | - | - | - | - | - | - | - | - | - | - | - | - | - | - | - | - | - | - | - | - | - | - | - | - | - | - | - | - | - | - | - | - | - | - | - | - | - | - | - | - | - | - | - | - | - | - | - | - | - | - | - | - | - | - | - | - | - | - | - | - | - | - | - | - | - | - | - | - | - | - | - | - | - | - | - | - | - | - | - | - | - | - | - | - | - | - | - | - | - | - | - | - | - | - | - | - | - | - | - | - | - | - | - | - | - | - | - | - | - | - | - | - | - | - | - | - | - | - | - | - | - | - | - | - | - | - | - | - | - | - | - | - | - | - | - | - | - | - | - | - | - | - | - | - | - | - | - | - | - | - | - | - | - | - | - | - | - | - | - | - | - | - | - | - | - | - | - | - | - | - | - | - | - | - | - | - | - | - | - | - | - | - | - | - | - | - | - | - | - | - | - | - | - | - | - | - | - | - | - | - | - | - | - | - | - | - | - | - | - | - | - | - | - | - | - | - | - | - | - | - | - | - | - | - | - | - | - | - | - | - | - | - | - | - | - | - | - | - | - | - | - | - | - | - | - | - | - | - | - | - | - | - | - | - | - | - | - | - | - | - | - | - | - | - | - | - | - | - | - | - | - | - | - | - | - | - | - | - | - | - | - | - | - | - | - | - | - | - | - | - | - | - | - | - | - | - | - | - | - | - | - | - | - | - | - | - | - | - | - | - | - | - | - | - | - | - | - | - | - | - | - | - | - | - | - | - | - | - | - | - | - | - | - | - | - | - | - | - | - | - | - | - | - | - | - | - | - | - | - | - | - | - | - | - | - | - | - | - | - | - | - | - | - | - | - | - | - | - | - | - | - | - | - | - | - | - | - | - | - | - | - | - | - | - | - | - | - | - | - | - | - | - | - | - | - | - | - | - | - | - | - | - | - | - | - | - | - | - | - | - | - | - | - | - | - | - | - | - | - | - | - | - | - | - | - | - | - | - | - | - | - | - | - | - | - | - | - | - | - | - | - | - | - | - | - | - | - | - | - | - | - | - | - | - | - | - | - | - | - | - | - | - | - | - | - | - | - | - | - | - | - | - | - | - | - | - | - | - | - | - | - | - | - | - | - | - | - | - | - | - | - | - | - | - | - | - | - | - | - | - | - | - | - | - | - | - | - | - | - | - | - | - | - | - | - | - | - | - | - | - | - | - | - | - | - | - | - | - | - | - | - | - | - | - | - | - | - | - | - | - | - | - | - | - | - | - | - | - | - | - | - | - | - | - | - | - | - | - | - | - | - | - | - | - | - | - | - | - | - | - | - | - | - | - | - | - | - | - | - | - | - | - | - | - | - | - | - | - | - | - | - | - | - | - | - | - | - | - | - | - | - | - | - | - | - | - | - | - | - | - | - | - | - | - | - | - | - | - | - | - | - | - | - | - | - | - | - | - | - | - | - | - | - | - | - | - | - | - | - | - | - | - | - | - | - | - | - | - | - | - | - | - | - | - | - | - | - | - | - | - | - | - | - | - | - | - | - | - | - | - | - | - | - | - | - | - | - | - | - | - | - | - | - | - | - | - | - | - | - | - | - | - | - | - | - | - | - | - | - | - | - | - | - | - | - | - | - | - | - | - | - | - | - | - | - | - | - | - | - | - | - | - | - | - | - | - | - | - | - | - | - | - | - | - | - | - | - | - | - | - | - | - | - | - | - | - | - | - | - | - | - | - | - | - | - | - | - | - | - | - | - | - | - | - | - | - | - | - | - | - | - | - | - | - | - | - | - | - | - | - | - | - | - | - | - | - | - | - | - | - | - | - | - | - | - | - | - | - | - | - | - | - | - | - | - | - | - | - | - | - | - | - | - | - | - | - | - | - | - | - | - | - | - | - | - | - | - | - | - | - | - | - | - | - | - | - | - | - | - | - | - | - | - | - | - | - | - | - | - | - | - | - | - | - | - | - | - | - | - | - | - | - | - | - | - | - | - | - | - | - | - | - | - | - | - | - | - | - | - | - | - | - | - | - | - | - | - | - | - | - | - | - | - | - | - | - | - | - | - | - | - | - | - | - | - | - | - | - | - | - | - | - | - | - | - | - | - | - | - | - | - | - | - | - | - | - | - | - | - | - | - | - | - | - | - | - | - | - | - | - | - | - | - | - | - | - | - | - | - | - | - | - | - | - | - | - | - | - | - | - | - | - | - | - | - | - | - | - | - | - | - | - | - | - | - | - | - | - | - | - | - | - | - | - | - | - | - | - | - | - | - | - | - | - | - | - | - | - | - | - | - | - | - | - | - | - | - | - | - | - | - | - | - | - | - | - | - | - | - | - | - | - | - | - | - | - | - | - | - | - | - | - | - | - | - | - | - | - | - | - | - | - | - | - | - | - | - | - | - | - | - | - | - | - | - | - | - | - | - | - | - | - | - | - | - | - | - | - | - | - | - | - | - | - | - | - | - | - | - | - | - | - | - | - | - | - | - | - | - | - | - | - | - | - | - | - | - | - | - | - | - | - | - | - | - | - | - | - | - | - | - | - | - | - | - | - | - | - | - | - | - | - | - | - | - | - | - | - | - | - | - | - | - | - | - | - | - | - | - | - | - | - | - | - | - | - | - | - | - | - | - | - | - | - | - | - | - | - | - | - | - | - | - | - | - | - | - | - | - | - | - | - | - | - | - | - | - | - | - | - | - | - | - | - | - | - | - | - | - | - | - | - | - | - | - | - | - | - | - | - | - | - | - | - | - | - | - | - | - | - | - | - | - | - | - | - | - | - | - | - | - | - | - | - | - | - | - | - | - | - | - | - | - | - | - | - | - | - | - | - | - | - | - | - | - | - | - | - | - | - | - | - | - | - | - | - | - | - | - | - | - | - | - | - | - | - | - | - | - | - | - | - | - | - | - | - | - | - | - | - | - | - | - | - | - | - | - | - | - | - | - | - | - | - | - | - | - | - | - | - | - | - | - | - | - | - | - | - | - | - | - | - | - | - | - | - | - | - | - | - | - | - | - | - | - | - | - | - | - | - | - | - | - | - | - | - | - | - | - | - | - | - | - | - | - | - | - | - | - | - | - | - | - | - | - | - | - | - | - | - | - | - | - | - | - | - | - | - | - | - | - | - | - | - | - | - | - | - | - | - | - | - | - | - | - | - | - | - | - | - | - | - | - | - | - | - | - | - | - | - | - | - | - | - | - | - | - | - | - | - | - | - | - | - | - | - | - | - | - | - | - | - | - | - | - | - | - | - | - | - | - | - | - | - | - | - | - | - | - | - | - | - | - | - | - | - | - | - | - | - | - | - | - | - | - | - | - | - | - | - | - | - | - | - | - | - | - | - | - | - | - | - | - | - | - | - | - | - | - | - | - | - | - | - | - | - | - | - | - | - | - | - | - | - | - | - | - | - | - | - | - | - | - | - | - | - | - | - | - | - | - | - | - | - | - | - | - | - | - | - | - | - | - | - | - | - | - | - | - | - | - | - | - | - | - | - | - | - | - | - | - | - | - | - | - | - | - | - | - | - | - | - | - | - | - | - | - | - | - | - | - | - | - | - | - | - | - | - | - | - | - | - | - | - | - | - | - | - | - | - | - | - | - | - | - | - | - | - | - | - | - | - | - | - | - | - | - | - | - | - | - | - | - | - | - | - | - | - | - | - | - | - | - | - | - | - | - | - | - | - | - | - | - | - | - | - | - | - | - | - | - | - | - | - | - | - | - | - | - | - | - | - | - | - | - | - | - | - | - | - | - | - | - | - | - | - | - | - | - | - | - | - | - | - | - | - | - | - | - | - | - | - | - | - | - | - | - | - | - | - | - | - | - | - | - | - | - | - | - | - | - | - | - | - | - | - | - | - | - | - | - | - | - | - | - | - | - | - | - | - | - | - | - | - | - | - | - | - | - | - | - | - | - | - | - | - | - | - | - | - | - | - | - | - | - | - | - | - | - | - | - | - | - | - | - | - | - | - | - | - | - | - | - | - | - | - | - | - | - | - | - | - | - | - | - | - | - | - | - | - | - | - | - | - | - | - | - | - | - | - | - | - | - | - | - | - | - | - | - | - | - | - | - | - | - | - | - | - | - | - | - | - | - | - | - | - | - | - | - | - | - | - | - | - | - | - | - | - |
| BIS d=0 cl.3 sym=1 env=1 | - | - | - | - | - | - | - | - | - | - | - | - | - | - | - | - | - | - | - | - | - | - | - | - | - | - | - | - | - | - | - | - | - | - | - | - | - | - | - | - | - | - | - | - | - | - | - | - | - | - | - | - | - | - | - | - | - | - | - | - | - | - | - | - | - | - | - | - | - | - | - | - | - | - | - | - | - | - | - | - | - | - | - | - | - | - | - | - | - | - | - | - | - | - | - | - | - | - | - | - | - | - | - | - | - | - | - | - | - | - | - | - | - | - | - | - | - | - | - | - | - | - | - | - | - | - | - | - | - | - | - | - | - | - | - | - | - | - | - | - | - | - | - | - | - | - | - | - | - | - | - | - | - | - | - | - | - | - | - | - | - | - | - | - | - | - | - | - | - | - | - | - | - | - | - | - | - | - | - | - | - | - | - | - | - | - | - | - | - | - | - | - | - | - | - | - | - | - | - | - | - | - | - | - | - | - | - | - | - | - | - | - | - | - | - | - | - | - | - | - | - | - | - | - | - | - | - | - | - | - | - | - | - | - | - | - | - | - | - | - | - | - | - | - | - | - | - | - | - | - | - | - | - | - | - | - | - | - | - | - | - | - | - | - | - | - | - | - | - | - | - | - | - | - | - | - | - | - | - | - | - | - | - | - | - | - | - | - | - | - | - | - | - | - | - | - | - | - | - | - | - | - | - | - | - | - | - | - | - | - | - | - | - | - | - | - | - | - | - | - | - | - | - | - | - | - | - | - | - | - | - | - | - | - | - | - | - | - | - | - | - | - | - | - | - | - | - | - | - | - | - | - | - | - | - | - | - | - | - | - | - | - | - | - | - | - | - | - | - | - | - | - | - | - | - | - | - | - | - | - | - | - | - | - | - | - | - | - | - | - | - | - | - | - | - | - | - | - | - | - | - | - | - | - | - | - | - | - | - | - | - | - | - | - | - | - | - | - | - | - | - | - | - | - | - | - | - | - | - | - | - | - | - | - | - | - | - | - | - | - | - | - | - | - | - | - | - | - | - | - | - | - | - | - | - | - | - | - | - | - | - | - | - | - | - | - | - | - | - | - | - | - | - | - | - | - | - | - | - | - | - | - | - | - | - | - | - | - | - | - | - | - | - | - | - | - | - | - | - | - | - | - | - | - | - | - | - | - | - | - | - | - | - | - | - | - | - | - | - | - | - | - | - | - | - | - | - | - | - | - | - | - | - | - | - | - | - | - | - | - | - | - | - | - | - | - | - | - | - | - | - | - | - | - | - | - | - | - | - | - | - | - | - | - | - | - | - | - | - | - | - | - | - | - | - | - | - | - | - | - | - | - | - | - | - | - | - | - | - | - | - | - | - | - | - | - | - | - | - | - | - | - | - | - | - | - | - | - | - | - | - | - | - | - | - | - | - | - | - | - | - | - | - | - | - | - | - | - | - | - | - | - | - | - | - | - | - | - | - | - | - | - | - | - | - | - | - | - | - | - | - | - | - | - | - | - | - | - | - | - | - | - | - | - | - | - | - | - | - | - | - | - | - | - | - | - | - | - | - | - | - | - | - | - | - | - | - | - | - | - | - | - | - | - | - | - | - | - | - | - | - | - | - | - | - | - | - | - | - | - | - | - | - | - | - | - | - | - | - | - | - | - | - | - | - | - | - | - | - | - | - | - | - | - | - | - | - | - | - | - | - | - | - | - | - | - | - | - | - | - | - | - | - | - | - | - | - | - | - | - | - | - | - | - | - | - | - | - | - | - | - | - | - | - | - | - | - | - | - | - | - | - | - | - | - | - | - | - | - | - | - | - | - | - | - | - | - | - | - | - | - | - | - | - | - | - | - | - | - | - | - | - | - | - | - | - | - | - | - | - | - | - | - | - | - | - | - | - | - | - | - | - | - | - | - | - | - | - | - | - | - | - | - | - | - | - | - | - | - | - | - | - | - | - | - | - | - | - | - | - | - | - | - | - | - | - | - | - | - | - | - | - | - | - | - | - | - | - | - | - | - | - | - | - | - | - | - | - | - | - | - | - | - | - | - | - | - | - | - | - | - | - | - | - | - | - | - | - | - | - | - | - | - | - | - | - | - | - | - | - | - | - | - | - | - | - | - | - | - | - | - | - | - | - | - | - | - | - | - | - | - | - | - | - | - | - | - | - | - | - | - | - | - | - | - | - | - | - | - | - | - | - | - | - | - | - | - | - | - | - | - | - | - | - | - | - | - | - | - | - | - | - | - | - | - | - | - | - | - | - | - | - | - | - | - | - | - | - | - | - | - | - | - | - | - | - | - | - | - | - | - | - | - | - | - | - | - | - | - | - | - | - | - | - | - | - | - | - | - | - | - | - | - | - | - | - | - | - | - | - | - | - | - | - | - | - | - | - | - | - | - | - | - | - | - | - | - | - | - | - | - | - | - | - | - | - | - | - | - | - | - | - | - | - | - | - | - | - | - | - | - | - | - | - | - | - | - | - | - | - | - | - | - | - | - | - | - | - | - | - | - | - | - | - | - | - | - | - | - | - | - | - | - | - | - | - | - | - | - | - | - | - | - | - | - | - | - | - | - | - | - | - | - | - | - | - | - | - | - | - | - | - | - | - | - | - | - | - | - | - | - | - | - | - | - | - | - | - | - | - | - | - | - | - | - | - | - | - | - | - | - | - | - | - | - | - | - | - | - | - | - | - | - | - | - | - | - | - | - | - | - | - | - | - | - | - | - | - | - | - | - | - | - | - | - | - | - | - | - | - | - | - | - | - | - | - | - | - | - | - | - | - | - | - | - | - | - | - | - | - | - | - | - | - | - | - | - | - | - | - | - | - | - | - | - | - | - | - | - | - | - | - | - | - | - | - | - | - | - | - | - | - | - | - | - | - | - | - | - | - | - | - | - | - | - | - | - | - | - | - | - | - | - | - | - | - | - | - | - | - | - | - | - | - | - | - | - | - | - | - | - | - | - | - | - | - | - | - | - | - | - | - | - | - | - | - | - | - | - | - | - | - | - | - | - | - | - | - | - | - | - | - | - | - | - | - | - | - | - | - | - | - | - | - | - | - | - | - | - | - | - | - | - | - | - | - | - | - | - | - | - | - | - | - | - | - | - | - | - | - | - | - | - | - | - | - | - | - | - | - | - | - | - | - | - | - | - | - | - | - | - | - | - | - | - | - | - | - | - | - | - | - | - | - | - | - | - | - | - | - | - | - | - | - | - | - | - | - | - | - | - | - | - | - | - | - | - | - | - | - | - | - | - | - | - | - | - | - | - | - | - | - | - | - | - | - | - | - | - | - | - | - | - | - | - | - | - | - | - | - | - | - | - | - | - | - | - | - | - | - | - | - | - | - | - | - | - | - | - | - | - | - | - | - | - | - | - | - | - | - | - | - | - | - | - | - | - | - | - | - | - | - | - | - | - | - | - | - | - | - | - | - | - | - | - | - | - | - | - | - | - | - | - | - | - | - | - | - | - | - | - | - | - | - | - | - | - | - | - | - | - | - | - | - | - | - | - | - | - | - | - | - | - | - | - | - | - | - | - | - | - | - | - | - | - | - | - | - | - | - | - | - | - | - | - | - | - | - | - | - | - | - | - | - | - | - | - | - | - | - | - | - | - | - | - | - | - | - | - | - | - | - | - | - | - | - | - | - | - | - | - | - | - | - | - | - | - | - | - | - | - | - | - | - | - | - | - | - | - | - | - | - | - | - | - | - | - | - | - | - | - | - | - | - | - | - | - | - | - | - | - | - | - | - | - | - | - | - | - | - | - | - | - | - | - | - | - | - | - | - | - | - | - | - | - | - | - | - | - | - | - | - | - | - | - | - | - | - | - | - | - | - | - | - | - | - | - | - | - | - | - | - | - | - | - | - | - | - | - | - | - | - | - | - | - | - | - | - | - | - | - | - | - | - | - | - | - | - | - | - | - | - | - | - | - | - | - | - | - | - | - | - | - | - | - | - | - | - | - | - | - | - | - | - | - | - | - | - | - | - | - | - | - | - | - | - | - | - | - | - | - | - | - | - | - | - | - | - | - | - | - | - | - | - | - | - | - | - | - | - | - | - | - | - | - | - | - | - | - | - | - | - | - | - | - | - | - | - | - | - | - | - | - | - | - | - | - | - | - | - | - | - | - | - | - | - | - | - | - | - | - | - | - | - | - | - | - | - | - | - | - | - | - | - | - | - | - | - | - | - | - | - | - | - | - | - | - | - | - | - | - | - | - | - | - | - | - | - | - | - | - | - | - | - | - | - | - | - | - | - | - | - | - | - | - | - | - | - | - | - | - | - | - | - | - | - | - | - | - | - | - | - | - | - | - | - | - | - | - | - | - | - | - | - | - | - | - | - | - | - | - | - | - | - | - | - | - | - | - | - | - | - | - | - | - | - | - | - | - | - | - | - | - | - | - | - | - | - | - | - | - | - | - | - | - | - | - | - | - | - | - | - | - | - | - | - | - | - | - | - | - | - | - | - | - | - | - | - | - | - | - | - | - | - | - | - | - | - | - | - | - | - | - | - | - | - | - | - | - | - | - | - | - | - | - | - | - | - | - | - | - | - | - | - | - | - | - | - | - | - | - | - | - | - | - | - | - | - | - | - | - | - | - | - | - | - | - | - | - | - | - | - | - | - | - | - | - | - | - | - | - | - | - | - | - | - | - | - | - | - | - | - | - | - | - | - | - | - | - | - | - | - | - | - | - | - | - | - | - | - | - | - | - | - | - | - | - | - | - | - | - | - | - | - | - | - | - | - | - | - | - | - | - | - | - | - | - | - | - | - | - | - | - | - | - | - | - | - | - | - | - | - | - | - | - | - | - | - | - | - | - | - | - | - | - | - | - | - | - | - | - | - | - | - | - | - | - | - | - | - | - | - | - | - | - | - | - | - | - | - | - | - | - | - | - | - | - | - | - | - | - | - | - | - | - | - | - | - | - | - | - | - | - | - | - | - | - | - | - | - | - | - | - | - | - | - | - | - | - | - | - | - | - | - | - | - | - | - | - | - | - | - | - | - | - | - | - | - | - | - | - | - | - | - | - | - | - | - | - | - | - | - | - | - | - | - | - | - | - | - | - | - | - | - | - | - | - | - | - | - | - | - | - | - | - | - | - | - | - | - | - | - | - | - | - | - | - | - | - | - | - | - | - | - | - | - | - | - | - | - | - | - | - | - | - | - | - | - | - | - | - | - | - | - | - | - | - | - | - | - | - | - | - | - | - | - | - | - | - | - | - | - | - | - | - | - | - | - | - | - | - | - | - | - | - | - | - | - | - | - | - | - | - | - | - | - | - | - | - | - | - | - | - | - | - | - | - | - | - | - | - | - | - | - | - | - | - | - | - | - | - | - | - | - | - | - | - | - | - | - | - | - | - | - | - | - | - | - | - | - | - | - | - | - | - | - | - | - | - | - | - | - | - | - | - | - | - | - | - | - | - | - | - | - | - | - | - | - | - | - | - | - | - | - | - | - | - | - | - | - | - | - | - | - | - | - | - | - | - | - | - | - | - | - | - | - | - | - | - | - | - | - | - | - | - | - | - | - | - | - | - | - | - | - | - | - | - | - | - | - | - | - | - | - | - | - | - | - | - | - | - | - | - | - | - | - | - | - | - | - | - | - | - | - | - | - | - | - | - | - | - | - | - | - | - | - | - | - | - | - | - | - | - | - | - | - | - | - | - | - | - | - | - | - | - | - | - | - | - | - | - | - | - | - | - | - | - | - | - | - | - | - | - | - | - | - | - | - | - | - | - | - | - | - | - | - | - | - | - | - | - | - | - | - | - | - | - | - | - | - | - | - | - | - | - | - | - | - | - | - | - | - | - | - | - | - | - | - | - | - | - | - | - | - | - | - | - | - | - | - | - | - | - | - | - | - | - | - | - | - | - | - | - | - | - | - | - | - | - | - | - | - | - | - | - | - | - | - | - | - | - | - | - | - | - | - | - | - | - | - | - | - | - | - | - | - | - | - | - | - | - | - | - | - | - | - | - | - | - | - | - | - | - | - | - | - | - | - | - | - | - | - | - | - | - | - | - | - | - | - | - | - | - | - | - | - | - | - | - | - | - | - | - | - | - | - | - | - | - | - | - | - | - | - | - | - | - | - | - | - | - | - | - | - | - | - | - | - | - | - | - | - | - | - | - | - | - | - | - | - | - | - | - | - | - | - | - | - | - | - | - | - | - | - | - | - | - | - | - | - | - | - | - | - | - | - | - | - | - | - | - | - | - | - | - | - | - | - | - | - | - | - | - | - | - | - | - | - | - | - | - | - | - | - | - | - | - | - | - | - | - | - | - | - | - | - | - | - | - | - | - | - | - | - | - | - | - | - | - | - | - | - | - | - | - | - | - | - | - | - | - | - | - | - | - | - | - | - | - | - | - | - | - | - | - | - | - | - | - | - | - | - | - | - | - | - | - | - | - | - | - | - | - | - | - | - | - | - | - | - | - | - | - | - | - | - | - | - | - | - | - | - | - | - | - | - | - | - | - | - | - | - | - | - | - | - | - | - | - | - | - | - | - | - | - | - | - | - | - | - | - | - | - | - | - | - | - | - | - | - | - | - | - | - | - | - | - | - | - | - | - | - | - | - | - | - | - | - | - | - | - | - | - | - | - | - | - | - | - | - | - | - | - | - | - | - | - | - | - | - | - | - | - | - | - | - | - | - | - | - | - | - | - | - | - | - | - | - | - | - | - | - | - | - | - | - | - | - | - | - | - | - | - | - | - | - | - | - | - | - | - | - | - | - | - | - | - | - | - | - | - | - | - | - | - | - | - | - | - | - | - | - | - | - | - | - | - | - | - | - | - | - | - | - | - | - | - | - | - | - | - | - | - | - | - | - | - | - | - | - | - | - | - | - | - | - | - | - | - | - | - | - | - | - | - | - | - | - | - | - | - | - | - | - | - | - | - | - | - | - | - | - | - | - | - | - | - | - | - | - | - | - | - | - | - | - | - | - | - | - | - | - | - | - | - | - | - | - | - | - | - | - | - | - | - | - | - | - | - | - | - | - | - | - | - | - | - | - | - | - | - | - | - | - | - | - | - | - | - | - | - | - | - | - | - | - | - | - | - | - | - | - | - | - | - | - | - | - | - | - | - | - | - | - | - | - | - | - | - | - | - | - | - | - | - | - | - | - | - | - | - | - | - | - | - | - | - | - | - | - | - | - | - | - | - | - | - | - | - | - | - | - | - | - | - | - | - | - | - | - | - | - | - | - | - | - | - | - | - | - | - | - | - | - | - | - | - | - | - | - | - | - | - | - | - | - | - | - | - | - | - | - | - | - | - | - | - | - | - | - | - | - | - | - | - | - | - | - | - | - | - | - | - | - | - | - | - | - | - | - | - | - | - | - | - | - | - | - | - | - | - | - | - | - | - | - | - | - | - | - | - | - | - | - | - | - | - | - | - | - | - | - | - | - | - | - | - | - | - | - | - | - | - | - | - | - | - | - | - | - | - | - | - | - | - | - | - | - | - | - | - | - | - | - | - | - | - | - | - | - | - | - | - | - | - | - | - | - | - | - | - | - | - | - | - | - | - | - | - | - | - | - | - | - | - | - | - | - | - | - | - | - | - | - | - | - | - | - | - | - | - | - | - | - | - | - | - | - | - | - | - | - | - | - | - | - | - | - | - | - | - | - | - | - | - | - | - | - | - | - | - | - | - | - | - | - | - | - | - | - | - | - | - | - | - | - | - | - | - | - | - | - | - | - | - | - | - | - | - | - | - | - | - | - | - | - | - | - | - | - | - | - | - | - | - | - | - | - | - | - | - | - | - | - | - | - | - | - | - | - | - | - | - | - | - | - | - | - | - | - | - | - | - | - | - | - | - | - | - | - | - | - | - | - | - | - | - | - | - | - | - | - | - | - | - | - | - | - | - | - | - | - | - | - | - | - | - | - | - | - | - | - | - | - | - | - | - | - | - | - | - | - | - | - | - | - | - | - | - | - | - | - | - | - | - | - | - | - | - | - | - | - | - | - | - | - | - | - | - | - | - | - | - | - | - | - | - | - | - | - | - | - | - | - | - | - | - | - | - | - | - | - | - | - | - | - | - | - | - | - | - | - | - | - | - | - | - | - | - | - | - | - | - | - | - | - | - | - | - | - | - | - | - | - | - | - | - | - | - | - | - | - | - | - | - | - | - | - | - | - | - | - | - | - | - | - | - | - | - | - | - | - | - | - | - | - | - | - | - | - | - | - | - | - | - | - | - | - | - | - | - | - | - | - | - | - | - | - | - | - | - | - | - | - | - | - | - | - | - | - | - | - | - | - | - | - | - | - | - | - | - | - | - | - | - | - | - | - | - | - | - | - | - | - | - | - | - | - | - | - | - | - | - | - | - | - | - | - | - | - | - | - | - | - | - | - | - | - | - | - | - | - | - | - | - | - | - | - | - | - | - | - | - | - | - | - | - | - | - | - | - | - | - | - | - | - | - | - | - | - | - | - | - | - | - | - | - | - | - | - | - | - | - | - | - | - | - | - | - | - | - | - | - | - | - | - | - | - | - | - | - | - | - | - | - | - | - | - | - | - | - | - | - | - | - | - | - | - | - | - | - | - | - | - | - | - | - | - | - | - | - | - | - | - | - | - | - | - | - | - | - | - | - | - | - | - | - | - | - | - | - | - | - | - | - | - | - | - | - | - | - | - | - | - | - | - | - | - | - | - | - | - | - | - | - | - | - | - | - | - | - | - | - | - | - | - | - | - | - | - | - | - | - | - | - | - | - | - | - | - | - | - | - | - | - | - | - | - | - | - | - | - | - | - | - | - | - | - | - | - | - | - | - | - | - | - | - | - | - | - | - | - | - | - | - | - | - | - | - | - | - | - | - | - | - | - | - | - | - | - | - | - | - | - | - | - | - | - | - | - | - | - | - | - | - | - | - | - | - | - | - | - | - | - | - | - | - | - | - | - | - | - | - | - | - | - | - | - | - | - | - | - | - | - | - | - | - | - | - | - | - | - | - | - | - | - | - | - | - | - | - | - | - | - | - | - | - | - | - | - | - | - | - | - | - | - | - | - | - | - | - | - | - | - | - | - | - | - | - | - | - | - | - | - | - | - | - | - | - | - | - | - | - | - | - | - | - | - | - | - | - | - | - | - | - | - | - | - | - | - | - | - | - | - | - | - | - | - | - | - | - | - | - | - | - | - | - | - | - | - | - | - | - | - | - | - | - | - | - | - | - | - | - | - | - | - | - | - | - | - | - | - | - | - | - | - | - | - | - | - | - | - | - | - | - | - | - | - | - | - | - | - | - | - | - | - | - | - | - | - | - | - | - | - | - | - | - | - | - | - | - | - | - | - | - | - | - | - | - | - | - | - | - | - | - | - | - | - | - | - | - | - | - | - | - | - | - | - | - | - | - | - | - | - | - | - | - | - | - | - | - | - | - | - | - | - | - | - | - | - | - | - | - | - | - | - | - | - | - | - | - | - | - | - | - | - | - | - | - | - | - | - | - | - | - | - | - | - | - | - | - | - | - | - | - | - | - | - | - | - | - | - | - | - | - | - | - | - | - | - | - | - | - | - | - | - | - | - | - | - | - | - | - | - | - | - | - | - | - | - | - | - | - | - | - | - | - | - | - | - | - | - | - | - | - | - | - | - | - | - | - | - | - | - | - | - | - | - | - | - | - | - | - | - | - | - | - | - | - | - | - | - | - | - | - | - | - | - | - | - | - | - | - | - | - | - | - | - | - | - | - | - | - | - | - | - | - | - | - | - | - | - | - | - | - | - | - | - | - | - | - | - | - | - | - | - | - | - | - | - | - | - | - | - | - | - | - | - | - | - | - | - | - | - | - | - | - | - | - | - | - | - | - | - | - | - | - | - | - | - | - | - | - | - | - | - | - | - | - | - | - | - | - | - | - | - | - | - | - | - | - | - | - | - | - | - | - | - | - | - | - | - | - | - | - | - | - | - | - | - | - | - | - | - | - | - | - | - | - | - | - | - | - | - | - | - | - | - | - | - | - | - | - | - | - | - | - | - | - | - | - | - | - | - | - | - | - | - | - | - | - | - | - | - | - | - | - | - | - | - | - | - | - | - | - | - | - | - | - | - | - | - | - | - | - | - | - | - | - | - | - | - | - | - | - | - | - | - | - | - | - | - | - | - | - | - | - | - | - | - | - | - | - | - | - | - | - | - | - | - | - | - | - | - | - | - | - | - | - | - | - | - | - | - | - | - | - | - | - | - | - | - | - | - | - | - | - | - | - | - | - | - | - | - | - | - | - | - | - | - | - | - | - | - | - | - | - | - | - | - | - | - | - | - | - | - | - | - | - | - | - | - | - | - | - | - | - | - | - | - | - | - | - | - | - | - | - | - | - | - | - | - | - | - | - | - | - | - | - | - | - | - | - | - | - | - | - | - | - | - | - | - | - | - | - | - | - | - | - | - | - | - | - | - | - | - | - | - | - | - | - | - | - | - | - | - | - | - | - | - | - | - | - | - | - | - | - | - | - | - | - | - | - | - | - | - | - | - | - | - | - | - | - | - | - | - | - | - | - | - | - | - | - | - | - | - | - | - | - | - | - | - | - | - | - | - | - | - | - | - | - | - | - | - | - | - | - | - | - | - | - | - | - | - | - | - | - | - | - | - | - | - | - | - | - | - | - | - | - | - | - | - | - | - | - | - | - | - | - | - | - | - | - | - | - | - | - | - | - | - | - | - | - | - | - | - | - | - | - | - | - | - | - | - | - | - | - | - | - | - | - | - | - | - | - | - | - | - | - | - | - | - | - | - | - | - | - | - | - | - | - | - | - | - | - | - | - | - | - | - | - | - | - | - | - | - | - | - | - | - | - | - | - | - | - | - | - | - | - | - | - | - | - | - | - | - | - | - | - | - | - | - | - | - | - | - | - | - | - | - | - | - | - | - | - | - | - | - | - | - | - | - | - | - | - | - | - | - | - | - | - | - | - | - | - | - | - | - | - | - | - | - | - | - | - | - | - | - | - | - | - | - | - | - | - | - | - | - | - | - | - | - | - | - | - | - | - | - | - | - | - | - | - | - | - | - | - | - | - | - | - | - | - | - | - | - | - | - | - | - | - | - | - | - | - | - | - | - | - | - | - | - | - | - | - | - | - | - | - | - | - | - | - | - | - | - | - | - | - | - | - | - | - | - | - | - | - | - | - | - | - | - | - | - | - | - | - | - | - | - | - | - | - | - | - | - | - | - | - | - | - | - | - | - | - | - | - | - | - | - | - | - | - | - | - | - | - | - | - | - | - | - | - | - | - | - | - | - | - | - | - | - | - | - | - | - | - | - | - | - | - | - | - | - | - | - | - | - | - | - | - | - | - | - | - | - | - | - | - | - | - | - | - | - | - | - | - | - | - | - | H | x | x | H | x | x | x | x | x | x | x | x | x | x | x | H | x | x | x | H | x | x | x | H | H | H | H | x | H | x | H | x | H | x | x | H | x | x | x | x | x | x | x | x | x | H | - | - | - | - | - | - | - | - | - | - | - | - | - | - | - | - | - | - | - | - | - | - | - | - | - | - | - | - | - | - | - | - | - | - | - | - | - | - | - | - | - | - | - | - | - | - | - | - | - | - | - | - | - | - | - | - | - | - | - | - | - | - | - | - | - | - | - | - | - | - | - | - | - | - | - | - | - | - | - | - | - | - | - | - | - | - | - | - | - | - | - | - | - | - | - | - | - | - | - | - | - | - | - | - | - | - | - | - | - | - | - | - | - | - | - | - | - | - | - | - | - | - | - | - | - | - | - | - | - | - | - | - | - | - | - | - | - | - | - | - | - | - | - | - | - | - | - | - | - | - | - | - | - | - | - | - | - | - | - | - | - | - | - | - | - | - | - | - | - | - | - | - | - | - | - | - | - | - | - | - | - | - | - | - | - | - | - | - | - | - | - | - | - | - | - | - | - | - | - | - | - | - | - | - | - | - | - | - | - | - | - | - | - | - | - | - | - | - | - | - | - | - | - | - | - | - | - | - | - | - | - | - | - | - | - | - | - | - | - | - | - | - | - | - | - | - | - | - | - | - | - | - | - | - | - | - | - | - | - | - | - | - | - | - | - | - | - | - | - | - | - | - | - | - | - | - | - | - | - | - | - | - | - | - | - | - | - | - | - | - | - | - | - | - | - | - | - | - | - | - | - | - | - | - | - | - | - | - | - | - | - | - | - | - | - | - | - | - | - | - | - | - | - | - | - | - | - | - | - | - | - | - | - | - | - | - | - | - | - | - | - | - | - | - | - | - | - | - | - | - | - | - |
| BIS d=0 cl.16 sym=1 env=1 | - | - | - | - | - | - | - | - | - | - | - | - | - | - | - | - | - | - | - | - | - | - | - | - | - | - | - | - | - | - | - | - | - | - | - | - | - | - | - | - | - | - | - | - | - | - | - | - | - | - | - | - | - | - | - | - | - | - | - | - | - | - | - | - | - | - | - | - | - | - | - | - | - | - | - | - | - | - | - | - | - | - | - | - | - | - | - | - | - | - | - | - | - | - | - | - | - | - | - | - | - | - | - | - | - | - | - | - | - | - | - | - | - | - | - | - | - | - | - | - | - | - | - | - | - | - | - | - | - | - | - | - | - | - | - | - | - | - | - | - | - | - | - | - | - | - | - | - | - | - | - | - | - | - | - | - | - | - | - | - | - | - | - | - | - | - | - | - | - | - | - | - | - | - | - | - | - | - | - | - | - | - | - | - | - | - | - | - | - | - | - | - | - | - | - | - | - | - | - | - | - | - | - | - | - | - | - | - | - | - | - | - | - | - | - | - | - | - | - | - | - | - | - | - | - | - | - | - | - | - | - | - | - | - | - | - | - | - | - | - | - | - | - | - | - | - | - | - | - | - | - | - | - | - | - | - | - | - | - | - | - | - | - | - | - | - | - | - | - | - | - | - | - | - | - | - | - | - | - | - | - | - | - | - | - | - | - | - | - | - | - | - | - | - | - | - | - | - | - | - | - | - | - | - | - | - | - | - | - | - | - | - | - | - | - | - | - | - | - | - | - | - | - | - | - | - | - | - | - | - | - | - | - | - | - | - | - | - | - | - | - | - | - | - | - | - | - | - | - | - | - | - | - | - | - | - | - | - | - | - | - | - | - | - | - | - | - | - | - | - | - | - | - | - | - | - | - | - | - | - | - | - | - | - | - | - | - | - | - | - | - | - | - | - | - | - | - | - | - | - | - | - | - | - | - | - | - | - | - | - | - | - | - | - | - | - | - | - | - | - | - | - | - | - | - | - | - | - | - | - | - | - | - | - | - | - | - | - | - | - | - | - | - | - | - | - | - | - | - | - | - | - | - | - | - | - | - | - | - | - | - | - | - | - | - | - | - | - | - | - | - | - | - | - | - | - | - | - | - | - | - | - | - | - | - | - | - | - | - | - | - | - | - | - | - | - | - | - | - | - | - | - | - | - | - | - | - | - | - | - | - | - | - | - | - | - | - | - | - | - | - | - | - | - | - | - | - | - | - | - | - | - | - | - | - | - | - | - | - | - | - | - | - | - | - | - | - | - | - | - | - | - | - | - | - | - | - | - | - | - | - | - | - | - | - | - | - | - | - | - | - | - | - | - | - | - | - | - | - | - | - | - | - | - | - | - | - | - | - | - | - | - | - | - | - | - | - | - | - | - | - | - | - | - | - | - | - | - | - | - | - | - | - | - | - | - | - | - | - | - | - | - | - | - | - | - | - | - | - | - | - | - | - | - | - | - | - | - | - | - | - | - | - | - | - | - | - | - | - | - | - | - | - | - | - | - | - | - | - | - | - | - | - | - | - | - | - | - | - | - | - | - | - | - | - | - | - | - | - | - | - | - | - | - | - | - | - | - | - | - | - | - | - | - | - | - | - | - | - | - | - | - | - | - | - | - | - | - | - | - | - | - | - | - | - | - | - | - | - | - | - | - | - | - | - | - | - | - | - | - | - | - | - | - | - | - | - | - | - | - | - | - | - | - | - | - | - | - | - | - | - | - | - | - | - | - | - | - | - | - | - | - | - | - | - | - | - | - | - | - | - | - | - | - | - | - | - | - | - | - | - | - | - | - | - | - | - | - | - | - | - | - | - | - | - | - | - | - | - | - | - | - | - | - | - | - | - | - | - | - | - | - | - | - | - | - | - | - | - | - | - | - | - | - | - | - | - | - | - | - | - | - | - | - | - | - | - | - | - | - | - | - | - | - | - | - | - | - | - | - | - | - | - | - | - | - | - | - | - | - | - | - | - | - | - | - | - | - | - | - | - | - | - | - | - | - | - | - | - | - | - | - | - | - | - | - | - | - | - | - | - | - | - | - | - | - | - | - | - | - | - | - | - | - | - | - | - | - | - | - | - | - | - | - | - | - | - | - | - | - | - | - | - | - | - | - | - | - | - | - | - | - | - | - | - | - | - | - | - | - | - | - | - | - | - | - | - | - | - | - | - | - | - | - | - | - | - | - | - | - | - | - | - | - | - | - | - | - | - | - | - | - | - | - | - | - | - | - | - | - | - | - | - | - | - | - | - | - | - | - | - | - | - | - | - | - | - | - | - | - | - | - | - | - | - | - | - | - | - | - | - | - | - | - | - | - | - | - | - | - | - | - | - | - | - | - | - | - | - | - | - | - | - | - | - | - | - | - | - | - | - | - | - | - | - | - | - | - | - | - | - | - | - | - | - | - | - | - | - | - | - | - | - | - | - | - | - | - | - | - | - | - | - | - | - | - | - | - | - | - | - | - | - | - | - | - | - | - | - | - | - | - | - | - | - | - | - | - | - | - | - | - | - | - | - | - | - | - | - | - | - | - | - | - | - | - | - | - | - | - | - | - | - | - | - | - | - | - | - | - | - | - | - | - | - | - | - | - | - | - | - | - | - | - | - | - | - | - | - | - | - | - | - | - | - | - | - | - | - | - | - | - | - | - | - | - | - | - | - | - | - | - | - | - | - | - | - | - | - | - | - | - | - | - | - | - | - | - | - | - | - | - | - | - | - | - | - | - | - | - | - | - | - | - | - | - | - | - | - | - | - | - | - | - | - | - | - | - | - | - | - | - | - | - | - | - | - | - | - | - | - | - | - | - | - | - | - | - | - | - | - | - | - | - | - | - | - | - | - | - | - | - | - | - | - | - | - | - | - | - | - | - | - | - | - | - | - | - | - | - | - | - | - | - | - | - | - | - | - | - | - | - | - | - | - | - | - | - | - | - | - | - | - | - | - | - | - | - | - | - | - | - | - | - | - | - | - | - | - | - | - | - | - | - | - | - | - | - | - | - | - | - | - | - | - | - | - | - | - | - | - | - | - | - | - | - | - | - | - | - | - | - | - | - | - | - | - | - | - | - | - | - | - | - | - | - | - | - | - | - | - | - | - | - | - | - | - | - | - | - | - | - | - | - | - | - | - | - | - | - | - | - | - | - | - | - | - | - | - | - | - | - | - | - | - | - | - | - | - | - | - | - | - | - | - | - | - | - | - | - | - | - | - | - | - | - | - | - | - | - | - | - | - | - | - | - | - | - | - | - | - | - | - | - | - | - | - | - | - | - | - | - | - | - | - | - | - | - | - | - | - | - | - | - | - | - | - | - | - | - | - | - | - | - | - | - | - | - | - | - | - | - | - | - | - | - | - | - | - | - | - | - | - | - | - | - | - | - | - | - | - | - | - | - | - | - | - | - | - | - | - | - | - | - | - | - | - | - | - | - | - | - | - | - | - | - | - | - | - | - | - | - | - | - | - | - | - | - | - | - | - | - | - | - | - | - | - | - | - | - | - | - | - | - | - | - | - | - | - | - | - | - | - | - | - | - | - | - | - | - | - | - | - | - | - | - | - | - | - | - | - | - | - | - | - | - | - | - | - | - | - | - | - | - | - | - | - | - | - | - | - | - | - | - | - | - | - | - | - | - | - | - | - | - | - | - | - | - | - | - | - | - | - | - | - | - | - | - | - | - | - | - | - | - | - | - | - | - | - | - | - | - | - | - | - | - | - | - | - | - | - | - | - | - | - | - | - | - | - | - | - | - | - | - | - | - | - | - | - | - | - | - | - | - | - | - | - | - | - | - | - | - | - | - | - | - | - | - | - | - | - | - | - | - | - | - | - | - | - | - | - | - | - | - | - | - | - | - | - | - | - | - | - | - | - | - | - | - | - | - | - | - | - | - | - | - | - | - | - | - | - | - | - | - | - | - | - | - | - | - | - | - | - | - | - | - | - | - | - | - | - | - | - | - | - | - | - | - | - | - | - | - | - | - | - | - | - | - | - | - | - | - | - | - | - | - | - | - | - | - | - | - | - | - | - | - | - | - | - | - | - | H | x | x | x | x | x | H | x | x | x | x | x | x | x | x | x | x | x | x | x | x | x | x | x | x | x | x | x | x | x | x | x | x | x | x | x | x | x | x | x | H | x | x | x | H | x | H | x | x | H | - | - | - | - | - | - | - | - | - | - | - | - | - | - | - | - | - | - | - | - | - | - | - | - | - | - | - | - | - | - | - | - | - | - | - | - | - | - | - | - | - | - | - | - | - | - | - | - | - | - | - | - | - | - | - | - | - | - | - | - | - | - | - | - | - | - | - | - | - | - | - | - | - | - | - | - | - | - | - | - | - | - | - | - | - | - | - | - | - | - | - | - | - | - | - | - | - | - | - | - | - | - | - | - | - | - | - | - | - | - | - | - | - | - | - | - | - | - | - | - | - | - | - | - | - | - | - | - | - | - | - | - | - | - | - | - | - | - | - | - | - | - | - | - | - | - | - | - | - | - | - | - | - | - | - | - | - | - | - | - | - | - | - | - | - | - | - | - | - | - | - | - | - | - | - | - | - | - | - | - | - | - | - | - | - | - | - | - | - | - | - | - | - | - | - | - | - | - | - | - | - | - | - | - | - | - | - | - | - | - | - | - | - | - | - | - | - | - | - | - | - | - | - | - | - | - | - | - | - | - | - | - | - | - | - | - | - | - | - | - | - | - | - | - | - | - | - | - | - | - | - | - | - | - | - | - | - | - | - | - | - | - | - | - | - | - | - | - | - | - | - | - | - | - | - | - | - | - | - | - | - | - | - | - | - | - | - | - | - | - | - | - | - | - | - | - | - | - | - | - | - | - | - | - | - | - | - | - | - | - | - | - | - | - | - | - | - | - | - | - | - | - | - | - | - | - | - | - | - | - | - | - | - | - | - | - | - | - | - | - | - | - | - | - | - | - | - | - | - | - | - | - | - | - | - | - | - | - | - | - | - | - | - | - | - | - | - | - | - | - | - | - | - | - | - | - | - | - | - | - | - | - | - | - | - | - | - | - | - | - | - | - | - | - | - | - | - | - | - | - | - | - | - | - | - | - | - | - | - | - | - | - | - | - | - | - | - | - | - | - | - | - | - | - | - | - | - | - | - | - | - | - | - | - | - | - | - | - | - | - | - | - | - | - | - | - | - | - | - | - | - | - | - | - | - | - | - | - | - | - | - | - | - | - | - | - | - | - | - | - | - | - | - | - | - | - | - | - | - | - | - | - | - | - | - | - | - | - | - | - | - | - | - | - | - | - | - | - | - | - | - | - | - | - | - | - | - | - | - | - | - | - | - | - | - | - | - | - | - | - | - | - | - | - | - | - | - | - | - | - | - | - | - | - | - | - | - | - | - | - | - | - | - | - | - | - | - | - | - | - | - | - | - | - | - | - | - | - | - | - | - | - | - | - | - | - | - | - | - | - | - | - | - | - | - | - | - | - | - | - | - | - | - | - | - | - | - | - | - | - | - | - | - | - | - | - | - | - | - | - | - | - | - | - | - | - | - | - | - | - | - | - | - | - | - | - | - | - | - | - | - | - | - | - | - | - | - | - | - | - | - | - | - | - | - | - | - | - | - | - | - | - | - | - | - | - | - | - | - | - | - | - | - | - | - | - | - | - | - | - | - | - | - | - | - | - | - | - | - | - | - | - | - | - | - | - | - | - | - | - | - | - | - | - | - | - | - | - | - | - | - | - | - | - | - | - | - | - | - | - | - | - | - | - | - | - | - | - | - | - | - | - | - | - | - | - | - | - | - | - | - | - | - | - | - | - | - | - | - | - | - | - | - | - | - | - | - | - | - | - | - | - | - | - | - | - | - | - | - | - | - | - | - | - | - | - | - | - | - | - | - | - | - | - | - | - | - | - | - | - | - | - | - | - | - | - | - | - | - | - | - | - | - | - | - | - | - | - | - | - | - | - | - | - | - | - | - | - | - | - | - | - | - | - | - | - | - | - | - | - | - | - | - | - | - | - | - | - | - | - | - | - | - | - | - | - | - | - | - | - | - | - | - | - | - | - | - | - | - | - | - | - | - | - | - | - | - | - | - | - | - | - | - | - | - | - | - | - | - | - | - | - | - | - | - | - | - | - | - | - | - | - | - | - | - | - | - | - | - | - | - | - | - | - | - | - | - | - | - | - | - | - | - | - | - | - | - | - | - | - | - | - | - | - | - | - | - | - | - | - | - | - | - | - | - | - | - | - | - | - | - | - | - | - | - | - | - | - | - | - | - | - | - | - | - | - | - | - | - | - | - | - | - | - | - | - | - | - | - | - | - | - | - | - | - | - | - | - | - | - | - | - | - | - | - | - | - | - | - | - | - | - | - | - | - | - | - | - | - | - | - | - | - | - | - | - | - | - | - | - | - | - | - | - | - | - | - | - | - | - | - | - | - | - | - | - | - | - | - | - | - | - | - | - | - | - | - | - | - | - | - | - | - | - | - | - | - | - | - | - | - | - | - | - | - | - | - | - | - | - | - | - | - | - | - | - | - | - | - | - | - | - | - | - | - | - | - | - | - | - | - | - | - | - | - | - | - | - | - | - | - | - | - | - | - | - | - | - | - | - | - | - | - | - | - | - | - | - | - | - | - | - | - | - | - | - | - | - | - | - | - | - | - | - | - | - | - | - | - | - | - | - | - | - | - | - | - | - | - | - | - | - | - | - | - | - | - | - | - | - | - | - | - | - | - | - | - | - | - | - | - | - | - | - | - | - | - | - | - | - | - | - | - | - | - | - | - | - | - | - | - | - | - | - | - | - | - | - | - | - | - | - | - | - | - | - | - | - | - | - | - | - | - | - | - | - | - | - | - | - | - | - | - | - | - | - | - | - | - | - | - | - | - | - | - | - | - | - | - | - | - | - | - | - | - | - | - | - | - | - | - | - | - | - | - | - | - | - | - | - | - | - | - | - | - | - | - | - | - | - | - | - | - | - | - | - | - | - | - | - | - | - | - | - | - | - | - | - | - | - | - | - | - | - | - | - | - | - | - | - | - | - | - | - | - | - | - | - | - | - | - | - | - | - | - | - | - | - | - | - | - | - | - | - | - | - | - | - | - | - | - | - | - | - | - | - | - | - | - | - | - | - | - | - | - | - | - | - | - | - | - | - | - | - | - | - | - | - | - | - | - | - | - | - | - | - | - | - | - | - | - | - | - | - | - | - | - | - | - | - | - | - | - | - | - | - | - | - | - | - | - | - | - | - | - | - | - | - | - | - | - | - | - | - | - | - | - | - | - | - | - | - | - | - | - | - | - | - | - | - | - | - | - | - | - | - | - | - | - | - | - | - | - | - | - | - | - | - | - | - | - | - | - | - | - | - | - | - | - | - | - | - | - | - | - | - | - | - | - | - | - | - | - | - | - | - | - | - | - | - | - | - | - | - | - | - | - | - | - | - | - | - | - | - | - | - | - | - | - | - | - | - | - | - | - | - | - | - | - | - | - | - | - | - | - | - | - | - | - | - | - | - | - | - | - | - | - | - | - | - | - | - | - | - | - | - | - | - | - | - | - | - | - | - | - | - | - | - | - | - | - | - | - | - | - | - | - | - | - | - | - | - | - | - | - | - | - | - | - | - | - | - | - | - | - | - | - | - | - | - | - | - | - | - | - | - | - | - | - | - | - | - | - | - | - | - | - | - | - | - | - | - | - | - | - | - | - | - | - | - | - | - | - | - | - | - | - | - | - | - | - | - | - | - | - | - | - | - | - | - | - | - | - | - | - | - | - | - | - | - | - | - | - | - | - | - | - | - | - | - | - | - | - | - | - | - | - | - | - | - | - | - | - | - | - | - | - | - | - | - | - | - | - | - | - | - | - | - | - | - | - | - | - | - | - | - | - | - | - | - | - | - | - | - | - | - | - | - | - | - | - | - | - | - | - | - | - | - | - | - | - | - | - | - | - | - | - | - | - | - | - | - | - | - | - | - | - | - | - | - | - | - | - | - | - | - | - | - | - | - | - | - | - | - | - | - | - | - | - | - | - | - | - | - | - | - | - | - | - | - | - | - | - | - | - | - | - | - | - | - | - | - | - | - | - | - | - | - | - | - | - | - | - | - | - | - | - | - | - | - | - | - | - | - | - | - | - | - | - | - | - | - | - | - | - | - | - | - | - | - | - | - | - | - | - | - | - | - | - | - | - | - | - | - | - | - | - | - | - | - | - | - | - | - | - | - | - | - | - | - | - | - | - | - | - | - | - | - | - | - | - | - | - | - | - | - | - | - | - | - | - | - | - | - | - | - | - | - | - | - | - | - | - | - | - | - | - | - | - | - | - | - | - | - | - | - | - | - | - | - | - | - | - | - | - | - | - | - | - | - | - | - | - | - | - | - | - | - | - | - | - | - | - | - | - | - | - | - | - | - | - | - | - | - | - | - | - | - | - | - | - | - | - | - | - | - | - | - | - | - | - | - | - | - | - | - | - | - | - | - | - | - | - | - | - | - | - | - | - | - | - | - | - | - | - | - | - | - | - | - | - | - | - | - | - | - | - | - | - | - | - | - | - | - | - | - | - | - | - | - | - | - | - | - | - | - | - | - | - | - | - | - | - | - | - | - | - | - | - | - | - | - | - | - | - | - | - | - | - | - | - | - | - | - | - | - | - | - | - | - | - | - | - | - | - | - | - | - | - | - | - | - | - | - | - | - | - | - | - | - | - | - | - | - | - | - | - | - | - | - | - | - | - | - | - | - | - | - | - | - | - | - | - | - | - | - | - | - | - | - | - | - | - | - | - | - | - | - | - | - | - | - | - | - | - | - | - | - | - | - | - | - | - | - | - | - | - | - | - | - | - | - | - | - | - | - | - | - | - | - | - | - | - | - | - | - | - | - | - | - | - | - | - | - | - | - | - | - | - | - | - | - | - | - | - | - | - | - | - | - | - | - | - | - | - | - | - | - | - | - | - | - | - | - | - | - | - | - | - | - | - | - | - | - | - | - | - | - | - | - | - | - | - | - | - | - | - | - | - | - | - | - | - | - | - | - | - | - | - | - | - | - | - | - | - | - | - | - | - | - | - | - | - | - | - | - | - | - | - | - | - | - | - | - | - | - | - | - | - | - | - | - | - | - | - | - | - | - | - | - | - | - | - | - | - | - | - | - | - | - | - | - | - | - | - | - | - | - | - | - | - | - | - | - | - | - | - | - | - | - | - | - | - | - | - | - | - | - | - | - | - | - | - | - | - | - | - | - | - | - | - | - | - | - | - | - | - | - | - | - | - | - | - | - | - | - | - | - | - | - | - | - | - | - | - | - | - | - | - | - | - | - | - | - | - | - | - | - | - | - | - | - | - | - | - | - | - | - | - | - | - | - | - | - | - | - | - | - | - | - | - | - | - | - | - | - | - | - | - | - | - | - | - | - | - | - | - | - | - | - | - | - | - | - | - | - | - | - | - | - | - | - | - | - | - | - | - | - | - | - | - | - | - | - | - | - | - | - | - | - | - | - | - | - | - | - | - | - | - | - | - | - | - | - | - | - | - | - | - | - | - | - | - | - | - | - | - | - | - | - | - | - | - | - | - | - | - | - | - | - | - | - | - | - | - | - | - | - | - | - | - | - | - | - | - | - | - | - | - | - | - | - | - | - | - | - | - | - | - | - | - | - | - | - | - | - | - | - | - | - | - | - | - | - | - | - | - | - | - | - | - | - | - | - | - | - | - | - | - | - | - | - | - | - | - | - | - | - | - | - | - | - | - | - | - | - | - | - | - | - | - | - | - | - | - | - | - | - | - | - | - | - | - | - | - | - | - | - | - | - | - | - | - | - | - | - | - | - | - | - | - | - | - | - | - | - | - | - | - | - | - | - | - | - | - | - | - | - | - | - | - | - | - | - | - | - | - | - | - | - | - | - | - | - | - | - | - | - | - | - | - | - | - | - | - | - | - | - | - | - | - | - | - | - | - | - | - | - | - | - | - | - | - | - | - | - | - | - | - | - | - | - | - | - | - | - | - | - | - | - | - | - | - | - | - | - | - | - | - | - | - | - | - | - | - | - | - | - | - | - | - | - | - | - | - | - | - | - | - | - | - | - | - | - | - | - | - | - | - | - | - | - | - | - | - | - | - | - | - | - | - | - | - | - | - | - | - | - | - | - | - | - | - | - | - | - | - | - | - | - | - | - | - | - | - | - | - | - | - | - | - | - | - | - | - | - | - | - | - | - | - | - | - | - | - | - | - | - | - | - | - | - | - | - | - | - | - | - | - | - | - | - | - | - | - | - | - | - | - | - | - | - | - | - | - | - | - | - | - | - | - | - | - | - | - | - | - | - | - | - | - | - | - | - | - | - | - | - | - | - | - | - | - | - | - | - | - | - | - | - | - | - | - | - | - | - | - | - | - | - | - | - | - | - | - | - | - | - | - | - | - | - | - | - | - | - | - | - | - | - | - | - | - | - | - | - | - | - | - | - | - | - | - | - | - | - | - | - | - | - | - | - | - | - | - | - | - | - | - | - | - | - | - | - | - | - | - | - | - | - | - | - | - | - | - | - | - | - | - | - | - | - | - | - | - | - | - | - | - | - | - | - | - | - | - | - | - | - | - | - | - | - | - | - | - | - | - | - | - | - | - | - | - | - | - | - | - | - | - | - | - | - | - | - | - | - | - | - | - | - | - | - | - | - | - | - | - | - | - | - | - | - | - | - | - | - | - | - | - | - | - | - | - | - | - | - | - | - | - | - | - | - | - | - | - | - | - | - | - | - | - | - | - | - | - | - | - | - | - | - | - | - | - | - | - | - | - | - | - | - | - | - | - | - | - | - | - | - | - | - | - | - | - | - | - | - | - | - | - | - | - | - | - | - | - | - | - | - | - | - | - | - | - | - | - | - | - | - | - | - | - | - | - | - | - | - | - | - | - | - | - | - | - | - | - | - | - | - | - | - | - | - | - | - | - | - | - | - | - | - | - | - | - | - | - | - | - | - | - | - | - | - | - | - | - | - | - | - | - | - | - | - | - | - | - | - | - | - | - | - | - | - | - | - | - | - | - | - | - | - | - | - | - | - | - | - | - | - | - | - | - | - | - | - | - | - | - | - | - | - | - | - | - | - | - | - | - | - | - | - | - | - | - | - | - | - | - | - | - | - | - | - | - | - | - | - | - | - | - | - | - | - | - | - | - | - | - | - | - | - | - | - | - | - | - | - | - | - | - | - | - | - | - | - | - | - | - | - | - | - | - | - | - | - | - | - | - | - | - | - | - | - | - | - | - | - | - | - | - | - | - | - | - | - | - | - | - | - | - | - | - | - | - | - | - | - | - | - | - | - | - | - | - | - | - | - | - | - | - | - | - | - | - | - | - | - | - | - | - | - | - | - | - | - | - | - | - | - | - | - | - | - | - | - | - | - | - | - | - | - | - | - | - | - | - | - | - | - | - | - | - | - | - | - | - | - | - | - | - | - | - | - | - | - | - | - | - | - | - | - | - | - | - | - | - | - | - | - | - | - | - | - | - | - | - | - | - | - | - | - | - | - | - | - | - | - | - | - | - | - | - | - | - | - | - | - | - | - | - | - | - | - | - | - | - | - | - | - | - | - | - | - | - | - | - | - | - | - | - | - | - | - | - | - | - | - | - | - | - | - | - | - | - | - | - | - | - | - | - | - | - | - | - | - | - | - | - | - | - | - | - | - | - | - | - | - | - | - | - | - | - | - | - | - | - | - | - | - | - | - | - | - | - | - | - | - | - | - | - | - | - | - | - | - | - | - | - | - | - | - | - | - | - | - | - | - | - | - | - | - | - | - | - | - | - | - | - | - | - | - | - | - | - | - | - | - | - | - | - | - | - | - | - | - | - | - | - | - | - | - | - | - | - | - | - | - | - | - | - | - | - | - | - | - | - | - | - | - | - | - | - | - | - | - | - | - | - | - | - | - | - | - | - | - | - | - | - | - | - | - | - | - | - | - | - | - | - | - | - | - | - | - | - | - | - | - | - | - | - | - | - | - | - | - | - | - | - | - | - | - | - | - | - | - | - | - | - | - | - | - | - | - | - | - | - | - | - | - | - | - | - | - | - | - | - | - | - | - | - | - | - | - | - | - | - | - | - | - | - | - | - | - | - | - | - | - | - | - | - | - | - | - | - | - | - | - | - | - | - | - | - | - | - | - | - | - | - | - | - | - | - | - | - | - | - | - | - | - | - | - | - | - | - | - | - | - | - | - | - | - | - | - | - | - | - | - | - | - | - | - | - | - | - | - | - | - | - | - | - | - | - | - | - | - | - | - | - |
| BIS d=0 cl.7 sym=1 env=1 | - | - | - | - | - | - | - | - | - | - | - | - | - | - | - | - | - | - | - | - | - | - | - | - | - | - | - | - | - | - | - | - | - | - | - | - | - | - | - | - | - | - | - | - | - | - | - | - | - | - | - | - | - | - | - | - | - | - | - | - | - | - | - | - | - | - | - | - | - | - | - | - | - | - | - | - | - | - | - | - | - | - | - | - | - | - | - | - | - | - | - | - | - | - | - | - | - | - | - | - | - | - | - | - | - | - | - | - | - | - | - | - | - | - | - | - | - | - | - | - | - | - | - | - | - | - | - | - | - | - | - | - | - | - | - | - | - | - | - | - | - | - | - | - | - | - | - | - | - | - | - | - | - | - | - | - | - | - | - | - | - | - | - | - | - | - | - | - | - | - | - | - | - | - | - | - | - | - | - | - | - | - | - | - | - | - | - | - | - | - | - | - | - | - | - | - | - | - | - | - | - | - | - | - | - | - | - | - | - | - | - | - | - | - | - | - | - | - | - | - | - | - | - | - | - | - | - | - | - | - | - | - | - | - | - | - | - | - | - | - | - | - | - | - | - | - | - | - | - | - | - | - | - | - | - | - | - | - | - | - | - | - | - | - | - | - | - | - | - | - | - | - | - | - | - | - | - | - | - | - | - | - | - | - | - | - | - | - | - | - | - | - | - | - | - | - | - | - | - | - | - | - | - | - | - | - | - | - | H | H | x | x | H | x | x | H | x | x | x | x | x | x | x | x | x | x | x | x | x | x | x | x | x | x | x | x | x | x | x | x | x | x | x | x | x | x | x | x | x | x | x | H | x | x | x | H | x | x | x | x | x | x | x | x | x | x | x | x | x | x | x | x | x | x | x | x | x | x | x | x | x | x | x | x | x | x | x | x | x | x | x | x | x | x | x | x | x | x | x | x | x | x | x | x | x | x | x | x | x | x | x | x | x | x | x | x | H | x | x | H | x | x | x | x | x | x | x | x | H | H | x | x | H | x | x | x | x | x | H | x | x | x | x | x | x | H | x | x | x | x | x | x | x | x | x | x | x | x | x | H | x | x | x | x | x | x | x | x | x | x | x | x | x | x | x | x | x | x | x | x | x | x | x | x | x | x | x | x | x | x | x | x | x | x | H | x | x | x | x | x | x | x | x | x | x | x | x | x | x | x | x | x | x | x | x | x | x | x | x | x | x | x | x | x | x | x | x | x | x | x | x | x | x | x | x | x | x | x | H | x | x | H | x | H | H | - | - | - | - | - | - | - | - | - | - | - | - | - | - | - | - | - | - | - | - | - | - | - | - | - | - | - | - | - | - | - | - | - | - | - | - | - | - | - | - | - | - | - | - | - | - | - | - | - | - | - | - | - | - | - | - | - | - | - | - | - | - | - | - | - | - | - | - | - | - | - | - | - | - | - | - | - | - | - | - | - | - | - | - | - | - | - | - | - | - | - | - | - | - | - | - | - | - | - | - | - | - | - | - | - | - | - | - | - | - | - | - | - | - | - | - | - | - | - | - | - | - | - | - | - | - | - | - | - | - | - | - | - | - | - | - | - | - | - | - | - | - | - | - | - | - | - | - | - | - | - | - | - | - | - | - | - | - | - | - | - | - | - | - | - | - | - | - | - | - | - | - | - | - | - | - | - | - | - | - | - | - | - | - | - | - | - | - | - | - | - | - | - | - | - | - | - | - | - | - | - | - | - | - | - | - | - | - | - | - | - | - | - | - | - | - | - | - | - | - | - | - | - | - | - | - | - | - | - | - | - | - | - | - | - | - | - | - | - | - | - | - | - | - | - | - | - | - | - | - | - | - | - | - | - | - | - | - | - | - | - | - | - | - | - | - | - | - | - | - | - | - | - | - | - | - | - | - | - | - | - | - | - | - | - | - | - | - | - | - | - | - | - | - | - | - | - | - | - | - | - | - | - | - | - | - | - | - | - | - | - | - | - | - | - | - | - | - | - | - | - | - | - | - | - | - | - | - | - | - | - | - | - | - | - | - | - | - | - | - | - | - | - | - | - | - | - | - | - | - | - | - | - | - | - | - | - | - | - | - | - | - | - | - | - | - | - | - | - | - | - | - | - | - | - | - | - | - | - | - | - | - | - | - | - | - | - | - | - | - | - | - | - | - | - | - | - | - | - | - | - | - | - | - | - | - | - | - | - | - | - | - | - | - | - | - | - | - | - | - | - | - | - | - | - | - | - | - | - | - | - | - | - | - | - | - | - | - | - | - | - | - | - | - | - | - | - | - | - | - | - | - | - | - | - | - | - | - | - | - | - | - | - | - | - | - | - | - | - | - | - | - | - | - | - | - | - | - | - | - | - | - | - | - | - | - | - | - | - | - | - | - | - | - | - | - | - | - | - | - | - | - | - | - | - | - | - | - | - | - | - | - | - | - | - | - | - | - | - | - | - | - | - | - | - | - | - | - | - | - | - | - | - | - | - | - | - | - | - | - | - | - | - | - | - | - | - | - | - | - | - | - | - | - | - | - | - | - | - | - | - | - | - | - | - | - | - | - | - | - | - | - | - | - | - | - | - | - | - | - | - | - | - | - | - | - | - | - | - | - | - | - | - | - | - | - | - | - | - | - | - | - | - | - | - | - | - | - | - | - | - | - | - | - | - | - | - | - | - | - | - | - | - | - | - | - | - | - | - | - | - | - | - | - | - | - | - | - | - | - | - | - | - | - | - | - | - | - | - | - | - | - | - | - | - | - | - | - | - | - | - | - | - | - | - | - | - | - | - | - | - | - | - | - | - | - | - | - | - | - | - | - | - | - | - | - | - | - | - | - | - | - | - | - | - | - | - | - | - | - | - | - | - | - | - | - | - | - | - | - | - | - | - | - | - | - | - | - | - | - | - | - | - | - | - | - | - | - | - | - | - | - | - | - | - | - | - | - | - | - | - | - | - | - | - | - | - | - | - | - | - | - | - | - | - | - | - | - | - | - | - | - | - | - | - | - | - | - | - | - | - | - | - | - | - | - | - | - | - | - | - | - | - | - | - | - | - | - | - | - | - | - | - | - | - | - | - | - | - | - | - | - | - | - | - | - | - | - | - | - | - | - | - | - | - | - | - | - | - | - | - | - | - | - | - | - | - | - | - | - | - | - | - | - | - | - | - | - | - | - | - | - | - | - | - | - | - | - | - | - | - | - | - | - | - | - | - | - | - | - | - | - | - | - | - | - | - | - | - | - | - | - | - | - | - | - | - | - | - | - | - | - | - | - | - | - | - | - | - | - | - | - | - | - | - | - | - | - | - | - | - | - | - | - | - | - | - | - | - | - | - | - | - | - | - | - | - | - | - | - | - | - | - | - | - | - | - | - | - | - | - | - | - | - | - | - | - | - | - | - | - | - | - | - | - | - | - | - | - | - | - | - | - | - | - | - | - | - | - | - | - | - | - | - | - | - | - | - | - | - | - | - | - | - | - | - | - | - | - | - | - | - | - | - | - | - | - | - | - | - | - | - | - | - | - | - | - | - | - | - | - | - | - | - | - | - | - | - | - | - | - | - | - | - | - | - | - | - | - | - | - | - | - | - | - | - | - | - | - | - | - | - | - | - | - | - | - | - | - | - | - | - | - | - | - | - | - | - | - | - | - | - | - | - | - | - | - | - | - | - | - | - | - | - | - | - | - | - | - | - | - | - | - | - | - | - | - | - | - | - | - | - | - | - | - | - | - | - | - | - | - | - | - | - | - | - | - | - | - | - | - | - | - | - | - | - | - | - | - | - | - | - | - | - | - | - | - | - | - | - | - | - | - | - | - | - | - | - | - | - | - | - | - | - | - | - | - | - | - | - | - | - | - | - | - | - | - | - | - | - | - | - | - | - | - | - | - | - | - | - | - | - | - | - | - | - | - | - | - | - | - | - | - | - | - | - | - | - | - | - | - | - | - | - | - | - | - | - | - | - | - | - | - | - | - | - | - | - | - | - | - | - | - | - | - | - | - | - | - | - | - | - | - | - | - | - | - | - | - | - | - | - | - | - | - | - | - | - | - | - | - | - | - | - | - | - | - | - | - | - | - | - | - | - | - | - | - | - | - | - | - | - | - | - | - | - | - | - | - | - | - | - | - | - | - | - | - | - | - | - | - | - | - | - | - | - | - | - | - | - | - | - | - | - | - | - | - | - | - | - | - | - | - | - | - | - | - | - | - | - | - | - | - | - | - | - | - | - | - | - | - | - | - | - | - | - | - | - | - | - | - | - | - | - | - | - | - | - | - | - | - | - | - | - | - | - | - | - | - | - | - | - | - | - | - | - | - | - | - | - | - | - | - | - | - | - | - | - | - | - | - | - | - | - | - | - | - | - | - | - | - | - | - | - | - | - | - | - | - | - | - | - | - | - | - | - | - | - | - | - | - | - | - | - | - | - | - | - | - | - | - | - | - | - | - | - | - | - | - | - | - | - | - | - | - | - | - | - | - | - | - | - | - | - | - | - | - | - | - | - | - | - | - | - | - | - | - | - | - | - | - | - | - | - | - | - | - | - | - | - | - | - | - | - | - | - | - | - | - | - | - | - | - | - | - | - | - | - | - | - | - | - | - | - | - | - | - | - | - | - | - | - | - | - | - | - | - | - | - | - | - | - | - | - | - | - | - | - | - | - | - | - | - | - | - | - | - | - | - | - | - | - | - | - | - | - | - | - | - | - | - | - | - | - | - | - | - | - | - | - | - | - | - | - | - | - | - | - | - | - | - | - | - | - | - | - | - | - | - | - | - | - | - | - | - | - | - | - | - | - | - | - | - | - | - | - | - | - | - | - | - | - | - | - | - | - | - | - | - | - | - | - | - | - | - | - | - | - | - | - | - | - | - | - | - | - | - | - | - | - | - | - | - | - | - | - | - | - | - | - | - | - | - | - | - | - | - | - | - | - | - | - | - | - | - | - | - | - | - | - | - | - | - | - | - | - | - | - | - | - | - | - | - | - | - | - | - | - | - | - | - | - | - | - | - | - | - | - | - | - | - | - | - | - | - | - | - | - | - | - | - | - | - | - | - | - | - | - | - | - | - | - | - | - | - | - | - | - | - | - | - | - | - | - | - | - | - | - | - | - | - | - | - | - | - | - | - | - | - | - | - | - | - | - | - | - | - | - | - | - | - | - | - | - | - | - | - | - | - | - | - | - | - | - | - | - | - | - | - | - | - | - | - | - | - | - | - | - | - | - | - | - | - | - | - | - | - | - | - | - | - | - | - | - | - | - | - | - | - | - | - | - | - | - | - | - | - | - | - | - | - | - | - | - | - | - | - | - | - | - | - | - | - | - | - | - | - | - | - | - | - | - | - | - | - | - | - | - | - | - | - | - | - | - | - | - | - | - | - | - | - | - | - | - | - | - | - | - | - | - | - | - | - | - | - | - | - | - | - | - | - | - | - | - | - | - | - | - | - | - | - | - | - | - | - | - | - | - | - | - | - | - | - | - | - | - | - | - | - | - | - | - | - | - | - | - | - | - | - | - | - | - | - | - | - | - | - | - | - | - | - | - | - | - | - | - | - | - | - | - | - | - | - | - | - | - | - | - | - | - | - | - | - | - | - | - | - | - | - | - | - | - | - | - | - | - | - | - | - | - | - | - | - | - | - | - | - | - | - | - | - | - | - | - | - | - | - | - | - | - | - | - | - | - | - | - | - | - | - | - | - | - | - | - | - | - | - | - | - | - | - | - | - | - | - | - | - | - | - | - | - | - | - | - | - | - | - | - | - | - | - | - | - | - | - | - | - | - | - | - | - | - | - | - | - | - | - | - | - | - | - | - | - | - | - | - | - | - | - | - | - | - | - | - | - | - | - | - | - | - | - | - | - | - | - | - | - | - | - | - | - | - | - | - | - | - | - | - | - | - | - | - | - | - | - | - | - | - | - | - | - | - | - | - | - | - | - | - | - | - | - | - | - | - | - | - | - | - | - | - | - | - | - | - | - | - | - | - | - | - | - | - | - | - | - | - | - | - | - | - | - | - | - | - | - | - | - | - | - | - | - | - | - | - | - | - | - | - | - | - | - | - | - | - | - | - | - | - | - | - | - | - | - | - | - | - | - | - | - | - | - | - | - | - | - | - | - | - | - | - | - | - | - | - | - | - | - | - | - | - | - | - | - | - | - | - | - | - | - | - | - | - | - | - | - | - | - | - | - | - | - | - | - | - | - | - | - | - | - | - | - | - | - | - | - | - | - | - | - | - | - | - | - | - | - | - | - | - | - | - | - | - | - | - | - | - | - | - | - | - | - | - | - | - | - | - | - | - | - | - | - | - | - | - | - | - | - | - | - | - | - | - | - | - | - | - | - | - | - | - | - | - | - | - | - | - | - | - | - | - | - | - | - | - | - | - | - | - | - | - | - | - | - | - | - | - | - | - | - | - | - | - | - | - | - | - | - | - | - | - | - | - | - | - | - | - | - | - | - | - | - | - | - | - | - | - | - | - | - | - | - | - | - | - | - | - | - | - | - | - | - | - | - | - | - | - | - | - | - | - | - | - | - | - | - | - | - | - | - | - | - | - | - | - | - | - | - | - | - | - | - | - | - | - | - | - | - | - | - | - | - | - | - | - | - | - | - | - | - | - | - | - | - | - | - | - | - | - | - | - | - | - | - | - | - | - | - | - | - | - | - | - | - | - | - | - | - | - | - | - | - | - | - | - | - | - | - | - | - | - | - | - | - | - | - | - | - | - | - | - | - | - | - | - | - | - | - | - | - | - | - | - | - | - | - | - | - | - | - | - | - | - | - | - | - | - | - | - | - | - | - | - | - | - | - | - | - | - | - | - | - | - | - | - | - | - | - | - | - | - | - | - | - | - | - | - | - | - | - | - | - | - | - | - | - | - | - | - | - | - | - | - | - | - | - | - | - | - | - | - | - | - | - | - | - | - | - | - | - | - | - | - | - | - | - | - | - | - | - | - | - | - | - | - | - | - | - | - | - | - | - | - | - | - | - | - | - | - | - | - | - | - | - | - | - | - | - | - | - | - | - | - | - | - | - | - | - | - | - | - | - | - | - | - | - | - | - | - | - | - | - | - | - | - | - | - | - | - | - | - | - | - | - | - | - | - | - | - | - | - | - | - | - | - | - | - | - | - | - | - | - | - | - | - | - | - | - | - | - | - | - | - | - | - | - | - | - | - | - | - | - | - | - | - | - | - | - | - | - | - | - | - | - | - | - | - | - | - | - | - | - | - | - | - | - | - | - | - | - | - | - | - | - | - | - | - | - | - | - | - | - | - | - | - | - | - | - | - | - | - | - | - | - | - | - | - | - | - | - | - | - | - | - | - | - | - | - | - | - | - | - | - | - | - | - | - | - | - | - | - | - | - | - | - | - | - | - | - | - | - | - | - | - | - | - | - | - | - | - | - | - | - | - | - | - | - | - | - | - | - | - | - | - | - | - | - | - | - | - | - | - | - | - | - | - | - | - | - | - | - | - | - | - | - | - | - | - | - | - | - | - | - | - | - | - | - | - | - | - | - | - | - | - | - | - | - | - | - | - | - | - | - | - | - | - | - | - | - | - | - | - | - | - | - | - | - | - | - | - | - | - | - | - | - | - | - | - | - | - | - | - | - | - | - | - | - | - | - | - | - | - | - | - | - | - | - | - | - | - | - | - | - | - | - | - | - | - | - | - | - | - | - | - | - | - | - | - | - | - | - | - | - | - | - | - | - | - | - | - | - | - | - | - | - | - | - | - | - | - | - | - | - | - | - | - | - | - | - | - | - | - | - | - | - | - | - | - | - | - | - | - | - | - | - | - | - | - | - | - | - | - | - | - | - | - | - | - | - | - | - | - | - | - | - | - | - | - | - | - | - | - | - | - | - | - | - | - | - | - | - | - | - | - | - | - | - | - | - | - | - | - | - | - | - | - | - | - | - | - | - | - | - | - | - | - | - | - | - | - | - | - | - | - | - | - | - | - | - | - | - | - | - | - | - | - | - | - | - | - | - | - | - | - | - | - | - | - | - | - | - | - | - | - | - | - | - | - | - | - | - | - | - | - | - | - | - | - | - | - | - | - | - | - | - | - | - | - | - | - | - | - | - | - | - | - | - | - | - | - | - | - | - | - | - | - | - | - | - | - | - | - | - | - | - | - | - | - | - | - | - | - | - | - | - | - | - | - | - | - | - | - | - | - | - | - | - | - | - | - | - | - | - | - | - | - | - | - | - | - | - | - | - | - | - | - | - | - | - | - | - | - | - | - | - | - | - | - | - | - | - | - | - | - | - | - | - | - | - | - | - | - | - | - | - | - | - | - | - | - | - | - | - | - | - | - | - | - | - | - | - | - | - | - | - | - | - | - | - | - | - | - | - | - | - | - | - | - | - | - | - | - | - | - | - | - | - | - | - | - | - | - | - | - | - | - | - | - | - | - | - | - | - | - | - | - | - | - | - | - | - | - | - | - | - | - | - | - | - | - | - | - | - | - | - | - | - | - | - | - | - | - | - | - | - | - | - | - | - | - | - | - | - | - | - | - | - | - | - | - | - | - | - | - | - | - | - | - | - | - | - | - | - | - | - | - | - | - | - | - | - | - | - | - | - | - | - | - | - | - | - | - | - | - | - | - | - | - | - | - | - | - | - | - | - | - | - | - | - | - | - | - | - | - | - | - | - | - | - | - | - | - | - | - | - | - | - | - | - | - | - | - | - | - | - | - | - | - | - | - | - | - | - | - | - | - | - | - | - | - | - | - | - | - | - | - | - | - | - | - | - | - | - | - | - | - | - | - | - | - | - | - | - | - | - | - | - | - | - | - | - | - | - | - | - | - | - | - | - | - | - | - | - | - | - | - | - | - | - | - | - | - | - | - | - | - | - | - | - | - | - | - | - | - | - | - | - | - | - | - | - | - | - | - | - | - | - | - | - | - | - | - | - | - | - | - | - | - | - | - | - | - | - | - | - | - | - | - | - | - | - | - | - | - | - | - | - | - | - | - | - | - | - | - | - | - | - | - | - | - | - | - | - | - | - | - | - | - | - | - | - | - | - | - | - | - | - | - | - | - | - | - | - | - | - | - | - | - | - | - | - | - | - | - | - | - | - | - | - | - | - | - | - | - | - | - | - | - | - | - | - | - | - | - | - | - | - | - | - | - | - | - | - | - | - | - | - | - | - | - | - | - | - | - | - | - | - | - | - | - | - | - | - | - | - | - | - | - | - | - | - | - | - | - | - | - | - | - | - | - | - | - | - | - | - | - | - | - | - | - | - | - | - | - | - | - | - | - | - | - | - | - | - | - | - | - | - | - | - | - | - | - | - | - | - | - | - | - | - | - | - | - | - | - | - | - | - | - | - | - | - | - | - | - | - | - | - | - | - | - | - | - | - | - | - | - | - | - | - | - | - | - | - | - | - | - | - | - | - | - | - | - | - | - | - | - | - | - | - | - | - | - | - | - | - | - | - | - | - | - | - | - | - | - | - | - | - | - | - | - | - | - | - | - | - | - | - | - | - | - | - | - | - | - | - | - | - | - | - | - | - | - | - | - | - | - | - | - | - | - | - | - | - | - | - | - | - | - | - | - | - | - | - | - | - | - | - | - | - | - | - | - | - | - | - | - | - | - | - | - | - | - | - | - | - | - | - | - | - | - | - | - | - | - | - | - | - | - | - | - | - | - | - | - | - | - | - | - | - | - | - | - | - | - | - | - | - | - | - | - | - | - | - | - | - | - | - | - | - | - | - | - | - | - | - | - | - | - | - | - | - | - | - | - | - | - | - | - | - | - | - | - | - | - | - | - | - | - | - | - | - | - | - | - | - | - | - | - | - | - | - | - | - | - | - | - | - | - | - | - | - | - | - | - | - | - | - | - | - | - | - | - | - | - | - | - | - | - | - | - | - | - | - | - | - | - | - | - | - | - | - | - | - | - | - | - | - | - | - | - | - | - | - | - | - | - | - | - | - | - | - | - | - | - | - | - | - | - | - | - | - | - | - | - | - | - | - | - | - | - | - | - | - | - | - | - | - | - | - | - | - | - | - | - | - | - | - | - | - | - | - | - | - | - | - | - | - | - | - | - | - | - | - | - | - | - | - | - | - | - | - | - | - | - | - | - | - | - | - | - | - | - | - | - | - | - | - | - | - | - | - | - | - | - | - | - | - | - | - | - | - | - | - | - | - | - | - | - | - | - | - | - | - | - | - | - | - | - | - | - | - | - | - | - | - | - | - | - | - | - | - | - | - | - | - | - | - | - | - | - | - | - | - | - | - | - | - | - | - | - | - | - | - | - | - | - | - | - | - | - | - | - | - | - | - | - | - | - | - | - | - | - | - | - | - | - | - | - | - | - | - | - | - | - | - | - | - | - | - | - | - | - | - | - | - | - | - | - | - | - | - | - | - | - | - | - | - | - | - | - | - | - | - | - | - | - | - | - | - | - | - | - | - | - | - | - | - | - | - | - | - | - | - | - | - | - | - | - | - | - | - | - | - | - | - | - | - | - | - | - | - | - | - | - | - | - | - | - | - | - | - | - | - | - | - | - | - | - | - | - | - | - | - | - | - | - | - | - | - | - | - | - | - | - | - | - | - | - | - | - | - | - | - | - | - | - | - | - | - | - | - | - | - | - | - | - | - | - | - | - | - | - | - | - | - | - | - | - | - | - | - | - | - | - | - | - | - | - | - | - | - | - | - | - | - | - | - | - | - | - | - | - | - | - | - | - | - | - | - | - | - | - | - | - | - | - | - | - | - | - | - | - | - | - | - | - | - | - | - | - | - | - | - | - | - | - | - | - | - | - | - | - | - | - | - | - | - | - | - | - | - | - | - | - | - | - | - | - | - | - | - | - | - | - | - | - | - | - | - | - | - | - | - | - | - | - | - | - | - | - | - | - | - | - | - | - | - | - | - | - | - | - | - | - | - | - | - | - | - | - | - | - | - | - | - | - | - | - | - | - | - | - | - | - | - | - | - | - | - | - | - | - | - | - | - | - | - | - | - | - | - | - | - | - | - | - | - | - | - | - | - | - | - | - | - | - | - | - | - | - | - | - | - | - | - | - | - | - | - | - | - | - | - | - | - | - | - | - | - | - | - | - | - | - | - | - | - | - | - | - | - | - | - | - | - | - | - | - | - | - | - | - | - | - | - | - | - | - | - | - | - | - | - | - | - | - | - | - | - | - | - | - | - | - | - | - | - | - | - | - | - | - | - | - | - | - | - | - | - | - | - | - | - | - | - | - | - | - | - | - | - | - | - | - | - | - | - | - | - | - | - | - | - | - | - | - | - | - | - | - | - | - | - | - | - | - | - | - | - | - | - | - | - | - | - | - | - | - | - | - | - | - | - | - | - | - | - | - | - | - | - | - | - | - | - | - | - | - | - | - | - | - | - | - | - | - | - | - | - | - | - | - | - | - | - | - | - | - | - | - | - | - | - | - | - | - | - | - | - | - | - | - | - | - | - | - | - | - | - | - | - | - | - | - | - | - | - | - | - | - | - | - | - | - | - | - | - | - | - | - | - | - | - | - | - | - | - | - | - | - | - | - | - | - | - | - | - | - | - | - | - | - | - | - | - | - | - | - | - | - | - | - | - | - | - | - | - | - | - | - | - | - | - | - | - | - | - | - | - | - | - | - | - | - | - | - | - | - | - | - | - | - | - | - | - | - | - | - | - | - | - | - | - | - | - | - | - | - | - | - | - | - | - | - | - | - | - | - | - | - | - | - | - | - | - | - | - | - | - | - | - | - | - | - | - | - | - | - | - | - | - | - | - | - | - | - | - | - | - | - | - | - | - | - | - | - | - | - | - | - | - | - | - | - | - | - | - | - | - | - | - | - | - | - | - | - | - | - | - | - | - | - | - | - | - | - | - | - | - | - | - | - | - | - | - | - | - | - | - | - | - | - | - | - | - | - | - | - | - | - | - | - | - | - | - | - | - | - | - | - | - | - | - | - | - | - | - | - | - | - | - | - | - | - | - | - | - | - | - | - | - | - | - | - | - | - | - | - | - | - | - | - | - | - | - | - | - | - | - | - | - | - | - | - | - | - | - | - | - | - | - | - | - | - | - | - | - | - | - | - | - | - | - | - | - | - | - | - | - | - | - | - | - | - | - | - | - | - | - | - | - | - | - | - | - | - | - | - | - | - | - | - | - | - | - | - | - | - | - | - | - | - | - | - | - | - | - | - | - | - | - | - | - | - | - | - | - | - | - | - | - | - | - | - | - | - | - | - | - | - | - | - | - | - | - | - | - | - | - | - | - | - | - | - | - | - | - | - | - | - | - | - | - | - |
| BIS d=0 cl.6 sym=1 env=1 | - | - | - | - | - | - | - | - | - | - | - | - | - | - | - | - | - | - | - | - | - | - | - | - | - | - | - | - | - | - | - | - | - | - | - | - | - | - | - | - | - | - | - | - | - | - | - | - | - | - | - | - | - | - | - | - | - | - | - | - | - | - | - | - | - | - | - | - | - | - | - | - | - | - | - | - | - | - | - | - | - | - | - | - | - | - | - | - | - | - | - | - | - | - | - | - | - | - | - | - | - | - | - | - | - | - | - | - | - | - | - | - | - | - | - | - | - | - | - | - | - | - | - | - | - | - | - | - | - | - | - | - | - | - | - | - | - | - | - | - | - | - | - | - | - | - | - | - | - | - | - | - | - | - | - | - | - | - | - | - | - | - | - | - | - | - | - | - | - | - | - | - | - | - | - | - | - | - | - | - | - | - | - | - | - | - | - | - | - | - | - | - | - | - | - | - | - | - | - | - | - | - | - | - | - | - | - | - | - | - | - | - | - | - | - | - | - | - | - | - | - | - | - | - | - | - | - | - | - | - | - | - | - | - | - | - | - | - | - | - | - | - | - | - | - | - | - | - | - | - | - | - | - | - | - | - | - | - | - | - | - | - | - | - | - | - | - | - | - | - | - | - | - | - | - | - | - | - | - | - | - | - | - | - | - | - | - | - | - | - | - | - | - | - | - | - | - | - | - | - | - | - | - | - | - | - | - | - | - | - | - | - | - | - | - | - | - | - | - | - | - | - | - | - | - | - | - | - | - | - | - | - | - | - | - | - | - | - | - | - | - | - | - | - | - | - | - | - | - | - | - | - | - | - | - | - | - | - | - | - | - | - | - | - | - | - | - | - | - | - | - | - | - | - | - | - | - | - | - | - | - | - | - | - | - | - | - | - | - | - | - | - | - | - | - | - | - | - | - | - | - | - | - | - | - | - | - | - | - | - | - | - | - | - | - | - | - | - | - | - | - | - | - | - | - | - | - | - | - | - | - | - | - | - | - | - | - | - | - | - | - | - | - | - | - | - | - | - | - | - | - | - | - | - | - | - | - | - | - | - | - | - | - | - | - | - | - | - | - | - | - | - | - | - | - | - | - | - | - | - | - | - | - | - | - | - | - | - | - | - | - | - | - | - | - | - | - | - | - | - | - | - | - | - | - | - | - | - | - | - | - | - | - | - | - | - | - | - | - | - | - | - | - | - | - | - | - | - | - | - | - | - | - | - | - | - | - | - | - | - | - | - | - | - | - | - | - | - | - | - | - | - | - | - | - | - | - | - | - | - | - | - | - | - | - | - | - | - | - | - | - | - | - | - | - | - | - | - | - | - | - | - | - | - | - | - | - | - | - | - | - | - | - | - | - | - | - | - | - | - | - | - | - | - | - | - | - | - | - | - | - | - | - | - | - | - | - | - | - | - | - | - | - | - | - | - | - | - | - | - | - | - | - | - | - | - | - | - | - | - | - | - | - | - | - | - | - | - | - | - | - | - | - | - | - | - | - | - | - | - | - | - | - | - | - | - | - | - | - | - | - | - | - | - | - | - | - | - | - | - | - | - | - | - | - | - | - | - | - | - | - | - | - | - | - | - | - | - | - | - | - | - | - | - | - | - | - | - | - | - | - | - | - | - | - | - | - | - | - | - | - | - | - | - | - | - | - | - | - | - | - | - | - | - | - | - | - | - | - | - | - | - | - | - | - | - | - | - | - | - | - | - | - | - | - | - | - | - | - | - | - | - | - | - | - | - | - | - | - | - | - | - | - | - | - | - | - | - | - | - | - | - | - | - | - | - | - | - | - | - | - | - | - | - | - | - | - | - | - | - | - | - | - | - | - | - | - | - | - | - | - | - | - | - | - | - | - | - | - | - | - | - | - | - | - | - | - | - | - | - | - | - | - | - | - | - | - | - | - | - | - | - | - | - | - | - | - | - | - | - | - | - | - | - | - | - | - | - | - | - | - | - | - | - | - | - | - | - | - | - | - | - | - | - | - | - | - | - | - | - | - | - | - | - | - | - | - | - | - | - | - | - | - | - | - | - | - | - | - | - | - | - | - | - | - | - | - | - | - | - | - | - | - | - | - | - | - | - | - | - | - | - | - | - | - | - | - | - | - | - | - | - | - | - | - | - | - | - | - | - | - | - | - | - | - | - | - | - | - | - | - | - | - | - | - | - | - | - | - | - | - | - | - | - | - | - | - | - | - | - | - | - | - | - | - | - | - | - | - | - | - | - | - | - | - | - | - | - | - | - | - | - | - | - | - | - | - | - | - | - | - | - | - | - | - | - | - | - | - | - | - | - | - | - | - | - | - | - | - | - | - | - | - | - | - | - | - | - | - | - | - | - | - | - | - | - | - | - | - | - | - | - | - | - | - | - | - | - | - | - | - | - | - | - | - | - | - | - | - | - | - | - | - | - | - | - | - | - | - | - | - | - | - | - | - | - | - | - | - | - | - | - | - | - | - | - | - | - | - | - | - | - | - | - | - | - | - | - | - | - | - | - | - | - | - | - | - | - | - | - | - | - | - | - | - | - | - | - | - | - | - | - | - | - | - | - | - | - | - | - | - | - | - | - | - | - | - | - | - | - | - | - | - | - | - | - | - | - | - | - | - | - | - | - | - | - | - | - | - | - | - | - | - | - | - | - | - | - | - | - | - | - | - | - | - | - | - | - | - | - | - | - | - | - | - | - | - | - | - | - | - | - | - | - | - | - | - | - | - | - | - | - | - | - | - | - | - | - | - | - | - | - | - | - | - | - | - | - | - | - | - | - | - | - | - | - | - | - | - | - | - | - | - | - | - | - | - | - | - | - | - | - | - | - | - | - | - | - | - | - | - | - | - | - | - | - | - | - | - | - | - | - | - | - | - | - | - | - | - | - | - | - | - | - | - | - | - | - | - | - | - | - | - | - | - | - | - | - | - | - | - | - | - | - | - | - | - | - | - | - | - | - | - | - | - | - | - | - | - | - | - | - | - | - | - | - | - | - | - | - | - | - | - | - | - | - | - | - | - | - | - | - | - | - | - | - | - | - | - | - | - | - | - | - | - | - | - | - | - | - | - | - | - | - | - | - | - | - | - | - | - | - | - | - | - | - | - | - | - | - | - | - | - | - | - | - | - | - | - | - | - | - | - | - | - | - | - | - | - | - | - | - | - | - | - | - | - | - | - | - | - | - | - | - | - | - | - | - | - | - | - | - | - | - | - | - | - | - | - | - | - | - | - | - | - | - | - | - | - | - | - | - | - | - | - | - | - | - | - | - | - | - | - | - | - | - | - | - | - | - | - | - | - | - | - | - | - | - | - | - | - | - | - | - | - | - | - | - | - | - | - | - | - | - | - | - | - | - | - | - | - | - | - | - | - | - | - | - | - | - | - | - | - | - | - | - | - | - | - | - | - | - | - | - | - | - | - | - | - | - | - | - | - | - | - | - | - | - | - | - | - | - | - | - | - | - | - | - | - | - | - | - | - | - | - | - | - | - | - | - | - | - | - | - | - | - | - | - | - | - | - | - | - | - | - | - | - | - | - | - | - | - | - | - | - | - | - | - | - | - | - | - | - | - | - | - | - | - | - | - | - | - | - | - | - | - | - | - | - | - | - | - | - | - | - | - | - | - | - | - | - | - | - | - | - | - | - | - | - | - | - | - | - | - | - | - | - | - | - | - | - | - | - | - | - | - | - | - | - | - | - | - | - | - | - | - | - | - | - | - | - | - | - | - | - | - | - | - | - | - | - | - | - | - | - | - | - | - | - | - | - | - | - | - | - | - | - | - | - | - | - | - | - | - | - | - | - | - | - | - | - | - | - | - | - | - | - | - | - | - | - | - | - | - | - | - | - | - | - | - | - | - | - | - | - | - | - | - | - | - | - | - | - | - | - | - | - | - | - | - | - | - | - | - | - | - | - | - | - | - | - | - | - | - | - | - | - | - | - | - | - | - | - | - | - | - | - | - | - | - | - | - | - | - | - | - | - | - | - | - | - | - | - | - | - | - | - | - | - | - | - | - | - | - | - | - | - | - | - | - | - | - | - | - | - | - | - | - | - | - | - | - | - | - | - | - | - | - | - | - | - | - | - | - | - | - | - | - | - | - | - | - | - | - | - | - | - | - | - | - | - | - | - | - | - | - | - | - | - | - | - | - | - | - | - | - | - | - | - | - | - | - | - | - | - | - | - | - | - | - | - | - | - | - | - | - | - | - | - | - | - | - | - | - | - | - | - | - | - | - | - | - | - | - | - | - | - | - | - | - | - | - | - | - | - | - | - | - | - | - | - | - | - | - | - | - | - | - | - | - | - | - | - | - | - | - | - | - | - | - | - | - | - | - | - | - | - | - | - | - | - | - | - | - | - | - | - | - | - | - | - | - | - | - | - | - | - | - | - | - | - | - | - | - | - | - | - | - | - | - | - | - | - | - | - | - | - | - | - | - | - | - | - | - | - | - | - | - | - | - | - | - | - | - | - | - | - | - | - | - | - | - | - | - | - | - | - | - | - | - | - | - | - | - | - | - | - | - | - | - | - | - | - | - | - | - | - | - | - | - | - | - | - | - | - | - | - | - | - | - | - | - | - | - | - | - | - | - | - | - | - | - | - | - | - | - | - | - | - | - | - | - | - | - | - | - | - | - | - | - | - | - | - | - | - | - | - | - | - | - | - | - | - | - | - | - | - | - | - | - | - | - | - | - | - | - | - | - | - | - | - | - | - | - | - | - | - | - | - | - | - | - | - | - | - | - | - | - | - | - | - | - | - | - | - | - | - | - | - | - | - | - | - | - | - | - | - | - | - | - | - | - | - | - | - | - | - | - | - | - | - | - | - | - | - | - | - | - | - | - | - | - | - | - | - | - | - | - | - | - | - | - | - | - | - | - | - | - | - | - | - | - | - | - | - | - | - | - | - | - | - | - | - | - | - | - | - | - | - | - | - | - | - | - | - | - | - | - | - | - | - | - | - | - | - | - | - | - | - | - | - | - | - | - | - | - | - | - | - | - | - | - | - | - | - | - | - | - | - | - | - | - | - | - | - | - | - | - | - | - | - | - | - | - | - | - | - | - | - | - | - | - | - | - | - | - | - | - | - | - | - | - | - | - | - | - | - | - | - | - | - | - | - | - | - | - | - | - | - | - | - | - | - | - | - | - | - | - | - | - | - | - | - | - | - | - | - | - | - | - | - | - | - | - | - | - | - | - | - | - | - | - | - | - | - | - | - | - | - | - | - | - | - | - | - | - | - | - | - | - | - | - | - | - | - | - | - | - | - | - | - | - | - | - | - | - | - | - | - | - | - | - | - | - | - | - | - | - | - | - | - | - | - | - | - | - | - | - | - | - | - | - | - | - | - | - | - | - | - | - | - | - | - | - | - | - | - | - | - | - | - | - | - | - | - | - | - | - | - | - | - | - | - | - | - | - | - | - | - | - | - | - | - | - | - | - | - | - | - | - | - | - | - | - | - | - | - | - | - | - | - | - | - | - | - | - | - | - | - | - | - | - | - | - | - | - | - | - | - | - | - | - | - | - | - | - | - | - | - | - | - | - | - | - | - | - | - | - | - | - | - | - | - | - | - | - | - | - | - | - | - | - | - | - | - | - | - | - | - | - | - | - | - | - | - | - | - | - | - | - | - | - | - | - | - | - | - | - | - | - | - | - | - | - | - | - | - | - | - | - | - | - | - | - | - | - | - | - | - | - | - | - | - | - | - | - | - | - | - | - | - | - | - | - | - | - | - | - | - | - | - | - | - | - | - | - | - | - | - | - | - | - | - | - | - | - | - | - | - | - | - | - | - | - | - | - | - | - | - | - | - | - | - | - | - | - | - | - | - | - | - | - | - | - | - | - | - | - | - | - | - | - | - | - | - | - | - | - | - | - | - | - | - | - | - | - | - | - | - | - | - | - | - | - | - | - | - | - | - | - | - | - | - | - | - | - | - | - | - | - | - | - | - | - | - | - | - | - | - | - | - | - | - | - | - | - | - | - | - | - | - | - | - | - | - | - | - | - | - | - | - | - | - | - | - | - | - | - | - | - | - | - | - | - | - | - | - | - | - | - | - | - | - | - | - | - | - | - | - | - | - | - | - | - | - | - | - | - | - | - | - | - | - | - | - | - | - | - | - | - | - | - | - | - | - | - | - | - | - | - | - | - | - | - | - | - | - | - | - | - | - | - | - | - | - | - | - | - | - | - | - | - | - | - | - | - | - | - | - | - | - | - | - | - | - | - | - | - | - | - | - | - | - | - | - | - | - | - | - | - | - | - | - | - | - | - | - | - | - | - | - | - | - | - | - | - | - | - | - | - | - | - | - | - | - | - | - | - | - | - | - | - | - | - | - | - | - | - | - | - | - | - | - | - | - | - | - | - | - | - | - | - | - | - | - | - | - | - | - | - | - | - | - | - | - | - | - | - | - | - | - | - | - | - | - | - | - | - | - | - | - | - | - | - | - | - | - | - | - | - | - | - | - | - | - | - | - | - | - | - | - | - | - | - | - | - | - | - | - | - | - | - | - | - | - | - | - | - | - | - | - | - | - | - | - | - | - | - | - | - | - | - | - | - | - | - | - | - | - | - | - | - | - | - | - | - | - | - | - | - | - | - | - | - | - | - | - | - | - | - | - | - | - | - | - | - | - | - | - | - | - | - | - | - | - | - | - | - | - | - | - | - | - | - | - | - | - | - | - | - | - | - | - | - | - | - | - | - | - | - | - | - | - | - | - | - | - | - | - | - | - | - | - | - | - | - | - | - | - | - | - | - | - | - | - | - | - | - | - | - | - | - | - | - | - | - | - | - | - | - | - | - | - | - | - | - | - | - | - | - | - | - | - | - | - | - | - | - | - | - | - | - | - | - | - | - | - | - | - | - | - | - | - | - | - | - | - | - | - | - | - | - | - | - | - | - | - | - | - | - | - | - | - | - | - | - | - | - | - | - | - | - | - | - | - | - | - | - | - | - | - | - | - | - | - | - | - | - | - | - | - | - | - | - | - | - | - | - | - | - | - | - | - | - | - | - | - | - | - | - | - | - | - | - | - | - | - | - | - | - | - | - | - | - | - | - | - | - | - | - | - | - | - | - | - | - | - | - | - | - | - | - | - | - | - | - | - | - | - | - | - | - | - | - | - | - | - | - | - | - | - | - | - | - | - | - | - | - | - | - | - | - | - | - | - | - | - | - | - | - | - | - | - | - | - | - | - | - | - | - | - | - | - | - | - | - | - | - | - | - | - | - | - | - | - | - | - | - | - | - | - | - | - | - | - | - | - | - | - | - | - | - | - | - | - | - | - | - | - | - | - | - | - | - | - | - | - | - | - | - | - | - | - | - | - | - | - | - | - | - | - | - | - | - | - | - | - | - | - | - | - | - | - | - | - | - | - | - | - | - | - | - | - | - | - | - | - | - | - | - | - | - | - | - | - | - | - | - | - | - | - | - | - | - | - | - | - | - | - | - | - | - | - | - | - | - | - | - | - | - | - | - | - | - | - | - | - | - | - | - | - | - | - | - | - | - | - | - | - | - | - | - | - | - | - | - | - | - | - | - | - | - | - | - | - | - | - | - | - | - | - | - | - | - | - | - | - | - | - | - | - | - | - | - | - | - | - | - | - | - | - | - | - | - | - | - | - | - | - | - | - | - | - | - | - | - | - | - | - | - | - | - | - | - | - | - | - | - | - | - | - | - | - | - | - | - | - | - | - | - | - | - | - | - | - | - | - | - | - | - | - | - | - | - | - | - | - | - | - | - | - | - | - | - | - | - | - | - | - | - | - | - | - | - | - | - | - | - | - | - | - | - | - | - | - | - | - | - | - | - | - | - | - | - | - | - | - | - | - | - | - | - | - | - | - | - | - | - | - | - | - | - | - | - | - | - | - | - | - | - | - | - | - | - | - | - | - | - | - | - | - | - | - | - | - | - | - | - | - | - | - | - | - | - | - | - | - | - | - | - | - | - | - | - | - | - | - | - | - | - | - | - | - | - | - | - | - | - | - | - | - | - | - | - | - | - | - | - | - | - | - | - | - | - | - | - | - | - | - | - | - | - | - | - | - | - | - | - | - | - | - | - | - | - | - | - | - | - | - | - | - | - | - | - | - | - | - | - | - | - | - | - | - | - | - | - | - | - | - | - | - | - | - | - | - | - | - | - | - | - | - | - | - | - | - | - | - | - | - | - | - | - | - | - | - | - | - | - | - | - | - | - | - | - | - | - | - | - | - | - | - | - | - | - | - | - | - | - | - | - | - | - | - | - | - | - | - | - | - | - | - | - | - | - | - | - | - | - | - | - | - | - | - | - | - | - | - | - | - | - | - | - | - | - | - | - | - | - | - | - | - | - | - | - | - | - | - | - | - | - | - | - | - | - | - | - | - | - | - | - | - | - | - | - | - | - | - | - | - | - | - | - | - | - | - | - | - | - | - | - | - | - | - | - | - | - | - | - | - | - | - | - | - | - | - | - | - | - | - | - | - | - | - | - | - | - | - | - | - | - | - | - | - | - | - | - | - | - | - | - | - | - | - | - | - | - | - | - | - | - | - | - | - | - | - | - | - | - | - | - | - | - | - | - | - | - | - | - | - | - | - | - | - | - | - | - | - | - | - | - | - | - | - | - | - | - | - | - | - | - | - | - | - | - | - | - | - | - | - | - | - | - | - | - | - | - | - | - | - | - | - | - | - | - | - | - | - | - | - | - | - | - | - | - | - | - | - | - | - | - | - | - | - | - | - | - | - | - | - | - | - | - | - | - | - | - | - | - | - | - | - | - | - | - | - | - | - | - | - | - | - | - | - | - | - | - | - | - | - | - | - | - | - | - | - | - | - | - | - | - | - | - | - | - | - | - | - | - | - | - | - | - | - | - | - | - | - | - | - | - | - | - | - | - | - | - | - | - | - | - | - | - | - | - | - | - | - | - | - | - | - | - | - | - | - | - | - | - | - | - | - | - | - | - | - | - | - | - | - | - | - | - | - | - | - | - | - | - | - | - | - | - | - | - | - | - | - | - | - | - | - | - | - | - | - | - | - | - | - | - | - | - | - | - | - | - | - | - | - | - | - | - | - | - | - | - | - | - | - | - | - | - | - | - | - | - | - | - | - | - | - | - | - | - | - | - | - | - | - | - | - | - | - | - | - | - | - | - | - | - | - | - | - | - | - | - | - | - | - | - | - | - | - | - | - | - | - | - | - | - | - | - | - | - | - | - | - | - | - | - | - | - | - | - | - | - | - | - | - | - | - | - | - | - | - | - | - | - | - | - | - | - | - | - | - | - | - | - | - | - | - | - | - | - | - | - | - | - | - | - | - | - | - | - | - | - | - | - | - | - | - | - | - | - | - | - | - | - | - | - | - | - | - | - | - | - | - | - | - | - | - | - | - | - | - | - | - | - | - | - | - | - | - | - | - | - | - | - | - | - | - | - | - | - | - | - | - | - | - | - | - | - | - | - | - | - | - | - | - | - | - | - | - | - | - | - | - | - | - | - | - | - | - | - | - | - | - | - | - | - | - | - | - | - | - | - | - | - | - | - | - | - | - | - | - | - | - | - | - | - | - | - | - | - | - | - | - | - | - | - | - | - | - | - | - | - | - | - | - | - | - | - | - | - | - | - | - | - | - | - | - | - | - | - | - | - | - | - | - | - | - | - | - | - | - | - | - | - | - | - | - | - | - | - | - | - | - | - | - | - | - | - | - | - | - | - | - | - | - | - | - | - | - | - | - | - | - | - | - | - | - | - | - | - | - | - | - | - | - | - | - | - | - | - | - | - | - | - | - | - | - | - | - | - | - | - | - | - | - | - | - | - | - | - | - | - | - | - | - | - | - | - | - | - | - | - | - | - | - | - | - | - | - | - | - | - | - | - | - | - | - | - | - | - | - | - | - | - | - | - | - | - | - | - | - | - | - | - | - | - | - | - | - | - | - | - | - | - | - | - | - | - | - | - | - | - | - | - | - | - | - | - | - | - | - | - | - | - | - | - | - | - | - | - | - | - | - | - | - | - | - | - | - | - | - | - | - | - | - | - | - | - | - | - | - | - | - | - | - | - | - | - | - | - | - | - | - | - | - | - | - | - | - | - | - | - | - | - | - | - | - | - | - | - | - | - | - | - | - | - | - | - | - | - | - | - | - | - | - | - | - | - | - | - | - | - | - | - | - | - | - | - | - | - | - | - | - | - | - | - | - | - | - | - | - | - | - | - | - | - | - | - | - | - | - | - | - | - | - | - | - | - | - | - | - | - | - | - | - | - | - | - | - | - | - | - | - | - | - | - | - | - | - | - | - | - | - | - | - | - | - | - | - | - | - | - | - | - | - | - | - | - | - | - | - | - | - | - | - | - | - | - | - | - | - | - | - | - | - | - | - | - | - | - | - | - | - | - | - | - | - | - | - | - | - | - | - | - | - | - | - | - | - | - | - | - | - | - | - | - | - | - | - | - | - | - | - | - | - | - | - | - | - | - | - | - | - | - | - | - | - | - | - | - | - | - | - | - | - | - | - | - | - | - | - | - | - | - | - | - | - | - | - | - | - | - | - | - | - | - | - | - | - | - | - | - | - | - | - | - | - | - | - | - | - | - | - | - | - | - | - | - | - | - | - | - | - | - | - | - | - | - | - | - | - | - | - | - | - | - | - | - | - | - | - | - | - | - | - | - | - | - | - | - | - | - | - | - | - | - | - | - | - | - | - | - | - | - | - | - | - | - | - | - | - | - | - | - | - | - | - | - | - | - | - | - | - | - | - | - | - | - | - | - | - | - | - | - | - | - | - | - | - | - | - | - | - | - | - | - | - | - | - | - | - | - | - | - | - | - | - | - | - | - | - | - | - | - | - | - | - | - | - | - | - | - | - | - | - | - | - | - | - | - | - | - | - | - | - | - | - | - | - | - | - | - | - | - | - | - | - | - | - | - | - | - | - | - | - | - | - | - | - | - | - | - | - | - | - | - | - | - | - | - | - | - | - | - | - | - | - | - | - | - | - | - | - | - | - | - | - | - | - | - | - | - | - | - | - | - | - | - | - | - | - | - | - | - | - | - | - | - | - | - | - | - | - | - | - | - | - | - | - | - | - | - | - | - | - | - | - | - | - | H | H | H | H | H | H | H | x | x | x | x | x | x | x | x | x | H | x | x | H | x | H | x | H | x | x | x | H | x | H | x | x | x | x | x | x | H | x | x | x | x | x | x | x | H | x | x | H | H | H | H | H | H | H | H | H | H | x | x | x | H | - | - | - | - | - | - | - | - | - | - | - | - | - | - | - | - | - | - | - | - | - | - | - | - | - | - | - | - | - | - | - | - | - | - | - | - | - | - | - | - | - | - | - | - | - | - | - | - | - | - | - | - | - | - | - | - | - | - | - | - | - | - | - | - | - | - | - | - | - | - | - | - | - | - | - | - | - | - | - | - | - | - | - | - | - | - | - | - | - | - | - | - | - | - | - | - | - | - | - | - | - | - | - | - | - | - | - | - | - | - | - | - | - | - | - | - | - | - | - | - | - | - | - | - | - | - | - | - | - | - | - | - | - | - | - | - | - | - | - | - | - | - | - | - | - | - | - | - | - | - | - | - | - | - | - | - | - | - | - | - | - | - | - | - | - | - | - | - | - | - | - | - | - | - | - | - | - | - | - | - | - | - | - | - | - | - | - | - | - | - | - | - | - | - | - | - | - | - | - | - | - | - | - | - | - | - | - | - | - | - | - | - | - | - | - | - | - | - | - | - | - | - | - | - | - | - | - | - | - | - | - | - | - | - | - | - | - | - | - | - | - | - | - | - | - | - | - | - | - | - | - | - | - | - | - | - | - | - | - | - | - | - | - | - | - | - | - | - | - | - | - | - | - | - | - | - | - | - | - | - | - | - | - | - | - | - | - | - | - | - | - | - | - | - | - | - | - | - | - | - | - | - | - | - | - | - | - | - | - | - | - | - | - | - | - | - | - | - | - | - | - | - | - | - | - | - | - | - | - | - | - | - | - | - | - | - | - | - | - | - | - | - | - | - | - | - | - | - | - | - | - | - | - | - | - | - | - | - | - | - | - | - | - | - | - | - | - | - | - | - | - | - | - | - | - | - | - | - | - | - | - | - | - | - | - | - | - | - | - | - | - | - | - | - | - | - | - | - | - | - | - | - | - | - | - | - | - | - | - | - | - | - | - | - | - | - | - | - | - | - | - | - | - | - | - | - | - | - | - | - | - | - | - | - | - | - | - | - | - | - | - | - | - | - | - | - | - | - | - | - | - | - | - | - |
| BIS d=0 cl.2 sym=1 env=1 | H | x | x | x | x | x | x | x | x | x | x | x | x | x | x | x | x | x | x | x | x | x | x | x | x | x | x | x | x | x | x | x | x | x | x | x | x | x | x | x | x | x | x | x | x | x | x | x | x | x | x | x | x | x | x | x | x | x | x | x | x | x | x | x | x | x | x | x | x | x | x | x | x | x | x | x | x | x | x | x | x | x | x | x | x | x | x | x | x | x | x | x | x | x | x | x | x | x | x | x | x | x | x | x | x | x | x | x | x | x | x | x | x | x | x | x | x | x | x | x | x | x | x | x | x | x | x | x | x | x | x | x | x | x | x | x | x | x | x | x | x | x | x | x | x | x | x | x | x | x | x | x | x | x | x | x | x | x | x | x | x | x | x | x | x | x | x | x | x | x | x | x | x | x | x | x | x | x | x | x | x | x | x | x | x | x | x | x | x | x | x | x | x | x | x | x | x | x | x | x | x | x | x | x | x | x | x | x | x | x | x | x | x | x | x | x | x | x | x | x | x | x | x | x | x | x | x | x | x | x | x | x | x | x | x | x | x | x | x | x | x | x | x | x | x | x | x | x | x | x | x | x | x | x | x | x | x | x | x | x | x | x | x | x | x | x | x | x | x | x | x | x | x | x | x | x | x | x | x | x | x | x | x | x | x | x | x | x | x | x | x | x | x | x | x | x | x | x | x | x | x | x | x | x | x | x | x | x | x | x | x | x | x | x | x | x | x | x | x | x | x | x | x | x | x | x | x | x | x | x | x | x | x | x | x | x | x | x | x | x | x | x | x | x | x | x | x | x | x | x | x | x | x | x | x | x | x | x | x | x | x | x | x | x | x | x | x | x | x | x | x | x | x | x | x | x | x | x | x | x | x | x | x | x | x | x | x | x | x | x | x | x | x | x | x | x | x | x | x | x | x | x | x | x | x | x | x | x | x | x | x | x | x | x | x | x | x | x | x | x | x | x | x | x | x | x | x | x | x | x | x | x | x | x | x | x | x | x | x | x | x | x | x | x | x | x | x | x | x | x | x | x | x | x | x | x | x | x | x | x | x | x | x | x | x | x | x | x | x | x | x | x | x | x | x | x | x | x | x | x | x | x | x | x | x | x | x | x | x | x | x | x | x | x | x | x | x | x | x | x | x | x | x | x | x | x | x | x | x | x | x | x | x | x | x | x | x | x | x | x | x | x | x | x | x | x | x | x | x | x | x | x | x | x | x | x | x | x | x | x | x | x | x | x | x | x | x | x | x | x | x | x | x | x | x | x | x | x | x | x | x | x | x | x | x | x | x | x | x | x | x | x | x | x | x | x | x | x | x | x | x | x | x | x | x | x | x | x | x | x | x | x | x | x | x | x | x | x | x | x | x | x | x | x | x | x | x | x | x | x | x | x | x | x | x | x | x | x | x | x | x | x | x | x | x | x | x | x | x | x | x | x | x | x | x | x | x | x | x | x | x | x | x | x | x | x | x | x | x | x | x | x | x | x | x | x | x | x | x | x | x | x | x | x | x | x | x | x | x | x | x | x | x | x | x | x | x | x | x | x | x | x | x | x | x | x | x | x | x | x | x | x | x | x | x | x | x | x | x | x | x | x | x | x | x | x | x | x | x | x | x | x | x | x | x | x | x | x | x | x | x | x | x | x | x | x | x | x | x | x | x | x | x | x | x | x | x | x | x | x | x | x | x | x | x | x | x | x | x | x | x | x | x | x | x | x | x | x | x | x | x | x | x | x | x | x | x | x | x | x | x | x | x | x | x | x | x | x | x | x | x | x | x | x | x | x | x | x | x | x | x | x | x | x | x | x | x | x | x | x | x | x | x | x | x | x | x | x | x | x | x | x | x | x | x | x | x | x | x | x | x | x | x | x | x | x | x | x | x | x | x | x | x | x | x | x | x | x | x | x | x | x | x | x | x | x | x | x | x | x | x | x | x | x | x | x | x | x | x | x | x | x | x | x | x | x | x | x | x | x | x | x | x | x | x | x | x | x | x | x | x | x | x | x | x | x | x | x | x | x | x | x | x | x | x | x | x | x | x | x | x | x | x | x | x | x | x | x | x | x | x | x | x | x | x | x | x | x | x | x | x | x | x | x | x | x | x | x | x | x | x | x | x | x | x | x | x | x | x | x | x | x | x | x | x | x | x | x | x | x | x | x | x | x | x | x | x | x | x | x | x | x | x | x | x | x | x | x | x | x | x | x | x | x | x | x | x | x | x | x | x | x | x | x | x | x | x | x | x | x | x | x | x | x | x | x | x | x | x | x | x | x | x | x | x | x | x | x | x | x | x | x | x | x | x | x | x | x | x | x | x | x | x | x | x | x | x | x | x | x | x | x | x | x | x | x | x | x | x | x | x | x | x | x | x | x | x | x | x | x | x | x | x | x | x | x | x | x | x | x | x | x | x | x | x | x | x | x | x | x | x | x | x | x | x | x | x | x | x | x | x | x | x | x | x | x | x | x | x | x | x | x | x | x | x | x | x | x | x | x | x | x | x | x | x | x | x | x | x | x | x | x | x | x | x | x | x | x | x | x | x | x | x | x | x | x | x | x | x | x | x | x | x | x | x | x | x | x | x | x | x | x | x | x | x | x | x | x | x | x | x | x | x | x | x | x | x | x | x | x | x | x | x | x | x | x | x | x | x | x | x | x | x | x | x | x | x | x | x | x | x | x | x | x | x | x | x | x | x | x | x | x | x | x | x | x | x | x | x | x | x | x | x | x | x | x | x | x | x | x | x | x | x | x | x | x | x | x | x | x | x | x | x | x | x | x | x | x | x | x | x | x | x | x | x | x | x | x | x | x | x | x | x | x | x | x | x | x | x | x | x | x | x | x | x | x | x | x | x | x | x | x | x | x | x | x | x | x | x | x | x | x | x | x | x | x | x | x | x | x | x | x | x | x | x | x | x | x | x | x | x | x | x | x | x | x | x | x | x | x | x | x | x | x | x | x | x | x | x | x | x | x | x | x | x | x | x | x | x | x | x | x | x | x | x | x | x | x | x | x | x | x | x | x | x | x | x | x | x | x | x | x | x | x | x | x | x | x | x | x | x | x | x | x | x | x | x | x | x | x | x | x | x | x | x | x | x | x | x | x | x | x | x | x | x | x | x | x | x | x | x | x | x | x | x | x | x | x | x | x | x | x | x | x | x | x | x | x | x | x | x | x | x | x | x | x | x | x | x | x | x | x | x | x | x | x | x | x | x | x | x | x | x | x | x | x | x | x | x | x | x | x | x | x | x | x | x | x | x | x | x | x | x | x | x | x | x | x | x | x | x | x | x | x | x | x | x | x | x | x | x | x | x | x | x | x | x | x | x | x | x | x | x | x | x | x | x | x | x | x | x | x | x | x | x | x | x | x | x | x | x | x | x | x | x | x | x | x | x | x | x | x | x | x | x | x | x | x | x | x | x | x | x | x | x | x | x | x | x | x | x | x | x | x | x | x | x | x | x | x | x | x | x | x | x | x | x | x | x | x | x | x | x | x | x | x | x | x | x | x | x | x | x | x | x | x | x | x | x | x | x | x | x | x | x | x | x | x | x | x | x | x | x | x | x | x | x | x | x | x | x | x | x | x | x | x | x | x | x | x | x | x | x | x | x | x | x | x | x | x | x | x | x | x | x | x | x | x | x | x | x | x | x | x | x | x | x | x | x | x | x | x | x | x | x | x | x | x | x | x | x | x | x | x | x | x | x | x | x | x | x | x | x | x | x | x | x | x | x | x | x | x | x | x | x | x | x | x | x | x | x | x | x | x | x | x | x | x | x | x | x | x | x | x | x | x | x | x | x | x | x | x | x | x | x | x | x | x | x | x | x | x | x | x | x | x | x | x | x | x | x | x | x | x | x | x | x | x | x | x | x | x | x | x | x | x | x | x | x | x | x | x | x | x | x | x | x | x | x | x | x | x | x | x | x | x | x | x | x | x | x | x | x | x | x | x | x | x | x | x | x | x | x | x | x | x | x | x | x | x | x | x | x | x | x | x | x | x | x | x | x | x | x | x | x | x | x | x | x | x | x | x | x | x | x | x | x | x | x | x | x | x | x | x | x | x | x | x | x | x | x | x | x | x | x | x | x | x | x | x | x | x | x | x | x | x | x | x | x | x | x | x | x | x | x | x | x | x | x | x | x | x | x | x | x | x | x | x | x | x | x | x | x | x | x | x | x | x | x | x | x | x | x | x | x | x | x | x | x | x | x | x | x | x | x | x | x | x | x | x | x | x | x | x | x | x | x | x | x | x | x | x | x | x | x | x | x | x | x | x | x | x | x | x | x | x | x | x | x | x | x | x | x | x | x | x | x | x | x | x | x | x | x | x | x | x | x | x | x | x | x | x | x | x | x | x | x | x | x | x | x | x | x | x | x | x | x | x | x | x | x | x | x | x | x | x | x | x | x | x | x | x | x | x | x | x | x | x | x | x | x | x | x | x | x | x | x | x | x | x | x | x | x | x | x | x | H | H | x | x | x | x | x | x | x | x | x | x | x | x | x | x | x | x | x | x | x | x | x | x | x | x | x | x | x | x | x | x | x | x | x | x | x | x | x | x | x | x | x | x | x | x | x | x | x | x | x | x | x | x | x | x | x | x | x | x | x | x | x | x | x | x | x | x | x | x | x | x | x | x | x | x | x | x | x | x | x | x | x | x | x | x | x | x | x | x | x | x | x | x | x | x | x | x | x | x | x | x | x | x | x | x | x | x | x | x | x | x | x | x | x | x | x | x | x | x | x | x | x | x | x | x | x | x | x | x | x | x | x | x | x | x | x | x | x | x | x | x | x | x | x | x | x | x | x | x | x | x | x | x | x | x | x | x | x | x | x | x | x | x | x | x | x | x | x | x | x | x | x | x | x | x | x | x | x | x | x | x | x | x | x | x | x | x | x | x | x | x | x | x | x | x | x | x | x | x | x | x | x | x | x | x | x | x | x | x | x | x | x | x | x | x | x | x | x | x | x | x | x | x | x | x | x | x | x | x | x | x | x | x | x | x | x | x | x | x | x | x | x | x | x | x | x | x | x | x | x | x | x | x | x | x | x | x | x | x | x | x | x | x | x | x | x | x | x | x | x | x | x | x | x | x | x | x | x | x | x | x | x | x | x | x | x | x | x | x | x | x | x | x | x | x | x | x | x | x | x | x | x | x | x | x | x | x | x | x | x | x | x | x | x | x | x | x | x | x | x | x | x | x | x | x | x | x | x | x | x | x | x | x | x | x | x | x | x | x | x | x | x | x | x | x | x | x | x | x | x | x | x | x | x | x | x | x | x | x | x | x | x | x | x | x | x | x | x | x | x | x | x | x | x | x | x | x | x | x | x | x | x | x | x | x | x | x | x | x | x | x | H | H | H | x | x | x | x | x | x | x | x | x | x | x | x | x | H | x | x | x | x | x | x | x | x | x | x | x | x | x | x | x | x | x | x | x | x | x | x | x | x | x | x | x | x | x | x | x | x | x | x | x | x | x | x | x | x | x | x | x | x | x | x | x | x | x | x | x | x | x | x | x | x | x | x | x | x | x | x | x | x | x | x | x | x | x | x | x | x | x | x | x | x | x | x | x | x | x | x | x | x | x | x | x | x | x | x | x | x | x | x | x | x | x | x | x | x | x | x | x | x | x | x | x | x | x | x | x | x | x | x | x | x | x | x | x | x | x | x | x | x | x | x | x | x | x | x | x | x | x | x | x | x | x | x | x | x | x | x | x | x | x | x | x | x | x | x | x | x | x | x | x | x | x | x | x | x | x | x | x | x | x | x | x | x | x | x | x | x | x | x | x | x | x | x | x | x | x | x | x | x | x | x | x | x | x | x | x | x | x | x | x | x | x | x | x | x | x | x | x | x | x | x | x | x | x | x | x | x | x | x | x | x | x | x | x | x | x | x | x | x | x | x | x | x | x | x | x | x | x | x | x | x | x | x | x | x | x | x | x | x | x | x | x | x | x | x | x | x | x | x | x | x | x | x | x | x | x | x | x | x | x | x | x | x | x | x | x | x | x | x | x | x | x | x | x | x | x | x | x | x | x | x | x | x | x | x | x | x | x | x | x | x | x | x | x | x | x | x | x | x | x | x | x | x | x | x | x | x | x | x | x | x | x | x | x | x | x | x | x | x | x | x | x | x | x | x | x | x | x | x | x | x | x | x | x | x | x | x | x | x | x | x | x | x | x | x | x | x | x | x | x | x | x | x | x | x | x | x | x | x | x | x | x | x | x | x | x | x | x | x | x | x | x | x | x | x | x | x | x | x | x | x | x | x | x | x | x | x | x | x | x | x | x | x | x | x | x | x | x | x | x | x | x | x | x | x | x | x | x | x | x | x | x | x | x | x | x | x | x | x | x | x | x | x | x | x | x | x | x | x | x | x | x | H | x | x | x | x | x | x | x | x | x | x | x | x | x | x | x | x | x | x | x | x | x | x | x | x | x | x | x | x | x | x | x | x | x | x | x | x | x | x | x | x | x | x | x | x | x | x | x | x | H | x | x | x | x | x | x | x | x | x | x | x | H | x | x | x | x | x | x | x | x | x | x | x | x | x | x | x | x | x | x | x | x | x | x | x | x | x | x | x | x | x | x | x | x | x | x | x | x | x | x | x | x | H | H | x | x | x | x | x | x | x | x | x | x | x | x | x | x | x | x | x | x | x | x | x | x | x | x | x | x | x | x | x | x | x | x | x | x | H | x | x | x | x | H | x | x | x | x | x | x | x | x | x | x | x | x | x | x | x | x | x | x | x | x | x | x | x | x | x | x | x | x | x | x | x | x | x | x | x | x | x | x | x | x | x | x | x | H | x | x | x | x | x | x | x | x | x | x | x | x | x | x | x | x | x | x | x | x | x | x | x | x | x | x | x | x | x | x | x | x | x | x | x | x | x | x | x | x | x | x | x | x | x | x | x | x | x | x | x | x | x | x | x | x | x | x | x | x | x | x | x | x | x | x | x | x | x | x | x | x | x | x | x | x | x | x | x | x | x | x | x | x | x | x | x | x | x | x | x | x | x | x | x | x | x | x | x | x | x | x | x | x | x | x | x | x | x | x | x | x | x | x | x | x | x | x | x | x | x | x | x | x | x | x | x | x | x | x | x | x | x | x | x | x | x | x | x | x | x | x | x | x | x | x | x | x | x | x | x | x | x | x | x | x | x | x | x | x | x | x | x | x | x | x | x | x | x | x | x | x | x | x | x | x | x | x | x | x | x | x | x | x | x | x | x | x | x | x | x | x | x | x | x | x | x | x | x | x | x | x | x | x | x | x | x | x | x | x | x | x | x | x | x | x | x | x | x | x | x | x | x | x | x | x | x | x | x | x | x | x | x | x | x | x | x | x | x | x | x | x | x | x | x | x | x | x | x | x | x | x | x | x | x | x | x | x | x | x | x | x | x | x | x | x | x | x | x | x | x | x | x | x | x | x | x | x | x | x | x | x | x | x | x | x | x | x | x | x | x | x | x | x | x | x | x | x | x | x | x | x | x | x | x | x | x | x | x | x | x | x | x | x | x | x | x | x | x | x | x | x | x | x | x | x | x | x | x | x | x | x | x | x | x | x | x | x | x | x | x | x | x | x | x | x | x | x | x | x | x | x | x | x | x | x | x | x | x | x | x | x | x | x | x | x | x | x | x | x | x | x | x | x | x | x | x | x | x | x | x | x | x | x | x | x | x | x | x | x | x | x | x | x | x | x | x | x | x | x | x | x | x | x | x | x | x | x | x | x | x | x | x | x | x | x | x | x | x | x | x | x | x | x | x | x | x | x | x | x | x | x | x | x | x | x | x | x | x | x | x | x | x | x | x | x | x | x | x | x | x | x | x | x | x | x | x | x | x | x | x | x | x | x | x | x | x | x | x | x | x | x | x | x | x | x | x | x | x | x | x | x | x | x | x | x | x | x | x | x | x | x | x | x | x | x | x | x | x | x | x | x | x | x | x | x | x | x | x | x | x | x | x | x | x | x | x | x | x | x | x | x | x | x | x | x | x | x | x | x | x | x | x | x | x | x | x | x | x | x | x | x | x | x | x | x | x | x | x | x | x | x | x | x | x | x | x | x | x | x | x | x | x | x | x | x | x | x | x | x | x | x | x | x | x | x | x | x | x | x | x | x | x | x | x | x | x | x | x | x | x | x | x | x | x | x | x | x | x | x | x | x | x | x | x | x | x | x | x | x | x | x | x | x | x | x | x | x | x | x | x | x | x | x | x | x | x | x | x | x | x | x | x | x | x | x | x | x | x | x | x | x | x | x | x | x | x | x | x | x | x | x | x | x | x | x | x | x | x | x | x | x | x | x | x | x | x | x | x | x | x | x | x | x | x | x | x | x | x | x | x | x | x | x | x | x | x | x | x | x | x | x | x | x | x | x | x | x | x | x | x | x | x | x | x | x | x | x | x | x | x | x | x | x | x | x | x | x | x | x | x | x | x | x | x | x | x | x | x | x | x | x | x | x | x | x | x | x | x | x | x | x | x | x | x | x | x | x | x | x | x | x | x | x | x | x | x | x | x | x | x | x | x | x | x | x | x | x | x | x | x | x | x | x | x | x | x | x | x | x | x | x | x | x | x | x | x | x | x | x | x | x | x | x | x | x | x | x | x | x | x | x | x | x | x | x | x | x | x | x | x | x | x | x | x | x | x | x | x | x | x | x | x | x | x | x | x | x | x | x | x | x | x | x | x | x | x | x | x | x | x | x | x | x | x | x | x | x | x | x | x | x | x | x | x | x | x | x | x | x | x | x | x | x | x | x | x | x | x | x | x | x | x | x | x | x | x | x | x | x | x | x | x | x | x | x | x | x | x | x | x | x | x | x | x | x | x | x | x | x | x | x | x | x | x | x | x | x | x | x | x | x | x | x | x | x | x | x | x | x | x | x | x | x | x | x | x | x | x | x | x | x | x | x | x | x | x | x | x | x | x | x | x | x | x | x | x | x | x | x | x | x | x | x | x | x | x | x | x | x | x | x | x | x | x | x | x | x | x | x | x | x | x | x | x | x | x | x | x | x | x | x | x | x | x | x | x | x | x | x | x | x | x | x | x | x | x | x | x | x | x | x | x | x | x | x | x | x | x | x | x | x | x | x | x | x | x | x | x | x | x | x | x | x | x | x | x | x | x | x | x | x | x | x | x | x | x | x | x | x | x | x | x | x | x | x | x | x | x | x | x | x | x | x | x | x | x | x | x | x | x | x | x | x | x | x | x | x | x | x | x | x | x | x | x | x | x | x | x | x | x | x | x | x | x | x | x | x | x | x | x | x | x | x | x | x | x | x | H | x | x | x | x | x | x | x | x | x | x | x | x | x | x | x | x | x | x | x | x | x | x | x | x | x | x | x | x | x | x | x | x | x | x | x | x | x | x | x | x | x | x | x | x | x | x | x | x | x | x | x | x | x | x | x | x | x | x | x | x | x | x | x | x | x | x | x | x | x | x | x | x | x | x | x | x | x | x | x | x | x | x | x | x | x | x | x | x | x | x | x | x | x | x | x | x | x | x | x | x | x | x | x | x | x | x | x | x | x | x | x | x | x | x | x | x | x | x | x | x | x | x | x | x | x | x | x | x | x | x | x | x | x | x | x | x | x | x | x | x | x | x | x | x | x | x | x | x | x | x | x | x | x | x | x | x | x | x | x | x | x | x | x | x | x | x | x | x | x | x | x | x | x | x | x | x | x | x | x | x | x | x | x | x | x | x | x | x | x | x | x | x | x | x | x | x | x | x | x | x | x | x | x | x | x | x | x | x | x | x | x | x | x | x | x | x | x | x | x | x | x | x | x | x | x | x | x | x | x | x | x | x | x | x | x | x | x | x | x | x | x | x | x | x | x | x | x | x | x | x | x | x | x | x | x | x | x | x | x | x | x | x | x | x | x | x | x | x | x | x | x | x | x | x | x | x | x | x | x | x | x | x | x | x | x | x | x | x | x | x | x | x | x | x | x | x | x | x | x | x | x | x | x | x | x | x | x | x | x | x | x | x | x | x | x | x | x | x | x | x | x | x | x | x | x | x | x | x | x | x | x | x | x | x | x | x | x | x | x | x | x | x | x | x | x | x | x | x | x | x | x | x | x | x | x | x | x | x | x | x | x | x | x | x | x | x | x | x | x | x | x | x | x | x | x | x | x | x | x | x | x | x | x | x | x | x | x | x | x | x | x | x | x | x | x | x | x | x | x | x | x | x | x | x | x | x | x | x | x | x | x | x | x | x | x | x | x | x | x | x | x | x | x | x | x | x | x | x | x | x | x | x | x | x | x | x | x | x | x | x | x | x | x | x | x | x | x | x | x | x | x | x | x | x | x | x | x | x | x | x | x | x | x | x | x | x | x | x | x | x | x | x | x | x | x | x | x | x | x | x | x | x | x | x | x | x | x | x | x | x | x | x | x | x | x | x | x | x | x | x | x | x | x | x | x | x | x | x | x | x | x | x | x | x | x | x | x | x | x | x | x | x | x | x | x | x | x | x | x | x | x | x | x | x | x | x | x | x | x | x | x | x | x | x | x | x | x | x | x | x | x | x | x | x | x | x | x | x | x | x | x | x | x | x | x | x | x | x | x | x | x | x | x | x | x | x | x | x | x | x | x | x | x | x | x | x | x | x | x | x | x | x | x | x | x | x | x | x | x | x | x | x | x | x | x | x | x | x | x | x | x | x | x | x | x | x | x | x | x | x | x | x | x | x | x | x | x | x | x | x | x | x | x | x | x | x | x | x | x | x | x | x | x | x | x | x | x | x | x | x | x | x | x | x | x | x | x | x | x | x | x | x | x | x | x | x | x | x | x | x | x | x | x | x | x | x | x | x | x | x | x | x | x | x | x | x | x | x | x | x | x | x | x | x | x | x | x | x | x | x | x | x | x | x | x | x | x | x | x | x | x | x | x | x | x | x | x | x | x | x | x | x | x | x | x | x | x | x | x | x | x | x | x | x | x | x | x | x | x | x | x | x | x | x | x | x | x | x | x | x | x | x | x | x | x | x | x | x | x | x | x | x | x | x | x | x | x | x | x | x | x | x | x | x | x | x | x | x | x | x | x | x | x | x | x | x | x | x | x | x | x | x | x | x | x | x | x | x | x | x | x | x | x | x | x | x | x | x | x | x | x | x | x | x | x | x | x | x | x | x | x | x | x | x | x | x | x | x | x | x | x | x | x | x | x | x | x | x | x | x | x | x | x | x | x | x | x | x | x | x | x | x | x | x | x | x | x | x | x | x | x | x | x | x | x | x | x | x | x | x | x | x | x | x | x | x | x | x | x | x | x | x | x | x | x | x | x | x | x | x | x | x | x | x | x | x | x | x | x | x | x | x | x | x | x | x | x | x | x | x | x | x | x | x | x | x | x | x | x | x | x | x | x | x | x | x | x | x | x | x | x | x | x | x | x | x | x | x | x | x | x | x | x | x | x | x | x | x | x | x | x | x | x | x | x | x | x | x | x | x | x | x | x | x | x | x | x | x | x | x | x | x | x | x | x | x | x | x | x | x | x | x | x | x | x | x | x | x | x | x | x | x | x | x | x | x | x | x | x | x | x | x | x | x | x | x | x | x | x | x | x | x | x | x | x | x | x | x | x | x | x | x | x | x | x | x | x | x | x | x | x | x | x | x | x | x | x | x | x | x | x | x | x | x | x | x | x | x | x | x | x | x | x | x | x | x | x | x | x | x | x | x | x | x | x | x | x | x | x | x | x | x | x | x | x | x | x | x | x | x | x | x | x | x | x | x | x | x | x | x | x | x | x | x | x | x | x | x | x | x | x | x | x | x | x | x | x | x | x | x | x | x | x | x | x | x | x | x | x | x | x | x | x | x | x | x | x | x | x | x | x | x | x | x | x | x | x | x | x | x | x | x | x | x | x | x | x | x | x | x | x | x | x | x | x | x | H | x | x | x | x | x | x | x | x | x | x | x | x | x | x | x | x | x | x | x | x | x | x | x | x | x | x | x | x | x | x | x | x | x | x | x | x | x | x | x | x | x | x | x | x | x | x | x | x | x | x | x | x | x | x | x | x | x | x | x | H | x | H | x | x | H | H | x | H | H | H | x | x | H | x | x | H | x | H | x | x | x | H | H | H | x | H | H | H | H | x | x | x | H | H | x | x | x | x | x | x | H | x | H | H | x | x | H | H | x | x | x | x | x | x | H | H | x | x | x | x | x | x | x | x | x | H | H | H | - | - | - | - | - | - | - | - | - | - | - | - | - | - | - | - | - | - | - | - | - | - | - | - | - | - | - | - | - | - | - | - | - | - |
| BIS d=0 cl.21 sym=0.55 env=0.83 | - | - | - | - | - | - | - | - | - | - | - | - | - | - | - | - | - | - | - | - | - | - | - | - | - | - | - | - | - | - | - | - | - | - | - | - | - | - | - | - | - | - | - | - | - | - | - | - | - | - | - | - | - | - | - | - | - | - | - | - | - | - | - | - | - | - | - | - | - | - | - | - | - | - | - | - | - | - | - | - | - | - | - | - | - | - | - | - | - | - | - | - | - | - | - | - | - | - | - | - | - | - | - | - | - | - | - | - | - | - | - | - | - | - | - | - | - | - | - | - | - | - | - | - | - | - | - | - | - | - | - | - | - | - | - | - | - | - | - | - | - | - | - | - | - | - | - | - | - | - | - | - | - | - | - | - | - | - | - | - | - | - | - | - | - | - | - | - | - | - | - | - | - | - | - | - | - | - | - | - | - | - | - | - | - | - | - | - | - | - | - | - | - | - | - | - | - | - | - | - | - | - | - | - | - | - | - | - | - | - | - | - | - | - | - | - | - | - | - | - | - | - | - | - | - | - | - | - | - | - | - | - | - | - | - | - | - | - | - | - | - | - | - | - | - | - | - | - | - | - | - | - | - | - | - | - | - | - | - | - | - | - | - | - | - | - | - | - | - | - | - | - | - | - | - | - | - | - | - | - | - | - | - | - | - | - | - | - | - | - | - | - | - | - | - | - | - | - | - | - | - | - | - | - | - | - | - | - | - | - | - | - | - | - | - | - | - | - | - | - | - | - | - | - | - | - | - | - | - | - | - | - | - | - | - | - | - | - | - | - | - | - | - | - | - | - | - | - | - | - | - | - | - | - | - | - | - | - | - | - | - | - | - | - | - | - | - | - | - | - | - | - | - | - | - | - | - | - | - | - | - | - | - | - | - | - | - | - | - | - | - | - | - | - | - | - | - | - | - | - | - | - | - | - | - | - | - | - | - | - | - | - | - | - | - | - | - | - | - | - | - | - | - | - | - | - | - | - | - | - | - | - | - | - | - | - | - | - | - | - | - | - | - | - | - | - | - | - | - | - | - | - | - | - | - | - | - | - | - | - | - | - | - | - | - | - | - | - | - | - | - | - | - | - | - | - | - | - | - | - | - | - | - | - | - | - | - | - | - | - | - | - | - | - | - | - | - | - | - | - | - | - | - | - | - | - | - | - | - | - | - | - | - | - | - | - | - | - | - | - | - | - | - | - | - | - | - | - | - | - | - | - | - | - | - | - | - | - | - | - | - | - | - | - | - | - | - | - | - | - | - | - | - | - | - | - | - | - | - | - | - | - | - | - | - | - | - | - | - | - | - | - | - | - | - | - | - | - | - | - | - | - | - | - | - | - | - | - | - | - | - | - | - | - | - | - | - | - | - | - | - | - | - | - | - | - | - | - | - | - | - | - | - | - | - | - | - | - | - | - | - | - | - | - | - | - | - | - | - | - | - | - | - | - | - | - | - | - | - | - | - | - | - | - | - | - | - | - | - | - | - | - | - | - | - | - | - | - | - | - | - | - | - | - | - | - | - | - | - | - | - | - | - | - | - | - | - | - | - | - | - | - | - | - | - | - | - | - | - | - | - | - | - | - | - | - | - | - | - | - | - | - | - | - | - | - | - | - | - | - | - | - | - | - | - | - | - | - | - | - | - | - | - | - | - | - | - | - | - | - | - | - | - | - | - | - | - | - | - | - | - | - | - | - | - | - | - | - | - | - | - | - | - | - | - | - | - | - | - | - | - | - | - | - | - | - | - | - | - | - | - | - | - | - | - | - | - | - | - | - | - | - | - | - | - | - | - | - | - | - | - | - | - | - | - | - | - | - | - | - | - | - | - | - | - | - | - | - | - | - | - | - | - | - | - | - | - | - | - | - | - | - | - | - | - | - | - | - | - | - | - | - | - | - | - | - | - | - | - | - | - | - | - | - | - | - | - | - | - | - | - | - | - | - | - | - | - | - | - | - | - | - | - | - | - | - | - | - | - | - | - | - | - | - | - | - | - | - | - | - | - | - | - | - | - | - | - | - | - | - | - | - | - | - | - | - | - | - | - | - | - | - | - | - | - | - | - | - | - | - | - | - | - | - | - | - | - | - | - | - | - | - | - | - | - | - | - | - | - | - | - | - | - | - | - | - | - | - | - | - | - | - | - | - | - | - | - | - | - | - | - | - | - | - | - | - | - | - | - | - | - | - | - | - | - | - | - | - | - | - | - | - | - | - | - | - | - | - | - | - | - | - | - | - | - | - | - | - | - | - | - | - | - | - | - | - | - | - | - | - | - | - | - | - | - | - | - | - | - | - | - | - | - | - | - | - | - | - | - | - | - | - | - | - | - | - | - | - | - | - | - | - | - | - | - | - | - | - | - | - | - | - | - | - | - | - | - | - | - | - | - | - | - | - | - | - | - | - | - | - | - | - | - | - | - | - | - | - | - | - | - | - | - | - | - | - | - | - | - | - | - | - | - | - | - | - | - | - | - | - | - | - | - | - | - | - | - | - | - | - | - | - | - | - | - | - | - | - | - | - | - | - | - | - | - | - | - | - | - | - | - | - | - | - | - | - | - | - | - | - | - | - | - | - | - | - | - | - | - | - | - | - | - | - | - | - | - | - | - | - | - | - | - | - | - | - | - | - | - | - | - | - | - | - | - | - | - | - | - | - | - | - | - | - | - | - | - | - | - | - | - | - | - | - | - | - | - | - | - | - | - | - | - | - | - | - | - | - | - | - | - | - | - | - | - | - | - | - | - | - | - | - | - | - | - | - | - | - | - | - | - | - | - | - | - | - | - | - | - | - | - | - | - | - | - | - | - | - | - | - | - | - | - | - | - | - | - | - | - | - | - | - | - | - | - | - | - | - | - | - | - | - | - | - | - | - | - | - | - | - | - | - | - | - | - | - | - | - | - | - | - | - | - | - | - | - | - | - | - | - | - | - | - | - | - | - | - | - | - | - | - | - | - | - | - | - | - | - | - | - | - | - | - | - | - | - | - | - | - | - | - | - | - | - | - | - | - | - | - | - | - | - | - | - | - | - | - | - | - | - | - | - | - | - | - | - | - | - | - | - | - | - | - | - | - | - | - | - | - | - | - | - | - | - | - | - | - | - | - | - | - | - | - | - | - | - | - | - | - | - | - | - | - | - | - | - | - | - | - | - | - | - | - | - | - | - | - | - | - | - | - | - | - | - | - | - | - | - | - | - | - | - | - | - | - | - | - | - | - | - | - | - | - | - | - | - | - | - | - | - | - | - | - | - | - | - | - | - | - | - | - | - | - | - | - | - | - | - | - | - | - | - | - | - | - | - | - | - | - | - | - | - | - | - | - | - | - | - | - | - | - | - | - | - | - | - | - | - | - | - | - | - | - | - | - | - | - | - | - | - | - | - | - | - | - | - | - | - | - | - | - | - | - | - | - | - | - | - | - | - | - | - | - | - | - | - | - | - | - | - | - | - | - | - | - | - | - | - | - | - | - | - | - | - | - | - | - | - | - | - | - | - | - | - | - | - | - | - | - | - | - | - | - | - | - | - | - | - | - | - | - | - | - | - | - | - | - | - | - | - | - | - | - | - | - | - | - | - | - | - | - | - | - | - | - | - | - | - | - | - | - | - | - | - | - | - | - | - | - | - | - | - | - | - | - | - | - | - | - | - | - | - | - | - | - | - | - | - | - | - | - | - | - | - | - | - | - | - | - | - | - | - | - | - | - | - | - | - | - | - | - | - | - | - | - | - | - | - | - | - | - | - | - | - | - | - | - | - | - | - | - | - | - | - | - | - | - | - | - | - | - | - | - | - | - | - | - | - | - | - | - | - | - | - | - | - | - | - | - | - | - | - | - | - | - | - | - | - | - | - | - | - | - | - | - | - | - | - | - | - | - | - | - | - | - | - | - | - | - | - | - | - | - | - | - | - | - | - | - | - | - | - | - | - | - | - | - | - | - | - | - | - | - | - | - | - | - | - | - | - | - | - | - | - | - | - | - | - | - | - | - | - | - | - | - | - | - | - | - | - | - | - | - | - | - | - | - | - | - | - | - | - | - | - | - | - | - | - | - | - | - | - | - | - | - | - | - | - | - | - | - | - | - | - | - | - | - | - | - | - | - | - | - | - | - | - | - | - | - | - | - | - | - | - | - | - | - | - | - | - | - | - | - | - | - | - | - | - | - | - | - | - | - | - | - | - | - | - | - | - | - | - | - | - | - | - | - | - | - | - | - | - | - | - | - | - | - | - | - | - | - | - | - | - | - | - | - | - | - | - | - | - | - | - | - | - | - | - | - | - | - | - | - | - | - | - | - | - | - | - | - | - | - | - | - | - | H | x | x | x | x | x | x | x | x | x | x | x | x | x | x | x | x | x | x | x | x | x | x | x | x | x | x | x | x | x | x | x | x | x | x | x | x | x | x | x | H | x | H | x | x | x | x | x | x | x | x | x | x | x | x | x | x | x | x | x | x | x | x | x | x | x | x | x | x | x | x | x | x | x | x | x | x | x | x | x | x | x | x | x | x | x | x | x | x | x | x | x | x | x | x | x | x | x | x | x | x | x | x | x | x | x | x | x | x | x | x | x | x | x | x | x | x | x | x | x | x | x | x | x | x | x | x | x | x | x | x | x | x | x | x | x | x | x | x | x | x | x | x | x | x | x | x | x | x | x | x | x | x | x | x | x | x | x | x | x | x | x | x | x | x | x | x | x | x | x | x | x | x | x | x | x | x | x | x | x | x | x | x | x | x | x | x | x | x | x | x | x | x | x | x | x | x | x | x | x | x | x | x | x | x | x | x | x | x | x | x | x | x | x | x | x | x | x | x | x | x | x | x | x | x | x | x | x | x | x | x | x | x | x | x | x | x | x | x | x | x | x | x | x | x | x | x | x | x | x | x | x | x | x | x | x | x | x | x | x | x | x | x | x | x | x | x | x | x | x | x | x | x | x | x | x | x | x | x | x | x | x | x | x | x | x | x | x | x | x | x | x | x | x | x | x | x | x | x | x | x | x | x | x | x | x | x | x | x | x | x | x | x | x | x | x | x | x | x | x | x | x | x | x | x | x | x | x | x | x | x | x | x | x | x | x | x | x | x | x | x | x | x | x | x | x | x | x | x | x | x | x | x | x | x | x | x | x | x | x | x | x | x | x | x | x | x | x | x | x | x | x | x | x | x | x | x | x | x | x | x | x | x | x | x | x | x | x | x | x | x | x | x | x | x | x | x | x | x | x | x | x | x | x | x | x | x | x | x | x | x | x | x | x | x | x | x | x | x | x | x | x | x | x | x | x | x | x | x | x | x | x | x | x | x | x | x | x | x | x | x | x | x | x | x | x | x | x | x | x | x | x | x | x | x | x | x | x | x | x | x | x | x | x | x | x | x | x | x | x | x | x | x | x | x | x | x | x | x | x | x | x | x | x | x | x | x | x | x | x | x | x | x | x | x | x | x | x | x | x | x | x | x | x | x | x | x | x | x | x | x | x | x | x | x | x | x | x | x | x | x | x | x | x | x | x | x | x | x | x | x | x | x | x | x | x | x | x | x | x | x | x | x | x | x | x | x | x | x | x | x | x | x | x | x | x | x | x | x | x | x | x | x | x | x | x | x | x | x | x | x | x | x | x | x | x | x | x | x | x | x | x | x | x | x | x | x | x | x | x | x | x | x | x | x | x | x | x | x | x | x | x | x | x | x | x | x | x | x | x | x | x | x | x | x | x | x | x | x | x | x | x | x | x | x | x | x | x | x | x | x | x | x | x | x | x | x | x | x | x | x | x | x | x | x | x | x | x | x | x | x | x | x | x | x | x | x | x | x | x | x | x | x | x | x | x | x | x | x | x | x | x | x | x | x | x | x | x | x | x | x | x | x | x | x | x | x | x | x | x | x | x | x | x | x | x | x | x | x | x | x | x | x | x | x | x | x | x | x | x | x | x | x | x | x | x | x | x | x | x | x | x | x | x | x | x | x | x | x | x | x | x | x | x | x | x | x | x | x | x | x | x | x | x | x | x | x | x | x | x | x | x | x | x | x | x | x | x | x | x | x | x | x | x | x | x | x | x | x | x | x | x | x | x | x | x | x | x | x | x | x | x | x | x | x | x | x | x | x | x | x | x | x | x | x | x | x | x | x | x | x | x | x | x | x | x | x | x | x | x | x | x | x | x | x | x | x | x | x | x | x | x | x | x | x | x | x | x | x | x | x | x | x | x | x | x | x | x | x | x | x | x | x | x | x | x | x | x | x | x | x | x | x | x | x | x | x | x | x | x | x | x | x | x | x | x | x | x | x | x | x | x | x | x | x | x | x | x | x | x | x | x | x | x | x | x | x | x | x | x | x | x | x | x | x | x | x | x | x | x | x | x | x | x | x | x | x | x | x | x | x | x | x | x | x | x | x | x | x | x | x | x | x | x | x | x | x | x | x | x | x | x | x | x | x | x | x | x | x | x | x | x | x | x | x | x | x | x | x | x | x | x | x | x | x | x | x | x | x | x | x | x | x | x | x | x | x | x | x | x | x | x | x | x | x | x | x | x | x | x | x | x | x | x | x | x | x | x | x | x | x | x | x | x | x | x | x | x | x | x | x | x | x | x | x | x | x | x | x | x | x | x | x | x | x | x | x | x | x | x | x | x | x | x | x | x | x | x | x | x | x | x | x | x | x | x | x | x | x | x | x | x | x | x | x | x | x | x | x | x | x | x | x | x | x | x | x | x | x | x | x | x | x | x | x | x | x | x | x | x | x | x | x | x | x | x | x | x | x | x | x | x | x | x | x | x | x | x | x | x | x | x | x | x | x | x | x | x | x | x | x | x | x | x | x | x | x | x | x | x | x | x | x | x | x | x | x | x | x | x | x | x | x | x | x | x | x | x | x | x | x | x | x | x | x | x | x | x | x | x | x | x | x | x | x | x | x | x | x | x | x | x | x | x | x | x | x | x | x | x | x | x | x | x | x | x | x | x | x | x | x | x | x | x | x | x | x | x | x | x | x | x | x | x | x | x | x | x | x | x | x | x | x | x | x | x | x | x | x | x | x | x | x | x | x | x | x | x | x | x | x | x | x | x | x | x | x | x | x | x | x | x | x | x | x | x | x | x | x | x | x | x | x | x | x | x | x | x | x | x | x | x | x | x | x | x | x | x | x | x | x | x | x | x | x | x | x | x | x | x | x | x | x | x | x | x | x | x | x | x | x | x | x | x | x | x | x | x | x | x | x | x | x | x | x | x | x | x | x | x | x | x | x | x | x | x | x | x | x | x | x | x | x | x | x | x | x | x | x | x | x | x | x | x | x | x | x | x | x | x | x | x | x | x | x | x | x | x | x | x | x | x | x | x | x | x | x | x | x | x | x | x | x | x | x | x | x | x | x | x | x | x | x | x | x | x | x | x | x | x | x | x | x | x | x | x | x | x | x | x | x | x | x | x | x | x | x | x | x | x | x | x | x | x | x | x | x | x | x | x | x | x | x | x | x | x | x | x | x | x | x | x | x | x | x | x | x | x | x | x | x | x | x | x | x | x | x | x | x | x | x | x | x | x | x | x | x | x | x | x | x | x | x | x | x | x | x | x | x | x | x | x | x | x | x | x | x | x | x | x | x | x | x | x | x | x | x | x | x | x | x | x | x | x | x | x | x | x | x | x | x | x | x | x | x | x | x | x | x | x | x | x | x | x | x | x | x | x | x | x | x | x | x | x | x | x | x | x | x | x | x | x | x | x | x | x | x | x | x | x | x | x | x | x | x | x | x | x | x | x | x | x | x | x | x | x | x | x | x | x | x | x | x | x | x | x | x | x | x | x | x | x | x | x | x | x | x | x | x | x | x | x | x | x | x | x | x | x | x | x | x | x | x | x | x | x | x | x | x | x | x | x | x | x | x | x | x | x | x | x | x | x | x | x | x | x | x | x | x | x | x | x | x | x | x | x | x | x | x | x | x | x | x | x | x | x | x | x | x | x | x | x | x | x | x | x | x | x | x | x | x | x | x | x | x | x | x | x | x | x | x | x | x | x | x | x | x | x | x | x | x | x | x | x | x | x | x | x | x | x | x | x | x | x | x | x | x | x | x | x | x | x | x | x | x | x | x | x | x | x | x | x | x | x | x | x | x | x | x | x | x | x | x | x | x | x | x | x | x | x | x | x | x | x | x | x | x | x | x | x | x | x | x | x | x | x | x | x | x | x | x | x | x | x | x | x | x | x | x | x | x | x | x | x | x | x | x | x | x | x | x | x | x | x | x | x | x | x | x | x | x | x | x | x | x | x | x | x | x | x | x | x | x | x | x | x | x | x | x | x | x | x | x | x | x | x | x | x | x | x | x | x | x | x | x | x | x | x | x | x | x | x | x | x | x | x | x | x | x | x | x | x | x | x | x | x | x | x | x | x | x | x | x | x | x | x | x | x | x | x | x | x | x | x | x | x | x | x | x | x | x | x | x | x | x | x | x | x | x | x | x | x | x | x | x | x | x | x | x | x | x | x | x | x | x | x | x | x | x | x | x | x | x | x | x | x | x | x | x | x | x | x | x | x | x | x | x | x | x | x | x | x | x | x | x | x | x | x | x | x | x | x | x | x | x | x | x | x | x | x | x | x | x | x | x | x | x | x | x | x | x | x | x | x | x | x | x | x | x | x | x | x | x | x | x | x | x | x | x | x | x | x | x | x | x | x | x | x | x | x | x | x | x | x | x | x | x | x | x | x | x | x | x | x | x | x | x | x | x | x | x | x | x | x | x | x | x | x | x | x | x | x | x | x | x | x | x | x | x | x | x | x | x | x | x | x | x | x | x | x | x | x | x | x | x | x | x | x | x | x | x | x | x | x | x | x | x | x | x | x | x | x | x | x | x | x | x | x | x | x | x | x | x | x | x | x | x | x | x | x | x | x | x | x | x | x | x | x | x | x | x | x | x | x | x | x | x | x | x | x | x | x | x | x | x | x | x | x | x | x | x | x | x | x | x | x | x | x | x | x | x | x | x | x | x | x | x | x | x | x | x | x | x | x | x | x | x | x | x | x | x | x | x | x | x | x | x | x | x | x | x | x | x | x | x | x | x | x | x | x | x | x | x | x | x | x | x | x | x | x | x | x | x | x | x | x | x | x | x | x | x | x | x | x | x | x | x | x | x | x | x | x | x | x | x | x | x | x | x | x | x | x | x | x | x | x | x | x | x | x | x | x | x | x | x | x | x | x | x | x | x | x | x | x | x | x | x | x | x | x | x | x | x | x | x | x | x | x | x | x | x | x | x | x | x | x | x | x | x | x | x | x | x | x | x | x | x | x | x | x | x | x | x | x | x | x | x | x | x | x | x | x | x | x | x | x | x | x | x | x | x | x | x | x | x | x | x | x | x | x | x | x | x | x | x | x | x | x | x | x | x | x | x | x | x | x | x | x | x | x | x | x | x | x | x | x | x | x | x | x | x | x | x | x | x | x | x | x | x | x | x | x | x | x | x | x | x | x | x | x | x | x | x | x | x | x | x | x | x | x | x | x | x | x | x | x | x | x | x | x | x | x | x | x | x | x | x | x | x | x | x | x | x | x | x | x | x | x | x | x | x | x | x | x | x | x | x | x | x | x | x | x | x | x | x | x | x | x | x | x | x | x | x | x | x | x | x | x | x | x | x | x | x | x | x | x | x | x | x | x | x | x | x | x | x | x | x | x | x | x | x | x | x | x | x | x | x | x | x | x | x | x | x | x | x | x | x | x | x | x | x | x | x | x | x | x | x | x | x | x | x | x | x | x | x | x | x | x | x | x | x | x | x | x | x | x | x | x | x | x | x | x | x | x | x | x | x | x | x | x | x | x | x | x | x | x | x | x | x | x | x | x | x | x | x | x | x | x | x | x | x | x | x | x | x | x | x | x | x | x | x | x | x | x | x | x | x | x | x | x | x | x | x | x | x | x | x | x | x | x | x | x | x | x | x | x | x | x | x | x | x | x | x | x | x | x | x | x | x | x | x | x | x | x | x | x | x | x | x | x | x | x | x | x | x | x | x | x | x | x | x | x | x | x | x | x | x | x | x | x | x | x | x | x | x | x | x | x | x | x | x | x | x | x | x | x | x | x | x | x | x | x | x | x | x | x | x | x | x | x | x | x | x | x | x | x | x | x | x | x | x | x | x | x | x | x | x | x | x | x | x | x | x | x | x | x | x | x | x | x | x | x | x | x | x | x | x | x | x | x | x | x | x | x | x | x | x | x | x | x | x | x | x | x | x | x | x | x | x | x | x | x | x | x | x | x | x | x | x | x | x | x | x | x | x | x | x | x | x | x | x | x | x | x | x | x | x | x | x | x | x | x | x | x | x | x | x | x | x | x | x | x | x | x | x | x | x | x | x | x | x | x | x | x | x | x | x | x | x | x | x | x | x | x | x | x | x | x | x | x | x | x | x | x | x | x | x | x | x | x | x | x | x | x | x | x | x | x | x | x | x | x | x | x | x | x | x | x | x | x | x | x | x | x | x | x | x | x | x | x | x | x | x | x | x | x | x | x | x | x | x | x | x | x | x | x | x | x | x | x | x | x | x | x | x | x | x | x | x | x | x | x | x | x | x | x | x | x | x | x | x | x | x | x | x | x | x | x | x | x | x | x | x | x | x | x | x | x | x | x | x | x | x | x | x | x | x | x | x | x | x | x | x | x | x | x | x | x | x | x | x | x | x | x | x | x | x | x | x | x | x | x | x | x | x | x | x | x | x | x | x | x | x | x | x | x | x | x | x | x | x | x | x | x | x | x | x | x | x | x | x | x | x | x | x | x | x | x | x | x | x | x | x | x | x | x | x | x | x | x | x | x | x | x | x | x | x | x | x | x | x | x | x | x | x | x | x | x | x | x | x | x | x | x | x | x | x | x | x | x | x | x | x | x | x | x | x | x | x | x | x | x | x | x | x | x | x | x | x | x | x | x | x | x | x | x | x | x | x | x | x | x | x | x | x | x | x | x | x | x | x | x | x | x | x | x | x | x | x | x | x | x | x | x | x | x | x | x | x | x | x | x | x | x | x | x | x | x | x | x | x | x | x | x | x | x | x | x | x | x | x | x | x | x | x | x | x | x | x | x | x | x | x | x | x | x | x | x | x | x | x | x | x | x | x | x | x | x | x | x | x | x | x | x | x | x | x | x | x | x | x | x | x | x | x | x | x | x | x | x | x | x | x | x | x | x | x | x | x | x | x | x | x | x | x | x | x | x | x | x | x | x | x | x | x | x | x | x | x | x | x | x | x | x | x | x | x | x | x | x | x | x | x | x | x | x | x | x | x | x | x | x | x | x | x | x | x | x | x | x | x | x | x | x | x | x | x | x | x | x | x | x | x | x | x | x | x | x | x | x | x | x | x | x | x | x | x | x | x | x | x | x | x | x | x | x | x | x | x | x | x | x | x | x | x | x | x | x | x | x | x | x | x | x | x | x | x | x | x | x | x | x | x | x | x | x | x | x | x | x | x | x | x | x | x | x | x | x | x | x | x | x | x | x | x | x | x | x | x | x | x | x | x | x | x | x | x | x | x | x | x | x | x | x | x | x | x | x | x | x | x | x | x | x | x | x | x | x | x | x | x | x | x | x | x | x | x | x | x | x | x | x | x | x | x | x | x | x | x | x | x | x | x | x | x | x | x | x | x | x | x | x | x | x | x | x | x | x | x | x | x | x | x | x | x | x | x | x | x | x | x | x | x | x | x | x | x | x | x | x | x | x | x | x | x | x | x | x | x | x | x | x | x | x | x | x | x | x | x | x | x | x | x | x | x | x | x | x | x | x | x | x | x | x | x | x | x | x | x | x | x | x | x | x | x | x | x | x | x | x | x | x | x | x | x | x | x | x | x | x | x | x | x | x | x | x | x | x | x | x | x | x | x | x | x | x | x | x | x | x | x | x | x | x | x | x | x | x | x | x | x | x | x | x | x | x | x | x | x | x | x | x | x | x | x | x | x | x | x | x | x | x | x | x | x | x | x | x | x | x | x | x | x | x | x | x | x | x | x | x | x | x | x | x | x | x | x | x | x | x | x | x | x | x | x | x | x | x | x | x | x | x | x | x | x | x | x | x | x | x | x | x | x | x | x | x | x | E | H | - | - | - | - | - | - | - | - | - | - | - | - | - | - | - | - | - | - | - | - | - | - | - | - | - | - | - | - | - | - | - | - | - | - | - | - | - | - | - | - | - | - | - | - | - | - | - | - | - | - | - | - | - | - | - | - | - | - | - | - | - | - | - | - | - | - | - | - | - | - | - | - | - | - | - | - | - | - | - | - | - | - | - |
| BIS d=0 cl.22 sym=0.55 env=0.65 | - | - | - | - | - | - | - | - | - | - | - | - | - | - | - | - | - | - | - | - | - | - | - | - | - | - | - | - | - | - | - | - | - | - | - | - | - | - | - | - | - | - | - | - | - | - | - | - | - | - | - | - | - | - | - | - | - | - | - | - | - | - | - | - | - | - | - | - | - | - | - | - | - | - | - | - | - | - | - | - | - | - | - | - | - | - | - | - | - | - | - | - | - | - | - | - | - | - | - | - | - | - | - | - | - | - | - | - | - | - | - | - | - | - | - | - | - | - | - | - | - | - | - | - | - | - | - | - | - | - | - | - | - | - | - | - | - | - | - | - | - | - | - | - | - | - | - | - | - | - | - | - | - | - | - | - | - | - | - | - | - | - | - | - | - | - | - | - | - | - | - | - | - | - | - | - | - | - | - | - | - | - | - | - | - | - | - | - | - | - | - | - | - | - | - | - | - | - | - | - | - | - | - | - | - | - | - | - | - | - | - | - | - | - | - | - | - | - | - | - | - | - | - | - | - | - | - | - | - | - | - | - | - | - | - | - | - | - | - | - | - | - | - | - | - | - | - | - | - | - | - | - | - | - | - | - | - | - | - | - | - | - | - | - | - | - | - | - | - | - | - | - | - | - | - | - | - | - | - | - | - | - | - | - | - | - | - | - | - | - | - | - | - | - | - | - | - | - | - | - | - | - | - | - | - | - | - | - | - | - | - | - | - | - | - | - | - | - | - | - | - | - | - | - | - | - | - | - | - | - | - | - | - | - | - | - | - | - | - | - | - | - | - | - | - | - | - | - | - | - | - | - | - | - | - | - | - | - | - | - | - | - | - | - | - | - | - | - | - | - | - | - | - | - | - | - | - | - | - | - | - | - | - | - | - | - | - | - | - | - | - | - | - | - | - | - | - | - | - | - | - | - | - | - | - | - | - | - | - | - | - | - | - | - | - | - | - | - | - | - | - | - | - | - | - | - | - | - | - | - | - | - | - | - | - | - | - | - | - | - | - | - | - | - | - | - | - | - | - | - | - | - | - | - | - | - | - | - | - | - | - | - | - | - | - | - | - | - | - | - | - | - | - | - | - | - | - | - | - | - | - | - | - | - | - | - | - | - | - | - | - | - | - | - | - | - | - | - | - | - | - | - | - | - | - | - | - | - | - | - | - | - | - | - | - | - | - | - | - | - | - | - | - | - | - | - | - | - | - | - | - | - | - | - | - | - | - | - | - | - | - | - | - | - | - | - | - | - | - | - | - | - | - | - | - | - | - | - | - | - | - | - | - | - | - | - | - | - | - | - | - | - | - | - | - | - | - | - | - | - | - | - | - | - | - | - | - | - | - | - | - | - | - | - | - | - | - | - | - | - | - | - | - | - | - | - | - | - | - | - | - | - | - | - | - | - | - | - | - | - | - | - | - | - | - | - | - | - | - | - | - | - | - | - | - | - | - | - | - | - | - | - | - | - | - | - | - | - | - | - | - | - | - | - | - | - | - | - | - | - | - | - | - | - | - | - | - | - | - | - | - | - | - | - | - | - | - | - | - | - | - | - | - | - | - | - | - | - | - | - | - | - | - | - | - | - | - | - | - | - | - | - | - | - | - | - | - | - | - | - | - | - | - | - | - | - | - | - | - | - | - | - | - | - | - | - | - | - | - | - | - | - | - | - | - | - | - | - | - | - | - | - | - | - | - | - | - | - | - | - | - | - | - | - | - | - | - | - | - | - | - | - | - | - | - | - | - | - | - | - | - | - | - | - | - | - | - | - | - | - | - | - | - | - | - | - | - | - | - | - | - | - | - | - | - | - | - | - | - | - | - | - | - | - | - | - | - | - | - | - | - | - | - | - | - | - | - | - | - | - | - | - | - | - | - | - | - | - | - | - | - | - | - | - | - | - | - | - | - | - | - | - | - | - | - | - | - | - | - | - | - | - | - | - | - | - | - | - | - | - | - | - | - | - | - | - | - | - | - | - | - | - | - | - | - | - | - | - | - | - | - | - | - | - | - | - | - | - | - | - | - | - | - | - | - | - | - | - | - | - | - | - | - | - | - | - | - | - | - | - | - | - | - | - | - | - | - | - | - | - | - | - | - | - | - | - | - | - | - | - | - | - | - | - | - | - | - | - | - | - | - | - | - | - | - | - | - | - | - | - | - | - | - | - | - | - | - | - | - | - | - | - | - | - | - | - | - | - | - | - | - | - | - | - | - | - | - | - | - | - | - | - | - | - | - | - | - | - | - | - | - | - | - | - | - | - | - | - | - | - | - | - | - | - | - | - | - | - | - | - | - | - | - | - | - | - | - | - | - | - | - | - | - | - | - | - | - | - | - | - | - | - | - | - | - | - | - | - | - | - | - | - | - | - | - | - | - | - | - | - | - | - | - | - | - | - | - | - | - | - | - | - | - | - | - | - | - | - | - | - | - | - | - | - | - | - | - | - | - | - | - | - | - | - | - | - | - | - | - | - | - | - | - | - | - | - | - | - | - | - | - | - | - | - | - | - | - | - | - | - | - | - | - | - | - | - | - | - | - | - | - | - | - | - | - | - | - | - | - | - | - | - | - | - | - | - | - | - | - | - | - | - | - | - | - | - | - | - | - | - | - | - | - | - | - | - | - | - | - | - | - | - | - | - | - | - | - | - | - | - | - | - | - | - | - | - | - | - | - | - | - | - | - | - | - | - | - | - | - | - | - | - | - | - | - | - | - | - | - | - | - | - | - | - | - | - | - | - | - | - | - | - | - | - | - | - | - | - | - | - | - | - | - | - | - | - | - | - | - | - | - | - | - | - | - | - | - | - | - | - | - | - | - | - | - | - | - | - | - | - | - | - | - | - | - | - | - | - | - | - | - | - | - | - | - | - | - | - | - | - | - | - | - | - | - | - | - | - | - | - | - | - | - | - | - | - | - | - | - | - | - | - | - | - | - | - | - | - | - | - | - | - | - | - | - | - | - | - | - | - | - | - | - | - | - | - | - | - | - | - | - | - | - | - | - | - | - | - | - | - | - | - | - | - | - | - | - | - | - | - | - | - | - | - | - | - | - | - | - | - | - | - | - | - | - | - | - | - | - | - | - | - | - | - | - | - | - | - | - | - | - | - | - | - | - | - | - | - | - | - | - | - | - | - | - | - | - | - | - | - | - | - | - | - | - | - | - | - | - | - | - | - | - | - | - | - | - | - | - | - | - | - | - | - | - | - | - | - | - | - | - | - | - | - | - | - | - | - | - | - | - | - | - | - | - | - | - | - | - | - | - | - | - | - | - | - | - | - | - | - | - | - | - | - | - | - | - | - | - | - | - | - | - | - | - | - | - | - | - | - | - | - | - | - | - | - | - | - | - | - | - | - | - | - | - | - | - | - | - | - | - | - | - | - | - | - | - | - | - | - | - | - | - | - | - | - | - | - | - | - | - | - | - | - | - | - | - | - | - | - | - | - | - | - | - | - | - | - | - | - | - | - | - | - | - | - | - | - | - | - | - | - | - | - | - | - | - | - | - | - | - | - | - | - | - | - | - | - | - | - | - | - | - | - | - | - | - | - | - | - | - | - | - | - | - | - | - | - | - | - | - | - | - | - | - | - | - | - | - | - | - | - | - | - | - | - | - | - | - | - | - | - | - | - | - | - | - | - | - | - | - | - | - | - | - | - | - | - | - | - | - | - | - | - | - | - | - | - | - | - | - | - | - | - | - | - | - | - | - | - | - | - | - | - | - | - | - | - | - | - | - | - | - | - | - | - | - | - | - | - | - | - | - | - | - | - | - | - | - | - | - | - | - | - | - | - | - | - | - | - | - | - | - | - | - | - | - | - | - | - | - | - | - | - | - | - | - | - | - | - | - | - | - | - | - | - | - | - | - | - | - | - | - | - | - | - | - | - | - | - | - | - | - | - | - | - | - | - | - | - | - | - | - | - | - | - | - | - | - | - | - | - | - | - | - | - | - | - | - | - | - | - | - | - | - | - | - | - | - | - | - | - | - | - | - | - | - | - | - | - | - | - | - | - | - | - | - | - | - | - | - | - | - | - | - | - | - | - | - | - | - | - | - | - | - | - | - | - | - | - | - | - | - | - | - | - | - | - | - | - | - | - | - | - | - | - | - | - | - | - | - | - | - | - | - | - | - | - | - | - | - | - | - | - | - | - | - | - | - | - | - | - | - | - | - | - | - | - | - | - | - | - | - | - | - | - | - | - | - | - | - | - | - | - | - | - | - | - | - | - | - | - | - | - | - | - | - | - | - | - | - | - | - | - | - | - | - | - | - | - | - | - | - | - | - | - | - | - | - | - | - | - | - | - | - | - | - | - | - | - | - | - | - | - | - | - | - | - | - | - | - | - | - | - | - | - | - | - | - | - | - | - | - | - | - | - | - | - | - | - | - | - | - | - | - | - | - | - | - | - | - | - | - | - | - | - | - | - | - | - | - | - | - | - | - | - | - | - | - | - | - | - | - | - | - | - | - | - | - | - | - | - | - | - | - | - | - | - | - | - | - | - | - | - | - | - | - | - | - | - | - | - | - | - | - | - | - | - | - | - | - | - | - | - | - | - | - | - | - | - | - | - | - | - | - | - | - | - | - | - | - | - | - | - | - | - | - | - | - | - | - | - | - | - | - | - | - | - | - | - | - | - | - | - | - | - | - | - | - | - | - | - | - | - | - | - | - | - | - | - | - | - | - | - | - | - | - | - | - | - | - | - | - | - | - | - | - | - | - | - | - | - | - | - | - | - | - | - | - | - | - | - | - | - | - | - | - | - | - | - | - | - | - | - | - | - | - | - | - | - | - | - | - | - | - | - | - | - | - | - | - | - | - | - | - | - | - | - | - | - | - | - | - | - | - | - | - | - | - | - | - | - | - | - | - | - | - | - | - | - | - | - | - | - | - | - | - | - | - | - | - | - | - | - | - | - | - | - | - | - | - | - | - | - | - | - | - | - | - | - | - | - | - | - | - | - | - | - | - | - | - | - | - | - | - | - | - | - | - | - | - | - | - | - | - | - | - | - | - | - | - | - | - | - | - | - | - | - | - | - | - | - | - | - | - | - | - | - | - | - | - | - | - | - | - | - | - | - | - | - | - | - | - | - | - | - | - | - | - | - | - | - | - | - | - | - | - | - | - | - | - | - | - | - | - | - | - | - | - | - | - | - | - | - | - | - | - | - | - | - | - | - | - | - | - | - | - | - | - | - | - | - | - | - | - | - | - | - | - | - | - | - | - | - | - | - | - | - | - | - | - | - | - | - | - | - | - | - | - | - | - | - | - | - | - | - | - | - | - | - | - | - | - | - | - | - | - | - | - | - | - | - | - | - | - | - | - | - | - | - | - | - | - | - | - | - | - | - | - | - | - | - | - | - | - | - | - | - | - | - | - | - | - | - | - | - | - | - | - | - | - | - | - | - | - | - | - | - | - | - | - | - | - | - | - | - | - | - | - | - | - | - | - | - | - | - | - | - | - | - | - | - | - | - | - | - | - | - | - | - | - | - | - | - | - | - | - | - | - | - | - | - | - | - | - | - | - | - | - | - | - | - | - | - | - | - | - | - | - | - | - | - | - | - | - | - | - | - | - | - | - | - | - | - | - | - | - | - | - | - | - | - | - | - | - | - | - | - | - | - | - | - | - | - | - | - | - | - | - | - | - | - | - | - | - | - | - | - | - | - | - | - | - | - | - | - | - | - | - | - | - | - | - | - | - | - | - | - | - | - | - | - | - | - | - | - | - | - | - | - | - | - | - | - | - | - | - | - | - | - | - | - | - | - | - | - | - | - | - | - | - | - | - | - | - | - | - | - | - | - | - | - | - | - | - | - | - | - | - | - | - | - | - | - | - | - | - | - | - | - | - | - | - | - | - | - | - | - | - | - | - | - | - | - | - | - | - | - | - | - | - | - | - | - | - | - | - | - | - | - | - | - | - | - | - | - | - | - | - | - | - | - | - | - | - | - | - | - | - | - | - | - | - | - | - | - | - | - | - | - | - | - | - | - | - | - | - | - | - | - | - | - | - | - | - | - | - | - | - | - | - | - | - | - | - | - | - | - | - | - | - | - | - | - | - | - | - | - | - | - | - | - | - | - | - | - | - | - | - | - | - | - | - | - | - | - | - | - | - | - | - | - | - | - | - | - | - | - | - | - | - | - | - | - | - | - | - | - | - | - | - | - | - | - | - | - | - | - | - | - | - | - | - | - | - | - | - | - | - | - | - | - | - | - | - | - | - | - | - | - | - | - | - | - | - | - | - | - | - | - | - | - | - | - | - | - | - | - | - | - | - | - | - | - | - | - | - | - | - | - | - | - | - | - | - | - | - | - | - | - | - | - | - | - | - | - | - | - | - | - | - | - | - | - | - | - | - | - | - | - | - | - | - | - | - | - | - | - | - | - | - | - | - | - | - | - | - | - | - | - | - | - | - | - | - | - | - | - | - | - | - | - | - | - | - | - | - | - | - | - | - | - | - | - | - | - | - | - | - | - | - | - | - | - | - | - | - | - | - | - | - | - | - | - | - | - | - | - | - | - | - | - | - | - | - | - | - | - | - | - | - | - | - | - | - | - | - | - | - | - | - | - | - | - | - | - | - | - | - | - | - | - | - | - | - | - | - | - | - | - | - | - | - | - | - | - | - | - | - | - | - | - | - | - | - | - | - | - | - | - | - | - | - | - | - | - | - | - | - | - | - | - | - | - | - | - | - | - | - | - | - | - | - | - | - | - | - | - | - | - | - | - | - | - | - | - | - | - | - | - | - | - | - | - | - | - | - | - | - | - | - | - | - | - | - | - | - | - | - | - | - | - | - | - | - | - | - | - | - | - | - | - | - | - | - | - | - | - | - | - | - | - | - | - | - | - | - | - | - | - | - | - | - | - | - | - | - | - | - | - | - | - | - | - | - | - | - | - | - | - | - | - | - | - | - | - | - | - | - | - | - | - | - | - | - | - | - | - | - | - | - | - | - | - | - | - | - | - | - | - | - | - | - | - | - | - | - | - | - | - | - | - | - | - | - | - | - | - | - | - | - | - | - | - | - | - | - | - | - | - | - | - | - | - | - | - | - | - | - | - | - | - | - | - | - | - | - | - | - | - | - | - | - | - | - | - | - | - | - | - | - | - | - | - | - | - | - | - | - | - | - | - | - | - | - | - | - | - | - | - | - | - | - | - | - | - | - | - | - | - | - | - | - | - | - | - | - | - | - | - | - | - | - | - | - | - | - | - | - | - | - | - | - | - | - | - | - | - | - | - | - | - | - | - | - | - | - | - | - | - | - | - | - | - | - | - | - | - | - | - | - | - | - | - | - | - | - | - | - | - | - | - | - | - | - | - | - | - | - | - | - | - | - | - | - | - | - | - | - | - | - | - | - | - | - | - | - | - | - | - | - | - | - | - | - | - | - | - | - | - | - | - | - | - | - | - | - | - | - | - | - | - | - | - | - | - | - | - | - | - | - | - | - | - | - | - | - | - | - | - | - | - | - | - | - | - | - | - | - | - | - | - | - | - | - | - | - | - | - | - | - | - | - | - | - | - | - | - | - | - | - | - | - | - | - | - | - | - | - | - | - | - | - | - | - | - | - | - | - | - | - | - | - | - | - | - | - | - | - | - | - | - | - | - | - | - | - | - | - | - | - | - | - | - | - | - | - | - | - | - | - | - | - | - | - | - | - | - | - | - | - | - | - | - | - | - | - | - | - | - | - | - | - | - | - | - | - | - | - | - | - | - | - | - | - | - | - | - | - | - | - | - | - | - | - | - | - | - | - | - | - | - | - | - | - | - | - | - | - | - | - | - | - | - | - | - | - | - | - | - | - | - | - | - | - | - | - | - | - | - | - | - | - | - | - | - | - | - | - | - | - | - | - | - | - | - | - | - | - | - | - | - | - | - | - | - | - | - | - | - | - | - | - | - | - | - | - | - | - | - | - | - | - | - | - | - | - | - | - | - | - | - | - | - | - | - | - | - | - | - | - | - | - | - | - | - | - | - | - | - | - | - | - | - | - | - | - | - | - | - | - | - | - | - | - | - | - | - | - | - | - | - | - | - | - | - | - | - | - | - | - | - | - | - | - | - | - | - | - | - | - | - | - | - | - | - | - | - | - | - | - | - | - | - | - | - | - | - | - | - | - | - | - | - | - | - | - | - | - | - | - | - | - | - | - | - | - | - | - | - | - | - | - | - | - | - | - | - | - | - | - | - | - | - | - | - | - | - | - | - | - | - | - | - | - | - | - | - | - | - | - | - | - | - | - | - | - | - | - | - | - | - | - | - | - | - | - | - | - | - | - | - | - | - | - | - | - | - | - | - | - | - | - | - | - | - | - | - | - | - | - | - | - | - | - | - | - | - | - | - | - | - | - | - | - | - | - | - | - | - | - | - | - | - | - | - | - | - | - | - | - | - | - | - | - | - | - | - | - | - | - | - | - | - | - | - | - | - | - | - | - | - | - | - | - | - | - | - | - | - | - | - | - | - | - | - | - | - | - | - | - | - | - | - | - | - | - | - | - | - | - | - | - | - | - | - | - | - | - | - | - | - | - | - | - | - | - | - | - | - | - | - | - | - | - | - | - | - | - | - | - | - | - | - | - | - | - | - | - | - | - | - | - | - | - | - | - | - | - | - | - | - | - | - | - | - | - | - | - | - | - | - | - | - | - | - | - | - | - | - | - | - | - | - | - | - | - | - | - | - | - | - | - | - | - | - | - | - | - | - | - | - | - | - | - | - | - | - | - | - | - | - | - | - | - | - | - | - | - | - | - | - | - | - | - | - | - | - | - | - | - | - | - | - | - | - | - | - | - | - | - | - | - | - | - | - | - | - | - | - | - | - | - | - | - | - | - | - | - | - | - | - | - | - | - | - | - | - | - | - | - | - | - | - | - | - | - | - | - | - | - | - | - | - | - | - | - | - | - | - | - | - | - | - | - | - | - | - | - | - | - | - | - | - | - | - | - | - | - | - | - | - | - | - | - | - | - | - | - | - | - | - | - | - | - | - | - | - | - | - | - | - | - | - | - | - | - | - | - | - | - | - | - | - | - | - | - | - | - | - | - | - | - | - | - | - | - | - | - | - | - | - | - | - | - | - | - | - | - | - | - | - | - | - | - | - | - | - | - | - | - | - | - | - | - | - | - | - | - | - | - | - | - | - | - | - | - | - | - | - | - | - | - | - | - | - | - | - | - | - | - | - | - | - | - | - | - | - | - | - | - | - | - | - | - | - | - | - | - | - | - | - | - | - | - | - | - | - | - | - | - | - | - | - | - | - | - | - | - | - | - | - | - | - | - | - | - | - | - | - | - | - | - | - | - | - | - | - | - | - | - | - | - | - | - | - | - | - | - | - | - | - | - | - | - | - | - | - | - | - | - | - | - | - | - | - | - | - | - | - | - | - | - | - | - | - | - | - | - | - | - | - | - | - | - | - | - | - | - | - | - | - | - | - | - | - | - | - | - | - | - | - | - | - | - | - | - | - | - | - | - | - | - | - | - | - | - | - | - | - | - | - | - | - | - | - | - | - | - | - | - | - | - | - | - | - | - | - | - | - | - | - | - | - | - | - | - | - | - | - | - | - | - | - | - | - | - | - | - | - | - | - | - | - | - | - | - | - | - | - | - | - | - | - | - | - | - | - | - | - | - | - | - | - | - | - | - | - | - | - | - | - | - | - | - | - | - | - | - | - | - | - | - | - | - | - | - | - | - | - | - | - | - | - | - | - | - | - | - | - | - | - | - | - | - | - | - | - | - | - | - | - | - | - | - | - | - | - | - | - | - | - | - | - | - | - | - | - | - | - | - | - | - | - | - | - | - | - | - | - | - | - | - | - | - | - | - | - | - | - | - | - | - | - | - | - | - | - | - | - | - | - | - | - | - | - | - | - | - | - | - | - | - | - | - | - | - | - | - | - | - | - | - | - | - | - | - | - | - | - | - | - | - | - | - | - | - | - | - | - | - | - | - | - | - | - | - | - | - | - | - | - | - | - | - | - | - | - | - | - | - | - | - | - | - | - | - | - | - | - | - | - | - | - | - | - | - | - | - | - | - | - | - | - | - | - | - | - | - | - | - | - | - | - | - | - | - | - | - | - | - | - | - | - | - | - | - | - | - | - | - | - | - | - | - | - | - | - | - | - | - | - | - | - | - | - | - | - | - | - | - | - | - | - | - | - | - | - | - | - | - | - | - | - | - | - | - | - | - | - | - | - | - | - | - | - | - | - | - | - | - | - | - | - | - | - | - | - | - | - | - | - | - | - | - | - | - | - | - | - | - | - | - | - | - | - | - | - | - | - | - | - | - | - | - | - | - | - | - | - | - | - | - | - | - | - | - | - | - | - | - | - | - | - | - | - | - | - | - | - | - | - | - | - | - | - | - | - | - | - | - | - | - | - | - | - | - | - | - | - | - | - | - | - | - | - | - | - | - | - | - | - | - | - | - | - | - | - | - | - | - | - | - | - | - | - | - | - | - | - | - | - | - | - | - | - | - | - | - | - | - | - | - | - | - | - | - | - | - | - | - | - | - | - | - | - | - | - | - | - | - | - | - | - | - | - | - | - | - | - | - | - | - | - | - | - | - | - | - | - | - | - | - | - | - | - | - | - | - | - | - | - | - | - | - | - | - | - | - | - | - | - | - | - | - | - | - | - | - | - | - | - | - | - | - | - | - | - | - | - | - | - | - | - | - | - | - | - | - | - | - | - | - | - | - | - | - | - | - | - | - | - | - | - | - | - | - | - | - | - | - | - | - | - | - | - | - | - | - | - | - | - | - | - | - | - | - | - | - | - | - | - | - | - | - | - | - | - | - | - | - | - | - | - | - | - | - | - | - | - | - | - | - | - | - | - | - | - | - | - | - | - | - | - | - | - | - | - | - | - | - | - | - | - | - | - | - | - | - | - | - | - | - | - | - | - | - | - | H | x | x | H | H | H | H | H | x | x | H | x | x | x | x | x | x | x | x | x | x | x | x | x | x | x | x | x | x | x | x | H | x | x | x | x | x | x | x | x | x | H | x | H | x | x | x | x | x | x | x | x | x | x | x | x | x | x | x | x | x | x | x | x | x | x | x | x | x | x | x | x | x | x | x | x | x | x | x | x | x | x | x | x | x | x | x | x | x | x | x | x | x | x | x | x | x | x | x | x | x | x | x | x | x | x | x | x | x | x | x | x | x | x | x | x | x | x | x | x | x | x | x | x | x | x | x | x | x | x | x | x | x | x | x | x | x | x | x | x | x | x | x | x | x | x | x | x | x | x | x | x | x | x | x | x | x | x | x | x | x | x | x | x | x | x | x | x | x | x | x | x | x | x | x | x | x | x | x | x | x | x | x | x | x | x | x | x | x | x | x | x | x | x | x | x | x | x | x | x | x | x | x | x | x | x | x | x | x | x | x | x | x | x | x | x | x | x | x | x | x | x | x | x | x | x | x | x | x | x | x | x | x | x | x | x | x | x | x | x | x | x | x | x | x | x | x | x | x | x | x | x | x | x | x | x | x | x | x | x | x | x | x | x | x | x | x | x | x | x | x | x | x | x | x | x | x | x | x | x | x | x | x | x | x | x | x | x | x | x | x | x | x | x | x | x | x | x | x | x | x | x | x | x | x | x | x | x | x | x | x | x | x | x | x | x | x | x | x | x | x | x | x | x | x | x | x | x | x | x | x | x | x | x | x | x | x | x | x | x | x | x | x | x | x | x | x | x | x | x | x | x | x | x | x | x | x | x | x | x | x | x | x | x | x | x | x | x | x | x | x | x | x | x | x | x | x | x | x | x | x | x | x | x | x | x | x | x | x | x | x | x | x | x | x | x | x | x | x | x | x | x | x | x | x | x | x | x | x | x | x | x | x | x | x | x | x | x | x | x | x | x | x | x | x | x | x | x | x | x | x | x | x | x | x | x | x | x | x | x | x | x | x | x | x | x | x | x | x | x | x | x | x | x | x | x | x | x | x | x | x | x | x | x | x | x | x | x | x | x | x | x | x | x | x | x | x | x | x | x | x | x | x | x | x | x | x | x | x | x | x | x | x | x | x | x | x | x | x | x | x | x | x | x | x | x | x | x | x | x | x | x | x | x | x | x | x | x | x | x | E | E | H | - | - | - | - | - | - | - | - | - | - | - | - | - | - | - | - | - | - | - | - | - | - | - | - | - | - | - | - | - | - | - | - | - | - | - | - | - | - | - | - | - | - | - | - | - | - | - | - | - | - | - | - | - | - | - | - | - | - | - | - | - | - | - | - | - | - | - |
