## Supplementary file 4-Output file BIS2Analyzer in table form with hits associated to dimensions and blocks for "Protein coevolution and physicochemical adaptation in the APAF-1/apoptosome: structural and functional implications": 9) Supplementary file 4-Output file BIS2Analyzer in table form with hits associated to dimensions and blocks.html

BIS cluster table


Clusters with env. score >= 0.5 and sym. score >= 0.5 :

| Dim | Cluster | Sym | Env | Pvalue | Hit patterns and blocks |
| --- | --- | --- | --- | --- | --- |
| 0 | 8 | 1 | 1 | 1.764537e-11 | Hit patterns: Positions: \| 4382 \| 4386 \| 4394 \| 4395 \| 4398 \| 4399; 60 sequences: \| L \| L \| A \| L \| F \| I; 38 sequences: \| L \| L \| A \| F \| F \| L; 9 sequences: \| - \| - \| - \| - \| - \| -; 7 sequences: \| L \| L \| A \| F \| F \| V; 2 sequences: \| L \| L \| A \| L \| F \| V; 1 sequence: \| L \| L \| A \| F \| F \| I; 1 sequence: \| L \| V \| A \| L \| F \| V; 1 sequence: \| L \| I \| A \| F \| L \| V All positions in cluster: 4382 4386 4394-4395 4398-4399 |
| 0 | 9 | 1 | 1 | 4.197756e-08 | Hit patterns: Positions: \| 419 \| 432; 115 sequences: \| E \| R; 2 sequences: \| - \| -; 2 sequences: \| K \| P All positions in cluster: 419-420 432-433 |
| 0 | 5 | 1 | 1 | 1.3966e-07 | Hit patterns: Positions: \| 4925 \| 4928 \| 4929; 112 sequences: \| H \| F \| P; 4 sequences: \| - \| - \| -; 2 sequences: \| H \| I \| P; 1 sequence: \| H \| L \| P All positions in cluster: 4925 4928-4929 |
| 0 | 4 | 1 | 1 | 1.248832e-06 | Hit patterns: Positions: \| 4952 \| 4973; 115 sequences: \| G \| G; 2 sequences: \| - \| -; 1 sequence: \| N \| R; 1 sequence: \| S \| R All positions in cluster: 4952 4973-4974 |
| 0 | 23 | 0.727273 | 0.502513 | 1.315132e-06 | Hit patterns: Positions: \| 1787 \| 1791; 113 sequences: \| F \| D; 2 sequences: \| - \| -; 2 sequences: \| Y \| D; 1 sequence: \| I \| G; 1 sequence: \| F \| G All positions in cluster: 1787-1788 1791 |
| 0 | 10 | 1 | 1 | 2.0002e-06 | Hit patterns: Positions: \| 4004 \| 4006 \| 4013; 98 sequences: \| V \| V \| L; 14 sequences: \| V \| I \| L; 2 sequences: \| - \| - \| -; 2 sequences: \| V \| L \| L; 1 sequence: \| A \| L \| I; 1 sequence: \| F \| V \| M; 1 sequence: \| I \| I \| L All positions in cluster: 4004 4006 4013 |
| 0 | 15 | 1 | 1 | 6.0006e-06 | Hit patterns: Positions: \| 2443 \| 2448 \| 2552; 98 sequences: \| A \| H \| I; 18 sequences: \| A \| H \| V; 3 sequences: \| - \| - \| - All positions in cluster: 2443 2448 2552 |
| 0 | 13 | 1 | 1 | 8.512087e-06 | Hit patterns: Positions: \| 104 \| 123 \| 251 \| 255 \| 261 \| 265 \| 266 \| 271 \| 275 \| 279; 87 sequences: \| R \| L \| L \| I \| I \| M \| I \| L \| E \| V; 7 sequences: \| R \| L \| L \| I \| I \| M \| V \| L \| E \| V; 4 sequences: \| K \| L \| L \| I \| I \| M \| I \| L \| E \| V; 3 sequences: \| R \| L \| L \| I \| V \| M \| I \| L \| E \| V; 3 sequences: \| R \| L \| L \| I \| L \| M \| I \| M \| E \| I; 3 sequences: \| - \| - \| - \| - \| - \| - \| - \| - \| - \| -; 2 sequences: \| R \| L \| L \| I \| L \| M \| I \| L \| E \| I; 2 sequences: \| R \| L \| L \| I \| I \| M \| I \| I \| E \| V; 1 sequence: \| R \| L \| L \| I \| I \| L \| I \| L \| D \| V; 1 sequence: \| R \| L \| L \| I \| L \| M \| I \| M \| E \| V; 1 sequence: \| R \| L \| L \| I \| L \| M \| V \| L \| E \| V; 1 sequence: \| R \| L \| L \| V \| I \| M \| I \| L \| E \| V; 1 sequence: \| R \| L \| L \| I \| L \| M \| I \| L \| E \| V; 1 sequence: \| R \| L \| I \| L \| V \| L \| L \| L \| D \| I; 1 sequence: \| R \| L \| I \| L \| I \| L \| I \| I \| E \| I; 1 sequence: \| R \| L \| L \| L \| I \| L \| I \| L \| D \| I All positions in cluster: 104 123 251 255 261 265-266 271 275 279 |
| 0 | 20 | 1 | 1 | 7.550589e-05 | Hit patterns: Positions: \| 1037 \| 1038 \| 1074 \| 1079 \| 1082 \| 1083; 41 sequences: \| I \| V \| Y \| L \| H \| M; 26 sequences: \| I \| I \| Y \| L \| H \| M; 12 sequences: \| V \| V \| Y \| L \| H \| M; 12 sequences: \| M \| V \| Y \| L \| H \| M; 11 sequences: \| M \| I \| Y \| L \| H \| M; 4 sequences: \| V \| I \| Y \| L \| H \| M; 3 sequences: \| L \| I \| Y \| L \| H \| M; 3 sequences: \| - \| - \| - \| - \| - \| -; 3 sequences: \| L \| V \| Y \| L \| H \| M; 1 sequence: \| A \| V \| Y \| L \| H \| M; 1 sequence: \| L \| V \| H \| M \| H \| L; 1 sequence: \| L \| V \| H \| L \| H \| L; 1 sequence: \| L \| V \| H \| L \| H \| M All positions in cluster: 1037-1038 1074 1079 1082-1083 |
| 0 | 1 | 1 | 1 | 0.0001473622 | Hit patterns: Positions: \| 5417 \| 5420; 115 sequences: \| K \| T; 2 sequences: \| - \| -; 1 sequence: \| K \| S; 1 sequence: \| R \| T All positions in cluster: 5417 5420 |
| 0 | 17 | 1 | 1 | 0.0001473622 | Hit patterns: Positions: \| 1634 \| 1700; 115 sequences: \| H \| L; 2 sequences: \| - \| -; 2 sequences: \| H \| I All positions in cluster: 1634 1700 |
| 0 | 12 | 1 | 1 | 0.0001796945 | Hit patterns: Positions: \| 2967 \| 5371 \| 5372; 104 sequences: \| T \| K \| N; 6 sequences: \| T \| K \| S; 2 sequences: \| S \| K \| N; 2 sequences: \| C \| I \| D; 2 sequences: \| T \| K \| T; 2 sequences: \| S \| K \| S; 1 sequence: \| C \| I \| E All positions in cluster: 2966-2967 5369-5374 |
| 0 | 11 | 1 | 1 | 0.0001831502 | Hit patterns: Positions: \| 3212 \| 3309 \| 3310 \| 3414 \| 3941; 103 sequences: \| S \| V \| L \| L \| Q; 10 sequences: \| S \| I \| L \| L \| Q; 2 sequences: \| - \| - \| - \| - \| -; 1 sequence: \| S \| V \| L \| L \| N; 1 sequence: \| T \| V \| L \| L \| Q; 1 sequence: \| T \| V \| L \| F \| Q; 1 sequence: \| S \| I \| F \| L \| Q All positions in cluster: 3212 3309-3310 3414 3941 |
| 0 | 19 | 1 | 1 | 0.0001941371 | Hit patterns: Positions: \| 1095 \| 1096 \| 1099 \| 1100 \| 1103 \| 1105 \| 1106 \| 1127 \| 1133 \| 1137; 100 sequences: \| E \| L \| L \| M \| L \| W \| I \| I \| Y \| L; 4 sequences: \| E \| L \| L \| M \| L \| W \| V \| I \| Y \| L; 4 sequences: \| E \| L \| L \| L \| L \| W \| I \| I \| Y \| L; 2 sequences: \| - \| - \| - \| - \| - \| - \| - \| - \| - \| -; 2 sequences: \| E \| L \| L \| M \| L \| W \| V \| I \| H \| L; 2 sequences: \| D \| L \| L \| L \| L \| W \| L \| I \| Y \| L; 1 sequence: \| E \| L \| L \| M \| M \| W \| I \| I \| Y \| L; 1 sequence: \| E \| L \| F \| L \| L \| W \| V \| I \| Y \| L; 1 sequence: \| D \| L \| I \| L \| L \| W \| V \| V \| H \| L; 1 sequence: \| E \| L \| L \| L \| I \| W \| L \| L \| Y \| V; 1 sequence: \| E \| L \| I \| L \| L \| W \| I \| L \| Y \| L All positions in cluster: 1095-1096 1099-1100 1103 1105-1106 1127 1133 1137 |
| 0 | 18 | 1 | 1 | 0.0002192982 | Hit patterns: Positions: \| 1261 \| 1269 \| 1309 \| 1321; 94 sequences: \| V \| A \| L \| W; 14 sequences: \| V \| A \| V \| W; 4 sequences: \| V \| A \| F \| L; 2 sequences: \| - \| - \| - \| -; 2 sequences: \| V \| A \| F \| W; 1 sequence: \| V \| A \| M \| W; 1 sequence: \| L \| A \| L \| W; 1 sequence: \| V \| V \| V \| W All positions in cluster: 1261 1269 1309 1321 |
| 0 | 14 | 1 | 1 | 0.0002287806 | Hit patterns: Positions: \| 2404 \| 2405 \| 2496 \| 2499 \| 2501 \| 2529 \| 2530 \| 2531 \| 2533; 92 sequences: \| L \| W \| V \| V \| F \| F \| L \| T \| S; 9 sequences: \| L \| W \| V \| V \| F \| L \| L \| S \| S; 4 sequences: \| L \| W \| V \| A \| F \| F \| L \| T \| S; 3 sequences: \| I \| W \| V \| V \| F \| F \| L \| T \| S; 3 sequences: \| I \| W \| V \| V \| F \| F \| V \| T \| S; 2 sequences: \| - \| - \| - \| - \| - \| - \| - \| - \| -; 1 sequence: \| L \| W \| V \| V \| F \| F \| M \| T \| S; 1 sequence: \| L \| W \| I \| V \| F \| F \| L \| T \| S; 1 sequence: \| L \| W \| V \| V \| F \| F \| I \| T \| S; 1 sequence: \| L \| W \| V \| V \| F \| F \| V \| T \| S; 1 sequence: \| V \| W \| V \| V \| F \| I \| L \| S \| S; 1 sequence: \| L \| W \| V \| V \| F \| L \| V \| S \| S All positions in cluster: 2404-2405 2496 2499 2501 2529-2531 2533 |
| 0 | 3 | 1 | 1 | 0.0003508772 | Hit patterns: Positions: \| 5031 \| 5034 \| 5046 \| 5050 \| 5054 \| 5055 \| 5056 \| 5057 \| 5059 \| 5061 \| 5063 \| 5066 \| 5076; 74 sequences: \| D \| D \| L \| P \| D \| I \| L \| V \| Y \| T \| P \| G \| V; 39 sequences: \| D \| D \| L \| P \| D \| I \| F \| V \| Y \| T \| P \| G \| V; 2 sequences: \| - \| - \| - \| - \| - \| - \| - \| - \| - \| - \| - \| - \| -; 1 sequence: \| D \| D \| V \| P \| D \| I \| L \| V \| Y \| T \| P \| G \| V; 1 sequence: \| D \| D \| L \| P \| D \| I \| L \| V \| Y \| T \| P \| G \| L; 1 sequence: \| E \| D \| L \| P \| D \| M \| L \| L \| Y \| T \| P \| G \| V; 1 sequence: \| D \| D \| V \| P \| D \| M \| L \| L \| Y \| T \| P \| G \| V All positions in cluster: 5031 5034 5046 5050 5054-5057 5059 5061 5063 5066 5076 |
| 0 | 16 | 1 | 1 | 0.0004262575 | Hit patterns: Positions: \| 1782 \| 1788 \| 1822 \| 1826 \| 1828 \| 1831; 67 sequences: \| V \| S \| L \| S \| D \| L; 32 sequences: \| V \| S \| V \| S \| D \| V; 4 sequences: \| V \| S \| I \| S \| D \| V; 4 sequences: \| V \| S \| L \| S \| D \| V; 4 sequences: \| V \| S \| V \| S \| D \| L; 2 sequences: \| - \| - \| - \| - \| - \| -; 1 sequence: \| V \| S \| I \| S \| D \| L; 1 sequence: \| I \| S \| L \| S \| D \| L; 1 sequence: \| V \| S \| M \| S \| D \| L; 1 sequence: \| V \| S \| V \| S \| D \| A; 1 sequence: \| I \| S \| I \| S \| D \| I; 1 sequence: \| I \| T \| I \| S \| D \| V All positions in cluster: 1782 1788 1822 1826 1828 1831 |
| 0 | 7 | 1 | 1 | 0.0009250694 | Hit patterns: Positions: \| 309 \| 310 \| 313 \| 316 \| 352 \| 356 \| 417 \| 420 \| 429 \| 430 \| 433 \| 439 \| 446 \| 460 \| 495 \| 539 \| 542 \| 544 \| 545; 45 sequences: \| L \| I \| I \| K \| L \| L \| L \| G \| V \| P \| P \| R \| I \| L \| L \| W \| I \| G \| M; 43 sequences: \| L \| I \| I \| K \| L \| L \| L \| G \| V \| P \| P \| R \| I \| L \| L \| W \| V \| G \| M; 8 sequences: \| L \| I \| L \| K \| L \| L \| L \| G \| V \| P \| P \| R \| I \| L \| L \| W \| V \| G \| M; 5 sequences: \| L \| I \| I \| K \| L \| L \| L \| G \| V \| P \| P \| R \| V \| L \| L \| W \| I \| G \| M; 2 sequences: \| L \| I \| V \| K \| L \| L \| L \| G \| V \| P \| P \| R \| V \| L \| L \| W \| V \| G \| M; 2 sequences: \| L \| L \| L \| K \| L \| L \| L \| G \| V \| P \| P \| R \| V \| L \| L \| W \| V \| G \| M; 2 sequences: \| - \| - \| - \| - \| - \| - \| - \| - \| - \| - \| - \| - \| - \| - \| - \| - \| - \| - \| -; 2 sequences: \| L \| L \| I \| K \| L \| L \| L \| G \| V \| P \| P \| R \| I \| L \| L \| W \| I \| G \| M; 1 sequence: \| L \| L \| V \| K \| L \| L \| L \| G \| V \| P \| P \| R \| I \| L \| L \| W \| I \| G \| M; 1 sequence: \| L \| I \| I \| K \| F \| L \| L \| G \| V \| P \| P \| R \| I \| L \| L \| W \| I \| G \| M; 1 sequence: \| L \| M \| I \| K \| L \| L \| L \| G \| V \| P \| P \| R \| I \| L \| L \| W \| I \| G \| M; 1 sequence: \| L \| L \| I \| K \| L \| L \| L \| G \| V \| P \| P \| R \| I \| L \| L \| W \| V \| G \| M; 1 sequence: \| L \| L \| L \| K \| L \| L \| L \| G \| V \| P \| P \| R \| V \| L \| L \| W \| I \| G \| M; 1 sequence: \| L \| V \| I \| K \| L \| L \| L \| G \| V \| P \| P \| R \| I \| L \| L \| W \| V \| G \| M; 1 sequence: \| L \| I \| L \| K \| L \| L \| L \| G \| I \| P \| P \| R \| A \| L \| L \| W \| V \| G \| M; 1 sequence: \| L \| L \| L \| R \| L \| L \| L \| G \| V \| P \| P \| R \| I \| L \| L \| W \| V \| G \| M; 1 sequence: \| F \| L \| L \| R \| L \| L \| L \| G \| V \| P \| P \| R \| I \| L \| L \| W \| L \| G \| L; 1 sequence: \| F \| L \| I \| K \| L \| L \| L \| G \| V \| P \| P \| R \| I \| L \| L \| W \| V \| G \| M All positions in cluster: 309-310 313 316 352 356 417 420 429-430 433 439 446 460 495 539 542 544-545 |
| 0 | 6 | 1 | 1 | 0.002150538 | Hit patterns: Positions: \| 4914 \| 4915 \| 4916 \| 4917 \| 4918 \| 4919 \| 4920 \| 4930 \| 4933 \| 4935 \| 4937 \| 4941 \| 4943 \| 4950 \| 4958 \| 4961 \| 4962 \| 4963 \| 4964 \| 4965 \| 4966 \| 4967 \| 4968 \| 4969 \| 4970 \| 4974; 29 sequences: \| V \| V \| I \| L \| S \| H \| G \| G \| V \| G \| D \| V \| V \| F \| L \| K \| P \| K \| L \| F \| F \| I \| Q \| A \| C \| G; 17 sequences: \| V \| V \| I \| L \| S \| H \| G \| G \| V \| G \| D \| V \| I \| F \| L \| K \| P \| K \| L \| F \| F \| I \| Q \| A \| C \| G; 16 sequences: \| V \| V \| I \| L \| S \| H \| G \| G \| V \| G \| D \| I \| I \| F \| L \| K \| P \| K \| L \| F \| F \| I \| Q \| A \| C \| G; 8 sequences: \| V \| V \| I \| L \| S \| H \| G \| G \| I \| G \| D \| I \| I \| F \| L \| K \| P \| K \| L \| F \| F \| I \| Q \| A \| C \| G; 8 sequences: \| V \| V \| I \| L \| S \| H \| G \| G \| I \| G \| D \| I \| V \| F \| L \| K \| P \| K \| L \| F \| F \| I \| Q \| A \| C \| G; 4 sequences: \| V \| V \| I \| L \| S \| H \| G \| G \| I \| G \| D \| V \| V \| F \| L \| K \| P \| K \| L \| F \| F \| I \| Q \| A \| C \| G; 3 sequences: \| V \| V \| V \| L \| S \| H \| G \| G \| V \| G \| D \| V \| I \| F \| L \| K \| P \| K \| L \| F \| F \| I \| Q \| A \| C \| G; 3 sequences: \| V \| V \| I \| L \| S \| H \| G \| G \| V \| G \| D \| V \| V \| L \| L \| K \| P \| K \| L \| F \| F \| I \| Q \| A \| C \| G; 3 sequences: \| V \| I \| M \| L \| S \| H \| G \| G \| V \| G \| D \| V \| V \| L \| L \| K \| P \| K \| L \| F \| F \| I \| Q \| A \| C \| G; 2 sequences: \| V \| V \| I \| L \| S \| H \| G \| G \| V \| G \| D \| I \| V \| F \| L \| K \| P \| K \| L \| F \| I \| I \| Q \| A \| C \| G; 2 sequences: \| V \| V \| I \| L \| S \| H \| G \| G \| V \| G \| D \| I \| V \| F \| L \| K \| P \| K \| L \| F \| F \| I \| Q \| A \| C \| G; 2 sequences: \| V \| V \| I \| L \| S \| H \| G \| G \| V \| G \| D \| I \| I \| F \| L \| K \| P \| K \| L \| F \| I \| I \| Q \| A \| C \| G; 2 sequences: \| V \| V \| V \| L \| S \| H \| G \| G \| V \| G \| D \| V \| V \| F \| L \| K \| P \| K \| L \| F \| F \| I \| Q \| A \| C \| G; 2 sequences: \| - \| - \| - \| - \| - \| - \| - \| - \| - \| - \| - \| - \| - \| - \| - \| - \| - \| - \| - \| - \| - \| - \| - \| - \| - \| -; 1 sequence: \| V \| V \| I \| L \| S \| H \| G \| G \| I \| G \| D \| L \| V \| F \| L \| K \| P \| K \| I \| F \| F \| I \| Q \| A \| C \| G; 1 sequence: \| V \| V \| L \| L \| S \| H \| G \| G \| I \| G \| D \| I \| V \| F \| L \| K \| P \| K \| L \| F \| F \| I \| Q \| A \| C \| G; 1 sequence: \| V \| V \| I \| L \| S \| H \| G \| G \| V \| G \| D \| L \| V \| F \| L \| K \| P \| K \| L \| F \| F \| I \| Q \| A \| C \| G; 1 sequence: \| V \| V \| I \| L \| S \| H \| G \| G \| V \| G \| D \| V \| L \| F \| L \| K \| P \| K \| L \| F \| F \| I \| Q \| A \| C \| G; 1 sequence: \| V \| I \| I \| L \| S \| H \| G \| G \| V \| G \| D \| V \| V \| F \| L \| K \| P \| K \| L \| F \| F \| I \| Q \| A \| C \| G; 1 sequence: \| V \| I \| M \| L \| S \| H \| G \| G \| V \| G \| D \| I \| V \| L \| L \| K \| P \| K \| L \| F \| F \| I \| Q \| A \| C \| G; 1 sequence: \| V \| A \| I \| L \| S \| H \| G \| G \| V \| G \| D \| V \| I \| F \| L \| K \| P \| K \| L \| F \| F \| I \| Q \| A \| C \| G; 1 sequence: \| V \| I \| I \| L \| S \| H \| G \| G \| V \| G \| D \| V \| I \| F \| L \| K \| P \| K \| L \| F \| F \| I \| Q \| A \| C \| G; 1 sequence: \| V \| A \| M \| L \| S \| H \| G \| G \| V \| G \| D \| V \| V \| L \| L \| K \| P \| K \| L \| F \| F \| L \| Q \| A \| C \| G; 1 sequence: \| I \| I \| I \| L \| S \| H \| G \| G \| V \| G \| D \| I \| V \| F \| L \| K \| P \| K \| L \| F \| F \| I \| Q \| A \| C \| G; 1 sequence: \| V \| I \| I \| L \| S \| H \| G \| G \| I \| G \| D \| V \| V \| F \| L \| K \| P \| K \| L \| F \| F \| I \| Q \| A \| C \| G; 1 sequence: \| V \| V \| L \| L \| S \| H \| G \| G \| I \| G \| D \| L \| V \| F \| L \| K \| P \| K \| L \| F \| F \| I \| Q \| A \| C \| G; 1 sequence: \| V \| V \| I \| L \| S \| H \| G \| G \| I \| G \| D \| I \| L \| F \| L \| K \| P \| K \| L \| F \| F \| I \| Q \| A \| C \| G; 1 sequence: \| V \| V \| V \| L \| S \| H \| G \| G \| I \| G \| D \| I \| V \| F \| L \| K \| P \| K \| L \| F \| F \| I \| Q \| A \| C \| G; 1 sequence: \| V \| A \| V \| L \| S \| H \| G \| G \| V \| G \| D \| L \| V \| F \| L \| K \| P \| K \| L \| F \| F \| F \| Q \| A \| C \| G; 1 sequence: \| V \| V \| I \| L \| S \| H \| G \| G \| V \| G \| D \| I \| V \| F \| L \| K \| P \| K \| I \| F \| F \| I \| Q \| A \| C \| G; 1 sequence: \| L \| A \| V \| L \| S \| H \| G \| G \| I \| G \| D \| L \| V \| F \| L \| K \| P \| K \| L \| F \| F \| I \| Q \| A \| C \| G; 1 sequence: \| F \| A \| I \| L \| T \| H \| G \| G \| V \| G \| D \| V \| I \| F \| L \| K \| P \| K \| L \| F \| F \| L \| Q \| A \| C \| G All positions in cluster: 4914-4920 4930 4933 4935 4937 4941 4943 4950 4958 4961-4970 4974 |
| 0 | 2 | 1 | 1 | 1 | Hit patterns: Positions: \| 1 \| 1977 \| 1978 \| 2369 \| 2370 \| 2371 \| 2385 \| 2823 \| 2872 \| 2884 \| 2925 \| 2926 \| 2961 \| 2966 \| 3010 \| 4109 \| 5266 \| 5326 \| 5328 \| 5331 \| 5332 \| 5334 \| 5335 \| 5336 \| 5339 \| 5342 \| 5344 \| 5348 \| 5349 \| 5350 \| 5352 \| 5353 \| 5354 \| 5355 \| 5359 \| 5360 \| 5367 \| 5369 \| 5370 \| 5373 \| 5374 \| 5381 \| 5382 \| 5392 \| 5393 \| 5394; 28 sequences: \| M \| I \| M \| L \| A \| V \| V \| G \| G \| I \| I \| L \| V \| F \| L \| M \| M \| K \| F \| K \| C \| Q \| C \| H \| E \| G \| H \| P \| N \| L \| G \| L \| F \| G \| G \| Q \| T \| A \| N \| K \| G \| T \| L \| Y \| I \| P; 21 sequences: \| M \| I \| L \| L \| V \| A \| V \| G \| G \| I \| V \| L \| V \| F \| L \| M \| M \| K \| F \| K \| C \| Q \| C \| H \| E \| G \| H \| P \| N \| L \| G \| L \| F \| G \| G \| Q \| T \| A \| N \| K \| G \| T \| L \| Y \| I \| P; 5 sequences: \| M \| I \| M \| L \| A \| L \| V \| G \| G \| I \| I \| L \| V \| F \| L \| M \| M \| K \| F \| K \| C \| Q \| C \| H \| E \| G \| H \| P \| N \| L \| G \| L \| F \| G \| G \| Q \| T \| A \| N \| K \| G \| T \| L \| Y \| I \| P; 3 sequences: \| M \| I \| M \| L \| A \| V \| V \| G \| G \| V \| I \| L \| V \| F \| L \| M \| M \| K \| F \| K \| C \| Q \| C \| H \| E \| G \| H \| P \| N \| L \| G \| L \| F \| G \| G \| Q \| T \| A \| N \| K \| G \| T \| L \| Y \| I \| P; 3 sequences: \| M \| L \| L \| L \| V \| A \| V \| G \| G \| I \| V \| L \| V \| F \| L \| M \| M \| K \| F \| K \| C \| Q \| C \| H \| E \| G \| H \| P \| N \| L \| G \| L \| F \| G \| G \| Q \| T \| A \| N \| K \| G \| T \| L \| Y \| I \| P; 3 sequences: \| M \| I \| V \| I \| V \| A \| V \| G \| G \| V \| V \| L \| V \| F \| L \| M \| M \| K \| F \| K \| C \| T \| C \| H \| E \| G \| H \| P \| N \| L \| G \| L \| F \| G \| G \| Q \| T \| A \| N \| K \| G \| T \| L \| Y \| I \| P; 2 sequences: \| M \| V \| M \| L \| V \| A \| V \| G \| G \| V \| V \| L \| V \| F \| L \| M \| M \| K \| F \| K \| C \| Q \| C \| H \| E \| G \| H \| P \| N \| L \| G \| L \| I \| G \| G \| Q \| T \| A \| N \| K \| G \| T \| L \| Y \| I \| P; 2 sequences: \| M \| I \| M \| L \| V \| A \| V \| G \| G \| V \| V \| L \| V \| F \| L \| M \| M \| K \| F \| K \| C \| Q \| C \| H \| E \| G \| H \| P \| N \| L \| G \| L \| I \| G \| G \| Q \| T \| A \| N \| K \| G \| T \| L \| Y \| I \| P; 2 sequences: \| M \| I \| L \| L \| V \| A \| I \| G \| G \| I \| V \| L \| V \| F \| L \| M \| M \| K \| F \| K \| C \| Q \| C \| H \| E \| G \| H \| P \| N \| L \| G \| L \| F \| G \| G \| Q \| T \| A \| N \| K \| G \| T \| L \| Y \| I \| P; 2 sequences: \| M \| V \| M \| L \| V \| A \| V \| G \| G \| V \| V \| L \| V \| F \| L \| M \| M \| K \| F \| K \| C \| Q \| C \| H \| E \| G \| H \| P \| N \| L \| G \| L \| F \| G \| G \| Q \| T \| A \| N \| K \| G \| T \| L \| Y \| I \| P; 2 sequences: \| M \| I \| I \| L \| L \| A \| V \| G \| G \| V \| V \| L \| V \| F \| L \| M \| M \| K \| F \| K \| C \| Q \| C \| H \| E \| G \| H \| P \| N \| L \| G \| L \| F \| G \| G \| Q \| T \| A \| N \| K \| G \| T \| L \| Y \| I \| P; 2 sequences: \| M \| I \| M \| L \| A \| V \| V \| G \| G \| I \| I \| L \| V \| F \| V \| M \| M \| K \| F \| K \| C \| Q \| C \| H \| E \| G \| H \| P \| N \| L \| G \| L \| F \| G \| G \| Q \| T \| A \| N \| K \| G \| T \| L \| Y \| I \| P; 2 sequences: \| M \| I \| L \| L \| A \| V \| V \| G \| G \| I \| I \| I \| V \| F \| L \| M \| M \| K \| F \| K \| C \| Q \| C \| H \| E \| G \| H \| P \| N \| L \| G \| L \| F \| G \| G \| Q \| T \| A \| N \| K \| G \| T \| L \| Y \| I \| P; 2 sequences: \| M \| I \| I \| L \| A \| V \| V \| G \| G \| I \| I \| I \| V \| F \| L \| M \| M \| K \| F \| K \| C \| Q \| C \| H \| E \| G \| H \| P \| N \| L \| G \| L \| F \| G \| G \| Q \| T \| A \| N \| K \| G \| T \| L \| Y \| I \| P; 2 sequences: \| M \| I \| M \| L \| A \| V \| V \| G \| G \| L \| I \| L \| V \| F \| L \| M \| M \| K \| F \| K \| C \| Q \| C \| H \| E \| G \| H \| P \| N \| L \| G \| L \| F \| G \| G \| Q \| T \| A \| N \| K \| G \| T \| L \| Y \| I \| P; 1 sequence: \| M \| I \| L \| L \| V \| A \| V \| G \| G \| V \| V \| L \| V \| F \| L \| M \| M \| K \| F \| K \| C \| Q \| C \| H \| E \| G \| H \| P \| N \| L \| G \| L \| F \| G \| G \| Q \| T \| A \| N \| K \| G \| T \| L \| Y \| I \| P; 1 sequence: \| M \| I \| V \| M \| V \| A \| V \| G \| G \| V \| V \| L \| V \| F \| L \| M \| M \| K \| F \| K \| C \| Q \| C \| H \| E \| G \| H \| P \| N \| L \| G \| L \| F \| G \| G \| Q \| T \| A \| N \| K \| G \| T \| L \| Y \| I \| P; 1 sequence: \| M \| V \| L \| L \| V \| A \| V \| G \| G \| I \| I \| L \| V \| F \| L \| M \| M \| K \| F \| K \| C \| Q \| C \| H \| E \| G \| H \| P \| N \| L \| G \| L \| F \| G \| G \| Q \| T \| A \| N \| K \| G \| T \| L \| Y \| I \| P; 1 sequence: \| M \| L \| I \| L \| V \| A \| V \| G \| G \| V \| V \| L \| V \| F \| L \| M \| M \| K \| F \| K \| C \| Q \| C \| H \| E \| G \| H \| P \| N \| L \| G \| L \| I \| G \| G \| Q \| T \| A \| N \| K \| G \| T \| L \| Y \| I \| P; 1 sequence: \| M \| I \| L \| L \| V \| A \| V \| G \| G \| L \| V \| L \| V \| F \| L \| M \| M \| K \| F \| K \| C \| Q \| C \| H \| E \| G \| H \| P \| N \| L \| G \| I \| F \| G \| G \| Q \| T \| A \| N \| K \| G \| T \| L \| Y \| I \| P; 1 sequence: \| M \| I \| L \| L \| A \| V \| V \| G \| G \| I \| I \| L \| V \| F \| L \| M \| M \| K \| F \| K \| C \| Q \| C \| H \| E \| G \| H \| P \| N \| L \| G \| L \| F \| G \| G \| Q \| T \| A \| N \| K \| G \| T \| L \| Y \| I \| P; 1 sequence: \| M \| I \| M \| L \| A \| V \| V \| G \| G \| I \| I \| L \| V \| F \| L \| M \| M \| R \| F \| K \| C \| Q \| C \| H \| E \| G \| H \| P \| N \| L \| G \| I \| F \| G \| G \| Q \| T \| A \| N \| K \| G \| T \| L \| Y \| I \| P; 1 sequence: \| M \| V \| M \| L \| A \| V \| V \| G \| G \| V \| I \| L \| V \| F \| L \| M \| M \| K \| F \| K \| C \| Q \| C \| H \| E \| G \| H \| P \| N \| L \| G \| L \| F \| G \| G \| Q \| T \| A \| N \| K \| G \| T \| L \| Y \| I \| P; 1 sequence: \| M \| I \| M \| L \| V \| I \| V \| G \| G \| I \| I \| L \| V \| F \| L \| M \| M \| K \| F \| K \| C \| Q \| C \| H \| E \| G \| H \| P \| N \| L \| G \| L \| F \| G \| G \| Q \| T \| A \| N \| K \| G \| T \| L \| Y \| I \| P; 1 sequence: \| M \| I \| L \| L \| V \| A \| V \| G \| G \| I \| I \| L \| V \| F \| L \| M \| M \| K \| F \| K \| C \| Q \| C \| H \| E \| G \| H \| P \| N \| L \| G \| L \| F \| G \| G \| Q \| T \| A \| N \| K \| G \| T \| L \| Y \| I \| P; 1 sequence: \| M \| I \| I \| L \| L \| A \| V \| G \| G \| L \| V \| L \| V \| F \| L \| M \| M \| K \| F \| K \| C \| Q \| C \| H \| E \| G \| H \| P \| N \| L \| G \| L \| F \| G \| G \| Q \| T \| A \| N \| K \| G \| T \| L \| Y \| I \| P; 1 sequence: \| M \| I \| M \| L \| A \| V \| V \| G \| G \| L \| I \| L \| I \| F \| L \| M \| M \| K \| F \| K \| C \| Q \| C \| H \| E \| G \| H \| P \| N \| L \| G \| L \| F \| G \| G \| Q \| T \| A \| N \| K \| G \| T \| L \| Y \| I \| P; 1 sequence: \| M \| I \| M \| L \| A \| V \| V \| G \| G \| I \| I \| L \| L \| F \| L \| M \| M \| K \| F \| K \| C \| Q \| C \| H \| E \| G \| H \| P \| N \| L \| G \| L \| F \| G \| G \| Q \| T \| A \| N \| K \| G \| T \| L \| Y \| I \| P; 1 sequence: \| M \| I \| M \| L \| A \| V \| V \| G \| G \| V \| V \| L \| V \| F \| L \| M \| M \| K \| F \| K \| C \| Q \| C \| H \| E \| G \| H \| P \| N \| L \| G \| L \| F \| G \| G \| Q \| T \| A \| N \| K \| G \| T \| L \| Y \| I \| P; 1 sequence: \| M \| M \| M \| L \| A \| V \| V \| G \| G \| I \| I \| L \| V \| F \| L \| M \| M \| K \| F \| K \| C \| Q \| C \| H \| E \| G \| H \| P \| N \| L \| G \| L \| F \| G \| G \| Q \| T \| A \| N \| K \| G \| T \| L \| Y \| I \| P; 1 sequence: \| M \| I \| I \| L \| L \| A \| V \| G \| G \| V \| V \| L \| V \| F \| L \| M \| M \| K \| F \| K \| C \| Q \| C \| H \| E \| G \| H \| P \| N \| L \| G \| M \| F \| G \| G \| Q \| T \| A \| N \| K \| G \| T \| L \| Y \| I \| P; 1 sequence: \| M \| I \| M \| L \| A \| V \| V \| G \| G \| I \| I \| F \| V \| F \| L \| M \| M \| K \| F \| K \| C \| Q \| C \| H \| E \| G \| H \| P \| N \| L \| G \| L \| F \| G \| G \| Q \| T \| A \| N \| K \| G \| T \| L \| Y \| I \| P; 1 sequence: \| M \| V \| I \| L \| V \| A \| V \| G \| G \| V \| V \| L \| V \| F \| L \| M \| M \| K \| F \| K \| C \| Q \| C \| H \| E \| G \| H \| P \| N \| L \| G \| L \| L \| G \| G \| Q \| T \| A \| N \| K \| G \| T \| L \| Y \| I \| P; 1 sequence: \| M \| I \| M \| L \| V \| V \| V \| G \| G \| L \| V \| L \| V \| F \| L \| M \| M \| K \| F \| K \| C \| Q \| C \| H \| E \| G \| H \| P \| N \| L \| G \| L \| F \| G \| G \| Q \| T \| A \| N \| K \| G \| T \| L \| Y \| I \| P; 1 sequence: \| M \| I \| I \| L \| V \| A \| V \| G \| G \| V \| V \| L \| V \| F \| L \| M \| M \| K \| F \| K \| C \| Q \| C \| H \| E \| G \| H \| P \| N \| L \| G \| L \| F \| G \| G \| Q \| T \| A \| N \| K \| G \| T \| L \| Y \| I \| P; 1 sequence: \| M \| I \| L \| L \| V \| A \| V \| G \| G \| I \| V \| L \| V \| F \| L \| M \| M \| K \| F \| K \| C \| Q \| C \| H \| E \| G \| H \| P \| N \| L \| G \| V \| F \| G \| G \| Q \| S \| A \| N \| K \| G \| T \| M \| Y \| I \| P; 1 sequence: \| M \| L \| M \| L \| A \| A \| V \| G \| G \| I \| I \| L \| V \| F \| L \| M \| M \| K \| F \| K \| C \| Q \| C \| H \| E \| G \| H \| P \| N \| L \| G \| L \| F \| G \| G \| Q \| T \| A \| N \| K \| G \| T \| L \| Y \| I \| P; 1 sequence: \| M \| I \| M \| L \| A \| V \| V \| G \| G \| I \| I \| L \| V \| F \| V \| M \| M \| K \| F \| K \| C \| Q \| C \| H \| E \| G \| H \| P \| N \| L \| G \| L \| F \| G \| G \| Q \| T \| A \| N \| K \| G \| S \| L \| H \| V \| P; 1 sequence: \| M \| I \| M \| I \| V \| A \| V \| G \| G \| V \| V \| F \| V \| F \| L \| M \| M \| K \| F \| K \| C \| Q \| C \| H \| E \| G \| H \| P \| N \| L \| G \| L \| F \| G \| G \| Q \| T \| A \| N \| K \| G \| T \| L \| Y \| I \| P; 1 sequence: \| M \| I \| M \| L \| A \| L \| V \| G \| G \| I \| I \| V \| V \| F \| L \| M \| M \| K \| F \| K \| C \| Q \| C \| H \| E \| G \| H \| P \| N \| L \| G \| L \| F \| G \| G \| Q \| T \| A \| N \| K \| G \| T \| L \| Y \| I \| P; 1 sequence: \| M \| I \| I \| L \| A \| V \| V \| G \| G \| I \| I \| L \| V \| F \| L \| M \| M \| K \| F \| K \| C \| Q \| C \| H \| E \| G \| H \| P \| N \| L \| G \| L \| F \| G \| G \| Q \| T \| A \| N \| K \| G \| T \| L \| Y \| I \| P; 1 sequence: \| M \| I \| I \| L \| A \| V \| V \| G \| G \| I \| V \| I \| V \| F \| L \| M \| M \| K \| F \| K \| C \| Q \| C \| H \| E \| G \| H \| P \| N \| L \| G \| L \| F \| G \| G \| Q \| T \| A \| N \| K \| G \| T \| L \| Y \| I \| P; 1 sequence: \| M \| I \| V \| M \| V \| A \| V \| G \| G \| V \| V \| I \| V \| F \| L \| M \| M \| K \| F \| K \| C \| Q \| C \| H \| E \| G \| H \| P \| N \| L \| G \| L \| F \| G \| G \| Q \| T \| A \| N \| K \| G \| T \| L \| Y \| I \| P; 1 sequence: \| M \| V \| I \| L \| L \| A \| V \| G \| G \| V \| V \| L \| V \| F \| L \| M \| M \| K \| F \| K \| C \| Q \| C \| H \| E \| G \| H \| P \| N \| L \| G \| L \| F \| G \| G \| Q \| T \| A \| N \| K \| G \| T \| L \| Y \| I \| P; 1 sequence: \| M \| I \| F \| L \| L \| A \| V \| G \| G \| V \| V \| L \| V \| F \| L \| M \| M \| K \| F \| K \| C \| Q \| C \| H \| E \| G \| H \| P \| N \| L \| G \| L \| F \| G \| G \| Q \| T \| A \| N \| K \| G \| T \| L \| Y \| I \| P; 1 sequence: \| M \| I \| M \| L \| A \| V \| V \| G \| G \| I \| V \| L \| V \| F \| V \| M \| M \| K \| F \| K \| C \| Q \| C \| H \| E \| G \| H \| P \| N \| L \| G \| L \| F \| G \| G \| Q \| T \| A \| N \| K \| G \| T \| L \| Y \| I \| P; 1 sequence: \| M \| L \| V \| L \| A \| V \| V \| G \| G \| I \| V \| L \| V \| F \| V \| M \| M \| K \| F \| K \| C \| Q \| C \| H \| E \| G \| H \| P \| N \| L \| G \| L \| F \| G \| G \| Q \| T \| A \| N \| K \| G \| T \| L \| Y \| I \| P; 1 sequence: \| M \| V \| L \| L \| V \| A \| V \| G \| G \| I \| V \| L \| V \| F \| L \| M \| M \| K \| F \| K \| C \| Q \| C \| H \| E \| G \| H \| P \| N \| L \| G \| L \| F \| G \| G \| Q \| T \| A \| N \| K \| G \| T \| L \| Y \| I \| P; 1 sequence: \| M \| I \| M \| L \| V \| V \| V \| G \| G \| I \| I \| L \| V \| F \| L \| M \| M \| K \| F \| K \| C \| Q \| C \| H \| E \| G \| H \| P \| N \| L \| G \| L \| F \| G \| G \| Q \| T \| A \| N \| K \| G \| T \| L \| Y \| I \| P; 1 sequence: \| M \| I \| M \| I \| A \| A \| V \| G \| G \| V \| I \| L \| V \| F \| L \| M \| M \| K \| F \| K \| C \| Q \| C \| H \| E \| G \| H \| P \| N \| L \| G \| L \| F \| G \| G \| Q \| T \| A \| N \| K \| G \| T \| L \| Y \| I \| P; 1 sequence: \| M \| V \| A \| L \| I \| A \| I \| G \| G \| V \| L \| L \| V \| F \| L \| M \| M \| K \| F \| K \| C \| Q \| C \| H \| E \| G \| H \| P \| N \| L \| G \| L \| F \| G \| G \| Q \| T \| A \| N \| K \| G \| T \| L \| Y \| I \| P; 1 sequence: \| M \| L \| I \| L \| L \| V \| V \| G \| G \| L \| V \| L \| V \| F \| L \| M \| M \| K \| F \| R \| C \| Q \| C \| H \| E \| G \| H \| P \| N \| L \| G \| L \| F \| G \| G \| Q \| T \| A \| N \| K \| G \| T \| L \| Y \| I \| P; 1 sequence: \| M \| V \| A \| I \| I \| A \| I \| G \| G \| V \| V \| A \| V \| F \| L \| M \| M \| K \| F \| K \| C \| Q \| C \| H \| D \| G \| H \| P \| N \| L \| G \| L \| W \| G \| G \| S \| T \| A \| N \| K \| G \| T \| L \| Y \| I \| P All positions in cluster: 1 1977-1978 2369-2371 2385 2823 2872 2884 2925-2926 2961 2966 3010 4109 5266 5326 5328 5331-5332 5334-5336 5339 5342 5344 5348-5350 5352-5355 5359-5360 5367 5369-5370 5373-5374 5381-5382 5392-5394 |
| 0 | 21 | 0.545455 | 0.829146 | 1 | Hit patterns: Positions: \| 1907 \| 1947 \| 1949 \| 5345; 98 sequences: \| F \| V \| V \| K; 5 sequences: \| F \| I \| V \| K; 2 sequences: \| F \| A \| P \| K; 2 sequences: \| F \| V \| P \| K; 2 sequences: \| F \| V \| L \| K; 1 sequence: \| F \| A \| V \| K; 1 sequence: \| L \| V \| V \| K; 1 sequence: \| F \| V \| V \| M; 1 sequence: \| I \| V \| V \| M; 1 sequence: \| F \| F \| L \| K; 1 sequence: \| F \| L \| V \| K; 1 sequence: \| I \| V \| V \| K; 1 sequence: \| - \| S \| L \| K; 1 sequence: \| F \| F \| V \| K; 1 sequence: \| - \| T \| V \| K All positions in cluster: 1907 1947 1949 5344-5345 |
| 0 | 22 | 0.545455 | 0.653266 | 1 | Hit patterns: Positions: \| 4839 \| 4842 \| 4843 \| 4844 \| 4845 \| 4846 \| 4849 \| 4870 \| 4880 \| 4882 \| 5361; 62 sequences: \| G \| L \| I \| I \| N \| N \| F \| L \| F \| V \| A; 21 sequences: \| G \| L \| I \| L \| N \| N \| F \| L \| F \| V \| A; 12 sequences: \| G \| L \| I \| I \| N \| N \| F \| M \| F \| V \| A; 6 sequences: \| G \| L \| I \| V \| N \| N \| F \| L \| F \| V \| A; 3 sequences: \| G \| L \| I \| F \| N \| N \| F \| L \| F \| V \| A; 2 sequences: \| G \| L \| I \| L \| N \| N \| F \| L \| F \| I \| A; 2 sequences: \| G \| F \| I \| I \| N \| N \| F \| L \| F \| V \| A; 2 sequences: \| - \| - \| - \| - \| - \| - \| - \| - \| - \| - \| A; 2 sequences: \| G \| L \| I \| I \| N \| N \| F \| L \| F \| I \| A; 1 sequence: \| G \| L \| I \| I \| N \| N \| F \| V \| F \| V \| S; 1 sequence: \| G \| L \| V \| I \| N \| N \| F \| L \| F \| V \| A; 1 sequence: \| G \| V \| I \| I \| N \| N \| F \| M \| F \| V \| A; 1 sequence: \| G \| I \| I \| I \| N \| N \| F \| L \| F \| V \| A; 1 sequence: \| G \| L \| I \| I \| N \| N \| F \| M \| L \| V \| A; 1 sequence: \| G \| F \| I \| V \| N \| N \| F \| L \| F \| V \| A; 1 sequence: \| G \| L \| I \| I \| N \| N \| F \| L \| F \| V \| S All positions in cluster: 4839 4842-4846 4849 4870 4880 4882 5359-5361 |

Table created with bis2html version 8.
