## Supplementary material for "Protein coevolution and physicochemical adaptation in the APAF-1/apoptosome: structural and functional implications": Figure 5-table supplement 1-Clusters of amino acids with coevolution signal

**Figure 5-table supplement 1. Clusters of amino acids with coevolution signal.** In the concatenation the numbering of the residues is followed continuously, not reflecting the real positions of the amino acids from the second protein, so for CASP9 and CYCS, the real positions are shown in parentheses.

| **Cluster** | **(Protein/Domain)** Amino acid/Position in concatenated sequence (real position) |
| --- | --- |
| **1** | **(CYCS)** K/1764 (100), T/1767 (103) |
|  | **(APAF-1/CARD)** M/1 |
|  | **(APAF-1/β-propeller 7)** I/813, M814, L854, A855, V856, V864 |
| **2** | **(APAF-1/β-propeller 8)** G/979, G/983, I/985, I/987, L/988, V/1006, F/1011, L/1018 |
|  | **(CASP9/CARD)** M/1249 (1) |
|  | **(CYCS)** M/1665 (1), K/1673 (9), F/1675 (11), K/1678 (14),C/1679 (15), Q/1681 (17), C/1682 (18), H/1683 (19), E/1686 (22), G/1689 (25), |
|  | H/1691 (27), P/1695 (31), N/1696 (32), L/1697 (33), G/1699 (35), L/1700 (36), F/1701 (37), G/1702 (38), G/1706 (42), Q/1707 (43),  T/1714 (50), A/1716 (52), N/1717 (53), K/1720 (56), G/1721 (57), T/1728 (64), L/1729 (65), Y/1739 (75), I/1740 (76), P/1741 (77) |
| **3** | **(CASP9/L2)** D/1575 (327), D/1578 (330) |
|  | **(CASP9/small subunit)** L/1583 (335), P/1584 (336), D/1588 (340), I/1589 (341), F/1590 (342), V/1591 (343), Y/1593 (345), T/1595  (347), P/1597 (349), G/1598 (350), V/1600 (352) |
| **4** | **(CASP9/Large subunit)** G/1517 (269), G/1536 (288) |
| **5** | **(CASP9/Large subunit)** H/1491 (243), F/1494 (246), P/1495 (247) |
| **6** | **(CASP9 Large subunit)** V/1480 (232), V/1481 (233), I/1482 (234), L/1483 (235), S/1484 (236), H/1485 (237), G/1486 (238), G/1496  (248), V/1498 (250), G/1500 (252), D/1502 (254), V/1506 (258), V/1508 (260) , F/1515 (267), L/1523 (275),  K/1526 (278), P/1527 (279), K/1528 (280), L/1529 (281), F/1530 (282), F/1531 (283), I/1532 (284), Q/1533 |
|  | (285), A/1534 (286), C/1535 (287), G/1537 (289) |
| **7** | **(APAF-1/CARD)** L/56, I/57, I/60, K/63, L/83, L/87 |
|  | **(APAF-1/NBD)** L/114, G/117, V/119, P/120, P/123, R/129, I/136, L/140, L/143, W/149, I/152, G/154, M/155 |
| **8** | **(CASP9/CARD)** L/1306 (58), L/1310 (62), A/1317 (69), L/1318 (70), F/1321 (73), I/1322 (74) |
| **9** | **(APAF-1/NBD)** E/116, R/122 |
| **10** | **(APAF-1/β-propeller 8)** V/1233, V/1235, L/1241 |
| **11** | **(APAF-1/β-propeller 8)** S/1060, V/1088, L/1089, L/1123, Q/1211 |
| **12** | **(APAF-1/β-propeller 8)** T/1012 |
|  | **(CYCS)** K/1718 (54), N/1719 (55) |
| **13** | **(APAF-1/CARD)** R/6, L/9, L/16, I/20, I/25, M/29, I/30, L/35, E/39, V/43 |
| **14** | **(APAF-1/β-propeller 7)** L/866, W/867, V/885, V/888, F/890, F/897, L/898, T/899, S/901 |
| **15** | **(APAF-1/β-propeller 7)** A/876, H/881, I/906 |
| **16** | **(APAF-1/β-propeller 7)** V/746, S/752, L/758, S/762, D/764, L/767 |
| **17** | **(APAF-1/β-propeller 7)** H/698, L/716 |
| **18** | **(APAF-1/HD2)** V/576, A/584 |
|  | **(APAF-1/L3)** L/595, W/597 |
| **19** | **(APAF-1/HD2)** E/499, L/500, A/503, L/504, L/507, W/509, I/510, I/523, Y/529, L/533 |
| **20** | **(APAF-1/HD2)** I/460, I/461, Y/482, L/487, H/490, M/491 |
| **21** | **(APAF-1/β-propeller 7)** F/787, V/799, V/801 |
|  | **(CYCS)** K/1692 (28) |
| **22** | **(CASP9/Large subunit)** G/1409 (161), L/1412 (164), I/1413 (165), I/1414 (166), N/1415 (167), N/1416 (168), F/1419 (171), L/1438  (190), F/1447 (199), V/1449 (201) |
|  | **(CYCS)** A/1708 (44) |
| **23** | **(APAF-1/β-propeller 7)** F/751, D/755 |
