## Supplementary material for "Protein coevolution and physicochemical adaptation in the APAF-1/apoptosome: structural and functional implications": Figure 5-figure supplement 1-Clusters of aminoacids with coevolution signal mapped in tridimensional structures

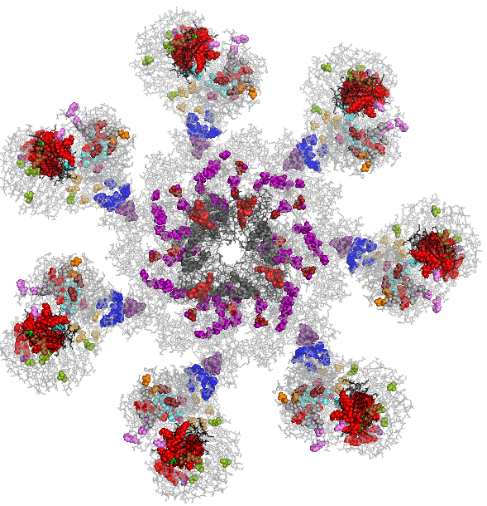

**Α**

**Regulatory region**

**CYCS**

**β- propeller 8**

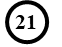

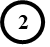

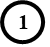

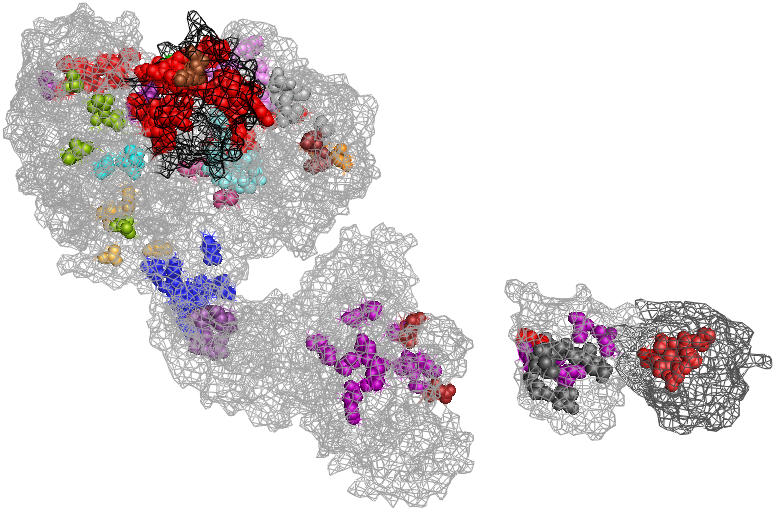

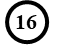

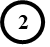

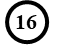

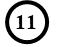

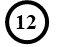

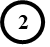

**β- propeller 7**

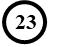

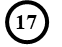

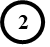

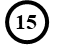

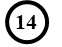

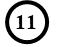

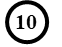

**CASP9**

**CARD**

**APAF-1**

**CARD**

**HD1**

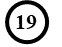

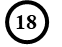

**p20/p10**

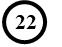

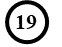

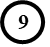

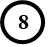

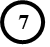

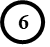

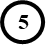

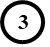

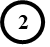

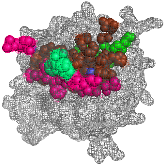

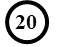

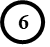

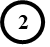

**HD2**

**ARM**

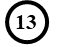

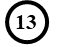

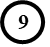

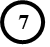

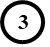

**WHD**

**Front view**

**L1**

**NBD**

**Regulatory region**

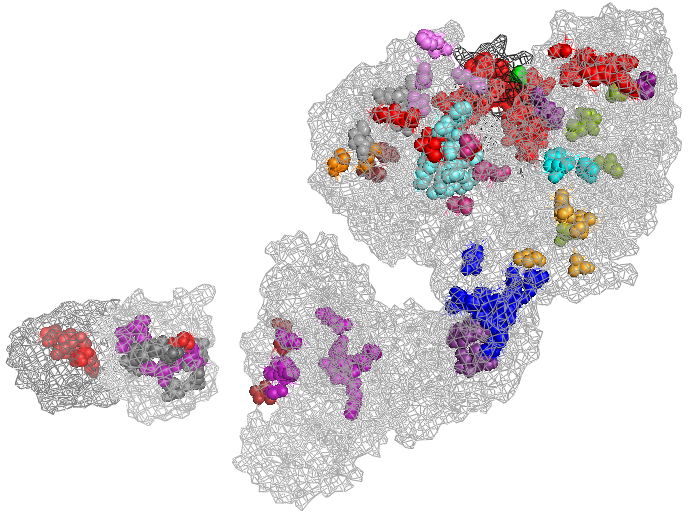

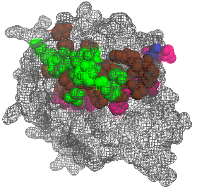

**Β**

**180°**

**CYCS**

**β- propeller 8**

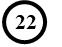

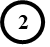

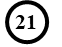

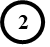

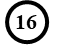

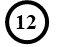

**β- propeller 7**

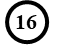

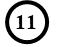

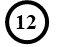

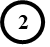

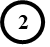

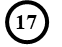

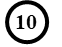

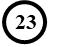

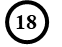

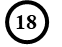

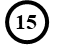

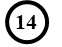

**CASP9**

**CARD**

**APAF-1**

**CARD**

**HD1**

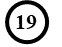

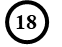

**p20/p10**

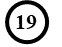

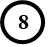

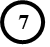

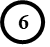

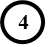

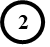

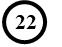

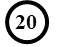

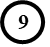

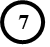

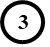

**HD2**

**ARM**

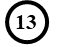

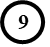

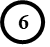

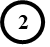

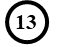

**L1**

**WHD**

**Back view**

**NBD**

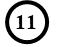

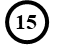

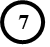

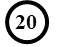

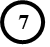

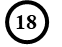

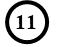

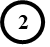

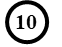

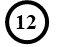

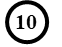

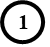

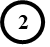

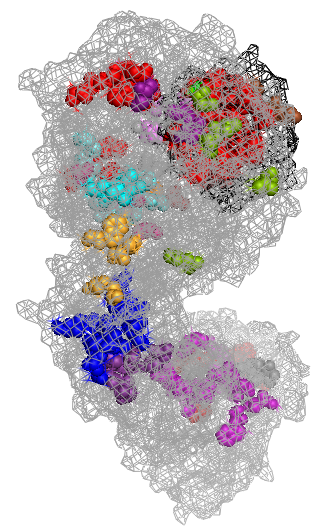

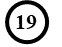

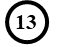

**β- propeller 8**

**Regulatory region**

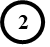

**C**

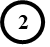

**CYCS**

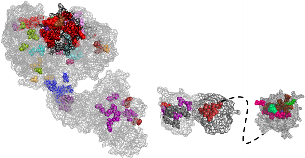

**90°**

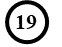

**HD2**

**ARM**

**APAF-1**

**CARD**

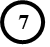

**Side views**

**D**

**Regulatory region**

**NBD**

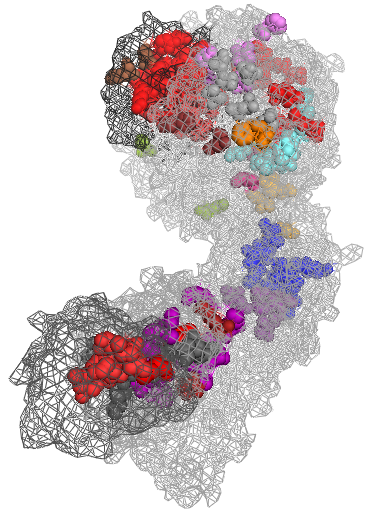

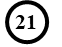

**CYCS**

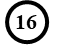

**β- propeller 7**

**APAF-1**

**CARD**

**90°**

**HD2**

**ARM**

**p20/p10**

**NBD**

**E**

**90°**

**Top view**

**β- propeller 8**

**β- propeller 7**

**NBD**

**CASP9**

**CARD**

**APAF-1**

**CARD**

**p20/p10**

**L1**

**CYCS**

**Regulatory region**

Cluster 6

Cluster 7

Cluster 8

Cluster 9

Cluster 10

Cluster 11

Cluster 12

Cluster 13

Cluster 14

Cluster 15

Cluster 20

Cluster 19

Cluster 18

Cluster 17

Cluster 16

Cluster 21

Cluster 22

Cluster 23

Cluster 1

Cluster 2

Cluster 3

Cluster 4

Cluster 5

**F**

**180°**

**Back view**

**Back view**

**C/287**

**H/237**

**H/237**

**C/287**

Cluster 22

Cluster 3

Cluster 4

Cluster 5

**G**

**Front view**

**180°**

**Back view**

**90°**

**90°**

**Side views**

**90°**

**Top view**

Cluster 22

Cluster 1

Cluster 2

Cluster 21

Cluster 12

**Figure 5-figure supplement 1-Clusters of aminoacids with coevolution signal mapped in tridimensional structures.** A single unit amplification is shown within the PDB 5JUY heptameter in “mesh” format, which allows you to see the residues mapped in the deep parts. The catalytic domain of CASP9 PDB 1JXQ was attached, representing L1 with intermittent lines. **A)** The unit is viewed from the front, **B)** by turning it 180° to the right showing a back view. **C)** 90° to the left. **D)** 90° to the right. **E)** 90° downwards showing a top view. The amino acids within each cluster were marked with a specific color. To facilitate the visualization of these groupings, the numbers of the clusters to which they belong were placed near their respective residues. **F)** Amplified visualization of the catalytic domain of CASP9 PDB 1JXQ, where the clusters with coevolution signal are shown as spheres with their respective colors shown in the legend. Active site C/287 and residue H/237 are marked. The protein domain is visualized from the front, turning it 180° to the right showing a back view. **G)** Enhanced visualization of CYCS PDB 1J3S, where the clusters with coevolution signal are shown as spheres with their respective colors shown in the legend. The protein is shown in a back view, in its original position when it is in interaction with APAF-1, making a 180° turn, showing the front part where the heme group can be seen as yellow sticks, 90° to the left , 90° to the right showing side views and 90° up showing the top.
