## Supplementary material for "Protein coevolution and physicochemical adaptation in the APAF-1/apoptosome: structural and functional implications": Figure 6-figure supplement 1-TreeSAAP sliding windows aminoacids under destabilizing positive selection

**A. Helical contact area (*Ca*)**

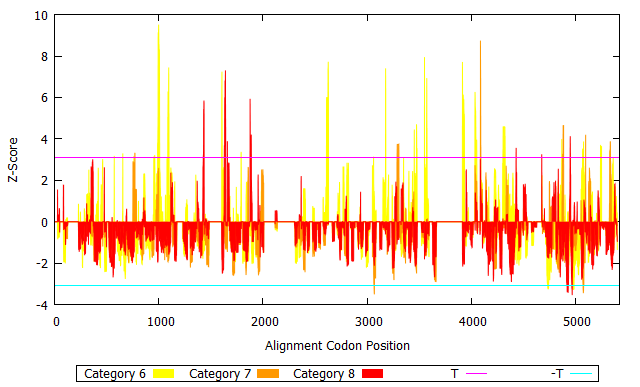

**B. Compressibility (*K0*)**

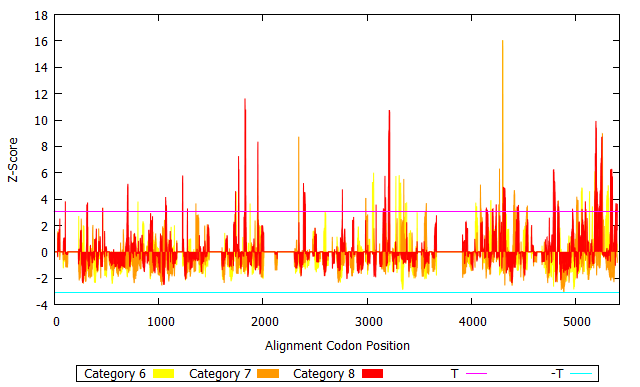

**C. Hydrophobicity by thermodynamic transfer (*Ht*)**

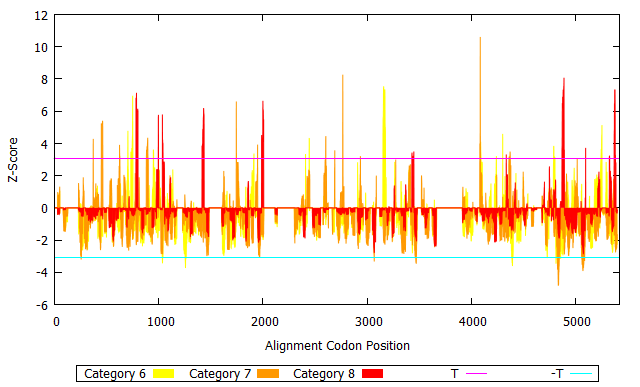

**D. Chromatographic index (*RF*)**

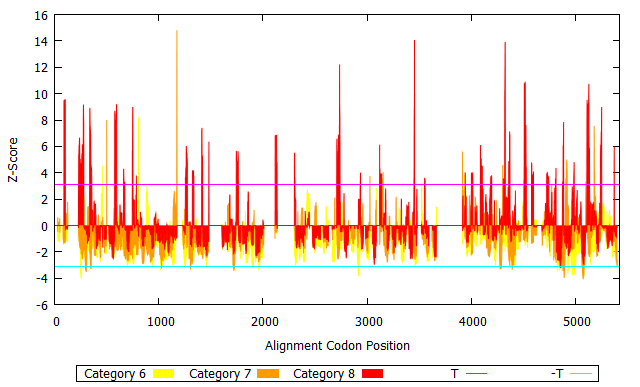

**E. Refractive index (*µ*)**

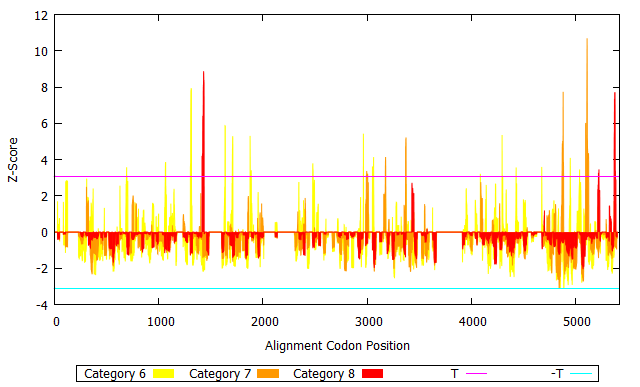

**F. Molecular weight (*Mw*)**

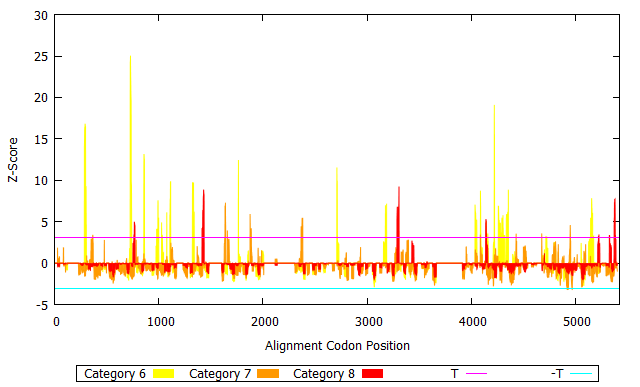

**G. Bulkiness (*Bl*)**

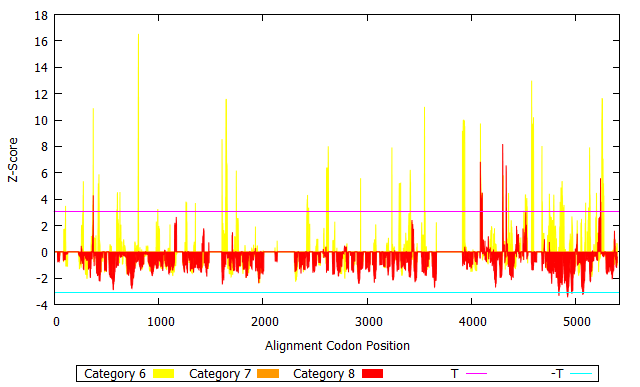

**H. Partial specific volume (*V0*)**

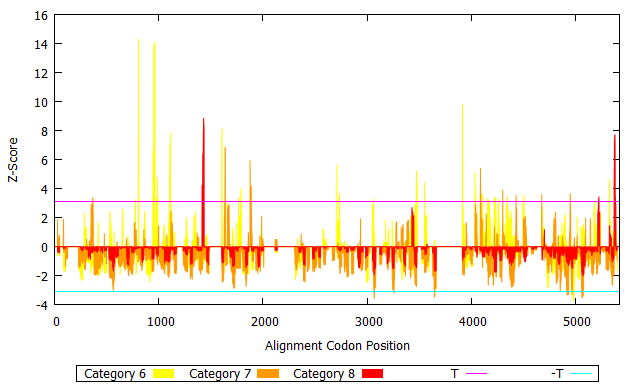

**I. Molecular volume (*Mv*)**

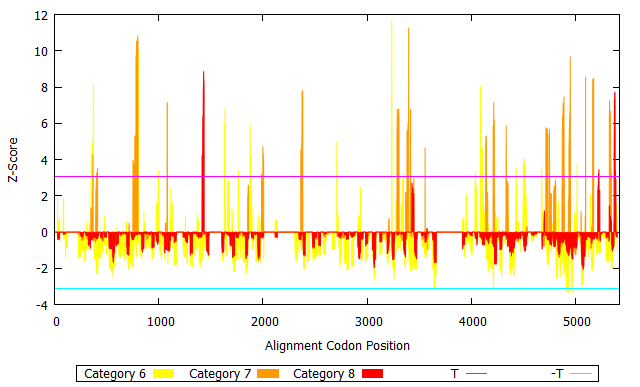

**Supplementary material 11. Amino acids under destabilizing positive selection shown by property.** **(A-I)** For the nine significant physicochemical properties, their respective sliding windows are shown, with the residues under destabilizing changes in the categories of magnitude 6, 7 and 8, colored yellow, orange and red, respectively. In each one the X-axis shows the 5,426 positions of the alignment length of the concatenated codon/protein sequences. APAF-1 CARD 1-259, L1 360-399, NBD 411-753, HD1 757-864, L2 865-909, WHD 910-1,025, HD2 1,026-1,271, L3 1,272-1,337, β propeller 7 1,338-2,595, L4 2,596-2,762, β propeller 8 2,763-4,056, CASP9 CARD 4,108-4,414, L1 4,415-4,825, p20 4,826-5,010, L2 5,011-5,033, p10 5,034-5,226 and CYCS 5,264-5,426. The Y-axis shows the z-score values. On this axis, the purple horizontal line in the positive quadrant and the turquoise blue horizontal line in the negative quadrant indicate the values +3.09 (T) and -3.09 (-T), respectively. Amino acids above T are under destabilizing changes with a positive tendency (increased property). Residues below –T are under destabilizing changes with a negative tendency (decrease in property).
